# Mechanism of catalytic apparatus of human chitotriosidase-1 and its dual inactivation mode by the *first-in-class* OATD-01 inhibitor

**DOI:** 10.1101/2024.12.18.628861

**Authors:** Dorota Niedzialek, Grzegorz Wieczorek, Katarzyna Drzewicka, Anna Antosiewicz, Mariusz Milewski, Agnieszka Bartoszewicz, Jacek Olczak, Zbigniew Zasłona

**Author notes:** Corresponding authors, Correspondence to Dorota Niedzialek and Zbigniew Zaslona.

## Abstract

Despite extensive research over the past three decades, there are still uncertainties regarding the catalytic mechanism of human chitotriosidase-1. To fill the gap, we reanalysed the structural information available for this enzyme. Based on the existing and new experimental data, complemented by multi-scale simulations, we modelled the full-length structure of human chitotriosidase-1 and proposed the general model of its catalytic mechanism. We have elucidated the catalytic role of the four highly conserved structural motifs present in glycoside hydrolases 18 family and demonstrated the impact of ions on achieving optimal catalytic conditions. Furthermore, we have identified distinct mechanical motions within the catalytic domain that collectively facilitate the catalysis. Finally, we demonstrate how subtle dynamical changes observed within the active site upon binding of the OATD-01 inhibitor correspond to long-range effects that are transmitted across enzyme subunits, leading to profound biological consequences.

## Main

Humans neither produce chitin nor utilize it as a nutrient source, yet the human genome encodes several proteins, which belong to the glycoside hydrolase 18 (**GH18**) family of chitinases (EC 3.2.2.14) – enzymes that break glycosidic bonds in β-1,4-*N*-acetylglucosamine (**GlcNAc**) [1][2]. GlcNAc is a widely occurring substance in nature that serves as a crucial structural sugar in various organisms. It is found in bacterial peptidoglycan and fungal chitin cell walls. GlcNAc-containing oligosaccharides are present in the extracellular matrix of animal cells, where they form glycans that, for example, decorate glycosylated proteins and glycolipids. Chitinases are expressed in a wide range of organisms from prokaryotes to eukaryotes. Humans produce three enzymatically active GH18 chitinases, Di-*N*- acetylchitobiase (**hDIAC**), chitotriosidase-1 (**hCHIT1**) and acidic mammalian chitinase (**hAMCase**) (**ExtendedData Fig. 1**). The rest are chitinase-like proteins which, due to mutations of the catalytic residues, bind but do not catalyse glycans [3].

The ability to degrade chitin suggests primordial role of hCHIT1 in innate defence mechanisms against GlcNAc-producing pathogens [4] (**ExtendedData Fig. 2A/E**). However, the role of hDIAC in the degradation of glycoproteins by hydrolysing the glycosidic bond in the (GlnNAc)_2_ core of asparagine-linked glycans, advocates additional roles for human chitinases, for example, through involvement in protein glycosylation [5]. GlcNAc-containing *N*-acetylglucosamine (**LacNAc**) is the main ligand for sugar-binding receptors [6] known to be modified by hCHIT1 [7] (**Fig. 1D**). Abnormal expressions of LacNAc are associated with cancers [8], autoimmune [9] inflammatory [10] and lysosomal storage diseases associated with overexpression of hCHIT1 [11]. Since hCHIT1 exhibits high transglycosylation activity independent of typical ‘transglycosylation conditions’ [12], it may play a harmful role in the human body by contributing to pathological glycosylation processes (**ExtendedData Fig. 2B/C**) [13][14]. For instance, a recent study demonstrated that increased expression and activity of hCHIT1 are linked to Alzheimer’s disease [15]. This result is consistent with previous findings that genetic inactivation of hCHIT1 is strongly associated with human longevity and several phenotypes of healthy ageing [16]. Other evidence suggests that some genetic variations that disrupt the enzymatic activity of hCHIT1 protect against the progression of non-alcoholic fatty liver disease [17] and reduce susceptibility to sarcoidosis [18]. In healthy individuals, macrophages produce low levels of hCHIT1. However, in response to pro-inflammatory signals, the secretion of hCHIT1 significantly increases [19][20]. Furthermore, GlcNAc oligomeric products generated by hCHIT1 function as molecular patterns that can be detected by toll-like receptors TLR9 in neutrophils and NOD-2 in macrophages [21] (**ExtendedData Fig. 2D**). This, in turn, may lead to induction of enhanced hCHIT1 expression, production of various cytokines and chemokines, and initiation of adaptive immune responses [22]. Hence, hCHIT1 seems to have different pathogenic roles depending on the associated diseases and activated cell types during the immune reaction [23].

**Figure 1.**
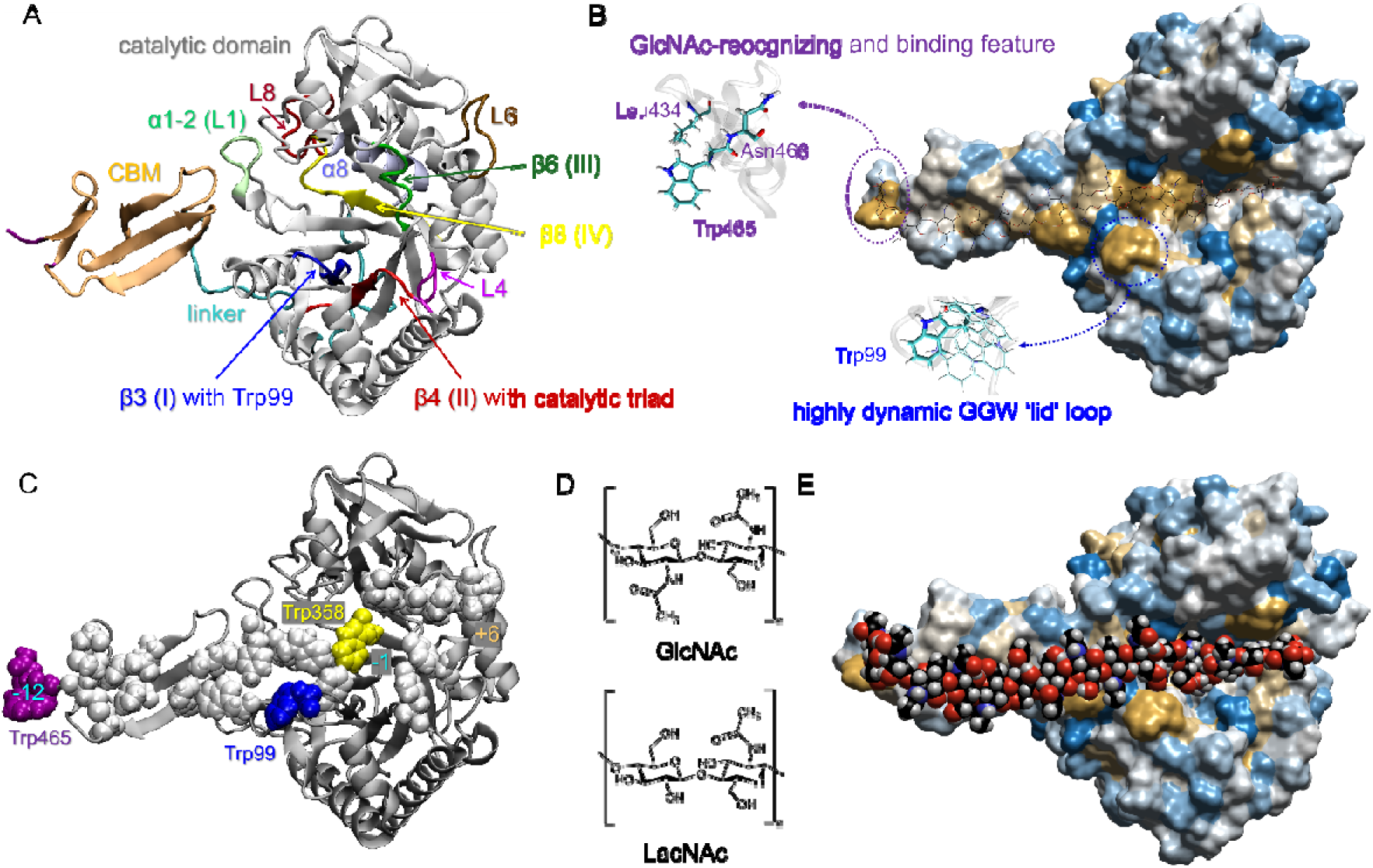
The model of the full-length hCHIT1. **A**) The cartoon representation of the 50-kDa hCHIT1 heterodimer. The distorted β-sandwich fold of the carbohydrate binding domain (CBM, residues 420–466 marked in orange, PDB ID: 6SO0) is attached to the catalytic domain by a proline-rich linker (residues 388–419 marked in cyan) and topped by the GlcNAc-recognizing feature (residues Trp465 and Asn466 marked in purple). The conserved motifs I (residues 90–99 within β3 strand), II (residues 132–142 within β4 strand), III (residues 210–216 within β6 strand) and IV (residues 354–364 within β8 strand) are marked in blue, red, green, and yellow, respectively. The L4 loop which extends from motif II (marked in magenta) contributes to the ‘flywheel’ mechanism for the enzyme-opening apparatus (see Fig. 4A). **B**) Two highly conserved tryptophane-containing structural motifs in the full-length hCHIT1. The first is the highly flexible tryptophan ‘lid’ at the entrance to the active site. The second is the platform-like feature, which is responsible for substrate recognition and binding. The surface representation of the hCHIT1 model coloured according to hydrophobic/hydrophilic (yellow/blue) character of amino acids. **C**) The cartoon representation of the full-length hCHIT1 model with the hydrophobic residues in the vdW representations. The key sugar binding sites: Trp99, Trp358 and Trp465 are coloured in purple blue, yellow and purple, respectively. Note that solvent-exposed aromatic residues arranged linearly at a distance equivalent to the spacing between (GlcNAc)2 units. **D**) The substrates of chitinases: β-1,4-*N*-acetylglucosamine (GlcNAc) and GlcNAc-containing *N*-acetylglucosamine (LacNAc). **E**) The same representation as in panel B with a docked model of (GlcNAc)8 substrate. GlcNAc oligosaccharides in vdW representation occupy the sugar-binding subsites, ranging from-12 to +4, where-n and +n indicate the nonreducing and reducing end, respectively (*i.e.*, the substrate and product occupying subunits).

**Figure 2.**
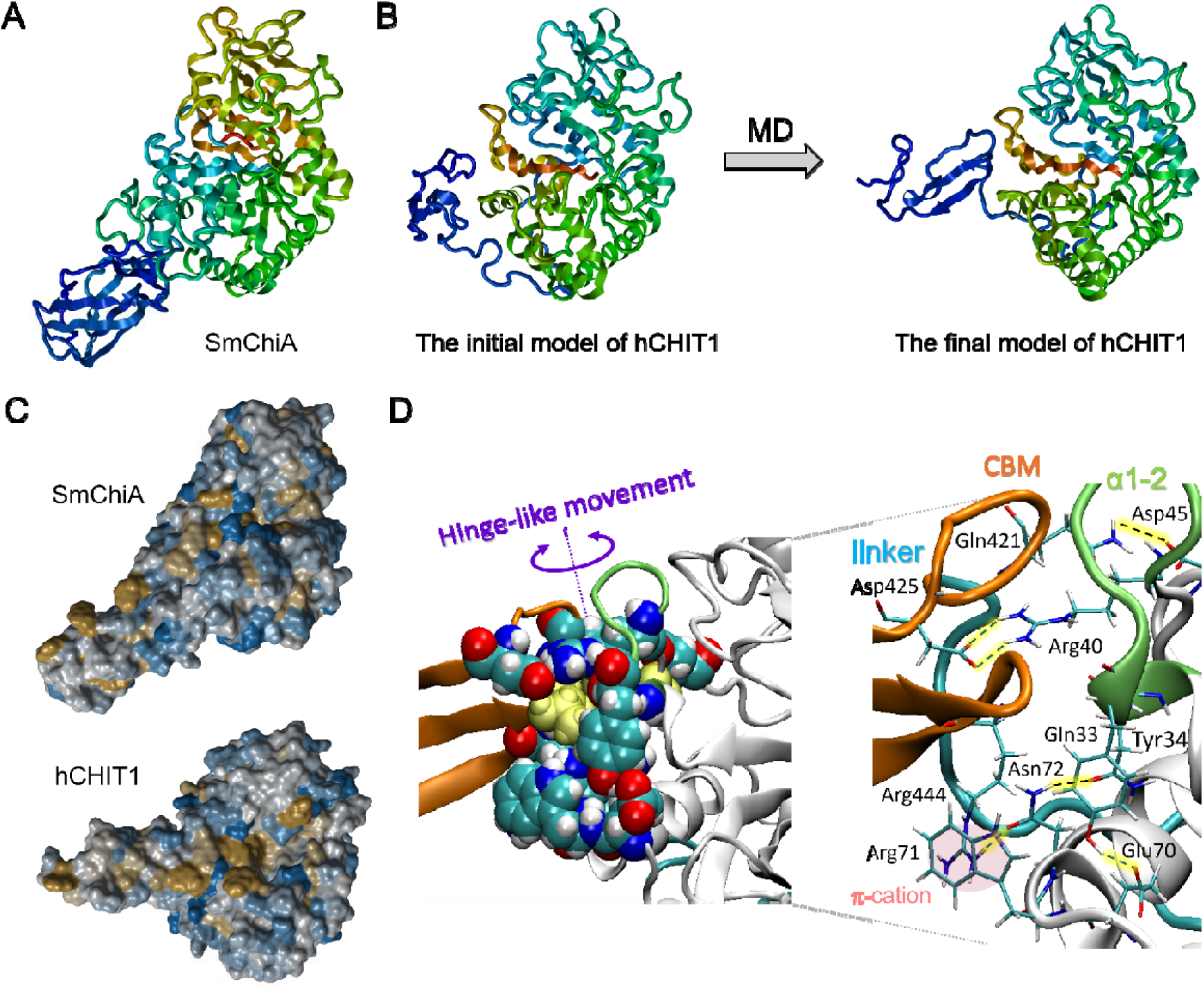
Modelling of the full-length hCHIT1 which, similarly to Chitinase A from *Serratia marcescens* (SmChiA), structurally belongs to the ‘bacterial-type’ chitinases. **A**) The rainbow cartoon representation of SmChiA (PDB ID: 5Z7M). **B)** The initial model of the full-length hCHIT1 was based on the 5HBF PDB entry with a structure of the unliganded hCHIT1 catalytic domain cocrystalised with CBM (without the linker). After modelling the missing linker, the structure was subjected to a series of extensive MD simulations until the final stable model of the immunoglobulin-like hCHIT1 heterodimer was obtained. **C**) The surface representations of the superimposed SmChiA structure (top) and the full-length hCHIT1 model (bottom). Both enzymes are coloured according to hydrophobic/hydrophilic (yellow/blue) characters of amino acids. Note the similar substrate binding grooves with hydrophobic residues arranged linearly that continue along the carbohydrate-binding domain, promoting efficient binding of polysaccharide substrates through CH-π interactions. **D)** The network of non-covalent interactions between the catalytic and CBM domains after assembly into the 50-kDa immunoglobulin-like heterodimer. The enzyme in cartoon representation is coloured as in Fig. 1A. The key residues shown in the left and right panels are in vdW and liquorice representations, respectively, and are coloured according to atom types, except for the hydrophobic residues that are in the core of the assembly (*i.e.*, Leu42 within the α1-2 subunit is accommodated between Ala441 and Ala442 on CBM) which are marked in yellow. The hydrophobic residues are not shown in the right panel for clarity. The attachment of the domains is tight, so that the heterodimer remains stable at room temperature. However, it is flexible enough to perform a slight hinge-like movement that allows the processed polysaccharide chain to slide towards the active site after each catalytic cycle.

hCHIT1 was the first mammalian chitinase discovered and characterized by *Aerts* and *co-workers* 30 years ago [24]. The full-length, 50-kDa hCHIT1 isoform, consists of the 39-kDa catalytic domain located at the N-terminus and the carbohydrate-binding module (**CBM**), connected by a 31-amino acid long linker (**Fig. 1A**). The experimental studies demonstrated that CBM is responsible for the high affinity of the full-length hCHIT1 towards GlcNAc-containing polysaccharides [25–27]. The GlcNAc identification and binding mechanism relies on a platform-like feature composed of adjacent tryptophan and asparagine that were shown to specifically bind acetylated glycans (**Fig. 1B**) [28][29]. The catalytic domain comprises the classic TIM-barrel fold with four regions highly conserved among all GH18 enzymes. The site-directed mutagenesis identified the catalytic triad within motif II, located on the β4 strand. The functions of the other three conserved motifs are still unknown [30]. The catalytic domain of hCHIT1 contains a long and deep substrate-binding cleft, which contains several solvent-exposed aromatic residues arranged in a linear fashion (**Fig. 1C**). These residues continue along the CBM and facilitate the efficient binding of polysaccharide substrates through CH–π interactions (**Fig. 1E**). A substantial number of possible contacts between GlcNAc substrate and the active site cleft provides strong binding that forces a distorted conformation of the-1 sugar and brings the susceptible glycosidic bond (*i.e.*, between-1 and +1 GlcNAc units) in proximity to the proton donor (**ExtendedData Fig. 3A/B**). The architecture of the catalytic groove and the active site in hCHIT1 resembles one from the most studied example of the GH18 family – *Chitinase A* from *Serratia marcescens* (**SmChiA**) – suggesting a common mechanism for hydrolysing GlcNAc from the reducing end (**Fig. S2A/C**) [31–33]. As both enzymes are classified as ‘bacterial-type’ chitinases [1], we expected full-length hCHIT1 to exhibit an analogous heterodimeric, immunoglobulin-like architecture. However, unlike SmChiA, which has firmly attached catalytic and carbohydrate-binding domains, the hCHIT1 heterodimer should exhibit an ‘elbow-bending’ dynamics thanks to its proline-rich linker [34].

Despite the wealth of crystallographic data for hCHIT1 collected during the last two decades, crystal structures of neither the full-length isoform of the enzyme nor a complex of its catalytic domain with a substrate have been solved to this day [35]. The lack of such critical structural information for hCHIT1 allowed for numerous interpretations of the existing data regarding possible enzymatic mechanisms [1][36]. Furthermore, conformational changes that are relevant to catalytic mechanisms are difficult to determine from static crystal structures alone. For example, despite the high conservation of tryptophan at the entrance of the active site in chitinases across species, the role of this residue has not been elucidated.

The objective of this study was to expand upon the existing understanding of the catalytic apparatus of hCHIT1 and fully comprehend the molecular mechanism of its inhibition by the *first-in-class* OATD-01 inhibitor, which is currently undergoing phase 2 clinical trials for the treatment of sarcoidosis [37–39].

To unravel the mechanism of the enzymatic apparatus of hCHIT1, we modelled the full-length enzyme based on the available structural information and experimental data obtained during our pre-clinical studies on hCHIT1 inhibitors (**Fig. 1**). Our extensive study involved classical and quantum mechanics/molecular mechanics (**QM/MM**) molecular dynamics (**MD**) simulations as well as experimental assays in different ionic conditions. By investigating conformational changes along the simulated enzymatic reaction trajectories, we discovered intrinsic local rearrangements within the active site of hCHIT1 that correlated with long-range inter-subunit motions. The observed sequence of multiscale conformational changes is essential for the substrate accommodation, creation of the intermediate states and product dissociation during the substrate-assisted catalytic process. Furthermore, we unravelled the catalytic roles of all four conserved motifs in GH18 family and discussed their coordinated involvement in the dynamic catalytic machinery of hCHIT1. Finally, we demonstrated that, by blocking individual components of the hCHIT1 catalytic apparatus, OATD-01 small molecule not only inactivates the enzyme, but, also, acts as an allosteric inhibitor of its interactions with potential biological partners involved in pathological immune responses.

## Results

Previous studies on chitinases did not focus on the catalytic role of the tryptophan at the entrance of the active site. Our first observation from MD simulations of unliganded hCHIT1 was the significant mobility of the Trp99 side chain located on a polyglycine loop (**GGW**) within the β3 strand, which alternately opened and closed the active site (**Fig. 1B**). The cycle of opening and closing the Trp99 ‘lid’ took only a few nanoseconds, which was of an order of magnitude faster than other local movements observed during simulations, and significantly faster than the catalytic turnover rate (∼1.5 seconds). Furthermore, we observed that the enzymatic activity of hCHIT1 is divided into several successive stages (**Fig. 3A**). In the initial, inactive state, the catalytic triad (*i.e.,* residues Asp136, Asp138 and Glu140 within the β4 strand) remains concealed within the active site cleft until the residues Leu135, Leu137, and Trp139, located within the hydrophobic core beneath the catalytic triad, begin to rotate through various rotamers and shift by exchanging van der Waals (**vdW**) contacts with residues from the hydrophobic core of the enzyme (**Fig. 3B/C**). The ‘flywheel’ of the β4 strand slide is the hinge-like movement of the adjacent loop, triggered by thermal fluctuations (**L4** in **Fig. 4A**). The shift of the β4 strand causes large conformational changes of its main chain that exposes the catalytic residues towards the bulk solvent. The final step in hCHIT1 activation, prior to substrate binding, is the rotation of the central residue of the catalytic triad (Asp138), driven by electrostatic interactions. hCHIT1 possesses a substantial dipole moment aligned with the active site which attracts polar GlcNAc-containing substrates and positive ions by the ‘torch of guidance’ effect [40] (**ExtendedData Fig. 4A**). Strong electrostatic attraction between the active site and the GlcNAc initiates and guides the substrate binding and stabilization of the positively charged intermediate state (*i.e.,* oxazolinium). Docking the substrate pushes the β4 strand back into the active site cleft. Further accommodation of the substrate involves its formation of many strong contacts with residues from-2 to +1 subsites of the binding cleft, which impose a distortion on the-1 sugar from the default chair into a (∼30 kJ/mol less energetically favourable) twisted boat conformation, allowing its 2- acetamido group to accommodate above Asp138 side chain (**Fig. 5A**).

**Figure 3.**
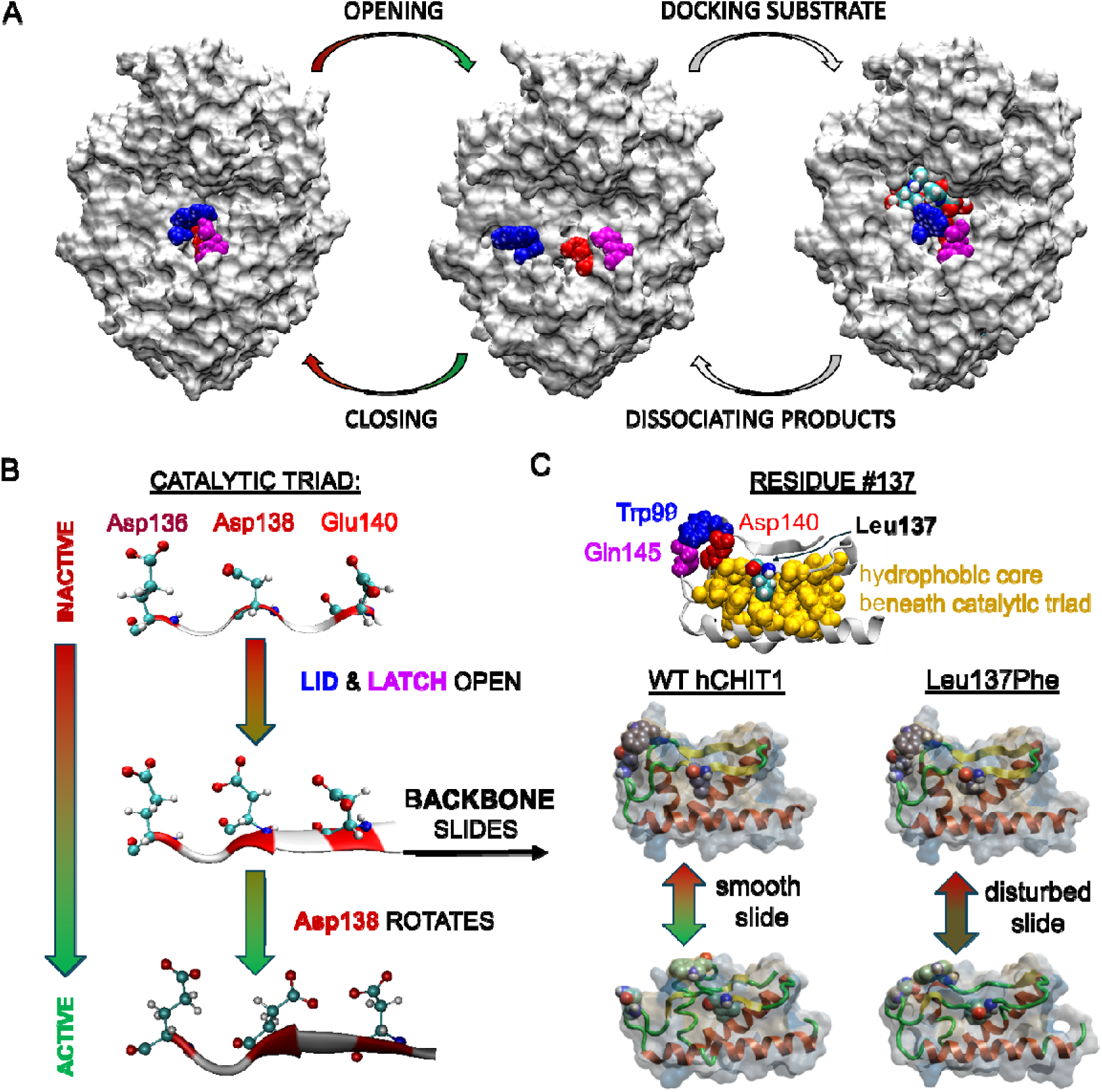
The multiscale activation of hCHIT1 enzyme. **A)** The three main conformations adopted by hCHIT1 during the catalytic cycle. The enzyme is represented as a white surface. Residues Trp99 (‘lid’), Glu140 (proton donor) and Gln145 (‘latch’) as well as the (GlcNAc)3 substrate are in the vdW representations, coloured in blue, red, magenta and according to the atom types, respectively. **B)** After opening the active site by ‘lid’ and ‘latch’ motifs, the main chain of the sliding β4 strand extends, while the side chain of Asp138 rotates towards Glu140, adopting an optimal position to bind 2-acetamido group of the-1 sugar. **C)** The lateral sections through the regions surrounding the β4 strands in hCHIT1 and its L137F mutant, both subjected to 300 nanosecond MD simulations. The β4 strand of the wild-type (WT) enzyme exhibits uniform sliding behaviour on adjacent hydrophobic surfaces, undergoing extensive conformational changes that result in the exposure of the active site towards the bulk aqueous solution prior to substrate binding. In the case of the L137F mutant, we observed distortions of the main chain within the catalytic triad, which resulted in a disruption of the β4 strand slide. This effect reduces enzymatic activity of the L137F mutant in comparison with the WT enzyme due to increased disorder (less β-sheet character) of the β4 strand in the case of the activated Leu137Phe mutated enzyme. The Trp99, Gln145 and Leu/Phe137 residues are in vdW representations while the rest is represented as cartoons and surfaces, coloured according to the hydrophobic/hydrophilic (yellow/blue) character of the amino acids. Note that motif II of the closest homologue of hCHIT1 (hAMCase) and its mouse version (mAMCase) differ only in this residue (*i.e.*, mAMCase, similarly to hCHIT1, possesses leucine while hAMCase phenylalanine (see ExtendedData Fig. 2).

**Figure 4.**
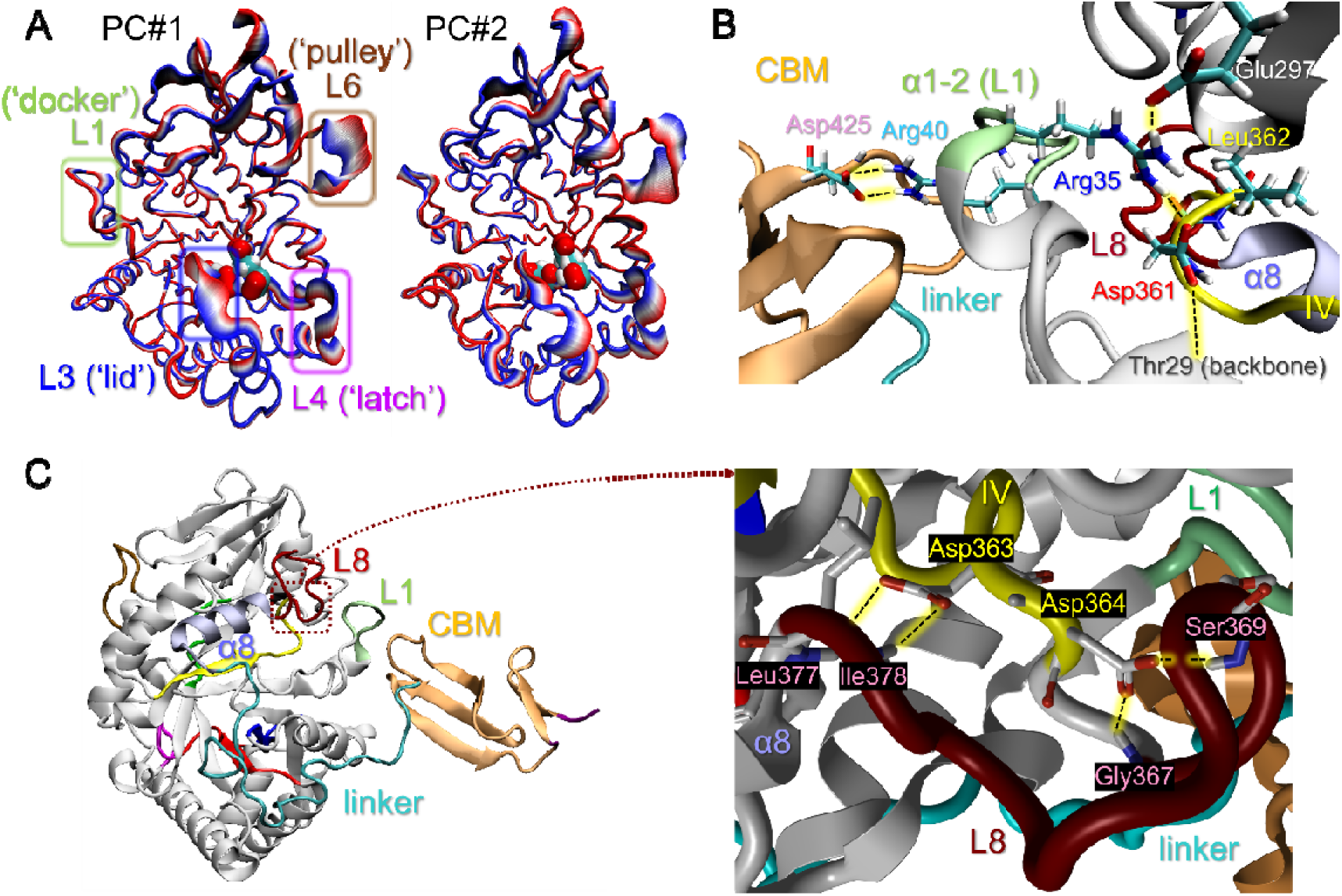
The dynamic features of the catalytic apparatus of hCHIT1. **A)** The two first modes of the principal component (PC) analysis indicate substantial dynamics of the loops emerging from the β3, β4 and β6 strands (L3, L4 and L6, respectively). Note that the ‘docker’ (L1) loop within the α1-2 subunit, which is involved in docking the CBM, also exhibits considerable dynamics. The coordinated movement of the loops L3 and L4 (harbouring the ‘lid’ and ‘latch’ residues, respectively) is required for opening/closing the active site. The substantial movement of the ‘pulley’ (L6) loop triggers the Tyr212-assisted pulling mechanism facilitating the substrate assisted GlcNAc catalysis (see Fig. 5). The enzyme is presented as a cartoon coloured in RWB colour scale, with the active site residues in vdW representation coloured by the atom types. **B**) The network of hydrogen bonds (marked as black dotted lines on a yellow background) connecting catalytic and carbohydrate-binding domains in the hCHIT1 heterodimer. The enzyme is in cartoon representation coloured as in Fig. 1A. The key residues are in licorice representation. Residue Asp361, which belongs to the conserved motif IV, is involved in the molecular ‘wire’ made of two salt-bridges (*i.e.*, Asp361–Arg35 and Arg40–Asp425) that promote optimal orientation and flexibility of the L1 loop (coloured in lime) for the assembly and stability of hCHIT1 heterodimer. **C**) The network of hydrogen bonds between conserved residues Asp363 and Asp364 and the L8 loop (dark red), which is adjacent to the α8 helix (violet) that serves as the anchor point of the proline-rich linker. These hydrogen interactions play pivotal role in mechanical signalling along the linker and the induction of its ‘elbow-like’ movements during assembly of the catalytic and carbohydrate-binding domains into heterodimer. The key residues are in licorice representation coloured according to the atom types.

**Figure 5.**
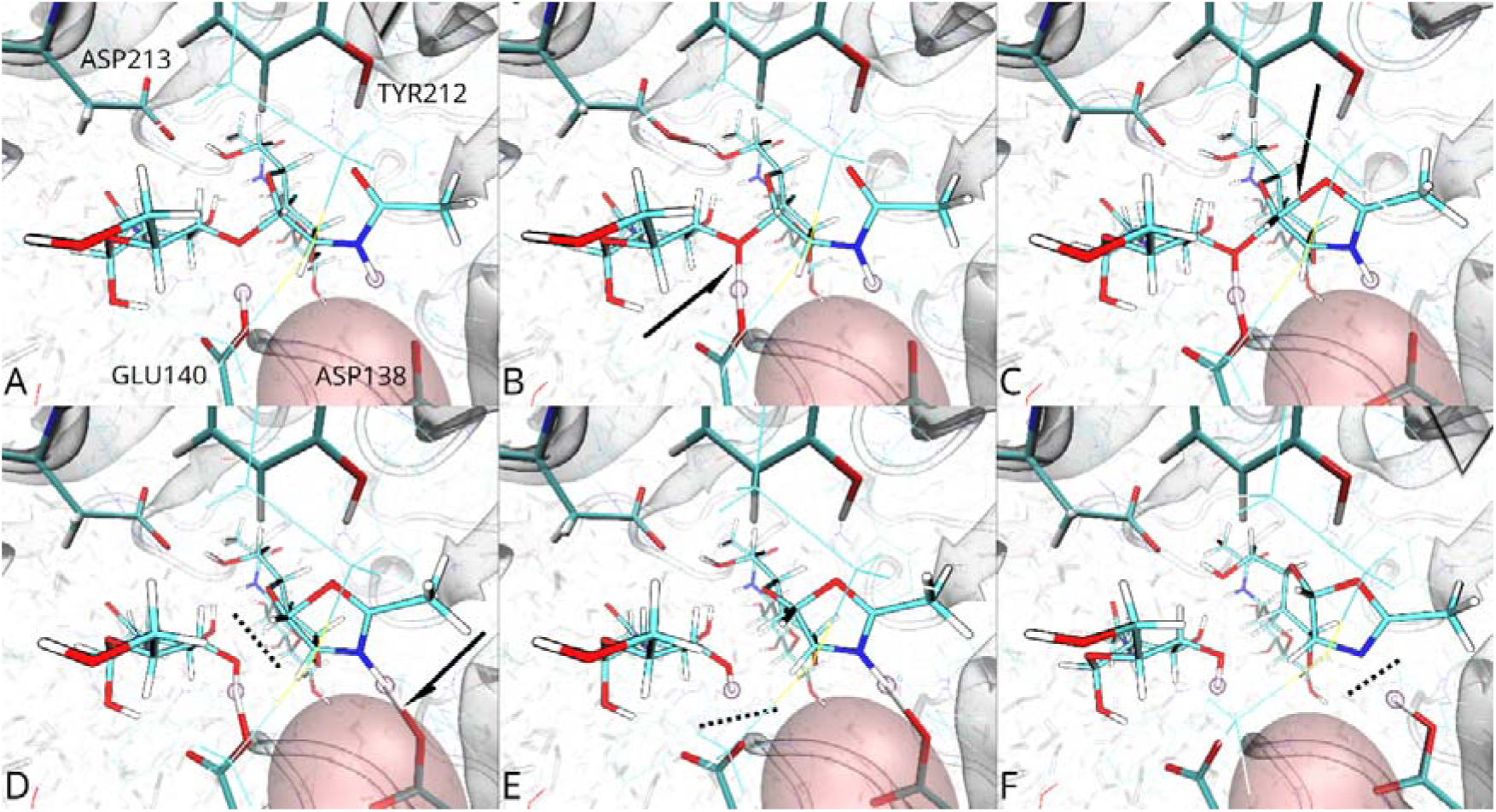
Anhydrous stages of the hCHIT1 catalytic process elucidated from QM/MM simulations. **A)** (GlcNAc)3 substrate gets optimally accommodated inside the active site with the glycosidic bond between-1 and +1 GlcNAc units in proximity to the protonated Glu140 residue. The 2-acetamido group of the-1 sugar is fixed in a correct conformation for substrate-assisted catalysis with the nitrogen and carbonyl groups forming tight hydrogen bonds with Asp138 and Tyr212 residues, respectively. **B)** The proton sharing between Glu140 and the susceptible glycosidic bond. The side chain of Tyr212 pushes the carbonyl oxygen towards the entrance of the active site, while the nitrogen of the 2-acetamido group, bound to Asp138 residue, is moving in the opposite direction; **C)** The carbonyl oxygen acts as a nucleophile and being pushed by Tyr212 finally reaches the C1 anomeric centre. **D)** The nucleophilic attack on the anomeric carbon creates covalent bicyclic oxazolinium intermediate and the glycosidic bond breaks; **E)** Residue Glu140 transfers its proton to the oxygen of the glycosylic bond to promote the leaving group departure; **F)** The proton between nitrogen and Asp138 oscillates during the formation/breakdown of the oxazolinium/oxazoline ring. Departing +1 sugar pops out of the binding cleft and releases the Trp99 ‘lid’ which opens, letting water molecule in the active site and thus initiating the water-dependent stage of the catalysis (see Fig. S6). Black arrows and dotted lines indicate emerging and broken bonds, respectively. The pink sphere represents potassium cation.

The MD simulations of the hCHIT1–(GlcNAc)_3_ model system showed that the GGW loop closes the active site with the Trp99 ‘lid’ upon substrate binding. Furthermore, the tryptophane ‘lid’ gets tightened by the 2-acetamido group of the +1 sugar and thereby the substrate locks itself in the active site from the aqueous environment (**Fig. 3A**). This strategy protects the unstable enzymatic reaction intermediate from decomposition by bulk water at the initial, anhydrous stage of the catalysis. The intermediate state is formed when the 2- acetamido group of the-1 sugar closes to form an oxazolinium/oxazoline ring [41], forced by Tyr212 side chain (within β6), which forms a hydrogen bond with the carbonyl oxygen of the-1 sugar and pushes the oxygen in the direction of the anomeric carbon (**Fig. 5C**). After the anhydrous stage of catalysis, the +1 sugar leaves the binding cleft. The untightened tryptophane ‘lid’ reopens, allowing water to enter the active site, starting the aqueous stage of the catalysis. Water breaks the oxazolinium/oxazoline ring and the-1 sugar detaches from the active site. After product release, the active site residues rearrange to their initial positions, completing the catalytic cycle (*vide infra*).

During MD simulations of unliganded hCHIT1, we observed correlated side chains rotation of Asp138 and the nearby Met210 with the motions of cations entering the active site from the bulk solvent. Two orientations (*i.e.*, facing either the Asp136 or Glu140 residues) of Asp138 were known from the crystal structures of unliganded hCHIT1 [36]. We have identified two regions of the highest cation occupancy, both associated with catalytic residues. The first was found between side chains of Asp138, Glu140 (β4 strand), Met210 (β6 strand), and Trp358 (β8 strand) (**ExtendedData Fig. 4B**). The second, broader region, was observed along the main chain, between residues Asp138 and Glu140, on the site of Met210. Significantly, we found excess densities in similar locations on the difference density maps in crystal structures of unliganded hCHIT1 (**ExtendedData Fig. 4C**) and in complex with OATD-01 inhibitor (**ExtendedData Fig. 3C**) – both identified as consistent with alkali metal ions. To increase the fidelity of these intriguing observations, we repeated the simulations with polarizable force field. Again, the simulated monovalent cations migrated to both previously identified regions. Furthermore, deficiency of cations (*i.e*., in simulations run at much lower than physiological ions concertation) led to a significant distortion of the catalytic triad with the Asp138 side chain leaning towards the main chain of the neighbouring β3 strand and forming hydrogen bonds with the main chain of Gly97 and Gly98 residues (*i.e.*, immobilizing the ‘lid’ loop) (**ExtendedData Fig. 4D**). This observation suggested a possible role for alkali metal ions in the stabilization of a pre-catalytic configuration of the active site.

Next, we examined the impact of the most prevalent cations in the body on the GlcNAc substrate. Quantum mechanical calculations of GlcNAc–cation systems suggest that sodium, potassium, calcium, and magnesium ions have high affinities for interacting with the amide group in agreement with previous experimental studies (**ExtendedData Fig. 3E**) [42]. The alkali metal ions (**M^+^**) polarize the entire GlcNAc molecule resulting in changes to the electrostatic potential charges distribution and the C2–N2 torsional potential energy profile (**ExtendedData Fig. 5**). In the twisted-boat conformation of GlcNAc, the 2-acetamido group can only adopt a *trans* conformation with the global minimum energy of C2–N2 torsion at 193 degrees. Upon polarization by M^+^, the 2-acetamido group adopts the *cis* orientation in the twisted-boat conformation, with the global minimum of the C2–N2 torsion at the angle optimal for the substrate-assisted GlcNAc hydrolysis. This result indicates involvement of alkali metal ions in achieving an optimal pre-catalytic conformation of the substrate.

To further investigate the contribution of M*^+^* to catalysis, we performed a series of steered QM/MM MD simulations of the CHIT1–(GlcNAc)_3_ model system (**Fig. 5**). As in classical MD simulations, the M^+^ immediately appeared between side chains of residues Asp138, Glu140, and Met210. From these simulations, we deduced that metal ions, in addition to presenting a particular conformation of GlcNAc-based substrates for catalysis, can be responsible for moderating the *pK_a_* value to facilitate the protonation cycling process (**ExtendedData Fig. 6**). The hydrophobic characters of the neighbouring Ala183, Met210, and Trp358 residues increase *pK_a_* of Asp138 to ∼10, leading to its protonation at the operational optimum pH for hCHIT1 of ∼7.5. Glu140, which protonates the glycosidic bond at the anhydrous stage of the catalysis, has *pK_a_* ∼6 before substrate binding, and stays deprotonated but stabilized by the hydrogen bond interaction with the adjacent tyrosine (**Tyr141**). To enable the substrate binding and initiate the first stage of hydrolysis, protonated Asp138 must transfer hydrogen to deprotonated Glu140. To enable proton transfer, the *pK_a_* value of Asp138 needs to drop significantly. The QM/MM simulations showed this happening after Met210 hides its methyl from the active site by rotation guided by the movement of M^+^ (*i.e.*, in line with the classical MD simulations and analysis of crystal structures). When the methyl moves away from Asp138, its *pK_a_* value drops enough to release the proton and transfer it to Glu140. It was previously reported that prior to the sugar binding, the *pK_a_* value of Glu140 increased from 6.2 to 8.7 [43]. M^+^, which acts as electrophile, by migrating along the main chain of the β4 strand closer to Glu140 can locally increase its p*Ka*. The key role of Asp138 in the hydrolysis is to share hydrogen with the nitrogen of the 2-acetamido group when the twisted boat-shaped-1 sugar accommodates the substrate in the active site. This 2-acetamido group forms with Asp138 and Tyr212 strong hydrogen bonds, which tighten up as the intermediate state approaches. The side chain of Tyr212 pushes the bound carbonyl oxygen of the 2-acetamido group towards the anomeric carbon during the movement of β6 strand (triggered by dynamic movement of the adjacent ‘pulley’ loop, **L6** in **Fig. 4A**), while the nitrogen of the 2-acetamido group bound to the side chain of Asp138, is pushed in the opposite direction. The conjugated transfer of protons through the 2-acetamido group (*i.e.*, between Tyr212 and the carbonyl oxygen and then between the nitrogen and Asp138), leading to the formation of the transition state, is promoted by the sulphur orbitals from the ∼3.7 Å distant Met210, which can be further enhanced by the ionic forces of the alkali metal ions [44]. The polarized carbonyl oxygen acts as a nucleophile that attacks the C1 anomeric centre and by creating oxazolinium intermediate, breaks the glycosidic bond. The simultaneous proton transfer from Glu140 to the oxygen of the glycosylic bond promotes departure of the +1 sugar. The leaving group releases the tryptophane ‘lid’, letting water enter the active site and start the aqueous stage of the hydrolysis. Deprotonated Glu140 at this stage acts as a general base facilitating the nucleophilic attack of a water molecule on the anomeric carbon. The broken oxazolinium/oxazoline ring reopens into the 2-acetamido group and the-1 sugar pops out of the active site. The dual role of Glu140 as a general acid/base during catalytic reaction places specific demands upon its protonation state, and hence its p*K_a_* values at each stage of catalysis. The presented QM/MM results suggest a possible contribution of M^+^ in regulating the conjugated protonation cycling of Asp138 and Glu140 residues during the catalytic reaction. However, the *pK_a_* shift due to alkali metal ions polarization is expected to be rather small and transient [45].

## Discussion

The multiscale simulations of this investigation revealed that all conserved motifs of hCHIT1 exhibit distinct dynamic behaviours and mechanical properties, which collectively facilitate catalysis. To better understand the enzymatic apparatus of hCHIT1, we simultaneously performed computational studies on its three closely related homologues (*i.e.*, mCHIT1, hAMCase and mAMCase, presented in **ExtendedData Fig. 1**), which exhibit different enzymatic efficiencies. The intrinsically flexible GGW loop within the first motif, is responsible for diffusion driven hinge-like movements of the Trp99 ‘lid’ that perpetually opens/closes the active site (**L3** ‘lid’ loop in **Fig. 4A**). Motif I is therefore directly involved in the activation of hCHIT1, allowing substrate binding and accommodation in the active site, protecting the unstable intermediate state from bulk water and promoting dissociation of the product from the binding cleft after each catalytic cycle. The importance of the tryptophane ‘lid’ structural feature is reflected by its high conservation in chitinases across diverse organisms. The MD simulations performed at pH 7.5 showed that the ‘lid’ remains closed for much longer for mAMCase than for hCHIT1, in agreement with the previously reported relative catalytic activity of both enzymes under the same conditions, which was lower for mAMCase [46]. The reason is a different residue X145 within the loop emerging from the β4 strand (**L4** ‘latch’ loop in **Fig. 4A**), which additionally locks the tryptophan ‘lid’ upon closing the active site. In hCHIT1 this non-covalent ‘latch’ residue is a neutral Gln145, whereas in mAMCase it is a positively charged Arg145. The energy of tryptophan-asparagine NH-π interaction is weak enough to be affected by solvation effects, thus can be easily overcome in physiological conditions by diffusion forces. At pH 7.5 tryptophan and arginine are bound together by a much stronger (*i.e.*, estimated as ∼90 kJ/mol) cation-π interaction, hence much more stable in physiological conditions [47]. We observed concerted hinge-like movements of the ‘lid’ and ‘latch’ loops that emerge from the adjacent β3 and β4 strands, opening the active site prior to hCHIT1 activation [48] (**Fig. 3A**). The dynamics of motifs I and II are therefore closely linked and should be synchronized for efficient catalysis. Concerted opening and closing of the ‘lid’ and ‘latch’ loops allow the β4 strand to slide smoothly outside/inside the active site cleft, present the catalytic residues for substrate binding, and isolate the active site from water during the anhydrous phase of catalysis, all at an efficient rate. Only a point mutation of residue X137 within the hydrophobic core, which distinguishes hAMCase from mAMCase enzymes, significantly alters their relative catalytic activity [49]. The lower enzymatic activity of mAMCase compared to hAMCase, especially at high substrate concentrations, is due to the L137F mutation, which distorts the main chain of the β4 strand and disrupts its sliding (**Fig. 3C**). Another position where a point mutation in, otherwise conserved, motif II occurs is residue X141. In hCHIT1, Tyr141 forms a bond with deprotonated Glu140 and thus completes the active site closure (**ExtendedData Fig. 6**). The significant decrease in enzymatic activity of mCHIT1 compared to hCHIT1, observed at high substrate concentrations, results from the combined effect of the Y141F and S144G mutations within the ‘latch’ loop. Phenylalanine prevents the formation of a hydrogen bond with Glu140, while glycine increases the flexibility of the ‘latch’ loop. The resulting structural changes decrease the effectiveness of the active site closure and increase the dynamics of the ‘latch’ loop. Consequently, the opening/closing motions of the ‘latch’ loop of mCHIT1 are much faster than the catalytic turnover rate. An excess of catalytically required motions is a known cause of substrate inhibition in enzymes, which are also observed in the case of mCHIT1 assays [49]. In contrast, the catalytic machinery of hCHIT1 maintains high catalytic activity even in the presence of excess substrate [39]. In conclusion, the conserved motif II is responsible for the presentation of the active site for substrate binding and, together with motif I, is directly involved in the mechanical activation of the enzyme.

The third conserved motif, within the β6 strand, serves a dual purpose in GlcNAc catalysis, playing both charge-transferring and mechanical roles. Residue Met210 plays a catalytic role through transient stabilization of a positively charged oxazolinium intermediate. Moreover, electrostatic interactions and dispersion forces between the lone electron pair on the sulphur on the methionine and the nitrogen on the 2-acetamido group facilitate the proton transfer reciprocally between the nitrogen and Asp138 during the formation and breakdown of the oxazoline ring. The side chains of Tyr212 and Asp213 together maintain the-1 sugar in the correct position for substrate-assisted catalysis by forming tight hydrogen bonds with the carbonyl oxygen and the alcohol hydroxyl group, respectively. The firm attachment of the-1 sugar allows Tyr212 side chain to effectively push the carbonyl oxygen towards the anomeric carbon and force the 2-acetamido group to form the intermediate oxazolinium/oxazoline ring (**Fig. 5**). The synchronized dynamics of the loops originating from the first three conserved motifs (L3, L4 and L6 in **Fig. 4A**) can be seen as a comprehensive ‘flywheel’ mechanism in the hCHIT1 apparatus.

Motif IV, within the β8 strand, is the longest conserved region in hCHIT1 and serves multiple purposes in GlcNAc catalysis. Interestingly, this entire motif is deleted in the most common mutation in the *CHIT1* gene, which causes irreversible inactivation of the enzyme. The QM/MM simulations showed that Trp358 side chain stabilizes both positively charged oxazolinium and neutral oxazoline intermediates during the enzymatic reaction through cation-π and π-π interactions, respectively. The high mobility in MD simulations, as well as the different conformations of the Trp358 side chain observed between the open and closed, unliganded and liganded hCHIT1 crystal structures, suggest a key role for this residue in accommodating the-1 sugar inside the active site, as well as in replacing the glycan chain along the binding cleft during processed hydrolysis and transglycosylation (**Fig. 1C**). The conformational changes of Trp358 are synchronized with the dynamics of the following three conserved aspartic acids (*i.e.*, Asp361, Asp363 and Asp364). These residues, by interacting with other hCHIT1 subunits, mediate the assembly of the catalytic and CBM domains into the immunoglobin-like heterodimer, and induce structural changes on the proline-rich linker that trigger the reciprocal hinge-like movements of the catalytic machinery, required for substrate polysaccharide processing (**Fig. 4C**). The salt-bridging between Asp361 and Arg35 promotes correct orientation and optimal flexibility of the α1-2 subunit for CBM docking (**Fig. 4B**). Residue Asp363 forms hydrogen bonds with the main chain of residues Leu377 and Ile378 located on the loop adjacent to the α8 helix, while Asp364 forms hydrogen bonds with the main chain of residues Gly367 and Ser369 hosted by the same loop (**L8** in **Fig. 1A** and **4C**). These interactions strengthen the anchor point of the proline-rich linker and are therefore directly involved in its ‘elbow-bending’ dynamics. This structural feature allows CBM to efficiently scan the environment, detect GlcNAc-containing substrates for modification more quickly, and navigate the enzyme to different regions of polymeric substrates and induce the hinge-like movements of the heterodimer during processive catalysis (**Fig. 2D**). In summary, motif IV is responsible for the correct positioning of the substrate within the binding cleft for the hydrolysis/transglycosylation process, the stabilization of intermediate states and the mechanical communication between the structural subunits of hCHIT1. The latter involves the assembly of the catalytic and CBM domains into a functional immunoglobin-like heterodimer, capable of processive catalysis and interaction with biological partners [14].

The multiscale simulations and the analysis of the difference density maps in crystal structures of unliganded hCHIT1 reveal a preferential region for alkali metal ions between residues Asp138, Glu140, Met210 and Trp358. These residues represent the conserved motifs surrounding the positively charged oxazolinium intermediate formed during the catalytic reaction. The presence of M^+^ facilitates the reorientation of the side chains of residues Asp138 and Met210, ensuring their optimal positioning for the binding of the 2-acetamido group of the-1 sugar. In this way, the entropic penalty of ordering the active site residues to create hCHIT1–glycan complex is paid by the previously bound cation. Moreover, the stabilized hCHIT1–M^+^ complex facilitates substrate binding for catalysis by moving the C2– N2 torsional potential energy global minimum to the angle optimal for the substrate-assisted GlcNAc hydrolysis. The addition of sodium/potassium ions to the biological assay resulted in higher Michaelis constant values without a higher catalytic activity proportional to the cation concentrations (**ExtendedData Fig. 7**), suggesting a cofactor-like involvement of alkali metal ions in the catalytic mechanism of hCHIT1 [50]. The addition of calcium and magnesium ions had inhibitory effects on hCHIT1. This can be explained by the high affinity of divalent metal ions (**M^2+^**) to the carboxyl groups of the catalytic triad and by the lower ligand exchange rates of M^2+^ compared to M^+^. Previous studies have suggested that the conserved tryptophan (Trp139) between the catalytic aspartate and glutamate is responsible for the ionization of catalytic residues and the formation of the transition state. It is possible that alkali metal ions occupying the nearby region contribute to this ionization effect. However, the high ionic mobility, low dehydration energy and small charge/size ratio of potassium/sodium ions contribute to their rapid association/dissociation kinetics and transient polarization. Consequently, the effects of M^+^ on the catalytic activity of hCHIT1 are not discernible in experimental assays.

The OATD-01 inhibitor acts as a three-branched lock that disrupts the dynamics of the sophisticated hCHIT1 catalytic machinery (**ExtendedData Fig. 8**). The first branch, the aminotriazole warhead, utilizes M^+^ within the electrostatic substrate attraction mechanism of hCHIT1 to efficiently navigate and anchor the inhibitor inside the polarized catalytic site (**ExtendedData Fig. 4F**). The second branch is methyl, which forcibly locks the ‘lid’ loop in a fixed position through CH-π interactions with Trp99 side chain. The third branch is 4- chlorobenzene, linked by a short aliphatic chain. The chlorobenzene moiety participates in hydrophobic and halogen interactions that link and stabilise the β8 strand and βB1/βB2 subunits. The short aliphatic chain strengthens the CH-π interactions with Trp99 side chain, thereby aiding in the immobilization of the ‘lid’ loop (**ExtendedData Fig. 8C**). The significant inhibitory potency of OATD-01 results from the interplay of simultaneous strong interactions between the three branches of the inhibitor and key active site subsections belonging to all four conserved hCHIT1 motifs, which are cooperatively involved in the dynamic catalytic apparatus. As a result, the inhibitor has two distinct and complementary modes of action (**ExtendedData Fig. 2F**). First, it deactivates the catalytic machinery by blocking the active site. Second, OATD-01 disrupts the communication between the hCHIT1 subunits by inducing structural perturbations in the regions responsible for the assembly of the catalytic and carbohydrate-binding domains into the immunoglobulin-like heterodimer (**Fig. 6**). The latter effect may have profound consequences such as impaired cooperativity with biological partners and thus dysregulated initiation of immune responses triggered by the immunoglobulin-like hCHIT1.

**Figure 6.**
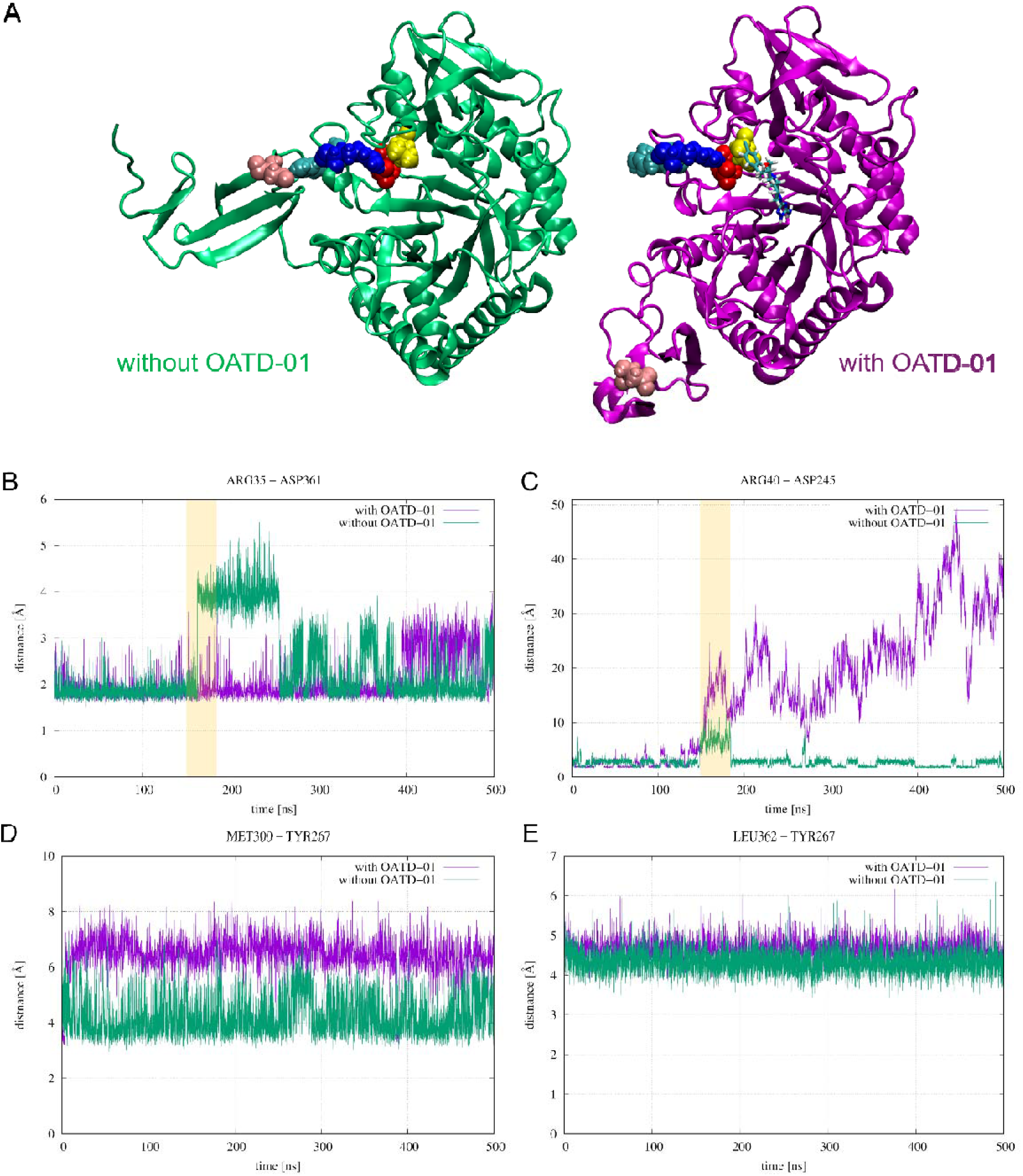
The effect of OATD-01 on the hCHIT1 heterodimer assembly and stability. **A**) Structures without and with OATD-01 ligand after 500 nanosecond MD simulations, coloured in green and magenta, respectively. The enzyme is represented as a cartoon, the OATD-01 inhibitor as licorice, and the key amino acids are in vdW representations. Yellow, red, blue, cyan, and pink colours correspond to Leu362, Asp361, Arg35, Arg40 and Asp425 residues, respectively. These amino acids are involved in the inter-subunit mechanical signal transduction between the catalytic domain (β8) and CBM through α1-2 subunit. The detailed view of the reciprocal orientation of these resides and OATD-01 is shown in Fig. 8C/D) The evolution of distances between the residues involved in the inter-subunit mechanical signalling: two salt bridges (Arg35–Asp361 and Arg40–Asp425) and residues involved in the hydrophobic sub-pocket accommodating the 4-chlorobenzene (branch III) of OATD-01 (Met300– Tyr267 and Leu362–Tyr267). OATD-01 exerts a strong interaction with residues Tyr267, Met300 and Leu362, thereby imposing rigidity on β8 **B**). The hydrophobic sub-pocket expands **D**/**E**) and tightens the hydrogen bonds between Asp361 and Arg35, which connect β8 with the α1-2 subunit. Consequently, the α1-2 structural feature becomes more rigid and induces stress on the adjacent salt bridge between Arg40 and Asp425, which connects the α1-2 and CBM subunits **C**). The Arg40**–**Asp425 salt bridge breaks, resulting in CBM dissociation. Increased rigidity of the α1-2 subunit, which contains the residue Arg40, prevents CBM from redocking. This change in the dynamic behaviour of Arg40 can be observed when comparing the α1-2 regions within the crystal structures from the 4WKA and 6ZE8 PDB entries (*i.e.*, without and with OATD-01 inhibitor).

## Supporting information

hCHIT1_OATD01_MD_500ns

hCHIT1_APO_MD_1000ns

## Methods

### Modelling dynamics of hCHIT1

Classical MD simulations of the catalytic domain were used for identification of the key dynamic features within the enzymatic apparatus of hCHIT1. Analysis of trajectories from MD simulations of hCHIT1 also highlighted the role of ions in this system. To better understand the hCHIT1 results, the same MD simulations were performed for each of the four homologues with known relative activities (ExtendedData Fig. 1). The simulated structures were obtained from RCSB Protein Data Bank [51] or predicted by AlphaFold [52]. Next, they were visually inspected and structurally analysed using Coot software [53]. The Amber ff99sb-ILDN force field [54] implemented in GROMACS molecular dynamics package [55] was used for all systems. The TIP4P [56] water was used to solvate the system and 0.150 M NaCl was added. After 150,000 steps of minimisation using steepest descent [57] and conjugated gradient [58] algorithms, 200 ps were run in the NVT ensemble with heating and position restraints on the protein and a further 200 ps of unrestrained NVT equilibration. After removal of the restraints and equilibration of 500,000 steps in the NPT ensemble, production runs of 500 ns were performed with a 1.0 fs time step. The simulation temperature was controlled at 310□K by the V-rescale thermostat [59], and the pressure of the system was kept constant at 1 atmosphere using the C-rescale barostat [60], in periodic boundary conditions. Both temperature and pressure were updated every 100 integration steps. Electrostatic interactions were described using the particle mesh Ewald (PME) algorithm [61]. Non-bonding interactions were calculated within the cut-off of 12 Å. Next, the simulated systems were converted from GROMACS to Tinker format using an in-house script and the simulations were continued with AMOEBA force field [62] implemented in Tinker 9 software [63]. The systems were again minimized with and gradually heated to 310 K under NVT conditions. After NVT and NpT equilibration runs (1 ns long with a 1 ps time step) production runs of 200 ns were performed. A Nosé–Hoover [64] thermostat and barostat were used. The trajectories from all MD simulations were visually examined and analysed using VMD 1.9.4a57 software [65]. To identify functionally relevant collective motions of hCHIT1, principal components analysis (PCA) of its MD trajectories was done in GROMACS.

### Model building of hCHIT1 with substrate

The experimentally resolved structures of catalytically active hCHIT1 contain only-2 and-1 units of the GlcNAc product. Therefore, the missing +1 sugar unit had to be modelled. The human YKL-39 pseudo-chitinase shows 52% sequence similarity to CHIT1 with root mean square deviation (RMSD) of 1.1 Å on the Cα atomic coordinates after optimal rigid body alignment. Due to lack of catalytic activity YKL-39 structures contain GlcNAc oligomers that could be used to model the missing +1 GlcNAc unit in the substrate. The structure of hCHIT1 (PDB ID: 4WKH) with the (GlcNAc)_4_ product was superimposed on the YKL-39 structure (PDB ID: 4P8W) with the (GlcNAc)_4_ with the structural alignment focused on the active site residues and the two same sugar units. The +1 unit from the YKL-39 structure was added to the (GlcNAc)_2_ product to build the model of the (GlcNAc)_3_ substrate. The resulting structure of hCHIT1 with substrate was subjected to the energy minimization with strong torsional restraints imposed on the dihedral angles of the-1 GlcNAc unit to maintain its twisted boat conformation. The structural optimisation of hCHIT1 with substrate was performed by successive runs of two energy minimisation algorithms: 30,000 steepest descent steps and another 30,000 conjugated gradient steps, using Amber03 [66] force field and TIP4P explicit water model implemented in GROMACS software. The optimization of the system was followed by a short (5 ns) relaxation run at 310 K, in which the torsional restraints retained in the-1 sugar.

### QM/MM calculations of hCHIT1 with substrate

To observe the substrate-assisted GlcNAc hydrolysis the system of hCHIT1 with the (GlcNAc)_3_ was divided into regions treated at the quantum (QM) and molecular mechanics (MM) levels of theory. The key active site residues: Asp138, Glu140, Met210 and Tyr212, the ion shared by Asp138 and Glu140, and the (GlcNAc)_3_ substrate was described at the QM DFT using PBE [67] functional and DZVP-MOLOPT basis set [68] level while the rest of the molecular system was treated with AMBER03 force field according to the default QM/MM settings of GROMACS with CP2K software interface [69]. To test the possible effect of metal ions on the catalytic process of hCHIT1, each of the sodium, potassium, calcium and magnesium cations was randomly inserted into the system a dozen times and then the energy of the system was minimized. After minimization, potassium ion tends to migrate to the region of excess density observed in the unliganded hCHIT1 structure, consistent with sodium/potassium ions being incorrectly modelled as water (ExtendedData Fig. 4C). This ion was included in the system and described at the quantum level during QM/MM simulations, which were carried out using PBE0 functional [70] and TZVP-MOLOPT basis set [68], truncated Coulomb potential [71], long range Coulomb correction [72], Grimme D3 dispersion correction with Becke-Johnson damping [73] and auxiliary density matrix method [74]. After fast (10 ps) heating and short (10 ps) equilibration under an NVT ensemble, the system was subjected to several short (∼40 ps) runs of QM/MM simulations.

The PCA analysis indicated that the distinct mechanical motions of the β6 strand observed in the classical MD simulations of hCHIT1 facilitate the substrate-assisted GlcNAc hydrolysis by the Tyr212 side chains pushing of the bound carbonyl oxygen of the 2-acetamido group towards the anomeric carbon, an adaptive bias was implemented in the QM/MM MD runs.

First, the distance between the carbonyl oxygen and the anomeric carbon from 3.2 (*i.e.*, in the *cis* conformation of the 2-acetamido group) to 1.43 Å (*i.e.*, the C–O covalent bond length) was sampled. Second, of the distance between the anomeric carbon and the oxygen of the glycosidic bond was changed from the default value of 1.3 to 4.3 Å. A decrease in the distance between the carbonyl oxygen and the anomeric carbon always corresponded to a spontaneous increase of the distance between the anomeric carbon and the oxygen of the glycosidic bond (and *vice versa*), which eventually led to the formation of the oxazolinium/oxazoline ring from the 2-acetamido group of the-1 sugar and the rupture of the glycosidic bond and dissociation of the +1 product. When exploring the free energy landscape of the substrate-assisted GlcNAc hydrolysis using accelerated weight histogram (AWH) method [75], it was impossible to obtain the free energy profile of the simulated process. This suggest that the activation energy required for hydrolysis is lower compared to the activation energy needed for the reverse reaction (∼60 kJ/mol), suggesting that the products of the catalytic reactions cannot easily revert to the original reactants.

The analysis of the elemental composition of the coordination sphere and the distances to the nearest atoms from the metal ions was performed using CheckMyMetal server [76].

### Model building of the full-length hCHIT1

The initial model of the full-length hCHIT1 was based on the 5HBF PDB entry with crystal structure of the unliganded hCHIT1 catalytic domain co-crystalized with CBM (without the proline-rich linker that connects the heterodimer). That structure had two mutual orientations of the CBM, and catalytic domains resolved. In the chain A, CBM was resolved across from the active site of hCHIT1. In chain B, CBM was found next to the α1-2 subunit, which is structurally closer to the mutual orientation of both domains observed in the immunoglobulin-like Chitinase A from Serratia marcescens (SmChiA, PDB ID: 5Z7M), that structurally belongs with hCHIT1 to the same type of ‘bacterial-type’ chitinases. The latter orientation was also proposed by the AlphaFold prediction therefore it became the initial conformation of the full-length hCHIT1. That initial structure would not allow for the processive catalysis therefore we further sampled the structure of the mutual arrangements within heterodimer in two ways. The first approach involved sampling hundreds of poses of CBM docked to the catalytic domain (without considering the linker) using the protein-protein docking method implemented in MOE (Molecular Operating Environment) molecular modelling programme [77]. These heterodimers were generated with and without the routine that addresses the hydrophobic complementarity of the complexed proteins. The 50 top scoring structures CBM–catalytic domain complexes were selected for visual inspection.

Their potential to perform processive hydrolysis of GlcNAc polymers from the reducing end was evaluated with reference to the sugar binding mode to CBM experimentally determined in the Ref. 26. This ability was further verified by molecular docking simulations of the (GlcNAc)_16_ oligomer to the most promising CBM–catalytic domain heterodimers using default parameters of the Extra Precision docking algorithm implemented in Flare molecular modelling programme [78]. The structure of the full-length hCHIT1 model obtained by this approach had a rich network of non-covalent interactions responsible for the tight binding of CBM to the catalytic domain in the immunoglobulin-like isoform of hCHIT1. The missing linker between the catalytic domain and CBM was manually modelled using the protein editing tools implemented in Flare and the resulting full-length hCHIT1 model was prepared for further simulations (e.g. addition of hydrogens, protonation of histidines, etc.) using Flare’s ProteinPrep protocol. ProteinPrep module was also used to calculate the *pKa* values of the polar amino acids at different stages of the enzyme. The system after a short (200 ns) equilibration under an NPT ensemble was subjected to 500 ns of MD simulation with default settings implemented in Flare (e.g. AMBER ff99SB force field, explicit TIP3P water solvent [79], AM1-BCC charges [80], PME for treating long-range electrostatics within the cut-off of 12 Å, etc.) except for the temperature (310 K), concentration of NaCl (0.150 M), the time step (1 fs) and hydrogen mass repartitioning option disabled. This simulation allowed the mutual orientation of the catalytic domain, CBM and the flexible linker that connects them into a heterodimer to be readjusted until the final stable model of the immunoglobulin-like hCHIT1 isoform was reached. The second approach took advantage of the flexibility of the proline-rich linker compared to SmChiA, in which the carbohydrate-binding module is rigidly attached to the catalytic domain. In this case, the initial structure of the full-length hCHIT1 was the starting point of dozens of 100 ns MD simulations performed at the room temperature until the most stable model of the immunoglobulin-like hCHIT1 was obtained with a mutual orientation of the catalytic domain and CBM optimal to perform processive hydrolysis of GlcNAc polymers from the reducing end. After less than 200 ns MD simulation, the final model obtained by the first approach (i.e. protein-protein docking of the catalytic domain and CBM with manually modelled linker) changed to the most stable structure simulated by the second approach (i.e. sampling conformational states of the full-length hCHIT1 structure predicted by AlphaFold), what increased our confidence in the proposed full-length hCHIT1 model. The simulation data analysis and visualization were performed using Flare. Despite differences in the simulation setup, the results obtained from MD calculations performed by Flare and GROMACS matched.

The structure of the full-length hCHIT1 with the (GlcNAc)_3_ substrate was superimposed on the structure with OATD-01 (PDB ID: 6ZE8) with the structural alignment focused on the active site residues. The substrate was replaced by the inhibitor and the hCHIT1–OATD-01 complex, after optimization, underwent MD simulations following the same protocol employed for the unliganded hCHIT1.

The calculations of the dipole moment of hCHIT1 were done using Protein Dipole Moments Server [81].

### DFT calculations

All Density Functional Theory (DFT) calculations in this work were performed using Gaussian 16 [82] software and visualised GaussView program [83]. The structure of the-1 GlcNAc unit in its twisted-boat conformation was extracted from a crystal structure of hCHIT1 with the (GlcNAc)_2_ product (PDB ID: 4WKH) and optimized using B3LYP functional with the dispersion correction added explicitly by Grimme’s method (third order) with Becke–Johnson damping [84] and 6-311++G(d,p) basis set [85]. The obtained global minimum energy trans conformation of the 2-acetamido group had the was the starting point for the potential energy scan of the C2–N2 torsion. The calculations were performed for GlcNAc molecule alone and in the presence of a metal cation (sodium, potassium, calcium or magnesium) in vacuum and in the implicit presence of a solvent using the polarizable continuum model (PCM) [86]. Electrostatic potential charges were calculated according to Merz-Kollman (MK) scheme [87]. The structure of the OATD-01 inhibitor was extracted from a crystal structure of hCHIT1 with the (GlcNAc)_2_ product (PDB ID: 6ZE8) and optimized following the same method employed for the unliganded hCHIT1.

### Enzymatic Assay

hCHIT1 was purified from FreeStyle™ 293-F cells using the previously described method [88]. Prior to the experiments, the enzyme was desalted with Amicon® Ultra 2 ml centrifugal filters (Merck Millipore) to replace the existing buffer with the assay buffer: 20 mM Tris, pH 7.4. The enzymatic assay was conducted in a 96-well plate (Greiner Bio-one, cat# 675076) and involved three sequential steps: (i) 15 µl of hCHIT1 at a concentration of 3x 0.25 nM was added to the wells in assay buffer, (ii) 15 µl of assay buffer containing different salts (NaCl, KCl, MgCl2, or CaCl2) at concentration of 3×10 mM was added to each well and incubated for 15 minutes, (iii) 15 µl of the fluorescent substrate, 4-methylumbelliferyl-β-D-N,N’,N’’-triacetylchitotriose, was introduced at various concentrations, creating a 2-fold serial dilution with 7 concentration points starting from 3x 25 µM. A control with no substrate was also included. After the final addition, the plate was centrifuged at 250 g for 5 minutes. The fluorescence signal was measured with excitation at 355 nm and emission at 460 nm using the kinetic mode of the The SpectraMax® i3x Multi-Mode Microplate Reader. Measurements were taken every 2 minutes at RT. Data analysis was performed using GraphPad Prism version 10.0.0, employing simple linear regression to calculate the initial reaction rate (Vo, represented by the slope of the fluorescence increase over time, [RFU/min]). The reaction rates were plotted against the substrate concentrations to determine Vmax [RFU/min] and Km [µM] values using the Michaelis-Menten model.

## Data availability

Materials described in this manuscript were either purchased commercially (with the manufacturer indicated) or derived internally at Molecure S.A. [88]. Atomic coordinates of the model of the full length hCHIT1 isoform as well as the model of the catalytic domain with (GlcNAc)_3_ substrate were deposited to the Supplementary Information in the PDB format. PDB accession codes for previously reported structures used in this study include 3FXY, 4WKA, 5HBF, 5Z7M, 6SO0 and 7CJ2. There are two movies deposited together with the manuscript. The hCHIT1_APO_MD_1000ns movie presents the dynamic behaviour of the apo hCHIT1 heterodimer during 1,000 ns MD simulation. The hCHIT1_OATD01_MD_500ns movie shows the immediate (within the first 200 ns) destabilising effect of the OATD-01 inhibitor on the otherwise stable hCHIT1 heterodimer. Source experimental and simulation data are provided on request (please, use the email address)

## Acknowledgements

Studies on OATD-01 were supported by three projects: (1) “Preclinical research and clinical trials of a first-in-class development candidate in the therapy of asthma and inflammatory bowel disease” (POIR.01.01.01-00-0168/15), acronym IBD, (2) “Development of a ‘first-in-class’ small molecule drug candidate for treatment of idiopathic pulmonary fibrosis through chitotriosidase inhibition” (POIR.01.01.01-00-0551/15), acronym IPF, both co-financed by European Union through the European Regional Development Fund within the Smart Growth Operational Programme and (3) “Preclinical and clinical development of drug candidate OATD-01, for the treatment of sarcoidosis patients” (MAZOWSZE/0128/19), acronym SARCO, as part of the “Path for Mazovia” competition co-financed by the National Centre for Research and Development from national funds. BAD was supported by the Ministry of Science and Higher Education, Poland (50//DW/2017/01/1). The development of the multiscale computational methodology used in the presented study was supported by OPUS grant no. 2018/31/B/ST4/03809 funded by the National Science Centre. We gratefully acknowledge Poland’s high-performance Infrastructure PLGrid ACC Cyfronet AGH for providing computer facilities and support within computational grant no PLG/2023/016104 and PLG/2023/016400.

## Author information

These authors contributed equally: Dorota Niedzialek, Grzegorz Wieczorek.

## Contributions

D.N. and G.W. conceptualized the studies, performed the computational part of the project and analysed acquired structural data and simulation results. K.D and A.A. designed and conducted enzymatic assays and analysed experimental results. D.N., G.W., M.M., K.D. and A.A. investigated, discussed and visualized the results. A.B., Z.Z. and J.O. reviewed the results. J.O. and Z.Z. supervised and administrated the project. The manuscript was written by D.N. and reviewed by all authors.

**Ethics declarations**

## Extended Data

**ExtendedData Fig1.**
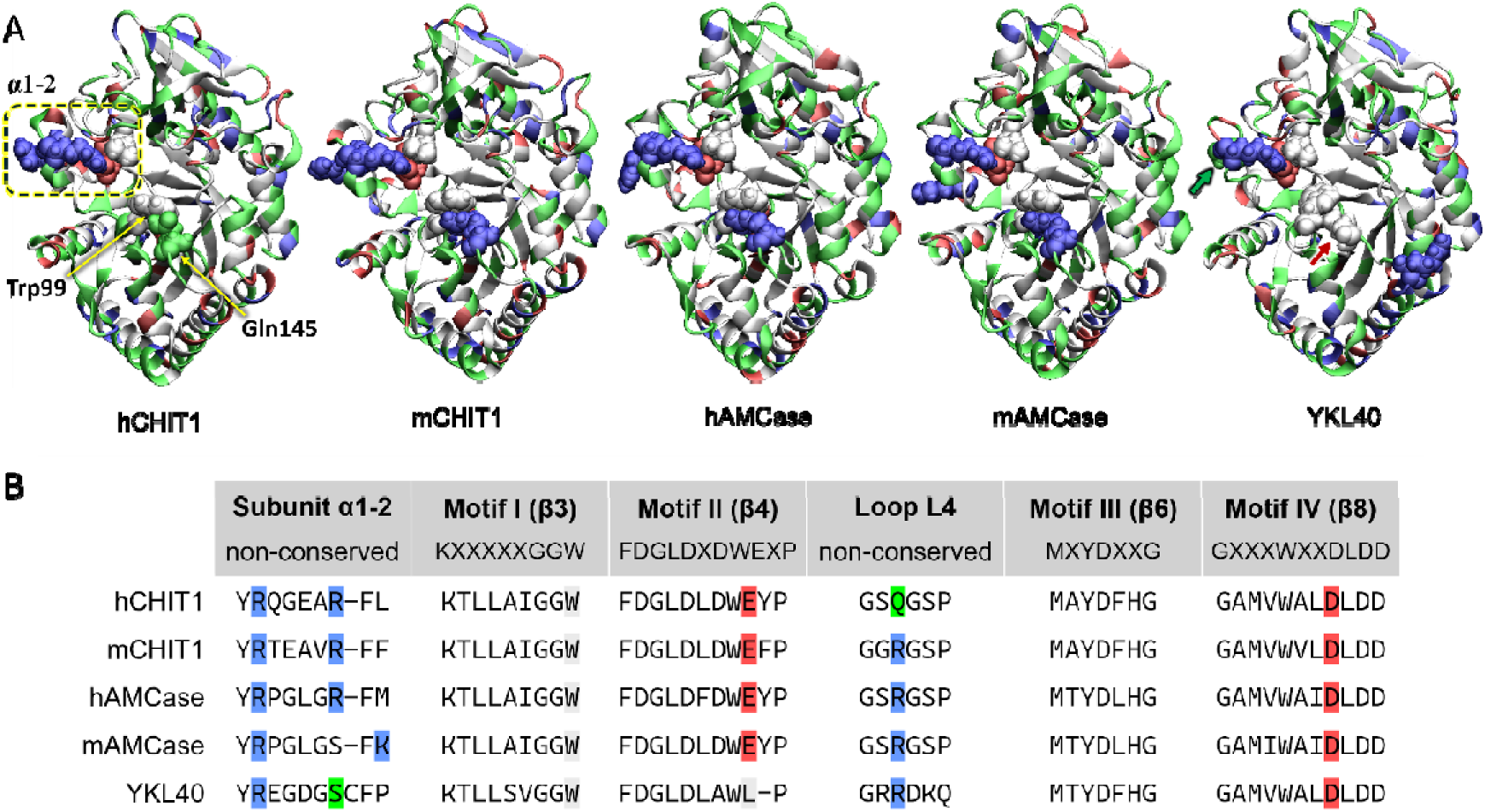
The four conserved motifs of hCHIT1, its closest homologues and catalytically inactive chitinase-like YKL40 protein. **A)** The catalytic domains of hCHIT1 (PDB ID: 4WKA), its mouse homologue (mCHIT1 (AlfaFold ID: AS-Q9D7Q1) as well as of human (hAMCase, PDB ID: 3FXY), mouse (mAMCase) homologues of Acidic Mammalian Chitinase and YKL40 (PDB ID: 7CJ2). All proteins comprise of similar classic TIM-barrel fold (*i.e.*, eight β strands tethered by eight α helices) despite substantial differences in composition (mCHIT1, hAMCase, mAMCase and YKL40 exhibit 78, 52, 50 and 52% similarity to hCHIT1). The colours of the cartoon representation correspond to hydrophobic (white), hydrophilic (green), acidic (red) and basic (blue) character of amino acids. The key residues at the entrance to the active site (Trp99 and X145) and residues of α1-2 subunit with the salt-bridge that connects it to the β8, together with conserved Leu362 are in vdW representation. Note, that only hCHIT1 has uncharged glutamine as residue X145 instead of arginine. The residues in a dotted circle are crucial for the inter-subunit mechanical signal transduction that trigger hCHIT1 heterodimer assembly/dissociation. The arrows indicate the main differences between YKL40 protein and the catalytically active chitinases: the red arrow indicates the E140A mutation within the catalytic triad while the green one emphasizes the lack of basic amino acid within the α1-2 subunit responsible for the heterodimer assembly. Note that only (h/m)CHIT1 and (h/m)AMCase homologs possess a carbohydrate binding domain, and other human chitinase-like proteins, such as YKL-40, consist only of the catalytic domain. **B)** Sequence comparison of the conserved motifs and key structural features in hCHIT1 and the four homologies pictured above. The Xs correspond to sequential differences within the conserved motifs. The key amino acids within each structural motif are marked in colour corresponding to its colour in the vdW representations above.

**ExtendedData Fig. 2.**
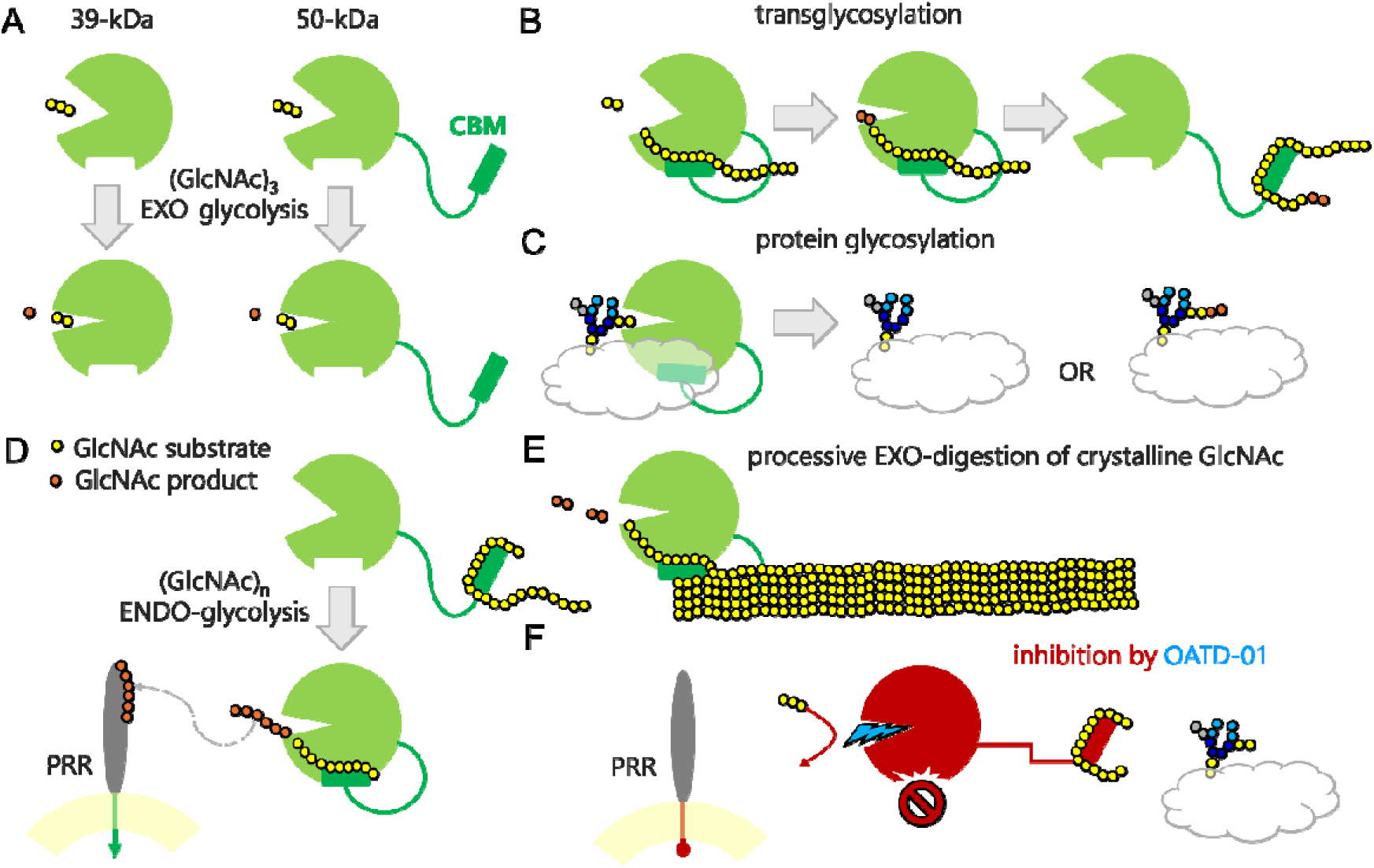
Schematic representation of hCHIT1 in its diverse modes of action. **A**) The full-length, 50-kDa (466 amino acids) isoform consists of the 39-kDa (387 amino acids) catalytic domain located at the N-terminus and the carbohydrate-binding module (CBM) connected by a linker (for details see Fig. 1). The 50-kDa hCHIT1 is initially produced by macrophages and stays dominant in the blood stream. The 39-kDa hCHIT1 consists of the catalytic domain only and is either cleaved post-translationally in the lysosome of macrophages or, less commonly, formed through differential RNA processing. Both isoforms can act as exochitinases that hydrolyse (GlcNAc)3 substrate which is used in biological assays to evaluate enzymatic activity of hCHIT1. **B**) Transglycosidase activity of hCHIT1 may be involved in **C**) pathological N-glycosylation processes. **D**) The full-length hCHIT1 can also act as an endochitinase able to degrade GlcNAc polymers. Product GlcNAc oligomers function as molecular patterns that can be detected by pattern recognition receptors (PRRs) and initiate adaptive immune responses. **E**) Exo-acting processive degradation of the crystalline (GlcNAc)n by hCHIT1 implies its primordial function as an innate defence mechanism against chitin-producing pathogens, such as fungi, nematodes, and insects. **F**) The *first-in-class* OATD-01 inhibitor not only blocks the active site but, most importantly, interrupts communication across hCHIT1 subunits by generating structural distortions in the regions responsible for the assembly of the 39-kDa and CBM into the immunoglobulin-like 50-kDa isoform. This leads to impaired cooperativity with biological partners of hCHIT1, for example in pathological glycosylation processes [14].

**ExtendedData Fig 3.**
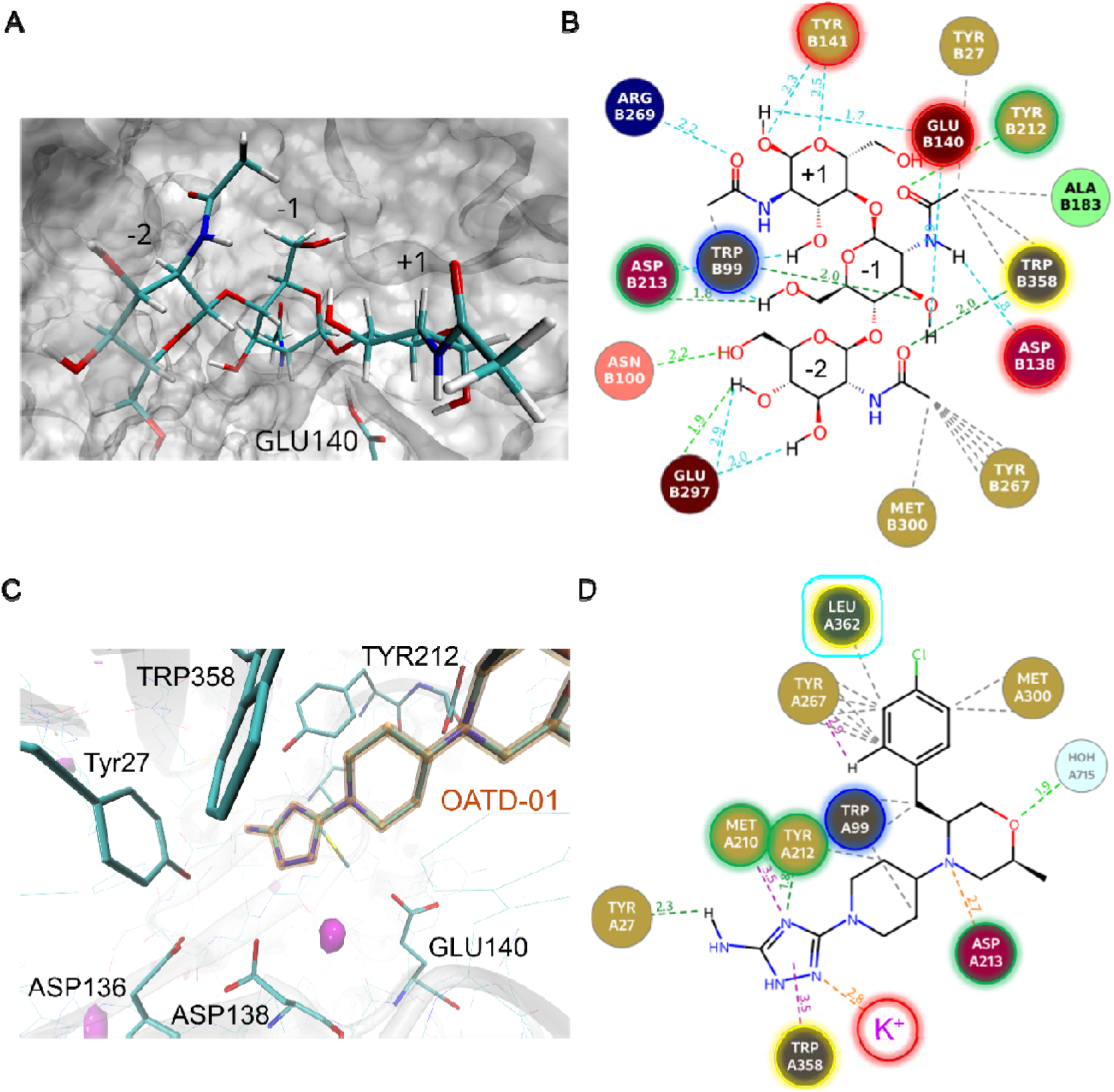
The substrate and OATD-01 inhibitor binding modes within the active site of hCHIT1. **A**) The white surface representation of the active site with the (GlcNAc)3 substrate (represented as licorice) modelled at the QM/MM level of theory. The strong interaction between the active site cleft and GlcNAc substrate forces a distorted conformation of the-1 sugar and brings the susceptible glycosidic bond (between-1 and +1 GlcNAc units) in proximity to the proton donor residue (Glu140). **B**) The two-dimensional interaction map of the binding mode of (GlcNAc)3 with the active site residues of hCHIT1. The colours of the frames around the symbols of the residues correspond to the colours of the conserved motifs assigned in Fig 1A. **C**) The electron density of the |Fo| − |Fc| map from the crystal structure of hCHIT1 complexed with OATD-01 (PDB ID: 6ZE8) contoured at 5σ and shown in magenta surface representation. The analysis of both structures, including the elemental composition of the coordination sphere and the distances to the nearest atoms, identified the excess density as consistent with potassium ion being incorrectly modelled as water. **D**) The two-dimensional interaction map of the binding mode of OATD-01 with the active site residues of hCHIT1. The colours of the frames around the symbols of the residues correspond to the colours of the conserved motifs assigned in Fig. 1A. The interaction with the alkali metal ion (here K+) mediates the interaction with the motif I. The cyan frame indicates the residue Leu362 that plays a key role in the second mode of hCHIT1 inactivation by the OATD-01 inhibitor (*i.e.*, dissociation of the heterodimer).

**ExtendedData Fig. 4.**
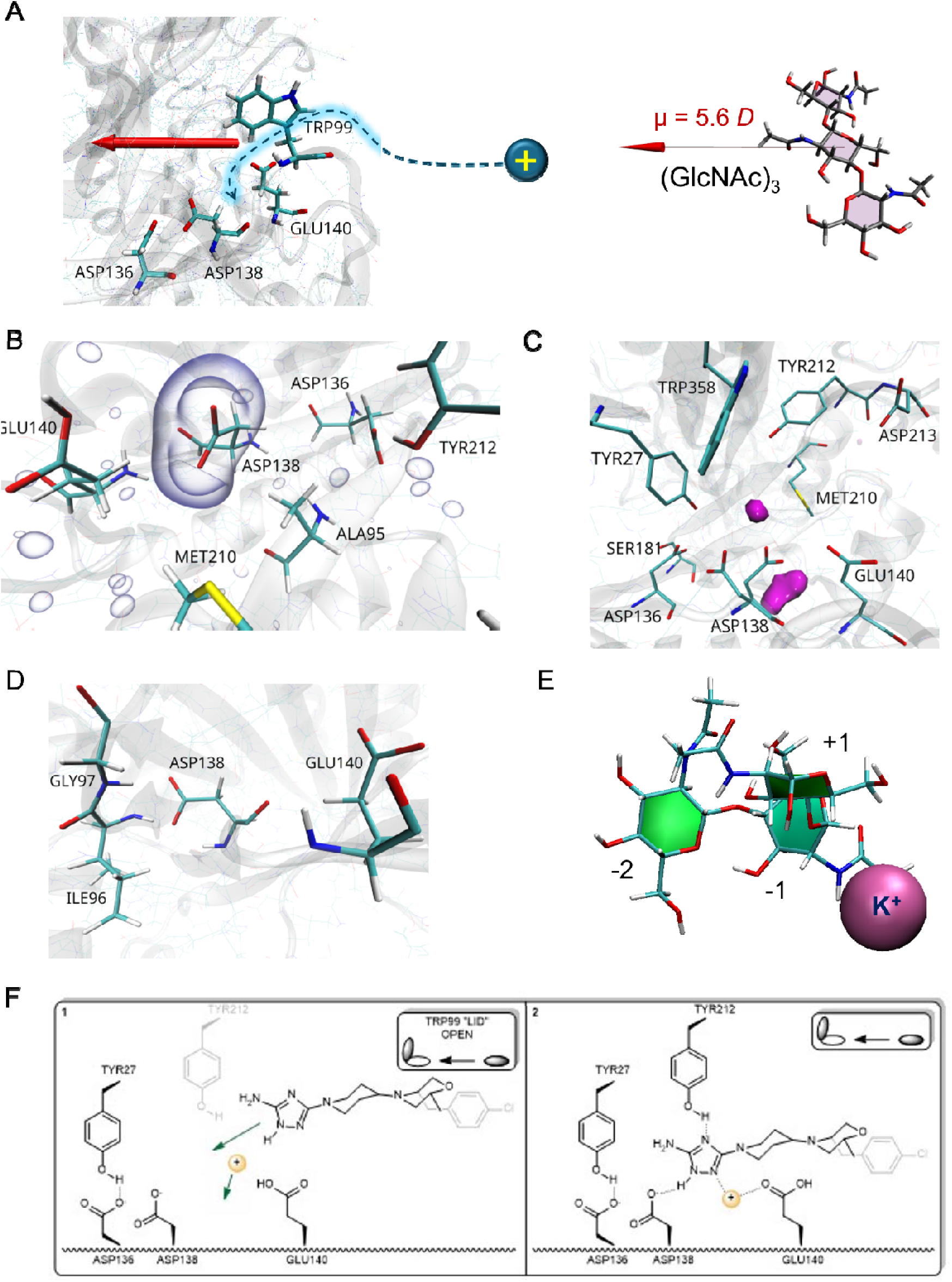
The electrostatics-based substrate/OATD-01 inhibitor attraction mechanism in hCHIT1. **A**) Electrostatic ‘torch of guidance’ effect in hCHIT1 which attracts (GlcNAc)3 substrate and positive ions towards the catalytic site. The overall enzyme charge distribution corresponds to the substantial dipole moment (μ = 242 *D*) indicated by red block arrow, which aligns with the active site along the β4 strand. The optimized structure and the dipole moment for (GlcNAc)3 were calculated at the DFT level of theory. The mutual alignment of the enzyme and substrate dipole moments guides the binding process. The enzyme is presented as a cartoon while the substrate and the key residues are in licorice representations. **B**) The three-dimensional isosurface map of electrostatic potential grid data, extracted from the QM/MM simulations of the unliganded hCHIT1, indicates a substantial negative charge above Asp138 responsible for electrostatic attraction of positive charges in this region. During substrate binding, M+ diffuses towards the space between Asp138 and Glu140. **C**) The electron density of the |Fo| − |Fc| map contoured at 5σ and shown in magenta surface representation extracted from the crystal structure of unliganded hCHIT1 (PDB ID: 5HBF). The excess density consistent with M+ being incorrectly modelled as water. The analysis of the elemental composition of the coordination sphere and the distances to the nearest atoms [76], identified the excess densities as consistent with sodium ions. **D**) A snapshot from an MD simulation with polarizable force-field performed on ion-depleted, unliganded CHIT1 system. In the absence of M+, Asp138 bends towards the main chain of the neighbouring β3 strand and forms a hydrogen bond with the main chain of Gly97 and Gly98 residues that immobilize the GGW loop. **E**) Preferential intermolecular arrangement of (GlcNAc)3 molecule (in green licorice representation) and potassium ion (in pink vdW representation) during the substrate binding process, derived from the QM/MM calculations of the hCHIT1 with (GlcNAc)3 model system. **F**) How OATD-01 exploits the ‘torch of guidance’ effect. Flowchart of the binding process of OATD-01, facilitated by the electrostatic attraction of the negatively charged triazole warhead with an alkali metal ion. The cation helps OATD-01 to adopt an optimal position in relation to the active site (*i.e.*, allowing optimal binding of the inhibitor warhead to Asp138 and Tyr212).

**ExtendedData Fig 5.**
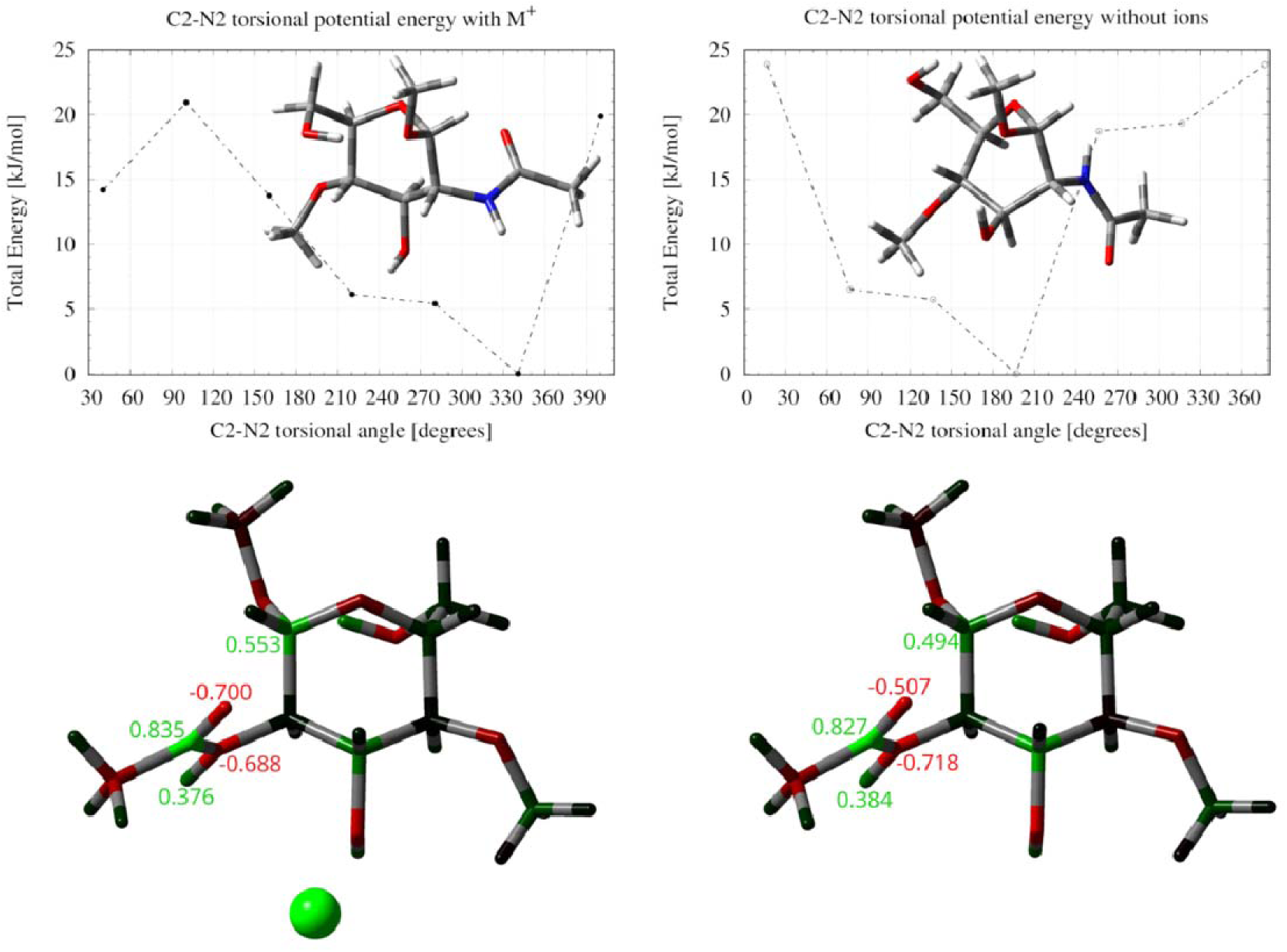
The role of monovalent metal ions in achieving an optimal pre-catalytic *cis* conformation of the 2-acetamido group of the-1 subunit of the GlcNAc substrate. **A)** The torsional potential of the C2–N2 bond with (left) and without (right) alkali metal (M+) ion. Note the difference in the position of global minimum between the two cases with the 2-acetamido group adapting *cis* and *trans* conformation with and without assistance of M+ (sodium) ion, respectively. M+ facilitates the correct orientation of the carbonyl oxygen of the 2-acetamido group for the nucleophilic attack of the carbonyl oxygen on the C1 anomeric centre. **B)** Electrostatic potential charges distribution along the 2-acetamido group and the anomeric carbon with (left) and without (right) M+ ion. Note the increase of the negative (red) and positive (green) charges on the carbonyl oxygen and the anomeric centre, respectively, in the case with M+ ion. The green sphere indicates M+ ion and the charges are in atomic units.

**ExtendedData Fig 6.**
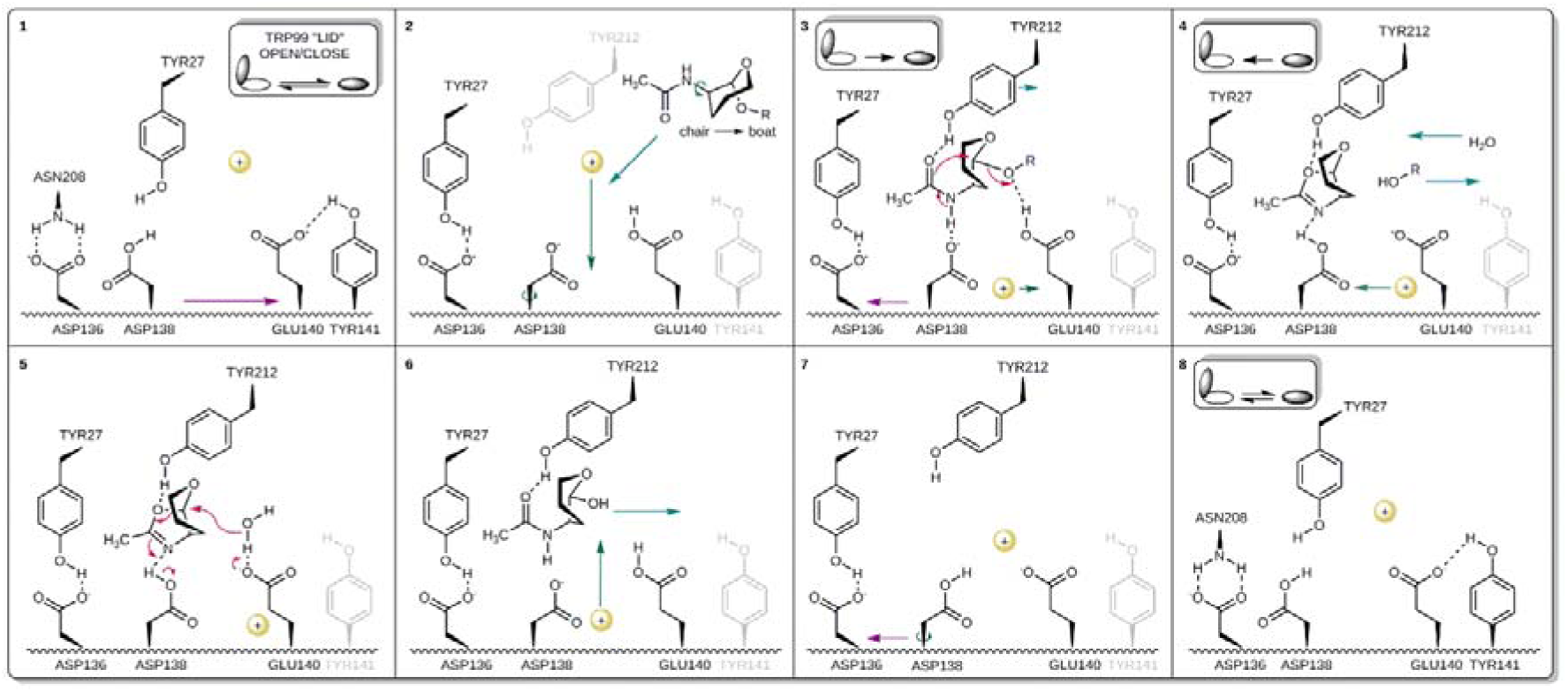
The revised mechanism of hCHIT1 catalysis based on multiscale simulations. For drawing clarity, the-2 sugar is not presented, the-1 sugar is presented without other substituents than the 2-acetamido group, and the +1 sugar is represented as “R”. The electron transfer is indicated by red arrows. The diffusive movements of the cation are indicated by green arrows. **Step 1**: In the inactive form of hCHIT1, Asp136 (β4 strand) interacts with Asn208 (β6 strand) by hydrogen bonds. Tyr27 is situated between Asp136 and Asp138 residues. The protonated Asp138 (*pKa* = ∼10) faces Asp136 while unprotonated Glu140 (*pKa* = 6) forms a hydrogen bond with the adjacent Tyr141. When ‘lid’ (β3 strand) and ‘latch’ (β4 strand) loops cooperatively open, the β4 strand slides towards the entrance of the active site cleft (purple arrow) and exposes Asp138 for binding the substrate; **Step 2**: The β4 strand stops when Asp136 forms hydrogen bond with Tyr27. The enzyme becomes active. The substrate, led by the electrostatic ‘torch of guidance’ effect enters the active site in the default chair conformation. Diffusive movement of M+ facilitate the rotation of Asp138 towards Glu140 as well as otherwise unfavourable, 180° rotation of the 2-acetamido group of the-1 sugar. Upon substrate binding, M+ diffuses towards Glu140 (*pKa* ↑) facilitating proton transfer from Asp138 (*pKa* ↓). **Step3**: The tight binding to the key residues of the active site cleft induces the boat-like conformation of the-1 sugar. The nitrogen and carbonyl oxygen from the 2-acetamido group of the-1 sugar form strong hydrogen bonds with the side chains of Asp138 and Tyr212, respectively. The binding substrate pushes the β4 strand in the direction of its momentum (purple arrow). Consequently, Asp138 pushes the hydrogen shared with the nitrogen from the 2-acetamido group in one direction while the carbonyl oxygen is pushed by Tyr212 in the opposite direction. Tthe general acid (Glu140) donates proton to the glycosidic bond between-1 and +1 sugar units, as the hydrogen bond between Asp138 and nitrogen of the 2-acetamido group tightens up. This process in coordinated by the movement of the M+ from Glu140 (*pKa* ↓) to Asp138 (*pKa* ↑). Pushed by Tyr212, carbonyl oxygen performs a nucleophilic attack on the anomeric carbon, forming the oxazolinium/oxazoline intermediate and in turn breaking the glycosidic bond; **Step 4**: Upon +1 sugar departure, the ‘lid’ gets released and opens the active site, allowing water molecule to enter. **Step 5**: Deprotonated Glu140 becomes a general base that facilitates the nucleophilic attack of the water molecule on the anomeric carbon. **Step 6**: The breaking intermediate springs when oxazolinium/oxazoline ring opens back into 2-acetamido group and consequently-1 sugar detaches from the active site, while M+ promotes the proton transfer between Glu140 and Asp138. **Step 7**: Detachment of the-1 sugar triggers the β4 strand to slide backward; **Step 8**: Glu140 and Asp136 again form hydrogen bonds with Tyr141 and Asn208, respectively. M+ returns above Asp138 facilitating its protonation (*pKa* ↑) and rotation towards Asp136 that closes the catalytic cycle.

**ExtendedData Fig. 7.**
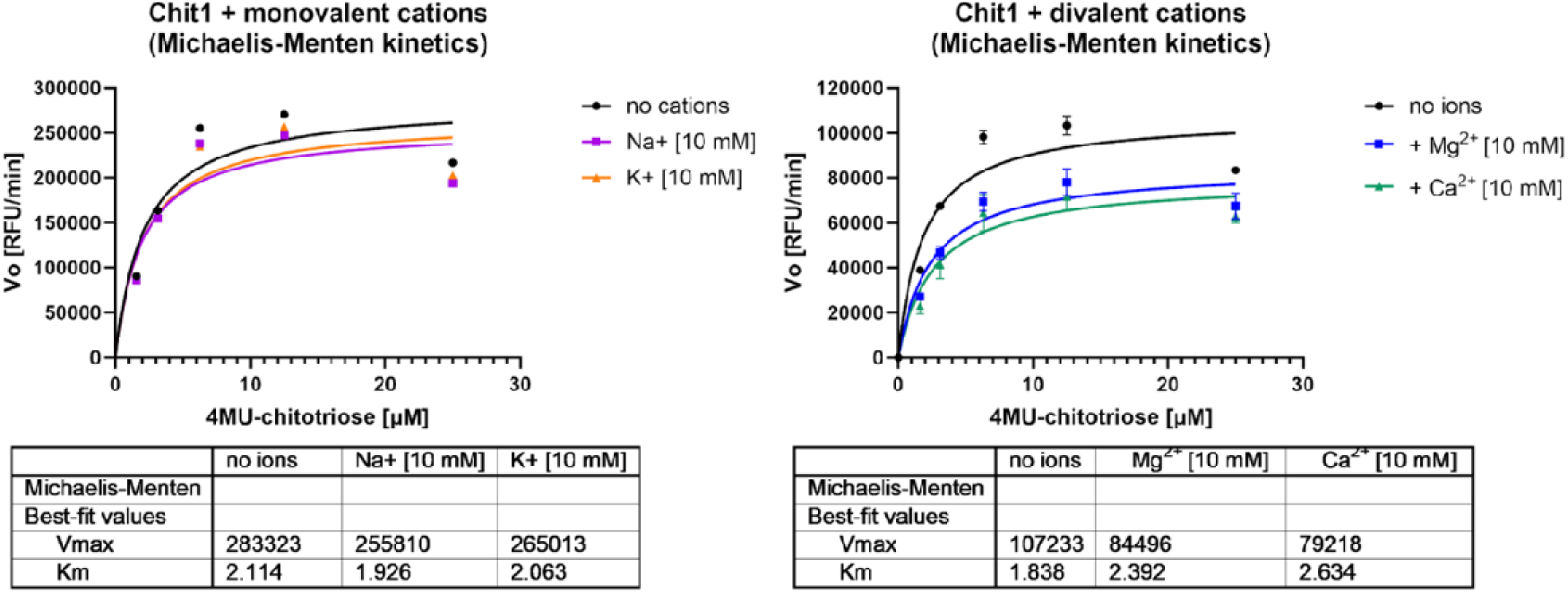
The influence of the most abundant ions in the human body (Na+, K+, Ca2+ and Mg2+) on (GlcNAc)3 substrate evaluated by the means of biological assays. Lower values of Michaelis constant (KM) indicate the tighter substrate binding when Na+ or K+ ions are added in the assay. The M+ requirement for substrate binding with the maximum rate of reaction (Vmax) is independent of [M+] indicate the type of Ia (cofactor-like) mechanism of hCHIT1 activation by alkali ions. Note that the ‘no ions’ system was prepared by a desalination process of the enzyme, which may not have been entirely successful. In such a case, the influence of adding ions may be less pronounced. Moreover, hCHIT1 might not respond well to ionic concentrations exceeding that of its natural habitat.

**ExtendedData Fig. 8.**
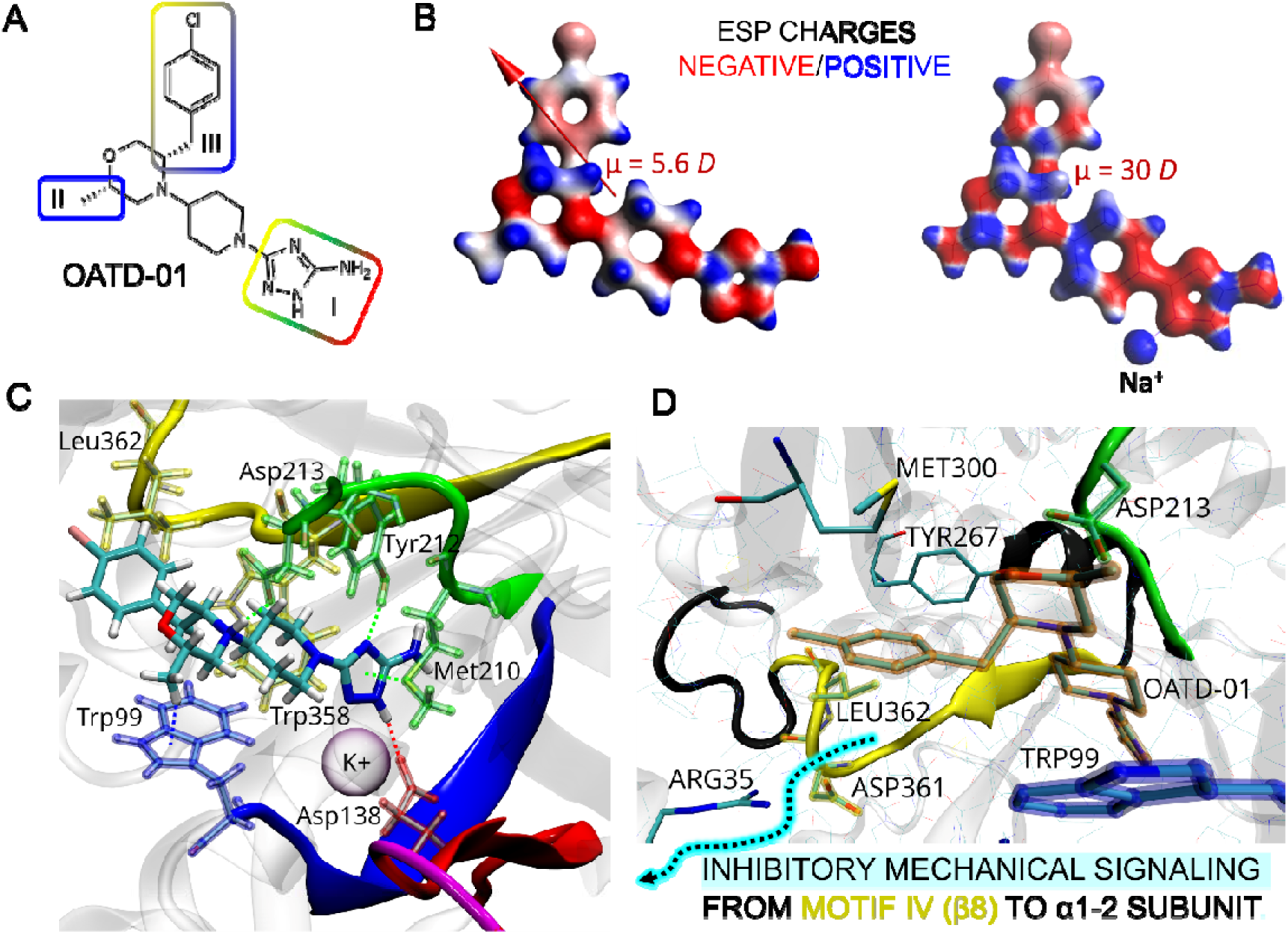
OATD-01 *first-in-class* hCHIT1 inhibitor. **A**) The structural features responsible for the high inhibitory potency against hCHIT1. The colour(s) of each frame correspond to the colour code of the four conserved motifs in Fig. 1A. B) The electrostatic potential (ESP) charges mapped on the electron density surface of OATD-01 alone (left) and interacting with sodium cation (right). The red/blue colours correspond to negative/positive values of charge. The dipole moment is represented as a red arrow and rotated with the parallel orientation of the dipole moment of hCHIT1. The dipole moment value drastically increased upon interaction with the cation but did not change direction (not presented). **C**) The binding mode of the OATD-01 electrostatically stabilized in the active site of CHIT1 by potassium ion, based on 6ZE8 PDB entry (see ExtendedData Fig. 3D). The enzyme is presented as a cartoon while the inhibitor and the key residues are in licorice representations. The OATD-01 inhibitor, by tightly binding the key residues involved in the catalysis, blocks the reciprocal movement of all four conserved motifs. **E)** The binding mode of the 4-chlorobenzene branch is responsible for the long-range effect of the OATD-01 inhibition. The 4-chlorobenzene moiety is locked inside a hydrophobic sub-pocket made up of residues Met300, Tyr276 and Leu362. The latter is next to the residue Asp361, which makes salt-bridge with Arg35, and therefore promotes optimal flexibility of α1-2 subunit and a correct orientation of the Arg40 residue for docking CBM. The allosteric inhibition of hCHIT1 by OATD-01 prevents the assembly of the catalytic and carbohydrate-binding domains into the immunoglobulin-like heterodimer. The black strand indicates the fragment within the conserved motif IV, which is removed by the most common genetic inactivation of the enzyme. OATD-01 has a comparable effect to this mutation.

