## SupplementaryInformation2 for "Mechanism of catalytic apparatus of human chitotriosidase-1 and its dual inactivation mode by the *first-in-class* OATD-01 inhibitor"

```
TITLE     Atomic coordinates of hCHIT1 catalytic domain with (GlcNAc)3 - model
CRYST1   68.708   50.012   46.311  90.00  90.00  90.00 P 1           1
MODEL        1
ATOM      1  N   ALA A  22      38.487  29.297  40.775  1.00  0.00           N
ATOM      2  H1  ALA A  22      39.368  28.783  40.767  1.00  0.00           H
ATOM      3  H2  ALA A  22      38.680  30.271  41.001  1.00  0.00           H
ATOM      4  H3  ALA A  22      37.977  28.937  41.578  1.00  0.00           H
ATOM      5  CA  ALA A  22      37.717  29.118  39.517  1.00  0.00           C
ATOM      6  HA  ALA A  22      37.676  28.048  39.299  1.00  0.00           H
ATOM      7  CB  ALA A  22      36.273  29.585  39.728  1.00  0.00           C
ATOM      8  HB1 ALA A  22      35.703  29.443  38.808  1.00  0.00           H
ATOM      9  HB2 ALA A  22      35.803  29.001  40.517  1.00  0.00           H
ATOM     10  HB3 ALA A  22      36.246  30.638  40.005  1.00  0.00           H
ATOM     11  C   ALA A  22      38.337  29.807  38.274  1.00  0.00           C
ATOM     12  O   ALA A  22      38.749  30.966  38.344  1.00  0.00           O
ATOM     13  N   LYS A  23      38.346  29.113  37.131  1.00  0.00           N
ATOM     14  H   LYS A  23      38.069  28.140  37.169  1.00  0.00           H
ATOM     15  CA  LYS A  23      38.533  29.697  35.792  1.00  0.00           C
ATOM     16  HA  LYS A  23      39.155  30.590  35.861  1.00  0.00           H
ATOM     17  CB  LYS A  23      39.214  28.670  34.860  1.00  0.00           C
ATOM     18  HB1 LYS A  23      38.740  27.695  34.987  1.00  0.00           H
ATOM     19  HB2 LYS A  23      39.048  28.985  33.830  1.00  0.00           H
ATOM     20  CG  LYS A  23      40.732  28.534  35.041  1.00  0.00           C
ATOM     21  HG1 LYS A  23      41.190  29.498  34.818  1.00  0.00           H
ATOM     22  HG2 LYS A  23      40.952  28.251  36.070  1.00  0.00           H
ATOM     23  CD  LYS A  23      41.302  27.471  34.082  1.00  0.00           C
ATOM     24  HD1 LYS A  23      40.875  27.621  33.090  1.00  0.00           H
ATOM     25  HD2 LYS A  23      41.003  26.481  34.428  1.00  0.00           H
ATOM     26  CE  LYS A  23      42.830  27.508  33.925  1.00  0.00           C
ATOM     27  HE1 LYS A  23      43.115  26.767  33.175  1.00  0.00           H
ATOM     28  HE2 LYS A  23      43.130  28.488  33.547  1.00  0.00           H
ATOM     29  NZ  LYS A  23      43.551  27.220  35.185  1.00  0.00           N
ATOM     30  HZ1 LYS A  23      43.452  27.990  35.839  1.00  0.00           H
ATOM     31  HZ2 LYS A  23      43.209  26.365  35.611  1.00  0.00           H
ATOM     32  HZ3 LYS A  23      44.545  27.092  35.003  1.00  0.00           H
ATOM     33  C   LYS A  23      37.188  30.102  35.153  1.00  0.00           C
ATOM     34  O   LYS A  23      36.170  29.441  35.372  1.00  0.00           O
ATOM     35  N   LEU A  24      37.222  31.090  34.254  1.00  0.00           N
ATOM     36  H   LEU A  24      38.084  31.616  34.155  1.00  0.00           H
ATOM     37  CA  LEU A  24      36.235  31.245  33.179  1.00  0.00           C
ATOM     38  HA  LEU A  24      35.591  30.367  33.159  1.00  0.00           H
ATOM     39  CB  LEU A  24      35.333  32.465  33.450  1.00  0.00           C
ATOM     40  HB1 LEU A  24      34.796  32.286  34.383  1.00  0.00           H
ATOM     41  HB2 LEU A  24      35.956  33.348  33.590  1.00  0.00           H
ATOM     42  CG  LEU A  24      34.302  32.786  32.346  1.00  0.00           C
ATOM     43  HG  LEU A  24      34.838  33.160  31.475  1.00  0.00           H
ATOM     44  CD1 LEU A  24      33.463  31.573  31.915  1.00  0.00           C
ATOM     45 HD11 LEU A  24      32.716  31.887  31.186  1.00  0.00           H
ATOM     46 HD12 LEU A  24      34.091  30.818  31.445  1.00  0.00           H
ATOM     47 HD13 LEU A  24      32.961  31.142  32.781  1.00  0.00           H
ATOM     48  CD2 LEU A  24      33.354  33.890  32.837  1.00  0.00           C
ATOM     49 HD21 LEU A  24      32.678  34.186  32.034  1.00  0.00           H
ATOM     50 HD22 LEU A  24      32.769  33.533  33.685  1.00  0.00           H
ATOM     51 HD23 LEU A  24      33.928  34.761  33.147  1.00  0.00           H
```

| | | | | | | | | | | |
|---|---|---|---|---|---|---|---|---|---|---|
| ATOM | 52 | C | LEU | A | 24 | 37.004 | 31.316 | 31.853 | 1.00 | 0.00 | C |
| ATOM | 53 | O | LEU | A | 24 | 37.762 | 32.263 | 31.640 | 1.00 | 0.00 | O |
| ATOM | 54 | N | VAL | A | 25 | 36.857 | 30.281 | 31.024 | 1.00 | 0.00 | N |
| ATOM | 55 | H | VAL | A | 25 | 36.219 | 29.540 | 31.317 | 1.00 | 0.00 | H |
| ATOM | 56 | CA | VAL | A | 25 | 37.619 | 30.064 | 29.781 | 1.00 | 0.00 | C |
| ATOM | 57 | HA | VAL | A | 25 | 38.472 | 30.741 | 29.774 | 1.00 | 0.00 | H |
| ATOM | 58 | CB | VAL | A | 25 | 38.168 | 28.618 | 29.692 | 1.00 | 0.00 | C |
| ATOM | 59 | HB | VAL | A | 25 | 37.333 | 27.925 | 29.600 | 1.00 | 0.00 | H |
| ATOM | 60 | CG1 | VAL | A | 25 | 39.071 | 28.453 | 28.463 | 1.00 | 0.00 | C |
| ATOM | 61 | HG11 | VAL | A | 25 | 39.440 | 27.429 | 28.412 | 1.00 | 0.00 | H |
| ATOM | 62 | HG12 | VAL | A | 25 | 38.513 | 28.657 | 27.548 | 1.00 | 0.00 | H |
| ATOM | 63 | HG13 | VAL | A | 25 | 39.916 | 29.140 | 28.529 | 1.00 | 0.00 | H |
| ATOM | 64 | CG2 | VAL | A | 25 | 38.956 | 28.215 | 30.951 | 1.00 | 0.00 | C |
| ATOM | 65 | HG21 | VAL | A | 25 | 39.393 | 27.224 | 30.816 | 1.00 | 0.00 | H |
| ATOM | 66 | HG22 | VAL | A | 25 | 39.753 | 28.931 | 31.147 | 1.00 | 0.00 | H |
| ATOM | 67 | HG23 | VAL | A | 25 | 38.287 | 28.172 | 31.810 | 1.00 | 0.00 | H |
| ATOM | 68 | C | VAL | A | 25 | 36.730 | 30.361 | 28.567 | 1.00 | 0.00 | C |
| ATOM | 69 | O | VAL | A | 25 | 35.775 | 29.634 | 28.298 | 1.00 | 0.00 | O |
| ATOM | 70 | N | CYS | A | 26 | 37.031 | 31.424 | 27.824 | 1.00 | 0.00 | N |
| ATOM | 71 | H | CYS | A | 26 | 37.841 | 31.984 | 28.089 | 1.00 | 0.00 | H |
| ATOM | 72 | CA | CYS | A | 26 | 36.169 | 31.925 | 26.746 | 1.00 | 0.00 | C |
| ATOM | 73 | HA | CYS | A | 26 | 35.238 | 31.359 | 26.703 | 1.00 | 0.00 | H |
| ATOM | 74 | CB | CYS | A | 26 | 35.813 | 33.376 | 27.071 | 1.00 | 0.00 | C |
| ATOM | 75 | HB1 | CYS | A | 26 | 36.735 | 33.952 | 27.166 | 1.00 | 0.00 | H |
| ATOM | 76 | HB2 | CYS | A | 26 | 35.228 | 33.797 | 26.255 | 1.00 | 0.00 | H |
| ATOM | 77 | SG | CYS | A | 26 | 34.861 | 33.539 | 28.601 | 1.00 | 0.00 | S |
| ATOM | 78 | C | CYS | A | 26 | 36.835 | 31.821 | 25.374 | 1.00 | 0.00 | C |
| ATOM | 79 | O | CYS | A | 26 | 37.918 | 32.367 | 25.172 | 1.00 | 0.00 | O |
| ATOM | 80 | N | TYR | A | 27 | 36.182 | 31.159 | 24.416 | 1.00 | 0.00 | N |
| ATOM | 81 | H | TYR | A | 27 | 35.298 | 30.716 | 24.632 | 1.00 | 0.00 | H |
| ATOM | 82 | CA | TYR | A | 27 | 36.648 | 31.110 | 23.026 | 1.00 | 0.00 | C |
| ATOM | 83 | HA | TYR | A | 27 | 37.738 | 31.135 | 23.021 | 1.00 | 0.00 | H |
| ATOM | 84 | CB | TYR | A | 27 | 36.233 | 29.787 | 22.366 | 1.00 | 0.00 | C |
| ATOM | 85 | HB1 | TYR | A | 27 | 35.163 | 29.631 | 22.518 | 1.00 | 0.00 | H |
| ATOM | 86 | HB2 | TYR | A | 27 | 36.390 | 29.879 | 21.290 | 1.00 | 0.00 | H |
| ATOM | 87 | CG | TYR | A | 27 | 36.997 | 28.559 | 22.841 | 1.00 | 0.00 | C |
| ATOM | 88 | CD1 | TYR | A | 27 | 38.194 | 28.174 | 22.200 | 1.00 | 0.00 | C |
| ATOM | 89 | HD1 | TYR | A | 27 | 38.594 | 28.781 | 21.400 | 1.00 | 0.00 | H |
| ATOM | 90 | CE1 | TYR | A | 27 | 38.862 | 26.991 | 22.582 | 1.00 | 0.00 | C |
| ATOM | 91 | HE1 | TYR | A | 27 | 39.769 | 26.685 | 22.079 | 1.00 | 0.00 | H |
| ATOM | 92 | CZ | TYR | A | 27 | 38.339 | 26.187 | 23.619 | 1.00 | 0.00 | C |
| ATOM | 93 | OH | TYR | A | 27 | 38.957 | 25.037 | 24.005 | 1.00 | 0.00 | O |
| ATOM | 94 | HH | TYR | A | 27 | 39.791 | 24.858 | 23.528 | 1.00 | 0.00 | H |
| ATOM | 95 | CE2 | TYR | A | 27 | 37.144 | 26.577 | 24.259 | 1.00 | 0.00 | C |
| ATOM | 96 | HE2 | TYR | A | 27 | 36.740 | 25.947 | 25.032 | 1.00 | 0.00 | H |
| ATOM | 97 | CD2 | TYR | A | 27 | 36.478 | 27.756 | 23.876 | 1.00 | 0.00 | C |
| ATOM | 98 | HD2 | TYR | A | 27 | 35.549 | 28.026 | 24.357 | 1.00 | 0.00 | H |
| ATOM | 99 | C | TYR | A | 27 | 36.151 | 32.321 | 22.209 | 1.00 | 0.00 | C |
| ATOM | 100 | O | TYR | A | 27 | 34.953 | 32.625 | 22.185 | 1.00 | 0.00 | O |
| ATOM | 101 | N | PHE | A | 28 | 37.088 | 32.979 | 21.523 | 1.00 | 0.00 | N |
| ATOM | 102 | H | PHE | A | 28 | 38.041 | 32.642 | 21.622 | 1.00 | 0.00 | H |
| ATOM | 103 | CA | PHE | A | 28 | 36.879 | 33.970 | 20.464 | 1.00 | 0.00 | C |
| ATOM | 104 | HA | PHE | A | 28 | 35.866 | 34.364 | 20.518 | 1.00 | 0.00 | H |
| ATOM | 105 | CB | PHE | A | 28 | 37.887 | 35.122 | 20.659 | 1.00 | 0.00 | C |

| | | | | | | | | | | | |
|---|---|---|---|---|---|---|---|---|---|---|---|
| ATOM | 106 | HB1 | PHE | A | 28 | 37.532 | 35.778 | 21.452 | 1.00 | 0.00 | H |
| ATOM | 107 | HB2 | PHE | A | 28 | 38.840 | 34.699 | 20.981 | 1.00 | 0.00 | H |
| ATOM | 108 | CG | PHE | A | 28 | 38.132 | 35.944 | 19.405 | 1.00 | 0.00 | C |
| ATOM | 109 | CD1 | PHE | A | 28 | 37.121 | 36.789 | 18.913 | 1.00 | 0.00 | C |
| ATOM | 110 | HD1 | PHE | A | 28 | 36.200 | 36.904 | 19.464 | 1.00 | 0.00 | H |
| ATOM | 111 | CE1 | PHE | A | 28 | 37.278 | 37.426 | 17.671 | 1.00 | 0.00 | C |
| ATOM | 112 | HE1 | PHE | A | 28 | 36.474 | 38.023 | 17.266 | 1.00 | 0.00 | H |
| ATOM | 113 | CZ | PHE | A | 28 | 38.457 | 37.247 | 16.927 | 1.00 | 0.00 | C |
| ATOM | 114 | HZ | PHE | A | 28 | 38.569 | 37.715 | 15.959 | 1.00 | 0.00 | H |
| ATOM | 115 | CE2 | PHE | A | 28 | 39.477 | 36.422 | 17.426 | 1.00 | 0.00 | C |
| ATOM | 116 | HE2 | PHE | A | 28 | 40.370 | 36.259 | 16.841 | 1.00 | 0.00 | H |
| ATOM | 117 | CD2 | PHE | A | 28 | 39.314 | 35.767 | 18.659 | 1.00 | 0.00 | C |
| ATOM | 118 | HD2 | PHE | A | 28 | 40.085 | 35.097 | 19.010 | 1.00 | 0.00 | H |
| ATOM | 119 | C | PHE | A | 28 | 37.088 | 33.302 | 19.094 | 1.00 | 0.00 | C |
| ATOM | 120 | O | PHE | A | 28 | 37.997 | 32.480 | 18.967 | 1.00 | 0.00 | O |
| ATOM | 121 | N | THR | A | 29 | 36.307 | 33.672 | 18.067 | 1.00 | 0.00 | N |
| ATOM | 122 | H | THR | A | 29 | 35.599 | 34.383 | 18.226 | 1.00 | 0.00 | H |
| ATOM | 123 | CA | THR | A | 29 | 36.432 | 33.123 | 16.703 | 1.00 | 0.00 | C |
| ATOM | 124 | HA | THR | A | 29 | 37.217 | 32.368 | 16.694 | 1.00 | 0.00 | H |
| ATOM | 125 | CB | THR | A | 29 | 35.147 | 32.416 | 16.255 | 1.00 | 0.00 | C |
| ATOM | 126 | HB | THR | A | 29 | 35.332 | 31.991 | 15.271 | 1.00 | 0.00 | H |
| ATOM | 127 | CG2 | THR | A | 29 | 34.724 | 31.288 | 17.199 | 1.00 | 0.00 | C |
| ATOM | 128 | HG21 | THR | A | 29 | 33.832 | 30.804 | 16.803 | 1.00 | 0.00 | H |
| ATOM | 129 | HG22 | THR | A | 29 | 35.521 | 30.549 | 17.270 | 1.00 | 0.00 | H |
| ATOM | 130 | HG23 | THR | A | 29 | 34.502 | 31.676 | 18.193 | 1.00 | 0.00 | H |
| ATOM | 131 | OG1 | THR | A | 29 | 34.080 | 33.321 | 16.149 | 1.00 | 0.00 | O |
| ATOM | 132 | HG1 | THR | A | 29 | 33.545 | 33.271 | 16.963 | 1.00 | 0.00 | H |
| ATOM | 133 | C | THR | A | 29 | 36.847 | 34.197 | 15.699 | 1.00 | 0.00 | C |
| ATOM | 134 | O | THR | A | 29 | 36.228 | 35.257 | 15.600 | 1.00 | 0.00 | O |
| ATOM | 135 | N | ASN | A | 30 | 37.887 | 33.922 | 14.906 | 1.00 | 0.00 | N |
| ATOM | 136 | H | ASN | A | 30 | 38.375 | 33.040 | 15.035 | 1.00 | 0.00 | H |
| ATOM | 137 | CA | ASN | A | 30 | 38.421 | 34.870 | 13.914 | 1.00 | 0.00 | C |
| ATOM | 138 | HA | ASN | A | 30 | 38.343 | 35.861 | 14.362 | 1.00 | 0.00 | H |
| ATOM | 139 | CB | ASN | A | 30 | 39.931 | 34.636 | 13.699 | 1.00 | 0.00 | C |
| ATOM | 140 | HB1 | ASN | A | 30 | 40.341 | 35.512 | 13.195 | 1.00 | 0.00 | H |
| ATOM | 141 | HB2 | ASN | A | 30 | 40.420 | 34.568 | 14.671 | 1.00 | 0.00 | H |
| ATOM | 142 | CG | ASN | A | 30 | 40.315 | 33.405 | 12.881 | 1.00 | 0.00 | C |
| ATOM | 143 | OD1 | ASN | A | 30 | 39.523 | 32.512 | 12.606 | 1.00 | 0.00 | O |
| ATOM | 144 | ND2 | ASN | A | 30 | 41.559 | 33.321 | 12.451 | 1.00 | 0.00 | N |
| ATOM | 145 | HD21 | ASN | A | 30 | 42.218 | 34.085 | 12.598 | 1.00 | 0.00 | H |
| ATOM | 146 | HD22 | ASN | A | 30 | 41.840 | 32.537 | 11.886 | 1.00 | 0.00 | H |
| ATOM | 147 | C | ASN | A | 30 | 37.607 | 34.959 | 12.599 | 1.00 | 0.00 | C |
| ATOM | 148 | O | ASN | A | 30 | 38.012 | 35.660 | 11.670 | 1.00 | 0.00 | O |
| ATOM | 149 | N | TRP | A | 31 | 36.450 | 34.285 | 12.511 | 1.00 | 0.00 | N |
| ATOM | 150 | H | TRP | A | 31 | 36.200 | 33.690 | 13.288 | 1.00 | 0.00 | H |
| ATOM | 151 | CA | TRP | A | 31 | 35.533 | 34.331 | 11.360 | 1.00 | 0.00 | C |
| ATOM | 152 | HA | TRP | A | 31 | 36.010 | 34.877 | 10.545 | 1.00 | 0.00 | H |
| ATOM | 153 | CB | TRP | A | 31 | 35.297 | 32.896 | 10.867 | 1.00 | 0.00 | C |
| ATOM | 154 | HB1 | TRP | A | 31 | 36.257 | 32.482 | 10.555 | 1.00 | 0.00 | H |
| ATOM | 155 | HB2 | TRP | A | 31 | 34.661 | 32.925 | 9.981 | 1.00 | 0.00 | H |
| ATOM | 156 | CG | TRP | A | 31 | 34.683 | 31.953 | 11.857 | 1.00 | 0.00 | C |
| ATOM | 157 | CD1 | TRP | A | 31 | 33.372 | 31.886 | 12.178 | 1.00 | 0.00 | C |
| ATOM | 158 | HD1 | TRP | A | 31 | 32.593 | 32.507 | 11.756 | 1.00 | 0.00 | H |
| ATOM | 159 | NE1 | TRP | A | 31 | 33.183 | 30.936 | 13.161 | 1.00 | 0.00 | N |

| | | | | | | | | | | | |
|---|---|---|---|---|---|---|---|---|---|---|---|
| ATOM | 160 | HE1 | TRP | A | 31 | 32.319 | 30.847 | 13.692 | 1.00 | 0.00 | H |
| ATOM | 161 | CE2 | TRP | A | 31 | 34.359 | 30.306 | 13.494 | 1.00 | 0.00 | C |
| ATOM | 162 | CZ2 | TRP | A | 31 | 34.680 | 29.309 | 14.426 | 1.00 | 0.00 | C |
| ATOM | 163 | HZ2 | TRP | A | 31 | 33.903 | 28.854 | 15.023 | 1.00 | 0.00 | H |
| ATOM | 164 | CH2 | TRP | A | 31 | 36.027 | 28.963 | 14.616 | 1.00 | 0.00 | C |
| ATOM | 165 | HH2 | TRP | A | 31 | 36.293 | 28.238 | 15.369 | 1.00 | 0.00 | H |
| ATOM | 166 | CZ3 | TRP | A | 31 | 37.027 | 29.598 | 13.858 | 1.00 | 0.00 | C |
| ATOM | 167 | HZ3 | TRP | A | 31 | 38.068 | 29.365 | 14.028 | 1.00 | 0.00 | H |
| ATOM | 168 | CE3 | TRP | A | 31 | 36.687 | 30.570 | 12.899 | 1.00 | 0.00 | C |
| ATOM | 169 | HE3 | TRP | A | 31 | 37.465 | 31.056 | 12.331 | 1.00 | 0.00 | H |
| ATOM | 170 | CD2 | TRP | A | 31 | 35.346 | 30.959 | 12.698 | 1.00 | 0.00 | C |
| ATOM | 171 | C | TRP | A | 31 | 34.193 | 35.048 | 11.631 | 1.00 | 0.00 | C |
| ATOM | 172 | O | TRP | A | 31 | 33.459 | 35.329 | 10.680 | 1.00 | 0.00 | O |
| ATOM | 173 | N | ALA | A | 32 | 33.851 | 35.376 | 12.885 | 1.00 | 0.00 | N |
| ATOM | 174 | H | ALA | A | 32 | 34.469 | 35.113 | 13.644 | 1.00 | 0.00 | H |
| ATOM | 175 | CA | ALA | A | 32 | 32.604 | 36.080 | 13.225 | 1.00 | 0.00 | C |
| ATOM | 176 | HA | ALA | A | 32 | 31.768 | 35.553 | 12.760 | 1.00 | 0.00 | H |
| ATOM | 177 | CB | ALA | A | 32 | 32.416 | 36.005 | 14.741 | 1.00 | 0.00 | C |
| ATOM | 178 | HB1 | ALA | A | 32 | 31.506 | 36.535 | 15.023 | 1.00 | 0.00 | H |
| ATOM | 179 | HB2 | ALA | A | 32 | 32.316 | 34.962 | 15.044 | 1.00 | 0.00 | H |
| ATOM | 180 | HB3 | ALA | A | 32 | 33.272 | 36.446 | 15.256 | 1.00 | 0.00 | H |
| ATOM | 181 | C | ALA | A | 32 | 32.548 | 37.534 | 12.705 | 1.00 | 0.00 | C |
| ATOM | 182 | O | ALA | A | 32 | 31.461 | 38.062 | 12.468 | 1.00 | 0.00 | O |
| ATOM | 183 | N | GLN | A | 33 | 33.708 | 38.146 | 12.447 | 1.00 | 0.00 | N |
| ATOM | 184 | H | GLN | A | 33 | 34.546 | 37.706 | 12.816 | 1.00 | 0.00 | H |
| ATOM | 185 | CA | GLN | A | 33 | 33.872 | 39.446 | 11.782 | 1.00 | 0.00 | C |
| ATOM | 186 | HA | GLN | A | 33 | 33.395 | 40.209 | 12.396 | 1.00 | 0.00 | H |
| ATOM | 187 | CB | GLN | A | 33 | 35.374 | 39.770 | 11.691 | 1.00 | 0.00 | C |
| ATOM | 188 | HB1 | GLN | A | 33 | 35.476 | 40.751 | 11.226 | 1.00 | 0.00 | H |
| ATOM | 189 | HB2 | GLN | A | 33 | 35.771 | 39.846 | 12.703 | 1.00 | 0.00 | H |
| ATOM | 190 | CG | GLN | A | 33 | 36.225 | 38.746 | 10.901 | 1.00 | 0.00 | C |
| ATOM | 191 | HG1 | GLN | A | 33 | 36.183 | 37.775 | 11.393 | 1.00 | 0.00 | H |
| ATOM | 192 | HG2 | GLN | A | 33 | 35.824 | 38.630 | 9.895 | 1.00 | 0.00 | H |
| ATOM | 193 | CD | GLN | A | 33 | 37.690 | 39.174 | 10.774 | 1.00 | 0.00 | C |
| ATOM | 194 | OE1 | GLN | A | 33 | 38.006 | 40.358 | 10.730 | 1.00 | 0.00 | O |
| ATOM | 195 | NE2 | GLN | A | 33 | 38.641 | 38.266 | 10.725 | 1.00 | 0.00 | N |
| ATOM | 196 | HE21 | GLN | A | 33 | 38.415 | 37.285 | 10.861 | 1.00 | 0.00 | H |
| ATOM | 197 | HE22 | GLN | A | 33 | 39.606 | 38.552 | 10.614 | 1.00 | 0.00 | H |
| ATOM | 198 | C | GLN | A | 33 | 33.225 | 39.553 | 10.389 | 1.00 | 0.00 | C |
| ATOM | 199 | O | GLN | A | 33 | 33.003 | 40.666 | 9.903 | 1.00 | 0.00 | O |
| ATOM | 200 | N | TYR | A | 34 | 32.927 | 38.419 | 9.747 | 1.00 | 0.00 | N |
| ATOM | 201 | H | TYR | A | 34 | 33.164 | 37.544 | 10.198 | 1.00 | 0.00 | H |
| ATOM | 202 | CA | TYR | A | 34 | 32.301 | 38.334 | 8.424 | 1.00 | 0.00 | C |
| ATOM | 203 | HA | TYR | A | 34 | 32.546 | 39.235 | 7.860 | 1.00 | 0.00 | H |
| ATOM | 204 | CB | TYR | A | 34 | 32.900 | 37.125 | 7.682 | 1.00 | 0.00 | C |
| ATOM | 205 | HB1 | TYR | A | 34 | 32.588 | 36.222 | 8.204 | 1.00 | 0.00 | H |
| ATOM | 206 | HB2 | TYR | A | 34 | 32.471 | 37.077 | 6.682 | 1.00 | 0.00 | H |
| ATOM | 207 | CG | TYR | A | 34 | 34.415 | 37.110 | 7.531 | 1.00 | 0.00 | C |
| ATOM | 208 | CD1 | TYR | A | 34 | 35.092 | 38.228 | 7.002 | 1.00 | 0.00 | C |
| ATOM | 209 | HD1 | TYR | A | 34 | 34.540 | 39.112 | 6.715 | 1.00 | 0.00 | H |
| ATOM | 210 | CE1 | TYR | A | 34 | 36.487 | 38.191 | 6.814 | 1.00 | 0.00 | C |
| ATOM | 211 | HE1 | TYR | A | 34 | 36.998 | 39.042 | 6.389 | 1.00 | 0.00 | H |
| ATOM | 212 | CZ | TYR | A | 34 | 37.216 | 37.026 | 7.142 | 1.00 | 0.00 | C |
| ATOM | 213 | OH | TYR | A | 34 | 38.555 | 36.979 | 6.907 | 1.00 | 0.00 | O |

| | | | | | | | | | | | |
|---|---|---|---|---|---|---|---|---|---|---|---|
| ATOM | 214 | HH | TYR | A | 34 | 38.942 | 36.096 | 7.041 | 1.00 | 0.00 | H |
| ATOM | 215 | CE2 | TYR | A | 34 | 36.543 | 35.909 | 7.683 | 1.00 | 0.00 | C |
| ATOM | 216 | HE2 | TYR | A | 34 | 37.095 | 35.013 | 7.929 | 1.00 | 0.00 | H |
| ATOM | 217 | CD2 | TYR | A | 34 | 35.148 | 35.956 | 7.874 | 1.00 | 0.00 | C |
| ATOM | 218 | HD2 | TYR | A | 34 | 34.638 | 35.086 | 8.261 | 1.00 | 0.00 | H |
| ATOM | 219 | C | TYR | A | 34 | 30.758 | 38.245 | 8.466 | 1.00 | 0.00 | C |
| ATOM | 220 | O | TYR | A | 34 | 30.122 | 38.209 | 7.406 | 1.00 | 0.00 | O |
| ATOM | 221 | N | ARG | A | 35 | 30.135 | 38.207 | 9.653 | 1.00 | 0.00 | N |
| ATOM | 222 | H | ARG | A | 35 | 30.706 | 38.252 | 10.490 | 1.00 | 0.00 | H |
| ATOM | 223 | CA | ARG | A | 35 | 28.669 | 38.271 | 9.814 | 1.00 | 0.00 | C |
| ATOM | 224 | HA | ARG | A | 35 | 28.208 | 37.666 | 9.036 | 1.00 | 0.00 | H |
| ATOM | 225 | CB | ARG | A | 35 | 28.246 | 37.689 | 11.170 | 1.00 | 0.00 | C |
| ATOM | 226 | HB1 | ARG | A | 35 | 28.794 | 38.182 | 11.973 | 1.00 | 0.00 | H |
| ATOM | 227 | HB2 | ARG | A | 35 | 27.178 | 37.853 | 11.323 | 1.00 | 0.00 | H |
| ATOM | 228 | CG | ARG | A | 35 | 28.520 | 36.181 | 11.191 | 1.00 | 0.00 | C |
| ATOM | 229 | HG1 | ARG | A | 35 | 27.933 | 35.693 | 10.412 | 1.00 | 0.00 | H |
| ATOM | 230 | HG2 | ARG | A | 35 | 29.579 | 36.001 | 10.999 | 1.00 | 0.00 | H |
| ATOM | 231 | CD | ARG | A | 35 | 28.170 | 35.566 | 12.541 | 1.00 | 0.00 | C |
| ATOM | 232 | HD1 | ARG | A | 35 | 28.712 | 36.114 | 13.312 | 1.00 | 0.00 | H |
| ATOM | 233 | HD2 | ARG | A | 35 | 27.097 | 35.645 | 12.724 | 1.00 | 0.00 | H |
| ATOM | 234 | NE | ARG | A | 35 | 28.582 | 34.158 | 12.546 | 1.00 | 0.00 | N |
| ATOM | 235 | HE | ARG | A | 35 | 28.696 | 33.713 | 11.642 | 1.00 | 0.00 | H |
| ATOM | 236 | CZ | ARG | A | 35 | 29.047 | 33.474 | 13.577 | 1.00 | 0.00 | C |
| ATOM | 237 | NH1 | ARG | A | 35 | 29.037 | 33.945 | 14.804 | 1.00 | 0.00 | N |
| ATOM | 238 | HH11 | ARG | A | 35 | 28.628 | 34.850 | 15.007 | 1.00 | 0.00 | H |
| ATOM | 239 | HH12 | ARG | A | 35 | 29.519 | 33.461 | 15.562 | 1.00 | 0.00 | H |
| ATOM | 240 | NH2 | ARG | A | 35 | 29.563 | 32.295 | 13.341 | 1.00 | 0.00 | N |
| ATOM | 241 | HH21 | ARG | A | 35 | 29.657 | 31.974 | 12.384 | 1.00 | 0.00 | H |
| ATOM | 242 | HH22 | ARG | A | 35 | 30.006 | 31.786 | 14.100 | 1.00 | 0.00 | H |
| ATOM | 243 | C | ARG | A | 35 | 28.162 | 39.707 | 9.646 | 1.00 | 0.00 | C |
| ATOM | 244 | O | ARG | A | 35 | 28.942 | 40.642 | 9.771 | 1.00 | 0.00 | O |
| ATOM | 245 | N | GLN | A | 36 | 26.881 | 39.903 | 9.342 | 1.00 | 0.00 | N |
| ATOM | 246 | H | GLN | A | 36 | 26.267 | 39.101 | 9.285 | 1.00 | 0.00 | H |
| ATOM | 247 | CA | GLN | A | 36 | 26.322 | 41.243 | 9.107 | 1.00 | 0.00 | C |
| ATOM | 248 | HA | GLN | A | 36 | 27.125 | 41.931 | 8.834 | 1.00 | 0.00 | H |
| ATOM | 249 | CB | GLN | A | 36 | 25.329 | 41.211 | 7.929 | 1.00 | 0.00 | C |
| ATOM | 250 | HB1 | GLN | A | 36 | 24.431 | 40.679 | 8.241 | 1.00 | 0.00 | H |
| ATOM | 251 | HB2 | GLN | A | 36 | 25.040 | 42.236 | 7.690 | 1.00 | 0.00 | H |
| ATOM | 252 | CG | GLN | A | 36 | 25.863 | 40.546 | 6.646 | 1.00 | 0.00 | C |
| ATOM | 253 | HG1 | GLN | A | 36 | 25.922 | 39.471 | 6.812 | 1.00 | 0.00 | H |
| ATOM | 254 | HG2 | GLN | A | 36 | 25.147 | 40.712 | 5.840 | 1.00 | 0.00 | H |
| ATOM | 255 | CD | GLN | A | 36 | 27.234 | 41.057 | 6.196 | 1.00 | 0.00 | C |
| ATOM | 256 | OE1 | GLN | A | 36 | 27.526 | 42.247 | 6.193 | 1.00 | 0.00 | O |
| ATOM | 257 | NE2 | GLN | A | 36 | 28.148 | 40.184 | 5.837 | 1.00 | 0.00 | N |
| ATOM | 258 | HE21 | GLN | A | 36 | 27.927 | 39.203 | 5.771 | 1.00 | 0.00 | H |
| ATOM | 259 | HE22 | GLN | A | 36 | 29.053 | 40.545 | 5.572 | 1.00 | 0.00 | H |
| ATOM | 260 | C | GLN | A | 36 | 25.662 | 41.831 | 10.365 | 1.00 | 0.00 | C |
| ATOM | 261 | O | GLN | A | 36 | 25.337 | 41.110 | 11.311 | 1.00 | 0.00 | O |
| ATOM | 262 | N | GLY | A | 37 | 25.444 | 43.151 | 10.357 | 1.00 | 0.00 | N |
| ATOM | 263 | H | GLY | A | 37 | 25.790 | 43.687 | 9.567 | 1.00 | 0.00 | H |
| ATOM | 264 | CA | GLY | A | 37 | 24.563 | 43.856 | 11.293 | 1.00 | 0.00 | C |
| ATOM | 265 | HA1 | GLY | A | 37 | 24.454 | 44.897 | 10.988 | 1.00 | 0.00 | H |
| ATOM | 266 | HA2 | GLY | A | 37 | 23.581 | 43.384 | 11.255 | 1.00 | 0.00 | H |
| ATOM | 267 | C | GLY | A | 37 | 25.033 | 43.832 | 12.746 | 1.00 | 0.00 | C |

| | | | | | | | | | | | |
|---|---|---|---|---|---|---|---|---|---|---|---|
| ATOM | 268 | O | GLY | A | 37 | 26.236 | 43.812 | 13.037 | 1.00 | 0.00 | O |
| ATOM | 269 | N | GLU | A | 38 | 24.070 | 43.835 | 13.665 | 1.00 | 0.00 | N |
| ATOM | 270 | H | GLU | A | 38 | 23.113 | 43.968 | 13.342 | 1.00 | 0.00 | H |
| ATOM | 271 | CA | GLU | A | 38 | 24.286 | 43.792 | 15.113 | 1.00 | 0.00 | C |
| ATOM | 272 | HA | GLU | A | 38 | 24.888 | 44.654 | 15.404 | 1.00 | 0.00 | H |
| ATOM | 273 | CB | GLU | A | 38 | 22.939 | 43.870 | 15.856 | 1.00 | 0.00 | C |
| ATOM | 274 | HB1 | GLU | A | 38 | 22.330 | 43.006 | 15.589 | 1.00 | 0.00 | H |
| ATOM | 275 | HB2 | GLU | A | 38 | 23.167 | 43.806 | 16.920 | 1.00 | 0.00 | H |
| ATOM | 276 | CG | GLU | A | 38 | 22.117 | 45.155 | 15.649 | 1.00 | 0.00 | C |
| ATOM | 277 | HG1 | GLU | A | 38 | 21.563 | 45.353 | 16.566 | 1.00 | 0.00 | H |
| ATOM | 278 | HG2 | GLU | A | 38 | 22.790 | 45.999 | 15.487 | 1.00 | 0.00 | H |
| ATOM | 279 | CD | GLU | A | 38 | 21.114 | 45.068 | 14.495 | 1.00 | 0.00 | C |
| ATOM | 280 | OE1 | GLU | A | 38 | 21.552 | 44.928 | 13.332 | 1.00 | 0.00 | O |
| ATOM | 281 | OE2 | GLU | A | 38 | 19.890 | 45.223 | 14.724 | 1.00 | 0.00 | O |
| ATOM | 282 | C | GLU | A | 38 | 25.031 | 42.520 | 15.559 | 1.00 | 0.00 | C |
| ATOM | 283 | O | GLU | A | 38 | 25.748 | 42.547 | 16.558 | 1.00 | 0.00 | O |
| ATOM | 284 | N | ALA | A | 39 | 24.914 | 41.424 | 14.800 | 1.00 | 0.00 | N |
| ATOM | 285 | H | ALA | A | 39 | 24.325 | 41.473 | 13.981 | 1.00 | 0.00 | H |
| ATOM | 286 | CA | ALA | A | 39 | 25.547 | 40.139 | 15.091 | 1.00 | 0.00 | C |
| ATOM | 287 | HA | ALA | A | 39 | 25.492 | 39.952 | 16.166 | 1.00 | 0.00 | H |
| ATOM | 288 | CB | ALA | A | 39 | 24.736 | 39.070 | 14.363 | 1.00 | 0.00 | C |
| ATOM | 289 | HB1 | ALA | A | 39 | 25.155 | 38.089 | 14.576 | 1.00 | 0.00 | H |
| ATOM | 290 | HB2 | ALA | A | 39 | 23.698 | 39.095 | 14.699 | 1.00 | 0.00 | H |
| ATOM | 291 | HB3 | ALA | A | 39 | 24.784 | 39.245 | 13.289 | 1.00 | 0.00 | H |
| ATOM | 292 | C | ALA | A | 39 | 27.032 | 40.054 | 14.690 | 1.00 | 0.00 | C |
| ATOM | 293 | O | ALA | A | 39 | 27.717 | 39.118 | 15.112 | 1.00 | 0.00 | O |
| ATOM | 294 | N | ARG | A | 40 | 27.549 | 41.005 | 13.895 | 1.00 | 0.00 | N |
| ATOM | 295 | H | ARG | A | 40 | 26.927 | 41.721 | 13.543 | 1.00 | 0.00 | H |
| ATOM | 296 | CA | ARG | A | 40 | 28.981 | 41.089 | 13.577 | 1.00 | 0.00 | C |
| ATOM | 297 | HA | ARG | A | 40 | 29.288 | 40.148 | 13.120 | 1.00 | 0.00 | H |
| ATOM | 298 | CB | ARG | A | 40 | 29.255 | 42.226 | 12.579 | 1.00 | 0.00 | C |
| ATOM | 299 | HB1 | ARG | A | 40 | 28.546 | 42.159 | 11.755 | 1.00 | 0.00 | H |
| ATOM | 300 | HB2 | ARG | A | 40 | 29.103 | 43.185 | 13.072 | 1.00 | 0.00 | H |
| ATOM | 301 | CG | ARG | A | 40 | 30.694 | 42.164 | 12.032 | 1.00 | 0.00 | C |
| ATOM | 302 | HG1 | ARG | A | 40 | 31.398 | 42.084 | 12.860 | 1.00 | 0.00 | H |
| ATOM | 303 | HG2 | ARG | A | 40 | 30.808 | 41.272 | 11.417 | 1.00 | 0.00 | H |
| ATOM | 304 | CD | ARG | A | 40 | 31.088 | 43.404 | 11.217 | 1.00 | 0.00 | C |
| ATOM | 305 | HD1 | ARG | A | 40 | 31.112 | 44.264 | 11.888 | 1.00 | 0.00 | H |
| ATOM | 306 | HD2 | ARG | A | 40 | 32.096 | 43.265 | 10.824 | 1.00 | 0.00 | H |
| ATOM | 307 | NE | ARG | A | 40 | 30.152 | 43.697 | 10.120 | 1.00 | 0.00 | N |
| ATOM | 308 | HE | ARG | A | 40 | 29.390 | 44.327 | 10.348 | 1.00 | 0.00 | H |
| ATOM | 309 | CZ | ARG | A | 40 | 30.173 | 43.199 | 8.891 | 1.00 | 0.00 | C |
| ATOM | 310 | NH1 | ARG | A | 40 | 31.057 | 42.307 | 8.505 | 1.00 | 0.00 | N |
| ATOM | 311 | HH11 | ARG | A | 40 | 31.673 | 41.874 | 9.180 | 1.00 | 0.00 | H |
| ATOM | 312 | HH12 | ARG | A | 40 | 30.995 | 41.930 | 7.570 | 1.00 | 0.00 | H |
| ATOM | 313 | NH2 | ARG | A | 40 | 29.267 | 43.580 | 8.020 | 1.00 | 0.00 | N |
| ATOM | 314 | HH21 | ARG | A | 40 | 28.523 | 44.217 | 8.284 | 1.00 | 0.00 | H |
| ATOM | 315 | HH22 | ARG | A | 40 | 29.169 | 43.113 | 7.129 | 1.00 | 0.00 | H |
| ATOM | 316 | C | ARG | A | 40 | 29.788 | 41.312 | 14.859 | 1.00 | 0.00 | C |
| ATOM | 317 | O | ARG | A | 40 | 29.621 | 42.346 | 15.514 | 1.00 | 0.00 | O |
| ATOM | 318 | N | PHE | A | 41 | 30.684 | 40.381 | 15.184 | 1.00 | 0.00 | N |
| ATOM | 319 | H | PHE | A | 41 | 30.761 | 39.571 | 14.586 | 1.00 | 0.00 | H |
| ATOM | 320 | CA | PHE | A | 41 | 31.616 | 40.490 | 16.307 | 1.00 | 0.00 | C |
| ATOM | 321 | HA | PHE | A | 41 | 31.188 | 41.137 | 17.076 | 1.00 | 0.00 | H |

| | | | | | | | | | | | |
|---|---|---|---|---|---|---|---|---|---|---|---|
| ATOM | 322 | CB | PHE | A | 41 | 31.813 | 39.094 | 16.911 | 1.00 | 0.00 | C |
| ATOM | 323 | HB1 | PHE | A | 41 | 30.841 | 38.604 | 16.998 | 1.00 | 0.00 | H |
| ATOM | 324 | HB2 | PHE | A | 41 | 32.426 | 38.495 | 16.237 | 1.00 | 0.00 | H |
| ATOM | 325 | CG | PHE | A | 41 | 32.455 | 39.110 | 18.281 | 1.00 | 0.00 | C |
| ATOM | 326 | CD1 | PHE | A | 41 | 33.855 | 39.155 | 18.417 | 1.00 | 0.00 | C |
| ATOM | 327 | HD1 | PHE | A | 41 | 34.483 | 39.177 | 17.538 | 1.00 | 0.00 | H |
| ATOM | 328 | CE1 | PHE | A | 41 | 34.436 | 39.191 | 19.697 | 1.00 | 0.00 | C |
| ATOM | 329 | HE1 | PHE | A | 41 | 35.509 | 39.239 | 19.801 | 1.00 | 0.00 | H |
| ATOM | 330 | CZ | PHE | A | 41 | 33.619 | 39.165 | 20.841 | 1.00 | 0.00 | C |
| ATOM | 331 | HZ | PHE | A | 41 | 34.056 | 39.189 | 21.826 | 1.00 | 0.00 | H |
| ATOM | 332 | CE2 | PHE | A | 41 | 32.222 | 39.113 | 20.704 | 1.00 | 0.00 | C |
| ATOM | 333 | HE2 | PHE | A | 41 | 31.586 | 39.086 | 21.578 | 1.00 | 0.00 | H |
| ATOM | 334 | CD2 | PHE | A | 41 | 31.643 | 39.094 | 19.427 | 1.00 | 0.00 | C |
| ATOM | 335 | HD2 | PHE | A | 41 | 30.569 | 39.064 | 19.331 | 1.00 | 0.00 | H |
| ATOM | 336 | C | PHE | A | 41 | 32.950 | 41.091 | 15.835 | 1.00 | 0.00 | C |
| ATOM | 337 | O | PHE | A | 41 | 33.416 | 40.752 | 14.748 | 1.00 | 0.00 | O |
| ATOM | 338 | N | LEU | A | 42 | 33.569 | 41.964 | 16.636 | 1.00 | 0.00 | N |
| ATOM | 339 | H | LEU | A | 42 | 33.152 | 42.180 | 17.535 | 1.00 | 0.00 | H |
| ATOM | 340 | CA | LEU | A | 42 | 34.820 | 42.668 | 16.309 | 1.00 | 0.00 | C |
| ATOM | 341 | HA | LEU | A | 42 | 35.409 | 42.050 | 15.631 | 1.00 | 0.00 | H |
| ATOM | 342 | CB | LEU | A | 42 | 34.466 | 44.013 | 15.632 | 1.00 | 0.00 | C |
| ATOM | 343 | HB1 | LEU | A | 42 | 33.749 | 44.530 | 16.270 | 1.00 | 0.00 | H |
| ATOM | 344 | HB2 | LEU | A | 42 | 35.353 | 44.634 | 15.589 | 1.00 | 0.00 | H |
| ATOM | 345 | CG | LEU | A | 42 | 33.890 | 43.941 | 14.202 | 1.00 | 0.00 | C |
| ATOM | 346 | HG | LEU | A | 42 | 32.979 | 43.344 | 14.204 | 1.00 | 0.00 | H |
| ATOM | 347 | CD1 | LEU | A | 42 | 33.522 | 45.356 | 13.736 | 1.00 | 0.00 | C |
| ATOM | 348 | HD11 | LEU | A | 42 | 33.086 | 45.309 | 12.738 | 1.00 | 0.00 | H |
| ATOM | 349 | HD12 | LEU | A | 42 | 32.788 | 45.794 | 14.411 | 1.00 | 0.00 | H |
| ATOM | 350 | HD13 | LEU | A | 42 | 34.409 | 45.990 | 13.709 | 1.00 | 0.00 | H |
| ATOM | 351 | CD2 | LEU | A | 42 | 34.886 | 43.340 | 13.197 | 1.00 | 0.00 | C |
| ATOM | 352 | HD21 | LEU | A | 42 | 34.449 | 43.341 | 12.198 | 1.00 | 0.00 | H |
| ATOM | 353 | HD22 | LEU | A | 42 | 35.806 | 43.926 | 13.184 | 1.00 | 0.00 | H |
| ATOM | 354 | HD23 | LEU | A | 42 | 35.121 | 42.311 | 13.462 | 1.00 | 0.00 | H |
| ATOM | 355 | C | LEU | A | 42 | 35.672 | 42.897 | 17.581 | 1.00 | 0.00 | C |
| ATOM | 356 | O | LEU | A | 42 | 35.143 | 42.739 | 18.684 | 1.00 | 0.00 | O |
| ATOM | 357 | N | PRO | A | 43 | 36.963 | 43.290 | 17.463 | 1.00 | 0.00 | N |
| ATOM | 358 | CD | PRO | A | 43 | 37.736 | 43.361 | 16.228 | 1.00 | 0.00 | C |
| ATOM | 359 | HD1 | PRO | A | 43 | 37.450 | 44.246 | 15.667 | 1.00 | 0.00 | H |
| ATOM | 360 | HD2 | PRO | A | 43 | 37.600 | 42.469 | 15.620 | 1.00 | 0.00 | H |
| ATOM | 361 | CG | PRO | A | 43 | 39.200 | 43.488 | 16.644 | 1.00 | 0.00 | C |
| ATOM | 362 | HG1 | PRO | A | 43 | 39.778 | 44.067 | 15.924 | 1.00 | 0.00 | H |
| ATOM | 363 | HG2 | PRO | A | 43 | 39.633 | 42.497 | 16.782 | 1.00 | 0.00 | H |
| ATOM | 364 | CB | PRO | A | 43 | 39.108 | 44.192 | 17.991 | 1.00 | 0.00 | C |
| ATOM | 365 | HB1 | PRO | A | 43 | 38.980 | 45.264 | 17.831 | 1.00 | 0.00 | H |
| ATOM | 366 | HB2 | PRO | A | 43 | 39.981 | 43.998 | 18.612 | 1.00 | 0.00 | H |
| ATOM | 367 | CA | PRO | A | 43 | 37.842 | 43.585 | 18.599 | 1.00 | 0.00 | C |
| ATOM | 368 | HA | PRO | A | 43 | 38.100 | 42.646 | 19.088 | 1.00 | 0.00 | H |
| ATOM | 369 | C | PRO | A | 43 | 37.239 | 44.527 | 19.641 | 1.00 | 0.00 | C |
| ATOM | 370 | O | PRO | A | 43 | 37.417 | 44.301 | 20.835 | 1.00 | 0.00 | O |
| ATOM | 371 | N | LYS | A | 44 | 36.482 | 45.543 | 19.205 | 1.00 | 0.00 | N |
| ATOM | 372 | H | LYS | A | 44 | 36.429 | 45.724 | 18.210 | 1.00 | 0.00 | H |
| ATOM | 373 | CA | LYS | A | 44 | 35.808 | 46.482 | 20.106 | 1.00 | 0.00 | C |
| ATOM | 374 | HA | LYS | A | 44 | 36.578 | 46.983 | 20.695 | 1.00 | 0.00 | H |
| ATOM | 375 | CB | LYS | A | 44 | 35.079 | 47.553 | 19.269 | 1.00 | 0.00 | C |

| | | | | | | | | | | | |
|---|---|---|---|---|---|---|---|---|---|---|---|
| ATOM | 376 | HB1 | LYS | A | 44 | 34.749 | 48.349 | 19.940 | 1.00 | 0.00 | H |
| ATOM | 377 | HB2 | LYS | A | 44 | 35.776 | 47.986 | 18.550 | 1.00 | 0.00 | H |
| ATOM | 378 | CG | LYS | A | 44 | 33.854 | 46.994 | 18.527 | 1.00 | 0.00 | C |
| ATOM | 379 | HG1 | LYS | A | 44 | 34.171 | 46.205 | 17.845 | 1.00 | 0.00 | H |
| ATOM | 380 | HG2 | LYS | A | 44 | 33.165 | 46.577 | 19.258 | 1.00 | 0.00 | H |
| ATOM | 381 | CD | LYS | A | 44 | 33.105 | 48.071 | 17.736 | 1.00 | 0.00 | C |
| ATOM | 382 | HD1 | LYS | A | 44 | 32.779 | 48.863 | 18.414 | 1.00 | 0.00 | H |
| ATOM | 383 | HD2 | LYS | A | 44 | 33.781 | 48.505 | 16.998 | 1.00 | 0.00 | H |
| ATOM | 384 | CE | LYS | A | 44 | 31.885 | 47.486 | 17.010 | 1.00 | 0.00 | C |
| ATOM | 385 | HE1 | LYS | A | 44 | 31.403 | 48.284 | 16.443 | 1.00 | 0.00 | H |
| ATOM | 386 | HE2 | LYS | A | 44 | 32.214 | 46.721 | 16.301 | 1.00 | 0.00 | H |
| ATOM | 387 | NZ | LYS | A | 44 | 30.900 | 46.889 | 17.943 | 1.00 | 0.00 | N |
| ATOM | 388 | HZ1 | LYS | A | 44 | 30.053 | 46.636 | 17.446 | 1.00 | 0.00 | H |
| ATOM | 389 | HZ2 | LYS | A | 44 | 31.263 | 46.037 | 18.366 | 1.00 | 0.00 | H |
| ATOM | 390 | HZ3 | LYS | A | 44 | 30.643 | 47.535 | 18.684 | 1.00 | 0.00 | H |
| ATOM | 391 | C | LYS | A | 44 | 34.858 | 45.819 | 21.125 | 1.00 | 0.00 | C |
| ATOM | 392 | O | LYS | A | 44 | 34.672 | 46.371 | 22.206 | 1.00 | 0.00 | O |
| ATOM | 393 | N | ASP | A | 45 | 34.263 | 44.662 | 20.809 | 1.00 | 0.00 | N |
| ATOM | 394 | H | ASP | A | 45 | 34.518 | 44.220 | 19.931 | 1.00 | 0.00 | H |
| ATOM | 395 | CA | ASP | A | 45 | 33.254 | 43.988 | 21.641 | 1.00 | 0.00 | C |
| ATOM | 396 | HA | ASP | A | 45 | 32.686 | 44.747 | 22.184 | 1.00 | 0.00 | H |
| ATOM | 397 | CB | ASP | A | 45 | 32.261 | 43.227 | 20.742 | 1.00 | 0.00 | C |
| ATOM | 398 | HB1 | ASP | A | 45 | 32.800 | 42.477 | 20.160 | 1.00 | 0.00 | H |
| ATOM | 399 | HB2 | ASP | A | 45 | 31.544 | 42.702 | 21.374 | 1.00 | 0.00 | H |
| ATOM | 400 | CG | ASP | A | 45 | 31.486 | 44.152 | 19.798 | 1.00 | 0.00 | C |
| ATOM | 401 | OD1 | ASP | A | 45 | 30.485 | 44.780 | 20.219 | 1.00 | 0.00 | O |
| ATOM | 402 | OD2 | ASP | A | 45 | 31.886 | 44.277 | 18.618 | 1.00 | 0.00 | O |
| ATOM | 403 | C | ASP | A | 45 | 33.872 | 43.051 | 22.700 | 1.00 | 0.00 | C |
| ATOM | 404 | O | ASP | A | 45 | 33.145 | 42.479 | 23.515 | 1.00 | 0.00 | O |
| ATOM | 405 | N | LEU | A | 46 | 35.202 | 42.894 | 22.721 | 1.00 | 0.00 | N |
| ATOM | 406 | H | LEU | A | 46 | 35.752 | 43.399 | 22.034 | 1.00 | 0.00 | H |
| ATOM | 407 | CA | LEU | A | 46 | 35.918 | 42.156 | 23.765 | 1.00 | 0.00 | C |
| ATOM | 408 | HA | LEU | A | 46 | 35.328 | 41.283 | 24.042 | 1.00 | 0.00 | H |
| ATOM | 409 | CB | LEU | A | 46 | 37.286 | 41.685 | 23.232 | 1.00 | 0.00 | C |
| ATOM | 410 | HB1 | LEU | A | 46 | 37.816 | 42.552 | 22.833 | 1.00 | 0.00 | H |
| ATOM | 411 | HB2 | LEU | A | 46 | 37.873 | 41.295 | 24.064 | 1.00 | 0.00 | H |
| ATOM | 412 | CG | LEU | A | 46 | 37.189 | 40.595 | 22.142 | 1.00 | 0.00 | C |
| ATOM | 413 | HG | LEU | A | 46 | 36.428 | 40.882 | 21.415 | 1.00 | 0.00 | H |
| ATOM | 414 | CD1 | LEU | A | 46 | 38.518 | 40.444 | 21.390 | 1.00 | 0.00 | C |
| ATOM | 415 | HD11 | LEU | A | 46 | 38.399 | 39.733 | 20.572 | 1.00 | 0.00 | H |
| ATOM | 416 | HD12 | LEU | A | 46 | 38.825 | 41.406 | 20.978 | 1.00 | 0.00 | H |
| ATOM | 417 | HD13 | LEU | A | 46 | 39.293 | 40.079 | 22.064 | 1.00 | 0.00 | H |
| ATOM | 418 | CD2 | LEU | A | 46 | 36.814 | 39.232 | 22.748 | 1.00 | 0.00 | C |
| ATOM | 419 | HD21 | LEU | A | 46 | 36.731 | 38.486 | 21.958 | 1.00 | 0.00 | H |
| ATOM | 420 | HD22 | LEU | A | 46 | 37.579 | 38.913 | 23.458 | 1.00 | 0.00 | H |
| ATOM | 421 | HD23 | LEU | A | 46 | 35.859 | 39.296 | 23.267 | 1.00 | 0.00 | H |
| ATOM | 422 | C | LEU | A | 46 | 36.069 | 43.014 | 25.030 | 1.00 | 0.00 | C |
| ATOM | 423 | O | LEU | A | 46 | 36.418 | 44.193 | 24.967 | 1.00 | 0.00 | O |
| ATOM | 424 | N | ASP | A | 47 | 35.813 | 42.423 | 26.193 | 1.00 | 0.00 | N |
| ATOM | 425 | H | ASP | A | 47 | 35.514 | 41.454 | 26.172 | 1.00 | 0.00 | H |
| ATOM | 426 | CA | ASP | A | 47 | 35.988 | 43.032 | 27.517 | 1.00 | 0.00 | C |
| ATOM | 427 | HA | ASP | A | 47 | 36.116 | 44.109 | 27.418 | 1.00 | 0.00 | H |
| ATOM | 428 | CB | ASP | A | 47 | 34.703 | 42.786 | 28.327 | 1.00 | 0.00 | C |
| ATOM | 429 | HB1 | ASP | A | 47 | 33.868 | 43.234 | 27.785 | 1.00 | 0.00 | H |

| | | | | | | | | | | | |
|---|---|---|---|---|---|---|---|---|---|---|---|
| ATOM | 430 | HB2 | ASP | A | 47 | 34.523 | 41.713 | 28.372 | 1.00 | 0.00 | H |
| ATOM | 431 | CG | ASP | A | 47 | 34.697 | 43.351 | 29.756 | 1.00 | 0.00 | C |
| ATOM | 432 | OD1 | ASP | A | 47 | 35.737 | 43.814 | 30.272 | 1.00 | 0.00 | O |
| ATOM | 433 | OD2 | ASP | A | 47 | 33.604 | 43.363 | 30.366 | 1.00 | 0.00 | O |
| ATOM | 434 | C | ASP | A | 47 | 37.217 | 42.407 | 28.207 | 1.00 | 0.00 | C |
| ATOM | 435 | O | ASP | A | 47 | 37.253 | 41.180 | 28.338 | 1.00 | 0.00 | O |
| ATOM | 436 | N | PRO | A | 48 | 38.210 | 43.197 | 28.666 | 1.00 | 0.00 | N |
| ATOM | 437 | CD | PRO | A | 48 | 38.308 | 44.646 | 28.531 | 1.00 | 0.00 | C |
| ATOM | 438 | HD1 | PRO | A | 48 | 37.407 | 45.148 | 28.881 | 1.00 | 0.00 | H |
| ATOM | 439 | HD2 | PRO | A | 48 | 38.497 | 44.900 | 27.487 | 1.00 | 0.00 | H |
| ATOM | 440 | CG | PRO | A | 48 | 39.498 | 45.068 | 29.389 | 1.00 | 0.00 | C |
| ATOM | 441 | HG1 | PRO | A | 48 | 39.166 | 45.250 | 30.410 | 1.00 | 0.00 | H |
| ATOM | 442 | HG2 | PRO | A | 48 | 39.999 | 45.948 | 28.984 | 1.00 | 0.00 | H |
| ATOM | 443 | CB | PRO | A | 48 | 40.397 | 43.836 | 29.361 | 1.00 | 0.00 | C |
| ATOM | 444 | HB1 | PRO | A | 48 | 41.041 | 43.795 | 30.237 | 1.00 | 0.00 | H |
| ATOM | 445 | HB2 | PRO | A | 48 | 41.002 | 43.851 | 28.454 | 1.00 | 0.00 | H |
| ATOM | 446 | CA | PRO | A | 48 | 39.412 | 42.662 | 29.297 | 1.00 | 0.00 | C |
| ATOM | 447 | HA | PRO | A | 48 | 39.836 | 41.893 | 28.651 | 1.00 | 0.00 | H |
| ATOM | 448 | C | PRO | A | 48 | 39.168 | 42.017 | 30.674 | 1.00 | 0.00 | C |
| ATOM | 449 | O | PRO | A | 48 | 40.068 | 41.345 | 31.166 | 1.00 | 0.00 | O |
| ATOM | 450 | N | SER | A | 49 | 37.989 | 42.165 | 31.295 | 1.00 | 0.00 | N |
| ATOM | 451 | H | SER | A | 49 | 37.290 | 42.793 | 30.895 | 1.00 | 0.00 | H |
| ATOM | 452 | CA | SER | A | 49 | 37.671 | 41.542 | 32.598 | 1.00 | 0.00 | C |
| ATOM | 453 | HA | SER | A | 49 | 38.390 | 40.744 | 32.790 | 1.00 | 0.00 | H |
| ATOM | 454 | CB | SER | A | 49 | 37.837 | 42.567 | 33.728 | 1.00 | 0.00 | C |
| ATOM | 455 | HB1 | SER | A | 49 | 37.714 | 42.067 | 34.687 | 1.00 | 0.00 | H |
| ATOM | 456 | HB2 | SER | A | 49 | 38.844 | 42.986 | 33.685 | 1.00 | 0.00 | H |
| ATOM | 457 | OG | SER | A | 49 | 36.889 | 43.612 | 33.629 | 1.00 | 0.00 | O |
| ATOM | 458 | HG | SER | A | 49 | 35.996 | 43.222 | 33.638 | 1.00 | 0.00 | H |
| ATOM | 459 | C | SER | A | 49 | 36.292 | 40.851 | 32.648 | 1.00 | 0.00 | C |
| ATOM | 460 | O | SER | A | 49 | 35.603 | 40.862 | 33.675 | 1.00 | 0.00 | O |
| ATOM | 461 | N | LEU | A | 50 | 35.858 | 40.250 | 31.533 | 1.00 | 0.00 | N |
| ATOM | 462 | H | LEU | A | 50 | 36.439 | 40.300 | 30.705 | 1.00 | 0.00 | H |
| ATOM | 463 | CA | LEU | A | 50 | 34.710 | 39.335 | 31.531 | 1.00 | 0.00 | C |
| ATOM | 464 | HA | LEU | A | 50 | 34.001 | 39.657 | 32.290 | 1.00 | 0.00 | H |
| ATOM | 465 | CB | LEU | A | 50 | 34.009 | 39.411 | 30.164 | 1.00 | 0.00 | C |
| ATOM | 466 | HB1 | LEU | A | 50 | 33.708 | 40.445 | 30.001 | 1.00 | 0.00 | H |
| ATOM | 467 | HB2 | LEU | A | 50 | 34.720 | 39.130 | 29.385 | 1.00 | 0.00 | H |
| ATOM | 468 | CG | LEU | A | 50 | 32.752 | 38.527 | 30.034 | 1.00 | 0.00 | C |
| ATOM | 469 | HG | LEU | A | 50 | 33.027 | 37.488 | 30.215 | 1.00 | 0.00 | H |
| ATOM | 470 | CD1 | LEU | A | 50 | 31.647 | 38.925 | 31.028 | 1.00 | 0.00 | C |
| ATOM | 471 | HD11 | LEU | A | 50 | 30.749 | 38.339 | 30.836 | 1.00 | 0.00 | H |
| ATOM | 472 | HD12 | LEU | A | 50 | 31.969 | 38.728 | 32.050 | 1.00 | 0.00 | H |
| ATOM | 473 | HD13 | LEU | A | 50 | 31.409 | 39.984 | 30.921 | 1.00 | 0.00 | H |
| ATOM | 474 | CD2 | LEU | A | 50 | 32.219 | 38.638 | 28.600 | 1.00 | 0.00 | C |
| ATOM | 475 | HD21 | LEU | A | 50 | 31.301 | 38.060 | 28.494 | 1.00 | 0.00 | H |
| ATOM | 476 | HD22 | LEU | A | 50 | 32.010 | 39.680 | 28.360 | 1.00 | 0.00 | H |
| ATOM | 477 | HD23 | LEU | A | 50 | 32.956 | 38.243 | 27.901 | 1.00 | 0.00 | H |
| ATOM | 478 | C | LEU | A | 50 | 35.149 | 37.912 | 31.905 | 1.00 | 0.00 | C |
| ATOM | 479 | O | LEU | A | 50 | 34.559 | 37.297 | 32.791 | 1.00 | 0.00 | O |
| ATOM | 480 | N | CYS | A | 51 | 36.198 | 37.399 | 31.264 | 1.00 | 0.00 | N |
| ATOM | 481 | H | CYS | A | 51 | 36.645 | 37.968 | 30.551 | 1.00 | 0.00 | H |
| ATOM | 482 | CA | CYS | A | 51 | 36.724 | 36.041 | 31.445 | 1.00 | 0.00 | C |
| ATOM | 483 | HA | CYS | A | 51 | 36.038 | 35.463 | 32.065 | 1.00 | 0.00 | H |

| | | | | | | | | | | | |
|---|---|---|---|---|---|---|---|---|---|---|---|
| ATOM | 484 | CB | CYS | A | 51 | 36.779 | 35.375 | 30.063 | 1.00 | 0.00 | C |
| ATOM | 485 | HB1 | CYS | A | 51 | 37.540 | 35.870 | 29.457 | 1.00 | 0.00 | H |
| ATOM | 486 | HB2 | CYS | A | 51 | 37.067 | 34.330 | 30.175 | 1.00 | 0.00 | H |
| ATOM | 487 | SG | CYS | A | 51 | 35.197 | 35.453 | 29.168 | 1.00 | 0.00 | S |
| ATOM | 488 | C | CYS | A | 51 | 38.094 | 36.045 | 32.149 | 1.00 | 0.00 | C |
| ATOM | 489 | O | CYS | A | 51 | 38.732 | 37.094 | 32.288 | 1.00 | 0.00 | O |
| ATOM | 490 | N | THR | A | 52 | 38.579 | 34.873 | 32.587 | 1.00 | 0.00 | N |
| ATOM | 491 | H | THR | A | 52 | 38.022 | 34.037 | 32.447 | 1.00 | 0.00 | H |
| ATOM | 492 | CA | THR | A | 52 | 39.918 | 34.731 | 33.199 | 1.00 | 0.00 | C |
| ATOM | 493 | HA | THR | A | 52 | 40.266 | 35.723 | 33.487 | 1.00 | 0.00 | H |
| ATOM | 494 | CB | THR | A | 52 | 39.917 | 33.909 | 34.499 | 1.00 | 0.00 | C |
| ATOM | 495 | HB | THR | A | 52 | 40.780 | 34.232 | 35.082 | 1.00 | 0.00 | H |
| ATOM | 496 | CG2 | THR | A | 52 | 38.678 | 34.137 | 35.356 | 1.00 | 0.00 | C |
| ATOM | 497 | HG21 | THR | A | 52 | 38.775 | 33.608 | 36.301 | 1.00 | 0.00 | H |
| ATOM | 498 | HG22 | THR | A | 52 | 38.568 | 35.201 | 35.553 | 1.00 | 0.00 | H |
| ATOM | 499 | HG23 | THR | A | 52 | 37.788 | 33.780 | 34.846 | 1.00 | 0.00 | H |
| ATOM | 500 | OG1 | THR | A | 52 | 40.051 | 32.521 | 34.288 | 1.00 | 0.00 | O |
| ATOM | 501 | HG1 | THR | A | 52 | 40.621 | 32.220 | 35.023 | 1.00 | 0.00 | H |
| ATOM | 502 | C | THR | A | 52 | 40.945 | 34.219 | 32.204 | 1.00 | 0.00 | C |
| ATOM | 503 | O | THR | A | 52 | 42.117 | 34.578 | 32.308 | 1.00 | 0.00 | O |
| ATOM | 504 | N | HID | A | 53 | 40.528 | 33.393 | 31.248 | 1.00 | 0.00 | N |
| ATOM | 505 | H | HID | A | 53 | 39.549 | 33.119 | 31.235 | 1.00 | 0.00 | H |
| ATOM | 506 | CA | HID | A | 53 | 41.343 | 32.941 | 30.121 | 1.00 | 0.00 | C |
| ATOM | 507 | HA | HID | A | 53 | 42.270 | 33.513 | 30.076 | 1.00 | 0.00 | H |
| ATOM | 508 | CB | HID | A | 53 | 41.691 | 31.449 | 30.287 | 1.00 | 0.00 | C |
| ATOM | 509 | HB1 | HID | A | 53 | 40.768 | 30.886 | 30.420 | 1.00 | 0.00 | H |
| ATOM | 510 | HB2 | HID | A | 53 | 42.156 | 31.102 | 29.363 | 1.00 | 0.00 | H |
| ATOM | 511 | CG | HID | A | 53 | 42.630 | 31.125 | 31.433 | 1.00 | 0.00 | C |
| ATOM | 512 | ND1 | HID | A | 53 | 42.530 | 31.573 | 32.730 | 1.00 | 0.00 | N |
| ATOM | 513 | HD1 | HID | A | 53 | 41.818 | 32.199 | 33.081 | 1.00 | 0.00 | H |
| ATOM | 514 | CE1 | HID | A | 53 | 43.572 | 31.099 | 33.428 | 1.00 | 0.00 | C |
| ATOM | 515 | HE1 | HID | A | 53 | 43.771 | 31.319 | 34.470 | 1.00 | 0.00 | H |
| ATOM | 516 | NE2 | HID | A | 53 | 44.337 | 30.315 | 32.650 | 1.00 | 0.00 | N |
| ATOM | 517 | CD2 | HID | A | 53 | 43.742 | 30.327 | 31.382 | 1.00 | 0.00 | C |
| ATOM | 518 | HD2 | HID | A | 53 | 44.104 | 29.802 | 30.510 | 1.00 | 0.00 | H |
| ATOM | 519 | C | HID | A | 53 | 40.566 | 33.186 | 28.813 | 1.00 | 0.00 | C |
| ATOM | 520 | O | HID | A | 53 | 39.366 | 32.911 | 28.739 | 1.00 | 0.00 | O |
| ATOM | 521 | N | LEU | A | 54 | 41.249 | 33.696 | 27.787 | 1.00 | 0.00 | N |
| ATOM | 522 | H | LEU | A | 54 | 42.240 | 33.880 | 27.911 | 1.00 | 0.00 | H |
| ATOM | 523 | CA | LEU | A | 54 | 40.693 | 33.993 | 26.465 | 1.00 | 0.00 | C |
| ATOM | 524 | HA | LEU | A | 54 | 39.630 | 33.750 | 26.443 | 1.00 | 0.00 | H |
| ATOM | 525 | CB | LEU | A | 54 | 40.874 | 35.505 | 26.219 | 1.00 | 0.00 | C |
| ATOM | 526 | HB1 | LEU | A | 54 | 40.313 | 36.041 | 26.986 | 1.00 | 0.00 | H |
| ATOM | 527 | HB2 | LEU | A | 54 | 41.930 | 35.750 | 26.349 | 1.00 | 0.00 | H |
| ATOM | 528 | CG | LEU | A | 54 | 40.428 | 36.032 | 24.841 | 1.00 | 0.00 | C |
| ATOM | 529 | HG | LEU | A | 54 | 41.011 | 35.535 | 24.067 | 1.00 | 0.00 | H |
| ATOM | 530 | CD1 | LEU | A | 54 | 38.940 | 35.776 | 24.568 | 1.00 | 0.00 | C |
| ATOM | 531 | HD11 | LEU | A | 54 | 38.646 | 36.263 | 23.637 | 1.00 | 0.00 | H |
| ATOM | 532 | HD12 | LEU | A | 54 | 38.758 | 34.709 | 24.464 | 1.00 | 0.00 | H |
| ATOM | 533 | HD13 | LEU | A | 54 | 38.337 | 36.171 | 25.386 | 1.00 | 0.00 | H |
| ATOM | 534 | CD2 | LEU | A | 54 | 40.705 | 37.542 | 24.769 | 1.00 | 0.00 | C |
| ATOM | 535 | HD21 | LEU | A | 54 | 40.403 | 37.931 | 23.796 | 1.00 | 0.00 | H |
| ATOM | 536 | HD22 | LEU | A | 54 | 40.147 | 38.065 | 25.546 | 1.00 | 0.00 | H |
| ATOM | 537 | HD23 | LEU | A | 54 | 41.771 | 37.731 | 24.902 | 1.00 | 0.00 | H |

| | | | | | | | | | | | |
|---|---|---|---|---|---|---|---|---|---|---|---|
| ATOM | 538 | C | LEU | A | 54 | 41.418 | 33.127 | 25.429 | 1.00 | 0.00 | C |
| ATOM | 539 | O | LEU | A | 54 | 42.647 | 33.146 | 25.374 | 1.00 | 0.00 | O |
| ATOM | 540 | N | ILE | A | 55 | 40.686 | 32.351 | 24.628 | 1.00 | 0.00 | N |
| ATOM | 541 | H | ILE | A | 55 | 39.672 | 32.388 | 24.707 | 1.00 | 0.00 | H |
| ATOM | 542 | CA | ILE | A | 55 | 41.272 | 31.417 | 23.652 | 1.00 | 0.00 | C |
| ATOM | 543 | HA | ILE | A | 55 | 42.351 | 31.489 | 23.725 | 1.00 | 0.00 | H |
| ATOM | 544 | CB | ILE | A | 55 | 40.916 | 29.942 | 23.967 | 1.00 | 0.00 | C |
| ATOM | 545 | HB | ILE | A | 55 | 39.859 | 29.780 | 23.756 | 1.00 | 0.00 | H |
| ATOM | 546 | CG2 | ILE | A | 55 | 41.747 | 29.032 | 23.039 | 1.00 | 0.00 | C |
| ATOM | 547 | HG21 | ILE | A | 55 | 41.471 | 27.988 | 23.182 | 1.00 | 0.00 | H |
| ATOM | 548 | HG22 | ILE | A | 55 | 41.556 | 29.274 | 21.995 | 1.00 | 0.00 | H |
| ATOM | 549 | HG23 | ILE | A | 55 | 42.811 | 29.148 | 23.246 | 1.00 | 0.00 | H |
| ATOM | 550 | CG1 | ILE | A | 55 | 41.162 | 29.618 | 25.464 | 1.00 | 0.00 | C |
| ATOM | 551 | HG11 | ILE | A | 55 | 42.094 | 30.079 | 25.783 | 1.00 | 0.00 | H |
| ATOM | 552 | HG12 | ILE | A | 55 | 40.361 | 30.071 | 26.049 | 1.00 | 0.00 | H |
| ATOM | 553 | CD | ILE | A | 55 | 41.222 | 28.127 | 25.831 | 1.00 | 0.00 | C |
| ATOM | 554 | HD1 | ILE | A | 55 | 41.352 | 28.024 | 26.907 | 1.00 | 0.00 | H |
| ATOM | 555 | HD2 | ILE | A | 55 | 40.301 | 27.629 | 25.528 | 1.00 | 0.00 | H |
| ATOM | 556 | HD3 | ILE | A | 55 | 42.076 | 27.647 | 25.353 | 1.00 | 0.00 | H |
| ATOM | 557 | C | ILE | A | 55 | 40.902 | 31.819 | 22.221 | 1.00 | 0.00 | C |
| ATOM | 558 | O | ILE | A | 55 | 39.732 | 32.003 | 21.900 | 1.00 | 0.00 | O |
| ATOM | 559 | N | TYR | A | 56 | 41.905 | 31.948 | 21.355 | 1.00 | 0.00 | N |
| ATOM | 560 | H | TYR | A | 56 | 42.845 | 31.777 | 21.698 | 1.00 | 0.00 | H |
| ATOM | 561 | CA | TYR | A | 56 | 41.763 | 32.348 | 19.953 | 1.00 | 0.00 | C |
| ATOM | 562 | HA | TYR | A | 56 | 40.904 | 33.013 | 19.855 | 1.00 | 0.00 | H |
| ATOM | 563 | CB | TYR | A | 56 | 43.035 | 33.133 | 19.581 | 1.00 | 0.00 | C |
| ATOM | 564 | HB1 | TYR | A | 56 | 43.246 | 33.831 | 20.390 | 1.00 | 0.00 | H |
| ATOM | 565 | HB2 | TYR | A | 56 | 43.882 | 32.450 | 19.547 | 1.00 | 0.00 | H |
| ATOM | 566 | CG | TYR | A | 56 | 42.996 | 33.926 | 18.286 | 1.00 | 0.00 | C |
| ATOM | 567 | CD1 | TYR | A | 56 | 43.049 | 33.284 | 17.030 | 1.00 | 0.00 | C |
| ATOM | 568 | HD1 | TYR | A | 56 | 43.092 | 32.206 | 16.970 | 1.00 | 0.00 | H |
| ATOM | 569 | CE1 | TYR | A | 56 | 43.089 | 34.046 | 15.844 | 1.00 | 0.00 | C |
| ATOM | 570 | HE1 | TYR | A | 56 | 43.147 | 33.555 | 14.884 | 1.00 | 0.00 | H |
| ATOM | 571 | CZ | TYR | A | 56 | 43.078 | 35.459 | 15.914 | 1.00 | 0.00 | C |
| ATOM | 572 | OH | TYR | A | 56 | 43.126 | 36.222 | 14.792 | 1.00 | 0.00 | O |
| ATOM | 573 | HH | TYR | A | 56 | 43.285 | 35.719 | 13.972 | 1.00 | 0.00 | H |
| ATOM | 574 | CE2 | TYR | A | 56 | 43.031 | 36.098 | 17.170 | 1.00 | 0.00 | C |
| ATOM | 575 | HE2 | TYR | A | 56 | 43.044 | 37.176 | 17.223 | 1.00 | 0.00 | H |
| ATOM | 576 | CD2 | TYR | A | 56 | 42.993 | 35.333 | 18.346 | 1.00 | 0.00 | C |
| ATOM | 577 | HD2 | TYR | A | 56 | 42.988 | 35.837 | 19.299 | 1.00 | 0.00 | H |
| ATOM | 578 | C | TYR | A | 56 | 41.546 | 31.125 | 19.039 | 1.00 | 0.00 | C |
| ATOM | 579 | O | TYR | A | 56 | 42.469 | 30.333 | 18.837 | 1.00 | 0.00 | O |
| ATOM | 580 | N | ALA | A | 57 | 40.348 | 30.971 | 18.466 | 1.00 | 0.00 | N |
| ATOM | 581 | H | ALA | A | 57 | 39.612 | 31.631 | 18.701 | 1.00 | 0.00 | H |
| ATOM | 582 | CA | ALA | A | 57 | 40.020 | 29.955 | 17.463 | 1.00 | 0.00 | C |
| ATOM | 583 | HA | ALA | A | 57 | 40.708 | 29.121 | 17.558 | 1.00 | 0.00 | H |
| ATOM | 584 | CB | ALA | A | 57 | 38.619 | 29.406 | 17.764 | 1.00 | 0.00 | C |
| ATOM | 585 | HB1 | ALA | A | 57 | 38.368 | 28.626 | 17.047 | 1.00 | 0.00 | H |
| ATOM | 586 | HB2 | ALA | A | 57 | 38.590 | 28.984 | 18.769 | 1.00 | 0.00 | H |
| ATOM | 587 | HB3 | ALA | A | 57 | 37.877 | 30.201 | 17.687 | 1.00 | 0.00 | H |
| ATOM | 588 | C | ALA | A | 57 | 40.105 | 30.552 | 16.038 | 1.00 | 0.00 | C |
| ATOM | 589 | O | ALA | A | 57 | 39.402 | 31.516 | 15.740 | 1.00 | 0.00 | O |
| ATOM | 590 | N | PHE | A | 58 | 40.925 | 30.040 | 15.113 | 1.00 | 0.00 | N |
| ATOM | 591 | H | PHE | A | 58 | 40.899 | 30.509 | 14.219 | 1.00 | 0.00 | H |

| | | | | | | | | | | | |
|---|---|---|---|---|---|---|---|---|---|---|---|
| ATOM | 592 | CA | PHE | A | 58 | 41.949 | 28.994 | 15.252 | 1.00 | 0.00 | C |
| ATOM | 593 | HA | PHE | A | 58 | 42.210 | 28.838 | 16.296 | 1.00 | 0.00 | H |
| ATOM | 594 | CB | PHE | A | 58 | 41.446 | 27.667 | 14.647 | 1.00 | 0.00 | C |
| ATOM | 595 | HB1 | PHE | A | 58 | 41.150 | 27.848 | 13.612 | 1.00 | 0.00 | H |
| ATOM | 596 | HB2 | PHE | A | 58 | 42.278 | 26.961 | 14.612 | 1.00 | 0.00 | H |
| ATOM | 597 | CG | PHE | A | 58 | 40.315 | 26.968 | 15.377 | 1.00 | 0.00 | C |
| ATOM | 598 | CD1 | PHE | A | 58 | 40.538 | 26.398 | 16.644 | 1.00 | 0.00 | C |
| ATOM | 599 | HD1 | PHE | A | 58 | 41.511 | 26.471 | 17.101 | 1.00 | 0.00 | H |
| ATOM | 600 | CE1 | PHE | A | 58 | 39.502 | 25.719 | 17.309 | 1.00 | 0.00 | C |
| ATOM | 601 | HE1 | PHE | A | 58 | 39.672 | 25.280 | 18.282 | 1.00 | 0.00 | H |
| ATOM | 602 | CZ | PHE | A | 58 | 38.242 | 25.593 | 16.702 | 1.00 | 0.00 | C |
| ATOM | 603 | HZ | PHE | A | 58 | 37.450 | 25.059 | 17.210 | 1.00 | 0.00 | H |
| ATOM | 604 | CE2 | PHE | A | 58 | 38.021 | 26.137 | 15.425 | 1.00 | 0.00 | C |
| ATOM | 605 | HE2 | PHE | A | 58 | 37.060 | 26.012 | 14.947 | 1.00 | 0.00 | H |
| ATOM | 606 | CD2 | PHE | A | 58 | 39.058 | 26.819 | 14.763 | 1.00 | 0.00 | C |
| ATOM | 607 | HD2 | PHE | A | 58 | 38.895 | 27.214 | 13.772 | 1.00 | 0.00 | H |
| ATOM | 608 | C | PHE | A | 58 | 43.250 | 29.393 | 14.537 | 1.00 | 0.00 | C |
| ATOM | 609 | O | PHE | A | 58 | 43.256 | 30.240 | 13.640 | 1.00 | 0.00 | O |
| ATOM | 610 | N | ALA | A | 59 | 44.343 | 28.727 | 14.911 | 1.00 | 0.00 | N |
| ATOM | 611 | H | ALA | A | 59 | 44.249 | 28.106 | 15.709 | 1.00 | 0.00 | H |
| ATOM | 612 | CA | ALA | A | 59 | 45.572 | 28.601 | 14.134 | 1.00 | 0.00 | C |
| ATOM | 613 | HA | ALA | A | 59 | 45.672 | 29.455 | 13.469 | 1.00 | 0.00 | H |
| ATOM | 614 | CB | ALA | A | 59 | 46.761 | 28.599 | 15.095 | 1.00 | 0.00 | C |
| ATOM | 615 | HB1 | ALA | A | 59 | 47.688 | 28.498 | 14.530 | 1.00 | 0.00 | H |
| ATOM | 616 | HB2 | ALA | A | 59 | 46.783 | 29.543 | 15.630 | 1.00 | 0.00 | H |
| ATOM | 617 | HB3 | ALA | A | 59 | 46.672 | 27.775 | 15.805 | 1.00 | 0.00 | H |
| ATOM | 618 | C | ALA | A | 59 | 45.563 | 27.333 | 13.269 | 1.00 | 0.00 | C |
| ATOM | 619 | O | ALA | A | 59 | 45.017 | 26.302 | 13.671 | 1.00 | 0.00 | O |
| ATOM | 620 | N | GLY | A | 60 | 46.219 | 27.420 | 12.108 | 1.00 | 0.00 | N |
| ATOM | 621 | H | GLY | A | 60 | 46.725 | 28.278 | 11.920 | 1.00 | 0.00 | H |
| ATOM | 622 | CA | GLY | A | 60 | 46.395 | 26.329 | 11.149 | 1.00 | 0.00 | C |
| ATOM | 623 | HA1 | GLY | A | 60 | 45.675 | 25.533 | 11.345 | 1.00 | 0.00 | H |
| ATOM | 624 | HA2 | GLY | A | 60 | 46.230 | 26.719 | 10.145 | 1.00 | 0.00 | H |
| ATOM | 625 | C | GLY | A | 60 | 47.803 | 25.732 | 11.187 | 1.00 | 0.00 | C |
| ATOM | 626 | O | GLY | A | 60 | 48.604 | 26.025 | 12.077 | 1.00 | 0.00 | O |
| ATOM | 627 | N | MET | A | 61 | 48.114 | 24.889 | 10.200 | 1.00 | 0.00 | N |
| ATOM | 628 | H | MET | A | 61 | 47.448 | 24.786 | 9.441 | 1.00 | 0.00 | H |
| ATOM | 629 | CA | MET | A | 61 | 49.387 | 24.163 | 10.108 | 1.00 | 0.00 | C |
| ATOM | 630 | HA | MET | A | 61 | 50.144 | 24.733 | 10.643 | 1.00 | 0.00 | H |
| ATOM | 631 | CB | MET | A | 61 | 49.227 | 22.795 | 10.788 | 1.00 | 0.00 | C |
| ATOM | 632 | HB1 | MET | A | 61 | 48.769 | 22.937 | 11.767 | 1.00 | 0.00 | H |
| ATOM | 633 | HB2 | MET | A | 61 | 48.562 | 22.177 | 10.185 | 1.00 | 0.00 | H |
| ATOM | 634 | CG | MET | A | 61 | 50.549 | 22.043 | 10.975 | 1.00 | 0.00 | C |
| ATOM | 635 | HG1 | MET | A | 61 | 51.125 | 22.092 | 10.051 | 1.00 | 0.00 | H |
| ATOM | 636 | HG2 | MET | A | 61 | 51.133 | 22.535 | 11.756 | 1.00 | 0.00 | H |
| ATOM | 637 | SD | MET | A | 61 | 50.341 | 20.291 | 11.394 | 1.00 | 0.00 | S |
| ATOM | 638 | CE | MET | A | 61 | 49.485 | 20.425 | 12.982 | 1.00 | 0.00 | C |
| ATOM | 639 | HE1 | MET | A | 61 | 49.356 | 19.429 | 13.403 | 1.00 | 0.00 | H |
| ATOM | 640 | HE2 | MET | A | 61 | 50.072 | 21.033 | 13.670 | 1.00 | 0.00 | H |
| ATOM | 641 | HE3 | MET | A | 61 | 48.501 | 20.873 | 12.845 | 1.00 | 0.00 | H |
| ATOM | 642 | C | MET | A | 61 | 49.834 | 24.000 | 8.651 | 1.00 | 0.00 | C |
| ATOM | 643 | O | MET | A | 61 | 49.085 | 23.458 | 7.830 | 1.00 | 0.00 | O |
| ATOM | 644 | N | THR | A | 62 | 51.070 | 24.399 | 8.333 | 1.00 | 0.00 | N |
| ATOM | 645 | H | THR | A | 62 | 51.644 | 24.781 | 9.083 | 1.00 | 0.00 | H |

| | | | | | | | | | | | |
|---|---|---|---|---|---|---|---|---|---|---|---|
| ATOM | 646 | CA | THR | A | 62 | 51.656 | 24.355 | 6.981 | 1.00 | 0.00 | C |
| ATOM | 647 | HA | THR | A | 62 | 51.033 | 23.760 | 6.320 | 1.00 | 0.00 | H |
| ATOM | 648 | CB | THR | A | 62 | 51.702 | 25.766 | 6.384 | 1.00 | 0.00 | C |
| ATOM | 649 | HB | THR | A | 62 | 52.339 | 26.412 | 6.989 | 1.00 | 0.00 | H |
| ATOM | 650 | CG2 | THR | A | 62 | 52.189 | 25.776 | 4.935 | 1.00 | 0.00 | C |
| ATOM | 651 | HG21 | THR | A | 62 | 52.135 | 26.791 | 4.541 | 1.00 | 0.00 | H |
| ATOM | 652 | HG22 | THR | A | 62 | 53.226 | 25.447 | 4.891 | 1.00 | 0.00 | H |
| ATOM | 653 | HG23 | THR | A | 62 | 51.573 | 25.120 | 4.322 | 1.00 | 0.00 | H |
| ATOM | 654 | OG1 | THR | A | 62 | 50.388 | 26.277 | 6.400 | 1.00 | 0.00 | O |
| ATOM | 655 | HG1 | THR | A | 62 | 50.264 | 26.654 | 7.288 | 1.00 | 0.00 | H |
| ATOM | 656 | C | THR | A | 62 | 53.033 | 23.706 | 7.055 | 1.00 | 0.00 | C |
| ATOM | 657 | O | THR | A | 62 | 53.894 | 24.179 | 7.792 | 1.00 | 0.00 | O |
| ATOM | 658 | N | ASN | A | 63 | 53.228 | 22.597 | 6.334 | 1.00 | 0.00 | N |
| ATOM | 659 | H | ASN | A | 63 | 52.461 | 22.263 | 5.768 | 1.00 | 0.00 | H |
| ATOM | 660 | CA | ASN | A | 63 | 54.445 | 21.761 | 6.376 | 1.00 | 0.00 | C |
| ATOM | 661 | HA | ASN | A | 63 | 54.218 | 20.839 | 5.841 | 1.00 | 0.00 | H |
| ATOM | 662 | CB | ASN | A | 63 | 55.620 | 22.446 | 5.651 | 1.00 | 0.00 | C |
| ATOM | 663 | HB1 | ASN | A | 63 | 55.893 | 23.355 | 6.185 | 1.00 | 0.00 | H |
| ATOM | 664 | HB2 | ASN | A | 63 | 56.486 | 21.784 | 5.661 | 1.00 | 0.00 | H |
| ATOM | 665 | CG | ASN | A | 63 | 55.306 | 22.775 | 4.202 | 1.00 | 0.00 | C |
| ATOM | 666 | OD1 | ASN | A | 63 | 54.923 | 21.909 | 3.419 | 1.00 | 0.00 | O |
| ATOM | 667 | ND2 | ASN | A | 63 | 55.447 | 24.028 | 3.817 | 1.00 | 0.00 | N |
| ATOM | 668 | HD21 | ASN | A | 63 | 55.832 | 24.710 | 4.460 | 1.00 | 0.00 | H |
| ATOM | 669 | HD22 | ASN | A | 63 | 55.131 | 24.286 | 2.892 | 1.00 | 0.00 | H |
| ATOM | 670 | C | ASN | A | 63 | 54.817 | 21.352 | 7.818 | 1.00 | 0.00 | C |
| ATOM | 671 | O | ASN | A | 63 | 55.942 | 21.567 | 8.271 | 1.00 | 0.00 | O |
| ATOM | 672 | N | HIS | A | 64 | 53.824 | 20.892 | 8.590 | 1.00 | 0.00 | N |
| ATOM | 673 | H | HIS | A | 64 | 52.937 | 20.711 | 8.136 | 1.00 | 0.00 | H |
| ATOM | 674 | CA | HIS | A | 64 | 53.891 | 20.570 | 10.028 | 1.00 | 0.00 | C |
| ATOM | 675 | HA | HIS | A | 64 | 52.929 | 20.141 | 10.305 | 1.00 | 0.00 | H |
| ATOM | 676 | CB | HIS | A | 64 | 54.953 | 19.494 | 10.313 | 1.00 | 0.00 | C |
| ATOM | 677 | HB1 | HIS | A | 64 | 55.941 | 19.928 | 10.204 | 1.00 | 0.00 | H |
| ATOM | 678 | HB2 | HIS | A | 64 | 54.857 | 19.179 | 11.352 | 1.00 | 0.00 | H |
| ATOM | 679 | CG | HIS | A | 64 | 54.891 | 18.262 | 9.457 | 1.00 | 0.00 | C |
| ATOM | 680 | ND1 | HIS | A | 64 | 53.773 | 17.454 | 9.248 | 1.00 | 0.00 | N |
| ATOM | 681 | CE1 | HIS | A | 64 | 54.225 | 16.389 | 8.567 | 1.00 | 0.00 | C |
| ATOM | 682 | HE1 | HIS | A | 64 | 53.632 | 15.531 | 8.293 | 1.00 | 0.00 | H |
| ATOM | 683 | NE2 | HIS | A | 64 | 55.529 | 16.520 | 8.284 | 1.00 | 0.00 | N |
| ATOM | 684 | HE2 | HIS | A | 64 | 56.099 | 15.827 | 7.800 | 1.00 | 0.00 | H |
| ATOM | 685 | CD2 | HIS | A | 64 | 55.970 | 17.685 | 8.861 | 1.00 | 0.00 | C |
| ATOM | 686 | HD2 | HIS | A | 64 | 56.982 | 18.053 | 8.885 | 1.00 | 0.00 | H |
| ATOM | 687 | C | HIS | A | 64 | 54.111 | 21.785 | 10.965 | 1.00 | 0.00 | C |
| ATOM | 688 | O | HIS | A | 64 | 53.916 | 21.651 | 12.169 | 1.00 | 0.00 | O |
| ATOM | 689 | N | GLN | A | 65 | 54.510 | 22.957 | 10.461 | 1.00 | 0.00 | N |
| ATOM | 690 | H | GLN | A | 65 | 54.602 | 23.044 | 9.457 | 1.00 | 0.00 | H |
| ATOM | 691 | CA | GLN | A | 65 | 54.744 | 24.158 | 11.271 | 1.00 | 0.00 | C |
| ATOM | 692 | HA | GLN | A | 65 | 55.168 | 23.856 | 12.228 | 1.00 | 0.00 | H |
| ATOM | 693 | CB | GLN | A | 65 | 55.742 | 25.102 | 10.570 | 1.00 | 0.00 | C |
| ATOM | 694 | HB1 | GLN | A | 65 | 55.890 | 25.985 | 11.194 | 1.00 | 0.00 | H |
| ATOM | 695 | HB2 | GLN | A | 65 | 55.320 | 25.433 | 9.622 | 1.00 | 0.00 | H |
| ATOM | 696 | CG | GLN | A | 65 | 57.113 | 24.453 | 10.311 | 1.00 | 0.00 | C |
| ATOM | 697 | HG1 | GLN | A | 65 | 56.994 | 23.588 | 9.664 | 1.00 | 0.00 | H |
| ATOM | 698 | HG2 | GLN | A | 65 | 57.529 | 24.120 | 11.263 | 1.00 | 0.00 | H |
| ATOM | 699 | CD | GLN | A | 65 | 58.104 | 25.400 | 9.635 | 1.00 | 0.00 | C |

| | | | | | | | | | | | |
|---|---|---|---|---|---|---|---|---|---|---|---|
| ATOM | 700 | OE1 | GLN | A | 65 | 57.935 | 25.807 | 8.489 | 1.00 | 0.00 | O |
| ATOM | 701 | NE2 | GLN | A | 65 | 59.174 | 25.772 | 10.302 | 1.00 | 0.00 | N |
| ATOM | 702 | HE21 | GLN | A | 65 | 59.401 | 25.355 | 11.195 | 1.00 | 0.00 | H |
| ATOM | 703 | HE22 | GLN | A | 65 | 59.822 | 26.424 | 9.876 | 1.00 | 0.00 | H |
| ATOM | 704 | C | GLN | A | 65 | 53.418 | 24.878 | 11.558 | 1.00 | 0.00 | C |
| ATOM | 705 | O | GLN | A | 65 | 52.488 | 24.806 | 10.755 | 1.00 | 0.00 | O |
| ATOM | 706 | N | LEU | A | 66 | 53.324 | 25.583 | 12.690 | 1.00 | 0.00 | N |
| ATOM | 707 | H | LEU | A | 66 | 54.128 | 25.615 | 13.306 | 1.00 | 0.00 | H |
| ATOM | 708 | CA | LEU | A | 66 | 52.147 | 26.386 | 13.058 | 1.00 | 0.00 | C |
| ATOM | 709 | HA | LEU | A | 66 | 51.254 | 25.770 | 12.934 | 1.00 | 0.00 | H |
| ATOM | 710 | CB | LEU | A | 66 | 52.303 | 26.781 | 14.542 | 1.00 | 0.00 | C |
| ATOM | 711 | HB1 | LEU | A | 66 | 52.533 | 25.888 | 15.126 | 1.00 | 0.00 | H |
| ATOM | 712 | HB2 | LEU | A | 66 | 53.161 | 27.452 | 14.619 | 1.00 | 0.00 | H |
| ATOM | 713 | CG | LEU | A | 66 | 51.079 | 27.473 | 15.178 | 1.00 | 0.00 | C |
| ATOM | 714 | HG | LEU | A | 66 | 50.760 | 28.305 | 14.549 | 1.00 | 0.00 | H |
| ATOM | 715 | CD1 | LEU | A | 66 | 49.904 | 26.510 | 15.382 | 1.00 | 0.00 | C |
| ATOM | 716 | HD11 | LEU | A | 66 | 49.067 | 27.040 | 15.832 | 1.00 | 0.00 | H |
| ATOM | 717 | HD12 | LEU | A | 66 | 49.584 | 26.090 | 14.432 | 1.00 | 0.00 | H |
| ATOM | 718 | HD13 | LEU | A | 66 | 50.200 | 25.705 | 16.054 | 1.00 | 0.00 | H |
| ATOM | 719 | CD2 | LEU | A | 66 | 51.465 | 28.028 | 16.551 | 1.00 | 0.00 | C |
| ATOM | 720 | HD21 | LEU | A | 66 | 50.637 | 28.602 | 16.960 | 1.00 | 0.00 | H |
| ATOM | 721 | HD22 | LEU | A | 66 | 51.714 | 27.217 | 17.235 | 1.00 | 0.00 | H |
| ATOM | 722 | HD23 | LEU | A | 66 | 52.323 | 28.683 | 16.444 | 1.00 | 0.00 | H |
| ATOM | 723 | C | LEU | A | 66 | 52.015 | 27.613 | 12.133 | 1.00 | 0.00 | C |
| ATOM | 724 | O | LEU | A | 66 | 53.025 | 28.244 | 11.812 | 1.00 | 0.00 | O |
| ATOM | 725 | N | SER | A | 67 | 50.803 | 27.992 | 11.714 | 1.00 | 0.00 | N |
| ATOM | 726 | H | SER | A | 67 | 49.992 | 27.437 | 11.976 | 1.00 | 0.00 | H |
| ATOM | 727 | CA | SER | A | 67 | 50.597 | 29.062 | 10.716 | 1.00 | 0.00 | C |
| ATOM | 728 | HA | SER | A | 67 | 51.361 | 29.824 | 10.872 | 1.00 | 0.00 | H |
| ATOM | 729 | CB | SER | A | 67 | 50.810 | 28.497 | 9.296 | 1.00 | 0.00 | C |
| ATOM | 730 | HB1 | SER | A | 67 | 50.685 | 29.300 | 8.571 | 1.00 | 0.00 | H |
| ATOM | 731 | HB2 | SER | A | 67 | 51.839 | 28.141 | 9.220 | 1.00 | 0.00 | H |
| ATOM | 732 | OG | SER | A | 67 | 49.942 | 27.418 | 8.967 | 1.00 | 0.00 | O |
| ATOM | 733 | HG | SER | A | 67 | 49.095 | 27.765 | 8.615 | 1.00 | 0.00 | H |
| ATOM | 734 | C | SER | A | 67 | 49.253 | 29.814 | 10.830 | 1.00 | 0.00 | C |
| ATOM | 735 | O | SER | A | 67 | 48.338 | 29.384 | 11.540 | 1.00 | 0.00 | O |
| ATOM | 736 | N | THR | A | 68 | 49.136 | 30.966 | 10.143 | 1.00 | 0.00 | N |
| ATOM | 737 | H | THR | A | 68 | 49.937 | 31.285 | 9.608 | 1.00 | 0.00 | H |
| ATOM | 738 | CA | THR | A | 68 | 47.920 | 31.811 | 10.116 | 1.00 | 0.00 | C |
| ATOM | 739 | HA | THR | A | 68 | 47.480 | 31.803 | 11.108 | 1.00 | 0.00 | H |
| ATOM | 740 | CB | THR | A | 68 | 48.233 | 33.281 | 9.812 | 1.00 | 0.00 | C |
| ATOM | 741 | HB | THR | A | 68 | 47.299 | 33.847 | 9.814 | 1.00 | 0.00 | H |
| ATOM | 742 | CG2 | THR | A | 68 | 49.155 | 33.911 | 10.855 | 1.00 | 0.00 | C |
| ATOM | 743 | HG21 | THR | A | 68 | 49.303 | 34.964 | 10.617 | 1.00 | 0.00 | H |
| ATOM | 744 | HG22 | THR | A | 68 | 48.695 | 33.836 | 11.841 | 1.00 | 0.00 | H |
| ATOM | 745 | HG23 | THR | A | 68 | 50.123 | 33.411 | 10.868 | 1.00 | 0.00 | H |
| ATOM | 746 | OG1 | THR | A | 68 | 48.834 | 33.410 | 8.544 | 1.00 | 0.00 | O |
| ATOM | 747 | HG1 | THR | A | 68 | 49.028 | 32.518 | 8.198 | 1.00 | 0.00 | H |
| ATOM | 748 | C | THR | A | 68 | 46.825 | 31.273 | 9.193 | 1.00 | 0.00 | C |
| ATOM | 749 | O | THR | A | 68 | 47.101 | 30.452 | 8.313 | 1.00 | 0.00 | O |
| ATOM | 750 | N | THR | A | 69 | 45.590 | 31.752 | 9.381 | 1.00 | 0.00 | N |
| ATOM | 751 | H | THR | A | 69 | 45.463 | 32.440 | 10.119 | 1.00 | 0.00 | H |
| ATOM | 752 | CA | THR | A | 69 | 44.404 | 31.387 | 8.587 | 1.00 | 0.00 | C |
| ATOM | 753 | HA | THR | A | 69 | 44.709 | 30.726 | 7.777 | 1.00 | 0.00 | H |

| | | | | | | | | | | |
|---|---|---|---|---|---|---|---|---|---|---|
| ATOM | 754 | CB | THR | A | 69 | 43.365 | 30.621 | 9.422 | 1.00 | 0.00 | C |
| ATOM | 755 | HB | THR | A | 69 | 42.472 | 30.471 | 8.811 | 1.00 | 0.00 | H |
| ATOM | 756 | CG2 | THR | A | 69 | 43.884 | 29.252 | 9.864 | 1.00 | 0.00 | C |
| ATOM | 757 | HG21 | THR | A | 69 | 43.113 | 28.734 | 10.436 | 1.00 | 0.00 | H |
| ATOM | 758 | HG22 | THR | A | 69 | 44.133 | 28.651 | 8.989 | 1.00 | 0.00 | H |
| ATOM | 759 | HG23 | THR | A | 69 | 44.771 | 29.361 | 10.488 | 1.00 | 0.00 | H |
| ATOM | 760 | OG1 | THR | A | 69 | 43.011 | 31.334 | 10.584 | 1.00 | 0.00 | O |
| ATOM | 761 | HG1 | THR | A | 69 | 43.791 | 31.329 | 11.168 | 1.00 | 0.00 | H |
| ATOM | 762 | C | THR | A | 69 | 43.775 | 32.608 | 7.926 | 1.00 | 0.00 | C |
| ATOM | 763 | O | THR | A | 69 | 43.526 | 32.551 | 6.720 | 1.00 | 0.00 | O |
| ATOM | 764 | N | GLU | A | 70 | 43.589 | 33.716 | 8.647 | 1.00 | 0.00 | N |
| ATOM | 765 | H | GLU | A | 70 | 43.885 | 33.718 | 9.621 | 1.00 | 0.00 | H |
| ATOM | 766 | CA | GLU | A | 70 | 43.016 | 34.960 | 8.121 | 1.00 | 0.00 | C |
| ATOM | 767 | HA | GLU | A | 70 | 42.589 | 34.764 | 7.140 | 1.00 | 0.00 | H |
| ATOM | 768 | CB | GLU | A | 70 | 41.862 | 35.469 | 9.004 | 1.00 | 0.00 | C |
| ATOM | 769 | HB1 | GLU | A | 70 | 42.247 | 35.725 | 9.992 | 1.00 | 0.00 | H |
| ATOM | 770 | HB2 | GLU | A | 70 | 41.460 | 36.378 | 8.554 | 1.00 | 0.00 | H |
| ATOM | 771 | CG | GLU | A | 70 | 40.711 | 34.462 | 9.165 | 1.00 | 0.00 | C |
| ATOM | 772 | HG1 | GLU | A | 70 | 41.043 | 33.652 | 9.813 | 1.00 | 0.00 | H |
| ATOM | 773 | HG2 | GLU | A | 70 | 39.873 | 34.960 | 9.652 | 1.00 | 0.00 | H |
| ATOM | 774 | CD | GLU | A | 70 | 40.232 | 33.859 | 7.844 | 1.00 | 0.00 | C |
| ATOM | 775 | OE1 | GLU | A | 70 | 39.856 | 34.600 | 6.904 | 1.00 | 0.00 | O |
| ATOM | 776 | OE2 | GLU | A | 70 | 40.224 | 32.615 | 7.720 | 1.00 | 0.00 | O |
| ATOM | 777 | C | GLU | A | 70 | 44.079 | 36.052 | 7.936 | 1.00 | 0.00 | C |
| ATOM | 778 | O | GLU | A | 70 | 45.031 | 36.176 | 8.709 | 1.00 | 0.00 | O |
| ATOM | 779 | N | TRP | A | 71 | 43.904 | 36.861 | 6.890 | 1.00 | 0.00 | N |
| ATOM | 780 | H | TRP | A | 71 | 43.095 | 36.695 | 6.311 | 1.00 | 0.00 | H |
| ATOM | 781 | CA | TRP | A | 71 | 44.758 | 37.984 | 6.479 | 1.00 | 0.00 | C |
| ATOM | 782 | HA | TRP | A | 71 | 45.697 | 37.592 | 6.084 | 1.00 | 0.00 | H |
| ATOM | 783 | CB | TRP | A | 71 | 44.017 | 38.725 | 5.346 | 1.00 | 0.00 | C |
| ATOM | 784 | HB1 | TRP | A | 71 | 44.611 | 39.591 | 5.049 | 1.00 | 0.00 | H |
| ATOM | 785 | HB2 | TRP | A | 71 | 43.959 | 38.065 | 4.479 | 1.00 | 0.00 | H |
| ATOM | 786 | CG | TRP | A | 71 | 42.622 | 39.186 | 5.679 | 1.00 | 0.00 | C |
| ATOM | 787 | CD1 | TRP | A | 71 | 41.532 | 38.389 | 5.743 | 1.00 | 0.00 | C |
| ATOM | 788 | HD1 | TRP | A | 71 | 41.517 | 37.330 | 5.529 | 1.00 | 0.00 | H |
| ATOM | 789 | NE1 | TRP | A | 71 | 40.445 | 39.103 | 6.190 | 1.00 | 0.00 | N |
| ATOM | 790 | HE1 | TRP | A | 71 | 39.547 | 38.664 | 6.375 | 1.00 | 0.00 | H |
| ATOM | 791 | CE2 | TRP | A | 71 | 40.756 | 40.427 | 6.390 | 1.00 | 0.00 | C |
| ATOM | 792 | CZ2 | TRP | A | 71 | 40.005 | 41.531 | 6.817 | 1.00 | 0.00 | C |
| ATOM | 793 | HZ2 | TRP | A | 71 | 38.973 | 41.409 | 7.112 | 1.00 | 0.00 | H |
| ATOM | 794 | CH2 | TRP | A | 71 | 40.617 | 42.793 | 6.864 | 1.00 | 0.00 | C |
| ATOM | 795 | HH2 | TRP | A | 71 | 40.059 | 43.655 | 7.204 | 1.00 | 0.00 | H |
| ATOM | 796 | CZ3 | TRP | A | 71 | 41.963 | 42.930 | 6.482 | 1.00 | 0.00 | C |
| ATOM | 797 | HZ3 | TRP | A | 71 | 42.434 | 43.901 | 6.521 | 1.00 | 0.00 | H |
| ATOM | 798 | CE3 | TRP | A | 71 | 42.714 | 41.808 | 6.079 | 1.00 | 0.00 | C |
| ATOM | 799 | HE3 | TRP | A | 71 | 43.753 | 41.927 | 5.813 | 1.00 | 0.00 | H |
| ATOM | 800 | CD2 | TRP | A | 71 | 42.135 | 40.523 | 6.027 | 1.00 | 0.00 | C |
| ATOM | 801 | C | TRP | A | 71 | 45.114 | 38.950 | 7.625 | 1.00 | 0.00 | C |
| ATOM | 802 | O | TRP | A | 71 | 46.245 | 39.444 | 7.698 | 1.00 | 0.00 | O |
| ATOM | 803 | N | ASN | A | 72 | 44.164 | 39.195 | 8.532 | 1.00 | 0.00 | N |
| ATOM | 804 | H | ASN | A | 72 | 43.268 | 38.740 | 8.416 | 1.00 | 0.00 | H |
| ATOM | 805 | CA | ASN | A | 72 | 44.315 | 40.091 | 9.675 | 1.00 | 0.00 | C |
| ATOM | 806 | HA | ASN | A | 72 | 45.136 | 40.770 | 9.450 | 1.00 | 0.00 | H |
| ATOM | 807 | CB | ASN | A | 72 | 43.074 | 40.998 | 9.802 | 1.00 | 0.00 | C |

| | | | | | | | | | | |
|---|---|---|---|---|---|---|---|---|---|---|
| ATOM | 808 | HB1 | ASN | A | 72 | 43.242 | 41.712 | 10.608 | 1.00 | 0.00 | H |
| ATOM | 809 | HB2 | ASN | A | 72 | 42.980 | 41.578 | 8.886 | 1.00 | 0.00 | H |
| ATOM | 810 | CG | ASN | A | 72 | 41.738 | 40.305 | 10.051 | 1.00 | 0.00 | C |
| ATOM | 811 | OD1 | ASN | A | 72 | 41.610 | 39.089 | 10.110 | 1.00 | 0.00 | O |
| ATOM | 812 | ND2 | ASN | A | 72 | 40.686 | 41.088 | 10.185 | 1.00 | 0.00 | N |
| ATOM | 813 | HD21 | ASN | A | 72 | 40.763 | 42.092 | 10.080 | 1.00 | 0.00 | H |
| ATOM | 814 | HD22 | ASN | A | 72 | 39.776 | 40.670 | 10.317 | 1.00 | 0.00 | H |
| ATOM | 815 | C | ASN | A | 72 | 44.720 | 39.399 | 10.995 | 1.00 | 0.00 | C |
| ATOM | 816 | O | ASN | A | 72 | 44.793 | 40.088 | 12.008 | 1.00 | 0.00 | O |
| ATOM | 817 | N | ASP | A | 73 | 45.068 | 38.102 | 11.020 | 1.00 | 0.00 | N |
| ATOM | 818 | H | ASP | A | 73 | 44.972 | 37.552 | 10.171 | 1.00 | 0.00 | H |
| ATOM | 819 | CA | ASP | A | 73 | 45.507 | 37.406 | 12.251 | 1.00 | 0.00 | C |
| ATOM | 820 | HA | ASP | A | 73 | 44.675 | 37.383 | 12.956 | 1.00 | 0.00 | H |
| ATOM | 821 | CB | ASP | A | 73 | 45.931 | 35.955 | 11.943 | 1.00 | 0.00 | C |
| ATOM | 822 | HB1 | ASP | A | 73 | 46.535 | 35.941 | 11.034 | 1.00 | 0.00 | H |
| ATOM | 823 | HB2 | ASP | A | 73 | 46.569 | 35.605 | 12.756 | 1.00 | 0.00 | H |
| ATOM | 824 | CG | ASP | A | 73 | 44.784 | 34.943 | 11.822 | 1.00 | 0.00 | C |
| ATOM | 825 | OD1 | ASP | A | 73 | 43.725 | 35.116 | 12.462 | 1.00 | 0.00 | O |
| ATOM | 826 | OD2 | ASP | A | 73 | 44.989 | 33.903 | 11.153 | 1.00 | 0.00 | O |
| ATOM | 827 | C | ASP | A | 73 | 46.668 | 38.133 | 12.962 | 1.00 | 0.00 | C |
| ATOM | 828 | O | ASP | A | 73 | 46.590 | 38.405 | 14.157 | 1.00 | 0.00 | O |
| ATOM | 829 | N | GLU | A | 74 | 47.708 | 38.537 | 12.222 | 1.00 | 0.00 | N |
| ATOM | 830 | H | GLU | A | 74 | 47.704 | 38.302 | 11.241 | 1.00 | 0.00 | H |
| ATOM | 831 | CA | GLU | A | 74 | 48.870 | 39.261 | 12.773 | 1.00 | 0.00 | C |
| ATOM | 832 | HA | GLU | A | 74 | 49.204 | 38.724 | 13.662 | 1.00 | 0.00 | H |
| ATOM | 833 | CB | GLU | A | 74 | 50.035 | 39.262 | 11.765 | 1.00 | 0.00 | C |
| ATOM | 834 | HB1 | GLU | A | 74 | 49.725 | 39.748 | 10.839 | 1.00 | 0.00 | H |
| ATOM | 835 | HB2 | GLU | A | 74 | 50.862 | 39.830 | 12.190 | 1.00 | 0.00 | H |
| ATOM | 836 | CG | GLU | A | 74 | 50.529 | 37.842 | 11.459 | 1.00 | 0.00 | C |
| ATOM | 837 | HG1 | GLU | A | 74 | 50.605 | 37.279 | 12.389 | 1.00 | 0.00 | H |
| ATOM | 838 | HG2 | GLU | A | 74 | 49.805 | 37.341 | 10.815 | 1.00 | 0.00 | H |
| ATOM | 839 | CD | GLU | A | 74 | 51.901 | 37.842 | 10.785 | 1.00 | 0.00 | C |
| ATOM | 840 | OE1 | GLU | A | 74 | 51.998 | 37.421 | 9.611 | 1.00 | 0.00 | O |
| ATOM | 841 | OE2 | GLU | A | 74 | 52.910 | 38.159 | 11.458 | 1.00 | 0.00 | O |
| ATOM | 842 | C | GLU | A | 74 | 48.555 | 40.700 | 13.237 | 1.00 | 0.00 | C |
| ATOM | 843 | O | GLU | A | 74 | 49.417 | 41.352 | 13.834 | 1.00 | 0.00 | O |
| ATOM | 844 | N | THR | A | 75 | 47.331 | 41.186 | 12.985 | 1.00 | 0.00 | N |
| ATOM | 845 | H | THR | A | 75 | 46.686 | 40.562 | 12.514 | 1.00 | 0.00 | H |
| ATOM | 846 | CA | THR | A | 75 | 46.778 | 42.457 | 13.482 | 1.00 | 0.00 | C |
| ATOM | 847 | HA | THR | A | 75 | 47.588 | 43.110 | 13.813 | 1.00 | 0.00 | H |
| ATOM | 848 | CB | THR | A | 75 | 46.019 | 43.161 | 12.349 | 1.00 | 0.00 | C |
| ATOM | 849 | HB | THR | A | 75 | 45.224 | 42.519 | 11.969 | 1.00 | 0.00 | H |
| ATOM | 850 | CG2 | THR | A | 75 | 45.400 | 44.489 | 12.779 | 1.00 | 0.00 | C |
| ATOM | 851 | HG21 | THR | A | 75 | 44.945 | 44.979 | 11.919 | 1.00 | 0.00 | H |
| ATOM | 852 | HG22 | THR | A | 75 | 44.625 | 44.320 | 13.527 | 1.00 | 0.00 | H |
| ATOM | 853 | HG23 | THR | A | 75 | 46.171 | 45.131 | 13.197 | 1.00 | 0.00 | H |
| ATOM | 854 | OG1 | THR | A | 75 | 46.927 | 43.449 | 11.308 | 1.00 | 0.00 | O |
| ATOM | 855 | HG1 | THR | A | 75 | 47.160 | 42.617 | 10.861 | 1.00 | 0.00 | H |
| ATOM | 856 | C | THR | A | 75 | 45.865 | 42.210 | 14.687 | 1.00 | 0.00 | C |
| ATOM | 857 | O | THR | A | 75 | 45.932 | 42.954 | 15.664 | 1.00 | 0.00 | O |
| ATOM | 858 | N | LEU | A | 76 | 45.052 | 41.149 | 14.655 | 1.00 | 0.00 | N |
| ATOM | 859 | H | LEU | A | 76 | 45.050 | 40.578 | 13.818 | 1.00 | 0.00 | H |
| ATOM | 860 | CA | LEU | A | 76 | 44.157 | 40.736 | 15.743 | 1.00 | 0.00 | C |
| ATOM | 861 | HA | LEU | A | 76 | 43.560 | 41.591 | 16.055 | 1.00 | 0.00 | H |

```
ATOM    862  CB  LEU A  76      43.222  39.625  15.224  1.00  0.00           C
ATOM    863  HB1 LEU A  76      43.837  38.836  14.788  1.00  0.00           H
ATOM    864  HB2 LEU A  76      42.687  39.195  16.071  1.00  0.00           H
ATOM    865  CG  LEU A  76      42.180  40.083  14.180  1.00  0.00           C
ATOM    866  HG  LEU A  76      42.680  40.610  13.368  1.00  0.00           H
ATOM    867  CD1 LEU A  76      41.458  38.867  13.583  1.00  0.00           C
ATOM    868 HD11 LEU A  76      40.706  39.195  12.865  1.00  0.00           H
ATOM    869 HD12 LEU A  76      42.176  38.231  13.064  1.00  0.00           H
ATOM    870 HD13 LEU A  76      40.970  38.292  14.372  1.00  0.00           H
ATOM    871  CD2 LEU A  76      41.142  41.033  14.793  1.00  0.00           C
ATOM    872 HD21 LEU A  76      40.401  41.293  14.040  1.00  0.00           H
ATOM    873 HD22 LEU A  76      40.643  40.551  15.635  1.00  0.00           H
ATOM    874 HD23 LEU A  76      41.620  41.952  15.128  1.00  0.00           H
ATOM    875  C   LEU A  76      44.934  40.283  16.986  1.00  0.00           C
ATOM    876  O   LEU A  76      44.573  40.661  18.099  1.00  0.00           O
ATOM    877  N   TYR A  77      46.064  39.590  16.818  1.00  0.00           N
ATOM    878  H   TYR A  77      46.281  39.244  15.887  1.00  0.00           H
ATOM    879  CA  TYR A  77      46.986  39.255  17.917  1.00  0.00           C
ATOM    880  HA  TYR A  77      46.444  38.680  18.667  1.00  0.00           H
ATOM    881  CB  TYR A  77      48.142  38.396  17.367  1.00  0.00           C
ATOM    882  HB1 TYR A  77      48.656  38.968  16.592  1.00  0.00           H
ATOM    883  HB2 TYR A  77      48.853  38.239  18.177  1.00  0.00           H
ATOM    884  CG  TYR A  77      47.790  37.021  16.806  1.00  0.00           C
ATOM    885  CD1 TYR A  77      46.696  36.287  17.307  1.00  0.00           C
ATOM    886  HD1 TYR A  77      46.065  36.693  18.082  1.00  0.00           H
ATOM    887  CE1 TYR A  77      46.407  35.007  16.805  1.00  0.00           C
ATOM    888  HE1 TYR A  77      45.573  34.447  17.197  1.00  0.00           H
ATOM    889  CZ  TYR A  77      47.199  34.443  15.787  1.00  0.00           C
ATOM    890  OH  TYR A  77      46.885  33.202  15.327  1.00  0.00           O
ATOM    891  HH  TYR A  77      47.491  32.840  14.669  1.00  0.00           H
ATOM    892  CE2 TYR A  77      48.311  35.160  15.291  1.00  0.00           C
ATOM    893  HE2 TYR A  77      48.934  34.730  14.521  1.00  0.00           H
ATOM    894  CD2 TYR A  77      48.602  36.441  15.806  1.00  0.00           C
ATOM    895  HD2 TYR A  77      49.450  36.988  15.422  1.00  0.00           H
ATOM    896  C   TYR A  77      47.533  40.494  18.660  1.00  0.00           C
ATOM    897  O   TYR A  77      47.962  40.396  19.809  1.00  0.00           O
ATOM    898  N   GLN A  78      47.486  41.678  18.045  1.00  0.00           N
ATOM    899  H   GLN A  78      47.097  41.720  17.111  1.00  0.00           H
ATOM    900  CA  GLN A  78      47.889  42.932  18.685  1.00  0.00           C
ATOM    901  HA  GLN A  78      48.678  42.732  19.413  1.00  0.00           H
ATOM    902  CB  GLN A  78      48.471  43.878  17.625  1.00  0.00           C
ATOM    903  HB1 GLN A  78      47.702  44.126  16.901  1.00  0.00           H
ATOM    904  HB2 GLN A  78      48.780  44.804  18.111  1.00  0.00           H
ATOM    905  CG  GLN A  78      49.685  43.260  16.899  1.00  0.00           C
ATOM    906  HG1 GLN A  78      50.485  43.112  17.625  1.00  0.00           H
ATOM    907  HG2 GLN A  78      49.429  42.287  16.481  1.00  0.00           H
ATOM    908  CD  GLN A  78      50.207  44.113  15.745  1.00  0.00           C
ATOM    909  OE1 GLN A  78      49.505  44.932  15.160  1.00  0.00           O
ATOM    910  NE2 GLN A  78      51.464  43.985  15.385  1.00  0.00           N
ATOM    911 HE21 GLN A  78      52.049  43.320  15.879  1.00  0.00           H
ATOM    912 HE22 GLN A  78      51.795  44.464  14.558  1.00  0.00           H
ATOM    913  C   GLN A  78      46.725  43.572  19.466  1.00  0.00           C
ATOM    914  O   GLN A  78      46.977  44.259  20.453  1.00  0.00           O
ATOM    915  N   GLU A  79      45.464  43.293  19.107  1.00  0.00           N
```

| | | | | | | | | | | |
|---|---|---|---|---|---|---|---|---|---|---|
| ATOM | 916 | H | GLU | A | 79 | 45.310 | 42.678 | 18.317 | 1.00 | 0.00 | H |
| ATOM | 917 | CA | GLU | A | 79 | 44.276 | 43.710 | 19.872 | 1.00 | 0.00 | C |
| ATOM | 918 | HA | GLU | A | 79 | 44.337 | 44.780 | 20.071 | 1.00 | 0.00 | H |
| ATOM | 919 | CB | GLU | A | 79 | 42.983 | 43.412 | 19.083 | 1.00 | 0.00 | C |
| ATOM | 920 | HB1 | GLU | A | 79 | 42.858 | 42.331 | 19.020 | 1.00 | 0.00 | H |
| ATOM | 921 | HB2 | GLU | A | 79 | 42.133 | 43.790 | 19.652 | 1.00 | 0.00 | H |
| ATOM | 922 | CG | GLU | A | 79 | 42.911 | 43.967 | 17.651 | 1.00 | 0.00 | C |
| ATOM | 923 | HG1 | GLU | A | 79 | 43.833 | 43.751 | 17.113 | 1.00 | 0.00 | H |
| ATOM | 924 | HG2 | GLU | A | 79 | 42.107 | 43.445 | 17.133 | 1.00 | 0.00 | H |
| ATOM | 925 | CD | GLU | A | 79 | 42.627 | 45.464 | 17.568 | 1.00 | 0.00 | C |
| ATOM | 926 | OE1 | GLU | A | 79 | 42.816 | 46.201 | 18.558 | 1.00 | 0.00 | O |
| ATOM | 927 | OE2 | GLU | A | 79 | 42.215 | 45.931 | 16.480 | 1.00 | 0.00 | O |
| ATOM | 928 | C | GLU | A | 79 | 44.201 | 42.969 | 21.219 | 1.00 | 0.00 | C |
| ATOM | 929 | O | GLU | A | 79 | 43.894 | 43.569 | 22.255 | 1.00 | 0.00 | O |
| ATOM | 930 | N | PHE | A | 80 | 44.528 | 41.668 | 21.203 | 1.00 | 0.00 | N |
| ATOM | 931 | H | PHE | A | 80 | 44.723 | 41.250 | 20.301 | 1.00 | 0.00 | H |
| ATOM | 932 | CA | PHE | A | 80 | 44.670 | 40.809 | 22.383 | 1.00 | 0.00 | C |
| ATOM | 933 | HA | PHE | A | 80 | 43.735 | 40.803 | 22.943 | 1.00 | 0.00 | H |
| ATOM | 934 | CB | PHE | A | 80 | 44.985 | 39.374 | 21.913 | 1.00 | 0.00 | C |
| ATOM | 935 | HB1 | PHE | A | 80 | 45.542 | 39.408 | 20.976 | 1.00 | 0.00 | H |
| ATOM | 936 | HB2 | PHE | A | 80 | 45.649 | 38.920 | 22.651 | 1.00 | 0.00 | H |
| ATOM | 937 | CG | PHE | A | 80 | 43.800 | 38.437 | 21.732 | 1.00 | 0.00 | C |
| ATOM | 938 | CD1 | PHE | A | 80 | 43.919 | 37.096 | 22.148 | 1.00 | 0.00 | C |
| ATOM | 939 | HD1 | PHE | A | 80 | 44.852 | 36.739 | 22.559 | 1.00 | 0.00 | H |
| ATOM | 940 | CE1 | PHE | A | 80 | 42.833 | 36.212 | 22.029 | 1.00 | 0.00 | C |
| ATOM | 941 | HE1 | PHE | A | 80 | 42.929 | 35.189 | 22.366 | 1.00 | 0.00 | H |
| ATOM | 942 | CZ | PHE | A | 80 | 41.627 | 36.657 | 21.462 | 1.00 | 0.00 | C |
| ATOM | 943 | HZ | PHE | A | 80 | 40.794 | 35.977 | 21.365 | 1.00 | 0.00 | H |
| ATOM | 944 | CE2 | PHE | A | 80 | 41.513 | 37.983 | 21.008 | 1.00 | 0.00 | C |
| ATOM | 945 | HE2 | PHE | A | 80 | 40.591 | 38.318 | 20.555 | 1.00 | 0.00 | H |
| ATOM | 946 | CD2 | PHE | A | 80 | 42.594 | 38.873 | 21.145 | 1.00 | 0.00 | C |
| ATOM | 947 | HD2 | PHE | A | 80 | 42.493 | 39.889 | 20.793 | 1.00 | 0.00 | H |
| ATOM | 948 | C | PHE | A | 80 | 45.762 | 41.322 | 23.335 | 1.00 | 0.00 | C |
| ATOM | 949 | O | PHE | A | 80 | 45.578 | 41.305 | 24.552 | 1.00 | 0.00 | O |
| ATOM | 950 | N | ASN | A | 81 | 46.873 | 41.840 | 22.801 | 1.00 | 0.00 | N |
| ATOM | 951 | H | ASN | A | 81 | 46.992 | 41.801 | 21.799 | 1.00 | 0.00 | H |
| ATOM | 952 | CA | ASN | A | 81 | 47.925 | 42.464 | 23.611 | 1.00 | 0.00 | C |
| ATOM | 953 | HA | ASN | A | 81 | 48.156 | 41.799 | 24.446 | 1.00 | 0.00 | H |
| ATOM | 954 | CB | ASN | A | 81 | 49.206 | 42.635 | 22.781 | 1.00 | 0.00 | C |
| ATOM | 955 | HB1 | ASN | A | 81 | 48.983 | 43.140 | 21.843 | 1.00 | 0.00 | H |
| ATOM | 956 | HB2 | ASN | A | 81 | 49.905 | 43.260 | 23.338 | 1.00 | 0.00 | H |
| ATOM | 957 | CG | ASN | A | 81 | 49.902 | 41.315 | 22.484 | 1.00 | 0.00 | C |
| ATOM | 958 | OD1 | ASN | A | 81 | 49.357 | 40.233 | 22.673 | 1.00 | 0.00 | O |
| ATOM | 959 | ND2 | ASN | A | 81 | 51.139 | 41.372 | 22.033 | 1.00 | 0.00 | N |
| ATOM | 960 | HD21 | ASN | A | 81 | 51.620 | 42.260 | 21.944 | 1.00 | 0.00 | H |
| ATOM | 961 | HD22 | ASN | A | 81 | 51.590 | 40.501 | 21.798 | 1.00 | 0.00 | H |
| ATOM | 962 | C | ASN | A | 81 | 47.480 | 43.800 | 24.226 | 1.00 | 0.00 | C |
| ATOM | 963 | O | ASN | A | 81 | 48.024 | 44.206 | 25.255 | 1.00 | 0.00 | O |
| ATOM | 964 | N | GLY | A | 82 | 46.459 | 44.458 | 23.663 | 1.00 | 0.00 | N |
| ATOM | 965 | H | GLY | A | 82 | 46.069 | 44.097 | 22.799 | 1.00 | 0.00 | H |
| ATOM | 966 | CA | GLY | A | 82 | 45.809 | 45.609 | 24.287 | 1.00 | 0.00 | C |
| ATOM | 967 | HA1 | GLY | A | 82 | 46.564 | 46.316 | 24.633 | 1.00 | 0.00 | H |
| ATOM | 968 | HA2 | GLY | A | 82 | 45.154 | 46.093 | 23.566 | 1.00 | 0.00 | H |
| ATOM | 969 | C | GLY | A | 82 | 44.984 | 45.164 | 25.492 | 1.00 | 0.00 | C |

| | | | | | | | | | | | |
|---|---|---|---|---|---|---|---|---|---|---|---|
| ATOM | 970 | O | GLY | A | 82 | 45.137 | 45.731 | 26.572 | 1.00 | 0.00 | O |
| ATOM | 971 | N | LEU | A | 83 | 44.191 | 44.094 | 25.356 | 1.00 | 0.00 | N |
| ATOM | 972 | H | LEU | A | 83 | 44.123 | 43.666 | 24.439 | 1.00 | 0.00 | H |
| ATOM | 973 | CA | LEU | A | 83 | 43.420 | 43.514 | 26.467 | 1.00 | 0.00 | C |
| ATOM | 974 | HA | LEU | A | 83 | 42.761 | 44.285 | 26.867 | 1.00 | 0.00 | H |
| ATOM | 975 | CB | LEU | A | 83 | 42.562 | 42.333 | 25.973 | 1.00 | 0.00 | C |
| ATOM | 976 | HB1 | LEU | A | 83 | 43.225 | 41.554 | 25.596 | 1.00 | 0.00 | H |
| ATOM | 977 | HB2 | LEU | A | 83 | 42.026 | 41.918 | 26.828 | 1.00 | 0.00 | H |
| ATOM | 978 | CG | LEU | A | 83 | 41.529 | 42.676 | 24.879 | 1.00 | 0.00 | C |
| ATOM | 979 | HG | LEU | A | 83 | 42.036 | 43.148 | 24.040 | 1.00 | 0.00 | H |
| ATOM | 980 | CD1 | LEU | A | 83 | 40.885 | 41.382 | 24.368 | 1.00 | 0.00 | C |
| ATOM | 981 | HD11 | LEU | A | 83 | 40.220 | 41.621 | 23.540 | 1.00 | 0.00 | H |
| ATOM | 982 | HD12 | LEU | A | 83 | 41.651 | 40.695 | 24.008 | 1.00 | 0.00 | H |
| ATOM | 983 | HD13 | LEU | A | 83 | 40.317 | 40.900 | 25.165 | 1.00 | 0.00 | H |
| ATOM | 984 | CD2 | LEU | A | 83 | 40.432 | 43.626 | 25.382 | 1.00 | 0.00 | C |
| ATOM | 985 | HD21 | LEU | A | 83 | 39.714 | 43.815 | 24.583 | 1.00 | 0.00 | H |
| ATOM | 986 | HD22 | LEU | A | 83 | 39.908 | 43.178 | 26.227 | 1.00 | 0.00 | H |
| ATOM | 987 | HD23 | LEU | A | 83 | 40.864 | 44.579 | 25.686 | 1.00 | 0.00 | H |
| ATOM | 988 | C | LEU | A | 83 | 44.332 | 43.096 | 27.631 | 1.00 | 0.00 | C |
| ATOM | 989 | O | LEU | A | 83 | 44.082 | 43.500 | 28.764 | 1.00 | 0.00 | O |
| ATOM | 990 | N | LYS | A | 84 | 45.435 | 42.383 | 27.368 | 1.00 | 0.00 | N |
| ATOM | 991 | H | LYS | A | 84 | 45.583 | 42.052 | 26.420 | 1.00 | 0.00 | H |
| ATOM | 992 | CA | LYS | A | 84 | 46.417 | 42.025 | 28.408 | 1.00 | 0.00 | C |
| ATOM | 993 | HA | LYS | A | 84 | 45.875 | 41.587 | 29.249 | 1.00 | 0.00 | H |
| ATOM | 994 | CB | LYS | A | 84 | 47.408 | 40.968 | 27.885 | 1.00 | 0.00 | C |
| ATOM | 995 | HB1 | LYS | A | 84 | 47.850 | 41.298 | 26.943 | 1.00 | 0.00 | H |
| ATOM | 996 | HB2 | LYS | A | 84 | 48.211 | 40.882 | 28.618 | 1.00 | 0.00 | H |
| ATOM | 997 | CG | LYS | A | 84 | 46.775 | 39.577 | 27.711 | 1.00 | 0.00 | C |
| ATOM | 998 | HG1 | LYS | A | 84 | 45.943 | 39.467 | 28.407 | 1.00 | 0.00 | H |
| ATOM | 999 | HG2 | LYS | A | 84 | 46.385 | 39.477 | 26.697 | 1.00 | 0.00 | H |
| ATOM | 1000 | CD | LYS | A | 84 | 47.778 | 38.447 | 28.002 | 1.00 | 0.00 | C |
| ATOM | 1001 | HD1 | LYS | A | 84 | 48.149 | 38.555 | 29.023 | 1.00 | 0.00 | H |
| ATOM | 1002 | HD2 | LYS | A | 84 | 47.254 | 37.493 | 27.935 | 1.00 | 0.00 | H |
| ATOM | 1003 | CE | LYS | A | 84 | 48.962 | 38.434 | 27.034 | 1.00 | 0.00 | C |
| ATOM | 1004 | HE1 | LYS | A | 84 | 48.575 | 38.268 | 26.026 | 1.00 | 0.00 | H |
| ATOM | 1005 | HE2 | LYS | A | 84 | 49.469 | 39.401 | 27.052 | 1.00 | 0.00 | H |
| ATOM | 1006 | NZ | LYS | A | 84 | 49.923 | 37.360 | 27.371 | 1.00 | 0.00 | N |
| ATOM | 1007 | HZ1 | LYS | A | 84 | 50.534 | 37.191 | 26.578 | 1.00 | 0.00 | H |
| ATOM | 1008 | HZ2 | LYS | A | 84 | 50.488 | 37.594 | 28.181 | 1.00 | 0.00 | H |
| ATOM | 1009 | HZ3 | LYS | A | 84 | 49.431 | 36.491 | 27.577 | 1.00 | 0.00 | H |
| ATOM | 1010 | C | LYS | A | 84 | 47.183 | 43.222 | 29.013 | 1.00 | 0.00 | C |
| ATOM | 1011 | O | LYS | A | 84 | 47.805 | 43.063 | 30.064 | 1.00 | 0.00 | O |
| ATOM | 1012 | N | LYS | A | 85 | 47.152 | 44.410 | 28.398 | 1.00 | 0.00 | N |
| ATOM | 1013 | H | LYS | A | 85 | 46.659 | 44.475 | 27.517 | 1.00 | 0.00 | H |
| ATOM | 1014 | CA | LYS | A | 85 | 47.721 | 45.645 | 28.963 | 1.00 | 0.00 | C |
| ATOM | 1015 | HA | LYS | A | 85 | 48.540 | 45.384 | 29.638 | 1.00 | 0.00 | H |
| ATOM | 1016 | CB | LYS | A | 85 | 48.329 | 46.466 | 27.813 | 1.00 | 0.00 | C |
| ATOM | 1017 | HB1 | LYS | A | 85 | 49.088 | 45.841 | 27.346 | 1.00 | 0.00 | H |
| ATOM | 1018 | HB2 | LYS | A | 85 | 47.563 | 46.688 | 27.070 | 1.00 | 0.00 | H |
| ATOM | 1019 | CG | LYS | A | 85 | 48.995 | 47.784 | 28.247 | 1.00 | 0.00 | C |
| ATOM | 1020 | HG1 | LYS | A | 85 | 48.217 | 48.510 | 28.489 | 1.00 | 0.00 | H |
| ATOM | 1021 | HG2 | LYS | A | 85 | 49.614 | 47.613 | 29.127 | 1.00 | 0.00 | H |
| ATOM | 1022 | CD | LYS | A | 85 | 49.886 | 48.331 | 27.118 | 1.00 | 0.00 | C |
| ATOM | 1023 | HD1 | LYS | A | 85 | 50.815 | 47.759 | 27.078 | 1.00 | 0.00 | H |

| | | | | | | | | | | | |
|---|---|---|---|---|---|---|---|---|---|---|---|
| ATOM | 1024 | HD2 | LYS | A | 85 | 49.369 | 48.187 | 26.168 | 1.00 | 0.00 | H |
| ATOM | 1025 | CE | LYS | A | 85 | 50.196 | 49.829 | 27.243 | 1.00 | 0.00 | C |
| ATOM | 1026 | HE1 | LYS | A | 85 | 50.705 | 50.157 | 26.333 | 1.00 | 0.00 | H |
| ATOM | 1027 | HE2 | LYS | A | 85 | 49.250 | 50.373 | 27.305 | 1.00 | 0.00 | H |
| ATOM | 1028 | NZ | LYS | A | 85 | 51.029 | 50.180 | 28.418 | 1.00 | 0.00 | N |
| ATOM | 1029 | HZ1 | LYS | A | 85 | 50.567 | 49.919 | 29.285 | 1.00 | 0.00 | H |
| ATOM | 1030 | HZ2 | LYS | A | 85 | 51.952 | 49.758 | 28.382 | 1.00 | 0.00 | H |
| ATOM | 1031 | HZ3 | LYS | A | 85 | 51.140 | 51.191 | 28.451 | 1.00 | 0.00 | H |
| ATOM | 1032 | C | LYS | A | 85 | 46.697 | 46.418 | 29.815 | 1.00 | 0.00 | C |
| ATOM | 1033 | O | LYS | A | 85 | 47.051 | 46.873 | 30.903 | 1.00 | 0.00 | O |
| ATOM | 1034 | N | MET | A | 86 | 45.426 | 46.489 | 29.393 | 1.00 | 0.00 | N |
| ATOM | 1035 | H | MET | A | 86 | 45.213 | 46.145 | 28.462 | 1.00 | 0.00 | H |
| ATOM | 1036 | CA | MET | A | 86 | 44.319 | 47.019 | 30.213 | 1.00 | 0.00 | C |
| ATOM | 1037 | HA | MET | A | 86 | 44.642 | 47.959 | 30.667 | 1.00 | 0.00 | H |
| ATOM | 1038 | CB | MET | A | 86 | 43.085 | 47.316 | 29.339 | 1.00 | 0.00 | C |
| ATOM | 1039 | HB1 | MET | A | 86 | 42.914 | 46.506 | 28.628 | 1.00 | 0.00 | H |
| ATOM | 1040 | HB2 | MET | A | 86 | 42.212 | 47.374 | 29.993 | 1.00 | 0.00 | H |
| ATOM | 1041 | CG | MET | A | 86 | 43.183 | 48.665 | 28.608 | 1.00 | 0.00 | C |
| ATOM | 1042 | HG1 | MET | A | 86 | 42.173 | 48.973 | 28.333 | 1.00 | 0.00 | H |
| ATOM | 1043 | HG2 | MET | A | 86 | 43.546 | 49.416 | 29.314 | 1.00 | 0.00 | H |
| ATOM | 1044 | SD | MET | A | 86 | 44.208 | 48.739 | 27.115 | 1.00 | 0.00 | S |
| ATOM | 1045 | CE | MET | A | 86 | 43.018 | 48.080 | 25.918 | 1.00 | 0.00 | C |
| ATOM | 1046 | HE1 | MET | A | 86 | 42.139 | 48.724 | 25.875 | 1.00 | 0.00 | H |
| ATOM | 1047 | HE2 | MET | A | 86 | 43.473 | 48.031 | 24.929 | 1.00 | 0.00 | H |
| ATOM | 1048 | HE3 | MET | A | 86 | 42.713 | 47.076 | 26.214 | 1.00 | 0.00 | H |
| ATOM | 1049 | C | MET | A | 86 | 43.945 | 46.101 | 31.393 | 1.00 | 0.00 | C |
| ATOM | 1050 | O | MET | A | 86 | 43.449 | 46.595 | 32.405 | 1.00 | 0.00 | O |
| ATOM | 1051 | N | ASN | A | 87 | 44.223 | 44.796 | 31.313 | 1.00 | 0.00 | N |
| ATOM | 1052 | H | ASN | A | 87 | 44.485 | 44.419 | 30.409 | 1.00 | 0.00 | H |
| ATOM | 1053 | CA | ASN | A | 87 | 44.144 | 43.861 | 32.441 | 1.00 | 0.00 | C |
| ATOM | 1054 | HA | ASN | A | 87 | 44.221 | 44.421 | 33.373 | 1.00 | 0.00 | H |
| ATOM | 1055 | CB | ASN | A | 87 | 42.770 | 43.171 | 32.454 | 1.00 | 0.00 | C |
| ATOM | 1056 | HB1 | ASN | A | 87 | 41.981 | 43.917 | 32.534 | 1.00 | 0.00 | H |
| ATOM | 1057 | HB2 | ASN | A | 87 | 42.633 | 42.628 | 31.518 | 1.00 | 0.00 | H |
| ATOM | 1058 | CG | ASN | A | 87 | 42.627 | 42.189 | 33.615 | 1.00 | 0.00 | C |
| ATOM | 1059 | OD1 | ASN | A | 87 | 43.362 | 42.224 | 34.597 | 1.00 | 0.00 | O |
| ATOM | 1060 | ND2 | ASN | A | 87 | 41.703 | 41.258 | 33.508 | 1.00 | 0.00 | N |
| ATOM | 1061 | HD21 | ASN | A | 87 | 41.127 | 41.217 | 32.672 | 1.00 | 0.00 | H |
| ATOM | 1062 | HD22 | ASN | A | 87 | 41.602 | 40.551 | 34.216 | 1.00 | 0.00 | H |
| ATOM | 1063 | C | ASN | A | 87 | 45.288 | 42.819 | 32.399 | 1.00 | 0.00 | C |
| ATOM | 1064 | O | ASN | A | 87 | 45.152 | 41.793 | 31.723 | 1.00 | 0.00 | O |
| ATOM | 1065 | N | PRO | A | 88 | 46.391 | 43.040 | 33.144 | 1.00 | 0.00 | N |
| ATOM | 1066 | CD | PRO | A | 88 | 46.738 | 44.296 | 33.794 | 1.00 | 0.00 | C |
| ATOM | 1067 | HD1 | PRO | A | 88 | 46.236 | 44.348 | 34.761 | 1.00 | 0.00 | H |
| ATOM | 1068 | HD2 | PRO | A | 88 | 46.473 | 45.160 | 33.183 | 1.00 | 0.00 | H |
| ATOM | 1069 | CG | PRO | A | 88 | 48.251 | 44.249 | 33.996 | 1.00 | 0.00 | C |
| ATOM | 1070 | HG1 | PRO | A | 88 | 48.571 | 44.853 | 34.846 | 1.00 | 0.00 | H |
| ATOM | 1071 | HG2 | PRO | A | 88 | 48.757 | 44.569 | 33.083 | 1.00 | 0.00 | H |
| ATOM | 1072 | CB | PRO | A | 88 | 48.495 | 42.762 | 34.229 | 1.00 | 0.00 | C |
| ATOM | 1073 | HB1 | PRO | A | 88 | 48.251 | 42.513 | 35.264 | 1.00 | 0.00 | H |
| ATOM | 1074 | HB2 | PRO | A | 88 | 49.521 | 42.478 | 34.000 | 1.00 | 0.00 | H |
| ATOM | 1075 | CA | PRO | A | 88 | 47.501 | 42.095 | 33.273 | 1.00 | 0.00 | C |
| ATOM | 1076 | HA | PRO | A | 88 | 47.977 | 41.991 | 32.299 | 1.00 | 0.00 | H |
| ATOM | 1077 | C | PRO | A | 88 | 47.126 | 40.692 | 33.768 | 1.00 | 0.00 | C |

| | | | | | | | | | | | |
|---|---|---|---|---|---|---|---|---|---|---|---|
| ATOM | 1078 | O | PRO | A | 88 | 47.864 | 39.749 | 33.489 | 1.00 | 0.00 | O |
| ATOM | 1079 | N | LYS | A | 89 | 45.995 | 40.510 | 34.468 | 1.00 | 0.00 | N |
| ATOM | 1080 | H | LYS | A | 89 | 45.398 | 41.298 | 34.690 | 1.00 | 0.00 | H |
| ATOM | 1081 | CA | LYS | A | 89 | 45.587 | 39.174 | 34.924 | 1.00 | 0.00 | C |
| ATOM | 1082 | HA | LYS | A | 89 | 46.445 | 38.695 | 35.398 | 1.00 | 0.00 | H |
| ATOM | 1083 | CB | LYS | A | 89 | 44.448 | 39.261 | 35.948 | 1.00 | 0.00 | C |
| ATOM | 1084 | HB1 | LYS | A | 89 | 43.588 | 39.740 | 35.479 | 1.00 | 0.00 | H |
| ATOM | 1085 | HB2 | LYS | A | 89 | 44.159 | 38.243 | 36.218 | 1.00 | 0.00 | H |
| ATOM | 1086 | CG | LYS | A | 89 | 44.818 | 40.002 | 37.237 | 1.00 | 0.00 | C |
| ATOM | 1087 | HG1 | LYS | A | 89 | 45.772 | 39.629 | 37.612 | 1.00 | 0.00 | H |
| ATOM | 1088 | HG2 | LYS | A | 89 | 44.918 | 41.068 | 37.029 | 1.00 | 0.00 | H |
| ATOM | 1089 | CD | LYS | A | 89 | 43.750 | 39.788 | 38.319 | 1.00 | 0.00 | C |
| ATOM | 1090 | HD1 | LYS | A | 89 | 43.776 | 38.746 | 38.640 | 1.00 | 0.00 | H |
| ATOM | 1091 | HD2 | LYS | A | 89 | 44.007 | 40.410 | 39.175 | 1.00 | 0.00 | H |
| ATOM | 1092 | CE | LYS | A | 89 | 42.337 | 40.129 | 37.820 | 1.00 | 0.00 | C |
| ATOM | 1093 | HE1 | LYS | A | 89 | 42.389 | 41.030 | 37.203 | 1.00 | 0.00 | H |
| ATOM | 1094 | HE2 | LYS | A | 89 | 41.967 | 39.314 | 37.192 | 1.00 | 0.00 | H |
| ATOM | 1095 | NZ | LYS | A | 89 | 41.389 | 40.379 | 38.927 | 1.00 | 0.00 | N |
| ATOM | 1096 | HZ1 | LYS | A | 89 | 40.475 | 40.625 | 38.554 | 1.00 | 0.00 | H |
| ATOM | 1097 | HZ2 | LYS | A | 89 | 41.275 | 39.564 | 39.524 | 1.00 | 0.00 | H |
| ATOM | 1098 | HZ3 | LYS | A | 89 | 41.711 | 41.159 | 39.489 | 1.00 | 0.00 | H |
| ATOM | 1099 | C | LYS | A | 89 | 45.121 | 38.252 | 33.789 | 1.00 | 0.00 | C |
| ATOM | 1100 | O | LYS | A | 89 | 45.158 | 37.028 | 33.973 | 1.00 | 0.00 | O |
| ATOM | 1101 | N | LEU | A | 90 | 44.625 | 38.800 | 32.674 | 1.00 | 0.00 | N |
| ATOM | 1102 | H | LEU | A | 90 | 44.693 | 39.804 | 32.566 | 1.00 | 0.00 | H |
| ATOM | 1103 | CA | LEU | A | 90 | 44.040 | 38.030 | 31.575 | 1.00 | 0.00 | C |
| ATOM | 1104 | HA | LEU | A | 90 | 43.288 | 37.356 | 31.988 | 1.00 | 0.00 | H |
| ATOM | 1105 | CB | LEU | A | 90 | 43.351 | 39.008 | 30.602 | 1.00 | 0.00 | C |
| ATOM | 1106 | HB1 | LEU | A | 90 | 42.624 | 39.606 | 31.156 | 1.00 | 0.00 | H |
| ATOM | 1107 | HB2 | LEU | A | 90 | 44.107 | 39.689 | 30.207 | 1.00 | 0.00 | H |
| ATOM | 1108 | CG | LEU | A | 90 | 42.628 | 38.337 | 29.414 | 1.00 | 0.00 | C |
| ATOM | 1109 | HG | LEU | A | 90 | 43.335 | 37.710 | 28.870 | 1.00 | 0.00 | H |
| ATOM | 1110 | CD1 | LEU | A | 90 | 41.453 | 37.460 | 29.876 | 1.00 | 0.00 | C |
| ATOM | 1111 | HD11 | LEU | A | 90 | 40.922 | 37.067 | 29.011 | 1.00 | 0.00 | H |
| ATOM | 1112 | HD12 | LEU | A | 90 | 41.816 | 36.620 | 30.464 | 1.00 | 0.00 | H |
| ATOM | 1113 | HD13 | LEU | A | 90 | 40.759 | 38.051 | 30.477 | 1.00 | 0.00 | H |
| ATOM | 1114 | CD2 | LEU | A | 90 | 42.125 | 39.417 | 28.448 | 1.00 | 0.00 | C |
| ATOM | 1115 | HD21 | LEU | A | 90 | 41.638 | 38.951 | 27.591 | 1.00 | 0.00 | H |
| ATOM | 1116 | HD22 | LEU | A | 90 | 41.413 | 40.070 | 28.953 | 1.00 | 0.00 | H |
| ATOM | 1117 | HD23 | LEU | A | 90 | 42.967 | 40.011 | 28.091 | 1.00 | 0.00 | H |
| ATOM | 1118 | C | LEU | A | 90 | 45.125 | 37.195 | 30.885 | 1.00 | 0.00 | C |
| ATOM | 1119 | O | LEU | A | 90 | 46.176 | 37.719 | 30.535 | 1.00 | 0.00 | O |
| ATOM | 1120 | N | LYS | A | 91 | 44.874 | 35.902 | 30.674 | 1.00 | 0.00 | N |
| ATOM | 1121 | H | LYS | A | 91 | 43.992 | 35.520 | 30.988 | 1.00 | 0.00 | H |
| ATOM | 1122 | CA | LYS | A | 91 | 45.799 | 35.005 | 29.964 | 1.00 | 0.00 | C |
| ATOM | 1123 | HA | LYS | A | 91 | 46.739 | 35.522 | 29.767 | 1.00 | 0.00 | H |
| ATOM | 1124 | CB | LYS | A | 91 | 46.095 | 33.773 | 30.839 | 1.00 | 0.00 | C |
| ATOM | 1125 | HB1 | LYS | A | 91 | 45.158 | 33.254 | 31.053 | 1.00 | 0.00 | H |
| ATOM | 1126 | HB2 | LYS | A | 91 | 46.734 | 33.088 | 30.279 | 1.00 | 0.00 | H |
| ATOM | 1127 | CG | LYS | A | 91 | 46.788 | 34.103 | 32.174 | 1.00 | 0.00 | C |
| ATOM | 1128 | HG1 | LYS | A | 91 | 46.148 | 34.752 | 32.772 | 1.00 | 0.00 | H |
| ATOM | 1129 | HG2 | LYS | A | 91 | 46.915 | 33.170 | 32.720 | 1.00 | 0.00 | H |
| ATOM | 1130 | CD | LYS | A | 91 | 48.163 | 34.767 | 31.997 | 1.00 | 0.00 | C |
| ATOM | 1131 | HD1 | LYS | A | 91 | 48.780 | 34.137 | 31.359 | 1.00 | 0.00 | H |

| | | | | | | | | | | | |
|---|---|---|---|---|---|---|---|---|---|---|---|
| ATOM | 1132 | HD2 | LYS | A | 91 | 48.048 | 35.746 | 31.528 | 1.00 | 0.00 | H |
| ATOM | 1133 | CE | LYS | A | 91 | 48.867 | 34.942 | 33.345 | 1.00 | 0.00 | C |
| ATOM | 1134 | HE1 | LYS | A | 91 | 48.253 | 35.587 | 33.978 | 1.00 | 0.00 | H |
| ATOM | 1135 | HE2 | LYS | A | 91 | 48.963 | 33.967 | 33.831 | 1.00 | 0.00 | H |
| ATOM | 1136 | NZ | LYS | A | 91 | 50.206 | 35.550 | 33.178 | 1.00 | 0.00 | N |
| ATOM | 1137 | HZ1 | LYS | A | 91 | 50.613 | 35.778 | 34.079 | 1.00 | 0.00 | H |
| ATOM | 1138 | HZ2 | LYS | A | 91 | 50.864 | 34.911 | 32.741 | 1.00 | 0.00 | H |
| ATOM | 1139 | HZ3 | LYS | A | 91 | 50.156 | 36.412 | 32.642 | 1.00 | 0.00 | H |
| ATOM | 1140 | C | LYS | A | 91 | 45.228 | 34.604 | 28.597 | 1.00 | 0.00 | C |
| ATOM | 1141 | O | LYS | A | 91 | 44.043 | 34.257 | 28.508 | 1.00 | 0.00 | O |
| ATOM | 1142 | N | THR | A | 92 | 46.050 | 34.655 | 27.542 | 1.00 | 0.00 | N |
| ATOM | 1143 | H | THR | A | 92 | 47.025 | 34.904 | 27.695 | 1.00 | 0.00 | H |
| ATOM | 1144 | CA | THR | A | 92 | 45.642 | 34.386 | 26.153 | 1.00 | 0.00 | C |
| ATOM | 1145 | HA | THR | A | 92 | 44.560 | 34.297 | 26.108 | 1.00 | 0.00 | H |
| ATOM | 1146 | CB | THR | A | 92 | 46.036 | 35.545 | 25.222 | 1.00 | 0.00 | C |
| ATOM | 1147 | HB | THR | A | 92 | 45.942 | 35.222 | 24.183 | 1.00 | 0.00 | H |
| ATOM | 1148 | CG2 | THR | A | 92 | 45.120 | 36.750 | 25.450 | 1.00 | 0.00 | C |
| ATOM | 1149 | HG21 | THR | A | 92 | 45.437 | 37.579 | 24.818 | 1.00 | 0.00 | H |
| ATOM | 1150 | HG22 | THR | A | 92 | 44.095 | 36.484 | 25.194 | 1.00 | 0.00 | H |
| ATOM | 1151 | HG23 | THR | A | 92 | 45.158 | 37.066 | 26.493 | 1.00 | 0.00 | H |
| ATOM | 1152 | OG1 | THR | A | 92 | 47.355 | 35.973 | 25.464 | 1.00 | 0.00 | O |
| ATOM | 1153 | HG1 | THR | A | 92 | 47.593 | 36.608 | 24.766 | 1.00 | 0.00 | H |
| ATOM | 1154 | C | THR | A | 92 | 46.225 | 33.061 | 25.680 | 1.00 | 0.00 | C |
| ATOM | 1155 | O | THR | A | 92 | 47.431 | 32.853 | 25.759 | 1.00 | 0.00 | O |
| ATOM | 1156 | N | LEU | A | 93 | 45.384 | 32.149 | 25.186 | 1.00 | 0.00 | N |
| ATOM | 1157 | H | LEU | A | 93 | 44.398 | 32.387 | 25.131 | 1.00 | 0.00 | H |
| ATOM | 1158 | CA | LEU | A | 93 | 45.818 | 30.879 | 24.587 | 1.00 | 0.00 | C |
| ATOM | 1159 | HA | LEU | A | 93 | 46.907 | 30.836 | 24.594 | 1.00 | 0.00 | H |
| ATOM | 1160 | CB | LEU | A | 93 | 45.303 | 29.655 | 25.372 | 1.00 | 0.00 | C |
| ATOM | 1161 | HB1 | LEU | A | 93 | 44.221 | 29.699 | 25.459 | 1.00 | 0.00 | H |
| ATOM | 1162 | HB2 | LEU | A | 93 | 45.536 | 28.773 | 24.772 | 1.00 | 0.00 | H |
| ATOM | 1163 | CG | LEU | A | 93 | 45.937 | 29.455 | 26.767 | 1.00 | 0.00 | C |
| ATOM | 1164 | HG | LEU | A | 93 | 46.982 | 29.758 | 26.730 | 1.00 | 0.00 | H |
| ATOM | 1165 | CD1 | LEU | A | 93 | 45.233 | 30.263 | 27.870 | 1.00 | 0.00 | C |
| ATOM | 1166 | HD11 | LEU | A | 93 | 45.709 | 30.058 | 28.829 | 1.00 | 0.00 | H |
| ATOM | 1167 | HD12 | LEU | A | 93 | 45.315 | 31.331 | 27.680 | 1.00 | 0.00 | H |
| ATOM | 1168 | HD13 | LEU | A | 93 | 44.180 | 29.988 | 27.924 | 1.00 | 0.00 | H |
| ATOM | 1169 | CD2 | LEU | A | 93 | 45.890 | 27.965 | 27.142 | 1.00 | 0.00 | C |
| ATOM | 1170 | HD21 | LEU | A | 93 | 46.314 | 27.815 | 28.135 | 1.00 | 0.00 | H |
| ATOM | 1171 | HD22 | LEU | A | 93 | 44.860 | 27.607 | 27.132 | 1.00 | 0.00 | H |
| ATOM | 1172 | HD23 | LEU | A | 93 | 46.471 | 27.381 | 26.428 | 1.00 | 0.00 | H |
| ATOM | 1173 | C | LEU | A | 93 | 45.395 | 30.808 | 23.116 | 1.00 | 0.00 | C |
| ATOM | 1174 | O | LEU | A | 93 | 44.413 | 31.427 | 22.708 | 1.00 | 0.00 | O |
| ATOM | 1175 | N | LEU | A | 94 | 46.143 | 30.058 | 22.310 | 1.00 | 0.00 | N |
| ATOM | 1176 | H | LEU | A | 94 | 46.942 | 29.577 | 22.706 | 1.00 | 0.00 | H |
| ATOM | 1177 | CA | LEU | A | 94 | 45.873 | 29.871 | 20.885 | 1.00 | 0.00 | C |
| ATOM | 1178 | HA | LEU | A | 94 | 45.116 | 30.584 | 20.556 | 1.00 | 0.00 | H |
| ATOM | 1179 | CB | LEU | A | 94 | 47.185 | 30.140 | 20.128 | 1.00 | 0.00 | C |
| ATOM | 1180 | HB1 | LEU | A | 94 | 47.561 | 31.121 | 20.426 | 1.00 | 0.00 | H |
| ATOM | 1181 | HB2 | LEU | A | 94 | 47.920 | 29.396 | 20.440 | 1.00 | 0.00 | H |
| ATOM | 1182 | CG | LEU | A | 94 | 47.076 | 30.113 | 18.593 | 1.00 | 0.00 | C |
| ATOM | 1183 | HG | LEU | A | 94 | 46.738 | 29.127 | 18.270 | 1.00 | 0.00 | H |
| ATOM | 1184 | CD1 | LEU | A | 94 | 46.092 | 31.168 | 18.068 | 1.00 | 0.00 | C |
| ATOM | 1185 | HD11 | LEU | A | 94 | 46.139 | 31.220 | 16.983 | 1.00 | 0.00 | H |

| | | | | | | | | | | | |
|---|---|---|---|---|---|---|---|---|---|---|---|
| ATOM | 1186 | HD12 | LEU | A | 94 | 45.076 | 30.892 | 18.340 | 1.00 | 0.00 | H |
| ATOM | 1187 | HD13 | LEU | A | 94 | 46.332 | 32.149 | 18.480 | 1.00 | 0.00 | H |
| ATOM | 1188 | CD2 | LEU | A | 94 | 48.477 | 30.357 | 18.018 | 1.00 | 0.00 | C |
| ATOM | 1189 | HD21 | LEU | A | 94 | 48.450 | 30.362 | 16.930 | 1.00 | 0.00 | H |
| ATOM | 1190 | HD22 | LEU | A | 94 | 48.863 | 31.313 | 18.368 | 1.00 | 0.00 | H |
| ATOM | 1191 | HD23 | LEU | A | 94 | 49.144 | 29.563 | 18.349 | 1.00 | 0.00 | H |
| ATOM | 1192 | C | LEU | A | 94 | 45.343 | 28.454 | 20.651 | 1.00 | 0.00 | C |
| ATOM | 1193 | O | LEU | A | 94 | 46.015 | 27.491 | 21.013 | 1.00 | 0.00 | O |
| ATOM | 1194 | N | ALA | A | 95 | 44.161 | 28.312 | 20.049 | 1.00 | 0.00 | N |
| ATOM | 1195 | H | ALA | A | 95 | 43.661 | 29.130 | 19.720 | 1.00 | 0.00 | H |
| ATOM | 1196 | CA | ALA | A | 95 | 43.614 | 27.003 | 19.697 | 1.00 | 0.00 | C |
| ATOM | 1197 | HA | ALA | A | 95 | 44.002 | 26.273 | 20.398 | 1.00 | 0.00 | H |
| ATOM | 1198 | CB | ALA | A | 95 | 42.090 | 27.015 | 19.851 | 1.00 | 0.00 | C |
| ATOM | 1199 | HB1 | ALA | A | 95 | 41.690 | 26.033 | 19.594 | 1.00 | 0.00 | H |
| ATOM | 1200 | HB2 | ALA | A | 95 | 41.829 | 27.223 | 20.886 | 1.00 | 0.00 | H |
| ATOM | 1201 | HB3 | ALA | A | 95 | 41.646 | 27.771 | 19.205 | 1.00 | 0.00 | H |
| ATOM | 1202 | C | ALA | A | 95 | 44.053 | 26.589 | 18.286 | 1.00 | 0.00 | C |
| ATOM | 1203 | O | ALA | A | 95 | 43.982 | 27.390 | 17.355 | 1.00 | 0.00 | O |
| ATOM | 1204 | N | ILE | A | 96 | 44.470 | 25.335 | 18.102 | 1.00 | 0.00 | N |
| ATOM | 1205 | H | ILE | A | 96 | 44.493 | 24.725 | 18.916 | 1.00 | 0.00 | H |
| ATOM | 1206 | CA | ILE | A | 96 | 44.783 | 24.764 | 16.777 | 1.00 | 0.00 | C |
| ATOM | 1207 | HA | ILE | A | 96 | 44.803 | 25.574 | 16.047 | 1.00 | 0.00 | H |
| ATOM | 1208 | CB | ILE | A | 96 | 46.197 | 24.128 | 16.760 | 1.00 | 0.00 | C |
| ATOM | 1209 | HB | ILE | A | 96 | 46.903 | 24.923 | 17.009 | 1.00 | 0.00 | H |
| ATOM | 1210 | CG2 | ILE | A | 96 | 46.384 | 23.008 | 17.803 | 1.00 | 0.00 | C |
| ATOM | 1211 | HG21 | ILE | A | 96 | 47.422 | 22.681 | 17.824 | 1.00 | 0.00 | H |
| ATOM | 1212 | HG22 | ILE | A | 96 | 46.142 | 23.380 | 18.796 | 1.00 | 0.00 | H |
| ATOM | 1213 | HG23 | ILE | A | 96 | 45.748 | 22.155 | 17.570 | 1.00 | 0.00 | H |
| ATOM | 1214 | CG1 | ILE | A | 96 | 46.539 | 23.634 | 15.334 | 1.00 | 0.00 | C |
| ATOM | 1215 | HG11 | ILE | A | 96 | 45.891 | 22.803 | 15.056 | 1.00 | 0.00 | H |
| ATOM | 1216 | HG12 | ILE | A | 96 | 46.367 | 24.446 | 14.632 | 1.00 | 0.00 | H |
| ATOM | 1217 | CD | ILE | A | 96 | 47.989 | 23.192 | 15.154 | 1.00 | 0.00 | C |
| ATOM | 1218 | HD1 | ILE | A | 96 | 48.185 | 23.021 | 14.096 | 1.00 | 0.00 | H |
| ATOM | 1219 | HD2 | ILE | A | 96 | 48.643 | 23.980 | 15.516 | 1.00 | 0.00 | H |
| ATOM | 1220 | HD3 | ILE | A | 96 | 48.183 | 22.271 | 15.701 | 1.00 | 0.00 | H |
| ATOM | 1221 | C | ILE | A | 96 | 43.689 | 23.797 | 16.311 | 1.00 | 0.00 | C |
| ATOM | 1222 | O | ILE | A | 96 | 43.276 | 22.919 | 17.074 | 1.00 | 0.00 | O |
| ATOM | 1223 | N | GLY | A | 97 | 43.276 | 23.928 | 15.041 | 1.00 | 0.00 | N |
| ATOM | 1224 | H | GLY | A | 97 | 43.664 | 24.689 | 14.490 | 1.00 | 0.00 | H |
| ATOM | 1225 | CA | GLY | A | 97 | 42.418 | 22.966 | 14.341 | 1.00 | 0.00 | C |
| ATOM | 1226 | HA1 | GLY | A | 97 | 42.826 | 22.805 | 13.345 | 1.00 | 0.00 | H |
| ATOM | 1227 | HA2 | GLY | A | 97 | 42.419 | 22.015 | 14.872 | 1.00 | 0.00 | H |
| ATOM | 1228 | C | GLY | A | 97 | 40.972 | 23.429 | 14.152 | 1.00 | 0.00 | C |
| ATOM | 1229 | O | GLY | A | 97 | 40.695 | 24.280 | 13.301 | 1.00 | 0.00 | O |
| ATOM | 1230 | N | GLY | A | 98 | 40.048 | 22.785 | 14.870 | 1.00 | 0.00 | N |
| ATOM | 1231 | H | GLY | A | 98 | 40.386 | 22.144 | 15.570 | 1.00 | 0.00 | H |
| ATOM | 1232 | CA | GLY | A | 98 | 38.591 | 22.878 | 14.713 | 1.00 | 0.00 | C |
| ATOM | 1233 | HA1 | GLY | A | 98 | 38.131 | 22.859 | 15.702 | 1.00 | 0.00 | H |
| ATOM | 1234 | HA2 | GLY | A | 98 | 38.318 | 23.809 | 14.219 | 1.00 | 0.00 | H |
| ATOM | 1235 | C | GLY | A | 98 | 37.982 | 21.727 | 13.907 | 1.00 | 0.00 | C |
| ATOM | 1236 | O | GLY | A | 98 | 38.684 | 20.929 | 13.285 | 1.00 | 0.00 | O |
| ATOM | 1237 | N | TRP | A | 99 | 36.651 | 21.647 | 13.910 | 1.00 | 0.00 | N |
| ATOM | 1238 | H | TRP | A | 99 | 36.151 | 22.345 | 14.439 | 1.00 | 0.00 | H |
| ATOM | 1239 | CA | TRP | A | 99 | 35.859 | 20.646 | 13.181 | 1.00 | 0.00 | C |

| | | | | | | | | | | | |
|---|---|---|---|---|---|---|---|---|---|---|---|
| ATOM | 1240 | HA | TRP | A | 99 | 36.097 | 19.654 | 13.563 | 1.00 | 0.00 | H |
| ATOM | 1241 | CB | TRP | A | 99 | 34.373 | 20.925 | 13.453 | 1.00 | 0.00 | C |
| ATOM | 1242 | HB1 | TRP | A | 99 | 34.149 | 20.634 | 14.478 | 1.00 | 0.00 | H |
| ATOM | 1243 | HB2 | TRP | A | 99 | 34.199 | 21.998 | 13.367 | 1.00 | 0.00 | H |
| ATOM | 1244 | CG | TRP | A | 99 | 33.399 | 20.247 | 12.536 | 1.00 | 0.00 | C |
| ATOM | 1245 | CD1 | TRP | A | 99 | 32.562 | 20.892 | 11.694 | 1.00 | 0.00 | C |
| ATOM | 1246 | HD1 | TRP | A | 99 | 32.494 | 21.968 | 11.585 | 1.00 | 0.00 | H |
| ATOM | 1247 | NE1 | TRP | A | 99 | 31.835 | 19.980 | 10.961 | 1.00 | 0.00 | N |
| ATOM | 1248 | HE1 | TRP | A | 99 | 31.172 | 20.258 | 10.241 | 1.00 | 0.00 | H |
| ATOM | 1249 | CE2 | TRP | A | 99 | 32.161 | 18.684 | 11.295 | 1.00 | 0.00 | C |
| ATOM | 1250 | CZ2 | TRP | A | 99 | 31.711 | 17.435 | 10.839 | 1.00 | 0.00 | C |
| ATOM | 1251 | HZ2 | TRP | A | 99 | 30.959 | 17.375 | 10.066 | 1.00 | 0.00 | H |
| ATOM | 1252 | CH2 | TRP | A | 99 | 32.269 | 16.267 | 11.385 | 1.00 | 0.00 | C |
| ATOM | 1253 | HH2 | TRP | A | 99 | 31.947 | 15.296 | 11.037 | 1.00 | 0.00 | H |
| ATOM | 1254 | CZ3 | TRP | A | 99 | 33.272 | 16.366 | 12.364 | 1.00 | 0.00 | C |
| ATOM | 1255 | HZ3 | TRP | A | 99 | 33.717 | 15.469 | 12.767 | 1.00 | 0.00 | H |
| ATOM | 1256 | CE3 | TRP | A | 99 | 33.723 | 17.625 | 12.803 | 1.00 | 0.00 | C |
| ATOM | 1257 | HE3 | TRP | A | 99 | 34.518 | 17.678 | 13.528 | 1.00 | 0.00 | H |
| ATOM | 1258 | CD2 | TRP | A | 99 | 33.168 | 18.820 | 12.298 | 1.00 | 0.00 | C |
| ATOM | 1259 | C | TRP | A | 99 | 36.153 | 20.626 | 11.670 | 1.00 | 0.00 | C |
| ATOM | 1260 | O | TRP | A | 99 | 36.439 | 19.561 | 11.115 | 1.00 | 0.00 | O |
| ATOM | 1261 | N | ASN | A | 100 | 36.133 | 21.794 | 11.015 | 1.00 | 0.00 | N |
| ATOM | 1262 | H | ASN | A | 100 | 35.914 | 22.630 | 11.540 | 1.00 | 0.00 | H |
| ATOM | 1263 | CA | ASN | A | 100 | 36.318 | 21.903 | 9.561 | 1.00 | 0.00 | C |
| ATOM | 1264 | HA | ASN | A | 100 | 35.601 | 21.244 | 9.067 | 1.00 | 0.00 | H |
| ATOM | 1265 | CB | ASN | A | 100 | 36.053 | 23.348 | 9.102 | 1.00 | 0.00 | C |
| ATOM | 1266 | HB1 | ASN | A | 100 | 36.660 | 24.033 | 9.693 | 1.00 | 0.00 | H |
| ATOM | 1267 | HB2 | ASN | A | 100 | 36.361 | 23.450 | 8.061 | 1.00 | 0.00 | H |
| ATOM | 1268 | CG | ASN | A | 100 | 34.599 | 23.795 | 9.187 | 1.00 | 0.00 | C |
| ATOM | 1269 | OD1 | ASN | A | 100 | 33.679 | 23.032 | 9.441 | 1.00 | 0.00 | O |
| ATOM | 1270 | ND2 | ASN | A | 100 | 34.345 | 25.062 | 8.937 | 1.00 | 0.00 | N |
| ATOM | 1271 | HD21 | ASN | A | 100 | 35.084 | 25.661 | 8.587 | 1.00 | 0.00 | H |
| ATOM | 1272 | HD22 | ASN | A | 100 | 33.403 | 25.410 | 9.028 | 1.00 | 0.00 | H |
| ATOM | 1273 | C | ASN | A | 100 | 37.725 | 21.481 | 9.101 | 1.00 | 0.00 | C |
| ATOM | 1274 | O | ASN | A | 100 | 37.883 | 21.004 | 7.976 | 1.00 | 0.00 | O |
| ATOM | 1275 | N | PHE | A | 101 | 38.741 | 21.645 | 9.959 | 1.00 | 0.00 | N |
| ATOM | 1276 | H | PHE | A | 101 | 38.520 | 22.027 | 10.870 | 1.00 | 0.00 | H |
| ATOM | 1277 | CA | PHE | A | 101 | 40.162 | 21.468 | 9.635 | 1.00 | 0.00 | C |
| ATOM | 1278 | HA | PHE | A | 101 | 40.424 | 22.163 | 8.836 | 1.00 | 0.00 | H |
| ATOM | 1279 | CB | PHE | A | 101 | 40.965 | 21.841 | 10.888 | 1.00 | 0.00 | C |
| ATOM | 1280 | HB1 | PHE | A | 101 | 40.662 | 22.839 | 11.197 | 1.00 | 0.00 | H |
| ATOM | 1281 | HB2 | PHE | A | 101 | 40.705 | 21.153 | 11.692 | 1.00 | 0.00 | H |
| ATOM | 1282 | CG | PHE | A | 101 | 42.474 | 21.856 | 10.737 | 1.00 | 0.00 | C |
| ATOM | 1283 | CD1 | PHE | A | 101 | 43.109 | 22.940 | 10.103 | 1.00 | 0.00 | C |
| ATOM | 1284 | HD1 | PHE | A | 101 | 42.519 | 23.740 | 9.684 | 1.00 | 0.00 | H |
| ATOM | 1285 | CE1 | PHE | A | 101 | 44.512 | 23.000 | 10.039 | 1.00 | 0.00 | C |
| ATOM | 1286 | HE1 | PHE | A | 101 | 44.991 | 23.844 | 9.566 | 1.00 | 0.00 | H |
| ATOM | 1287 | CZ | PHE | A | 101 | 45.284 | 21.976 | 10.612 | 1.00 | 0.00 | C |
| ATOM | 1288 | HZ | PHE | A | 101 | 46.360 | 22.028 | 10.581 | 1.00 | 0.00 | H |
| ATOM | 1289 | CE2 | PHE | A | 101 | 44.655 | 20.887 | 11.236 | 1.00 | 0.00 | C |
| ATOM | 1290 | HE2 | PHE | A | 101 | 45.248 | 20.100 | 11.681 | 1.00 | 0.00 | H |
| ATOM | 1291 | CD2 | PHE | A | 101 | 43.251 | 20.825 | 11.297 | 1.00 | 0.00 | C |
| ATOM | 1292 | HD2 | PHE | A | 101 | 42.772 | 19.994 | 11.792 | 1.00 | 0.00 | H |
| ATOM | 1293 | C | PHE | A | 101 | 40.531 | 20.059 | 9.145 | 1.00 | 0.00 | C |

| | | | | | | | | | | | |
|---|---|---|---|---|---|---|---|---|---|---|---|
| ATOM | 1294 | O | PHE | A | 101 | 41.474 | 19.928 | 8.366 | 1.00 | 0.00 | O |
| ATOM | 1295 | N | GLY | A | 102 | 39.801 | 19.021 | 9.567 | 1.00 | 0.00 | N |
| ATOM | 1296 | H | GLY | A | 102 | 39.041 | 19.200 | 10.208 | 1.00 | 0.00 | H |
| ATOM | 1297 | CA | GLY | A | 102 | 40.080 | 17.628 | 9.202 | 1.00 | 0.00 | C |
| ATOM | 1298 | HA1 | GLY | A | 102 | 39.170 | 17.045 | 9.320 | 1.00 | 0.00 | H |
| ATOM | 1299 | HA2 | GLY | A | 102 | 40.401 | 17.572 | 8.162 | 1.00 | 0.00 | H |
| ATOM | 1300 | C | GLY | A | 102 | 41.171 | 16.994 | 10.069 | 1.00 | 0.00 | C |
| ATOM | 1301 | O | GLY | A | 102 | 41.799 | 17.663 | 10.894 | 1.00 | 0.00 | O |
| ATOM | 1302 | N | THR | A | 103 | 41.426 | 15.693 | 9.879 | 1.00 | 0.00 | N |
| ATOM | 1303 | H | THR | A | 103 | 40.890 | 15.208 | 9.165 | 1.00 | 0.00 | H |
| ATOM | 1304 | CA | THR | A | 103 | 42.420 | 14.930 | 10.662 | 1.00 | 0.00 | C |
| ATOM | 1305 | HA | THR | A | 103 | 42.564 | 15.432 | 11.620 | 1.00 | 0.00 | H |
| ATOM | 1306 | CB | THR | A | 103 | 41.926 | 13.518 | 10.983 | 1.00 | 0.00 | C |
| ATOM | 1307 | HB | THR | A | 103 | 42.720 | 13.009 | 11.532 | 1.00 | 0.00 | H |
| ATOM | 1308 | CG2 | THR | A | 103 | 40.668 | 13.521 | 11.846 | 1.00 | 0.00 | C |
| ATOM | 1309 | HG21 | THR | A | 103 | 40.400 | 12.499 | 12.106 | 1.00 | 0.00 | H |
| ATOM | 1310 | HG22 | THR | A | 103 | 40.851 | 14.083 | 12.758 | 1.00 | 0.00 | H |
| ATOM | 1311 | HG23 | THR | A | 103 | 39.834 | 13.979 | 11.318 | 1.00 | 0.00 | H |
| ATOM | 1312 | OG1 | THR | A | 103 | 41.672 | 12.801 | 9.802 | 1.00 | 0.00 | O |
| ATOM | 1313 | HG1 | THR | A | 103 | 40.734 | 12.890 | 9.576 | 1.00 | 0.00 | H |
| ATOM | 1314 | C | THR | A | 103 | 43.803 | 14.856 | 10.020 | 1.00 | 0.00 | C |
| ATOM | 1315 | O | THR | A | 103 | 44.781 | 14.931 | 10.759 | 1.00 | 0.00 | O |
| ATOM | 1316 | N | GLN | A | 104 | 43.909 | 14.794 | 8.691 | 1.00 | 0.00 | N |
| ATOM | 1317 | H | GLN | A | 104 | 43.042 | 14.781 | 8.169 | 1.00 | 0.00 | H |
| ATOM | 1318 | CA | GLN | A | 104 | 45.133 | 14.404 | 7.969 | 1.00 | 0.00 | C |
| ATOM | 1319 | HA | GLN | A | 104 | 45.322 | 13.356 | 8.210 | 1.00 | 0.00 | H |
| ATOM | 1320 | CB | GLN | A | 104 | 44.869 | 14.512 | 6.455 | 1.00 | 0.00 | C |
| ATOM | 1321 | HB1 | GLN | A | 104 | 43.892 | 14.082 | 6.229 | 1.00 | 0.00 | H |
| ATOM | 1322 | HB2 | GLN | A | 104 | 44.850 | 15.565 | 6.171 | 1.00 | 0.00 | H |
| ATOM | 1323 | CG | GLN | A | 104 | 45.917 | 13.792 | 5.590 | 1.00 | 0.00 | C |
| ATOM | 1324 | HG1 | GLN | A | 104 | 45.693 | 13.987 | 4.543 | 1.00 | 0.00 | H |
| ATOM | 1325 | HG2 | GLN | A | 104 | 46.914 | 14.181 | 5.800 | 1.00 | 0.00 | H |
| ATOM | 1326 | CD | GLN | A | 104 | 45.886 | 12.282 | 5.806 | 1.00 | 0.00 | C |
| ATOM | 1327 | OE1 | GLN | A | 104 | 44.965 | 11.597 | 5.369 | 1.00 | 0.00 | O |
| ATOM | 1328 | NE2 | GLN | A | 104 | 46.832 | 11.724 | 6.527 | 1.00 | 0.00 | N |
| ATOM | 1329 | HE21 | GLN | A | 104 | 47.640 | 12.255 | 6.840 | 1.00 | 0.00 | H |
| ATOM | 1330 | HE22 | GLN | A | 104 | 46.747 | 10.731 | 6.730 | 1.00 | 0.00 | H |
| ATOM | 1331 | C | GLN | A | 104 | 46.420 | 15.164 | 8.349 | 1.00 | 0.00 | C |
| ATOM | 1332 | O | GLN | A | 104 | 47.437 | 14.521 | 8.596 | 1.00 | 0.00 | O |
| ATOM | 1333 | N | LYS | A | 105 | 46.415 | 16.503 | 8.420 | 1.00 | 0.00 | N |
| ATOM | 1334 | H | LYS | A | 105 | 45.564 | 17.001 | 8.184 | 1.00 | 0.00 | H |
| ATOM | 1335 | CA | LYS | A | 105 | 47.609 | 17.291 | 8.783 | 1.00 | 0.00 | C |
| ATOM | 1336 | HA | LYS | A | 105 | 48.401 | 17.082 | 8.062 | 1.00 | 0.00 | H |
| ATOM | 1337 | CB | LYS | A | 105 | 47.295 | 18.797 | 8.757 | 1.00 | 0.00 | C |
| ATOM | 1338 | HB1 | LYS | A | 105 | 46.396 | 18.999 | 9.339 | 1.00 | 0.00 | H |
| ATOM | 1339 | HB2 | LYS | A | 105 | 48.124 | 19.322 | 9.236 | 1.00 | 0.00 | H |
| ATOM | 1340 | CG | LYS | A | 105 | 47.130 | 19.373 | 7.345 | 1.00 | 0.00 | C |
| ATOM | 1341 | HG1 | LYS | A | 105 | 48.037 | 19.179 | 6.771 | 1.00 | 0.00 | H |
| ATOM | 1342 | HG2 | LYS | A | 105 | 46.282 | 18.907 | 6.843 | 1.00 | 0.00 | H |
| ATOM | 1343 | CD | LYS | A | 105 | 46.905 | 20.887 | 7.446 | 1.00 | 0.00 | C |
| ATOM | 1344 | HD1 | LYS | A | 105 | 45.966 | 21.085 | 7.964 | 1.00 | 0.00 | H |
| ATOM | 1345 | HD2 | LYS | A | 105 | 47.718 | 21.322 | 8.025 | 1.00 | 0.00 | H |
| ATOM | 1346 | CE | LYS | A | 105 | 46.877 | 21.523 | 6.055 | 1.00 | 0.00 | C |
| ATOM | 1347 | HE1 | LYS | A | 105 | 47.793 | 21.243 | 5.527 | 1.00 | 0.00 | H |

```
ATOM   1348  HE2 LYS A 105      46.029  21.124   5.492  1.00  0.00           H
ATOM   1349  NZ  LYS A 105      46.786  23.001   6.123  1.00  0.00           N
ATOM   1350  HZ1 LYS A 105      46.932  23.409   5.206  1.00  0.00           H
ATOM   1351  HZ2 LYS A 105      45.878  23.299   6.467  1.00  0.00           H
ATOM   1352  HZ3 LYS A 105      47.495  23.397   6.733  1.00  0.00           H
ATOM   1353  C   LYS A 105      48.178  16.918  10.162  1.00  0.00           C
ATOM   1354  O   LYS A 105      49.393  16.787  10.310  1.00  0.00           O
ATOM   1355  N   PHE A 106      47.316  16.731  11.169  1.00  0.00           N
ATOM   1356  H   PHE A 106      46.338  16.899  10.989  1.00  0.00           H
ATOM   1357  CA  PHE A 106      47.717  16.235  12.485  1.00  0.00           C
ATOM   1358  HA  PHE A 106      48.503  16.879  12.880  1.00  0.00           H
ATOM   1359  CB  PHE A 106      46.531  16.261  13.459  1.00  0.00           C
ATOM   1360  HB1 PHE A 106      45.704  15.690  13.041  1.00  0.00           H
ATOM   1361  HB2 PHE A 106      46.863  15.738  14.349  1.00  0.00           H
ATOM   1362  CG  PHE A 106      46.023  17.606  13.948  1.00  0.00           C
ATOM   1363  CD1 PHE A 106      46.853  18.429  14.734  1.00  0.00           C
ATOM   1364  HD1 PHE A 106      47.873  18.136  14.932  1.00  0.00           H
ATOM   1365  CE1 PHE A 106      46.345  19.611  15.303  1.00  0.00           C
ATOM   1366  HE1 PHE A 106      46.979  20.229  15.921  1.00  0.00           H
ATOM   1367  CZ  PHE A 106      45.003  19.972  15.095  1.00  0.00           C
ATOM   1368  HZ  PHE A 106      44.602  20.869  15.548  1.00  0.00           H
ATOM   1369  CE2 PHE A 106      44.172  19.153  14.313  1.00  0.00           C
ATOM   1370  HE2 PHE A 106      43.139  19.426  14.159  1.00  0.00           H
ATOM   1371  CD2 PHE A 106      44.677  17.969  13.748  1.00  0.00           C
ATOM   1372  HD2 PHE A 106      44.017  17.327  13.185  1.00  0.00           H
ATOM   1373  C   PHE A 106      48.297  14.810  12.399  1.00  0.00           C
ATOM   1374  O   PHE A 106      49.389  14.591  12.927  1.00  0.00           O
ATOM   1375  N   THR A 107      47.607  13.878  11.715  1.00  0.00           N
ATOM   1376  H   THR A 107      46.714  14.148  11.316  1.00  0.00           H
ATOM   1377  CA  THR A 107      48.038  12.478  11.504  1.00  0.00           C
ATOM   1378  HA  THR A 107      48.077  11.964  12.465  1.00  0.00           H
ATOM   1379  CB  THR A 107      47.026  11.744  10.612  1.00  0.00           C
ATOM   1380  HB  THR A 107      46.932  12.267   9.659  1.00  0.00           H
ATOM   1381  CG2 THR A 107      47.400  10.289  10.331  1.00  0.00           C
ATOM   1382 HG21 THR A 107      46.606   9.816   9.756  1.00  0.00           H
ATOM   1383 HG22 THR A 107      48.319  10.241   9.746  1.00  0.00           H
ATOM   1384 HG23 THR A 107      47.537   9.746  11.265  1.00  0.00           H
ATOM   1385  OG1 THR A 107      45.771  11.723  11.254  1.00  0.00           O
ATOM   1386  HG1 THR A 107      45.863  11.121  12.028  1.00  0.00           H
ATOM   1387  C   THR A 107      49.442  12.410  10.904  1.00  0.00           C
ATOM   1388  O   THR A 107      50.285  11.684  11.429  1.00  0.00           O
ATOM   1389  N   ASP A 108      49.704  13.203   9.862  1.00  0.00           N
ATOM   1390  H   ASP A 108      48.928  13.725   9.463  1.00  0.00           H
ATOM   1391  CA  ASP A 108      51.022  13.353   9.228  1.00  0.00           C
ATOM   1392  HA  ASP A 108      51.380  12.380   8.889  1.00  0.00           H
ATOM   1393  CB  ASP A 108      50.912  14.308   8.021  1.00  0.00           C
ATOM   1394  HB1 ASP A 108      50.466  15.246   8.353  1.00  0.00           H
ATOM   1395  HB2 ASP A 108      51.924  14.530   7.688  1.00  0.00           H
ATOM   1396  CG  ASP A 108      50.160  13.827   6.778  1.00  0.00           C
ATOM   1397  OD1 ASP A 108      49.628  12.689   6.745  1.00  0.00           O
ATOM   1398  OD2 ASP A 108      50.177  14.614   5.794  1.00  0.00           O
ATOM   1399  C   ASP A 108      52.069  13.936  10.201  1.00  0.00           C
ATOM   1400  O   ASP A 108      53.204  13.456  10.273  1.00  0.00           O
ATOM   1401  N   MET A 109      51.691  14.978  10.953  1.00  0.00           N
```

```
ATOM   1402  H   MET A 109      50.759  15.351  10.804  1.00  0.00           H
ATOM   1403  CA  MET A 109      52.580  15.689  11.880  1.00  0.00           C
ATOM   1404  HA  MET A 109      53.484  15.955  11.339  1.00  0.00           H
ATOM   1405  CB  MET A 109      51.908  16.999  12.323  1.00  0.00           C
ATOM   1406  HB1 MET A 109      51.707  17.586  11.427  1.00  0.00           H
ATOM   1407  HB2 MET A 109      50.958  16.790  12.813  1.00  0.00           H
ATOM   1408  CG  MET A 109      52.789  17.847  13.251  1.00  0.00           C
ATOM   1409  HG1 MET A 109      53.828  17.748  12.933  1.00  0.00           H
ATOM   1410  HG2 MET A 109      52.512  18.894  13.122  1.00  0.00           H
ATOM   1411  SD  MET A 109      52.695  17.455  15.020  1.00  0.00           S
ATOM   1412  CE  MET A 109      51.113  18.234  15.414  1.00  0.00           C
ATOM   1413  HE1 MET A 109      50.959  18.207  16.491  1.00  0.00           H
ATOM   1414  HE2 MET A 109      51.129  19.276  15.093  1.00  0.00           H
ATOM   1415  HE3 MET A 109      50.305  17.700  14.917  1.00  0.00           H
ATOM   1416  C   MET A 109      53.023  14.818  13.056  1.00  0.00           C
ATOM   1417  O   MET A 109      54.189  14.866  13.440  1.00  0.00           O
ATOM   1418  N   VAL A 110      52.146  13.968  13.587  1.00  0.00           N
ATOM   1419  H   VAL A 110      51.185  14.005  13.244  1.00  0.00           H
ATOM   1420  CA  VAL A 110      52.513  13.079  14.703  1.00  0.00           C
ATOM   1421  HA  VAL A 110      53.175  13.646  15.353  1.00  0.00           H
ATOM   1422  CB  VAL A 110      51.311  12.681  15.576  1.00  0.00           C
ATOM   1423  HB  VAL A 110      51.716  12.072  16.379  1.00  0.00           H
ATOM   1424  CG1 VAL A 110      50.680  13.905  16.247  1.00  0.00           C
ATOM   1425 HG11 VAL A 110      49.950  13.579  16.987  1.00  0.00           H
ATOM   1426 HG12 VAL A 110      51.459  14.474  16.756  1.00  0.00           H
ATOM   1427 HG13 VAL A 110      50.190  14.544  15.512  1.00  0.00           H
ATOM   1428  CG2 VAL A 110      50.267  11.852  14.814  1.00  0.00           C
ATOM   1429 HG21 VAL A 110      49.556  11.419  15.512  1.00  0.00           H
ATOM   1430 HG22 VAL A 110      49.728  12.470  14.106  1.00  0.00           H
ATOM   1431 HG23 VAL A 110      50.750  11.047  14.270  1.00  0.00           H
ATOM   1432  C   VAL A 110      53.298  11.825  14.284  1.00  0.00           C
ATOM   1433  O   VAL A 110      53.845  11.145  15.154  1.00  0.00           O
ATOM   1434  N   ALA A 111      53.369  11.502  12.984  1.00  0.00           N
ATOM   1435  H   ALA A 111      52.902  12.106  12.315  1.00  0.00           H
ATOM   1436  CA  ALA A 111      53.898  10.228  12.483  1.00  0.00           C
ATOM   1437  HA  ALA A 111      53.332   9.418  12.946  1.00  0.00           H
ATOM   1438  CB  ALA A 111      53.643  10.178  10.971  1.00  0.00           C
ATOM   1439  HB1 ALA A 111      53.960   9.214  10.574  1.00  0.00           H
ATOM   1440  HB2 ALA A 111      52.578  10.305  10.765  1.00  0.00           H
ATOM   1441  HB3 ALA A 111      54.200  10.967  10.464  1.00  0.00           H
ATOM   1442  C   ALA A 111      55.385   9.970  12.806  1.00  0.00           C
ATOM   1443  O   ALA A 111      55.790   8.809  12.926  1.00  0.00           O
ATOM   1444  N   THR A 112      56.195  11.026  12.972  1.00  0.00           N
ATOM   1445  H   THR A 112      55.776  11.947  12.934  1.00  0.00           H
ATOM   1446  CA  THR A 112      57.610  10.943  13.380  1.00  0.00           C
ATOM   1447  HA  THR A 112      57.776   9.991  13.880  1.00  0.00           H
ATOM   1448  CB  THR A 112      58.591  11.021  12.200  1.00  0.00           C
ATOM   1449  HB  THR A 112      59.609  10.982  12.593  1.00  0.00           H
ATOM   1450  CG2 THR A 112      58.427   9.877  11.206  1.00  0.00           C
ATOM   1451 HG21 THR A 112      59.204   9.945  10.444  1.00  0.00           H
ATOM   1452 HG22 THR A 112      58.529   8.922  11.722  1.00  0.00           H
ATOM   1453 HG23 THR A 112      57.453   9.918  10.719  1.00  0.00           H
ATOM   1454  OG1 THR A 112      58.428  12.229  11.503  1.00  0.00           O
ATOM   1455  HG1 THR A 112      57.619  12.153  10.981  1.00  0.00           H
```

```
ATOM   1456  C   THR A 112      57.941  12.024  14.393  1.00  0.00           C
ATOM   1457  O   THR A 112      57.322  13.086  14.402  1.00  0.00           O
ATOM   1458  N   ALA A 113      58.943  11.770  15.233  1.00  0.00           N
ATOM   1459  H   ALA A 113      59.434  10.887  15.156  1.00  0.00           H
ATOM   1460  CA  ALA A 113      59.392  12.739  16.232  1.00  0.00           C
ATOM   1461  HA  ALA A 113      58.538  13.043  16.840  1.00  0.00           H
ATOM   1462  CB  ALA A 113      60.408  12.054  17.140  1.00  0.00           C
ATOM   1463  HB1 ALA A 113      60.711  12.737  17.935  1.00  0.00           H
ATOM   1464  HB2 ALA A 113      59.941  11.176  17.577  1.00  0.00           H
ATOM   1465  HB3 ALA A 113      61.290  11.753  16.575  1.00  0.00           H
ATOM   1466  C   ALA A 113      59.958  14.016  15.600  1.00  0.00           C
ATOM   1467  O   ALA A 113      59.771  15.100  16.149  1.00  0.00           O
ATOM   1468  N   ASN A 114      60.599  13.925  14.432  1.00  0.00           N
ATOM   1469  H   ASN A 114      60.805  13.003  14.066  1.00  0.00           H
ATOM   1470  CA  ASN A 114      61.026  15.098  13.671  1.00  0.00           C
ATOM   1471  HA  ASN A 114      61.628  15.735  14.320  1.00  0.00           H
ATOM   1472  CB  ASN A 114      61.914  14.660  12.503  1.00  0.00           C
ATOM   1473  HB1 ASN A 114      62.846  14.247  12.888  1.00  0.00           H
ATOM   1474  HB2 ASN A 114      61.399  13.884  11.946  1.00  0.00           H
ATOM   1475  CG  ASN A 114      62.237  15.825  11.575  1.00  0.00           C
ATOM   1476  OD1 ASN A 114      62.964  16.747  11.925  1.00  0.00           O
ATOM   1477  ND2 ASN A 114      61.690  15.841  10.377  1.00  0.00           N
ATOM   1478 HD21 ASN A 114      61.123  15.065  10.061  1.00  0.00           H
ATOM   1479 HD22 ASN A 114      61.875  16.642   9.786  1.00  0.00           H
ATOM   1480  C   ASN A 114      59.818  15.923  13.196  1.00  0.00           C
ATOM   1481  O   ASN A 114      59.817  17.136  13.369  1.00  0.00           O
ATOM   1482  N   ASN A 115      58.749  15.296  12.690  1.00  0.00           N
ATOM   1483  H   ASN A 115      58.762  14.289  12.588  1.00  0.00           H
ATOM   1484  CA  ASN A 115      57.531  16.026  12.312  1.00  0.00           C
ATOM   1485  HA  ASN A 115      57.807  16.816  11.611  1.00  0.00           H
ATOM   1486  CB  ASN A 115      56.551  15.088  11.595  1.00  0.00           C
ATOM   1487  HB1 ASN A 115      56.327  14.237  12.235  1.00  0.00           H
ATOM   1488  HB2 ASN A 115      55.630  15.639  11.420  1.00  0.00           H
ATOM   1489  CG  ASN A 115      57.012  14.563  10.242  1.00  0.00           C
ATOM   1490  OD1 ASN A 115      58.074  14.892   9.726  1.00  0.00           O
ATOM   1491  ND2 ASN A 115      56.210  13.716   9.623  1.00  0.00           N
ATOM   1492 HD21 ASN A 115      55.276  13.540   9.978  1.00  0.00           H
ATOM   1493 HD22 ASN A 115      56.446  13.477   8.665  1.00  0.00           H
ATOM   1494  C   ASN A 115      56.874  16.712  13.534  1.00  0.00           C
ATOM   1495  O   ASN A 115      56.463  17.871  13.436  1.00  0.00           O
ATOM   1496  N   ARG A 116      56.886  16.068  14.713  1.00  0.00           N
ATOM   1497  H   ARG A 116      57.160  15.091  14.719  1.00  0.00           H
ATOM   1498  CA  ARG A 116      56.467  16.696  15.979  1.00  0.00           C
ATOM   1499  HA  ARG A 116      55.483  17.139  15.821  1.00  0.00           H
ATOM   1500  CB  ARG A 116      56.334  15.653  17.101  1.00  0.00           C
ATOM   1501  HB1 ARG A 116      57.279  15.126  17.232  1.00  0.00           H
ATOM   1502  HB2 ARG A 116      56.100  16.177  18.030  1.00  0.00           H
ATOM   1503  CG  ARG A 116      55.211  14.645  16.815  1.00  0.00           C
ATOM   1504  HG1 ARG A 116      54.317  15.195  16.518  1.00  0.00           H
ATOM   1505  HG2 ARG A 116      55.501  13.993  15.993  1.00  0.00           H
ATOM   1506  CD  ARG A 116      54.844  13.785  18.030  1.00  0.00           C
ATOM   1507  HD1 ARG A 116      54.445  14.438  18.808  1.00  0.00           H
ATOM   1508  HD2 ARG A 116      54.052  13.090  17.747  1.00  0.00           H
ATOM   1509  NE  ARG A 116      55.992  13.048  18.586  1.00  0.00           N
```

```
ATOM   1510  HE  ARG A 116      56.431  13.464  19.408  1.00  0.00           H
ATOM   1511  CZ  ARG A 116      56.398  11.824  18.277  1.00  0.00           C
ATOM   1512  NH1 ARG A 116      55.881  11.135  17.282  1.00  0.00           N
ATOM   1513 HH11 ARG A 116      55.165  11.528  16.689  1.00  0.00           H
ATOM   1514 HH12 ARG A 116      56.201  10.190  17.097  1.00  0.00           H
ATOM   1515  NH2 ARG A 116      57.341  11.264  19.001  1.00  0.00           N
ATOM   1516 HH21 ARG A 116      57.765  11.775  19.775  1.00  0.00           H
ATOM   1517 HH22 ARG A 116      57.571  10.286  18.878  1.00  0.00           H
ATOM   1518  C   ARG A 116      57.402  17.837  16.410  1.00  0.00           C
ATOM   1519  O   ARG A 116      56.922  18.889  16.826  1.00  0.00           O
ATOM   1520  N   GLN A 117      58.721  17.686  16.277  1.00  0.00           N
ATOM   1521  H   GLN A 117      59.069  16.775  15.997  1.00  0.00           H
ATOM   1522  CA  GLN A 117      59.702  18.746  16.539  1.00  0.00           C
ATOM   1523  HA  GLN A 117      59.595  19.087  17.570  1.00  0.00           H
ATOM   1524  CB  GLN A 117      61.120  18.178  16.369  1.00  0.00           C
ATOM   1525  HB1 GLN A 117      61.258  17.374  17.091  1.00  0.00           H
ATOM   1526  HB2 GLN A 117      61.218  17.765  15.368  1.00  0.00           H
ATOM   1527  CG  GLN A 117      62.235  19.216  16.586  1.00  0.00           C
ATOM   1528  HG1 GLN A 117      62.063  20.079  15.944  1.00  0.00           H
ATOM   1529  HG2 GLN A 117      62.223  19.540  17.626  1.00  0.00           H
ATOM   1530  CD  GLN A 117      63.619  18.666  16.244  1.00  0.00           C
ATOM   1531  OE1 GLN A 117      64.525  18.625  17.064  1.00  0.00           O
ATOM   1532  NE2 GLN A 117      63.845  18.264  15.013  1.00  0.00           N
ATOM   1533 HE21 GLN A 117      63.136  18.384  14.302  1.00  0.00           H
ATOM   1534 HE22 GLN A 117      64.737  17.838  14.794  1.00  0.00           H
ATOM   1535  C   GLN A 117      59.455  19.963  15.638  1.00  0.00           C
ATOM   1536  O   GLN A 117      59.431  21.075  16.154  1.00  0.00           O
ATOM   1537  N   THR A 118      59.201  19.764  14.336  1.00  0.00           N
ATOM   1538  H   THR A 118      59.301  18.823  13.968  1.00  0.00           H
ATOM   1539  CA  THR A 118      58.831  20.817  13.374  1.00  0.00           C
ATOM   1540  HA  THR A 118      59.663  21.514  13.270  1.00  0.00           H
ATOM   1541  CB  THR A 118      58.546  20.180  12.007  1.00  0.00           C
ATOM   1542  HB  THR A 118      57.700  19.498  12.084  1.00  0.00           H
ATOM   1543  CG2 THR A 118      58.239  21.218  10.938  1.00  0.00           C
ATOM   1544 HG21 THR A 118      58.174  20.731   9.967  1.00  0.00           H
ATOM   1545 HG22 THR A 118      57.285  21.693  11.156  1.00  0.00           H
ATOM   1546 HG23 THR A 118      59.024  21.971  10.916  1.00  0.00           H
ATOM   1547  OG1 THR A 118      59.677  19.463  11.573  1.00  0.00           O
ATOM   1548  HG1 THR A 118      59.451  19.025  10.735  1.00  0.00           H
ATOM   1549  C   THR A 118      57.631  21.624  13.866  1.00  0.00           C
ATOM   1550  O   THR A 118      57.644  22.856  13.808  1.00  0.00           O
ATOM   1551  N   PHE A 119      56.609  20.950  14.403  1.00  0.00           N
ATOM   1552  H   PHE A 119      56.629  19.939  14.381  1.00  0.00           H
ATOM   1553  CA  PHE A 119      55.474  21.614  15.038  1.00  0.00           C
ATOM   1554  HA  PHE A 119      55.100  22.378  14.355  1.00  0.00           H
ATOM   1555  CB  PHE A 119      54.345  20.603  15.277  1.00  0.00           C
ATOM   1556  HB1 PHE A 119      54.058  20.169  14.321  1.00  0.00           H
ATOM   1557  HB2 PHE A 119      54.696  19.790  15.910  1.00  0.00           H
ATOM   1558  CG  PHE A 119      53.121  21.237  15.904  1.00  0.00           C
ATOM   1559  CD1 PHE A 119      52.425  22.238  15.204  1.00  0.00           C
ATOM   1560  HD1 PHE A 119      52.740  22.531  14.214  1.00  0.00           H
ATOM   1561  CE1 PHE A 119      51.333  22.885  15.797  1.00  0.00           C
ATOM   1562  HE1 PHE A 119      50.826  23.665  15.252  1.00  0.00           H
ATOM   1563  CZ  PHE A 119      50.917  22.526  17.091  1.00  0.00           C
```

| | | | | | | | | | | | |
|---|---|---|---|---|---|---|---|---|---|---|---|
| ATOM | 1564 | HZ | PHE | A | 119 | 50.078 | 23.030 | 17.550 | 1.00 | 0.00 | H |
| ATOM | 1565 | CE2 | PHE | A | 119 | 51.601 | 21.516 | 17.791 | 1.00 | 0.00 | C |
| ATOM | 1566 | HE2 | PHE | A | 119 | 51.286 | 21.243 | 18.788 | 1.00 | 0.00 | H |
| ATOM | 1567 | CD2 | PHE | A | 119 | 52.707 | 20.876 | 17.202 | 1.00 | 0.00 | C |
| ATOM | 1568 | HD2 | PHE | A | 119 | 53.243 | 20.116 | 17.751 | 1.00 | 0.00 | H |
| ATOM | 1569 | C | PHE | A | 119 | 55.875 | 22.334 | 16.332 | 1.00 | 0.00 | C |
| ATOM | 1570 | O | PHE | A | 119 | 55.706 | 23.545 | 16.427 | 1.00 | 0.00 | O |
| ATOM | 1571 | N | VAL | A | 120 | 56.431 | 21.612 | 17.306 | 1.00 | 0.00 | N |
| ATOM | 1572 | H | VAL | A | 120 | 56.593 | 20.626 | 17.107 | 1.00 | 0.00 | H |
| ATOM | 1573 | CA | VAL | A | 120 | 56.751 | 22.120 | 18.655 | 1.00 | 0.00 | C |
| ATOM | 1574 | HA | VAL | A | 120 | 55.816 | 22.433 | 19.123 | 1.00 | 0.00 | H |
| ATOM | 1575 | CB | VAL | A | 120 | 57.368 | 20.992 | 19.520 | 1.00 | 0.00 | C |
| ATOM | 1576 | HB | VAL | A | 120 | 58.185 | 20.533 | 18.958 | 1.00 | 0.00 | H |
| ATOM | 1577 | CG1 | VAL | A | 120 | 57.930 | 21.490 | 20.860 | 1.00 | 0.00 | C |
| ATOM | 1578 | HG11 | VAL | A | 120 | 58.280 | 20.637 | 21.441 | 1.00 | 0.00 | H |
| ATOM | 1579 | HG12 | VAL | A | 120 | 58.775 | 22.158 | 20.697 | 1.00 | 0.00 | H |
| ATOM | 1580 | HG13 | VAL | A | 120 | 57.155 | 22.014 | 21.421 | 1.00 | 0.00 | H |
| ATOM | 1581 | CG2 | VAL | A | 120 | 56.338 | 19.890 | 19.840 | 1.00 | 0.00 | C |
| ATOM | 1582 | HG21 | VAL | A | 120 | 56.844 | 19.022 | 20.264 | 1.00 | 0.00 | H |
| ATOM | 1583 | HG22 | VAL | A | 120 | 55.614 | 20.255 | 20.563 | 1.00 | 0.00 | H |
| ATOM | 1584 | HG23 | VAL | A | 120 | 55.798 | 19.579 | 18.949 | 1.00 | 0.00 | H |
| ATOM | 1585 | C | VAL | A | 120 | 57.687 | 23.339 | 18.596 | 1.00 | 0.00 | C |
| ATOM | 1586 | O | VAL | A | 120 | 57.400 | 24.373 | 19.200 | 1.00 | 0.00 | O |
| ATOM | 1587 | N | ASN | A | 121 | 58.784 | 23.247 | 17.838 | 1.00 | 0.00 | N |
| ATOM | 1588 | H | ASN | A | 121 | 58.958 | 22.382 | 17.335 | 1.00 | 0.00 | H |
| ATOM | 1589 | CA | ASN | A | 121 | 59.810 | 24.288 | 17.767 | 1.00 | 0.00 | C |
| ATOM | 1590 | HA | ASN | A | 121 | 60.081 | 24.579 | 18.783 | 1.00 | 0.00 | H |
| ATOM | 1591 | CB | ASN | A | 121 | 61.069 | 23.728 | 17.078 | 1.00 | 0.00 | C |
| ATOM | 1592 | HB1 | ASN | A | 121 | 60.805 | 23.301 | 16.110 | 1.00 | 0.00 | H |
| ATOM | 1593 | HB2 | ASN | A | 121 | 61.754 | 24.554 | 16.891 | 1.00 | 0.00 | H |
| ATOM | 1594 | CG | ASN | A | 121 | 61.823 | 22.681 | 17.902 | 1.00 | 0.00 | C |
| ATOM | 1595 | OD1 | ASN | A | 121 | 61.394 | 22.217 | 18.956 | 1.00 | 0.00 | O |
| ATOM | 1596 | ND2 | ASN | A | 121 | 63.001 | 22.289 | 17.452 | 1.00 | 0.00 | N |
| ATOM | 1597 | HD21 | ASN | A | 121 | 63.360 | 22.649 | 16.578 | 1.00 | 0.00 | H |
| ATOM | 1598 | HD22 | ASN | A | 121 | 63.548 | 21.662 | 18.021 | 1.00 | 0.00 | H |
| ATOM | 1599 | C | ASN | A | 121 | 59.306 | 25.551 | 17.046 | 1.00 | 0.00 | C |
| ATOM | 1600 | O | ASN | A | 121 | 59.649 | 26.664 | 17.449 | 1.00 | 0.00 | O |
| ATOM | 1601 | N | SER | A | 122 | 58.456 | 25.413 | 16.022 | 1.00 | 0.00 | N |
| ATOM | 1602 | H | SER | A | 122 | 58.171 | 24.480 | 15.752 | 1.00 | 0.00 | H |
| ATOM | 1603 | CA | SER | A | 122 | 57.815 | 26.572 | 15.383 | 1.00 | 0.00 | C |
| ATOM | 1604 | HA | SER | A | 122 | 58.560 | 27.360 | 15.256 | 1.00 | 0.00 | H |
| ATOM | 1605 | CB | SER | A | 122 | 57.303 | 26.227 | 13.974 | 1.00 | 0.00 | C |
| ATOM | 1606 | HB1 | SER | A | 122 | 56.848 | 27.120 | 13.543 | 1.00 | 0.00 | H |
| ATOM | 1607 | HB2 | SER | A | 122 | 58.150 | 25.942 | 13.346 | 1.00 | 0.00 | H |
| ATOM | 1608 | OG | SER | A | 122 | 56.348 | 25.180 | 13.965 | 1.00 | 0.00 | O |
| ATOM | 1609 | HG | SER | A | 122 | 56.831 | 24.325 | 13.968 | 1.00 | 0.00 | H |
| ATOM | 1610 | C | SER | A | 122 | 56.702 | 27.176 | 16.258 | 1.00 | 0.00 | C |
| ATOM | 1611 | O | SER | A | 122 | 56.615 | 28.401 | 16.359 | 1.00 | 0.00 | O |
| ATOM | 1612 | N | ALA | A | 123 | 55.896 | 26.357 | 16.946 | 1.00 | 0.00 | N |
| ATOM | 1613 | H | ALA | A | 123 | 55.982 | 25.355 | 16.795 | 1.00 | 0.00 | H |
| ATOM | 1614 | CA | ALA | A | 123 | 54.816 | 26.812 | 17.824 | 1.00 | 0.00 | C |
| ATOM | 1615 | HA | ALA | A | 123 | 54.158 | 27.463 | 17.250 | 1.00 | 0.00 | H |
| ATOM | 1616 | CB | ALA | A | 123 | 54.013 | 25.587 | 18.284 | 1.00 | 0.00 | C |
| ATOM | 1617 | HB1 | ALA | A | 123 | 53.194 | 25.906 | 18.929 | 1.00 | 0.00 | H |

| | | | | | | | | | | | |
|---|---|---|---|---|---|---|---|---|---|---|---|
| ATOM | 1618 | HB2 | ALA | A | 123 | 53.601 | 25.063 | 17.421 | 1.00 | 0.00 | H |
| ATOM | 1619 | HB3 | ALA | A | 123 | 54.653 | 24.902 | 18.844 | 1.00 | 0.00 | H |
| ATOM | 1620 | C | ALA | A | 123 | 55.322 | 27.641 | 19.016 | 1.00 | 0.00 | C |
| ATOM | 1621 | O | ALA | A | 123 | 54.777 | 28.710 | 19.287 | 1.00 | 0.00 | O |
| ATOM | 1622 | N | ILE | A | 124 | 56.392 | 27.200 | 19.688 | 1.00 | 0.00 | N |
| ATOM | 1623 | H | ILE | A | 124 | 56.748 | 26.281 | 19.445 | 1.00 | 0.00 | H |
| ATOM | 1624 | CA | ILE | A | 124 | 56.985 | 27.920 | 20.833 | 1.00 | 0.00 | C |
| ATOM | 1625 | HA | ILE | A | 124 | 56.213 | 28.080 | 21.589 | 1.00 | 0.00 | H |
| ATOM | 1626 | CB | ILE | A | 124 | 58.111 | 27.063 | 21.464 | 1.00 | 0.00 | C |
| ATOM | 1627 | HB | ILE | A | 124 | 58.765 | 26.708 | 20.666 | 1.00 | 0.00 | H |
| ATOM | 1628 | CG2 | ILE | A | 124 | 58.965 | 27.879 | 22.451 | 1.00 | 0.00 | C |
| ATOM | 1629 | HG21 | ILE | A | 124 | 59.749 | 27.252 | 22.873 | 1.00 | 0.00 | H |
| ATOM | 1630 | HG22 | ILE | A | 124 | 59.460 | 28.706 | 21.942 | 1.00 | 0.00 | H |
| ATOM | 1631 | HG23 | ILE | A | 124 | 58.342 | 28.271 | 23.256 | 1.00 | 0.00 | H |
| ATOM | 1632 | CG1 | ILE | A | 124 | 57.502 | 25.837 | 22.187 | 1.00 | 0.00 | C |
| ATOM | 1633 | HG11 | ILE | A | 124 | 56.956 | 26.170 | 23.072 | 1.00 | 0.00 | H |
| ATOM | 1634 | HG12 | ILE | A | 124 | 56.786 | 25.344 | 21.531 | 1.00 | 0.00 | H |
| ATOM | 1635 | CD | ILE | A | 124 | 58.536 | 24.784 | 22.607 | 1.00 | 0.00 | C |
| ATOM | 1636 | HD1 | ILE | A | 124 | 58.016 | 23.892 | 22.954 | 1.00 | 0.00 | H |
| ATOM | 1637 | HD2 | ILE | A | 124 | 59.163 | 24.516 | 21.755 | 1.00 | 0.00 | H |
| ATOM | 1638 | HD3 | ILE | A | 124 | 59.160 | 25.158 | 23.418 | 1.00 | 0.00 | H |
| ATOM | 1639 | C | ILE | A | 124 | 57.448 | 29.324 | 20.416 | 1.00 | 0.00 | C |
| ATOM | 1640 | O | ILE | A | 124 | 57.083 | 30.309 | 21.067 | 1.00 | 0.00 | O |
| ATOM | 1641 | N | ARG | A | 125 | 58.170 | 29.433 | 19.289 | 1.00 | 0.00 | N |
| ATOM | 1642 | H | ARG | A | 125 | 58.416 | 28.586 | 18.797 | 1.00 | 0.00 | H |
| ATOM | 1643 | CA | ARG | A | 125 | 58.565 | 30.724 | 18.709 | 1.00 | 0.00 | C |
| ATOM | 1644 | HA | ARG | A | 125 | 59.216 | 31.234 | 19.420 | 1.00 | 0.00 | H |
| ATOM | 1645 | CB | ARG | A | 125 | 59.334 | 30.526 | 17.390 | 1.00 | 0.00 | C |
| ATOM | 1646 | HB1 | ARG | A | 125 | 58.859 | 29.752 | 16.787 | 1.00 | 0.00 | H |
| ATOM | 1647 | HB2 | ARG | A | 125 | 59.299 | 31.462 | 16.829 | 1.00 | 0.00 | H |
| ATOM | 1648 | CG | ARG | A | 125 | 60.808 | 30.176 | 17.622 | 1.00 | 0.00 | C |
| ATOM | 1649 | HG1 | ARG | A | 125 | 61.246 | 30.927 | 18.280 | 1.00 | 0.00 | H |
| ATOM | 1650 | HG2 | ARG | A | 125 | 60.892 | 29.203 | 18.103 | 1.00 | 0.00 | H |
| ATOM | 1651 | CD | ARG | A | 125 | 61.601 | 30.181 | 16.308 | 1.00 | 0.00 | C |
| ATOM | 1652 | HD1 | ARG | A | 125 | 61.350 | 31.078 | 15.739 | 1.00 | 0.00 | H |
| ATOM | 1653 | HD2 | ARG | A | 125 | 62.663 | 30.239 | 16.545 | 1.00 | 0.00 | H |
| ATOM | 1654 | NE | ARG | A | 125 | 61.353 | 28.994 | 15.473 | 1.00 | 0.00 | N |
| ATOM | 1655 | HE | ARG | A | 125 | 60.702 | 29.096 | 14.702 | 1.00 | 0.00 | H |
| ATOM | 1656 | CZ | ARG | A | 125 | 62.062 | 27.872 | 15.506 | 1.00 | 0.00 | C |
| ATOM | 1657 | NH1 | ARG | A | 125 | 61.883 | 26.971 | 14.569 | 1.00 | 0.00 | N |
| ATOM | 1658 | HH11 | ARG | A | 125 | 61.239 | 27.171 | 13.809 | 1.00 | 0.00 | H |
| ATOM | 1659 | HH12 | ARG | A | 125 | 62.518 | 26.188 | 14.515 | 1.00 | 0.00 | H |
| ATOM | 1660 | NH2 | ARG | A | 125 | 62.969 | 27.622 | 16.427 | 1.00 | 0.00 | N |
| ATOM | 1661 | HH21 | ARG | A | 125 | 63.165 | 28.292 | 17.164 | 1.00 | 0.00 | H |
| ATOM | 1662 | HH22 | ARG | A | 125 | 63.533 | 26.790 | 16.352 | 1.00 | 0.00 | H |
| ATOM | 1663 | C | ARG | A | 125 | 57.356 | 31.633 | 18.465 | 1.00 | 0.00 | C |
| ATOM | 1664 | O | ARG | A | 125 | 57.409 | 32.805 | 18.845 | 1.00 | 0.00 | O |
| ATOM | 1665 | N | PHE | A | 126 | 56.288 | 31.085 | 17.874 | 1.00 | 0.00 | N |
| ATOM | 1666 | H | PHE | A | 126 | 56.348 | 30.103 | 17.634 | 1.00 | 0.00 | H |
| ATOM | 1667 | CA | PHE | A | 126 | 55.053 | 31.780 | 17.499 | 1.00 | 0.00 | C |
| ATOM | 1668 | HA | PHE | A | 126 | 55.302 | 32.604 | 16.828 | 1.00 | 0.00 | H |
| ATOM | 1669 | CB | PHE | A | 126 | 54.170 | 30.770 | 16.742 | 1.00 | 0.00 | C |
| ATOM | 1670 | HB1 | PHE | A | 126 | 54.810 | 30.131 | 16.138 | 1.00 | 0.00 | H |
| ATOM | 1671 | HB2 | PHE | A | 126 | 53.683 | 30.121 | 17.467 | 1.00 | 0.00 | H |

| | | | | | | | | | | | |
|---|---|---|---|---|---|---|---|---|---|---|---|
| ATOM | 1672 | CG | PHE | A | 126 | 53.111 | 31.314 | 15.800 | 1.00 | 0.00 | C |
| ATOM | 1673 | CD1 | PHE | A | 126 | 51.805 | 31.571 | 16.258 | 1.00 | 0.00 | C |
| ATOM | 1674 | HD1 | PHE | A | 126 | 51.564 | 31.473 | 17.307 | 1.00 | 0.00 | H |
| ATOM | 1675 | CE1 | PHE | A | 126 | 50.797 | 31.933 | 15.346 | 1.00 | 0.00 | C |
| ATOM | 1676 | HE1 | PHE | A | 126 | 49.794 | 32.121 | 15.702 | 1.00 | 0.00 | H |
| ATOM | 1677 | CZ | PHE | A | 126 | 51.086 | 32.038 | 13.974 | 1.00 | 0.00 | C |
| ATOM | 1678 | HZ | PHE | A | 126 | 50.305 | 32.311 | 13.280 | 1.00 | 0.00 | H |
| ATOM | 1679 | CE2 | PHE | A | 126 | 52.389 | 31.787 | 13.513 | 1.00 | 0.00 | C |
| ATOM | 1680 | HE2 | PHE | A | 126 | 52.618 | 31.868 | 12.460 | 1.00 | 0.00 | H |
| ATOM | 1681 | CD2 | PHE | A | 126 | 53.398 | 31.429 | 14.425 | 1.00 | 0.00 | C |
| ATOM | 1682 | HD2 | PHE | A | 126 | 54.395 | 31.229 | 14.063 | 1.00 | 0.00 | H |
| ATOM | 1683 | C | PHE | A | 126 | 54.336 | 32.358 | 18.729 | 1.00 | 0.00 | C |
| ATOM | 1684 | O | PHE | A | 126 | 54.043 | 33.553 | 18.759 | 1.00 | 0.00 | O |
| ATOM | 1685 | N | LEU | A | 127 | 54.105 | 31.549 | 19.775 | 1.00 | 0.00 | N |
| ATOM | 1686 | H | LEU | A | 127 | 54.391 | 30.575 | 19.710 | 1.00 | 0.00 | H |
| ATOM | 1687 | CA | LEU | A | 127 | 53.421 | 32.014 | 20.989 | 1.00 | 0.00 | C |
| ATOM | 1688 | HA | LEU | A | 127 | 52.468 | 32.460 | 20.698 | 1.00 | 0.00 | H |
| ATOM | 1689 | CB | LEU | A | 127 | 53.154 | 30.851 | 21.965 | 1.00 | 0.00 | C |
| ATOM | 1690 | HB1 | LEU | A | 127 | 54.109 | 30.411 | 22.260 | 1.00 | 0.00 | H |
| ATOM | 1691 | HB2 | LEU | A | 127 | 52.706 | 31.279 | 22.864 | 1.00 | 0.00 | H |
| ATOM | 1692 | CG | LEU | A | 127 | 52.236 | 29.715 | 21.464 | 1.00 | 0.00 | C |
| ATOM | 1693 | HG | LEU | A | 127 | 52.797 | 29.094 | 20.771 | 1.00 | 0.00 | H |
| ATOM | 1694 | CD1 | LEU | A | 127 | 51.838 | 28.848 | 22.666 | 1.00 | 0.00 | C |
| ATOM | 1695 | HD11 | LEU | A | 127 | 51.250 | 27.998 | 22.326 | 1.00 | 0.00 | H |
| ATOM | 1696 | HD12 | LEU | A | 127 | 52.727 | 28.486 | 23.180 | 1.00 | 0.00 | H |
| ATOM | 1697 | HD13 | LEU | A | 127 | 51.246 | 29.435 | 23.368 | 1.00 | 0.00 | H |
| ATOM | 1698 | CD2 | LEU | A | 127 | 50.960 | 30.176 | 20.744 | 1.00 | 0.00 | C |
| ATOM | 1699 | HD21 | LEU | A | 127 | 50.389 | 29.301 | 20.434 | 1.00 | 0.00 | H |
| ATOM | 1700 | HD22 | LEU | A | 127 | 50.342 | 30.777 | 21.408 | 1.00 | 0.00 | H |
| ATOM | 1701 | HD23 | LEU | A | 127 | 51.210 | 30.751 | 19.853 | 1.00 | 0.00 | H |
| ATOM | 1702 | C | LEU | A | 127 | 54.212 | 33.127 | 21.685 | 1.00 | 0.00 | C |
| ATOM | 1703 | O | LEU | A | 127 | 53.640 | 34.175 | 21.994 | 1.00 | 0.00 | O |
| ATOM | 1704 | N | ARG | A | 128 | 55.524 | 32.945 | 21.885 | 1.00 | 0.00 | N |
| ATOM | 1705 | H | ARG | A | 128 | 55.948 | 32.067 | 21.596 | 1.00 | 0.00 | H |
| ATOM | 1706 | CA | ARG | A | 128 | 56.368 | 33.975 | 22.505 | 1.00 | 0.00 | C |
| ATOM | 1707 | HA | ARG | A | 128 | 55.914 | 34.250 | 23.456 | 1.00 | 0.00 | H |
| ATOM | 1708 | CB | ARG | A | 128 | 57.779 | 33.428 | 22.776 | 1.00 | 0.00 | C |
| ATOM | 1709 | HB1 | ARG | A | 128 | 58.188 | 33.004 | 21.857 | 1.00 | 0.00 | H |
| ATOM | 1710 | HB2 | ARG | A | 128 | 58.423 | 34.258 | 23.071 | 1.00 | 0.00 | H |
| ATOM | 1711 | CG | ARG | A | 128 | 57.838 | 32.364 | 23.887 | 1.00 | 0.00 | C |
| ATOM | 1712 | HG1 | ARG | A | 128 | 57.316 | 31.464 | 23.562 | 1.00 | 0.00 | H |
| ATOM | 1713 | HG2 | ARG | A | 128 | 58.885 | 32.108 | 24.042 | 1.00 | 0.00 | H |
| ATOM | 1714 | CD | ARG | A | 128 | 57.232 | 32.840 | 25.217 | 1.00 | 0.00 | C |
| ATOM | 1715 | HD1 | ARG | A | 128 | 57.546 | 33.865 | 25.411 | 1.00 | 0.00 | H |
| ATOM | 1716 | HD2 | ARG | A | 128 | 56.145 | 32.813 | 25.137 | 1.00 | 0.00 | H |
| ATOM | 1717 | NE | ARG | A | 128 | 57.666 | 32.001 | 26.340 | 1.00 | 0.00 | N |
| ATOM | 1718 | HE | ARG | A | 128 | 58.507 | 31.437 | 26.195 | 1.00 | 0.00 | H |
| ATOM | 1719 | CZ | ARG | A | 128 | 57.137 | 31.945 | 27.554 | 1.00 | 0.00 | C |
| ATOM | 1720 | NH1 | ARG | A | 128 | 56.094 | 32.650 | 27.936 | 1.00 | 0.00 | N |
| ATOM | 1721 | HH11 | ARG | A | 128 | 55.623 | 33.275 | 27.292 | 1.00 | 0.00 | H |
| ATOM | 1722 | HH12 | ARG | A | 128 | 55.752 | 32.530 | 28.881 | 1.00 | 0.00 | H |
| ATOM | 1723 | NH2 | ARG | A | 128 | 57.674 | 31.146 | 28.435 | 1.00 | 0.00 | N |
| ATOM | 1724 | HH21 | ARG | A | 128 | 58.424 | 30.516 | 28.137 | 1.00 | 0.00 | H |
| ATOM | 1725 | HH22 | ARG | A | 128 | 57.263 | 31.063 | 29.357 | 1.00 | 0.00 | H |

```
ATOM   1726  C   ARG A 128      56.415  35.263  21.671  1.00  0.00           C
ATOM   1727  O   ARG A 128      56.268  36.347  22.244  1.00  0.00           O
ATOM   1728  N   LYS A 129      56.526  35.160  20.341  1.00  0.00           N
ATOM   1729  H   LYS A 129      56.625  34.235  19.940  1.00  0.00           H
ATOM   1730  CA  LYS A 129      56.465  36.293  19.405  1.00  0.00           C
ATOM   1731  HA  LYS A 129      57.296  36.969  19.613  1.00  0.00           H
ATOM   1732  CB  LYS A 129      56.633  35.758  17.968  1.00  0.00           C
ATOM   1733  HB1 LYS A 129      57.638  35.346  17.863  1.00  0.00           H
ATOM   1734  HB2 LYS A 129      55.918  34.955  17.796  1.00  0.00           H
ATOM   1735  CG  LYS A 129      56.423  36.827  16.884  1.00  0.00           C
ATOM   1736  HG1 LYS A 129      55.427  37.261  16.980  1.00  0.00           H
ATOM   1737  HG2 LYS A 129      57.166  37.614  17.003  1.00  0.00           H
ATOM   1738  CD  LYS A 129      56.544  36.217  15.485  1.00  0.00           C
ATOM   1739  HD1 LYS A 129      57.568  35.875  15.323  1.00  0.00           H
ATOM   1740  HD2 LYS A 129      55.865  35.365  15.407  1.00  0.00           H
ATOM   1741  CE  LYS A 129      56.162  37.261  14.432  1.00  0.00           C
ATOM   1742  HE1 LYS A 129      55.145  37.607  14.638  1.00  0.00           H
ATOM   1743  HE2 LYS A 129      56.840  38.117  14.514  1.00  0.00           H
ATOM   1744  NZ  LYS A 129      56.229  36.688  13.070  1.00  0.00           N
ATOM   1745  HZ1 LYS A 129      57.192  36.464  12.835  1.00  0.00           H
ATOM   1746  HZ2 LYS A 129      55.692  35.825  13.017  1.00  0.00           H
ATOM   1747  HZ3 LYS A 129      55.874  37.335  12.375  1.00  0.00           H
ATOM   1748  C   LYS A 129      55.172  37.109  19.578  1.00  0.00           C
ATOM   1749  O   LYS A 129      55.233  38.294  19.897  1.00  0.00           O
ATOM   1750  N   TYR A 130      53.997  36.497  19.407  1.00  0.00           N
ATOM   1751  H   TYR A 130      53.987  35.517  19.139  1.00  0.00           H
ATOM   1752  CA  TYR A 130      52.729  37.240  19.417  1.00  0.00           C
ATOM   1753  HA  TYR A 130      52.896  38.190  18.909  1.00  0.00           H
ATOM   1754  CB  TYR A 130      51.684  36.479  18.593  1.00  0.00           C
ATOM   1755  HB1 TYR A 130      51.575  35.475  19.004  1.00  0.00           H
ATOM   1756  HB2 TYR A 130      50.724  36.983  18.689  1.00  0.00           H
ATOM   1757  CG  TYR A 130      52.031  36.396  17.116  1.00  0.00           C
ATOM   1758  CD1 TYR A 130      52.052  37.560  16.318  1.00  0.00           C
ATOM   1759  HD1 TYR A 130      51.791  38.516  16.750  1.00  0.00           H
ATOM   1760  CE1 TYR A 130      52.410  37.484  14.956  1.00  0.00           C
ATOM   1761  HE1 TYR A 130      52.416  38.376  14.345  1.00  0.00           H
ATOM   1762  CZ  TYR A 130      52.768  36.238  14.394  1.00  0.00           C
ATOM   1763  OH  TYR A 130      53.140  36.141  13.090  1.00  0.00           O
ATOM   1764  HH  TYR A 130      53.050  36.987  12.604  1.00  0.00           H
ATOM   1765  CE2 TYR A 130      52.752  35.079  15.192  1.00  0.00           C
ATOM   1766  HE2 TYR A 130      53.025  34.129  14.763  1.00  0.00           H
ATOM   1767  CD2 TYR A 130      52.367  35.158  16.540  1.00  0.00           C
ATOM   1768  HD2 TYR A 130      52.321  34.258  17.135  1.00  0.00           H
ATOM   1769  C   TYR A 130      52.221  37.611  20.826  1.00  0.00           C
ATOM   1770  O   TYR A 130      51.438  38.550  20.942  1.00  0.00           O
ATOM   1771  N   SER A 131      52.701  36.926  21.873  1.00  0.00           N
ATOM   1772  H   SER A 131      53.280  36.125  21.644  1.00  0.00           H
ATOM   1773  CA  SER A 131      52.508  37.195  23.318  1.00  0.00           C
ATOM   1774  HA  SER A 131      53.422  36.890  23.823  1.00  0.00           H
ATOM   1775  CB  SER A 131      52.328  38.682  23.659  1.00  0.00           C
ATOM   1776  HB1 SER A 131      53.009  39.271  23.048  1.00  0.00           H
ATOM   1777  HB2 SER A 131      51.306  38.983  23.447  1.00  0.00           H
ATOM   1778  OG  SER A 131      52.610  38.920  25.025  1.00  0.00           O
ATOM   1779  HG  SER A 131      52.707  39.884  25.157  1.00  0.00           H
```

```
ATOM   1780  C    SER A 131      51.397  36.326  23.931  1.00  0.00           C
ATOM   1781  O    SER A 131      50.563  36.786  24.715  1.00  0.00           O
ATOM   1782  N    PHE A 132      51.345  35.052  23.548  1.00  0.00           N
ATOM   1783  H    PHE A 132      52.053  34.712  22.909  1.00  0.00           H
ATOM   1784  CA   PHE A 132      50.432  34.068  24.121  1.00  0.00           C
ATOM   1785  HA   PHE A 132      49.522  34.569  24.457  1.00  0.00           H
ATOM   1786  CB   PHE A 132      50.027  33.066  23.034  1.00  0.00           C
ATOM   1787  HB1  PHE A 132      50.905  32.789  22.454  1.00  0.00           H
ATOM   1788  HB2  PHE A 132      49.644  32.160  23.507  1.00  0.00           H
ATOM   1789  CG   PHE A 132      48.959  33.608  22.104  1.00  0.00           C
ATOM   1790  CD1  PHE A 132      47.612  33.280  22.333  1.00  0.00           C
ATOM   1791  HD1  PHE A 132      47.361  32.631  23.150  1.00  0.00           H
ATOM   1792  CE1  PHE A 132      46.597  33.803  21.516  1.00  0.00           C
ATOM   1793  HE1  PHE A 132      45.563  33.551  21.702  1.00  0.00           H
ATOM   1794  CZ   PHE A 132      46.928  34.675  20.468  1.00  0.00           C
ATOM   1795  HZ   PHE A 132      46.147  35.103  19.857  1.00  0.00           H
ATOM   1796  CE2  PHE A 132      48.273  35.017  20.242  1.00  0.00           C
ATOM   1797  HE2  PHE A 132      48.522  35.729  19.467  1.00  0.00           H
ATOM   1798  CD2  PHE A 132      49.289  34.481  21.053  1.00  0.00           C
ATOM   1799  HD2  PHE A 132      50.317  34.763  20.883  1.00  0.00           H
ATOM   1800  C    PHE A 132      51.044  33.387  25.353  1.00  0.00           C
ATOM   1801  O    PHE A 132      52.264  33.298  25.510  1.00  0.00           O
ATOM   1802  N    ASP A 133      50.162  32.907  26.224  1.00  0.00           N
ATOM   1803  H    ASP A 133      49.177  33.066  26.022  1.00  0.00           H
ATOM   1804  CA   ASP A 133      50.452  32.201  27.471  1.00  0.00           C
ATOM   1805  HA   ASP A 133      51.479  32.405  27.781  1.00  0.00           H
ATOM   1806  CB   ASP A 133      49.501  32.732  28.555  1.00  0.00           C
ATOM   1807  HB1  ASP A 133      48.467  32.541  28.262  1.00  0.00           H
ATOM   1808  HB2  ASP A 133      49.687  32.186  29.478  1.00  0.00           H
ATOM   1809  CG   ASP A 133      49.694  34.222  28.834  1.00  0.00           C
ATOM   1810  OD1  ASP A 133      50.723  34.581  29.457  1.00  0.00           O
ATOM   1811  OD2  ASP A 133      48.810  35.025  28.458  1.00  0.00           O
ATOM   1812  C    ASP A 133      50.297  30.673  27.346  1.00  0.00           C
ATOM   1813  O    ASP A 133      50.639  29.955  28.283  1.00  0.00           O
ATOM   1814  N    GLY A 134      49.790  30.152  26.218  1.00  0.00           N
ATOM   1815  H    GLY A 134      49.499  30.785  25.487  1.00  0.00           H
ATOM   1816  CA   GLY A 134      49.625  28.708  26.005  1.00  0.00           C
ATOM   1817  HA1  GLY A 134      50.608  28.237  26.029  1.00  0.00           H
ATOM   1818  HA2  GLY A 134      49.069  28.287  26.843  1.00  0.00           H
ATOM   1819  C    GLY A 134      48.929  28.288  24.709  1.00  0.00           C
ATOM   1820  O    GLY A 134      48.531  29.125  23.896  1.00  0.00           O
ATOM   1821  N    LEU A 135      48.796  26.971  24.531  1.00  0.00           N
ATOM   1822  H    LEU A 135      49.121  26.370  25.284  1.00  0.00           H
ATOM   1823  CA   LEU A 135      48.211  26.301  23.362  1.00  0.00           C
ATOM   1824  HA   LEU A 135      47.868  27.046  22.645  1.00  0.00           H
ATOM   1825  CB   LEU A 135      49.314  25.425  22.724  1.00  0.00           C
ATOM   1826  HB1  LEU A 135      50.233  26.002  22.637  1.00  0.00           H
ATOM   1827  HB2  LEU A 135      49.518  24.599  23.409  1.00  0.00           H
ATOM   1828  CG   LEU A 135      48.969  24.846  21.335  1.00  0.00           C
ATOM   1829  HG   LEU A 135      47.961  24.430  21.351  1.00  0.00           H
ATOM   1830  CD1  LEU A 135      49.055  25.916  20.235  1.00  0.00           C
ATOM   1831 HD11  LEU A 135      48.764  25.483  19.277  1.00  0.00           H
ATOM   1832 HD12  LEU A 135      48.388  26.745  20.457  1.00  0.00           H
ATOM   1833 HD13  LEU A 135      50.074  26.298  20.159  1.00  0.00           H
```

```
ATOM   1834  CD2 LEU A 135      49.945  23.711  20.993  1.00  0.00           C
ATOM   1835 HD21 LEU A 135      49.680  23.274  20.030  1.00  0.00           H
ATOM   1836 HD22 LEU A 135      50.966  24.091  20.944  1.00  0.00           H
ATOM   1837 HD23 LEU A 135      49.889  22.933  21.754  1.00  0.00           H
ATOM   1838  C   LEU A 135      47.014  25.431  23.784  1.00  0.00           C
ATOM   1839  O   LEU A 135      47.108  24.712  24.777  1.00  0.00           O
ATOM   1840  N   ASP A 136      45.929  25.447  23.005  1.00  0.00           N
ATOM   1841  H   ASP A 136      45.925  26.074  22.206  1.00  0.00           H
ATOM   1842  CA  ASP A 136      44.780  24.541  23.147  1.00  0.00           C
ATOM   1843  HA  ASP A 136      44.930  23.914  24.024  1.00  0.00           H
ATOM   1844  CB  ASP A 136      43.493  25.352  23.383  1.00  0.00           C
ATOM   1845  HB1 ASP A 136      43.657  26.041  24.213  1.00  0.00           H
ATOM   1846  HB2 ASP A 136      43.268  25.949  22.500  1.00  0.00           H
ATOM   1847  CG  ASP A 136      42.296  24.456  23.724  1.00  0.00           C
ATOM   1848  OD1 ASP A 136      42.326  23.829  24.810  1.00  0.00           O
ATOM   1849  OD2 ASP A 136      41.334  24.407  22.927  1.00  0.00           O
ATOM   1850  C   ASP A 136      44.649  23.626  21.915  1.00  0.00           C
ATOM   1851  O   ASP A 136      44.622  24.085  20.771  1.00  0.00           O
ATOM   1852  N   LEU A 137      44.587  22.311  22.132  1.00  0.00           N
ATOM   1853  H   LEU A 137      44.691  21.985  23.090  1.00  0.00           H
ATOM   1854  CA  LEU A 137      44.529  21.324  21.048  1.00  0.00           C
ATOM   1855  HA  LEU A 137      45.018  21.732  20.162  1.00  0.00           H
ATOM   1856  CB  LEU A 137      45.293  20.055  21.482  1.00  0.00           C
ATOM   1857  HB1 LEU A 137      44.788  19.627  22.348  1.00  0.00           H
ATOM   1858  HB2 LEU A 137      45.239  19.328  20.670  1.00  0.00           H
ATOM   1859  CG  LEU A 137      46.778  20.269  21.853  1.00  0.00           C
ATOM   1860  HG  LEU A 137      46.850  20.959  22.695  1.00  0.00           H
ATOM   1861  CD1 LEU A 137      47.375  18.925  22.291  1.00  0.00           C
ATOM   1862 HD11 LEU A 137      48.414  19.058  22.592  1.00  0.00           H
ATOM   1863 HD12 LEU A 137      46.815  18.528  23.138  1.00  0.00           H
ATOM   1864 HD13 LEU A 137      47.331  18.216  21.465  1.00  0.00           H
ATOM   1865  CD2 LEU A 137      47.593  20.840  20.682  1.00  0.00           C
ATOM   1866 HD21 LEU A 137      48.648  20.872  20.949  1.00  0.00           H
ATOM   1867 HD22 LEU A 137      47.468  20.218  19.796  1.00  0.00           H
ATOM   1868 HD23 LEU A 137      47.268  21.857  20.464  1.00  0.00           H
ATOM   1869  C   LEU A 137      43.078  20.999  20.662  1.00  0.00           C
ATOM   1870  O   LEU A 137      42.373  20.364  21.441  1.00  0.00           O
ATOM   1871  N   ASP A 138      42.634  21.334  19.375  1.00  0.00           N
ATOM   1872  H   ASP A 138      43.188  21.943  18.764  1.00  0.00           H
ATOM   1873  CA  ASP A 138      41.271  21.028  18.928  1.00  0.00           C
ATOM   1874  HA  ASP A 138      40.798  20.407  19.704  1.00  0.00           H
ATOM   1875  CB  ASP A 138      40.352  22.227  18.724  1.00  0.00           C
ATOM   1876  HB1 ASP A 138      40.462  22.916  19.574  1.00  0.00           H
ATOM   1877  HB2 ASP A 138      40.639  22.791  17.826  1.00  0.00           H
ATOM   1878  CG  ASP A 138      38.867  21.822  18.618  1.00  0.00           C
ATOM   1879  OD1 ASP A 138      38.591  20.570  18.664  1.00  0.00           O
ATOM   1880  OD2 ASP A 138      38.031  22.759  18.517  1.00  0.00           O
ATOM   1881  C   ASP A 138      41.404  20.124  17.700  1.00  0.00           C
ATOM   1882  O   ASP A 138      41.182  20.506  16.553  1.00  0.00           O
ATOM   1883  N   TRP A 139      41.807  18.834  17.936  1.00  0.00           N
ATOM   1884  H   TRP A 139      42.046  18.517  18.889  1.00  0.00           H
ATOM   1885  CA  TRP A 139      41.768  17.790  16.903  1.00  0.00           C
ATOM   1886  HA  TRP A 139      41.908  18.228  15.914  1.00  0.00           H
ATOM   1887  CB  TRP A 139      42.948  16.833  17.151  1.00  0.00           C
```

| | | | | | | | | | | | |
|---|---|---|---|---|---|---|---|---|---|---|---|
| ATOM | 1888 | HB1 | TRP | A | 139 | 43.849 | 17.446 | 17.220 | 1.00 | 0.00 | H |
| ATOM | 1889 | HB2 | TRP | A | 139 | 42.810 | 16.360 | 18.124 | 1.00 | 0.00 | H |
| ATOM | 1890 | CG | TRP | A | 139 | 43.254 | 15.745 | 16.154 | 1.00 | 0.00 | C |
| ATOM | 1891 | CD1 | TRP | A | 139 | 42.475 | 15.285 | 15.145 | 1.00 | 0.00 | C |
| ATOM | 1892 | HD1 | TRP | A | 139 | 41.498 | 15.648 | 14.878 | 1.00 | 0.00 | H |
| ATOM | 1893 | NE1 | TRP | A | 139 | 43.105 | 14.248 | 14.492 | 1.00 | 0.00 | N |
| ATOM | 1894 | HE1 | TRP | A | 139 | 42.675 | 13.706 | 13.755 | 1.00 | 0.00 | H |
| ATOM | 1895 | CE2 | TRP | A | 139 | 44.364 | 14.031 | 14.990 | 1.00 | 0.00 | C |
| ATOM | 1896 | CZ2 | TRP | A | 139 | 45.395 | 13.162 | 14.619 | 1.00 | 0.00 | C |
| ATOM | 1897 | HZ2 | TRP | A | 139 | 45.260 | 12.485 | 13.791 | 1.00 | 0.00 | H |
| ATOM | 1898 | CH2 | TRP | A | 139 | 46.616 | 13.219 | 15.306 | 1.00 | 0.00 | C |
| ATOM | 1899 | HH2 | TRP | A | 139 | 47.434 | 12.582 | 14.994 | 1.00 | 0.00 | H |
| ATOM | 1900 | CZ3 | TRP | A | 139 | 46.774 | 14.125 | 16.372 | 1.00 | 0.00 | C |
| ATOM | 1901 | HZ3 | TRP | A | 139 | 47.720 | 14.188 | 16.894 | 1.00 | 0.00 | H |
| ATOM | 1902 | CE3 | TRP | A | 139 | 45.725 | 14.988 | 16.741 | 1.00 | 0.00 | C |
| ATOM | 1903 | HE3 | TRP | A | 139 | 45.878 | 15.704 | 17.535 | 1.00 | 0.00 | H |
| ATOM | 1904 | CD2 | TRP | A | 139 | 44.491 | 14.969 | 16.056 | 1.00 | 0.00 | C |
| ATOM | 1905 | C | TRP | A | 139 | 40.404 | 17.077 | 16.949 | 1.00 | 0.00 | C |
| ATOM | 1906 | O | TRP | A | 139 | 40.000 | 16.602 | 18.003 | 1.00 | 0.00 | O |
| ATOM | 1907 | N | GLH | A | 140 | 39.659 | 16.985 | 15.768 | 1.00 | 0.00 | N |
| ATOM | 1908 | H | GLH | A | 140 | 39.968 | 17.566 | 14.977 | 1.00 | 0.00 | H |
| ATOM | 1909 | CA | GLH | A | 140 | 38.393 | 16.253 | 15.660 | 1.00 | 0.00 | C |
| ATOM | 1910 | HA | GLH | A | 140 | 38.189 | 15.795 | 16.636 | 1.00 | 0.00 | H |
| ATOM | 1911 | CB | GLH | A | 140 | 37.207 | 17.146 | 15.328 | 1.00 | 0.00 | C |
| ATOM | 1912 | HB1 | GLH | A | 140 | 37.417 | 17.719 | 14.413 | 1.00 | 0.00 | H |
| ATOM | 1913 | HB2 | GLH | A | 140 | 36.334 | 16.512 | 15.118 | 1.00 | 0.00 | H |
| ATOM | 1914 | CG | GLH | A | 140 | 36.883 | 18.071 | 16.489 | 1.00 | 0.00 | C |
| ATOM | 1915 | HG1 | GLH | A | 140 | 36.957 | 17.497 | 17.433 | 1.00 | 0.00 | H |
| ATOM | 1916 | HG2 | GLH | A | 140 | 37.574 | 18.919 | 16.587 | 1.00 | 0.00 | H |
| ATOM | 1917 | CD | GLH | A | 140 | 35.464 | 18.585 | 16.480 | 1.00 | 0.00 | C |
| ATOM | 1918 | OE1 | GLH | A | 140 | 34.511 | 17.980 | 15.983 | 1.00 | 0.00 | O |
| ATOM | 1919 | OE2 | GLH | A | 140 | 35.364 | 19.738 | 17.122 | 1.00 | 0.00 | O |
| ATOM | 1920 | HE2 | GLH | A | 140 | 34.384 | 20.041 | 17.201 | 1.00 | 0.00 | H |
| ATOM | 1921 | C | GLH | A | 140 | 38.580 | 15.135 | 14.630 | 1.00 | 0.00 | C |
| ATOM | 1922 | O | GLH | A | 140 | 38.235 | 15.310 | 13.461 | 1.00 | 0.00 | O |
| ATOM | 1923 | N | TYR | A | 141 | 39.138 | 13.933 | 14.960 | 1.00 | 0.00 | N |
| ATOM | 1924 | H | TYR | A | 141 | 39.256 | 13.207 | 14.227 | 1.00 | 0.00 | H |
| ATOM | 1925 | CA | TYR | A | 141 | 39.647 | 13.513 | 16.281 | 1.00 | 0.00 | C |
| ATOM | 1926 | HA | TYR | A | 141 | 39.984 | 14.369 | 16.858 | 1.00 | 0.00 | H |
| ATOM | 1927 | CB | TYR | A | 141 | 38.542 | 12.769 | 17.058 | 1.00 | 0.00 | C |
| ATOM | 1928 | HB1 | TYR | A | 141 | 38.284 | 11.876 | 16.496 | 1.00 | 0.00 | H |
| ATOM | 1929 | HB2 | TYR | A | 141 | 38.943 | 12.428 | 18.012 | 1.00 | 0.00 | H |
| ATOM | 1930 | CG | TYR | A | 141 | 37.270 | 13.547 | 17.335 | 1.00 | 0.00 | C |
| ATOM | 1931 | CD1 | TYR | A | 141 | 36.224 | 13.543 | 16.391 | 1.00 | 0.00 | C |
| ATOM | 1932 | HD1 | TYR | A | 141 | 36.321 | 12.970 | 15.480 | 1.00 | 0.00 | H |
| ATOM | 1933 | CE1 | TYR | A | 141 | 35.057 | 14.293 | 16.624 | 1.00 | 0.00 | C |
| ATOM | 1934 | HE1 | TYR | A | 141 | 34.250 | 14.291 | 15.905 | 1.00 | 0.00 | H |
| ATOM | 1935 | CZ | TYR | A | 141 | 34.920 | 15.035 | 17.816 | 1.00 | 0.00 | C |
| ATOM | 1936 | OH | TYR | A | 141 | 33.769 | 15.723 | 18.048 | 1.00 | 0.00 | O |
| ATOM | 1937 | HH | TYR | A | 141 | 33.733 | 16.146 | 18.908 | 1.00 | 0.00 | H |
| ATOM | 1938 | CE2 | TYR | A | 141 | 35.970 | 15.044 | 18.763 | 1.00 | 0.00 | C |
| ATOM | 1939 | HE2 | TYR | A | 141 | 35.869 | 15.618 | 19.669 | 1.00 | 0.00 | H |
| ATOM | 1940 | CD2 | TYR | A | 141 | 37.141 | 14.300 | 18.520 | 1.00 | 0.00 | C |
| ATOM | 1941 | HD2 | TYR | A | 141 | 37.947 | 14.312 | 19.241 | 1.00 | 0.00 | H |

| | | | | | | | | | | | |
|---|---|---|---|---|---|---|---|---|---|---|---|
| ATOM | 1942 | C | TYR | A | 141 | 40.840 | 12.553 | 16.095 | 1.00 | 0.00 | C |
| ATOM | 1943 | O | TYR | A | 141 | 40.942 | 11.955 | 15.021 | 1.00 | 0.00 | O |
| ATOM | 1944 | N | PRO | A | 142 | 41.705 | 12.339 | 17.105 | 1.00 | 0.00 | N |
| ATOM | 1945 | CD | PRO | A | 142 | 41.815 | 13.075 | 18.359 | 1.00 | 0.00 | C |
| ATOM | 1946 | HD1 | PRO | A | 142 | 40.842 | 13.319 | 18.786 | 1.00 | 0.00 | H |
| ATOM | 1947 | HD2 | PRO | A | 142 | 42.387 | 13.987 | 18.191 | 1.00 | 0.00 | H |
| ATOM | 1948 | CG | PRO | A | 142 | 42.585 | 12.159 | 19.309 | 1.00 | 0.00 | C |
| ATOM | 1949 | HG1 | PRO | A | 142 | 41.885 | 11.512 | 19.835 | 1.00 | 0.00 | H |
| ATOM | 1950 | HG2 | PRO | A | 142 | 43.181 | 12.724 | 20.025 | 1.00 | 0.00 | H |
| ATOM | 1951 | CB | PRO | A | 142 | 43.455 | 11.326 | 18.365 | 1.00 | 0.00 | C |
| ATOM | 1952 | HB1 | PRO | A | 142 | 43.717 | 10.362 | 18.802 | 1.00 | 0.00 | H |
| ATOM | 1953 | HB2 | PRO | A | 142 | 44.356 | 11.887 | 18.112 | 1.00 | 0.00 | H |
| ATOM | 1954 | CA | PRO | A | 142 | 42.585 | 11.178 | 17.112 | 1.00 | 0.00 | C |
| ATOM | 1955 | HA | PRO | A | 142 | 43.220 | 11.173 | 16.224 | 1.00 | 0.00 | H |
| ATOM | 1956 | C | PRO | A | 142 | 41.715 | 9.914 | 17.136 | 1.00 | 0.00 | C |
| ATOM | 1957 | O | PRO | A | 142 | 40.800 | 9.808 | 17.950 | 1.00 | 0.00 | O |
| ATOM | 1958 | N | GLY | A | 143 | 41.959 | 8.981 | 16.215 | 1.00 | 0.00 | N |
| ATOM | 1959 | H | GLY | A | 143 | 42.670 | 9.170 | 15.512 | 1.00 | 0.00 | H |
| ATOM | 1960 | CA | GLY | A | 143 | 41.276 | 7.687 | 16.133 | 1.00 | 0.00 | C |
| ATOM | 1961 | HA1 | GLY | A | 143 | 41.861 | 7.033 | 15.487 | 1.00 | 0.00 | H |
| ATOM | 1962 | HA2 | GLY | A | 143 | 41.218 | 7.255 | 17.132 | 1.00 | 0.00 | H |
| ATOM | 1963 | C | GLY | A | 143 | 39.851 | 7.711 | 15.570 | 1.00 | 0.00 | C |
| ATOM | 1964 | O | GLY | A | 143 | 39.200 | 6.672 | 15.592 | 1.00 | 0.00 | O |
| ATOM | 1965 | N | SER | A | 144 | 39.337 | 8.836 | 15.065 | 1.00 | 0.00 | N |
| ATOM | 1966 | H | SER | A | 144 | 39.940 | 9.645 | 14.994 | 1.00 | 0.00 | H |
| ATOM | 1967 | CA | SER | A | 144 | 37.996 | 8.911 | 14.445 | 1.00 | 0.00 | C |
| ATOM | 1968 | HA | SER | A | 144 | 37.741 | 7.934 | 14.037 | 1.00 | 0.00 | H |
| ATOM | 1969 | CB | SER | A | 144 | 36.913 | 9.270 | 15.484 | 1.00 | 0.00 | C |
| ATOM | 1970 | HB1 | SER | A | 144 | 37.098 | 10.267 | 15.877 | 1.00 | 0.00 | H |
| ATOM | 1971 | HB2 | SER | A | 144 | 35.941 | 9.288 | 14.993 | 1.00 | 0.00 | H |
| ATOM | 1972 | OG | SER | A | 144 | 36.840 | 8.353 | 16.563 | 1.00 | 0.00 | O |
| ATOM | 1973 | HG | SER | A | 144 | 37.435 | 8.678 | 17.273 | 1.00 | 0.00 | H |
| ATOM | 1974 | C | SER | A | 144 | 37.954 | 9.912 | 13.272 | 1.00 | 0.00 | C |
| ATOM | 1975 | O | SER | A | 144 | 38.924 | 10.634 | 13.043 | 1.00 | 0.00 | O |
| ATOM | 1976 | N | GLN | A | 145 | 36.843 | 9.982 | 12.519 | 1.00 | 0.00 | N |
| ATOM | 1977 | H | GLN | A | 145 | 36.074 | 9.352 | 12.734 | 1.00 | 0.00 | H |
| ATOM | 1978 | CA | GLN | A | 145 | 36.625 | 10.973 | 11.442 | 1.00 | 0.00 | C |
| ATOM | 1979 | HA | GLN | A | 145 | 35.701 | 10.701 | 10.929 | 1.00 | 0.00 | H |
| ATOM | 1980 | CB | GLN | A | 145 | 36.431 | 12.384 | 12.054 | 1.00 | 0.00 | C |
| ATOM | 1981 | HB1 | GLN | A | 145 | 37.136 | 12.542 | 12.870 | 1.00 | 0.00 | H |
| ATOM | 1982 | HB2 | GLN | A | 145 | 36.629 | 13.156 | 11.311 | 1.00 | 0.00 | H |
| ATOM | 1983 | CG | GLN | A | 145 | 34.998 | 12.597 | 12.573 | 1.00 | 0.00 | C |
| ATOM | 1984 | HG1 | GLN | A | 145 | 34.732 | 11.787 | 13.254 | 1.00 | 0.00 | H |
| ATOM | 1985 | HG2 | GLN | A | 145 | 34.960 | 13.530 | 13.133 | 1.00 | 0.00 | H |
| ATOM | 1986 | CD | GLN | A | 145 | 33.980 | 12.677 | 11.430 | 1.00 | 0.00 | C |
| ATOM | 1987 | OE1 | GLN | A | 145 | 34.176 | 13.356 | 10.432 | 1.00 | 0.00 | O |
| ATOM | 1988 | NE2 | GLN | A | 145 | 32.882 | 11.958 | 11.493 | 1.00 | 0.00 | N |
| ATOM | 1989 | HE21 | GLN | A | 145 | 32.666 | 11.382 | 12.300 | 1.00 | 0.00 | H |
| ATOM | 1990 | HE22 | GLN | A | 145 | 32.262 | 12.003 | 10.701 | 1.00 | 0.00 | H |
| ATOM | 1991 | C | GLN | A | 145 | 37.687 | 10.918 | 10.316 | 1.00 | 0.00 | C |
| ATOM | 1992 | O | GLN | A | 145 | 37.960 | 11.922 | 9.651 | 1.00 | 0.00 | O |
| ATOM | 1993 | N | GLY | A | 146 | 38.277 | 9.736 | 10.090 | 1.00 | 0.00 | N |
| ATOM | 1994 | H | GLY | A | 146 | 38.028 | 8.960 | 10.692 | 1.00 | 0.00 | H |
| ATOM | 1995 | CA | GLY | A | 146 | 39.301 | 9.459 | 9.073 | 1.00 | 0.00 | C |

| | | | | | | | | | | | |
|---|---|---|---|---|---|---|---|---|---|---|---|
| ATOM | 1996 | HA1 | GLY | A | 146 | 39.014 | 8.564 | 8.522 | 1.00 | 0.00 | H |
| ATOM | 1997 | HA2 | GLY | A | 146 | 39.388 | 10.296 | 8.381 | 1.00 | 0.00 | H |
| ATOM | 1998 | C | GLY | A | 146 | 40.685 | 9.188 | 9.664 | 1.00 | 0.00 | C |
| ATOM | 1999 | O | GLY | A | 146 | 41.514 | 8.550 | 9.019 | 1.00 | 0.00 | O |
| ATOM | 2000 | N | SER | A | 147 | 40.935 | 9.610 | 10.904 | 1.00 | 0.00 | N |
| ATOM | 2001 | H | SER | A | 147 | 40.183 | 10.058 | 11.418 | 1.00 | 0.00 | H |
| ATOM | 2002 | CA | SER | A | 147 | 42.196 | 9.339 | 11.604 | 1.00 | 0.00 | C |
| ATOM | 2003 | HA | SER | A | 147 | 43.010 | 9.672 | 10.966 | 1.00 | 0.00 | H |
| ATOM | 2004 | CB | SER | A | 147 | 42.241 | 10.173 | 12.886 | 1.00 | 0.00 | C |
| ATOM | 2005 | HB1 | SER | A | 147 | 42.359 | 11.224 | 12.636 | 1.00 | 0.00 | H |
| ATOM | 2006 | HB2 | SER | A | 147 | 41.291 | 10.044 | 13.400 | 1.00 | 0.00 | H |
| ATOM | 2007 | OG | SER | A | 147 | 43.251 | 9.792 | 13.792 | 1.00 | 0.00 | O |
| ATOM | 2008 | HG | SER | A | 147 | 44.165 | 9.943 | 13.447 | 1.00 | 0.00 | H |
| ATOM | 2009 | C | SER | A | 147 | 42.363 | 7.836 | 11.921 | 1.00 | 0.00 | C |
| ATOM | 2010 | O | SER | A | 147 | 41.424 | 7.232 | 12.459 | 1.00 | 0.00 | O |
| ATOM | 2011 | N | PRO | A | 148 | 43.528 | 7.230 | 11.604 | 1.00 | 0.00 | N |
| ATOM | 2012 | CD | PRO | A | 148 | 44.639 | 7.816 | 10.862 | 1.00 | 0.00 | C |
| ATOM | 2013 | HD1 | PRO | A | 148 | 44.963 | 8.757 | 11.303 | 1.00 | 0.00 | H |
| ATOM | 2014 | HD2 | PRO | A | 148 | 44.339 | 7.970 | 9.824 | 1.00 | 0.00 | H |
| ATOM | 2015 | CG | PRO | A | 148 | 45.779 | 6.803 | 10.921 | 1.00 | 0.00 | C |
| ATOM | 2016 | HG1 | PRO | A | 148 | 46.383 | 6.994 | 11.809 | 1.00 | 0.00 | H |
| ATOM | 2017 | HG2 | PRO | A | 148 | 46.395 | 6.835 | 10.022 | 1.00 | 0.00 | H |
| ATOM | 2018 | CB | PRO | A | 148 | 45.042 | 5.472 | 11.062 | 1.00 | 0.00 | C |
| ATOM | 2019 | HB1 | PRO | A | 148 | 45.664 | 4.706 | 11.526 | 1.00 | 0.00 | H |
| ATOM | 2020 | HB2 | PRO | A | 148 | 44.705 | 5.139 | 10.079 | 1.00 | 0.00 | H |
| ATOM | 2021 | CA | PRO | A | 148 | 43.826 | 5.837 | 11.920 | 1.00 | 0.00 | C |
| ATOM | 2022 | HA | PRO | A | 148 | 42.983 | 5.206 | 11.637 | 1.00 | 0.00 | H |
| ATOM | 2023 | C | PRO | A | 148 | 44.103 | 5.664 | 13.417 | 1.00 | 0.00 | C |
| ATOM | 2024 | O | PRO | A | 148 | 44.572 | 6.579 | 14.090 | 1.00 | 0.00 | O |
| ATOM | 2025 | N | ALA | A | 149 | 43.838 | 4.466 | 13.944 | 1.00 | 0.00 | N |
| ATOM | 2026 | H | ALA | A | 149 | 43.500 | 3.734 | 13.334 | 1.00 | 0.00 | H |
| ATOM | 2027 | CA | ALA | A | 149 | 43.797 | 4.209 | 15.386 | 1.00 | 0.00 | C |
| ATOM | 2028 | HA | ALA | A | 149 | 43.117 | 4.938 | 15.830 | 1.00 | 0.00 | H |
| ATOM | 2029 | CB | ALA | A | 149 | 43.192 | 2.818 | 15.583 | 1.00 | 0.00 | C |
| ATOM | 2030 | HB1 | ALA | A | 149 | 43.144 | 2.567 | 16.642 | 1.00 | 0.00 | H |
| ATOM | 2031 | HB2 | ALA | A | 149 | 42.186 | 2.788 | 15.163 | 1.00 | 0.00 | H |
| ATOM | 2032 | HB3 | ALA | A | 149 | 43.801 | 2.071 | 15.087 | 1.00 | 0.00 | H |
| ATOM | 2033 | C | ALA | A | 149 | 45.146 | 4.378 | 16.117 | 1.00 | 0.00 | C |
| ATOM | 2034 | O | ALA | A | 149 | 45.148 | 4.589 | 17.335 | 1.00 | 0.00 | O |
| ATOM | 2035 | N | VAL | A | 150 | 46.267 | 4.365 | 15.386 | 1.00 | 0.00 | N |
| ATOM | 2036 | H | VAL | A | 150 | 46.151 | 4.091 | 14.419 | 1.00 | 0.00 | H |
| ATOM | 2037 | CA | VAL | A | 150 | 47.612 | 4.764 | 15.869 | 1.00 | 0.00 | C |
| ATOM | 2038 | HA | VAL | A | 150 | 47.892 | 4.057 | 16.648 | 1.00 | 0.00 | H |
| ATOM | 2039 | CB | VAL | A | 150 | 48.700 | 4.707 | 14.767 | 1.00 | 0.00 | C |
| ATOM | 2040 | HB | VAL | A | 150 | 49.643 | 5.052 | 15.198 | 1.00 | 0.00 | H |
| ATOM | 2041 | CG1 | VAL | A | 150 | 48.925 | 3.274 | 14.281 | 1.00 | 0.00 | C |
| ATOM | 2042 | HG11 | VAL | A | 150 | 49.787 | 3.240 | 13.614 | 1.00 | 0.00 | H |
| ATOM | 2043 | HG12 | VAL | A | 150 | 49.114 | 2.617 | 15.130 | 1.00 | 0.00 | H |
| ATOM | 2044 | HG13 | VAL | A | 150 | 48.044 | 2.936 | 13.739 | 1.00 | 0.00 | H |
| ATOM | 2045 | CG2 | VAL | A | 150 | 48.386 | 5.595 | 13.549 | 1.00 | 0.00 | C |
| ATOM | 2046 | HG21 | VAL | A | 150 | 49.169 | 5.483 | 12.799 | 1.00 | 0.00 | H |
| ATOM | 2047 | HG22 | VAL | A | 150 | 47.435 | 5.303 | 13.107 | 1.00 | 0.00 | H |
| ATOM | 2048 | HG23 | VAL | A | 150 | 48.343 | 6.643 | 13.842 | 1.00 | 0.00 | H |
| ATOM | 2049 | C | VAL | A | 150 | 47.655 | 6.173 | 16.480 | 1.00 | 0.00 | C |

| | | | | | | | | | | | |
|---|---|---|---|---|---|---|---|---|---|---|---|
| ATOM | 2050 | O | VAL | A | 150 | 48.501 | 6.451 | 17.332 | 1.00 | 0.00 | O |
| ATOM | 2051 | N | ASP | A | 151 | 46.739 | 7.060 | 16.074 | 1.00 | 0.00 | N |
| ATOM | 2052 | H | ASP | A | 151 | 46.066 | 6.791 | 15.362 | 1.00 | 0.00 | H |
| ATOM | 2053 | CA | ASP | A | 151 | 46.696 | 8.440 | 16.552 | 1.00 | 0.00 | C |
| ATOM | 2054 | HA | ASP | A | 151 | 47.703 | 8.836 | 16.503 | 1.00 | 0.00 | H |
| ATOM | 2055 | CB | ASP | A | 151 | 45.850 | 9.310 | 15.606 | 1.00 | 0.00 | C |
| ATOM | 2056 | HB1 | ASP | A | 151 | 44.889 | 8.819 | 15.468 | 1.00 | 0.00 | H |
| ATOM | 2057 | HB2 | ASP | A | 151 | 45.658 | 10.268 | 16.091 | 1.00 | 0.00 | H |
| ATOM | 2058 | CG | ASP | A | 151 | 46.502 | 9.610 | 14.237 | 1.00 | 0.00 | C |
| ATOM | 2059 | OD1 | ASP | A | 151 | 47.735 | 9.455 | 14.083 | 1.00 | 0.00 | O |
| ATOM | 2060 | OD2 | ASP | A | 151 | 45.767 | 10.069 | 13.326 | 1.00 | 0.00 | O |
| ATOM | 2061 | C | ASP | A | 151 | 46.250 | 8.550 | 18.026 | 1.00 | 0.00 | C |
| ATOM | 2062 | O | ASP | A | 151 | 46.450 | 9.600 | 18.631 | 1.00 | 0.00 | O |
| ATOM | 2063 | N | LYS | A | 152 | 45.730 | 7.484 | 18.655 | 1.00 | 0.00 | N |
| ATOM | 2064 | H | LYS | A | 152 | 45.595 | 6.634 | 18.120 | 1.00 | 0.00 | H |
| ATOM | 2065 | CA | LYS | A | 152 | 45.461 | 7.469 | 20.104 | 1.00 | 0.00 | C |
| ATOM | 2066 | HA | LYS | A | 152 | 44.904 | 8.376 | 20.348 | 1.00 | 0.00 | H |
| ATOM | 2067 | CB | LYS | A | 152 | 44.567 | 6.266 | 20.461 | 1.00 | 0.00 | C |
| ATOM | 2068 | HB1 | LYS | A | 152 | 43.638 | 6.366 | 19.902 | 1.00 | 0.00 | H |
| ATOM | 2069 | HB2 | LYS | A | 152 | 45.048 | 5.338 | 20.158 | 1.00 | 0.00 | H |
| ATOM | 2070 | CG | LYS | A | 152 | 44.239 | 6.187 | 21.963 | 1.00 | 0.00 | C |
| ATOM | 2071 | HG1 | LYS | A | 152 | 45.151 | 5.970 | 22.520 | 1.00 | 0.00 | H |
| ATOM | 2072 | HG2 | LYS | A | 152 | 43.844 | 7.144 | 22.299 | 1.00 | 0.00 | H |
| ATOM | 2073 | CD | LYS | A | 152 | 43.208 | 5.091 | 22.262 | 1.00 | 0.00 | C |
| ATOM | 2074 | HD1 | LYS | A | 152 | 42.314 | 5.257 | 21.662 | 1.00 | 0.00 | H |
| ATOM | 2075 | HD2 | LYS | A | 152 | 43.633 | 4.121 | 21.997 | 1.00 | 0.00 | H |
| ATOM | 2076 | CE | LYS | A | 152 | 42.826 | 5.098 | 23.747 | 1.00 | 0.00 | C |
| ATOM | 2077 | HE1 | LYS | A | 152 | 43.724 | 4.914 | 24.344 | 1.00 | 0.00 | H |
| ATOM | 2078 | HE2 | LYS | A | 152 | 42.427 | 6.080 | 24.016 | 1.00 | 0.00 | H |
| ATOM | 2079 | NZ | LYS | A | 152 | 41.821 | 4.057 | 24.058 | 1.00 | 0.00 | N |
| ATOM | 2080 | HZ1 | LYS | A | 152 | 40.980 | 4.168 | 23.502 | 1.00 | 0.00 | H |
| ATOM | 2081 | HZ2 | LYS | A | 152 | 42.208 | 3.135 | 23.880 | 1.00 | 0.00 | H |
| ATOM | 2082 | HZ3 | LYS | A | 152 | 41.556 | 4.097 | 25.039 | 1.00 | 0.00 | H |
| ATOM | 2083 | C | LYS | A | 152 | 46.763 | 7.539 | 20.927 | 1.00 | 0.00 | C |
| ATOM | 2084 | O | LYS | A | 152 | 46.942 | 8.494 | 21.686 | 1.00 | 0.00 | O |
| ATOM | 2085 | N | GLU | A | 153 | 47.694 | 6.594 | 20.733 | 1.00 | 0.00 | N |
| ATOM | 2086 | H | GLU | A | 153 | 47.477 | 5.837 | 20.091 | 1.00 | 0.00 | H |
| ATOM | 2087 | CA | GLU | A | 153 | 49.034 | 6.618 | 21.350 | 1.00 | 0.00 | C |
| ATOM | 2088 | HA | GLU | A | 153 | 48.935 | 6.644 | 22.436 | 1.00 | 0.00 | H |
| ATOM | 2089 | CB | GLU | A | 153 | 49.840 | 5.364 | 20.949 | 1.00 | 0.00 | C |
| ATOM | 2090 | HB1 | GLU | A | 153 | 49.688 | 5.165 | 19.887 | 1.00 | 0.00 | H |
| ATOM | 2091 | HB2 | GLU | A | 153 | 50.902 | 5.579 | 21.074 | 1.00 | 0.00 | H |
| ATOM | 2092 | CG | GLU | A | 153 | 49.502 | 4.099 | 21.758 | 1.00 | 0.00 | C |
| ATOM | 2093 | HG1 | GLU | A | 153 | 48.678 | 4.299 | 22.445 | 1.00 | 0.00 | H |
| ATOM | 2094 | HG2 | GLU | A | 153 | 49.184 | 3.313 | 21.072 | 1.00 | 0.00 | H |
| ATOM | 2095 | CD | GLU | A | 153 | 50.710 | 3.603 | 22.559 | 1.00 | 0.00 | C |
| ATOM | 2096 | OE1 | GLU | A | 153 | 51.231 | 2.497 | 22.287 | 1.00 | 0.00 | O |
| ATOM | 2097 | OE2 | GLU | A | 153 | 51.155 | 4.328 | 23.483 | 1.00 | 0.00 | O |
| ATOM | 2098 | C | GLU | A | 153 | 49.835 | 7.860 | 20.930 | 1.00 | 0.00 | C |
| ATOM | 2099 | O | GLU | A | 153 | 50.592 | 8.420 | 21.728 | 1.00 | 0.00 | O |
| ATOM | 2100 | N | ARG | A | 154 | 49.700 | 8.306 | 19.676 | 1.00 | 0.00 | N |
| ATOM | 2101 | H | ARG | A | 154 | 49.113 | 7.793 | 19.028 | 1.00 | 0.00 | H |
| ATOM | 2102 | CA | ARG | A | 154 | 50.449 | 9.466 | 19.185 | 1.00 | 0.00 | C |
| ATOM | 2103 | HA | ARG | A | 154 | 51.479 | 9.376 | 19.526 | 1.00 | 0.00 | H |

| | | | | | | | | | | | |
|---|---|---|---|---|---|---|---|---|---|---|---|
| ATOM | 2104 | CB | ARG | A | 154 | 50.493 | 9.460 | 17.655 | 1.00 | 0.00 | C |
| ATOM | 2105 | HB1 | ARG | A | 154 | 49.494 | 9.343 | 17.258 | 1.00 | 0.00 | H |
| ATOM | 2106 | HB2 | ARG | A | 154 | 50.880 | 10.416 | 17.330 | 1.00 | 0.00 | H |
| ATOM | 2107 | CG | ARG | A | 154 | 51.412 | 8.359 | 17.107 | 1.00 | 0.00 | C |
| ATOM | 2108 | HG1 | ARG | A | 154 | 52.423 | 8.535 | 17.480 | 1.00 | 0.00 | H |
| ATOM | 2109 | HG2 | ARG | A | 154 | 51.078 | 7.383 | 17.461 | 1.00 | 0.00 | H |
| ATOM | 2110 | CD | ARG | A | 154 | 51.443 | 8.348 | 15.574 | 1.00 | 0.00 | C |
| ATOM | 2111 | HD1 | ARG | A | 154 | 50.461 | 8.053 | 15.197 | 1.00 | 0.00 | H |
| ATOM | 2112 | HD2 | ARG | A | 154 | 51.679 | 9.348 | 15.215 | 1.00 | 0.00 | H |
| ATOM | 2113 | NE | ARG | A | 154 | 52.484 | 7.426 | 15.104 | 1.00 | 0.00 | N |
| ATOM | 2114 | HE | ARG | A | 154 | 53.247 | 7.246 | 15.749 | 1.00 | 0.00 | H |
| ATOM | 2115 | CZ | ARG | A | 154 | 52.542 | 6.755 | 13.965 | 1.00 | 0.00 | C |
| ATOM | 2116 | NH1 | ARG | A | 154 | 51.666 | 6.910 | 13.000 | 1.00 | 0.00 | N |
| ATOM | 2117 | HH11 | ARG | A | 154 | 50.953 | 7.631 | 13.060 | 1.00 | 0.00 | H |
| ATOM | 2118 | HH12 | ARG | A | 154 | 51.728 | 6.337 | 12.170 | 1.00 | 0.00 | H |
| ATOM | 2119 | NH2 | ARG | A | 154 | 53.519 | 5.898 | 13.784 | 1.00 | 0.00 | N |
| ATOM | 2120 | HH21 | ARG | A | 154 | 54.236 | 5.786 | 14.494 | 1.00 | 0.00 | H |
| ATOM | 2121 | HH22 | ARG | A | 154 | 53.543 | 5.350 | 12.932 | 1.00 | 0.00 | H |
| ATOM | 2122 | C | ARG | A | 154 | 49.936 | 10.799 | 19.753 | 1.00 | 0.00 | C |
| ATOM | 2123 | O | ARG | A | 154 | 50.748 | 11.701 | 19.957 | 1.00 | 0.00 | O |
| ATOM | 2124 | N | PHE | A | 155 | 48.652 | 10.916 | 20.120 | 1.00 | 0.00 | N |
| ATOM | 2125 | H | PHE | A | 155 | 48.015 | 10.161 | 19.905 | 1.00 | 0.00 | H |
| ATOM | 2126 | CA | PHE | A | 155 | 48.103 | 12.129 | 20.744 | 1.00 | 0.00 | C |
| ATOM | 2127 | HA | PHE | A | 155 | 48.399 | 12.994 | 20.146 | 1.00 | 0.00 | H |
| ATOM | 2128 | CB | PHE | A | 155 | 46.563 | 12.063 | 20.767 | 1.00 | 0.00 | C |
| ATOM | 2129 | HB1 | PHE | A | 155 | 46.211 | 11.883 | 19.752 | 1.00 | 0.00 | H |
| ATOM | 2130 | HB2 | PHE | A | 155 | 46.238 | 11.219 | 21.373 | 1.00 | 0.00 | H |
| ATOM | 2131 | CG | PHE | A | 155 | 45.890 | 13.329 | 21.272 | 1.00 | 0.00 | C |
| ATOM | 2132 | CD1 | PHE | A | 155 | 45.465 | 14.308 | 20.353 | 1.00 | 0.00 | C |
| ATOM | 2133 | HD1 | PHE | A | 155 | 45.590 | 14.133 | 19.295 | 1.00 | 0.00 | H |
| ATOM | 2134 | CE1 | PHE | A | 155 | 44.914 | 15.521 | 20.804 | 1.00 | 0.00 | C |
| ATOM | 2135 | HE1 | PHE | A | 155 | 44.621 | 16.276 | 20.089 | 1.00 | 0.00 | H |
| ATOM | 2136 | CZ | PHE | A | 155 | 44.777 | 15.758 | 22.184 | 1.00 | 0.00 | C |
| ATOM | 2137 | HZ | PHE | A | 155 | 44.377 | 16.696 | 22.542 | 1.00 | 0.00 | H |
| ATOM | 2138 | CE2 | PHE | A | 155 | 45.173 | 14.772 | 23.105 | 1.00 | 0.00 | C |
| ATOM | 2139 | HE2 | PHE | A | 155 | 45.065 | 14.949 | 24.167 | 1.00 | 0.00 | H |
| ATOM | 2140 | CD2 | PHE | A | 155 | 45.724 | 13.559 | 22.651 | 1.00 | 0.00 | C |
| ATOM | 2141 | HD2 | PHE | A | 155 | 46.036 | 12.814 | 23.369 | 1.00 | 0.00 | H |
| ATOM | 2142 | C | PHE | A | 155 | 48.686 | 12.301 | 22.150 | 1.00 | 0.00 | C |
| ATOM | 2143 | O | PHE | A | 155 | 49.100 | 13.402 | 22.530 | 1.00 | 0.00 | O |
| ATOM | 2144 | N | THR | A | 156 | 48.792 | 11.189 | 22.885 | 1.00 | 0.00 | N |
| ATOM | 2145 | H | THR | A | 156 | 48.388 | 10.335 | 22.513 | 1.00 | 0.00 | H |
| ATOM | 2146 | CA | THR | A | 156 | 49.423 | 11.094 | 24.204 | 1.00 | 0.00 | C |
| ATOM | 2147 | HA | THR | A | 156 | 48.890 | 11.734 | 24.905 | 1.00 | 0.00 | H |
| ATOM | 2148 | CB | THR | A | 156 | 49.349 | 9.650 | 24.694 | 1.00 | 0.00 | C |
| ATOM | 2149 | HB | THR | A | 156 | 50.027 | 9.023 | 24.119 | 1.00 | 0.00 | H |
| ATOM | 2150 | CG2 | THR | A | 156 | 49.695 | 9.564 | 26.170 | 1.00 | 0.00 | C |
| ATOM | 2151 | HG21 | THR | A | 156 | 49.588 | 8.535 | 26.490 | 1.00 | 0.00 | H |
| ATOM | 2152 | HG22 | THR | A | 156 | 50.729 | 9.867 | 26.325 | 1.00 | 0.00 | H |
| ATOM | 2153 | HG23 | THR | A | 156 | 49.032 | 10.204 | 26.752 | 1.00 | 0.00 | H |
| ATOM | 2154 | OG1 | THR | A | 156 | 48.048 | 9.162 | 24.515 | 1.00 | 0.00 | O |
| ATOM | 2155 | HG1 | THR | A | 156 | 48.098 | 8.182 | 24.630 | 1.00 | 0.00 | H |
| ATOM | 2156 | C | THR | A | 156 | 50.874 | 11.550 | 24.154 | 1.00 | 0.00 | C |
| ATOM | 2157 | O | THR | A | 156 | 51.251 | 12.448 | 24.909 | 1.00 | 0.00 | O |

| | | | | | | | | | | | |
|---|---|---|---|---|---|---|---|---|---|---|---|
| ATOM | 2158 | N | THR | A | 157 | 51.659 | 10.989 | 23.224 | 1.00 | 0.00 | N |
| ATOM | 2159 | H | THR | A | 157 | 51.273 | 10.217 | 22.689 | 1.00 | 0.00 | H |
| ATOM | 2160 | CA | THR | A | 157 | 53.045 | 11.398 | 22.960 | 1.00 | 0.00 | C |
| ATOM | 2161 | HA | THR | A | 157 | 53.628 | 11.302 | 23.878 | 1.00 | 0.00 | H |
| ATOM | 2162 | CB | THR | A | 157 | 53.670 | 10.478 | 21.909 | 1.00 | 0.00 | C |
| ATOM | 2163 | HB | THR | A | 157 | 52.946 | 10.281 | 21.117 | 1.00 | 0.00 | H |
| ATOM | 2164 | CG2 | THR | A | 157 | 54.943 | 11.027 | 21.278 | 1.00 | 0.00 | C |
| ATOM | 2165 | HG21 | THR | A | 157 | 55.397 | 10.258 | 20.654 | 1.00 | 0.00 | H |
| ATOM | 2166 | HG22 | THR | A | 157 | 54.700 | 11.887 | 20.658 | 1.00 | 0.00 | H |
| ATOM | 2167 | HG23 | THR | A | 157 | 55.648 | 11.335 | 22.050 | 1.00 | 0.00 | H |
| ATOM | 2168 | OG1 | THR | A | 157 | 54.038 | 9.270 | 22.539 | 1.00 | 0.00 | O |
| ATOM | 2169 | HG1 | THR | A | 157 | 53.352 | 9.043 | 23.199 | 1.00 | 0.00 | H |
| ATOM | 2170 | C | THR | A | 157 | 53.121 | 12.874 | 22.578 | 1.00 | 0.00 | C |
| ATOM | 2171 | O | THR | A | 157 | 53.936 | 13.576 | 23.156 | 1.00 | 0.00 | O |
| ATOM | 2172 | N | LEU | A | 158 | 52.260 | 13.391 | 21.694 | 1.00 | 0.00 | N |
| ATOM | 2173 | H | LEU | A | 158 | 51.628 | 12.769 | 21.202 | 1.00 | 0.00 | H |
| ATOM | 2174 | CA | LEU | A | 158 | 52.256 | 14.818 | 21.339 | 1.00 | 0.00 | C |
| ATOM | 2175 | HA | LEU | A | 158 | 53.261 | 15.076 | 21.004 | 1.00 | 0.00 | H |
| ATOM | 2176 | CB | LEU | A | 158 | 51.283 | 15.050 | 20.163 | 1.00 | 0.00 | C |
| ATOM | 2177 | HB1 | LEU | A | 158 | 51.631 | 14.478 | 19.301 | 1.00 | 0.00 | H |
| ATOM | 2178 | HB2 | LEU | A | 158 | 50.306 | 14.659 | 20.438 | 1.00 | 0.00 | H |
| ATOM | 2179 | CG | LEU | A | 158 | 51.119 | 16.531 | 19.747 | 1.00 | 0.00 | C |
| ATOM | 2180 | HG | LEU | A | 158 | 50.778 | 17.102 | 20.610 | 1.00 | 0.00 | H |
| ATOM | 2181 | CD1 | LEU | A | 158 | 52.437 | 17.141 | 19.239 | 1.00 | 0.00 | C |
| ATOM | 2182 | HD11 | LEU | A | 158 | 52.268 | 18.172 | 18.929 | 1.00 | 0.00 | H |
| ATOM | 2183 | HD12 | LEU | A | 158 | 53.187 | 17.143 | 20.029 | 1.00 | 0.00 | H |
| ATOM | 2184 | HD13 | LEU | A | 158 | 52.812 | 16.571 | 18.388 | 1.00 | 0.00 | H |
| ATOM | 2185 | CD2 | LEU | A | 158 | 50.050 | 16.680 | 18.655 | 1.00 | 0.00 | C |
| ATOM | 2186 | HD21 | LEU | A | 158 | 49.830 | 17.737 | 18.503 | 1.00 | 0.00 | H |
| ATOM | 2187 | HD22 | LEU | A | 158 | 50.401 | 16.252 | 17.720 | 1.00 | 0.00 | H |
| ATOM | 2188 | HD23 | LEU | A | 158 | 49.131 | 16.175 | 18.949 | 1.00 | 0.00 | H |
| ATOM | 2189 | C | LEU | A | 158 | 51.981 | 15.724 | 22.555 | 1.00 | 0.00 | C |
| ATOM | 2190 | O | LEU | A | 158 | 52.681 | 16.719 | 22.715 | 1.00 | 0.00 | O |
| ATOM | 2191 | N | VAL | A | 159 | 51.048 | 15.362 | 23.443 | 1.00 | 0.00 | N |
| ATOM | 2192 | H | VAL | A | 159 | 50.491 | 14.540 | 23.230 | 1.00 | 0.00 | H |
| ATOM | 2193 | CA | VAL | A | 159 | 50.818 | 16.086 | 24.714 | 1.00 | 0.00 | C |
| ATOM | 2194 | HA | VAL | A | 159 | 50.604 | 17.126 | 24.465 | 1.00 | 0.00 | H |
| ATOM | 2195 | CB | VAL | A | 159 | 49.599 | 15.517 | 25.489 | 1.00 | 0.00 | C |
| ATOM | 2196 | HB | VAL | A | 159 | 49.653 | 14.429 | 25.473 | 1.00 | 0.00 | H |
| ATOM | 2197 | CG1 | VAL | A | 159 | 49.543 | 15.961 | 26.960 | 1.00 | 0.00 | C |
| ATOM | 2198 | HG11 | VAL | A | 159 | 48.642 | 15.575 | 27.433 | 1.00 | 0.00 | H |
| ATOM | 2199 | HG12 | VAL | A | 159 | 50.396 | 15.559 | 27.500 | 1.00 | 0.00 | H |
| ATOM | 2200 | HG13 | VAL | A | 159 | 49.549 | 17.048 | 27.029 | 1.00 | 0.00 | H |
| ATOM | 2201 | CG2 | VAL | A | 159 | 48.279 | 15.933 | 24.823 | 1.00 | 0.00 | C |
| ATOM | 2202 | HG21 | VAL | A | 159 | 47.433 | 15.536 | 25.385 | 1.00 | 0.00 | H |
| ATOM | 2203 | HG22 | VAL | A | 159 | 48.205 | 17.020 | 24.781 | 1.00 | 0.00 | H |
| ATOM | 2204 | HG23 | VAL | A | 159 | 48.233 | 15.533 | 23.816 | 1.00 | 0.00 | H |
| ATOM | 2205 | C | VAL | A | 159 | 52.076 | 16.068 | 25.601 | 1.00 | 0.00 | C |
| ATOM | 2206 | O | VAL | A | 159 | 52.442 | 17.093 | 26.173 | 1.00 | 0.00 | O |
| ATOM | 2207 | N | GLN | A | 160 | 52.745 | 14.916 | 25.706 | 1.00 | 0.00 | N |
| ATOM | 2208 | H | GLN | A | 160 | 52.405 | 14.113 | 25.189 | 1.00 | 0.00 | H |
| ATOM | 2209 | CA | GLN | A | 160 | 53.959 | 14.735 | 26.507 | 1.00 | 0.00 | C |
| ATOM | 2210 | HA | GLN | A | 160 | 53.791 | 15.120 | 27.514 | 1.00 | 0.00 | H |
| ATOM | 2211 | CB | GLN | A | 160 | 54.230 | 13.224 | 26.576 | 1.00 | 0.00 | C |

| | | | | | | | | | | |
|---|---|---|---|---|---|---|---|---|---|---|
| ATOM | 2212 | HB1 | GLN | A | 160 | 53.303 | 12.707 | 26.823 | 1.00 | 0.00 | H |
| ATOM | 2213 | HB2 | GLN | A | 160 | 54.551 | 12.883 | 25.592 | 1.00 | 0.00 | H |
| ATOM | 2214 | CG | GLN | A | 160 | 55.280 | 12.805 | 27.613 | 1.00 | 0.00 | C |
| ATOM | 2215 | HG1 | GLN | A | 160 | 56.223 | 13.312 | 27.426 | 1.00 | 0.00 | H |
| ATOM | 2216 | HG2 | GLN | A | 160 | 54.933 | 13.085 | 28.609 | 1.00 | 0.00 | H |
| ATOM | 2217 | CD | GLN | A | 160 | 55.521 | 11.298 | 27.553 | 1.00 | 0.00 | C |
| ATOM | 2218 | OE1 | GLN | A | 160 | 55.714 | 10.714 | 26.491 | 1.00 | 0.00 | O |
| ATOM | 2219 | NE2 | GLN | A | 160 | 55.467 | 10.591 | 28.660 | 1.00 | 0.00 | N |
| ATOM | 2220 | HE21 | GLN | A | 160 | 55.239 | 11.030 | 29.545 | 1.00 | 0.00 | H |
| ATOM | 2221 | HE22 | GLN | A | 160 | 55.647 | 9.601 | 28.606 | 1.00 | 0.00 | H |
| ATOM | 2222 | C | GLN | A | 160 | 55.159 | 15.492 | 25.910 | 1.00 | 0.00 | C |
| ATOM | 2223 | O | GLN | A | 160 | 56.021 | 15.975 | 26.640 | 1.00 | 0.00 | O |
| ATOM | 2224 | N | ASP | A | 161 | 55.237 | 15.601 | 24.585 | 1.00 | 0.00 | N |
| ATOM | 2225 | H | ASP | A | 161 | 54.556 | 15.087 | 24.031 | 1.00 | 0.00 | H |
| ATOM | 2226 | CA | ASP | A | 161 | 56.285 | 16.311 | 23.850 | 1.00 | 0.00 | C |
| ATOM | 2227 | HA | ASP | A | 161 | 57.257 | 16.091 | 24.282 | 1.00 | 0.00 | H |
| ATOM | 2228 | CB | ASP | A | 161 | 56.295 | 15.841 | 22.387 | 1.00 | 0.00 | C |
| ATOM | 2229 | HB1 | ASP | A | 161 | 55.285 | 15.902 | 21.980 | 1.00 | 0.00 | H |
| ATOM | 2230 | HB2 | ASP | A | 161 | 56.917 | 16.530 | 21.814 | 1.00 | 0.00 | H |
| ATOM | 2231 | CG | ASP | A | 161 | 56.846 | 14.425 | 22.162 | 1.00 | 0.00 | C |
| ATOM | 2232 | OD1 | ASP | A | 161 | 57.140 | 13.691 | 23.141 | 1.00 | 0.00 | O |
| ATOM | 2233 | OD2 | ASP | A | 161 | 57.021 | 14.101 | 20.962 | 1.00 | 0.00 | O |
| ATOM | 2234 | C | ASP | A | 161 | 56.113 | 17.828 | 23.938 | 1.00 | 0.00 | C |
| ATOM | 2235 | O | ASP | A | 161 | 57.104 | 18.533 | 24.138 | 1.00 | 0.00 | O |
| ATOM | 2236 | N | LEU | A | 162 | 54.876 | 18.326 | 23.842 | 1.00 | 0.00 | N |
| ATOM | 2237 | H | LEU | A | 162 | 54.115 | 17.681 | 23.646 | 1.00 | 0.00 | H |
| ATOM | 2238 | CA | LEU | A | 162 | 54.550 | 19.733 | 24.079 | 1.00 | 0.00 | C |
| ATOM | 2239 | HA | LEU | A | 162 | 55.197 | 20.359 | 23.464 | 1.00 | 0.00 | H |
| ATOM | 2240 | CB | LEU | A | 162 | 53.078 | 19.984 | 23.693 | 1.00 | 0.00 | C |
| ATOM | 2241 | HB1 | LEU | A | 162 | 52.469 | 19.200 | 24.145 | 1.00 | 0.00 | H |
| ATOM | 2242 | HB2 | LEU | A | 162 | 52.756 | 20.941 | 24.108 | 1.00 | 0.00 | H |
| ATOM | 2243 | CG | LEU | A | 162 | 52.830 | 20.001 | 22.169 | 1.00 | 0.00 | C |
| ATOM | 2244 | HG | LEU | A | 162 | 53.394 | 19.193 | 21.702 | 1.00 | 0.00 | H |
| ATOM | 2245 | CD1 | LEU | A | 162 | 51.342 | 19.783 | 21.864 | 1.00 | 0.00 | C |
| ATOM | 2246 | HD11 | LEU | A | 162 | 51.189 | 19.727 | 20.786 | 1.00 | 0.00 | H |
| ATOM | 2247 | HD12 | LEU | A | 162 | 50.999 | 18.852 | 22.316 | 1.00 | 0.00 | H |
| ATOM | 2248 | HD13 | LEU | A | 162 | 50.756 | 20.609 | 22.263 | 1.00 | 0.00 | H |
| ATOM | 2249 | CD2 | LEU | A | 162 | 53.262 | 21.336 | 21.542 | 1.00 | 0.00 | C |
| ATOM | 2250 | HD21 | LEU | A | 162 | 53.203 | 21.269 | 20.456 | 1.00 | 0.00 | H |
| ATOM | 2251 | HD22 | LEU | A | 162 | 52.607 | 22.137 | 21.882 | 1.00 | 0.00 | H |
| ATOM | 2252 | HD23 | LEU | A | 162 | 54.285 | 21.580 | 21.825 | 1.00 | 0.00 | H |
| ATOM | 2253 | C | LEU | A | 162 | 54.848 | 20.110 | 25.535 | 1.00 | 0.00 | C |
| ATOM | 2254 | O | LEU | A | 162 | 55.607 | 21.046 | 25.768 | 1.00 | 0.00 | O |
| ATOM | 2255 | N | ALA | A | 163 | 54.336 | 19.359 | 26.516 | 1.00 | 0.00 | N |
| ATOM | 2256 | H | ALA | A | 163 | 53.742 | 18.578 | 26.260 | 1.00 | 0.00 | H |
| ATOM | 2257 | CA | ALA | A | 163 | 54.526 | 19.650 | 27.941 | 1.00 | 0.00 | C |
| ATOM | 2258 | HA | ALA | A | 163 | 54.115 | 20.641 | 28.144 | 1.00 | 0.00 | H |
| ATOM | 2259 | CB | ALA | A | 163 | 53.731 | 18.622 | 28.756 | 1.00 | 0.00 | C |
| ATOM | 2260 | HB1 | ALA | A | 163 | 53.833 | 18.834 | 29.821 | 1.00 | 0.00 | H |
| ATOM | 2261 | HB2 | ALA | A | 163 | 52.675 | 18.675 | 28.488 | 1.00 | 0.00 | H |
| ATOM | 2262 | HB3 | ALA | A | 163 | 54.104 | 17.615 | 28.559 | 1.00 | 0.00 | H |
| ATOM | 2263 | C | ALA | A | 163 | 56.009 | 19.684 | 28.358 | 1.00 | 0.00 | C |
| ATOM | 2264 | O | ALA | A | 163 | 56.437 | 20.647 | 29.006 | 1.00 | 0.00 | O |
| ATOM | 2265 | N | ASN | A | 164 | 56.799 | 18.680 | 27.952 | 1.00 | 0.00 | N |

| | | | | | | | | | | | |
|---|---|---|---|---|---|---|---|---|---|---|---|
| ATOM | 2266 | H | ASN | A | 164 | 56.380 | 17.904 | 27.449 | 1.00 | 0.00 | H |
| ATOM | 2267 | CA | ASN | A | 164 | 58.231 | 18.610 | 28.269 | 1.00 | 0.00 | C |
| ATOM | 2268 | HA | ASN | A | 164 | 58.352 | 18.790 | 29.337 | 1.00 | 0.00 | H |
| ATOM | 2269 | CB | ASN | A | 164 | 58.771 | 17.197 | 27.963 | 1.00 | 0.00 | C |
| ATOM | 2270 | HB1 | ASN | A | 164 | 58.496 | 16.916 | 26.946 | 1.00 | 0.00 | H |
| ATOM | 2271 | HB2 | ASN | A | 164 | 59.860 | 17.218 | 28.010 | 1.00 | 0.00 | H |
| ATOM | 2272 | CG | ASN | A | 164 | 58.299 | 16.112 | 28.940 | 1.00 | 0.00 | C |
| ATOM | 2273 | OD1 | ASN | A | 164 | 57.846 | 16.375 | 30.048 | 1.00 | 0.00 | O |
| ATOM | 2274 | ND2 | ASN | A | 164 | 58.435 | 14.847 | 28.587 | 1.00 | 0.00 | N |
| ATOM | 2275 | HD21 | ASN | A | 164 | 58.795 | 14.593 | 27.679 | 1.00 | 0.00 | H |
| ATOM | 2276 | HD22 | ASN | A | 164 | 58.120 | 14.131 | 29.230 | 1.00 | 0.00 | H |
| ATOM | 2277 | C | ASN | A | 164 | 59.051 | 19.703 | 27.548 | 1.00 | 0.00 | C |
| ATOM | 2278 | O | ASN | A | 164 | 60.020 | 20.216 | 28.123 | 1.00 | 0.00 | O |
| ATOM | 2279 | N | ALA | A | 165 | 58.656 | 20.115 | 26.333 | 1.00 | 0.00 | N |
| ATOM | 2280 | H | ALA | A | 165 | 57.856 | 19.663 | 25.903 | 1.00 | 0.00 | H |
| ATOM | 2281 | CA | ALA | A | 165 | 59.301 | 21.209 | 25.595 | 1.00 | 0.00 | C |
| ATOM | 2282 | HA | ALA | A | 165 | 60.385 | 21.082 | 25.647 | 1.00 | 0.00 | H |
| ATOM | 2283 | CB | ALA | A | 165 | 58.878 | 21.140 | 24.126 | 1.00 | 0.00 | C |
| ATOM | 2284 | HB1 | ALA | A | 165 | 59.360 | 21.941 | 23.564 | 1.00 | 0.00 | H |
| ATOM | 2285 | HB2 | ALA | A | 165 | 59.187 | 20.193 | 23.693 | 1.00 | 0.00 | H |
| ATOM | 2286 | HB3 | ALA | A | 165 | 57.795 | 21.243 | 24.035 | 1.00 | 0.00 | H |
| ATOM | 2287 | C | ALA | A | 165 | 58.965 | 22.592 | 26.168 | 1.00 | 0.00 | C |
| ATOM | 2288 | O | ALA | A | 165 | 59.853 | 23.443 | 26.253 | 1.00 | 0.00 | O |
| ATOM | 2289 | N | PHE | A | 166 | 57.709 | 22.801 | 26.585 | 1.00 | 0.00 | N |
| ATOM | 2290 | H | PHE | A | 166 | 57.023 | 22.069 | 26.423 | 1.00 | 0.00 | H |
| ATOM | 2291 | CA | PHE | A | 166 | 57.229 | 24.047 | 27.183 | 1.00 | 0.00 | C |
| ATOM | 2292 | HA | PHE | A | 166 | 57.480 | 24.882 | 26.526 | 1.00 | 0.00 | H |
| ATOM | 2293 | CB | PHE | A | 166 | 55.697 | 24.008 | 27.362 | 1.00 | 0.00 | C |
| ATOM | 2294 | HB1 | PHE | A | 166 | 55.431 | 23.103 | 27.910 | 1.00 | 0.00 | H |
| ATOM | 2295 | HB2 | PHE | A | 166 | 55.421 | 24.854 | 27.993 | 1.00 | 0.00 | H |
| ATOM | 2296 | CG | PHE | A | 166 | 54.811 | 24.083 | 26.120 | 1.00 | 0.00 | C |
| ATOM | 2297 | CD1 | PHE | A | 166 | 55.326 | 24.004 | 24.807 | 1.00 | 0.00 | C |
| ATOM | 2298 | HD1 | PHE | A | 166 | 56.384 | 23.896 | 24.634 | 1.00 | 0.00 | H |
| ATOM | 2299 | CE1 | PHE | A | 166 | 54.462 | 24.032 | 23.699 | 1.00 | 0.00 | C |
| ATOM | 2300 | HE1 | PHE | A | 166 | 54.867 | 23.940 | 22.700 | 1.00 | 0.00 | H |
| ATOM | 2301 | CZ | PHE | A | 166 | 53.077 | 24.172 | 23.887 | 1.00 | 0.00 | C |
| ATOM | 2302 | HZ | PHE | A | 166 | 52.410 | 24.197 | 23.037 | 1.00 | 0.00 | H |
| ATOM | 2303 | CE2 | PHE | A | 166 | 52.557 | 24.269 | 25.187 | 1.00 | 0.00 | C |
| ATOM | 2304 | HE2 | PHE | A | 166 | 51.492 | 24.373 | 25.337 | 1.00 | 0.00 | H |
| ATOM | 2305 | CD2 | PHE | A | 166 | 53.419 | 24.215 | 26.296 | 1.00 | 0.00 | C |
| ATOM | 2306 | HD2 | PHE | A | 166 | 53.005 | 24.271 | 27.292 | 1.00 | 0.00 | H |
| ATOM | 2307 | C | PHE | A | 166 | 57.928 | 24.299 | 28.521 | 1.00 | 0.00 | C |
| ATOM | 2308 | O | PHE | A | 166 | 58.332 | 25.433 | 28.780 | 1.00 | 0.00 | O |
| ATOM | 2309 | N | GLN | A | 167 | 58.150 | 23.265 | 29.348 | 1.00 | 0.00 | N |
| ATOM | 2310 | H | GLN | A | 167 | 57.750 | 22.355 | 29.134 | 1.00 | 0.00 | H |
| ATOM | 2311 | CA | GLN | A | 167 | 59.014 | 23.429 | 30.518 | 1.00 | 0.00 | C |
| ATOM | 2312 | HA | GLN | A | 167 | 58.655 | 24.295 | 31.077 | 1.00 | 0.00 | H |
| ATOM | 2313 | CB | GLN | A | 167 | 58.967 | 22.219 | 31.470 | 1.00 | 0.00 | C |
| ATOM | 2314 | HB1 | GLN | A | 167 | 57.943 | 22.077 | 31.821 | 1.00 | 0.00 | H |
| ATOM | 2315 | HB2 | GLN | A | 167 | 59.298 | 21.318 | 30.951 | 1.00 | 0.00 | H |
| ATOM | 2316 | CG | GLN | A | 167 | 59.894 | 22.500 | 32.675 | 1.00 | 0.00 | C |
| ATOM | 2317 | HG1 | GLN | A | 167 | 60.929 | 22.492 | 32.338 | 1.00 | 0.00 | H |
| ATOM | 2318 | HG2 | GLN | A | 167 | 59.671 | 23.492 | 33.070 | 1.00 | 0.00 | H |
| ATOM | 2319 | CD | GLN | A | 167 | 59.812 | 21.514 | 33.831 | 1.00 | 0.00 | C |

| | | | | | | | | | | | |
|---|---|---|---|---|---|---|---|---|---|---|---|
| ATOM | 2320 | OE1 | GLN | A | 167 | 59.230 | 20.436 | 33.752 | 1.00 | 0.00 | O |
| ATOM | 2321 | NE2 | GLN | A | 167 | 60.423 | 21.834 | 34.948 | 1.00 | 0.00 | N |
| ATOM | 2322 | HE21 | GLN | A | 167 | 60.939 | 22.707 | 35.022 | 1.00 | 0.00 | H |
| ATOM | 2323 | HE22 | GLN | A | 167 | 60.333 | 21.221 | 35.752 | 1.00 | 0.00 | H |
| ATOM | 2324 | C | GLN | A | 167 | 60.451 | 23.770 | 30.108 | 1.00 | 0.00 | C |
| ATOM | 2325 | O | GLN | A | 167 | 61.014 | 24.684 | 30.699 | 1.00 | 0.00 | O |
| ATOM | 2326 | N | GLN | A | 168 | 61.057 | 23.095 | 29.123 | 1.00 | 0.00 | N |
| ATOM | 2327 | H | GLN | A | 168 | 60.571 | 22.347 | 28.644 | 1.00 | 0.00 | H |
| ATOM | 2328 | CA | GLN | A | 168 | 62.461 | 23.359 | 28.790 | 1.00 | 0.00 | C |
| ATOM | 2329 | HA | GLN | A | 168 | 63.028 | 23.263 | 29.716 | 1.00 | 0.00 | H |
| ATOM | 2330 | CB | GLN | A | 168 | 62.987 | 22.318 | 27.784 | 1.00 | 0.00 | C |
| ATOM | 2331 | HB1 | GLN | A | 168 | 62.692 | 21.320 | 28.112 | 1.00 | 0.00 | H |
| ATOM | 2332 | HB2 | GLN | A | 168 | 62.538 | 22.503 | 26.807 | 1.00 | 0.00 | H |
| ATOM | 2333 | CG | GLN | A | 168 | 64.523 | 22.347 | 27.645 | 1.00 | 0.00 | C |
| ATOM | 2334 | HG1 | GLN | A | 168 | 64.818 | 21.639 | 26.870 | 1.00 | 0.00 | H |
| ATOM | 2335 | HG2 | GLN | A | 168 | 64.847 | 23.338 | 27.330 | 1.00 | 0.00 | H |
| ATOM | 2336 | CD | GLN | A | 168 | 65.236 | 21.986 | 28.949 | 1.00 | 0.00 | C |
| ATOM | 2337 | OE1 | GLN | A | 168 | 65.427 | 22.804 | 29.838 | 1.00 | 0.00 | O |
| ATOM | 2338 | NE2 | GLN | A | 168 | 65.614 | 20.747 | 29.169 | 1.00 | 0.00 | N |
| ATOM | 2339 | HE21 | GLN | A | 168 | 65.587 | 20.052 | 28.437 | 1.00 | 0.00 | H |
| ATOM | 2340 | HE22 | GLN | A | 168 | 66.028 | 20.543 | 30.066 | 1.00 | 0.00 | H |
| ATOM | 2341 | C | GLN | A | 168 | 62.688 | 24.804 | 28.314 | 1.00 | 0.00 | C |
| ATOM | 2342 | O | GLN | A | 168 | 63.667 | 25.423 | 28.728 | 1.00 | 0.00 | O |
| ATOM | 2343 | N | GLU | A | 169 | 61.763 | 25.388 | 27.543 | 1.00 | 0.00 | N |
| ATOM | 2344 | H | GLU | A | 169 | 60.984 | 24.836 | 27.198 | 1.00 | 0.00 | H |
| ATOM | 2345 | CA | GLU | A | 169 | 61.822 | 26.815 | 27.200 | 1.00 | 0.00 | C |
| ATOM | 2346 | HA | GLU | A | 169 | 62.796 | 27.006 | 26.753 | 1.00 | 0.00 | H |
| ATOM | 2347 | CB | GLU | A | 169 | 60.775 | 27.148 | 26.125 | 1.00 | 0.00 | C |
| ATOM | 2348 | HB1 | GLU | A | 169 | 60.950 | 26.481 | 25.280 | 1.00 | 0.00 | H |
| ATOM | 2349 | HB2 | GLU | A | 169 | 59.769 | 26.943 | 26.493 | 1.00 | 0.00 | H |
| ATOM | 2350 | CG | GLU | A | 169 | 60.891 | 28.595 | 25.603 | 1.00 | 0.00 | C |
| ATOM | 2351 | HG1 | GLU | A | 169 | 61.941 | 28.890 | 25.576 | 1.00 | 0.00 | H |
| ATOM | 2352 | HG2 | GLU | A | 169 | 60.528 | 28.619 | 24.578 | 1.00 | 0.00 | H |
| ATOM | 2353 | CD | GLU | A | 169 | 60.096 | 29.641 | 26.386 | 1.00 | 0.00 | C |
| ATOM | 2354 | OE1 | GLU | A | 169 | 59.308 | 29.284 | 27.289 | 1.00 | 0.00 | O |
| ATOM | 2355 | OE2 | GLU | A | 169 | 60.210 | 30.844 | 26.055 | 1.00 | 0.00 | O |
| ATOM | 2356 | C | GLU | A | 169 | 61.710 | 27.710 | 28.448 | 1.00 | 0.00 | C |
| ATOM | 2357 | O | GLU | A | 169 | 62.439 | 28.695 | 28.563 | 1.00 | 0.00 | O |
| ATOM | 2358 | N | ALA | A | 170 | 60.885 | 27.352 | 29.438 | 1.00 | 0.00 | N |
| ATOM | 2359 | H | ALA | A | 170 | 60.328 | 26.515 | 29.325 | 1.00 | 0.00 | H |
| ATOM | 2360 | CA | ALA | A | 170 | 60.770 | 28.099 | 30.695 | 1.00 | 0.00 | C |
| ATOM | 2361 | HA | ALA | A | 170 | 60.651 | 29.159 | 30.456 | 1.00 | 0.00 | H |
| ATOM | 2362 | CB | ALA | A | 170 | 59.509 | 27.633 | 31.428 | 1.00 | 0.00 | C |
| ATOM | 2363 | HB1 | ALA | A | 170 | 59.331 | 28.275 | 32.292 | 1.00 | 0.00 | H |
| ATOM | 2364 | HB2 | ALA | A | 170 | 58.658 | 27.703 | 30.755 | 1.00 | 0.00 | H |
| ATOM | 2365 | HB3 | ALA | A | 170 | 59.621 | 26.603 | 31.766 | 1.00 | 0.00 | H |
| ATOM | 2366 | C | ALA | A | 170 | 62.019 | 27.973 | 31.588 | 1.00 | 0.00 | C |
| ATOM | 2367 | O | ALA | A | 170 | 62.348 | 28.917 | 32.307 | 1.00 | 0.00 | O |
| ATOM | 2368 | N | GLN | A | 171 | 62.732 | 26.842 | 31.545 | 1.00 | 0.00 | N |
| ATOM | 2369 | H | GLN | A | 171 | 62.374 | 26.083 | 30.972 | 1.00 | 0.00 | H |
| ATOM | 2370 | CA | GLN | A | 171 | 63.997 | 26.663 | 32.271 | 1.00 | 0.00 | C |
| ATOM | 2371 | HA | GLN | A | 171 | 63.903 | 27.087 | 33.274 | 1.00 | 0.00 | H |
| ATOM | 2372 | CB | GLN | A | 171 | 64.359 | 25.168 | 32.385 | 1.00 | 0.00 | C |
| ATOM | 2373 | HB1 | GLN | A | 171 | 64.484 | 24.771 | 31.378 | 1.00 | 0.00 | H |

| | | | | | | | | | | | |
|---|---|---|---|---|---|---|---|---|---|---|---|
| ATOM | 2374 | HB2 | GLN | A | 171 | 65.325 | 25.085 | 32.883 | 1.00 | 0.00 | H |
| ATOM | 2375 | CG | GLN | A | 171 | 63.366 | 24.253 | 33.126 | 1.00 | 0.00 | C |
| ATOM | 2376 | HG1 | GLN | A | 171 | 62.446 | 24.204 | 32.560 | 1.00 | 0.00 | H |
| ATOM | 2377 | HG2 | GLN | A | 171 | 63.792 | 23.250 | 33.150 | 1.00 | 0.00 | H |
| ATOM | 2378 | CD | GLN | A | 171 | 63.008 | 24.639 | 34.561 | 1.00 | 0.00 | C |
| ATOM | 2379 | OE1 | GLN | A | 171 | 61.888 | 24.411 | 35.013 | 1.00 | 0.00 | O |
| ATOM | 2380 | NE2 | GLN | A | 171 | 63.914 | 25.197 | 35.335 | 1.00 | 0.00 | N |
| ATOM | 2381 | HE21 | GLN | A | 171 | 64.860 | 25.361 | 35.025 | 1.00 | 0.00 | H |
| ATOM | 2382 | HE22 | GLN | A | 171 | 63.680 | 25.277 | 36.316 | 1.00 | 0.00 | H |
| ATOM | 2383 | C | GLN | A | 171 | 65.164 | 27.395 | 31.586 | 1.00 | 0.00 | C |
| ATOM | 2384 | O | GLN | A | 171 | 66.048 | 27.907 | 32.282 | 1.00 | 0.00 | O |
| ATOM | 2385 | N | THR | A | 172 | 65.184 | 27.428 | 30.243 | 1.00 | 0.00 | N |
| ATOM | 2386 | H | THR | A | 172 | 64.437 | 26.955 | 29.744 | 1.00 | 0.00 | H |
| ATOM | 2387 | CA | THR | A | 172 | 66.274 | 28.010 | 29.437 | 1.00 | 0.00 | C |
| ATOM | 2388 | HA | THR | A | 172 | 67.203 | 27.906 | 29.992 | 1.00 | 0.00 | H |
| ATOM | 2389 | CB | THR | A | 172 | 66.460 | 27.272 | 28.107 | 1.00 | 0.00 | C |
| ATOM | 2390 | HB | THR | A | 172 | 67.167 | 27.831 | 27.494 | 1.00 | 0.00 | H |
| ATOM | 2391 | CG2 | THR | A | 172 | 67.018 | 25.863 | 28.310 | 1.00 | 0.00 | C |
| ATOM | 2392 | HG21 | THR | A | 172 | 67.155 | 25.383 | 27.342 | 1.00 | 0.00 | H |
| ATOM | 2393 | HG22 | THR | A | 172 | 67.986 | 25.923 | 28.808 | 1.00 | 0.00 | H |
| ATOM | 2394 | HG23 | THR | A | 172 | 66.339 | 25.263 | 28.916 | 1.00 | 0.00 | H |
| ATOM | 2395 | OG1 | THR | A | 172 | 65.242 | 27.185 | 27.412 | 1.00 | 0.00 | O |
| ATOM | 2396 | HG1 | THR | A | 172 | 64.719 | 26.472 | 27.826 | 1.00 | 0.00 | H |
| ATOM | 2397 | C | THR | A | 172 | 66.115 | 29.504 | 29.183 | 1.00 | 0.00 | C |
| ATOM | 2398 | O | THR | A | 172 | 67.113 | 30.206 | 29.049 | 1.00 | 0.00 | O |
| ATOM | 2399 | N | SER | A | 173 | 64.890 | 30.020 | 29.156 | 1.00 | 0.00 | N |
| ATOM | 2400 | H | SER | A | 173 | 64.099 | 29.385 | 29.118 | 1.00 | 0.00 | H |
| ATOM | 2401 | CA | SER | A | 173 | 64.644 | 31.467 | 29.140 | 1.00 | 0.00 | C |
| ATOM | 2402 | HA | SER | A | 173 | 65.453 | 31.960 | 28.602 | 1.00 | 0.00 | H |
| ATOM | 2403 | CB | SER | A | 173 | 63.341 | 31.785 | 28.390 | 1.00 | 0.00 | C |
| ATOM | 2404 | HB1 | SER | A | 173 | 63.239 | 32.867 | 28.316 | 1.00 | 0.00 | H |
| ATOM | 2405 | HB2 | SER | A | 173 | 63.396 | 31.376 | 27.380 | 1.00 | 0.00 | H |
| ATOM | 2406 | OG | SER | A | 173 | 62.202 | 31.254 | 29.046 | 1.00 | 0.00 | O |
| ATOM | 2407 | HG | SER | A | 173 | 62.127 | 30.312 | 28.773 | 1.00 | 0.00 | H |
| ATOM | 2408 | C | SER | A | 173 | 64.615 | 32.075 | 30.553 | 1.00 | 0.00 | C |
| ATOM | 2409 | O | SER | A | 173 | 65.225 | 33.122 | 30.782 | 1.00 | 0.00 | O |
| ATOM | 2410 | N | GLY | A | 174 | 63.945 | 31.423 | 31.513 | 1.00 | 0.00 | N |
| ATOM | 2411 | H | GLY | A | 174 | 63.522 | 30.535 | 31.275 | 1.00 | 0.00 | H |
| ATOM | 2412 | CA | GLY | A | 174 | 63.601 | 31.988 | 32.828 | 1.00 | 0.00 | C |
| ATOM | 2413 | HA1 | GLY | A | 174 | 63.596 | 31.185 | 33.564 | 1.00 | 0.00 | H |
| ATOM | 2414 | HA2 | GLY | A | 174 | 64.333 | 32.744 | 33.115 | 1.00 | 0.00 | H |
| ATOM | 2415 | C | GLY | A | 174 | 62.214 | 32.645 | 32.869 | 1.00 | 0.00 | C |
| ATOM | 2416 | O | GLY | A | 174 | 61.828 | 33.193 | 33.900 | 1.00 | 0.00 | O |
| ATOM | 2417 | N | LYS | A | 175 | 61.449 | 32.607 | 31.773 | 1.00 | 0.00 | N |
| ATOM | 2418 | H | LYS | A | 175 | 61.818 | 32.136 | 30.955 | 1.00 | 0.00 | H |
| ATOM | 2419 | CA | LYS | A | 175 | 60.096 | 33.182 | 31.672 | 1.00 | 0.00 | C |
| ATOM | 2420 | HA | LYS | A | 175 | 60.099 | 34.182 | 32.109 | 1.00 | 0.00 | H |
| ATOM | 2421 | CB | LYS | A | 175 | 59.714 | 33.280 | 30.187 | 1.00 | 0.00 | C |
| ATOM | 2422 | HB1 | LYS | A | 175 | 59.793 | 32.284 | 29.748 | 1.00 | 0.00 | H |
| ATOM | 2423 | HB2 | LYS | A | 175 | 58.677 | 33.609 | 30.095 | 1.00 | 0.00 | H |
| ATOM | 2424 | CG | LYS | A | 175 | 60.593 | 34.253 | 29.396 | 1.00 | 0.00 | C |
| ATOM | 2425 | HG1 | LYS | A | 175 | 60.399 | 35.272 | 29.734 | 1.00 | 0.00 | H |
| ATOM | 2426 | HG2 | LYS | A | 175 | 61.647 | 34.025 | 29.556 | 1.00 | 0.00 | H |
| ATOM | 2427 | CD | LYS | A | 175 | 60.277 | 34.131 | 27.903 | 1.00 | 0.00 | C |

| | | | | | | | | | | | |
|---|---|---|---|---|---|---|---|---|---|---|---|
| ATOM | 2428 | HD1 | LYS | A | 175 | 60.420 | 33.097 | 27.582 | 1.00 | 0.00 | H |
| ATOM | 2429 | HD2 | LYS | A | 175 | 59.242 | 34.427 | 27.723 | 1.00 | 0.00 | H |
| ATOM | 2430 | CE | LYS | A | 175 | 61.223 | 35.041 | 27.124 | 1.00 | 0.00 | C |
| ATOM | 2431 | HE1 | LYS | A | 175 | 61.107 | 36.064 | 27.489 | 1.00 | 0.00 | H |
| ATOM | 2432 | HE2 | LYS | A | 175 | 62.254 | 34.727 | 27.307 | 1.00 | 0.00 | H |
| ATOM | 2433 | NZ | LYS | A | 175 | 60.939 | 35.001 | 25.677 | 1.00 | 0.00 | N |
| ATOM | 2434 | HZ1 | LYS | A | 175 | 61.031 | 34.054 | 25.320 | 1.00 | 0.00 | H |
| ATOM | 2435 | HZ2 | LYS | A | 175 | 60.005 | 35.347 | 25.481 | 1.00 | 0.00 | H |
| ATOM | 2436 | HZ3 | LYS | A | 175 | 61.596 | 35.575 | 25.160 | 1.00 | 0.00 | H |
| ATOM | 2437 | C | LYS | A | 175 | 59.030 | 32.354 | 32.413 | 1.00 | 0.00 | C |
| ATOM | 2438 | O | LYS | A | 175 | 59.249 | 31.183 | 32.732 | 1.00 | 0.00 | O |
| ATOM | 2439 | N | GLU | A | 176 | 57.844 | 32.936 | 32.608 | 1.00 | 0.00 | N |
| ATOM | 2440 | H | GLU | A | 176 | 57.731 | 33.906 | 32.336 | 1.00 | 0.00 | H |
| ATOM | 2441 | CA | GLU | A | 176 | 56.638 | 32.210 | 33.015 | 1.00 | 0.00 | C |
| ATOM | 2442 | HA | GLU | A | 176 | 56.837 | 31.703 | 33.959 | 1.00 | 0.00 | H |
| ATOM | 2443 | CB | GLU | A | 176 | 55.512 | 33.227 | 33.244 | 1.00 | 0.00 | C |
| ATOM | 2444 | HB1 | GLU | A | 176 | 55.873 | 33.988 | 33.937 | 1.00 | 0.00 | H |
| ATOM | 2445 | HB2 | GLU | A | 176 | 55.274 | 33.715 | 32.298 | 1.00 | 0.00 | H |
| ATOM | 2446 | CG | GLU | A | 176 | 54.238 | 32.614 | 33.831 | 1.00 | 0.00 | C |
| ATOM | 2447 | HG1 | GLU | A | 176 | 53.827 | 31.878 | 33.140 | 1.00 | 0.00 | H |
| ATOM | 2448 | HG2 | GLU | A | 176 | 54.473 | 32.117 | 34.773 | 1.00 | 0.00 | H |
| ATOM | 2449 | CD | GLU | A | 176 | 53.211 | 33.714 | 34.071 | 1.00 | 0.00 | C |
| ATOM | 2450 | OE1 | GLU | A | 176 | 52.641 | 34.227 | 33.075 | 1.00 | 0.00 | O |
| ATOM | 2451 | OE2 | GLU | A | 176 | 53.045 | 34.137 | 35.244 | 1.00 | 0.00 | O |
| ATOM | 2452 | C | GLU | A | 176 | 56.259 | 31.161 | 31.951 | 1.00 | 0.00 | C |
| ATOM | 2453 | O | GLU | A | 176 | 56.185 | 31.478 | 30.760 | 1.00 | 0.00 | O |
| ATOM | 2454 | N | ARG | A | 177 | 56.052 | 29.909 | 32.372 | 1.00 | 0.00 | N |
| ATOM | 2455 | H | ARG | A | 177 | 56.109 | 29.728 | 33.368 | 1.00 | 0.00 | H |
| ATOM | 2456 | CA | ARG | A | 177 | 55.877 | 28.750 | 31.487 | 1.00 | 0.00 | C |
| ATOM | 2457 | HA | ARG | A | 177 | 56.726 | 28.730 | 30.806 | 1.00 | 0.00 | H |
| ATOM | 2458 | CB | ARG | A | 177 | 55.902 | 27.461 | 32.335 | 1.00 | 0.00 | C |
| ATOM | 2459 | HB1 | ARG | A | 177 | 56.799 | 27.461 | 32.957 | 1.00 | 0.00 | H |
| ATOM | 2460 | HB2 | ARG | A | 177 | 55.032 | 27.448 | 32.991 | 1.00 | 0.00 | H |
| ATOM | 2461 | CG | ARG | A | 177 | 55.903 | 26.187 | 31.476 | 1.00 | 0.00 | C |
| ATOM | 2462 | HG1 | ARG | A | 177 | 55.035 | 26.188 | 30.829 | 1.00 | 0.00 | H |
| ATOM | 2463 | HG2 | ARG | A | 177 | 56.789 | 26.193 | 30.844 | 1.00 | 0.00 | H |
| ATOM | 2464 | CD | ARG | A | 177 | 55.895 | 24.880 | 32.279 | 1.00 | 0.00 | C |
| ATOM | 2465 | HD1 | ARG | A | 177 | 55.941 | 24.047 | 31.577 | 1.00 | 0.00 | H |
| ATOM | 2466 | HD2 | ARG | A | 177 | 56.796 | 24.843 | 32.895 | 1.00 | 0.00 | H |
| ATOM | 2467 | NE | ARG | A | 177 | 54.720 | 24.711 | 33.153 | 1.00 | 0.00 | N |
| ATOM | 2468 | HE | ARG | A | 177 | 54.912 | 24.602 | 34.140 | 1.00 | 0.00 | H |
| ATOM | 2469 | CZ | ARG | A | 177 | 53.456 | 24.481 | 32.806 | 1.00 | 0.00 | C |
| ATOM | 2470 | NH1 | ARG | A | 177 | 52.987 | 24.526 | 31.580 | 1.00 | 0.00 | N |
| ATOM | 2471 | HH11 | ARG | A | 177 | 53.542 | 24.846 | 30.798 | 1.00 | 0.00 | H |
| ATOM | 2472 | HH12 | ARG | A | 177 | 52.023 | 24.246 | 31.418 | 1.00 | 0.00 | H |
| ATOM | 2473 | NH2 | ARG | A | 177 | 52.600 | 24.179 | 33.745 | 1.00 | 0.00 | N |
| ATOM | 2474 | HH21 | ARG | A | 177 | 52.915 | 24.065 | 34.699 | 1.00 | 0.00 | H |
| ATOM | 2475 | HH22 | ARG | A | 177 | 51.625 | 24.012 | 33.513 | 1.00 | 0.00 | H |
| ATOM | 2476 | C | ARG | A | 177 | 54.598 | 28.840 | 30.640 | 1.00 | 0.00 | C |
| ATOM | 2477 | O | ARG | A | 177 | 53.530 | 29.168 | 31.167 | 1.00 | 0.00 | O |
| ATOM | 2478 | N | LEU | A | 178 | 54.699 | 28.485 | 29.353 | 1.00 | 0.00 | N |
| ATOM | 2479 | H | LEU | A | 178 | 55.624 | 28.254 | 29.005 | 1.00 | 0.00 | H |
| ATOM | 2480 | CA | LEU | A | 178 | 53.570 | 28.287 | 28.435 | 1.00 | 0.00 | C |
| ATOM | 2481 | HA | LEU | A | 178 | 52.946 | 29.179 | 28.450 | 1.00 | 0.00 | H |

```
ATOM   2482  CB  LEU A 178      54.099  28.053  27.003  1.00  0.00           C
ATOM   2483  HB1 LEU A 178      54.790  27.208  27.031  1.00  0.00           H
ATOM   2484  HB2 LEU A 178      53.263  27.766  26.363  1.00  0.00           H
ATOM   2485  CG  LEU A 178      54.817  29.257  26.357  1.00  0.00           C
ATOM   2486  HG  LEU A 178      55.590  29.618  27.034  1.00  0.00           H
ATOM   2487  CD1 LEU A 178      55.503  28.814  25.056  1.00  0.00           C
ATOM   2488 HD11 LEU A 178      56.015  29.661  24.599  1.00  0.00           H
ATOM   2489 HD12 LEU A 178      56.241  28.041  25.274  1.00  0.00           H
ATOM   2490 HD13 LEU A 178      54.769  28.418  24.356  1.00  0.00           H
ATOM   2491  CD2 LEU A 178      53.846  30.411  26.065  1.00  0.00           C
ATOM   2492 HD21 LEU A 178      54.371  31.215  25.550  1.00  0.00           H
ATOM   2493 HD22 LEU A 178      53.024  30.064  25.437  1.00  0.00           H
ATOM   2494 HD23 LEU A 178      53.442  30.808  26.996  1.00  0.00           H
ATOM   2495  C   LEU A 178      52.701  27.096  28.872  1.00  0.00           C
ATOM   2496  O   LEU A 178      53.222  26.027  29.195  1.00  0.00           O
ATOM   2497  N   LEU A 179      51.380  27.276  28.868  1.00  0.00           N
ATOM   2498  H   LEU A 179      51.030  28.187  28.592  1.00  0.00           H
ATOM   2499  CA  LEU A 179      50.391  26.266  29.260  1.00  0.00           C
ATOM   2500  HA  LEU A 179      50.835  25.602  30.001  1.00  0.00           H
ATOM   2501  CB  LEU A 179      49.167  26.961  29.888  1.00  0.00           C
ATOM   2502  HB1 LEU A 179      48.651  27.523  29.108  1.00  0.00           H
ATOM   2503  HB2 LEU A 179      48.483  26.188  30.243  1.00  0.00           H
ATOM   2504  CG  LEU A 179      49.465  27.925  31.055  1.00  0.00           C
ATOM   2505  HG  LEU A 179      50.019  28.786  30.679  1.00  0.00           H
ATOM   2506  CD1 LEU A 179      48.131  28.428  31.622  1.00  0.00           C
ATOM   2507 HD11 LEU A 179      48.318  29.099  32.458  1.00  0.00           H
ATOM   2508 HD12 LEU A 179      47.578  28.967  30.851  1.00  0.00           H
ATOM   2509 HD13 LEU A 179      47.532  27.587  31.972  1.00  0.00           H
ATOM   2510  CD2 LEU A 179      50.293  27.274  32.170  1.00  0.00           C
ATOM   2511 HD21 LEU A 179      50.431  27.986  32.982  1.00  0.00           H
ATOM   2512 HD22 LEU A 179      49.778  26.395  32.549  1.00  0.00           H
ATOM   2513 HD23 LEU A 179      51.278  26.991  31.799  1.00  0.00           H
ATOM   2514  C   LEU A 179      49.955  25.393  28.076  1.00  0.00           C
ATOM   2515  O   LEU A 179      50.038  25.814  26.921  1.00  0.00           O
ATOM   2516  N   LEU A 180      49.433  24.200  28.371  1.00  0.00           N
ATOM   2517  H   LEU A 180      49.419  23.905  29.340  1.00  0.00           H
ATOM   2518  CA  LEU A 180      48.880  23.279  27.374  1.00  0.00           C
ATOM   2519  HA  LEU A 180      48.770  23.800  26.423  1.00  0.00           H
ATOM   2520  CB  LEU A 180      49.879  22.118  27.201  1.00  0.00           C
ATOM   2521  HB1 LEU A 180      50.859  22.526  26.952  1.00  0.00           H
ATOM   2522  HB2 LEU A 180      49.970  21.599  28.158  1.00  0.00           H
ATOM   2523  CG  LEU A 180      49.480  21.096  26.120  1.00  0.00           C
ATOM   2524  HG  LEU A 180      48.473  20.733  26.317  1.00  0.00           H
ATOM   2525  CD1 LEU A 180      49.497  21.697  24.708  1.00  0.00           C
ATOM   2526 HD11 LEU A 180      49.233  20.927  23.985  1.00  0.00           H
ATOM   2527 HD12 LEU A 180      48.761  22.495  24.627  1.00  0.00           H
ATOM   2528 HD13 LEU A 180      50.487  22.089  24.474  1.00  0.00           H
ATOM   2529  CD2 LEU A 180      50.442  19.903  26.199  1.00  0.00           C
ATOM   2530 HD21 LEU A 180      50.192  19.176  25.427  1.00  0.00           H
ATOM   2531 HD22 LEU A 180      51.468  20.242  26.065  1.00  0.00           H
ATOM   2532 HD23 LEU A 180      50.359  19.423  27.174  1.00  0.00           H
ATOM   2533  C   LEU A 180      47.495  22.773  27.798  1.00  0.00           C
ATOM   2534  O   LEU A 180      47.361  22.144  28.846  1.00  0.00           O
ATOM   2535  N   SER A 181      46.476  22.989  26.974  1.00  0.00           N
```

| | | | | | | | | | | | |
|---|---|---|---|---|---|---|---|---|---|---|---|
| ATOM | 2536 | H | SER | A | 181 | 46.628 | 23.537 | 26.127 | 1.00 | 0.00 | H |
| ATOM | 2537 | CA | SER | A | 181 | 45.123 | 22.456 | 27.176 | 1.00 | 0.00 | C |
| ATOM | 2538 | HA | SER | A | 181 | 45.150 | 21.718 | 27.976 | 1.00 | 0.00 | H |
| ATOM | 2539 | CB | SER | A | 181 | 44.172 | 23.582 | 27.630 | 1.00 | 0.00 | C |
| ATOM | 2540 | HB1 | SER | A | 181 | 43.195 | 23.161 | 27.872 | 1.00 | 0.00 | H |
| ATOM | 2541 | HB2 | SER | A | 181 | 44.576 | 24.028 | 28.540 | 1.00 | 0.00 | H |
| ATOM | 2542 | OG | SER | A | 181 | 44.000 | 24.604 | 26.665 | 1.00 | 0.00 | O |
| ATOM | 2543 | HG | SER | A | 181 | 43.393 | 24.283 | 25.958 | 1.00 | 0.00 | H |
| ATOM | 2544 | C | SER | A | 181 | 44.596 | 21.707 | 25.938 | 1.00 | 0.00 | C |
| ATOM | 2545 | O | SER | A | 181 | 45.268 | 21.614 | 24.907 | 1.00 | 0.00 | O |
| ATOM | 2546 | N | ALA | A | 182 | 43.384 | 21.148 | 26.023 | 1.00 | 0.00 | N |
| ATOM | 2547 | H | ALA | A | 182 | 42.855 | 21.244 | 26.878 | 1.00 | 0.00 | H |
| ATOM | 2548 | CA | ALA | A | 182 | 42.728 | 20.494 | 24.893 | 1.00 | 0.00 | C |
| ATOM | 2549 | HA | ALA | A | 182 | 43.035 | 21.000 | 23.977 | 1.00 | 0.00 | H |
| ATOM | 2550 | CB | ALA | A | 182 | 43.191 | 19.030 | 24.827 | 1.00 | 0.00 | C |
| ATOM | 2551 | HB1 | ALA | A | 182 | 42.733 | 18.538 | 23.968 | 1.00 | 0.00 | H |
| ATOM | 2552 | HB2 | ALA | A | 182 | 44.277 | 18.989 | 24.727 | 1.00 | 0.00 | H |
| ATOM | 2553 | HB3 | ALA | A | 182 | 42.897 | 18.507 | 25.738 | 1.00 | 0.00 | H |
| ATOM | 2554 | C | ALA | A | 182 | 41.200 | 20.593 | 24.972 | 1.00 | 0.00 | C |
| ATOM | 2555 | O | ALA | A | 182 | 40.604 | 20.368 | 26.027 | 1.00 | 0.00 | O |
| ATOM | 2556 | N | ALA | A | 183 | 40.565 | 20.843 | 23.825 | 1.00 | 0.00 | N |
| ATOM | 2557 | H | ALA | A | 183 | 41.127 | 21.071 | 23.014 | 1.00 | 0.00 | H |
| ATOM | 2558 | CA | ALA | A | 183 | 39.128 | 20.694 | 23.629 | 1.00 | 0.00 | C |
| ATOM | 2559 | HA | ALA | A | 183 | 38.601 | 21.074 | 24.503 | 1.00 | 0.00 | H |
| ATOM | 2560 | CB | ALA | A | 183 | 38.725 | 21.551 | 22.423 | 1.00 | 0.00 | C |
| ATOM | 2561 | HB1 | ALA | A | 183 | 37.658 | 21.439 | 22.231 | 1.00 | 0.00 | H |
| ATOM | 2562 | HB2 | ALA | A | 183 | 38.944 | 22.600 | 22.626 | 1.00 | 0.00 | H |
| ATOM | 2563 | HB3 | ALA | A | 183 | 39.279 | 21.245 | 21.536 | 1.00 | 0.00 | H |
| ATOM | 2564 | C | ALA | A | 183 | 38.785 | 19.205 | 23.441 | 1.00 | 0.00 | C |
| ATOM | 2565 | O | ALA | A | 183 | 39.015 | 18.650 | 22.363 | 1.00 | 0.00 | O |
| ATOM | 2566 | N | VAL | A | 184 | 38.251 | 18.548 | 24.473 | 1.00 | 0.00 | N |
| ATOM | 2567 | H | VAL | A | 184 | 38.071 | 19.090 | 25.320 | 1.00 | 0.00 | H |
| ATOM | 2568 | CA | VAL | A | 184 | 38.041 | 17.081 | 24.505 | 1.00 | 0.00 | C |
| ATOM | 2569 | HA | VAL | A | 184 | 38.571 | 16.664 | 23.650 | 1.00 | 0.00 | H |
| ATOM | 2570 | CB | VAL | A | 184 | 38.668 | 16.427 | 25.756 | 1.00 | 0.00 | C |
| ATOM | 2571 | HB | VAL | A | 184 | 38.530 | 15.357 | 25.639 | 1.00 | 0.00 | H |
| ATOM | 2572 | CG1 | VAL | A | 184 | 40.183 | 16.686 | 25.816 | 1.00 | 0.00 | C |
| ATOM | 2573 | HG11 | VAL | A | 184 | 40.629 | 16.100 | 26.620 | 1.00 | 0.00 | H |
| ATOM | 2574 | HG12 | VAL | A | 184 | 40.648 | 16.397 | 24.874 | 1.00 | 0.00 | H |
| ATOM | 2575 | HG13 | VAL | A | 184 | 40.382 | 17.740 | 26.002 | 1.00 | 0.00 | H |
| ATOM | 2576 | CG2 | VAL | A | 184 | 38.001 | 16.853 | 27.072 | 1.00 | 0.00 | C |
| ATOM | 2577 | HG21 | VAL | A | 184 | 38.458 | 16.317 | 27.902 | 1.00 | 0.00 | H |
| ATOM | 2578 | HG22 | VAL | A | 184 | 38.121 | 17.923 | 27.230 | 1.00 | 0.00 | H |
| ATOM | 2579 | HG23 | VAL | A | 184 | 36.942 | 16.602 | 27.060 | 1.00 | 0.00 | H |
| ATOM | 2580 | C | VAL | A | 184 | 36.556 | 16.674 | 24.383 | 1.00 | 0.00 | C |
| ATOM | 2581 | O | VAL | A | 184 | 35.694 | 17.429 | 24.842 | 1.00 | 0.00 | O |
| ATOM | 2582 | N | PRO | A | 185 | 36.236 | 15.519 | 23.754 | 1.00 | 0.00 | N |
| ATOM | 2583 | CD | PRO | A | 185 | 37.170 | 14.609 | 23.101 | 1.00 | 0.00 | C |
| ATOM | 2584 | HD1 | PRO | A | 185 | 37.937 | 14.257 | 23.791 | 1.00 | 0.00 | H |
| ATOM | 2585 | HD2 | PRO | A | 185 | 37.632 | 15.116 | 22.254 | 1.00 | 0.00 | H |
| ATOM | 2586 | CG | PRO | A | 185 | 36.344 | 13.426 | 22.597 | 1.00 | 0.00 | C |
| ATOM | 2587 | HG1 | PRO | A | 185 | 36.296 | 12.658 | 23.369 | 1.00 | 0.00 | H |
| ATOM | 2588 | HG2 | PRO | A | 185 | 36.751 | 13.015 | 21.674 | 1.00 | 0.00 | H |
| ATOM | 2589 | CB | PRO | A | 185 | 34.961 | 14.034 | 22.382 | 1.00 | 0.00 | C |

```
ATOM   2590  HB1 PRO A 185      34.174  13.281  22.443  1.00  0.00           H
ATOM   2591  HB2 PRO A 185      34.923  14.537  21.418  1.00  0.00           H
ATOM   2592  CA  PRO A 185      34.865  15.072  23.503  1.00  0.00           C
ATOM   2593  HA  PRO A 185      34.257  15.909  23.162  1.00  0.00           H
ATOM   2594  C   PRO A 185      34.199  14.463  24.739  1.00  0.00           C
ATOM   2595  O   PRO A 185      34.863  13.988  25.655  1.00  0.00           O
ATOM   2596  N   ALA A 186      32.865  14.460  24.734  1.00  0.00           N
ATOM   2597  H   ALA A 186      32.394  14.904  23.949  1.00  0.00           H
ATOM   2598  CA  ALA A 186      32.027  14.043  25.856  1.00  0.00           C
ATOM   2599  HA  ALA A 186      32.615  14.069  26.775  1.00  0.00           H
ATOM   2600  CB  ALA A 186      30.921  15.100  25.957  1.00  0.00           C
ATOM   2601  HB1 ALA A 186      30.277  14.875  26.798  1.00  0.00           H
ATOM   2602  HB2 ALA A 186      31.363  16.085  26.105  1.00  0.00           H
ATOM   2603  HB3 ALA A 186      30.318  15.106  25.047  1.00  0.00           H
ATOM   2604  C   ALA A 186      31.445  12.615  25.743  1.00  0.00           C
ATOM   2605  O   ALA A 186      30.946  12.088  26.742  1.00  0.00           O
ATOM   2606  N   GLY A 187      31.489  11.989  24.560  1.00  0.00           N
ATOM   2607  H   GLY A 187      31.882  12.490  23.772  1.00  0.00           H
ATOM   2608  CA  GLY A 187      30.899  10.673  24.272  1.00  0.00           C
ATOM   2609  HA1 GLY A 187      30.131  10.436  25.008  1.00  0.00           H
ATOM   2610  HA2 GLY A 187      30.430  10.703  23.290  1.00  0.00           H
ATOM   2611  C   GLY A 187      31.916   9.531  24.228  1.00  0.00           C
ATOM   2612  O   GLY A 187      33.034   9.678  23.722  1.00  0.00           O
ATOM   2613  N   GLN A 188      31.512   8.367  24.741  1.00  0.00           N
ATOM   2614  H   GLN A 188      30.568   8.316  25.110  1.00  0.00           H
ATOM   2615  CA  GLN A 188      32.357   7.176  24.855  1.00  0.00           C
ATOM   2616  HA  GLN A 188      33.212   7.437  25.482  1.00  0.00           H
ATOM   2617  CB  GLN A 188      31.605   6.054  25.591  1.00  0.00           C
ATOM   2618  HB1 GLN A 188      32.312   5.250  25.802  1.00  0.00           H
ATOM   2619  HB2 GLN A 188      31.263   6.441  26.551  1.00  0.00           H
ATOM   2620  CG  GLN A 188      30.405   5.449  24.837  1.00  0.00           C
ATOM   2621  HG1 GLN A 188      29.742   6.244  24.498  1.00  0.00           H
ATOM   2622  HG2 GLN A 188      30.754   4.895  23.966  1.00  0.00           H
ATOM   2623  CD  GLN A 188      29.633   4.500  25.750  1.00  0.00           C
ATOM   2624  OE1 GLN A 188      30.063   3.393  26.056  1.00  0.00           O
ATOM   2625  NE2 GLN A 188      28.503   4.902  26.284  1.00  0.00           N
ATOM   2626 HE21 GLN A 188      28.081   5.775  25.992  1.00  0.00           H
ATOM   2627 HE22 GLN A 188      28.116   4.340  27.032  1.00  0.00           H
ATOM   2628  C   GLN A 188      32.948   6.681  23.526  1.00  0.00           C
ATOM   2629  O   GLN A 188      34.031   6.099  23.567  1.00  0.00           O
ATOM   2630  N   THR A 189      32.318   6.917  22.360  1.00  0.00           N
ATOM   2631  H   THR A 189      31.415   7.389  22.366  1.00  0.00           H
ATOM   2632  CA  THR A 189      32.896   6.535  21.057  1.00  0.00           C
ATOM   2633  HA  THR A 189      33.151   5.480  21.082  1.00  0.00           H
ATOM   2634  CB  THR A 189      31.911   6.732  19.901  1.00  0.00           C
ATOM   2635  HB  THR A 189      31.672   7.789  19.799  1.00  0.00           H
ATOM   2636  CG2 THR A 189      32.477   6.217  18.577  1.00  0.00           C
ATOM   2637 HG21 THR A 189      31.722   6.294  17.795  1.00  0.00           H
ATOM   2638 HG22 THR A 189      33.337   6.817  18.282  1.00  0.00           H
ATOM   2639 HG23 THR A 189      32.781   5.174  18.676  1.00  0.00           H
ATOM   2640  OG1 THR A 189      30.721   6.021  20.147  1.00  0.00           O
ATOM   2641  HG1 THR A 189      30.942   5.158  20.549  1.00  0.00           H
ATOM   2642  C   THR A 189      34.200   7.285  20.820  1.00  0.00           C
ATOM   2643  O   THR A 189      35.256   6.655  20.766  1.00  0.00           O
```

| | | | | | | | | | | | |
|---|---|---|---|---|---|---|---|---|---|---|---|
| ATOM | 2644 | N | TYR | A | 190 | 34.157 | 8.620 | 20.731 | 1.00 | 0.00 | N |
| ATOM | 2645 | H | TYR | A | 190 | 33.255 | 9.086 | 20.777 | 1.00 | 0.00 | H |
| ATOM | 2646 | CA | TYR | A | 190 | 35.363 | 9.419 | 20.478 | 1.00 | 0.00 | C |
| ATOM | 2647 | HA | TYR | A | 190 | 35.869 | 8.997 | 19.610 | 1.00 | 0.00 | H |
| ATOM | 2648 | CB | TYR | A | 190 | 35.008 | 10.885 | 20.164 | 1.00 | 0.00 | C |
| ATOM | 2649 | HB1 | TYR | A | 190 | 34.509 | 11.299 | 21.041 | 1.00 | 0.00 | H |
| ATOM | 2650 | HB2 | TYR | A | 190 | 35.942 | 11.433 | 20.032 | 1.00 | 0.00 | H |
| ATOM | 2651 | CG | TYR | A | 190 | 34.151 | 11.181 | 18.939 | 1.00 | 0.00 | C |
| ATOM | 2652 | CD1 | TYR | A | 190 | 34.177 | 10.364 | 17.789 | 1.00 | 0.00 | C |
| ATOM | 2653 | HD1 | TYR | A | 190 | 34.811 | 9.492 | 17.754 | 1.00 | 0.00 | H |
| ATOM | 2654 | CE1 | TYR | A | 190 | 33.368 | 10.668 | 16.675 | 1.00 | 0.00 | C |
| ATOM | 2655 | HE1 | TYR | A | 190 | 33.400 | 10.033 | 15.801 | 1.00 | 0.00 | H |
| ATOM | 2656 | CZ | TYR | A | 190 | 32.509 | 11.790 | 16.706 | 1.00 | 0.00 | C |
| ATOM | 2657 | OH | TYR | A | 190 | 31.706 | 12.088 | 15.644 | 1.00 | 0.00 | O |
| ATOM | 2658 | HH | TYR | A | 190 | 31.796 | 11.442 | 14.921 | 1.00 | 0.00 | H |
| ATOM | 2659 | CE2 | TYR | A | 190 | 32.483 | 12.609 | 17.855 | 1.00 | 0.00 | C |
| ATOM | 2660 | HE2 | TYR | A | 190 | 31.841 | 13.478 | 17.880 | 1.00 | 0.00 | H |
| ATOM | 2661 | CD2 | TYR | A | 190 | 33.309 | 12.312 | 18.955 | 1.00 | 0.00 | C |
| ATOM | 2662 | HD2 | TYR | A | 190 | 33.285 | 12.960 | 19.822 | 1.00 | 0.00 | H |
| ATOM | 2663 | C | TYR | A | 190 | 36.369 | 9.359 | 21.643 | 1.00 | 0.00 | C |
| ATOM | 2664 | O | TYR | A | 190 | 37.576 | 9.376 | 21.385 | 1.00 | 0.00 | O |
| ATOM | 2665 | N | VAL | A | 191 | 35.899 | 9.262 | 22.895 | 1.00 | 0.00 | N |
| ATOM | 2666 | H | VAL | A | 191 | 34.886 | 9.272 | 23.020 | 1.00 | 0.00 | H |
| ATOM | 2667 | CA | VAL | A | 191 | 36.770 | 9.173 | 24.086 | 1.00 | 0.00 | C |
| ATOM | 2668 | HA | VAL | A | 191 | 37.489 | 9.993 | 24.024 | 1.00 | 0.00 | H |
| ATOM | 2669 | CB | VAL | A | 191 | 35.981 | 9.349 | 25.409 | 1.00 | 0.00 | C |
| ATOM | 2670 | HB | VAL | A | 191 | 35.136 | 8.660 | 25.401 | 1.00 | 0.00 | H |
| ATOM | 2671 | CG1 | VAL | A | 191 | 36.842 | 9.061 | 26.657 | 1.00 | 0.00 | C |
| ATOM | 2672 | HG11 | VAL | A | 191 | 36.295 | 9.326 | 27.562 | 1.00 | 0.00 | H |
| ATOM | 2673 | HG12 | VAL | A | 191 | 37.091 | 8.001 | 26.712 | 1.00 | 0.00 | H |
| ATOM | 2674 | HG13 | VAL | A | 191 | 37.761 | 9.647 | 26.620 | 1.00 | 0.00 | H |
| ATOM | 2675 | CG2 | VAL | A | 191 | 35.448 | 10.787 | 25.532 | 1.00 | 0.00 | C |
| ATOM | 2676 | HG21 | VAL | A | 191 | 34.799 | 10.870 | 26.405 | 1.00 | 0.00 | H |
| ATOM | 2677 | HG22 | VAL | A | 191 | 36.280 | 11.479 | 25.638 | 1.00 | 0.00 | H |
| ATOM | 2678 | HG23 | VAL | A | 191 | 34.881 | 11.073 | 24.650 | 1.00 | 0.00 | H |
| ATOM | 2679 | C | VAL | A | 191 | 37.565 | 7.860 | 24.099 | 1.00 | 0.00 | C |
| ATOM | 2680 | O | VAL | A | 191 | 38.795 | 7.901 | 24.160 | 1.00 | 0.00 | O |
| ATOM | 2681 | N | ASP | A | 192 | 36.905 | 6.696 | 24.006 | 1.00 | 0.00 | N |
| ATOM | 2682 | H | ASP | A | 192 | 35.890 | 6.687 | 23.910 | 1.00 | 0.00 | H |
| ATOM | 2683 | CA | ASP | A | 192 | 37.594 | 5.397 | 24.058 | 1.00 | 0.00 | C |
| ATOM | 2684 | HA | ASP | A | 192 | 38.319 | 5.426 | 24.867 | 1.00 | 0.00 | H |
| ATOM | 2685 | CB | ASP | A | 192 | 36.615 | 4.260 | 24.407 | 1.00 | 0.00 | C |
| ATOM | 2686 | HB1 | ASP | A | 192 | 35.788 | 4.267 | 23.702 | 1.00 | 0.00 | H |
| ATOM | 2687 | HB2 | ASP | A | 192 | 37.134 | 3.309 | 24.286 | 1.00 | 0.00 | H |
| ATOM | 2688 | CG | ASP | A | 192 | 36.099 | 4.318 | 25.853 | 1.00 | 0.00 | C |
| ATOM | 2689 | OD1 | ASP | A | 192 | 36.920 | 4.520 | 26.782 | 1.00 | 0.00 | O |
| ATOM | 2690 | OD2 | ASP | A | 192 | 34.887 | 4.104 | 26.092 | 1.00 | 0.00 | O |
| ATOM | 2691 | C | ASP | A | 192 | 38.393 | 5.094 | 22.775 | 1.00 | 0.00 | C |
| ATOM | 2692 | O | ASP | A | 192 | 39.276 | 4.236 | 22.798 | 1.00 | 0.00 | O |
| ATOM | 2693 | N | ALA | A | 193 | 38.135 | 5.799 | 21.665 | 1.00 | 0.00 | N |
| ATOM | 2694 | H | ALA | A | 193 | 37.344 | 6.433 | 21.680 | 1.00 | 0.00 | H |
| ATOM | 2695 | CA | ALA | A | 193 | 38.911 | 5.682 | 20.423 | 1.00 | 0.00 | C |
| ATOM | 2696 | HA | ALA | A | 193 | 39.291 | 4.663 | 20.330 | 1.00 | 0.00 | H |
| ATOM | 2697 | CB | ALA | A | 193 | 37.972 | 5.937 | 19.237 | 1.00 | 0.00 | C |

| | | | | | | | | | | | |
|---|---|---|---|---|---|---|---|---|---|---|---|
| ATOM | 2698 | HB1 | ALA | A | 193 | 38.526 | 5.854 | 18.301 | 1.00 | 0.00 | H |
| ATOM | 2699 | HB2 | ALA | A | 193 | 37.175 | 5.195 | 19.227 | 1.00 | 0.00 | H |
| ATOM | 2700 | HB3 | ALA | A | 193 | 37.535 | 6.933 | 19.302 | 1.00 | 0.00 | H |
| ATOM | 2701 | C | ALA | A | 193 | 40.135 | 6.615 | 20.343 | 1.00 | 0.00 | C |
| ATOM | 2702 | O | ALA | A | 193 | 41.057 | 6.310 | 19.586 | 1.00 | 0.00 | O |
| ATOM | 2703 | N | GLY | A | 194 | 40.164 | 7.738 | 21.077 | 1.00 | 0.00 | N |
| ATOM | 2704 | H | GLY | A | 194 | 39.372 | 7.932 | 21.676 | 1.00 | 0.00 | H |
| ATOM | 2705 | CA | GLY | A | 194 | 41.125 | 8.824 | 20.812 | 1.00 | 0.00 | C |
| ATOM | 2706 | HA1 | GLY | A | 194 | 41.765 | 8.576 | 19.965 | 1.00 | 0.00 | H |
| ATOM | 2707 | HA2 | GLY | A | 194 | 40.544 | 9.706 | 20.549 | 1.00 | 0.00 | H |
| ATOM | 2708 | C | GLY | A | 194 | 42.033 | 9.268 | 21.954 | 1.00 | 0.00 | C |
| ATOM | 2709 | O | GLY | A | 194 | 43.179 | 9.630 | 21.698 | 1.00 | 0.00 | O |
| ATOM | 2710 | N | TYR | A | 195 | 41.540 | 9.284 | 23.194 | 1.00 | 0.00 | N |
| ATOM | 2711 | H | TYR | A | 195 | 40.607 | 8.927 | 23.354 | 1.00 | 0.00 | H |
| ATOM | 2712 | CA | TYR | A | 195 | 42.173 | 10.041 | 24.285 | 1.00 | 0.00 | C |
| ATOM | 2713 | HA | TYR | A | 195 | 43.067 | 10.549 | 23.921 | 1.00 | 0.00 | H |
| ATOM | 2714 | CB | TYR | A | 195 | 41.199 | 11.134 | 24.758 | 1.00 | 0.00 | C |
| ATOM | 2715 | HB1 | TYR | A | 195 | 40.251 | 10.674 | 25.046 | 1.00 | 0.00 | H |
| ATOM | 2716 | HB2 | TYR | A | 195 | 41.617 | 11.598 | 25.652 | 1.00 | 0.00 | H |
| ATOM | 2717 | CG | TYR | A | 195 | 40.928 | 12.223 | 23.728 | 1.00 | 0.00 | C |
| ATOM | 2718 | CD1 | TYR | A | 195 | 39.984 | 12.017 | 22.700 | 1.00 | 0.00 | C |
| ATOM | 2719 | HD1 | TYR | A | 195 | 39.437 | 11.086 | 22.650 | 1.00 | 0.00 | H |
| ATOM | 2720 | CE1 | TYR | A | 195 | 39.757 | 13.014 | 21.732 | 1.00 | 0.00 | C |
| ATOM | 2721 | HE1 | TYR | A | 195 | 39.046 | 12.843 | 20.938 | 1.00 | 0.00 | H |
| ATOM | 2722 | CZ | TYR | A | 195 | 40.445 | 14.246 | 21.809 | 1.00 | 0.00 | C |
| ATOM | 2723 | OH | TYR | A | 195 | 40.190 | 15.228 | 20.899 | 1.00 | 0.00 | O |
| ATOM | 2724 | HH | TYR | A | 195 | 40.719 | 16.021 | 21.054 | 1.00 | 0.00 | H |
| ATOM | 2725 | CE2 | TYR | A | 195 | 41.383 | 14.457 | 22.843 | 1.00 | 0.00 | C |
| ATOM | 2726 | HE2 | TYR | A | 195 | 41.913 | 15.396 | 22.912 | 1.00 | 0.00 | H |
| ATOM | 2727 | CD2 | TYR | A | 195 | 41.627 | 13.445 | 23.791 | 1.00 | 0.00 | C |
| ATOM | 2728 | HD2 | TYR | A | 195 | 42.352 | 13.610 | 24.576 | 1.00 | 0.00 | H |
| ATOM | 2729 | C | TYR | A | 195 | 42.599 | 9.133 | 25.449 | 1.00 | 0.00 | C |
| ATOM | 2730 | O | TYR | A | 195 | 41.759 | 8.521 | 26.107 | 1.00 | 0.00 | O |
| ATOM | 2731 | N | GLU | A | 196 | 43.900 | 9.043 | 25.730 | 1.00 | 0.00 | N |
| ATOM | 2732 | H | GLU | A | 196 | 44.559 | 9.554 | 25.148 | 1.00 | 0.00 | H |
| ATOM | 2733 | CA | GLU | A | 196 | 44.425 | 8.304 | 26.883 | 1.00 | 0.00 | C |
| ATOM | 2734 | HA | GLU | A | 196 | 43.783 | 7.445 | 27.063 | 1.00 | 0.00 | H |
| ATOM | 2735 | CB | GLU | A | 196 | 45.821 | 7.741 | 26.606 | 1.00 | 0.00 | C |
| ATOM | 2736 | HB1 | GLU | A | 196 | 46.518 | 8.558 | 26.435 | 1.00 | 0.00 | H |
| ATOM | 2737 | HB2 | GLU | A | 196 | 46.146 | 7.192 | 27.488 | 1.00 | 0.00 | H |
| ATOM | 2738 | CG | GLU | A | 196 | 45.816 | 6.786 | 25.404 | 1.00 | 0.00 | C |
| ATOM | 2739 | HG1 | GLU | A | 196 | 44.988 | 6.088 | 25.495 | 1.00 | 0.00 | H |
| ATOM | 2740 | HG2 | GLU | A | 196 | 45.688 | 7.354 | 24.481 | 1.00 | 0.00 | H |
| ATOM | 2741 | CD | GLU | A | 196 | 47.116 | 6.003 | 25.337 | 1.00 | 0.00 | C |
| ATOM | 2742 | OE1 | GLU | A | 196 | 48.128 | 6.593 | 24.907 | 1.00 | 0.00 | O |
| ATOM | 2743 | OE2 | GLU | A | 196 | 47.151 | 4.833 | 25.787 | 1.00 | 0.00 | O |
| ATOM | 2744 | C | GLU | A | 196 | 44.378 | 9.202 | 28.126 | 1.00 | 0.00 | C |
| ATOM | 2745 | O | GLU | A | 196 | 45.350 | 9.876 | 28.470 | 1.00 | 0.00 | O |
| ATOM | 2746 | N | VAL | A | 197 | 43.198 | 9.244 | 28.746 | 1.00 | 0.00 | N |
| ATOM | 2747 | H | VAL | A | 197 | 42.517 | 8.540 | 28.459 | 1.00 | 0.00 | H |
| ATOM | 2748 | CA | VAL | A | 197 | 42.741 | 10.279 | 29.684 | 1.00 | 0.00 | C |
| ATOM | 2749 | HA | VAL | A | 197 | 42.724 | 11.225 | 29.141 | 1.00 | 0.00 | H |
| ATOM | 2750 | CB | VAL | A | 197 | 41.298 | 9.982 | 30.164 | 1.00 | 0.00 | C |
| ATOM | 2751 | HB | VAL | A | 197 | 41.296 | 9.014 | 30.661 | 1.00 | 0.00 | H |

```
ATOM   2752  CG1 VAL A 197      40.784  11.050  31.144  1.00  0.00           C
ATOM   2753 HG11 VAL A 197      39.750  10.845  31.422  1.00  0.00           H
ATOM   2754 HG12 VAL A 197      41.383  11.059  32.054  1.00  0.00           H
ATOM   2755 HG13 VAL A 197      40.840  12.029  30.673  1.00  0.00           H
ATOM   2756  CG2 VAL A 197      40.310   9.891  28.986  1.00  0.00           C
ATOM   2757 HG21 VAL A 197      39.292   9.773  29.358  1.00  0.00           H
ATOM   2758 HG22 VAL A 197      40.366  10.793  28.375  1.00  0.00           H
ATOM   2759 HG23 VAL A 197      40.541   9.024  28.370  1.00  0.00           H
ATOM   2760  C   VAL A 197      43.683  10.430  30.884  1.00  0.00           C
ATOM   2761  O   VAL A 197      44.067  11.551  31.211  1.00  0.00           O
ATOM   2762  N   ASP A 198      44.086   9.323  31.513  1.00  0.00           N
ATOM   2763  H   ASP A 198      43.745   8.433  31.178  1.00  0.00           H
ATOM   2764  CA  ASP A 198      44.968   9.316  32.688  1.00  0.00           C
ATOM   2765  HA  ASP A 198      44.660  10.110  33.372  1.00  0.00           H
ATOM   2766  CB  ASP A 198      44.791   7.977  33.410  1.00  0.00           C
ATOM   2767  HB1 ASP A 198      43.742   7.846  33.665  1.00  0.00           H
ATOM   2768  HB2 ASP A 198      45.079   7.169  32.736  1.00  0.00           H
ATOM   2769  CG  ASP A 198      45.604   7.875  34.699  1.00  0.00           C
ATOM   2770  OD1 ASP A 198      45.162   8.409  35.741  1.00  0.00           O
ATOM   2771  OD2 ASP A 198      46.629   7.154  34.685  1.00  0.00           O
ATOM   2772  C   ASP A 198      46.444   9.548  32.322  1.00  0.00           C
ATOM   2773  O   ASP A 198      47.206  10.071  33.133  1.00  0.00           O
ATOM   2774  N   LYS A 199      46.863   9.212  31.094  1.00  0.00           N
ATOM   2775  H   LYS A 199      46.181   8.847  30.437  1.00  0.00           H
ATOM   2776  CA  LYS A 199      48.203   9.563  30.610  1.00  0.00           C
ATOM   2777  HA  LYS A 199      48.926   9.348  31.394  1.00  0.00           H
ATOM   2778  CB  LYS A 199      48.580   8.738  29.369  1.00  0.00           C
ATOM   2779  HB1 LYS A 199      47.824   8.885  28.598  1.00  0.00           H
ATOM   2780  HB2 LYS A 199      49.527   9.130  28.995  1.00  0.00           H
ATOM   2781  CG  LYS A 199      48.766   7.230  29.621  1.00  0.00           C
ATOM   2782  HG1 LYS A 199      49.518   7.077  30.395  1.00  0.00           H
ATOM   2783  HG2 LYS A 199      47.819   6.789  29.936  1.00  0.00           H
ATOM   2784  CD  LYS A 199      49.238   6.561  28.324  1.00  0.00           C
ATOM   2785  HD1 LYS A 199      48.510   6.787  27.548  1.00  0.00           H
ATOM   2786  HD2 LYS A 199      50.201   6.989  28.037  1.00  0.00           H
ATOM   2787  CE  LYS A 199      49.401   5.042  28.409  1.00  0.00           C
ATOM   2788  HE1 LYS A 199      50.243   4.814  29.069  1.00  0.00           H
ATOM   2789  HE2 LYS A 199      48.496   4.603  28.839  1.00  0.00           H
ATOM   2790  NZ  LYS A 199      49.643   4.485  27.056  1.00  0.00           N
ATOM   2791  HZ1 LYS A 199      48.807   4.571  26.477  1.00  0.00           H
ATOM   2792  HZ2 LYS A 199      50.376   5.010  26.584  1.00  0.00           H
ATOM   2793  HZ3 LYS A 199      49.933   3.517  27.078  1.00  0.00           H
ATOM   2794  C   LYS A 199      48.328  11.071  30.332  1.00  0.00           C
ATOM   2795  O   LYS A 199      49.291  11.693  30.791  1.00  0.00           O
ATOM   2796  N   ILE A 200      47.360  11.676  29.628  1.00  0.00           N
ATOM   2797  H   ILE A 200      46.601  11.095  29.268  1.00  0.00           H
ATOM   2798  CA  ILE A 200      47.384  13.122  29.310  1.00  0.00           C
ATOM   2799  HA  ILE A 200      48.405  13.384  29.019  1.00  0.00           H
ATOM   2800  CB  ILE A 200      46.475  13.471  28.104  1.00  0.00           C
ATOM   2801  HB  ILE A 200      46.618  14.531  27.892  1.00  0.00           H
ATOM   2802  CG2 ILE A 200      46.955  12.703  26.857  1.00  0.00           C
ATOM   2803 HG21 ILE A 200      46.479  13.099  25.961  1.00  0.00           H
ATOM   2804 HG22 ILE A 200      48.032  12.824  26.747  1.00  0.00           H
ATOM   2805 HG23 ILE A 200      46.722  11.642  26.944  1.00  0.00           H
```

```
ATOM   2806  CG1 ILE A 200      44.968  13.282  28.395  1.00  0.00           C
ATOM   2807 HG11 ILE A 200      44.794  12.280  28.771  1.00  0.00           H
ATOM   2808 HG12 ILE A 200      44.674  13.985  29.174  1.00  0.00           H
ATOM   2809  CD  ILE A 200      44.043  13.505  27.191  1.00  0.00           C
ATOM   2810  HD1 ILE A 200      43.006  13.464  27.525  1.00  0.00           H
ATOM   2811  HD2 ILE A 200      44.236  14.481  26.746  1.00  0.00           H
ATOM   2812  HD3 ILE A 200      44.196  12.724  26.447  1.00  0.00           H
ATOM   2813  C   ILE A 200      47.091  14.004  30.534  1.00  0.00           C
ATOM   2814  O   ILE A 200      47.651  15.093  30.634  1.00  0.00           O
ATOM   2815  N   ALA A 201      46.301  13.510  31.496  1.00  0.00           N
ATOM   2816  H   ALA A 201      45.848  12.622  31.319  1.00  0.00           H
ATOM   2817  CA  ALA A 201      45.932  14.192  32.745  1.00  0.00           C
ATOM   2818  HA  ALA A 201      45.246  15.012  32.524  1.00  0.00           H
ATOM   2819  CB  ALA A 201      45.224  13.163  33.636  1.00  0.00           C
ATOM   2820  HB1 ALA A 201      45.185  13.528  34.658  1.00  0.00           H
ATOM   2821  HB2 ALA A 201      44.208  12.985  33.290  1.00  0.00           H
ATOM   2822  HB3 ALA A 201      45.774  12.223  33.652  1.00  0.00           H
ATOM   2823  C   ALA A 201      47.117  14.776  33.526  1.00  0.00           C
ATOM   2824  O   ALA A 201      46.995  15.828  34.158  1.00  0.00           O
ATOM   2825  N   GLN A 202      48.258  14.088  33.505  1.00  0.00           N
ATOM   2826  H   GLN A 202      48.285  13.237  32.964  1.00  0.00           H
ATOM   2827  CA  GLN A 202      49.447  14.496  34.250  1.00  0.00           C
ATOM   2828  HA  GLN A 202      49.152  14.794  35.258  1.00  0.00           H
ATOM   2829  CB  GLN A 202      50.431  13.315  34.358  1.00  0.00           C
ATOM   2830  HB1 GLN A 202      50.974  13.221  33.416  1.00  0.00           H
ATOM   2831  HB2 GLN A 202      51.162  13.550  35.133  1.00  0.00           H
ATOM   2832  CG  GLN A 202      49.802  11.943  34.666  1.00  0.00           C
ATOM   2833  HG1 GLN A 202      49.283  11.592  33.775  1.00  0.00           H
ATOM   2834  HG2 GLN A 202      50.606  11.237  34.859  1.00  0.00           H
ATOM   2835  CD  GLN A 202      48.836  11.897  35.855  1.00  0.00           C
ATOM   2836  OE1 GLN A 202      48.944  12.641  36.828  1.00  0.00           O
ATOM   2837  NE2 GLN A 202      47.847  11.035  35.799  1.00  0.00           N
ATOM   2838 HE21 GLN A 202      47.719  10.452  34.979  1.00  0.00           H
ATOM   2839 HE22 GLN A 202      47.304  10.842  36.631  1.00  0.00           H
ATOM   2840  C   GLN A 202      50.136  15.701  33.592  1.00  0.00           C
ATOM   2841  O   GLN A 202      50.669  16.553  34.300  1.00  0.00           O
ATOM   2842  N   ASN A 203      50.098  15.770  32.255  1.00  0.00           N
ATOM   2843  H   ASN A 203      49.519  15.096  31.770  1.00  0.00           H
ATOM   2844  CA  ASN A 203      50.911  16.657  31.415  1.00  0.00           C
ATOM   2845  HA  ASN A 203      51.768  17.019  31.987  1.00  0.00           H
ATOM   2846  CB  ASN A 203      51.420  15.819  30.228  1.00  0.00           C
ATOM   2847  HB1 ASN A 203      50.556  15.389  29.723  1.00  0.00           H
ATOM   2848  HB2 ASN A 203      51.936  16.465  29.520  1.00  0.00           H
ATOM   2849  CG  ASN A 203      52.373  14.687  30.611  1.00  0.00           C
ATOM   2850  OD1 ASN A 203      53.171  14.779  31.535  1.00  0.00           O
ATOM   2851  ND2 ASN A 203      52.334  13.578  29.896  1.00  0.00           N
ATOM   2852 HD21 ASN A 203      51.666  13.466  29.140  1.00  0.00           H
ATOM   2853 HD22 ASN A 203      52.945  12.818  30.155  1.00  0.00           H
ATOM   2854  C   ASN A 203      50.144  17.889  30.883  1.00  0.00           C
ATOM   2855  O   ASN A 203      50.759  18.913  30.566  1.00  0.00           O
ATOM   2856  N   LEU A 204      48.811  17.812  30.765  1.00  0.00           N
ATOM   2857  H   LEU A 204      48.362  16.929  30.987  1.00  0.00           H
ATOM   2858  CA  LEU A 204      47.955  18.969  30.472  1.00  0.00           C
ATOM   2859  HA  LEU A 204      48.406  19.558  29.672  1.00  0.00           H
```

| | | | | | | | | | | | |
|---|---|---|---|---|---|---|---|---|---|---|---|
| ATOM | 2860 | CB | LEU | A | 204 | 46.552 | 18.499 | 30.029 | 1.00 | 0.00 | C |
| ATOM | 2861 | HB1 | LEU | A | 204 | 46.143 | 17.852 | 30.808 | 1.00 | 0.00 | H |
| ATOM | 2862 | HB2 | LEU | A | 204 | 45.900 | 19.372 | 29.952 | 1.00 | 0.00 | H |
| ATOM | 2863 | CG | LEU | A | 204 | 46.511 | 17.746 | 28.683 | 1.00 | 0.00 | C |
| ATOM | 2864 | HG | LEU | A | 204 | 47.169 | 16.881 | 28.739 | 1.00 | 0.00 | H |
| ATOM | 2865 | CD1 | LEU | A | 204 | 45.091 | 17.242 | 28.398 | 1.00 | 0.00 | C |
| ATOM | 2866 | HD11 | LEU | A | 204 | 45.085 | 16.662 | 27.475 | 1.00 | 0.00 | H |
| ATOM | 2867 | HD12 | LEU | A | 204 | 44.752 | 16.604 | 29.215 | 1.00 | 0.00 | H |
| ATOM | 2868 | HD13 | LEU | A | 204 | 44.410 | 18.087 | 28.287 | 1.00 | 0.00 | H |
| ATOM | 2869 | CD2 | LEU | A | 204 | 46.963 | 18.627 | 27.512 | 1.00 | 0.00 | C |
| ATOM | 2870 | HD21 | LEU | A | 204 | 46.839 | 18.091 | 26.571 | 1.00 | 0.00 | H |
| ATOM | 2871 | HD22 | LEU | A | 204 | 46.373 | 19.544 | 27.478 | 1.00 | 0.00 | H |
| ATOM | 2872 | HD23 | LEU | A | 204 | 48.015 | 18.876 | 27.626 | 1.00 | 0.00 | H |
| ATOM | 2873 | C | LEU | A | 204 | 47.863 | 19.890 | 31.695 | 1.00 | 0.00 | C |
| ATOM | 2874 | O | LEU | A | 204 | 47.898 | 19.431 | 32.835 | 1.00 | 0.00 | O |
| ATOM | 2875 | N | ASP | A | 205 | 47.712 | 21.191 | 31.465 | 1.00 | 0.00 | N |
| ATOM | 2876 | H | ASP | A | 205 | 47.684 | 21.505 | 30.498 | 1.00 | 0.00 | H |
| ATOM | 2877 | CA | ASP | A | 205 | 47.476 | 22.190 | 32.510 | 1.00 | 0.00 | C |
| ATOM | 2878 | HA | ASP | A | 205 | 48.012 | 21.899 | 33.414 | 1.00 | 0.00 | H |
| ATOM | 2879 | CB | ASP | A | 205 | 48.023 | 23.543 | 32.035 | 1.00 | 0.00 | C |
| ATOM | 2880 | HB1 | ASP | A | 205 | 47.807 | 23.702 | 30.978 | 1.00 | 0.00 | H |
| ATOM | 2881 | HB2 | ASP | A | 205 | 47.539 | 24.342 | 32.599 | 1.00 | 0.00 | H |
| ATOM | 2882 | CG | ASP | A | 205 | 49.522 | 23.620 | 32.300 | 1.00 | 0.00 | C |
| ATOM | 2883 | OD1 | ASP | A | 205 | 49.868 | 23.829 | 33.487 | 1.00 | 0.00 | O |
| ATOM | 2884 | OD2 | ASP | A | 205 | 50.340 | 23.474 | 31.361 | 1.00 | 0.00 | O |
| ATOM | 2885 | C | ASP | A | 205 | 45.992 | 22.290 | 32.886 | 1.00 | 0.00 | C |
| ATOM | 2886 | O | ASP | A | 205 | 45.664 | 22.401 | 34.070 | 1.00 | 0.00 | O |
| ATOM | 2887 | N | PHE | A | 206 | 45.101 | 22.179 | 31.897 | 1.00 | 0.00 | N |
| ATOM | 2888 | H | PHE | A | 206 | 45.453 | 22.080 | 30.953 | 1.00 | 0.00 | H |
| ATOM | 2889 | CA | PHE | A | 206 | 43.653 | 22.090 | 32.075 | 1.00 | 0.00 | C |
| ATOM | 2890 | HA | PHE | A | 206 | 43.481 | 21.403 | 32.902 | 1.00 | 0.00 | H |
| ATOM | 2891 | CB | PHE | A | 206 | 43.052 | 23.448 | 32.490 | 1.00 | 0.00 | C |
| ATOM | 2892 | HB1 | PHE | A | 206 | 41.993 | 23.309 | 32.703 | 1.00 | 0.00 | H |
| ATOM | 2893 | HB2 | PHE | A | 206 | 43.508 | 23.759 | 33.431 | 1.00 | 0.00 | H |
| ATOM | 2894 | CG | PHE | A | 206 | 43.195 | 24.587 | 31.498 | 1.00 | 0.00 | C |
| ATOM | 2895 | CD1 | PHE | A | 206 | 44.355 | 25.386 | 31.501 | 1.00 | 0.00 | C |
| ATOM | 2896 | HD1 | PHE | A | 206 | 45.168 | 25.161 | 32.178 | 1.00 | 0.00 | H |
| ATOM | 2897 | CE1 | PHE | A | 206 | 44.457 | 26.489 | 30.635 | 1.00 | 0.00 | C |
| ATOM | 2898 | HE1 | PHE | A | 206 | 45.350 | 27.097 | 30.639 | 1.00 | 0.00 | H |
| ATOM | 2899 | CZ | PHE | A | 206 | 43.396 | 26.801 | 29.767 | 1.00 | 0.00 | C |
| ATOM | 2900 | HZ | PHE | A | 206 | 43.473 | 27.646 | 29.098 | 1.00 | 0.00 | H |
| ATOM | 2901 | CE2 | PHE | A | 206 | 42.238 | 26.006 | 29.760 | 1.00 | 0.00 | C |
| ATOM | 2902 | HE2 | PHE | A | 206 | 41.422 | 26.245 | 29.094 | 1.00 | 0.00 | H |
| ATOM | 2903 | CD2 | PHE | A | 206 | 42.140 | 24.896 | 30.617 | 1.00 | 0.00 | C |
| ATOM | 2904 | HD2 | PHE | A | 206 | 41.246 | 24.288 | 30.598 | 1.00 | 0.00 | H |
| ATOM | 2905 | C | PHE | A | 206 | 42.977 | 21.468 | 30.844 | 1.00 | 0.00 | C |
| ATOM | 2906 | O | PHE | A | 206 | 43.559 | 21.398 | 29.764 | 1.00 | 0.00 | O |
| ATOM | 2907 | N | VAL | A | 207 | 41.756 | 20.971 | 31.031 | 1.00 | 0.00 | N |
| ATOM | 2908 | H | VAL | A | 207 | 41.353 | 21.084 | 31.958 | 1.00 | 0.00 | H |
| ATOM | 2909 | CA | VAL | A | 207 | 41.026 | 20.124 | 30.078 | 1.00 | 0.00 | C |
| ATOM | 2910 | HA | VAL | A | 207 | 41.572 | 20.075 | 29.134 | 1.00 | 0.00 | H |
| ATOM | 2911 | CB | VAL | A | 207 | 40.880 | 18.686 | 30.637 | 1.00 | 0.00 | C |
| ATOM | 2912 | HB | VAL | A | 207 | 40.336 | 18.730 | 31.584 | 1.00 | 0.00 | H |
| ATOM | 2913 | CG1 | VAL | A | 207 | 40.097 | 17.781 | 29.677 | 1.00 | 0.00 | C |

| | | | | | | | | | | | |
|---|---|---|---|---|---|---|---|---|---|---|---|
| ATOM | 2914 | HG11 | VAL | A | 207 | 40.045 | 16.769 | 30.081 | 1.00 | 0.00 | H |
| ATOM | 2915 | HG12 | VAL | A | 207 | 39.079 | 18.150 | 29.558 | 1.00 | 0.00 | H |
| ATOM | 2916 | HG13 | VAL | A | 207 | 40.590 | 17.755 | 28.704 | 1.00 | 0.00 | H |
| ATOM | 2917 | CG2 | VAL | A | 207 | 42.255 | 18.058 | 30.918 | 1.00 | 0.00 | C |
| ATOM | 2918 | HG21 | VAL | A | 207 | 42.139 | 17.016 | 31.212 | 1.00 | 0.00 | H |
| ATOM | 2919 | HG22 | VAL | A | 207 | 42.871 | 18.106 | 30.022 | 1.00 | 0.00 | H |
| ATOM | 2920 | HG23 | VAL | A | 207 | 42.758 | 18.587 | 31.729 | 1.00 | 0.00 | H |
| ATOM | 2921 | C | VAL | A | 207 | 39.653 | 20.743 | 29.822 | 1.00 | 0.00 | C |
| ATOM | 2922 | O | VAL | A | 207 | 38.856 | 20.873 | 30.750 | 1.00 | 0.00 | O |
| ATOM | 2923 | N | ASN | A | 208 | 39.387 | 21.128 | 28.572 | 1.00 | 0.00 | N |
| ATOM | 2924 | H | ASN | A | 208 | 40.072 | 20.931 | 27.848 | 1.00 | 0.00 | H |
| ATOM | 2925 | CA | ASN | A | 208 | 38.164 | 21.816 | 28.156 | 1.00 | 0.00 | C |
| ATOM | 2926 | HA | ASN | A | 208 | 37.711 | 22.309 | 29.018 | 1.00 | 0.00 | H |
| ATOM | 2927 | CB | ASN | A | 208 | 38.531 | 22.907 | 27.131 | 1.00 | 0.00 | C |
| ATOM | 2928 | HB1 | ASN | A | 208 | 39.033 | 22.459 | 26.275 | 1.00 | 0.00 | H |
| ATOM | 2929 | HB2 | ASN | A | 208 | 37.626 | 23.387 | 26.768 | 1.00 | 0.00 | H |
| ATOM | 2930 | CG | ASN | A | 208 | 39.429 | 23.986 | 27.731 | 1.00 | 0.00 | C |
| ATOM | 2931 | OD1 | ASN | A | 208 | 39.128 | 24.541 | 28.782 | 1.00 | 0.00 | O |
| ATOM | 2932 | ND2 | ASN | A | 208 | 40.552 | 24.299 | 27.107 | 1.00 | 0.00 | N |
| ATOM | 2933 | HD21 | ASN | A | 208 | 40.803 | 23.892 | 26.209 | 1.00 | 0.00 | H |
| ATOM | 2934 | HD22 | ASN | A | 208 | 41.150 | 24.999 | 27.509 | 1.00 | 0.00 | H |
| ATOM | 2935 | C | ASN | A | 208 | 37.150 | 20.796 | 27.605 | 1.00 | 0.00 | C |
| ATOM | 2936 | O | ASN | A | 208 | 37.276 | 20.321 | 26.470 | 1.00 | 0.00 | O |
| ATOM | 2937 | N | LEU | A | 209 | 36.152 | 20.424 | 28.414 | 1.00 | 0.00 | N |
| ATOM | 2938 | H | LEU | A | 209 | 36.080 | 20.853 | 29.335 | 1.00 | 0.00 | H |
| ATOM | 2939 | CA | LEU | A | 209 | 35.172 | 19.388 | 28.069 | 1.00 | 0.00 | C |
| ATOM | 2940 | HA | LEU | A | 209 | 35.678 | 18.609 | 27.498 | 1.00 | 0.00 | H |
| ATOM | 2941 | CB | LEU | A | 209 | 34.648 | 18.761 | 29.376 | 1.00 | 0.00 | C |
| ATOM | 2942 | HB1 | LEU | A | 209 | 35.500 | 18.387 | 29.945 | 1.00 | 0.00 | H |
| ATOM | 2943 | HB2 | LEU | A | 209 | 34.173 | 19.546 | 29.969 | 1.00 | 0.00 | H |
| ATOM | 2944 | CG | LEU | A | 209 | 33.637 | 17.610 | 29.193 | 1.00 | 0.00 | C |
| ATOM | 2945 | HG | LEU | A | 209 | 32.778 | 17.976 | 28.632 | 1.00 | 0.00 | H |
| ATOM | 2946 | CD1 | LEU | A | 209 | 34.234 | 16.420 | 28.427 | 1.00 | 0.00 | C |
| ATOM | 2947 | HD11 | LEU | A | 209 | 33.525 | 15.593 | 28.417 | 1.00 | 0.00 | H |
| ATOM | 2948 | HD12 | LEU | A | 209 | 34.443 | 16.701 | 27.395 | 1.00 | 0.00 | H |
| ATOM | 2949 | HD13 | LEU | A | 209 | 35.157 | 16.092 | 28.907 | 1.00 | 0.00 | H |
| ATOM | 2950 | CD2 | LEU | A | 209 | 33.140 | 17.148 | 30.571 | 1.00 | 0.00 | C |
| ATOM | 2951 | HD21 | LEU | A | 209 | 32.378 | 16.379 | 30.451 | 1.00 | 0.00 | H |
| ATOM | 2952 | HD22 | LEU | A | 209 | 33.966 | 16.739 | 31.154 | 1.00 | 0.00 | H |
| ATOM | 2953 | HD23 | LEU | A | 209 | 32.701 | 17.988 | 31.109 | 1.00 | 0.00 | H |
| ATOM | 2954 | C | LEU | A | 209 | 34.049 | 19.967 | 27.199 | 1.00 | 0.00 | C |
| ATOM | 2955 | O | LEU | A | 209 | 33.259 | 20.785 | 27.668 | 1.00 | 0.00 | O |
| ATOM | 2956 | N | MET | A | 210 | 33.946 | 19.509 | 25.947 | 1.00 | 0.00 | N |
| ATOM | 2957 | H | MET | A | 210 | 34.623 | 18.821 | 25.639 | 1.00 | 0.00 | H |
| ATOM | 2958 | CA | MET | A | 210 | 32.905 | 19.899 | 24.980 | 1.00 | 0.00 | C |
| ATOM | 2959 | HA | MET | A | 210 | 32.746 | 20.978 | 25.038 | 1.00 | 0.00 | H |
| ATOM | 2960 | CB | MET | A | 210 | 33.361 | 19.562 | 23.547 | 1.00 | 0.00 | C |
| ATOM | 2961 | HB1 | MET | A | 210 | 33.519 | 18.486 | 23.471 | 1.00 | 0.00 | H |
| ATOM | 2962 | HB2 | MET | A | 210 | 32.563 | 19.838 | 22.854 | 1.00 | 0.00 | H |
| ATOM | 2963 | CG | MET | A | 210 | 34.640 | 20.292 | 23.117 | 1.00 | 0.00 | C |
| ATOM | 2964 | HG1 | MET | A | 210 | 34.465 | 21.365 | 23.216 | 1.00 | 0.00 | H |
| ATOM | 2965 | HG2 | MET | A | 210 | 35.460 | 20.025 | 23.784 | 1.00 | 0.00 | H |
| ATOM | 2966 | SD | MET | A | 210 | 35.144 | 19.967 | 21.404 | 1.00 | 0.00 | S |
| ATOM | 2967 | CE | MET | A | 210 | 35.696 | 18.249 | 21.513 | 1.00 | 0.00 | C |

| | | | | | | | | | | | |
|---|---|---|---|---|---|---|---|---|---|---|---|
| ATOM | 2968 | HE1 | MET | A | 210 | 36.582 | 18.198 | 22.139 | 1.00 | 0.00 | H |
| ATOM | 2969 | HE2 | MET | A | 210 | 34.919 | 17.627 | 21.954 | 1.00 | 0.00 | H |
| ATOM | 2970 | HE3 | MET | A | 210 | 35.930 | 17.882 | 20.515 | 1.00 | 0.00 | H |
| ATOM | 2971 | C | MET | A | 210 | 31.562 | 19.228 | 25.313 | 1.00 | 0.00 | C |
| ATOM | 2972 | O | MET | A | 210 | 31.085 | 18.357 | 24.582 | 1.00 | 0.00 | O |
| ATOM | 2973 | N | ALA | A | 211 | 30.974 | 19.574 | 26.461 | 1.00 | 0.00 | N |
| ATOM | 2974 | H | ALA | A | 211 | 31.444 | 20.277 | 27.021 | 1.00 | 0.00 | H |
| ATOM | 2975 | CA | ALA | A | 211 | 29.739 | 18.988 | 26.986 | 1.00 | 0.00 | C |
| ATOM | 2976 | HA | ALA | A | 211 | 29.761 | 17.913 | 26.787 | 1.00 | 0.00 | H |
| ATOM | 2977 | CB | ALA | A | 211 | 29.726 | 19.185 | 28.507 | 1.00 | 0.00 | C |
| ATOM | 2978 | HB1 | ALA | A | 211 | 28.924 | 18.589 | 28.939 | 1.00 | 0.00 | H |
| ATOM | 2979 | HB2 | ALA | A | 211 | 30.666 | 18.852 | 28.944 | 1.00 | 0.00 | H |
| ATOM | 2980 | HB3 | ALA | A | 211 | 29.570 | 20.238 | 28.749 | 1.00 | 0.00 | H |
| ATOM | 2981 | C | ALA | A | 211 | 28.489 | 19.536 | 26.265 | 1.00 | 0.00 | C |
| ATOM | 2982 | O | ALA | A | 211 | 27.574 | 20.074 | 26.870 | 1.00 | 0.00 | O |
| ATOM | 2983 | N | TYR | A | 212 | 28.458 | 19.394 | 24.879 | 1.00 | 0.00 | N |
| ATOM | 2984 | H | TYR | A | 212 | 29.263 | 18.945 | 24.434 | 1.00 | 0.00 | H |
| ATOM | 2985 | CA | TYR | A | 212 | 27.504 | 20.039 | 24.015 | 1.00 | 0.00 | C |
| ATOM | 2986 | HA | TYR | A | 212 | 26.479 | 19.871 | 24.388 | 1.00 | 0.00 | H |
| ATOM | 2987 | CB | TYR | A | 212 | 27.685 | 21.585 | 23.863 | 1.00 | 0.00 | C |
| ATOM | 2988 | HB1 | TYR | A | 212 | 27.063 | 21.929 | 23.020 | 1.00 | 0.00 | H |
| ATOM | 2989 | HB2 | TYR | A | 212 | 27.260 | 22.046 | 24.768 | 1.00 | 0.00 | H |
| ATOM | 2990 | CG | TYR | A | 212 | 29.103 | 22.079 | 23.706 | 1.00 | 0.00 | C |
| ATOM | 2991 | CD1 | TYR | A | 212 | 29.828 | 21.917 | 22.514 | 1.00 | 0.00 | C |
| ATOM | 2992 | HD1 | TYR | A | 212 | 29.379 | 21.383 | 21.674 | 1.00 | 0.00 | H |
| ATOM | 2993 | CE1 | TYR | A | 212 | 31.103 | 22.461 | 22.353 | 1.00 | 0.00 | C |
| ATOM | 2994 | HE1 | TYR | A | 212 | 31.627 | 22.360 | 21.402 | 1.00 | 0.00 | H |
| ATOM | 2995 | CZ | TYR | A | 212 | 31.694 | 23.203 | 23.394 | 1.00 | 0.00 | C |
| ATOM | 2996 | OH | TYR | A | 212 | 32.910 | 23.799 | 23.293 | 1.00 | 0.00 | O |
| ATOM | 2997 | HH | TYR | A | 212 | 33.270 | 23.756 | 22.352 | 1.00 | 0.00 | H |
| ATOM | 2998 | CE2 | TYR | A | 212 | 30.994 | 23.341 | 24.605 | 1.00 | 0.00 | C |
| ATOM | 2999 | HE2 | TYR | A | 212 | 31.443 | 23.942 | 25.397 | 1.00 | 0.00 | H |
| ATOM | 3000 | CD2 | TYR | A | 212 | 29.725 | 22.781 | 24.750 | 1.00 | 0.00 | C |
| ATOM | 3001 | HD2 | TYR | A | 212 | 29.190 | 22.917 | 25.694 | 1.00 | 0.00 | H |
| ATOM | 3002 | C | TYR | A | 212 | 27.576 | 19.276 | 22.681 | 1.00 | 0.00 | C |
| ATOM | 3003 | O | TYR | A | 212 | 28.484 | 18.460 | 22.451 | 1.00 | 0.00 | O |
| ATOM | 3004 | N | ASP | A | 213 | 26.574 | 19.513 | 21.808 | 1.00 | 0.00 | N |
| ATOM | 3005 | H | ASP | A | 213 | 25.873 | 20.240 | 21.967 | 1.00 | 0.00 | H |
| ATOM | 3006 | CA | ASP | A | 213 | 26.435 | 18.756 | 20.563 | 1.00 | 0.00 | C |
| ATOM | 3007 | HA | ASP | A | 213 | 25.519 | 19.156 | 20.094 | 1.00 | 0.00 | H |
| ATOM | 3008 | CB | ASP | A | 213 | 27.602 | 18.959 | 19.595 | 1.00 | 0.00 | C |
| ATOM | 3009 | HB1 | ASP | A | 213 | 28.514 | 18.508 | 20.001 | 1.00 | 0.00 | H |
| ATOM | 3010 | HB2 | ASP | A | 213 | 27.389 | 18.417 | 18.662 | 1.00 | 0.00 | H |
| ATOM | 3011 | CG | ASP | A | 213 | 27.872 | 20.422 | 19.259 | 1.00 | 0.00 | C |
| ATOM | 3012 | OD1 | ASP | A | 213 | 27.014 | 21.290 | 19.566 | 1.00 | 0.00 | O |
| ATOM | 3013 | OD2 | ASP | A | 213 | 28.982 | 20.646 | 18.652 | 1.00 | 0.00 | O |
| ATOM | 3014 | C | ASP | A | 213 | 26.138 | 17.275 | 20.822 | 1.00 | 0.00 | C |
| ATOM | 3015 | O | ASP | A | 213 | 26.454 | 16.428 | 19.989 | 1.00 | 0.00 | O |
| ATOM | 3016 | N | PHE | A | 214 | 25.475 | 16.915 | 21.978 | 1.00 | 0.00 | N |
| ATOM | 3017 | H | PHE | A | 214 | 25.183 | 17.624 | 22.670 | 1.00 | 0.00 | H |
| ATOM | 3018 | CA | PHE | A | 214 | 25.177 | 15.515 | 22.324 | 1.00 | 0.00 | C |
| ATOM | 3019 | HA | PHE | A | 214 | 26.089 | 14.920 | 22.252 | 1.00 | 0.00 | H |
| ATOM | 3020 | CB | PHE | A | 214 | 24.672 | 15.455 | 23.769 | 1.00 | 0.00 | C |
| ATOM | 3021 | HB1 | PHE | A | 214 | 23.908 | 16.219 | 23.924 | 1.00 | 0.00 | H |

| | | | | | | | | | | |
|---|---|---|---|---|---|---|---|---|---|---|
| ATOM | 3022 | HB2 | PHE | A | 214 | 24.175 | 14.498 | 23.916 | 1.00 | 0.00 | H |
| ATOM | 3023 | CG | PHE | A | 214 | 25.740 | 15.568 | 24.840 | 1.00 | 0.00 | C |
| ATOM | 3024 | CD1 | PHE | A | 214 | 26.630 | 14.503 | 25.070 | 1.00 | 0.00 | C |
| ATOM | 3025 | HD1 | PHE | A | 214 | 26.603 | 13.628 | 24.436 | 1.00 | 0.00 | H |
| ATOM | 3026 | CE1 | PHE | A | 214 | 27.528 | 14.557 | 26.151 | 1.00 | 0.00 | C |
| ATOM | 3027 | HE1 | PHE | A | 214 | 28.188 | 13.723 | 26.342 | 1.00 | 0.00 | H |
| ATOM | 3028 | CZ | PHE | A | 214 | 27.545 | 15.682 | 26.997 | 1.00 | 0.00 | C |
| ATOM | 3029 | HZ | PHE | A | 214 | 28.217 | 15.718 | 27.841 | 1.00 | 0.00 | H |
| ATOM | 3030 | CE2 | PHE | A | 214 | 26.672 | 16.754 | 26.756 | 1.00 | 0.00 | C |
| ATOM | 3031 | HE2 | PHE | A | 214 | 26.675 | 17.612 | 27.412 | 1.00 | 0.00 | H |
| ATOM | 3032 | CD2 | PHE | A | 214 | 25.782 | 16.698 | 25.671 | 1.00 | 0.00 | C |
| ATOM | 3033 | HD2 | PHE | A | 214 | 25.111 | 17.513 | 25.481 | 1.00 | 0.00 | H |
| ATOM | 3034 | C | PHE | A | 214 | 24.138 | 14.868 | 21.392 | 1.00 | 0.00 | C |
| ATOM | 3035 | O | PHE | A | 214 | 24.329 | 13.741 | 20.935 | 1.00 | 0.00 | O |
| ATOM | 3036 | N | HIS | A | 215 | 23.075 | 15.599 | 21.075 | 1.00 | 0.00 | N |
| ATOM | 3037 | H | HIS | A | 215 | 22.990 | 16.511 | 21.495 | 1.00 | 0.00 | H |
| ATOM | 3038 | CA | HIS | A | 215 | 22.143 | 15.287 | 19.993 | 1.00 | 0.00 | C |
| ATOM | 3039 | HA | HIS | A | 215 | 22.346 | 14.291 | 19.595 | 1.00 | 0.00 | H |
| ATOM | 3040 | CB | HIS | A | 215 | 20.717 | 15.310 | 20.557 | 1.00 | 0.00 | C |
| ATOM | 3041 | HB1 | HIS | A | 215 | 20.547 | 16.242 | 21.089 | 1.00 | 0.00 | H |
| ATOM | 3042 | HB2 | HIS | A | 215 | 19.998 | 15.269 | 19.739 | 1.00 | 0.00 | H |
| ATOM | 3043 | CG | HIS | A | 215 | 20.463 | 14.137 | 21.469 | 1.00 | 0.00 | C |
| ATOM | 3044 | ND1 | HIS | A | 215 | 20.709 | 12.819 | 21.160 | 1.00 | 0.00 | N |
| ATOM | 3045 | HD1 | HIS | A | 215 | 21.008 | 12.473 | 20.250 | 1.00 | 0.00 | H |
| ATOM | 3046 | CE1 | HIS | A | 215 | 20.396 | 12.075 | 22.229 | 1.00 | 0.00 | C |
| ATOM | 3047 | HE1 | HIS | A | 215 | 20.463 | 10.999 | 22.275 | 1.00 | 0.00 | H |
| ATOM | 3048 | NE2 | HIS | A | 215 | 19.909 | 12.849 | 23.212 | 1.00 | 0.00 | N |
| ATOM | 3049 | CD2 | HIS | A | 215 | 19.987 | 14.167 | 22.752 | 1.00 | 0.00 | C |
| ATOM | 3050 | HD2 | HIS | A | 215 | 19.725 | 15.051 | 23.316 | 1.00 | 0.00 | H |
| ATOM | 3051 | C | HIS | A | 215 | 22.338 | 16.275 | 18.834 | 1.00 | 0.00 | C |
| ATOM | 3052 | O | HIS | A | 215 | 22.822 | 17.391 | 19.054 | 1.00 | 0.00 | O |
| ATOM | 3053 | N | GLY | A | 216 | 21.967 | 15.887 | 17.607 | 1.00 | 0.00 | N |
| ATOM | 3054 | H | GLY | A | 216 | 21.492 | 14.994 | 17.499 | 1.00 | 0.00 | H |
| ATOM | 3055 | CA | GLY | A | 216 | 22.257 | 16.676 | 16.401 | 1.00 | 0.00 | C |
| ATOM | 3056 | HA1 | GLY | A | 216 | 22.100 | 17.734 | 16.615 | 1.00 | 0.00 | H |
| ATOM | 3057 | HA2 | GLY | A | 216 | 23.311 | 16.562 | 16.161 | 1.00 | 0.00 | H |
| ATOM | 3058 | C | GLY | A | 216 | 21.449 | 16.306 | 15.160 | 1.00 | 0.00 | C |
| ATOM | 3059 | O | GLY | A | 216 | 20.718 | 15.321 | 15.143 | 1.00 | 0.00 | O |
| ATOM | 3060 | N | SER | A | 217 | 21.595 | 17.105 | 14.102 | 1.00 | 0.00 | N |
| ATOM | 3061 | H | SER | A | 217 | 22.239 | 17.886 | 14.196 | 1.00 | 0.00 | H |
| ATOM | 3062 | CA | SER | A | 217 | 20.831 | 17.004 | 12.842 | 1.00 | 0.00 | C |
| ATOM | 3063 | HA | SER | A | 217 | 19.772 | 17.134 | 13.067 | 1.00 | 0.00 | H |
| ATOM | 3064 | CB | SER | A | 217 | 21.254 | 18.126 | 11.879 | 1.00 | 0.00 | C |
| ATOM | 3065 | HB1 | SER | A | 217 | 20.528 | 18.174 | 11.068 | 1.00 | 0.00 | H |
| ATOM | 3066 | HB2 | SER | A | 217 | 21.241 | 19.079 | 12.410 | 1.00 | 0.00 | H |
| ATOM | 3067 | OG | SER | A | 217 | 22.548 | 17.935 | 11.324 | 1.00 | 0.00 | O |
| ATOM | 3068 | HG | SER | A | 217 | 22.429 | 17.513 | 10.443 | 1.00 | 0.00 | H |
| ATOM | 3069 | C | SER | A | 217 | 20.952 | 15.650 | 12.133 | 1.00 | 0.00 | C |
| ATOM | 3070 | O | SER | A | 217 | 20.090 | 15.277 | 11.340 | 1.00 | 0.00 | O |
| ATOM | 3071 | N | TRP | A | 218 | 21.996 | 14.890 | 12.460 | 1.00 | 0.00 | N |
| ATOM | 3072 | H | TRP | A | 218 | 22.651 | 15.275 | 13.124 | 1.00 | 0.00 | H |
| ATOM | 3073 | CA | TRP | A | 218 | 22.246 | 13.524 | 12.022 | 1.00 | 0.00 | C |
| ATOM | 3074 | HA | TRP | A | 218 | 22.090 | 13.473 | 10.942 | 1.00 | 0.00 | H |
| ATOM | 3075 | CB | TRP | A | 218 | 23.728 | 13.245 | 12.316 | 1.00 | 0.00 | C |

```
ATOM   3076  HB1 TRP A 218      23.957  12.211  12.062  1.00  0.00           H
ATOM   3077  HB2 TRP A 218      24.323  13.878  11.659  1.00  0.00           H
ATOM   3078  CG  TRP A 218      24.188  13.489  13.723  1.00  0.00           C
ATOM   3079  CD1 TRP A 218      24.062  12.613  14.740  1.00  0.00           C
ATOM   3080  HD1 TRP A 218      23.610  11.634  14.659  1.00  0.00           H
ATOM   3081  NE1 TRP A 218      24.595  13.148  15.894  1.00  0.00           N
ATOM   3082  HE1 TRP A 218      24.605  12.639  16.776  1.00  0.00           H
ATOM   3083  CE2 TRP A 218      25.125  14.402  15.672  1.00  0.00           C
ATOM   3084  CZ2 TRP A 218      25.786  15.328  16.494  1.00  0.00           C
ATOM   3085  HZ2 TRP A 218      25.956  15.107  17.538  1.00  0.00           H
ATOM   3086  CH2 TRP A 218      26.228  16.543  15.942  1.00  0.00           C
ATOM   3087  HH2 TRP A 218      26.748  17.260  16.563  1.00  0.00           H
ATOM   3088  CZ3 TRP A 218      25.995  16.820  14.584  1.00  0.00           C
ATOM   3089  HZ3 TRP A 218      26.343  17.753  14.158  1.00  0.00           H
ATOM   3090  CE3 TRP A 218      25.312  15.893  13.772  1.00  0.00           C
ATOM   3091  HE3 TRP A 218      25.138  16.124  12.730  1.00  0.00           H
ATOM   3092  CD2 TRP A 218      24.870  14.654  14.289  1.00  0.00           C
ATOM   3093  C   TRP A 218      21.313  12.467  12.656  1.00  0.00           C
ATOM   3094  O   TRP A 218      21.456  11.284  12.335  1.00  0.00           O
ATOM   3095  N   GLU A 219      20.393  12.854  13.551  1.00  0.00           N
ATOM   3096  H   GLU A 219      20.373  13.835  13.810  1.00  0.00           H
ATOM   3097  CA  GLU A 219      19.459  11.964  14.256  1.00  0.00           C
ATOM   3098  HA  GLU A 219      19.717  10.929  14.043  1.00  0.00           H
ATOM   3099  CB  GLU A 219      19.550  12.182  15.772  1.00  0.00           C
ATOM   3100  HB1 GLU A 219      19.192  13.186  16.005  1.00  0.00           H
ATOM   3101  HB2 GLU A 219      18.890  11.466  16.265  1.00  0.00           H
ATOM   3102  CG  GLU A 219      20.964  12.012  16.338  1.00  0.00           C
ATOM   3103  HG1 GLU A 219      21.340  11.018  16.092  1.00  0.00           H
ATOM   3104  HG2 GLU A 219      21.624  12.763  15.908  1.00  0.00           H
ATOM   3105  CD  GLU A 219      20.951  12.204  17.846  1.00  0.00           C
ATOM   3106  OE1 GLU A 219      20.445  13.240  18.316  1.00  0.00           O
ATOM   3107  OE2 GLU A 219      21.388  11.298  18.596  1.00  0.00           O
ATOM   3108  C   GLU A 219      17.999  12.197  13.837  1.00  0.00           C
ATOM   3109  O   GLU A 219      17.530  13.331  13.743  1.00  0.00           O
ATOM   3110  N   LYS A 220      17.222  11.127  13.651  1.00  0.00           N
ATOM   3111  H   LYS A 220      17.642  10.211  13.763  1.00  0.00           H
ATOM   3112  CA  LYS A 220      15.845  11.199  13.122  1.00  0.00           C
ATOM   3113  HA  LYS A 220      15.839  11.973  12.354  1.00  0.00           H
ATOM   3114  CB  LYS A 220      15.503   9.869  12.422  1.00  0.00           C
ATOM   3115  HB1 LYS A 220      14.519   9.957  11.959  1.00  0.00           H
ATOM   3116  HB2 LYS A 220      16.228   9.703  11.623  1.00  0.00           H
ATOM   3117  CG  LYS A 220      15.505   8.643  13.350  1.00  0.00           C
ATOM   3118  HG1 LYS A 220      16.518   8.480  13.720  1.00  0.00           H
ATOM   3119  HG2 LYS A 220      14.840   8.806  14.197  1.00  0.00           H
ATOM   3120  CD  LYS A 220      15.040   7.401  12.580  1.00  0.00           C
ATOM   3121  HD1 LYS A 220      13.964   7.455  12.404  1.00  0.00           H
ATOM   3122  HD2 LYS A 220      15.557   7.369  11.620  1.00  0.00           H
ATOM   3123  CE  LYS A 220      15.379   6.140  13.377  1.00  0.00           C
ATOM   3124  HE1 LYS A 220      16.423   6.207  13.686  1.00  0.00           H
ATOM   3125  HE2 LYS A 220      14.767   6.095  14.280  1.00  0.00           H
ATOM   3126  NZ  LYS A 220      15.200   4.908  12.580  1.00  0.00           N
ATOM   3127  HZ1 LYS A 220      15.625   4.130  13.073  1.00  0.00           H
ATOM   3128  HZ2 LYS A 220      14.216   4.706  12.426  1.00  0.00           H
ATOM   3129  HZ3 LYS A 220      15.670   5.005  11.684  1.00  0.00           H
```

| | | | | | | | | | | | |
|---|---|---|---|---|---|---|---|---|---|---|---|
| ATOM | 3130 | C | LYS | A | 220 | 14.754 | 11.636 | 14.128 | 1.00 | 0.00 | C |
| ATOM | 3131 | O | LYS | A | 220 | 13.562 | 11.478 | 13.846 | 1.00 | 0.00 | O |
| ATOM | 3132 | N | VAL | A | 221 | 15.134 | 12.156 | 15.296 | 1.00 | 0.00 | N |
| ATOM | 3133 | H | VAL | A | 221 | 16.114 | 12.396 | 15.383 | 1.00 | 0.00 | H |
| ATOM | 3134 | CA | VAL | A | 221 | 14.232 | 12.486 | 16.420 | 1.00 | 0.00 | C |
| ATOM | 3135 | HA | VAL | A | 221 | 13.284 | 12.796 | 15.986 | 1.00 | 0.00 | H |
| ATOM | 3136 | CB | VAL | A | 221 | 13.964 | 11.294 | 17.376 | 1.00 | 0.00 | C |
| ATOM | 3137 | HB | VAL | A | 221 | 13.438 | 11.678 | 18.250 | 1.00 | 0.00 | H |
| ATOM | 3138 | CG1 | VAL | A | 221 | 13.041 | 10.247 | 16.743 | 1.00 | 0.00 | C |
| ATOM | 3139 | HG11 | VAL | A | 221 | 12.784 | 9.495 | 17.488 | 1.00 | 0.00 | H |
| ATOM | 3140 | HG12 | VAL | A | 221 | 12.124 | 10.725 | 16.396 | 1.00 | 0.00 | H |
| ATOM | 3141 | HG13 | VAL | A | 221 | 13.535 | 9.754 | 15.907 | 1.00 | 0.00 | H |
| ATOM | 3142 | CG2 | VAL | A | 221 | 15.254 | 10.612 | 17.867 | 1.00 | 0.00 | C |
| ATOM | 3143 | HG21 | VAL | A | 221 | 15.005 | 9.820 | 18.574 | 1.00 | 0.00 | H |
| ATOM | 3144 | HG22 | VAL | A | 221 | 15.802 | 10.176 | 17.030 | 1.00 | 0.00 | H |
| ATOM | 3145 | HG23 | VAL | A | 221 | 15.892 | 11.335 | 18.375 | 1.00 | 0.00 | H |
| ATOM | 3146 | C | VAL | A | 221 | 14.752 | 13.654 | 17.259 | 1.00 | 0.00 | C |
| ATOM | 3147 | O | VAL | A | 221 | 15.966 | 13.870 | 17.330 | 1.00 | 0.00 | O |
| ATOM | 3148 | N | THR | A | 222 | 13.829 | 14.379 | 17.905 | 1.00 | 0.00 | N |
| ATOM | 3149 | H | THR | A | 222 | 12.849 | 14.142 | 17.777 | 1.00 | 0.00 | H |
| ATOM | 3150 | CA | THR | A | 222 | 14.137 | 15.515 | 18.783 | 1.00 | 0.00 | C |
| ATOM | 3151 | HA | THR | A | 222 | 14.860 | 16.140 | 18.261 | 1.00 | 0.00 | H |
| ATOM | 3152 | CB | THR | A | 222 | 12.904 | 16.391 | 19.042 | 1.00 | 0.00 | C |
| ATOM | 3153 | HB | THR | A | 222 | 13.179 | 17.179 | 19.742 | 1.00 | 0.00 | H |
| ATOM | 3154 | CG2 | THR | A | 222 | 12.374 | 17.035 | 17.760 | 1.00 | 0.00 | C |
| ATOM | 3155 | HG21 | THR | A | 222 | 11.509 | 17.652 | 17.995 | 1.00 | 0.00 | H |
| ATOM | 3156 | HG22 | THR | A | 222 | 13.150 | 17.662 | 17.318 | 1.00 | 0.00 | H |
| ATOM | 3157 | HG23 | THR | A | 222 | 12.076 | 16.270 | 17.042 | 1.00 | 0.00 | H |
| ATOM | 3158 | OG1 | THR | A | 222 | 11.875 | 15.615 | 19.597 | 1.00 | 0.00 | O |
| ATOM | 3159 | HG1 | THR | A | 222 | 11.179 | 16.208 | 19.934 | 1.00 | 0.00 | H |
| ATOM | 3160 | C | THR | A | 222 | 14.795 | 15.077 | 20.094 | 1.00 | 0.00 | C |
| ATOM | 3161 | O | THR | A | 222 | 14.447 | 14.055 | 20.689 | 1.00 | 0.00 | O |
| ATOM | 3162 | N | GLY | A | 223 | 15.765 | 15.879 | 20.528 | 1.00 | 0.00 | N |
| ATOM | 3163 | H | GLY | A | 223 | 15.937 | 16.705 | 19.970 | 1.00 | 0.00 | H |
| ATOM | 3164 | CA | GLY | A | 223 | 16.657 | 15.653 | 21.670 | 1.00 | 0.00 | C |
| ATOM | 3165 | HA1 | GLY | A | 223 | 16.062 | 15.514 | 22.572 | 1.00 | 0.00 | H |
| ATOM | 3166 | HA2 | GLY | A | 223 | 17.227 | 14.736 | 21.521 | 1.00 | 0.00 | H |
| ATOM | 3167 | C | GLY | A | 223 | 17.631 | 16.817 | 21.871 | 1.00 | 0.00 | C |
| ATOM | 3168 | O | GLY | A | 223 | 17.986 | 17.516 | 20.917 | 1.00 | 0.00 | O |
| ATOM | 3169 | N | HIS | A | 224 | 18.039 | 17.068 | 23.112 | 1.00 | 0.00 | N |
| ATOM | 3170 | H | HIS | A | 224 | 17.719 | 16.441 | 23.850 | 1.00 | 0.00 | H |
| ATOM | 3171 | CA | HIS | A | 224 | 18.705 | 18.312 | 23.520 | 1.00 | 0.00 | C |
| ATOM | 3172 | HA | HIS | A | 224 | 18.216 | 19.125 | 22.981 | 1.00 | 0.00 | H |
| ATOM | 3173 | CB | HIS | A | 224 | 18.407 | 18.527 | 25.008 | 1.00 | 0.00 | C |
| ATOM | 3174 | HB1 | HIS | A | 224 | 17.363 | 18.282 | 25.205 | 1.00 | 0.00 | H |
| ATOM | 3175 | HB2 | HIS | A | 224 | 19.025 | 17.858 | 25.605 | 1.00 | 0.00 | H |
| ATOM | 3176 | CG | HIS | A | 224 | 18.638 | 19.946 | 25.434 | 1.00 | 0.00 | C |
| ATOM | 3177 | ND1 | HIS | A | 224 | 19.875 | 20.497 | 25.765 | 1.00 | 0.00 | N |
| ATOM | 3178 | CE1 | HIS | A | 224 | 19.642 | 21.796 | 26.005 | 1.00 | 0.00 | C |
| ATOM | 3179 | HE1 | HIS | A | 224 | 20.389 | 22.523 | 26.302 | 1.00 | 0.00 | H |
| ATOM | 3180 | NE2 | HIS | A | 224 | 18.338 | 22.075 | 25.841 | 1.00 | 0.00 | N |
| ATOM | 3181 | HE2 | HIS | A | 224 | 17.921 | 22.994 | 25.990 | 1.00 | 0.00 | H |
| ATOM | 3182 | CD2 | HIS | A | 224 | 17.690 | 20.924 | 25.464 | 1.00 | 0.00 | C |
| ATOM | 3183 | HD2 | HIS | A | 224 | 16.638 | 20.815 | 25.243 | 1.00 | 0.00 | H |

```
ATOM   3184  C   HIS A 224      20.224  18.413  23.206  1.00  0.00           C
ATOM   3185  O   HIS A 224      20.970  17.436  23.251  1.00  0.00           O
ATOM   3186  N   ASN A 225      20.700  19.632  22.927  1.00  0.00           N
ATOM   3187  H   ASN A 225      20.035  20.389  22.868  1.00  0.00           H
ATOM   3188  CA  ASN A 225      22.101  19.948  22.619  1.00  0.00           C
ATOM   3189  HA  ASN A 225      22.431  19.317  21.791  1.00  0.00           H
ATOM   3190  CB  ASN A 225      22.148  21.417  22.160  1.00  0.00           C
ATOM   3191  HB1 ASN A 225      21.541  21.540  21.262  1.00  0.00           H
ATOM   3192  HB2 ASN A 225      21.719  22.048  22.940  1.00  0.00           H
ATOM   3193  CG  ASN A 225      23.563  21.897  21.851  1.00  0.00           C
ATOM   3194  OD1 ASN A 225      24.343  21.210  21.209  1.00  0.00           O
ATOM   3195  ND2 ASN A 225      23.945  23.067  22.321  1.00  0.00           N
ATOM   3196 HD21 ASN A 225      23.314  23.613  22.903  1.00  0.00           H
ATOM   3197 HD22 ASN A 225      24.853  23.422  22.073  1.00  0.00           H
ATOM   3198  C   ASN A 225      23.068  19.720  23.803  1.00  0.00           C
ATOM   3199  O   ASN A 225      24.165  19.197  23.620  1.00  0.00           O
ATOM   3200  N   SER A 226      22.680  20.099  25.021  1.00  0.00           N
ATOM   3201  H   SER A 226      21.742  20.476  25.123  1.00  0.00           H
ATOM   3202  CA  SER A 226      23.521  20.049  26.227  1.00  0.00           C
ATOM   3203  HA  SER A 226      24.234  19.238  26.132  1.00  0.00           H
ATOM   3204  CB  SER A 226      24.348  21.343  26.303  1.00  0.00           C
ATOM   3205  HB1 SER A 226      24.851  21.487  25.347  1.00  0.00           H
ATOM   3206  HB2 SER A 226      23.675  22.186  26.466  1.00  0.00           H
ATOM   3207  OG  SER A 226      25.328  21.312  27.330  1.00  0.00           O
ATOM   3208  HG  SER A 226      26.091  20.759  27.050  1.00  0.00           H
ATOM   3209  C   SER A 226      22.707  19.753  27.509  1.00  0.00           C
ATOM   3210  O   SER A 226      22.658  20.589  28.417  1.00  0.00           O
ATOM   3211  N   PRO A 227      22.050  18.578  27.599  1.00  0.00           N
ATOM   3212  CD  PRO A 227      21.988  17.542  26.572  1.00  0.00           C
ATOM   3213  HD1 PRO A 227      22.978  17.264  26.215  1.00  0.00           H
ATOM   3214  HD2 PRO A 227      21.373  17.891  25.746  1.00  0.00           H
ATOM   3215  CG  PRO A 227      21.317  16.335  27.213  1.00  0.00           C
ATOM   3216  HG1 PRO A 227      22.075  15.697  27.658  1.00  0.00           H
ATOM   3217  HG2 PRO A 227      20.715  15.774  26.495  1.00  0.00           H
ATOM   3218  CB  PRO A 227      20.463  16.961  28.310  1.00  0.00           C
ATOM   3219  HB1 PRO A 227      20.303  16.257  29.125  1.00  0.00           H
ATOM   3220  HB2 PRO A 227      19.513  17.290  27.888  1.00  0.00           H
ATOM   3221  CA  PRO A 227      21.268  18.178  28.767  1.00  0.00           C
ATOM   3222  HA  PRO A 227      20.586  18.979  29.054  1.00  0.00           H
ATOM   3223  C   PRO A 227      22.166  17.827  29.959  1.00  0.00           C
ATOM   3224  O   PRO A 227      23.273  17.314  29.776  1.00  0.00           O
ATOM   3225  N   LEU A 228      21.681  18.066  31.187  1.00  0.00           N
ATOM   3226  H   LEU A 228      20.765  18.492  31.293  1.00  0.00           H
ATOM   3227  CA  LEU A 228      22.443  17.749  32.400  1.00  0.00           C
ATOM   3228  HA  LEU A 228      23.463  18.068  32.225  1.00  0.00           H
ATOM   3229  CB  LEU A 228      21.906  18.560  33.596  1.00  0.00           C
ATOM   3230  HB1 LEU A 228      21.892  19.613  33.315  1.00  0.00           H
ATOM   3231  HB2 LEU A 228      20.881  18.251  33.803  1.00  0.00           H
ATOM   3232  CG  LEU A 228      22.734  18.412  34.894  1.00  0.00           C
ATOM   3233  HG  LEU A 228      22.664  17.382  35.242  1.00  0.00           H
ATOM   3234  CD1 LEU A 228      24.222  18.754  34.717  1.00  0.00           C
ATOM   3235 HD11 LEU A 228      24.715  18.780  35.689  1.00  0.00           H
ATOM   3236 HD12 LEU A 228      24.710  17.985  34.122  1.00  0.00           H
ATOM   3237 HD13 LEU A 228      24.333  19.725  34.233  1.00  0.00           H
```

```
ATOM   3238  CD2 LEU A 228      22.144  19.329  35.974  1.00  0.00           C
ATOM   3239 HD21 LEU A 228      22.680  19.190  36.914  1.00  0.00           H
ATOM   3240 HD22 LEU A 228      22.228  20.371  35.665  1.00  0.00           H
ATOM   3241 HD23 LEU A 228      21.096  19.085  36.132  1.00  0.00           H
ATOM   3242  C   LEU A 228      22.492  16.238  32.666  1.00  0.00           C
ATOM   3243  O   LEU A 228      23.578  15.679  32.768  1.00  0.00           O
ATOM   3244  N   TYR A 229      21.330  15.586  32.730  1.00  0.00           N
ATOM   3245  H   TYR A 229      20.499  16.126  32.555  1.00  0.00           H
ATOM   3246  CA  TYR A 229      21.159  14.143  32.978  1.00  0.00           C
ATOM   3247  HA  TYR A 229      22.130  13.672  33.144  1.00  0.00           H
ATOM   3248  CB  TYR A 229      20.304  13.976  34.249  1.00  0.00           C
ATOM   3249  HB1 TYR A 229      19.316  14.401  34.067  1.00  0.00           H
ATOM   3250  HB2 TYR A 229      20.168  12.914  34.455  1.00  0.00           H
ATOM   3251  CG  TYR A 229      20.908  14.610  35.486  1.00  0.00           C
ATOM   3252  CD1 TYR A 229      22.106  14.093  36.011  1.00  0.00           C
ATOM   3253  HD1 TYR A 229      22.585  13.248  35.533  1.00  0.00           H
ATOM   3254  CE1 TYR A 229      22.705  14.695  37.130  1.00  0.00           C
ATOM   3255  HE1 TYR A 229      23.645  14.317  37.504  1.00  0.00           H
ATOM   3256  CZ  TYR A 229      22.101  15.816  37.739  1.00  0.00           C
ATOM   3257  OH  TYR A 229      22.702  16.393  38.816  1.00  0.00           O
ATOM   3258  HH  TYR A 229      22.294  17.228  39.088  1.00  0.00           H
ATOM   3259  CE2 TYR A 229      20.886  16.325  37.227  1.00  0.00           C
ATOM   3260  HE2 TYR A 229      20.420  17.181  37.692  1.00  0.00           H
ATOM   3261  CD2 TYR A 229      20.295  15.723  36.098  1.00  0.00           C
ATOM   3262  HD2 TYR A 229      19.377  16.119  35.693  1.00  0.00           H
ATOM   3263  C   TYR A 229      20.494  13.426  31.786  1.00  0.00           C
ATOM   3264  O   TYR A 229      20.155  14.067  30.788  1.00  0.00           O
ATOM   3265  N   LYS A 230      20.232  12.115  31.887  1.00  0.00           N
ATOM   3266  H   LYS A 230      20.539  11.605  32.709  1.00  0.00           H
ATOM   3267  CA  LYS A 230      19.344  11.441  30.926  1.00  0.00           C
ATOM   3268  HA  LYS A 230      19.680  11.703  29.923  1.00  0.00           H
ATOM   3269  CB  LYS A 230      19.438   9.904  31.045  1.00  0.00           C
ATOM   3270  HB1 LYS A 230      19.062   9.498  30.105  1.00  0.00           H
ATOM   3271  HB2 LYS A 230      20.483   9.607  31.130  1.00  0.00           H
ATOM   3272  CG  LYS A 230      18.634   9.273  32.202  1.00  0.00           C
ATOM   3273  HG1 LYS A 230      19.170   9.405  33.142  1.00  0.00           H
ATOM   3274  HG2 LYS A 230      17.665   9.760  32.297  1.00  0.00           H
ATOM   3275  CD  LYS A 230      18.372   7.780  31.952  1.00  0.00           C
ATOM   3276  HD1 LYS A 230      18.057   7.632  30.918  1.00  0.00           H
ATOM   3277  HD2 LYS A 230      19.286   7.218  32.131  1.00  0.00           H
ATOM   3278  CE  LYS A 230      17.256   7.286  32.876  1.00  0.00           C
ATOM   3279  HE1 LYS A 230      17.597   7.348  33.913  1.00  0.00           H
ATOM   3280  HE2 LYS A 230      16.396   7.948  32.759  1.00  0.00           H
ATOM   3281  NZ  LYS A 230      16.842   5.899  32.564  1.00  0.00           N
ATOM   3282  HZ1 LYS A 230      17.579   5.237  32.787  1.00  0.00           H
ATOM   3283  HZ2 LYS A 230      16.027   5.650  33.115  1.00  0.00           H
ATOM   3284  HZ3 LYS A 230      16.611   5.794  31.579  1.00  0.00           H
ATOM   3285  C   LYS A 230      17.890  11.937  31.038  1.00  0.00           C
ATOM   3286  O   LYS A 230      17.486  12.461  32.084  1.00  0.00           O
ATOM   3287  N   ARG A 231      17.070  11.695  30.016  1.00  0.00           N
ATOM   3288  H   ARG A 231      17.475  11.284  29.175  1.00  0.00           H
ATOM   3289  CA  ARG A 231      15.606  11.713  30.121  1.00  0.00           C
ATOM   3290  HA  ARG A 231      15.305  12.504  30.808  1.00  0.00           H
ATOM   3291  CB  ARG A 231      15.042  12.038  28.731  1.00  0.00           C
```

| | | | | | | | | | | | |
|---|---|---|---|---|---|---|---|---|---|---|---|
| ATOM | 3292 | HB1 | ARG | A | 231 | 15.557 | 12.915 | 28.333 | 1.00 | 0.00 | H |
| ATOM | 3293 | HB2 | ARG | A | 231 | 15.215 | 11.203 | 28.055 | 1.00 | 0.00 | H |
| ATOM | 3294 | CG | ARG | A | 231 | 13.549 | 12.341 | 28.796 | 1.00 | 0.00 | C |
| ATOM | 3295 | HG1 | ARG | A | 231 | 13.011 | 11.454 | 29.129 | 1.00 | 0.00 | H |
| ATOM | 3296 | HG2 | ARG | A | 231 | 13.387 | 13.152 | 29.508 | 1.00 | 0.00 | H |
| ATOM | 3297 | CD | ARG | A | 231 | 13.000 | 12.761 | 27.435 | 1.00 | 0.00 | C |
| ATOM | 3298 | HD1 | ARG | A | 231 | 13.549 | 13.635 | 27.083 | 1.00 | 0.00 | H |
| ATOM | 3299 | HD2 | ARG | A | 231 | 13.147 | 11.946 | 26.726 | 1.00 | 0.00 | H |
| ATOM | 3300 | NE | ARG | A | 231 | 11.577 | 13.100 | 27.577 | 1.00 | 0.00 | N |
| ATOM | 3301 | HE | ARG | A | 231 | 11.308 | 13.485 | 28.476 | 1.00 | 0.00 | H |
| ATOM | 3302 | CZ | ARG | A | 231 | 10.573 | 12.709 | 26.807 | 1.00 | 0.00 | C |
| ATOM | 3303 | NH1 | ARG | A | 231 | 9.351 | 13.077 | 27.110 | 1.00 | 0.00 | N |
| ATOM | 3304 | HH11 | ARG | A | 231 | 9.173 | 13.556 | 27.980 | 1.00 | 0.00 | H |
| ATOM | 3305 | HH12 | ARG | A | 231 | 8.585 | 12.633 | 26.626 | 1.00 | 0.00 | H |
| ATOM | 3306 | NH2 | ARG | A | 231 | 10.729 | 11.937 | 25.756 | 1.00 | 0.00 | N |
| ATOM | 3307 | HH21 | ARG | A | 231 | 11.651 | 11.644 | 25.451 | 1.00 | 0.00 | H |
| ATOM | 3308 | HH22 | ARG | A | 231 | 9.909 | 11.569 | 25.300 | 1.00 | 0.00 | H |
| ATOM | 3309 | C | ARG | A | 231 | 15.109 | 10.361 | 30.672 | 1.00 | 0.00 | C |
| ATOM | 3310 | O | ARG | A | 231 | 15.748 | 9.336 | 30.432 | 1.00 | 0.00 | O |
| ATOM | 3311 | N | GLN | A | 232 | 14.000 | 10.303 | 31.419 | 1.00 | 0.00 | N |
| ATOM | 3312 | H | GLN | A | 232 | 13.491 | 11.159 | 31.638 | 1.00 | 0.00 | H |
| ATOM | 3313 | CA | GLN | A | 232 | 13.533 | 9.025 | 31.989 | 1.00 | 0.00 | C |
| ATOM | 3314 | HA | GLN | A | 232 | 14.359 | 8.577 | 32.538 | 1.00 | 0.00 | H |
| ATOM | 3315 | CB | GLN | A | 232 | 12.371 | 9.234 | 32.975 | 1.00 | 0.00 | C |
| ATOM | 3316 | HB1 | GLN | A | 232 | 11.631 | 9.897 | 32.526 | 1.00 | 0.00 | H |
| ATOM | 3317 | HB2 | GLN | A | 232 | 11.896 | 8.270 | 33.161 | 1.00 | 0.00 | H |
| ATOM | 3318 | CG | GLN | A | 232 | 12.799 | 9.813 | 34.335 | 1.00 | 0.00 | C |
| ATOM | 3319 | HG1 | GLN | A | 232 | 13.149 | 10.832 | 34.197 | 1.00 | 0.00 | H |
| ATOM | 3320 | HG2 | GLN | A | 232 | 11.928 | 9.844 | 34.989 | 1.00 | 0.00 | H |
| ATOM | 3321 | CD | GLN | A | 232 | 13.899 | 9.006 | 35.026 | 1.00 | 0.00 | C |
| ATOM | 3322 | OE1 | GLN | A | 232 | 13.704 | 7.907 | 35.531 | 1.00 | 0.00 | O |
| ATOM | 3323 | NE2 | GLN | A | 232 | 15.129 | 9.471 | 35.025 | 1.00 | 0.00 | N |
| ATOM | 3324 | HE21 | GLN | A | 232 | 15.815 | 8.981 | 35.582 | 1.00 | 0.00 | H |
| ATOM | 3325 | HE22 | GLN | A | 232 | 15.333 | 10.411 | 34.700 | 1.00 | 0.00 | H |
| ATOM | 3326 | C | GLN | A | 232 | 13.194 | 7.973 | 30.921 | 1.00 | 0.00 | C |
| ATOM | 3327 | O | GLN | A | 232 | 13.533 | 6.810 | 31.119 | 1.00 | 0.00 | O |
| ATOM | 3328 | N | GLU | A | 233 | 12.657 | 8.363 | 29.760 | 1.00 | 0.00 | N |
| ATOM | 3329 | H | GLU | A | 233 | 12.228 | 9.285 | 29.718 | 1.00 | 0.00 | H |
| ATOM | 3330 | CA | GLU | A | 233 | 12.430 | 7.481 | 28.603 | 1.00 | 0.00 | C |
| ATOM | 3331 | HA | GLU | A | 233 | 11.842 | 6.630 | 28.946 | 1.00 | 0.00 | H |
| ATOM | 3332 | CB | GLU | A | 233 | 11.598 | 8.238 | 27.541 | 1.00 | 0.00 | C |
| ATOM | 3333 | HB1 | GLU | A | 233 | 12.065 | 9.198 | 27.311 | 1.00 | 0.00 | H |
| ATOM | 3334 | HB2 | GLU | A | 233 | 11.604 | 7.649 | 26.625 | 1.00 | 0.00 | H |
| ATOM | 3335 | CG | GLU | A | 233 | 10.115 | 8.443 | 27.892 | 1.00 | 0.00 | C |
| ATOM | 3336 | HG1 | GLU | A | 233 | 9.553 | 8.409 | 26.957 | 1.00 | 0.00 | H |
| ATOM | 3337 | HG2 | GLU | A | 233 | 9.757 | 7.618 | 28.504 | 1.00 | 0.00 | H |
| ATOM | 3338 | CD | GLU | A | 233 | 9.791 | 9.770 | 28.576 | 1.00 | 0.00 | C |
| ATOM | 3339 | OE1 | GLU | A | 233 | 10.468 | 10.153 | 29.564 | 1.00 | 0.00 | O |
| ATOM | 3340 | OE2 | GLU | A | 233 | 8.820 | 10.422 | 28.126 | 1.00 | 0.00 | O |
| ATOM | 3341 | C | GLU | A | 233 | 13.708 | 6.879 | 27.953 | 1.00 | 0.00 | C |
| ATOM | 3342 | O | GLU | A | 233 | 13.590 | 6.161 | 26.952 | 1.00 | 0.00 | O |
| ATOM | 3343 | N | GLU | A | 234 | 14.921 | 7.161 | 28.447 | 1.00 | 0.00 | N |
| ATOM | 3344 | H | GLU | A | 234 | 14.978 | 7.723 | 29.288 | 1.00 | 0.00 | H |
| ATOM | 3345 | CA | GLU | A | 234 | 16.194 | 6.714 | 27.851 | 1.00 | 0.00 | C |

| | | | | | | | | | | | |
|---|---|---|---|---|---|---|---|---|---|---|---|
| ATOM | 3346 | HA | GLU | A | 234 | 16.003 | 6.364 | 26.839 | 1.00 | 0.00 | H |
| ATOM | 3347 | CB | GLU | A | 234 | 17.182 | 7.889 | 27.733 | 1.00 | 0.00 | C |
| ATOM | 3348 | HB1 | GLU | A | 234 | 17.334 | 8.331 | 28.715 | 1.00 | 0.00 | H |
| ATOM | 3349 | HB2 | GLU | A | 234 | 18.144 | 7.520 | 27.374 | 1.00 | 0.00 | H |
| ATOM | 3350 | CG | GLU | A | 234 | 16.681 | 8.948 | 26.743 | 1.00 | 0.00 | C |
| ATOM | 3351 | HG1 | GLU | A | 234 | 16.607 | 8.491 | 25.755 | 1.00 | 0.00 | H |
| ATOM | 3352 | HG2 | GLU | A | 234 | 15.687 | 9.284 | 27.035 | 1.00 | 0.00 | H |
| ATOM | 3353 | CD | GLU | A | 234 | 17.598 | 10.165 | 26.659 | 1.00 | 0.00 | C |
| ATOM | 3354 | OE1 | GLU | A | 234 | 18.196 | 10.566 | 27.682 | 1.00 | 0.00 | O |
| ATOM | 3355 | OE2 | GLU | A | 234 | 17.688 | 10.754 | 25.557 | 1.00 | 0.00 | O |
| ATOM | 3356 | C | GLU | A | 234 | 16.861 | 5.551 | 28.608 | 1.00 | 0.00 | C |
| ATOM | 3357 | O | GLU | A | 234 | 16.624 | 5.321 | 29.797 | 1.00 | 0.00 | O |
| ATOM | 3358 | N | SER | A | 235 | 17.750 | 4.839 | 27.919 | 1.00 | 0.00 | N |
| ATOM | 3359 | H | SER | A | 235 | 17.894 | 5.082 | 26.939 | 1.00 | 0.00 | H |
| ATOM | 3360 | CA | SER | A | 235 | 18.562 | 3.720 | 28.419 | 1.00 | 0.00 | C |
| ATOM | 3361 | HA | SER | A | 235 | 18.865 | 3.934 | 29.444 | 1.00 | 0.00 | H |
| ATOM | 3362 | CB | SER | A | 235 | 17.714 | 2.429 | 28.453 | 1.00 | 0.00 | C |
| ATOM | 3363 | HB1 | SER | A | 235 | 18.273 | 1.636 | 28.944 | 1.00 | 0.00 | H |
| ATOM | 3364 | HB2 | SER | A | 235 | 16.828 | 2.621 | 29.062 | 1.00 | 0.00 | H |
| ATOM | 3365 | OG | SER | A | 235 | 17.287 | 1.970 | 27.175 | 1.00 | 0.00 | O |
| ATOM | 3366 | HG | SER | A | 235 | 18.011 | 1.490 | 26.730 | 1.00 | 0.00 | H |
| ATOM | 3367 | C | SER | A | 235 | 19.861 | 3.573 | 27.590 | 1.00 | 0.00 | C |
| ATOM | 3368 | O | SER | A | 235 | 20.195 | 4.458 | 26.797 | 1.00 | 0.00 | O |
| ATOM | 3369 | N | GLY | A | 236 | 20.616 | 2.478 | 27.758 | 1.00 | 0.00 | N |
| ATOM | 3370 | H | GLY | A | 236 | 20.345 | 1.797 | 28.460 | 1.00 | 0.00 | H |
| ATOM | 3371 | CA | GLY | A | 236 | 21.761 | 2.140 | 26.898 | 1.00 | 0.00 | C |
| ATOM | 3372 | HA1 | GLY | A | 236 | 22.174 | 1.179 | 27.196 | 1.00 | 0.00 | H |
| ATOM | 3373 | HA2 | GLY | A | 236 | 21.414 | 2.061 | 25.868 | 1.00 | 0.00 | H |
| ATOM | 3374 | C | GLY | A | 236 | 22.885 | 3.175 | 26.930 | 1.00 | 0.00 | C |
| ATOM | 3375 | O | GLY | A | 236 | 23.134 | 3.803 | 27.960 | 1.00 | 0.00 | O |
| ATOM | 3376 | N | ALA | A | 237 | 23.579 | 3.350 | 25.803 | 1.00 | 0.00 | N |
| ATOM | 3377 | H | ALA | A | 237 | 23.330 | 2.795 | 24.991 | 1.00 | 0.00 | H |
| ATOM | 3378 | CA | ALA | A | 237 | 24.584 | 4.402 | 25.629 | 1.00 | 0.00 | C |
| ATOM | 3379 | HA | ALA | A | 237 | 25.223 | 4.434 | 26.514 | 1.00 | 0.00 | H |
| ATOM | 3380 | CB | ALA | A | 237 | 25.449 | 4.037 | 24.416 | 1.00 | 0.00 | C |
| ATOM | 3381 | HB1 | ALA | A | 237 | 26.216 | 4.795 | 24.265 | 1.00 | 0.00 | H |
| ATOM | 3382 | HB2 | ALA | A | 237 | 25.941 | 3.079 | 24.589 | 1.00 | 0.00 | H |
| ATOM | 3383 | HB3 | ALA | A | 237 | 24.832 | 3.973 | 23.518 | 1.00 | 0.00 | H |
| ATOM | 3384 | C | ALA | A | 237 | 23.950 | 5.800 | 25.476 | 1.00 | 0.00 | C |
| ATOM | 3385 | O | ALA | A | 237 | 24.583 | 6.800 | 25.832 | 1.00 | 0.00 | O |
| ATOM | 3386 | N | ALA | A | 238 | 22.702 | 5.879 | 24.993 | 1.00 | 0.00 | N |
| ATOM | 3387 | H | ALA | A | 238 | 22.242 | 5.018 | 24.711 | 1.00 | 0.00 | H |
| ATOM | 3388 | CA | ALA | A | 238 | 21.949 | 7.127 | 24.855 | 1.00 | 0.00 | C |
| ATOM | 3389 | HA | ALA | A | 238 | 22.513 | 7.797 | 24.210 | 1.00 | 0.00 | H |
| ATOM | 3390 | CB | ALA | A | 238 | 20.610 | 6.826 | 24.173 | 1.00 | 0.00 | C |
| ATOM | 3391 | HB1 | ALA | A | 238 | 20.039 | 7.750 | 24.067 | 1.00 | 0.00 | H |
| ATOM | 3392 | HB2 | ALA | A | 238 | 20.781 | 6.404 | 23.182 | 1.00 | 0.00 | H |
| ATOM | 3393 | HB3 | ALA | A | 238 | 20.033 | 6.122 | 24.771 | 1.00 | 0.00 | H |
| ATOM | 3394 | C | ALA | A | 238 | 21.759 | 7.847 | 26.199 | 1.00 | 0.00 | C |
| ATOM | 3395 | O | ALA | A | 238 | 21.914 | 9.065 | 26.263 | 1.00 | 0.00 | O |
| ATOM | 3396 | N | ALA | A | 239 | 21.548 | 7.100 | 27.288 | 1.00 | 0.00 | N |
| ATOM | 3397 | H | ALA | A | 239 | 21.367 | 6.110 | 27.146 | 1.00 | 0.00 | H |
| ATOM | 3398 | CA | ALA | A | 239 | 21.506 | 7.612 | 28.664 | 1.00 | 0.00 | C |
| ATOM | 3399 | HA | ALA | A | 239 | 20.756 | 8.405 | 28.706 | 1.00 | 0.00 | H |

```
ATOM   3400  CB  ALA A 239      21.042   6.467  29.569  1.00  0.00           C
ATOM   3401  HB1 ALA A 239      21.009   6.814  30.601  1.00  0.00           H
ATOM   3402  HB2 ALA A 239      20.046   6.143  29.272  1.00  0.00           H
ATOM   3403  HB3 ALA A 239      21.739   5.629  29.503  1.00  0.00           H
ATOM   3404  C   ALA A 239      22.825   8.244  29.174  1.00  0.00           C
ATOM   3405  O   ALA A 239      22.864   8.779  30.285  1.00  0.00           O
ATOM   3406  N   SER A 240      23.901   8.224  28.383  1.00  0.00           N
ATOM   3407  H   SER A 240      23.839   7.704  27.517  1.00  0.00           H
ATOM   3408  CA  SER A 240      25.171   8.908  28.679  1.00  0.00           C
ATOM   3409  HA  SER A 240      25.123   9.328  29.681  1.00  0.00           H
ATOM   3410  CB  SER A 240      26.321   7.893  28.679  1.00  0.00           C
ATOM   3411  HB1 SER A 240      26.338   7.345  27.737  1.00  0.00           H
ATOM   3412  HB2 SER A 240      27.269   8.417  28.798  1.00  0.00           H
ATOM   3413  OG  SER A 240      26.166   6.990  29.756  1.00  0.00           O
ATOM   3414  HG  SER A 240      27.028   6.551  29.917  1.00  0.00           H
ATOM   3415  C   SER A 240      25.476  10.096  27.741  1.00  0.00           C
ATOM   3416  O   SER A 240      26.572  10.664  27.794  1.00  0.00           O
ATOM   3417  N   LEU A 241      24.524  10.503  26.891  1.00  0.00           N
ATOM   3418  H   LEU A 241      23.637  10.009  26.891  1.00  0.00           H
ATOM   3419  CA  LEU A 241      24.631  11.668  26.000  1.00  0.00           C
ATOM   3420  HA  LEU A 241      25.670  11.783  25.694  1.00  0.00           H
ATOM   3421  CB  LEU A 241      23.772  11.425  24.741  1.00  0.00           C
ATOM   3422  HB1 LEU A 241      22.732  11.306  25.053  1.00  0.00           H
ATOM   3423  HB2 LEU A 241      23.815  12.306  24.104  1.00  0.00           H
ATOM   3424  CG  LEU A 241      24.194  10.205  23.899  1.00  0.00           C
ATOM   3425  HG  LEU A 241      24.203   9.316  24.528  1.00  0.00           H
ATOM   3426  CD1 LEU A 241      23.163   9.986  22.785  1.00  0.00           C
ATOM   3427 HD11 LEU A 241      23.392   9.066  22.249  1.00  0.00           H
ATOM   3428 HD12 LEU A 241      22.170   9.884  23.222  1.00  0.00           H
ATOM   3429 HD13 LEU A 241      23.166  10.829  22.093  1.00  0.00           H
ATOM   3430  CD2 LEU A 241      25.592  10.374  23.281  1.00  0.00           C
ATOM   3431 HD21 LEU A 241      25.822   9.516  22.649  1.00  0.00           H
ATOM   3432 HD22 LEU A 241      25.631  11.283  22.680  1.00  0.00           H
ATOM   3433 HD23 LEU A 241      26.347  10.427  24.065  1.00  0.00           H
ATOM   3434  C   LEU A 241      24.268  12.971  26.740  1.00  0.00           C
ATOM   3435  O   LEU A 241      23.399  13.730  26.323  1.00  0.00           O
ATOM   3436  N   ASN A 242      24.931  13.221  27.869  1.00  0.00           N
ATOM   3437  H   ASN A 242      25.666  12.579  28.137  1.00  0.00           H
ATOM   3438  CA  ASN A 242      24.610  14.313  28.790  1.00  0.00           C
ATOM   3439  HA  ASN A 242      24.337  15.184  28.194  1.00  0.00           H
ATOM   3440  CB  ASN A 242      23.382  13.929  29.635  1.00  0.00           C
ATOM   3441  HB1 ASN A 242      23.199  14.684  30.395  1.00  0.00           H
ATOM   3442  HB2 ASN A 242      22.506  13.907  28.987  1.00  0.00           H
ATOM   3443  CG  ASN A 242      23.513  12.570  30.308  1.00  0.00           C
ATOM   3444  OD1 ASN A 242      24.251  12.383  31.269  1.00  0.00           O
ATOM   3445  ND2 ASN A 242      22.800  11.579  29.813  1.00  0.00           N
ATOM   3446 HD21 ASN A 242      22.171  11.727  29.026  1.00  0.00           H
ATOM   3447 HD22 ASN A 242      22.840  10.658  30.225  1.00  0.00           H
ATOM   3448  C   ASN A 242      25.813  14.737  29.653  1.00  0.00           C
ATOM   3449  O   ASN A 242      26.796  14.003  29.790  1.00  0.00           O
ATOM   3450  N   VAL A 243      25.738  15.951  30.210  1.00  0.00           N
ATOM   3451  H   VAL A 243      24.860  16.456  30.091  1.00  0.00           H
ATOM   3452  CA  VAL A 243      26.835  16.617  30.940  1.00  0.00           C
ATOM   3453  HA  VAL A 243      27.668  16.749  30.248  1.00  0.00           H
```

```
ATOM   3454  CB  VAL A 243      26.398  18.013  31.448  1.00  0.00           C
ATOM   3455  HB  VAL A 243      25.466  17.884  31.984  1.00  0.00           H
ATOM   3456  CG1 VAL A 243      27.405  18.645  32.424  1.00  0.00           C
ATOM   3457 HG11 VAL A 243      27.138  19.684  32.621  1.00  0.00           H
ATOM   3458 HG12 VAL A 243      27.400  18.110  33.375  1.00  0.00           H
ATOM   3459 HG13 VAL A 243      28.405  18.596  31.997  1.00  0.00           H
ATOM   3460  CG2 VAL A 243      26.134  19.003  30.298  1.00  0.00           C
ATOM   3461 HG21 VAL A 243      25.624  19.886  30.685  1.00  0.00           H
ATOM   3462 HG22 VAL A 243      27.071  19.318  29.846  1.00  0.00           H
ATOM   3463 HG23 VAL A 243      25.509  18.554  29.528  1.00  0.00           H
ATOM   3464  C   VAL A 243      27.317  15.767  32.123  1.00  0.00           C
ATOM   3465  O   VAL A 243      28.523  15.608  32.305  1.00  0.00           O
ATOM   3466  N   ASP A 244      26.397  15.195  32.903  1.00  0.00           N
ATOM   3467  H   ASP A 244      25.410  15.359  32.715  1.00  0.00           H
ATOM   3468  CA  ASP A 244      26.718  14.389  34.080  1.00  0.00           C
ATOM   3469  HA  ASP A 244      27.312  14.996  34.761  1.00  0.00           H
ATOM   3470  CB  ASP A 244      25.415  13.999  34.784  1.00  0.00           C
ATOM   3471  HB1 ASP A 244      24.939  14.899  35.176  1.00  0.00           H
ATOM   3472  HB2 ASP A 244      24.739  13.534  34.064  1.00  0.00           H
ATOM   3473  CG  ASP A 244      25.670  13.024  35.929  1.00  0.00           C
ATOM   3474  OD1 ASP A 244      26.337  13.413  36.912  1.00  0.00           O
ATOM   3475  OD2 ASP A 244      25.214  11.862  35.835  1.00  0.00           O
ATOM   3476  C   ASP A 244      27.546  13.149  33.726  1.00  0.00           C
ATOM   3477  O   ASP A 244      28.607  12.936  34.313  1.00  0.00           O
ATOM   3478  N   ALA A 245      27.110  12.375  32.726  1.00  0.00           N
ATOM   3479  H   ALA A 245      26.233  12.609  32.276  1.00  0.00           H
ATOM   3480  CA  ALA A 245      27.864  11.221  32.239  1.00  0.00           C
ATOM   3481  HA  ALA A 245      28.025  10.526  33.066  1.00  0.00           H
ATOM   3482  CB  ALA A 245      27.044  10.519  31.159  1.00  0.00           C
ATOM   3483  HB1 ALA A 245      27.623   9.692  30.748  1.00  0.00           H
ATOM   3484  HB2 ALA A 245      26.132  10.121  31.598  1.00  0.00           H
ATOM   3485  HB3 ALA A 245      26.794  11.221  30.364  1.00  0.00           H
ATOM   3486  C   ALA A 245      29.241  11.629  31.706  1.00  0.00           C
ATOM   3487  O   ALA A 245      30.232  10.958  31.988  1.00  0.00           O
ATOM   3488  N   ALA A 246      29.325  12.756  30.993  1.00  0.00           N
ATOM   3489  H   ALA A 246      28.462  13.241  30.767  1.00  0.00           H
ATOM   3490  CA  ALA A 246      30.586  13.279  30.474  1.00  0.00           C
ATOM   3491  HA  ALA A 246      31.077  12.488  29.905  1.00  0.00           H
ATOM   3492  CB  ALA A 246      30.263  14.425  29.516  1.00  0.00           C
ATOM   3493  HB1 ALA A 246      31.179  14.765  29.034  1.00  0.00           H
ATOM   3494  HB2 ALA A 246      29.567  14.062  28.760  1.00  0.00           H
ATOM   3495  HB3 ALA A 246      29.807  15.259  30.050  1.00  0.00           H
ATOM   3496  C   ALA A 246      31.556  13.700  31.591  1.00  0.00           C
ATOM   3497  O   ALA A 246      32.736  13.364  31.522  1.00  0.00           O
ATOM   3498  N   VAL A 247      31.064  14.383  32.629  1.00  0.00           N
ATOM   3499  H   VAL A 247      30.076  14.628  32.601  1.00  0.00           H
ATOM   3500  CA  VAL A 247      31.871  14.808  33.786  1.00  0.00           C
ATOM   3501  HA  VAL A 247      32.780  15.275  33.401  1.00  0.00           H
ATOM   3502  CB  VAL A 247      31.130  15.864  34.641  1.00  0.00           C
ATOM   3503  HB  VAL A 247      30.106  15.526  34.805  1.00  0.00           H
ATOM   3504  CG1 VAL A 247      31.789  16.098  36.014  1.00  0.00           C
ATOM   3505 HG11 VAL A 247      31.290  16.911  36.538  1.00  0.00           H
ATOM   3506 HG12 VAL A 247      31.703  15.205  36.634  1.00  0.00           H
ATOM   3507 HG13 VAL A 247      32.843  16.350  35.887  1.00  0.00           H
```

| | | | | | | | | | | | |
|---|---|---|---|---|---|---|---|---|---|---|---|
| ATOM | 3508 | CG2 | VAL | A | 247 | 31.094 | 17.208 | 33.887 | 1.00 | 0.00 | C |
| ATOM | 3509 | HG21 | VAL | A | 247 | 30.515 | 17.939 | 34.450 | 1.00 | 0.00 | H |
| ATOM | 3510 | HG22 | VAL | A | 247 | 32.107 | 17.589 | 33.752 | 1.00 | 0.00 | H |
| ATOM | 3511 | HG23 | VAL | A | 247 | 30.633 | 17.086 | 32.907 | 1.00 | 0.00 | H |
| ATOM | 3512 | C | VAL | A | 247 | 32.298 | 13.600 | 34.629 | 1.00 | 0.00 | C |
| ATOM | 3513 | O | VAL | A | 247 | 33.479 | 13.489 | 34.952 | 1.00 | 0.00 | O |
| ATOM | 3514 | N | GLN | A | 248 | 31.382 | 12.676 | 34.954 | 1.00 | 0.00 | N |
| ATOM | 3515 | H | GLN | A | 248 | 30.415 | 12.806 | 34.665 | 1.00 | 0.00 | H |
| ATOM | 3516 | CA | GLN | A | 248 | 31.733 | 11.456 | 35.688 | 1.00 | 0.00 | C |
| ATOM | 3517 | HA | GLN | A | 248 | 32.255 | 11.737 | 36.605 | 1.00 | 0.00 | H |
| ATOM | 3518 | CB | GLN | A | 248 | 30.476 | 10.652 | 36.080 | 1.00 | 0.00 | C |
| ATOM | 3519 | HB1 | GLN | A | 248 | 29.822 | 10.536 | 35.214 | 1.00 | 0.00 | H |
| ATOM | 3520 | HB2 | GLN | A | 248 | 30.798 | 9.657 | 36.394 | 1.00 | 0.00 | H |
| ATOM | 3521 | CG | GLN | A | 248 | 29.691 | 11.281 | 37.250 | 1.00 | 0.00 | C |
| ATOM | 3522 | HG1 | GLN | A | 248 | 30.376 | 11.503 | 38.068 | 1.00 | 0.00 | H |
| ATOM | 3523 | HG2 | GLN | A | 248 | 29.235 | 12.216 | 36.923 | 1.00 | 0.00 | H |
| ATOM | 3524 | CD | GLN | A | 248 | 28.607 | 10.337 | 37.782 | 1.00 | 0.00 | C |
| ATOM | 3525 | OE1 | GLN | A | 248 | 28.865 | 9.401 | 38.534 | 1.00 | 0.00 | O |
| ATOM | 3526 | NE2 | GLN | A | 248 | 27.357 | 10.521 | 37.426 | 1.00 | 0.00 | N |
| ATOM | 3527 | HE21 | GLN | A | 248 | 27.064 | 11.333 | 36.893 | 1.00 | 0.00 | H |
| ATOM | 3528 | HE22 | GLN | A | 248 | 26.697 | 9.781 | 37.617 | 1.00 | 0.00 | H |
| ATOM | 3529 | C | GLN | A | 248 | 32.730 | 10.600 | 34.892 | 1.00 | 0.00 | C |
| ATOM | 3530 | O | GLN | A | 248 | 33.699 | 10.131 | 35.483 | 1.00 | 0.00 | O |
| ATOM | 3531 | N | GLN | A | 249 | 32.562 | 10.458 | 33.570 | 1.00 | 0.00 | N |
| ATOM | 3532 | H | GLN | A | 249 | 31.732 | 10.851 | 33.136 | 1.00 | 0.00 | H |
| ATOM | 3533 | CA | GLN | A | 249 | 33.507 | 9.752 | 32.698 | 1.00 | 0.00 | C |
| ATOM | 3534 | HA | GLN | A | 249 | 33.597 | 8.721 | 33.047 | 1.00 | 0.00 | H |
| ATOM | 3535 | CB | GLN | A | 249 | 32.973 | 9.740 | 31.249 | 1.00 | 0.00 | C |
| ATOM | 3536 | HB1 | GLN | A | 249 | 32.055 | 9.151 | 31.220 | 1.00 | 0.00 | H |
| ATOM | 3537 | HB2 | GLN | A | 249 | 32.744 | 10.762 | 30.948 | 1.00 | 0.00 | H |
| ATOM | 3538 | CG | GLN | A | 249 | 33.977 | 9.145 | 30.243 | 1.00 | 0.00 | C |
| ATOM | 3539 | HG1 | GLN | A | 249 | 34.866 | 9.774 | 30.207 | 1.00 | 0.00 | H |
| ATOM | 3540 | HG2 | GLN | A | 249 | 34.263 | 8.155 | 30.593 | 1.00 | 0.00 | H |
| ATOM | 3541 | CD | GLN | A | 249 | 33.462 | 9.018 | 28.806 | 1.00 | 0.00 | C |
| ATOM | 3542 | OE1 | GLN | A | 249 | 33.898 | 8.156 | 28.054 | 1.00 | 0.00 | O |
| ATOM | 3543 | NE2 | GLN | A | 249 | 32.557 | 9.860 | 28.351 | 1.00 | 0.00 | N |
| ATOM | 3544 | HE21 | GLN | A | 249 | 32.169 | 10.576 | 28.940 | 1.00 | 0.00 | H |
| ATOM | 3545 | HE22 | GLN | A | 249 | 32.251 | 9.749 | 27.402 | 1.00 | 0.00 | H |
| ATOM | 3546 | C | GLN | A | 249 | 34.911 | 10.358 | 32.770 | 1.00 | 0.00 | C |
| ATOM | 3547 | O | GLN | A | 249 | 35.859 | 9.632 | 33.055 | 1.00 | 0.00 | O |
| ATOM | 3548 | N | TRP | A | 250 | 35.078 | 11.660 | 32.509 | 1.00 | 0.00 | N |
| ATOM | 3549 | H | TRP | A | 250 | 34.274 | 12.235 | 32.276 | 1.00 | 0.00 | H |
| ATOM | 3550 | CA | TRP | A | 250 | 36.414 | 12.263 | 32.460 | 1.00 | 0.00 | C |
| ATOM | 3551 | HA | TRP | A | 250 | 37.046 | 11.618 | 31.846 | 1.00 | 0.00 | H |
| ATOM | 3552 | CB | TRP | A | 250 | 36.351 | 13.629 | 31.760 | 1.00 | 0.00 | C |
| ATOM | 3553 | HB1 | TRP | A | 250 | 35.429 | 14.149 | 32.027 | 1.00 | 0.00 | H |
| ATOM | 3554 | HB2 | TRP | A | 250 | 37.186 | 14.241 | 32.103 | 1.00 | 0.00 | H |
| ATOM | 3555 | CG | TRP | A | 250 | 36.461 | 13.515 | 30.268 | 1.00 | 0.00 | C |
| ATOM | 3556 | CD1 | TRP | A | 250 | 35.453 | 13.214 | 29.420 | 1.00 | 0.00 | C |
| ATOM | 3557 | HD1 | TRP | A | 250 | 34.422 | 13.070 | 29.713 | 1.00 | 0.00 | H |
| ATOM | 3558 | NE1 | TRP | A | 250 | 35.947 | 13.096 | 28.135 | 1.00 | 0.00 | N |
| ATOM | 3559 | HE1 | TRP | A | 250 | 35.365 | 12.977 | 27.311 | 1.00 | 0.00 | H |
| ATOM | 3560 | CE2 | TRP | A | 250 | 37.314 | 13.257 | 28.097 | 1.00 | 0.00 | C |
| ATOM | 3561 | CZ2 | TRP | A | 250 | 38.266 | 13.170 | 27.070 | 1.00 | 0.00 | C |

| | | | | | | | | | | | |
|---|---|---|---|---|---|---|---|---|---|---|---|
| ATOM | 3562 | HZ2 | TRP | A | 250 | 37.967 | 12.904 | 26.068 | 1.00 | 0.00 | H |
| ATOM | 3563 | CH2 | TRP | A | 250 | 39.610 | 13.461 | 27.355 | 1.00 | 0.00 | C |
| ATOM | 3564 | HH2 | TRP | A | 250 | 40.355 | 13.413 | 26.574 | 1.00 | 0.00 | H |
| ATOM | 3565 | CZ3 | TRP | A | 250 | 39.977 | 13.860 | 28.651 | 1.00 | 0.00 | C |
| ATOM | 3566 | HZ3 | TRP | A | 250 | 41.006 | 14.112 | 28.870 | 1.00 | 0.00 | H |
| ATOM | 3567 | CE3 | TRP | A | 250 | 39.011 | 13.933 | 29.671 | 1.00 | 0.00 | C |
| ATOM | 3568 | HE3 | TRP | A | 250 | 39.306 | 14.254 | 30.654 | 1.00 | 0.00 | H |
| ATOM | 3569 | CD2 | TRP | A | 250 | 37.661 | 13.605 | 29.434 | 1.00 | 0.00 | C |
| ATOM | 3570 | C | TRP | A | 250 | 37.102 | 12.308 | 33.833 | 1.00 | 0.00 | C |
| ATOM | 3571 | O | TRP | A | 250 | 38.326 | 12.159 | 33.892 | 1.00 | 0.00 | O |
| ATOM | 3572 | N | LEU | A | 251 | 36.345 | 12.443 | 34.931 | 1.00 | 0.00 | N |
| ATOM | 3573 | H | LEU | A | 251 | 35.344 | 12.552 | 34.808 | 1.00 | 0.00 | H |
| ATOM | 3574 | CA | LEU | A | 251 | 36.875 | 12.310 | 36.294 | 1.00 | 0.00 | C |
| ATOM | 3575 | HA | LEU | A | 251 | 37.769 | 12.926 | 36.396 | 1.00 | 0.00 | H |
| ATOM | 3576 | CB | LEU | A | 251 | 35.823 | 12.767 | 37.325 | 1.00 | 0.00 | C |
| ATOM | 3577 | HB1 | LEU | A | 251 | 34.889 | 12.238 | 37.128 | 1.00 | 0.00 | H |
| ATOM | 3578 | HB2 | LEU | A | 251 | 36.166 | 12.461 | 38.315 | 1.00 | 0.00 | H |
| ATOM | 3579 | CG | LEU | A | 251 | 35.541 | 14.282 | 37.378 | 1.00 | 0.00 | C |
| ATOM | 3580 | HG | LEU | A | 251 | 35.221 | 14.624 | 36.397 | 1.00 | 0.00 | H |
| ATOM | 3581 | CD1 | LEU | A | 251 | 34.399 | 14.536 | 38.374 | 1.00 | 0.00 | C |
| ATOM | 3582 | HD11 | LEU | A | 251 | 34.157 | 15.598 | 38.403 | 1.00 | 0.00 | H |
| ATOM | 3583 | HD12 | LEU | A | 251 | 33.509 | 13.987 | 38.064 | 1.00 | 0.00 | H |
| ATOM | 3584 | HD13 | LEU | A | 251 | 34.690 | 14.210 | 39.373 | 1.00 | 0.00 | H |
| ATOM | 3585 | CD2 | LEU | A | 251 | 36.776 | 15.098 | 37.789 | 1.00 | 0.00 | C |
| ATOM | 3586 | HD21 | LEU | A | 251 | 36.500 | 16.144 | 37.919 | 1.00 | 0.00 | H |
| ATOM | 3587 | HD22 | LEU | A | 251 | 37.185 | 14.719 | 38.726 | 1.00 | 0.00 | H |
| ATOM | 3588 | HD23 | LEU | A | 251 | 37.538 | 15.044 | 37.013 | 1.00 | 0.00 | H |
| ATOM | 3589 | C | LEU | A | 251 | 37.310 | 10.868 | 36.587 | 1.00 | 0.00 | C |
| ATOM | 3590 | O | LEU | A | 251 | 38.440 | 10.656 | 37.028 | 1.00 | 0.00 | O |
| ATOM | 3591 | N | GLN | A | 252 | 36.451 | 9.876 | 36.339 | 1.00 | 0.00 | N |
| ATOM | 3592 | H | GLN | A | 252 | 35.542 | 10.100 | 35.939 | 1.00 | 0.00 | H |
| ATOM | 3593 | CA | GLN | A | 252 | 36.732 | 8.474 | 36.664 | 1.00 | 0.00 | C |
| ATOM | 3594 | HA | GLN | A | 252 | 37.087 | 8.417 | 37.695 | 1.00 | 0.00 | H |
| ATOM | 3595 | CB | GLN | A | 252 | 35.442 | 7.648 | 36.532 | 1.00 | 0.00 | C |
| ATOM | 3596 | HB1 | GLN | A | 252 | 34.967 | 7.865 | 35.575 | 1.00 | 0.00 | H |
| ATOM | 3597 | HB2 | GLN | A | 252 | 35.691 | 6.588 | 36.535 | 1.00 | 0.00 | H |
| ATOM | 3598 | CG | GLN | A | 252 | 34.454 | 7.926 | 37.680 | 1.00 | 0.00 | C |
| ATOM | 3599 | HG1 | GLN | A | 252 | 34.285 | 8.997 | 37.786 | 1.00 | 0.00 | H |
| ATOM | 3600 | HG2 | GLN | A | 252 | 33.500 | 7.458 | 37.436 | 1.00 | 0.00 | H |
| ATOM | 3601 | CD | GLN | A | 252 | 34.948 | 7.364 | 39.011 | 1.00 | 0.00 | C |
| ATOM | 3602 | OE1 | GLN | A | 252 | 35.488 | 8.064 | 39.863 | 1.00 | 0.00 | O |
| ATOM | 3603 | NE2 | GLN | A | 252 | 34.814 | 6.076 | 39.221 | 1.00 | 0.00 | N |
| ATOM | 3604 | HE21 | GLN | A | 252 | 34.415 | 5.485 | 38.513 | 1.00 | 0.00 | H |
| ATOM | 3605 | HE22 | GLN | A | 252 | 35.143 | 5.681 | 40.095 | 1.00 | 0.00 | H |
| ATOM | 3606 | C | GLN | A | 252 | 37.852 | 7.891 | 35.796 | 1.00 | 0.00 | C |
| ATOM | 3607 | O | GLN | A | 252 | 38.697 | 7.153 | 36.304 | 1.00 | 0.00 | O |
| ATOM | 3608 | N | LYS | A | 253 | 37.940 | 8.279 | 34.523 | 1.00 | 0.00 | N |
| ATOM | 3609 | H | LYS | A | 253 | 37.192 | 8.849 | 34.132 | 1.00 | 0.00 | H |
| ATOM | 3610 | CA | LYS | A | 253 | 39.053 | 7.921 | 33.637 | 1.00 | 0.00 | C |
| ATOM | 3611 | HA | LYS | A | 253 | 39.306 | 6.876 | 33.824 | 1.00 | 0.00 | H |
| ATOM | 3612 | CB | LYS | A | 253 | 38.593 | 8.019 | 32.168 | 1.00 | 0.00 | C |
| ATOM | 3613 | HB1 | LYS | A | 253 | 38.203 | 9.017 | 31.967 | 1.00 | 0.00 | H |
| ATOM | 3614 | HB2 | LYS | A | 253 | 39.448 | 7.844 | 31.515 | 1.00 | 0.00 | H |
| ATOM | 3615 | CG | LYS | A | 253 | 37.525 | 6.958 | 31.842 | 1.00 | 0.00 | C |

| | | | | | | | | | | |
|---|---|---|---|---|---|---|---|---|---|---|
| ATOM | 3616 | HG1 | LYS | A | 253 | 37.933 | 5.968 | 32.056 | 1.00 | 0.00 | H |
| ATOM | 3617 | HG2 | LYS | A | 253 | 36.647 | 7.111 | 32.471 | 1.00 | 0.00 | H |
| ATOM | 3618 | CD | LYS | A | 253 | 37.092 | 7.008 | 30.371 | 1.00 | 0.00 | C |
| ATOM | 3619 | HD1 | LYS | A | 253 | 36.677 | 7.992 | 30.151 | 1.00 | 0.00 | H |
| ATOM | 3620 | HD2 | LYS | A | 253 | 37.960 | 6.840 | 29.733 | 1.00 | 0.00 | H |
| ATOM | 3621 | CE | LYS | A | 253 | 36.039 | 5.918 | 30.121 | 1.00 | 0.00 | C |
| ATOM | 3622 | HE1 | LYS | A | 253 | 36.486 | 4.942 | 30.338 | 1.00 | 0.00 | H |
| ATOM | 3623 | HE2 | LYS | A | 253 | 35.209 | 6.064 | 30.818 | 1.00 | 0.00 | H |
| ATOM | 3624 | NZ | LYS | A | 253 | 35.522 | 5.931 | 28.732 | 1.00 | 0.00 | N |
| ATOM | 3625 | HZ1 | LYS | A | 253 | 34.790 | 5.240 | 28.621 | 1.00 | 0.00 | H |
| ATOM | 3626 | HZ2 | LYS | A | 253 | 35.113 | 6.828 | 28.482 | 1.00 | 0.00 | H |
| ATOM | 3627 | HZ3 | LYS | A | 253 | 36.245 | 5.682 | 28.061 | 1.00 | 0.00 | H |
| ATOM | 3628 | C | LYS | A | 253 | 40.354 | 8.704 | 33.939 | 1.00 | 0.00 | C |
| ATOM | 3629 | O | LYS | A | 253 | 41.323 | 8.569 | 33.194 | 1.00 | 0.00 | O |
| ATOM | 3630 | N | GLY | A | 254 | 40.424 | 9.469 | 35.039 | 1.00 | 0.00 | N |
| ATOM | 3631 | H | GLY | A | 254 | 39.588 | 9.538 | 35.607 | 1.00 | 0.00 | H |
| ATOM | 3632 | CA | GLY | A | 254 | 41.687 | 9.934 | 35.639 | 1.00 | 0.00 | C |
| ATOM | 3633 | HA1 | GLY | A | 254 | 41.631 | 9.763 | 36.715 | 1.00 | 0.00 | H |
| ATOM | 3634 | HA2 | GLY | A | 254 | 42.528 | 9.364 | 35.241 | 1.00 | 0.00 | H |
| ATOM | 3635 | C | GLY | A | 254 | 42.019 | 11.416 | 35.466 | 1.00 | 0.00 | C |
| ATOM | 3636 | O | GLY | A | 254 | 43.147 | 11.803 | 35.757 | 1.00 | 0.00 | O |
| ATOM | 3637 | N | THR | A | 255 | 41.083 | 12.265 | 35.019 | 1.00 | 0.00 | N |
| ATOM | 3638 | H | THR | A | 255 | 40.157 | 11.910 | 34.818 | 1.00 | 0.00 | H |
| ATOM | 3639 | CA | THR | A | 255 | 41.323 | 13.722 | 34.946 | 1.00 | 0.00 | C |
| ATOM | 3640 | HA | THR | A | 255 | 42.338 | 13.873 | 34.586 | 1.00 | 0.00 | H |
| ATOM | 3641 | CB | THR | A | 255 | 40.392 | 14.440 | 33.971 | 1.00 | 0.00 | C |
| ATOM | 3642 | HB | THR | A | 255 | 39.376 | 14.481 | 34.370 | 1.00 | 0.00 | H |
| ATOM | 3643 | CG2 | THR | A | 255 | 40.876 | 15.858 | 33.667 | 1.00 | 0.00 | C |
| ATOM | 3644 | HG21 | THR | A | 255 | 40.252 | 16.297 | 32.890 | 1.00 | 0.00 | H |
| ATOM | 3645 | HG22 | THR | A | 255 | 40.811 | 16.477 | 34.560 | 1.00 | 0.00 | H |
| ATOM | 3646 | HG23 | THR | A | 255 | 41.911 | 15.836 | 33.323 | 1.00 | 0.00 | H |
| ATOM | 3647 | OG1 | THR | A | 255 | 40.399 | 13.726 | 32.768 | 1.00 | 0.00 | O |
| ATOM | 3648 | HG1 | THR | A | 255 | 39.803 | 12.972 | 32.896 | 1.00 | 0.00 | H |
| ATOM | 3649 | C | THR | A | 255 | 41.156 | 14.334 | 36.337 | 1.00 | 0.00 | C |
| ATOM | 3650 | O | THR | A | 255 | 40.102 | 14.112 | 36.939 | 1.00 | 0.00 | O |
| ATOM | 3651 | N | PRO | A | 256 | 42.123 | 15.121 | 36.845 | 1.00 | 0.00 | N |
| ATOM | 3652 | CD | PRO | A | 256 | 43.476 | 15.269 | 36.339 | 1.00 | 0.00 | C |
| ATOM | 3653 | HD1 | PRO | A | 256 | 43.492 | 15.985 | 35.517 | 1.00 | 0.00 | H |
| ATOM | 3654 | HD2 | PRO | A | 256 | 43.870 | 14.307 | 36.029 | 1.00 | 0.00 | H |
| ATOM | 3655 | CG | PRO | A | 256 | 44.303 | 15.782 | 37.509 | 1.00 | 0.00 | C |
| ATOM | 3656 | HG1 | PRO | A | 256 | 45.141 | 16.393 | 37.179 | 1.00 | 0.00 | H |
| ATOM | 3657 | HG2 | PRO | A | 256 | 44.653 | 14.934 | 38.097 | 1.00 | 0.00 | H |
| ATOM | 3658 | CB | PRO | A | 256 | 43.298 | 16.585 | 38.324 | 1.00 | 0.00 | C |
| ATOM | 3659 | HB1 | PRO | A | 256 | 43.251 | 17.604 | 37.942 | 1.00 | 0.00 | H |
| ATOM | 3660 | HB2 | PRO | A | 256 | 43.571 | 16.588 | 39.376 | 1.00 | 0.00 | H |
| ATOM | 3661 | CA | PRO | A | 256 | 41.966 | 15.864 | 38.084 | 1.00 | 0.00 | C |
| ATOM | 3662 | HA | PRO | A | 256 | 41.786 | 15.150 | 38.885 | 1.00 | 0.00 | H |
| ATOM | 3663 | C | PRO | A | 256 | 40.804 | 16.853 | 37.990 | 1.00 | 0.00 | C |
| ATOM | 3664 | O | PRO | A | 256 | 40.675 | 17.570 | 36.997 | 1.00 | 0.00 | O |
| ATOM | 3665 | N | ALA | A | 257 | 39.990 | 16.936 | 39.046 | 1.00 | 0.00 | N |
| ATOM | 3666 | H | ALA | A | 257 | 40.154 | 16.326 | 39.836 | 1.00 | 0.00 | H |
| ATOM | 3667 | CA | ALA | A | 257 | 38.917 | 17.927 | 39.146 | 1.00 | 0.00 | C |
| ATOM | 3668 | HA | ALA | A | 257 | 38.250 | 17.812 | 38.290 | 1.00 | 0.00 | H |
| ATOM | 3669 | CB | ALA | A | 257 | 38.126 | 17.649 | 40.429 | 1.00 | 0.00 | C |

| | | | | | | | | | | |
|---|---|---|---|---|---|---|---|---|---|---|
| ATOM | 3670 | HB1 | ALA | A | 257 | 37.278 | 18.331 | 40.492 | 1.00 | 0.00 | H |
| ATOM | 3671 | HB2 | ALA | A | 257 | 37.751 | 16.624 | 40.424 | 1.00 | 0.00 | H |
| ATOM | 3672 | HB3 | ALA | A | 257 | 38.767 | 17.797 | 41.298 | 1.00 | 0.00 | H |
| ATOM | 3673 | C | ALA | A | 257 | 39.461 | 19.368 | 39.108 | 1.00 | 0.00 | C |
| ATOM | 3674 | O | ALA | A | 257 | 38.815 | 20.259 | 38.564 | 1.00 | 0.00 | O |
| ATOM | 3675 | N | SER | A | 258 | 40.681 | 19.588 | 39.602 | 1.00 | 0.00 | N |
| ATOM | 3676 | H | SER | A | 258 | 41.159 | 18.808 | 40.046 | 1.00 | 0.00 | H |
| ATOM | 3677 | CA | SER | A | 258 | 41.377 | 20.880 | 39.523 | 1.00 | 0.00 | C |
| ATOM | 3678 | HA | SER | A | 258 | 40.714 | 21.655 | 39.906 | 1.00 | 0.00 | H |
| ATOM | 3679 | CB | SER | A | 258 | 42.621 | 20.826 | 40.426 | 1.00 | 0.00 | C |
| ATOM | 3680 | HB1 | SER | A | 258 | 42.943 | 21.842 | 40.650 | 1.00 | 0.00 | H |
| ATOM | 3681 | HB2 | SER | A | 258 | 42.366 | 20.342 | 41.370 | 1.00 | 0.00 | H |
| ATOM | 3682 | OG | SER | A | 258 | 43.692 | 20.133 | 39.800 | 1.00 | 0.00 | O |
| ATOM | 3683 | HG | SER | A | 258 | 44.049 | 19.467 | 40.431 | 1.00 | 0.00 | H |
| ATOM | 3684 | C | SER | A | 258 | 41.768 | 21.286 | 38.084 | 1.00 | 0.00 | C |
| ATOM | 3685 | O | SER | A | 258 | 41.991 | 22.472 | 37.810 | 1.00 | 0.00 | O |
| ATOM | 3686 | N | LYS | A | 259 | 41.855 | 20.316 | 37.161 | 1.00 | 0.00 | N |
| ATOM | 3687 | H | LYS | A | 259 | 41.666 | 19.369 | 37.470 | 1.00 | 0.00 | H |
| ATOM | 3688 | CA | LYS | A | 259 | 42.186 | 20.510 | 35.743 | 1.00 | 0.00 | C |
| ATOM | 3689 | HA | LYS | A | 259 | 42.620 | 21.502 | 35.599 | 1.00 | 0.00 | H |
| ATOM | 3690 | CB | LYS | A | 259 | 43.231 | 19.458 | 35.311 | 1.00 | 0.00 | C |
| ATOM | 3691 | HB1 | LYS | A | 259 | 42.848 | 18.455 | 35.506 | 1.00 | 0.00 | H |
| ATOM | 3692 | HB2 | LYS | A | 259 | 43.385 | 19.551 | 34.235 | 1.00 | 0.00 | H |
| ATOM | 3693 | CG | LYS | A | 259 | 44.595 | 19.625 | 36.005 | 1.00 | 0.00 | C |
| ATOM | 3694 | HG1 | LYS | A | 259 | 44.880 | 20.675 | 35.975 | 1.00 | 0.00 | H |
| ATOM | 3695 | HG2 | LYS | A | 259 | 44.518 | 19.317 | 37.047 | 1.00 | 0.00 | H |
| ATOM | 3696 | CD | LYS | A | 259 | 45.680 | 18.796 | 35.300 | 1.00 | 0.00 | C |
| ATOM | 3697 | HD1 | LYS | A | 259 | 45.366 | 17.753 | 35.248 | 1.00 | 0.00 | H |
| ATOM | 3698 | HD2 | LYS | A | 259 | 45.797 | 19.171 | 34.283 | 1.00 | 0.00 | H |
| ATOM | 3699 | CE | LYS | A | 259 | 47.026 | 18.884 | 36.036 | 1.00 | 0.00 | C |
| ATOM | 3700 | HE1 | LYS | A | 259 | 47.302 | 19.933 | 36.153 | 1.00 | 0.00 | H |
| ATOM | 3701 | HE2 | LYS | A | 259 | 46.912 | 18.447 | 37.032 | 1.00 | 0.00 | H |
| ATOM | 3702 | NZ | LYS | A | 259 | 48.103 | 18.179 | 35.301 | 1.00 | 0.00 | N |
| ATOM | 3703 | HZ1 | LYS | A | 259 | 48.992 | 18.227 | 35.787 | 1.00 | 0.00 | H |
| ATOM | 3704 | HZ2 | LYS | A | 259 | 47.853 | 17.207 | 35.149 | 1.00 | 0.00 | H |
| ATOM | 3705 | HZ3 | LYS | A | 259 | 48.233 | 18.598 | 34.384 | 1.00 | 0.00 | H |
| ATOM | 3706 | C | LYS | A | 259 | 40.960 | 20.433 | 34.811 | 1.00 | 0.00 | C |
| ATOM | 3707 | O | LYS | A | 259 | 41.023 | 20.962 | 33.699 | 1.00 | 0.00 | O |
| ATOM | 3708 | N | LEU | A | 260 | 39.865 | 19.772 | 35.206 | 1.00 | 0.00 | N |
| ATOM | 3709 | H | LEU | A | 260 | 39.873 | 19.341 | 36.122 | 1.00 | 0.00 | H |
| ATOM | 3710 | CA | LEU | A | 260 | 38.677 | 19.588 | 34.362 | 1.00 | 0.00 | C |
| ATOM | 3711 | HA | LEU | A | 260 | 39.016 | 19.439 | 33.337 | 1.00 | 0.00 | H |
| ATOM | 3712 | CB | LEU | A | 260 | 37.931 | 18.313 | 34.807 | 1.00 | 0.00 | C |
| ATOM | 3713 | HB1 | LEU | A | 260 | 38.636 | 17.482 | 34.828 | 1.00 | 0.00 | H |
| ATOM | 3714 | HB2 | LEU | A | 260 | 37.574 | 18.462 | 35.827 | 1.00 | 0.00 | H |
| ATOM | 3715 | CG | LEU | A | 260 | 36.736 | 17.917 | 33.910 | 1.00 | 0.00 | C |
| ATOM | 3716 | HG | LEU | A | 260 | 36.006 | 18.727 | 33.907 | 1.00 | 0.00 | H |
| ATOM | 3717 | CD1 | LEU | A | 260 | 37.159 | 17.633 | 32.459 | 1.00 | 0.00 | C |
| ATOM | 3718 | HD11 | LEU | A | 260 | 36.307 | 17.254 | 31.895 | 1.00 | 0.00 | H |
| ATOM | 3719 | HD12 | LEU | A | 260 | 37.506 | 18.546 | 31.976 | 1.00 | 0.00 | H |
| ATOM | 3720 | HD13 | LEU | A | 260 | 37.952 | 16.885 | 32.436 | 1.00 | 0.00 | H |
| ATOM | 3721 | CD2 | LEU | A | 260 | 36.054 | 16.666 | 34.482 | 1.00 | 0.00 | C |
| ATOM | 3722 | HD21 | LEU | A | 260 | 35.192 | 16.398 | 33.870 | 1.00 | 0.00 | H |
| ATOM | 3723 | HD22 | LEU | A | 260 | 36.753 | 15.829 | 34.501 | 1.00 | 0.00 | H |

```
ATOM   3724 HD23 LEU A 260      35.707  16.869  35.494  1.00  0.00           H
ATOM   3725  C   LEU A 260      37.785  20.839  34.383  1.00  0.00           C
ATOM   3726  O   LEU A 260      37.375  21.304  35.446  1.00  0.00           O
ATOM   3727  N   ILE A 261      37.485  21.380  33.201  1.00  0.00           N
ATOM   3728  H   ILE A 261      37.847  20.936  32.363  1.00  0.00           H
ATOM   3729  CA  ILE A 261      36.707  22.610  32.985  1.00  0.00           C
ATOM   3730  HA  ILE A 261      36.381  23.015  33.944  1.00  0.00           H
ATOM   3731  CB  ILE A 261      37.576  23.671  32.258  1.00  0.00           C
ATOM   3732  HB  ILE A 261      37.688  23.357  31.218  1.00  0.00           H
ATOM   3733  CG2 ILE A 261      36.856  25.025  32.248  1.00  0.00           C
ATOM   3734 HG21 ILE A 261      37.419  25.738  31.647  1.00  0.00           H
ATOM   3735 HG22 ILE A 261      35.866  24.907  31.814  1.00  0.00           H
ATOM   3736 HG23 ILE A 261      36.755  25.415  33.258  1.00  0.00           H
ATOM   3737  CG1 ILE A 261      39.009  23.853  32.813  1.00  0.00           C
ATOM   3738 HG11 ILE A 261      39.561  22.927  32.666  1.00  0.00           H
ATOM   3739 HG12 ILE A 261      39.515  24.617  32.222  1.00  0.00           H
ATOM   3740  CD  ILE A 261      39.114  24.252  34.292  1.00  0.00           C
ATOM   3741  HD1 ILE A 261      40.162  24.238  34.591  1.00  0.00           H
ATOM   3742  HD2 ILE A 261      38.718  25.255  34.442  1.00  0.00           H
ATOM   3743  HD3 ILE A 261      38.568  23.551  34.918  1.00  0.00           H
ATOM   3744  C   ILE A 261      35.463  22.263  32.149  1.00  0.00           C
ATOM   3745  O   ILE A 261      35.587  21.663  31.078  1.00  0.00           O
ATOM   3746  N   LEU A 262      34.269  22.626  32.632  1.00  0.00           N
ATOM   3747  H   LEU A 262      34.239  23.174  33.482  1.00  0.00           H
ATOM   3748  CA  LEU A 262      32.989  22.259  32.009  1.00  0.00           C
ATOM   3749  HA  LEU A 262      33.109  21.290  31.522  1.00  0.00           H
ATOM   3750  CB  LEU A 262      31.927  22.121  33.121  1.00  0.00           C
ATOM   3751  HB1 LEU A 262      32.210  21.290  33.769  1.00  0.00           H
ATOM   3752  HB2 LEU A 262      31.933  23.033  33.720  1.00  0.00           H
ATOM   3753  CG  LEU A 262      30.483  21.893  32.619  1.00  0.00           C
ATOM   3754  HG  LEU A 262      30.170  22.756  32.031  1.00  0.00           H
ATOM   3755  CD1 LEU A 262      30.357  20.643  31.738  1.00  0.00           C
ATOM   3756 HD11 LEU A 262      29.335  20.572  31.368  1.00  0.00           H
ATOM   3757 HD12 LEU A 262      31.016  20.705  30.876  1.00  0.00           H
ATOM   3758 HD13 LEU A 262      30.604  19.750  32.313  1.00  0.00           H
ATOM   3759  CD2 LEU A 262      29.533  21.776  33.819  1.00  0.00           C
ATOM   3760 HD21 LEU A 262      28.506  21.669  33.468  1.00  0.00           H
ATOM   3761 HD22 LEU A 262      29.796  20.909  34.427  1.00  0.00           H
ATOM   3762 HD23 LEU A 262      29.597  22.675  34.431  1.00  0.00           H
ATOM   3763  C   LEU A 262      32.564  23.275  30.938  1.00  0.00           C
ATOM   3764  O   LEU A 262      32.408  24.454  31.252  1.00  0.00           O
ATOM   3765  N   GLY A 263      32.331  22.811  29.704  1.00  0.00           N
ATOM   3766  H   GLY A 263      32.516  21.836  29.512  1.00  0.00           H
ATOM   3767  CA  GLY A 263      31.902  23.628  28.563  1.00  0.00           C
ATOM   3768  HA1 GLY A 263      32.453  24.565  28.569  1.00  0.00           H
ATOM   3769  HA2 GLY A 263      32.128  23.102  27.637  1.00  0.00           H
ATOM   3770  C   GLY A 263      30.407  23.949  28.536  1.00  0.00           C
ATOM   3771  O   GLY A 263      29.560  23.075  28.741  1.00  0.00           O
ATOM   3772  N   MET A 264      30.097  25.204  28.209  1.00  0.00           N
ATOM   3773  H   MET A 264      30.867  25.861  28.123  1.00  0.00           H
ATOM   3774  CA  MET A 264      28.750  25.778  28.123  1.00  0.00           C
ATOM   3775  HA  MET A 264      28.011  25.032  28.403  1.00  0.00           H
ATOM   3776  CB  MET A 264      28.637  26.949  29.120  1.00  0.00           C
ATOM   3777  HB1 MET A 264      29.234  27.785  28.751  1.00  0.00           H
```

| | | | | | | | | | | | |
|---|---|---|---|---|---|---|---|---|---|---|---|
| ATOM | 3778 | HB2 | MET | A | 264 | 27.597 | 27.269 | 29.164 | 1.00 | 0.00 | H |
| ATOM | 3779 | CG | MET | A | 264 | 29.105 | 26.641 | 30.551 | 1.00 | 0.00 | C |
| ATOM | 3780 | HG1 | MET | A | 264 | 30.136 | 26.287 | 30.523 | 1.00 | 0.00 | H |
| ATOM | 3781 | HG2 | MET | A | 264 | 29.110 | 27.579 | 31.102 | 1.00 | 0.00 | H |
| ATOM | 3782 | SD | MET | A | 264 | 28.114 | 25.450 | 31.485 | 1.00 | 0.00 | S |
| ATOM | 3783 | CE | MET | A | 264 | 29.095 | 25.467 | 33.006 | 1.00 | 0.00 | C |
| ATOM | 3784 | HE1 | MET | A | 264 | 28.659 | 24.783 | 33.733 | 1.00 | 0.00 | H |
| ATOM | 3785 | HE2 | MET | A | 264 | 30.116 | 25.154 | 32.783 | 1.00 | 0.00 | H |
| ATOM | 3786 | HE3 | MET | A | 264 | 29.108 | 26.474 | 33.422 | 1.00 | 0.00 | H |
| ATOM | 3787 | C | MET | A | 264 | 28.485 | 26.293 | 26.688 | 1.00 | 0.00 | C |
| ATOM | 3788 | O | MET | A | 264 | 29.319 | 27.041 | 26.172 | 1.00 | 0.00 | O |
| ATOM | 3789 | N | PRO | A | 265 | 27.368 | 25.932 | 26.027 | 1.00 | 0.00 | N |
| ATOM | 3790 | CD | PRO | A | 265 | 26.469 | 24.846 | 26.397 | 1.00 | 0.00 | C |
| ATOM | 3791 | HD1 | PRO | A | 265 | 25.903 | 25.094 | 27.296 | 1.00 | 0.00 | H |
| ATOM | 3792 | HD2 | PRO | A | 265 | 27.045 | 23.933 | 26.550 | 1.00 | 0.00 | H |
| ATOM | 3793 | CG | PRO | A | 265 | 25.518 | 24.653 | 25.215 | 1.00 | 0.00 | C |
| ATOM | 3794 | HG1 | PRO | A | 265 | 24.589 | 25.187 | 25.405 | 1.00 | 0.00 | H |
| ATOM | 3795 | HG2 | PRO | A | 265 | 25.316 | 23.604 | 25.021 | 1.00 | 0.00 | H |
| ATOM | 3796 | CB | PRO | A | 265 | 26.253 | 25.296 | 24.041 | 1.00 | 0.00 | C |
| ATOM | 3797 | HB1 | PRO | A | 265 | 25.561 | 25.649 | 23.276 | 1.00 | 0.00 | H |
| ATOM | 3798 | HB2 | PRO | A | 265 | 26.958 | 24.578 | 23.619 | 1.00 | 0.00 | H |
| ATOM | 3799 | CA | PRO | A | 265 | 27.029 | 26.439 | 24.699 | 1.00 | 0.00 | C |
| ATOM | 3800 | HA | PRO | A | 265 | 27.928 | 26.642 | 24.116 | 1.00 | 0.00 | H |
| ATOM | 3801 | C | PRO | A | 265 | 26.172 | 27.711 | 24.790 | 1.00 | 0.00 | C |
| ATOM | 3802 | O | PRO | A | 265 | 25.090 | 27.689 | 25.375 | 1.00 | 0.00 | O |
| ATOM | 3803 | N | THR | A | 266 | 26.606 | 28.810 | 24.161 | 1.00 | 0.00 | N |
| ATOM | 3804 | H | THR | A | 266 | 27.514 | 28.796 | 23.704 | 1.00 | 0.00 | H |
| ATOM | 3805 | CA | THR | A | 266 | 25.804 | 30.046 | 24.022 | 1.00 | 0.00 | C |
| ATOM | 3806 | HA | THR | A | 266 | 25.117 | 30.106 | 24.864 | 1.00 | 0.00 | H |
| ATOM | 3807 | CB | THR | A | 266 | 26.688 | 31.300 | 24.089 | 1.00 | 0.00 | C |
| ATOM | 3808 | HB | THR | A | 266 | 26.107 | 32.178 | 23.798 | 1.00 | 0.00 | H |
| ATOM | 3809 | CG2 | THR | A | 266 | 27.219 | 31.508 | 25.506 | 1.00 | 0.00 | C |
| ATOM | 3810 | HG21 | THR | A | 266 | 27.932 | 32.330 | 25.521 | 1.00 | 0.00 | H |
| ATOM | 3811 | HG22 | THR | A | 266 | 26.399 | 31.741 | 26.180 | 1.00 | 0.00 | H |
| ATOM | 3812 | HG23 | THR | A | 266 | 27.711 | 30.602 | 25.851 | 1.00 | 0.00 | H |
| ATOM | 3813 | OG1 | THR | A | 266 | 27.807 | 31.166 | 23.252 | 1.00 | 0.00 | O |
| ATOM | 3814 | HG1 | THR | A | 266 | 28.180 | 32.048 | 23.118 | 1.00 | 0.00 | H |
| ATOM | 3815 | C | THR | A | 266 | 24.900 | 30.000 | 22.786 | 1.00 | 0.00 | C |
| ATOM | 3816 | O | THR | A | 266 | 24.733 | 30.979 | 22.068 | 1.00 | 0.00 | O |
| ATOM | 3817 | N | TYR | A | 267 | 24.304 | 28.835 | 22.526 | 1.00 | 0.00 | N |
| ATOM | 3818 | H | TYR | A | 267 | 24.470 | 28.073 | 23.169 | 1.00 | 0.00 | H |
| ATOM | 3819 | CA | TYR | A | 267 | 23.522 | 28.544 | 21.317 | 1.00 | 0.00 | C |
| ATOM | 3820 | HA | TYR | A | 267 | 22.906 | 29.416 | 21.088 | 1.00 | 0.00 | H |
| ATOM | 3821 | CB | TYR | A | 267 | 24.476 | 28.316 | 20.126 | 1.00 | 0.00 | C |
| ATOM | 3822 | HB1 | TYR | A | 267 | 23.890 | 28.173 | 19.219 | 1.00 | 0.00 | H |
| ATOM | 3823 | HB2 | TYR | A | 267 | 25.072 | 29.216 | 19.972 | 1.00 | 0.00 | H |
| ATOM | 3824 | CG | TYR | A | 267 | 25.410 | 27.131 | 20.285 | 1.00 | 0.00 | C |
| ATOM | 3825 | CD1 | TYR | A | 267 | 26.659 | 27.304 | 20.911 | 1.00 | 0.00 | C |
| ATOM | 3826 | HD1 | TYR | A | 267 | 26.957 | 28.284 | 21.260 | 1.00 | 0.00 | H |
| ATOM | 3827 | CE1 | TYR | A | 267 | 27.506 | 26.201 | 21.112 | 1.00 | 0.00 | C |
| ATOM | 3828 | HE1 | TYR | A | 267 | 28.446 | 26.322 | 21.630 | 1.00 | 0.00 | H |
| ATOM | 3829 | CZ | TYR | A | 267 | 27.114 | 24.922 | 20.672 | 1.00 | 0.00 | C |
| ATOM | 3830 | OH | TYR | A | 267 | 27.932 | 23.863 | 20.893 | 1.00 | 0.00 | O |
| ATOM | 3831 | HH | TYR | A | 267 | 27.575 | 23.044 | 20.544 | 1.00 | 0.00 | H |

| | | | | | | | | | | | |
|---|---|---|---|---|---|---|---|---|---|---|---|
| ATOM | 3832 | CE2 | TYR | A | 267 | 25.869 | 24.744 | 20.028 | 1.00 | 0.00 | C |
| ATOM | 3833 | HE2 | TYR | A | 267 | 25.567 | 23.761 | 19.697 | 1.00 | 0.00 | H |
| ATOM | 3834 | CD2 | TYR | A | 267 | 25.020 | 25.851 | 19.839 | 1.00 | 0.00 | C |
| ATOM | 3835 | HD2 | TYR | A | 267 | 24.059 | 25.717 | 19.363 | 1.00 | 0.00 | H |
| ATOM | 3836 | C | TYR | A | 267 | 22.567 | 27.349 | 21.495 | 1.00 | 0.00 | C |
| ATOM | 3837 | O | TYR | A | 267 | 22.716 | 26.529 | 22.412 | 1.00 | 0.00 | O |
| ATOM | 3838 | N | GLY | A | 268 | 21.601 | 27.241 | 20.577 | 1.00 | 0.00 | N |
| ATOM | 3839 | H | GLY | A | 268 | 21.584 | 27.944 | 19.841 | 1.00 | 0.00 | H |
| ATOM | 3840 | CA | GLY | A | 268 | 20.681 | 26.112 | 20.426 | 1.00 | 0.00 | C |
| ATOM | 3841 | HA1 | GLY | A | 268 | 20.813 | 25.396 | 21.236 | 1.00 | 0.00 | H |
| ATOM | 3842 | HA2 | GLY | A | 268 | 19.660 | 26.489 | 20.435 | 1.00 | 0.00 | H |
| ATOM | 3843 | C | GLY | A | 268 | 20.860 | 25.386 | 19.095 | 1.00 | 0.00 | C |
| ATOM | 3844 | O | GLY | A | 268 | 21.549 | 25.870 | 18.196 | 1.00 | 0.00 | O |
| ATOM | 3845 | N | ARG | A | 269 | 20.216 | 24.225 | 18.953 | 1.00 | 0.00 | N |
| ATOM | 3846 | H | ARG | A | 269 | 19.668 | 23.885 | 19.737 | 1.00 | 0.00 | H |
| ATOM | 3847 | CA | ARG | A | 269 | 20.222 | 23.432 | 17.718 | 1.00 | 0.00 | C |
| ATOM | 3848 | HA | ARG | A | 269 | 20.678 | 24.023 | 16.929 | 1.00 | 0.00 | H |
| ATOM | 3849 | CB | ARG | A | 269 | 21.063 | 22.166 | 17.892 | 1.00 | 0.00 | C |
| ATOM | 3850 | HB1 | ARG | A | 269 | 20.785 | 21.681 | 18.826 | 1.00 | 0.00 | H |
| ATOM | 3851 | HB2 | ARG | A | 269 | 20.851 | 21.493 | 17.060 | 1.00 | 0.00 | H |
| ATOM | 3852 | CG | ARG | A | 269 | 22.567 | 22.478 | 17.899 | 1.00 | 0.00 | C |
| ATOM | 3853 | HG1 | ARG | A | 269 | 22.798 | 23.180 | 17.097 | 1.00 | 0.00 | H |
| ATOM | 3854 | HG2 | ARG | A | 269 | 22.843 | 22.935 | 18.850 | 1.00 | 0.00 | H |
| ATOM | 3855 | CD | ARG | A | 269 | 23.388 | 21.205 | 17.679 | 1.00 | 0.00 | C |
| ATOM | 3856 | HD1 | ARG | A | 269 | 23.239 | 20.535 | 18.528 | 1.00 | 0.00 | H |
| ATOM | 3857 | HD2 | ARG | A | 269 | 23.041 | 20.697 | 16.779 | 1.00 | 0.00 | H |
| ATOM | 3858 | NE | ARG | A | 269 | 24.817 | 21.531 | 17.573 | 1.00 | 0.00 | N |
| ATOM | 3859 | HE | ARG | A | 269 | 25.284 | 21.705 | 18.452 | 1.00 | 0.00 | H |
| ATOM | 3860 | CZ | ARG | A | 269 | 25.566 | 21.602 | 16.478 | 1.00 | 0.00 | C |
| ATOM | 3861 | NH1 | ARG | A | 269 | 25.117 | 21.321 | 15.277 | 1.00 | 0.00 | N |
| ATOM | 3862 | HH11 | ARG | A | 269 | 24.206 | 20.895 | 15.168 | 1.00 | 0.00 | H |
| ATOM | 3863 | HH12 | ARG | A | 269 | 25.742 | 21.338 | 14.488 | 1.00 | 0.00 | H |
| ATOM | 3864 | NH2 | ARG | A | 269 | 26.822 | 21.955 | 16.577 | 1.00 | 0.00 | N |
| ATOM | 3865 | HH21 | ARG | A | 269 | 27.204 | 22.166 | 17.484 | 1.00 | 0.00 | H |
| ATOM | 3866 | HH22 | ARG | A | 269 | 27.446 | 21.840 | 15.797 | 1.00 | 0.00 | H |
| ATOM | 3867 | C | ARG | A | 269 | 18.812 | 23.095 | 17.237 | 1.00 | 0.00 | C |
| ATOM | 3868 | O | ARG | A | 269 | 17.945 | 22.710 | 18.019 | 1.00 | 0.00 | O |
| ATOM | 3869 | N | SER | A | 270 | 18.611 | 23.268 | 15.936 | 1.00 | 0.00 | N |
| ATOM | 3870 | H | SER | A | 270 | 19.421 | 23.493 | 15.371 | 1.00 | 0.00 | H |
| ATOM | 3871 | CA | SER | A | 270 | 17.327 | 23.468 | 15.252 | 1.00 | 0.00 | C |
| ATOM | 3872 | HA | SER | A | 270 | 16.522 | 23.563 | 15.982 | 1.00 | 0.00 | H |
| ATOM | 3873 | CB | SER | A | 270 | 17.427 | 24.782 | 14.455 | 1.00 | 0.00 | C |
| ATOM | 3874 | HB1 | SER | A | 270 | 16.603 | 24.843 | 13.746 | 1.00 | 0.00 | H |
| ATOM | 3875 | HB2 | SER | A | 270 | 17.361 | 25.623 | 15.145 | 1.00 | 0.00 | H |
| ATOM | 3876 | OG | SER | A | 270 | 18.671 | 24.847 | 13.766 | 1.00 | 0.00 | O |
| ATOM | 3877 | HG | SER | A | 270 | 18.671 | 25.585 | 13.119 | 1.00 | 0.00 | H |
| ATOM | 3878 | C | SER | A | 270 | 16.986 | 22.317 | 14.289 | 1.00 | 0.00 | C |
| ATOM | 3879 | O | SER | A | 270 | 17.841 | 21.842 | 13.538 | 1.00 | 0.00 | O |
| ATOM | 3880 | N | PHE | A | 271 | 15.732 | 21.852 | 14.300 | 1.00 | 0.00 | N |
| ATOM | 3881 | H | PHE | A | 271 | 15.070 | 22.280 | 14.940 | 1.00 | 0.00 | H |
| ATOM | 3882 | CA | PHE | A | 271 | 15.275 | 20.638 | 13.610 | 1.00 | 0.00 | C |
| ATOM | 3883 | HA | PHE | A | 271 | 15.993 | 20.338 | 12.847 | 1.00 | 0.00 | H |
| ATOM | 3884 | CB | PHE | A | 271 | 15.136 | 19.493 | 14.631 | 1.00 | 0.00 | C |
| ATOM | 3885 | HB1 | PHE | A | 271 | 14.495 | 19.824 | 15.450 | 1.00 | 0.00 | H |

```
ATOM   3886  HB2 PHE A 271      14.632  18.663  14.135  1.00  0.00           H
ATOM   3887  CG  PHE A 271      16.413  18.951  15.235  1.00  0.00           C
ATOM   3888  CD1 PHE A 271      16.832  17.640  14.933  1.00  0.00           C
ATOM   3889  HD1 PHE A 271      16.254  17.022  14.260  1.00  0.00           H
ATOM   3890  CE1 PHE A 271      17.994  17.119  15.527  1.00  0.00           C
ATOM   3891  HE1 PHE A 271      18.290  16.099  15.319  1.00  0.00           H
ATOM   3892  CZ  PHE A 271      18.760  17.918  16.393  1.00  0.00           C
ATOM   3893  HZ  PHE A 271      19.646  17.518  16.862  1.00  0.00           H
ATOM   3894  CE2 PHE A 271      18.352  19.230  16.680  1.00  0.00           C
ATOM   3895  HE2 PHE A 271      18.928  19.846  17.354  1.00  0.00           H
ATOM   3896  CD2 PHE A 271      17.160  19.729  16.139  1.00  0.00           C
ATOM   3897  HD2 PHE A 271      16.817  20.709  16.434  1.00  0.00           H
ATOM   3898  C   PHE A 271      13.913  20.868  12.936  1.00  0.00           C
ATOM   3899  O   PHE A 271      13.036  21.514  13.521  1.00  0.00           O
ATOM   3900  N   THR A 272      13.703  20.274  11.752  1.00  0.00           N
ATOM   3901  H   THR A 272      14.439  19.685  11.375  1.00  0.00           H
ATOM   3902  CA  THR A 272      12.401  20.266  11.064  1.00  0.00           C
ATOM   3903  HA  THR A 272      11.827  21.137  11.381  1.00  0.00           H
ATOM   3904  CB  THR A 272      12.561  20.359   9.543  1.00  0.00           C
ATOM   3905  HB  THR A 272      13.143  19.514   9.168  1.00  0.00           H
ATOM   3906  CG2 THR A 272      11.188  20.375   8.872  1.00  0.00           C
ATOM   3907 HG21 THR A 272      11.290  20.585   7.812  1.00  0.00           H
ATOM   3908 HG22 THR A 272      10.704  19.405   8.976  1.00  0.00           H
ATOM   3909 HG23 THR A 272      10.562  21.137   9.333  1.00  0.00           H
ATOM   3910  OG1 THR A 272      13.200  21.573   9.205  1.00  0.00           O
ATOM   3911  HG1 THR A 272      14.163  21.425   9.264  1.00  0.00           H
ATOM   3912  C   THR A 272      11.614  19.011  11.443  1.00  0.00           C
ATOM   3913  O   THR A 272      12.089  17.893  11.244  1.00  0.00           O
ATOM   3914  N   LEU A 273      10.391  19.209  11.939  1.00  0.00           N
ATOM   3915  H   LEU A 273      10.084  20.168  12.042  1.00  0.00           H
ATOM   3916  CA  LEU A 273       9.438  18.166  12.326  1.00  0.00           C
ATOM   3917  HA  LEU A 273       9.970  17.385  12.870  1.00  0.00           H
ATOM   3918  CB  LEU A 273       8.349  18.781  13.231  1.00  0.00           C
ATOM   3919  HB1 LEU A 273       7.917  19.638  12.712  1.00  0.00           H
ATOM   3920  HB2 LEU A 273       7.547  18.055  13.364  1.00  0.00           H
ATOM   3921  CG  LEU A 273       8.824  19.224  14.627  1.00  0.00           C
ATOM   3922  HG  LEU A 273       9.678  19.892  14.520  1.00  0.00           H
ATOM   3923  CD1 LEU A 273       7.710  20.018  15.322  1.00  0.00           C
ATOM   3924 HD11 LEU A 273       8.047  20.332  16.309  1.00  0.00           H
ATOM   3925 HD12 LEU A 273       7.468  20.907  14.740  1.00  0.00           H
ATOM   3926 HD13 LEU A 273       6.815  19.403  15.423  1.00  0.00           H
ATOM   3927  CD2 LEU A 273       9.225  18.019  15.496  1.00  0.00           C
ATOM   3928 HD21 LEU A 273       9.494  18.354  16.499  1.00  0.00           H
ATOM   3929 HD22 LEU A 273       8.401  17.315  15.561  1.00  0.00           H
ATOM   3930 HD23 LEU A 273      10.073  17.491  15.065  1.00  0.00           H
ATOM   3931  C   LEU A 273       8.770  17.525  11.106  1.00  0.00           C
ATOM   3932  O   LEU A 273       8.454  18.204  10.126  1.00  0.00           O
ATOM   3933  N   ALA A 274       8.454  16.233  11.208  1.00  0.00           N
ATOM   3934  H   ALA A 274       8.713  15.733  12.050  1.00  0.00           H
ATOM   3935  CA  ALA A 274       7.618  15.539  10.225  1.00  0.00           C
ATOM   3936  HA  ALA A 274       8.037  15.696   9.231  1.00  0.00           H
ATOM   3937  CB  ALA A 274       7.656  14.043  10.554  1.00  0.00           C
ATOM   3938  HB1 ALA A 274       7.096  13.487   9.800  1.00  0.00           H
ATOM   3939  HB2 ALA A 274       8.683  13.679  10.572  1.00  0.00           H
```

| | | | | | | | | | | | |
|---|---|---|---|---|---|---|---|---|---|---|---|
| ATOM | 3940 | HB3 | ALA | A | 274 | 7.200 | 13.871 | 11.528 | 1.00 | 0.00 | H |
| ATOM | 3941 | C | ALA | A | 274 | 6.171 | 16.070 | 10.195 | 1.00 | 0.00 | C |
| ATOM | 3942 | O | ALA | A | 274 | 5.535 | 16.087 | 9.138 | 1.00 | 0.00 | O |
| ATOM | 3943 | N | SER | A | 275 | 5.655 | 16.540 | 11.330 | 1.00 | 0.00 | N |
| ATOM | 3944 | H | SER | A | 275 | 6.203 | 16.453 | 12.182 | 1.00 | 0.00 | H |
| ATOM | 3945 | CA | SER | A | 275 | 4.310 | 17.103 | 11.466 | 1.00 | 0.00 | C |
| ATOM | 3946 | HA | SER | A | 275 | 4.050 | 17.638 | 10.552 | 1.00 | 0.00 | H |
| ATOM | 3947 | CB | SER | A | 275 | 3.310 | 15.952 | 11.684 | 1.00 | 0.00 | C |
| ATOM | 3948 | HB1 | SER | A | 275 | 3.756 | 15.010 | 11.358 | 1.00 | 0.00 | H |
| ATOM | 3949 | HB2 | SER | A | 275 | 3.081 | 15.862 | 12.748 | 1.00 | 0.00 | H |
| ATOM | 3950 | OG | SER | A | 275 | 2.110 | 16.137 | 10.950 | 1.00 | 0.00 | O |
| ATOM | 3951 | HG | SER | A | 275 | 2.325 | 16.042 | 9.993 | 1.00 | 0.00 | H |
| ATOM | 3952 | C | SER | A | 275 | 4.264 | 18.100 | 12.638 | 1.00 | 0.00 | C |
| ATOM | 3953 | O | SER | A | 275 | 4.833 | 17.838 | 13.697 | 1.00 | 0.00 | O |
| ATOM | 3954 | N | SER | A | 276 | 3.572 | 19.235 | 12.498 | 1.00 | 0.00 | N |
| ATOM | 3955 | H | SER | A | 276 | 3.026 | 19.396 | 11.652 | 1.00 | 0.00 | H |
| ATOM | 3956 | CA | SER | A | 276 | 3.480 | 20.266 | 13.551 | 1.00 | 0.00 | C |
| ATOM | 3957 | HA | SER | A | 276 | 4.466 | 20.388 | 14.004 | 1.00 | 0.00 | H |
| ATOM | 3958 | CB | SER | A | 276 | 3.090 | 21.617 | 12.924 | 1.00 | 0.00 | C |
| ATOM | 3959 | HB1 | SER | A | 276 | 3.159 | 22.400 | 13.680 | 1.00 | 0.00 | H |
| ATOM | 3960 | HB2 | SER | A | 276 | 3.797 | 21.852 | 12.126 | 1.00 | 0.00 | H |
| ATOM | 3961 | OG | SER | A | 276 | 1.775 | 21.600 | 12.389 | 1.00 | 0.00 | O |
| ATOM | 3962 | HG | SER | A | 276 | 1.150 | 21.750 | 13.131 | 1.00 | 0.00 | H |
| ATOM | 3963 | C | SER | A | 276 | 2.525 | 19.876 | 14.702 | 1.00 | 0.00 | C |
| ATOM | 3964 | O | SER | A | 276 | 2.249 | 20.674 | 15.597 | 1.00 | 0.00 | O |
| ATOM | 3965 | N | SER | A | 277 | 1.983 | 18.661 | 14.666 | 1.00 | 0.00 | N |
| ATOM | 3966 | H | SER | A | 277 | 2.197 | 18.081 | 13.866 | 1.00 | 0.00 | H |
| ATOM | 3967 | CA | SER | A | 277 | 1.154 | 18.010 | 15.687 | 1.00 | 0.00 | C |
| ATOM | 3968 | HA | SER | A | 277 | 0.708 | 18.763 | 16.338 | 1.00 | 0.00 | H |
| ATOM | 3969 | CB | SER | A | 277 | 0.021 | 17.260 | 14.966 | 1.00 | 0.00 | C |
| ATOM | 3970 | HB1 | SER | A | 277 | -0.463 | 16.560 | 15.649 | 1.00 | 0.00 | H |
| ATOM | 3971 | HB2 | SER | A | 277 | -0.722 | 17.992 | 14.646 | 1.00 | 0.00 | H |
| ATOM | 3972 | OG | SER | A | 277 | 0.491 | 16.569 | 13.813 | 1.00 | 0.00 | O |
| ATOM | 3973 | HG | SER | A | 277 | -0.111 | 16.787 | 13.067 | 1.00 | 0.00 | H |
| ATOM | 3974 | C | SER | A | 277 | 1.952 | 17.063 | 16.610 | 1.00 | 0.00 | C |
| ATOM | 3975 | O | SER | A | 277 | 1.371 | 16.434 | 17.500 | 1.00 | 0.00 | O |
| ATOM | 3976 | N | ASP | A | 278 | 3.279 | 16.972 | 16.452 | 1.00 | 0.00 | N |
| ATOM | 3977 | H | ASP | A | 278 | 3.717 | 17.471 | 15.686 | 1.00 | 0.00 | H |
| ATOM | 3978 | CA | ASP | A | 278 | 4.153 | 16.300 | 17.417 | 1.00 | 0.00 | C |
| ATOM | 3979 | HA | ASP | A | 278 | 3.612 | 16.223 | 18.356 | 1.00 | 0.00 | H |
| ATOM | 3980 | CB | ASP | A | 278 | 4.475 | 14.872 | 16.945 | 1.00 | 0.00 | C |
| ATOM | 3981 | HB1 | ASP | A | 278 | 3.555 | 14.395 | 16.603 | 1.00 | 0.00 | H |
| ATOM | 3982 | HB2 | ASP | A | 278 | 5.167 | 14.910 | 16.104 | 1.00 | 0.00 | H |
| ATOM | 3983 | CG | ASP | A | 278 | 5.062 | 14.004 | 18.062 | 1.00 | 0.00 | C |
| ATOM | 3984 | OD1 | ASP | A | 278 | 4.717 | 14.237 | 19.241 | 1.00 | 0.00 | O |
| ATOM | 3985 | OD2 | ASP | A | 278 | 5.806 | 13.048 | 17.737 | 1.00 | 0.00 | O |
| ATOM | 3986 | C | ASP | A | 278 | 5.407 | 17.130 | 17.714 | 1.00 | 0.00 | C |
| ATOM | 3987 | O | ASP | A | 278 | 6.339 | 17.184 | 16.913 | 1.00 | 0.00 | O |
| ATOM | 3988 | N | THR | A | 279 | 5.388 | 17.795 | 18.877 | 1.00 | 0.00 | N |
| ATOM | 3989 | H | THR | A | 279 | 4.538 | 17.702 | 19.423 | 1.00 | 0.00 | H |
| ATOM | 3990 | CA | THR | A | 279 | 6.366 | 18.798 | 19.345 | 1.00 | 0.00 | C |
| ATOM | 3991 | HA | THR | A | 279 | 7.131 | 18.920 | 18.580 | 1.00 | 0.00 | H |
| ATOM | 3992 | CB | THR | A | 279 | 5.674 | 20.158 | 19.551 | 1.00 | 0.00 | C |
| ATOM | 3993 | HB | THR | A | 279 | 6.375 | 20.854 | 20.012 | 1.00 | 0.00 | H |

| | | | | | | | | | | | |
|---|---|---|---|---|---|---|---|---|---|---|---|
| ATOM | 3994 | CG2 | THR | A | 279 | 5.218 | 20.764 | 18.225 | 1.00 | 0.00 | C |
| ATOM | 3995 | HG21 | THR | A | 279 | 4.830 | 21.766 | 18.403 | 1.00 | 0.00 | H |
| ATOM | 3996 | HG22 | THR | A | 279 | 6.065 | 20.845 | 17.547 | 1.00 | 0.00 | H |
| ATOM | 3997 | HG23 | THR | A | 279 | 4.443 | 20.149 | 17.765 | 1.00 | 0.00 | H |
| ATOM | 3998 | OG1 | THR | A | 279 | 4.535 | 20.025 | 20.376 | 1.00 | 0.00 | O |
| ATOM | 3999 | HG1 | THR | A | 279 | 4.022 | 20.846 | 20.286 | 1.00 | 0.00 | H |
| ATOM | 4000 | C | THR | A | 279 | 7.112 | 18.382 | 20.618 | 1.00 | 0.00 | C |
| ATOM | 4001 | O | THR | A | 279 | 7.872 | 19.177 | 21.171 | 1.00 | 0.00 | O |
| ATOM | 4002 | N | ARG | A | 280 | 6.936 | 17.143 | 21.089 | 1.00 | 0.00 | N |
| ATOM | 4003 | H | ARG | A | 280 | 6.316 | 16.528 | 20.581 | 1.00 | 0.00 | H |
| ATOM | 4004 | CA | ARG | A | 280 | 7.640 | 16.609 | 22.265 | 1.00 | 0.00 | C |
| ATOM | 4005 | HA | ARG | A | 280 | 7.523 | 17.330 | 23.075 | 1.00 | 0.00 | H |
| ATOM | 4006 | CB | ARG | A | 280 | 6.965 | 15.299 | 22.717 | 1.00 | 0.00 | C |
| ATOM | 4007 | HB1 | ARG | A | 280 | 7.380 | 15.009 | 23.682 | 1.00 | 0.00 | H |
| ATOM | 4008 | HB2 | ARG | A | 280 | 5.898 | 15.480 | 22.858 | 1.00 | 0.00 | H |
| ATOM | 4009 | CG | ARG | A | 280 | 7.152 | 14.132 | 21.734 | 1.00 | 0.00 | C |
| ATOM | 4010 | HG1 | ARG | A | 280 | 6.660 | 14.365 | 20.795 | 1.00 | 0.00 | H |
| ATOM | 4011 | HG2 | ARG | A | 280 | 8.209 | 13.971 | 21.535 | 1.00 | 0.00 | H |
| ATOM | 4012 | CD | ARG | A | 280 | 6.554 | 12.849 | 22.313 | 1.00 | 0.00 | C |
| ATOM | 4013 | HD1 | ARG | A | 280 | 6.937 | 12.692 | 23.322 | 1.00 | 0.00 | H |
| ATOM | 4014 | HD2 | ARG | A | 280 | 5.472 | 12.969 | 22.377 | 1.00 | 0.00 | H |
| ATOM | 4015 | NE | ARG | A | 280 | 6.850 | 11.679 | 21.473 | 1.00 | 0.00 | N |
| ATOM | 4016 | HE | ARG | A | 280 | 6.275 | 11.560 | 20.645 | 1.00 | 0.00 | H |
| ATOM | 4017 | CZ | ARG | A | 280 | 7.701 | 10.697 | 21.740 | 1.00 | 0.00 | C |
| ATOM | 4018 | NH1 | ARG | A | 280 | 8.613 | 10.789 | 22.680 | 1.00 | 0.00 | N |
| ATOM | 4019 | HH11 | ARG | A | 280 | 8.764 | 11.676 | 23.136 | 1.00 | 0.00 | H |
| ATOM | 4020 | HH12 | ARG | A | 280 | 9.278 | 10.038 | 22.837 | 1.00 | 0.00 | H |
| ATOM | 4021 | NH2 | ARG | A | 280 | 7.626 | 9.585 | 21.051 | 1.00 | 0.00 | N |
| ATOM | 4022 | HH21 | ARG | A | 280 | 6.918 | 9.503 | 20.334 | 1.00 | 0.00 | H |
| ATOM | 4023 | HH22 | ARG | A | 280 | 8.121 | 8.759 | 21.363 | 1.00 | 0.00 | H |
| ATOM | 4024 | C | ARG | A | 280 | 9.152 | 16.415 | 22.036 | 1.00 | 0.00 | C |
| ATOM | 4025 | O | ARG | A | 280 | 9.651 | 16.494 | 20.913 | 1.00 | 0.00 | O |
| ATOM | 4026 | N | VAL | A | 281 | 9.876 | 16.069 | 23.099 | 1.00 | 0.00 | N |
| ATOM | 4027 | H | VAL | A | 281 | 9.403 | 16.079 | 24.002 | 1.00 | 0.00 | H |
| ATOM | 4028 | CA | VAL | A | 281 | 11.182 | 15.393 | 23.026 | 1.00 | 0.00 | C |
| ATOM | 4029 | HA | VAL | A | 281 | 11.795 | 15.897 | 22.278 | 1.00 | 0.00 | H |
| ATOM | 4030 | CB | VAL | A | 281 | 11.922 | 15.460 | 24.383 | 1.00 | 0.00 | C |
| ATOM | 4031 | HB | VAL | A | 281 | 11.300 | 14.985 | 25.142 | 1.00 | 0.00 | H |
| ATOM | 4032 | CG1 | VAL | A | 281 | 13.269 | 14.728 | 24.327 | 1.00 | 0.00 | C |
| ATOM | 4033 | HG11 | VAL | A | 281 | 13.823 | 14.901 | 25.249 | 1.00 | 0.00 | H |
| ATOM | 4034 | HG12 | VAL | A | 281 | 13.105 | 13.657 | 24.221 | 1.00 | 0.00 | H |
| ATOM | 4035 | HG13 | VAL | A | 281 | 13.859 | 15.093 | 23.486 | 1.00 | 0.00 | H |
| ATOM | 4036 | CG2 | VAL | A | 281 | 12.175 | 16.913 | 24.826 | 1.00 | 0.00 | C |
| ATOM | 4037 | HG21 | VAL | A | 281 | 12.613 | 16.914 | 25.824 | 1.00 | 0.00 | H |
| ATOM | 4038 | HG22 | VAL | A | 281 | 12.862 | 17.403 | 24.136 | 1.00 | 0.00 | H |
| ATOM | 4039 | HG23 | VAL | A | 281 | 11.244 | 17.476 | 24.868 | 1.00 | 0.00 | H |
| ATOM | 4040 | C | VAL | A | 281 | 10.949 | 13.933 | 22.599 | 1.00 | 0.00 | C |
| ATOM | 4041 | O | VAL | A | 281 | 10.057 | 13.271 | 23.133 | 1.00 | 0.00 | O |
| ATOM | 4042 | N | GLY | A | 282 | 11.713 | 13.448 | 21.612 | 1.00 | 0.00 | N |
| ATOM | 4043 | H | GLY | A | 282 | 12.436 | 14.050 | 21.226 | 1.00 | 0.00 | H |
| ATOM | 4044 | CA | GLY | A | 282 | 11.540 | 12.134 | 20.970 | 1.00 | 0.00 | C |
| ATOM | 4045 | HA1 | GLY | A | 282 | 12.510 | 11.803 | 20.599 | 1.00 | 0.00 | H |
| ATOM | 4046 | HA2 | GLY | A | 282 | 11.170 | 11.411 | 21.697 | 1.00 | 0.00 | H |
| ATOM | 4047 | C | GLY | A | 282 | 10.577 | 12.134 | 19.772 | 1.00 | 0.00 | C |

| | | | | | | | | | | | |
|---|---|---|---|---|---|---|---|---|---|---|---|
| ATOM | 4048 | O | GLY | A | 282 | 10.267 | 11.068 | 19.237 | 1.00 | 0.00 | O |
| ATOM | 4049 | N | ALA | A | 283 | 10.102 | 13.307 | 19.341 | 1.00 | 0.00 | N |
| ATOM | 4050 | H | ALA | A | 283 | 10.492 | 14.144 | 19.757 | 1.00 | 0.00 | H |
| ATOM | 4051 | CA | ALA | A | 283 | 9.229 | 13.482 | 18.178 | 1.00 | 0.00 | C |
| ATOM | 4052 | HA | ALA | A | 283 | 8.406 | 12.773 | 18.267 | 1.00 | 0.00 | H |
| ATOM | 4053 | CB | ALA | A | 283 | 8.637 | 14.898 | 18.205 | 1.00 | 0.00 | C |
| ATOM | 4054 | HB1 | ALA | A | 283 | 7.952 | 15.024 | 17.366 | 1.00 | 0.00 | H |
| ATOM | 4055 | HB2 | ALA | A | 283 | 8.085 | 15.060 | 19.127 | 1.00 | 0.00 | H |
| ATOM | 4056 | HB3 | ALA | A | 283 | 9.431 | 15.642 | 18.125 | 1.00 | 0.00 | H |
| ATOM | 4057 | C | ALA | A | 283 | 9.995 | 13.249 | 16.855 | 1.00 | 0.00 | C |
| ATOM | 4058 | O | ALA | A | 283 | 11.160 | 13.648 | 16.771 | 1.00 | 0.00 | O |
| ATOM | 4059 | N | PRO | A | 284 | 9.370 | 12.656 | 15.815 | 1.00 | 0.00 | N |
| ATOM | 4060 | CD | PRO | A | 284 | 8.094 | 11.957 | 15.857 | 1.00 | 0.00 | C |
| ATOM | 4061 | HD1 | PRO | A | 284 | 7.290 | 12.681 | 15.720 | 1.00 | 0.00 | H |
| ATOM | 4062 | HD2 | PRO | A | 284 | 7.956 | 11.400 | 16.783 | 1.00 | 0.00 | H |
| ATOM | 4063 | CG | PRO | A | 284 | 8.115 | 10.984 | 14.682 | 1.00 | 0.00 | C |
| ATOM | 4064 | HG1 | PRO | A | 284 | 7.111 | 10.744 | 14.332 | 1.00 | 0.00 | H |
| ATOM | 4065 | HG2 | PRO | A | 284 | 8.655 | 10.077 | 14.960 | 1.00 | 0.00 | H |
| ATOM | 4066 | CB | PRO | A | 284 | 8.911 | 11.766 | 13.645 | 1.00 | 0.00 | C |
| ATOM | 4067 | HB1 | PRO | A | 284 | 8.252 | 12.492 | 13.171 | 1.00 | 0.00 | H |
| ATOM | 4068 | HB2 | PRO | A | 284 | 9.355 | 11.116 | 12.894 | 1.00 | 0.00 | H |
| ATOM | 4069 | CA | PRO | A | 284 | 9.969 | 12.483 | 14.493 | 1.00 | 0.00 | C |
| ATOM | 4070 | HA | PRO | A | 284 | 10.836 | 11.832 | 14.602 | 1.00 | 0.00 | H |
| ATOM | 4071 | C | PRO | A | 284 | 10.394 | 13.799 | 13.831 | 1.00 | 0.00 | C |
| ATOM | 4072 | O | PRO | A | 284 | 9.594 | 14.729 | 13.686 | 1.00 | 0.00 | O |
| ATOM | 4073 | N | ALA | A | 285 | 11.648 | 13.831 | 13.368 | 1.00 | 0.00 | N |
| ATOM | 4074 | H | ALA | A | 285 | 12.244 | 13.033 | 13.557 | 1.00 | 0.00 | H |
| ATOM | 4075 | CA | ALA | A | 285 | 12.221 | 14.907 | 12.559 | 1.00 | 0.00 | C |
| ATOM | 4076 | HA | ALA | A | 285 | 11.515 | 15.734 | 12.496 | 1.00 | 0.00 | H |
| ATOM | 4077 | CB | ALA | A | 285 | 13.487 | 15.419 | 13.262 | 1.00 | 0.00 | C |
| ATOM | 4078 | HB1 | ALA | A | 285 | 13.905 | 16.259 | 12.705 | 1.00 | 0.00 | H |
| ATOM | 4079 | HB2 | ALA | A | 285 | 13.234 | 15.763 | 14.267 | 1.00 | 0.00 | H |
| ATOM | 4080 | HB3 | ALA | A | 285 | 14.233 | 14.626 | 13.329 | 1.00 | 0.00 | H |
| ATOM | 4081 | C | ALA | A | 285 | 12.523 | 14.420 | 11.129 | 1.00 | 0.00 | C |
| ATOM | 4082 | O | ALA | A | 285 | 12.938 | 13.269 | 10.931 | 1.00 | 0.00 | O |
| ATOM | 4083 | N | THR | A | 286 | 12.343 | 15.292 | 10.129 | 1.00 | 0.00 | N |
| ATOM | 4084 | H | THR | A | 286 | 12.003 | 16.222 | 10.362 | 1.00 | 0.00 | H |
| ATOM | 4085 | CA | THR | A | 286 | 12.702 | 15.039 | 8.717 | 1.00 | 0.00 | C |
| ATOM | 4086 | HA | THR | A | 286 | 12.788 | 13.965 | 8.561 | 1.00 | 0.00 | H |
| ATOM | 4087 | CB | THR | A | 286 | 11.619 | 15.542 | 7.756 | 1.00 | 0.00 | C |
| ATOM | 4088 | HB | THR | A | 286 | 11.987 | 15.499 | 6.728 | 1.00 | 0.00 | H |
| ATOM | 4089 | CG2 | THR | A | 286 | 10.384 | 14.652 | 7.850 | 1.00 | 0.00 | C |
| ATOM | 4090 | HG21 | THR | A | 286 | 9.612 | 15.025 | 7.179 | 1.00 | 0.00 | H |
| ATOM | 4091 | HG22 | THR | A | 286 | 10.636 | 13.634 | 7.552 | 1.00 | 0.00 | H |
| ATOM | 4092 | HG23 | THR | A | 286 | 10.007 | 14.641 | 8.869 | 1.00 | 0.00 | H |
| ATOM | 4093 | OG1 | THR | A | 286 | 11.253 | 16.866 | 8.063 | 1.00 | 0.00 | O |
| ATOM | 4094 | HG1 | THR | A | 286 | 11.903 | 17.457 | 7.633 | 1.00 | 0.00 | H |
| ATOM | 4095 | C | THR | A | 286 | 14.061 | 15.611 | 8.334 | 1.00 | 0.00 | C |
| ATOM | 4096 | O | THR | A | 286 | 14.545 | 15.290 | 7.248 | 1.00 | 0.00 | O |
| ATOM | 4097 | N | GLY | A | 287 | 14.707 | 16.382 | 9.212 | 1.00 | 0.00 | N |
| ATOM | 4098 | H | GLY | A | 287 | 14.236 | 16.645 | 10.066 | 1.00 | 0.00 | H |
| ATOM | 4099 | CA | GLY | A | 287 | 16.091 | 16.830 | 9.044 | 1.00 | 0.00 | C |
| ATOM | 4100 | HA1 | GLY | A | 287 | 16.765 | 16.040 | 9.377 | 1.00 | 0.00 | H |
| ATOM | 4101 | HA2 | GLY | A | 287 | 16.291 | 16.998 | 7.989 | 1.00 | 0.00 | H |

```
ATOM   4102  C   GLY A 287      16.424  18.088   9.837  1.00  0.00           C
ATOM   4103  O   GLY A 287      15.789  18.390  10.849  1.00  0.00           O
ATOM   4104  N   SER A 288      17.430  18.832   9.383  1.00  0.00           N
ATOM   4105  H   SER A 288      17.845  18.607   8.483  1.00  0.00           H
ATOM   4106  CA  SER A 288      17.855  20.078  10.028  1.00  0.00           C
ATOM   4107  HA  SER A 288      18.005  19.875  11.088  1.00  0.00           H
ATOM   4108  CB  SER A 288      19.207  20.547   9.458  1.00  0.00           C
ATOM   4109  HB1 SER A 288      19.470  21.515   9.887  1.00  0.00           H
ATOM   4110  HB2 SER A 288      19.972  19.830   9.751  1.00  0.00           H
ATOM   4111  OG  SER A 288      19.211  20.636   8.041  1.00  0.00           O
ATOM   4112  HG  SER A 288      19.117  21.578   7.787  1.00  0.00           H
ATOM   4113  C   SER A 288      16.787  21.182   9.938  1.00  0.00           C
ATOM   4114  O   SER A 288      15.895  21.156   9.080  1.00  0.00           O
ATOM   4115  N   GLY A 289      16.864  22.162  10.841  1.00  0.00           N
ATOM   4116  H   GLY A 289      17.563  22.110  11.575  1.00  0.00           H
ATOM   4117  CA  GLY A 289      15.981  23.327  10.851  1.00  0.00           C
ATOM   4118  HA1 GLY A 289      14.957  23.012  10.651  1.00  0.00           H
ATOM   4119  HA2 GLY A 289      16.013  23.810  11.826  1.00  0.00           H
ATOM   4120  C   GLY A 289      16.363  24.373   9.803  1.00  0.00           C
ATOM   4121  O   GLY A 289      17.439  24.315   9.194  1.00  0.00           O
ATOM   4122  N   THR A 290      15.471  25.344   9.607  1.00  0.00           N
ATOM   4123  H   THR A 290      14.594  25.287  10.116  1.00  0.00           H
ATOM   4124  CA  THR A 290      15.679  26.514   8.746  1.00  0.00           C
ATOM   4125  HA  THR A 290      15.871  26.154   7.740  1.00  0.00           H
ATOM   4126  CB  THR A 290      14.395  27.348   8.707  1.00  0.00           C
ATOM   4127  HB  THR A 290      14.146  27.688   9.714  1.00  0.00           H
ATOM   4128  CG2 THR A 290      14.488  28.554   7.777  1.00  0.00           C
ATOM   4129 HG21 THR A 290      13.504  29.012   7.686  1.00  0.00           H
ATOM   4130 HG22 THR A 290      15.181  29.287   8.184  1.00  0.00           H
ATOM   4131 HG23 THR A 290      14.826  28.250   6.788  1.00  0.00           H
ATOM   4132  OG1 THR A 290      13.358  26.528   8.226  1.00  0.00           O
ATOM   4133  HG1 THR A 290      13.587  26.261   7.318  1.00  0.00           H
ATOM   4134  C   THR A 290      16.876  27.336   9.255  1.00  0.00           C
ATOM   4135  O   THR A 290      16.996  27.517  10.470  1.00  0.00           O
ATOM   4136  N   PRO A 291      17.775  27.829   8.377  1.00  0.00           N
ATOM   4137  CD  PRO A 291      17.859  27.539   6.953  1.00  0.00           C
ATOM   4138  HD1 PRO A 291      17.179  28.193   6.407  1.00  0.00           H
ATOM   4139  HD2 PRO A 291      17.643  26.494   6.732  1.00  0.00           H
ATOM   4140  CG  PRO A 291      19.299  27.846   6.559  1.00  0.00           C
ATOM   4141  HG1 PRO A 291      19.367  28.150   5.515  1.00  0.00           H
ATOM   4142  HG2 PRO A 291      19.926  26.975   6.754  1.00  0.00           H
ATOM   4143  CB  PRO A 291      19.693  28.975   7.506  1.00  0.00           C
ATOM   4144  HB1 PRO A 291      19.370  29.928   7.085  1.00  0.00           H
ATOM   4145  HB2 PRO A 291      20.767  28.987   7.691  1.00  0.00           H
ATOM   4146  CA  PRO A 291      18.908  28.658   8.783  1.00  0.00           C
ATOM   4147  HA  PRO A 291      19.544  28.068   9.445  1.00  0.00           H
ATOM   4148  C   PRO A 291      18.481  29.942   9.501  1.00  0.00           C
ATOM   4149  O   PRO A 291      17.429  30.519   9.210  1.00  0.00           O
ATOM   4150  N   GLY A 292      19.340  30.408  10.408  1.00  0.00           N
ATOM   4151  H   GLY A 292      20.191  29.882  10.583  1.00  0.00           H
ATOM   4152  CA  GLY A 292      19.185  31.656  11.151  1.00  0.00           C
ATOM   4153  HA1 GLY A 292      18.151  31.742  11.484  1.00  0.00           H
ATOM   4154  HA2 GLY A 292      19.843  31.648  12.022  1.00  0.00           H
ATOM   4155  C   GLY A 292      19.550  32.888  10.314  1.00  0.00           C
```

| | | | | | | | | | | | |
|---|---|---|---|---|---|---|---|---|---|---|---|
| ATOM | 4156 | O | GLY | A | 292 | 20.260 | 32.764 | 9.304 | 1.00 | 0.00 | O |
| ATOM | 4157 | N | PRO | A | 293 | 19.060 | 34.080 | 10.703 | 1.00 | 0.00 | N |
| ATOM | 4158 | CD | PRO | A | 293 | 18.215 | 34.314 | 11.867 | 1.00 | 0.00 | C |
| ATOM | 4159 | HD1 | PRO | A | 293 | 18.614 | 33.835 | 12.762 | 1.00 | 0.00 | H |
| ATOM | 4160 | HD2 | PRO | A | 293 | 17.207 | 33.949 | 11.665 | 1.00 | 0.00 | H |
| ATOM | 4161 | CG | PRO | A | 293 | 18.183 | 35.826 | 12.052 | 1.00 | 0.00 | C |
| ATOM | 4162 | HG1 | PRO | A | 293 | 19.037 | 36.136 | 12.651 | 1.00 | 0.00 | H |
| ATOM | 4163 | HG2 | PRO | A | 293 | 17.250 | 36.158 | 12.506 | 1.00 | 0.00 | H |
| ATOM | 4164 | CB | PRO | A | 293 | 18.334 | 36.341 | 10.622 | 1.00 | 0.00 | C |
| ATOM | 4165 | HB1 | PRO | A | 293 | 18.762 | 37.342 | 10.607 | 1.00 | 0.00 | H |
| ATOM | 4166 | HB2 | PRO | A | 293 | 17.358 | 36.340 | 10.135 | 1.00 | 0.00 | H |
| ATOM | 4167 | CA | PRO | A | 293 | 19.242 | 35.311 | 9.938 | 1.00 | 0.00 | C |
| ATOM | 4168 | HA | PRO | A | 293 | 18.890 | 35.159 | 8.917 | 1.00 | 0.00 | H |
| ATOM | 4169 | C | PRO | A | 293 | 20.702 | 35.785 | 9.880 | 1.00 | 0.00 | C |
| ATOM | 4170 | O | PRO | A | 293 | 21.079 | 36.441 | 8.906 | 1.00 | 0.00 | O |
| ATOM | 4171 | N | PHE | A | 294 | 21.527 | 35.455 | 10.882 | 1.00 | 0.00 | N |
| ATOM | 4172 | H | PHE | A | 294 | 21.168 | 34.889 | 11.652 | 1.00 | 0.00 | H |
| ATOM | 4173 | CA | PHE | A | 294 | 22.904 | 35.934 | 11.012 | 1.00 | 0.00 | C |
| ATOM | 4174 | HA | PHE | A | 294 | 23.137 | 36.638 | 10.218 | 1.00 | 0.00 | H |
| ATOM | 4175 | CB | PHE | A | 294 | 23.030 | 36.675 | 12.344 | 1.00 | 0.00 | C |
| ATOM | 4176 | HB1 | PHE | A | 294 | 22.758 | 35.984 | 13.141 | 1.00 | 0.00 | H |
| ATOM | 4177 | HB2 | PHE | A | 294 | 24.075 | 36.948 | 12.493 | 1.00 | 0.00 | H |
| ATOM | 4178 | CG | PHE | A | 294 | 22.192 | 37.937 | 12.452 | 1.00 | 0.00 | C |
| ATOM | 4179 | CD1 | PHE | A | 294 | 22.557 | 39.090 | 11.730 | 1.00 | 0.00 | C |
| ATOM | 4180 | HD1 | PHE | A | 294 | 23.417 | 39.074 | 11.076 | 1.00 | 0.00 | H |
| ATOM | 4181 | CE1 | PHE | A | 294 | 21.831 | 40.283 | 11.888 | 1.00 | 0.00 | C |
| ATOM | 4182 | HE1 | PHE | A | 294 | 22.132 | 41.170 | 11.348 | 1.00 | 0.00 | H |
| ATOM | 4183 | CZ | PHE | A | 294 | 20.724 | 40.325 | 12.752 | 1.00 | 0.00 | C |
| ATOM | 4184 | HZ | PHE | A | 294 | 20.171 | 41.246 | 12.881 | 1.00 | 0.00 | H |
| ATOM | 4185 | CE2 | PHE | A | 294 | 20.338 | 39.169 | 13.451 | 1.00 | 0.00 | C |
| ATOM | 4186 | HE2 | PHE | A | 294 | 19.487 | 39.200 | 14.116 | 1.00 | 0.00 | H |
| ATOM | 4187 | CD2 | PHE | A | 294 | 21.075 | 37.980 | 13.308 | 1.00 | 0.00 | C |
| ATOM | 4188 | HD2 | PHE | A | 294 | 20.787 | 37.103 | 13.870 | 1.00 | 0.00 | H |
| ATOM | 4189 | C | PHE | A | 294 | 23.950 | 34.814 | 10.944 | 1.00 | 0.00 | C |
| ATOM | 4190 | O | PHE | A | 294 | 24.995 | 35.030 | 10.320 | 1.00 | 0.00 | O |
| ATOM | 4191 | N | THR | A | 295 | 23.688 | 33.649 | 11.556 | 1.00 | 0.00 | N |
| ATOM | 4192 | H | THR | A | 295 | 22.815 | 33.561 | 12.080 | 1.00 | 0.00 | H |
| ATOM | 4193 | CA | THR | A | 295 | 24.620 | 32.502 | 11.570 | 1.00 | 0.00 | C |
| ATOM | 4194 | HA | THR | A | 295 | 25.635 | 32.870 | 11.706 | 1.00 | 0.00 | H |
| ATOM | 4195 | CB | THR | A | 295 | 24.308 | 31.553 | 12.732 | 1.00 | 0.00 | C |
| ATOM | 4196 | HB | THR | A | 295 | 24.938 | 30.670 | 12.623 | 1.00 | 0.00 | H |
| ATOM | 4197 | CG2 | THR | A | 295 | 24.623 | 32.181 | 14.089 | 1.00 | 0.00 | C |
| ATOM | 4198 | HG21 | THR | A | 295 | 24.376 | 31.471 | 14.877 | 1.00 | 0.00 | H |
| ATOM | 4199 | HG22 | THR | A | 295 | 25.685 | 32.418 | 14.148 | 1.00 | 0.00 | H |
| ATOM | 4200 | HG23 | THR | A | 295 | 24.046 | 33.094 | 14.235 | 1.00 | 0.00 | H |
| ATOM | 4201 | OG1 | THR | A | 295 | 22.954 | 31.158 | 12.683 | 1.00 | 0.00 | O |
| ATOM | 4202 | HG1 | THR | A | 295 | 22.461 | 31.607 | 13.417 | 1.00 | 0.00 | H |
| ATOM | 4203 | C | THR | A | 295 | 24.623 | 31.726 | 10.257 | 1.00 | 0.00 | C |
| ATOM | 4204 | O | THR | A | 295 | 25.676 | 31.261 | 9.823 | 1.00 | 0.00 | O |
| ATOM | 4205 | N | LYS | A | 296 | 23.463 | 31.610 | 9.604 | 1.00 | 0.00 | N |
| ATOM | 4206 | H | LYS | A | 296 | 22.644 | 31.961 | 10.082 | 1.00 | 0.00 | H |
| ATOM | 4207 | CA | LYS | A | 296 | 23.277 | 31.076 | 8.245 | 1.00 | 0.00 | C |
| ATOM | 4208 | HA | LYS | A | 296 | 22.200 | 31.024 | 8.082 | 1.00 | 0.00 | H |
| ATOM | 4209 | CB | LYS | A | 296 | 23.855 | 32.066 | 7.206 | 1.00 | 0.00 | C |

| | | | | | | | | | | | |
|---|---|---|---|---|---|---|---|---|---|---|---|
| ATOM | 4210 | HB1 | LYS | A | 296 | 24.914 | 32.242 | 7.405 | 1.00 | 0.00 | H |
| ATOM | 4211 | HB2 | LYS | A | 296 | 23.770 | 31.619 | 6.215 | 1.00 | 0.00 | H |
| ATOM | 4212 | CG | LYS | A | 296 | 23.105 | 33.406 | 7.171 | 1.00 | 0.00 | C |
| ATOM | 4213 | HG1 | LYS | A | 296 | 22.036 | 33.208 | 7.090 | 1.00 | 0.00 | H |
| ATOM | 4214 | HG2 | LYS | A | 296 | 23.290 | 33.966 | 8.088 | 1.00 | 0.00 | H |
| ATOM | 4215 | CD | LYS | A | 296 | 23.552 | 34.236 | 5.959 | 1.00 | 0.00 | C |
| ATOM | 4216 | HD1 | LYS | A | 296 | 24.584 | 34.565 | 6.100 | 1.00 | 0.00 | H |
| ATOM | 4217 | HD2 | LYS | A | 296 | 23.498 | 33.619 | 5.060 | 1.00 | 0.00 | H |
| ATOM | 4218 | CE | LYS | A | 296 | 22.636 | 35.449 | 5.777 | 1.00 | 0.00 | C |
| ATOM | 4219 | HE1 | LYS | A | 296 | 21.598 | 35.107 | 5.739 | 1.00 | 0.00 | H |
| ATOM | 4220 | HE2 | LYS | A | 296 | 22.744 | 36.111 | 6.638 | 1.00 | 0.00 | H |
| ATOM | 4221 | NZ | LYS | A | 296 | 22.954 | 36.190 | 4.535 | 1.00 | 0.00 | N |
| ATOM | 4222 | HZ1 | LYS | A | 296 | 23.909 | 36.535 | 4.576 | 1.00 | 0.00 | H |
| ATOM | 4223 | HZ2 | LYS | A | 296 | 22.872 | 35.601 | 3.710 | 1.00 | 0.00 | H |
| ATOM | 4224 | HZ3 | LYS | A | 296 | 22.351 | 37.000 | 4.450 | 1.00 | 0.00 | H |
| ATOM | 4225 | C | LYS | A | 296 | 23.774 | 29.624 | 8.018 | 1.00 | 0.00 | C |
| ATOM | 4226 | O | LYS | A | 296 | 23.960 | 29.227 | 6.861 | 1.00 | 0.00 | O |
| ATOM | 4227 | N | GLU | A | 297 | 23.971 | 28.824 | 9.069 | 1.00 | 0.00 | N |
| ATOM | 4228 | H | GLU | A | 297 | 23.733 | 29.201 | 9.981 | 1.00 | 0.00 | H |
| ATOM | 4229 | CA | GLU | A | 297 | 24.296 | 27.393 | 9.008 | 1.00 | 0.00 | C |
| ATOM | 4230 | HA | GLU | A | 297 | 24.426 | 27.100 | 7.967 | 1.00 | 0.00 | H |
| ATOM | 4231 | CB | GLU | A | 297 | 25.643 | 27.129 | 9.709 | 1.00 | 0.00 | C |
| ATOM | 4232 | HB1 | GLU | A | 297 | 26.412 | 27.597 | 9.099 | 1.00 | 0.00 | H |
| ATOM | 4233 | HB2 | GLU | A | 297 | 25.658 | 27.597 | 10.693 | 1.00 | 0.00 | H |
| ATOM | 4234 | CG | GLU | A | 297 | 25.973 | 25.636 | 9.860 | 1.00 | 0.00 | C |
| ATOM | 4235 | HG1 | GLU | A | 297 | 25.452 | 25.237 | 10.731 | 1.00 | 0.00 | H |
| ATOM | 4236 | HG2 | GLU | A | 297 | 25.607 | 25.107 | 8.978 | 1.00 | 0.00 | H |
| ATOM | 4237 | CD | GLU | A | 297 | 27.472 | 25.346 | 9.997 | 1.00 | 0.00 | C |
| ATOM | 4238 | OE1 | GLU | A | 297 | 27.926 | 24.342 | 9.396 | 1.00 | 0.00 | O |
| ATOM | 4239 | OE2 | GLU | A | 297 | 28.223 | 26.079 | 10.678 | 1.00 | 0.00 | O |
| ATOM | 4240 | C | GLU | A | 297 | 23.142 | 26.550 | 9.581 | 1.00 | 0.00 | C |
| ATOM | 4241 | O | GLU | A | 297 | 22.737 | 26.704 | 10.735 | 1.00 | 0.00 | O |
| ATOM | 4242 | N | GLY | A | 298 | 22.598 | 25.644 | 8.764 | 1.00 | 0.00 | N |
| ATOM | 4243 | H | GLY | A | 298 | 22.930 | 25.596 | 7.807 | 1.00 | 0.00 | H |
| ATOM | 4244 | CA | GLY | A | 298 | 21.450 | 24.815 | 9.130 | 1.00 | 0.00 | C |
| ATOM | 4245 | HA1 | GLY | A | 298 | 20.635 | 25.463 | 9.457 | 1.00 | 0.00 | H |
| ATOM | 4246 | HA2 | GLY | A | 298 | 21.121 | 24.239 | 8.267 | 1.00 | 0.00 | H |
| ATOM | 4247 | C | GLY | A | 298 | 21.779 | 23.842 | 10.260 | 1.00 | 0.00 | C |
| ATOM | 4248 | O | GLY | A | 298 | 22.746 | 23.075 | 10.178 | 1.00 | 0.00 | O |
| ATOM | 4249 | N | GLY | A | 299 | 20.946 | 23.848 | 11.304 | 1.00 | 0.00 | N |
| ATOM | 4250 | H | GLY | A | 299 | 20.210 | 24.547 | 11.313 | 1.00 | 0.00 | H |
| ATOM | 4251 | CA | GLY | A | 299 | 21.047 | 22.957 | 12.467 | 1.00 | 0.00 | C |
| ATOM | 4252 | HA1 | GLY | A | 299 | 20.048 | 22.596 | 12.704 | 1.00 | 0.00 | H |
| ATOM | 4253 | HA2 | GLY | A | 299 | 21.690 | 22.105 | 12.247 | 1.00 | 0.00 | H |
| ATOM | 4254 | C | GLY | A | 299 | 21.569 | 23.621 | 13.743 | 1.00 | 0.00 | C |
| ATOM | 4255 | O | GLY | A | 299 | 21.491 | 22.995 | 14.797 | 1.00 | 0.00 | O |
| ATOM | 4256 | N | MET | A | 300 | 22.042 | 24.872 | 13.688 | 1.00 | 0.00 | N |
| ATOM | 4257 | H | MET | A | 300 | 22.057 | 25.338 | 12.787 | 1.00 | 0.00 | H |
| ATOM | 4258 | CA | MET | A | 300 | 22.357 | 25.702 | 14.864 | 1.00 | 0.00 | C |
| ATOM | 4259 | HA | MET | A | 300 | 21.908 | 25.253 | 15.747 | 1.00 | 0.00 | H |
| ATOM | 4260 | CB | MET | A | 300 | 23.883 | 25.744 | 15.098 | 1.00 | 0.00 | C |
| ATOM | 4261 | HB1 | MET | A | 300 | 24.209 | 24.728 | 15.322 | 1.00 | 0.00 | H |
| ATOM | 4262 | HB2 | MET | A | 300 | 24.395 | 26.059 | 14.189 | 1.00 | 0.00 | H |
| ATOM | 4263 | CG | MET | A | 300 | 24.312 | 26.650 | 16.270 | 1.00 | 0.00 | C |

| | | | | | | | | | | | |
|---|---|---|---|---|---|---|---|---|---|---|---|
| ATOM | 4264 | HG1 | MET | A | 300 | 23.576 | 26.572 | 17.067 | 1.00 | 0.00 | H |
| ATOM | 4265 | HG2 | MET | A | 300 | 25.237 | 26.269 | 16.687 | 1.00 | 0.00 | H |
| ATOM | 4266 | SD | MET | A | 300 | 24.568 | 28.412 | 15.907 | 1.00 | 0.00 | S |
| ATOM | 4267 | CE | MET | A | 300 | 26.259 | 28.382 | 15.267 | 1.00 | 0.00 | C |
| ATOM | 4268 | HE1 | MET | A | 300 | 26.526 | 29.370 | 14.891 | 1.00 | 0.00 | H |
| ATOM | 4269 | HE2 | MET | A | 300 | 26.945 | 28.129 | 16.075 | 1.00 | 0.00 | H |
| ATOM | 4270 | HE3 | MET | A | 300 | 26.357 | 27.653 | 14.461 | 1.00 | 0.00 | H |
| ATOM | 4271 | C | MET | A | 300 | 21.741 | 27.097 | 14.717 | 1.00 | 0.00 | C |
| ATOM | 4272 | O | MET | A | 300 | 21.633 | 27.610 | 13.601 | 1.00 | 0.00 | O |
| ATOM | 4273 | N | LEU | A | 301 | 21.356 | 27.697 | 15.847 | 1.00 | 0.00 | N |
| ATOM | 4274 | H | LEU | A | 301 | 21.495 | 27.195 | 16.718 | 1.00 | 0.00 | H |
| ATOM | 4275 | CA | LEU | A | 301 | 20.937 | 29.093 | 15.988 | 1.00 | 0.00 | C |
| ATOM | 4276 | HA | LEU | A | 301 | 21.256 | 29.678 | 15.122 | 1.00 | 0.00 | H |
| ATOM | 4277 | CB | LEU | A | 301 | 19.404 | 29.154 | 16.130 | 1.00 | 0.00 | C |
| ATOM | 4278 | HB1 | LEU | A | 301 | 19.124 | 28.579 | 17.015 | 1.00 | 0.00 | H |
| ATOM | 4279 | HB2 | LEU | A | 301 | 19.121 | 30.191 | 16.306 | 1.00 | 0.00 | H |
| ATOM | 4280 | CG | LEU | A | 301 | 18.600 | 28.629 | 14.923 | 1.00 | 0.00 | C |
| ATOM | 4281 | HG | LEU | A | 301 | 18.950 | 27.634 | 14.652 | 1.00 | 0.00 | H |
| ATOM | 4282 | CD1 | LEU | A | 301 | 17.120 | 28.507 | 15.305 | 1.00 | 0.00 | C |
| ATOM | 4283 | HD11 | LEU | A | 301 | 16.548 | 28.124 | 14.460 | 1.00 | 0.00 | H |
| ATOM | 4284 | HD12 | LEU | A | 301 | 17.011 | 27.819 | 16.144 | 1.00 | 0.00 | H |
| ATOM | 4285 | HD13 | LEU | A | 301 | 16.732 | 29.483 | 15.592 | 1.00 | 0.00 | H |
| ATOM | 4286 | CD2 | LEU | A | 301 | 18.737 | 29.551 | 13.706 | 1.00 | 0.00 | C |
| ATOM | 4287 | HD21 | LEU | A | 301 | 18.134 | 29.168 | 12.882 | 1.00 | 0.00 | H |
| ATOM | 4288 | HD22 | LEU | A | 301 | 18.399 | 30.557 | 13.957 | 1.00 | 0.00 | H |
| ATOM | 4289 | HD23 | LEU | A | 301 | 19.777 | 29.596 | 13.384 | 1.00 | 0.00 | H |
| ATOM | 4290 | C | LEU | A | 301 | 21.609 | 29.679 | 17.241 | 1.00 | 0.00 | C |
| ATOM | 4291 | O | LEU | A | 301 | 21.628 | 29.025 | 18.287 | 1.00 | 0.00 | O |
| ATOM | 4292 | N | ALA | A | 302 | 22.155 | 30.893 | 17.153 | 1.00 | 0.00 | N |
| ATOM | 4293 | H | ALA | A | 302 | 22.057 | 31.397 | 16.271 | 1.00 | 0.00 | H |
| ATOM | 4294 | CA | ALA | A | 302 | 22.786 | 31.577 | 18.291 | 1.00 | 0.00 | C |
| ATOM | 4295 | HA | ALA | A | 302 | 23.495 | 30.896 | 18.765 | 1.00 | 0.00 | H |
| ATOM | 4296 | CB | ALA | A | 302 | 23.556 | 32.791 | 17.761 | 1.00 | 0.00 | C |
| ATOM | 4297 | HB1 | ALA | A | 302 | 24.061 | 33.294 | 18.585 | 1.00 | 0.00 | H |
| ATOM | 4298 | HB2 | ALA | A | 302 | 24.304 | 32.473 | 17.037 | 1.00 | 0.00 | H |
| ATOM | 4299 | HB3 | ALA | A | 302 | 22.871 | 33.491 | 17.284 | 1.00 | 0.00 | H |
| ATOM | 4300 | C | ALA | A | 302 | 21.754 | 31.984 | 19.360 | 1.00 | 0.00 | C |
| ATOM | 4301 | O | ALA | A | 302 | 20.564 | 32.115 | 19.057 | 1.00 | 0.00 | O |
| ATOM | 4302 | N | TYR | A | 303 | 22.186 | 32.268 | 20.597 | 1.00 | 0.00 | N |
| ATOM | 4303 | H | TYR | A | 303 | 23.179 | 32.204 | 20.814 | 1.00 | 0.00 | H |
| ATOM | 4304 | CA | TYR | A | 303 | 21.276 | 32.733 | 21.657 | 1.00 | 0.00 | C |
| ATOM | 4305 | HA | TYR | A | 303 | 20.577 | 31.927 | 21.883 | 1.00 | 0.00 | H |
| ATOM | 4306 | CB | TYR | A | 303 | 22.071 | 33.052 | 22.933 | 1.00 | 0.00 | C |
| ATOM | 4307 | HB1 | TYR | A | 303 | 22.652 | 32.173 | 23.212 | 1.00 | 0.00 | H |
| ATOM | 4308 | HB2 | TYR | A | 303 | 22.775 | 33.859 | 22.727 | 1.00 | 0.00 | H |
| ATOM | 4309 | CG | TYR | A | 303 | 21.203 | 33.433 | 24.119 | 1.00 | 0.00 | C |
| ATOM | 4310 | CD1 | TYR | A | 303 | 20.482 | 32.437 | 24.805 | 1.00 | 0.00 | C |
| ATOM | 4311 | HD1 | TYR | A | 303 | 20.571 | 31.405 | 24.495 | 1.00 | 0.00 | H |
| ATOM | 4312 | CE1 | TYR | A | 303 | 19.688 | 32.771 | 25.919 | 1.00 | 0.00 | C |
| ATOM | 4313 | HE1 | TYR | A | 303 | 19.159 | 31.999 | 26.459 | 1.00 | 0.00 | H |
| ATOM | 4314 | CZ | TYR | A | 303 | 19.578 | 34.117 | 26.329 | 1.00 | 0.00 | C |
| ATOM | 4315 | OH | TYR | A | 303 | 18.768 | 34.440 | 27.374 | 1.00 | 0.00 | O |
| ATOM | 4316 | HH | TYR | A | 303 | 18.724 | 35.399 | 27.523 | 1.00 | 0.00 | H |
| ATOM | 4317 | CE2 | TYR | A | 303 | 20.298 | 35.118 | 25.641 | 1.00 | 0.00 | C |

```
ATOM   4318  HE2 TYR A 303      20.224  36.149  25.955  1.00  0.00           H
ATOM   4319  CD2 TYR A 303      21.113  34.773  24.545  1.00  0.00           C
ATOM   4320  HD2 TYR A 303      21.673  35.542  24.029  1.00  0.00           H
ATOM   4321  C   TYR A 303      20.449  33.944  21.188  1.00  0.00           C
ATOM   4322  O   TYR A 303      19.216  33.910  21.216  1.00  0.00           O
ATOM   4323  N   TYR A 304      21.109  34.951  20.604  1.00  0.00           N
ATOM   4324  H   TYR A 304      22.121  34.892  20.580  1.00  0.00           H
ATOM   4325  CA  TYR A 304      20.467  36.137  20.022  1.00  0.00           C
ATOM   4326  HA  TYR A 304      19.820  36.561  20.788  1.00  0.00           H
ATOM   4327  CB  TYR A 304      21.541  37.193  19.714  1.00  0.00           C
ATOM   4328  HB1 TYR A 304      21.051  38.123  19.428  1.00  0.00           H
ATOM   4329  HB2 TYR A 304      22.101  37.399  20.627  1.00  0.00           H
ATOM   4330  CG  TYR A 304      22.509  36.817  18.611  1.00  0.00           C
ATOM   4331  CD1 TYR A 304      22.101  36.872  17.264  1.00  0.00           C
ATOM   4332  HD1 TYR A 304      21.106  37.200  17.007  1.00  0.00           H
ATOM   4333  CE1 TYR A 304      22.984  36.492  16.242  1.00  0.00           C
ATOM   4334  HE1 TYR A 304      22.664  36.521  15.217  1.00  0.00           H
ATOM   4335  CZ  TYR A 304      24.294  36.076  16.552  1.00  0.00           C
ATOM   4336  OH  TYR A 304      25.137  35.707  15.549  1.00  0.00           O
ATOM   4337  HH  TYR A 304      26.048  35.614  15.854  1.00  0.00           H
ATOM   4338  CE2 TYR A 304      24.713  36.060  17.899  1.00  0.00           C
ATOM   4339  HE2 TYR A 304      25.710  35.754  18.156  1.00  0.00           H
ATOM   4340  CD2 TYR A 304      23.819  36.415  18.924  1.00  0.00           C
ATOM   4341  HD2 TYR A 304      24.146  36.374  19.955  1.00  0.00           H
ATOM   4342  C   TYR A 304      19.560  35.863  18.797  1.00  0.00           C
ATOM   4343  O   TYR A 304      18.872  36.782  18.344  1.00  0.00           O
ATOM   4344  N   GLU A 305      19.531  34.642  18.251  1.00  0.00           N
ATOM   4345  H   GLU A 305      20.154  33.931  18.624  1.00  0.00           H
ATOM   4346  CA  GLU A 305      18.594  34.229  17.195  1.00  0.00           C
ATOM   4347  HA  GLU A 305      18.240  35.114  16.666  1.00  0.00           H
ATOM   4348  CB  GLU A 305      19.308  33.341  16.151  1.00  0.00           C
ATOM   4349  HB1 GLU A 305      19.775  32.495  16.651  1.00  0.00           H
ATOM   4350  HB2 GLU A 305      18.572  32.950  15.450  1.00  0.00           H
ATOM   4351  CG  GLU A 305      20.364  34.119  15.348  1.00  0.00           C
ATOM   4352  HG1 GLU A 305      19.883  34.966  14.857  1.00  0.00           H
ATOM   4353  HG2 GLU A 305      21.116  34.503  16.033  1.00  0.00           H
ATOM   4354  CD  GLU A 305      21.077  33.264  14.296  1.00  0.00           C
ATOM   4355  OE1 GLU A 305      21.548  32.159  14.643  1.00  0.00           O
ATOM   4356  OE2 GLU A 305      21.215  33.715  13.134  1.00  0.00           O
ATOM   4357  C   GLU A 305      17.333  33.533  17.757  1.00  0.00           C
ATOM   4358  O   GLU A 305      16.343  33.446  17.027  1.00  0.00           O
ATOM   4359  N   VAL A 306      17.314  33.081  19.024  1.00  0.00           N
ATOM   4360  H   VAL A 306      18.177  33.148  19.560  1.00  0.00           H
ATOM   4361  CA  VAL A 306      16.168  32.344  19.624  1.00  0.00           C
ATOM   4362  HA  VAL A 306      15.325  32.432  18.940  1.00  0.00           H
ATOM   4363  CB  VAL A 306      16.464  30.830  19.771  1.00  0.00           C
ATOM   4364  HB  VAL A 306      15.584  30.348  20.199  1.00  0.00           H
ATOM   4365  CG1 VAL A 306      16.692  30.183  18.399  1.00  0.00           C
ATOM   4366 HG11 VAL A 306      16.823  29.108  18.513  1.00  0.00           H
ATOM   4367 HG12 VAL A 306      15.829  30.365  17.758  1.00  0.00           H
ATOM   4368 HG13 VAL A 306      17.582  30.600  17.932  1.00  0.00           H
ATOM   4369  CG2 VAL A 306      17.651  30.531  20.706  1.00  0.00           C
ATOM   4370 HG21 VAL A 306      17.782  29.453  20.801  1.00  0.00           H
ATOM   4371 HG22 VAL A 306      18.567  30.965  20.310  1.00  0.00           H
```

| | | | | | | | | | | | |
|---|---|---|---|---|---|---|---|---|---|---|---|
| ATOM | 4372 | HG23 | VAL | A | 306 | 17.462 | 30.943 | 21.696 | 1.00 | 0.00 | H |
| ATOM | 4373 | C | VAL | A | 306 | 15.652 | 32.882 | 20.974 | 1.00 | 0.00 | C |
| ATOM | 4374 | O | VAL | A | 306 | 14.619 | 32.417 | 21.458 | 1.00 | 0.00 | O |
| ATOM | 4375 | N | CYS | A | 307 | 16.325 | 33.850 | 21.603 | 1.00 | 0.00 | N |
| ATOM | 4376 | H | CYS | A | 307 | 17.184 | 34.198 | 21.193 | 1.00 | 0.00 | H |
| ATOM | 4377 | CA | CYS | A | 307 | 15.966 | 34.354 | 22.939 | 1.00 | 0.00 | C |
| ATOM | 4378 | HA | CYS | A | 307 | 15.823 | 33.510 | 23.609 | 1.00 | 0.00 | H |
| ATOM | 4379 | CB | CYS | A | 307 | 17.138 | 35.181 | 23.465 | 1.00 | 0.00 | C |
| ATOM | 4380 | HB1 | CYS | A | 307 | 16.897 | 35.539 | 24.465 | 1.00 | 0.00 | H |
| ATOM | 4381 | HB2 | CYS | A | 307 | 18.016 | 34.539 | 23.546 | 1.00 | 0.00 | H |
| ATOM | 4382 | SG | CYS | A | 307 | 17.562 | 36.602 | 22.433 | 1.00 | 0.00 | S |
| ATOM | 4383 | C | CYS | A | 307 | 14.658 | 35.155 | 22.993 | 1.00 | 0.00 | C |
| ATOM | 4384 | O | CYS | A | 307 | 13.970 | 35.141 | 24.008 | 1.00 | 0.00 | O |
| ATOM | 4385 | N | SER | A | 308 | 14.277 | 35.812 | 21.902 | 1.00 | 0.00 | N |
| ATOM | 4386 | H | SER | A | 308 | 14.926 | 35.809 | 21.117 | 1.00 | 0.00 | H |
| ATOM | 4387 | CA | SER | A | 308 | 13.016 | 36.544 | 21.722 | 1.00 | 0.00 | C |
| ATOM | 4388 | HA | SER | A | 308 | 12.339 | 36.345 | 22.555 | 1.00 | 0.00 | H |
| ATOM | 4389 | CB | SER | A | 308 | 13.311 | 38.054 | 21.687 | 1.00 | 0.00 | C |
| ATOM | 4390 | HB1 | SER | A | 308 | 14.215 | 38.244 | 21.107 | 1.00 | 0.00 | H |
| ATOM | 4391 | HB2 | SER | A | 308 | 12.482 | 38.581 | 21.213 | 1.00 | 0.00 | H |
| ATOM | 4392 | OG | SER | A | 308 | 13.448 | 38.587 | 22.992 | 1.00 | 0.00 | O |
| ATOM | 4393 | HG | SER | A | 308 | 14.090 | 38.075 | 23.519 | 1.00 | 0.00 | H |
| ATOM | 4394 | C | SER | A | 308 | 12.293 | 36.063 | 20.447 | 1.00 | 0.00 | C |
| ATOM | 4395 | O | SER | A | 308 | 11.853 | 36.863 | 19.616 | 1.00 | 0.00 | O |
| ATOM | 4396 | N | TRP | A | 309 | 12.228 | 34.741 | 20.240 | 1.00 | 0.00 | N |
| ATOM | 4397 | H | TRP | A | 309 | 12.618 | 34.127 | 20.943 | 1.00 | 0.00 | H |
| ATOM | 4398 | CA | TRP | A | 309 | 11.692 | 34.122 | 19.023 | 1.00 | 0.00 | C |
| ATOM | 4399 | HA | TRP | A | 309 | 12.181 | 34.602 | 18.177 | 1.00 | 0.00 | H |
| ATOM | 4400 | CB | TRP | A | 309 | 12.088 | 32.636 | 19.010 | 1.00 | 0.00 | C |
| ATOM | 4401 | HB1 | TRP | A | 309 | 13.115 | 32.557 | 19.357 | 1.00 | 0.00 | H |
| ATOM | 4402 | HB2 | TRP | A | 309 | 11.468 | 32.096 | 19.727 | 1.00 | 0.00 | H |
| ATOM | 4403 | CG | TRP | A | 309 | 12.048 | 31.917 | 17.693 | 1.00 | 0.00 | C |
| ATOM | 4404 | CD1 | TRP | A | 309 | 12.273 | 32.465 | 16.477 | 1.00 | 0.00 | C |
| ATOM | 4405 | HD1 | TRP | A | 309 | 12.495 | 33.509 | 16.288 | 1.00 | 0.00 | H |
| ATOM | 4406 | NE1 | TRP | A | 309 | 12.221 | 31.487 | 15.508 | 1.00 | 0.00 | N |
| ATOM | 4407 | HE1 | TRP | A | 309 | 12.389 | 31.675 | 14.526 | 1.00 | 0.00 | H |
| ATOM | 4408 | CE2 | TRP | A | 309 | 11.972 | 30.244 | 16.043 | 1.00 | 0.00 | C |
| ATOM | 4409 | CZ2 | TRP | A | 309 | 11.855 | 28.962 | 15.484 | 1.00 | 0.00 | C |
| ATOM | 4410 | HZ2 | TRP | A | 309 | 11.945 | 28.820 | 14.417 | 1.00 | 0.00 | H |
| ATOM | 4411 | CH2 | TRP | A | 309 | 11.650 | 27.863 | 16.336 | 1.00 | 0.00 | C |
| ATOM | 4412 | HH2 | TRP | A | 309 | 11.576 | 26.863 | 15.932 | 1.00 | 0.00 | H |
| ATOM | 4413 | CZ3 | TRP | A | 309 | 11.573 | 28.063 | 17.725 | 1.00 | 0.00 | C |
| ATOM | 4414 | HZ3 | TRP | A | 309 | 11.451 | 27.215 | 18.381 | 1.00 | 0.00 | H |
| ATOM | 4415 | CE3 | TRP | A | 309 | 11.689 | 29.356 | 18.269 | 1.00 | 0.00 | C |
| ATOM | 4416 | HE3 | TRP | A | 309 | 11.656 | 29.493 | 19.336 | 1.00 | 0.00 | H |
| ATOM | 4417 | CD2 | TRP | A | 309 | 11.877 | 30.485 | 17.445 | 1.00 | 0.00 | C |
| ATOM | 4418 | C | TRP | A | 309 | 10.172 | 34.351 | 18.894 | 1.00 | 0.00 | C |
| ATOM | 4419 | O | TRP | A | 309 | 9.371 | 33.861 | 19.696 | 1.00 | 0.00 | O |
| ATOM | 4420 | N | LYS | A | 310 | 9.765 | 35.136 | 17.893 | 1.00 | 0.00 | N |
| ATOM | 4421 | H | LYS | A | 310 | 10.451 | 35.459 | 17.223 | 1.00 | 0.00 | H |
| ATOM | 4422 | CA | LYS | A | 310 | 8.408 | 35.692 | 17.790 | 1.00 | 0.00 | C |
| ATOM | 4423 | HA | LYS | A | 310 | 8.116 | 36.081 | 18.767 | 1.00 | 0.00 | H |
| ATOM | 4424 | CB | LYS | A | 310 | 8.437 | 36.875 | 16.802 | 1.00 | 0.00 | C |
| ATOM | 4425 | HB1 | LYS | A | 310 | 8.968 | 36.578 | 15.896 | 1.00 | 0.00 | H |

| | | | | | | | | | | | |
|---|---|---|---|---|---|---|---|---|---|---|---|
| ATOM | 4426 | HB2 | LYS | A | 310 | 7.421 | 37.154 | 16.521 | 1.00 | 0.00 | H |
| ATOM | 4427 | CG | LYS | A | 310 | 9.121 | 38.089 | 17.458 | 1.00 | 0.00 | C |
| ATOM | 4428 | HG1 | LYS | A | 310 | 8.389 | 38.601 | 18.080 | 1.00 | 0.00 | H |
| ATOM | 4429 | HG2 | LYS | A | 310 | 9.917 | 37.737 | 18.108 | 1.00 | 0.00 | H |
| ATOM | 4430 | CD | LYS | A | 310 | 9.726 | 39.082 | 16.452 | 1.00 | 0.00 | C |
| ATOM | 4431 | HD1 | LYS | A | 310 | 10.436 | 38.556 | 15.811 | 1.00 | 0.00 | H |
| ATOM | 4432 | HD2 | LYS | A | 310 | 8.930 | 39.481 | 15.820 | 1.00 | 0.00 | H |
| ATOM | 4433 | CE | LYS | A | 310 | 10.446 | 40.255 | 17.146 | 1.00 | 0.00 | C |
| ATOM | 4434 | HE1 | LYS | A | 310 | 10.893 | 40.892 | 16.376 | 1.00 | 0.00 | H |
| ATOM | 4435 | HE2 | LYS | A | 310 | 9.703 | 40.854 | 17.679 | 1.00 | 0.00 | H |
| ATOM | 4436 | NZ | LYS | A | 310 | 11.495 | 39.806 | 18.101 | 1.00 | 0.00 | N |
| ATOM | 4437 | HZ1 | LYS | A | 310 | 12.171 | 39.196 | 17.653 | 1.00 | 0.00 | H |
| ATOM | 4438 | HZ2 | LYS | A | 310 | 11.969 | 40.599 | 18.523 | 1.00 | 0.00 | H |
| ATOM | 4439 | HZ3 | LYS | A | 310 | 11.066 | 39.295 | 18.867 | 1.00 | 0.00 | H |
| ATOM | 4440 | C | LYS | A | 310 | 7.381 | 34.612 | 17.418 | 1.00 | 0.00 | C |
| ATOM | 4441 | O | LYS | A | 310 | 7.602 | 33.829 | 16.492 | 1.00 | 0.00 | O |
| ATOM | 4442 | N | GLY | A | 311 | 6.291 | 34.508 | 18.186 | 1.00 | 0.00 | N |
| ATOM | 4443 | H | GLY | A | 311 | 6.181 | 35.173 | 18.944 | 1.00 | 0.00 | H |
| ATOM | 4444 | CA | GLY | A | 311 | 5.224 | 33.517 | 17.977 | 1.00 | 0.00 | C |
| ATOM | 4445 | HA1 | GLY | A | 311 | 4.357 | 33.804 | 18.573 | 1.00 | 0.00 | H |
| ATOM | 4446 | HA2 | GLY | A | 311 | 4.952 | 33.507 | 16.921 | 1.00 | 0.00 | H |
| ATOM | 4447 | C | GLY | A | 311 | 5.596 | 32.080 | 18.369 | 1.00 | 0.00 | C |
| ATOM | 4448 | O | GLY | A | 311 | 4.865 | 31.154 | 18.018 | 1.00 | 0.00 | O |
| ATOM | 4449 | N | ALA | A | 312 | 6.725 | 31.862 | 19.048 | 1.00 | 0.00 | N |
| ATOM | 4450 | H | ALA | A | 312 | 7.266 | 32.667 | 19.328 | 1.00 | 0.00 | H |
| ATOM | 4451 | CA | ALA | A | 312 | 7.180 | 30.548 | 19.512 | 1.00 | 0.00 | C |
| ATOM | 4452 | HA | ALA | A | 312 | 6.878 | 29.789 | 18.788 | 1.00 | 0.00 | H |
| ATOM | 4453 | CB | ALA | A | 312 | 8.711 | 30.581 | 19.548 | 1.00 | 0.00 | C |
| ATOM | 4454 | HB1 | ALA | A | 312 | 9.093 | 29.578 | 19.725 | 1.00 | 0.00 | H |
| ATOM | 4455 | HB2 | ALA | A | 312 | 9.103 | 30.937 | 18.595 | 1.00 | 0.00 | H |
| ATOM | 4456 | HB3 | ALA | A | 312 | 9.051 | 31.243 | 20.345 | 1.00 | 0.00 | H |
| ATOM | 4457 | C | ALA | A | 312 | 6.570 | 30.143 | 20.875 | 1.00 | 0.00 | C |
| ATOM | 4458 | O | ALA | A | 312 | 5.913 | 30.956 | 21.535 | 1.00 | 0.00 | O |
| ATOM | 4459 | N | THR | A | 313 | 6.791 | 28.888 | 21.288 | 1.00 | 0.00 | N |
| ATOM | 4460 | H | THR | A | 313 | 7.318 | 28.277 | 20.672 | 1.00 | 0.00 | H |
| ATOM | 4461 | CA | THR | A | 313 | 6.256 | 28.289 | 22.524 | 1.00 | 0.00 | C |
| ATOM | 4462 | HA | THR | A | 313 | 5.850 | 29.066 | 23.170 | 1.00 | 0.00 | H |
| ATOM | 4463 | CB | THR | A | 313 | 5.110 | 27.333 | 22.176 | 1.00 | 0.00 | C |
| ATOM | 4464 | HB | THR | A | 313 | 5.472 | 26.549 | 21.509 | 1.00 | 0.00 | H |
| ATOM | 4465 | CG2 | THR | A | 313 | 4.489 | 26.695 | 23.416 | 1.00 | 0.00 | C |
| ATOM | 4466 | HG21 | THR | A | 313 | 3.626 | 26.097 | 23.124 | 1.00 | 0.00 | H |
| ATOM | 4467 | HG22 | THR | A | 313 | 5.209 | 26.042 | 23.909 | 1.00 | 0.00 | H |
| ATOM | 4468 | HG23 | THR | A | 313 | 4.165 | 27.470 | 24.111 | 1.00 | 0.00 | H |
| ATOM | 4469 | OG1 | THR | A | 313 | 4.088 | 28.067 | 21.539 | 1.00 | 0.00 | O |
| ATOM | 4470 | HG1 | THR | A | 313 | 4.243 | 28.073 | 20.588 | 1.00 | 0.00 | H |
| ATOM | 4471 | C | THR | A | 313 | 7.372 | 27.574 | 23.284 | 1.00 | 0.00 | C |
| ATOM | 4472 | O | THR | A | 313 | 7.987 | 26.651 | 22.753 | 1.00 | 0.00 | O |
| ATOM | 4473 | N | LYS | A | 314 | 7.650 | 28.022 | 24.514 | 1.00 | 0.00 | N |
| ATOM | 4474 | H | LYS | A | 314 | 7.043 | 28.744 | 24.888 | 1.00 | 0.00 | H |
| ATOM | 4475 | CA | LYS | A | 314 | 8.782 | 27.607 | 25.362 | 1.00 | 0.00 | C |
| ATOM | 4476 | HA | LYS | A | 314 | 9.455 | 26.990 | 24.767 | 1.00 | 0.00 | H |
| ATOM | 4477 | CB | LYS | A | 314 | 9.549 | 28.886 | 25.784 | 1.00 | 0.00 | C |
| ATOM | 4478 | HB1 | LYS | A | 314 | 9.820 | 29.429 | 24.879 | 1.00 | 0.00 | H |
| ATOM | 4479 | HB2 | LYS | A | 314 | 8.874 | 29.521 | 26.361 | 1.00 | 0.00 | H |

| | | | | | | | | | | |
|---|---|---|---|---|---|---|---|---|---|---|
| ATOM | 4480 | CG | LYS | A | 314 | 10.833 | 28.661 | 26.610 | 1.00 | 0.00 | C |
| ATOM | 4481 | HG1 | LYS | A | 314 | 10.561 | 28.251 | 27.584 | 1.00 | 0.00 | H |
| ATOM | 4482 | HG2 | LYS | A | 314 | 11.474 | 27.944 | 26.094 | 1.00 | 0.00 | H |
| ATOM | 4483 | CD | LYS | A | 314 | 11.622 | 29.966 | 26.836 | 1.00 | 0.00 | C |
| ATOM | 4484 | HD1 | LYS | A | 314 | 12.002 | 30.328 | 25.879 | 1.00 | 0.00 | H |
| ATOM | 4485 | HD2 | LYS | A | 314 | 10.956 | 30.723 | 27.255 | 1.00 | 0.00 | H |
| ATOM | 4486 | CE | LYS | A | 314 | 12.790 | 29.727 | 27.810 | 1.00 | 0.00 | C |
| ATOM | 4487 | HE1 | LYS | A | 314 | 12.377 | 29.409 | 28.771 | 1.00 | 0.00 | H |
| ATOM | 4488 | HE2 | LYS | A | 314 | 13.410 | 28.912 | 27.434 | 1.00 | 0.00 | H |
| ATOM | 4489 | NZ | LYS | A | 314 | 13.633 | 30.933 | 28.019 | 1.00 | 0.00 | N |
| ATOM | 4490 | HZ1 | LYS | A | 314 | 14.045 | 31.277 | 27.155 | 1.00 | 0.00 | H |
| ATOM | 4491 | HZ2 | LYS | A | 314 | 13.102 | 31.695 | 28.433 | 1.00 | 0.00 | H |
| ATOM | 4492 | HZ3 | LYS | A | 314 | 14.399 | 30.725 | 28.657 | 1.00 | 0.00 | H |
| ATOM | 4493 | C | LYS | A | 314 | 8.341 | 26.762 | 26.579 | 1.00 | 0.00 | C |
| ATOM | 4494 | O | LYS | A | 314 | 7.221 | 26.886 | 27.082 | 1.00 | 0.00 | O |
| ATOM | 4495 | N | GLN | A | 315 | 9.252 | 25.922 | 27.078 | 1.00 | 0.00 | N |
| ATOM | 4496 | H | GLN | A | 315 | 10.147 | 25.857 | 26.605 | 1.00 | 0.00 | H |
| ATOM | 4497 | CA | GLN | A | 315 | 9.135 | 25.107 | 28.293 | 1.00 | 0.00 | C |
| ATOM | 4498 | HA | GLN | A | 315 | 8.814 | 25.748 | 29.115 | 1.00 | 0.00 | H |
| ATOM | 4499 | CB | GLN | A | 315 | 8.080 | 23.999 | 28.119 | 1.00 | 0.00 | C |
| ATOM | 4500 | HB1 | GLN | A | 315 | 8.029 | 23.419 | 29.041 | 1.00 | 0.00 | H |
| ATOM | 4501 | HB2 | GLN | A | 315 | 7.095 | 24.447 | 27.984 | 1.00 | 0.00 | H |
| ATOM | 4502 | CG | GLN | A | 315 | 8.373 | 23.031 | 26.962 | 1.00 | 0.00 | C |
| ATOM | 4503 | HG1 | GLN | A | 315 | 9.438 | 23.009 | 26.739 | 1.00 | 0.00 | H |
| ATOM | 4504 | HG2 | GLN | A | 315 | 8.068 | 22.035 | 27.277 | 1.00 | 0.00 | H |
| ATOM | 4505 | CD | GLN | A | 315 | 7.593 | 23.370 | 25.700 | 1.00 | 0.00 | C |
| ATOM | 4506 | OE1 | GLN | A | 315 | 6.415 | 23.057 | 25.603 | 1.00 | 0.00 | O |
| ATOM | 4507 | NE2 | GLN | A | 315 | 8.194 | 24.015 | 24.721 | 1.00 | 0.00 | N |
| ATOM | 4508 | HE21 | GLN | A | 315 | 9.182 | 24.223 | 24.766 | 1.00 | 0.00 | H |
| ATOM | 4509 | HE22 | GLN | A | 315 | 7.643 | 24.230 | 23.902 | 1.00 | 0.00 | H |
| ATOM | 4510 | C | GLN | A | 315 | 10.518 | 24.537 | 28.665 | 1.00 | 0.00 | C |
| ATOM | 4511 | O | GLN | A | 315 | 11.506 | 24.896 | 28.023 | 1.00 | 0.00 | O |
| ATOM | 4512 | N | ARG | A | 316 | 10.604 | 23.639 | 29.661 | 1.00 | 0.00 | N |
| ATOM | 4513 | H | ARG | A | 316 | 9.760 | 23.371 | 30.153 | 1.00 | 0.00 | H |
| ATOM | 4514 | CA | ARG | A | 316 | 11.868 | 23.028 | 30.108 | 1.00 | 0.00 | C |
| ATOM | 4515 | HA | ARG | A | 316 | 12.615 | 23.179 | 29.337 | 1.00 | 0.00 | H |
| ATOM | 4516 | CB | ARG | A | 316 | 12.373 | 23.686 | 31.411 | 1.00 | 0.00 | C |
| ATOM | 4517 | HB1 | ARG | A | 316 | 11.735 | 23.370 | 32.235 | 1.00 | 0.00 | H |
| ATOM | 4518 | HB2 | ARG | A | 316 | 13.389 | 23.340 | 31.609 | 1.00 | 0.00 | H |
| ATOM | 4519 | CG | ARG | A | 316 | 12.363 | 25.223 | 31.373 | 1.00 | 0.00 | C |
| ATOM | 4520 | HG1 | ARG | A | 316 | 12.890 | 25.577 | 30.488 | 1.00 | 0.00 | H |
| ATOM | 4521 | HG2 | ARG | A | 316 | 11.327 | 25.548 | 31.307 | 1.00 | 0.00 | H |
| ATOM | 4522 | CD | ARG | A | 316 | 12.956 | 25.874 | 32.631 | 1.00 | 0.00 | C |
| ATOM | 4523 | HD1 | ARG | A | 316 | 12.476 | 26.842 | 32.784 | 1.00 | 0.00 | H |
| ATOM | 4524 | HD2 | ARG | A | 316 | 12.731 | 25.259 | 33.505 | 1.00 | 0.00 | H |
| ATOM | 4525 | NE | ARG | A | 316 | 14.410 | 26.074 | 32.518 | 1.00 | 0.00 | N |
| ATOM | 4526 | HE | ARG | A | 316 | 15.011 | 25.449 | 33.042 | 1.00 | 0.00 | H |
| ATOM | 4527 | CZ | ARG | A | 316 | 15.018 | 27.088 | 31.913 | 1.00 | 0.00 | C |
| ATOM | 4528 | NH1 | ARG | A | 316 | 16.324 | 27.142 | 31.950 | 1.00 | 0.00 | N |
| ATOM | 4529 | HH11 | ARG | A | 316 | 16.828 | 26.482 | 32.530 | 1.00 | 0.00 | H |
| ATOM | 4530 | HH12 | ARG | A | 316 | 16.796 | 27.968 | 31.585 | 1.00 | 0.00 | H |
| ATOM | 4531 | NH2 | ARG | A | 316 | 14.379 | 28.045 | 31.275 | 1.00 | 0.00 | N |
| ATOM | 4532 | HH21 | ARG | A | 316 | 13.367 | 28.035 | 31.211 | 1.00 | 0.00 | H |
| ATOM | 4533 | HH22 | ARG | A | 316 | 14.903 | 28.833 | 30.899 | 1.00 | 0.00 | H |

| | | | | | | | | | | | |
|---|---|---|---|---|---|---|---|---|---|---|---|
| ATOM | 4534 | C | ARG | A | 316 | 11.738 | 21.515 | 30.296 | 1.00 | 0.00 | C |
| ATOM | 4535 | O | ARG | A | 316 | 10.730 | 21.028 | 30.808 | 1.00 | 0.00 | O |
| ATOM | 4536 | N | ILE | A | 317 | 12.769 | 20.755 | 29.921 | 1.00 | 0.00 | N |
| ATOM | 4537 | H | ILE | A | 317 | 13.579 | 21.229 | 29.539 | 1.00 | 0.00 | H |
| ATOM | 4538 | CA | ILE | A | 317 | 12.847 | 19.300 | 30.147 | 1.00 | 0.00 | C |
| ATOM | 4539 | HA | ILE | A | 317 | 11.892 | 18.851 | 29.870 | 1.00 | 0.00 | H |
| ATOM | 4540 | CB | ILE | A | 317 | 13.944 | 18.624 | 29.282 | 1.00 | 0.00 | C |
| ATOM | 4541 | HB | ILE | A | 317 | 14.921 | 18.920 | 29.665 | 1.00 | 0.00 | H |
| ATOM | 4542 | CG2 | ILE | A | 317 | 13.793 | 17.096 | 29.428 | 1.00 | 0.00 | C |
| ATOM | 4543 | HG21 | ILE | A | 317 | 14.589 | 16.581 | 28.895 | 1.00 | 0.00 | H |
| ATOM | 4544 | HG22 | ILE | A | 317 | 13.854 | 16.798 | 30.474 | 1.00 | 0.00 | H |
| ATOM | 4545 | HG23 | ILE | A | 317 | 12.834 | 16.773 | 29.021 | 1.00 | 0.00 | H |
| ATOM | 4546 | CG1 | ILE | A | 317 | 13.888 | 19.049 | 27.796 | 1.00 | 0.00 | C |
| ATOM | 4547 | HG11 | ILE | A | 317 | 12.911 | 18.797 | 27.384 | 1.00 | 0.00 | H |
| ATOM | 4548 | HG12 | ILE | A | 317 | 14.025 | 20.127 | 27.736 | 1.00 | 0.00 | H |
| ATOM | 4549 | CD | ILE | A | 317 | 14.975 | 18.428 | 26.910 | 1.00 | 0.00 | C |
| ATOM | 4550 | HD1 | ILE | A | 317 | 14.943 | 18.894 | 25.925 | 1.00 | 0.00 | H |
| ATOM | 4551 | HD2 | ILE | A | 317 | 15.959 | 18.595 | 27.350 | 1.00 | 0.00 | H |
| ATOM | 4552 | HD3 | ILE | A | 317 | 14.809 | 17.359 | 26.787 | 1.00 | 0.00 | H |
| ATOM | 4553 | C | ILE | A | 317 | 13.087 | 19.050 | 31.642 | 1.00 | 0.00 | C |
| ATOM | 4554 | O | ILE | A | 317 | 14.197 | 19.284 | 32.122 | 1.00 | 0.00 | O |
| ATOM | 4555 | N | GLN | A | 318 | 12.069 | 18.575 | 32.364 | 1.00 | 0.00 | N |
| ATOM | 4556 | H | GLN | A | 318 | 11.179 | 18.433 | 31.894 | 1.00 | 0.00 | H |
| ATOM | 4557 | CA | GLN | A | 318 | 12.078 | 18.446 | 33.830 | 1.00 | 0.00 | C |
| ATOM | 4558 | HA | GLN | A | 318 | 12.303 | 19.428 | 34.253 | 1.00 | 0.00 | H |
| ATOM | 4559 | CB | GLN | A | 318 | 10.672 | 18.037 | 34.313 | 1.00 | 0.00 | C |
| ATOM | 4560 | HB1 | GLN | A | 318 | 10.630 | 18.144 | 35.398 | 1.00 | 0.00 | H |
| ATOM | 4561 | HB2 | GLN | A | 318 | 9.952 | 18.737 | 33.888 | 1.00 | 0.00 | H |
| ATOM | 4562 | CG | GLN | A | 318 | 10.248 | 16.598 | 33.950 | 1.00 | 0.00 | C |
| ATOM | 4563 | HG1 | GLN | A | 318 | 10.406 | 16.417 | 32.887 | 1.00 | 0.00 | H |
| ATOM | 4564 | HG2 | GLN | A | 318 | 10.863 | 15.893 | 34.508 | 1.00 | 0.00 | H |
| ATOM | 4565 | CD | GLN | A | 318 | 8.773 | 16.330 | 34.258 | 1.00 | 0.00 | C |
| ATOM | 4566 | OE1 | GLN | A | 318 | 7.908 | 17.170 | 34.047 | 1.00 | 0.00 | O |
| ATOM | 4567 | NE2 | GLN | A | 318 | 8.411 | 15.163 | 34.737 | 1.00 | 0.00 | N |
| ATOM | 4568 | HE21 | GLN | A | 318 | 9.087 | 14.425 | 34.894 | 1.00 | 0.00 | H |
| ATOM | 4569 | HE22 | GLN | A | 318 | 7.424 | 15.018 | 34.915 | 1.00 | 0.00 | H |
| ATOM | 4570 | C | GLN | A | 318 | 13.151 | 17.483 | 34.359 | 1.00 | 0.00 | C |
| ATOM | 4571 | O | GLN | A | 318 | 13.620 | 17.637 | 35.487 | 1.00 | 0.00 | O |
| ATOM | 4572 | N | ASP | A | 319 | 13.551 | 16.505 | 33.543 | 1.00 | 0.00 | N |
| ATOM | 4573 | H | ASP | A | 319 | 13.057 | 16.398 | 32.670 | 1.00 | 0.00 | H |
| ATOM | 4574 | CA | ASP | A | 319 | 14.606 | 15.538 | 33.846 | 1.00 | 0.00 | C |
| ATOM | 4575 | HA | ASP | A | 319 | 14.534 | 15.230 | 34.888 | 1.00 | 0.00 | H |
| ATOM | 4576 | CB | ASP | A | 319 | 14.426 | 14.312 | 32.934 | 1.00 | 0.00 | C |
| ATOM | 4577 | HB1 | ASP | A | 319 | 14.345 | 14.644 | 31.897 | 1.00 | 0.00 | H |
| ATOM | 4578 | HB2 | ASP | A | 319 | 15.318 | 13.693 | 33.010 | 1.00 | 0.00 | H |
| ATOM | 4579 | CG | ASP | A | 319 | 13.223 | 13.435 | 33.278 | 1.00 | 0.00 | C |
| ATOM | 4580 | OD1 | ASP | A | 319 | 12.764 | 12.714 | 32.360 | 1.00 | 0.00 | O |
| ATOM | 4581 | OD2 | ASP | A | 319 | 12.792 | 13.455 | 34.457 | 1.00 | 0.00 | O |
| ATOM | 4582 | C | ASP | A | 319 | 16.008 | 16.115 | 33.617 | 1.00 | 0.00 | C |
| ATOM | 4583 | O | ASP | A | 319 | 16.920 | 15.878 | 34.410 | 1.00 | 0.00 | O |
| ATOM | 4584 | N | GLN | A | 320 | 16.180 | 16.860 | 32.525 | 1.00 | 0.00 | N |
| ATOM | 4585 | H | GLN | A | 320 | 15.358 | 17.097 | 31.991 | 1.00 | 0.00 | H |
| ATOM | 4586 | CA | GLN | A | 320 | 17.490 | 17.242 | 31.992 | 1.00 | 0.00 | C |
| ATOM | 4587 | HA | GLN | A | 320 | 18.258 | 16.616 | 32.448 | 1.00 | 0.00 | H |

| | | | | | | | | | | | |
|---|---|---|---|---|---|---|---|---|---|---|---|
| ATOM | 4588 | CB | GLN | A | 320 | 17.511 | 16.981 | 30.479 | 1.00 | 0.00 | C |
| ATOM | 4589 | HB1 | GLN | A | 320 | 16.788 | 17.621 | 29.976 | 1.00 | 0.00 | H |
| ATOM | 4590 | HB2 | GLN | A | 320 | 18.500 | 17.238 | 30.112 | 1.00 | 0.00 | H |
| ATOM | 4591 | CG | GLN | A | 320 | 17.212 | 15.509 | 30.144 | 1.00 | 0.00 | C |
| ATOM | 4592 | HG1 | GLN | A | 320 | 17.806 | 14.869 | 30.790 | 1.00 | 0.00 | H |
| ATOM | 4593 | HG2 | GLN | A | 320 | 16.165 | 15.288 | 30.344 | 1.00 | 0.00 | H |
| ATOM | 4594 | CD | GLN | A | 320 | 17.490 | 15.162 | 28.684 | 1.00 | 0.00 | C |
| ATOM | 4595 | OE1 | GLN | A | 320 | 16.805 | 15.629 | 27.788 | 1.00 | 0.00 | O |
| ATOM | 4596 | NE2 | GLN | A | 320 | 18.486 | 14.352 | 28.403 | 1.00 | 0.00 | N |
| ATOM | 4597 | HE21 | GLN | A | 320 | 19.139 | 14.060 | 29.116 | 1.00 | 0.00 | H |
| ATOM | 4598 | HE22 | GLN | A | 320 | 18.617 | 14.046 | 27.436 | 1.00 | 0.00 | H |
| ATOM | 4599 | C | GLN | A | 320 | 17.871 | 18.691 | 32.325 | 1.00 | 0.00 | C |
| ATOM | 4600 | O | GLN | A | 320 | 19.007 | 19.095 | 32.089 | 1.00 | 0.00 | O |
| ATOM | 4601 | N | LYS | A | 321 | 16.940 | 19.465 | 32.895 | 1.00 | 0.00 | N |
| ATOM | 4602 | H | LYS | A | 321 | 16.000 | 19.081 | 32.962 | 1.00 | 0.00 | H |
| ATOM | 4603 | CA | LYS | A | 321 | 17.172 | 20.735 | 33.603 | 1.00 | 0.00 | C |
| ATOM | 4604 | HA | LYS | A | 321 | 16.226 | 21.005 | 34.074 | 1.00 | 0.00 | H |
| ATOM | 4605 | CB | LYS | A | 321 | 18.216 | 20.574 | 34.735 | 1.00 | 0.00 | C |
| ATOM | 4606 | HB1 | LYS | A | 321 | 19.217 | 20.607 | 34.305 | 1.00 | 0.00 | H |
| ATOM | 4607 | HB2 | LYS | A | 321 | 18.116 | 21.435 | 35.396 | 1.00 | 0.00 | H |
| ATOM | 4608 | CG | LYS | A | 321 | 18.137 | 19.303 | 35.601 | 1.00 | 0.00 | C |
| ATOM | 4609 | HG1 | LYS | A | 321 | 18.498 | 18.452 | 35.022 | 1.00 | 0.00 | H |
| ATOM | 4610 | HG2 | LYS | A | 321 | 18.812 | 19.435 | 36.447 | 1.00 | 0.00 | H |
| ATOM | 4611 | CD | LYS | A | 321 | 16.743 | 18.973 | 36.143 | 1.00 | 0.00 | C |
| ATOM | 4612 | HD1 | LYS | A | 321 | 16.354 | 19.824 | 36.699 | 1.00 | 0.00 | H |
| ATOM | 4613 | HD2 | LYS | A | 321 | 16.066 | 18.745 | 35.320 | 1.00 | 0.00 | H |
| ATOM | 4614 | CE | LYS | A | 321 | 16.852 | 17.753 | 37.061 | 1.00 | 0.00 | C |
| ATOM | 4615 | HE1 | LYS | A | 321 | 17.298 | 16.928 | 36.501 | 1.00 | 0.00 | H |
| ATOM | 4616 | HE2 | LYS | A | 321 | 17.515 | 17.991 | 37.899 | 1.00 | 0.00 | H |
| ATOM | 4617 | NZ | LYS | A | 321 | 15.532 | 17.337 | 37.577 | 1.00 | 0.00 | N |
| ATOM | 4618 | HZ1 | LYS | A | 321 | 15.610 | 16.506 | 38.160 | 1.00 | 0.00 | H |
| ATOM | 4619 | HZ2 | LYS | A | 321 | 15.116 | 18.059 | 38.144 | 1.00 | 0.00 | H |
| ATOM | 4620 | HZ3 | LYS | A | 321 | 14.891 | 17.135 | 36.814 | 1.00 | 0.00 | H |
| ATOM | 4621 | C | LYS | A | 321 | 17.499 | 21.900 | 32.646 | 1.00 | 0.00 | C |
| ATOM | 4622 | O | LYS | A | 321 | 18.242 | 22.821 | 32.998 | 1.00 | 0.00 | O |
| ATOM | 4623 | N | VAL | A | 322 | 16.974 | 21.835 | 31.422 | 1.00 | 0.00 | N |
| ATOM | 4624 | H | VAL | A | 322 | 16.383 | 21.033 | 31.254 | 1.00 | 0.00 | H |
| ATOM | 4625 | CA | VAL | A | 322 | 17.307 | 22.701 | 30.269 | 1.00 | 0.00 | C |
| ATOM | 4626 | HA | VAL | A | 322 | 17.741 | 23.623 | 30.653 | 1.00 | 0.00 | H |
| ATOM | 4627 | CB | VAL | A | 322 | 18.337 | 22.022 | 29.333 | 1.00 | 0.00 | C |
| ATOM | 4628 | HB | VAL | A | 322 | 18.421 | 22.639 | 28.443 | 1.00 | 0.00 | H |
| ATOM | 4629 | CG1 | VAL | A | 322 | 19.731 | 21.963 | 29.968 | 1.00 | 0.00 | C |
| ATOM | 4630 | HG11 | VAL | A | 322 | 20.443 | 21.585 | 29.237 | 1.00 | 0.00 | H |
| ATOM | 4631 | HG12 | VAL | A | 322 | 20.043 | 22.962 | 30.267 | 1.00 | 0.00 | H |
| ATOM | 4632 | HG13 | VAL | A | 322 | 19.730 | 21.305 | 30.834 | 1.00 | 0.00 | H |
| ATOM | 4633 | CG2 | VAL | A | 322 | 17.904 | 20.604 | 28.907 | 1.00 | 0.00 | C |
| ATOM | 4634 | HG21 | VAL | A | 322 | 18.670 | 20.155 | 28.279 | 1.00 | 0.00 | H |
| ATOM | 4635 | HG22 | VAL | A | 322 | 17.762 | 19.962 | 29.774 | 1.00 | 0.00 | H |
| ATOM | 4636 | HG23 | VAL | A | 322 | 16.973 | 20.648 | 28.342 | 1.00 | 0.00 | H |
| ATOM | 4637 | C | VAL | A | 322 | 16.056 | 23.055 | 29.439 | 1.00 | 0.00 | C |
| ATOM | 4638 | O | VAL | A | 322 | 15.119 | 22.251 | 29.432 | 1.00 | 0.00 | O |
| ATOM | 4639 | N | PRO | A | 323 | 16.007 | 24.220 | 28.758 | 1.00 | 0.00 | N |
| ATOM | 4640 | CD | PRO | A | 323 | 16.992 | 25.297 | 28.824 | 1.00 | 0.00 | C |
| ATOM | 4641 | HD1 | PRO | A | 323 | 17.968 | 24.968 | 28.472 | 1.00 | 0.00 | H |

| | | | | | | | | | | | |
|---|---|---|---|---|---|---|---|---|---|---|---|
| ATOM | 4642 | HD2 | PRO | A | 323 | 17.074 | 25.650 | 29.851 | 1.00 | 0.00 | H |
| ATOM | 4643 | CG | PRO | A | 323 | 16.456 | 26.429 | 27.943 | 1.00 | 0.00 | C |
| ATOM | 4644 | HG1 | PRO | A | 323 | 16.818 | 26.321 | 26.922 | 1.00 | 0.00 | H |
| ATOM | 4645 | HG2 | PRO | A | 323 | 16.721 | 27.409 | 28.340 | 1.00 | 0.00 | H |
| ATOM | 4646 | CB | PRO | A | 323 | 14.948 | 26.193 | 27.970 | 1.00 | 0.00 | C |
| ATOM | 4647 | HB1 | PRO | A | 323 | 14.443 | 26.619 | 27.107 | 1.00 | 0.00 | H |
| ATOM | 4648 | HB2 | PRO | A | 323 | 14.532 | 26.613 | 28.886 | 1.00 | 0.00 | H |
| ATOM | 4649 | CA | PRO | A | 323 | 14.833 | 24.669 | 28.012 | 1.00 | 0.00 | C |
| ATOM | 4650 | HA | PRO | A | 323 | 13.937 | 24.431 | 28.571 | 1.00 | 0.00 | H |
| ATOM | 4651 | C | PRO | A | 323 | 14.711 | 24.051 | 26.606 | 1.00 | 0.00 | C |
| ATOM | 4652 | O | PRO | A | 323 | 15.695 | 23.629 | 26.000 | 1.00 | 0.00 | O |
| ATOM | 4653 | N | TYR | A | 324 | 13.480 | 24.045 | 26.075 | 1.00 | 0.00 | N |
| ATOM | 4654 | H | TYR | A | 324 | 12.716 | 24.395 | 26.650 | 1.00 | 0.00 | H |
| ATOM | 4655 | CA | TYR | A | 324 | 13.157 | 23.713 | 24.678 | 1.00 | 0.00 | C |
| ATOM | 4656 | HA | TYR | A | 324 | 14.027 | 23.989 | 24.081 | 1.00 | 0.00 | H |
| ATOM | 4657 | CB | TYR | A | 324 | 12.965 | 22.196 | 24.483 | 1.00 | 0.00 | C |
| ATOM | 4658 | HB1 | TYR | A | 324 | 12.982 | 21.995 | 23.412 | 1.00 | 0.00 | H |
| ATOM | 4659 | HB2 | TYR | A | 324 | 13.837 | 21.687 | 24.896 | 1.00 | 0.00 | H |
| ATOM | 4660 | CG | TYR | A | 324 | 11.719 | 21.540 | 25.069 | 1.00 | 0.00 | C |
| ATOM | 4661 | CD1 | TYR | A | 324 | 11.654 | 21.257 | 26.446 | 1.00 | 0.00 | C |
| ATOM | 4662 | HD1 | TYR | A | 324 | 12.444 | 21.605 | 27.093 | 1.00 | 0.00 | H |
| ATOM | 4663 | CE1 | TYR | A | 324 | 10.583 | 20.506 | 26.971 | 1.00 | 0.00 | C |
| ATOM | 4664 | HE1 | TYR | A | 324 | 10.540 | 20.273 | 28.023 | 1.00 | 0.00 | H |
| ATOM | 4665 | CZ | TYR | A | 324 | 9.547 | 20.056 | 26.123 | 1.00 | 0.00 | C |
| ATOM | 4666 | OH | TYR | A | 324 | 8.505 | 19.354 | 26.654 | 1.00 | 0.00 | O |
| ATOM | 4667 | HH | TYR | A | 324 | 7.805 | 19.160 | 26.018 | 1.00 | 0.00 | H |
| ATOM | 4668 | CE2 | TYR | A | 324 | 9.585 | 20.375 | 24.747 | 1.00 | 0.00 | C |
| ATOM | 4669 | HE2 | TYR | A | 324 | 8.792 | 20.047 | 24.089 | 1.00 | 0.00 | H |
| ATOM | 4670 | CD2 | TYR | A | 324 | 10.673 | 21.105 | 24.227 | 1.00 | 0.00 | C |
| ATOM | 4671 | HD2 | TYR | A | 324 | 10.712 | 21.313 | 23.168 | 1.00 | 0.00 | H |
| ATOM | 4672 | C | TYR | A | 324 | 11.978 | 24.549 | 24.133 | 1.00 | 0.00 | C |
| ATOM | 4673 | O | TYR | A | 324 | 11.047 | 24.909 | 24.866 | 1.00 | 0.00 | O |
| ATOM | 4674 | N | ILE | A | 325 | 12.038 | 24.914 | 22.848 | 1.00 | 0.00 | N |
| ATOM | 4675 | H | ILE | A | 325 | 12.755 | 24.496 | 22.262 | 1.00 | 0.00 | H |
| ATOM | 4676 | CA | ILE | A | 325 | 11.171 | 25.915 | 22.202 | 1.00 | 0.00 | C |
| ATOM | 4677 | HA | ILE | A | 325 | 10.213 | 25.905 | 22.715 | 1.00 | 0.00 | H |
| ATOM | 4678 | CB | ILE | A | 325 | 11.792 | 27.329 | 22.366 | 1.00 | 0.00 | C |
| ATOM | 4679 | HB | ILE | A | 325 | 12.012 | 27.466 | 23.425 | 1.00 | 0.00 | H |
| ATOM | 4680 | CG2 | ILE | A | 325 | 13.113 | 27.485 | 21.587 | 1.00 | 0.00 | C |
| ATOM | 4681 | HG21 | ILE | A | 325 | 13.645 | 28.377 | 21.917 | 1.00 | 0.00 | H |
| ATOM | 4682 | HG22 | ILE | A | 325 | 13.760 | 26.628 | 21.764 | 1.00 | 0.00 | H |
| ATOM | 4683 | HG23 | ILE | A | 325 | 12.919 | 27.566 | 20.518 | 1.00 | 0.00 | H |
| ATOM | 4684 | CG1 | ILE | A | 325 | 10.795 | 28.439 | 21.982 | 1.00 | 0.00 | C |
| ATOM | 4685 | HG11 | ILE | A | 325 | 10.528 | 28.338 | 20.933 | 1.00 | 0.00 | H |
| ATOM | 4686 | HG12 | ILE | A | 325 | 9.894 | 28.317 | 22.575 | 1.00 | 0.00 | H |
| ATOM | 4687 | CD | ILE | A | 325 | 11.315 | 29.858 | 22.239 | 1.00 | 0.00 | C |
| ATOM | 4688 | HD1 | ILE | A | 325 | 10.520 | 30.575 | 22.043 | 1.00 | 0.00 | H |
| ATOM | 4689 | HD2 | ILE | A | 325 | 11.623 | 29.952 | 23.280 | 1.00 | 0.00 | H |
| ATOM | 4690 | HD3 | ILE | A | 325 | 12.156 | 30.091 | 21.587 | 1.00 | 0.00 | H |
| ATOM | 4691 | C | ILE | A | 325 | 10.885 | 25.558 | 20.735 | 1.00 | 0.00 | C |
| ATOM | 4692 | O | ILE | A | 325 | 11.788 | 25.167 | 19.990 | 1.00 | 0.00 | O |
| ATOM | 4693 | N | PHE | A | 326 | 9.630 | 25.708 | 20.309 | 1.00 | 0.00 | N |
| ATOM | 4694 | H | PHE | A | 326 | 8.929 | 26.027 | 20.968 | 1.00 | 0.00 | H |
| ATOM | 4695 | CA | PHE | A | 326 | 9.171 | 25.350 | 18.963 | 1.00 | 0.00 | C |

| | | | | | | | | | | | |
|---|---|---|---|---|---|---|---|---|---|---|---|
| ATOM | 4696 | HA | PHE | A | 326 | 10.034 | 25.302 | 18.298 | 1.00 | 0.00 | H |
| ATOM | 4697 | CB | PHE | A | 326 | 8.548 | 23.940 | 18.989 | 1.00 | 0.00 | C |
| ATOM | 4698 | HB1 | PHE | A | 326 | 8.084 | 23.731 | 18.024 | 1.00 | 0.00 | H |
| ATOM | 4699 | HB2 | PHE | A | 326 | 9.362 | 23.224 | 19.103 | 1.00 | 0.00 | H |
| ATOM | 4700 | CG | PHE | A | 326 | 7.527 | 23.670 | 20.082 | 1.00 | 0.00 | C |
| ATOM | 4701 | CD1 | PHE | A | 326 | 6.223 | 24.190 | 19.987 | 1.00 | 0.00 | C |
| ATOM | 4702 | HD1 | PHE | A | 326 | 5.946 | 24.807 | 19.146 | 1.00 | 0.00 | H |
| ATOM | 4703 | CE1 | PHE | A | 326 | 5.273 | 23.907 | 20.985 | 1.00 | 0.00 | C |
| ATOM | 4704 | HE1 | PHE | A | 326 | 4.273 | 24.312 | 20.911 | 1.00 | 0.00 | H |
| ATOM | 4705 | CZ | PHE | A | 326 | 5.623 | 23.102 | 22.082 | 1.00 | 0.00 | C |
| ATOM | 4706 | HZ | PHE | A | 326 | 4.896 | 22.887 | 22.854 | 1.00 | 0.00 | H |
| ATOM | 4707 | CE2 | PHE | A | 326 | 6.920 | 22.571 | 22.175 | 1.00 | 0.00 | C |
| ATOM | 4708 | HE2 | PHE | A | 326 | 7.182 | 21.935 | 23.008 | 1.00 | 0.00 | H |
| ATOM | 4709 | CD2 | PHE | A | 326 | 7.873 | 22.861 | 21.183 | 1.00 | 0.00 | C |
| ATOM | 4710 | HD2 | PHE | A | 326 | 8.868 | 22.447 | 21.264 | 1.00 | 0.00 | H |
| ATOM | 4711 | C | PHE | A | 326 | 8.238 | 26.408 | 18.353 | 1.00 | 0.00 | C |
| ATOM | 4712 | O | PHE | A | 326 | 7.664 | 27.242 | 19.061 | 1.00 | 0.00 | O |
| ATOM | 4713 | N | ARG | A | 327 | 8.104 | 26.371 | 17.021 | 1.00 | 0.00 | N |
| ATOM | 4714 | H | ARG | A | 327 | 8.615 | 25.642 | 16.527 | 1.00 | 0.00 | H |
| ATOM | 4715 | CA | ARG | A | 327 | 7.309 | 27.298 | 16.210 | 1.00 | 0.00 | C |
| ATOM | 4716 | HA | ARG | A | 327 | 6.378 | 27.499 | 16.743 | 1.00 | 0.00 | H |
| ATOM | 4717 | CB | ARG | A | 327 | 8.070 | 28.629 | 16.048 | 1.00 | 0.00 | C |
| ATOM | 4718 | HB1 | ARG | A | 327 | 8.399 | 28.979 | 17.027 | 1.00 | 0.00 | H |
| ATOM | 4719 | HB2 | ARG | A | 327 | 8.955 | 28.474 | 15.433 | 1.00 | 0.00 | H |
| ATOM | 4720 | CG | ARG | A | 327 | 7.173 | 29.707 | 15.424 | 1.00 | 0.00 | C |
| ATOM | 4721 | HG1 | ARG | A | 327 | 6.823 | 29.380 | 14.446 | 1.00 | 0.00 | H |
| ATOM | 4722 | HG2 | ARG | A | 327 | 6.307 | 29.832 | 16.073 | 1.00 | 0.00 | H |
| ATOM | 4723 | CD | ARG | A | 327 | 7.859 | 31.072 | 15.283 | 1.00 | 0.00 | C |
| ATOM | 4724 | HD1 | ARG | A | 327 | 7.095 | 31.807 | 15.026 | 1.00 | 0.00 | H |
| ATOM | 4725 | HD2 | ARG | A | 327 | 8.304 | 31.350 | 16.240 | 1.00 | 0.00 | H |
| ATOM | 4726 | NE | ARG | A | 327 | 8.891 | 31.083 | 14.233 | 1.00 | 0.00 | N |
| ATOM | 4727 | HE | ARG | A | 327 | 9.044 | 30.212 | 13.737 | 1.00 | 0.00 | H |
| ATOM | 4728 | CZ | ARG | A | 327 | 9.641 | 32.120 | 13.877 | 1.00 | 0.00 | C |
| ATOM | 4729 | NH1 | ARG | A | 327 | 10.548 | 31.968 | 12.936 | 1.00 | 0.00 | N |
| ATOM | 4730 | HH11 | ARG | A | 327 | 10.671 | 31.073 | 12.471 | 1.00 | 0.00 | H |
| ATOM | 4731 | HH12 | ARG | A | 327 | 11.146 | 32.732 | 12.662 | 1.00 | 0.00 | H |
| ATOM | 4732 | NH2 | ARG | A | 327 | 9.522 | 33.299 | 14.454 | 1.00 | 0.00 | N |
| ATOM | 4733 | HH21 | ARG | A | 327 | 8.797 | 33.456 | 15.145 | 1.00 | 0.00 | H |
| ATOM | 4734 | HH22 | ARG | A | 327 | 10.149 | 34.065 | 14.242 | 1.00 | 0.00 | H |
| ATOM | 4735 | C | ARG | A | 327 | 6.969 | 26.657 | 14.855 | 1.00 | 0.00 | C |
| ATOM | 4736 | O | ARG | A | 327 | 7.859 | 26.390 | 14.042 | 1.00 | 0.00 | O |
| ATOM | 4737 | N | ASP | A | 328 | 5.680 | 26.412 | 14.617 | 1.00 | 0.00 | N |
| ATOM | 4738 | H | ASP | A | 328 | 5.029 | 26.665 | 15.353 | 1.00 | 0.00 | H |
| ATOM | 4739 | CA | ASP | A | 328 | 5.141 | 25.606 | 13.511 | 1.00 | 0.00 | C |
| ATOM | 4740 | HA | ASP | A | 328 | 4.117 | 25.335 | 13.774 | 1.00 | 0.00 | H |
| ATOM | 4741 | CB | ASP | A | 328 | 5.059 | 26.439 | 12.213 | 1.00 | 0.00 | C |
| ATOM | 4742 | HB1 | ASP | A | 328 | 6.039 | 26.859 | 11.978 | 1.00 | 0.00 | H |
| ATOM | 4743 | HB2 | ASP | A | 328 | 4.761 | 25.788 | 11.391 | 1.00 | 0.00 | H |
| ATOM | 4744 | CG | ASP | A | 328 | 4.018 | 27.558 | 12.306 | 1.00 | 0.00 | C |
| ATOM | 4745 | OD1 | ASP | A | 328 | 2.922 | 27.321 | 12.866 | 1.00 | 0.00 | O |
| ATOM | 4746 | OD2 | ASP | A | 328 | 4.266 | 28.673 | 11.797 | 1.00 | 0.00 | O |
| ATOM | 4747 | C | ASP | A | 328 | 5.876 | 24.255 | 13.391 | 1.00 | 0.00 | C |
| ATOM | 4748 | O | ASP | A | 328 | 5.871 | 23.483 | 14.351 | 1.00 | 0.00 | O |
| ATOM | 4749 | N | ASN | A | 329 | 6.511 | 23.961 | 12.253 | 1.00 | 0.00 | N |

| | | | | | | | | | | |
|---|---|---|---|---|---|---|---|---|---|---|
| ATOM | 4750 | H | ASN | A | 329 | 6.464 | 24.622 | 11.489 | 1.00 | 0.00 | H |
| ATOM | 4751 | CA | ASN | A | 329 | 7.295 | 22.739 | 12.043 | 1.00 | 0.00 | C |
| ATOM | 4752 | HA | ASN | A | 329 | 6.817 | 21.939 | 12.609 | 1.00 | 0.00 | H |
| ATOM | 4753 | CB | ASN | A | 329 | 7.229 | 22.337 | 10.557 | 1.00 | 0.00 | C |
| ATOM | 4754 | HB1 | ASN | A | 329 | 7.559 | 21.300 | 10.475 | 1.00 | 0.00 | H |
| ATOM | 4755 | HB2 | ASN | A | 329 | 6.197 | 22.373 | 10.208 | 1.00 | 0.00 | H |
| ATOM | 4756 | CG | ASN | A | 329 | 8.101 | 23.188 | 9.633 | 1.00 | 0.00 | C |
| ATOM | 4757 | OD1 | ASN | A | 329 | 8.435 | 24.337 | 9.908 | 1.00 | 0.00 | O |
| ATOM | 4758 | ND2 | ASN | A | 329 | 8.530 | 22.637 | 8.517 | 1.00 | 0.00 | N |
| ATOM | 4759 | HD21 | ASN | A | 329 | 8.282 | 21.678 | 8.288 | 1.00 | 0.00 | H |
| ATOM | 4760 | HD22 | ASN | A | 329 | 9.125 | 23.173 | 7.898 | 1.00 | 0.00 | H |
| ATOM | 4761 | C | ASN | A | 329 | 8.757 | 22.824 | 12.542 | 1.00 | 0.00 | C |
| ATOM | 4762 | O | ASN | A | 329 | 9.520 | 21.885 | 12.321 | 1.00 | 0.00 | O |
| ATOM | 4763 | N | GLN | A | 330 | 9.184 | 23.926 | 13.167 | 1.00 | 0.00 | N |
| ATOM | 4764 | H | GLN | A | 330 | 8.509 | 24.651 | 13.386 | 1.00 | 0.00 | H |
| ATOM | 4765 | CA | GLN | A | 330 | 10.555 | 24.107 | 13.659 | 1.00 | 0.00 | C |
| ATOM | 4766 | HA | GLN | A | 330 | 11.214 | 23.370 | 13.196 | 1.00 | 0.00 | H |
| ATOM | 4767 | CB | GLN | A | 330 | 11.076 | 25.509 | 13.293 | 1.00 | 0.00 | C |
| ATOM | 4768 | HB1 | GLN | A | 330 | 10.449 | 26.264 | 13.765 | 1.00 | 0.00 | H |
| ATOM | 4769 | HB2 | GLN | A | 330 | 12.082 | 25.620 | 13.700 | 1.00 | 0.00 | H |
| ATOM | 4770 | CG | GLN | A | 330 | 11.130 | 25.776 | 11.777 | 1.00 | 0.00 | C |
| ATOM | 4771 | HG1 | GLN | A | 330 | 10.123 | 25.762 | 11.363 | 1.00 | 0.00 | H |
| ATOM | 4772 | HG2 | GLN | A | 330 | 11.544 | 26.768 | 11.608 | 1.00 | 0.00 | H |
| ATOM | 4773 | CD | GLN | A | 330 | 12.013 | 24.769 | 11.047 | 1.00 | 0.00 | C |
| ATOM | 4774 | OE1 | GLN | A | 330 | 13.210 | 24.694 | 11.289 | 1.00 | 0.00 | O |
| ATOM | 4775 | NE2 | GLN | A | 330 | 11.469 | 23.933 | 10.191 | 1.00 | 0.00 | N |
| ATOM | 4776 | HE21 | GLN | A | 330 | 10.497 | 24.027 | 9.931 | 1.00 | 0.00 | H |
| ATOM | 4777 | HE22 | GLN | A | 330 | 12.078 | 23.265 | 9.738 | 1.00 | 0.00 | H |
| ATOM | 4778 | C | GLN | A | 330 | 10.626 | 23.876 | 15.174 | 1.00 | 0.00 | C |
| ATOM | 4779 | O | GLN | A | 330 | 9.728 | 24.270 | 15.915 | 1.00 | 0.00 | O |
| ATOM | 4780 | N | TRP | A | 331 | 11.710 | 23.245 | 15.631 | 1.00 | 0.00 | N |
| ATOM | 4781 | H | TRP | A | 331 | 12.412 | 22.956 | 14.960 | 1.00 | 0.00 | H |
| ATOM | 4782 | CA | TRP | A | 331 | 11.912 | 22.794 | 17.012 | 1.00 | 0.00 | C |
| ATOM | 4783 | HA | TRP | A | 331 | 11.240 | 23.327 | 17.685 | 1.00 | 0.00 | H |
| ATOM | 4784 | CB | TRP | A | 331 | 11.591 | 21.290 | 17.042 | 1.00 | 0.00 | C |
| ATOM | 4785 | HB1 | TRP | A | 331 | 10.665 | 21.130 | 16.487 | 1.00 | 0.00 | H |
| ATOM | 4786 | HB2 | TRP | A | 331 | 12.383 | 20.762 | 16.513 | 1.00 | 0.00 | H |
| ATOM | 4787 | CG | TRP | A | 331 | 11.422 | 20.641 | 18.380 | 1.00 | 0.00 | C |
| ATOM | 4788 | CD1 | TRP | A | 331 | 10.227 | 20.356 | 18.940 | 1.00 | 0.00 | C |
| ATOM | 4789 | HD1 | TRP | A | 331 | 9.263 | 20.575 | 18.496 | 1.00 | 0.00 | H |
| ATOM | 4790 | NE1 | TRP | A | 331 | 10.415 | 19.723 | 20.147 | 1.00 | 0.00 | N |
| ATOM | 4791 | HE1 | TRP | A | 331 | 9.635 | 19.399 | 20.721 | 1.00 | 0.00 | H |
| ATOM | 4792 | CE2 | TRP | A | 331 | 11.749 | 19.575 | 20.447 | 1.00 | 0.00 | C |
| ATOM | 4793 | CZ2 | TRP | A | 331 | 12.423 | 19.062 | 21.563 | 1.00 | 0.00 | C |
| ATOM | 4794 | HZ2 | TRP | A | 331 | 11.865 | 18.649 | 22.390 | 1.00 | 0.00 | H |
| ATOM | 4795 | CH2 | TRP | A | 331 | 13.826 | 19.107 | 21.594 | 1.00 | 0.00 | C |
| ATOM | 4796 | HH2 | TRP | A | 331 | 14.364 | 18.714 | 22.446 | 1.00 | 0.00 | H |
| ATOM | 4797 | CZ3 | TRP | A | 331 | 14.529 | 19.657 | 20.507 | 1.00 | 0.00 | C |
| ATOM | 4798 | HZ3 | TRP | A | 331 | 15.609 | 19.683 | 20.527 | 1.00 | 0.00 | H |
| ATOM | 4799 | CE3 | TRP | A | 331 | 13.842 | 20.164 | 19.386 | 1.00 | 0.00 | C |
| ATOM | 4800 | HE3 | TRP | A | 331 | 14.406 | 20.575 | 18.561 | 1.00 | 0.00 | H |
| ATOM | 4801 | CD2 | TRP | A | 331 | 12.430 | 20.152 | 19.329 | 1.00 | 0.00 | C |
| ATOM | 4802 | C | TRP | A | 331 | 13.366 | 23.043 | 17.435 | 1.00 | 0.00 | C |
| ATOM | 4803 | O | TRP | A | 331 | 14.273 | 22.708 | 16.671 | 1.00 | 0.00 | O |

| | | | | | | | | | | | |
|---|---|---|---|---|---|---|---|---|---|---|---|
| ATOM | 4804 | N | VAL | A | 332 | 13.605 | 23.601 | 18.627 | 1.00 | 0.00 | N |
| ATOM | 4805 | H | VAL | A | 332 | 12.813 | 23.944 | 19.184 | 1.00 | 0.00 | H |
| ATOM | 4806 | CA | VAL | A | 332 | 14.957 | 23.953 | 19.097 | 1.00 | 0.00 | C |
| ATOM | 4807 | HA | VAL | A | 332 | 15.678 | 23.452 | 18.457 | 1.00 | 0.00 | H |
| ATOM | 4808 | CB | VAL | A | 332 | 15.223 | 25.477 | 18.993 | 1.00 | 0.00 | C |
| ATOM | 4809 | HB | VAL | A | 332 | 14.600 | 25.987 | 19.722 | 1.00 | 0.00 | H |
| ATOM | 4810 | CG1 | VAL | A | 332 | 16.693 | 25.816 | 19.301 | 1.00 | 0.00 | C |
| ATOM | 4811 | HG11 | VAL | A | 332 | 16.843 | 26.894 | 19.263 | 1.00 | 0.00 | H |
| ATOM | 4812 | HG12 | VAL | A | 332 | 16.959 | 25.477 | 20.299 | 1.00 | 0.00 | H |
| ATOM | 4813 | HG13 | VAL | A | 332 | 17.351 | 25.342 | 18.573 | 1.00 | 0.00 | H |
| ATOM | 4814 | CG2 | VAL | A | 332 | 14.887 | 26.049 | 17.602 | 1.00 | 0.00 | C |
| ATOM | 4815 | HG21 | VAL | A | 332 | 15.111 | 27.116 | 17.578 | 1.00 | 0.00 | H |
| ATOM | 4816 | HG22 | VAL | A | 332 | 15.473 | 25.543 | 16.835 | 1.00 | 0.00 | H |
| ATOM | 4817 | HG23 | VAL | A | 332 | 13.827 | 25.926 | 17.385 | 1.00 | 0.00 | H |
| ATOM | 4818 | C | VAL | A | 332 | 15.190 | 23.468 | 20.534 | 1.00 | 0.00 | C |
| ATOM | 4819 | O | VAL | A | 332 | 14.365 | 23.690 | 21.422 | 1.00 | 0.00 | O |
| ATOM | 4820 | N | GLY | A | 333 | 16.346 | 22.831 | 20.760 | 1.00 | 0.00 | N |
| ATOM | 4821 | H | GLY | A | 333 | 16.958 | 22.700 | 19.960 | 1.00 | 0.00 | H |
| ATOM | 4822 | CA | GLY | A | 333 | 16.911 | 22.555 | 22.089 | 1.00 | 0.00 | C |
| ATOM | 4823 | HA1 | GLY | A | 333 | 16.163 | 22.694 | 22.871 | 1.00 | 0.00 | H |
| ATOM | 4824 | HA2 | GLY | A | 333 | 17.289 | 21.534 | 22.125 | 1.00 | 0.00 | H |
| ATOM | 4825 | C | GLY | A | 333 | 18.095 | 23.486 | 22.348 | 1.00 | 0.00 | C |
| ATOM | 4826 | O | GLY | A | 333 | 18.978 | 23.588 | 21.497 | 1.00 | 0.00 | O |
| ATOM | 4827 | N | PHE | A | 334 | 18.108 | 24.175 | 23.492 | 1.00 | 0.00 | N |
| ATOM | 4828 | H | PHE | A | 334 | 17.371 | 24.017 | 24.167 | 1.00 | 0.00 | H |
| ATOM | 4829 | CA | PHE | A | 334 | 19.000 | 25.313 | 23.757 | 1.00 | 0.00 | C |
| ATOM | 4830 | HA | PHE | A | 334 | 19.974 | 25.107 | 23.312 | 1.00 | 0.00 | H |
| ATOM | 4831 | CB | PHE | A | 334 | 18.414 | 26.564 | 23.074 | 1.00 | 0.00 | C |
| ATOM | 4832 | HB1 | PHE | A | 334 | 19.191 | 27.325 | 23.016 | 1.00 | 0.00 | H |
| ATOM | 4833 | HB2 | PHE | A | 334 | 18.142 | 26.325 | 22.049 | 1.00 | 0.00 | H |
| ATOM | 4834 | CG | PHE | A | 334 | 17.196 | 27.155 | 23.761 | 1.00 | 0.00 | C |
| ATOM | 4835 | CD1 | PHE | A | 334 | 15.949 | 26.507 | 23.693 | 1.00 | 0.00 | C |
| ATOM | 4836 | HD1 | PHE | A | 334 | 15.840 | 25.587 | 23.140 | 1.00 | 0.00 | H |
| ATOM | 4837 | CE1 | PHE | A | 334 | 14.835 | 27.058 | 24.348 | 1.00 | 0.00 | C |
| ATOM | 4838 | HE1 | PHE | A | 334 | 13.887 | 26.552 | 24.311 | 1.00 | 0.00 | H |
| ATOM | 4839 | CZ | PHE | A | 334 | 14.951 | 28.270 | 25.044 | 1.00 | 0.00 | C |
| ATOM | 4840 | HZ | PHE | A | 334 | 14.086 | 28.707 | 25.513 | 1.00 | 0.00 | H |
| ATOM | 4841 | CE2 | PHE | A | 334 | 16.199 | 28.903 | 25.139 | 1.00 | 0.00 | C |
| ATOM | 4842 | HE2 | PHE | A | 334 | 16.303 | 29.810 | 25.713 | 1.00 | 0.00 | H |
| ATOM | 4843 | CD2 | PHE | A | 334 | 17.316 | 28.353 | 24.489 | 1.00 | 0.00 | C |
| ATOM | 4844 | HD2 | PHE | A | 334 | 18.271 | 28.850 | 24.553 | 1.00 | 0.00 | H |
| ATOM | 4845 | C | PHE | A | 334 | 19.241 | 25.548 | 25.261 | 1.00 | 0.00 | C |
| ATOM | 4846 | O | PHE | A | 334 | 18.713 | 24.820 | 26.098 | 1.00 | 0.00 | O |
| ATOM | 4847 | N | ASP | A | 335 | 20.018 | 26.576 | 25.605 | 1.00 | 0.00 | N |
| ATOM | 4848 | H | ASP | A | 335 | 20.404 | 27.150 | 24.862 | 1.00 | 0.00 | H |
| ATOM | 4849 | CA | ASP | A | 335 | 20.320 | 26.991 | 26.981 | 1.00 | 0.00 | C |
| ATOM | 4850 | HA | ASP | A | 335 | 19.766 | 26.373 | 27.688 | 1.00 | 0.00 | H |
| ATOM | 4851 | CB | ASP | A | 335 | 21.822 | 26.804 | 27.241 | 1.00 | 0.00 | C |
| ATOM | 4852 | HB1 | ASP | A | 335 | 22.394 | 27.014 | 26.335 | 1.00 | 0.00 | H |
| ATOM | 4853 | HB2 | ASP | A | 335 | 22.135 | 27.523 | 27.999 | 1.00 | 0.00 | H |
| ATOM | 4854 | CG | ASP | A | 335 | 22.148 | 25.408 | 27.765 | 1.00 | 0.00 | C |
| ATOM | 4855 | OD1 | ASP | A | 335 | 22.548 | 24.524 | 26.973 | 1.00 | 0.00 | O |
| ATOM | 4856 | OD2 | ASP | A | 335 | 22.036 | 25.213 | 28.998 | 1.00 | 0.00 | O |
| ATOM | 4857 | C | ASP | A | 335 | 19.921 | 28.454 | 27.250 | 1.00 | 0.00 | C |

| | | | | | | | | | | | |
|---|---|---|---|---|---|---|---|---|---|---|---|
| ATOM | 4858 | O | ASP | A | 335 | 19.926 | 29.289 | 26.344 | 1.00 | 0.00 | O |
| ATOM | 4859 | N | ASP | A | 336 | 19.627 | 28.779 | 28.512 | 1.00 | 0.00 | N |
| ATOM | 4860 | H | ASP | A | 336 | 19.751 | 28.075 | 29.236 | 1.00 | 0.00 | H |
| ATOM | 4861 | CA | ASP | A | 336 | 19.345 | 30.143 | 28.970 | 1.00 | 0.00 | C |
| ATOM | 4862 | HA | ASP | A | 336 | 19.843 | 30.821 | 28.280 | 1.00 | 0.00 | H |
| ATOM | 4863 | CB | ASP | A | 336 | 17.830 | 30.442 | 28.852 | 1.00 | 0.00 | C |
| ATOM | 4864 | HB1 | ASP | A | 336 | 17.715 | 31.516 | 28.723 | 1.00 | 0.00 | H |
| ATOM | 4865 | HB2 | ASP | A | 336 | 17.454 | 29.992 | 27.932 | 1.00 | 0.00 | H |
| ATOM | 4866 | CG | ASP | A | 336 | 16.918 | 30.008 | 30.009 | 1.00 | 0.00 | C |
| ATOM | 4867 | OD1 | ASP | A | 336 | 17.394 | 29.509 | 31.052 | 1.00 | 0.00 | O |
| ATOM | 4868 | OD2 | ASP | A | 336 | 15.684 | 30.213 | 29.882 | 1.00 | 0.00 | O |
| ATOM | 4869 | C | ASP | A | 336 | 19.974 | 30.429 | 30.351 | 1.00 | 0.00 | C |
| ATOM | 4870 | O | ASP | A | 336 | 20.673 | 29.577 | 30.904 | 1.00 | 0.00 | O |
| ATOM | 4871 | N | VAL | A | 337 | 19.735 | 31.617 | 30.921 | 1.00 | 0.00 | N |
| ATOM | 4872 | H | VAL | A | 337 | 19.120 | 32.277 | 30.444 | 1.00 | 0.00 | H |
| ATOM | 4873 | CA | VAL | A | 337 | 20.370 | 32.059 | 32.180 | 1.00 | 0.00 | C |
| ATOM | 4874 | HA | VAL | A | 337 | 21.445 | 32.108 | 31.998 | 1.00 | 0.00 | H |
| ATOM | 4875 | CB | VAL | A | 337 | 19.890 | 33.487 | 32.549 | 1.00 | 0.00 | C |
| ATOM | 4876 | HB | VAL | A | 337 | 18.809 | 33.531 | 32.418 | 1.00 | 0.00 | H |
| ATOM | 4877 | CG1 | VAL | A | 337 | 20.178 | 33.928 | 33.992 | 1.00 | 0.00 | C |
| ATOM | 4878 | HG11 | VAL | A | 337 | 19.845 | 34.956 | 34.142 | 1.00 | 0.00 | H |
| ATOM | 4879 | HG12 | VAL | A | 337 | 19.636 | 33.295 | 34.692 | 1.00 | 0.00 | H |
| ATOM | 4880 | HG13 | VAL | A | 337 | 21.245 | 33.872 | 34.194 | 1.00 | 0.00 | H |
| ATOM | 4881 | CG2 | VAL | A | 337 | 20.556 | 34.511 | 31.610 | 1.00 | 0.00 | C |
| ATOM | 4882 | HG21 | VAL | A | 337 | 20.167 | 35.508 | 31.810 | 1.00 | 0.00 | H |
| ATOM | 4883 | HG22 | VAL | A | 337 | 21.635 | 34.513 | 31.768 | 1.00 | 0.00 | H |
| ATOM | 4884 | HG23 | VAL | A | 337 | 20.358 | 34.268 | 30.567 | 1.00 | 0.00 | H |
| ATOM | 4885 | C | VAL | A | 337 | 20.157 | 31.068 | 33.340 | 1.00 | 0.00 | C |
| ATOM | 4886 | O | VAL | A | 337 | 21.061 | 30.913 | 34.159 | 1.00 | 0.00 | O |
| ATOM | 4887 | N | GLU | A | 338 | 19.023 | 30.358 | 33.406 | 1.00 | 0.00 | N |
| ATOM | 4888 | H | GLU | A | 338 | 18.362 | 30.411 | 32.635 | 1.00 | 0.00 | H |
| ATOM | 4889 | CA | GLU | A | 338 | 18.728 | 29.439 | 34.514 | 1.00 | 0.00 | C |
| ATOM | 4890 | HA | GLU | A | 338 | 19.124 | 29.872 | 35.431 | 1.00 | 0.00 | H |
| ATOM | 4891 | CB | GLU | A | 338 | 17.206 | 29.330 | 34.692 | 1.00 | 0.00 | C |
| ATOM | 4892 | HB1 | GLU | A | 338 | 16.769 | 30.327 | 34.638 | 1.00 | 0.00 | H |
| ATOM | 4893 | HB2 | GLU | A | 338 | 16.798 | 28.733 | 33.883 | 1.00 | 0.00 | H |
| ATOM | 4894 | CG | GLU | A | 338 | 16.817 | 28.708 | 36.041 | 1.00 | 0.00 | C |
| ATOM | 4895 | HG1 | GLU | A | 338 | 17.443 | 27.839 | 36.239 | 1.00 | 0.00 | H |
| ATOM | 4896 | HG2 | GLU | A | 338 | 16.992 | 29.436 | 36.835 | 1.00 | 0.00 | H |
| ATOM | 4897 | CD | GLU | A | 338 | 15.364 | 28.245 | 36.095 | 1.00 | 0.00 | C |
| ATOM | 4898 | OE1 | GLU | A | 338 | 15.095 | 27.296 | 36.874 | 1.00 | 0.00 | O |
| ATOM | 4899 | OE2 | GLU | A | 338 | 14.498 | 28.781 | 35.366 | 1.00 | 0.00 | O |
| ATOM | 4900 | C | GLU | A | 338 | 19.382 | 28.055 | 34.325 | 1.00 | 0.00 | C |
| ATOM | 4901 | O | GLU | A | 338 | 19.772 | 27.409 | 35.304 | 1.00 | 0.00 | O |
| ATOM | 4902 | N | SER | A | 339 | 19.575 | 27.586 | 33.087 | 1.00 | 0.00 | N |
| ATOM | 4903 | H | SER | A | 339 | 19.367 | 28.178 | 32.286 | 1.00 | 0.00 | H |
| ATOM | 4904 | CA | SER | A | 339 | 20.344 | 26.354 | 32.862 | 1.00 | 0.00 | C |
| ATOM | 4905 | HA | SER | A | 339 | 20.068 | 25.628 | 33.625 | 1.00 | 0.00 | H |
| ATOM | 4906 | CB | SER | A | 339 | 20.006 | 25.697 | 31.519 | 1.00 | 0.00 | C |
| ATOM | 4907 | HB1 | SER | A | 339 | 20.587 | 24.779 | 31.432 | 1.00 | 0.00 | H |
| ATOM | 4908 | HB2 | SER | A | 339 | 18.947 | 25.436 | 31.513 | 1.00 | 0.00 | H |
| ATOM | 4909 | OG | SER | A | 339 | 20.279 | 26.518 | 30.405 | 1.00 | 0.00 | O |
| ATOM | 4910 | HG | SER | A | 339 | 20.975 | 26.060 | 29.872 | 1.00 | 0.00 | H |
| ATOM | 4911 | C | SER | A | 339 | 21.849 | 26.597 | 33.038 | 1.00 | 0.00 | C |

| | | | | | | | | | | | |
|---|---|---|---|---|---|---|---|---|---|---|---|
| ATOM | 4912 | O | SER | A | 339 | 22.516 | 25.793 | 33.688 | 1.00 | 0.00 | O |
| ATOM | 4913 | N | PHE | A | 340 | 22.374 | 27.745 | 32.591 | 1.00 | 0.00 | N |
| ATOM | 4914 | H | PHE | A | 340 | 21.782 | 28.366 | 32.045 | 1.00 | 0.00 | H |
| ATOM | 4915 | CA | PHE | A | 340 | 23.746 | 28.179 | 32.883 | 1.00 | 0.00 | C |
| ATOM | 4916 | HA | PHE | A | 340 | 24.440 | 27.433 | 32.492 | 1.00 | 0.00 | H |
| ATOM | 4917 | CB | PHE | A | 340 | 24.021 | 29.521 | 32.176 | 1.00 | 0.00 | C |
| ATOM | 4918 | HB1 | PHE | A | 340 | 23.095 | 30.091 | 32.079 | 1.00 | 0.00 | H |
| ATOM | 4919 | HB2 | PHE | A | 340 | 24.685 | 30.110 | 32.810 | 1.00 | 0.00 | H |
| ATOM | 4920 | CG | PHE | A | 340 | 24.674 | 29.402 | 30.807 | 1.00 | 0.00 | C |
| ATOM | 4921 | CD1 | PHE | A | 340 | 24.222 | 28.461 | 29.860 | 1.00 | 0.00 | C |
| ATOM | 4922 | HD1 | PHE | A | 340 | 23.377 | 27.824 | 30.080 | 1.00 | 0.00 | H |
| ATOM | 4923 | CE1 | PHE | A | 340 | 24.864 | 28.348 | 28.614 | 1.00 | 0.00 | C |
| ATOM | 4924 | HE1 | PHE | A | 340 | 24.515 | 27.626 | 27.891 | 1.00 | 0.00 | H |
| ATOM | 4925 | CZ | PHE | A | 340 | 25.954 | 29.176 | 28.301 | 1.00 | 0.00 | C |
| ATOM | 4926 | HZ | PHE | A | 340 | 26.437 | 29.092 | 27.338 | 1.00 | 0.00 | H |
| ATOM | 4927 | CE2 | PHE | A | 340 | 26.411 | 30.114 | 29.240 | 1.00 | 0.00 | C |
| ATOM | 4928 | HE2 | PHE | A | 340 | 27.240 | 30.761 | 28.995 | 1.00 | 0.00 | H |
| ATOM | 4929 | CD2 | PHE | A | 340 | 25.760 | 30.236 | 30.481 | 1.00 | 0.00 | C |
| ATOM | 4930 | HD2 | PHE | A | 340 | 26.092 | 30.983 | 31.185 | 1.00 | 0.00 | H |
| ATOM | 4931 | C | PHE | A | 340 | 24.003 | 28.267 | 34.397 | 1.00 | 0.00 | C |
| ATOM | 4932 | O | PHE | A | 340 | 25.035 | 27.788 | 34.880 | 1.00 | 0.00 | O |
| ATOM | 4933 | N | LYS | A | 341 | 23.063 | 28.823 | 35.172 | 1.00 | 0.00 | N |
| ATOM | 4934 | H | LYS | A | 341 | 22.260 | 29.250 | 34.721 | 1.00 | 0.00 | H |
| ATOM | 4935 | CA | LYS | A | 341 | 23.109 | 28.807 | 36.642 | 1.00 | 0.00 | C |
| ATOM | 4936 | HA | LYS | A | 341 | 24.017 | 29.305 | 36.988 | 1.00 | 0.00 | H |
| ATOM | 4937 | CB | LYS | A | 341 | 21.893 | 29.563 | 37.204 | 1.00 | 0.00 | C |
| ATOM | 4938 | HB1 | LYS | A | 341 | 21.002 | 29.292 | 36.639 | 1.00 | 0.00 | H |
| ATOM | 4939 | HB2 | LYS | A | 341 | 21.734 | 29.277 | 38.244 | 1.00 | 0.00 | H |
| ATOM | 4940 | CG | LYS | A | 341 | 22.109 | 31.080 | 37.150 | 1.00 | 0.00 | C |
| ATOM | 4941 | HG1 | LYS | A | 341 | 22.811 | 31.363 | 37.927 | 1.00 | 0.00 | H |
| ATOM | 4942 | HG2 | LYS | A | 341 | 22.524 | 31.362 | 36.185 | 1.00 | 0.00 | H |
| ATOM | 4943 | CD | LYS | A | 341 | 20.803 | 31.835 | 37.379 | 1.00 | 0.00 | C |
| ATOM | 4944 | HD1 | LYS | A | 341 | 20.080 | 31.525 | 36.626 | 1.00 | 0.00 | H |
| ATOM | 4945 | HD2 | LYS | A | 341 | 20.414 | 31.588 | 38.365 | 1.00 | 0.00 | H |
| ATOM | 4946 | CE | LYS | A | 341 | 21.035 | 33.343 | 37.263 | 1.00 | 0.00 | C |
| ATOM | 4947 | HE1 | LYS | A | 341 | 21.829 | 33.644 | 37.950 | 1.00 | 0.00 | H |
| ATOM | 4948 | HE2 | LYS | A | 341 | 21.365 | 33.569 | 36.248 | 1.00 | 0.00 | H |
| ATOM | 4949 | NZ | LYS | A | 341 | 19.798 | 34.095 | 37.570 | 1.00 | 0.00 | N |
| ATOM | 4950 | HZ1 | LYS | A | 341 | 19.010 | 33.695 | 37.070 | 1.00 | 0.00 | H |
| ATOM | 4951 | HZ2 | LYS | A | 341 | 19.595 | 34.052 | 38.561 | 1.00 | 0.00 | H |
| ATOM | 4952 | HZ3 | LYS | A | 341 | 19.900 | 35.071 | 37.308 | 1.00 | 0.00 | H |
| ATOM | 4953 | C | LYS | A | 341 | 23.176 | 27.370 | 37.173 | 1.00 | 0.00 | C |
| ATOM | 4954 | O | LYS | A | 341 | 24.169 | 27.003 | 37.800 | 1.00 | 0.00 | O |
| ATOM | 4955 | N | THR | A | 342 | 22.186 | 26.532 | 36.837 | 1.00 | 0.00 | N |
| ATOM | 4956 | H | THR | A | 342 | 21.443 | 26.918 | 36.264 | 1.00 | 0.00 | H |
| ATOM | 4957 | CA | THR | A | 342 | 22.075 | 25.115 | 37.252 | 1.00 | 0.00 | C |
| ATOM | 4958 | HA | THR | A | 342 | 21.912 | 25.075 | 38.331 | 1.00 | 0.00 | H |
| ATOM | 4959 | CB | THR | A | 342 | 20.855 | 24.489 | 36.564 | 1.00 | 0.00 | C |
| ATOM | 4960 | HB | THR | A | 342 | 20.991 | 24.502 | 35.481 | 1.00 | 0.00 | H |
| ATOM | 4961 | CG2 | THR | A | 342 | 20.602 | 23.050 | 37.010 | 1.00 | 0.00 | C |
| ATOM | 4962 | HG21 | THR | A | 342 | 19.684 | 22.685 | 36.554 | 1.00 | 0.00 | H |
| ATOM | 4963 | HG22 | THR | A | 342 | 21.422 | 22.409 | 36.689 | 1.00 | 0.00 | H |
| ATOM | 4964 | HG23 | THR | A | 342 | 20.509 | 23.001 | 38.094 | 1.00 | 0.00 | H |
| ATOM | 4965 | OG1 | THR | A | 342 | 19.708 | 25.244 | 36.908 | 1.00 | 0.00 | O |

| | | | | | | | | | | | |
|---|---|---|---|---|---|---|---|---|---|---|---|
| ATOM | 4966 | HG1 | THR | A | 342 | 19.652 | 26.002 | 36.287 | 1.00 | 0.00 | H |
| ATOM | 4967 | C | THR | A | 342 | 23.354 | 24.314 | 36.968 | 1.00 | 0.00 | C |
| ATOM | 4968 | O | THR | A | 342 | 23.791 | 23.529 | 37.812 | 1.00 | 0.00 | O |
| ATOM | 4969 | N | LYS | A | 343 | 23.997 | 24.552 | 35.822 | 1.00 | 0.00 | N |
| ATOM | 4970 | H | LYS | A | 343 | 23.534 | 25.138 | 35.133 | 1.00 | 0.00 | H |
| ATOM | 4971 | CA | LYS | A | 343 | 25.260 | 23.914 | 35.423 | 1.00 | 0.00 | C |
| ATOM | 4972 | HA | LYS | A | 343 | 25.195 | 22.842 | 35.618 | 1.00 | 0.00 | H |
| ATOM | 4973 | CB | LYS | A | 343 | 25.450 | 24.122 | 33.910 | 1.00 | 0.00 | C |
| ATOM | 4974 | HB1 | LYS | A | 343 | 25.315 | 25.178 | 33.668 | 1.00 | 0.00 | H |
| ATOM | 4975 | HB2 | LYS | A | 343 | 26.466 | 23.836 | 33.634 | 1.00 | 0.00 | H |
| ATOM | 4976 | CG | LYS | A | 343 | 24.476 | 23.266 | 33.082 | 1.00 | 0.00 | C |
| ATOM | 4977 | HG1 | LYS | A | 343 | 24.769 | 22.219 | 33.169 | 1.00 | 0.00 | H |
| ATOM | 4978 | HG2 | LYS | A | 343 | 23.461 | 23.371 | 33.462 | 1.00 | 0.00 | H |
| ATOM | 4979 | CD | LYS | A | 343 | 24.498 | 23.684 | 31.608 | 1.00 | 0.00 | C |
| ATOM | 4980 | HD1 | LYS | A | 343 | 24.102 | 24.697 | 31.508 | 1.00 | 0.00 | H |
| ATOM | 4981 | HD2 | LYS | A | 343 | 25.528 | 23.669 | 31.252 | 1.00 | 0.00 | H |
| ATOM | 4982 | CE | LYS | A | 343 | 23.649 | 22.716 | 30.778 | 1.00 | 0.00 | C |
| ATOM | 4983 | HE1 | LYS | A | 343 | 24.012 | 21.699 | 30.947 | 1.00 | 0.00 | H |
| ATOM | 4984 | HE2 | LYS | A | 343 | 22.609 | 22.771 | 31.110 | 1.00 | 0.00 | H |
| ATOM | 4985 | NZ | LYS | A | 343 | 23.730 | 23.018 | 29.332 | 1.00 | 0.00 | N |
| ATOM | 4986 | HZ1 | LYS | A | 343 | 23.302 | 22.269 | 28.796 | 1.00 | 0.00 | H |
| ATOM | 4987 | HZ2 | LYS | A | 343 | 23.223 | 23.872 | 29.102 | 1.00 | 0.00 | H |
| ATOM | 4988 | HZ3 | LYS | A | 343 | 24.694 | 23.105 | 29.033 | 1.00 | 0.00 | H |
| ATOM | 4989 | C | LYS | A | 343 | 26.485 | 24.421 | 36.208 | 1.00 | 0.00 | C |
| ATOM | 4990 | O | LYS | A | 343 | 27.367 | 23.624 | 36.530 | 1.00 | 0.00 | O |
| ATOM | 4991 | N | VAL | A | 344 | 26.546 | 25.708 | 36.564 | 1.00 | 0.00 | N |
| ATOM | 4992 | H | VAL | A | 344 | 25.790 | 26.320 | 36.268 | 1.00 | 0.00 | H |
| ATOM | 4993 | CA | VAL | A | 344 | 27.603 | 26.238 | 37.450 | 1.00 | 0.00 | C |
| ATOM | 4994 | HA | VAL | A | 344 | 28.542 | 25.785 | 37.136 | 1.00 | 0.00 | H |
| ATOM | 4995 | CB | VAL | A | 344 | 27.776 | 27.767 | 37.303 | 1.00 | 0.00 | C |
| ATOM | 4996 | HB | VAL | A | 344 | 26.798 | 28.242 | 37.377 | 1.00 | 0.00 | H |
| ATOM | 4997 | CG1 | VAL | A | 344 | 28.708 | 28.368 | 38.368 | 1.00 | 0.00 | C |
| ATOM | 4998 | HG11 | VAL | A | 344 | 28.924 | 29.410 | 38.136 | 1.00 | 0.00 | H |
| ATOM | 4999 | HG12 | VAL | A | 344 | 28.231 | 28.335 | 39.345 | 1.00 | 0.00 | H |
| ATOM | 5000 | HG13 | VAL | A | 344 | 29.641 | 27.806 | 38.403 | 1.00 | 0.00 | H |
| ATOM | 5001 | CG2 | VAL | A | 344 | 28.390 | 28.073 | 35.922 | 1.00 | 0.00 | C |
| ATOM | 5002 | HG21 | VAL | A | 344 | 28.545 | 29.146 | 35.810 | 1.00 | 0.00 | H |
| ATOM | 5003 | HG22 | VAL | A | 344 | 29.354 | 27.574 | 35.819 | 1.00 | 0.00 | H |
| ATOM | 5004 | HG23 | VAL | A | 344 | 27.729 | 27.730 | 35.127 | 1.00 | 0.00 | H |
| ATOM | 5005 | C | VAL | A | 344 | 27.378 | 25.807 | 38.908 | 1.00 | 0.00 | C |
| ATOM | 5006 | O | VAL | A | 344 | 28.340 | 25.477 | 39.597 | 1.00 | 0.00 | O |
| ATOM | 5007 | N | SER | A | 345 | 26.129 | 25.698 | 39.364 | 1.00 | 0.00 | N |
| ATOM | 5008 | H | SER | A | 345 | 25.365 | 26.104 | 38.828 | 1.00 | 0.00 | H |
| ATOM | 5009 | CA | SER | A | 345 | 25.790 | 25.083 | 40.655 | 1.00 | 0.00 | C |
| ATOM | 5010 | HA | SER | A | 345 | 26.273 | 25.640 | 41.458 | 1.00 | 0.00 | H |
| ATOM | 5011 | CB | SER | A | 345 | 24.269 | 25.162 | 40.888 | 1.00 | 0.00 | C |
| ATOM | 5012 | HB1 | SER | A | 345 | 23.752 | 24.568 | 40.135 | 1.00 | 0.00 | H |
| ATOM | 5013 | HB2 | SER | A | 345 | 24.037 | 24.755 | 41.873 | 1.00 | 0.00 | H |
| ATOM | 5014 | OG | SER | A | 345 | 23.814 | 26.508 | 40.816 | 1.00 | 0.00 | O |
| ATOM | 5015 | HG | SER | A | 345 | 22.838 | 26.540 | 40.896 | 1.00 | 0.00 | H |
| ATOM | 5016 | C | SER | A | 345 | 26.308 | 23.631 | 40.721 | 1.00 | 0.00 | C |
| ATOM | 5017 | O | SER | A | 345 | 26.959 | 23.251 | 41.696 | 1.00 | 0.00 | O |
| ATOM | 5018 | N | TYR | A | 346 | 26.132 | 22.848 | 39.646 | 1.00 | 0.00 | N |
| ATOM | 5019 | H | TYR | A | 346 | 25.590 | 23.214 | 38.873 | 1.00 | 0.00 | H |

| | | | | | | | | | | | |
|---|---|---|---|---|---|---|---|---|---|---|---|
| ATOM | 5020 | CA | TYR | A | 346 | 26.686 | 21.492 | 39.517 | 1.00 | 0.00 | C |
| ATOM | 5021 | HA | TYR | A | 346 | 26.350 | 20.898 | 40.367 | 1.00 | 0.00 | H |
| ATOM | 5022 | CB | TYR | A | 346 | 26.123 | 20.840 | 38.241 | 1.00 | 0.00 | C |
| ATOM | 5023 | HB1 | TYR | A | 346 | 25.037 | 20.808 | 38.314 | 1.00 | 0.00 | H |
| ATOM | 5024 | HB2 | TYR | A | 346 | 26.364 | 21.463 | 37.383 | 1.00 | 0.00 | H |
| ATOM | 5025 | CG | TYR | A | 346 | 26.624 | 19.434 | 37.955 | 1.00 | 0.00 | C |
| ATOM | 5026 | CD1 | TYR | A | 346 | 25.891 | 18.309 | 38.385 | 1.00 | 0.00 | C |
| ATOM | 5027 | HD1 | TYR | A | 346 | 24.974 | 18.442 | 38.943 | 1.00 | 0.00 | H |
| ATOM | 5028 | CE1 | TYR | A | 346 | 26.338 | 17.008 | 38.078 | 1.00 | 0.00 | C |
| ATOM | 5029 | HE1 | TYR | A | 346 | 25.764 | 16.149 | 38.397 | 1.00 | 0.00 | H |
| ATOM | 5030 | CZ | TYR | A | 346 | 27.543 | 16.827 | 37.361 | 1.00 | 0.00 | C |
| ATOM | 5031 | OH | TYR | A | 346 | 28.016 | 15.582 | 37.079 | 1.00 | 0.00 | O |
| ATOM | 5032 | HH | TYR | A | 346 | 27.367 | 14.871 | 37.213 | 1.00 | 0.00 | H |
| ATOM | 5033 | CE2 | TYR | A | 346 | 28.277 | 17.956 | 36.937 | 1.00 | 0.00 | C |
| ATOM | 5034 | HE2 | TYR | A | 346 | 29.184 | 17.816 | 36.374 | 1.00 | 0.00 | H |
| ATOM | 5035 | CD2 | TYR | A | 346 | 27.812 | 19.253 | 37.221 | 1.00 | 0.00 | C |
| ATOM | 5036 | HD2 | TYR | A | 346 | 28.367 | 20.109 | 36.866 | 1.00 | 0.00 | H |
| ATOM | 5037 | C | TYR | A | 346 | 28.227 | 21.465 | 39.552 | 1.00 | 0.00 | C |
| ATOM | 5038 | O | TYR | A | 346 | 28.801 | 20.678 | 40.307 | 1.00 | 0.00 | O |
| ATOM | 5039 | N | LEU | A | 347 | 28.925 | 22.329 | 38.797 | 1.00 | 0.00 | N |
| ATOM | 5040 | H | LEU | A | 347 | 28.419 | 22.956 | 38.180 | 1.00 | 0.00 | H |
| ATOM | 5041 | CA | LEU | A | 347 | 30.401 | 22.319 | 38.764 | 1.00 | 0.00 | C |
| ATOM | 5042 | HA | LEU | A | 347 | 30.697 | 21.281 | 38.599 | 1.00 | 0.00 | H |
| ATOM | 5043 | CB | LEU | A | 347 | 30.920 | 23.114 | 37.540 | 1.00 | 0.00 | C |
| ATOM | 5044 | HB1 | LEU | A | 347 | 31.824 | 22.628 | 37.182 | 1.00 | 0.00 | H |
| ATOM | 5045 | HB2 | LEU | A | 347 | 30.184 | 22.993 | 36.744 | 1.00 | 0.00 | H |
| ATOM | 5046 | CG | LEU | A | 347 | 31.247 | 24.618 | 37.687 | 1.00 | 0.00 | C |
| ATOM | 5047 | HG | LEU | A | 347 | 30.486 | 25.100 | 38.295 | 1.00 | 0.00 | H |
| ATOM | 5048 | CD1 | LEU | A | 347 | 32.632 | 24.881 | 38.308 | 1.00 | 0.00 | C |
| ATOM | 5049 | HD11 | LEU | A | 347 | 32.819 | 25.954 | 38.352 | 1.00 | 0.00 | H |
| ATOM | 5050 | HD12 | LEU | A | 347 | 32.691 | 24.494 | 39.322 | 1.00 | 0.00 | H |
| ATOM | 5051 | HD13 | LEU | A | 347 | 33.411 | 24.415 | 37.705 | 1.00 | 0.00 | H |
| ATOM | 5052 | CD2 | LEU | A | 347 | 31.240 | 25.262 | 36.292 | 1.00 | 0.00 | C |
| ATOM | 5053 | HD21 | LEU | A | 347 | 31.446 | 26.329 | 36.377 | 1.00 | 0.00 | H |
| ATOM | 5054 | HD22 | LEU | A | 347 | 31.997 | 24.800 | 35.658 | 1.00 | 0.00 | H |
| ATOM | 5055 | HD23 | LEU | A | 347 | 30.264 | 25.133 | 35.823 | 1.00 | 0.00 | H |
| ATOM | 5056 | C | LEU | A | 347 | 31.020 | 22.695 | 40.123 | 1.00 | 0.00 | C |
| ATOM | 5057 | O | LEU | A | 347 | 32.015 | 22.092 | 40.529 | 1.00 | 0.00 | O |
| ATOM | 5058 | N | LYS | A | 348 | 30.383 | 23.611 | 40.864 | 1.00 | 0.00 | N |
| ATOM | 5059 | H | LYS | A | 348 | 29.582 | 24.078 | 40.453 | 1.00 | 0.00 | H |
| ATOM | 5060 | CA | LYS | A | 348 | 30.735 | 23.966 | 42.251 | 1.00 | 0.00 | C |
| ATOM | 5061 | HA | LYS | A | 348 | 31.815 | 24.111 | 42.327 | 1.00 | 0.00 | H |
| ATOM | 5062 | CB | LYS | A | 348 | 30.035 | 25.282 | 42.634 | 1.00 | 0.00 | C |
| ATOM | 5063 | HB1 | LYS | A | 348 | 28.966 | 25.172 | 42.444 | 1.00 | 0.00 | H |
| ATOM | 5064 | HB2 | LYS | A | 348 | 30.172 | 25.451 | 43.704 | 1.00 | 0.00 | H |
| ATOM | 5065 | CG | LYS | A | 348 | 30.584 | 26.515 | 41.895 | 1.00 | 0.00 | C |
| ATOM | 5066 | HG1 | LYS | A | 348 | 31.657 | 26.589 | 42.077 | 1.00 | 0.00 | H |
| ATOM | 5067 | HG2 | LYS | A | 348 | 30.432 | 26.404 | 40.822 | 1.00 | 0.00 | H |
| ATOM | 5068 | CD | LYS | A | 348 | 29.920 | 27.820 | 42.365 | 1.00 | 0.00 | C |
| ATOM | 5069 | HD1 | LYS | A | 348 | 30.171 | 27.998 | 43.413 | 1.00 | 0.00 | H |
| ATOM | 5070 | HD2 | LYS | A | 348 | 30.321 | 28.642 | 41.769 | 1.00 | 0.00 | H |
| ATOM | 5071 | CE | LYS | A | 348 | 28.394 | 27.770 | 42.207 | 1.00 | 0.00 | C |
| ATOM | 5072 | HE1 | LYS | A | 348 | 28.154 | 27.475 | 41.182 | 1.00 | 0.00 | H |
| ATOM | 5073 | HE2 | LYS | A | 348 | 27.982 | 27.013 | 42.879 | 1.00 | 0.00 | H |

| | | | | | | | | | | | |
|---|---|---|---|---|---|---|---|---|---|---|---|
| ATOM | 5074 | NZ | LYS | A | 348 | 27.751 | 29.067 | 42.501 | 1.00 | 0.00 | N |
| ATOM | 5075 | HZ1 | LYS | A | 348 | 27.808 | 29.291 | 43.492 | 1.00 | 0.00 | H |
| ATOM | 5076 | HZ2 | LYS | A | 348 | 28.180 | 29.819 | 41.971 | 1.00 | 0.00 | H |
| ATOM | 5077 | HZ3 | LYS | A | 348 | 26.771 | 29.009 | 42.240 | 1.00 | 0.00 | H |
| ATOM | 5078 | C | LYS | A | 348 | 30.375 | 22.876 | 43.281 | 1.00 | 0.00 | C |
| ATOM | 5079 | O | LYS | A | 348 | 30.862 | 22.927 | 44.410 | 1.00 | 0.00 | O |
| ATOM | 5080 | N | GLN | A | 349 | 29.533 | 21.898 | 42.933 | 1.00 | 0.00 | N |
| ATOM | 5081 | H | GLN | A | 349 | 29.110 | 21.927 | 42.016 | 1.00 | 0.00 | H |
| ATOM | 5082 | CA | GLN | A | 349 | 29.294 | 20.722 | 43.773 | 1.00 | 0.00 | C |
| ATOM | 5083 | HA | GLN | A | 349 | 29.336 | 21.017 | 44.825 | 1.00 | 0.00 | H |
| ATOM | 5084 | CB | GLN | A | 349 | 27.894 | 20.139 | 43.489 | 1.00 | 0.00 | C |
| ATOM | 5085 | HB1 | GLN | A | 349 | 27.173 | 20.955 | 43.496 | 1.00 | 0.00 | H |
| ATOM | 5086 | HB2 | GLN | A | 349 | 27.861 | 19.684 | 42.499 | 1.00 | 0.00 | H |
| ATOM | 5087 | CG | GLN | A | 349 | 27.435 | 19.113 | 44.540 | 1.00 | 0.00 | C |
| ATOM | 5088 | HG1 | GLN | A | 349 | 27.711 | 19.471 | 45.529 | 1.00 | 0.00 | H |
| ATOM | 5089 | HG2 | GLN | A | 349 | 26.347 | 19.045 | 44.513 | 1.00 | 0.00 | H |
| ATOM | 5090 | CD | GLN | A | 349 | 27.998 | 17.709 | 44.313 | 1.00 | 0.00 | C |
| ATOM | 5091 | OE1 | GLN | A | 349 | 27.571 | 16.983 | 43.420 | 1.00 | 0.00 | O |
| ATOM | 5092 | NE2 | GLN | A | 349 | 28.956 | 17.267 | 45.097 | 1.00 | 0.00 | N |
| ATOM | 5093 | HE21 | GLN | A | 349 | 29.304 | 17.859 | 45.846 | 1.00 | 0.00 | H |
| ATOM | 5094 | HE22 | GLN | A | 349 | 29.439 | 16.409 | 44.885 | 1.00 | 0.00 | H |
| ATOM | 5095 | C | GLN | A | 349 | 30.414 | 19.704 | 43.539 | 1.00 | 0.00 | C |
| ATOM | 5096 | O | GLN | A | 349 | 31.080 | 19.298 | 44.486 | 1.00 | 0.00 | O |
| ATOM | 5097 | N | LYS | A | 350 | 30.661 | 19.325 | 42.280 | 1.00 | 0.00 | N |
| ATOM | 5098 | H | LYS | A | 350 | 30.082 | 19.706 | 41.538 | 1.00 | 0.00 | H |
| ATOM | 5099 | CA | LYS | A | 350 | 31.641 | 18.290 | 41.921 | 1.00 | 0.00 | C |
| ATOM | 5100 | HA | LYS | A | 350 | 31.488 | 17.432 | 42.577 | 1.00 | 0.00 | H |
| ATOM | 5101 | CB | LYS | A | 350 | 31.401 | 17.857 | 40.460 | 1.00 | 0.00 | C |
| ATOM | 5102 | HB1 | LYS | A | 350 | 31.546 | 18.723 | 39.812 | 1.00 | 0.00 | H |
| ATOM | 5103 | HB2 | LYS | A | 350 | 32.150 | 17.111 | 40.188 | 1.00 | 0.00 | H |
| ATOM | 5104 | CG | LYS | A | 350 | 30.013 | 17.267 | 40.161 | 1.00 | 0.00 | C |
| ATOM | 5105 | HG1 | LYS | A | 350 | 29.247 | 18.023 | 40.335 | 1.00 | 0.00 | H |
| ATOM | 5106 | HG2 | LYS | A | 350 | 29.986 | 17.000 | 39.104 | 1.00 | 0.00 | H |
| ATOM | 5107 | CD | LYS | A | 350 | 29.680 | 16.018 | 40.988 | 1.00 | 0.00 | C |
| ATOM | 5108 | HD1 | LYS | A | 350 | 30.448 | 15.260 | 40.829 | 1.00 | 0.00 | H |
| ATOM | 5109 | HD2 | LYS | A | 350 | 29.644 | 16.274 | 42.046 | 1.00 | 0.00 | H |
| ATOM | 5110 | CE | LYS | A | 350 | 28.321 | 15.472 | 40.541 | 1.00 | 0.00 | C |
| ATOM | 5111 | HE1 | LYS | A | 350 | 27.585 | 16.281 | 40.549 | 1.00 | 0.00 | H |
| ATOM | 5112 | HE2 | LYS | A | 350 | 28.411 | 15.113 | 39.513 | 1.00 | 0.00 | H |
| ATOM | 5113 | NZ | LYS | A | 350 | 27.845 | 14.370 | 41.405 | 1.00 | 0.00 | N |
| ATOM | 5114 | HZ1 | LYS | A | 350 | 27.089 | 13.867 | 40.944 | 1.00 | 0.00 | H |
| ATOM | 5115 | HZ2 | LYS | A | 350 | 28.577 | 13.701 | 41.615 | 1.00 | 0.00 | H |
| ATOM | 5116 | HZ3 | LYS | A | 350 | 27.487 | 14.707 | 42.291 | 1.00 | 0.00 | H |
| ATOM | 5117 | C | LYS | A | 350 | 33.115 | 18.695 | 42.133 | 1.00 | 0.00 | C |
| ATOM | 5118 | O | LYS | A | 350 | 33.982 | 17.821 | 42.089 | 1.00 | 0.00 | O |
| ATOM | 5119 | N | GLY | A | 351 | 33.412 | 19.983 | 42.348 | 1.00 | 0.00 | N |
| ATOM | 5120 | H | GLY | A | 351 | 32.646 | 20.641 | 42.360 | 1.00 | 0.00 | H |
| ATOM | 5121 | CA | GLY | A | 351 | 34.764 | 20.480 | 42.652 | 1.00 | 0.00 | C |
| ATOM | 5122 | HA1 | GLY | A | 351 | 34.677 | 21.377 | 43.264 | 1.00 | 0.00 | H |
| ATOM | 5123 | HA2 | GLY | A | 351 | 35.314 | 19.720 | 43.210 | 1.00 | 0.00 | H |
| ATOM | 5124 | C | GLY | A | 351 | 35.599 | 20.841 | 41.420 | 1.00 | 0.00 | C |
| ATOM | 5125 | O | GLY | A | 351 | 36.831 | 20.823 | 41.492 | 1.00 | 0.00 | O |
| ATOM | 5126 | N | LEU | A | 352 | 34.950 | 21.139 | 40.287 | 1.00 | 0.00 | N |
| ATOM | 5127 | H | LEU | A | 352 | 33.943 | 21.235 | 40.330 | 1.00 | 0.00 | H |

| | | | | | | | | | | | |
|---|---|---|---|---|---|---|---|---|---|---|---|
| ATOM | 5128 | CA | LEU | A | 352 | 35.621 | 21.373 | 39.002 | 1.00 | 0.00 | C |
| ATOM | 5129 | HA | LEU | A | 352 | 36.386 | 20.605 | 38.876 | 1.00 | 0.00 | H |
| ATOM | 5130 | CB | LEU | A | 352 | 34.620 | 21.248 | 37.837 | 1.00 | 0.00 | C |
| ATOM | 5131 | HB1 | LEU | A | 352 | 33.874 | 22.034 | 37.942 | 1.00 | 0.00 | H |
| ATOM | 5132 | HB2 | LEU | A | 352 | 35.160 | 21.430 | 36.907 | 1.00 | 0.00 | H |
| ATOM | 5133 | CG | LEU | A | 352 | 33.894 | 19.892 | 37.713 | 1.00 | 0.00 | C |
| ATOM | 5134 | HG | LEU | A | 352 | 33.174 | 19.806 | 38.527 | 1.00 | 0.00 | H |
| ATOM | 5135 | CD1 | LEU | A | 352 | 33.127 | 19.842 | 36.382 | 1.00 | 0.00 | C |
| ATOM | 5136 | HD11 | LEU | A | 352 | 32.566 | 18.912 | 36.314 | 1.00 | 0.00 | H |
| ATOM | 5137 | HD12 | LEU | A | 352 | 32.424 | 20.669 | 36.318 | 1.00 | 0.00 | H |
| ATOM | 5138 | HD13 | LEU | A | 352 | 33.823 | 19.898 | 35.543 | 1.00 | 0.00 | H |
| ATOM | 5139 | CD2 | LEU | A | 352 | 34.849 | 18.692 | 37.782 | 1.00 | 0.00 | C |
| ATOM | 5140 | HD21 | LEU | A | 352 | 34.304 | 17.775 | 37.559 | 1.00 | 0.00 | H |
| ATOM | 5141 | HD22 | LEU | A | 352 | 35.653 | 18.817 | 37.059 | 1.00 | 0.00 | H |
| ATOM | 5142 | HD23 | LEU | A | 352 | 35.271 | 18.601 | 38.783 | 1.00 | 0.00 | H |
| ATOM | 5143 | C | LEU | A | 352 | 36.360 | 22.718 | 38.953 | 1.00 | 0.00 | C |
| ATOM | 5144 | O | LEU | A | 352 | 35.994 | 23.681 | 39.637 | 1.00 | 0.00 | O |
| ATOM | 5145 | N | GLY | A | 353 | 37.382 | 22.798 | 38.095 | 1.00 | 0.00 | N |
| ATOM | 5146 | H | GLY | A | 353 | 37.604 | 21.965 | 37.556 | 1.00 | 0.00 | H |
| ATOM | 5147 | CA | GLY | A | 353 | 38.322 | 23.923 | 38.003 | 1.00 | 0.00 | C |
| ATOM | 5148 | HA1 | GLY | A | 353 | 38.652 | 24.174 | 39.008 | 1.00 | 0.00 | H |
| ATOM | 5149 | HA2 | GLY | A | 353 | 39.180 | 23.616 | 37.405 | 1.00 | 0.00 | H |
| ATOM | 5150 | C | GLY | A | 353 | 37.753 | 25.204 | 37.388 | 1.00 | 0.00 | C |
| ATOM | 5151 | O | GLY | A | 353 | 38.458 | 26.213 | 37.350 | 1.00 | 0.00 | O |
| ATOM | 5152 | N | GLY | A | 354 | 36.496 | 25.190 | 36.933 | 1.00 | 0.00 | N |
| ATOM | 5153 | H | GLY | A | 354 | 35.997 | 24.313 | 36.978 | 1.00 | 0.00 | H |
| ATOM | 5154 | CA | GLY | A | 354 | 35.777 | 26.350 | 36.398 | 1.00 | 0.00 | C |
| ATOM | 5155 | HA1 | GLY | A | 354 | 35.136 | 26.760 | 37.179 | 1.00 | 0.00 | H |
| ATOM | 5156 | HA2 | GLY | A | 354 | 36.485 | 27.121 | 36.105 | 1.00 | 0.00 | H |
| ATOM | 5157 | C | GLY | A | 354 | 34.889 | 26.022 | 35.200 | 1.00 | 0.00 | C |
| ATOM | 5158 | O | GLY | A | 354 | 34.668 | 24.853 | 34.877 | 1.00 | 0.00 | O |
| ATOM | 5159 | N | ALA | A | 355 | 34.397 | 27.073 | 34.541 | 1.00 | 0.00 | N |
| ATOM | 5160 | H | ALA | A | 355 | 34.688 | 27.998 | 34.838 | 1.00 | 0.00 | H |
| ATOM | 5161 | CA | ALA | A | 355 | 33.590 | 26.975 | 33.323 | 1.00 | 0.00 | C |
| ATOM | 5162 | HA | ALA | A | 355 | 33.247 | 25.945 | 33.200 | 1.00 | 0.00 | H |
| ATOM | 5163 | CB | ALA | A | 355 | 32.346 | 27.863 | 33.460 | 1.00 | 0.00 | C |
| ATOM | 5164 | HB1 | ALA | A | 355 | 31.712 | 27.742 | 32.579 | 1.00 | 0.00 | H |
| ATOM | 5165 | HB2 | ALA | A | 355 | 31.774 | 27.576 | 34.341 | 1.00 | 0.00 | H |
| ATOM | 5166 | HB3 | ALA | A | 355 | 32.639 | 28.909 | 33.540 | 1.00 | 0.00 | H |
| ATOM | 5167 | C | ALA | A | 355 | 34.394 | 27.350 | 32.067 | 1.00 | 0.00 | C |
| ATOM | 5168 | O | ALA | A | 355 | 35.252 | 28.238 | 32.083 | 1.00 | 0.00 | O |
| ATOM | 5169 | N | MET | A | 356 | 34.064 | 26.691 | 30.963 | 1.00 | 0.00 | N |
| ATOM | 5170 | H | MET | A | 356 | 33.320 | 26.000 | 31.040 | 1.00 | 0.00 | H |
| ATOM | 5171 | CA | MET | A | 356 | 34.473 | 27.012 | 29.600 | 1.00 | 0.00 | C |
| ATOM | 5172 | HA | MET | A | 356 | 35.139 | 27.864 | 29.630 | 1.00 | 0.00 | H |
| ATOM | 5173 | CB | MET | A | 356 | 35.240 | 25.808 | 29.017 | 1.00 | 0.00 | C |
| ATOM | 5174 | HB1 | MET | A | 356 | 36.168 | 25.699 | 29.578 | 1.00 | 0.00 | H |
| ATOM | 5175 | HB2 | MET | A | 356 | 34.663 | 24.894 | 29.157 | 1.00 | 0.00 | H |
| ATOM | 5176 | CG | MET | A | 356 | 35.619 | 25.944 | 27.540 | 1.00 | 0.00 | C |
| ATOM | 5177 | HG1 | MET | A | 356 | 35.913 | 26.974 | 27.334 | 1.00 | 0.00 | H |
| ATOM | 5178 | HG2 | MET | A | 356 | 36.499 | 25.328 | 27.355 | 1.00 | 0.00 | H |
| ATOM | 5179 | SD | MET | A | 356 | 34.322 | 25.442 | 26.376 | 1.00 | 0.00 | S |
| ATOM | 5180 | CE | MET | A | 356 | 34.605 | 23.653 | 26.349 | 1.00 | 0.00 | C |
| ATOM | 5181 | HE1 | MET | A | 356 | 34.598 | 23.250 | 27.361 | 1.00 | 0.00 | H |

| | | | | | | | | | | | |
|---|---|---|---|---|---|---|---|---|---|---|---|
| ATOM | 5182 | HE2 | MET | A | 356 | 33.816 | 23.175 | 25.769 | 1.00 | 0.00 | H |
| ATOM | 5183 | HE3 | MET | A | 356 | 35.558 | 23.442 | 25.866 | 1.00 | 0.00 | H |
| ATOM | 5184 | C | MET | A | 356 | 33.219 | 27.405 | 28.808 | 1.00 | 0.00 | C |
| ATOM | 5185 | O | MET | A | 356 | 32.129 | 26.947 | 29.139 | 1.00 | 0.00 | O |
| ATOM | 5186 | N | VAL | A | 357 | 33.340 | 28.258 | 27.793 | 1.00 | 0.00 | N |
| ATOM | 5187 | H | VAL | A | 357 | 34.255 | 28.644 | 27.593 | 1.00 | 0.00 | H |
| ATOM | 5188 | CA | VAL | A | 357 | 32.199 | 28.701 | 26.982 | 1.00 | 0.00 | C |
| ATOM | 5189 | HA | VAL | A | 357 | 31.461 | 27.902 | 27.023 | 1.00 | 0.00 | H |
| ATOM | 5190 | CB | VAL | A | 357 | 31.524 | 29.954 | 27.592 | 1.00 | 0.00 | C |
| ATOM | 5191 | HB | VAL | A | 357 | 31.225 | 29.695 | 28.608 | 1.00 | 0.00 | H |
| ATOM | 5192 | CG1 | VAL | A | 357 | 32.452 | 31.175 | 27.702 | 1.00 | 0.00 | C |
| ATOM | 5193 | HG11 | VAL | A | 357 | 31.934 | 31.983 | 28.220 | 1.00 | 0.00 | H |
| ATOM | 5194 | HG12 | VAL | A | 357 | 33.340 | 30.919 | 28.278 | 1.00 | 0.00 | H |
| ATOM | 5195 | HG13 | VAL | A | 357 | 32.747 | 31.527 | 26.713 | 1.00 | 0.00 | H |
| ATOM | 5196 | CG2 | VAL | A | 357 | 30.251 | 30.347 | 26.834 | 1.00 | 0.00 | C |
| ATOM | 5197 | HG21 | VAL | A | 357 | 29.729 | 31.133 | 27.376 | 1.00 | 0.00 | H |
| ATOM | 5198 | HG22 | VAL | A | 357 | 30.491 | 30.703 | 25.831 | 1.00 | 0.00 | H |
| ATOM | 5199 | HG23 | VAL | A | 357 | 29.584 | 29.491 | 26.765 | 1.00 | 0.00 | H |
| ATOM | 5200 | C | VAL | A | 357 | 32.565 | 28.875 | 25.506 | 1.00 | 0.00 | C |
| ATOM | 5201 | O | VAL | A | 357 | 33.550 | 29.539 | 25.171 | 1.00 | 0.00 | O |
| ATOM | 5202 | N | TRP | A | 358 | 31.739 | 28.273 | 24.644 | 1.00 | 0.00 | N |
| ATOM | 5203 | H | TRP | A | 358 | 30.958 | 27.754 | 25.038 | 1.00 | 0.00 | H |
| ATOM | 5204 | CA | TRP | A | 358 | 31.757 | 28.404 | 23.187 | 1.00 | 0.00 | C |
| ATOM | 5205 | HA | TRP | A | 358 | 32.602 | 29.015 | 22.880 | 1.00 | 0.00 | H |
| ATOM | 5206 | CB | TRP | A | 358 | 31.954 | 27.019 | 22.552 | 1.00 | 0.00 | C |
| ATOM | 5207 | HB1 | TRP | A | 358 | 32.839 | 26.561 | 22.995 | 1.00 | 0.00 | H |
| ATOM | 5208 | HB2 | TRP | A | 358 | 31.103 | 26.381 | 22.796 | 1.00 | 0.00 | H |
| ATOM | 5209 | CG | TRP | A | 358 | 32.142 | 27.035 | 21.063 | 1.00 | 0.00 | C |
| ATOM | 5210 | CD1 | TRP | A | 358 | 31.155 | 27.036 | 20.139 | 1.00 | 0.00 | C |
| ATOM | 5211 | HD1 | TRP | A | 358 | 30.096 | 27.019 | 20.359 | 1.00 | 0.00 | H |
| ATOM | 5212 | NE1 | TRP | A | 358 | 31.702 | 27.070 | 18.873 | 1.00 | 0.00 | N |
| ATOM | 5213 | HE1 | TRP | A | 358 | 31.151 | 27.051 | 18.021 | 1.00 | 0.00 | H |
| ATOM | 5214 | CE2 | TRP | A | 358 | 33.076 | 27.126 | 18.911 | 1.00 | 0.00 | C |
| ATOM | 5215 | CZ2 | TRP | A | 358 | 34.060 | 27.209 | 17.916 | 1.00 | 0.00 | C |
| ATOM | 5216 | HZ2 | TRP | A | 358 | 33.777 | 27.243 | 16.873 | 1.00 | 0.00 | H |
| ATOM | 5217 | CH2 | TRP | A | 358 | 35.412 | 27.238 | 18.294 | 1.00 | 0.00 | C |
| ATOM | 5218 | HH2 | TRP | A | 358 | 36.185 | 27.304 | 17.543 | 1.00 | 0.00 | H |
| ATOM | 5219 | CZ3 | TRP | A | 358 | 35.757 | 27.175 | 19.654 | 1.00 | 0.00 | C |
| ATOM | 5220 | HZ3 | TRP | A | 358 | 36.799 | 27.183 | 19.942 | 1.00 | 0.00 | H |
| ATOM | 5221 | CE3 | TRP | A | 358 | 34.756 | 27.102 | 20.642 | 1.00 | 0.00 | C |
| ATOM | 5222 | HE3 | TRP | A | 358 | 35.028 | 27.053 | 21.684 | 1.00 | 0.00 | H |
| ATOM | 5223 | CD2 | TRP | A | 358 | 33.389 | 27.082 | 20.301 | 1.00 | 0.00 | C |
| ATOM | 5224 | C | TRP | A | 358 | 30.432 | 29.052 | 22.733 | 1.00 | 0.00 | C |
| ATOM | 5225 | O | TRP | A | 358 | 29.374 | 28.436 | 22.846 | 1.00 | 0.00 | O |
| ATOM | 5226 | N | ALA | A | 359 | 30.413 | 30.295 | 22.249 | 1.00 | 0.00 | N |
| ATOM | 5227 | H | ALA | A | 359 | 29.479 | 30.658 | 22.095 | 1.00 | 0.00 | H |
| ATOM | 5228 | CA | ALA | A | 359 | 31.520 | 31.239 | 22.058 | 1.00 | 0.00 | C |
| ATOM | 5229 | HA | ALA | A | 359 | 32.311 | 31.046 | 22.784 | 1.00 | 0.00 | H |
| ATOM | 5230 | CB | ALA | A | 359 | 32.087 | 31.040 | 20.642 | 1.00 | 0.00 | C |
| ATOM | 5231 | HB1 | ALA | A | 359 | 32.939 | 31.701 | 20.478 | 1.00 | 0.00 | H |
| ATOM | 5232 | HB2 | ALA | A | 359 | 32.417 | 30.009 | 20.509 | 1.00 | 0.00 | H |
| ATOM | 5233 | HB3 | ALA | A | 359 | 31.322 | 31.269 | 19.897 | 1.00 | 0.00 | H |
| ATOM | 5234 | C | ALA | A | 359 | 31.067 | 32.692 | 22.297 | 1.00 | 0.00 | C |
| ATOM | 5235 | O | ALA | A | 359 | 29.869 | 32.978 | 22.392 | 1.00 | 0.00 | O |

| | | | | | | | | | | | |
|---|---|---|---|---|---|---|---|---|---|---|---|
| ATOM | 5236 | N | LEU | A | 360 | 32.037 | 33.612 | 22.385 | 1.00 | 0.00 | N |
| ATOM | 5237 | H | LEU | A | 360 | 32.997 | 33.306 | 22.262 | 1.00 | 0.00 | H |
| ATOM | 5238 | CA | LEU | A | 360 | 31.806 | 35.043 | 22.647 | 1.00 | 0.00 | C |
| ATOM | 5239 | HA | LEU | A | 360 | 31.251 | 35.151 | 23.579 | 1.00 | 0.00 | H |
| ATOM | 5240 | CB | LEU | A | 360 | 33.175 | 35.744 | 22.770 | 1.00 | 0.00 | C |
| ATOM | 5241 | HB1 | LEU | A | 360 | 33.783 | 35.459 | 21.911 | 1.00 | 0.00 | H |
| ATOM | 5242 | HB2 | LEU | A | 360 | 33.028 | 36.825 | 22.720 | 1.00 | 0.00 | H |
| ATOM | 5243 | CG | LEU | A | 360 | 33.939 | 35.413 | 24.068 | 1.00 | 0.00 | C |
| ATOM | 5244 | HG | LEU | A | 360 | 33.866 | 34.344 | 24.273 | 1.00 | 0.00 | H |
| ATOM | 5245 | CD1 | LEU | A | 360 | 35.422 | 35.766 | 23.910 | 1.00 | 0.00 | C |
| ATOM | 5246 | HD11 | LEU | A | 360 | 35.960 | 35.525 | 24.827 | 1.00 | 0.00 | H |
| ATOM | 5247 | HD12 | LEU | A | 360 | 35.853 | 35.188 | 23.095 | 1.00 | 0.00 | H |
| ATOM | 5248 | HD13 | LEU | A | 360 | 35.533 | 36.828 | 23.694 | 1.00 | 0.00 | H |
| ATOM | 5249 | CD2 | LEU | A | 360 | 33.370 | 36.189 | 25.266 | 1.00 | 0.00 | C |
| ATOM | 5250 | HD21 | LEU | A | 360 | 33.895 | 35.905 | 26.178 | 1.00 | 0.00 | H |
| ATOM | 5251 | HD22 | LEU | A | 360 | 33.486 | 37.261 | 25.106 | 1.00 | 0.00 | H |
| ATOM | 5252 | HD23 | LEU | A | 360 | 32.313 | 35.969 | 25.389 | 1.00 | 0.00 | H |
| ATOM | 5253 | C | LEU | A | 360 | 30.945 | 35.723 | 21.571 | 1.00 | 0.00 | C |
| ATOM | 5254 | O | LEU | A | 360 | 30.237 | 36.675 | 21.876 | 1.00 | 0.00 | O |
| ATOM | 5255 | N | ASP | A | 361 | 30.979 | 35.221 | 20.336 | 1.00 | 0.00 | N |
| ATOM | 5256 | H | ASP | A | 361 | 31.608 | 34.448 | 20.162 | 1.00 | 0.00 | H |
| ATOM | 5257 | CA | ASP | A | 361 | 30.189 | 35.712 | 19.201 | 1.00 | 0.00 | C |
| ATOM | 5258 | HA | ASP | A | 361 | 30.051 | 36.787 | 19.320 | 1.00 | 0.00 | H |
| ATOM | 5259 | CB | ASP | A | 361 | 30.982 | 35.480 | 17.904 | 1.00 | 0.00 | C |
| ATOM | 5260 | HB1 | ASP | A | 361 | 30.393 | 35.818 | 17.051 | 1.00 | 0.00 | H |
| ATOM | 5261 | HB2 | ASP | A | 361 | 31.879 | 36.100 | 17.929 | 1.00 | 0.00 | H |
| ATOM | 5262 | CG | ASP | A | 361 | 31.389 | 34.018 | 17.698 | 1.00 | 0.00 | C |
| ATOM | 5263 | OD1 | ASP | A | 361 | 30.720 | 33.300 | 16.924 | 1.00 | 0.00 | O |
| ATOM | 5264 | OD2 | ASP | A | 361 | 32.421 | 33.621 | 18.291 | 1.00 | 0.00 | O |
| ATOM | 5265 | C | ASP | A | 361 | 28.773 | 35.114 | 19.089 | 1.00 | 0.00 | C |
| ATOM | 5266 | O | ASP | A | 361 | 28.077 | 35.450 | 18.131 | 1.00 | 0.00 | O |
| ATOM | 5267 | N | LEU | A | 362 | 28.352 | 34.237 | 20.013 | 1.00 | 0.00 | N |
| ATOM | 5268 | H | LEU | A | 362 | 28.986 | 34.009 | 20.768 | 1.00 | 0.00 | H |
| ATOM | 5269 | CA | LEU | A | 362 | 27.040 | 33.562 | 19.984 | 1.00 | 0.00 | C |
| ATOM | 5270 | HA | LEU | A | 362 | 26.498 | 33.858 | 19.088 | 1.00 | 0.00 | H |
| ATOM | 5271 | CB | LEU | A | 362 | 27.261 | 32.036 | 19.920 | 1.00 | 0.00 | C |
| ATOM | 5272 | HB1 | LEU | A | 362 | 27.782 | 31.733 | 20.823 | 1.00 | 0.00 | H |
| ATOM | 5273 | HB2 | LEU | A | 362 | 26.285 | 31.547 | 19.920 | 1.00 | 0.00 | H |
| ATOM | 5274 | CG | LEU | A | 362 | 28.069 | 31.523 | 18.708 | 1.00 | 0.00 | C |
| ATOM | 5275 | HG | LEU | A | 362 | 29.045 | 32.008 | 18.700 | 1.00 | 0.00 | H |
| ATOM | 5276 | CD1 | LEU | A | 362 | 28.304 | 30.011 | 18.851 | 1.00 | 0.00 | C |
| ATOM | 5277 | HD11 | LEU | A | 362 | 28.919 | 29.654 | 18.025 | 1.00 | 0.00 | H |
| ATOM | 5278 | HD12 | LEU | A | 362 | 28.821 | 29.800 | 19.787 | 1.00 | 0.00 | H |
| ATOM | 5279 | HD13 | LEU | A | 362 | 27.352 | 29.480 | 18.840 | 1.00 | 0.00 | H |
| ATOM | 5280 | CD2 | LEU | A | 362 | 27.373 | 31.799 | 17.367 | 1.00 | 0.00 | C |
| ATOM | 5281 | HD21 | LEU | A | 362 | 27.979 | 31.405 | 16.550 | 1.00 | 0.00 | H |
| ATOM | 5282 | HD22 | LEU | A | 362 | 26.395 | 31.317 | 17.348 | 1.00 | 0.00 | H |
| ATOM | 5283 | HD23 | LEU | A | 362 | 27.252 | 32.871 | 17.216 | 1.00 | 0.00 | H |
| ATOM | 5284 | C | LEU | A | 362 | 26.108 | 33.956 | 21.152 | 1.00 | 0.00 | C |
| ATOM | 5285 | O | LEU | A | 362 | 24.890 | 33.876 | 20.998 | 1.00 | 0.00 | O |
| ATOM | 5286 | N | ASP | A | 363 | 26.656 | 34.442 | 22.275 | 1.00 | 0.00 | N |
| ATOM | 5287 | H | ASP | A | 363 | 27.655 | 34.559 | 22.283 | 1.00 | 0.00 | H |
| ATOM | 5288 | CA | ASP | A | 363 | 25.888 | 35.167 | 23.307 | 1.00 | 0.00 | C |
| ATOM | 5289 | HA | ASP | A | 363 | 25.011 | 34.572 | 23.572 | 1.00 | 0.00 | H |

| | | | | | | | | | | | |
|---|---|---|---|---|---|---|---|---|---|---|---|
| ATOM | 5290 | CB | ASP | A | 363 | 26.768 | 35.341 | 24.569 | 1.00 | 0.00 | C |
| ATOM | 5291 | HB1 | ASP | A | 363 | 27.275 | 34.397 | 24.766 | 1.00 | 0.00 | H |
| ATOM | 5292 | HB2 | ASP | A | 363 | 27.539 | 36.084 | 24.357 | 1.00 | 0.00 | H |
| ATOM | 5293 | CG | ASP | A | 363 | 26.029 | 35.742 | 25.861 | 1.00 | 0.00 | C |
| ATOM | 5294 | OD1 | ASP | A | 363 | 24.780 | 35.777 | 25.884 | 1.00 | 0.00 | O |
| ATOM | 5295 | OD2 | ASP | A | 363 | 26.713 | 36.007 | 26.880 | 1.00 | 0.00 | O |
| ATOM | 5296 | C | ASP | A | 363 | 25.404 | 36.524 | 22.745 | 1.00 | 0.00 | C |
| ATOM | 5297 | O | ASP | A | 363 | 25.924 | 36.998 | 21.734 | 1.00 | 0.00 | O |
| ATOM | 5298 | N | ASP | A | 364 | 24.444 | 37.190 | 23.396 | 1.00 | 0.00 | N |
| ATOM | 5299 | H | ASP | A | 364 | 24.108 | 36.790 | 24.269 | 1.00 | 0.00 | H |
| ATOM | 5300 | CA | ASP | A | 364 | 23.999 | 38.538 | 22.991 | 1.00 | 0.00 | C |
| ATOM | 5301 | HA | ASP | A | 364 | 24.013 | 38.572 | 21.900 | 1.00 | 0.00 | H |
| ATOM | 5302 | CB | ASP | A | 364 | 22.538 | 38.798 | 23.406 | 1.00 | 0.00 | C |
| ATOM | 5303 | HB1 | ASP | A | 364 | 21.939 | 37.943 | 23.094 | 1.00 | 0.00 | H |
| ATOM | 5304 | HB2 | ASP | A | 364 | 22.476 | 38.873 | 24.491 | 1.00 | 0.00 | H |
| ATOM | 5305 | CG | ASP | A | 364 | 21.926 | 40.063 | 22.776 | 1.00 | 0.00 | C |
| ATOM | 5306 | OD1 | ASP | A | 364 | 22.557 | 40.696 | 21.901 | 1.00 | 0.00 | O |
| ATOM | 5307 | OD2 | ASP | A | 364 | 20.789 | 40.429 | 23.165 | 1.00 | 0.00 | O |
| ATOM | 5308 | C | ASP | A | 364 | 24.997 | 39.609 | 23.475 | 1.00 | 0.00 | C |
| ATOM | 5309 | O | ASP | A | 364 | 24.703 | 40.445 | 24.333 | 1.00 | 0.00 | O |
| ATOM | 5310 | N | PHE | A | 365 | 26.217 | 39.555 | 22.931 | 1.00 | 0.00 | N |
| ATOM | 5311 | H | PHE | A | 365 | 26.381 | 38.797 | 22.275 | 1.00 | 0.00 | H |
| ATOM | 5312 | CA | PHE | A | 365 | 27.360 | 40.404 | 23.275 | 1.00 | 0.00 | C |
| ATOM | 5313 | HA | PHE | A | 365 | 27.608 | 40.238 | 24.323 | 1.00 | 0.00 | H |
| ATOM | 5314 | CB | PHE | A | 365 | 28.573 | 39.970 | 22.433 | 1.00 | 0.00 | C |
| ATOM | 5315 | HB1 | PHE | A | 365 | 29.453 | 40.498 | 22.802 | 1.00 | 0.00 | H |
| ATOM | 5316 | HB2 | PHE | A | 365 | 28.750 | 38.904 | 22.589 | 1.00 | 0.00 | H |
| ATOM | 5317 | CG | PHE | A | 365 | 28.450 | 40.236 | 20.942 | 1.00 | 0.00 | C |
| ATOM | 5318 | CD1 | PHE | A | 365 | 28.831 | 41.487 | 20.415 | 1.00 | 0.00 | C |
| ATOM | 5319 | HD1 | PHE | A | 365 | 29.220 | 42.254 | 21.069 | 1.00 | 0.00 | H |
| ATOM | 5320 | CE1 | PHE | A | 365 | 28.681 | 41.752 | 19.043 | 1.00 | 0.00 | C |
| ATOM | 5321 | HE1 | PHE | A | 365 | 28.957 | 42.717 | 18.645 | 1.00 | 0.00 | H |
| ATOM | 5322 | CZ | PHE | A | 365 | 28.147 | 40.768 | 18.195 | 1.00 | 0.00 | C |
| ATOM | 5323 | HZ | PHE | A | 365 | 28.004 | 40.983 | 17.148 | 1.00 | 0.00 | H |
| ATOM | 5324 | CE2 | PHE | A | 365 | 27.786 | 39.511 | 18.710 | 1.00 | 0.00 | C |
| ATOM | 5325 | HE2 | PHE | A | 365 | 27.379 | 38.751 | 18.058 | 1.00 | 0.00 | H |
| ATOM | 5326 | CD2 | PHE | A | 365 | 27.943 | 39.244 | 20.081 | 1.00 | 0.00 | C |
| ATOM | 5327 | HD2 | PHE | A | 365 | 27.672 | 38.274 | 20.471 | 1.00 | 0.00 | H |
| ATOM | 5328 | C | PHE | A | 365 | 27.081 | 41.905 | 23.104 | 1.00 | 0.00 | C |
| ATOM | 5329 | O | PHE | A | 365 | 27.729 | 42.716 | 23.758 | 1.00 | 0.00 | O |
| ATOM | 5330 | N | ALA | A | 366 | 26.117 | 42.278 | 22.253 | 1.00 | 0.00 | N |
| ATOM | 5331 | H | ALA | A | 366 | 25.644 | 41.540 | 21.750 | 1.00 | 0.00 | H |
| ATOM | 5332 | CA | ALA | A | 366 | 25.709 | 43.661 | 21.993 | 1.00 | 0.00 | C |
| ATOM | 5333 | HA | ALA | A | 366 | 26.476 | 44.337 | 22.367 | 1.00 | 0.00 | H |
| ATOM | 5334 | CB | ALA | A | 366 | 25.627 | 43.832 | 20.473 | 1.00 | 0.00 | C |
| ATOM | 5335 | HB1 | ALA | A | 366 | 25.429 | 44.875 | 20.234 | 1.00 | 0.00 | H |
| ATOM | 5336 | HB2 | ALA | A | 366 | 26.577 | 43.545 | 20.020 | 1.00 | 0.00 | H |
| ATOM | 5337 | HB3 | ALA | A | 366 | 24.834 | 43.209 | 20.061 | 1.00 | 0.00 | H |
| ATOM | 5338 | C | ALA | A | 366 | 24.391 | 44.070 | 22.685 | 1.00 | 0.00 | C |
| ATOM | 5339 | O | ALA | A | 366 | 23.988 | 45.237 | 22.603 | 1.00 | 0.00 | O |
| ATOM | 5340 | N | GLY | A | 367 | 23.704 | 43.128 | 23.343 | 1.00 | 0.00 | N |
| ATOM | 5341 | H | GLY | A | 367 | 24.076 | 42.184 | 23.355 | 1.00 | 0.00 | H |
| ATOM | 5342 | CA | GLY | A | 367 | 22.415 | 43.338 | 24.011 | 1.00 | 0.00 | C |
| ATOM | 5343 | HA1 | GLY | A | 367 | 22.151 | 42.420 | 24.526 | 1.00 | 0.00 | H |

| | | | | | | | | | | |
|---|---|---|---|---|---|---|---|---|---|---|
| ATOM | 5344 | HA2 | GLY | A | 367 | 22.503 | 44.147 | 24.735 | 1.00 | 0.00 | H |
| ATOM | 5345 | C | GLY | A | 367 | 21.269 | 43.676 | 23.056 | 1.00 | 0.00 | C |
| ATOM | 5346 | O | GLY | A | 367 | 20.296 | 44.308 | 23.468 | 1.00 | 0.00 | O |
| ATOM | 5347 | N | PHE | A | 368 | 21.393 | 43.329 | 21.772 | 1.00 | 0.00 | N |
| ATOM | 5348 | H | PHE | A | 368 | 22.131 | 42.674 | 21.532 | 1.00 | 0.00 | H |
| ATOM | 5349 | CA | PHE | A | 368 | 20.514 | 43.824 | 20.714 | 1.00 | 0.00 | C |
| ATOM | 5350 | HA | PHE | A | 368 | 20.236 | 44.851 | 20.960 | 1.00 | 0.00 | H |
| ATOM | 5351 | CB | PHE | A | 368 | 21.281 | 43.872 | 19.382 | 1.00 | 0.00 | C |
| ATOM | 5352 | HB1 | PHE | A | 368 | 20.804 | 44.630 | 18.760 | 1.00 | 0.00 | H |
| ATOM | 5353 | HB2 | PHE | A | 368 | 22.300 | 44.220 | 19.559 | 1.00 | 0.00 | H |
| ATOM | 5354 | CG | PHE | A | 368 | 21.328 | 42.580 | 18.579 | 1.00 | 0.00 | C |
| ATOM | 5355 | CD1 | PHE | A | 368 | 22.342 | 41.628 | 18.798 | 1.00 | 0.00 | C |
| ATOM | 5356 | HD1 | PHE | A | 368 | 23.072 | 41.780 | 19.578 | 1.00 | 0.00 | H |
| ATOM | 5357 | CE1 | PHE | A | 368 | 22.422 | 40.481 | 17.989 | 1.00 | 0.00 | C |
| ATOM | 5358 | HE1 | PHE | A | 368 | 23.213 | 39.762 | 18.153 | 1.00 | 0.00 | H |
| ATOM | 5359 | CZ | PHE | A | 368 | 21.485 | 40.275 | 16.962 | 1.00 | 0.00 | C |
| ATOM | 5360 | HZ | PHE | A | 368 | 21.559 | 39.408 | 16.323 | 1.00 | 0.00 | H |
| ATOM | 5361 | CE2 | PHE | A | 368 | 20.454 | 41.206 | 16.757 | 1.00 | 0.00 | C |
| ATOM | 5362 | HE2 | PHE | A | 368 | 19.731 | 41.050 | 15.968 | 1.00 | 0.00 | H |
| ATOM | 5363 | CD2 | PHE | A | 368 | 20.380 | 42.357 | 17.560 | 1.00 | 0.00 | C |
| ATOM | 5364 | HD2 | PHE | A | 368 | 19.607 | 43.085 | 17.372 | 1.00 | 0.00 | H |
| ATOM | 5365 | C | PHE | A | 368 | 19.214 | 43.023 | 20.587 | 1.00 | 0.00 | C |
| ATOM | 5366 | O | PHE | A | 368 | 18.188 | 43.586 | 20.207 | 1.00 | 0.00 | O |
| ATOM | 5367 | N | SER | A | 369 | 19.221 | 41.730 | 20.909 | 1.00 | 0.00 | N |
| ATOM | 5368 | H | SER | A | 369 | 20.071 | 41.337 | 21.314 | 1.00 | 0.00 | H |
| ATOM | 5369 | CA | SER | A | 369 | 18.097 | 40.835 | 20.597 | 1.00 | 0.00 | C |
| ATOM | 5370 | HA | SER | A | 369 | 17.427 | 41.300 | 19.875 | 1.00 | 0.00 | H |
| ATOM | 5371 | CB | SER | A | 369 | 18.644 | 39.566 | 19.946 | 1.00 | 0.00 | C |
| ATOM | 5372 | HB1 | SER | A | 369 | 19.278 | 39.842 | 19.102 | 1.00 | 0.00 | H |
| ATOM | 5373 | HB2 | SER | A | 369 | 19.243 | 39.016 | 20.673 | 1.00 | 0.00 | H |
| ATOM | 5374 | OG | SER | A | 369 | 17.587 | 38.747 | 19.480 | 1.00 | 0.00 | O |
| ATOM | 5375 | HG | SER | A | 369 | 18.003 | 37.948 | 19.095 | 1.00 | 0.00 | H |
| ATOM | 5376 | C | SER | A | 369 | 17.253 | 40.459 | 21.818 | 1.00 | 0.00 | C |
| ATOM | 5377 | O | SER | A | 369 | 16.020 | 40.440 | 21.732 | 1.00 | 0.00 | O |
| ATOM | 5378 | N | CYS | A | 370 | 17.904 | 40.154 | 22.944 | 1.00 | 0.00 | N |
| ATOM | 5379 | H | CYS | A | 370 | 18.923 | 40.210 | 22.945 | 1.00 | 0.00 | H |
| ATOM | 5380 | CA | CYS | A | 370 | 17.273 | 39.494 | 24.093 | 1.00 | 0.00 | C |
| ATOM | 5381 | HA | CYS | A | 370 | 16.395 | 38.949 | 23.746 | 1.00 | 0.00 | H |
| ATOM | 5382 | CB | CYS | A | 370 | 18.263 | 38.478 | 24.680 | 1.00 | 0.00 | C |
| ATOM | 5383 | HB1 | CYS | A | 370 | 19.030 | 39.006 | 25.243 | 1.00 | 0.00 | H |
| ATOM | 5384 | HB2 | CYS | A | 370 | 17.727 | 37.832 | 25.378 | 1.00 | 0.00 | H |
| ATOM | 5385 | SG | CYS | A | 370 | 19.095 | 37.439 | 23.452 | 1.00 | 0.00 | S |
| ATOM | 5386 | C | CYS | A | 370 | 16.795 | 40.459 | 25.181 | 1.00 | 0.00 | C |
| ATOM | 5387 | O | CYS | A | 370 | 16.063 | 40.043 | 26.078 | 1.00 | 0.00 | O |
| ATOM | 5388 | N | ASN | A | 371 | 17.214 | 41.727 | 25.122 | 1.00 | 0.00 | N |
| ATOM | 5389 | H | ASN | A | 371 | 17.784 | 41.991 | 24.331 | 1.00 | 0.00 | H |
| ATOM | 5390 | CA | ASN | A | 371 | 16.825 | 42.798 | 26.047 | 1.00 | 0.00 | C |
| ATOM | 5391 | HA | ASN | A | 371 | 17.356 | 43.687 | 25.708 | 1.00 | 0.00 | H |
| ATOM | 5392 | CB | ASN | A | 371 | 15.311 | 43.081 | 25.895 | 1.00 | 0.00 | C |
| ATOM | 5393 | HB1 | ASN | A | 371 | 14.968 | 42.732 | 24.920 | 1.00 | 0.00 | H |
| ATOM | 5394 | HB2 | ASN | A | 371 | 14.747 | 42.540 | 26.655 | 1.00 | 0.00 | H |
| ATOM | 5395 | CG | ASN | A | 371 | 14.994 | 44.570 | 25.975 | 1.00 | 0.00 | C |
| ATOM | 5396 | OD1 | ASN | A | 371 | 14.612 | 45.195 | 24.988 | 1.00 | 0.00 | O |
| ATOM | 5397 | ND2 | ASN | A | 371 | 15.178 | 45.186 | 27.126 | 1.00 | 0.00 | N |

| | | | | | | | | | | | |
|---|---|---|---|---|---|---|---|---|---|---|---|
| ATOM | 5398 | HD21 | ASN | A | 371 | 15.062 | 46.188 | 27.162 | 1.00 | 0.00 | H |
| ATOM | 5399 | HD22 | ASN | A | 371 | 15.493 | 44.665 | 27.932 | 1.00 | 0.00 | H |
| ATOM | 5400 | C | ASN | A | 371 | 17.306 | 42.543 | 27.502 | 1.00 | 0.00 | C |
| ATOM | 5401 | O | ASN | A | 371 | 16.670 | 42.970 | 28.469 | 1.00 | 0.00 | O |
| ATOM | 5402 | N | GLN | A | 372 | 18.411 | 41.805 | 27.667 | 1.00 | 0.00 | N |
| ATOM | 5403 | H | GLN | A | 372 | 18.902 | 41.516 | 26.825 | 1.00 | 0.00 | H |
| ATOM | 5404 | CA | GLN | A | 372 | 19.032 | 41.421 | 28.945 | 1.00 | 0.00 | C |
| ATOM | 5405 | HA | GLN | A | 372 | 18.418 | 41.772 | 29.776 | 1.00 | 0.00 | H |
| ATOM | 5406 | CB | GLN | A | 372 | 19.136 | 39.877 | 29.016 | 1.00 | 0.00 | C |
| ATOM | 5407 | HB1 | GLN | A | 372 | 19.651 | 39.519 | 28.123 | 1.00 | 0.00 | H |
| ATOM | 5408 | HB2 | GLN | A | 372 | 19.730 | 39.599 | 29.887 | 1.00 | 0.00 | H |
| ATOM | 5409 | CG | GLN | A | 372 | 17.799 | 39.130 | 29.146 | 1.00 | 0.00 | C |
| ATOM | 5410 | HG1 | GLN | A | 372 | 17.310 | 39.428 | 30.075 | 1.00 | 0.00 | H |
| ATOM | 5411 | HG2 | GLN | A | 372 | 17.146 | 39.398 | 28.318 | 1.00 | 0.00 | H |
| ATOM | 5412 | CD | GLN | A | 372 | 17.989 | 37.608 | 29.141 | 1.00 | 0.00 | C |
| ATOM | 5413 | OE1 | GLN | A | 372 | 17.928 | 36.951 | 28.108 | 1.00 | 0.00 | O |
| ATOM | 5414 | NE2 | GLN | A | 372 | 18.240 | 36.971 | 30.267 | 1.00 | 0.00 | N |
| ATOM | 5415 | HE21 | GLN | A | 372 | 18.339 | 37.461 | 31.151 | 1.00 | 0.00 | H |
| ATOM | 5416 | HE22 | GLN | A | 372 | 18.295 | 35.963 | 30.237 | 1.00 | 0.00 | H |
| ATOM | 5417 | C | GLN | A | 372 | 20.426 | 42.072 | 29.128 | 1.00 | 0.00 | C |
| ATOM | 5418 | O | GLN | A | 372 | 21.225 | 41.596 | 29.937 | 1.00 | 0.00 | O |
| ATOM | 5419 | N | GLY | A | 373 | 20.762 | 43.126 | 28.371 | 1.00 | 0.00 | N |
| ATOM | 5420 | H | GLY | A | 373 | 20.067 | 43.517 | 27.741 | 1.00 | 0.00 | H |
| ATOM | 5421 | CA | GLY | A | 373 | 22.091 | 43.758 | 28.363 | 1.00 | 0.00 | C |
| ATOM | 5422 | HA1 | GLY | A | 373 | 22.009 | 44.760 | 27.941 | 1.00 | 0.00 | H |
| ATOM | 5423 | HA2 | GLY | A | 373 | 22.471 | 43.831 | 29.383 | 1.00 | 0.00 | H |
| ATOM | 5424 | C | GLY | A | 373 | 23.117 | 42.979 | 27.531 | 1.00 | 0.00 | C |
| ATOM | 5425 | O | GLY | A | 373 | 22.788 | 41.956 | 26.927 | 1.00 | 0.00 | O |
| ATOM | 5426 | N | ARG | A | 374 | 24.368 | 43.457 | 27.507 | 1.00 | 0.00 | N |
| ATOM | 5427 | H | ARG | A | 374 | 24.572 | 44.289 | 28.042 | 1.00 | 0.00 | H |
| ATOM | 5428 | CA | ARG | A | 374 | 25.490 | 42.769 | 26.842 | 1.00 | 0.00 | C |
| ATOM | 5429 | HA | ARG | A | 374 | 25.185 | 42.510 | 25.828 | 1.00 | 0.00 | H |
| ATOM | 5430 | CB | ARG | A | 374 | 26.731 | 43.678 | 26.755 | 1.00 | 0.00 | C |
| ATOM | 5431 | HB1 | ARG | A | 374 | 26.973 | 44.074 | 27.743 | 1.00 | 0.00 | H |
| ATOM | 5432 | HB2 | ARG | A | 374 | 27.571 | 43.062 | 26.428 | 1.00 | 0.00 | H |
| ATOM | 5433 | CG | ARG | A | 374 | 26.570 | 44.840 | 25.761 | 1.00 | 0.00 | C |
| ATOM | 5434 | HG1 | ARG | A | 374 | 26.111 | 44.463 | 24.849 | 1.00 | 0.00 | H |
| ATOM | 5435 | HG2 | ARG | A | 374 | 25.916 | 45.602 | 26.185 | 1.00 | 0.00 | H |
| ATOM | 5436 | CD | ARG | A | 374 | 27.935 | 45.457 | 25.410 | 1.00 | 0.00 | C |
| ATOM | 5437 | HD1 | ARG | A | 374 | 28.317 | 45.989 | 26.284 | 1.00 | 0.00 | H |
| ATOM | 5438 | HD2 | ARG | A | 374 | 28.638 | 44.661 | 25.175 | 1.00 | 0.00 | H |
| ATOM | 5439 | NE | ARG | A | 374 | 27.830 | 46.398 | 24.279 | 1.00 | 0.00 | N |
| ATOM | 5440 | HE | ARG | A | 374 | 27.362 | 47.270 | 24.495 | 1.00 | 0.00 | H |
| ATOM | 5441 | CZ | ARG | A | 374 | 28.308 | 46.241 | 23.045 | 1.00 | 0.00 | C |
| ATOM | 5442 | NH1 | ARG | A | 374 | 28.072 | 47.158 | 22.136 | 1.00 | 0.00 | N |
| ATOM | 5443 | HH11 | ARG | A | 374 | 27.554 | 47.995 | 22.374 | 1.00 | 0.00 | H |
| ATOM | 5444 | HH12 | ARG | A | 374 | 28.363 | 47.000 | 21.178 | 1.00 | 0.00 | H |
| ATOM | 5445 | NH2 | ARG | A | 374 | 29.007 | 45.191 | 22.670 | 1.00 | 0.00 | N |
| ATOM | 5446 | HH21 | ARG | A | 374 | 29.232 | 44.457 | 23.338 | 1.00 | 0.00 | H |
| ATOM | 5447 | HH22 | ARG | A | 374 | 29.436 | 45.139 | 21.752 | 1.00 | 0.00 | H |
| ATOM | 5448 | C | ARG | A | 374 | 25.863 | 41.469 | 27.570 | 1.00 | 0.00 | C |
| ATOM | 5449 | O | ARG | A | 374 | 25.855 | 41.427 | 28.801 | 1.00 | 0.00 | O |
| ATOM | 5450 | N | TYR | A | 375 | 26.222 | 40.422 | 26.826 | 1.00 | 0.00 | N |
| ATOM | 5451 | H | TYR | A | 375 | 26.081 | 40.505 | 25.826 | 1.00 | 0.00 | H |

| | | | | | | | | | | | |
|---|---|---|---|---|---|---|---|---|---|---|---|
| ATOM | 5452 | CA | TYR | A | 375 | 26.717 | 39.128 | 27.336 | 1.00 | 0.00 | C |
| ATOM | 5453 | HA | TYR | A | 375 | 26.697 | 38.426 | 26.503 | 1.00 | 0.00 | H |
| ATOM | 5454 | CB | TYR | A | 375 | 28.186 | 39.269 | 27.791 | 1.00 | 0.00 | C |
| ATOM | 5455 | HB1 | TYR | A | 375 | 28.258 | 40.074 | 28.521 | 1.00 | 0.00 | H |
| ATOM | 5456 | HB2 | TYR | A | 375 | 28.482 | 38.353 | 28.301 | 1.00 | 0.00 | H |
| ATOM | 5457 | CG | TYR | A | 375 | 29.218 | 39.482 | 26.706 | 1.00 | 0.00 | C |
| ATOM | 5458 | CD1 | TYR | A | 375 | 29.537 | 38.427 | 25.828 | 1.00 | 0.00 | C |
| ATOM | 5459 | HD1 | TYR | A | 375 | 28.982 | 37.500 | 25.873 | 1.00 | 0.00 | H |
| ATOM | 5460 | CE1 | TYR | A | 375 | 30.591 | 38.570 | 24.906 | 1.00 | 0.00 | C |
| ATOM | 5461 | HE1 | TYR | A | 375 | 30.829 | 37.767 | 24.223 | 1.00 | 0.00 | H |
| ATOM | 5462 | CZ | TYR | A | 375 | 31.327 | 39.774 | 24.858 | 1.00 | 0.00 | C |
| ATOM | 5463 | OH | TYR | A | 375 | 32.361 | 39.909 | 23.984 | 1.00 | 0.00 | O |
| ATOM | 5464 | HH | TYR | A | 375 | 32.631 | 40.835 | 23.876 | 1.00 | 0.00 | H |
| ATOM | 5465 | CE2 | TYR | A | 375 | 31.001 | 40.831 | 25.734 | 1.00 | 0.00 | C |
| ATOM | 5466 | HE2 | TYR | A | 375 | 31.572 | 41.748 | 25.718 | 1.00 | 0.00 | H |
| ATOM | 5467 | CD2 | TYR | A | 375 | 29.946 | 40.685 | 26.653 | 1.00 | 0.00 | C |
| ATOM | 5468 | HD2 | TYR | A | 375 | 29.720 | 41.489 | 27.338 | 1.00 | 0.00 | H |
| ATOM | 5469 | C | TYR | A | 375 | 25.841 | 38.496 | 28.457 | 1.00 | 0.00 | C |
| ATOM | 5470 | O | TYR | A | 375 | 26.364 | 38.199 | 29.540 | 1.00 | 0.00 | O |
| ATOM | 5471 | N | PRO | A | 376 | 24.516 | 38.322 | 28.270 | 1.00 | 0.00 | N |
| ATOM | 5472 | CD | PRO | A | 376 | 23.763 | 38.556 | 27.047 | 1.00 | 0.00 | C |
| ATOM | 5473 | HD1 | PRO | A | 376 | 24.274 | 38.164 | 26.170 | 1.00 | 0.00 | H |
| ATOM | 5474 | HD2 | PRO | A | 376 | 23.594 | 39.624 | 26.923 | 1.00 | 0.00 | H |
| ATOM | 5475 | CG | PRO | A | 376 | 22.423 | 37.854 | 27.254 | 1.00 | 0.00 | C |
| ATOM | 5476 | HG1 | PRO | A | 376 | 22.513 | 36.798 | 26.998 | 1.00 | 0.00 | H |
| ATOM | 5477 | HG2 | PRO | A | 376 | 21.623 | 38.325 | 26.682 | 1.00 | 0.00 | H |
| ATOM | 5478 | CB | PRO | A | 376 | 22.208 | 38.007 | 28.758 | 1.00 | 0.00 | C |
| ATOM | 5479 | HB1 | PRO | A | 376 | 21.529 | 37.250 | 29.153 | 1.00 | 0.00 | H |
| ATOM | 5480 | HB2 | PRO | A | 376 | 21.827 | 39.007 | 28.962 | 1.00 | 0.00 | H |
| ATOM | 5481 | CA | PRO | A | 376 | 23.617 | 37.901 | 29.345 | 1.00 | 0.00 | C |
| ATOM | 5482 | HA | PRO | A | 376 | 23.703 | 38.600 | 30.177 | 1.00 | 0.00 | H |
| ATOM | 5483 | C | PRO | A | 376 | 23.913 | 36.497 | 29.887 | 1.00 | 0.00 | C |
| ATOM | 5484 | O | PRO | A | 376 | 23.683 | 36.258 | 31.077 | 1.00 | 0.00 | O |
| ATOM | 5485 | N | LEU | A | 377 | 24.456 | 35.589 | 29.064 | 1.00 | 0.00 | N |
| ATOM | 5486 | H | LEU | A | 377 | 24.610 | 35.824 | 28.082 | 1.00 | 0.00 | H |
| ATOM | 5487 | CA | LEU | A | 377 | 24.893 | 34.267 | 29.514 | 1.00 | 0.00 | C |
| ATOM | 5488 | HA | LEU | A | 377 | 24.224 | 33.908 | 30.295 | 1.00 | 0.00 | H |
| ATOM | 5489 | CB | LEU | A | 377 | 24.842 | 33.285 | 28.330 | 1.00 | 0.00 | C |
| ATOM | 5490 | HB1 | LEU | A | 377 | 25.510 | 33.638 | 27.546 | 1.00 | 0.00 | H |
| ATOM | 5491 | HB2 | LEU | A | 377 | 25.227 | 32.332 | 28.676 | 1.00 | 0.00 | H |
| ATOM | 5492 | CG | LEU | A | 377 | 23.449 | 33.034 | 27.716 | 1.00 | 0.00 | C |
| ATOM | 5493 | HG | LEU | A | 377 | 23.057 | 33.965 | 27.305 | 1.00 | 0.00 | H |
| ATOM | 5494 | CD1 | LEU | A | 377 | 23.587 | 32.017 | 26.573 | 1.00 | 0.00 | C |
| ATOM | 5495 | HD11 | LEU | A | 377 | 22.605 | 31.742 | 26.196 | 1.00 | 0.00 | H |
| ATOM | 5496 | HD12 | LEU | A | 377 | 24.163 | 32.457 | 25.759 | 1.00 | 0.00 | H |
| ATOM | 5497 | HD13 | LEU | A | 377 | 24.081 | 31.113 | 26.929 | 1.00 | 0.00 | H |
| ATOM | 5498 | CD2 | LEU | A | 377 | 22.447 | 32.497 | 28.749 | 1.00 | 0.00 | C |
| ATOM | 5499 | HD21 | LEU | A | 377 | 21.492 | 32.313 | 28.262 | 1.00 | 0.00 | H |
| ATOM | 5500 | HD22 | LEU | A | 377 | 22.815 | 31.568 | 29.186 | 1.00 | 0.00 | H |
| ATOM | 5501 | HD23 | LEU | A | 377 | 22.287 | 33.231 | 29.536 | 1.00 | 0.00 | H |
| ATOM | 5502 | C | LEU | A | 377 | 26.295 | 34.310 | 30.138 | 1.00 | 0.00 | C |
| ATOM | 5503 | O | LEU | A | 377 | 26.515 | 33.724 | 31.198 | 1.00 | 0.00 | O |
| ATOM | 5504 | N | ILE | A | 378 | 27.256 | 35.011 | 29.532 | 1.00 | 0.00 | N |
| ATOM | 5505 | H | ILE | A | 378 | 27.034 | 35.447 | 28.631 | 1.00 | 0.00 | H |

| | | | | | | | | | | | |
|---|---|---|---|---|---|---|---|---|---|---|---|
| ATOM | 5506 | CA | ILE | A | 378 | 28.653 | 34.993 | 30.013 | 1.00 | 0.00 | C |
| ATOM | 5507 | HA | ILE | A | 378 | 28.871 | 33.984 | 30.364 | 1.00 | 0.00 | H |
| ATOM | 5508 | CB | ILE | A | 378 | 29.637 | 35.260 | 28.845 | 1.00 | 0.00 | C |
| ATOM | 5509 | HB | ILE | A | 378 | 29.419 | 36.233 | 28.405 | 1.00 | 0.00 | H |
| ATOM | 5510 | CG2 | ILE | A | 378 | 31.095 | 35.246 | 29.346 | 1.00 | 0.00 | C |
| ATOM | 5511 | HG21 | ILE | A | 378 | 31.780 | 35.499 | 28.538 | 1.00 | 0.00 | H |
| ATOM | 5512 | HG22 | ILE | A | 378 | 31.234 | 35.984 | 30.133 | 1.00 | 0.00 | H |
| ATOM | 5513 | HG23 | ILE | A | 378 | 31.350 | 34.261 | 29.739 | 1.00 | 0.00 | H |
| ATOM | 5514 | CG1 | ILE | A | 378 | 29.456 | 34.162 | 27.764 | 1.00 | 0.00 | C |
| ATOM | 5515 | HG11 | ILE | A | 378 | 29.690 | 33.196 | 28.209 | 1.00 | 0.00 | H |
| ATOM | 5516 | HG12 | ILE | A | 378 | 28.418 | 34.126 | 27.438 | 1.00 | 0.00 | H |
| ATOM | 5517 | CD | ILE | A | 378 | 30.307 | 34.337 | 26.501 | 1.00 | 0.00 | C |
| ATOM | 5518 | HD1 | ILE | A | 378 | 30.024 | 33.582 | 25.767 | 1.00 | 0.00 | H |
| ATOM | 5519 | HD2 | ILE | A | 378 | 30.131 | 35.324 | 26.074 | 1.00 | 0.00 | H |
| ATOM | 5520 | HD3 | ILE | A | 378 | 31.364 | 34.209 | 26.731 | 1.00 | 0.00 | H |
| ATOM | 5521 | C | ILE | A | 378 | 28.853 | 35.872 | 31.265 | 1.00 | 0.00 | C |
| ATOM | 5522 | O | ILE | A | 378 | 29.609 | 35.481 | 32.155 | 1.00 | 0.00 | O |
| ATOM | 5523 | N | GLN | A | 379 | 28.108 | 36.972 | 31.437 | 1.00 | 0.00 | N |
| ATOM | 5524 | H | GLN | A | 379 | 27.484 | 37.270 | 30.692 | 1.00 | 0.00 | H |
| ATOM | 5525 | CA | GLN | A | 379 | 28.060 | 37.678 | 32.729 | 1.00 | 0.00 | C |
| ATOM | 5526 | HA | GLN | A | 379 | 29.080 | 37.913 | 33.037 | 1.00 | 0.00 | H |
| ATOM | 5527 | CB | GLN | A | 379 | 27.262 | 38.995 | 32.646 | 1.00 | 0.00 | C |
| ATOM | 5528 | HB1 | GLN | A | 379 | 26.313 | 38.833 | 32.133 | 1.00 | 0.00 | H |
| ATOM | 5529 | HB2 | GLN | A | 379 | 27.033 | 39.307 | 33.666 | 1.00 | 0.00 | H |
| ATOM | 5530 | CG | GLN | A | 379 | 28.033 | 40.144 | 31.974 | 1.00 | 0.00 | C |
| ATOM | 5531 | HG1 | GLN | A | 379 | 29.061 | 40.152 | 32.334 | 1.00 | 0.00 | H |
| ATOM | 5532 | HG2 | GLN | A | 379 | 28.051 | 39.988 | 30.896 | 1.00 | 0.00 | H |
| ATOM | 5533 | CD | GLN | A | 379 | 27.419 | 41.510 | 32.298 | 1.00 | 0.00 | C |
| ATOM | 5534 | OE1 | GLN | A | 379 | 27.448 | 41.975 | 33.431 | 1.00 | 0.00 | O |
| ATOM | 5535 | NE2 | GLN | A | 379 | 26.838 | 42.208 | 31.350 | 1.00 | 0.00 | N |
| ATOM | 5536 | HE21 | GLN | A | 379 | 26.716 | 41.825 | 30.421 | 1.00 | 0.00 | H |
| ATOM | 5537 | HE22 | GLN | A | 379 | 26.408 | 43.080 | 31.605 | 1.00 | 0.00 | H |
| ATOM | 5538 | C | GLN | A | 379 | 27.475 | 36.790 | 33.841 | 1.00 | 0.00 | C |
| ATOM | 5539 | O | GLN | A | 379 | 27.936 | 36.876 | 34.982 | 1.00 | 0.00 | O |
| ATOM | 5540 | N | THR | A | 380 | 26.516 | 35.909 | 33.519 | 1.00 | 0.00 | N |
| ATOM | 5541 | H | THR | A | 380 | 26.189 | 35.882 | 32.562 | 1.00 | 0.00 | H |
| ATOM | 5542 | CA | THR | A | 380 | 25.953 | 34.940 | 34.472 | 1.00 | 0.00 | C |
| ATOM | 5543 | HA | THR | A | 380 | 25.661 | 35.478 | 35.373 | 1.00 | 0.00 | H |
| ATOM | 5544 | CB | THR | A | 380 | 24.697 | 34.281 | 33.886 | 1.00 | 0.00 | C |
| ATOM | 5545 | HB | THR | A | 380 | 24.880 | 33.928 | 32.874 | 1.00 | 0.00 | H |
| ATOM | 5546 | CG2 | THR | A | 380 | 24.175 | 33.105 | 34.706 | 1.00 | 0.00 | C |
| ATOM | 5547 | HG21 | THR | A | 380 | 23.219 | 32.774 | 34.303 | 1.00 | 0.00 | H |
| ATOM | 5548 | HG22 | THR | A | 380 | 24.869 | 32.273 | 34.629 | 1.00 | 0.00 | H |
| ATOM | 5549 | HG23 | THR | A | 380 | 24.063 | 33.382 | 35.753 | 1.00 | 0.00 | H |
| ATOM | 5550 | OG1 | THR | A | 380 | 23.692 | 35.265 | 33.856 | 1.00 | 0.00 | O |
| ATOM | 5551 | HG1 | THR | A | 380 | 23.619 | 35.586 | 32.943 | 1.00 | 0.00 | H |
| ATOM | 5552 | C | THR | A | 380 | 27.006 | 33.934 | 34.924 | 1.00 | 0.00 | C |
| ATOM | 5553 | O | THR | A | 380 | 27.129 | 33.712 | 36.128 | 1.00 | 0.00 | O |
| ATOM | 5554 | N | LEU | A | 381 | 27.826 | 33.395 | 34.009 | 1.00 | 0.00 | N |
| ATOM | 5555 | H | LEU | A | 381 | 27.675 | 33.631 | 33.034 | 1.00 | 0.00 | H |
| ATOM | 5556 | CA | LEU | A | 381 | 28.983 | 32.561 | 34.374 | 1.00 | 0.00 | C |
| ATOM | 5557 | HA | LEU | A | 381 | 28.632 | 31.692 | 34.932 | 1.00 | 0.00 | H |
| ATOM | 5558 | CB | LEU | A | 381 | 29.754 | 32.088 | 33.121 | 1.00 | 0.00 | C |
| ATOM | 5559 | HB1 | LEU | A | 381 | 30.121 | 32.962 | 32.583 | 1.00 | 0.00 | H |

```
ATOM   5560  HB2 LEU A 381      30.627  31.527  33.460  1.00  0.00           H
ATOM   5561  CG  LEU A 381      28.972  31.201  32.132  1.00  0.00           C
ATOM   5562  HG  LEU A 381      28.123  31.766  31.756  1.00  0.00           H
ATOM   5563  CD1 LEU A 381      29.868  30.842  30.938  1.00  0.00           C
ATOM   5564 HD11 LEU A 381      29.304  30.249  30.218  1.00  0.00           H
ATOM   5565 HD12 LEU A 381      30.213  31.752  30.446  1.00  0.00           H
ATOM   5566 HD13 LEU A 381      30.729  30.264  31.273  1.00  0.00           H
ATOM   5567  CD2 LEU A 381      28.458  29.911  32.786  1.00  0.00           C
ATOM   5568 HD21 LEU A 381      27.957  29.292  32.042  1.00  0.00           H
ATOM   5569 HD22 LEU A 381      29.289  29.350  33.217  1.00  0.00           H
ATOM   5570 HD23 LEU A 381      27.737  30.148  33.563  1.00  0.00           H
ATOM   5571  C   LEU A 381      29.926  33.309  35.327  1.00  0.00           C
ATOM   5572  O   LEU A 381      30.263  32.777  36.385  1.00  0.00           O
ATOM   5573  N   ARG A 382      30.306  34.551  34.994  1.00  0.00           N
ATOM   5574  H   ARG A 382      30.004  34.924  34.099  1.00  0.00           H
ATOM   5575  CA  ARG A 382      31.183  35.368  35.843  1.00  0.00           C
ATOM   5576  HA  ARG A 382      32.110  34.810  35.986  1.00  0.00           H
ATOM   5577  CB  ARG A 382      31.536  36.682  35.119  1.00  0.00           C
ATOM   5578  HB1 ARG A 382      31.909  36.450  34.120  1.00  0.00           H
ATOM   5579  HB2 ARG A 382      30.638  37.285  35.013  1.00  0.00           H
ATOM   5580  CG  ARG A 382      32.621  37.476  35.871  1.00  0.00           C
ATOM   5581  HG1 ARG A 382      32.315  37.643  36.900  1.00  0.00           H
ATOM   5582  HG2 ARG A 382      33.533  36.879  35.886  1.00  0.00           H
ATOM   5583  CD  ARG A 382      32.953  38.834  35.239  1.00  0.00           C
ATOM   5584  HD1 ARG A 382      33.798  39.270  35.775  1.00  0.00           H
ATOM   5585  HD2 ARG A 382      33.253  38.678  34.207  1.00  0.00           H
ATOM   5586  NE  ARG A 382      31.819  39.773  35.264  1.00  0.00           N
ATOM   5587  HE  ARG A 382      30.957  39.468  35.709  1.00  0.00           H
ATOM   5588  CZ  ARG A 382      31.779  40.954  34.662  1.00  0.00           C
ATOM   5589  NH1 ARG A 382      32.820  41.475  34.049  1.00  0.00           N
ATOM   5590 HH11 ARG A 382      33.734  41.035  34.075  1.00  0.00           H
ATOM   5591 HH12 ARG A 382      32.748  42.392  33.624  1.00  0.00           H
ATOM   5592  NH2 ARG A 382      30.663  41.641  34.664  1.00  0.00           N
ATOM   5593 HH21 ARG A 382      29.872  41.317  35.210  1.00  0.00           H
ATOM   5594 HH22 ARG A 382      30.617  42.527  34.175  1.00  0.00           H
ATOM   5595  C   ARG A 382      30.576  35.610  37.239  1.00  0.00           C
ATOM   5596  O   ARG A 382      31.298  35.514  38.232  1.00  0.00           O
ATOM   5597  N   GLN A 383      29.268  35.876  37.349  1.00  0.00           N
ATOM   5598  H   GLN A 383      28.717  35.959  36.500  1.00  0.00           H
ATOM   5599  CA  GLN A 383      28.613  36.116  38.640  1.00  0.00           C
ATOM   5600  HA  GLN A 383      29.235  36.807  39.213  1.00  0.00           H
ATOM   5601  CB  GLN A 383      27.238  36.777  38.440  1.00  0.00           C
ATOM   5602  HB1 GLN A 383      27.379  37.715  37.906  1.00  0.00           H
ATOM   5603  HB2 GLN A 383      26.608  36.129  37.829  1.00  0.00           H
ATOM   5604  CG  GLN A 383      26.505  37.068  39.769  1.00  0.00           C
ATOM   5605  HG1 GLN A 383      25.568  37.580  39.546  1.00  0.00           H
ATOM   5606  HG2 GLN A 383      26.252  36.129  40.263  1.00  0.00           H
ATOM   5607  CD  GLN A 383      27.309  37.946  40.729  1.00  0.00           C
ATOM   5608  OE1 GLN A 383      27.628  39.085  40.436  1.00  0.00           O
ATOM   5609  NE2 GLN A 383      27.708  37.478  41.896  1.00  0.00           N
ATOM   5610 HE21 GLN A 383      27.575  36.505  42.149  1.00  0.00           H
ATOM   5611 HE22 GLN A 383      28.368  38.047  42.405  1.00  0.00           H
ATOM   5612  C   GLN A 383      28.488  34.841  39.477  1.00  0.00           C
ATOM   5613  O   GLN A 383      28.866  34.859  40.645  1.00  0.00           O
```

| | | | | | | | | | | | |
|---|---|---|---|---|---|---|---|---|---|---|---|
| ATOM | 5614 | N | GLU | A | 384 | 27.988 | 33.732 | 38.928 | 1.00 | 0.00 | N |
| ATOM | 5615 | H | GLU | A | 384 | 27.682 | 33.745 | 37.956 | 1.00 | 0.00 | H |
| ATOM | 5616 | CA | GLU | A | 384 | 27.776 | 32.500 | 39.705 | 1.00 | 0.00 | C |
| ATOM | 5617 | HA | GLU | A | 384 | 27.226 | 32.753 | 40.613 | 1.00 | 0.00 | H |
| ATOM | 5618 | CB | GLU | A | 384 | 26.918 | 31.522 | 38.891 | 1.00 | 0.00 | C |
| ATOM | 5619 | HB1 | GLU | A | 384 | 27.400 | 31.367 | 37.925 | 1.00 | 0.00 | H |
| ATOM | 5620 | HB2 | GLU | A | 384 | 26.891 | 30.563 | 39.403 | 1.00 | 0.00 | H |
| ATOM | 5621 | CG | GLU | A | 384 | 25.474 | 32.002 | 38.648 | 1.00 | 0.00 | C |
| ATOM | 5622 | HG1 | GLU | A | 384 | 25.480 | 33.019 | 38.254 | 1.00 | 0.00 | H |
| ATOM | 5623 | HG2 | GLU | A | 384 | 25.046 | 31.362 | 37.880 | 1.00 | 0.00 | H |
| ATOM | 5624 | CD | GLU | A | 384 | 24.557 | 31.932 | 39.876 | 1.00 | 0.00 | C |
| ATOM | 5625 | OE1 | GLU | A | 384 | 24.421 | 30.840 | 40.476 | 1.00 | 0.00 | O |
| ATOM | 5626 | OE2 | GLU | A | 384 | 23.890 | 32.943 | 40.201 | 1.00 | 0.00 | O |
| ATOM | 5627 | C | GLU | A | 384 | 29.106 | 31.851 | 40.149 | 1.00 | 0.00 | C |
| ATOM | 5628 | O | GLU | A | 384 | 29.158 | 31.194 | 41.195 | 1.00 | 0.00 | O |
| ATOM | 5629 | N | LEU | A | 385 | 30.202 | 32.093 | 39.414 | 1.00 | 0.00 | N |
| ATOM | 5630 | H | LEU | A | 385 | 30.083 | 32.569 | 38.525 | 1.00 | 0.00 | H |
| ATOM | 5631 | CA | LEU | A | 385 | 31.576 | 31.775 | 39.833 | 1.00 | 0.00 | C |
| ATOM | 5632 | HA | LEU | A | 385 | 31.558 | 30.863 | 40.427 | 1.00 | 0.00 | H |
| ATOM | 5633 | CB | LEU | A | 385 | 32.446 | 31.540 | 38.582 | 1.00 | 0.00 | C |
| ATOM | 5634 | HB1 | LEU | A | 385 | 32.412 | 32.439 | 37.965 | 1.00 | 0.00 | H |
| ATOM | 5635 | HB2 | LEU | A | 385 | 33.478 | 31.395 | 38.902 | 1.00 | 0.00 | H |
| ATOM | 5636 | CG | LEU | A | 385 | 32.051 | 30.321 | 37.724 | 1.00 | 0.00 | C |
| ATOM | 5637 | HG | LEU | A | 385 | 31.002 | 30.396 | 37.439 | 1.00 | 0.00 | H |
| ATOM | 5638 | CD1 | LEU | A | 385 | 32.894 | 30.310 | 36.443 | 1.00 | 0.00 | C |
| ATOM | 5639 | HD11 | LEU | A | 385 | 32.620 | 29.450 | 35.836 | 1.00 | 0.00 | H |
| ATOM | 5640 | HD12 | LEU | A | 385 | 32.710 | 31.218 | 35.868 | 1.00 | 0.00 | H |
| ATOM | 5641 | HD13 | LEU | A | 385 | 33.953 | 30.249 | 36.692 | 1.00 | 0.00 | H |
| ATOM | 5642 | CD2 | LEU | A | 385 | 32.266 | 28.999 | 38.477 | 1.00 | 0.00 | C |
| ATOM | 5643 | HD21 | LEU | A | 385 | 32.024 | 28.164 | 37.820 | 1.00 | 0.00 | H |
| ATOM | 5644 | HD22 | LEU | A | 385 | 33.303 | 28.910 | 38.799 | 1.00 | 0.00 | H |
| ATOM | 5645 | HD23 | LEU | A | 385 | 31.616 | 28.947 | 39.349 | 1.00 | 0.00 | H |
| ATOM | 5646 | C | LEU | A | 385 | 32.214 | 32.855 | 40.732 | 1.00 | 0.00 | C |
| ATOM | 5647 | O | LEU | A | 385 | 33.312 | 32.644 | 41.244 | 1.00 | 0.00 | O |
| ATOM | 5648 | N | SER | A | 386 | 31.565 | 34.006 | 40.920 | 1.00 | 0.00 | N |
| ATOM | 5649 | H | SER | A | 386 | 30.643 | 34.087 | 40.505 | 1.00 | 0.00 | H |
| ATOM | 5650 | CA | SER | A | 386 | 31.998 | 35.150 | 41.754 | 1.00 | 0.00 | C |
| ATOM | 5651 | HA | SER | A | 386 | 31.205 | 35.898 | 41.716 | 1.00 | 0.00 | H |
| ATOM | 5652 | CB | SER | A | 386 | 32.126 | 34.749 | 43.235 | 1.00 | 0.00 | C |
| ATOM | 5653 | HB1 | SER | A | 386 | 32.980 | 34.088 | 43.374 | 1.00 | 0.00 | H |
| ATOM | 5654 | HB2 | SER | A | 386 | 32.289 | 35.648 | 43.830 | 1.00 | 0.00 | H |
| ATOM | 5655 | OG | SER | A | 386 | 30.945 | 34.104 | 43.694 | 1.00 | 0.00 | O |
| ATOM | 5656 | HG | SER | A | 386 | 31.028 | 33.145 | 43.512 | 1.00 | 0.00 | H |
| ATOM | 5657 | C | SER | A | 386 | 33.261 | 35.867 | 41.218 | 1.00 | 0.00 | C |
| ATOM | 5658 | O | SER | A | 386 | 33.914 | 36.647 | 41.915 | 1.00 | 0.00 | O |
| ATOM | 5659 | N | LEU | A | 387 | 33.615 | 35.647 | 39.947 | 1.00 | 0.00 | N |
| ATOM | 5660 | H | LEU | A | 387 | 32.972 | 35.095 | 39.388 | 1.00 | 0.00 | H |
| ATOM | 5661 | CA | LEU | A | 387 | 34.820 | 36.151 | 39.268 | 1.00 | 0.00 | C |
| ATOM | 5662 | HA | LEU | A | 387 | 35.649 | 36.150 | 39.974 | 1.00 | 0.00 | H |
| ATOM | 5663 | CB | LEU | A | 387 | 35.156 | 35.181 | 38.112 | 1.00 | 0.00 | C |
| ATOM | 5664 | HB1 | LEU | A | 387 | 34.299 | 35.134 | 37.439 | 1.00 | 0.00 | H |
| ATOM | 5665 | HB2 | LEU | A | 387 | 35.999 | 35.576 | 37.543 | 1.00 | 0.00 | H |
| ATOM | 5666 | CG | LEU | A | 387 | 35.523 | 33.751 | 38.561 | 1.00 | 0.00 | C |
| ATOM | 5667 | HG | LEU | A | 387 | 34.727 | 33.359 | 39.188 | 1.00 | 0.00 | H |

```
ATOM   5668  CD1 LEU A 387      35.638  32.835  37.335  1.00  0.00           C
ATOM   5669 HD11 LEU A 387      35.873  31.818  37.649  1.00  0.00           H
ATOM   5670 HD12 LEU A 387      34.693  32.826  36.794  1.00  0.00           H
ATOM   5671 HD13 LEU A 387      36.420  33.195  36.672  1.00  0.00           H
ATOM   5672  CD2 LEU A 387      36.829  33.720  39.367  1.00  0.00           C
ATOM   5673 HD21 LEU A 387      37.094  32.695  39.613  1.00  0.00           H
ATOM   5674 HD22 LEU A 387      37.635  34.164  38.786  1.00  0.00           H
ATOM   5675 HD23 LEU A 387      36.712  34.267  40.302  1.00  0.00           H
ATOM   5676  C   LEU A 387      34.687  37.616  38.800  1.00  0.00           C
ATOM   5677  O   LEU A 387      35.344  38.033  37.847  1.00  0.00           O
ATOM   5678  N   VAL A 388      33.820  38.392  39.448  1.00  0.00           N
ATOM   5679  H   VAL A 388      33.438  38.015  40.307  1.00  0.00           H
ATOM   5680  CA  VAL A 388      33.474  39.774  39.077  1.00  0.00           C
ATOM   5681  HA  VAL A 388      33.490  39.840  37.992  1.00  0.00           H
ATOM   5682  CB  VAL A 388      32.046  40.140  39.544  1.00  0.00           C
ATOM   5683  HB  VAL A 388      32.020  40.099  40.631  1.00  0.00           H
ATOM   5684  CG1 VAL A 388      31.658  41.551  39.082  1.00  0.00           C
ATOM   5685 HG11 VAL A 388      30.655  41.793  39.434  1.00  0.00           H
ATOM   5686 HG12 VAL A 388      32.345  42.290  39.487  1.00  0.00           H
ATOM   5687 HG13 VAL A 388      31.688  41.604  37.993  1.00  0.00           H
ATOM   5688  CG2 VAL A 388      30.991  39.150  39.021  1.00  0.00           C
ATOM   5689 HG21 VAL A 388      30.006  39.444  39.378  1.00  0.00           H
ATOM   5690 HG22 VAL A 388      30.980  39.153  37.933  1.00  0.00           H
ATOM   5691 HG23 VAL A 388      31.198  38.143  39.383  1.00  0.00           H
ATOM   5692  C   VAL A 388      34.493  40.747  39.705  1.00  0.00           C
ATOM   5693  O   VAL A 388      34.753  40.616  40.904  1.00  0.00           O
ATOM   5694  N   PRO A 389      35.061  41.717  38.956  1.00  0.00           N
ATOM   5695  CD  PRO A 389      34.941  41.887  37.515  1.00  0.00           C
ATOM   5696  HD1 PRO A 389      34.026  42.436  37.287  1.00  0.00           H
ATOM   5697  HD2 PRO A 389      34.958  40.933  36.989  1.00  0.00           H
ATOM   5698  CG  PRO A 389      36.157  42.708  37.100  1.00  0.00           C
ATOM   5699  HG1 PRO A 389      35.951  43.310  36.215  1.00  0.00           H
ATOM   5700  HG2 PRO A 389      36.998  42.038  36.928  1.00  0.00           H
ATOM   5701  CB  PRO A 389      36.436  43.579  38.322  1.00  0.00           C
ATOM   5702  HB1 PRO A 389      35.842  44.491  38.249  1.00  0.00           H
ATOM   5703  HB2 PRO A 389      37.495  43.829  38.401  1.00  0.00           H
ATOM   5704  CA  PRO A 389      35.969  42.723  39.508  1.00  0.00           C
ATOM   5705  HA  PRO A 389      36.832  42.218  39.943  1.00  0.00           H
ATOM   5706  C   PRO A 389      35.283  43.596  40.565  1.00  0.00           C
ATOM   5707  O   PRO A 389      34.185  44.100  40.337  1.00  0.00           O
ATOM   5708  N   ARG A 390      35.942  43.814  41.705  1.00  0.00           N
ATOM   5709  H   ARG A 390      36.848  43.382  41.825  1.00  0.00           H
ATOM   5710  CA  ARG A 390      35.554  44.854  42.679  1.00  0.00           C
ATOM   5711  HA  ARG A 390      34.469  44.962  42.674  1.00  0.00           H
ATOM   5712  CB  ARG A 390      35.966  44.453  44.112  1.00  0.00           C
ATOM   5713  HB1 ARG A 390      37.053  44.473  44.202  1.00  0.00           H
ATOM   5714  HB2 ARG A 390      35.553  45.189  44.805  1.00  0.00           H
ATOM   5715  CG  ARG A 390      35.455  43.056  44.506  1.00  0.00           C
ATOM   5716  HG1 ARG A 390      34.396  42.979  44.256  1.00  0.00           H
ATOM   5717  HG2 ARG A 390      35.995  42.307  43.924  1.00  0.00           H
ATOM   5718  CD  ARG A 390      35.635  42.729  45.998  1.00  0.00           C
ATOM   5719  HD1 ARG A 390      36.685  42.858  46.268  1.00  0.00           H
ATOM   5720  HD2 ARG A 390      35.023  43.407  46.594  1.00  0.00           H
ATOM   5721  NE  ARG A 390      35.228  41.338  46.249  1.00  0.00           N
```

| | | | | | | | | | | |
|---|---|---|---|---|---|---|---|---|---|---|
| ATOM | 5722 | HE | ARG | A | 390 | 34.748 | 40.891 | 45.475 | 1.00 | 0.00 | H |
| ATOM | 5723 | CZ | ARG | A | 390 | 35.442 | 40.580 | 47.315 | 1.00 | 0.00 | C |
| ATOM | 5724 | NH1 | ARG | A | 390 | 35.965 | 41.032 | 48.432 | 1.00 | 0.00 | N |
| ATOM | 5725 | HH11 | ARG | A | 390 | 36.086 | 42.026 | 48.576 | 1.00 | 0.00 | H |
| ATOM | 5726 | HH12 | ARG | A | 390 | 36.119 | 40.400 | 49.203 | 1.00 | 0.00 | H |
| ATOM | 5727 | NH2 | ARG | A | 390 | 35.117 | 39.310 | 47.265 | 1.00 | 0.00 | N |
| ATOM | 5728 | HH21 | ARG | A | 390 | 34.757 | 38.916 | 46.409 | 1.00 | 0.00 | H |
| ATOM | 5729 | HH22 | ARG | A | 390 | 35.279 | 38.684 | 48.046 | 1.00 | 0.00 | H |
| ATOM | 5730 | C | ARG | A | 390 | 36.129 | 46.230 | 42.293 | 1.00 | 0.00 | C |
| ATOM | 5731 | OC1 | ARG | A | 390 | 35.639 | 47.255 | 42.817 | 1.00 | 0.00 | O |
| ATOM | 5732 | OC2 | ARG | A | 390 | 37.097 | 46.280 | 41.502 | 1.00 | 0.00 | O |
| ATOM | 5733 | C | UNL | B | 391 | 30.678 | 17.654 | 16.357 | 1.00 | 0.00 | C |
| ATOM | 5734 | C1 | UNL | B | 391 | 30.291 | 19.027 | 15.764 | 1.00 | 0.00 | C |
| ATOM | 5735 | C2 | UNL | B | 391 | 31.425 | 20.049 | 15.977 | 1.00 | 0.00 | C |
| ATOM | 5736 | C3 | UNL | B | 391 | 31.828 | 20.085 | 17.459 | 1.00 | 0.00 | C |
| ATOM | 5737 | C4 | UNL | B | 391 | 31.977 | 18.698 | 18.120 | 1.00 | 0.00 | C |
| ATOM | 5738 | C5 | UNL | B | 391 | 31.918 | 18.774 | 19.650 | 1.00 | 0.00 | C |
| ATOM | 5739 | C6 | UNL | B | 391 | 28.899 | 19.419 | 13.715 | 1.00 | 0.00 | C |
| ATOM | 5740 | C7 | UNL | B | 391 | 28.696 | 18.992 | 12.276 | 1.00 | 0.00 | C |
| ATOM | 5741 | O | UNL | B | 391 | 30.932 | 17.772 | 17.744 | 1.00 | 0.00 | O |
| ATOM | 5742 | O1 | UNL | B | 391 | 31.076 | 21.346 | 15.498 | 1.00 | 0.00 | O |
| ATOM | 5743 | O2 | UNL | B | 391 | 33.104 | 20.793 | 17.524 | 1.00 | 0.00 | O |
| ATOM | 5744 | O3 | UNL | B | 391 | 32.907 | 17.922 | 20.278 | 1.00 | 0.00 | O |
| ATOM | 5745 | O4 | UNL | B | 391 | 28.129 | 20.236 | 14.252 | 1.00 | 0.00 | O |
| ATOM | 5746 | N | UNL | B | 391 | 29.952 | 18.844 | 14.359 | 1.00 | 0.00 | N |
| ATOM | 5747 | O5 | UNL | B | 391 | 31.713 | 17.002 | 15.635 | 1.00 | 0.00 | O |
| ATOM | 5748 | C8 | UNL | B | 391 | 33.359 | 21.722 | 18.681 | 1.00 | 0.00 | C |
| ATOM | 5749 | C9 | UNL | B | 391 | 34.524 | 22.600 | 18.192 | 1.00 | 0.00 | C |
| ATOM | 5750 | C10 | UNL | B | 391 | 34.051 | 23.666 | 17.150 | 1.00 | 0.00 | C |
| ATOM | 5751 | C11 | UNL | B | 391 | 32.524 | 23.775 | 17.080 | 1.00 | 0.00 | C |
| ATOM | 5752 | C12 | UNL | B | 391 | 31.883 | 23.644 | 18.464 | 1.00 | 0.00 | C |
| ATOM | 5753 | C13 | UNL | B | 391 | 30.367 | 23.733 | 18.451 | 1.00 | 0.00 | C |
| ATOM | 5754 | C14 | UNL | B | 391 | 35.075 | 23.548 | 20.462 | 1.00 | 0.00 | C |
| ATOM | 5755 | C15 | UNL | B | 391 | 36.228 | 23.897 | 21.372 | 1.00 | 0.00 | C |
| ATOM | 5756 | N1 | UNL | B | 391 | 35.416 | 23.088 | 19.238 | 1.00 | 0.00 | N |
| ATOM | 5757 | O6 | UNL | B | 391 | 34.588 | 23.320 | 15.849 | 1.00 | 0.00 | O |
| ATOM | 5758 | O7 | UNL | B | 391 | 32.055 | 24.998 | 16.475 | 1.00 | 0.00 | O |
| ATOM | 5759 | O8 | UNL | B | 391 | 32.186 | 22.340 | 19.029 | 1.00 | 0.00 | O |
| ATOM | 5760 | O9 | UNL | B | 391 | 29.745 | 22.817 | 17.535 | 1.00 | 0.00 | O |
| ATOM | 5761 | O10 | UNL | B | 391 | 33.884 | 23.668 | 20.833 | 1.00 | 0.00 | O |
| ATOM | 5762 | C16 | UNL | B | 391 | 31.874 | 24.934 | 15.085 | 1.00 | 0.00 | C |
| ATOM | 5763 | C17 | UNL | B | 391 | 30.687 | 25.802 | 14.628 | 1.00 | 0.00 | C |
| ATOM | 5764 | C18 | UNL | B | 391 | 30.579 | 25.801 | 13.078 | 1.00 | 0.00 | C |
| ATOM | 5765 | C19 | UNL | B | 391 | 31.950 | 26.093 | 12.427 | 1.00 | 0.00 | C |
| ATOM | 5766 | C20 | UNL | B | 391 | 33.027 | 25.191 | 13.039 | 1.00 | 0.00 | C |
| ATOM | 5767 | C21 | UNL | B | 391 | 34.450 | 25.455 | 12.589 | 1.00 | 0.00 | C |
| ATOM | 5768 | C22 | UNL | B | 391 | 28.862 | 26.005 | 16.292 | 1.00 | 0.00 | C |
| ATOM | 5769 | C23 | UNL | B | 391 | 27.697 | 25.278 | 16.922 | 1.00 | 0.00 | C |
| ATOM | 5770 | N2 | UNL | B | 391 | 29.452 | 25.347 | 15.243 | 1.00 | 0.00 | N |
| ATOM | 5771 | O11 | UNL | B | 391 | 29.602 | 26.754 | 12.698 | 1.00 | 0.00 | O |
| ATOM | 5772 | O12 | UNL | B | 391 | 31.912 | 25.984 | 11.002 | 1.00 | 0.00 | O |
| ATOM | 5773 | O13 | UNL | B | 391 | 33.056 | 25.415 | 14.458 | 1.00 | 0.00 | O |
| ATOM | 5774 | O14 | UNL | B | 391 | 35.355 | 24.514 | 13.187 | 1.00 | 0.00 | O |
| ATOM | 5775 | O15 | UNL | B | 391 | 29.249 | 27.108 | 16.694 | 1.00 | 0.00 | O |

```
ATOM   5776  H   UNL B 391      29.812  16.978  16.284  1.00  0.00           H
ATOM   5777  H1  UNL B 391      29.395  19.385  16.293  1.00  0.00           H
ATOM   5778  H2  UNL B 391      32.291  19.728  15.375  1.00  0.00           H
ATOM   5779  H3  UNL B 391      31.076  20.658  18.011  1.00  0.00           H
ATOM   5780  H4  UNL B 391      32.953  18.275  17.828  1.00  0.00           H
ATOM   5781  H5  UNL B 391      30.925  18.430  19.969  1.00  0.00           H
ATOM   5782  H6  UNL B 391      32.023  19.806  20.006  1.00  0.00           H
ATOM   5783  H7  UNL B 391      29.077  19.779  11.610  1.00  0.00           H
ATOM   5784  H8  UNL B 391      29.196  18.050  12.025  1.00  0.00           H
ATOM   5785  H9  UNL B 391      27.619  18.890  12.093  1.00  0.00           H
ATOM   5786  H10 UNL B 391      30.534  21.803  16.197  1.00  0.00           H
ATOM   5787  H11 UNL B 391      33.642  18.506  20.545  1.00  0.00           H
ATOM   5788  H12 UNL B 391      32.584  17.435  15.786  1.00  0.00           H
ATOM   5789  H13 UNL B 391      35.181  21.939  17.615  1.00  0.00           H
ATOM   5790  H14 UNL B 391      34.487  24.638  17.410  1.00  0.00           H
ATOM   5791  H15 UNL B 391      32.164  22.929  16.474  1.00  0.00           H
ATOM   5792  H16 UNL B 391      32.283  24.401  19.158  1.00  0.00           H
ATOM   5793  H17 UNL B 391      30.011  23.544  19.474  1.00  0.00           H
ATOM   5794  H18 UNL B 391      30.089  24.761  18.186  1.00  0.00           H
ATOM   5795  H19 UNL B 391      36.178  24.960  21.634  1.00  0.00           H
ATOM   5796  H20 UNL B 391      36.118  23.338  22.311  1.00  0.00           H
ATOM   5797  H21 UNL B 391      37.206  23.689  20.923  1.00  0.00           H
ATOM   5798  H22 UNL B 391      34.017  22.604  15.511  1.00  0.00           H
ATOM   5799  H23 UNL B 391      29.386  22.010  18.077  1.00  0.00           H
ATOM   5800  H24 UNL B 391      31.703  23.878  14.777  1.00  0.00           H
ATOM   5801  H25 UNL B 391      30.856  26.831  14.977  1.00  0.00           H
ATOM   5802  H26 UNL B 391      30.285  24.778  12.753  1.00  0.00           H
ATOM   5803  H27 UNL B 391      32.213  27.141  12.654  1.00  0.00           H
ATOM   5804  H28 UNL B 391      32.779  24.128  12.832  1.00  0.00           H
ATOM   5805  H29 UNL B 391      34.712  26.492  12.868  1.00  0.00           H
ATOM   5806  H30 UNL B 391      34.533  25.357  11.502  1.00  0.00           H
ATOM   5807  H31 UNL B 391      26.963  24.979  16.159  1.00  0.00           H
ATOM   5808  H32 UNL B 391      28.057  24.363  17.411  1.00  0.00           H
ATOM   5809  H33 UNL B 391      27.234  25.929  17.669  1.00  0.00           H
ATOM   5810  H34 UNL B 391      29.095  26.451  11.883  1.00  0.00           H
ATOM   5811  H35 UNL B 391      31.329  25.218  10.746  1.00  0.00           H
ATOM   5812  H36 UNL B 391      35.076  24.432  14.128  1.00  0.00           H
ATOM   5813  H37 UNL B 391      33.673  21.108  19.534  1.00  0.00           H
ATOM   5814  H38 UNL B 391      30.568  18.227  13.828  1.00  0.00           H
ATOM   5815  H39 UNL B 391      36.433  22.935  19.017  1.00  0.00           H
ATOM   5816  H40 UNL B 391      29.227  24.356  15.154  1.00  0.00           H
ATOM   5817  K      K   392      38.897  18.408  19.770  1.00  0.00           K
TER
ENDMDL
```
