## SupplementaryInformation1 for "Mechanism of catalytic apparatus of human chitotriosidase-1 and its dual inactivation mode by the *first-in-class* OATD-01 inhibitor"

```
TITLE     Atomic coordinates of the full length isoform of hCHIT1 - model
CRYST1   60.857   67.465   66.703  90.00  90.00  90.00 P 1           1
MODEL        1
ATOM      1  N   ALA A  22      25.940  36.521  51.237  1.00  0.00           N
ATOM      2  H1  ALA A  22      25.869  35.825  50.522  1.00  0.00           H
ATOM      3  H2  ALA A  22      26.878  36.550  51.582  1.00  0.00           H
ATOM      4  H3  ALA A  22      25.316  36.295  51.985  1.00  0.00           H
ATOM      5  CA  ALA A  22      25.582  37.835  50.680  1.00  0.00           C
ATOM      6  HA  ALA A  22      24.644  37.711  50.357  1.00  0.00           H
ATOM      7  CB  ALA A  22      25.695  38.896  51.788  1.00  0.00           C
ATOM      8  HB1 ALA A  22      25.453  39.793  51.417  1.00  0.00           H
ATOM      9  HB2 ALA A  22      25.072  38.665  52.535  1.00  0.00           H
ATOM     10  HB3 ALA A  22      26.634  38.920  52.132  1.00  0.00           H
ATOM     11  C   ALA A  22      26.353  38.252  49.442  1.00  0.00           C
ATOM     12  O   ALA A  22      26.718  39.401  49.231  1.00  0.00           O
ATOM     13  N   LYS A  23      26.573  37.219  48.611  1.00  0.00           N
ATOM     14  H   LYS A  23      26.295  36.298  48.884  1.00  0.00           H
ATOM     15  CA  LYS A  23      27.209  37.424  47.322  1.00  0.00           C
ATOM     16  HA  LYS A  23      27.230  38.414  47.186  1.00  0.00           H
ATOM     17  CB  LYS A  23      28.536  36.707  47.437  1.00  0.00           C
ATOM     18  HB1 LYS A  23      28.396  35.738  47.233  1.00  0.00           H
ATOM     19  HB2 LYS A  23      28.874  36.806  48.373  1.00  0.00           H
ATOM     20  CG  LYS A  23      29.566  37.296  46.453  1.00  0.00           C
ATOM     21  HG1 LYS A  23      29.697  38.261  46.682  1.00  0.00           H
ATOM     22  HG2 LYS A  23      29.190  37.221  45.529  1.00  0.00           H
ATOM     23  CD  LYS A  23      30.942  36.627  46.452  1.00  0.00           C
ATOM     24  HD1 LYS A  23      31.596  37.256  46.032  1.00  0.00           H
ATOM     25  HD2 LYS A  23      30.884  35.792  45.905  1.00  0.00           H
ATOM     26  CE  LYS A  23      31.465  36.253  47.836  1.00  0.00           C
ATOM     27  HE1 LYS A  23      32.267  35.662  47.744  1.00  0.00           H
ATOM     28  HE2 LYS A  23      30.752  35.771  48.346  1.00  0.00           H
ATOM     29  NZ  LYS A  23      31.859  37.437  48.613  1.00  0.00           N
ATOM     30  HZ1 LYS A  23      32.195  37.150  49.510  1.00  0.00           H
ATOM     31  HZ2 LYS A  23      32.582  37.929  48.128  1.00  0.00           H
ATOM     32  HZ3 LYS A  23      31.068  38.038  48.730  1.00  0.00           H
ATOM     33  C   LYS A  23      26.485  36.753  46.137  1.00  0.00           C
ATOM     34  O   LYS A  23      25.866  35.747  46.353  1.00  0.00           O
ATOM     35  N   LEU A  24      26.622  37.325  44.934  1.00  0.00           N
ATOM     36  H   LEU A  24      27.216  38.129  44.890  1.00  0.00           H
ATOM     37  CA  LEU A  24      25.993  36.902  43.658  1.00  0.00           C
ATOM     38  HA  LEU A  24      25.866  35.910  43.644  1.00  0.00           H
ATOM     39  CB  LEU A  24      24.593  37.427  43.517  1.00  0.00           C
ATOM     40  HB1 LEU A  24      24.654  38.403  43.725  1.00  0.00           H
ATOM     41  HB2 LEU A  24      24.064  36.967  44.230  1.00  0.00           H
ATOM     42  CG  LEU A  24      23.768  37.319  42.246  1.00  0.00           C
ATOM     43  HG  LEU A  24      24.322  37.885  41.635  1.00  0.00           H
ATOM     44  CD1 LEU A  24      23.737  35.967  41.592  1.00  0.00           C
ATOM     45 HD11 LEU A  24      23.170  36.008  40.770  1.00  0.00           H
ATOM     46 HD12 LEU A  24      24.667  35.696  41.343  1.00  0.00           H
ATOM     47 HD13 LEU A  24      23.354  35.298  42.229  1.00  0.00           H
ATOM     48  CD2 LEU A  24      22.322  37.897  42.348  1.00  0.00           C
ATOM     49 HD21 LEU A  24      21.855  37.784  41.471  1.00  0.00           H
ATOM     50 HD22 LEU A  24      21.817  37.409  43.060  1.00  0.00           H
ATOM     51 HD23 LEU A  24      22.367  38.869  42.579  1.00  0.00           H
```

| | | | | | | | | | | | |
|---|---|---|---|---|---|---|---|---|---|---|---|
| ATOM | 52 | C | LEU | A | 24 | 26.922 | 37.190 | 42.540 | 1.00 | 0.00 | C |
| ATOM | 53 | O | LEU | A | 24 | 26.929 | 38.294 | 41.884 | 1.00 | 0.00 | O |
| ATOM | 54 | N | VAL | A | 25 | 27.734 | 36.195 | 42.118 | 1.00 | 0.00 | N |
| ATOM | 55 | H | VAL | A | 25 | 27.760 | 35.357 | 42.663 | 1.00 | 0.00 | H |
| ATOM | 56 | CA | VAL | A | 25 | 28.582 | 36.252 | 40.917 | 1.00 | 0.00 | C |
| ATOM | 57 | HA | VAL | A | 25 | 28.882 | 37.196 | 40.782 | 1.00 | 0.00 | H |
| ATOM | 58 | CB | VAL | A | 25 | 29.832 | 35.334 | 41.142 | 1.00 | 0.00 | C |
| ATOM | 59 | HB | VAL | A | 25 | 29.465 | 34.409 | 41.242 | 1.00 | 0.00 | H |
| ATOM | 60 | CG1 | VAL | A | 25 | 30.783 | 35.676 | 39.987 | 1.00 | 0.00 | C |
| ATOM | 61 | HG11 | VAL | A | 25 | 31.613 | 35.123 | 40.064 | 1.00 | 0.00 | H |
| ATOM | 62 | HG12 | VAL | A | 25 | 30.333 | 35.482 | 39.116 | 1.00 | 0.00 | H |
| ATOM | 63 | HG13 | VAL | A | 25 | 31.023 | 36.646 | 40.029 | 1.00 | 0.00 | H |
| ATOM | 64 | CG2 | VAL | A | 25 | 30.687 | 35.477 | 42.370 | 1.00 | 0.00 | C |
| ATOM | 65 | HG21 | VAL | A | 25 | 31.432 | 34.810 | 42.337 | 1.00 | 0.00 | H |
| ATOM | 66 | HG22 | VAL | A | 25 | 31.068 | 36.401 | 42.405 | 1.00 | 0.00 | H |
| ATOM | 67 | HG23 | VAL | A | 25 | 30.130 | 35.315 | 43.185 | 1.00 | 0.00 | H |
| ATOM | 68 | C | VAL | A | 25 | 27.755 | 35.888 | 39.682 | 1.00 | 0.00 | C |
| ATOM | 69 | O | VAL | A | 25 | 26.812 | 35.075 | 39.696 | 1.00 | 0.00 | O |
| ATOM | 70 | N | CYS | A | 26 | 28.048 | 36.513 | 38.589 | 1.00 | 0.00 | N |
| ATOM | 71 | H | CYS | A | 26 | 28.724 | 37.248 | 38.628 | 1.00 | 0.00 | H |
| ATOM | 72 | CA | CYS | A | 26 | 27.413 | 36.183 | 37.248 | 1.00 | 0.00 | C |
| ATOM | 73 | HA | CYS | A | 26 | 27.179 | 35.211 | 37.261 | 1.00 | 0.00 | H |
| ATOM | 74 | CB | CYS | A | 26 | 26.205 | 37.134 | 37.112 | 1.00 | 0.00 | C |
| ATOM | 75 | HB1 | CYS | A | 26 | 25.731 | 36.808 | 36.294 | 1.00 | 0.00 | H |
| ATOM | 76 | HB2 | CYS | A | 26 | 26.619 | 38.025 | 36.924 | 1.00 | 0.00 | H |
| ATOM | 77 | SG | CYS | A | 26 | 25.087 | 37.241 | 38.479 | 1.00 | 0.00 | S |
| ATOM | 78 | C | CYS | A | 26 | 28.353 | 36.429 | 36.023 | 1.00 | 0.00 | C |
| ATOM | 79 | O | CYS | A | 26 | 29.023 | 37.443 | 36.098 | 1.00 | 0.00 | O |
| ATOM | 80 | N | TYR | A | 27 | 28.538 | 35.422 | 35.152 | 1.00 | 0.00 | N |
| ATOM | 81 | H | TYR | A | 27 | 27.999 | 34.590 | 35.286 | 1.00 | 0.00 | H |
| ATOM | 82 | CA | TYR | A | 27 | 29.475 | 35.450 | 34.014 | 1.00 | 0.00 | C |
| ATOM | 83 | HA | TYR | A | 27 | 30.203 | 36.056 | 34.334 | 1.00 | 0.00 | H |
| ATOM | 84 | CB | TYR | A | 27 | 29.998 | 34.035 | 33.710 | 1.00 | 0.00 | C |
| ATOM | 85 | HB1 | TYR | A | 27 | 30.479 | 34.062 | 32.834 | 1.00 | 0.00 | H |
| ATOM | 86 | HB2 | TYR | A | 27 | 29.212 | 33.420 | 33.640 | 1.00 | 0.00 | H |
| ATOM | 87 | CG | TYR | A | 27 | 30.949 | 33.446 | 34.736 | 1.00 | 0.00 | C |
| ATOM | 88 | CD1 | TYR | A | 27 | 30.464 | 32.580 | 35.734 | 1.00 | 0.00 | C |
| ATOM | 89 | HD1 | TYR | A | 27 | 29.484 | 32.391 | 35.799 | 1.00 | 0.00 | H |
| ATOM | 90 | CE1 | TYR | A | 27 | 31.357 | 31.980 | 36.639 | 1.00 | 0.00 | C |
| ATOM | 91 | HE1 | TYR | A | 27 | 31.009 | 31.354 | 37.337 | 1.00 | 0.00 | H |
| ATOM | 92 | CZ | TYR | A | 27 | 32.734 | 32.252 | 36.572 | 1.00 | 0.00 | C |
| ATOM | 93 | OH | TYR | A | 27 | 33.559 | 31.688 | 37.491 | 1.00 | 0.00 | O |
| ATOM | 94 | HH | TYR | A | 27 | 34.501 | 31.974 | 37.313 | 1.00 | 0.00 | H |
| ATOM | 95 | CE2 | TYR | A | 27 | 33.224 | 33.112 | 35.567 | 1.00 | 0.00 | C |
| ATOM | 96 | HE2 | TYR | A | 27 | 34.204 | 33.304 | 35.508 | 1.00 | 0.00 | H |
| ATOM | 97 | CD2 | TYR | A | 27 | 32.333 | 33.701 | 34.646 | 1.00 | 0.00 | C |
| ATOM | 98 | HD2 | TYR | A | 27 | 32.684 | 34.301 | 33.927 | 1.00 | 0.00 | H |
| ATOM | 99 | C | TYR | A | 27 | 28.861 | 36.038 | 32.728 | 1.00 | 0.00 | C |
| ATOM | 100 | O | TYR | A | 27 | 27.763 | 35.658 | 32.320 | 1.00 | 0.00 | O |
| ATOM | 101 | N | PHE | A | 28 | 29.614 | 36.901 | 32.024 | 1.00 | 0.00 | N |
| ATOM | 102 | H | PHE | A | 28 | 30.448 | 37.222 | 32.472 | 1.00 | 0.00 | H |
| ATOM | 103 | CA | PHE | A | 28 | 29.368 | 37.416 | 30.713 | 1.00 | 0.00 | C |
| ATOM | 104 | HA | PHE | A | 28 | 28.411 | 37.299 | 30.448 | 1.00 | 0.00 | H |
| ATOM | 105 | CB | PHE | A | 28 | 29.719 | 38.891 | 30.790 | 1.00 | 0.00 | C |

| | | | | | | | | | | | |
|---|---|---|---|---|---|---|---|---|---|---|---|
| ATOM | 106 | HB1 | PHE | A | 28 | 30.646 | 38.984 | 31.154 | 1.00 | 0.00 | H |
| ATOM | 107 | HB2 | PHE | A | 28 | 29.073 | 39.346 | 31.403 | 1.00 | 0.00 | H |
| ATOM | 108 | CG | PHE | A | 28 | 29.655 | 39.555 | 29.422 | 1.00 | 0.00 | C |
| ATOM | 109 | CD1 | PHE | A | 28 | 28.581 | 40.387 | 28.983 | 1.00 | 0.00 | C |
| ATOM | 110 | HD1 | PHE | A | 28 | 27.804 | 40.561 | 29.588 | 1.00 | 0.00 | H |
| ATOM | 111 | CE1 | PHE | A | 28 | 28.613 | 40.940 | 27.748 | 1.00 | 0.00 | C |
| ATOM | 112 | HE1 | PHE | A | 28 | 27.849 | 41.525 | 27.475 | 1.00 | 0.00 | H |
| ATOM | 113 | CZ | PHE | A | 28 | 29.650 | 40.742 | 26.830 | 1.00 | 0.00 | C |
| ATOM | 114 | HZ | PHE | A | 28 | 29.596 | 41.098 | 25.897 | 1.00 | 0.00 | H |
| ATOM | 115 | CE2 | PHE | A | 28 | 30.766 | 40.025 | 27.273 | 1.00 | 0.00 | C |
| ATOM | 116 | HE2 | PHE | A | 28 | 31.561 | 39.946 | 26.672 | 1.00 | 0.00 | H |
| ATOM | 117 | CD2 | PHE | A | 28 | 30.803 | 39.429 | 28.503 | 1.00 | 0.00 | C |
| ATOM | 118 | HD2 | PHE | A | 28 | 31.609 | 38.907 | 28.780 | 1.00 | 0.00 | H |
| ATOM | 119 | C | PHE | A | 28 | 30.139 | 36.623 | 29.703 | 1.00 | 0.00 | C |
| ATOM | 120 | O | PHE | A | 28 | 31.388 | 36.570 | 29.688 | 1.00 | 0.00 | O |
| ATOM | 121 | N | THR | A | 29 | 29.476 | 35.992 | 28.719 | 1.00 | 0.00 | N |
| ATOM | 122 | H | THR | A | 29 | 28.477 | 36.030 | 28.702 | 1.00 | 0.00 | H |
| ATOM | 123 | CA | THR | A | 29 | 30.189 | 35.240 | 27.656 | 1.00 | 0.00 | C |
| ATOM | 124 | HA | THR | A | 29 | 31.058 | 35.012 | 28.095 | 1.00 | 0.00 | H |
| ATOM | 125 | CB | THR | A | 29 | 29.503 | 33.918 | 27.274 | 1.00 | 0.00 | C |
| ATOM | 126 | HB | THR | A | 29 | 30.069 | 33.480 | 26.575 | 1.00 | 0.00 | H |
| ATOM | 127 | CG2 | THR | A | 29 | 29.295 | 32.994 | 28.475 | 1.00 | 0.00 | C |
| ATOM | 128 | HG21 | THR | A | 29 | 28.847 | 32.152 | 28.174 | 1.00 | 0.00 | H |
| ATOM | 129 | HG22 | THR | A | 29 | 30.181 | 32.773 | 28.883 | 1.00 | 0.00 | H |
| ATOM | 130 | HG23 | THR | A | 29 | 28.721 | 33.453 | 29.153 | 1.00 | 0.00 | H |
| ATOM | 131 | OG1 | THR | A | 29 | 28.281 | 34.096 | 26.610 | 1.00 | 0.00 | O |
| ATOM | 132 | HG1 | THR | A | 29 | 27.888 | 33.203 | 26.390 | 1.00 | 0.00 | H |
| ATOM | 133 | C | THR | A | 29 | 30.489 | 36.098 | 26.420 | 1.00 | 0.00 | C |
| ATOM | 134 | O | THR | A | 29 | 29.586 | 36.584 | 25.734 | 1.00 | 0.00 | O |
| ATOM | 135 | N | ASN | A | 30 | 31.769 | 36.247 | 26.049 | 1.00 | 0.00 | N |
| ATOM | 136 | H | ASN | A | 30 | 32.479 | 35.769 | 26.566 | 1.00 | 0.00 | H |
| ATOM | 137 | CA | ASN | A | 30 | 32.168 | 37.088 | 24.910 | 1.00 | 0.00 | C |
| ATOM | 138 | HA | ASN | A | 30 | 31.538 | 37.859 | 24.999 | 1.00 | 0.00 | H |
| ATOM | 139 | CB | ASN | A | 30 | 33.613 | 37.621 | 25.061 | 1.00 | 0.00 | C |
| ATOM | 140 | HB1 | ASN | A | 30 | 33.755 | 37.868 | 26.020 | 1.00 | 0.00 | H |
| ATOM | 141 | HB2 | ASN | A | 30 | 33.705 | 38.437 | 24.490 | 1.00 | 0.00 | H |
| ATOM | 142 | CG | ASN | A | 30 | 34.728 | 36.662 | 24.668 | 1.00 | 0.00 | C |
| ATOM | 143 | OD1 | ASN | A | 30 | 34.550 | 35.462 | 24.561 | 1.00 | 0.00 | O |
| ATOM | 144 | ND2 | ASN | A | 30 | 35.932 | 37.142 | 24.451 | 1.00 | 0.00 | N |
| ATOM | 145 | HD21 | ASN | A | 30 | 36.104 | 38.123 | 24.545 | 1.00 | 0.00 | H |
| ATOM | 146 | HD22 | ASN | A | 30 | 36.676 | 36.527 | 24.192 | 1.00 | 0.00 | H |
| ATOM | 147 | C | ASN | A | 30 | 31.855 | 36.499 | 23.516 | 1.00 | 0.00 | C |
| ATOM | 148 | O | ASN | A | 30 | 32.272 | 37.082 | 22.524 | 1.00 | 0.00 | O |
| ATOM | 149 | N | TRP | A | 31 | 31.047 | 35.437 | 23.405 | 1.00 | 0.00 | N |
| ATOM | 150 | H | TRP | A | 31 | 30.747 | 34.987 | 24.246 | 1.00 | 0.00 | H |
| ATOM | 151 | CA | TRP | A | 31 | 30.571 | 34.889 | 22.123 | 1.00 | 0.00 | C |
| ATOM | 152 | HA | TRP | A | 31 | 30.983 | 35.492 | 21.440 | 1.00 | 0.00 | H |
| ATOM | 153 | CB | TRP | A | 31 | 31.103 | 33.463 | 21.919 | 1.00 | 0.00 | C |
| ATOM | 154 | HB1 | TRP | A | 31 | 32.101 | 33.505 | 21.866 | 1.00 | 0.00 | H |
| ATOM | 155 | HB2 | TRP | A | 31 | 30.738 | 33.112 | 21.057 | 1.00 | 0.00 | H |
| ATOM | 156 | CG | TRP | A | 31 | 30.762 | 32.464 | 22.978 | 1.00 | 0.00 | C |
| ATOM | 157 | CD1 | TRP | A | 31 | 29.838 | 31.484 | 22.872 | 1.00 | 0.00 | C |
| ATOM | 158 | HD1 | TRP | A | 31 | 29.260 | 31.321 | 22.072 | 1.00 | 0.00 | H |
| ATOM | 159 | NE1 | TRP | A | 31 | 29.812 | 30.742 | 24.034 | 1.00 | 0.00 | N |

| | | | | | | | | | | | |
|---|---|---|---|---|---|---|---|---|---|---|---|
| ATOM | 160 | HE1 | TRP | A | 31 | 29.219 | 29.951 | 24.182 | 1.00 | 0.00 | H |
| ATOM | 161 | CE2 | TRP | A | 31 | 30.704 | 31.225 | 24.964 | 1.00 | 0.00 | C |
| ATOM | 162 | CZ2 | TRP | A | 31 | 31.047 | 30.833 | 26.262 | 1.00 | 0.00 | C |
| ATOM | 163 | HZ2 | TRP | A | 31 | 30.547 | 30.094 | 26.713 | 1.00 | 0.00 | H |
| ATOM | 164 | CH2 | TRP | A | 31 | 32.100 | 31.485 | 26.928 | 1.00 | 0.00 | C |
| ATOM | 165 | HH2 | TRP | A | 31 | 32.369 | 31.196 | 27.847 | 1.00 | 0.00 | H |
| ATOM | 166 | CZ3 | TRP | A | 31 | 32.771 | 32.544 | 26.295 | 1.00 | 0.00 | C |
| ATOM | 167 | HZ3 | TRP | A | 31 | 33.513 | 33.018 | 26.769 | 1.00 | 0.00 | H |
| ATOM | 168 | CE3 | TRP | A | 31 | 32.402 | 32.941 | 24.996 | 1.00 | 0.00 | C |
| ATOM | 169 | HE3 | TRP | A | 31 | 32.890 | 33.697 | 24.559 | 1.00 | 0.00 | H |
| ATOM | 170 | CD2 | TRP | A | 31 | 31.361 | 32.298 | 24.298 | 1.00 | 0.00 | C |
| ATOM | 171 | C | TRP | A | 31 | 29.054 | 35.022 | 21.889 | 1.00 | 0.00 | C |
| ATOM | 172 | O | TRP | A | 31 | 28.610 | 34.853 | 20.754 | 1.00 | 0.00 | O |
| ATOM | 173 | N | ALA | A | 32 | 28.275 | 35.513 | 22.862 | 1.00 | 0.00 | N |
| ATOM | 174 | H | ALA | A | 32 | 28.669 | 35.616 | 23.775 | 1.00 | 0.00 | H |
| ATOM | 175 | CA | ALA | A | 32 | 26.868 | 35.910 | 22.660 | 1.00 | 0.00 | C |
| ATOM | 176 | HA | ALA | A | 32 | 26.415 | 35.127 | 22.234 | 1.00 | 0.00 | H |
| ATOM | 177 | CB | ALA | A | 32 | 26.295 | 36.236 | 24.029 | 1.00 | 0.00 | C |
| ATOM | 178 | HB1 | ALA | A | 32 | 25.338 | 36.511 | 23.933 | 1.00 | 0.00 | H |
| ATOM | 179 | HB2 | ALA | A | 32 | 26.353 | 35.428 | 24.615 | 1.00 | 0.00 | H |
| ATOM | 180 | HB3 | ALA | A | 32 | 26.817 | 36.984 | 24.439 | 1.00 | 0.00 | H |
| ATOM | 181 | C | ALA | A | 32 | 26.686 | 37.109 | 21.703 | 1.00 | 0.00 | C |
| ATOM | 182 | O | ALA | A | 32 | 25.613 | 37.233 | 21.041 | 1.00 | 0.00 | O |
| ATOM | 183 | N | GLN | A | 33 | 27.758 | 37.900 | 21.503 | 1.00 | 0.00 | N |
| ATOM | 184 | H | GLN | A | 33 | 28.512 | 37.860 | 22.158 | 1.00 | 0.00 | H |
| ATOM | 185 | CA | GLN | A | 33 | 27.865 | 38.819 | 20.362 | 1.00 | 0.00 | C |
| ATOM | 186 | HA | GLN | A | 33 | 27.018 | 39.340 | 20.462 | 1.00 | 0.00 | H |
| ATOM | 187 | CB | GLN | A | 33 | 29.110 | 39.713 | 20.495 | 1.00 | 0.00 | C |
| ATOM | 188 | HB1 | GLN | A | 33 | 29.064 | 40.182 | 21.377 | 1.00 | 0.00 | H |
| ATOM | 189 | HB2 | GLN | A | 33 | 29.089 | 40.388 | 19.758 | 1.00 | 0.00 | H |
| ATOM | 190 | CG | GLN | A | 33 | 30.451 | 38.957 | 20.419 | 1.00 | 0.00 | C |
| ATOM | 191 | HG1 | GLN | A | 33 | 30.573 | 38.609 | 19.490 | 1.00 | 0.00 | H |
| ATOM | 192 | HG2 | GLN | A | 33 | 30.427 | 38.191 | 21.062 | 1.00 | 0.00 | H |
| ATOM | 193 | CD | GLN | A | 33 | 31.647 | 39.844 | 20.766 | 1.00 | 0.00 | C |
| ATOM | 194 | OE1 | GLN | A | 33 | 32.128 | 40.625 | 19.960 | 1.00 | 0.00 | O |
| ATOM | 195 | NE2 | GLN | A | 33 | 32.116 | 39.824 | 21.997 | 1.00 | 0.00 | N |
| ATOM | 196 | HE21 | GLN | A | 33 | 31.690 | 39.236 | 22.685 | 1.00 | 0.00 | H |
| ATOM | 197 | HE22 | GLN | A | 33 | 32.898 | 40.397 | 22.244 | 1.00 | 0.00 | H |
| ATOM | 198 | C | GLN | A | 33 | 27.813 | 38.176 | 18.960 | 1.00 | 0.00 | C |
| ATOM | 199 | O | GLN | A | 33 | 27.374 | 38.822 | 18.008 | 1.00 | 0.00 | O |
| ATOM | 200 | N | TYR | A | 34 | 28.211 | 36.910 | 18.811 | 1.00 | 0.00 | N |
| ATOM | 201 | H | TYR | A | 34 | 28.542 | 36.421 | 19.618 | 1.00 | 0.00 | H |
| ATOM | 202 | CA | TYR | A | 34 | 28.189 | 36.196 | 17.526 | 1.00 | 0.00 | C |
| ATOM | 203 | HA | TYR | A | 34 | 28.231 | 36.930 | 16.848 | 1.00 | 0.00 | H |
| ATOM | 204 | CB | TYR | A | 34 | 29.394 | 35.269 | 17.422 | 1.00 | 0.00 | C |
| ATOM | 205 | HB1 | TYR | A | 34 | 29.404 | 34.875 | 16.503 | 1.00 | 0.00 | H |
| ATOM | 206 | HB2 | TYR | A | 34 | 29.283 | 34.539 | 18.096 | 1.00 | 0.00 | H |
| ATOM | 207 | CG | TYR | A | 34 | 30.741 | 35.916 | 17.661 | 1.00 | 0.00 | C |
| ATOM | 208 | CD1 | TYR | A | 34 | 31.195 | 36.992 | 16.869 | 1.00 | 0.00 | C |
| ATOM | 209 | HD1 | TYR | A | 34 | 30.586 | 37.409 | 16.195 | 1.00 | 0.00 | H |
| ATOM | 210 | CE1 | TYR | A | 34 | 32.509 | 37.478 | 17.030 | 1.00 | 0.00 | C |
| ATOM | 211 | HE1 | TYR | A | 34 | 32.837 | 38.233 | 16.463 | 1.00 | 0.00 | H |
| ATOM | 212 | CZ | TYR | A | 34 | 33.359 | 36.895 | 17.994 | 1.00 | 0.00 | C |
| ATOM | 213 | OH | TYR | A | 34 | 34.665 | 37.248 | 18.070 | 1.00 | 0.00 | O |

| | | | | | | | | | | | |
|---|---|---|---|---|---|---|---|---|---|---|---|
| ATOM | 214 | HH | TYR | A | 34 | 35.106 | 36.736 | 18.807 | 1.00 | 0.00 | H |
| ATOM | 215 | CE2 | TYR | A | 34 | 32.865 | 35.875 | 18.830 | 1.00 | 0.00 | C |
| ATOM | 216 | HE2 | TYR | A | 34 | 33.440 | 35.510 | 19.562 | 1.00 | 0.00 | H |
| ATOM | 217 | CD2 | TYR | A | 34 | 31.575 | 35.369 | 18.643 | 1.00 | 0.00 | C |
| ATOM | 218 | HD2 | TYR | A | 34 | 31.248 | 34.615 | 19.212 | 1.00 | 0.00 | H |
| ATOM | 219 | C | TYR | A | 34 | 26.904 | 35.397 | 17.280 | 1.00 | 0.00 | C |
| ATOM | 220 | O | TYR | A | 34 | 26.682 | 34.918 | 16.166 | 1.00 | 0.00 | O |
| ATOM | 221 | N | ARG | A | 35 | 26.024 | 35.300 | 18.284 | 1.00 | 0.00 | N |
| ATOM | 222 | H | ARG | A | 35 | 26.340 | 35.479 | 19.216 | 1.00 | 0.00 | H |
| ATOM | 223 | CA | ARG | A | 35 | 24.615 | 34.942 | 18.078 | 1.00 | 0.00 | C |
| ATOM | 224 | HA | ARG | A | 35 | 24.582 | 34.096 | 17.546 | 1.00 | 0.00 | H |
| ATOM | 225 | CB | ARG | A | 35 | 23.934 | 34.680 | 19.435 | 1.00 | 0.00 | C |
| ATOM | 226 | HB1 | ARG | A | 35 | 22.942 | 34.646 | 19.313 | 1.00 | 0.00 | H |
| ATOM | 227 | HB2 | ARG | A | 35 | 24.166 | 35.413 | 20.075 | 1.00 | 0.00 | H |
| ATOM | 228 | CG | ARG | A | 35 | 24.434 | 33.333 | 19.988 | 1.00 | 0.00 | C |
| ATOM | 229 | HG1 | ARG | A | 35 | 25.433 | 33.344 | 20.031 | 1.00 | 0.00 | H |
| ATOM | 230 | HG2 | ARG | A | 35 | 24.135 | 32.596 | 19.382 | 1.00 | 0.00 | H |
| ATOM | 231 | CD | ARG | A | 35 | 23.883 | 33.076 | 21.386 | 1.00 | 0.00 | C |
| ATOM | 232 | HD1 | ARG | A | 35 | 22.942 | 32.744 | 21.314 | 1.00 | 0.00 | H |
| ATOM | 233 | HD2 | ARG | A | 35 | 23.896 | 33.929 | 21.907 | 1.00 | 0.00 | H |
| ATOM | 234 | NE | ARG | A | 35 | 24.687 | 32.057 | 22.128 | 1.00 | 0.00 | N |
| ATOM | 235 | HE | ARG | A | 35 | 24.928 | 31.244 | 21.597 | 1.00 | 0.00 | H |
| ATOM | 236 | CZ | ARG | A | 35 | 25.135 | 32.059 | 23.389 | 1.00 | 0.00 | C |
| ATOM | 237 | NH1 | ARG | A | 35 | 24.687 | 32.842 | 24.350 | 1.00 | 0.00 | N |
| ATOM | 238 | HH11 | ARG | A | 35 | 23.957 | 33.498 | 24.161 | 1.00 | 0.00 | H |
| ATOM | 239 | HH12 | ARG | A | 35 | 25.078 | 32.780 | 25.268 | 1.00 | 0.00 | H |
| ATOM | 240 | NH2 | ARG | A | 35 | 26.094 | 31.240 | 23.721 | 1.00 | 0.00 | N |
| ATOM | 241 | HH21 | ARG | A | 35 | 26.483 | 30.623 | 23.037 | 1.00 | 0.00 | H |
| ATOM | 242 | HH22 | ARG | A | 35 | 26.440 | 31.229 | 24.659 | 1.00 | 0.00 | H |
| ATOM | 243 | C | ARG | A | 35 | 23.934 | 36.047 | 17.271 | 1.00 | 0.00 | C |
| ATOM | 244 | O | ARG | A | 35 | 24.435 | 37.170 | 17.191 | 1.00 | 0.00 | O |
| ATOM | 245 | N | GLN | A | 36 | 22.826 | 35.732 | 16.608 | 1.00 | 0.00 | N |
| ATOM | 246 | H | GLN | A | 36 | 22.454 | 34.807 | 16.687 | 1.00 | 0.00 | H |
| ATOM | 247 | CA | GLN | A | 36 | 22.117 | 36.743 | 15.738 | 1.00 | 0.00 | C |
| ATOM | 248 | HA | GLN | A | 36 | 22.601 | 37.618 | 15.739 | 1.00 | 0.00 | H |
| ATOM | 249 | CB | GLN | A | 36 | 22.038 | 36.209 | 14.296 | 1.00 | 0.00 | C |
| ATOM | 250 | HB1 | GLN | A | 36 | 21.617 | 36.916 | 13.728 | 1.00 | 0.00 | H |
| ATOM | 251 | HB2 | GLN | A | 36 | 21.457 | 35.395 | 14.300 | 1.00 | 0.00 | H |
| ATOM | 252 | CG | GLN | A | 36 | 23.392 | 35.828 | 13.662 | 1.00 | 0.00 | C |
| ATOM | 253 | HG1 | GLN | A | 36 | 24.060 | 36.523 | 13.927 | 1.00 | 0.00 | H |
| ATOM | 254 | HG2 | GLN | A | 36 | 23.278 | 35.836 | 12.669 | 1.00 | 0.00 | H |
| ATOM | 255 | CD | GLN | A | 36 | 23.969 | 34.460 | 14.050 | 1.00 | 0.00 | C |
| ATOM | 256 | OE1 | GLN | A | 36 | 23.370 | 33.619 | 14.716 | 1.00 | 0.00 | O |
| ATOM | 257 | NE2 | GLN | A | 36 | 25.171 | 34.172 | 13.610 | 1.00 | 0.00 | N |
| ATOM | 258 | HE21 | GLN | A | 36 | 25.669 | 34.832 | 13.048 | 1.00 | 0.00 | H |
| ATOM | 259 | HE22 | GLN | A | 36 | 25.589 | 33.292 | 13.837 | 1.00 | 0.00 | H |
| ATOM | 260 | C | GLN | A | 36 | 20.801 | 37.048 | 16.308 | 1.00 | 0.00 | C |
| ATOM | 261 | O | GLN | A | 36 | 20.141 | 36.259 | 16.977 | 1.00 | 0.00 | O |
| ATOM | 262 | N | GLY | A | 37 | 20.450 | 38.347 | 16.140 | 1.00 | 0.00 | N |
| ATOM | 263 | H | GLY | A | 37 | 21.146 | 38.965 | 15.774 | 1.00 | 0.00 | H |
| ATOM | 264 | CA | GLY | A | 37 | 19.113 | 38.935 | 16.454 | 1.00 | 0.00 | C |
| ATOM | 265 | HA1 | GLY | A | 37 | 18.427 | 38.421 | 15.939 | 1.00 | 0.00 | H |
| ATOM | 266 | HA2 | GLY | A | 37 | 19.119 | 39.886 | 16.145 | 1.00 | 0.00 | H |
| ATOM | 267 | C | GLY | A | 37 | 18.762 | 38.889 | 17.963 | 1.00 | 0.00 | C |

| | | | | | | | | | | | |
|---|---|---|---|---|---|---|---|---|---|---|---|
| ATOM | 268 | O | GLY | A | 37 | 19.512 | 39.338 | 18.856 | 1.00 | 0.00 | O |
| ATOM | 269 | N | GLU | A | 38 | 17.590 | 38.277 | 18.307 | 1.00 | 0.00 | N |
| ATOM | 270 | H | GLU | A | 38 | 17.101 | 37.787 | 17.585 | 1.00 | 0.00 | H |
| ATOM | 271 | CA | GLU | A | 38 | 17.006 | 38.277 | 19.605 | 1.00 | 0.00 | C |
| ATOM | 272 | HA | GLU | A | 38 | 16.913 | 39.254 | 19.795 | 1.00 | 0.00 | H |
| ATOM | 273 | CB | GLU | A | 38 | 15.698 | 37.525 | 19.500 | 1.00 | 0.00 | C |
| ATOM | 274 | HB1 | GLU | A | 38 | 15.314 | 37.522 | 20.423 | 1.00 | 0.00 | H |
| ATOM | 275 | HB2 | GLU | A | 38 | 15.944 | 36.592 | 19.238 | 1.00 | 0.00 | H |
| ATOM | 276 | CG | GLU | A | 38 | 14.567 | 37.975 | 18.546 | 1.00 | 0.00 | C |
| ATOM | 277 | HG1 | GLU | A | 38 | 13.709 | 37.561 | 18.850 | 1.00 | 0.00 | H |
| ATOM | 278 | HG2 | GLU | A | 38 | 14.782 | 37.657 | 17.623 | 1.00 | 0.00 | H |
| ATOM | 279 | CD | GLU | A | 38 | 14.406 | 39.529 | 18.527 | 1.00 | 0.00 | C |
| ATOM | 280 | OE1 | GLU | A | 38 | 14.322 | 40.088 | 19.606 | 1.00 | 0.00 | O |
| ATOM | 281 | OE2 | GLU | A | 38 | 14.445 | 40.167 | 17.431 | 1.00 | 0.00 | O |
| ATOM | 282 | C | GLU | A | 38 | 17.863 | 37.626 | 20.736 | 1.00 | 0.00 | C |
| ATOM | 283 | O | GLU | A | 38 | 17.753 | 37.950 | 21.895 | 1.00 | 0.00 | O |
| ATOM | 284 | N | ALA | A | 39 | 18.721 | 36.734 | 20.281 | 1.00 | 0.00 | N |
| ATOM | 285 | H | ALA | A | 39 | 18.738 | 36.590 | 19.292 | 1.00 | 0.00 | H |
| ATOM | 286 | CA | ALA | A | 39 | 19.629 | 35.952 | 21.060 | 1.00 | 0.00 | C |
| ATOM | 287 | HA | ALA | A | 39 | 19.273 | 35.907 | 21.993 | 1.00 | 0.00 | H |
| ATOM | 288 | CB | ALA | A | 39 | 19.644 | 34.637 | 20.362 | 1.00 | 0.00 | C |
| ATOM | 289 | HB1 | ALA | A | 39 | 20.261 | 34.013 | 20.841 | 1.00 | 0.00 | H |
| ATOM | 290 | HB2 | ALA | A | 39 | 18.721 | 34.253 | 20.355 | 1.00 | 0.00 | H |
| ATOM | 291 | HB3 | ALA | A | 39 | 19.960 | 34.762 | 19.422 | 1.00 | 0.00 | H |
| ATOM | 292 | C | ALA | A | 39 | 21.066 | 36.554 | 21.111 | 1.00 | 0.00 | C |
| ATOM | 293 | O | ALA | A | 39 | 21.880 | 36.108 | 21.951 | 1.00 | 0.00 | O |
| ATOM | 294 | N | ARG | A | 40 | 21.403 | 37.530 | 20.264 | 1.00 | 0.00 | N |
| ATOM | 295 | H | ARG | A | 40 | 20.814 | 37.672 | 19.468 | 1.00 | 0.00 | H |
| ATOM | 296 | CA | ARG | A | 40 | 22.559 | 38.404 | 20.406 | 1.00 | 0.00 | C |
| ATOM | 297 | HA | ARG | A | 40 | 23.297 | 37.738 | 20.511 | 1.00 | 0.00 | H |
| ATOM | 298 | CB | ARG | A | 40 | 22.810 | 39.343 | 19.189 | 1.00 | 0.00 | C |
| ATOM | 299 | HB1 | ARG | A | 40 | 22.200 | 40.133 | 19.254 | 1.00 | 0.00 | H |
| ATOM | 300 | HB2 | ARG | A | 40 | 22.619 | 38.843 | 18.344 | 1.00 | 0.00 | H |
| ATOM | 301 | CG | ARG | A | 40 | 24.263 | 39.847 | 19.143 | 1.00 | 0.00 | C |
| ATOM | 302 | HG1 | ARG | A | 40 | 24.880 | 39.060 | 19.134 | 1.00 | 0.00 | H |
| ATOM | 303 | HG2 | ARG | A | 40 | 24.441 | 40.401 | 19.956 | 1.00 | 0.00 | H |
| ATOM | 304 | CD | ARG | A | 40 | 24.515 | 40.703 | 17.883 | 1.00 | 0.00 | C |
| ATOM | 305 | HD1 | ARG | A | 40 | 25.239 | 41.366 | 18.074 | 1.00 | 0.00 | H |
| ATOM | 306 | HD2 | ARG | A | 40 | 23.675 | 41.188 | 17.641 | 1.00 | 0.00 | H |
| ATOM | 307 | NE | ARG | A | 40 | 24.935 | 39.901 | 16.697 | 1.00 | 0.00 | N |
| ATOM | 308 | HE | ARG | A | 40 | 24.912 | 38.902 | 16.743 | 1.00 | 0.00 | H |
| ATOM | 309 | CZ | ARG | A | 40 | 25.335 | 40.491 | 15.593 | 1.00 | 0.00 | C |
| ATOM | 310 | NH1 | ARG | A | 40 | 26.027 | 39.837 | 14.707 | 1.00 | 0.00 | N |
| ATOM | 311 | HH11 | ARG | A | 40 | 26.259 | 38.877 | 14.863 | 1.00 | 0.00 | H |
| ATOM | 312 | HH12 | ARG | A | 40 | 26.327 | 40.294 | 13.870 | 1.00 | 0.00 | H |
| ATOM | 313 | NH2 | ARG | A | 40 | 25.061 | 41.753 | 15.399 | 1.00 | 0.00 | N |
| ATOM | 314 | HH21 | ARG | A | 40 | 24.549 | 42.265 | 16.089 | 1.00 | 0.00 | H |
| ATOM | 315 | HH22 | ARG | A | 40 | 25.363 | 42.207 | 14.561 | 1.00 | 0.00 | H |
| ATOM | 316 | C | ARG | A | 40 | 22.435 | 39.268 | 21.619 | 1.00 | 0.00 | C |
| ATOM | 317 | O | ARG | A | 40 | 21.463 | 39.896 | 21.846 | 1.00 | 0.00 | O |
| ATOM | 318 | N | PHE | A | 41 | 23.501 | 39.395 | 22.426 | 1.00 | 0.00 | N |
| ATOM | 319 | H | PHE | A | 41 | 24.384 | 39.067 | 22.091 | 1.00 | 0.00 | H |
| ATOM | 320 | CA | PHE | A | 41 | 23.450 | 39.980 | 23.758 | 1.00 | 0.00 | C |
| ATOM | 321 | HA | PHE | A | 41 | 22.641 | 40.566 | 23.810 | 1.00 | 0.00 | H |

| | | | | | | | | | | | |
|---|---|---|---|---|---|---|---|---|---|---|---|
| ATOM | 322 | CB | PHE | A | 41 | 23.256 | 38.919 | 24.775 | 1.00 | 0.00 | C |
| ATOM | 323 | HB1 | PHE | A | 41 | 23.971 | 38.235 | 24.633 | 1.00 | 0.00 | H |
| ATOM | 324 | HB2 | PHE | A | 41 | 22.361 | 38.506 | 24.608 | 1.00 | 0.00 | H |
| ATOM | 325 | CG | PHE | A | 41 | 23.303 | 39.327 | 26.206 | 1.00 | 0.00 | C |
| ATOM | 326 | CD1 | PHE | A | 41 | 24.505 | 39.202 | 26.925 | 1.00 | 0.00 | C |
| ATOM | 327 | HD1 | PHE | A | 41 | 25.351 | 38.962 | 26.448 | 1.00 | 0.00 | H |
| ATOM | 328 | CE1 | PHE | A | 41 | 24.516 | 39.410 | 28.282 | 1.00 | 0.00 | C |
| ATOM | 329 | HE1 | PHE | A | 41 | 25.329 | 39.159 | 28.807 | 1.00 | 0.00 | H |
| ATOM | 330 | CZ | PHE | A | 41 | 23.435 | 39.955 | 28.937 | 1.00 | 0.00 | C |
| ATOM | 331 | HZ | PHE | A | 41 | 23.512 | 40.240 | 29.892 | 1.00 | 0.00 | H |
| ATOM | 332 | CE2 | PHE | A | 41 | 22.244 | 40.102 | 28.257 | 1.00 | 0.00 | C |
| ATOM | 333 | HE2 | PHE | A | 41 | 21.453 | 40.497 | 28.724 | 1.00 | 0.00 | H |
| ATOM | 334 | CD2 | PHE | A | 41 | 22.132 | 39.699 | 26.905 | 1.00 | 0.00 | C |
| ATOM | 335 | HD2 | PHE | A | 41 | 21.241 | 39.678 | 26.452 | 1.00 | 0.00 | H |
| ATOM | 336 | C | PHE | A | 41 | 24.781 | 40.766 | 23.996 | 1.00 | 0.00 | C |
| ATOM | 337 | O | PHE | A | 41 | 25.832 | 40.126 | 23.758 | 1.00 | 0.00 | O |
| ATOM | 338 | N | LEU | A | 42 | 24.784 | 42.024 | 24.383 | 1.00 | 0.00 | N |
| ATOM | 339 | H | LEU | A | 42 | 23.904 | 42.374 | 24.704 | 1.00 | 0.00 | H |
| ATOM | 340 | CA | LEU | A | 42 | 25.908 | 43.002 | 24.412 | 1.00 | 0.00 | C |
| ATOM | 341 | HA | LEU | A | 42 | 26.782 | 42.544 | 24.249 | 1.00 | 0.00 | H |
| ATOM | 342 | CB | LEU | A | 42 | 25.661 | 44.059 | 23.284 | 1.00 | 0.00 | C |
| ATOM | 343 | HB1 | LEU | A | 42 | 26.401 | 44.731 | 23.312 | 1.00 | 0.00 | H |
| ATOM | 344 | HB2 | LEU | A | 42 | 24.789 | 44.515 | 23.462 | 1.00 | 0.00 | H |
| ATOM | 345 | CG | LEU | A | 42 | 25.620 | 43.359 | 21.834 | 1.00 | 0.00 | C |
| ATOM | 346 | HG | LEU | A | 42 | 24.866 | 42.707 | 21.762 | 1.00 | 0.00 | H |
| ATOM | 347 | CD1 | LEU | A | 42 | 25.322 | 44.424 | 20.849 | 1.00 | 0.00 | C |
| ATOM | 348 | HD11 | LEU | A | 42 | 25.288 | 44.029 | 19.931 | 1.00 | 0.00 | H |
| ATOM | 349 | HD12 | LEU | A | 42 | 24.439 | 44.841 | 21.066 | 1.00 | 0.00 | H |
| ATOM | 350 | HD13 | LEU | A | 42 | 26.038 | 45.121 | 20.885 | 1.00 | 0.00 | H |
| ATOM | 351 | CD2 | LEU | A | 42 | 26.860 | 42.444 | 21.463 | 1.00 | 0.00 | C |
| ATOM | 352 | HD21 | LEU | A | 42 | 26.731 | 42.063 | 20.548 | 1.00 | 0.00 | H |
| ATOM | 353 | HD22 | LEU | A | 42 | 27.696 | 42.992 | 21.481 | 1.00 | 0.00 | H |
| ATOM | 354 | HD23 | LEU | A | 42 | 26.935 | 41.700 | 22.126 | 1.00 | 0.00 | H |
| ATOM | 355 | C | LEU | A | 42 | 26.043 | 43.570 | 25.800 | 1.00 | 0.00 | C |
| ATOM | 356 | O | LEU | A | 42 | 25.077 | 43.675 | 26.495 | 1.00 | 0.00 | O |
| ATOM | 357 | N | PRO | A | 43 | 27.191 | 44.165 | 26.275 | 1.00 | 0.00 | N |
| ATOM | 358 | CD | PRO | A | 43 | 28.451 | 44.041 | 25.657 | 1.00 | 0.00 | C |
| ATOM | 359 | HD1 | PRO | A | 43 | 28.460 | 44.623 | 24.844 | 1.00 | 0.00 | H |
| ATOM | 360 | HD2 | PRO | A | 43 | 28.574 | 43.087 | 25.385 | 1.00 | 0.00 | H |
| ATOM | 361 | CG | PRO | A | 43 | 29.517 | 44.463 | 26.630 | 1.00 | 0.00 | C |
| ATOM | 362 | HG1 | PRO | A | 43 | 30.251 | 44.950 | 26.156 | 1.00 | 0.00 | H |
| ATOM | 363 | HG2 | PRO | A | 43 | 29.896 | 43.667 | 27.101 | 1.00 | 0.00 | H |
| ATOM | 364 | CB | PRO | A | 43 | 28.700 | 45.408 | 27.607 | 1.00 | 0.00 | C |
| ATOM | 365 | HB1 | PRO | A | 43 | 28.726 | 46.360 | 27.303 | 1.00 | 0.00 | H |
| ATOM | 366 | HB2 | PRO | A | 43 | 29.041 | 45.348 | 28.545 | 1.00 | 0.00 | H |
| ATOM | 367 | CA | PRO | A | 43 | 27.279 | 44.885 | 27.541 | 1.00 | 0.00 | C |
| ATOM | 368 | HA | PRO | A | 43 | 27.049 | 44.162 | 28.192 | 1.00 | 0.00 | H |
| ATOM | 369 | C | PRO | A | 43 | 26.257 | 46.059 | 27.861 | 1.00 | 0.00 | C |
| ATOM | 370 | O | PRO | A | 43 | 26.018 | 46.324 | 29.061 | 1.00 | 0.00 | O |
| ATOM | 371 | N | LYS | A | 44 | 25.621 | 46.695 | 26.868 | 1.00 | 0.00 | N |
| ATOM | 372 | H | LYS | A | 44 | 25.965 | 46.575 | 25.937 | 1.00 | 0.00 | H |
| ATOM | 373 | CA | LYS | A | 44 | 24.457 | 47.552 | 27.063 | 1.00 | 0.00 | C |
| ATOM | 374 | HA | LYS | A | 44 | 24.896 | 48.254 | 27.623 | 1.00 | 0.00 | H |
| ATOM | 375 | CB | LYS | A | 44 | 23.901 | 48.167 | 25.793 | 1.00 | 0.00 | C |

| | | | | | | | | | | | |
|---|---|---|---|---|---|---|---|---|---|---|---|
| ATOM | 376 | HB1 | LYS | A | 44 | 23.792 | 47.407 | 25.152 | 1.00 | 0.00 | H |
| ATOM | 377 | HB2 | LYS | A | 44 | 24.613 | 48.785 | 25.459 | 1.00 | 0.00 | H |
| ATOM | 378 | CG | LYS | A | 44 | 22.598 | 48.958 | 25.731 | 1.00 | 0.00 | C |
| ATOM | 379 | HG1 | LYS | A | 44 | 21.837 | 48.323 | 25.861 | 1.00 | 0.00 | H |
| ATOM | 380 | HG2 | LYS | A | 44 | 22.529 | 49.381 | 24.828 | 1.00 | 0.00 | H |
| ATOM | 381 | CD | LYS | A | 44 | 22.522 | 50.063 | 26.812 | 1.00 | 0.00 | C |
| ATOM | 382 | HD1 | LYS | A | 44 | 23.448 | 50.410 | 26.962 | 1.00 | 0.00 | H |
| ATOM | 383 | HD2 | LYS | A | 44 | 22.182 | 49.645 | 27.655 | 1.00 | 0.00 | H |
| ATOM | 384 | CE | LYS | A | 44 | 21.578 | 51.273 | 26.424 | 1.00 | 0.00 | C |
| ATOM | 385 | HE1 | LYS | A | 44 | 20.624 | 50.986 | 26.511 | 1.00 | 0.00 | H |
| ATOM | 386 | HE2 | LYS | A | 44 | 21.761 | 51.542 | 25.478 | 1.00 | 0.00 | H |
| ATOM | 387 | NZ | LYS | A | 44 | 21.736 | 52.509 | 27.263 | 1.00 | 0.00 | N |
| ATOM | 388 | HZ1 | LYS | A | 44 | 21.101 | 53.212 | 26.943 | 1.00 | 0.00 | H |
| ATOM | 389 | HZ2 | LYS | A | 44 | 21.538 | 52.291 | 28.219 | 1.00 | 0.00 | H |
| ATOM | 390 | HZ3 | LYS | A | 44 | 22.674 | 52.846 | 27.186 | 1.00 | 0.00 | H |
| ATOM | 391 | C | LYS | A | 44 | 23.291 | 46.902 | 27.816 | 1.00 | 0.00 | C |
| ATOM | 392 | O | LYS | A | 44 | 22.555 | 47.493 | 28.554 | 1.00 | 0.00 | O |
| ATOM | 393 | N | ASP | A | 45 | 23.124 | 45.583 | 27.505 | 1.00 | 0.00 | N |
| ATOM | 394 | H | ASP | A | 45 | 23.792 | 45.150 | 26.900 | 1.00 | 0.00 | H |
| ATOM | 395 | CA | ASP | A | 45 | 22.011 | 44.766 | 28.014 | 1.00 | 0.00 | C |
| ATOM | 396 | HA | ASP | A | 45 | 21.234 | 45.389 | 27.921 | 1.00 | 0.00 | H |
| ATOM | 397 | CB | ASP | A | 45 | 21.844 | 43.530 | 27.121 | 1.00 | 0.00 | C |
| ATOM | 398 | HB1 | ASP | A | 45 | 21.074 | 42.998 | 27.473 | 1.00 | 0.00 | H |
| ATOM | 399 | HB2 | ASP | A | 45 | 22.682 | 42.988 | 27.186 | 1.00 | 0.00 | H |
| ATOM | 400 | CG | ASP | A | 45 | 21.581 | 43.815 | 25.637 | 1.00 | 0.00 | C |
| ATOM | 401 | OD1 | ASP | A | 45 | 21.965 | 42.939 | 24.803 | 1.00 | 0.00 | O |
| ATOM | 402 | OD2 | ASP | A | 45 | 20.988 | 44.926 | 25.351 | 1.00 | 0.00 | O |
| ATOM | 403 | C | ASP | A | 45 | 21.975 | 44.455 | 29.525 | 1.00 | 0.00 | C |
| ATOM | 404 | O | ASP | A | 45 | 20.989 | 43.967 | 30.085 | 1.00 | 0.00 | O |
| ATOM | 405 | N | LEU | A | 46 | 23.105 | 44.668 | 30.171 | 1.00 | 0.00 | N |
| ATOM | 406 | H | LEU | A | 46 | 23.849 | 45.084 | 29.648 | 1.00 | 0.00 | H |
| ATOM | 407 | CA | LEU | A | 46 | 23.379 | 44.360 | 31.565 | 1.00 | 0.00 | C |
| ATOM | 408 | HA | LEU | A | 46 | 23.056 | 43.443 | 31.800 | 1.00 | 0.00 | H |
| ATOM | 409 | CB | LEU | A | 46 | 24.865 | 44.333 | 31.772 | 1.00 | 0.00 | C |
| ATOM | 410 | HB1 | LEU | A | 46 | 25.019 | 44.239 | 32.756 | 1.00 | 0.00 | H |
| ATOM | 411 | HB2 | LEU | A | 46 | 25.217 | 45.216 | 31.462 | 1.00 | 0.00 | H |
| ATOM | 412 | CG | LEU | A | 46 | 25.654 | 43.217 | 31.050 | 1.00 | 0.00 | C |
| ATOM | 413 | HG | LEU | A | 46 | 25.362 | 43.255 | 30.094 | 1.00 | 0.00 | H |
| ATOM | 414 | CD1 | LEU | A | 46 | 27.193 | 43.409 | 31.077 | 1.00 | 0.00 | C |
| ATOM | 415 | HD11 | LEU | A | 46 | 27.632 | 42.652 | 30.593 | 1.00 | 0.00 | H |
| ATOM | 416 | HD12 | LEU | A | 46 | 27.429 | 44.272 | 30.630 | 1.00 | 0.00 | H |
| ATOM | 417 | HD13 | LEU | A | 46 | 27.510 | 43.427 | 32.025 | 1.00 | 0.00 | H |
| ATOM | 418 | CD2 | LEU | A | 46 | 25.325 | 41.854 | 31.679 | 1.00 | 0.00 | C |
| ATOM | 419 | HD21 | LEU | A | 46 | 25.838 | 41.137 | 31.208 | 1.00 | 0.00 | H |
| ATOM | 420 | HD22 | LEU | A | 46 | 25.578 | 41.864 | 32.647 | 1.00 | 0.00 | H |
| ATOM | 421 | HD23 | LEU | A | 46 | 24.345 | 41.675 | 31.592 | 1.00 | 0.00 | H |
| ATOM | 422 | C | LEU | A | 46 | 22.608 | 45.331 | 32.431 | 1.00 | 0.00 | C |
| ATOM | 423 | O | LEU | A | 46 | 22.379 | 46.489 | 32.032 | 1.00 | 0.00 | O |
| ATOM | 424 | N | ASP | A | 47 | 22.312 | 44.954 | 33.696 | 1.00 | 0.00 | N |
| ATOM | 425 | H | ASP | A | 47 | 22.564 | 44.025 | 33.967 | 1.00 | 0.00 | H |
| ATOM | 426 | CA | ASP | A | 47 | 21.662 | 45.780 | 34.692 | 1.00 | 0.00 | C |
| ATOM | 427 | HA | ASP | A | 47 | 21.294 | 46.642 | 34.343 | 1.00 | 0.00 | H |
| ATOM | 428 | CB | ASP | A | 47 | 20.487 | 44.909 | 35.187 | 1.00 | 0.00 | C |
| ATOM | 429 | HB1 | ASP | A | 47 | 20.850 | 44.017 | 35.455 | 1.00 | 0.00 | H |

| | | | | | | | | | | | |
|---|---|---|---|---|---|---|---|---|---|---|---|
| ATOM | 430 | HB2 | ASP | A | 47 | 19.843 | 44.793 | 34.431 | 1.00 | 0.00 | H |
| ATOM | 431 | CG | ASP | A | 47 | 19.719 | 45.501 | 36.390 | 1.00 | 0.00 | C |
| ATOM | 432 | OD1 | ASP | A | 47 | 18.498 | 45.662 | 36.382 | 1.00 | 0.00 | O |
| ATOM | 433 | OD2 | ASP | A | 47 | 20.321 | 45.818 | 37.376 | 1.00 | 0.00 | O |
| ATOM | 434 | C | ASP | A | 47 | 22.742 | 46.178 | 35.719 | 1.00 | 0.00 | C |
| ATOM | 435 | O | ASP | A | 47 | 23.458 | 45.285 | 36.112 | 1.00 | 0.00 | O |
| ATOM | 436 | N | PRO | A | 48 | 22.817 | 47.403 | 36.198 | 1.00 | 0.00 | N |
| ATOM | 437 | CD | PRO | A | 48 | 21.992 | 48.513 | 35.751 | 1.00 | 0.00 | C |
| ATOM | 438 | HD1 | PRO | A | 48 | 21.125 | 48.526 | 36.250 | 1.00 | 0.00 | H |
| ATOM | 439 | HD2 | PRO | A | 48 | 21.811 | 48.440 | 34.770 | 1.00 | 0.00 | H |
| ATOM | 440 | CG | PRO | A | 48 | 22.846 | 49.720 | 36.077 | 1.00 | 0.00 | C |
| ATOM | 441 | HG1 | PRO | A | 48 | 22.263 | 50.506 | 36.281 | 1.00 | 0.00 | H |
| ATOM | 442 | HG2 | PRO | A | 48 | 23.440 | 49.938 | 35.302 | 1.00 | 0.00 | H |
| ATOM | 443 | CB | PRO | A | 48 | 23.684 | 49.409 | 37.266 | 1.00 | 0.00 | C |
| ATOM | 444 | HB1 | PRO | A | 48 | 23.233 | 49.728 | 38.100 | 1.00 | 0.00 | H |
| ATOM | 445 | HB2 | PRO | A | 48 | 24.579 | 49.848 | 37.182 | 1.00 | 0.00 | H |
| ATOM | 446 | CA | PRO | A | 48 | 23.801 | 47.867 | 37.237 | 1.00 | 0.00 | C |
| ATOM | 447 | HA | PRO | A | 48 | 24.686 | 47.513 | 36.935 | 1.00 | 0.00 | H |
| ATOM | 448 | C | PRO | A | 48 | 23.734 | 47.256 | 38.646 | 1.00 | 0.00 | C |
| ATOM | 449 | O | PRO | A | 48 | 24.660 | 47.442 | 39.418 | 1.00 | 0.00 | O |
| ATOM | 450 | N | SER | A | 49 | 22.634 | 46.623 | 39.053 | 1.00 | 0.00 | N |
| ATOM | 451 | H | SER | A | 49 | 21.924 | 46.448 | 38.371 | 1.00 | 0.00 | H |
| ATOM | 452 | CA | SER | A | 49 | 22.379 | 46.156 | 40.445 | 1.00 | 0.00 | C |
| ATOM | 453 | HA | SER | A | 49 | 23.322 | 45.958 | 40.713 | 1.00 | 0.00 | H |
| ATOM | 454 | CB | SER | A | 49 | 21.786 | 47.275 | 41.337 | 1.00 | 0.00 | C |
| ATOM | 455 | HB1 | SER | A | 49 | 22.438 | 48.023 | 41.456 | 1.00 | 0.00 | H |
| ATOM | 456 | HB2 | SER | A | 49 | 21.521 | 46.918 | 42.233 | 1.00 | 0.00 | H |
| ATOM | 457 | OG | SER | A | 49 | 20.645 | 47.840 | 40.793 | 1.00 | 0.00 | O |
| ATOM | 458 | HG | SER | A | 49 | 20.303 | 48.554 | 41.404 | 1.00 | 0.00 | H |
| ATOM | 459 | C | SER | A | 49 | 21.639 | 44.796 | 40.638 | 1.00 | 0.00 | C |
| ATOM | 460 | O | SER | A | 49 | 21.163 | 44.553 | 41.758 | 1.00 | 0.00 | O |
| ATOM | 461 | N | LEU | A | 50 | 21.529 | 43.954 | 39.645 | 1.00 | 0.00 | N |
| ATOM | 462 | H | LEU | A | 50 | 21.846 | 44.325 | 38.772 | 1.00 | 0.00 | H |
| ATOM | 463 | CA | LEU | A | 50 | 21.045 | 42.602 | 39.552 | 1.00 | 0.00 | C |
| ATOM | 464 | HA | LEU | A | 50 | 20.155 | 42.605 | 40.008 | 1.00 | 0.00 | H |
| ATOM | 465 | CB | LEU | A | 50 | 20.872 | 42.227 | 38.046 | 1.00 | 0.00 | C |
| ATOM | 466 | HB1 | LEU | A | 50 | 21.729 | 42.417 | 37.568 | 1.00 | 0.00 | H |
| ATOM | 467 | HB2 | LEU | A | 50 | 20.142 | 42.789 | 37.658 | 1.00 | 0.00 | H |
| ATOM | 468 | CG | LEU | A | 50 | 20.520 | 40.793 | 37.759 | 1.00 | 0.00 | C |
| ATOM | 469 | HG | LEU | A | 50 | 21.177 | 40.176 | 38.193 | 1.00 | 0.00 | H |
| ATOM | 470 | CD1 | LEU | A | 50 | 19.071 | 40.662 | 38.276 | 1.00 | 0.00 | C |
| ATOM | 471 | HD11 | LEU | A | 50 | 18.747 | 39.728 | 38.126 | 1.00 | 0.00 | H |
| ATOM | 472 | HD12 | LEU | A | 50 | 19.046 | 40.870 | 39.254 | 1.00 | 0.00 | H |
| ATOM | 473 | HD13 | LEU | A | 50 | 18.482 | 41.303 | 37.783 | 1.00 | 0.00 | H |
| ATOM | 474 | CD2 | LEU | A | 50 | 20.514 | 40.526 | 36.227 | 1.00 | 0.00 | C |
| ATOM | 475 | HD21 | LEU | A | 50 | 20.278 | 39.569 | 36.056 | 1.00 | 0.00 | H |
| ATOM | 476 | HD22 | LEU | A | 50 | 19.839 | 41.119 | 35.788 | 1.00 | 0.00 | H |
| ATOM | 477 | HD23 | LEU | A | 50 | 21.421 | 40.718 | 35.852 | 1.00 | 0.00 | H |
| ATOM | 478 | C | LEU | A | 50 | 22.085 | 41.621 | 40.272 | 1.00 | 0.00 | C |
| ATOM | 479 | O | LEU | A | 50 | 21.732 | 40.920 | 41.280 | 1.00 | 0.00 | O |
| ATOM | 480 | N | CYS | A | 51 | 23.366 | 41.690 | 39.913 | 1.00 | 0.00 | N |
| ATOM | 481 | H | CYS | A | 51 | 23.582 | 42.316 | 39.164 | 1.00 | 0.00 | H |
| ATOM | 482 | CA | CYS | A | 51 | 24.531 | 40.932 | 40.506 | 1.00 | 0.00 | C |
| ATOM | 483 | HA | CYS | A | 51 | 24.194 | 40.216 | 41.117 | 1.00 | 0.00 | H |

| | | | | | | | | | | |
|---|---|---|---|---|---|---|---|---|---|---|
| ATOM | 484 | CB | CYS | A | 51 | 25.316 | 40.252 | 39.372 | 1.00 | 0.00 | C |
| ATOM | 485 | HB1 | CYS | A | 51 | 26.020 | 39.740 | 39.864 | 1.00 | 0.00 | H |
| ATOM | 486 | HB2 | CYS | A | 51 | 25.737 | 41.025 | 38.898 | 1.00 | 0.00 | H |
| ATOM | 487 | SG | CYS | A | 51 | 24.368 | 39.169 | 38.210 | 1.00 | 0.00 | S |
| ATOM | 488 | C | CYS | A | 51 | 25.348 | 41.783 | 41.440 | 1.00 | 0.00 | C |
| ATOM | 489 | O | CYS | A | 51 | 25.347 | 43.024 | 41.402 | 1.00 | 0.00 | O |
| ATOM | 490 | N | THR | A | 52 | 26.123 | 41.153 | 42.369 | 1.00 | 0.00 | N |
| ATOM | 491 | H | THR | A | 52 | 26.082 | 40.159 | 42.475 | 1.00 | 0.00 | H |
| ATOM | 492 | CA | THR | A | 52 | 27.037 | 41.973 | 43.229 | 1.00 | 0.00 | C |
| ATOM | 493 | HA | THR | A | 52 | 26.578 | 42.860 | 43.277 | 1.00 | 0.00 | H |
| ATOM | 494 | CB | THR | A | 52 | 27.230 | 41.346 | 44.657 | 1.00 | 0.00 | C |
| ATOM | 495 | HB | THR | A | 52 | 28.009 | 41.785 | 45.105 | 1.00 | 0.00 | H |
| ATOM | 496 | CG2 | THR | A | 52 | 25.844 | 41.406 | 45.436 | 1.00 | 0.00 | C |
| ATOM | 497 | HG21 | THR | A | 52 | 25.956 | 41.009 | 46.347 | 1.00 | 0.00 | H |
| ATOM | 498 | HG22 | THR | A | 52 | 25.550 | 42.358 | 45.521 | 1.00 | 0.00 | H |
| ATOM | 499 | HG23 | THR | A | 52 | 25.154 | 40.889 | 44.929 | 1.00 | 0.00 | H |
| ATOM | 500 | OG1 | THR | A | 52 | 27.714 | 40.042 | 44.676 | 1.00 | 0.00 | O |
| ATOM | 501 | HG1 | THR | A | 52 | 27.802 | 39.733 | 45.623 | 1.00 | 0.00 | H |
| ATOM | 502 | C | THR | A | 52 | 28.436 | 42.111 | 42.590 | 1.00 | 0.00 | C |
| ATOM | 503 | O | THR | A | 52 | 29.166 | 43.063 | 42.844 | 1.00 | 0.00 | O |
| ATOM | 504 | N | HIS | A | 53 | 28.752 | 41.089 | 41.742 | 1.00 | 0.00 | N |
| ATOM | 505 | H | HIS | A | 53 | 28.058 | 40.374 | 41.663 | 1.00 | 0.00 | H |
| ATOM | 506 | CA | HIS | A | 53 | 29.956 | 40.880 | 40.928 | 1.00 | 0.00 | C |
| ATOM | 507 | HA | HIS | A | 53 | 30.323 | 41.793 | 40.750 | 1.00 | 0.00 | H |
| ATOM | 508 | CB | HIS | A | 53 | 30.966 | 39.870 | 41.737 | 1.00 | 0.00 | C |
| ATOM | 509 | HB1 | HIS | A | 53 | 31.841 | 39.806 | 41.257 | 1.00 | 0.00 | H |
| ATOM | 510 | HB2 | HIS | A | 53 | 30.556 | 38.960 | 41.805 | 1.00 | 0.00 | H |
| ATOM | 511 | CG | HIS | A | 53 | 31.317 | 40.238 | 43.128 | 1.00 | 0.00 | C |
| ATOM | 512 | ND1 | HIS | A | 53 | 30.441 | 40.151 | 44.150 | 1.00 | 0.00 | N |
| ATOM | 513 | HD1 | HIS | A | 53 | 29.463 | 39.948 | 44.113 | 1.00 | 0.00 | H |
| ATOM | 514 | CE1 | HIS | A | 53 | 31.203 | 40.404 | 45.236 | 1.00 | 0.00 | C |
| ATOM | 515 | HE1 | HIS | A | 53 | 30.903 | 40.279 | 46.182 | 1.00 | 0.00 | H |
| ATOM | 516 | NE2 | HIS | A | 53 | 32.392 | 40.837 | 44.883 | 1.00 | 0.00 | N |
| ATOM | 517 | CD2 | HIS | A | 53 | 32.584 | 40.630 | 43.540 | 1.00 | 0.00 | C |
| ATOM | 518 | HD2 | HIS | A | 53 | 33.419 | 40.734 | 42.999 | 1.00 | 0.00 | H |
| ATOM | 519 | C | HIS | A | 53 | 29.600 | 40.314 | 39.571 | 1.00 | 0.00 | C |
| ATOM | 520 | O | HIS | A | 53 | 28.885 | 39.356 | 39.458 | 1.00 | 0.00 | O |
| ATOM | 521 | N | LEU | A | 54 | 30.111 | 40.945 | 38.461 | 1.00 | 0.00 | N |
| ATOM | 522 | H | LEU | A | 54 | 30.583 | 41.818 | 38.587 | 1.00 | 0.00 | H |
| ATOM | 523 | CA | LEU | A | 54 | 30.003 | 40.418 | 37.138 | 1.00 | 0.00 | C |
| ATOM | 524 | HA | LEU | A | 54 | 29.317 | 39.690 | 37.138 | 1.00 | 0.00 | H |
| ATOM | 525 | CB | LEU | A | 54 | 29.616 | 41.590 | 36.162 | 1.00 | 0.00 | C |
| ATOM | 526 | HB1 | LEU | A | 54 | 29.725 | 41.263 | 35.223 | 1.00 | 0.00 | H |
| ATOM | 527 | HB2 | LEU | A | 54 | 30.243 | 42.352 | 36.325 | 1.00 | 0.00 | H |
| ATOM | 528 | CG | LEU | A | 54 | 28.165 | 42.133 | 36.307 | 1.00 | 0.00 | C |
| ATOM | 529 | HG | LEU | A | 54 | 28.081 | 42.652 | 37.158 | 1.00 | 0.00 | H |
| ATOM | 530 | CD1 | LEU | A | 54 | 27.833 | 42.999 | 35.122 | 1.00 | 0.00 | C |
| ATOM | 531 | HD11 | LEU | A | 54 | 26.901 | 43.348 | 35.215 | 1.00 | 0.00 | H |
| ATOM | 532 | HD12 | LEU | A | 54 | 28.473 | 43.766 | 35.080 | 1.00 | 0.00 | H |
| ATOM | 533 | HD13 | LEU | A | 54 | 27.905 | 42.459 | 34.284 | 1.00 | 0.00 | H |
| ATOM | 534 | CD2 | LEU | A | 54 | 27.168 | 41.055 | 36.399 | 1.00 | 0.00 | C |
| ATOM | 535 | HD21 | LEU | A | 54 | 26.255 | 41.452 | 36.491 | 1.00 | 0.00 | H |
| ATOM | 536 | HD22 | LEU | A | 54 | 27.206 | 40.494 | 35.572 | 1.00 | 0.00 | H |
| ATOM | 537 | HD23 | LEU | A | 54 | 27.365 | 40.486 | 37.197 | 1.00 | 0.00 | H |

| | | | | | | | | | | | |
|---|---|---|---|---|---|---|---|---|---|---|---|
| ATOM | 538 | C | LEU | A | 54 | 31.362 | 39.866 | 36.791 | 1.00 | 0.00 | C |
| ATOM | 539 | O | LEU | A | 54 | 32.360 | 40.458 | 37.191 | 1.00 | 0.00 | O |
| ATOM | 540 | N | ILE | A | 55 | 31.443 | 38.799 | 36.060 | 1.00 | 0.00 | N |
| ATOM | 541 | H | ILE | A | 55 | 30.633 | 38.241 | 35.879 | 1.00 | 0.00 | H |
| ATOM | 542 | CA | ILE | A | 55 | 32.771 | 38.420 | 35.494 | 1.00 | 0.00 | C |
| ATOM | 543 | HA | ILE | A | 55 | 33.347 | 39.152 | 35.857 | 1.00 | 0.00 | H |
| ATOM | 544 | CB | ILE | A | 55 | 33.268 | 37.041 | 35.988 | 1.00 | 0.00 | C |
| ATOM | 545 | HB | ILE | A | 55 | 32.706 | 36.322 | 35.578 | 1.00 | 0.00 | H |
| ATOM | 546 | CG2 | ILE | A | 55 | 34.728 | 36.812 | 35.553 | 1.00 | 0.00 | C |
| ATOM | 547 | HG21 | ILE | A | 55 | 35.036 | 35.918 | 35.878 | 1.00 | 0.00 | H |
| ATOM | 548 | HG22 | ILE | A | 55 | 34.787 | 36.842 | 34.555 | 1.00 | 0.00 | H |
| ATOM | 549 | HG23 | ILE | A | 55 | 35.308 | 37.527 | 35.943 | 1.00 | 0.00 | H |
| ATOM | 550 | CG1 | ILE | A | 55 | 33.111 | 36.923 | 37.505 | 1.00 | 0.00 | C |
| ATOM | 551 | HG11 | ILE | A | 55 | 32.134 | 36.830 | 37.697 | 1.00 | 0.00 | H |
| ATOM | 552 | HG12 | ILE | A | 55 | 33.450 | 37.774 | 37.907 | 1.00 | 0.00 | H |
| ATOM | 553 | CD | ILE | A | 55 | 33.832 | 35.763 | 38.205 | 1.00 | 0.00 | C |
| ATOM | 554 | HD1 | ILE | A | 55 | 33.648 | 35.801 | 39.187 | 1.00 | 0.00 | H |
| ATOM | 555 | HD2 | ILE | A | 55 | 33.501 | 34.894 | 37.837 | 1.00 | 0.00 | H |
| ATOM | 556 | HD3 | ILE | A | 55 | 34.817 | 35.838 | 38.047 | 1.00 | 0.00 | H |
| ATOM | 557 | C | ILE | A | 55 | 32.801 | 38.402 | 33.929 | 1.00 | 0.00 | C |
| ATOM | 558 | O | ILE | A | 55 | 31.813 | 37.925 | 33.395 | 1.00 | 0.00 | O |
| ATOM | 559 | N | TYR | A | 56 | 33.752 | 38.971 | 33.234 | 1.00 | 0.00 | N |
| ATOM | 560 | H | TYR | A | 56 | 34.420 | 39.541 | 33.713 | 1.00 | 0.00 | H |
| ATOM | 561 | CA | TYR | A | 56 | 33.885 | 38.803 | 31.730 | 1.00 | 0.00 | C |
| ATOM | 562 | HA | TYR | A | 56 | 32.937 | 38.814 | 31.413 | 1.00 | 0.00 | H |
| ATOM | 563 | CB | TYR | A | 56 | 34.636 | 39.944 | 31.090 | 1.00 | 0.00 | C |
| ATOM | 564 | HB1 | TYR | A | 56 | 35.576 | 39.905 | 31.428 | 1.00 | 0.00 | H |
| ATOM | 565 | HB2 | TYR | A | 56 | 34.205 | 40.792 | 31.399 | 1.00 | 0.00 | H |
| ATOM | 566 | CG | TYR | A | 56 | 34.680 | 39.962 | 29.551 | 1.00 | 0.00 | C |
| ATOM | 567 | CD1 | TYR | A | 56 | 35.635 | 39.258 | 28.821 | 1.00 | 0.00 | C |
| ATOM | 568 | HD1 | TYR | A | 56 | 36.277 | 38.630 | 29.261 | 1.00 | 0.00 | H |
| ATOM | 569 | CE1 | TYR | A | 56 | 35.647 | 39.483 | 27.404 | 1.00 | 0.00 | C |
| ATOM | 570 | HE1 | TYR | A | 56 | 36.250 | 38.923 | 26.835 | 1.00 | 0.00 | H |
| ATOM | 571 | CZ | TYR | A | 56 | 34.828 | 40.480 | 26.766 | 1.00 | 0.00 | C |
| ATOM | 572 | OH | TYR | A | 56 | 34.917 | 40.718 | 25.495 | 1.00 | 0.00 | O |
| ATOM | 573 | HH | TYR | A | 56 | 34.271 | 41.439 | 25.245 | 1.00 | 0.00 | H |
| ATOM | 574 | CE2 | TYR | A | 56 | 33.956 | 41.220 | 27.576 | 1.00 | 0.00 | C |
| ATOM | 575 | HE2 | TYR | A | 56 | 33.461 | 42.000 | 27.194 | 1.00 | 0.00 | H |
| ATOM | 576 | CD2 | TYR | A | 56 | 33.777 | 40.869 | 28.918 | 1.00 | 0.00 | C |
| ATOM | 577 | HD2 | TYR | A | 56 | 33.014 | 41.255 | 29.436 | 1.00 | 0.00 | H |
| ATOM | 578 | C | TYR | A | 56 | 34.637 | 37.478 | 31.324 | 1.00 | 0.00 | C |
| ATOM | 579 | O | TYR | A | 56 | 35.728 | 37.235 | 31.862 | 1.00 | 0.00 | O |
| ATOM | 580 | N | ALA | A | 57 | 34.095 | 36.659 | 30.425 | 1.00 | 0.00 | N |
| ATOM | 581 | H | ALA | A | 57 | 33.202 | 36.905 | 30.049 | 1.00 | 0.00 | H |
| ATOM | 582 | CA | ALA | A | 57 | 34.717 | 35.424 | 29.951 | 1.00 | 0.00 | C |
| ATOM | 583 | HA | ALA | A | 57 | 35.555 | 35.264 | 30.472 | 1.00 | 0.00 | H |
| ATOM | 584 | CB | ALA | A | 57 | 33.745 | 34.264 | 30.210 | 1.00 | 0.00 | C |
| ATOM | 585 | HB1 | ALA | A | 57 | 34.154 | 33.409 | 29.892 | 1.00 | 0.00 | H |
| ATOM | 586 | HB2 | ALA | A | 57 | 33.557 | 34.199 | 31.190 | 1.00 | 0.00 | H |
| ATOM | 587 | HB3 | ALA | A | 57 | 32.891 | 34.429 | 29.717 | 1.00 | 0.00 | H |
| ATOM | 588 | C | ALA | A | 57 | 35.099 | 35.580 | 28.465 | 1.00 | 0.00 | C |
| ATOM | 589 | O | ALA | A | 57 | 34.208 | 35.672 | 27.623 | 1.00 | 0.00 | O |
| ATOM | 590 | N | PHE | A | 58 | 36.381 | 35.644 | 28.079 | 1.00 | 0.00 | N |
| ATOM | 591 | H | PHE | A | 58 | 36.525 | 35.808 | 27.103 | 1.00 | 0.00 | H |

| | | | | | | | | | | | |
|---|---|---|---|---|---|---|---|---|---|---|---|
| ATOM | 592 | CA | PHE | A | 58 | 37.614 | 35.508 | 28.883 | 1.00 | 0.00 | C |
| ATOM | 593 | HA | PHE | A | 58 | 37.353 | 35.841 | 29.789 | 1.00 | 0.00 | H |
| ATOM | 594 | CB | PHE | A | 58 | 38.083 | 34.039 | 28.884 | 1.00 | 0.00 | C |
| ATOM | 595 | HB1 | PHE | A | 58 | 38.807 | 33.955 | 29.569 | 1.00 | 0.00 | H |
| ATOM | 596 | HB2 | PHE | A | 58 | 38.457 | 33.842 | 27.978 | 1.00 | 0.00 | H |
| ATOM | 597 | CG | PHE | A | 58 | 37.070 | 32.950 | 29.184 | 1.00 | 0.00 | C |
| ATOM | 598 | CD1 | PHE | A | 58 | 36.970 | 32.404 | 30.477 | 1.00 | 0.00 | C |
| ATOM | 599 | HD1 | PHE | A | 58 | 37.499 | 32.799 | 31.228 | 1.00 | 0.00 | H |
| ATOM | 600 | CE1 | PHE | A | 58 | 36.127 | 31.302 | 30.715 | 1.00 | 0.00 | C |
| ATOM | 601 | HE1 | PHE | A | 58 | 36.050 | 30.923 | 31.637 | 1.00 | 0.00 | H |
| ATOM | 602 | CZ | PHE | A | 58 | 35.394 | 30.734 | 29.659 | 1.00 | 0.00 | C |
| ATOM | 603 | HZ | PHE | A | 58 | 34.818 | 29.933 | 29.823 | 1.00 | 0.00 | H |
| ATOM | 604 | CE2 | PHE | A | 58 | 35.472 | 31.293 | 28.374 | 1.00 | 0.00 | C |
| ATOM | 605 | HE2 | PHE | A | 58 | 34.930 | 30.908 | 27.628 | 1.00 | 0.00 | H |
| ATOM | 606 | CD2 | PHE | A | 58 | 36.312 | 32.394 | 28.136 | 1.00 | 0.00 | C |
| ATOM | 607 | HD2 | PHE | A | 58 | 36.372 | 32.785 | 27.217 | 1.00 | 0.00 | H |
| ATOM | 608 | C | PHE | A | 58 | 38.781 | 36.327 | 28.289 | 1.00 | 0.00 | C |
| ATOM | 609 | O | PHE | A | 58 | 38.781 | 36.581 | 27.082 | 1.00 | 0.00 | O |
| ATOM | 610 | N | ALA | A | 59 | 39.852 | 36.557 | 29.061 | 1.00 | 0.00 | N |
| ATOM | 611 | H | ALA | A | 59 | 39.723 | 36.666 | 30.047 | 1.00 | 0.00 | H |
| ATOM | 612 | CA | ALA | A | 59 | 41.209 | 36.654 | 28.509 | 1.00 | 0.00 | C |
| ATOM | 613 | HA | ALA | A | 59 | 41.107 | 36.952 | 27.560 | 1.00 | 0.00 | H |
| ATOM | 614 | CB | ALA | A | 59 | 41.973 | 37.703 | 29.315 | 1.00 | 0.00 | C |
| ATOM | 615 | HB1 | ALA | A | 59 | 42.903 | 37.787 | 28.956 | 1.00 | 0.00 | H |
| ATOM | 616 | HB2 | ALA | A | 59 | 41.507 | 38.585 | 29.242 | 1.00 | 0.00 | H |
| ATOM | 617 | HB3 | ALA | A | 59 | 42.010 | 37.424 | 30.275 | 1.00 | 0.00 | H |
| ATOM | 618 | C | ALA | A | 59 | 41.943 | 35.289 | 28.468 | 1.00 | 0.00 | C |
| ATOM | 619 | O | ALA | A | 59 | 41.648 | 34.398 | 29.267 | 1.00 | 0.00 | O |
| ATOM | 620 | N | GLY | A | 60 | 42.948 | 35.158 | 27.590 | 1.00 | 0.00 | N |
| ATOM | 621 | H | GLY | A | 60 | 43.050 | 35.869 | 26.894 | 1.00 | 0.00 | H |
| ATOM | 622 | CA | GLY | A | 60 | 43.914 | 34.044 | 27.574 | 1.00 | 0.00 | C |
| ATOM | 623 | HA1 | GLY | A | 60 | 43.989 | 33.694 | 26.640 | 1.00 | 0.00 | H |
| ATOM | 624 | HA2 | GLY | A | 60 | 43.581 | 33.318 | 28.176 | 1.00 | 0.00 | H |
| ATOM | 625 | C | GLY | A | 60 | 45.255 | 34.495 | 28.041 | 1.00 | 0.00 | C |
| ATOM | 626 | O | GLY | A | 60 | 45.389 | 35.304 | 28.950 | 1.00 | 0.00 | O |
| ATOM | 627 | N | MET | A | 61 | 46.332 | 33.969 | 27.348 | 1.00 | 0.00 | N |
| ATOM | 628 | H | MET | A | 61 | 46.160 | 33.316 | 26.610 | 1.00 | 0.00 | H |
| ATOM | 629 | CA | MET | A | 61 | 47.689 | 34.330 | 27.654 | 1.00 | 0.00 | C |
| ATOM | 630 | HA | MET | A | 61 | 47.662 | 35.254 | 28.037 | 1.00 | 0.00 | H |
| ATOM | 631 | CB | MET | A | 61 | 48.258 | 33.314 | 28.724 | 1.00 | 0.00 | C |
| ATOM | 632 | HB1 | MET | A | 61 | 47.760 | 33.467 | 29.578 | 1.00 | 0.00 | H |
| ATOM | 633 | HB2 | MET | A | 61 | 49.225 | 33.530 | 28.858 | 1.00 | 0.00 | H |
| ATOM | 634 | CG | MET | A | 61 | 48.182 | 31.790 | 28.436 | 1.00 | 0.00 | C |
| ATOM | 635 | HG1 | MET | A | 61 | 48.939 | 31.613 | 27.807 | 1.00 | 0.00 | H |
| ATOM | 636 | HG2 | MET | A | 61 | 47.314 | 31.666 | 27.956 | 1.00 | 0.00 | H |
| ATOM | 637 | SD | MET | A | 61 | 48.286 | 30.754 | 29.945 | 1.00 | 0.00 | S |
| ATOM | 638 | CE | MET | A | 61 | 49.943 | 31.209 | 30.546 | 1.00 | 0.00 | C |
| ATOM | 639 | HE1 | MET | A | 61 | 50.142 | 30.707 | 31.388 | 1.00 | 0.00 | H |
| ATOM | 640 | HE2 | MET | A | 61 | 50.624 | 30.978 | 29.851 | 1.00 | 0.00 | H |
| ATOM | 641 | HE3 | MET | A | 61 | 49.973 | 32.192 | 30.729 | 1.00 | 0.00 | H |
| ATOM | 642 | C | MET | A | 61 | 48.516 | 34.353 | 26.329 | 1.00 | 0.00 | C |
| ATOM | 643 | O | MET | A | 61 | 48.315 | 33.598 | 25.370 | 1.00 | 0.00 | O |
| ATOM | 644 | N | THR | A | 62 | 49.617 | 35.144 | 26.278 | 1.00 | 0.00 | N |
| ATOM | 645 | H | THR | A | 62 | 49.748 | 35.825 | 26.998 | 1.00 | 0.00 | H |

| | | | | | | | | | | | |
|---|---|---|---|---|---|---|---|---|---|---|---|
| ATOM | 646 | CA | THR | A | 62 | 50.643 | 35.046 | 25.198 | 1.00 | 0.00 | C |
| ATOM | 647 | HA | THR | A | 62 | 50.831 | 34.074 | 25.056 | 1.00 | 0.00 | H |
| ATOM | 648 | CB | THR | A | 62 | 50.113 | 35.686 | 23.925 | 1.00 | 0.00 | C |
| ATOM | 649 | HB | THR | A | 62 | 49.690 | 36.545 | 24.212 | 1.00 | 0.00 | H |
| ATOM | 650 | CG2 | THR | A | 62 | 51.208 | 35.785 | 22.891 | 1.00 | 0.00 | C |
| ATOM | 651 | HG21 | THR | A | 62 | 50.845 | 36.207 | 22.060 | 1.00 | 0.00 | H |
| ATOM | 652 | HG22 | THR | A | 62 | 51.956 | 36.344 | 23.250 | 1.00 | 0.00 | H |
| ATOM | 653 | HG23 | THR | A | 62 | 51.549 | 34.869 | 22.678 | 1.00 | 0.00 | H |
| ATOM | 654 | OG1 | THR | A | 62 | 49.129 | 34.926 | 23.252 | 1.00 | 0.00 | O |
| ATOM | 655 | HG1 | THR | A | 62 | 48.835 | 35.414 | 22.430 | 1.00 | 0.00 | H |
| ATOM | 656 | C | THR | A | 62 | 51.913 | 35.785 | 25.607 | 1.00 | 0.00 | C |
| ATOM | 657 | O | THR | A | 62 | 51.822 | 36.921 | 26.036 | 1.00 | 0.00 | O |
| ATOM | 658 | N | ASN | A | 63 | 53.073 | 35.212 | 25.287 | 1.00 | 0.00 | N |
| ATOM | 659 | H | ASN | A | 63 | 53.014 | 34.375 | 24.744 | 1.00 | 0.00 | H |
| ATOM | 660 | CA | ASN | A | 63 | 54.414 | 35.655 | 25.631 | 1.00 | 0.00 | C |
| ATOM | 661 | HA | ASN | A | 63 | 55.037 | 34.951 | 25.291 | 1.00 | 0.00 | H |
| ATOM | 662 | CB | ASN | A | 63 | 54.797 | 36.925 | 24.883 | 1.00 | 0.00 | C |
| ATOM | 663 | HB1 | ASN | A | 63 | 55.731 | 37.164 | 25.148 | 1.00 | 0.00 | H |
| ATOM | 664 | HB2 | ASN | A | 63 | 54.173 | 37.650 | 25.175 | 1.00 | 0.00 | H |
| ATOM | 665 | CG | ASN | A | 63 | 54.763 | 36.899 | 23.387 | 1.00 | 0.00 | C |
| ATOM | 666 | OD1 | ASN | A | 63 | 54.766 | 35.914 | 22.678 | 1.00 | 0.00 | O |
| ATOM | 667 | ND2 | ASN | A | 63 | 54.542 | 38.013 | 22.757 | 1.00 | 0.00 | N |
| ATOM | 668 | HD21 | ASN | A | 63 | 54.400 | 38.859 | 23.271 | 1.00 | 0.00 | H |
| ATOM | 669 | HD22 | ASN | A | 63 | 54.514 | 38.025 | 21.757 | 1.00 | 0.00 | H |
| ATOM | 670 | C | ASN | A | 63 | 54.572 | 35.732 | 27.160 | 1.00 | 0.00 | C |
| ATOM | 671 | O | ASN | A | 63 | 55.126 | 36.668 | 27.771 | 1.00 | 0.00 | O |
| ATOM | 672 | N | HIS | A | 64 | 54.039 | 34.668 | 27.832 | 1.00 | 0.00 | N |
| ATOM | 673 | H | HIS | A | 64 | 53.697 | 33.901 | 27.290 | 1.00 | 0.00 | H |
| ATOM | 674 | CA | HIS | A | 64 | 53.931 | 34.569 | 29.319 | 1.00 | 0.00 | C |
| ATOM | 675 | HA | HIS | A | 64 | 53.317 | 33.795 | 29.478 | 1.00 | 0.00 | H |
| ATOM | 676 | CB | HIS | A | 64 | 55.325 | 34.346 | 29.948 | 1.00 | 0.00 | C |
| ATOM | 677 | HB1 | HIS | A | 64 | 55.206 | 34.069 | 30.901 | 1.00 | 0.00 | H |
| ATOM | 678 | HB2 | HIS | A | 64 | 55.834 | 35.206 | 29.912 | 1.00 | 0.00 | H |
| ATOM | 679 | CG | HIS | A | 64 | 56.172 | 33.306 | 29.290 | 1.00 | 0.00 | C |
| ATOM | 680 | ND1 | HIS | A | 64 | 55.871 | 32.001 | 29.174 | 1.00 | 0.00 | N |
| ATOM | 681 | CE1 | HIS | A | 64 | 56.894 | 31.382 | 28.519 | 1.00 | 0.00 | C |
| ATOM | 682 | HE1 | HIS | A | 64 | 56.964 | 30.407 | 28.310 | 1.00 | 0.00 | H |
| ATOM | 683 | NE2 | HIS | A | 64 | 57.818 | 32.362 | 28.198 | 1.00 | 0.00 | N |
| ATOM | 684 | HE2 | HIS | A | 64 | 58.679 | 32.198 | 27.717 | 1.00 | 0.00 | H |
| ATOM | 685 | CD2 | HIS | A | 64 | 57.379 | 33.612 | 28.640 | 1.00 | 0.00 | C |
| ATOM | 686 | HD2 | HIS | A | 64 | 57.816 | 34.504 | 28.523 | 1.00 | 0.00 | H |
| ATOM | 687 | C | HIS | A | 64 | 53.253 | 35.732 | 29.964 | 1.00 | 0.00 | C |
| ATOM | 688 | O | HIS | A | 64 | 53.394 | 35.859 | 31.173 | 1.00 | 0.00 | O |
| ATOM | 689 | N | GLN | A | 65 | 52.572 | 36.569 | 29.266 | 1.00 | 0.00 | N |
| ATOM | 690 | H | GLN | A | 65 | 52.667 | 36.534 | 28.271 | 1.00 | 0.00 | H |
| ATOM | 691 | CA | GLN | A | 65 | 51.668 | 37.569 | 29.830 | 1.00 | 0.00 | C |
| ATOM | 692 | HA | GLN | A | 65 | 51.816 | 37.620 | 30.818 | 1.00 | 0.00 | H |
| ATOM | 693 | CB | GLN | A | 65 | 52.031 | 38.896 | 29.112 | 1.00 | 0.00 | C |
| ATOM | 694 | HB1 | GLN | A | 65 | 51.341 | 39.570 | 29.377 | 1.00 | 0.00 | H |
| ATOM | 695 | HB2 | GLN | A | 65 | 51.968 | 38.723 | 28.129 | 1.00 | 0.00 | H |
| ATOM | 696 | CG | GLN | A | 65 | 53.434 | 39.532 | 29.385 | 1.00 | 0.00 | C |
| ATOM | 697 | HG1 | GLN | A | 65 | 54.128 | 38.905 | 29.031 | 1.00 | 0.00 | H |
| ATOM | 698 | HG2 | GLN | A | 65 | 53.541 | 39.623 | 30.375 | 1.00 | 0.00 | H |
| ATOM | 699 | CD | GLN | A | 65 | 53.753 | 40.893 | 28.797 | 1.00 | 0.00 | C |

| | | | | | | | | | | | |
|---|---|---|---|---|---|---|---|---|---|---|---|
| ATOM | 700 | OE1 | GLN | A | 65 | 52.865 | 41.625 | 28.385 | 1.00 | 0.00 | O |
| ATOM | 701 | NE2 | GLN | A | 65 | 54.976 | 41.186 | 28.580 | 1.00 | 0.00 | N |
| ATOM | 702 | HE21 | GLN | A | 65 | 55.695 | 40.525 | 28.797 | 1.00 | 0.00 | H |
| ATOM | 703 | HE22 | GLN | A | 65 | 55.216 | 42.076 | 28.193 | 1.00 | 0.00 | H |
| ATOM | 704 | C | GLN | A | 65 | 50.178 | 37.166 | 29.628 | 1.00 | 0.00 | C |
| ATOM | 705 | O | GLN | A | 65 | 49.824 | 36.154 | 28.944 | 1.00 | 0.00 | O |
| ATOM | 706 | N | LEU | A | 66 | 49.267 | 37.973 | 30.161 | 1.00 | 0.00 | N |
| ATOM | 707 | H | LEU | A | 66 | 49.590 | 38.605 | 30.866 | 1.00 | 0.00 | H |
| ATOM | 708 | CA | LEU | A | 66 | 47.823 | 38.040 | 29.825 | 1.00 | 0.00 | C |
| ATOM | 709 | HA | LEU | A | 66 | 47.447 | 37.127 | 29.983 | 1.00 | 0.00 | H |
| ATOM | 710 | CB | LEU | A | 66 | 47.066 | 39.030 | 30.775 | 1.00 | 0.00 | C |
| ATOM | 711 | HB1 | LEU | A | 66 | 46.442 | 39.614 | 30.256 | 1.00 | 0.00 | H |
| ATOM | 712 | HB2 | LEU | A | 66 | 47.712 | 39.600 | 31.282 | 1.00 | 0.00 | H |
| ATOM | 713 | CG | LEU | A | 66 | 46.275 | 38.216 | 31.743 | 1.00 | 0.00 | C |
| ATOM | 714 | HG | LEU | A | 66 | 45.782 | 37.609 | 31.120 | 1.00 | 0.00 | H |
| ATOM | 715 | CD1 | LEU | A | 66 | 47.166 | 37.447 | 32.796 | 1.00 | 0.00 | C |
| ATOM | 716 | HD11 | LEU | A | 66 | 46.577 | 36.923 | 33.412 | 1.00 | 0.00 | H |
| ATOM | 717 | HD12 | LEU | A | 66 | 47.785 | 36.824 | 32.317 | 1.00 | 0.00 | H |
| ATOM | 718 | HD13 | LEU | A | 66 | 47.700 | 38.106 | 33.326 | 1.00 | 0.00 | H |
| ATOM | 719 | CD2 | LEU | A | 66 | 45.271 | 39.005 | 32.514 | 1.00 | 0.00 | C |
| ATOM | 720 | HD21 | LEU | A | 66 | 44.780 | 38.399 | 33.140 | 1.00 | 0.00 | H |
| ATOM | 721 | HD22 | LEU | A | 66 | 45.738 | 39.718 | 33.038 | 1.00 | 0.00 | H |
| ATOM | 722 | HD23 | LEU | A | 66 | 44.622 | 39.427 | 31.881 | 1.00 | 0.00 | H |
| ATOM | 723 | C | LEU | A | 66 | 47.699 | 38.485 | 28.303 | 1.00 | 0.00 | C |
| ATOM | 724 | O | LEU | A | 66 | 48.611 | 38.981 | 27.701 | 1.00 | 0.00 | O |
| ATOM | 725 | N | SER | A | 67 | 46.613 | 38.117 | 27.638 | 1.00 | 0.00 | N |
| ATOM | 726 | H | SER | A | 67 | 45.981 | 37.501 | 28.108 | 1.00 | 0.00 | H |
| ATOM | 727 | CA | SER | A | 67 | 46.249 | 38.530 | 26.272 | 1.00 | 0.00 | C |
| ATOM | 728 | HA | SER | A | 67 | 46.501 | 39.496 | 26.217 | 1.00 | 0.00 | H |
| ATOM | 729 | CB | SER | A | 67 | 47.018 | 37.639 | 25.282 | 1.00 | 0.00 | C |
| ATOM | 730 | HB1 | SER | A | 67 | 46.523 | 36.786 | 25.116 | 1.00 | 0.00 | H |
| ATOM | 731 | HB2 | SER | A | 67 | 47.932 | 37.431 | 25.630 | 1.00 | 0.00 | H |
| ATOM | 732 | OG | SER | A | 67 | 47.174 | 38.300 | 24.025 | 1.00 | 0.00 | O |
| ATOM | 733 | HG | SER | A | 67 | 47.675 | 37.706 | 23.396 | 1.00 | 0.00 | H |
| ATOM | 734 | C | SER | A | 67 | 44.744 | 38.408 | 25.977 | 1.00 | 0.00 | C |
| ATOM | 735 | O | SER | A | 67 | 44.011 | 37.748 | 26.647 | 1.00 | 0.00 | O |
| ATOM | 736 | N | THR | A | 68 | 44.227 | 39.096 | 24.915 | 1.00 | 0.00 | N |
| ATOM | 737 | H | THR | A | 68 | 44.859 | 39.677 | 24.401 | 1.00 | 0.00 | H |
| ATOM | 738 | CA | THR | A | 68 | 42.822 | 39.063 | 24.454 | 1.00 | 0.00 | C |
| ATOM | 739 | HA | THR | A | 68 | 42.249 | 39.136 | 25.270 | 1.00 | 0.00 | H |
| ATOM | 740 | CB | THR | A | 68 | 42.656 | 40.278 | 23.502 | 1.00 | 0.00 | C |
| ATOM | 741 | HB | THR | A | 68 | 41.710 | 40.258 | 23.178 | 1.00 | 0.00 | H |
| ATOM | 742 | CG2 | THR | A | 68 | 43.002 | 41.587 | 24.155 | 1.00 | 0.00 | C |
| ATOM | 743 | HG21 | THR | A | 68 | 42.877 | 42.331 | 23.498 | 1.00 | 0.00 | H |
| ATOM | 744 | HG22 | THR | A | 68 | 42.404 | 41.735 | 24.943 | 1.00 | 0.00 | H |
| ATOM | 745 | HG23 | THR | A | 68 | 43.955 | 41.567 | 24.458 | 1.00 | 0.00 | H |
| ATOM | 746 | OG1 | THR | A | 68 | 43.381 | 40.210 | 22.321 | 1.00 | 0.00 | O |
| ATOM | 747 | HG1 | THR | A | 68 | 43.211 | 41.032 | 21.777 | 1.00 | 0.00 | H |
| ATOM | 748 | C | THR | A | 68 | 42.413 | 37.709 | 23.762 | 1.00 | 0.00 | C |
| ATOM | 749 | O | THR | A | 68 | 43.292 | 36.878 | 23.503 | 1.00 | 0.00 | O |
| ATOM | 750 | N | THR | A | 69 | 41.131 | 37.424 | 23.616 | 1.00 | 0.00 | N |
| ATOM | 751 | H | THR | A | 69 | 40.458 | 38.082 | 23.954 | 1.00 | 0.00 | H |
| ATOM | 752 | CA | THR | A | 69 | 40.644 | 36.195 | 22.985 | 1.00 | 0.00 | C |
| ATOM | 753 | HA | THR | A | 69 | 41.441 | 35.619 | 22.805 | 1.00 | 0.00 | H |

| | | | | | | | | | | | |
|---|---|---|---|---|---|---|---|---|---|---|---|
| ATOM | 754 | CB | THR | A | 69 | 39.700 | 35.428 | 23.925 | 1.00 | 0.00 | C |
| ATOM | 755 | HB | THR | A | 69 | 38.868 | 35.970 | 24.041 | 1.00 | 0.00 | H |
| ATOM | 756 | CG2 | THR | A | 69 | 39.408 | 34.021 | 23.414 | 1.00 | 0.00 | C |
| ATOM | 757 | HG21 | THR | A | 69 | 38.793 | 33.556 | 24.051 | 1.00 | 0.00 | H |
| ATOM | 758 | HG22 | THR | A | 69 | 38.976 | 34.076 | 22.514 | 1.00 | 0.00 | H |
| ATOM | 759 | HG23 | THR | A | 69 | 40.263 | 33.508 | 23.343 | 1.00 | 0.00 | H |
| ATOM | 760 | OG1 | THR | A | 69 | 40.272 | 35.270 | 25.205 | 1.00 | 0.00 | O |
| ATOM | 761 | HG1 | THR | A | 69 | 39.638 | 34.769 | 25.794 | 1.00 | 0.00 | H |
| ATOM | 762 | C | THR | A | 69 | 39.974 | 36.547 | 21.668 | 1.00 | 0.00 | C |
| ATOM | 763 | O | THR | A | 69 | 40.399 | 36.074 | 20.621 | 1.00 | 0.00 | O |
| ATOM | 764 | N | GLU | A | 70 | 39.044 | 37.502 | 21.699 | 1.00 | 0.00 | N |
| ATOM | 765 | H | GLU | A | 70 | 38.920 | 38.008 | 22.553 | 1.00 | 0.00 | H |
| ATOM | 766 | CA | GLU | A | 70 | 38.193 | 37.857 | 20.561 | 1.00 | 0.00 | C |
| ATOM | 767 | HA | GLU | A | 70 | 38.455 | 37.252 | 19.809 | 1.00 | 0.00 | H |
| ATOM | 768 | CB | GLU | A | 70 | 36.725 | 37.629 | 20.948 | 1.00 | 0.00 | C |
| ATOM | 769 | HB1 | GLU | A | 70 | 36.152 | 37.866 | 20.164 | 1.00 | 0.00 | H |
| ATOM | 770 | HB2 | GLU | A | 70 | 36.504 | 38.230 | 21.716 | 1.00 | 0.00 | H |
| ATOM | 771 | CG | GLU | A | 70 | 36.415 | 36.179 | 21.358 | 1.00 | 0.00 | C |
| ATOM | 772 | HG1 | GLU | A | 70 | 35.457 | 36.121 | 21.640 | 1.00 | 0.00 | H |
| ATOM | 773 | HG2 | GLU | A | 70 | 37.005 | 35.927 | 22.125 | 1.00 | 0.00 | H |
| ATOM | 774 | CD | GLU | A | 70 | 36.648 | 35.186 | 20.218 | 1.00 | 0.00 | C |
| ATOM | 775 | OE1 | GLU | A | 70 | 35.662 | 34.920 | 19.507 | 1.00 | 0.00 | O |
| ATOM | 776 | OE2 | GLU | A | 70 | 37.792 | 34.719 | 20.040 | 1.00 | 0.00 | O |
| ATOM | 777 | C | GLU | A | 70 | 38.437 | 39.301 | 20.105 | 1.00 | 0.00 | C |
| ATOM | 778 | O | GLU | A | 70 | 38.757 | 40.169 | 20.917 | 1.00 | 0.00 | O |
| ATOM | 779 | N | TRP | A | 71 | 38.249 | 39.598 | 18.811 | 1.00 | 0.00 | N |
| ATOM | 780 | H | TRP | A | 71 | 37.787 | 38.936 | 18.221 | 1.00 | 0.00 | H |
| ATOM | 781 | CA | TRP | A | 71 | 38.707 | 40.879 | 18.234 | 1.00 | 0.00 | C |
| ATOM | 782 | HA | TRP | A | 71 | 39.668 | 40.924 | 18.505 | 1.00 | 0.00 | H |
| ATOM | 783 | CB | TRP | A | 71 | 38.580 | 40.838 | 16.700 | 1.00 | 0.00 | C |
| ATOM | 784 | HB1 | TRP | A | 71 | 38.712 | 39.892 | 16.405 | 1.00 | 0.00 | H |
| ATOM | 785 | HB2 | TRP | A | 71 | 39.303 | 41.411 | 16.315 | 1.00 | 0.00 | H |
| ATOM | 786 | CG | TRP | A | 71 | 37.285 | 41.306 | 16.095 | 1.00 | 0.00 | C |
| ATOM | 787 | CD1 | TRP | A | 71 | 36.111 | 40.632 | 16.082 | 1.00 | 0.00 | C |
| ATOM | 788 | HD1 | TRP | A | 71 | 35.959 | 39.732 | 16.491 | 1.00 | 0.00 | H |
| ATOM | 789 | NE1 | TRP | A | 71 | 35.153 | 41.376 | 15.418 | 1.00 | 0.00 | N |
| ATOM | 790 | HE1 | TRP | A | 71 | 34.216 | 41.068 | 15.253 | 1.00 | 0.00 | H |
| ATOM | 791 | CE2 | TRP | A | 71 | 35.652 | 42.595 | 15.013 | 1.00 | 0.00 | C |
| ATOM | 792 | CZ2 | TRP | A | 71 | 35.082 | 43.695 | 14.359 | 1.00 | 0.00 | C |
| ATOM | 793 | HZ2 | TRP | A | 71 | 34.108 | 43.697 | 14.131 | 1.00 | 0.00 | H |
| ATOM | 794 | CH2 | TRP | A | 71 | 35.891 | 44.795 | 14.027 | 1.00 | 0.00 | C |
| ATOM | 795 | HH2 | TRP | A | 71 | 35.497 | 45.582 | 13.553 | 1.00 | 0.00 | H |
| ATOM | 796 | CZ3 | TRP | A | 71 | 37.256 | 44.785 | 14.364 | 1.00 | 0.00 | C |
| ATOM | 797 | HZ3 | TRP | A | 71 | 37.835 | 45.562 | 14.116 | 1.00 | 0.00 | H |
| ATOM | 798 | CE3 | TRP | A | 71 | 37.813 | 43.686 | 15.050 | 1.00 | 0.00 | C |
| ATOM | 799 | HE3 | TRP | A | 71 | 38.782 | 43.700 | 15.296 | 1.00 | 0.00 | H |
| ATOM | 800 | CD2 | TRP | A | 71 | 37.026 | 42.565 | 15.396 | 1.00 | 0.00 | C |
| ATOM | 801 | C | TRP | A | 71 | 38.055 | 42.143 | 18.842 | 1.00 | 0.00 | C |
| ATOM | 802 | O | TRP | A | 71 | 38.652 | 43.222 | 18.819 | 1.00 | 0.00 | O |
| ATOM | 803 | N | ASN | A | 72 | 36.869 | 41.992 | 19.449 | 1.00 | 0.00 | N |
| ATOM | 804 | H | ASN | A | 72 | 36.466 | 41.077 | 19.452 | 1.00 | 0.00 | H |
| ATOM | 805 | CA | ASN | A | 72 | 36.116 | 43.065 | 20.109 | 1.00 | 0.00 | C |
| ATOM | 806 | HA | ASN | A | 72 | 36.419 | 43.904 | 19.657 | 1.00 | 0.00 | H |
| ATOM | 807 | CB | ASN | A | 72 | 34.645 | 42.822 | 19.903 | 1.00 | 0.00 | C |

| | | | | | | | | | | | |
|---|---|---|---|---|---|---|---|---|---|---|---|
| ATOM | 808 | HB1 | ASN | A | 72 | 34.134 | 43.470 | 20.468 | 1.00 | 0.00 | H |
| ATOM | 809 | HB2 | ASN | A | 72 | 34.434 | 41.889 | 20.194 | 1.00 | 0.00 | H |
| ATOM | 810 | CG | ASN | A | 72 | 34.246 | 43.002 | 18.427 | 1.00 | 0.00 | C |
| ATOM | 811 | OD1 | ASN | A | 72 | 34.844 | 43.792 | 17.712 | 1.00 | 0.00 | O |
| ATOM | 812 | ND2 | ASN | A | 72 | 33.138 | 42.433 | 18.026 | 1.00 | 0.00 | N |
| ATOM | 813 | HD21 | ASN | A | 72 | 32.585 | 41.905 | 18.671 | 1.00 | 0.00 | H |
| ATOM | 814 | HD22 | ASN | A | 72 | 32.844 | 42.526 | 17.075 | 1.00 | 0.00 | H |
| ATOM | 815 | C | ASN | A | 72 | 36.406 | 43.196 | 21.614 | 1.00 | 0.00 | C |
| ATOM | 816 | O | ASN | A | 72 | 35.901 | 44.178 | 22.266 | 1.00 | 0.00 | O |
| ATOM | 817 | N | ASP | A | 73 | 37.218 | 42.299 | 22.282 | 1.00 | 0.00 | N |
| ATOM | 818 | H | ASP | A | 73 | 37.525 | 41.504 | 21.760 | 1.00 | 0.00 | H |
| ATOM | 819 | CA | ASP | A | 73 | 37.672 | 42.382 | 23.648 | 1.00 | 0.00 | C |
| ATOM | 820 | HA | ASP | A | 73 | 36.838 | 42.096 | 24.119 | 1.00 | 0.00 | H |
| ATOM | 821 | CB | ASP | A | 73 | 38.807 | 41.435 | 23.946 | 1.00 | 0.00 | C |
| ATOM | 822 | HB1 | ASP | A | 73 | 39.250 | 41.710 | 24.800 | 1.00 | 0.00 | H |
| ATOM | 823 | HB2 | ASP | A | 73 | 39.471 | 41.472 | 23.199 | 1.00 | 0.00 | H |
| ATOM | 824 | CG | ASP | A | 73 | 38.274 | 40.019 | 24.083 | 1.00 | 0.00 | C |
| ATOM | 825 | OD1 | ASP | A | 73 | 37.044 | 39.772 | 24.201 | 1.00 | 0.00 | O |
| ATOM | 826 | OD2 | ASP | A | 73 | 39.134 | 39.137 | 24.273 | 1.00 | 0.00 | O |
| ATOM | 827 | C | ASP | A | 73 | 38.029 | 43.768 | 24.130 | 1.00 | 0.00 | C |
| ATOM | 828 | O | ASP | A | 73 | 37.454 | 44.247 | 25.078 | 1.00 | 0.00 | O |
| ATOM | 829 | N | GLU | A | 74 | 39.154 | 44.373 | 23.686 | 1.00 | 0.00 | N |
| ATOM | 830 | H | GLU | A | 74 | 39.622 | 43.988 | 22.891 | 1.00 | 0.00 | H |
| ATOM | 831 | CA | GLU | A | 74 | 39.710 | 45.554 | 24.312 | 1.00 | 0.00 | C |
| ATOM | 832 | HA | GLU | A | 74 | 39.826 | 45.217 | 25.247 | 1.00 | 0.00 | H |
| ATOM | 833 | CB | GLU | A | 74 | 41.124 | 45.878 | 23.970 | 1.00 | 0.00 | C |
| ATOM | 834 | HB1 | GLU | A | 74 | 41.693 | 45.104 | 24.248 | 1.00 | 0.00 | H |
| ATOM | 835 | HB2 | GLU | A | 74 | 41.388 | 46.690 | 24.490 | 1.00 | 0.00 | H |
| ATOM | 836 | CG | GLU | A | 74 | 41.315 | 46.145 | 22.457 | 1.00 | 0.00 | C |
| ATOM | 837 | HG1 | GLU | A | 74 | 40.586 | 46.758 | 22.152 | 1.00 | 0.00 | H |
| ATOM | 838 | HG2 | GLU | A | 74 | 41.241 | 45.273 | 21.973 | 1.00 | 0.00 | H |
| ATOM | 839 | CD | GLU | A | 74 | 42.617 | 46.770 | 22.037 | 1.00 | 0.00 | C |
| ATOM | 840 | OE1 | GLU | A | 74 | 43.004 | 47.798 | 22.611 | 1.00 | 0.00 | O |
| ATOM | 841 | OE2 | GLU | A | 74 | 43.264 | 46.279 | 21.038 | 1.00 | 0.00 | O |
| ATOM | 842 | C | GLU | A | 74 | 38.714 | 46.813 | 24.164 | 1.00 | 0.00 | C |
| ATOM | 843 | O | GLU | A | 74 | 38.852 | 47.843 | 24.781 | 1.00 | 0.00 | O |
| ATOM | 844 | N | THR | A | 75 | 37.721 | 46.648 | 23.268 | 1.00 | 0.00 | N |
| ATOM | 845 | H | THR | A | 75 | 37.829 | 45.927 | 22.584 | 1.00 | 0.00 | H |
| ATOM | 846 | CA | THR | A | 75 | 36.493 | 47.450 | 23.223 | 1.00 | 0.00 | C |
| ATOM | 847 | HA | THR | A | 75 | 36.789 | 48.389 | 23.400 | 1.00 | 0.00 | H |
| ATOM | 848 | CB | THR | A | 75 | 35.865 | 47.452 | 21.821 | 1.00 | 0.00 | C |
| ATOM | 849 | HB | THR | A | 75 | 35.455 | 46.557 | 21.648 | 1.00 | 0.00 | H |
| ATOM | 850 | CG2 | THR | A | 75 | 34.870 | 48.538 | 21.679 | 1.00 | 0.00 | C |
| ATOM | 851 | HG21 | THR | A | 75 | 34.479 | 48.514 | 20.759 | 1.00 | 0.00 | H |
| ATOM | 852 | HG22 | THR | A | 75 | 34.142 | 48.413 | 22.353 | 1.00 | 0.00 | H |
| ATOM | 853 | HG23 | THR | A | 75 | 35.316 | 49.420 | 21.829 | 1.00 | 0.00 | H |
| ATOM | 854 | OG1 | THR | A | 75 | 36.773 | 47.549 | 20.789 | 1.00 | 0.00 | O |
| ATOM | 855 | HG1 | THR | A | 75 | 36.286 | 47.544 | 19.915 | 1.00 | 0.00 | H |
| ATOM | 856 | C | THR | A | 75 | 35.514 | 47.046 | 24.353 | 1.00 | 0.00 | C |
| ATOM | 857 | O | THR | A | 75 | 35.005 | 47.866 | 25.065 | 1.00 | 0.00 | O |
| ATOM | 858 | N | LEU | A | 76 | 35.123 | 45.806 | 24.385 | 1.00 | 0.00 | N |
| ATOM | 859 | H | LEU | A | 76 | 35.588 | 45.132 | 23.811 | 1.00 | 0.00 | H |
| ATOM | 860 | CA | LEU | A | 76 | 34.006 | 45.370 | 25.251 | 1.00 | 0.00 | C |
| ATOM | 861 | HA | LEU | A | 76 | 33.333 | 46.101 | 25.134 | 1.00 | 0.00 | H |

| | | | | | | | | | | | |
|---|---|---|---|---|---|---|---|---|---|---|---|
| ATOM | 862 | CB | LEU | A | 76 | 33.441 | 44.029 | 24.789 | 1.00 | 0.00 | C |
| ATOM | 863 | HB1 | LEU | A | 76 | 32.708 | 43.782 | 25.423 | 1.00 | 0.00 | H |
| ATOM | 864 | HB2 | LEU | A | 76 | 34.179 | 43.358 | 24.855 | 1.00 | 0.00 | H |
| ATOM | 865 | CG | LEU | A | 76 | 32.869 | 43.939 | 23.374 | 1.00 | 0.00 | C |
| ATOM | 866 | HG | LEU | A | 76 | 33.659 | 44.084 | 22.779 | 1.00 | 0.00 | H |
| ATOM | 867 | CD1 | LEU | A | 76 | 32.117 | 42.550 | 23.185 | 1.00 | 0.00 | C |
| ATOM | 868 | HD11 | LEU | A | 76 | 31.745 | 42.496 | 22.258 | 1.00 | 0.00 | H |
| ATOM | 869 | HD12 | LEU | A | 76 | 32.762 | 41.799 | 23.330 | 1.00 | 0.00 | H |
| ATOM | 870 | HD13 | LEU | A | 76 | 31.372 | 42.482 | 23.848 | 1.00 | 0.00 | H |
| ATOM | 871 | CD2 | LEU | A | 76 | 31.868 | 45.021 | 23.037 | 1.00 | 0.00 | C |
| ATOM | 872 | HD21 | LEU | A | 76 | 31.542 | 44.894 | 22.100 | 1.00 | 0.00 | H |
| ATOM | 873 | HD22 | LEU | A | 76 | 31.095 | 44.968 | 23.669 | 1.00 | 0.00 | H |
| ATOM | 874 | HD23 | LEU | A | 76 | 32.305 | 45.917 | 23.119 | 1.00 | 0.00 | H |
| ATOM | 875 | C | LEU | A | 76 | 34.402 | 45.245 | 26.744 | 1.00 | 0.00 | C |
| ATOM | 876 | O | LEU | A | 76 | 33.458 | 45.371 | 27.575 | 1.00 | 0.00 | O |
| ATOM | 877 | N | TYR | A | 77 | 35.657 | 44.972 | 27.141 | 1.00 | 0.00 | N |
| ATOM | 878 | H | TYR | A | 77 | 36.284 | 44.657 | 26.429 | 1.00 | 0.00 | H |
| ATOM | 879 | CA | TYR | A | 77 | 36.236 | 45.084 | 28.535 | 1.00 | 0.00 | C |
| ATOM | 880 | HA | TYR | A | 77 | 35.871 | 44.360 | 29.120 | 1.00 | 0.00 | H |
| ATOM | 881 | CB | TYR | A | 77 | 37.730 | 44.858 | 28.522 | 1.00 | 0.00 | C |
| ATOM | 882 | HB1 | TYR | A | 77 | 38.028 | 44.985 | 29.468 | 1.00 | 0.00 | H |
| ATOM | 883 | HB2 | TYR | A | 77 | 38.107 | 45.589 | 27.953 | 1.00 | 0.00 | H |
| ATOM | 884 | CG | TYR | A | 77 | 38.402 | 43.587 | 28.050 | 1.00 | 0.00 | C |
| ATOM | 885 | CD1 | TYR | A | 77 | 37.761 | 42.358 | 28.241 | 1.00 | 0.00 | C |
| ATOM | 886 | HD1 | TYR | A | 77 | 36.812 | 42.333 | 28.556 | 1.00 | 0.00 | H |
| ATOM | 887 | CE1 | TYR | A | 77 | 38.450 | 41.155 | 27.990 | 1.00 | 0.00 | C |
| ATOM | 888 | HE1 | TYR | A | 77 | 37.950 | 40.290 | 28.040 | 1.00 | 0.00 | H |
| ATOM | 889 | CZ | TYR | A | 77 | 39.820 | 41.134 | 27.671 | 1.00 | 0.00 | C |
| ATOM | 890 | OH | TYR | A | 77 | 40.427 | 40.000 | 27.205 | 1.00 | 0.00 | O |
| ATOM | 891 | HH | TYR | A | 77 | 41.395 | 40.185 | 27.037 | 1.00 | 0.00 | H |
| ATOM | 892 | CE2 | TYR | A | 77 | 40.490 | 42.377 | 27.504 | 1.00 | 0.00 | C |
| ATOM | 893 | HE2 | TYR | A | 77 | 41.455 | 42.399 | 27.244 | 1.00 | 0.00 | H |
| ATOM | 894 | CD2 | TYR | A | 77 | 39.779 | 43.582 | 27.709 | 1.00 | 0.00 | C |
| ATOM | 895 | HD2 | TYR | A | 77 | 40.259 | 44.454 | 27.611 | 1.00 | 0.00 | H |
| ATOM | 896 | C | TYR | A | 77 | 35.844 | 46.460 | 29.103 | 1.00 | 0.00 | C |
| ATOM | 897 | O | TYR | A | 77 | 35.376 | 46.601 | 30.257 | 1.00 | 0.00 | O |
| ATOM | 898 | N | GLN | A | 78 | 36.024 | 47.555 | 28.295 | 1.00 | 0.00 | N |
| ATOM | 899 | H | GLN | A | 78 | 36.349 | 47.443 | 27.356 | 1.00 | 0.00 | H |
| ATOM | 900 | CA | GLN | A | 78 | 35.742 | 48.851 | 28.810 | 1.00 | 0.00 | C |
| ATOM | 901 | HA | GLN | A | 78 | 36.168 | 48.835 | 29.714 | 1.00 | 0.00 | H |
| ATOM | 902 | CB | GLN | A | 78 | 36.332 | 49.855 | 27.804 | 1.00 | 0.00 | C |
| ATOM | 903 | HB1 | GLN | A | 78 | 36.343 | 50.747 | 28.257 | 1.00 | 0.00 | H |
| ATOM | 904 | HB2 | GLN | A | 78 | 35.707 | 49.885 | 27.024 | 1.00 | 0.00 | H |
| ATOM | 905 | CG | GLN | A | 78 | 37.737 | 49.635 | 27.233 | 1.00 | 0.00 | C |
| ATOM | 906 | HG1 | GLN | A | 78 | 37.785 | 50.043 | 26.321 | 1.00 | 0.00 | H |
| ATOM | 907 | HG2 | GLN | A | 78 | 37.912 | 48.652 | 27.169 | 1.00 | 0.00 | H |
| ATOM | 908 | CD | GLN | A | 78 | 38.808 | 50.279 | 28.129 | 1.00 | 0.00 | C |
| ATOM | 909 | OE1 | GLN | A | 78 | 39.712 | 51.081 | 27.609 | 1.00 | 0.00 | O |
| ATOM | 910 | NE2 | GLN | A | 78 | 38.779 | 50.289 | 29.372 | 1.00 | 0.00 | N |
| ATOM | 911 | HE21 | GLN | A | 78 | 38.019 | 49.857 | 29.858 | 1.00 | 0.00 | H |
| ATOM | 912 | HE22 | GLN | A | 78 | 39.515 | 50.729 | 29.886 | 1.00 | 0.00 | H |
| ATOM | 913 | C | GLN | A | 78 | 34.196 | 49.168 | 28.901 | 1.00 | 0.00 | C |
| ATOM | 914 | O | GLN | A | 78 | 33.766 | 49.874 | 29.835 | 1.00 | 0.00 | O |
| ATOM | 915 | N | GLU | A | 79 | 33.366 | 48.678 | 27.943 | 1.00 | 0.00 | N |

| | | | | | | | | | | |
|---|---|---|---|---|---|---|---|---|---|---|
| ATOM | 916 | H | GLU | A | 79 | 33.771 | 48.237 | 27.142 | 1.00 | 0.00 | H |
| ATOM | 917 | CA | GLU | A | 79 | 31.905 | 48.766 | 28.029 | 1.00 | 0.00 | C |
| ATOM | 918 | HA | GLU | A | 79 | 31.658 | 49.714 | 28.229 | 1.00 | 0.00 | H |
| ATOM | 919 | CB | GLU | A | 79 | 31.320 | 48.329 | 26.656 | 1.00 | 0.00 | C |
| ATOM | 920 | HB1 | GLU | A | 79 | 30.321 | 48.318 | 26.706 | 1.00 | 0.00 | H |
| ATOM | 921 | HB2 | GLU | A | 79 | 31.652 | 47.414 | 26.426 | 1.00 | 0.00 | H |
| ATOM | 922 | CG | GLU | A | 79 | 31.670 | 49.191 | 25.536 | 1.00 | 0.00 | C |
| ATOM | 923 | HG1 | GLU | A | 79 | 31.560 | 48.654 | 24.700 | 1.00 | 0.00 | H |
| ATOM | 924 | HG2 | GLU | A | 79 | 32.631 | 49.447 | 25.644 | 1.00 | 0.00 | H |
| ATOM | 925 | CD | GLU | A | 79 | 30.808 | 50.454 | 25.453 | 1.00 | 0.00 | C |
| ATOM | 926 | OE1 | GLU | A | 79 | 30.431 | 51.020 | 26.506 | 1.00 | 0.00 | O |
| ATOM | 927 | OE2 | GLU | A | 79 | 30.456 | 50.864 | 24.294 | 1.00 | 0.00 | O |
| ATOM | 928 | C | GLU | A | 79 | 31.366 | 47.995 | 29.214 | 1.00 | 0.00 | C |
| ATOM | 929 | O | GLU | A | 79 | 30.442 | 48.462 | 29.857 | 1.00 | 0.00 | O |
| ATOM | 930 | N | PHE | A | 80 | 31.872 | 46.781 | 29.455 | 1.00 | 0.00 | N |
| ATOM | 931 | H | PHE | A | 80 | 32.478 | 46.401 | 28.756 | 1.00 | 0.00 | H |
| ATOM | 932 | CA | PHE | A | 80 | 31.626 | 45.939 | 30.655 | 1.00 | 0.00 | C |
| ATOM | 933 | HA | PHE | A | 80 | 30.658 | 45.695 | 30.719 | 1.00 | 0.00 | H |
| ATOM | 934 | CB | PHE | A | 80 | 32.390 | 44.631 | 30.481 | 1.00 | 0.00 | C |
| ATOM | 935 | HB1 | PHE | A | 80 | 33.357 | 44.866 | 30.378 | 1.00 | 0.00 | H |
| ATOM | 936 | HB2 | PHE | A | 80 | 32.059 | 44.206 | 29.638 | 1.00 | 0.00 | H |
| ATOM | 937 | CG | PHE | A | 80 | 32.319 | 43.572 | 31.550 | 1.00 | 0.00 | C |
| ATOM | 938 | CD1 | PHE | A | 80 | 31.144 | 42.822 | 31.687 | 1.00 | 0.00 | C |
| ATOM | 939 | HD1 | PHE | A | 80 | 30.374 | 42.922 | 31.056 | 1.00 | 0.00 | H |
| ATOM | 940 | CE1 | PHE | A | 80 | 31.111 | 41.875 | 32.819 | 1.00 | 0.00 | C |
| ATOM | 941 | HE1 | PHE | A | 80 | 30.303 | 41.304 | 32.965 | 1.00 | 0.00 | H |
| ATOM | 942 | CZ | PHE | A | 80 | 32.189 | 41.773 | 33.678 | 1.00 | 0.00 | C |
| ATOM | 943 | HZ | PHE | A | 80 | 32.137 | 41.121 | 34.435 | 1.00 | 0.00 | H |
| ATOM | 944 | CE2 | PHE | A | 80 | 33.350 | 42.542 | 33.535 | 1.00 | 0.00 | C |
| ATOM | 945 | HE2 | PHE | A | 80 | 34.125 | 42.452 | 34.160 | 1.00 | 0.00 | H |
| ATOM | 946 | CD2 | PHE | A | 80 | 33.361 | 43.451 | 32.458 | 1.00 | 0.00 | C |
| ATOM | 947 | HD2 | PHE | A | 80 | 34.161 | 44.039 | 32.342 | 1.00 | 0.00 | H |
| ATOM | 948 | C | PHE | A | 80 | 31.973 | 46.747 | 31.939 | 1.00 | 0.00 | C |
| ATOM | 949 | O | PHE | A | 80 | 31.171 | 46.871 | 32.891 | 1.00 | 0.00 | O |
| ATOM | 950 | N | ASN | A | 81 | 33.215 | 47.232 | 32.069 | 1.00 | 0.00 | N |
| ATOM | 951 | H | ASN | A | 81 | 33.863 | 47.076 | 31.323 | 1.00 | 0.00 | H |
| ATOM | 952 | CA | ASN | A | 81 | 33.690 | 47.996 | 33.270 | 1.00 | 0.00 | C |
| ATOM | 953 | HA | ASN | A | 81 | 33.313 | 47.487 | 34.043 | 1.00 | 0.00 | H |
| ATOM | 954 | CB | ASN | A | 81 | 35.281 | 48.055 | 33.270 | 1.00 | 0.00 | C |
| ATOM | 955 | HB1 | ASN | A | 81 | 35.619 | 48.603 | 34.035 | 1.00 | 0.00 | H |
| ATOM | 956 | HB2 | ASN | A | 81 | 35.625 | 48.434 | 32.411 | 1.00 | 0.00 | H |
| ATOM | 957 | CG | ASN | A | 81 | 35.775 | 46.621 | 33.421 | 1.00 | 0.00 | C |
| ATOM | 958 | OD1 | ASN | A | 81 | 35.239 | 45.692 | 34.038 | 1.00 | 0.00 | O |
| ATOM | 959 | ND2 | ASN | A | 81 | 36.950 | 46.383 | 32.836 | 1.00 | 0.00 | N |
| ATOM | 960 | HD21 | ASN | A | 81 | 37.425 | 47.118 | 32.352 | 1.00 | 0.00 | H |
| ATOM | 961 | HD22 | ASN | A | 81 | 37.355 | 45.470 | 32.881 | 1.00 | 0.00 | H |
| ATOM | 962 | C | ASN | A | 81 | 33.113 | 49.429 | 33.366 | 1.00 | 0.00 | C |
| ATOM | 963 | O | ASN | A | 81 | 33.040 | 49.999 | 34.464 | 1.00 | 0.00 | O |
| ATOM | 964 | N | GLY | A | 82 | 32.602 | 49.958 | 32.234 | 1.00 | 0.00 | N |
| ATOM | 965 | H | GLY | A | 82 | 32.797 | 49.501 | 31.366 | 1.00 | 0.00 | H |
| ATOM | 966 | CA | GLY | A | 82 | 31.807 | 51.123 | 32.213 | 1.00 | 0.00 | C |
| ATOM | 967 | HA1 | GLY | A | 82 | 31.790 | 51.407 | 31.254 | 1.00 | 0.00 | H |
| ATOM | 968 | HA2 | GLY | A | 82 | 32.318 | 51.794 | 32.751 | 1.00 | 0.00 | H |
| ATOM | 969 | C | GLY | A | 82 | 30.365 | 51.036 | 32.744 | 1.00 | 0.00 | C |

| | | | | | | | | | | | |
|---|---|---|---|---|---|---|---|---|---|---|---|
| ATOM | 970 | O | GLY | A | 82 | 29.895 | 52.102 | 33.151 | 1.00 | 0.00 | O |
| ATOM | 971 | N | LEU | A | 83 | 29.769 | 49.836 | 32.935 | 1.00 | 0.00 | N |
| ATOM | 972 | H | LEU | A | 83 | 30.169 | 49.045 | 32.471 | 1.00 | 0.00 | H |
| ATOM | 973 | CA | LEU | A | 83 | 28.596 | 49.604 | 33.759 | 1.00 | 0.00 | C |
| ATOM | 974 | HA | LEU | A | 83 | 27.835 | 50.039 | 33.277 | 1.00 | 0.00 | H |
| ATOM | 975 | CB | LEU | A | 83 | 28.370 | 48.111 | 33.857 | 1.00 | 0.00 | C |
| ATOM | 976 | HB1 | LEU | A | 83 | 29.096 | 47.720 | 34.422 | 1.00 | 0.00 | H |
| ATOM | 977 | HB2 | LEU | A | 83 | 28.421 | 47.725 | 32.936 | 1.00 | 0.00 | H |
| ATOM | 978 | CG | LEU | A | 83 | 26.975 | 47.786 | 34.490 | 1.00 | 0.00 | C |
| ATOM | 979 | HG | LEU | A | 83 | 27.005 | 48.250 | 35.376 | 1.00 | 0.00 | H |
| ATOM | 980 | CD1 | LEU | A | 83 | 25.739 | 48.258 | 33.763 | 1.00 | 0.00 | C |
| ATOM | 981 | HD11 | LEU | A | 83 | 24.925 | 47.984 | 34.276 | 1.00 | 0.00 | H |
| ATOM | 982 | HD12 | LEU | A | 83 | 25.761 | 49.254 | 33.681 | 1.00 | 0.00 | H |
| ATOM | 983 | HD13 | LEU | A | 83 | 25.712 | 47.849 | 32.851 | 1.00 | 0.00 | H |
| ATOM | 984 | CD2 | LEU | A | 83 | 26.773 | 46.323 | 34.578 | 1.00 | 0.00 | C |
| ATOM | 985 | HD21 | LEU | A | 83 | 25.879 | 46.134 | 34.984 | 1.00 | 0.00 | H |
| ATOM | 986 | HD22 | LEU | A | 83 | 26.814 | 45.925 | 33.661 | 1.00 | 0.00 | H |
| ATOM | 987 | HD23 | LEU | A | 83 | 27.490 | 45.921 | 35.148 | 1.00 | 0.00 | H |
| ATOM | 988 | C | LEU | A | 83 | 28.633 | 50.303 | 35.108 | 1.00 | 0.00 | C |
| ATOM | 989 | O | LEU | A | 83 | 27.558 | 50.887 | 35.450 | 1.00 | 0.00 | O |
| ATOM | 990 | N | LYS | A | 84 | 29.788 | 50.340 | 35.832 | 1.00 | 0.00 | N |
| ATOM | 991 | H | LYS | A | 84 | 30.633 | 50.020 | 35.403 | 1.00 | 0.00 | H |
| ATOM | 992 | CA | LYS | A | 84 | 29.843 | 50.820 | 37.188 | 1.00 | 0.00 | C |
| ATOM | 993 | HA | LYS | A | 84 | 29.058 | 50.413 | 37.656 | 1.00 | 0.00 | H |
| ATOM | 994 | CB | LYS | A | 84 | 31.239 | 50.369 | 37.721 | 1.00 | 0.00 | C |
| ATOM | 995 | HB1 | LYS | A | 84 | 31.326 | 50.688 | 38.665 | 1.00 | 0.00 | H |
| ATOM | 996 | HB2 | LYS | A | 84 | 31.940 | 50.804 | 37.156 | 1.00 | 0.00 | H |
| ATOM | 997 | CG | LYS | A | 84 | 31.510 | 48.841 | 37.714 | 1.00 | 0.00 | C |
| ATOM | 998 | HG1 | LYS | A | 84 | 31.508 | 48.500 | 36.774 | 1.00 | 0.00 | H |
| ATOM | 999 | HG2 | LYS | A | 84 | 30.804 | 48.369 | 38.241 | 1.00 | 0.00 | H |
| ATOM | 1000 | CD | LYS | A | 84 | 32.858 | 48.549 | 38.335 | 1.00 | 0.00 | C |
| ATOM | 1001 | HD1 | LYS | A | 84 | 32.856 | 47.605 | 38.666 | 1.00 | 0.00 | H |
| ATOM | 1002 | HD2 | LYS | A | 84 | 32.993 | 49.172 | 39.105 | 1.00 | 0.00 | H |
| ATOM | 1003 | CE | LYS | A | 84 | 34.019 | 48.710 | 37.399 | 1.00 | 0.00 | C |
| ATOM | 1004 | HE1 | LYS | A | 84 | 34.061 | 49.658 | 37.082 | 1.00 | 0.00 | H |
| ATOM | 1005 | HE2 | LYS | A | 84 | 33.900 | 48.101 | 36.615 | 1.00 | 0.00 | H |
| ATOM | 1006 | NZ | LYS | A | 84 | 35.290 | 48.388 | 38.033 | 1.00 | 0.00 | N |
| ATOM | 1007 | HZ1 | LYS | A | 84 | 36.032 | 48.509 | 37.373 | 1.00 | 0.00 | H |
| ATOM | 1008 | HZ2 | LYS | A | 84 | 35.436 | 48.995 | 38.814 | 1.00 | 0.00 | H |
| ATOM | 1009 | HZ3 | LYS | A | 84 | 35.275 | 47.439 | 38.347 | 1.00 | 0.00 | H |
| ATOM | 1010 | C | LYS | A | 84 | 29.642 | 52.342 | 37.331 | 1.00 | 0.00 | C |
| ATOM | 1011 | O | LYS | A | 84 | 29.454 | 52.812 | 38.450 | 1.00 | 0.00 | O |
| ATOM | 1012 | N | LYS | A | 85 | 29.715 | 53.162 | 36.253 | 1.00 | 0.00 | N |
| ATOM | 1013 | H | LYS | A | 85 | 30.070 | 52.799 | 35.392 | 1.00 | 0.00 | H |
| ATOM | 1014 | CA | LYS | A | 85 | 29.281 | 54.593 | 36.311 | 1.00 | 0.00 | C |
| ATOM | 1015 | HA | LYS | A | 85 | 29.836 | 55.116 | 36.957 | 1.00 | 0.00 | H |
| ATOM | 1016 | CB | LYS | A | 85 | 29.573 | 55.212 | 34.969 | 1.00 | 0.00 | C |
| ATOM | 1017 | HB1 | LYS | A | 85 | 29.071 | 54.697 | 34.274 | 1.00 | 0.00 | H |
| ATOM | 1018 | HB2 | LYS | A | 85 | 30.555 | 55.134 | 34.798 | 1.00 | 0.00 | H |
| ATOM | 1019 | CG | LYS | A | 85 | 29.140 | 56.726 | 34.935 | 1.00 | 0.00 | C |
| ATOM | 1020 | HG1 | LYS | A | 85 | 29.479 | 57.181 | 35.759 | 1.00 | 0.00 | H |
| ATOM | 1021 | HG2 | LYS | A | 85 | 28.142 | 56.779 | 34.911 | 1.00 | 0.00 | H |
| ATOM | 1022 | CD | LYS | A | 85 | 29.713 | 57.465 | 33.681 | 1.00 | 0.00 | C |
| ATOM | 1023 | HD1 | LYS | A | 85 | 29.243 | 58.343 | 33.593 | 1.00 | 0.00 | H |

| | | | | | | | | | | | |
|---|---|---|---|---|---|---|---|---|---|---|---|
| ATOM | 1024 | HD2 | LYS | A | 85 | 29.526 | 56.904 | 32.874 | 1.00 | 0.00 | H |
| ATOM | 1025 | CE | LYS | A | 85 | 31.206 | 57.759 | 33.681 | 1.00 | 0.00 | C |
| ATOM | 1026 | HE1 | LYS | A | 85 | 31.712 | 56.900 | 33.760 | 1.00 | 0.00 | H |
| ATOM | 1027 | HE2 | LYS | A | 85 | 31.426 | 58.348 | 34.458 | 1.00 | 0.00 | H |
| ATOM | 1028 | NZ | LYS | A | 85 | 31.679 | 58.432 | 32.478 | 1.00 | 0.00 | N |
| ATOM | 1029 | HZ1 | LYS | A | 85 | 32.664 | 58.591 | 32.550 | 1.00 | 0.00 | H |
| ATOM | 1030 | HZ2 | LYS | A | 85 | 31.492 | 57.859 | 31.680 | 1.00 | 0.00 | H |
| ATOM | 1031 | HZ3 | LYS | A | 85 | 31.206 | 59.308 | 32.378 | 1.00 | 0.00 | H |
| ATOM | 1032 | C | LYS | A | 85 | 27.810 | 54.703 | 36.844 | 1.00 | 0.00 | C |
| ATOM | 1033 | O | LYS | A | 85 | 27.528 | 55.672 | 37.535 | 1.00 | 0.00 | O |
| ATOM | 1034 | N | MET | A | 86 | 26.875 | 53.833 | 36.415 | 1.00 | 0.00 | N |
| ATOM | 1035 | H | MET | A | 86 | 27.165 | 53.124 | 35.772 | 1.00 | 0.00 | H |
| ATOM | 1036 | CA | MET | A | 86 | 25.496 | 53.844 | 36.808 | 1.00 | 0.00 | C |
| ATOM | 1037 | HA | MET | A | 86 | 25.325 | 54.829 | 36.795 | 1.00 | 0.00 | H |
| ATOM | 1038 | CB | MET | A | 86 | 24.656 | 53.068 | 35.779 | 1.00 | 0.00 | C |
| ATOM | 1039 | HB1 | MET | A | 86 | 23.686 | 53.236 | 35.957 | 1.00 | 0.00 | H |
| ATOM | 1040 | HB2 | MET | A | 86 | 24.847 | 52.091 | 35.873 | 1.00 | 0.00 | H |
| ATOM | 1041 | CG | MET | A | 86 | 24.945 | 53.465 | 34.356 | 1.00 | 0.00 | C |
| ATOM | 1042 | HG1 | MET | A | 86 | 25.944 | 53.441 | 34.311 | 1.00 | 0.00 | H |
| ATOM | 1043 | HG2 | MET | A | 86 | 24.633 | 54.414 | 34.316 | 1.00 | 0.00 | H |
| ATOM | 1044 | SD | MET | A | 86 | 24.207 | 52.478 | 33.060 | 1.00 | 0.00 | S |
| ATOM | 1045 | CE | MET | A | 86 | 22.421 | 52.646 | 33.431 | 1.00 | 0.00 | C |
| ATOM | 1046 | HE1 | MET | A | 86 | 21.891 | 52.127 | 32.760 | 1.00 | 0.00 | H |
| ATOM | 1047 | HE2 | MET | A | 86 | 22.236 | 52.292 | 34.348 | 1.00 | 0.00 | H |
| ATOM | 1048 | HE3 | MET | A | 86 | 22.161 | 53.611 | 33.387 | 1.00 | 0.00 | H |
| ATOM | 1049 | C | MET | A | 86 | 25.197 | 53.248 | 38.248 | 1.00 | 0.00 | C |
| ATOM | 1050 | O | MET | A | 86 | 24.031 | 53.292 | 38.697 | 1.00 | 0.00 | O |
| ATOM | 1051 | N | ASN | A | 87 | 26.225 | 52.648 | 38.893 | 1.00 | 0.00 | N |
| ATOM | 1052 | H | ASN | A | 87 | 27.089 | 52.586 | 38.394 | 1.00 | 0.00 | H |
| ATOM | 1053 | CA | ASN | A | 87 | 26.207 | 52.073 | 40.259 | 1.00 | 0.00 | C |
| ATOM | 1054 | HA | ASN | A | 87 | 25.761 | 52.784 | 40.803 | 1.00 | 0.00 | H |
| ATOM | 1055 | CB | ASN | A | 87 | 25.337 | 50.789 | 40.405 | 1.00 | 0.00 | C |
| ATOM | 1056 | HB1 | ASN | A | 87 | 25.572 | 50.140 | 39.681 | 1.00 | 0.00 | H |
| ATOM | 1057 | HB2 | ASN | A | 87 | 24.369 | 51.030 | 40.336 | 1.00 | 0.00 | H |
| ATOM | 1058 | CG | ASN | A | 87 | 25.561 | 50.114 | 41.733 | 1.00 | 0.00 | C |
| ATOM | 1059 | OD1 | ASN | A | 87 | 25.831 | 50.742 | 42.709 | 1.00 | 0.00 | O |
| ATOM | 1060 | ND2 | ASN | A | 87 | 25.413 | 48.831 | 41.921 | 1.00 | 0.00 | N |
| ATOM | 1061 | HD21 | ASN | A | 87 | 25.137 | 48.243 | 41.161 | 1.00 | 0.00 | H |
| ATOM | 1062 | HD22 | ASN | A | 87 | 25.575 | 48.436 | 42.825 | 1.00 | 0.00 | H |
| ATOM | 1063 | C | ASN | A | 87 | 27.676 | 51.797 | 40.726 | 1.00 | 0.00 | C |
| ATOM | 1064 | O | ASN | A | 87 | 28.227 | 50.731 | 40.362 | 1.00 | 0.00 | O |
| ATOM | 1065 | N | PRO | A | 88 | 28.331 | 52.690 | 41.526 | 1.00 | 0.00 | N |
| ATOM | 1066 | CD | PRO | A | 88 | 27.753 | 53.933 | 42.095 | 1.00 | 0.00 | C |
| ATOM | 1067 | HD1 | PRO | A | 88 | 26.895 | 53.736 | 42.569 | 1.00 | 0.00 | H |
| ATOM | 1068 | HD2 | PRO | A | 88 | 27.588 | 54.605 | 41.373 | 1.00 | 0.00 | H |
| ATOM | 1069 | CG | PRO | A | 88 | 28.742 | 54.448 | 43.043 | 1.00 | 0.00 | C |
| ATOM | 1070 | HG1 | PRO | A | 88 | 28.539 | 54.115 | 43.964 | 1.00 | 0.00 | H |
| ATOM | 1071 | HG2 | PRO | A | 88 | 28.731 | 55.448 | 43.040 | 1.00 | 0.00 | H |
| ATOM | 1072 | CB | PRO | A | 88 | 30.086 | 53.859 | 42.473 | 1.00 | 0.00 | C |
| ATOM | 1073 | HB1 | PRO | A | 88 | 30.763 | 53.735 | 43.199 | 1.00 | 0.00 | H |
| ATOM | 1074 | HB2 | PRO | A | 88 | 30.466 | 54.449 | 41.760 | 1.00 | 0.00 | H |
| ATOM | 1075 | CA | PRO | A | 88 | 29.719 | 52.516 | 41.882 | 1.00 | 0.00 | C |
| ATOM | 1076 | HA | PRO | A | 88 | 30.277 | 52.299 | 41.081 | 1.00 | 0.00 | H |
| ATOM | 1077 | C | PRO | A | 88 | 29.973 | 51.297 | 42.747 | 1.00 | 0.00 | C |

```
ATOM   1078  O   PRO A  88      31.157  50.924  42.843  1.00  0.00           O
ATOM   1079  N   LYS A  89      28.913  50.611  43.280  1.00  0.00           N
ATOM   1080  H   LYS A  89      27.982  50.900  43.057  1.00  0.00           H
ATOM   1081  CA  LYS A  89      29.115  49.452  44.178  1.00  0.00           C
ATOM   1082  HA  LYS A  89      29.997  49.544  44.640  1.00  0.00           H
ATOM   1083  CB  LYS A  89      27.861  49.427  45.155  1.00  0.00           C
ATOM   1084  HB1 LYS A  89      27.086  49.032  44.662  1.00  0.00           H
ATOM   1085  HB2 LYS A  89      27.646  50.368  45.415  1.00  0.00           H
ATOM   1086  CG  LYS A  89      27.985  48.642  46.457  1.00  0.00           C
ATOM   1087  HG1 LYS A  89      28.457  47.782  46.262  1.00  0.00           H
ATOM   1088  HG2 LYS A  89      27.064  48.448  46.796  1.00  0.00           H
ATOM   1089  CD  LYS A  89      28.769  49.407  47.547  1.00  0.00           C
ATOM   1090  HD1 LYS A  89      28.283  50.234  47.829  1.00  0.00           H
ATOM   1091  HD2 LYS A  89      29.685  49.651  47.229  1.00  0.00           H
ATOM   1092  CE  LYS A  89      28.831  48.372  48.698  1.00  0.00           C
ATOM   1093  HE1 LYS A  89      29.275  47.539  48.368  1.00  0.00           H
ATOM   1094  HE2 LYS A  89      27.902  48.158  48.998  1.00  0.00           H
ATOM   1095  NZ  LYS A  89      29.563  48.783  49.890  1.00  0.00           N
ATOM   1096  HZ1 LYS A  89      29.541  48.046  50.565  1.00  0.00           H
ATOM   1097  HZ2 LYS A  89      30.512  48.983  49.646  1.00  0.00           H
ATOM   1098  HZ3 LYS A  89      29.138  49.602  50.276  1.00  0.00           H
ATOM   1099  C   LYS A  89      29.224  48.178  43.393  1.00  0.00           C
ATOM   1100  O   LYS A  89      29.740  47.239  43.937  1.00  0.00           O
ATOM   1101  N   LEU A  90      28.848  48.162  42.151  1.00  0.00           N
ATOM   1102  H   LEU A  90      28.472  48.981  41.717  1.00  0.00           H
ATOM   1103  CA  LEU A  90      28.993  46.883  41.388  1.00  0.00           C
ATOM   1104  HA  LEU A  90      28.542  46.213  41.977  1.00  0.00           H
ATOM   1105  CB  LEU A  90      28.251  46.949  39.991  1.00  0.00           C
ATOM   1106  HB1 LEU A  90      28.647  47.701  39.464  1.00  0.00           H
ATOM   1107  HB2 LEU A  90      27.283  47.135  40.161  1.00  0.00           H
ATOM   1108  CG  LEU A  90      28.331  45.667  39.107  1.00  0.00           C
ATOM   1109  HG  LEU A  90      29.260  45.363  38.895  1.00  0.00           H
ATOM   1110  CD1 LEU A  90      27.644  44.529  39.873  1.00  0.00           C
ATOM   1111 HD11 LEU A  90      27.680  43.691  39.328  1.00  0.00           H
ATOM   1112 HD12 LEU A  90      28.114  44.381  40.743  1.00  0.00           H
ATOM   1113 HD13 LEU A  90      26.690  44.773  40.047  1.00  0.00           H
ATOM   1114  CD2 LEU A  90      27.693  45.983  37.777  1.00  0.00           C
ATOM   1115 HD21 LEU A  90      27.731  45.175  37.189  1.00  0.00           H
ATOM   1116 HD22 LEU A  90      26.739  46.247  37.919  1.00  0.00           H
ATOM   1117 HD23 LEU A  90      28.186  46.736  37.341  1.00  0.00           H
ATOM   1118  C   LEU A  90      30.542  46.521  41.158  1.00  0.00           C
ATOM   1119  O   LEU A  90      31.388  47.329  40.851  1.00  0.00           O
ATOM   1120  N   LYS A  91      30.936  45.257  41.363  1.00  0.00           N
ATOM   1121  H   LYS A  91      30.286  44.616  41.772  1.00  0.00           H
ATOM   1122  CA  LYS A  91      32.321  44.751  41.005  1.00  0.00           C
ATOM   1123  HA  LYS A  91      32.883  45.578  40.992  1.00  0.00           H
ATOM   1124  CB  LYS A  91      32.946  43.824  42.042  1.00  0.00           C
ATOM   1125  HB1 LYS A  91      33.693  43.335  41.591  1.00  0.00           H
ATOM   1126  HB2 LYS A  91      32.241  43.172  42.321  1.00  0.00           H
ATOM   1127  CG  LYS A  91      33.512  44.515  43.322  1.00  0.00           C
ATOM   1128  HG1 LYS A  91      34.118  45.254  43.029  1.00  0.00           H
ATOM   1129  HG2 LYS A  91      34.039  43.834  43.831  1.00  0.00           H
ATOM   1130  CD  LYS A  91      32.560  45.128  44.313  1.00  0.00           C
ATOM   1131  HD1 LYS A  91      32.161  45.944  43.894  1.00  0.00           H
```

| | | | | | | | | | | | |
|---|---|---|---|---|---|---|---|---|---|---|---|
| ATOM | 1132 | HD2 | LYS | A | 91 | 33.084 | 45.391 | 45.123 | 1.00 | 0.00 | H |
| ATOM | 1133 | CE | LYS | A | 91 | 31.425 | 44.166 | 44.733 | 1.00 | 0.00 | C |
| ATOM | 1134 | HE1 | LYS | A | 91 | 31.769 | 43.519 | 45.413 | 1.00 | 0.00 | H |
| ATOM | 1135 | HE2 | LYS | A | 91 | 31.097 | 43.665 | 43.932 | 1.00 | 0.00 | H |
| ATOM | 1136 | NZ | LYS | A | 91 | 30.268 | 44.852 | 45.326 | 1.00 | 0.00 | N |
| ATOM | 1137 | HZ1 | LYS | A | 91 | 29.572 | 44.178 | 45.575 | 1.00 | 0.00 | H |
| ATOM | 1138 | HZ2 | LYS | A | 91 | 30.558 | 45.349 | 46.144 | 1.00 | 0.00 | H |
| ATOM | 1139 | HZ3 | LYS | A | 91 | 29.886 | 45.496 | 44.663 | 1.00 | 0.00 | H |
| ATOM | 1140 | C | LYS | A | 91 | 32.345 | 44.089 | 39.627 | 1.00 | 0.00 | C |
| ATOM | 1141 | O | LYS | A | 91 | 31.340 | 43.433 | 39.297 | 1.00 | 0.00 | O |
| ATOM | 1142 | N | THR | A | 92 | 33.411 | 44.146 | 38.935 | 1.00 | 0.00 | N |
| ATOM | 1143 | H | THR | A | 92 | 34.217 | 44.597 | 39.320 | 1.00 | 0.00 | H |
| ATOM | 1144 | CA | THR | A | 92 | 33.496 | 43.538 | 37.516 | 1.00 | 0.00 | C |
| ATOM | 1145 | HA | THR | A | 92 | 32.809 | 42.821 | 37.398 | 1.00 | 0.00 | H |
| ATOM | 1146 | CB | THR | A | 92 | 33.256 | 44.680 | 36.489 | 1.00 | 0.00 | C |
| ATOM | 1147 | HB | THR | A | 92 | 33.471 | 44.261 | 35.607 | 1.00 | 0.00 | H |
| ATOM | 1148 | CG2 | THR | A | 92 | 31.826 | 45.256 | 36.634 | 1.00 | 0.00 | C |
| ATOM | 1149 | HG21 | THR | A | 92 | 31.692 | 45.988 | 35.966 | 1.00 | 0.00 | H |
| ATOM | 1150 | HG22 | THR | A | 92 | 31.157 | 44.531 | 36.473 | 1.00 | 0.00 | H |
| ATOM | 1151 | HG23 | THR | A | 92 | 31.706 | 45.621 | 37.557 | 1.00 | 0.00 | H |
| ATOM | 1152 | OG1 | THR | A | 92 | 34.155 | 45.813 | 36.512 | 1.00 | 0.00 | O |
| ATOM | 1153 | HG1 | THR | A | 92 | 33.886 | 46.466 | 35.804 | 1.00 | 0.00 | H |
| ATOM | 1154 | C | THR | A | 92 | 34.903 | 42.917 | 37.310 | 1.00 | 0.00 | C |
| ATOM | 1155 | O | THR | A | 92 | 35.856 | 43.601 | 37.041 | 1.00 | 0.00 | O |
| ATOM | 1156 | N | LEU | A | 93 | 35.008 | 41.582 | 37.384 | 1.00 | 0.00 | N |
| ATOM | 1157 | H | LEU | A | 93 | 34.159 | 41.088 | 37.573 | 1.00 | 0.00 | H |
| ATOM | 1158 | CA | LEU | A | 93 | 36.217 | 40.748 | 37.222 | 1.00 | 0.00 | C |
| ATOM | 1159 | HA | LEU | A | 93 | 36.936 | 41.401 | 37.459 | 1.00 | 0.00 | H |
| ATOM | 1160 | CB | LEU | A | 93 | 36.178 | 39.526 | 38.160 | 1.00 | 0.00 | C |
| ATOM | 1161 | HB1 | LEU | A | 93 | 37.052 | 39.050 | 38.068 | 1.00 | 0.00 | H |
| ATOM | 1162 | HB2 | LEU | A | 93 | 35.439 | 38.930 | 37.845 | 1.00 | 0.00 | H |
| ATOM | 1163 | CG | LEU | A | 93 | 35.945 | 39.839 | 39.680 | 1.00 | 0.00 | C |
| ATOM | 1164 | HG | LEU | A | 93 | 36.459 | 40.668 | 39.899 | 1.00 | 0.00 | H |
| ATOM | 1165 | CD1 | LEU | A | 93 | 34.410 | 39.978 | 40.022 | 1.00 | 0.00 | C |
| ATOM | 1166 | HD11 | LEU | A | 93 | 34.302 | 40.177 | 40.996 | 1.00 | 0.00 | H |
| ATOM | 1167 | HD12 | LEU | A | 93 | 34.015 | 40.723 | 39.484 | 1.00 | 0.00 | H |
| ATOM | 1168 | HD13 | LEU | A | 93 | 33.941 | 39.123 | 39.802 | 1.00 | 0.00 | H |
| ATOM | 1169 | CD2 | LEU | A | 93 | 36.441 | 38.712 | 40.514 | 1.00 | 0.00 | C |
| ATOM | 1170 | HD21 | LEU | A | 93 | 36.289 | 38.920 | 41.480 | 1.00 | 0.00 | H |
| ATOM | 1171 | HD22 | LEU | A | 93 | 35.949 | 37.876 | 40.271 | 1.00 | 0.00 | H |
| ATOM | 1172 | HD23 | LEU | A | 93 | 37.419 | 38.582 | 40.351 | 1.00 | 0.00 | H |
| ATOM | 1173 | C | LEU | A | 93 | 36.481 | 40.242 | 35.798 | 1.00 | 0.00 | C |
| ATOM | 1174 | O | LEU | A | 93 | 35.575 | 40.263 | 34.914 | 1.00 | 0.00 | O |
| ATOM | 1175 | N | LEU | A | 94 | 37.637 | 39.660 | 35.554 | 1.00 | 0.00 | N |
| ATOM | 1176 | H | LEU | A | 94 | 38.312 | 39.673 | 36.291 | 1.00 | 0.00 | H |
| ATOM | 1177 | CA | LEU | A | 94 | 38.038 | 38.985 | 34.291 | 1.00 | 0.00 | C |
| ATOM | 1178 | HA | LEU | A | 94 | 37.290 | 39.089 | 33.636 | 1.00 | 0.00 | H |
| ATOM | 1179 | CB | LEU | A | 94 | 39.449 | 39.598 | 33.882 | 1.00 | 0.00 | C |
| ATOM | 1180 | HB1 | LEU | A | 94 | 40.129 | 39.364 | 34.576 | 1.00 | 0.00 | H |
| ATOM | 1181 | HB2 | LEU | A | 94 | 39.374 | 40.593 | 33.811 | 1.00 | 0.00 | H |
| ATOM | 1182 | CG | LEU | A | 94 | 40.034 | 39.173 | 32.622 | 1.00 | 0.00 | C |
| ATOM | 1183 | HG | LEU | A | 94 | 40.123 | 38.178 | 32.665 | 1.00 | 0.00 | H |
| ATOM | 1184 | CD1 | LEU | A | 94 | 39.098 | 39.570 | 31.459 | 1.00 | 0.00 | C |
| ATOM | 1185 | HD11 | LEU | A | 94 | 39.501 | 39.277 | 30.592 | 1.00 | 0.00 | H |

| | | | | | | | | | | | |
|---|---|---|---|---|---|---|---|---|---|---|---|
| ATOM | 1186 | HD12 | LEU | A | 94 | 38.210 | 39.127 | 31.580 | 1.00 | 0.00 | H |
| ATOM | 1187 | HD13 | LEU | A | 94 | 38.977 | 40.563 | 31.452 | 1.00 | 0.00 | H |
| ATOM | 1188 | CD2 | LEU | A | 94 | 41.424 | 39.814 | 32.462 | 1.00 | 0.00 | C |
| ATOM | 1189 | HD21 | LEU | A | 94 | 41.828 | 39.522 | 31.595 | 1.00 | 0.00 | H |
| ATOM | 1190 | HD22 | LEU | A | 94 | 41.335 | 40.810 | 32.470 | 1.00 | 0.00 | H |
| ATOM | 1191 | HD23 | LEU | A | 94 | 42.013 | 39.526 | 33.217 | 1.00 | 0.00 | H |
| ATOM | 1192 | C | LEU | A | 94 | 38.304 | 37.519 | 34.555 | 1.00 | 0.00 | C |
| ATOM | 1193 | O | LEU | A | 94 | 39.222 | 37.145 | 35.342 | 1.00 | 0.00 | O |
| ATOM | 1194 | N | ALA | A | 95 | 37.574 | 36.634 | 33.881 | 1.00 | 0.00 | N |
| ATOM | 1195 | H | ALA | A | 95 | 36.780 | 36.956 | 33.366 | 1.00 | 0.00 | H |
| ATOM | 1196 | CA | ALA | A | 95 | 37.892 | 35.212 | 33.868 | 1.00 | 0.00 | C |
| ATOM | 1197 | HA | ALA | A | 95 | 38.198 | 34.993 | 34.795 | 1.00 | 0.00 | H |
| ATOM | 1198 | CB | ALA | A | 95 | 36.647 | 34.376 | 33.548 | 1.00 | 0.00 | C |
| ATOM | 1199 | HB1 | ALA | A | 95 | 36.889 | 33.406 | 33.545 | 1.00 | 0.00 | H |
| ATOM | 1200 | HB2 | ALA | A | 95 | 35.946 | 34.542 | 34.242 | 1.00 | 0.00 | H |
| ATOM | 1201 | HB3 | ALA | A | 95 | 36.294 | 34.635 | 32.649 | 1.00 | 0.00 | H |
| ATOM | 1202 | C | ALA | A | 95 | 39.030 | 34.930 | 32.887 | 1.00 | 0.00 | C |
| ATOM | 1203 | O | ALA | A | 95 | 39.102 | 35.523 | 31.805 | 1.00 | 0.00 | O |
| ATOM | 1204 | N | ILE | A | 96 | 39.861 | 33.956 | 33.246 | 1.00 | 0.00 | N |
| ATOM | 1205 | H | ILE | A | 96 | 39.770 | 33.583 | 34.169 | 1.00 | 0.00 | H |
| ATOM | 1206 | CA | ILE | A | 96 | 40.896 | 33.395 | 32.384 | 1.00 | 0.00 | C |
| ATOM | 1207 | HA | ILE | A | 96 | 41.029 | 34.027 | 31.620 | 1.00 | 0.00 | H |
| ATOM | 1208 | CB | ILE | A | 96 | 42.227 | 33.292 | 33.154 | 1.00 | 0.00 | C |
| ATOM | 1209 | HB | ILE | A | 96 | 42.068 | 32.708 | 33.950 | 1.00 | 0.00 | H |
| ATOM | 1210 | CG2 | ILE | A | 96 | 43.307 | 32.679 | 32.237 | 1.00 | 0.00 | C |
| ATOM | 1211 | HG21 | ILE | A | 96 | 44.170 | 32.612 | 32.737 | 1.00 | 0.00 | H |
| ATOM | 1212 | HG22 | ILE | A | 96 | 43.019 | 31.767 | 31.946 | 1.00 | 0.00 | H |
| ATOM | 1213 | HG23 | ILE | A | 96 | 43.431 | 33.261 | 31.433 | 1.00 | 0.00 | H |
| ATOM | 1214 | CG1 | ILE | A | 96 | 42.693 | 34.594 | 33.761 | 1.00 | 0.00 | C |
| ATOM | 1215 | HG11 | ILE | A | 96 | 43.605 | 34.423 | 34.134 | 1.00 | 0.00 | H |
| ATOM | 1216 | HG12 | ILE | A | 96 | 42.061 | 34.801 | 34.508 | 1.00 | 0.00 | H |
| ATOM | 1217 | CD | ILE | A | 96 | 42.783 | 35.847 | 32.864 | 1.00 | 0.00 | C |
| ATOM | 1218 | HD1 | ILE | A | 96 | 43.105 | 36.623 | 33.407 | 1.00 | 0.00 | H |
| ATOM | 1219 | HD2 | ILE | A | 96 | 43.423 | 35.676 | 32.115 | 1.00 | 0.00 | H |
| ATOM | 1220 | HD3 | ILE | A | 96 | 41.879 | 36.054 | 32.489 | 1.00 | 0.00 | H |
| ATOM | 1221 | C | ILE | A | 96 | 40.413 | 32.048 | 31.826 | 1.00 | 0.00 | C |
| ATOM | 1222 | O | ILE | A | 96 | 39.949 | 31.190 | 32.575 | 1.00 | 0.00 | O |
| ATOM | 1223 | N | GLY | A | 97 | 40.489 | 31.877 | 30.506 | 1.00 | 0.00 | N |
| ATOM | 1224 | H | GLY | A | 97 | 40.942 | 32.590 | 29.971 | 1.00 | 0.00 | H |
| ATOM | 1225 | CA | GLY | A | 97 | 39.956 | 30.718 | 29.783 | 1.00 | 0.00 | C |
| ATOM | 1226 | HA1 | GLY | A | 97 | 39.050 | 30.511 | 30.152 | 1.00 | 0.00 | H |
| ATOM | 1227 | HA2 | GLY | A | 97 | 40.567 | 29.943 | 29.944 | 1.00 | 0.00 | H |
| ATOM | 1228 | C | GLY | A | 97 | 39.857 | 31.001 | 28.282 | 1.00 | 0.00 | C |
| ATOM | 1229 | O | GLY | A | 97 | 40.593 | 31.840 | 27.766 | 1.00 | 0.00 | O |
| ATOM | 1230 | N | GLY | A | 98 | 38.927 | 30.355 | 27.572 | 1.00 | 0.00 | N |
| ATOM | 1231 | H | GLY | A | 98 | 38.312 | 29.724 | 28.045 | 1.00 | 0.00 | H |
| ATOM | 1232 | CA | GLY | A | 98 | 38.776 | 30.538 | 26.125 | 1.00 | 0.00 | C |
| ATOM | 1233 | HA1 | GLY | A | 98 | 38.785 | 31.522 | 25.945 | 1.00 | 0.00 | H |
| ATOM | 1234 | HA2 | GLY | A | 98 | 37.887 | 30.160 | 25.867 | 1.00 | 0.00 | H |
| ATOM | 1235 | C | GLY | A | 98 | 39.874 | 29.861 | 25.299 | 1.00 | 0.00 | C |
| ATOM | 1236 | O | GLY | A | 98 | 40.866 | 29.370 | 25.837 | 1.00 | 0.00 | O |
| ATOM | 1237 | N | TRP | A | 99 | 39.723 | 29.857 | 23.971 | 1.00 | 0.00 | N |
| ATOM | 1238 | H | TRP | A | 99 | 38.957 | 30.367 | 23.580 | 1.00 | 0.00 | H |
| ATOM | 1239 | CA | TRP | A | 99 | 40.628 | 29.140 | 23.052 | 1.00 | 0.00 | C |

```
ATOM   1240  HA  TRP A  99      40.578  28.192  23.367  1.00  0.00           H
ATOM   1241  CB  TRP A  99      40.115  29.306  21.611  1.00  0.00           C
ATOM   1242  HB1 TRP A  99      39.140  29.082  21.596  1.00  0.00           H
ATOM   1243  HB2 TRP A  99      40.613  28.667  21.024  1.00  0.00           H
ATOM   1244  CG  TRP A  99      40.268  30.676  21.011  1.00  0.00           C
ATOM   1245  CD1 TRP A  99      39.298  31.614  20.914  1.00  0.00           C
ATOM   1246  HD1 TRP A  99      38.353  31.496  21.218  1.00  0.00           H
ATOM   1247  NE1 TRP A  99      39.808  32.759  20.332  1.00  0.00           N
ATOM   1248  HE1 TRP A  99      39.272  33.583  20.147  1.00  0.00           H
ATOM   1249  CE2 TRP A  99      41.149  32.624  20.038  1.00  0.00           C
ATOM   1250  CZ2 TRP A  99      42.113  33.489  19.497  1.00  0.00           C
ATOM   1251  HZ2 TRP A  99      41.868  34.427  19.253  1.00  0.00           H
ATOM   1252  CH2 TRP A  99      43.425  33.026  19.301  1.00  0.00           C
ATOM   1253  HH2 TRP A  99      44.122  33.637  18.926  1.00  0.00           H
ATOM   1254  CZ3 TRP A  99      43.756  31.701  19.634  1.00  0.00           C
ATOM   1255  HZ3 TRP A  99      44.686  31.367  19.481  1.00  0.00           H
ATOM   1256  CE3 TRP A  99      42.785  30.841  20.182  1.00  0.00           C
ATOM   1257  HE3 TRP A  99      43.035  29.900  20.409  1.00  0.00           H
ATOM   1258  CD2 TRP A  99      41.462  31.283  20.417  1.00  0.00           C
ATOM   1259  C   TRP A  99      42.114  29.548  23.145  1.00  0.00           C
ATOM   1260  O   TRP A  99      42.992  28.770  22.777  1.00  0.00           O
ATOM   1261  N   ASN A 100      42.411  30.760  23.630  1.00  0.00           N
ATOM   1262  H   ASN A 100      41.657  31.340  23.939  1.00  0.00           H
ATOM   1263  CA  ASN A 100      43.774  31.288  23.735  1.00  0.00           C
ATOM   1264  HA  ASN A 100      44.353  30.778  23.099  1.00  0.00           H
ATOM   1265  CB  ASN A 100      43.734  32.766  23.277  1.00  0.00           C
ATOM   1266  HB1 ASN A 100      43.376  33.317  24.031  1.00  0.00           H
ATOM   1267  HB2 ASN A 100      43.118  32.834  22.492  1.00  0.00           H
ATOM   1268  CG  ASN A 100      45.079  33.364  22.863  1.00  0.00           C
ATOM   1269  OD1 ASN A 100      45.916  32.725  22.231  1.00  0.00           O
ATOM   1270  ND2 ASN A 100      45.314  34.527  23.345  1.00  0.00           N
ATOM   1271 HD21 ASN A 100      44.641  34.967  23.939  1.00  0.00           H
ATOM   1272 HD22 ASN A 100      46.170  34.995  23.127  1.00  0.00           H
ATOM   1273  C   ASN A 100      44.426  30.999  25.122  1.00  0.00           C
ATOM   1274  O   ASN A 100      45.601  31.288  25.341  1.00  0.00           O
ATOM   1275  N   PHE A 101      43.741  30.349  26.062  1.00  0.00           N
ATOM   1276  H   PHE A 101      42.746  30.297  25.972  1.00  0.00           H
ATOM   1277  CA  PHE A 101      44.370  29.702  27.227  1.00  0.00           C
ATOM   1278  HA  PHE A 101      45.230  30.176  27.415  1.00  0.00           H
ATOM   1279  CB  PHE A 101      43.421  29.830  28.424  1.00  0.00           C
ATOM   1280  HB1 PHE A 101      42.612  29.269  28.247  1.00  0.00           H
ATOM   1281  HB2 PHE A 101      43.150  30.789  28.511  1.00  0.00           H
ATOM   1282  CG  PHE A 101      44.011  29.391  29.749  1.00  0.00           C
ATOM   1283  CD1 PHE A 101      43.465  28.325  30.476  1.00  0.00           C
ATOM   1284  HD1 PHE A 101      42.694  27.803  30.111  1.00  0.00           H
ATOM   1285  CE1 PHE A 101      44.017  28.003  31.723  1.00  0.00           C
ATOM   1286  HE1 PHE A 101      43.675  27.208  32.224  1.00  0.00           H
ATOM   1287  CZ  PHE A 101      45.046  28.785  32.279  1.00  0.00           C
ATOM   1288  HZ  PHE A 101      45.417  28.518  33.168  1.00  0.00           H
ATOM   1289  CE2 PHE A 101      45.562  29.887  31.667  1.00  0.00           C
ATOM   1290  HE2 PHE A 101      46.218  30.484  32.129  1.00  0.00           H
ATOM   1291  CD2 PHE A 101      45.114  30.153  30.313  1.00  0.00           C
ATOM   1292  HD2 PHE A 101      45.564  30.860  29.767  1.00  0.00           H
ATOM   1293  C   PHE A 101      44.691  28.235  26.921  1.00  0.00           C
```

| | | | | | | | | | | | |
|---|---|---|---|---|---|---|---|---|---|---|---|
| ATOM | 1294 | O | PHE | A | 101 | 44.202 | 27.647 | 25.962 | 1.00 | 0.00 | O |
| ATOM | 1295 | N | GLY | A | 102 | 45.401 | 27.603 | 27.847 | 1.00 | 0.00 | N |
| ATOM | 1296 | H | GLY | A | 102 | 45.939 | 28.149 | 28.489 | 1.00 | 0.00 | H |
| ATOM | 1297 | CA | GLY | A | 102 | 45.433 | 26.152 | 27.972 | 1.00 | 0.00 | C |
| ATOM | 1298 | HA1 | GLY | A | 102 | 45.545 | 25.760 | 27.059 | 1.00 | 0.00 | H |
| ATOM | 1299 | HA2 | GLY | A | 102 | 44.562 | 25.850 | 28.360 | 1.00 | 0.00 | H |
| ATOM | 1300 | C | GLY | A | 102 | 46.550 | 25.693 | 28.848 | 1.00 | 0.00 | C |
| ATOM | 1301 | O | GLY | A | 102 | 47.540 | 26.311 | 29.279 | 1.00 | 0.00 | O |
| ATOM | 1302 | N | THR | A | 103 | 46.413 | 24.413 | 29.241 | 1.00 | 0.00 | N |
| ATOM | 1303 | H | THR | A | 103 | 45.637 | 23.899 | 28.876 | 1.00 | 0.00 | H |
| ATOM | 1304 | CA | THR | A | 103 | 47.320 | 23.710 | 30.170 | 1.00 | 0.00 | C |
| ATOM | 1305 | HA | THR | A | 103 | 47.340 | 24.252 | 31.010 | 1.00 | 0.00 | H |
| ATOM | 1306 | CB | THR | A | 103 | 46.742 | 22.326 | 30.463 | 1.00 | 0.00 | C |
| ATOM | 1307 | HB | THR | A | 103 | 45.759 | 22.437 | 30.609 | 1.00 | 0.00 | H |
| ATOM | 1308 | CG2 | THR | A | 103 | 46.852 | 21.303 | 29.382 | 1.00 | 0.00 | C |
| ATOM | 1309 | HG21 | THR | A | 103 | 46.441 | 20.446 | 29.691 | 1.00 | 0.00 | H |
| ATOM | 1310 | HG22 | THR | A | 103 | 46.373 | 21.627 | 28.566 | 1.00 | 0.00 | H |
| ATOM | 1311 | HG23 | THR | A | 103 | 47.816 | 21.151 | 29.164 | 1.00 | 0.00 | H |
| ATOM | 1312 | OG1 | THR | A | 103 | 47.304 | 21.824 | 31.651 | 1.00 | 0.00 | O |
| ATOM | 1313 | HG1 | THR | A | 103 | 46.925 | 20.918 | 31.842 | 1.00 | 0.00 | H |
| ATOM | 1314 | C | THR | A | 103 | 48.777 | 23.650 | 29.658 | 1.00 | 0.00 | C |
| ATOM | 1315 | O | THR | A | 103 | 49.668 | 23.782 | 30.520 | 1.00 | 0.00 | O |
| ATOM | 1316 | N | GLN | A | 104 | 49.076 | 23.608 | 28.361 | 1.00 | 0.00 | N |
| ATOM | 1317 | H | GLN | A | 104 | 48.338 | 23.553 | 27.689 | 1.00 | 0.00 | H |
| ATOM | 1318 | CA | GLN | A | 104 | 50.461 | 23.641 | 27.903 | 1.00 | 0.00 | C |
| ATOM | 1319 | HA | GLN | A | 104 | 50.993 | 23.078 | 28.536 | 1.00 | 0.00 | H |
| ATOM | 1320 | CB | GLN | A | 104 | 50.519 | 22.949 | 26.510 | 1.00 | 0.00 | C |
| ATOM | 1321 | HB1 | GLN | A | 104 | 50.005 | 22.094 | 26.576 | 1.00 | 0.00 | H |
| ATOM | 1322 | HB2 | GLN | A | 104 | 51.478 | 22.747 | 26.313 | 1.00 | 0.00 | H |
| ATOM | 1323 | CG | GLN | A | 104 | 49.916 | 23.827 | 25.312 | 1.00 | 0.00 | C |
| ATOM | 1324 | HG1 | GLN | A | 104 | 50.097 | 23.402 | 24.425 | 1.00 | 0.00 | H |
| ATOM | 1325 | HG2 | GLN | A | 104 | 50.289 | 24.755 | 25.320 | 1.00 | 0.00 | H |
| ATOM | 1326 | CD | GLN | A | 104 | 48.413 | 23.888 | 25.542 | 1.00 | 0.00 | C |
| ATOM | 1327 | OE1 | GLN | A | 104 | 47.839 | 25.020 | 25.730 | 1.00 | 0.00 | O |
| ATOM | 1328 | NE2 | GLN | A | 104 | 47.703 | 22.782 | 25.736 | 1.00 | 0.00 | N |
| ATOM | 1329 | HE21 | GLN | A | 104 | 48.156 | 21.890 | 25.732 | 1.00 | 0.00 | H |
| ATOM | 1330 | HE22 | GLN | A | 104 | 46.716 | 22.841 | 25.886 | 1.00 | 0.00 | H |
| ATOM | 1331 | C | GLN | A | 104 | 51.128 | 25.064 | 28.053 | 1.00 | 0.00 | C |
| ATOM | 1332 | O | GLN | A | 104 | 52.231 | 25.125 | 28.625 | 1.00 | 0.00 | O |
| ATOM | 1333 | N | LYS | A | 105 | 50.382 | 26.205 | 27.774 | 1.00 | 0.00 | N |
| ATOM | 1334 | H | LYS | A | 105 | 49.501 | 26.077 | 27.319 | 1.00 | 0.00 | H |
| ATOM | 1335 | CA | LYS | A | 105 | 50.780 | 27.578 | 28.091 | 1.00 | 0.00 | C |
| ATOM | 1336 | HA | LYS | A | 105 | 51.696 | 27.574 | 27.690 | 1.00 | 0.00 | H |
| ATOM | 1337 | CB | LYS | A | 105 | 49.849 | 28.648 | 27.469 | 1.00 | 0.00 | C |
| ATOM | 1338 | HB1 | LYS | A | 105 | 50.213 | 29.550 | 27.702 | 1.00 | 0.00 | H |
| ATOM | 1339 | HB2 | LYS | A | 105 | 48.940 | 28.541 | 27.872 | 1.00 | 0.00 | H |
| ATOM | 1340 | CG | LYS | A | 105 | 49.668 | 28.623 | 25.971 | 1.00 | 0.00 | C |
| ATOM | 1341 | HG1 | LYS | A | 105 | 49.152 | 27.795 | 25.753 | 1.00 | 0.00 | H |
| ATOM | 1342 | HG2 | LYS | A | 105 | 50.581 | 28.562 | 25.568 | 1.00 | 0.00 | H |
| ATOM | 1343 | CD | LYS | A | 105 | 48.940 | 29.824 | 25.330 | 1.00 | 0.00 | C |
| ATOM | 1344 | HD1 | LYS | A | 105 | 49.501 | 30.646 | 25.426 | 1.00 | 0.00 | H |
| ATOM | 1345 | HD2 | LYS | A | 105 | 48.061 | 29.965 | 25.786 | 1.00 | 0.00 | H |
| ATOM | 1346 | CE | LYS | A | 105 | 48.698 | 29.552 | 23.846 | 1.00 | 0.00 | C |
| ATOM | 1347 | HE1 | LYS | A | 105 | 48.307 | 28.637 | 23.740 | 1.00 | 0.00 | H |

```
ATOM   1348  HE2 LYS A 105      49.569  29.601  23.358  1.00  0.00           H
ATOM   1349  NZ  LYS A 105      47.740  30.569  23.246  1.00  0.00           N
ATOM   1350  HZ1 LYS A 105      47.602  30.365  22.277  1.00  0.00           H
ATOM   1351  HZ2 LYS A 105      46.863  30.524  23.724  1.00  0.00           H
ATOM   1352  HZ3 LYS A 105      48.125  31.487  23.342  1.00  0.00           H
ATOM   1353  C   LYS A 105      50.836  27.887  29.587  1.00  0.00           C
ATOM   1354  O   LYS A 105      51.803  28.514  30.013  1.00  0.00           O
ATOM   1355  N   PHE A 106      49.917  27.405  30.412  1.00  0.00           N
ATOM   1356  H   PHE A 106      49.142  26.914  30.013  1.00  0.00           H
ATOM   1357  CA  PHE A 106      49.963  27.546  31.882  1.00  0.00           C
ATOM   1358  HA  PHE A 106      50.179  28.501  32.087  1.00  0.00           H
ATOM   1359  CB  PHE A 106      48.593  27.219  32.524  1.00  0.00           C
ATOM   1360  HB1 PHE A 106      48.430  26.233  32.489  1.00  0.00           H
ATOM   1361  HB2 PHE A 106      47.866  27.695  32.029  1.00  0.00           H
ATOM   1362  CG  PHE A 106      48.515  27.633  33.935  1.00  0.00           C
ATOM   1363  CD1 PHE A 106      47.938  28.926  34.239  1.00  0.00           C
ATOM   1364  HD1 PHE A 106      47.663  29.555  33.512  1.00  0.00           H
ATOM   1365  CE1 PHE A 106      47.781  29.259  35.579  1.00  0.00           C
ATOM   1366  HE1 PHE A 106      47.267  30.080  35.826  1.00  0.00           H
ATOM   1367  CZ  PHE A 106      48.331  28.463  36.586  1.00  0.00           C
ATOM   1368  HZ  PHE A 106      48.312  28.794  37.529  1.00  0.00           H
ATOM   1369  CE2 PHE A 106      48.903  27.228  36.320  1.00  0.00           C
ATOM   1370  HE2 PHE A 106      49.178  26.614  37.060  1.00  0.00           H
ATOM   1371  CD2 PHE A 106      49.089  26.857  34.919  1.00  0.00           C
ATOM   1372  HD2 PHE A 106      49.627  26.049  34.677  1.00  0.00           H
ATOM   1373  C   PHE A 106      51.087  26.722  32.433  1.00  0.00           C
ATOM   1374  O   PHE A 106      51.741  27.200  33.422  1.00  0.00           O
ATOM   1375  N   THR A 107      51.490  25.655  31.709  1.00  0.00           N
ATOM   1376  H   THR A 107      50.944  25.321  30.941  1.00  0.00           H
ATOM   1377  CA  THR A 107      52.781  24.983  32.088  1.00  0.00           C
ATOM   1378  HA  THR A 107      52.855  25.078  33.081  1.00  0.00           H
ATOM   1379  CB  THR A 107      52.816  23.445  31.763  1.00  0.00           C
ATOM   1380  HB  THR A 107      52.965  23.365  30.777  1.00  0.00           H
ATOM   1381  CG2 THR A 107      53.830  22.640  32.512  1.00  0.00           C
ATOM   1382 HG21 THR A 107      53.768  21.682  32.232  1.00  0.00           H
ATOM   1383 HG22 THR A 107      54.746  22.987  32.311  1.00  0.00           H
ATOM   1384 HG23 THR A 107      53.654  22.713  33.494  1.00  0.00           H
ATOM   1385  OG1 THR A 107      51.609  22.834  32.194  1.00  0.00           O
ATOM   1386  HG1 THR A 107      51.637  21.857  31.985  1.00  0.00           H
ATOM   1387  C   THR A 107      53.946  25.732  31.564  1.00  0.00           C
ATOM   1388  O   THR A 107      54.931  25.649  32.267  1.00  0.00           O
ATOM   1389  N   ASP A 108      53.906  26.416  30.406  1.00  0.00           N
ATOM   1390  H   ASP A 108      53.086  26.411  29.833  1.00  0.00           H
ATOM   1391  CA  ASP A 108      55.107  27.173  30.006  1.00  0.00           C
ATOM   1392  HA  ASP A 108      55.771  26.425  30.005  1.00  0.00           H
ATOM   1393  CB  ASP A 108      55.007  27.847  28.646  1.00  0.00           C
ATOM   1394  HB1 ASP A 108      55.804  28.443  28.546  1.00  0.00           H
ATOM   1395  HB2 ASP A 108      54.174  28.401  28.644  1.00  0.00           H
ATOM   1396  CG  ASP A 108      54.959  26.863  27.433  1.00  0.00           C
ATOM   1397  OD1 ASP A 108      54.506  27.336  26.397  1.00  0.00           O
ATOM   1398  OD2 ASP A 108      55.416  25.728  27.487  1.00  0.00           O
ATOM   1399  C   ASP A 108      55.468  28.315  30.968  1.00  0.00           C
ATOM   1400  O   ASP A 108      56.639  28.650  31.139  1.00  0.00           O
ATOM   1401  N   MET A 109      54.383  28.844  31.623  1.00  0.00           N
```

| | | | | | | | | | | | |
|---|---|---|---|---|---|---|---|---|---|---|---|
| ATOM | 1402 | H | MET | A | 109 | 53.487 | 28.412 | 31.515 | 1.00 | 0.00 | H |
| ATOM | 1403 | CA | MET | A | 109 | 54.508 | 30.007 | 32.462 | 1.00 | 0.00 | C |
| ATOM | 1404 | HA | MET | A | 109 | 55.303 | 30.502 | 32.111 | 1.00 | 0.00 | H |
| ATOM | 1405 | CB | MET | A | 109 | 53.205 | 30.851 | 32.304 | 1.00 | 0.00 | C |
| ATOM | 1406 | HB1 | MET | A | 109 | 52.423 | 30.236 | 32.403 | 1.00 | 0.00 | H |
| ATOM | 1407 | HB2 | MET | A | 109 | 53.203 | 31.249 | 31.387 | 1.00 | 0.00 | H |
| ATOM | 1408 | CG | MET | A | 109 | 53.019 | 32.005 | 33.312 | 1.00 | 0.00 | C |
| ATOM | 1409 | HG1 | MET | A | 109 | 52.352 | 32.596 | 32.858 | 1.00 | 0.00 | H |
| ATOM | 1410 | HG2 | MET | A | 109 | 53.914 | 32.451 | 33.312 | 1.00 | 0.00 | H |
| ATOM | 1411 | SD | MET | A | 109 | 52.478 | 31.546 | 35.026 | 1.00 | 0.00 | S |
| ATOM | 1412 | CE | MET | A | 109 | 50.981 | 30.580 | 34.790 | 1.00 | 0.00 | C |
| ATOM | 1413 | HE1 | MET | A | 109 | 50.626 | 30.293 | 35.680 | 1.00 | 0.00 | H |
| ATOM | 1414 | HE2 | MET | A | 109 | 50.295 | 31.137 | 34.321 | 1.00 | 0.00 | H |
| ATOM | 1415 | HE3 | MET | A | 109 | 51.189 | 29.773 | 34.238 | 1.00 | 0.00 | H |
| ATOM | 1416 | C | MET | A | 109 | 54.939 | 29.705 | 33.899 | 1.00 | 0.00 | C |
| ATOM | 1417 | O | MET | A | 109 | 55.692 | 30.413 | 34.544 | 1.00 | 0.00 | O |
| ATOM | 1418 | N | VAL | A | 110 | 54.309 | 28.676 | 34.424 | 1.00 | 0.00 | N |
| ATOM | 1419 | H | VAL | A | 110 | 53.865 | 28.015 | 33.819 | 1.00 | 0.00 | H |
| ATOM | 1420 | CA | VAL | A | 110 | 54.242 | 28.476 | 35.860 | 1.00 | 0.00 | C |
| ATOM | 1421 | HA | VAL | A | 110 | 54.300 | 29.385 | 36.273 | 1.00 | 0.00 | H |
| ATOM | 1422 | CB | VAL | A | 110 | 52.853 | 27.838 | 36.203 | 1.00 | 0.00 | C |
| ATOM | 1423 | HB | VAL | A | 110 | 52.244 | 28.069 | 35.445 | 1.00 | 0.00 | H |
| ATOM | 1424 | CG1 | VAL | A | 110 | 52.770 | 26.281 | 36.249 | 1.00 | 0.00 | C |
| ATOM | 1425 | HG11 | VAL | A | 110 | 51.837 | 26.004 | 36.477 | 1.00 | 0.00 | H |
| ATOM | 1426 | HG12 | VAL | A | 110 | 53.020 | 25.907 | 35.356 | 1.00 | 0.00 | H |
| ATOM | 1427 | HG13 | VAL | A | 110 | 53.400 | 25.934 | 36.944 | 1.00 | 0.00 | H |
| ATOM | 1428 | CG2 | VAL | A | 110 | 52.442 | 28.417 | 37.614 | 1.00 | 0.00 | C |
| ATOM | 1429 | HG21 | VAL | A | 110 | 51.556 | 28.040 | 37.884 | 1.00 | 0.00 | H |
| ATOM | 1430 | HG22 | VAL | A | 110 | 53.132 | 28.164 | 38.292 | 1.00 | 0.00 | H |
| ATOM | 1431 | HG23 | VAL | A | 110 | 52.378 | 29.413 | 37.560 | 1.00 | 0.00 | H |
| ATOM | 1432 | C | VAL | A | 110 | 55.489 | 27.747 | 36.423 | 1.00 | 0.00 | C |
| ATOM | 1433 | O | VAL | A | 110 | 55.846 | 27.947 | 37.611 | 1.00 | 0.00 | O |
| ATOM | 1434 | N | ALA | A | 111 | 56.228 | 26.970 | 35.652 | 1.00 | 0.00 | N |
| ATOM | 1435 | H | ALA | A | 111 | 55.926 | 26.870 | 34.704 | 1.00 | 0.00 | H |
| ATOM | 1436 | CA | ALA | A | 111 | 57.446 | 26.235 | 36.031 | 1.00 | 0.00 | C |
| ATOM | 1437 | HA | ALA | A | 111 | 57.311 | 26.078 | 37.009 | 1.00 | 0.00 | H |
| ATOM | 1438 | CB | ALA | A | 111 | 57.452 | 24.994 | 35.149 | 1.00 | 0.00 | C |
| ATOM | 1439 | HB1 | ALA | A | 111 | 58.260 | 24.443 | 35.355 | 1.00 | 0.00 | H |
| ATOM | 1440 | HB2 | ALA | A | 111 | 56.628 | 24.456 | 35.325 | 1.00 | 0.00 | H |
| ATOM | 1441 | HB3 | ALA | A | 111 | 57.472 | 25.268 | 34.188 | 1.00 | 0.00 | H |
| ATOM | 1442 | C | ALA | A | 111 | 58.829 | 26.940 | 36.056 | 1.00 | 0.00 | C |
| ATOM | 1443 | O | ALA | A | 111 | 59.805 | 26.260 | 36.342 | 1.00 | 0.00 | O |
| ATOM | 1444 | N | THR | A | 112 | 58.891 | 28.232 | 35.682 | 1.00 | 0.00 | N |
| ATOM | 1445 | H | THR | A | 112 | 58.034 | 28.710 | 35.488 | 1.00 | 0.00 | H |
| ATOM | 1446 | CA | THR | A | 112 | 60.139 | 28.972 | 35.544 | 1.00 | 0.00 | C |
| ATOM | 1447 | HA | THR | A | 112 | 60.819 | 28.440 | 36.049 | 1.00 | 0.00 | H |
| ATOM | 1448 | CB | THR | A | 112 | 60.565 | 29.135 | 34.071 | 1.00 | 0.00 | C |
| ATOM | 1449 | HB | THR | A | 112 | 59.831 | 29.533 | 33.521 | 1.00 | 0.00 | H |
| ATOM | 1450 | CG2 | THR | A | 112 | 61.814 | 29.951 | 33.873 | 1.00 | 0.00 | C |
| ATOM | 1451 | HG21 | THR | A | 112 | 62.022 | 30.010 | 32.897 | 1.00 | 0.00 | H |
| ATOM | 1452 | HG22 | THR | A | 112 | 61.673 | 30.871 | 34.240 | 1.00 | 0.00 | H |
| ATOM | 1453 | HG23 | THR | A | 112 | 62.576 | 29.515 | 34.351 | 1.00 | 0.00 | H |
| ATOM | 1454 | OG1 | THR | A | 112 | 60.912 | 27.848 | 33.672 | 1.00 | 0.00 | O |
| ATOM | 1455 | HG1 | THR | A | 112 | 61.204 | 27.863 | 32.716 | 1.00 | 0.00 | H |

| | | | | | | | | | | | |
|---|---|---|---|---|---|---|---|---|---|---|---|
| ATOM | 1456 | C | THR | A | 112 | 59.901 | 30.378 | 36.091 | 1.00 | 0.00 | C |
| ATOM | 1457 | O | THR | A | 112 | 59.131 | 31.143 | 35.556 | 1.00 | 0.00 | O |
| ATOM | 1458 | N | ALA | A | 113 | 60.668 | 30.729 | 37.125 | 1.00 | 0.00 | N |
| ATOM | 1459 | H | ALA | A | 113 | 61.386 | 30.087 | 37.394 | 1.00 | 0.00 | H |
| ATOM | 1460 | CA | ALA | A | 113 | 60.557 | 31.988 | 37.918 | 1.00 | 0.00 | C |
| ATOM | 1461 | HA | ALA | A | 113 | 59.757 | 31.918 | 38.514 | 1.00 | 0.00 | H |
| ATOM | 1462 | CB | ALA | A | 113 | 61.844 | 32.165 | 38.710 | 1.00 | 0.00 | C |
| ATOM | 1463 | HB1 | ALA | A | 113 | 61.790 | 33.005 | 39.250 | 1.00 | 0.00 | H |
| ATOM | 1464 | HB2 | ALA | A | 113 | 61.969 | 31.384 | 39.321 | 1.00 | 0.00 | H |
| ATOM | 1465 | HB3 | ALA | A | 113 | 62.618 | 32.223 | 38.080 | 1.00 | 0.00 | H |
| ATOM | 1466 | C | ALA | A | 113 | 60.362 | 33.175 | 37.023 | 1.00 | 0.00 | C |
| ATOM | 1467 | O | ALA | A | 113 | 59.386 | 33.872 | 37.224 | 1.00 | 0.00 | O |
| ATOM | 1468 | N | ASN | A | 114 | 61.242 | 33.512 | 36.044 | 1.00 | 0.00 | N |
| ATOM | 1469 | H | ASN | A | 114 | 62.050 | 32.938 | 35.910 | 1.00 | 0.00 | H |
| ATOM | 1470 | CA | ASN | A | 114 | 61.064 | 34.667 | 35.183 | 1.00 | 0.00 | C |
| ATOM | 1471 | HA | ASN | A | 114 | 61.063 | 35.406 | 35.856 | 1.00 | 0.00 | H |
| ATOM | 1472 | CB | ASN | A | 114 | 62.231 | 34.824 | 34.180 | 1.00 | 0.00 | C |
| ATOM | 1473 | HB1 | ASN | A | 114 | 62.184 | 34.093 | 33.499 | 1.00 | 0.00 | H |
| ATOM | 1474 | HB2 | ASN | A | 114 | 63.100 | 34.766 | 34.671 | 1.00 | 0.00 | H |
| ATOM | 1475 | CG | ASN | A | 114 | 62.147 | 36.180 | 33.467 | 1.00 | 0.00 | C |
| ATOM | 1476 | OD1 | ASN | A | 114 | 62.289 | 37.300 | 34.191 | 1.00 | 0.00 | O |
| ATOM | 1477 | ND2 | ASN | A | 114 | 62.088 | 36.316 | 32.232 | 1.00 | 0.00 | N |
| ATOM | 1478 | HD21 | ASN | A | 114 | 62.094 | 35.510 | 31.639 | 1.00 | 0.00 | H |
| ATOM | 1479 | HD22 | ASN | A | 114 | 62.035 | 37.231 | 31.832 | 1.00 | 0.00 | H |
| ATOM | 1480 | C | ASN | A | 114 | 59.720 | 34.575 | 34.332 | 1.00 | 0.00 | C |
| ATOM | 1481 | O | ASN | A | 114 | 59.100 | 35.604 | 34.072 | 1.00 | 0.00 | O |
| ATOM | 1482 | N | ASN | A | 115 | 59.241 | 33.446 | 33.858 | 1.00 | 0.00 | N |
| ATOM | 1483 | H | ASN | A | 115 | 59.760 | 32.613 | 34.049 | 1.00 | 0.00 | H |
| ATOM | 1484 | CA | ASN | A | 115 | 58.014 | 33.304 | 33.075 | 1.00 | 0.00 | C |
| ATOM | 1485 | HA | ASN | A | 115 | 58.034 | 33.985 | 32.343 | 1.00 | 0.00 | H |
| ATOM | 1486 | CB | ASN | A | 115 | 57.908 | 31.930 | 32.420 | 1.00 | 0.00 | C |
| ATOM | 1487 | HB1 | ASN | A | 115 | 57.121 | 31.965 | 31.804 | 1.00 | 0.00 | H |
| ATOM | 1488 | HB2 | ASN | A | 115 | 57.729 | 31.275 | 33.154 | 1.00 | 0.00 | H |
| ATOM | 1489 | CG | ASN | A | 115 | 59.136 | 31.413 | 31.602 | 1.00 | 0.00 | C |
| ATOM | 1490 | OD1 | ASN | A | 115 | 60.287 | 31.874 | 31.675 | 1.00 | 0.00 | O |
| ATOM | 1491 | ND2 | ASN | A | 115 | 58.886 | 30.347 | 30.905 | 1.00 | 0.00 | N |
| ATOM | 1492 | HD21 | ASN | A | 115 | 57.974 | 29.937 | 30.927 | 1.00 | 0.00 | H |
| ATOM | 1493 | HD22 | ASN | A | 115 | 59.605 | 29.935 | 30.346 | 1.00 | 0.00 | H |
| ATOM | 1494 | C | ASN | A | 115 | 56.862 | 33.502 | 34.057 | 1.00 | 0.00 | C |
| ATOM | 1495 | O | ASN | A | 115 | 55.933 | 34.194 | 33.780 | 1.00 | 0.00 | O |
| ATOM | 1496 | N | ARG | A | 116 | 56.920 | 32.893 | 35.257 | 1.00 | 0.00 | N |
| ATOM | 1497 | H | ARG | A | 116 | 57.727 | 32.349 | 35.487 | 1.00 | 0.00 | H |
| ATOM | 1498 | CA | ARG | A | 116 | 55.851 | 33.004 | 36.220 | 1.00 | 0.00 | C |
| ATOM | 1499 | HA | ARG | A | 116 | 55.022 | 32.684 | 35.762 | 1.00 | 0.00 | H |
| ATOM | 1500 | CB | ARG | A | 116 | 56.187 | 32.159 | 37.398 | 1.00 | 0.00 | C |
| ATOM | 1501 | HB1 | ARG | A | 116 | 56.867 | 32.649 | 37.943 | 1.00 | 0.00 | H |
| ATOM | 1502 | HB2 | ARG | A | 116 | 56.583 | 31.305 | 37.060 | 1.00 | 0.00 | H |
| ATOM | 1503 | CG | ARG | A | 116 | 55.036 | 31.816 | 38.290 | 1.00 | 0.00 | C |
| ATOM | 1504 | HG1 | ARG | A | 116 | 54.367 | 31.300 | 37.755 | 1.00 | 0.00 | H |
| ATOM | 1505 | HG2 | ARG | A | 116 | 54.622 | 32.668 | 38.610 | 1.00 | 0.00 | H |
| ATOM | 1506 | CD | ARG | A | 116 | 55.430 | 30.994 | 39.488 | 1.00 | 0.00 | C |
| ATOM | 1507 | HD1 | ARG | A | 116 | 56.013 | 30.237 | 39.195 | 1.00 | 0.00 | H |
| ATOM | 1508 | HD2 | ARG | A | 116 | 54.609 | 30.631 | 39.928 | 1.00 | 0.00 | H |
| ATOM | 1509 | NE | ARG | A | 116 | 56.159 | 31.749 | 40.479 | 1.00 | 0.00 | N |

```
ATOM   1510  HE  ARG A 116      55.708  32.583  40.796  1.00  0.00           H
ATOM   1511  CZ  ARG A 116      57.305  31.507  41.016  1.00  0.00           C
ATOM   1512  NH1 ARG A 116      57.867  30.378  40.765  1.00  0.00           N
ATOM   1513 HH11 ARG A 116      57.414  29.716  40.167  1.00  0.00           H
ATOM   1514 HH12 ARG A 116      58.757  30.162  41.167  1.00  0.00           H
ATOM   1515  NH2 ARG A 116      57.910  32.523  41.652  1.00  0.00           N
ATOM   1516 HH21 ARG A 116      57.470  33.420  41.694  1.00  0.00           H
ATOM   1517 HH22 ARG A 116      58.801  32.383  42.084  1.00  0.00           H
ATOM   1518  C   ARG A 116      55.754  34.465  36.728  1.00  0.00           C
ATOM   1519  O   ARG A 116      54.618  34.906  36.873  1.00  0.00           O
ATOM   1520  N   GLN A 117      56.853  35.218  36.805  1.00  0.00           N
ATOM   1521  H   GLN A 117      57.738  34.862  36.506  1.00  0.00           H
ATOM   1522  CA  GLN A 117      56.735  36.566  37.332  1.00  0.00           C
ATOM   1523  HA  GLN A 117      55.864  36.594  37.823  1.00  0.00           H
ATOM   1524  CB  GLN A 117      57.983  36.770  38.199  1.00  0.00           C
ATOM   1525  HB1 GLN A 117      58.132  37.754  38.291  1.00  0.00           H
ATOM   1526  HB2 GLN A 117      58.758  36.359  37.719  1.00  0.00           H
ATOM   1527  CG  GLN A 117      57.854  36.120  39.644  1.00  0.00           C
ATOM   1528  HG1 GLN A 117      58.776  35.955  39.995  1.00  0.00           H
ATOM   1529  HG2 GLN A 117      57.371  35.249  39.552  1.00  0.00           H
ATOM   1530  CD  GLN A 117      57.093  36.961  40.715  1.00  0.00           C
ATOM   1531  OE1 GLN A 117      57.189  38.204  40.770  1.00  0.00           O
ATOM   1532  NE2 GLN A 117      56.223  36.372  41.532  1.00  0.00           N
ATOM   1533 HE21 GLN A 117      56.056  35.389  41.460  1.00  0.00           H
ATOM   1534 HE22 GLN A 117      55.736  36.913  42.218  1.00  0.00           H
ATOM   1535  C   GLN A 117      56.595  37.671  36.301  1.00  0.00           C
ATOM   1536  O   GLN A 117      56.093  38.719  36.596  1.00  0.00           O
ATOM   1537  N   THR A 118      56.797  37.359  35.012  1.00  0.00           N
ATOM   1538  H   THR A 118      57.345  36.548  34.808  1.00  0.00           H
ATOM   1539  CA  THR A 118      56.265  38.132  33.880  1.00  0.00           C
ATOM   1540  HA  THR A 118      56.629  39.060  33.956  1.00  0.00           H
ATOM   1541  CB  THR A 118      56.824  37.548  32.607  1.00  0.00           C
ATOM   1542  HB  THR A 118      56.677  36.560  32.648  1.00  0.00           H
ATOM   1543  CG2 THR A 118      56.156  38.036  31.324  1.00  0.00           C
ATOM   1544 HG21 THR A 118      56.589  37.599  30.536  1.00  0.00           H
ATOM   1545 HG22 THR A 118      55.184  37.801  31.344  1.00  0.00           H
ATOM   1546 HG23 THR A 118      56.257  39.028  31.252  1.00  0.00           H
ATOM   1547  OG1 THR A 118      58.246  37.791  32.562  1.00  0.00           O
ATOM   1548  HG1 THR A 118      58.623  37.406  31.720  1.00  0.00           H
ATOM   1549  C   THR A 118      54.667  38.194  33.915  1.00  0.00           C
ATOM   1550  O   THR A 118      54.079  39.281  33.804  1.00  0.00           O
ATOM   1551  N   PHE A 119      54.038  37.076  34.250  1.00  0.00           N
ATOM   1552  H   PHE A 119      54.594  36.263  34.421  1.00  0.00           H
ATOM   1553  CA  PHE A 119      52.582  36.948  34.388  1.00  0.00           C
ATOM   1554  HA  PHE A 119      52.263  37.248  33.489  1.00  0.00           H
ATOM   1555  CB  PHE A 119      52.192  35.480  34.734  1.00  0.00           C
ATOM   1556  HB1 PHE A 119      52.223  35.380  35.728  1.00  0.00           H
ATOM   1557  HB2 PHE A 119      52.870  34.875  34.317  1.00  0.00           H
ATOM   1558  CG  PHE A 119      50.836  34.973  34.293  1.00  0.00           C
ATOM   1559  CD1 PHE A 119      50.555  34.876  32.948  1.00  0.00           C
ATOM   1560  HD1 PHE A 119      51.255  35.122  32.278  1.00  0.00           H
ATOM   1561  CE1 PHE A 119      49.316  34.442  32.515  1.00  0.00           C
ATOM   1562  HE1 PHE A 119      49.155  34.358  31.532  1.00  0.00           H
ATOM   1563  CZ  PHE A 119      48.240  34.103  33.427  1.00  0.00           C
```

| | | | | | | | | | | | |
|---|---|---|---|---|---|---|---|---|---|---|---|
| ATOM | 1564 | HZ | PHE | A | 119 | 47.364 | 33.744 | 33.107 | 1.00 | 0.00 | H |
| ATOM | 1565 | CE2 | PHE | A | 119 | 48.509 | 34.312 | 34.787 | 1.00 | 0.00 | C |
| ATOM | 1566 | HE2 | PHE | A | 119 | 47.793 | 34.143 | 35.464 | 1.00 | 0.00 | H |
| ATOM | 1567 | CD2 | PHE | A | 119 | 49.783 | 34.760 | 35.212 | 1.00 | 0.00 | C |
| ATOM | 1568 | HD2 | PHE | A | 119 | 49.942 | 34.930 | 36.185 | 1.00 | 0.00 | H |
| ATOM | 1569 | C | PHE | A | 119 | 51.970 | 37.861 | 35.462 | 1.00 | 0.00 | C |
| ATOM | 1570 | O | PHE | A | 119 | 50.990 | 38.541 | 35.153 | 1.00 | 0.00 | O |
| ATOM | 1571 | N | VAL | A | 120 | 52.356 | 37.783 | 36.759 | 1.00 | 0.00 | N |
| ATOM | 1572 | H | VAL | A | 120 | 53.091 | 37.143 | 36.983 | 1.00 | 0.00 | H |
| ATOM | 1573 | CA | VAL | A | 120 | 51.783 | 38.560 | 37.857 | 1.00 | 0.00 | C |
| ATOM | 1574 | HA | VAL | A | 120 | 50.787 | 38.487 | 37.806 | 1.00 | 0.00 | H |
| ATOM | 1575 | CB | VAL | A | 120 | 52.246 | 37.913 | 39.188 | 1.00 | 0.00 | C |
| ATOM | 1576 | HB | VAL | A | 120 | 51.717 | 38.381 | 39.896 | 1.00 | 0.00 | H |
| ATOM | 1577 | CG1 | VAL | A | 120 | 52.090 | 36.394 | 39.025 | 1.00 | 0.00 | C |
| ATOM | 1578 | HG11 | VAL | A | 120 | 52.380 | 35.937 | 39.866 | 1.00 | 0.00 | H |
| ATOM | 1579 | HG12 | VAL | A | 120 | 51.132 | 36.175 | 38.840 | 1.00 | 0.00 | H |
| ATOM | 1580 | HG13 | VAL | A | 120 | 52.657 | 36.081 | 38.263 | 1.00 | 0.00 | H |
| ATOM | 1581 | CG2 | VAL | A | 120 | 53.630 | 38.109 | 39.658 | 1.00 | 0.00 | C |
| ATOM | 1582 | HG21 | VAL | A | 120 | 53.759 | 37.628 | 40.525 | 1.00 | 0.00 | H |
| ATOM | 1583 | HG22 | VAL | A | 120 | 54.267 | 37.747 | 38.978 | 1.00 | 0.00 | H |
| ATOM | 1584 | HG23 | VAL | A | 120 | 53.802 | 39.085 | 39.790 | 1.00 | 0.00 | H |
| ATOM | 1585 | C | VAL | A | 120 | 52.024 | 40.078 | 37.690 | 1.00 | 0.00 | C |
| ATOM | 1586 | O | VAL | A | 120 | 51.121 | 40.853 | 37.999 | 1.00 | 0.00 | O |
| ATOM | 1587 | N | ASN | A | 121 | 53.194 | 40.452 | 37.161 | 1.00 | 0.00 | N |
| ATOM | 1588 | H | ASN | A | 121 | 53.891 | 39.749 | 37.020 | 1.00 | 0.00 | H |
| ATOM | 1589 | CA | ASN | A | 121 | 53.506 | 41.786 | 36.788 | 1.00 | 0.00 | C |
| ATOM | 1590 | HA | ASN | A | 121 | 53.243 | 42.288 | 37.612 | 1.00 | 0.00 | H |
| ATOM | 1591 | CB | ASN | A | 121 | 54.985 | 41.906 | 36.492 | 1.00 | 0.00 | C |
| ATOM | 1592 | HB1 | ASN | A | 121 | 55.177 | 42.767 | 36.021 | 1.00 | 0.00 | H |
| ATOM | 1593 | HB2 | ASN | A | 121 | 55.290 | 41.141 | 35.925 | 1.00 | 0.00 | H |
| ATOM | 1594 | CG | ASN | A | 121 | 55.679 | 41.875 | 37.833 | 1.00 | 0.00 | C |
| ATOM | 1595 | OD1 | ASN | A | 121 | 55.435 | 42.706 | 38.708 | 1.00 | 0.00 | O |
| ATOM | 1596 | ND2 | ASN | A | 121 | 56.536 | 40.951 | 38.094 | 1.00 | 0.00 | N |
| ATOM | 1597 | HD21 | ASN | A | 121 | 56.743 | 40.253 | 37.409 | 1.00 | 0.00 | H |
| ATOM | 1598 | HD22 | ASN | A | 121 | 56.995 | 40.931 | 38.982 | 1.00 | 0.00 | H |
| ATOM | 1599 | C | ASN | A | 121 | 52.703 | 42.304 | 35.570 | 1.00 | 0.00 | C |
| ATOM | 1600 | O | ASN | A | 121 | 52.325 | 43.484 | 35.586 | 1.00 | 0.00 | O |
| ATOM | 1601 | N | SER | A | 122 | 52.307 | 41.364 | 34.714 | 1.00 | 0.00 | N |
| ATOM | 1602 | H | SER | A | 122 | 52.657 | 40.436 | 34.842 | 1.00 | 0.00 | H |
| ATOM | 1603 | CA | SER | A | 122 | 51.401 | 41.603 | 33.609 | 1.00 | 0.00 | C |
| ATOM | 1604 | HA | SER | A | 122 | 51.727 | 42.444 | 33.177 | 1.00 | 0.00 | H |
| ATOM | 1605 | CB | SER | A | 122 | 51.489 | 40.405 | 32.650 | 1.00 | 0.00 | C |
| ATOM | 1606 | HB1 | SER | A | 122 | 51.427 | 39.537 | 33.143 | 1.00 | 0.00 | H |
| ATOM | 1607 | HB2 | SER | A | 122 | 52.334 | 40.426 | 32.116 | 1.00 | 0.00 | H |
| ATOM | 1608 | OG | SER | A | 122 | 50.379 | 40.502 | 31.754 | 1.00 | 0.00 | O |
| ATOM | 1609 | HG | SER | A | 122 | 50.402 | 39.736 | 31.111 | 1.00 | 0.00 | H |
| ATOM | 1610 | C | SER | A | 122 | 49.915 | 41.772 | 34.115 | 1.00 | 0.00 | C |
| ATOM | 1611 | O | SER | A | 122 | 49.255 | 42.795 | 33.826 | 1.00 | 0.00 | O |
| ATOM | 1612 | N | ALA | A | 123 | 49.458 | 40.882 | 34.987 | 1.00 | 0.00 | N |
| ATOM | 1613 | H | ALA | A | 123 | 50.056 | 40.129 | 35.263 | 1.00 | 0.00 | H |
| ATOM | 1614 | CA | ALA | A | 123 | 48.084 | 40.961 | 35.573 | 1.00 | 0.00 | C |
| ATOM | 1615 | HA | ALA | A | 123 | 47.442 | 40.883 | 34.810 | 1.00 | 0.00 | H |
| ATOM | 1616 | CB | ALA | A | 123 | 48.023 | 39.845 | 36.625 | 1.00 | 0.00 | C |
| ATOM | 1617 | HB1 | ALA | A | 123 | 47.122 | 39.844 | 37.058 | 1.00 | 0.00 | H |

| | | | | | | | | | | | |
|---|---|---|---|---|---|---|---|---|---|---|---|
| ATOM | 1618 | HB2 | ALA | A | 123 | 48.182 | 38.962 | 36.184 | 1.00 | 0.00 | H |
| ATOM | 1619 | HB3 | ALA | A | 123 | 48.726 | 40.002 | 37.319 | 1.00 | 0.00 | H |
| ATOM | 1620 | C | ALA | A | 123 | 47.700 | 42.269 | 36.210 | 1.00 | 0.00 | C |
| ATOM | 1621 | O | ALA | A | 123 | 46.694 | 42.901 | 35.879 | 1.00 | 0.00 | O |
| ATOM | 1622 | N | ILE | A | 124 | 48.544 | 42.817 | 37.051 | 1.00 | 0.00 | N |
| ATOM | 1623 | H | ILE | A | 124 | 49.417 | 42.366 | 37.236 | 1.00 | 0.00 | H |
| ATOM | 1624 | CA | ILE | A | 124 | 48.230 | 44.075 | 37.721 | 1.00 | 0.00 | C |
| ATOM | 1625 | HA | ILE | A | 124 | 47.345 | 43.861 | 38.135 | 1.00 | 0.00 | H |
| ATOM | 1626 | CB | ILE | A | 124 | 49.138 | 44.450 | 38.856 | 1.00 | 0.00 | C |
| ATOM | 1627 | HB | ILE | A | 124 | 48.927 | 45.351 | 39.235 | 1.00 | 0.00 | H |
| ATOM | 1628 | CG2 | ILE | A | 124 | 48.878 | 43.506 | 40.002 | 1.00 | 0.00 | C |
| ATOM | 1629 | HG21 | ILE | A | 124 | 49.475 | 43.741 | 40.769 | 1.00 | 0.00 | H |
| ATOM | 1630 | HG22 | ILE | A | 124 | 47.923 | 43.583 | 40.287 | 1.00 | 0.00 | H |
| ATOM | 1631 | HG23 | ILE | A | 124 | 49.064 | 42.568 | 39.710 | 1.00 | 0.00 | H |
| ATOM | 1632 | CG1 | ILE | A | 124 | 50.576 | 44.615 | 38.359 | 1.00 | 0.00 | C |
| ATOM | 1633 | HG11 | ILE | A | 124 | 50.556 | 45.185 | 37.538 | 1.00 | 0.00 | H |
| ATOM | 1634 | HG12 | ILE | A | 124 | 50.929 | 43.709 | 38.127 | 1.00 | 0.00 | H |
| ATOM | 1635 | CD | ILE | A | 124 | 51.537 | 45.262 | 39.373 | 1.00 | 0.00 | C |
| ATOM | 1636 | HD1 | ILE | A | 124 | 52.449 | 45.332 | 38.970 | 1.00 | 0.00 | H |
| ATOM | 1637 | HD2 | ILE | A | 124 | 51.206 | 46.176 | 39.609 | 1.00 | 0.00 | H |
| ATOM | 1638 | HD3 | ILE | A | 124 | 51.579 | 44.699 | 40.198 | 1.00 | 0.00 | H |
| ATOM | 1639 | C | ILE | A | 124 | 48.195 | 45.245 | 36.700 | 1.00 | 0.00 | C |
| ATOM | 1640 | O | ILE | A | 124 | 47.594 | 46.259 | 36.958 | 1.00 | 0.00 | O |
| ATOM | 1641 | N | ARG | A | 125 | 48.874 | 45.143 | 35.573 | 1.00 | 0.00 | N |
| ATOM | 1642 | H | ARG | A | 125 | 49.449 | 44.336 | 35.437 | 1.00 | 0.00 | H |
| ATOM | 1643 | CA | ARG | A | 125 | 48.828 | 46.160 | 34.501 | 1.00 | 0.00 | C |
| ATOM | 1644 | HA | ARG | A | 125 | 48.813 | 47.074 | 34.907 | 1.00 | 0.00 | H |
| ATOM | 1645 | CB | ARG | A | 125 | 50.115 | 46.055 | 33.657 | 1.00 | 0.00 | C |
| ATOM | 1646 | HB1 | ARG | A | 125 | 49.971 | 46.517 | 32.782 | 1.00 | 0.00 | H |
| ATOM | 1647 | HB2 | ARG | A | 125 | 50.322 | 45.090 | 33.495 | 1.00 | 0.00 | H |
| ATOM | 1648 | CG | ARG | A | 125 | 51.309 | 46.688 | 34.335 | 1.00 | 0.00 | C |
| ATOM | 1649 | HG1 | ARG | A | 125 | 51.642 | 46.076 | 35.052 | 1.00 | 0.00 | H |
| ATOM | 1650 | HG2 | ARG | A | 125 | 51.032 | 47.559 | 34.742 | 1.00 | 0.00 | H |
| ATOM | 1651 | CD | ARG | A | 125 | 52.416 | 46.915 | 33.246 | 1.00 | 0.00 | C |
| ATOM | 1652 | HD1 | ARG | A | 125 | 53.123 | 47.520 | 33.613 | 1.00 | 0.00 | H |
| ATOM | 1653 | HD2 | ARG | A | 125 | 52.005 | 47.335 | 32.437 | 1.00 | 0.00 | H |
| ATOM | 1654 | NE | ARG | A | 125 | 53.087 | 45.631 | 32.809 | 1.00 | 0.00 | N |
| ATOM | 1655 | HE | ARG | A | 125 | 52.695 | 45.112 | 32.049 | 1.00 | 0.00 | H |
| ATOM | 1656 | CZ | ARG | A | 125 | 54.165 | 45.180 | 33.396 | 1.00 | 0.00 | C |
| ATOM | 1657 | NH1 | ARG | A | 125 | 54.726 | 45.706 | 34.445 | 1.00 | 0.00 | N |
| ATOM | 1658 | HH11 | ARG | A | 125 | 54.331 | 46.522 | 34.867 | 1.00 | 0.00 | H |
| ATOM | 1659 | HH12 | ARG | A | 125 | 55.552 | 45.294 | 34.830 | 1.00 | 0.00 | H |
| ATOM | 1660 | NH2 | ARG | A | 125 | 54.718 | 44.135 | 32.865 | 1.00 | 0.00 | N |
| ATOM | 1661 | HH21 | ARG | A | 125 | 54.318 | 43.718 | 32.049 | 1.00 | 0.00 | H |
| ATOM | 1662 | HH22 | ARG | A | 125 | 55.545 | 43.746 | 33.271 | 1.00 | 0.00 | H |
| ATOM | 1663 | C | ARG | A | 125 | 47.516 | 46.014 | 33.709 | 1.00 | 0.00 | C |
| ATOM | 1664 | O | ARG | A | 125 | 46.861 | 46.984 | 33.385 | 1.00 | 0.00 | O |
| ATOM | 1665 | N | PHE | A | 126 | 47.222 | 44.781 | 33.258 | 1.00 | 0.00 | N |
| ATOM | 1666 | H | PHE | A | 126 | 47.831 | 44.045 | 33.554 | 1.00 | 0.00 | H |
| ATOM | 1667 | CA | PHE | A | 126 | 46.096 | 44.386 | 32.371 | 1.00 | 0.00 | C |
| ATOM | 1668 | HA | PHE | A | 126 | 46.233 | 44.828 | 31.485 | 1.00 | 0.00 | H |
| ATOM | 1669 | CB | PHE | A | 126 | 46.040 | 42.870 | 32.094 | 1.00 | 0.00 | C |
| ATOM | 1670 | HB1 | PHE | A | 126 | 45.571 | 42.453 | 32.872 | 1.00 | 0.00 | H |
| ATOM | 1671 | HB2 | PHE | A | 126 | 46.986 | 42.548 | 32.064 | 1.00 | 0.00 | H |

| | | | | | | | | | | | |
|---|---|---|---|---|---|---|---|---|---|---|---|
| ATOM | 1672 | CG | PHE | A | 126 | 45.308 | 42.402 | 30.757 | 1.00 | 0.00 | C |
| ATOM | 1673 | CD1 | PHE | A | 126 | 43.938 | 41.955 | 30.776 | 1.00 | 0.00 | C |
| ATOM | 1674 | HD1 | PHE | A | 126 | 43.414 | 41.993 | 31.627 | 1.00 | 0.00 | H |
| ATOM | 1675 | CE1 | PHE | A | 126 | 43.340 | 41.471 | 29.616 | 1.00 | 0.00 | C |
| ATOM | 1676 | HE1 | PHE | A | 126 | 42.367 | 41.242 | 29.606 | 1.00 | 0.00 | H |
| ATOM | 1677 | CZ | PHE | A | 126 | 44.138 | 41.304 | 28.433 | 1.00 | 0.00 | C |
| ATOM | 1678 | HZ | PHE | A | 126 | 43.712 | 40.954 | 27.599 | 1.00 | 0.00 | H |
| ATOM | 1679 | CE2 | PHE | A | 126 | 45.530 | 41.629 | 28.427 | 1.00 | 0.00 | C |
| ATOM | 1680 | HE2 | PHE | A | 126 | 46.099 | 41.471 | 27.620 | 1.00 | 0.00 | H |
| ATOM | 1681 | CD2 | PHE | A | 126 | 46.076 | 42.191 | 29.628 | 1.00 | 0.00 | C |
| ATOM | 1682 | HD2 | PHE | A | 126 | 47.045 | 42.438 | 29.649 | 1.00 | 0.00 | H |
| ATOM | 1683 | C | PHE | A | 126 | 44.831 | 44.878 | 33.022 | 1.00 | 0.00 | C |
| ATOM | 1684 | O | PHE | A | 126 | 44.159 | 45.699 | 32.472 | 1.00 | 0.00 | O |
| ATOM | 1685 | N | LEU | A | 127 | 44.500 | 44.415 | 34.269 | 1.00 | 0.00 | N |
| ATOM | 1686 | H | LEU | A | 127 | 45.165 | 43.840 | 34.745 | 1.00 | 0.00 | H |
| ATOM | 1687 | CA | LEU | A | 127 | 43.244 | 44.705 | 34.946 | 1.00 | 0.00 | C |
| ATOM | 1688 | HA | LEU | A | 127 | 42.555 | 44.534 | 34.241 | 1.00 | 0.00 | H |
| ATOM | 1689 | CB | LEU | A | 127 | 43.133 | 43.770 | 36.139 | 1.00 | 0.00 | C |
| ATOM | 1690 | HB1 | LEU | A | 127 | 42.212 | 43.902 | 36.506 | 1.00 | 0.00 | H |
| ATOM | 1691 | HB2 | LEU | A | 127 | 43.806 | 44.093 | 36.804 | 1.00 | 0.00 | H |
| ATOM | 1692 | CG | LEU | A | 127 | 43.347 | 42.237 | 35.977 | 1.00 | 0.00 | C |
| ATOM | 1693 | HG | LEU | A | 127 | 44.213 | 42.040 | 35.516 | 1.00 | 0.00 | H |
| ATOM | 1694 | CD1 | LEU | A | 127 | 43.265 | 41.653 | 37.397 | 1.00 | 0.00 | C |
| ATOM | 1695 | HD11 | LEU | A | 127 | 43.396 | 40.662 | 37.358 | 1.00 | 0.00 | H |
| ATOM | 1696 | HD12 | LEU | A | 127 | 43.978 | 42.061 | 37.967 | 1.00 | 0.00 | H |
| ATOM | 1697 | HD13 | LEU | A | 127 | 42.368 | 41.855 | 37.789 | 1.00 | 0.00 | H |
| ATOM | 1698 | CD2 | LEU | A | 127 | 42.273 | 41.599 | 35.124 | 1.00 | 0.00 | C |
| ATOM | 1699 | HD21 | LEU | A | 127 | 42.448 | 40.618 | 35.046 | 1.00 | 0.00 | H |
| ATOM | 1700 | HD22 | LEU | A | 127 | 41.379 | 41.745 | 35.548 | 1.00 | 0.00 | H |
| ATOM | 1701 | HD23 | LEU | A | 127 | 42.281 | 42.012 | 34.213 | 1.00 | 0.00 | H |
| ATOM | 1702 | C | LEU | A | 127 | 42.955 | 46.142 | 35.306 | 1.00 | 0.00 | C |
| ATOM | 1703 | O | LEU | A | 127 | 41.900 | 46.701 | 34.912 | 1.00 | 0.00 | O |
| ATOM | 1704 | N | ARG | A | 128 | 43.975 | 46.934 | 35.596 | 1.00 | 0.00 | N |
| ATOM | 1705 | H | ARG | A | 128 | 44.865 | 46.484 | 35.674 | 1.00 | 0.00 | H |
| ATOM | 1706 | CA | ARG | A | 128 | 43.974 | 48.342 | 35.812 | 1.00 | 0.00 | C |
| ATOM | 1707 | HA | ARG | A | 128 | 43.164 | 48.364 | 36.398 | 1.00 | 0.00 | H |
| ATOM | 1708 | CB | ARG | A | 128 | 45.198 | 48.883 | 36.659 | 1.00 | 0.00 | C |
| ATOM | 1709 | HB1 | ARG | A | 128 | 45.179 | 49.883 | 36.678 | 1.00 | 0.00 | H |
| ATOM | 1710 | HB2 | ARG | A | 128 | 46.056 | 48.573 | 36.248 | 1.00 | 0.00 | H |
| ATOM | 1711 | CG | ARG | A | 128 | 45.154 | 48.381 | 38.098 | 1.00 | 0.00 | C |
| ATOM | 1712 | HG1 | ARG | A | 128 | 46.031 | 48.580 | 38.535 | 1.00 | 0.00 | H |
| ATOM | 1713 | HG2 | ARG | A | 128 | 45.003 | 47.393 | 38.089 | 1.00 | 0.00 | H |
| ATOM | 1714 | CD | ARG | A | 128 | 44.060 | 49.021 | 38.918 | 1.00 | 0.00 | C |
| ATOM | 1715 | HD1 | ARG | A | 128 | 43.160 | 48.735 | 38.590 | 1.00 | 0.00 | H |
| ATOM | 1716 | HD2 | ARG | A | 128 | 44.127 | 50.018 | 38.880 | 1.00 | 0.00 | H |
| ATOM | 1717 | NE | ARG | A | 128 | 44.251 | 48.564 | 40.302 | 1.00 | 0.00 | N |
| ATOM | 1718 | HE | ARG | A | 128 | 45.190 | 48.397 | 40.603 | 1.00 | 0.00 | H |
| ATOM | 1719 | CZ | ARG | A | 128 | 43.287 | 48.355 | 41.182 | 1.00 | 0.00 | C |
| ATOM | 1720 | NH1 | ARG | A | 128 | 42.040 | 48.527 | 40.902 | 1.00 | 0.00 | N |
| ATOM | 1721 | HH11 | ARG | A | 128 | 41.770 | 48.830 | 39.988 | 1.00 | 0.00 | H |
| ATOM | 1722 | HH12 | ARG | A | 128 | 41.343 | 48.357 | 41.599 | 1.00 | 0.00 | H |
| ATOM | 1723 | NH2 | ARG | A | 128 | 43.611 | 48.038 | 42.425 | 1.00 | 0.00 | N |
| ATOM | 1724 | HH21 | ARG | A | 128 | 44.573 | 47.960 | 42.686 | 1.00 | 0.00 | H |
| ATOM | 1725 | HH22 | ARG | A | 128 | 42.894 | 47.877 | 43.103 | 1.00 | 0.00 | H |

```
ATOM   1726  C   ARG A 128      43.741  49.182  34.464  1.00  0.00           C
ATOM   1727  O   ARG A 128      43.057  50.183  34.466  1.00  0.00           O
ATOM   1728  N   LYS A 129      44.449  48.803  33.382  1.00  0.00           N
ATOM   1729  H   LYS A 129      45.104  48.051  33.449  1.00  0.00           H
ATOM   1730  CA  LYS A 129      44.252  49.509  32.086  1.00  0.00           C
ATOM   1731  HA  LYS A 129      44.419  50.485  32.223  1.00  0.00           H
ATOM   1732  CB  LYS A 129      45.266  48.888  31.129  1.00  0.00           C
ATOM   1733  HB1 LYS A 129      45.053  47.914  31.051  1.00  0.00           H
ATOM   1734  HB2 LYS A 129      46.174  49.000  31.534  1.00  0.00           H
ATOM   1735  CG  LYS A 129      45.287  49.499  29.705  1.00  0.00           C
ATOM   1736  HG1 LYS A 129      45.449  50.483  29.783  1.00  0.00           H
ATOM   1737  HG2 LYS A 129      44.397  49.341  29.278  1.00  0.00           H
ATOM   1738  CD  LYS A 129      46.340  48.940  28.775  1.00  0.00           C
ATOM   1739  HD1 LYS A 129      46.118  47.976  28.627  1.00  0.00           H
ATOM   1740  HD2 LYS A 129      47.217  49.011  29.250  1.00  0.00           H
ATOM   1741  CE  LYS A 129      46.438  49.683  27.368  1.00  0.00           C
ATOM   1742  HE1 LYS A 129      47.298  49.439  26.920  1.00  0.00           H
ATOM   1743  HE2 LYS A 129      46.405  50.672  27.510  1.00  0.00           H
ATOM   1744  NZ  LYS A 129      45.360  49.367  26.396  1.00  0.00           N
ATOM   1745  HZ1 LYS A 129      45.512  49.880  25.551  1.00  0.00           H
ATOM   1746  HZ2 LYS A 129      45.368  48.387  26.197  1.00  0.00           H
ATOM   1747  HZ3 LYS A 129      44.475  49.621  26.786  1.00  0.00           H
ATOM   1748  C   LYS A 129      42.806  49.464  31.554  1.00  0.00           C
ATOM   1749  O   LYS A 129      42.350  50.468  30.956  1.00  0.00           O
ATOM   1750  N   TYR A 130      42.055  48.363  31.849  1.00  0.00           N
ATOM   1751  H   TYR A 130      42.485  47.648  32.401  1.00  0.00           H
ATOM   1752  CA  TYR A 130      40.677  48.143  31.426  1.00  0.00           C
ATOM   1753  HA  TYR A 130      40.643  48.762  30.642  1.00  0.00           H
ATOM   1754  CB  TYR A 130      40.404  46.712  30.903  1.00  0.00           C
ATOM   1755  HB1 TYR A 130      39.433  46.646  30.673  1.00  0.00           H
ATOM   1756  HB2 TYR A 130      40.617  46.068  31.638  1.00  0.00           H
ATOM   1757  CG  TYR A 130      41.249  46.328  29.636  1.00  0.00           C
ATOM   1758  CD1 TYR A 130      41.100  47.020  28.424  1.00  0.00           C
ATOM   1759  HD1 TYR A 130      40.438  47.764  28.334  1.00  0.00           H
ATOM   1760  CE1 TYR A 130      41.913  46.634  27.334  1.00  0.00           C
ATOM   1761  HE1 TYR A 130      41.771  47.094  26.457  1.00  0.00           H
ATOM   1762  CZ  TYR A 130      42.900  45.652  27.406  1.00  0.00           C
ATOM   1763  OH  TYR A 130      43.568  45.209  26.296  1.00  0.00           O
ATOM   1764  HH  TYR A 130      44.230  44.509  26.563  1.00  0.00           H
ATOM   1765  CE2 TYR A 130      43.072  45.010  28.648  1.00  0.00           C
ATOM   1766  HE2 TYR A 130      43.771  44.304  28.758  1.00  0.00           H
ATOM   1767  CD2 TYR A 130      42.250  45.372  29.733  1.00  0.00           C
ATOM   1768  HD2 TYR A 130      42.394  44.920  30.613  1.00  0.00           H
ATOM   1769  C   TYR A 130      39.611  48.637  32.434  1.00  0.00           C
ATOM   1770  O   TYR A 130      38.451  48.914  32.114  1.00  0.00           O
ATOM   1771  N   SER A 131      40.028  48.740  33.663  1.00  0.00           N
ATOM   1772  H   SER A 131      41.001  48.573  33.821  1.00  0.00           H
ATOM   1773  CA  SER A 131      39.195  49.086  34.876  1.00  0.00           C
ATOM   1774  HA  SER A 131      39.867  49.370  35.560  1.00  0.00           H
ATOM   1775  CB  SER A 131      38.351  50.318  34.666  1.00  0.00           C
ATOM   1776  HB1 SER A 131      38.001  50.664  35.536  1.00  0.00           H
ATOM   1777  HB2 SER A 131      37.586  50.127  34.051  1.00  0.00           H
ATOM   1778  OG  SER A 131      39.163  51.320  34.077  1.00  0.00           O
ATOM   1779  HG  SER A 131      38.619  52.146  33.930  1.00  0.00           H
```

| | | | | | | | | | | | |
|---|---|---|---|---|---|---|---|---|---|---|---|
| ATOM | 1780 | C | SER | A | 131 | 38.450 | 47.880 | 35.530 | 1.00 | 0.00 | C |
| ATOM | 1781 | O | SER | A | 131 | 37.352 | 47.987 | 36.122 | 1.00 | 0.00 | O |
| ATOM | 1782 | N | PHE | A | 132 | 39.064 | 46.653 | 35.490 | 1.00 | 0.00 | N |
| ATOM | 1783 | H | PHE | A | 132 | 39.870 | 46.564 | 34.904 | 1.00 | 0.00 | H |
| ATOM | 1784 | CA | PHE | A | 132 | 38.644 | 45.470 | 36.226 | 1.00 | 0.00 | C |
| ATOM | 1785 | HA | PHE | A | 132 | 37.666 | 45.611 | 36.071 | 1.00 | 0.00 | H |
| ATOM | 1786 | CB | PHE | A | 132 | 39.188 | 44.177 | 35.624 | 1.00 | 0.00 | C |
| ATOM | 1787 | HB1 | PHE | A | 132 | 38.941 | 43.431 | 36.243 | 1.00 | 0.00 | H |
| ATOM | 1788 | HB2 | PHE | A | 132 | 40.184 | 44.261 | 35.584 | 1.00 | 0.00 | H |
| ATOM | 1789 | CG | PHE | A | 132 | 38.679 | 43.815 | 34.207 | 1.00 | 0.00 | C |
| ATOM | 1790 | CD1 | PHE | A | 132 | 37.376 | 43.283 | 34.032 | 1.00 | 0.00 | C |
| ATOM | 1791 | HD1 | PHE | A | 132 | 36.768 | 43.195 | 34.821 | 1.00 | 0.00 | H |
| ATOM | 1792 | CE1 | PHE | A | 132 | 36.925 | 42.873 | 32.744 | 1.00 | 0.00 | C |
| ATOM | 1793 | HE1 | PHE | A | 132 | 36.005 | 42.504 | 32.615 | 1.00 | 0.00 | H |
| ATOM | 1794 | CZ | PHE | A | 132 | 37.786 | 42.999 | 31.690 | 1.00 | 0.00 | C |
| ATOM | 1795 | HZ | PHE | A | 132 | 37.446 | 42.858 | 30.760 | 1.00 | 0.00 | H |
| ATOM | 1796 | CE2 | PHE | A | 132 | 39.152 | 43.326 | 31.887 | 1.00 | 0.00 | C |
| ATOM | 1797 | HE2 | PHE | A | 132 | 39.798 | 43.259 | 31.127 | 1.00 | 0.00 | H |
| ATOM | 1798 | CD2 | PHE | A | 132 | 39.587 | 43.737 | 33.132 | 1.00 | 0.00 | C |
| ATOM | 1799 | HD2 | PHE | A | 132 | 40.547 | 43.979 | 33.271 | 1.00 | 0.00 | H |
| ATOM | 1800 | C | PHE | A | 132 | 38.897 | 45.404 | 37.757 | 1.00 | 0.00 | C |
| ATOM | 1801 | O | PHE | A | 132 | 39.958 | 45.871 | 38.152 | 1.00 | 0.00 | O |
| ATOM | 1802 | N | ASP | A | 133 | 37.939 | 44.892 | 38.491 | 1.00 | 0.00 | N |
| ATOM | 1803 | H | ASP | A | 133 | 37.094 | 44.680 | 37.999 | 1.00 | 0.00 | H |
| ATOM | 1804 | CA | ASP | A | 133 | 37.919 | 44.582 | 39.929 | 1.00 | 0.00 | C |
| ATOM | 1805 | HA | ASP | A | 133 | 38.608 | 45.222 | 40.268 | 1.00 | 0.00 | H |
| ATOM | 1806 | CB | ASP | A | 133 | 36.505 | 44.865 | 40.521 | 1.00 | 0.00 | C |
| ATOM | 1807 | HB1 | ASP | A | 133 | 36.461 | 44.549 | 41.469 | 1.00 | 0.00 | H |
| ATOM | 1808 | HB2 | ASP | A | 133 | 35.807 | 44.396 | 39.980 | 1.00 | 0.00 | H |
| ATOM | 1809 | CG | ASP | A | 133 | 36.206 | 46.326 | 40.502 | 1.00 | 0.00 | C |
| ATOM | 1810 | OD1 | ASP | A | 133 | 37.097 | 47.167 | 40.727 | 1.00 | 0.00 | O |
| ATOM | 1811 | OD2 | ASP | A | 133 | 34.993 | 46.599 | 40.389 | 1.00 | 0.00 | O |
| ATOM | 1812 | C | ASP | A | 133 | 38.446 | 43.184 | 40.349 | 1.00 | 0.00 | C |
| ATOM | 1813 | O | ASP | A | 133 | 38.745 | 42.999 | 41.549 | 1.00 | 0.00 | O |
| ATOM | 1814 | N | GLY | A | 134 | 38.774 | 42.203 | 39.465 | 1.00 | 0.00 | N |
| ATOM | 1815 | H | GLY | A | 134 | 38.562 | 42.323 | 38.495 | 1.00 | 0.00 | H |
| ATOM | 1816 | CA | GLY | A | 134 | 39.459 | 40.925 | 39.929 | 1.00 | 0.00 | C |
| ATOM | 1817 | HA1 | GLY | A | 134 | 38.820 | 40.397 | 40.488 | 1.00 | 0.00 | H |
| ATOM | 1818 | HA2 | GLY | A | 134 | 40.261 | 41.165 | 40.476 | 1.00 | 0.00 | H |
| ATOM | 1819 | C | GLY | A | 134 | 39.911 | 40.060 | 38.804 | 1.00 | 0.00 | C |
| ATOM | 1820 | O | GLY | A | 134 | 39.733 | 40.399 | 37.614 | 1.00 | 0.00 | O |
| ATOM | 1821 | N | LEU | A | 135 | 40.480 | 38.896 | 39.085 | 1.00 | 0.00 | N |
| ATOM | 1822 | H | LEU | A | 135 | 40.671 | 38.751 | 40.056 | 1.00 | 0.00 | H |
| ATOM | 1823 | CA | LEU | A | 135 | 40.867 | 37.813 | 38.235 | 1.00 | 0.00 | C |
| ATOM | 1824 | HA | LEU | A | 135 | 40.568 | 38.038 | 37.308 | 1.00 | 0.00 | H |
| ATOM | 1825 | CB | LEU | A | 135 | 42.421 | 37.652 | 38.444 | 1.00 | 0.00 | C |
| ATOM | 1826 | HB1 | LEU | A | 135 | 42.566 | 37.224 | 39.336 | 1.00 | 0.00 | H |
| ATOM | 1827 | HB2 | LEU | A | 135 | 42.825 | 38.567 | 38.443 | 1.00 | 0.00 | H |
| ATOM | 1828 | CG | LEU | A | 135 | 43.239 | 36.839 | 37.460 | 1.00 | 0.00 | C |
| ATOM | 1829 | HG | LEU | A | 135 | 42.783 | 35.962 | 37.310 | 1.00 | 0.00 | H |
| ATOM | 1830 | CD1 | LEU | A | 135 | 43.323 | 37.472 | 36.026 | 1.00 | 0.00 | C |
| ATOM | 1831 | HD11 | LEU | A | 135 | 43.875 | 36.885 | 35.434 | 1.00 | 0.00 | H |
| ATOM | 1832 | HD12 | LEU | A | 135 | 42.402 | 37.558 | 35.645 | 1.00 | 0.00 | H |
| ATOM | 1833 | HD13 | LEU | A | 135 | 43.745 | 38.377 | 36.085 | 1.00 | 0.00 | H |

```
ATOM   1834  CD2 LEU A 135      44.611  36.620  38.120  1.00  0.00           C
ATOM   1835 HD21 LEU A 135      45.194  36.085  37.508  1.00  0.00           H
ATOM   1836 HD22 LEU A 135      45.040  37.505  38.299  1.00  0.00           H
ATOM   1837 HD23 LEU A 135      44.491  36.127  38.982  1.00  0.00           H
ATOM   1838  C   LEU A 135      40.179  36.530  38.664  1.00  0.00           C
ATOM   1839  O   LEU A 135      40.213  36.239  39.831  1.00  0.00           O
ATOM   1840  N   ASP A 136      39.632  35.720  37.768  1.00  0.00           N
ATOM   1841  H   ASP A 136      39.653  36.012  36.812  1.00  0.00           H
ATOM   1842  CA  ASP A 136      39.000  34.432  38.064  1.00  0.00           C
ATOM   1843  HA  ASP A 136      39.132  34.254  39.039  1.00  0.00           H
ATOM   1844  CB  ASP A 136      37.491  34.510  37.810  1.00  0.00           C
ATOM   1845  HB1 ASP A 136      37.337  34.673  36.835  1.00  0.00           H
ATOM   1846  HB2 ASP A 136      37.115  35.271  38.339  1.00  0.00           H
ATOM   1847  CG  ASP A 136      36.762  33.226  38.213  1.00  0.00           C
ATOM   1848  OD1 ASP A 136      35.881  32.785  37.448  1.00  0.00           O
ATOM   1849  OD2 ASP A 136      37.027  32.728  39.332  1.00  0.00           O
ATOM   1850  C   ASP A 136      39.698  33.308  37.287  1.00  0.00           C
ATOM   1851  O   ASP A 136      39.903  33.411  36.076  1.00  0.00           O
ATOM   1852  N   LEU A 137      40.164  32.297  38.018  1.00  0.00           N
ATOM   1853  H   LEU A 137      39.824  32.213  38.955  1.00  0.00           H
ATOM   1854  CA  LEU A 137      41.134  31.297  37.563  1.00  0.00           C
ATOM   1855  HA  LEU A 137      41.410  31.587  36.647  1.00  0.00           H
ATOM   1856  CB  LEU A 137      42.292  31.275  38.576  1.00  0.00           C
ATOM   1857  HB1 LEU A 137      42.830  30.449  38.407  1.00  0.00           H
ATOM   1858  HB2 LEU A 137      41.895  31.233  39.493  1.00  0.00           H
ATOM   1859  CG  LEU A 137      43.254  32.454  38.554  1.00  0.00           C
ATOM   1860  HG  LEU A 137      42.739  33.262  38.841  1.00  0.00           H
ATOM   1861  CD1 LEU A 137      44.395  32.058  39.474  1.00  0.00           C
ATOM   1862 HD11 LEU A 137      45.069  32.796  39.505  1.00  0.00           H
ATOM   1863 HD12 LEU A 137      44.039  31.892  40.394  1.00  0.00           H
ATOM   1864 HD13 LEU A 137      44.828  31.225  39.129  1.00  0.00           H
ATOM   1865  CD2 LEU A 137      43.876  32.773  37.165  1.00  0.00           C
ATOM   1866 HD21 LEU A 137      44.491  33.557  37.247  1.00  0.00           H
ATOM   1867 HD22 LEU A 137      44.390  31.979  36.840  1.00  0.00           H
ATOM   1868 HD23 LEU A 137      43.147  32.987  36.515  1.00  0.00           H
ATOM   1869  C   LEU A 137      40.539  29.891  37.446  1.00  0.00           C
ATOM   1870  O   LEU A 137      39.811  29.463  38.335  1.00  0.00           O
ATOM   1871  N   ASP A 138      40.992  29.131  36.447  1.00  0.00           N
ATOM   1872  H   ASP A 138      41.552  29.576  35.748  1.00  0.00           H
ATOM   1873  CA  ASP A 138      40.734  27.695  36.292  1.00  0.00           C
ATOM   1874  HA  ASP A 138      40.664  27.300  37.208  1.00  0.00           H
ATOM   1875  CB  ASP A 138      39.407  27.506  35.547  1.00  0.00           C
ATOM   1876  HB1 ASP A 138      39.557  27.717  34.581  1.00  0.00           H
ATOM   1877  HB2 ASP A 138      38.740  28.145  35.928  1.00  0.00           H
ATOM   1878  CG  ASP A 138      38.824  26.094  35.642  1.00  0.00           C
ATOM   1879  OD1 ASP A 138      37.772  25.907  34.989  1.00  0.00           O
ATOM   1880  OD2 ASP A 138      39.378  25.258  36.398  1.00  0.00           O
ATOM   1881  C   ASP A 138      41.905  27.024  35.545  1.00  0.00           C
ATOM   1882  O   ASP A 138      42.635  27.691  34.804  1.00  0.00           O
ATOM   1883  N   TRP A 139      42.143  25.732  35.778  1.00  0.00           N
ATOM   1884  H   TRP A 139      41.531  25.239  36.396  1.00  0.00           H
ATOM   1885  CA  TRP A 139      43.262  25.000  35.169  1.00  0.00           C
ATOM   1886  HA  TRP A 139      43.279  25.311  34.219  1.00  0.00           H
ATOM   1887  CB  TRP A 139      44.580  25.330  35.915  1.00  0.00           C
```

```
ATOM   1888  HB1 TRP A 139      44.474  25.059  36.872  1.00  0.00           H
ATOM   1889  HB2 TRP A 139      44.730  26.317  35.863  1.00  0.00           H
ATOM   1890  CG  TRP A 139      45.838  24.686  35.431  1.00  0.00           C
ATOM   1891  CD1 TRP A 139      45.989  24.435  34.138  1.00  0.00           C
ATOM   1892  HD1 TRP A 139      45.326  24.667  33.426  1.00  0.00           H
ATOM   1893  NE1 TRP A 139      47.207  23.802  33.919  1.00  0.00           N
ATOM   1894  HE1 TRP A 139      47.560  23.539  33.021  1.00  0.00           H
ATOM   1895  CE2 TRP A 139      47.870  23.584  35.134  1.00  0.00           C
ATOM   1896  CZ2 TRP A 139      49.215  23.169  35.444  1.00  0.00           C
ATOM   1897  HZ2 TRP A 139      49.825  22.805  34.740  1.00  0.00           H
ATOM   1898  CH2 TRP A 139      49.641  23.303  36.818  1.00  0.00           C
ATOM   1899  HH2 TRP A 139      50.557  23.007  37.089  1.00  0.00           H
ATOM   1900  CZ3 TRP A 139      48.752  23.860  37.791  1.00  0.00           C
ATOM   1901  HZ3 TRP A 139      49.064  24.005  38.730  1.00  0.00           H
ATOM   1902  CE3 TRP A 139      47.483  24.192  37.437  1.00  0.00           C
ATOM   1903  HE3 TRP A 139      46.858  24.478  38.164  1.00  0.00           H
ATOM   1904  CD2 TRP A 139      46.994  24.163  36.145  1.00  0.00           C
ATOM   1905  C   TRP A 139      43.038  23.489  35.184  1.00  0.00           C
ATOM   1906  O   TRP A 139      42.843  22.920  36.261  1.00  0.00           O
ATOM   1907  N   GLU A 140      43.240  22.812  34.043  1.00  0.00           N
ATOM   1908  H   GLU A 140      43.421  23.304  33.191  1.00  0.00           H
ATOM   1909  CA  GLU A 140      43.193  21.340  34.054  1.00  0.00           C
ATOM   1910  HA  GLU A 140      42.794  21.113  34.942  1.00  0.00           H
ATOM   1911  CB  GLU A 140      42.364  20.869  32.838  1.00  0.00           C
ATOM   1912  HB1 GLU A 140      42.451  19.875  32.770  1.00  0.00           H
ATOM   1913  HB2 GLU A 140      42.748  21.294  32.018  1.00  0.00           H
ATOM   1914  CG  GLU A 140      40.866  21.206  32.882  1.00  0.00           C
ATOM   1915  HG1 GLU A 140      40.550  21.109  33.826  1.00  0.00           H
ATOM   1916  HG2 GLU A 140      40.381  20.554  32.299  1.00  0.00           H
ATOM   1917  CD  GLU A 140      40.525  22.625  32.404  1.00  0.00           C
ATOM   1918  OE1 GLU A 140      41.275  23.172  31.562  1.00  0.00           O
ATOM   1919  OE2 GLU A 140      39.486  23.139  32.860  1.00  0.00           O
ATOM   1920  C   GLU A 140      44.565  20.675  33.967  1.00  0.00           C
ATOM   1921  O   GLU A 140      45.155  20.605  32.863  1.00  0.00           O
ATOM   1922  N   TYR A 141      45.179  20.191  35.058  1.00  0.00           N
ATOM   1923  H   TYR A 141      46.113  19.864  34.916  1.00  0.00           H
ATOM   1924  CA  TYR A 141      44.640  20.068  36.513  1.00  0.00           C
ATOM   1925  HA  TYR A 141      43.969  20.787  36.696  1.00  0.00           H
ATOM   1926  CB  TYR A 141      43.939  18.689  36.745  1.00  0.00           C
ATOM   1927  HB1 TYR A 141      43.670  18.612  37.705  1.00  0.00           H
ATOM   1928  HB2 TYR A 141      44.575  17.952  36.517  1.00  0.00           H
ATOM   1929  CG  TYR A 141      42.720  18.456  35.946  1.00  0.00           C
ATOM   1930  CD1 TYR A 141      41.475  18.875  36.467  1.00  0.00           C
ATOM   1931  HD1 TYR A 141      41.436  19.318  37.363  1.00  0.00           H
ATOM   1932  CE1 TYR A 141      40.289  18.674  35.735  1.00  0.00           C
ATOM   1933  HE1 TYR A 141      39.412  18.979  36.106  1.00  0.00           H
ATOM   1934  CZ  TYR A 141      40.344  18.040  34.476  1.00  0.00           C
ATOM   1935  OH  TYR A 141      39.217  17.843  33.747  1.00  0.00           O
ATOM   1936  HH  TYR A 141      39.450  17.383  32.890  1.00  0.00           H
ATOM   1937  CE2 TYR A 141      41.586  17.621  33.964  1.00  0.00           C
ATOM   1938  HE2 TYR A 141      41.628  17.147  33.085  1.00  0.00           H
ATOM   1939  CD2 TYR A 141      42.769  17.867  34.685  1.00  0.00           C
ATOM   1940  HD2 TYR A 141      43.652  17.617  34.289  1.00  0.00           H
ATOM   1941  C   TYR A 141      45.815  20.238  37.459  1.00  0.00           C
```

| | | | | | | | | | | | |
|---|---|---|---|---|---|---|---|---|---|---|---|
| ATOM | 1942 | O | TYR | A | 141 | 46.825 | 19.512 | 37.362 | 1.00 | 0.00 | O |
| ATOM | 1943 | N | PRO | A | 142 | 45.596 | 20.871 | 38.640 | 1.00 | 0.00 | N |
| ATOM | 1944 | CD | PRO | A | 142 | 44.573 | 21.817 | 38.893 | 1.00 | 0.00 | C |
| ATOM | 1945 | HD1 | PRO | A | 142 | 43.693 | 21.355 | 38.779 | 1.00 | 0.00 | H |
| ATOM | 1946 | HD2 | PRO | A | 142 | 44.656 | 22.547 | 38.215 | 1.00 | 0.00 | H |
| ATOM | 1947 | CG | PRO | A | 142 | 44.716 | 22.370 | 40.305 | 1.00 | 0.00 | C |
| ATOM | 1948 | HG1 | PRO | A | 142 | 44.243 | 21.798 | 40.976 | 1.00 | 0.00 | H |
| ATOM | 1949 | HG2 | PRO | A | 142 | 44.379 | 23.309 | 40.370 | 1.00 | 0.00 | H |
| ATOM | 1950 | CB | PRO | A | 142 | 46.217 | 22.307 | 40.483 | 1.00 | 0.00 | C |
| ATOM | 1951 | HB1 | PRO | A | 142 | 46.461 | 22.252 | 41.451 | 1.00 | 0.00 | H |
| ATOM | 1952 | HB2 | PRO | A | 142 | 46.660 | 23.103 | 40.072 | 1.00 | 0.00 | H |
| ATOM | 1953 | CA | PRO | A | 142 | 46.548 | 20.982 | 39.713 | 1.00 | 0.00 | C |
| ATOM | 1954 | HA | PRO | A | 142 | 47.447 | 20.970 | 39.276 | 1.00 | 0.00 | H |
| ATOM | 1955 | C | PRO | A | 142 | 46.559 | 19.785 | 40.634 | 1.00 | 0.00 | C |
| ATOM | 1956 | O | PRO | A | 142 | 45.909 | 19.654 | 41.687 | 1.00 | 0.00 | O |
| ATOM | 1957 | N | GLY | A | 143 | 47.351 | 18.741 | 40.165 | 1.00 | 0.00 | N |
| ATOM | 1958 | H | GLY | A | 143 | 47.662 | 18.779 | 39.215 | 1.00 | 0.00 | H |
| ATOM | 1959 | CA | GLY | A | 143 | 47.747 | 17.620 | 40.955 | 1.00 | 0.00 | C |
| ATOM | 1960 | HA1 | GLY | A | 143 | 48.063 | 17.975 | 41.835 | 1.00 | 0.00 | H |
| ATOM | 1961 | HA2 | GLY | A | 143 | 48.508 | 17.178 | 40.481 | 1.00 | 0.00 | H |
| ATOM | 1962 | C | GLY | A | 143 | 46.553 | 16.628 | 41.146 | 1.00 | 0.00 | C |
| ATOM | 1963 | O | GLY | A | 143 | 46.809 | 15.562 | 41.771 | 1.00 | 0.00 | O |
| ATOM | 1964 | N | SER | A | 144 | 45.394 | 16.787 | 40.491 | 1.00 | 0.00 | N |
| ATOM | 1965 | H | SER | A | 144 | 45.309 | 17.559 | 39.861 | 1.00 | 0.00 | H |
| ATOM | 1966 | CA | SER | A | 144 | 44.255 | 15.898 | 40.647 | 1.00 | 0.00 | C |
| ATOM | 1967 | HA | SER | A | 144 | 44.439 | 15.430 | 41.511 | 1.00 | 0.00 | H |
| ATOM | 1968 | CB | SER | A | 144 | 42.981 | 16.679 | 40.749 | 1.00 | 0.00 | C |
| ATOM | 1969 | HB1 | SER | A | 144 | 43.014 | 17.320 | 41.515 | 1.00 | 0.00 | H |
| ATOM | 1970 | HB2 | SER | A | 144 | 42.195 | 16.071 | 40.857 | 1.00 | 0.00 | H |
| ATOM | 1971 | OG | SER | A | 144 | 42.774 | 17.413 | 39.614 | 1.00 | 0.00 | O |
| ATOM | 1972 | HG | SER | A | 144 | 41.920 | 17.927 | 39.700 | 1.00 | 0.00 | H |
| ATOM | 1973 | C | SER | A | 144 | 44.120 | 14.913 | 39.436 | 1.00 | 0.00 | C |
| ATOM | 1974 | O | SER | A | 144 | 44.765 | 15.145 | 38.368 | 1.00 | 0.00 | O |
| ATOM | 1975 | N | GLN | A | 145 | 43.266 | 13.839 | 39.494 | 1.00 | 0.00 | N |
| ATOM | 1976 | H | GLN | A | 145 | 42.786 | 13.688 | 40.358 | 1.00 | 0.00 | H |
| ATOM | 1977 | CA | GLN | A | 145 | 42.992 | 12.892 | 38.407 | 1.00 | 0.00 | C |
| ATOM | 1978 | HA | GLN | A | 145 | 42.621 | 12.141 | 38.954 | 1.00 | 0.00 | H |
| ATOM | 1979 | CB | GLN | A | 145 | 41.887 | 13.404 | 37.447 | 1.00 | 0.00 | C |
| ATOM | 1980 | HB1 | GLN | A | 145 | 41.085 | 13.611 | 38.008 | 1.00 | 0.00 | H |
| ATOM | 1981 | HB2 | GLN | A | 145 | 41.672 | 12.654 | 36.822 | 1.00 | 0.00 | H |
| ATOM | 1982 | CG | GLN | A | 145 | 42.148 | 14.646 | 36.574 | 1.00 | 0.00 | C |
| ATOM | 1983 | HG1 | GLN | A | 145 | 42.435 | 15.394 | 37.172 | 1.00 | 0.00 | H |
| ATOM | 1984 | HG2 | GLN | A | 145 | 41.293 | 14.893 | 36.119 | 1.00 | 0.00 | H |
| ATOM | 1985 | CD | GLN | A | 145 | 43.199 | 14.504 | 35.502 | 1.00 | 0.00 | C |
| ATOM | 1986 | OE1 | GLN | A | 145 | 42.906 | 14.095 | 34.392 | 1.00 | 0.00 | O |
| ATOM | 1987 | NE2 | GLN | A | 145 | 44.468 | 14.822 | 35.720 | 1.00 | 0.00 | N |
| ATOM | 1988 | HE21 | GLN | A | 145 | 44.750 | 15.167 | 36.615 | 1.00 | 0.00 | H |
| ATOM | 1989 | HE22 | GLN | A | 145 | 45.142 | 14.717 | 34.989 | 1.00 | 0.00 | H |
| ATOM | 1990 | C | GLN | A | 145 | 44.194 | 12.184 | 37.733 | 1.00 | 0.00 | C |
| ATOM | 1991 | O | GLN | A | 145 | 44.034 | 11.804 | 36.586 | 1.00 | 0.00 | O |
| ATOM | 1992 | N | GLY | A | 146 | 45.356 | 12.204 | 38.397 | 1.00 | 0.00 | N |
| ATOM | 1993 | H | GLY | A | 146 | 45.374 | 12.616 | 39.308 | 1.00 | 0.00 | H |
| ATOM | 1994 | CA | GLY | A | 146 | 46.635 | 11.640 | 37.845 | 1.00 | 0.00 | C |
| ATOM | 1995 | HA1 | GLY | A | 146 | 46.443 | 11.260 | 36.940 | 1.00 | 0.00 | H |

```
ATOM   1996  HA2 GLY A 146      46.942  10.909  38.455  1.00  0.00           H
ATOM   1997  C   GLY A 146      47.748  12.687  37.721  1.00  0.00           C
ATOM   1998  O   GLY A 146      48.942  12.306  37.920  1.00  0.00           O
ATOM   1999  N   SER A 147      47.445  13.940  37.357  1.00  0.00           N
ATOM   2000  H   SER A 147      46.507  14.139  37.074  1.00  0.00           H
ATOM   2001  CA  SER A 147      48.420  15.033  37.353  1.00  0.00           C
ATOM   2002  HA  SER A 147      48.962  14.907  36.522  1.00  0.00           H
ATOM   2003  CB  SER A 147      47.663  16.345  37.324  1.00  0.00           C
ATOM   2004  HB1 SER A 147      47.260  16.531  38.220  1.00  0.00           H
ATOM   2005  HB2 SER A 147      46.940  16.311  36.634  1.00  0.00           H
ATOM   2006  OG  SER A 147      48.423  17.451  37.017  1.00  0.00           O
ATOM   2007  HG  SER A 147      47.843  18.266  37.021  1.00  0.00           H
ATOM   2008  C   SER A 147      49.407  14.987  38.519  1.00  0.00           C
ATOM   2009  O   SER A 147      48.920  14.681  39.600  1.00  0.00           O
ATOM   2010  N   PRO A 148      50.723  15.135  38.447  1.00  0.00           N
ATOM   2011  CD  PRO A 148      51.419  15.363  37.203  1.00  0.00           C
ATOM   2012  HD1 PRO A 148      50.886  15.972  36.616  1.00  0.00           H
ATOM   2013  HD2 PRO A 148      51.574  14.496  36.730  1.00  0.00           H
ATOM   2014  CG  PRO A 148      52.710  15.997  37.627  1.00  0.00           C
ATOM   2015  HG1 PRO A 148      52.586  16.977  37.784  1.00  0.00           H
ATOM   2016  HG2 PRO A 148      53.415  15.858  36.932  1.00  0.00           H
ATOM   2017  CB  PRO A 148      53.027  15.222  38.953  1.00  0.00           C
ATOM   2018  HB1 PRO A 148      53.642  15.741  39.547  1.00  0.00           H
ATOM   2019  HB2 PRO A 148      53.421  14.321  38.770  1.00  0.00           H
ATOM   2020  CA  PRO A 148      51.642  15.092  39.585  1.00  0.00           C
ATOM   2021  HA  PRO A 148      51.460  14.244  40.083  1.00  0.00           H
ATOM   2022  C   PRO A 148      51.455  16.216  40.593  1.00  0.00           C
ATOM   2023  O   PRO A 148      51.144  17.362  40.281  1.00  0.00           O
ATOM   2024  N   ALA A 149      52.013  15.996  41.757  1.00  0.00           N
ATOM   2025  H   ALA A 149      52.475  15.118  41.882  1.00  0.00           H
ATOM   2026  CA  ALA A 149      52.021  16.895  42.861  1.00  0.00           C
ATOM   2027  HA  ALA A 149      51.065  17.178  42.934  1.00  0.00           H
ATOM   2028  CB  ALA A 149      52.375  16.174  44.142  1.00  0.00           C
ATOM   2029  HB1 ALA A 149      52.374  16.824  44.902  1.00  0.00           H
ATOM   2030  HB2 ALA A 149      51.702  15.456  44.317  1.00  0.00           H
ATOM   2031  HB3 ALA A 149      53.283  15.765  44.054  1.00  0.00           H
ATOM   2032  C   ALA A 149      52.832  18.173  42.596  1.00  0.00           C
ATOM   2033  O   ALA A 149      52.646  19.167  43.331  1.00  0.00           O
ATOM   2034  N   VAL A 150      53.695  18.204  41.569  1.00  0.00           N
ATOM   2035  H   VAL A 150      53.922  17.332  41.135  1.00  0.00           H
ATOM   2036  CA  VAL A 150      54.316  19.376  41.045  1.00  0.00           C
ATOM   2037  HA  VAL A 150      54.683  19.791  41.878  1.00  0.00           H
ATOM   2038  CB  VAL A 150      55.514  19.000  40.068  1.00  0.00           C
ATOM   2039  HB  VAL A 150      56.012  18.187  40.368  1.00  0.00           H
ATOM   2040  CG1 VAL A 150      55.050  18.593  38.712  1.00  0.00           C
ATOM   2041 HG11 VAL A 150      55.841  18.369  38.142  1.00  0.00           H
ATOM   2042 HG12 VAL A 150      54.457  17.792  38.789  1.00  0.00           H
ATOM   2043 HG13 VAL A 150      54.541  19.345  38.294  1.00  0.00           H
ATOM   2044  CG2 VAL A 150      56.448  20.220  40.008  1.00  0.00           C
ATOM   2045 HG21 VAL A 150      57.217  20.021  39.401  1.00  0.00           H
ATOM   2046 HG22 VAL A 150      55.943  21.010  39.659  1.00  0.00           H
ATOM   2047 HG23 VAL A 150      56.791  20.422  40.925  1.00  0.00           H
ATOM   2048  C   VAL A 150      53.297  20.424  40.494  1.00  0.00           C
ATOM   2049  O   VAL A 150      53.475  21.606  40.672  1.00  0.00           O
```

```
ATOM   2050  N   ASP A 151      52.253  19.976  39.833  1.00  0.00           N
ATOM   2051  H   ASP A 151      52.205  19.002  39.612  1.00  0.00           H
ATOM   2052  CA  ASP A 151      51.172  20.839  39.413  1.00  0.00           C
ATOM   2053  HA  ASP A 151      51.575  21.608  38.917  1.00  0.00           H
ATOM   2054  CB  ASP A 151      50.358  20.126  38.417  1.00  0.00           C
ATOM   2055  HB1 ASP A 151      49.466  20.576  38.371  1.00  0.00           H
ATOM   2056  HB2 ASP A 151      50.241  19.185  38.733  1.00  0.00           H
ATOM   2057  CG  ASP A 151      50.964  20.105  37.041  1.00  0.00           C
ATOM   2058  OD1 ASP A 151      50.294  19.542  36.127  1.00  0.00           O
ATOM   2059  OD2 ASP A 151      52.118  20.639  36.824  1.00  0.00           O
ATOM   2060  C   ASP A 151      50.352  21.358  40.670  1.00  0.00           C
ATOM   2061  O   ASP A 151      49.682  22.450  40.616  1.00  0.00           O
ATOM   2062  N   LYS A 152      50.259  20.603  41.769  1.00  0.00           N
ATOM   2063  H   LYS A 152      50.646  19.681  41.746  1.00  0.00           H
ATOM   2064  CA  LYS A 152      49.628  21.053  42.987  1.00  0.00           C
ATOM   2065  HA  LYS A 152      48.716  21.355  42.708  1.00  0.00           H
ATOM   2066  CB  LYS A 152      49.273  19.954  43.963  1.00  0.00           C
ATOM   2067  HB1 LYS A 152      50.115  19.508  44.266  1.00  0.00           H
ATOM   2068  HB2 LYS A 152      48.691  19.286  43.498  1.00  0.00           H
ATOM   2069  CG  LYS A 152      48.540  20.470  45.173  1.00  0.00           C
ATOM   2070  HG1 LYS A 152      47.682  20.892  44.879  1.00  0.00           H
ATOM   2071  HG2 LYS A 152      49.108  21.153  45.632  1.00  0.00           H
ATOM   2072  CD  LYS A 152      48.223  19.398  46.146  1.00  0.00           C
ATOM   2073  HD1 LYS A 152      47.720  18.692  45.648  1.00  0.00           H
ATOM   2074  HD2 LYS A 152      47.629  19.802  46.841  1.00  0.00           H
ATOM   2075  CE  LYS A 152      49.477  18.759  46.837  1.00  0.00           C
ATOM   2076  HE1 LYS A 152      50.184  19.442  47.022  1.00  0.00           H
ATOM   2077  HE2 LYS A 152      49.864  18.019  46.287  1.00  0.00           H
ATOM   2078  NZ  LYS A 152      49.043  18.208  48.091  1.00  0.00           N
ATOM   2079  HZ1 LYS A 152      49.822  17.791  48.559  1.00  0.00           H
ATOM   2080  HZ2 LYS A 152      48.662  18.936  48.661  1.00  0.00           H
ATOM   2081  HZ3 LYS A 152      48.343  17.514  47.925  1.00  0.00           H
ATOM   2082  C   LYS A 152      50.419  22.248  43.555  1.00  0.00           C
ATOM   2083  O   LYS A 152      49.885  23.291  43.824  1.00  0.00           O
ATOM   2084  N   GLU A 153      51.749  22.131  43.617  1.00  0.00           N
ATOM   2085  H   GLU A 153      52.155  21.268  43.317  1.00  0.00           H
ATOM   2086  CA  GLU A 153      52.696  23.229  44.117  1.00  0.00           C
ATOM   2087  HA  GLU A 153      52.246  23.571  44.942  1.00  0.00           H
ATOM   2088  CB  GLU A 153      54.110  22.664  44.460  1.00  0.00           C
ATOM   2089  HB1 GLU A 153      54.671  22.630  43.633  1.00  0.00           H
ATOM   2090  HB2 GLU A 153      54.023  21.744  44.842  1.00  0.00           H
ATOM   2091  CG  GLU A 153      54.809  23.538  45.475  1.00  0.00           C
ATOM   2092  HG1 GLU A 153      54.709  24.484  45.167  1.00  0.00           H
ATOM   2093  HG2 GLU A 153      55.776  23.284  45.465  1.00  0.00           H
ATOM   2094  CD  GLU A 153      54.298  23.446  46.934  1.00  0.00           C
ATOM   2095  OE1 GLU A 153      54.600  22.505  47.624  1.00  0.00           O
ATOM   2096  OE2 GLU A 153      53.633  24.345  47.502  1.00  0.00           O
ATOM   2097  C   GLU A 153      52.797  24.461  43.233  1.00  0.00           C
ATOM   2098  O   GLU A 153      52.686  25.563  43.736  1.00  0.00           O
ATOM   2099  N   ARG A 154      52.679  24.254  41.894  1.00  0.00           N
ATOM   2100  H   ARG A 154      52.596  23.303  41.596  1.00  0.00           H
ATOM   2101  CA  ARG A 154      52.660  25.275  40.839  1.00  0.00           C
ATOM   2102  HA  ARG A 154      53.453  25.871  40.970  1.00  0.00           H
ATOM   2103  CB  ARG A 154      52.713  24.558  39.511  1.00  0.00           C
```

| | | | | | | | | | | | |
|---|---|---|---|---|---|---|---|---|---|---|---|
| ATOM | 2104 | HB1 | ARG | A | 154 | 52.310 | 25.149 | 38.812 | 1.00 | 0.00 | H |
| ATOM | 2105 | HB2 | ARG | A | 154 | 52.181 | 23.714 | 39.582 | 1.00 | 0.00 | H |
| ATOM | 2106 | CG | ARG | A | 154 | 54.170 | 24.221 | 39.126 | 1.00 | 0.00 | C |
| ATOM | 2107 | HG1 | ARG | A | 154 | 54.632 | 23.748 | 39.876 | 1.00 | 0.00 | H |
| ATOM | 2108 | HG2 | ARG | A | 154 | 54.675 | 25.053 | 38.895 | 1.00 | 0.00 | H |
| ATOM | 2109 | CD | ARG | A | 154 | 54.081 | 23.310 | 37.910 | 1.00 | 0.00 | C |
| ATOM | 2110 | HD1 | ARG | A | 154 | 53.728 | 23.834 | 37.135 | 1.00 | 0.00 | H |
| ATOM | 2111 | HD2 | ARG | A | 154 | 53.457 | 22.555 | 38.113 | 1.00 | 0.00 | H |
| ATOM | 2112 | NE | ARG | A | 154 | 55.341 | 22.717 | 37.484 | 1.00 | 0.00 | N |
| ATOM | 2113 | HE | ARG | A | 154 | 56.162 | 23.119 | 37.890 | 1.00 | 0.00 | H |
| ATOM | 2114 | CZ | ARG | A | 154 | 55.559 | 21.712 | 36.628 | 1.00 | 0.00 | C |
| ATOM | 2115 | NH1 | ARG | A | 154 | 54.608 | 21.166 | 35.925 | 1.00 | 0.00 | N |
| ATOM | 2116 | HH11 | ARG | A | 154 | 53.668 | 21.496 | 36.013 | 1.00 | 0.00 | H |
| ATOM | 2117 | HH12 | ARG | A | 154 | 54.818 | 20.416 | 35.297 | 1.00 | 0.00 | H |
| ATOM | 2118 | NH2 | ARG | A | 154 | 56.812 | 21.456 | 36.375 | 1.00 | 0.00 | N |
| ATOM | 2119 | HH21 | ARG | A | 154 | 57.531 | 21.997 | 36.811 | 1.00 | 0.00 | H |
| ATOM | 2120 | HH22 | ARG | A | 154 | 57.054 | 20.719 | 35.745 | 1.00 | 0.00 | H |
| ATOM | 2121 | C | ARG | A | 154 | 51.469 | 26.172 | 40.927 | 1.00 | 0.00 | C |
| ATOM | 2122 | O | ARG | A | 154 | 51.507 | 27.382 | 40.781 | 1.00 | 0.00 | O |
| ATOM | 2123 | N | PHE | A | 155 | 50.275 | 25.630 | 41.246 | 1.00 | 0.00 | N |
| ATOM | 2124 | H | PHE | A | 155 | 50.201 | 24.633 | 41.276 | 1.00 | 0.00 | H |
| ATOM | 2125 | CA | PHE | A | 155 | 49.046 | 26.447 | 41.560 | 1.00 | 0.00 | C |
| ATOM | 2126 | HA | PHE | A | 155 | 48.993 | 27.195 | 40.898 | 1.00 | 0.00 | H |
| ATOM | 2127 | CB | PHE | A | 155 | 47.821 | 25.552 | 41.397 | 1.00 | 0.00 | C |
| ATOM | 2128 | HB1 | PHE | A | 155 | 47.748 | 24.971 | 42.208 | 1.00 | 0.00 | H |
| ATOM | 2129 | HB2 | PHE | A | 155 | 47.957 | 24.980 | 40.588 | 1.00 | 0.00 | H |
| ATOM | 2130 | CG | PHE | A | 155 | 46.510 | 26.224 | 41.231 | 1.00 | 0.00 | C |
| ATOM | 2131 | CD1 | PHE | A | 155 | 46.298 | 27.036 | 40.036 | 1.00 | 0.00 | C |
| ATOM | 2132 | HD1 | PHE | A | 155 | 47.024 | 27.175 | 39.363 | 1.00 | 0.00 | H |
| ATOM | 2133 | CE1 | PHE | A | 155 | 45.000 | 27.626 | 39.871 | 1.00 | 0.00 | C |
| ATOM | 2134 | HE1 | PHE | A | 155 | 44.772 | 28.182 | 39.072 | 1.00 | 0.00 | H |
| ATOM | 2135 | CZ | PHE | A | 155 | 44.074 | 27.383 | 40.885 | 1.00 | 0.00 | C |
| ATOM | 2136 | HZ | PHE | A | 155 | 43.198 | 27.862 | 40.822 | 1.00 | 0.00 | H |
| ATOM | 2137 | CE2 | PHE | A | 155 | 44.257 | 26.516 | 42.005 | 1.00 | 0.00 | C |
| ATOM | 2138 | HE2 | PHE | A | 155 | 43.502 | 26.287 | 42.620 | 1.00 | 0.00 | H |
| ATOM | 2139 | CD2 | PHE | A | 155 | 45.545 | 26.002 | 42.200 | 1.00 | 0.00 | C |
| ATOM | 2140 | HD2 | PHE | A | 155 | 45.765 | 25.483 | 43.026 | 1.00 | 0.00 | H |
| ATOM | 2141 | C | PHE | A | 155 | 49.113 | 27.157 | 42.924 | 1.00 | 0.00 | C |
| ATOM | 2142 | O | PHE | A | 155 | 48.764 | 28.341 | 43.102 | 1.00 | 0.00 | O |
| ATOM | 2143 | N | THR | A | 156 | 49.639 | 26.431 | 43.919 | 1.00 | 0.00 | N |
| ATOM | 2144 | H | THR | A | 156 | 49.922 | 25.487 | 43.747 | 1.00 | 0.00 | H |
| ATOM | 2145 | CA | THR | A | 156 | 49.807 | 27.017 | 45.284 | 1.00 | 0.00 | C |
| ATOM | 2146 | HA | THR | A | 156 | 48.879 | 27.212 | 45.603 | 1.00 | 0.00 | H |
| ATOM | 2147 | CB | THR | A | 156 | 50.387 | 26.009 | 46.281 | 1.00 | 0.00 | C |
| ATOM | 2148 | HB | THR | A | 156 | 51.308 | 25.713 | 46.028 | 1.00 | 0.00 | H |
| ATOM | 2149 | CG2 | THR | A | 156 | 50.435 | 26.577 | 47.676 | 1.00 | 0.00 | C |
| ATOM | 2150 | HG21 | THR | A | 156 | 50.817 | 25.896 | 48.300 | 1.00 | 0.00 | H |
| ATOM | 2151 | HG22 | THR | A | 156 | 51.010 | 27.395 | 47.683 | 1.00 | 0.00 | H |
| ATOM | 2152 | HG23 | THR | A | 156 | 49.510 | 26.817 | 47.970 | 1.00 | 0.00 | H |
| ATOM | 2153 | OG1 | THR | A | 156 | 49.585 | 24.862 | 46.249 | 1.00 | 0.00 | O |
| ATOM | 2154 | HG1 | THR | A | 156 | 49.944 | 24.186 | 46.893 | 1.00 | 0.00 | H |
| ATOM | 2155 | C | THR | A | 156 | 50.690 | 28.339 | 45.207 | 1.00 | 0.00 | C |
| ATOM | 2156 | O | THR | A | 156 | 50.369 | 29.435 | 45.641 | 1.00 | 0.00 | O |
| ATOM | 2157 | N | THR | A | 157 | 51.811 | 28.165 | 44.474 | 1.00 | 0.00 | N |

```
ATOM   2158  H   THR A 157      51.958  27.289  44.014  1.00  0.00           H
ATOM   2159  CA  THR A 157      52.818  29.219  44.331  1.00  0.00           C
ATOM   2160  HA  THR A 157      52.969  29.518  45.273  1.00  0.00           H
ATOM   2161  CB  THR A 157      54.079  28.663  43.714  1.00  0.00           C
ATOM   2162  HB  THR A 157      53.838  28.287  42.820  1.00  0.00           H
ATOM   2163  CG2 THR A 157      55.220  29.717  43.532  1.00  0.00           C
ATOM   2164 HG21 THR A 157      56.018  29.277  43.120  1.00  0.00           H
ATOM   2165 HG22 THR A 157      54.901  30.454  42.936  1.00  0.00           H
ATOM   2166 HG23 THR A 157      55.471  30.094  44.424  1.00  0.00           H
ATOM   2167  OG1 THR A 157      54.509  27.537  44.429  1.00  0.00           O
ATOM   2168  HG1 THR A 157      55.344  27.177  44.013  1.00  0.00           H
ATOM   2169  C   THR A 157      52.346  30.397  43.444  1.00  0.00           C
ATOM   2170  O   THR A 157      52.754  31.550  43.717  1.00  0.00           O
ATOM   2171  N   LEU A 158      51.524  30.199  42.404  1.00  0.00           N
ATOM   2172  H   LEU A 158      51.276  29.262  42.158  1.00  0.00           H
ATOM   2173  CA  LEU A 158      50.993  31.269  41.640  1.00  0.00           C
ATOM   2174  HA  LEU A 158      51.785  31.868  41.525  1.00  0.00           H
ATOM   2175  CB  LEU A 158      50.456  30.729  40.272  1.00  0.00           C
ATOM   2176  HB1 LEU A 158      49.587  30.257  40.419  1.00  0.00           H
ATOM   2177  HB2 LEU A 158      51.120  30.091  39.882  1.00  0.00           H
ATOM   2178  CG  LEU A 158      50.230  31.874  39.274  1.00  0.00           C
ATOM   2179  HG  LEU A 158      49.745  32.566  39.809  1.00  0.00           H
ATOM   2180  CD1 LEU A 158      51.423  32.664  38.842  1.00  0.00           C
ATOM   2181 HD11 LEU A 158      51.138  33.373  38.197  1.00  0.00           H
ATOM   2182 HD12 LEU A 158      51.847  33.091  39.641  1.00  0.00           H
ATOM   2183 HD13 LEU A 158      52.083  32.056  38.400  1.00  0.00           H
ATOM   2184  CD2 LEU A 158      49.371  31.429  38.090  1.00  0.00           C
ATOM   2185 HD21 LEU A 158      49.245  32.197  37.462  1.00  0.00           H
ATOM   2186 HD22 LEU A 158      49.827  30.679  37.611  1.00  0.00           H
ATOM   2187 HD23 LEU A 158      48.480  31.120  38.422  1.00  0.00           H
ATOM   2188  C   LEU A 158      49.827  31.992  42.414  1.00  0.00           C
ATOM   2189  O   LEU A 158      49.780  33.238  42.446  1.00  0.00           O
ATOM   2190  N   VAL A 159      48.896  31.189  43.039  1.00  0.00           N
ATOM   2191  H   VAL A 159      48.922  30.202  42.879  1.00  0.00           H
ATOM   2192  CA  VAL A 159      47.855  31.744  43.943  1.00  0.00           C
ATOM   2193  HA  VAL A 159      47.190  32.227  43.373  1.00  0.00           H
ATOM   2194  CB  VAL A 159      47.035  30.674  44.676  1.00  0.00           C
ATOM   2195  HB  VAL A 159      47.723  30.006  44.958  1.00  0.00           H
ATOM   2196  CG1 VAL A 159      46.329  31.129  45.937  1.00  0.00           C
ATOM   2197 HG11 VAL A 159      45.827  30.360  46.334  1.00  0.00           H
ATOM   2198 HG12 VAL A 159      47.004  31.462  46.595  1.00  0.00           H
ATOM   2199 HG13 VAL A 159      45.690  31.865  45.714  1.00  0.00           H
ATOM   2200  CG2 VAL A 159      46.001  30.042  43.771  1.00  0.00           C
ATOM   2201 HG21 VAL A 159      45.488  29.352  44.282  1.00  0.00           H
ATOM   2202 HG22 VAL A 159      45.373  30.746  43.439  1.00  0.00           H
ATOM   2203 HG23 VAL A 159      46.458  29.610  42.994  1.00  0.00           H
ATOM   2204  C   VAL A 159      48.588  32.676  44.933  1.00  0.00           C
ATOM   2205  O   VAL A 159      48.078  33.777  45.205  1.00  0.00           O
ATOM   2206  N   GLN A 160      49.739  32.254  45.461  1.00  0.00           N
ATOM   2207  H   GLN A 160      50.085  31.345  45.227  1.00  0.00           H
ATOM   2208  CA  GLN A 160      50.513  33.114  46.390  1.00  0.00           C
ATOM   2209  HA  GLN A 160      49.857  33.545  47.010  1.00  0.00           H
ATOM   2210  CB  GLN A 160      51.477  32.159  47.072  1.00  0.00           C
ATOM   2211  HB1 GLN A 160      52.218  31.963  46.430  1.00  0.00           H
```

```
ATOM   2212  HB2 GLN A 160      50.980  31.315  47.274  1.00  0.00           H
ATOM   2213  CG  GLN A 160      52.092  32.690  48.382  1.00  0.00           C
ATOM   2214  HG1 GLN A 160      51.350  32.798  49.043  1.00  0.00           H
ATOM   2215  HG2 GLN A 160      52.497  33.583  48.186  1.00  0.00           H
ATOM   2216  CD  GLN A 160      53.167  31.857  49.058  1.00  0.00           C
ATOM   2217  OE1 GLN A 160      53.490  30.781  48.706  1.00  0.00           O
ATOM   2218  NE2 GLN A 160      53.765  32.364  50.130  1.00  0.00           N
ATOM   2219 HE21 GLN A 160      53.503  33.268  50.468  1.00  0.00           H
ATOM   2220 HE22 GLN A 160      54.477  31.842  50.599  1.00  0.00           H
ATOM   2221  C   GLN A 160      51.212  34.345  45.797  1.00  0.00           C
ATOM   2222  O   GLN A 160      51.326  35.379  46.417  1.00  0.00           O
ATOM   2223  N   ASP A 161      51.698  34.224  44.593  1.00  0.00           N
ATOM   2224  H   ASP A 161      51.625  33.330  44.150  1.00  0.00           H
ATOM   2225  CA  ASP A 161      52.333  35.288  43.854  1.00  0.00           C
ATOM   2226  HA  ASP A 161      52.969  35.729  44.487  1.00  0.00           H
ATOM   2227  CB  ASP A 161      53.044  34.639  42.613  1.00  0.00           C
ATOM   2228  HB1 ASP A 161      53.138  35.326  41.892  1.00  0.00           H
ATOM   2229  HB2 ASP A 161      52.489  33.878  42.278  1.00  0.00           H
ATOM   2230  CG  ASP A 161      54.411  34.095  42.871  1.00  0.00           C
ATOM   2231  OD1 ASP A 161      55.122  33.721  41.892  1.00  0.00           O
ATOM   2232  OD2 ASP A 161      54.860  34.052  44.042  1.00  0.00           O
ATOM   2233  C   ASP A 161      51.286  36.327  43.389  1.00  0.00           C
ATOM   2234  O   ASP A 161      51.597  37.544  43.353  1.00  0.00           O
ATOM   2235  N   LEU A 162      50.000  35.938  43.029  1.00  0.00           N
ATOM   2236  H   LEU A 162      49.735  34.975  43.081  1.00  0.00           H
ATOM   2237  CA  LEU A 162      49.068  36.905  42.590  1.00  0.00           C
ATOM   2238  HA  LEU A 162      49.566  37.554  42.015  1.00  0.00           H
ATOM   2239  CB  LEU A 162      47.966  36.218  41.784  1.00  0.00           C
ATOM   2240  HB1 LEU A 162      47.171  36.824  41.757  1.00  0.00           H
ATOM   2241  HB2 LEU A 162      47.721  35.367  42.248  1.00  0.00           H
ATOM   2242  CG  LEU A 162      48.439  35.902  40.316  1.00  0.00           C
ATOM   2243  HG  LEU A 162      49.424  35.727  40.300  1.00  0.00           H
ATOM   2244  CD1 LEU A 162      47.563  34.688  39.813  1.00  0.00           C
ATOM   2245 HD11 LEU A 162      47.829  34.450  38.879  1.00  0.00           H
ATOM   2246 HD12 LEU A 162      47.709  33.901  40.412  1.00  0.00           H
ATOM   2247 HD13 LEU A 162      46.597  34.945  39.829  1.00  0.00           H
ATOM   2248  CD2 LEU A 162      48.412  37.044  39.374  1.00  0.00           C
ATOM   2249 HD21 LEU A 162      48.728  36.742  38.475  1.00  0.00           H
ATOM   2250 HD22 LEU A 162      47.478  37.392  39.299  1.00  0.00           H
ATOM   2251 HD23 LEU A 162      49.012  37.769  39.712  1.00  0.00           H
ATOM   2252  C   LEU A 162      48.493  37.589  43.816  1.00  0.00           C
ATOM   2253  O   LEU A 162      48.113  38.764  43.692  1.00  0.00           O
ATOM   2254  N   ALA A 163      48.340  36.936  44.988  1.00  0.00           N
ATOM   2255  H   ALA A 163      48.517  35.952  45.004  1.00  0.00           H
ATOM   2256  CA  ALA A 163      47.938  37.570  46.212  1.00  0.00           C
ATOM   2257  HA  ALA A 163      47.086  38.083  46.106  1.00  0.00           H
ATOM   2258  CB  ALA A 163      47.681  36.420  47.148  1.00  0.00           C
ATOM   2259  HB1 ALA A 163      47.392  36.773  48.038  1.00  0.00           H
ATOM   2260  HB2 ALA A 163      46.961  35.838  46.771  1.00  0.00           H
ATOM   2261  HB3 ALA A 163      48.519  35.886  47.257  1.00  0.00           H
ATOM   2262  C   ALA A 163      48.931  38.669  46.642  1.00  0.00           C
ATOM   2263  O   ALA A 163      48.430  39.782  47.008  1.00  0.00           O
ATOM   2264  N   ASN A 164      50.230  38.328  46.540  1.00  0.00           N
ATOM   2265  H   ASN A 164      50.470  37.381  46.324  1.00  0.00           H
```

| | | | | | | | | | | | |
|---|---|---|---|---|---|---|---|---|---|---|---|
| ATOM | 2266 | CA | ASN | A | 164 | 51.301 | 39.309 | 46.737 | 1.00 | 0.00 | C |
| ATOM | 2267 | HA | ASN | A | 164 | 51.106 | 39.568 | 47.683 | 1.00 | 0.00 | H |
| ATOM | 2268 | CB | ASN | A | 164 | 52.738 | 38.696 | 46.541 | 1.00 | 0.00 | C |
| ATOM | 2269 | HB1 | ASN | A | 164 | 52.874 | 38.494 | 45.571 | 1.00 | 0.00 | H |
| ATOM | 2270 | HB2 | ASN | A | 164 | 52.804 | 37.850 | 47.070 | 1.00 | 0.00 | H |
| ATOM | 2271 | CG | ASN | A | 164 | 53.882 | 39.603 | 46.984 | 1.00 | 0.00 | C |
| ATOM | 2272 | OD1 | ASN | A | 164 | 53.881 | 40.822 | 46.846 | 1.00 | 0.00 | O |
| ATOM | 2273 | ND2 | ASN | A | 164 | 54.954 | 39.020 | 47.458 | 1.00 | 0.00 | N |
| ATOM | 2274 | HD21 | ASN | A | 164 | 54.996 | 38.023 | 47.517 | 1.00 | 0.00 | H |
| ATOM | 2275 | HD22 | ASN | A | 164 | 55.731 | 39.572 | 47.761 | 1.00 | 0.00 | H |
| ATOM | 2276 | C | ASN | A | 164 | 51.192 | 40.563 | 45.886 | 1.00 | 0.00 | C |
| ATOM | 2277 | O | ASN | A | 164 | 50.739 | 41.551 | 46.456 | 1.00 | 0.00 | O |
| ATOM | 2278 | N | ALA | A | 165 | 51.288 | 40.473 | 44.519 | 1.00 | 0.00 | N |
| ATOM | 2279 | H | ALA | A | 165 | 51.513 | 39.568 | 44.157 | 1.00 | 0.00 | H |
| ATOM | 2280 | CA | ALA | A | 165 | 51.108 | 41.507 | 43.544 | 1.00 | 0.00 | C |
| ATOM | 2281 | HA | ALA | A | 165 | 51.883 | 42.138 | 43.580 | 1.00 | 0.00 | H |
| ATOM | 2282 | CB | ALA | A | 165 | 50.998 | 40.743 | 42.188 | 1.00 | 0.00 | C |
| ATOM | 2283 | HB1 | ALA | A | 165 | 50.869 | 41.400 | 41.445 | 1.00 | 0.00 | H |
| ATOM | 2284 | HB2 | ALA | A | 165 | 51.837 | 40.221 | 42.030 | 1.00 | 0.00 | H |
| ATOM | 2285 | HB3 | ALA | A | 165 | 50.218 | 40.118 | 42.220 | 1.00 | 0.00 | H |
| ATOM | 2286 | C | ALA | A | 165 | 49.882 | 42.364 | 43.871 | 1.00 | 0.00 | C |
| ATOM | 2287 | O | ALA | A | 165 | 49.991 | 43.562 | 44.014 | 1.00 | 0.00 | O |
| ATOM | 2288 | N | PHE | A | 166 | 48.695 | 41.781 | 44.021 | 1.00 | 0.00 | N |
| ATOM | 2289 | H | PHE | A | 166 | 48.665 | 40.785 | 43.932 | 1.00 | 0.00 | H |
| ATOM | 2290 | CA | PHE | A | 166 | 47.459 | 42.443 | 44.297 | 1.00 | 0.00 | C |
| ATOM | 2291 | HA | PHE | A | 166 | 47.404 | 43.160 | 43.602 | 1.00 | 0.00 | H |
| ATOM | 2292 | CB | PHE | A | 166 | 46.280 | 41.434 | 44.189 | 1.00 | 0.00 | C |
| ATOM | 2293 | HB1 | PHE | A | 166 | 45.481 | 41.850 | 44.624 | 1.00 | 0.00 | H |
| ATOM | 2294 | HB2 | PHE | A | 166 | 46.539 | 40.607 | 44.689 | 1.00 | 0.00 | H |
| ATOM | 2295 | CG | PHE | A | 166 | 45.869 | 41.009 | 42.753 | 1.00 | 0.00 | C |
| ATOM | 2296 | CD1 | PHE | A | 166 | 46.589 | 41.297 | 41.591 | 1.00 | 0.00 | C |
| ATOM | 2297 | HD1 | PHE | A | 166 | 47.448 | 41.796 | 41.701 | 1.00 | 0.00 | H |
| ATOM | 2298 | CE1 | PHE | A | 166 | 46.223 | 40.956 | 40.308 | 1.00 | 0.00 | C |
| ATOM | 2299 | HE1 | PHE | A | 166 | 46.812 | 41.144 | 39.522 | 1.00 | 0.00 | H |
| ATOM | 2300 | CZ | PHE | A | 166 | 45.002 | 40.342 | 40.172 | 1.00 | 0.00 | C |
| ATOM | 2301 | HZ | PHE | A | 166 | 44.645 | 40.186 | 39.251 | 1.00 | 0.00 | H |
| ATOM | 2302 | CE2 | PHE | A | 166 | 44.236 | 39.925 | 41.267 | 1.00 | 0.00 | C |
| ATOM | 2303 | HE2 | PHE | A | 166 | 43.380 | 39.426 | 41.128 | 1.00 | 0.00 | H |
| ATOM | 2304 | CD2 | PHE | A | 166 | 44.688 | 40.218 | 42.562 | 1.00 | 0.00 | C |
| ATOM | 2305 | HD2 | PHE | A | 166 | 44.185 | 39.874 | 43.355 | 1.00 | 0.00 | H |
| ATOM | 2306 | C | PHE | A | 166 | 47.474 | 43.179 | 45.587 | 1.00 | 0.00 | C |
| ATOM | 2307 | O | PHE | A | 166 | 47.009 | 44.324 | 45.566 | 1.00 | 0.00 | O |
| ATOM | 2308 | N | GLN | A | 167 | 47.892 | 42.619 | 46.759 | 1.00 | 0.00 | N |
| ATOM | 2309 | H | GLN | A | 167 | 48.025 | 41.628 | 46.750 | 1.00 | 0.00 | H |
| ATOM | 2310 | CA | GLN | A | 167 | 48.170 | 43.298 | 48.033 | 1.00 | 0.00 | C |
| ATOM | 2311 | HA | GLN | A | 167 | 47.278 | 43.642 | 48.327 | 1.00 | 0.00 | H |
| ATOM | 2312 | CB | GLN | A | 167 | 48.788 | 42.296 | 49.059 | 1.00 | 0.00 | C |
| ATOM | 2313 | HB1 | GLN | A | 167 | 49.517 | 41.780 | 48.610 | 1.00 | 0.00 | H |
| ATOM | 2314 | HB2 | GLN | A | 167 | 48.075 | 41.666 | 49.367 | 1.00 | 0.00 | H |
| ATOM | 2315 | CG | GLN | A | 167 | 49.383 | 42.867 | 50.278 | 1.00 | 0.00 | C |
| ATOM | 2316 | HG1 | GLN | A | 167 | 50.133 | 43.466 | 49.996 | 1.00 | 0.00 | H |
| ATOM | 2317 | HG2 | GLN | A | 167 | 49.747 | 42.110 | 50.820 | 1.00 | 0.00 | H |
| ATOM | 2318 | CD | GLN | A | 167 | 48.445 | 43.656 | 51.139 | 1.00 | 0.00 | C |
| ATOM | 2319 | OE1 | GLN | A | 167 | 47.244 | 43.735 | 50.925 | 1.00 | 0.00 | O |

```
ATOM   2320  NE2 GLN A 167      48.952  44.448  52.070  1.00  0.00           N
ATOM   2321 HE21 GLN A 167      49.943  44.511  52.188  1.00  0.00           H
ATOM   2322 HE22 GLN A 167      48.344  44.984  52.656  1.00  0.00           H
ATOM   2323  C   GLN A 167      49.139  44.474  47.850  1.00  0.00           C
ATOM   2324  O   GLN A 167      48.832  45.629  48.168  1.00  0.00           O
ATOM   2325  N   GLN A 168      50.315  44.223  47.244  1.00  0.00           N
ATOM   2326  H   GLN A 168      50.451  43.299  46.888  1.00  0.00           H
ATOM   2327  CA  GLN A 168      51.397  45.157  47.059  1.00  0.00           C
ATOM   2328  HA  GLN A 168      51.449  45.562  47.972  1.00  0.00           H
ATOM   2329  CB  GLN A 168      52.676  44.373  46.597  1.00  0.00           C
ATOM   2330  HB1 GLN A 168      52.519  44.039  45.668  1.00  0.00           H
ATOM   2331  HB2 GLN A 168      52.806  43.596  47.213  1.00  0.00           H
ATOM   2332  CG  GLN A 168      53.995  45.192  46.587  1.00  0.00           C
ATOM   2333  HG1 GLN A 168      53.972  45.812  45.803  1.00  0.00           H
ATOM   2334  HG2 GLN A 168      54.758  44.554  46.485  1.00  0.00           H
ATOM   2335  CD  GLN A 168      54.274  46.036  47.823  1.00  0.00           C
ATOM   2336  OE1 GLN A 168      54.091  45.572  48.963  1.00  0.00           O
ATOM   2337  NE2 GLN A 168      54.625  47.286  47.661  1.00  0.00           N
ATOM   2338 HE21 GLN A 168      54.705  47.668  46.740  1.00  0.00           H
ATOM   2339 HE22 GLN A 168      54.813  47.860  48.458  1.00  0.00           H
ATOM   2340  C   GLN A 168      51.128  46.316  46.077  1.00  0.00           C
ATOM   2341  O   GLN A 168      51.658  47.417  46.233  1.00  0.00           O
ATOM   2342  N   GLU A 169      50.154  46.130  45.140  1.00  0.00           N
ATOM   2343  H   GLU A 169      49.795  45.204  45.024  1.00  0.00           H
ATOM   2344  CA  GLU A 169      49.591  47.189  44.287  1.00  0.00           C
ATOM   2345  HA  GLU A 169      50.371  47.793  44.121  1.00  0.00           H
ATOM   2346  CB  GLU A 169      49.066  46.626  42.920  1.00  0.00           C
ATOM   2347  HB1 GLU A 169      48.275  46.053  43.134  1.00  0.00           H
ATOM   2348  HB2 GLU A 169      49.799  46.057  42.547  1.00  0.00           H
ATOM   2349  CG  GLU A 169      48.620  47.611  41.769  1.00  0.00           C
ATOM   2350  HG1 GLU A 169      48.578  47.116  40.901  1.00  0.00           H
ATOM   2351  HG2 GLU A 169      49.279  48.360  41.694  1.00  0.00           H
ATOM   2352  CD  GLU A 169      47.231  48.209  42.058  1.00  0.00           C
ATOM   2353  OE1 GLU A 169      46.502  47.771  42.974  1.00  0.00           O
ATOM   2354  OE2 GLU A 169      46.806  49.002  41.258  1.00  0.00           O
ATOM   2355  C   GLU A 169      48.673  48.130  44.988  1.00  0.00           C
ATOM   2356  O   GLU A 169      48.592  49.351  44.844  1.00  0.00           O
ATOM   2357  N   ALA A 170      47.907  47.568  45.940  1.00  0.00           N
ATOM   2358  H   ALA A 170      47.902  46.572  46.027  1.00  0.00           H
ATOM   2359  CA  ALA A 170      47.119  48.322  46.812  1.00  0.00           C
ATOM   2360  HA  ALA A 170      46.689  48.993  46.208  1.00  0.00           H
ATOM   2361  CB  ALA A 170      46.008  47.459  47.366  1.00  0.00           C
ATOM   2362  HB1 ALA A 170      45.445  48.001  47.990  1.00  0.00           H
ATOM   2363  HB2 ALA A 170      45.442  47.121  46.614  1.00  0.00           H
ATOM   2364  HB3 ALA A 170      46.403  46.684  47.860  1.00  0.00           H
ATOM   2365  C   ALA A 170      47.995  49.075  47.908  1.00  0.00           C
ATOM   2366  O   ALA A 170      47.673  50.228  48.236  1.00  0.00           O
ATOM   2367  N   GLN A 171      49.147  48.579  48.256  1.00  0.00           N
ATOM   2368  H   GLN A 171      49.354  47.622  48.053  1.00  0.00           H
ATOM   2369  CA  GLN A 171      50.129  49.393  48.931  1.00  0.00           C
ATOM   2370  HA  GLN A 171      49.617  49.900  49.625  1.00  0.00           H
ATOM   2371  CB  GLN A 171      51.193  48.437  49.536  1.00  0.00           C
ATOM   2372  HB1 GLN A 171      51.967  48.987  49.851  1.00  0.00           H
ATOM   2373  HB2 GLN A 171      51.498  47.812  48.818  1.00  0.00           H
```

| | | | | | | | | | | | |
|---|---|---|---|---|---|---|---|---|---|---|---|
| ATOM | 2374 | CG | GLN | A | 171 | 50.648 | 47.593 | 50.743 | 1.00 | 0.00 | C |
| ATOM | 2375 | HG1 | GLN | A | 171 | 51.394 | 47.073 | 51.159 | 1.00 | 0.00 | H |
| ATOM | 2376 | HG2 | GLN | A | 171 | 49.939 | 46.963 | 50.425 | 1.00 | 0.00 | H |
| ATOM | 2377 | CD | GLN | A | 171 | 50.046 | 48.509 | 51.800 | 1.00 | 0.00 | C |
| ATOM | 2378 | OE1 | GLN | A | 171 | 48.904 | 48.905 | 51.693 | 1.00 | 0.00 | O |
| ATOM | 2379 | NE2 | GLN | A | 171 | 50.859 | 48.900 | 52.758 | 1.00 | 0.00 | N |
| ATOM | 2380 | HE21 | GLN | A | 171 | 51.811 | 48.595 | 52.761 | 1.00 | 0.00 | H |
| ATOM | 2381 | HE22 | GLN | A | 171 | 50.525 | 49.503 | 53.483 | 1.00 | 0.00 | H |
| ATOM | 2382 | C | GLN | A | 171 | 50.697 | 50.519 | 48.120 | 1.00 | 0.00 | C |
| ATOM | 2383 | O | GLN | A | 171 | 51.460 | 51.259 | 48.634 | 1.00 | 0.00 | O |
| ATOM | 2384 | N | THR | A | 172 | 50.372 | 50.576 | 46.778 | 1.00 | 0.00 | N |
| ATOM | 2385 | H | THR | A | 172 | 49.693 | 49.928 | 46.431 | 1.00 | 0.00 | H |
| ATOM | 2386 | CA | THR | A | 172 | 50.972 | 51.545 | 45.814 | 1.00 | 0.00 | C |
| ATOM | 2387 | HA | THR | A | 172 | 51.627 | 52.107 | 46.319 | 1.00 | 0.00 | H |
| ATOM | 2388 | CB | THR | A | 172 | 51.807 | 50.839 | 44.759 | 1.00 | 0.00 | C |
| ATOM | 2389 | HB | THR | A | 172 | 51.167 | 50.211 | 44.316 | 1.00 | 0.00 | H |
| ATOM | 2390 | CG2 | THR | A | 172 | 52.486 | 51.735 | 43.747 | 1.00 | 0.00 | C |
| ATOM | 2391 | HG21 | THR | A | 172 | 53.006 | 51.175 | 43.102 | 1.00 | 0.00 | H |
| ATOM | 2392 | HG22 | THR | A | 172 | 51.795 | 52.260 | 43.250 | 1.00 | 0.00 | H |
| ATOM | 2393 | HG23 | THR | A | 172 | 53.105 | 52.362 | 44.219 | 1.00 | 0.00 | H |
| ATOM | 2394 | OG1 | THR | A | 172 | 52.822 | 50.080 | 45.311 | 1.00 | 0.00 | O |
| ATOM | 2395 | HG1 | THR | A | 172 | 53.342 | 49.637 | 44.581 | 1.00 | 0.00 | H |
| ATOM | 2396 | C | THR | A | 172 | 49.927 | 52.526 | 45.281 | 1.00 | 0.00 | C |
| ATOM | 2397 | O | THR | A | 172 | 49.947 | 53.713 | 45.625 | 1.00 | 0.00 | O |
| ATOM | 2398 | N | SER | A | 173 | 48.982 | 52.004 | 44.446 | 1.00 | 0.00 | N |
| ATOM | 2399 | H | SER | A | 173 | 48.981 | 51.019 | 44.272 | 1.00 | 0.00 | H |
| ATOM | 2400 | CA | SER | A | 173 | 48.000 | 52.815 | 43.816 | 1.00 | 0.00 | C |
| ATOM | 2401 | HA | SER | A | 173 | 48.529 | 53.539 | 43.373 | 1.00 | 0.00 | H |
| ATOM | 2402 | CB | SER | A | 173 | 47.182 | 51.882 | 42.876 | 1.00 | 0.00 | C |
| ATOM | 2403 | HB1 | SER | A | 173 | 47.790 | 51.296 | 42.341 | 1.00 | 0.00 | H |
| ATOM | 2404 | HB2 | SER | A | 173 | 46.606 | 52.417 | 42.258 | 1.00 | 0.00 | H |
| ATOM | 2405 | OG | SER | A | 173 | 46.319 | 51.020 | 43.546 | 1.00 | 0.00 | O |
| ATOM | 2406 | HG | SER | A | 173 | 45.830 | 50.455 | 42.882 | 1.00 | 0.00 | H |
| ATOM | 2407 | C | SER | A | 173 | 47.004 | 53.421 | 44.822 | 1.00 | 0.00 | C |
| ATOM | 2408 | O | SER | A | 173 | 46.429 | 54.492 | 44.608 | 1.00 | 0.00 | O |
| ATOM | 2409 | N | GLY | A | 174 | 46.725 | 52.737 | 45.953 | 1.00 | 0.00 | N |
| ATOM | 2410 | H | GLY | A | 174 | 47.214 | 51.872 | 46.068 | 1.00 | 0.00 | H |
| ATOM | 2411 | CA | GLY | A | 174 | 45.824 | 53.084 | 47.008 | 1.00 | 0.00 | C |
| ATOM | 2412 | HA1 | GLY | A | 174 | 46.010 | 54.039 | 47.240 | 1.00 | 0.00 | H |
| ATOM | 2413 | HA2 | GLY | A | 174 | 46.057 | 52.500 | 47.786 | 1.00 | 0.00 | H |
| ATOM | 2414 | C | GLY | A | 174 | 44.321 | 52.898 | 46.611 | 1.00 | 0.00 | C |
| ATOM | 2415 | O | GLY | A | 174 | 43.412 | 53.208 | 47.443 | 1.00 | 0.00 | O |
| ATOM | 2416 | N | LYS | A | 175 | 44.113 | 52.202 | 45.458 | 1.00 | 0.00 | N |
| ATOM | 2417 | H | LYS | A | 175 | 44.924 | 51.907 | 44.952 | 1.00 | 0.00 | H |
| ATOM | 2418 | CA | LYS | A | 175 | 42.858 | 51.860 | 44.913 | 1.00 | 0.00 | C |
| ATOM | 2419 | HA | LYS | A | 175 | 42.309 | 52.690 | 45.009 | 1.00 | 0.00 | H |
| ATOM | 2420 | CB | LYS | A | 175 | 42.990 | 51.620 | 43.421 | 1.00 | 0.00 | C |
| ATOM | 2421 | HB1 | LYS | A | 175 | 42.105 | 51.276 | 43.108 | 1.00 | 0.00 | H |
| ATOM | 2422 | HB2 | LYS | A | 175 | 43.683 | 50.909 | 43.306 | 1.00 | 0.00 | H |
| ATOM | 2423 | CG | LYS | A | 175 | 43.390 | 52.855 | 42.527 | 1.00 | 0.00 | C |
| ATOM | 2424 | HG1 | LYS | A | 175 | 44.289 | 53.196 | 42.800 | 1.00 | 0.00 | H |
| ATOM | 2425 | HG2 | LYS | A | 175 | 42.712 | 53.583 | 42.628 | 1.00 | 0.00 | H |
| ATOM | 2426 | CD | LYS | A | 175 | 43.453 | 52.464 | 41.064 | 1.00 | 0.00 | C |
| ATOM | 2427 | HD1 | LYS | A | 175 | 42.736 | 51.790 | 40.886 | 1.00 | 0.00 | H |

| | | | | | | | | | | | |
|---|---|---|---|---|---|---|---|---|---|---|---|
| ATOM | 2428 | HD2 | LYS | A | 175 | 44.348 | 52.055 | 40.884 | 1.00 | 0.00 | H |
| ATOM | 2429 | CE | LYS | A | 175 | 43.253 | 53.672 | 40.130 | 1.00 | 0.00 | C |
| ATOM | 2430 | HE1 | LYS | A | 175 | 43.983 | 54.341 | 40.272 | 1.00 | 0.00 | H |
| ATOM | 2431 | HE2 | LYS | A | 175 | 42.368 | 54.102 | 40.308 | 1.00 | 0.00 | H |
| ATOM | 2432 | NZ | LYS | A | 175 | 43.282 | 53.262 | 38.725 | 1.00 | 0.00 | N |
| ATOM | 2433 | HZ1 | LYS | A | 175 | 43.150 | 54.063 | 38.141 | 1.00 | 0.00 | H |
| ATOM | 2434 | HZ2 | LYS | A | 175 | 44.165 | 52.841 | 38.519 | 1.00 | 0.00 | H |
| ATOM | 2435 | HZ3 | LYS | A | 175 | 42.550 | 52.602 | 38.555 | 1.00 | 0.00 | H |
| ATOM | 2436 | C | LYS | A | 175 | 42.167 | 50.705 | 45.738 | 1.00 | 0.00 | C |
| ATOM | 2437 | O | LYS | A | 175 | 42.688 | 50.167 | 46.676 | 1.00 | 0.00 | O |
| ATOM | 2438 | N | GLU | A | 176 | 40.986 | 50.253 | 45.247 | 1.00 | 0.00 | N |
| ATOM | 2439 | H | GLU | A | 176 | 40.506 | 50.761 | 44.532 | 1.00 | 0.00 | H |
| ATOM | 2440 | CA | GLU | A | 176 | 40.459 | 49.084 | 45.749 | 1.00 | 0.00 | C |
| ATOM | 2441 | HA | GLU | A | 176 | 40.613 | 49.349 | 46.701 | 1.00 | 0.00 | H |
| ATOM | 2442 | CB | GLU | A | 176 | 38.964 | 48.889 | 45.377 | 1.00 | 0.00 | C |
| ATOM | 2443 | HB1 | GLU | A | 176 | 38.740 | 47.914 | 45.373 | 1.00 | 0.00 | H |
| ATOM | 2444 | HB2 | GLU | A | 176 | 38.787 | 49.275 | 44.472 | 1.00 | 0.00 | H |
| ATOM | 2445 | CG | GLU | A | 176 | 38.071 | 49.578 | 46.371 | 1.00 | 0.00 | C |
| ATOM | 2446 | HG1 | GLU | A | 176 | 38.450 | 49.463 | 47.289 | 1.00 | 0.00 | H |
| ATOM | 2447 | HG2 | GLU | A | 176 | 37.159 | 49.169 | 46.333 | 1.00 | 0.00 | H |
| ATOM | 2448 | CD | GLU | A | 176 | 37.979 | 51.056 | 46.040 | 1.00 | 0.00 | C |
| ATOM | 2449 | OE1 | GLU | A | 176 | 38.405 | 51.936 | 46.787 | 1.00 | 0.00 | O |
| ATOM | 2450 | OE2 | GLU | A | 176 | 37.469 | 51.350 | 44.926 | 1.00 | 0.00 | O |
| ATOM | 2451 | C | GLU | A | 176 | 41.240 | 47.789 | 45.469 | 1.00 | 0.00 | C |
| ATOM | 2452 | O | GLU | A | 176 | 41.667 | 47.598 | 44.361 | 1.00 | 0.00 | O |
| ATOM | 2453 | N | ARG | A | 177 | 41.394 | 46.787 | 46.411 | 1.00 | 0.00 | N |
| ATOM | 2454 | H | ARG | A | 177 | 40.869 | 46.866 | 47.259 | 1.00 | 0.00 | H |
| ATOM | 2455 | CA | ARG | A | 177 | 42.242 | 45.652 | 46.267 | 1.00 | 0.00 | C |
| ATOM | 2456 | HA | ARG | A | 177 | 43.074 | 46.128 | 45.984 | 1.00 | 0.00 | H |
| ATOM | 2457 | CB | ARG | A | 177 | 42.509 | 44.823 | 47.541 | 1.00 | 0.00 | C |
| ATOM | 2458 | HB1 | ARG | A | 177 | 41.876 | 44.049 | 47.524 | 1.00 | 0.00 | H |
| ATOM | 2459 | HB2 | ARG | A | 177 | 42.296 | 45.411 | 48.322 | 1.00 | 0.00 | H |
| ATOM | 2460 | CG | ARG | A | 177 | 43.973 | 44.257 | 47.735 | 1.00 | 0.00 | C |
| ATOM | 2461 | HG1 | ARG | A | 177 | 44.255 | 44.436 | 48.678 | 1.00 | 0.00 | H |
| ATOM | 2462 | HG2 | ARG | A | 177 | 44.581 | 44.743 | 47.107 | 1.00 | 0.00 | H |
| ATOM | 2463 | CD | ARG | A | 177 | 44.242 | 42.799 | 47.506 | 1.00 | 0.00 | C |
| ATOM | 2464 | HD1 | ARG | A | 177 | 45.233 | 42.665 | 47.499 | 1.00 | 0.00 | H |
| ATOM | 2465 | HD2 | ARG | A | 177 | 43.865 | 42.551 | 46.613 | 1.00 | 0.00 | H |
| ATOM | 2466 | NE | ARG | A | 177 | 43.701 | 41.845 | 48.464 | 1.00 | 0.00 | N |
| ATOM | 2467 | HE | ARG | A | 177 | 44.116 | 41.863 | 49.373 | 1.00 | 0.00 | H |
| ATOM | 2468 | CZ | ARG | A | 177 | 42.708 | 40.933 | 48.290 | 1.00 | 0.00 | C |
| ATOM | 2469 | NH1 | ARG | A | 177 | 42.150 | 40.698 | 47.066 | 1.00 | 0.00 | N |
| ATOM | 2470 | HH11 | ARG | A | 177 | 42.471 | 41.204 | 46.265 | 1.00 | 0.00 | H |
| ATOM | 2471 | HH12 | ARG | A | 177 | 41.421 | 40.020 | 46.971 | 1.00 | 0.00 | H |
| ATOM | 2472 | NH2 | ARG | A | 177 | 42.194 | 40.216 | 49.250 | 1.00 | 0.00 | N |
| ATOM | 2473 | HH21 | ARG | A | 177 | 42.530 | 40.320 | 50.186 | 1.00 | 0.00 | H |
| ATOM | 2474 | HH22 | ARG | A | 177 | 41.464 | 39.562 | 49.050 | 1.00 | 0.00 | H |
| ATOM | 2475 | C | ARG | A | 177 | 41.710 | 44.709 | 45.228 | 1.00 | 0.00 | C |
| ATOM | 2476 | O | ARG | A | 177 | 40.515 | 44.295 | 45.216 | 1.00 | 0.00 | O |
| ATOM | 2477 | N | LEU | A | 178 | 42.531 | 44.395 | 44.264 | 1.00 | 0.00 | N |
| ATOM | 2478 | H | LEU | A | 178 | 43.468 | 44.743 | 44.300 | 1.00 | 0.00 | H |
| ATOM | 2479 | CA | LEU | A | 178 | 42.138 | 43.554 | 43.132 | 1.00 | 0.00 | C |
| ATOM | 2480 | HA | LEU | A | 178 | 41.344 | 44.000 | 42.719 | 1.00 | 0.00 | H |
| ATOM | 2481 | CB | LEU | A | 178 | 43.227 | 43.435 | 42.038 | 1.00 | 0.00 | C |

| | | | | | | | | | | | |
|---|---|---|---|---|---|---|---|---|---|---|---|
| ATOM | 2482 | HB1 | LEU | A | 178 | 42.937 | 42.762 | 41.357 | 1.00 | 0.00 | H |
| ATOM | 2483 | HB2 | LEU | A | 178 | 44.084 | 43.137 | 42.458 | 1.00 | 0.00 | H |
| ATOM | 2484 | CG | LEU | A | 178 | 43.474 | 44.757 | 41.331 | 1.00 | 0.00 | C |
| ATOM | 2485 | HG | LEU | A | 178 | 43.465 | 45.528 | 41.968 | 1.00 | 0.00 | H |
| ATOM | 2486 | CD1 | LEU | A | 178 | 44.799 | 44.615 | 40.672 | 1.00 | 0.00 | C |
| ATOM | 2487 | HD11 | LEU | A | 178 | 45.021 | 45.460 | 40.185 | 1.00 | 0.00 | H |
| ATOM | 2488 | HD12 | LEU | A | 178 | 45.498 | 44.435 | 41.364 | 1.00 | 0.00 | H |
| ATOM | 2489 | HD13 | LEU | A | 178 | 44.769 | 43.854 | 40.023 | 1.00 | 0.00 | H |
| ATOM | 2490 | CD2 | LEU | A | 178 | 42.409 | 45.046 | 40.311 | 1.00 | 0.00 | C |
| ATOM | 2491 | HD21 | LEU | A | 178 | 42.602 | 45.921 | 39.866 | 1.00 | 0.00 | H |
| ATOM | 2492 | HD22 | LEU | A | 178 | 42.398 | 44.318 | 39.625 | 1.00 | 0.00 | H |
| ATOM | 2493 | HD23 | LEU | A | 178 | 41.518 | 45.091 | 40.763 | 1.00 | 0.00 | H |
| ATOM | 2494 | C | LEU | A | 178 | 41.789 | 42.195 | 43.683 | 1.00 | 0.00 | C |
| ATOM | 2495 | O | LEU | A | 178 | 42.600 | 41.503 | 44.361 | 1.00 | 0.00 | O |
| ATOM | 2496 | N | LEU | A | 179 | 40.573 | 41.726 | 43.433 | 1.00 | 0.00 | N |
| ATOM | 2497 | H | LEU | A | 179 | 39.943 | 42.289 | 42.899 | 1.00 | 0.00 | H |
| ATOM | 2498 | CA | LEU | A | 179 | 40.110 | 40.404 | 43.912 | 1.00 | 0.00 | C |
| ATOM | 2499 | HA | LEU | A | 179 | 40.453 | 40.398 | 44.851 | 1.00 | 0.00 | H |
| ATOM | 2500 | CB | LEU | A | 179 | 38.563 | 40.252 | 43.790 | 1.00 | 0.00 | C |
| ATOM | 2501 | HB1 | LEU | A | 179 | 38.334 | 39.307 | 44.023 | 1.00 | 0.00 | H |
| ATOM | 2502 | HB2 | LEU | A | 179 | 38.318 | 40.429 | 42.837 | 1.00 | 0.00 | H |
| ATOM | 2503 | CG | LEU | A | 179 | 37.685 | 41.185 | 44.685 | 1.00 | 0.00 | C |
| ATOM | 2504 | HG | LEU | A | 179 | 37.974 | 42.105 | 44.419 | 1.00 | 0.00 | H |
| ATOM | 2505 | CD1 | LEU | A | 179 | 36.194 | 41.119 | 44.399 | 1.00 | 0.00 | C |
| ATOM | 2506 | HD11 | LEU | A | 179 | 35.709 | 41.743 | 45.012 | 1.00 | 0.00 | H |
| ATOM | 2507 | HD12 | LEU | A | 179 | 36.024 | 41.385 | 43.450 | 1.00 | 0.00 | H |
| ATOM | 2508 | HD13 | LEU | A | 179 | 35.867 | 40.185 | 44.547 | 1.00 | 0.00 | H |
| ATOM | 2509 | CD2 | LEU | A | 179 | 37.929 | 41.011 | 46.205 | 1.00 | 0.00 | C |
| ATOM | 2510 | HD21 | LEU | A | 179 | 37.336 | 41.637 | 46.712 | 1.00 | 0.00 | H |
| ATOM | 2511 | HD22 | LEU | A | 179 | 37.723 | 40.069 | 46.470 | 1.00 | 0.00 | H |
| ATOM | 2512 | HD23 | LEU | A | 179 | 38.885 | 41.215 | 46.415 | 1.00 | 0.00 | H |
| ATOM | 2513 | C | LEU | A | 179 | 40.686 | 39.243 | 43.123 | 1.00 | 0.00 | C |
| ATOM | 2514 | O | LEU | A | 179 | 40.820 | 39.122 | 41.910 | 1.00 | 0.00 | O |
| ATOM | 2515 | N | LEU | A | 180 | 40.883 | 38.157 | 43.882 | 1.00 | 0.00 | N |
| ATOM | 2516 | H | LEU | A | 180 | 40.659 | 38.203 | 44.855 | 1.00 | 0.00 | H |
| ATOM | 2517 | CA | LEU | A | 180 | 41.416 | 36.902 | 43.338 | 1.00 | 0.00 | C |
| ATOM | 2518 | HA | LEU | A | 180 | 41.475 | 36.988 | 42.343 | 1.00 | 0.00 | H |
| ATOM | 2519 | CB | LEU | A | 180 | 42.793 | 36.635 | 43.929 | 1.00 | 0.00 | C |
| ATOM | 2520 | HB1 | LEU | A | 180 | 42.685 | 36.547 | 44.919 | 1.00 | 0.00 | H |
| ATOM | 2521 | HB2 | LEU | A | 180 | 43.368 | 37.428 | 43.726 | 1.00 | 0.00 | H |
| ATOM | 2522 | CG | LEU | A | 180 | 43.524 | 35.393 | 43.431 | 1.00 | 0.00 | C |
| ATOM | 2523 | HG | LEU | A | 180 | 42.894 | 34.630 | 43.574 | 1.00 | 0.00 | H |
| ATOM | 2524 | CD1 | LEU | A | 180 | 43.933 | 35.424 | 41.956 | 1.00 | 0.00 | C |
| ATOM | 2525 | HD11 | LEU | A | 180 | 44.403 | 34.573 | 41.722 | 1.00 | 0.00 | H |
| ATOM | 2526 | HD12 | LEU | A | 180 | 43.117 | 35.522 | 41.386 | 1.00 | 0.00 | H |
| ATOM | 2527 | HD13 | LEU | A | 180 | 44.546 | 36.198 | 41.796 | 1.00 | 0.00 | H |
| ATOM | 2528 | CD2 | LEU | A | 180 | 44.788 | 35.260 | 44.194 | 1.00 | 0.00 | C |
| ATOM | 2529 | HD21 | LEU | A | 180 | 45.283 | 34.449 | 43.881 | 1.00 | 0.00 | H |
| ATOM | 2530 | HD22 | LEU | A | 180 | 45.353 | 36.072 | 44.048 | 1.00 | 0.00 | H |
| ATOM | 2531 | HD23 | LEU | A | 180 | 44.581 | 35.169 | 45.168 | 1.00 | 0.00 | H |
| ATOM | 2532 | C | LEU | A | 180 | 40.484 | 35.786 | 43.680 | 1.00 | 0.00 | C |
| ATOM | 2533 | O | LEU | A | 180 | 40.176 | 35.377 | 44.837 | 1.00 | 0.00 | O |
| ATOM | 2534 | N | SER | A | 181 | 39.918 | 35.211 | 42.590 | 1.00 | 0.00 | N |
| ATOM | 2535 | H | SER | A | 181 | 40.245 | 35.535 | 41.702 | 1.00 | 0.00 | H |

| | | | | | | | | | | | |
|---|---|---|---|---|---|---|---|---|---|---|---|
| ATOM | 2536 | CA | SER | A | 181 | 38.907 | 34.194 | 42.557 | 1.00 | 0.00 | C |
| ATOM | 2537 | HA | SER | A | 181 | 38.781 | 33.886 | 43.500 | 1.00 | 0.00 | H |
| ATOM | 2538 | CB | SER | A | 181 | 37.675 | 34.845 | 41.829 | 1.00 | 0.00 | C |
| ATOM | 2539 | HB1 | SER | A | 181 | 37.967 | 35.257 | 40.966 | 1.00 | 0.00 | H |
| ATOM | 2540 | HB2 | SER | A | 181 | 37.265 | 35.547 | 42.412 | 1.00 | 0.00 | H |
| ATOM | 2541 | OG | SER | A | 181 | 36.598 | 34.037 | 41.468 | 1.00 | 0.00 | O |
| ATOM | 2542 | HG | SER | A | 181 | 35.900 | 34.594 | 41.018 | 1.00 | 0.00 | H |
| ATOM | 2543 | C | SER | A | 181 | 39.329 | 33.019 | 41.745 | 1.00 | 0.00 | C |
| ATOM | 2544 | O | SER | A | 181 | 40.187 | 33.190 | 40.916 | 1.00 | 0.00 | O |
| ATOM | 2545 | N | ALA | A | 182 | 38.661 | 31.831 | 42.061 | 1.00 | 0.00 | N |
| ATOM | 2546 | H | ALA | A | 182 | 37.946 | 31.818 | 42.760 | 1.00 | 0.00 | H |
| ATOM | 2547 | CA | ALA | A | 182 | 39.029 | 30.610 | 41.349 | 1.00 | 0.00 | C |
| ATOM | 2548 | HA | ALA | A | 182 | 39.296 | 30.896 | 40.429 | 1.00 | 0.00 | H |
| ATOM | 2549 | CB | ALA | A | 182 | 40.276 | 29.922 | 41.971 | 1.00 | 0.00 | C |
| ATOM | 2550 | HB1 | ALA | A | 182 | 40.492 | 29.093 | 41.454 | 1.00 | 0.00 | H |
| ATOM | 2551 | HB2 | ALA | A | 182 | 41.054 | 30.549 | 41.936 | 1.00 | 0.00 | H |
| ATOM | 2552 | HB3 | ALA | A | 182 | 40.084 | 29.681 | 42.922 | 1.00 | 0.00 | H |
| ATOM | 2553 | C | ALA | A | 182 | 37.804 | 29.706 | 41.209 | 1.00 | 0.00 | C |
| ATOM | 2554 | O | ALA | A | 182 | 37.129 | 29.374 | 42.186 | 1.00 | 0.00 | O |
| ATOM | 2555 | N | ALA | A | 183 | 37.511 | 29.344 | 39.969 | 1.00 | 0.00 | N |
| ATOM | 2556 | H | ALA | A | 183 | 38.059 | 29.740 | 39.232 | 1.00 | 0.00 | H |
| ATOM | 2557 | CA | ALA | A | 183 | 36.462 | 28.422 | 39.596 | 1.00 | 0.00 | C |
| ATOM | 2558 | HA | ALA | A | 183 | 35.713 | 28.497 | 40.255 | 1.00 | 0.00 | H |
| ATOM | 2559 | CB | ALA | A | 183 | 35.968 | 28.830 | 38.215 | 1.00 | 0.00 | C |
| ATOM | 2560 | HB1 | ALA | A | 183 | 35.239 | 28.210 | 37.925 | 1.00 | 0.00 | H |
| ATOM | 2561 | HB2 | ALA | A | 183 | 35.616 | 29.765 | 38.250 | 1.00 | 0.00 | H |
| ATOM | 2562 | HB3 | ALA | A | 183 | 36.725 | 28.782 | 37.563 | 1.00 | 0.00 | H |
| ATOM | 2563 | C | ALA | A | 183 | 36.983 | 26.984 | 39.657 | 1.00 | 0.00 | C |
| ATOM | 2564 | O | ALA | A | 183 | 38.171 | 26.733 | 39.454 | 1.00 | 0.00 | O |
| ATOM | 2565 | N | VAL | A | 184 | 36.120 | 26.045 | 40.054 | 1.00 | 0.00 | N |
| ATOM | 2566 | H | VAL | A | 184 | 35.150 | 26.274 | 40.129 | 1.00 | 0.00 | H |
| ATOM | 2567 | CA | VAL | A | 184 | 36.562 | 24.684 | 40.383 | 1.00 | 0.00 | C |
| ATOM | 2568 | HA | VAL | A | 184 | 37.364 | 24.560 | 39.799 | 1.00 | 0.00 | H |
| ATOM | 2569 | CB | VAL | A | 184 | 36.941 | 24.583 | 41.870 | 1.00 | 0.00 | C |
| ATOM | 2570 | HB | VAL | A | 184 | 37.235 | 23.628 | 41.907 | 1.00 | 0.00 | H |
| ATOM | 2571 | CG1 | VAL | A | 184 | 38.085 | 25.520 | 42.340 | 1.00 | 0.00 | C |
| ATOM | 2572 | HG11 | VAL | A | 184 | 38.253 | 25.378 | 43.315 | 1.00 | 0.00 | H |
| ATOM | 2573 | HG12 | VAL | A | 184 | 38.917 | 25.314 | 41.825 | 1.00 | 0.00 | H |
| ATOM | 2574 | HG13 | VAL | A | 184 | 37.822 | 26.472 | 42.183 | 1.00 | 0.00 | H |
| ATOM | 2575 | CG2 | VAL | A | 184 | 35.794 | 24.786 | 42.839 | 1.00 | 0.00 | C |
| ATOM | 2576 | HG21 | VAL | A | 184 | 36.131 | 24.703 | 43.777 | 1.00 | 0.00 | H |
| ATOM | 2577 | HG22 | VAL | A | 184 | 35.401 | 25.696 | 42.705 | 1.00 | 0.00 | H |
| ATOM | 2578 | HG23 | VAL | A | 184 | 35.092 | 24.093 | 42.675 | 1.00 | 0.00 | H |
| ATOM | 2579 | C | VAL | A | 184 | 35.534 | 23.593 | 40.019 | 1.00 | 0.00 | C |
| ATOM | 2580 | O | VAL | A | 184 | 34.340 | 23.821 | 40.203 | 1.00 | 0.00 | O |
| ATOM | 2581 | N | PRO | A | 185 | 35.945 | 22.396 | 39.552 | 1.00 | 0.00 | N |
| ATOM | 2582 | CD | PRO | A | 185 | 37.321 | 21.978 | 39.343 | 1.00 | 0.00 | C |
| ATOM | 2583 | HD1 | PRO | A | 185 | 37.847 | 22.145 | 40.177 | 1.00 | 0.00 | H |
| ATOM | 2584 | HD2 | PRO | A | 185 | 37.715 | 22.504 | 38.589 | 1.00 | 0.00 | H |
| ATOM | 2585 | CG | PRO | A | 185 | 37.272 | 20.487 | 39.010 | 1.00 | 0.00 | C |
| ATOM | 2586 | HG1 | PRO | A | 185 | 37.340 | 19.930 | 39.838 | 1.00 | 0.00 | H |
| ATOM | 2587 | HG2 | PRO | A | 185 | 38.004 | 20.235 | 38.377 | 1.00 | 0.00 | H |
| ATOM | 2588 | CB | PRO | A | 185 | 35.900 | 20.334 | 38.354 | 1.00 | 0.00 | C |
| ATOM | 2589 | HB1 | PRO | A | 185 | 35.559 | 19.399 | 38.449 | 1.00 | 0.00 | H |

```
ATOM   2590  HB2 PRO A 185      35.936  20.579  37.385  1.00  0.00           H
ATOM   2591  CA  PRO A 185      35.033  21.322  39.142  1.00  0.00           C
ATOM   2592  HA  PRO A 185      34.302  21.725  38.591  1.00  0.00           H
ATOM   2593  C   PRO A 185      34.328  20.644  40.329  1.00  0.00           C
ATOM   2594  O   PRO A 185      34.827  20.634  41.456  1.00  0.00           O
ATOM   2595  N   ALA A 186      33.170  20.031  40.064  1.00  0.00           N
ATOM   2596  H   ALA A 186      32.885  19.944  39.109  1.00  0.00           H
ATOM   2597  CA  ALA A 186      32.297  19.480  41.105  1.00  0.00           C
ATOM   2598  HA  ALA A 186      32.690  19.798  41.968  1.00  0.00           H
ATOM   2599  CB  ALA A 186      30.893  20.056  40.878  1.00  0.00           C
ATOM   2600  HB1 ALA A 186      30.270  19.699  41.574  1.00  0.00           H
ATOM   2601  HB2 ALA A 186      30.928  21.053  40.942  1.00  0.00           H
ATOM   2602  HB3 ALA A 186      30.567  19.791  39.971  1.00  0.00           H
ATOM   2603  C   ALA A 186      32.320  17.942  41.258  1.00  0.00           C
ATOM   2604  O   ALA A 186      32.001  17.452  42.340  1.00  0.00           O
ATOM   2605  N   GLY A 187      32.712  17.166  40.239  1.00  0.00           N
ATOM   2606  H   GLY A 187      32.961  17.604  39.375  1.00  0.00           H
ATOM   2607  CA  GLY A 187      32.794  15.700  40.331  1.00  0.00           C
ATOM   2608  HA1 GLY A 187      32.856  15.344  39.399  1.00  0.00           H
ATOM   2609  HA2 GLY A 187      31.948  15.378  40.755  1.00  0.00           H
ATOM   2610  C   GLY A 187      33.996  15.203  41.149  1.00  0.00           C
ATOM   2611  O   GLY A 187      35.124  15.643  40.930  1.00  0.00           O
ATOM   2612  N   GLN A 188      33.790  14.205  42.020  1.00  0.00           N
ATOM   2613  H   GLN A 188      32.888  13.774  42.035  1.00  0.00           H
ATOM   2614  CA  GLN A 188      34.816  13.702  42.960  1.00  0.00           C
ATOM   2615  HA  GLN A 188      35.144  14.513  43.444  1.00  0.00           H
ATOM   2616  CB  GLN A 188      34.151  12.701  43.929  1.00  0.00           C
ATOM   2617  HB1 GLN A 188      33.806  11.925  43.401  1.00  0.00           H
ATOM   2618  HB2 GLN A 188      33.389  13.157  44.388  1.00  0.00           H
ATOM   2619  CG  GLN A 188      35.075  12.159  44.997  1.00  0.00           C
ATOM   2620  HG1 GLN A 188      34.548  12.059  45.841  1.00  0.00           H
ATOM   2621  HG2 GLN A 188      35.808  12.824  45.139  1.00  0.00           H
ATOM   2622  CD  GLN A 188      35.699  10.796  44.632  1.00  0.00           C
ATOM   2623  OE1 GLN A 188      35.801  10.429  43.510  1.00  0.00           O
ATOM   2624  NE2 GLN A 188      36.208  10.036  45.537  1.00  0.00           N
ATOM   2625 HE21 GLN A 188      36.203  10.326  46.494  1.00  0.00           H
ATOM   2626 HE22 GLN A 188      36.608   9.156  45.283  1.00  0.00           H
ATOM   2627  C   GLN A 188      36.067  13.111  42.281  1.00  0.00           C
ATOM   2628  O   GLN A 188      37.171  13.253  42.806  1.00  0.00           O
ATOM   2629  N   THR A 189      35.929  12.592  41.057  1.00  0.00           N
ATOM   2630  H   THR A 189      35.014  12.575  40.654  1.00  0.00           H
ATOM   2631  CA  THR A 189      37.045  12.044  40.267  1.00  0.00           C
ATOM   2632  HA  THR A 189      37.501  11.422  40.904  1.00  0.00           H
ATOM   2633  CB  THR A 189      36.503  11.295  39.040  1.00  0.00           C
ATOM   2634  HB  THR A 189      35.853  10.610  39.369  1.00  0.00           H
ATOM   2635  CG2 THR A 189      35.841  12.206  38.000  1.00  0.00           C
ATOM   2636 HG21 THR A 189      35.513  11.654  37.233  1.00  0.00           H
ATOM   2637 HG22 THR A 189      35.070  12.685  38.419  1.00  0.00           H
ATOM   2638 HG23 THR A 189      36.508  12.874  37.669  1.00  0.00           H
ATOM   2639  OG1 THR A 189      37.535  10.596  38.392  1.00  0.00           O
ATOM   2640  HG1 THR A 189      37.164  10.115  37.597  1.00  0.00           H
ATOM   2641  C   THR A 189      38.083  13.091  39.838  1.00  0.00           C
ATOM   2642  O   THR A 189      39.178  12.717  39.427  1.00  0.00           O
ATOM   2643  N   TYR A 190      37.764  14.388  39.919  1.00  0.00           N
```

```
ATOM   2644  H    TYR A 190      36.832  14.616  40.200  1.00  0.00           H
ATOM   2645  CA   TYR A 190      38.675  15.502  39.628  1.00  0.00           C
ATOM   2646  HA   TYR A 190      39.416  15.102  39.088  1.00  0.00           H
ATOM   2647  CB   TYR A 190      37.913  16.585  38.842  1.00  0.00           C
ATOM   2648  HB1  TYR A 190      38.541  17.342  38.660  1.00  0.00           H
ATOM   2649  HB2  TYR A 190      37.155  16.910  39.408  1.00  0.00           H
ATOM   2650  CG   TYR A 190      37.334  16.142  37.514  1.00  0.00           C
ATOM   2651  CD1  TYR A 190      38.199  15.875  36.438  1.00  0.00           C
ATOM   2652  HD1  TYR A 190      39.188  15.963  36.559  1.00  0.00           H
ATOM   2653  CE1  TYR A 190      37.676  15.484  35.193  1.00  0.00           C
ATOM   2654  HE1  TYR A 190      38.295  15.293  34.431  1.00  0.00           H
ATOM   2655  CZ   TYR A 190      36.283  15.359  35.012  1.00  0.00           C
ATOM   2656  OH   TYR A 190      35.820  14.976  33.792  1.00  0.00           O
ATOM   2657  HH   TYR A 190      34.821  14.931  33.814  1.00  0.00           H
ATOM   2658  CE2  TYR A 190      35.413  15.633  36.092  1.00  0.00           C
ATOM   2659  HE2  TYR A 190      34.424  15.547  35.972  1.00  0.00           H
ATOM   2660  CD2  TYR A 190      35.940  16.027  37.339  1.00  0.00           C
ATOM   2661  HD2  TYR A 190      35.323  16.225  38.101  1.00  0.00           H
ATOM   2662  C    TYR A 190      39.274  16.112  40.905  1.00  0.00           C
ATOM   2663  O    TYR A 190      39.977  17.111  40.883  1.00  0.00           O
ATOM   2664  N    VAL A 191      38.895  15.662  42.113  1.00  0.00           N
ATOM   2665  H    VAL A 191      38.539  14.731  42.189  1.00  0.00           H
ATOM   2666  CA   VAL A 191      38.982  16.472  43.306  1.00  0.00           C
ATOM   2667  HA   VAL A 191      39.657  17.163  43.049  1.00  0.00           H
ATOM   2668  CB   VAL A 191      37.619  17.222  43.628  1.00  0.00           C
ATOM   2669  HB   VAL A 191      36.875  16.554  43.603  1.00  0.00           H
ATOM   2670  CG1  VAL A 191      37.708  17.894  45.041  1.00  0.00           C
ATOM   2671 HG11 VAL A 191      36.849  18.365  45.241  1.00  0.00           H
ATOM   2672 HG12 VAL A 191      37.872  17.192  45.734  1.00  0.00           H
ATOM   2673 HG13 VAL A 191      38.460  18.553  45.050  1.00  0.00           H
ATOM   2674  CG2  VAL A 191      37.192  18.221  42.571  1.00  0.00           C
ATOM   2675 HG21 VAL A 191      36.331  18.648  42.845  1.00  0.00           H
ATOM   2676 HG22 VAL A 191      37.896  18.924  42.473  1.00  0.00           H
ATOM   2677 HG23 VAL A 191      37.067  17.750  41.698  1.00  0.00           H
ATOM   2678  C    VAL A 191      39.492  15.570  44.535  1.00  0.00           C
ATOM   2679  O    VAL A 191      40.008  16.085  45.471  1.00  0.00           O
ATOM   2680  N    ASP A 192      39.450  14.223  44.481  1.00  0.00           N
ATOM   2681  H    ASP A 192      39.023  13.773  43.697  1.00  0.00           H
ATOM   2682  CA   ASP A 192      40.018  13.417  45.550  1.00  0.00           C
ATOM   2683  HA   ASP A 192      39.542  13.825  46.329  1.00  0.00           H
ATOM   2684  CB   ASP A 192      39.679  11.901  45.403  1.00  0.00           C
ATOM   2685  HB1  ASP A 192      38.683  11.811  45.387  1.00  0.00           H
ATOM   2686  HB2  ASP A 192      40.043  11.428  46.205  1.00  0.00           H
ATOM   2687  CG   ASP A 192      40.238  11.185  44.139  1.00  0.00           C
ATOM   2688  OD1  ASP A 192      39.982  10.018  43.948  1.00  0.00           O
ATOM   2689  OD2  ASP A 192      41.242  11.643  43.592  1.00  0.00           O
ATOM   2690  C    ASP A 192      41.561  13.582  45.726  1.00  0.00           C
ATOM   2691  O    ASP A 192      42.131  13.112  46.723  1.00  0.00           O
ATOM   2692  N    ALA A 193      42.302  14.190  44.820  1.00  0.00           N
ATOM   2693  H    ALA A 193      41.824  14.500  43.998  1.00  0.00           H
ATOM   2694  CA   ALA A 193      43.745  14.468  44.870  1.00  0.00           C
ATOM   2695  HA   ALA A 193      44.163  14.174  45.730  1.00  0.00           H
ATOM   2696  CB   ALA A 193      44.461  13.614  43.755  1.00  0.00           C
ATOM   2697  HB1  ALA A 193      45.444  13.794  43.778  1.00  0.00           H
```

| | | | | | | | | | | |
|---|---|---|---|---|---|---|---|---|---|---|
| ATOM | 2698 | HB2 | ALA | A | 193 | 44.296 | 12.642 | 43.922 | 1.00 | 0.00 | H |
| ATOM | 2699 | HB3 | ALA | A | 193 | 44.097 | 13.865 | 42.858 | 1.00 | 0.00 | H |
| ATOM | 2700 | C | ALA | A | 193 | 43.832 | 16.050 | 44.717 | 1.00 | 0.00 | C |
| ATOM | 2701 | O | ALA | A | 193 | 42.758 | 16.605 | 44.502 | 1.00 | 0.00 | O |
| ATOM | 2702 | N | GLY | A | 194 | 45.051 | 16.653 | 44.830 | 1.00 | 0.00 | N |
| ATOM | 2703 | H | GLY | A | 194 | 45.789 | 16.176 | 45.308 | 1.00 | 0.00 | H |
| ATOM | 2704 | CA | GLY | A | 194 | 45.299 | 17.954 | 44.280 | 1.00 | 0.00 | C |
| ATOM | 2705 | HA1 | GLY | A | 194 | 44.933 | 17.929 | 43.350 | 1.00 | 0.00 | H |
| ATOM | 2706 | HA2 | GLY | A | 194 | 46.294 | 18.048 | 44.245 | 1.00 | 0.00 | H |
| ATOM | 2707 | C | GLY | A | 194 | 44.732 | 19.166 | 44.981 | 1.00 | 0.00 | C |
| ATOM | 2708 | O | GLY | A | 194 | 44.232 | 18.949 | 46.090 | 1.00 | 0.00 | O |
| ATOM | 2709 | N | TYR | A | 195 | 44.826 | 20.368 | 44.431 | 1.00 | 0.00 | N |
| ATOM | 2710 | H | TYR | A | 195 | 45.267 | 20.421 | 43.535 | 1.00 | 0.00 | H |
| ATOM | 2711 | CA | TYR | A | 195 | 44.340 | 21.647 | 45.016 | 1.00 | 0.00 | C |
| ATOM | 2712 | HA | TYR | A | 195 | 44.961 | 22.253 | 44.518 | 1.00 | 0.00 | H |
| ATOM | 2713 | CB | TYR | A | 195 | 42.865 | 21.984 | 44.720 | 1.00 | 0.00 | C |
| ATOM | 2714 | HB1 | TYR | A | 195 | 42.680 | 22.862 | 45.162 | 1.00 | 0.00 | H |
| ATOM | 2715 | HB2 | TYR | A | 195 | 42.317 | 21.269 | 45.155 | 1.00 | 0.00 | H |
| ATOM | 2716 | CG | TYR | A | 195 | 42.330 | 22.112 | 43.299 | 1.00 | 0.00 | C |
| ATOM | 2717 | CD1 | TYR | A | 195 | 41.985 | 20.949 | 42.580 | 1.00 | 0.00 | C |
| ATOM | 2718 | HD1 | TYR | A | 195 | 42.258 | 20.061 | 42.950 | 1.00 | 0.00 | H |
| ATOM | 2719 | CE1 | TYR | A | 195 | 41.282 | 20.980 | 41.380 | 1.00 | 0.00 | C |
| ATOM | 2720 | HE1 | TYR | A | 195 | 41.011 | 20.132 | 40.925 | 1.00 | 0.00 | H |
| ATOM | 2721 | CZ | TYR | A | 195 | 40.962 | 22.224 | 40.823 | 1.00 | 0.00 | C |
| ATOM | 2722 | OH | TYR | A | 195 | 40.396 | 22.260 | 39.589 | 1.00 | 0.00 | O |
| ATOM | 2723 | HH | TYR | A | 195 | 40.227 | 23.210 | 39.328 | 1.00 | 0.00 | H |
| ATOM | 2724 | CE2 | TYR | A | 195 | 41.254 | 23.380 | 41.511 | 1.00 | 0.00 | C |
| ATOM | 2725 | HE2 | TYR | A | 195 | 40.964 | 24.258 | 41.131 | 1.00 | 0.00 | H |
| ATOM | 2726 | CD2 | TYR | A | 195 | 41.922 | 23.360 | 42.686 | 1.00 | 0.00 | C |
| ATOM | 2727 | HD2 | TYR | A | 195 | 42.139 | 24.224 | 43.141 | 1.00 | 0.00 | H |
| ATOM | 2728 | C | TYR | A | 195 | 44.538 | 21.817 | 46.532 | 1.00 | 0.00 | C |
| ATOM | 2729 | O | TYR | A | 195 | 43.783 | 21.412 | 47.381 | 1.00 | 0.00 | O |
| ATOM | 2730 | N | GLU | A | 196 | 45.706 | 22.336 | 46.991 | 1.00 | 0.00 | N |
| ATOM | 2731 | H | GLU | A | 196 | 46.365 | 22.618 | 46.294 | 1.00 | 0.00 | H |
| ATOM | 2732 | CA | GLU | A | 196 | 46.112 | 22.530 | 48.367 | 1.00 | 0.00 | C |
| ATOM | 2733 | HA | GLU | A | 196 | 45.932 | 21.634 | 48.774 | 1.00 | 0.00 | H |
| ATOM | 2734 | CB | GLU | A | 196 | 47.619 | 22.688 | 48.552 | 1.00 | 0.00 | C |
| ATOM | 2735 | HB1 | GLU | A | 196 | 47.866 | 23.605 | 48.238 | 1.00 | 0.00 | H |
| ATOM | 2736 | HB2 | GLU | A | 196 | 48.066 | 22.006 | 47.974 | 1.00 | 0.00 | H |
| ATOM | 2737 | CG | GLU | A | 196 | 48.157 | 22.519 | 49.984 | 1.00 | 0.00 | C |
| ATOM | 2738 | HG1 | GLU | A | 196 | 47.506 | 22.904 | 50.638 | 1.00 | 0.00 | H |
| ATOM | 2739 | HG2 | GLU | A | 196 | 49.036 | 22.988 | 50.073 | 1.00 | 0.00 | H |
| ATOM | 2740 | CD | GLU | A | 196 | 48.350 | 21.058 | 50.290 | 1.00 | 0.00 | C |
| ATOM | 2741 | OE1 | GLU | A | 196 | 47.802 | 20.163 | 49.583 | 1.00 | 0.00 | O |
| ATOM | 2742 | OE2 | GLU | A | 196 | 48.832 | 20.705 | 51.376 | 1.00 | 0.00 | O |
| ATOM | 2743 | C | GLU | A | 196 | 45.198 | 23.498 | 49.135 | 1.00 | 0.00 | C |
| ATOM | 2744 | O | GLU | A | 196 | 45.665 | 24.571 | 49.584 | 1.00 | 0.00 | O |
| ATOM | 2745 | N | VAL | A | 197 | 43.929 | 23.199 | 49.363 | 1.00 | 0.00 | N |
| ATOM | 2746 | H | VAL | A | 197 | 43.605 | 22.336 | 48.976 | 1.00 | 0.00 | H |
| ATOM | 2747 | CA | VAL | A | 197 | 42.932 | 23.986 | 50.121 | 1.00 | 0.00 | C |
| ATOM | 2748 | HA | VAL | A | 197 | 42.899 | 24.779 | 49.513 | 1.00 | 0.00 | H |
| ATOM | 2749 | CB | VAL | A | 197 | 41.532 | 23.403 | 50.131 | 1.00 | 0.00 | C |
| ATOM | 2750 | HB | VAL | A | 197 | 41.735 | 22.529 | 50.572 | 1.00 | 0.00 | H |
| ATOM | 2751 | CG1 | VAL | A | 197 | 40.498 | 24.283 | 50.938 | 1.00 | 0.00 | C |

```
ATOM   2752 HG11 VAL A 197      39.597  23.850  50.909  1.00  0.00           H
ATOM   2753 HG12 VAL A 197      40.799  24.363  51.888  1.00  0.00           H
ATOM   2754 HG13 VAL A 197      40.443  25.193  50.528  1.00  0.00           H
ATOM   2755  CG2 VAL A 197      40.896  23.100  48.748  1.00  0.00           C
ATOM   2756 HG21 VAL A 197      39.980  22.722  48.878  1.00  0.00           H
ATOM   2757 HG22 VAL A 197      40.835  23.945  48.217  1.00  0.00           H
ATOM   2758 HG23 VAL A 197      41.463  22.438  48.258  1.00  0.00           H
ATOM   2759  C   VAL A 197      43.368  24.569  51.473  1.00  0.00           C
ATOM   2760  O   VAL A 197      43.155  25.802  51.666  1.00  0.00           O
ATOM   2761  N   ASP A 198      44.087  23.814  52.233  1.00  0.00           N
ATOM   2762  H   ASP A 198      44.174  22.843  52.012  1.00  0.00           H
ATOM   2763  CA  ASP A 198      44.782  24.344  53.417  1.00  0.00           C
ATOM   2764  HA  ASP A 198      43.994  24.610  53.972  1.00  0.00           H
ATOM   2765  CB  ASP A 198      45.743  23.236  54.070  1.00  0.00           C
ATOM   2766  HB1 ASP A 198      46.140  23.577  54.922  1.00  0.00           H
ATOM   2767  HB2 ASP A 198      46.476  22.993  53.435  1.00  0.00           H
ATOM   2768  CG  ASP A 198      44.941  21.975  54.384  1.00  0.00           C
ATOM   2769  OD1 ASP A 198      45.599  20.971  54.622  1.00  0.00           O
ATOM   2770  OD2 ASP A 198      43.707  21.987  54.452  1.00  0.00           O
ATOM   2771  C   ASP A 198      45.526  25.641  53.237  1.00  0.00           C
ATOM   2772  O   ASP A 198      45.635  26.408  54.201  1.00  0.00           O
ATOM   2773  N   LYS A 199      46.224  25.843  52.052  1.00  0.00           N
ATOM   2774  H   LYS A 199      46.145  25.145  51.340  1.00  0.00           H
ATOM   2775  CA  LYS A 199      47.060  26.997  51.777  1.00  0.00           C
ATOM   2776  HA  LYS A 199      47.222  27.349  52.699  1.00  0.00           H
ATOM   2777  CB  LYS A 199      48.375  26.582  51.061  1.00  0.00           C
ATOM   2778  HB1 LYS A 199      48.886  27.386  50.757  1.00  0.00           H
ATOM   2779  HB2 LYS A 199      48.184  25.992  50.276  1.00  0.00           H
ATOM   2780  CG  LYS A 199      49.162  25.813  52.142  1.00  0.00           C
ATOM   2781  HG1 LYS A 199      48.671  24.971  52.364  1.00  0.00           H
ATOM   2782  HG2 LYS A 199      49.233  26.383  52.960  1.00  0.00           H
ATOM   2783  CD  LYS A 199      50.561  25.440  51.707  1.00  0.00           C
ATOM   2784  HD1 LYS A 199      50.497  24.977  50.823  1.00  0.00           H
ATOM   2785  HD2 LYS A 199      50.938  24.811  52.386  1.00  0.00           H
ATOM   2786  CE  LYS A 199      51.504  26.658  51.570  1.00  0.00           C
ATOM   2787  HE1 LYS A 199      51.231  27.198  50.774  1.00  0.00           H
ATOM   2788  HE2 LYS A 199      52.443  26.336  51.448  1.00  0.00           H
ATOM   2789  NZ  LYS A 199      51.508  27.569  52.727  1.00  0.00           N
ATOM   2790  HZ1 LYS A 199      52.141  28.324  52.555  1.00  0.00           H
ATOM   2791  HZ2 LYS A 199      50.587  27.932  52.869  1.00  0.00           H
ATOM   2792  HZ3 LYS A 199      51.799  27.070  53.543  1.00  0.00           H
ATOM   2793  C   LYS A 199      46.356  28.078  50.938  1.00  0.00           C
ATOM   2794  O   LYS A 199      46.216  29.207  51.351  1.00  0.00           O
ATOM   2795  N   ILE A 200      45.728  27.637  49.815  1.00  0.00           N
ATOM   2796  H   ILE A 200      45.688  26.651  49.652  1.00  0.00           H
ATOM   2797  CA  ILE A 200      45.112  28.521  48.840  1.00  0.00           C
ATOM   2798  HA  ILE A 200      45.755  29.252  48.611  1.00  0.00           H
ATOM   2799  CB  ILE A 200      44.858  27.713  47.564  1.00  0.00           C
ATOM   2800  HB  ILE A 200      44.499  28.405  46.938  1.00  0.00           H
ATOM   2801  CG2 ILE A 200      46.198  27.039  47.049  1.00  0.00           C
ATOM   2802 HG21 ILE A 200      46.010  26.517  46.217  1.00  0.00           H
ATOM   2803 HG22 ILE A 200      46.874  27.749  46.852  1.00  0.00           H
ATOM   2804 HG23 ILE A 200      46.555  26.426  47.754  1.00  0.00           H
ATOM   2805  CG1 ILE A 200      43.743  26.655  47.638  1.00  0.00           C
```

| | | | | | | | | | | | |
|---|---|---|---|---|---|---|---|---|---|---|---|
| ATOM | 2806 | HG11 | ILE | A | 200 | 42.893 | 27.134 | 47.857 | 1.00 | 0.00 | H |
| ATOM | 2807 | HG12 | ILE | A | 200 | 43.975 | 26.030 | 48.383 | 1.00 | 0.00 | H |
| ATOM | 2808 | CD | ILE | A | 200 | 43.482 | 25.800 | 46.380 | 1.00 | 0.00 | C |
| ATOM | 2809 | HD1 | ILE | A | 200 | 42.738 | 25.157 | 46.562 | 1.00 | 0.00 | H |
| ATOM | 2810 | HD2 | ILE | A | 200 | 43.228 | 26.396 | 45.618 | 1.00 | 0.00 | H |
| ATOM | 2811 | HD3 | ILE | A | 200 | 44.311 | 25.292 | 46.144 | 1.00 | 0.00 | H |
| ATOM | 2812 | C | ILE | A | 200 | 43.914 | 29.262 | 49.439 | 1.00 | 0.00 | C |
| ATOM | 2813 | O | ILE | A | 200 | 43.724 | 30.413 | 49.123 | 1.00 | 0.00 | O |
| ATOM | 2814 | N | ALA | A | 201 | 43.161 | 28.729 | 50.337 | 1.00 | 0.00 | N |
| ATOM | 2815 | H | ALA | A | 201 | 43.372 | 27.803 | 50.651 | 1.00 | 0.00 | H |
| ATOM | 2816 | CA | ALA | A | 201 | 41.979 | 29.440 | 50.932 | 1.00 | 0.00 | C |
| ATOM | 2817 | HA | ALA | A | 201 | 41.524 | 29.830 | 50.131 | 1.00 | 0.00 | H |
| ATOM | 2818 | CB | ALA | A | 201 | 41.032 | 28.465 | 51.639 | 1.00 | 0.00 | C |
| ATOM | 2819 | HB1 | ALA | A | 201 | 40.259 | 28.970 | 52.022 | 1.00 | 0.00 | H |
| ATOM | 2820 | HB2 | ALA | A | 201 | 40.696 | 27.790 | 50.982 | 1.00 | 0.00 | H |
| ATOM | 2821 | HB3 | ALA | A | 201 | 41.523 | 27.998 | 52.375 | 1.00 | 0.00 | H |
| ATOM | 2822 | C | ALA | A | 201 | 42.303 | 30.647 | 51.804 | 1.00 | 0.00 | C |
| ATOM | 2823 | O | ALA | A | 201 | 41.394 | 31.479 | 51.980 | 1.00 | 0.00 | O |
| ATOM | 2824 | N | GLN | A | 202 | 43.580 | 30.880 | 52.203 | 1.00 | 0.00 | N |
| ATOM | 2825 | H | GLN | A | 202 | 44.214 | 30.115 | 52.095 | 1.00 | 0.00 | H |
| ATOM | 2826 | CA | GLN | A | 202 | 44.164 | 32.130 | 52.783 | 1.00 | 0.00 | C |
| ATOM | 2827 | HA | GLN | A | 202 | 43.557 | 32.467 | 53.502 | 1.00 | 0.00 | H |
| ATOM | 2828 | CB | GLN | A | 202 | 45.625 | 31.677 | 53.163 | 1.00 | 0.00 | C |
| ATOM | 2829 | HB1 | GLN | A | 202 | 46.157 | 31.605 | 52.319 | 1.00 | 0.00 | H |
| ATOM | 2830 | HB2 | GLN | A | 202 | 45.568 | 30.779 | 53.599 | 1.00 | 0.00 | H |
| ATOM | 2831 | CG | GLN | A | 202 | 46.422 | 32.580 | 54.112 | 1.00 | 0.00 | C |
| ATOM | 2832 | HG1 | GLN | A | 202 | 45.900 | 32.697 | 54.957 | 1.00 | 0.00 | H |
| ATOM | 2833 | HG2 | GLN | A | 202 | 46.552 | 33.470 | 53.674 | 1.00 | 0.00 | H |
| ATOM | 2834 | CD | GLN | A | 202 | 47.800 | 31.989 | 54.457 | 1.00 | 0.00 | C |
| ATOM | 2835 | OE1 | GLN | A | 202 | 48.002 | 30.790 | 54.888 | 1.00 | 0.00 | O |
| ATOM | 2836 | NE2 | GLN | A | 202 | 48.769 | 32.545 | 54.095 | 1.00 | 0.00 | N |
| ATOM | 2837 | HE21 | GLN | A | 202 | 48.692 | 33.392 | 53.569 | 1.00 | 0.00 | H |
| ATOM | 2838 | HE22 | GLN | A | 202 | 49.671 | 32.171 | 54.313 | 1.00 | 0.00 | H |
| ATOM | 2839 | C | GLN | A | 202 | 44.299 | 33.271 | 51.785 | 1.00 | 0.00 | C |
| ATOM | 2840 | O | GLN | A | 202 | 44.339 | 34.406 | 52.246 | 1.00 | 0.00 | O |
| ATOM | 2841 | N | ASN | A | 203 | 44.512 | 32.965 | 50.518 | 1.00 | 0.00 | N |
| ATOM | 2842 | H | ASN | A | 203 | 44.403 | 32.010 | 50.244 | 1.00 | 0.00 | H |
| ATOM | 2843 | CA | ASN | A | 203 | 44.882 | 33.892 | 49.530 | 1.00 | 0.00 | C |
| ATOM | 2844 | HA | ASN | A | 203 | 45.012 | 34.728 | 50.063 | 1.00 | 0.00 | H |
| ATOM | 2845 | CB | ASN | A | 203 | 46.115 | 33.439 | 48.854 | 1.00 | 0.00 | C |
| ATOM | 2846 | HB1 | ASN | A | 203 | 46.155 | 33.908 | 47.972 | 1.00 | 0.00 | H |
| ATOM | 2847 | HB2 | ASN | A | 203 | 46.023 | 32.455 | 48.705 | 1.00 | 0.00 | H |
| ATOM | 2848 | CG | ASN | A | 203 | 47.418 | 33.670 | 49.547 | 1.00 | 0.00 | C |
| ATOM | 2849 | OD1 | ASN | A | 203 | 47.643 | 34.673 | 50.222 | 1.00 | 0.00 | O |
| ATOM | 2850 | ND2 | ASN | A | 203 | 48.333 | 32.749 | 49.384 | 1.00 | 0.00 | N |
| ATOM | 2851 | HD21 | ASN | A | 203 | 48.139 | 31.946 | 48.820 | 1.00 | 0.00 | H |
| ATOM | 2852 | HD22 | ASN | A | 203 | 49.226 | 32.848 | 49.823 | 1.00 | 0.00 | H |
| ATOM | 2853 | C | ASN | A | 203 | 43.823 | 34.007 | 48.411 | 1.00 | 0.00 | C |
| ATOM | 2854 | O | ASN | A | 203 | 43.638 | 35.081 | 47.815 | 1.00 | 0.00 | O |
| ATOM | 2855 | N | LEU | A | 204 | 43.035 | 32.849 | 48.181 | 1.00 | 0.00 | N |
| ATOM | 2856 | H | LEU | A | 204 | 43.306 | 31.958 | 48.545 | 1.00 | 0.00 | H |
| ATOM | 2857 | CA | LEU | A | 204 | 41.809 | 33.005 | 47.390 | 1.00 | 0.00 | C |
| ATOM | 2858 | HA | LEU | A | 204 | 42.023 | 33.585 | 46.604 | 1.00 | 0.00 | H |
| ATOM | 2859 | CB | LEU | A | 204 | 41.380 | 31.617 | 46.950 | 1.00 | 0.00 | C |

| | | | | | | | | | | | |
|---|---|---|---|---|---|---|---|---|---|---|---|
| ATOM | 2860 | HB1 | LEU | A | 204 | 40.493 | 31.709 | 46.498 | 1.00 | 0.00 | H |
| ATOM | 2861 | HB2 | LEU | A | 204 | 41.278 | 31.061 | 47.775 | 1.00 | 0.00 | H |
| ATOM | 2862 | CG | LEU | A | 204 | 42.355 | 30.888 | 45.981 | 1.00 | 0.00 | C |
| ATOM | 2863 | HG | LEU | A | 204 | 43.250 | 30.922 | 46.425 | 1.00 | 0.00 | H |
| ATOM | 2864 | CD1 | LEU | A | 204 | 41.883 | 29.439 | 45.768 | 1.00 | 0.00 | C |
| ATOM | 2865 | HD11 | LEU | A | 204 | 42.511 | 28.972 | 45.145 | 1.00 | 0.00 | H |
| ATOM | 2866 | HD12 | LEU | A | 204 | 41.868 | 28.961 | 46.646 | 1.00 | 0.00 | H |
| ATOM | 2867 | HD13 | LEU | A | 204 | 40.964 | 29.442 | 45.374 | 1.00 | 0.00 | H |
| ATOM | 2868 | CD2 | LEU | A | 204 | 42.509 | 31.559 | 44.583 | 1.00 | 0.00 | C |
| ATOM | 2869 | HD21 | LEU | A | 204 | 43.150 | 31.029 | 44.028 | 1.00 | 0.00 | H |
| ATOM | 2870 | HD22 | LEU | A | 204 | 41.619 | 31.589 | 44.128 | 1.00 | 0.00 | H |
| ATOM | 2871 | HD23 | LEU | A | 204 | 42.857 | 32.490 | 44.696 | 1.00 | 0.00 | H |
| ATOM | 2872 | C | LEU | A | 204 | 40.701 | 33.708 | 48.179 | 1.00 | 0.00 | C |
| ATOM | 2873 | O | LEU | A | 204 | 40.411 | 33.394 | 49.321 | 1.00 | 0.00 | O |
| ATOM | 2874 | N | ASP | A | 205 | 40.034 | 34.658 | 47.556 | 1.00 | 0.00 | N |
| ATOM | 2875 | H | ASP | A | 205 | 40.173 | 34.786 | 46.574 | 1.00 | 0.00 | H |
| ATOM | 2876 | CA | ASP | A | 205 | 39.096 | 35.531 | 48.262 | 1.00 | 0.00 | C |
| ATOM | 2877 | HA | ASP | A | 205 | 39.430 | 35.703 | 49.189 | 1.00 | 0.00 | H |
| ATOM | 2878 | CB | ASP | A | 205 | 38.990 | 36.828 | 47.504 | 1.00 | 0.00 | C |
| ATOM | 2879 | HB1 | ASP | A | 205 | 38.078 | 37.211 | 47.651 | 1.00 | 0.00 | H |
| ATOM | 2880 | HB2 | ASP | A | 205 | 39.120 | 36.641 | 46.530 | 1.00 | 0.00 | H |
| ATOM | 2881 | CG | ASP | A | 205 | 39.989 | 37.849 | 47.913 | 1.00 | 0.00 | C |
| ATOM | 2882 | OD1 | ASP | A | 205 | 40.302 | 38.023 | 49.094 | 1.00 | 0.00 | O |
| ATOM | 2883 | OD2 | ASP | A | 205 | 40.610 | 38.425 | 46.990 | 1.00 | 0.00 | O |
| ATOM | 2884 | C | ASP | A | 205 | 37.712 | 34.822 | 48.192 | 1.00 | 0.00 | C |
| ATOM | 2885 | O | ASP | A | 205 | 37.068 | 34.808 | 49.234 | 1.00 | 0.00 | O |
| ATOM | 2886 | N | PHE | A | 206 | 37.388 | 34.140 | 47.082 | 1.00 | 0.00 | N |
| ATOM | 2887 | H | PHE | A | 206 | 37.977 | 34.276 | 46.286 | 1.00 | 0.00 | H |
| ATOM | 2888 | CA | PHE | A | 206 | 36.234 | 33.194 | 46.909 | 1.00 | 0.00 | C |
| ATOM | 2889 | HA | PHE | A | 206 | 36.157 | 32.802 | 47.826 | 1.00 | 0.00 | H |
| ATOM | 2890 | CB | PHE | A | 206 | 34.894 | 33.870 | 46.585 | 1.00 | 0.00 | C |
| ATOM | 2891 | HB1 | PHE | A | 206 | 34.556 | 34.262 | 47.441 | 1.00 | 0.00 | H |
| ATOM | 2892 | HB2 | PHE | A | 206 | 34.274 | 33.147 | 46.281 | 1.00 | 0.00 | H |
| ATOM | 2893 | CG | PHE | A | 206 | 34.760 | 34.963 | 45.567 | 1.00 | 0.00 | C |
| ATOM | 2894 | CD1 | PHE | A | 206 | 34.433 | 34.619 | 44.247 | 1.00 | 0.00 | C |
| ATOM | 2895 | HD1 | PHE | A | 206 | 34.308 | 33.666 | 43.970 | 1.00 | 0.00 | H |
| ATOM | 2896 | CE1 | PHE | A | 206 | 34.288 | 35.692 | 43.319 | 1.00 | 0.00 | C |
| ATOM | 2897 | HE1 | PHE | A | 206 | 34.157 | 35.482 | 42.350 | 1.00 | 0.00 | H |
| ATOM | 2898 | CZ | PHE | A | 206 | 34.324 | 37.053 | 43.734 | 1.00 | 0.00 | C |
| ATOM | 2899 | HZ | PHE | A | 206 | 34.227 | 37.779 | 43.054 | 1.00 | 0.00 | H |
| ATOM | 2900 | CE2 | PHE | A | 206 | 34.491 | 37.383 | 45.065 | 1.00 | 0.00 | C |
| ATOM | 2901 | HE2 | PHE | A | 206 | 34.468 | 38.330 | 45.386 | 1.00 | 0.00 | H |
| ATOM | 2902 | CD2 | PHE | A | 206 | 34.701 | 36.274 | 45.960 | 1.00 | 0.00 | C |
| ATOM | 2903 | HD2 | PHE | A | 206 | 34.811 | 36.475 | 46.933 | 1.00 | 0.00 | H |
| ATOM | 2904 | C | PHE | A | 206 | 36.521 | 32.087 | 45.854 | 1.00 | 0.00 | C |
| ATOM | 2905 | O | PHE | A | 206 | 37.325 | 32.203 | 44.891 | 1.00 | 0.00 | O |
| ATOM | 2906 | N | VAL | A | 207 | 35.779 | 30.963 | 46.065 | 1.00 | 0.00 | N |
| ATOM | 2907 | H | VAL | A | 207 | 35.140 | 30.986 | 46.834 | 1.00 | 0.00 | H |
| ATOM | 2908 | CA | VAL | A | 207 | 35.821 | 29.693 | 45.263 | 1.00 | 0.00 | C |
| ATOM | 2909 | HA | VAL | A | 207 | 36.530 | 29.766 | 44.561 | 1.00 | 0.00 | H |
| ATOM | 2910 | CB | VAL | A | 207 | 36.155 | 28.542 | 46.212 | 1.00 | 0.00 | C |
| ATOM | 2911 | HB | VAL | A | 207 | 35.404 | 28.605 | 46.869 | 1.00 | 0.00 | H |
| ATOM | 2912 | CG1 | VAL | A | 207 | 36.241 | 27.227 | 45.466 | 1.00 | 0.00 | C |
| ATOM | 2913 | HG11 | VAL | A | 207 | 36.460 | 26.493 | 46.109 | 1.00 | 0.00 | H |

| | | | | | | | | | | | |
|---|---|---|---|---|---|---|---|---|---|---|---|
| ATOM | 2914 | HG12 | VAL | A | 207 | 35.363 | 27.035 | 45.027 | 1.00 | 0.00 | H |
| ATOM | 2915 | HG13 | VAL | A | 207 | 36.956 | 27.285 | 44.770 | 1.00 | 0.00 | H |
| ATOM | 2916 | CG2 | VAL | A | 207 | 37.579 | 28.607 | 46.879 | 1.00 | 0.00 | C |
| ATOM | 2917 | HG21 | VAL | A | 207 | 37.703 | 27.816 | 47.478 | 1.00 | 0.00 | H |
| ATOM | 2918 | HG22 | VAL | A | 207 | 38.281 | 28.599 | 46.167 | 1.00 | 0.00 | H |
| ATOM | 2919 | HG23 | VAL | A | 207 | 37.658 | 29.447 | 47.415 | 1.00 | 0.00 | H |
| ATOM | 2920 | C | VAL | A | 207 | 34.475 | 29.465 | 44.556 | 1.00 | 0.00 | C |
| ATOM | 2921 | O | VAL | A | 207 | 33.478 | 29.051 | 45.215 | 1.00 | 0.00 | O |
| ATOM | 2922 | N | ASN | A | 208 | 34.457 | 29.580 | 43.230 | 1.00 | 0.00 | N |
| ATOM | 2923 | H | ASN | A | 208 | 35.304 | 29.842 | 42.768 | 1.00 | 0.00 | H |
| ATOM | 2924 | CA | ASN | A | 208 | 33.270 | 29.345 | 42.410 | 1.00 | 0.00 | C |
| ATOM | 2925 | HA | ASN | A | 208 | 32.468 | 29.561 | 42.967 | 1.00 | 0.00 | H |
| ATOM | 2926 | CB | ASN | A | 208 | 33.310 | 30.298 | 41.198 | 1.00 | 0.00 | C |
| ATOM | 2927 | HB1 | ASN | A | 208 | 32.492 | 30.148 | 40.642 | 1.00 | 0.00 | H |
| ATOM | 2928 | HB2 | ASN | A | 208 | 34.124 | 30.095 | 40.654 | 1.00 | 0.00 | H |
| ATOM | 2929 | CG | ASN | A | 208 | 33.355 | 31.764 | 41.584 | 1.00 | 0.00 | C |
| ATOM | 2930 | OD1 | ASN | A | 208 | 32.660 | 32.216 | 42.481 | 1.00 | 0.00 | O |
| ATOM | 2931 | ND2 | ASN | A | 208 | 34.185 | 32.531 | 40.922 | 1.00 | 0.00 | N |
| ATOM | 2932 | HD21 | ASN | A | 208 | 34.754 | 32.143 | 40.197 | 1.00 | 0.00 | H |
| ATOM | 2933 | HD22 | ASN | A | 208 | 34.249 | 33.505 | 41.141 | 1.00 | 0.00 | H |
| ATOM | 2934 | C | ASN | A | 208 | 33.162 | 27.856 | 42.025 | 1.00 | 0.00 | C |
| ATOM | 2935 | O | ASN | A | 208 | 33.614 | 27.442 | 40.956 | 1.00 | 0.00 | O |
| ATOM | 2936 | N | LEU | A | 209 | 32.561 | 27.031 | 42.889 | 1.00 | 0.00 | N |
| ATOM | 2937 | H | LEU | A | 209 | 32.324 | 27.383 | 43.794 | 1.00 | 0.00 | H |
| ATOM | 2938 | CA | LEU | A | 209 | 32.232 | 25.640 | 42.578 | 1.00 | 0.00 | C |
| ATOM | 2939 | HA | LEU | A | 209 | 33.088 | 25.154 | 42.403 | 1.00 | 0.00 | H |
| ATOM | 2940 | CB | LEU | A | 209 | 31.540 | 25.039 | 43.778 | 1.00 | 0.00 | C |
| ATOM | 2941 | HB1 | LEU | A | 209 | 30.766 | 25.634 | 43.995 | 1.00 | 0.00 | H |
| ATOM | 2942 | HB2 | LEU | A | 209 | 32.197 | 25.059 | 44.532 | 1.00 | 0.00 | H |
| ATOM | 2943 | CG | LEU | A | 209 | 31.014 | 23.564 | 43.599 | 1.00 | 0.00 | C |
| ATOM | 2944 | HG | LEU | A | 209 | 30.410 | 23.614 | 42.803 | 1.00 | 0.00 | H |
| ATOM | 2945 | CD1 | LEU | A | 209 | 32.107 | 22.549 | 43.321 | 1.00 | 0.00 | C |
| ATOM | 2946 | HD11 | LEU | A | 209 | 31.701 | 21.641 | 43.220 | 1.00 | 0.00 | H |
| ATOM | 2947 | HD12 | LEU | A | 209 | 32.585 | 22.796 | 42.478 | 1.00 | 0.00 | H |
| ATOM | 2948 | HD13 | LEU | A | 209 | 32.756 | 22.541 | 44.082 | 1.00 | 0.00 | H |
| ATOM | 2949 | CD2 | LEU | A | 209 | 30.213 | 23.036 | 44.776 | 1.00 | 0.00 | C |
| ATOM | 2950 | HD21 | LEU | A | 209 | 29.917 | 22.100 | 44.585 | 1.00 | 0.00 | H |
| ATOM | 2951 | HD22 | LEU | A | 209 | 30.783 | 23.044 | 45.598 | 1.00 | 0.00 | H |
| ATOM | 2952 | HD23 | LEU | A | 209 | 29.411 | 23.616 | 44.920 | 1.00 | 0.00 | H |
| ATOM | 2953 | C | LEU | A | 209 | 31.329 | 25.568 | 41.326 | 1.00 | 0.00 | C |
| ATOM | 2954 | O | LEU | A | 209 | 30.177 | 26.004 | 41.339 | 1.00 | 0.00 | O |
| ATOM | 2955 | N | MET | A | 210 | 31.830 | 24.956 | 40.257 | 1.00 | 0.00 | N |
| ATOM | 2956 | H | MET | A | 210 | 32.765 | 24.611 | 40.335 | 1.00 | 0.00 | H |
| ATOM | 2957 | CA | MET | A | 210 | 31.153 | 24.736 | 38.978 | 1.00 | 0.00 | C |
| ATOM | 2958 | HA | MET | A | 210 | 30.616 | 25.558 | 38.789 | 1.00 | 0.00 | H |
| ATOM | 2959 | CB | MET | A | 210 | 32.197 | 24.544 | 37.862 | 1.00 | 0.00 | C |
| ATOM | 2960 | HB1 | MET | A | 210 | 31.722 | 24.367 | 37.000 | 1.00 | 0.00 | H |
| ATOM | 2961 | HB2 | MET | A | 210 | 32.776 | 23.762 | 38.090 | 1.00 | 0.00 | H |
| ATOM | 2962 | CG | MET | A | 210 | 33.085 | 25.783 | 37.691 | 1.00 | 0.00 | C |
| ATOM | 2963 | HG1 | MET | A | 210 | 33.603 | 25.840 | 38.545 | 1.00 | 0.00 | H |
| ATOM | 2964 | HG2 | MET | A | 210 | 32.442 | 26.547 | 37.639 | 1.00 | 0.00 | H |
| ATOM | 2965 | SD | MET | A | 210 | 34.172 | 25.720 | 36.248 | 1.00 | 0.00 | S |
| ATOM | 2966 | CE | MET | A | 210 | 35.497 | 24.602 | 36.779 | 1.00 | 0.00 | C |
| ATOM | 2967 | HE1 | MET | A | 210 | 36.162 | 24.498 | 36.039 | 1.00 | 0.00 | H |

| | | | | | | | | | | | |
|---|---|---|---|---|---|---|---|---|---|---|---|
| ATOM | 2968 | HE2 | MET | A | 210 | 35.951 | 24.982 | 37.585 | 1.00 | 0.00 | H |
| ATOM | 2969 | HE3 | MET | A | 210 | 35.109 | 23.709 | 37.006 | 1.00 | 0.00 | H |
| ATOM | 2970 | C | MET | A | 210 | 30.188 | 23.550 | 39.081 | 1.00 | 0.00 | C |
| ATOM | 2971 | O | MET | A | 210 | 30.439 | 22.464 | 38.559 | 1.00 | 0.00 | O |
| ATOM | 2972 | N | ALA | A | 211 | 29.086 | 23.737 | 39.811 | 1.00 | 0.00 | N |
| ATOM | 2973 | H | ALA | A | 211 | 28.947 | 24.636 | 40.226 | 1.00 | 0.00 | H |
| ATOM | 2974 | CA | ALA | A | 211 | 28.067 | 22.709 | 40.044 | 1.00 | 0.00 | C |
| ATOM | 2975 | HA | ALA | A | 211 | 28.552 | 21.836 | 40.099 | 1.00 | 0.00 | H |
| ATOM | 2976 | CB | ALA | A | 211 | 27.400 | 22.971 | 41.398 | 1.00 | 0.00 | C |
| ATOM | 2977 | HB1 | ALA | A | 211 | 26.703 | 22.274 | 41.566 | 1.00 | 0.00 | H |
| ATOM | 2978 | HB2 | ALA | A | 211 | 28.089 | 22.934 | 42.122 | 1.00 | 0.00 | H |
| ATOM | 2979 | HB3 | ALA | A | 211 | 26.973 | 23.875 | 41.390 | 1.00 | 0.00 | H |
| ATOM | 2980 | C | ALA | A | 211 | 27.093 | 22.539 | 38.852 | 1.00 | 0.00 | C |
| ATOM | 2981 | O | ALA | A | 211 | 25.880 | 22.377 | 39.015 | 1.00 | 0.00 | O |
| ATOM | 2982 | N | TYR | A | 212 | 27.649 | 22.573 | 37.644 | 1.00 | 0.00 | N |
| ATOM | 2983 | H | TYR | A | 212 | 28.631 | 22.757 | 37.615 | 1.00 | 0.00 | H |
| ATOM | 2984 | CA | TYR | A | 212 | 26.993 | 22.373 | 36.356 | 1.00 | 0.00 | C |
| ATOM | 2985 | HA | TYR | A | 212 | 26.070 | 22.051 | 36.568 | 1.00 | 0.00 | H |
| ATOM | 2986 | CB | TYR | A | 212 | 26.890 | 23.701 | 35.578 | 1.00 | 0.00 | C |
| ATOM | 2987 | HB1 | TYR | A | 212 | 25.965 | 24.055 | 35.714 | 1.00 | 0.00 | H |
| ATOM | 2988 | HB2 | TYR | A | 212 | 27.028 | 23.491 | 34.610 | 1.00 | 0.00 | H |
| ATOM | 2989 | CG | TYR | A | 212 | 27.846 | 24.834 | 35.920 | 1.00 | 0.00 | C |
| ATOM | 2990 | CD1 | TYR | A | 212 | 29.133 | 24.877 | 35.350 | 1.00 | 0.00 | C |
| ATOM | 2991 | HD1 | TYR | A | 212 | 29.468 | 24.102 | 34.814 | 1.00 | 0.00 | H |
| ATOM | 2992 | CE1 | TYR | A | 212 | 29.950 | 26.011 | 35.533 | 1.00 | 0.00 | C |
| ATOM | 2993 | HE1 | TYR | A | 212 | 30.865 | 26.038 | 35.130 | 1.00 | 0.00 | H |
| ATOM | 2994 | CZ | TYR | A | 212 | 29.478 | 27.107 | 36.284 | 1.00 | 0.00 | C |
| ATOM | 2995 | OH | TYR | A | 212 | 30.249 | 28.217 | 36.411 | 1.00 | 0.00 | O |
| ATOM | 2996 | HH | TYR | A | 212 | 29.770 | 28.894 | 36.970 | 1.00 | 0.00 | H |
| ATOM | 2997 | CE2 | TYR | A | 212 | 28.204 | 27.047 | 36.891 | 1.00 | 0.00 | C |
| ATOM | 2998 | HE2 | TYR | A | 212 | 27.878 | 27.812 | 37.447 | 1.00 | 0.00 | H |
| ATOM | 2999 | CD2 | TYR | A | 212 | 27.394 | 25.909 | 36.713 | 1.00 | 0.00 | C |
| ATOM | 3000 | HD2 | TYR | A | 212 | 26.496 | 25.863 | 37.150 | 1.00 | 0.00 | H |
| ATOM | 3001 | C | TYR | A | 212 | 27.727 | 21.294 | 35.551 | 1.00 | 0.00 | C |
| ATOM | 3002 | O | TYR | A | 212 | 28.785 | 20.806 | 35.950 | 1.00 | 0.00 | O |
| ATOM | 3003 | N | ASP | A | 213 | 27.157 | 20.911 | 34.409 | 1.00 | 0.00 | N |
| ATOM | 3004 | H | ASP | A | 213 | 26.275 | 21.318 | 34.171 | 1.00 | 0.00 | H |
| ATOM | 3005 | CA | ASP | A | 213 | 27.729 | 19.938 | 33.479 | 1.00 | 0.00 | C |
| ATOM | 3006 | HA | ASP | A | 213 | 27.023 | 19.817 | 32.781 | 1.00 | 0.00 | H |
| ATOM | 3007 | CB | ASP | A | 213 | 28.991 | 20.518 | 32.803 | 1.00 | 0.00 | C |
| ATOM | 3008 | HB1 | ASP | A | 213 | 29.166 | 19.995 | 31.969 | 1.00 | 0.00 | H |
| ATOM | 3009 | HB2 | ASP | A | 213 | 29.760 | 20.409 | 33.432 | 1.00 | 0.00 | H |
| ATOM | 3010 | CG | ASP | A | 213 | 28.898 | 22.004 | 32.421 | 1.00 | 0.00 | C |
| ATOM | 3011 | OD1 | ASP | A | 213 | 29.899 | 22.725 | 32.639 | 1.00 | 0.00 | O |
| ATOM | 3012 | OD2 | ASP | A | 213 | 27.860 | 22.422 | 31.842 | 1.00 | 0.00 | O |
| ATOM | 3013 | C | ASP | A | 213 | 27.936 | 18.539 | 34.111 | 1.00 | 0.00 | C |
| ATOM | 3014 | O | ASP | A | 213 | 28.745 | 17.739 | 33.637 | 1.00 | 0.00 | O |
| ATOM | 3015 | N | PHE | A | 214 | 27.203 | 18.208 | 35.187 | 1.00 | 0.00 | N |
| ATOM | 3016 | H | PHE | A | 214 | 26.556 | 18.885 | 35.538 | 1.00 | 0.00 | H |
| ATOM | 3017 | CA | PHE | A | 214 | 27.291 | 16.911 | 35.885 | 1.00 | 0.00 | C |
| ATOM | 3018 | HA | PHE | A | 214 | 28.247 | 16.852 | 36.173 | 1.00 | 0.00 | H |
| ATOM | 3019 | CB | PHE | A | 214 | 26.333 | 16.893 | 37.083 | 1.00 | 0.00 | C |
| ATOM | 3020 | HB1 | PHE | A | 214 | 26.130 | 15.940 | 37.306 | 1.00 | 0.00 | H |
| ATOM | 3021 | HB2 | PHE | A | 214 | 25.490 | 17.360 | 36.816 | 1.00 | 0.00 | H |

| | | | | | | | | | | | |
|---|---|---|---|---|---|---|---|---|---|---|---|
| ATOM | 3022 | CG | PHE | A | 214 | 26.849 | 17.560 | 38.343 | 1.00 | 0.00 | C |
| ATOM | 3023 | CD1 | PHE | A | 214 | 26.290 | 18.767 | 38.788 | 1.00 | 0.00 | C |
| ATOM | 3024 | HD1 | PHE | A | 214 | 25.639 | 19.257 | 38.208 | 1.00 | 0.00 | H |
| ATOM | 3025 | CE1 | PHE | A | 214 | 26.639 | 19.291 | 40.044 | 1.00 | 0.00 | C |
| ATOM | 3026 | HE1 | PHE | A | 214 | 26.210 | 20.133 | 40.371 | 1.00 | 0.00 | H |
| ATOM | 3027 | CZ | PHE | A | 214 | 27.590 | 18.632 | 40.840 | 1.00 | 0.00 | C |
| ATOM | 3028 | HZ | PHE | A | 214 | 27.829 | 18.997 | 41.740 | 1.00 | 0.00 | H |
| ATOM | 3029 | CE2 | PHE | A | 214 | 28.205 | 17.459 | 40.371 | 1.00 | 0.00 | C |
| ATOM | 3030 | HE2 | PHE | A | 214 | 28.908 | 17.007 | 40.921 | 1.00 | 0.00 | H |
| ATOM | 3031 | CD2 | PHE | A | 214 | 27.829 | 16.922 | 39.125 | 1.00 | 0.00 | C |
| ATOM | 3032 | HD2 | PHE | A | 214 | 28.261 | 16.083 | 38.794 | 1.00 | 0.00 | H |
| ATOM | 3033 | C | PHE | A | 214 | 26.981 | 15.715 | 34.977 | 1.00 | 0.00 | C |
| ATOM | 3034 | O | PHE | A | 214 | 27.577 | 14.655 | 35.142 | 1.00 | 0.00 | O |
| ATOM | 3035 | N | HIS | A | 215 | 26.088 | 15.909 | 34.005 | 1.00 | 0.00 | N |
| ATOM | 3036 | H | HIS | A | 215 | 25.618 | 16.792 | 34.004 | 1.00 | 0.00 | H |
| ATOM | 3037 | CA | HIS | A | 215 | 25.718 | 14.975 | 32.939 | 1.00 | 0.00 | C |
| ATOM | 3038 | HA | HIS | A | 215 | 26.503 | 14.382 | 32.761 | 1.00 | 0.00 | H |
| ATOM | 3039 | CB | HIS | A | 215 | 24.525 | 14.126 | 33.404 | 1.00 | 0.00 | C |
| ATOM | 3040 | HB1 | HIS | A | 215 | 24.238 | 13.548 | 32.640 | 1.00 | 0.00 | H |
| ATOM | 3041 | HB2 | HIS | A | 215 | 23.780 | 14.746 | 33.651 | 1.00 | 0.00 | H |
| ATOM | 3042 | CG | HIS | A | 215 | 24.772 | 13.219 | 34.591 | 1.00 | 0.00 | C |
| ATOM | 3043 | ND1 | HIS | A | 215 | 25.711 | 12.207 | 34.661 | 1.00 | 0.00 | N |
| ATOM | 3044 | HD1 | HIS | A | 215 | 26.380 | 11.978 | 33.954 | 1.00 | 0.00 | H |
| ATOM | 3045 | CE1 | HIS | A | 215 | 25.570 | 11.583 | 35.846 | 1.00 | 0.00 | C |
| ATOM | 3046 | HE1 | HIS | A | 215 | 26.115 | 10.806 | 36.161 | 1.00 | 0.00 | H |
| ATOM | 3047 | NE2 | HIS | A | 215 | 24.574 | 12.176 | 36.542 | 1.00 | 0.00 | N |
| ATOM | 3048 | CD2 | HIS | A | 215 | 24.069 | 13.213 | 35.769 | 1.00 | 0.00 | C |
| ATOM | 3049 | HD2 | HIS | A | 215 | 23.331 | 13.838 | 36.023 | 1.00 | 0.00 | H |
| ATOM | 3050 | C | HIS | A | 215 | 25.359 | 15.782 | 31.676 | 1.00 | 0.00 | C |
| ATOM | 3051 | O | HIS | A | 215 | 25.107 | 16.987 | 31.760 | 1.00 | 0.00 | O |
| ATOM | 3052 | N | GLY | A | 216 | 25.322 | 15.141 | 30.509 | 1.00 | 0.00 | N |
| ATOM | 3053 | H | GLY | A | 216 | 25.498 | 14.157 | 30.503 | 1.00 | 0.00 | H |
| ATOM | 3054 | CA | GLY | A | 216 | 25.036 | 15.804 | 29.232 | 1.00 | 0.00 | C |
| ATOM | 3055 | HA1 | GLY | A | 216 | 25.642 | 16.596 | 29.155 | 1.00 | 0.00 | H |
| ATOM | 3056 | HA2 | GLY | A | 216 | 24.085 | 16.113 | 29.253 | 1.00 | 0.00 | H |
| ATOM | 3057 | C | GLY | A | 216 | 25.248 | 14.877 | 28.035 | 1.00 | 0.00 | C |
| ATOM | 3058 | O | GLY | A | 216 | 25.397 | 13.675 | 28.205 | 1.00 | 0.00 | O |
| ATOM | 3059 | N | SER | A | 217 | 25.282 | 15.427 | 26.820 | 1.00 | 0.00 | N |
| ATOM | 3060 | H | SER | A | 217 | 25.417 | 16.417 | 26.771 | 1.00 | 0.00 | H |
| ATOM | 3061 | CA | SER | A | 217 | 25.138 | 14.701 | 25.537 | 1.00 | 0.00 | C |
| ATOM | 3062 | HA | SER | A | 217 | 24.243 | 14.255 | 25.560 | 1.00 | 0.00 | H |
| ATOM | 3063 | CB | SER | A | 217 | 25.322 | 15.754 | 24.454 | 1.00 | 0.00 | C |
| ATOM | 3064 | HB1 | SER | A | 217 | 25.149 | 15.365 | 23.549 | 1.00 | 0.00 | H |
| ATOM | 3065 | HB2 | SER | A | 217 | 26.247 | 16.133 | 24.482 | 1.00 | 0.00 | H |
| ATOM | 3066 | OG | SER | A | 217 | 24.411 | 16.821 | 24.648 | 1.00 | 0.00 | O |
| ATOM | 3067 | HG | SER | A | 217 | 24.542 | 17.506 | 23.931 | 1.00 | 0.00 | H |
| ATOM | 3068 | C | SER | A | 217 | 26.056 | 13.543 | 25.305 | 1.00 | 0.00 | C |
| ATOM | 3069 | O | SER | A | 217 | 25.812 | 12.727 | 24.420 | 1.00 | 0.00 | O |
| ATOM | 3070 | N | TRP | A | 218 | 27.194 | 13.509 | 26.007 | 1.00 | 0.00 | N |
| ATOM | 3071 | H | TRP | A | 218 | 27.345 | 14.257 | 26.654 | 1.00 | 0.00 | H |
| ATOM | 3072 | CA | TRP | A | 218 | 28.245 | 12.486 | 25.925 | 1.00 | 0.00 | C |
| ATOM | 3073 | HA | TRP | A | 218 | 28.333 | 12.273 | 24.952 | 1.00 | 0.00 | H |
| ATOM | 3074 | CB | TRP | A | 218 | 29.516 | 13.089 | 26.546 | 1.00 | 0.00 | C |
| ATOM | 3075 | HB1 | TRP | A | 218 | 29.788 | 13.887 | 26.009 | 1.00 | 0.00 | H |

| | | | | | | | | | | | |
|---|---|---|---|---|---|---|---|---|---|---|---|
| ATOM | 3076 | HB2 | TRP | A | 218 | 30.244 | 12.404 | 26.517 | 1.00 | 0.00 | H |
| ATOM | 3077 | CG | TRP | A | 218 | 29.358 | 13.530 | 27.976 | 1.00 | 0.00 | C |
| ATOM | 3078 | CD1 | TRP | A | 218 | 29.470 | 12.725 | 29.056 | 1.00 | 0.00 | C |
| ATOM | 3079 | HD1 | TRP | A | 218 | 29.736 | 11.761 | 29.028 | 1.00 | 0.00 | H |
| ATOM | 3080 | NE1 | TRP | A | 218 | 29.163 | 13.429 | 30.204 | 1.00 | 0.00 | N |
| ATOM | 3081 | HE1 | TRP | A | 218 | 29.182 | 13.041 | 31.126 | 1.00 | 0.00 | H |
| ATOM | 3082 | CE2 | TRP | A | 218 | 28.828 | 14.733 | 29.917 | 1.00 | 0.00 | C |
| ATOM | 3083 | CZ2 | TRP | A | 218 | 28.435 | 15.818 | 30.713 | 1.00 | 0.00 | C |
| ATOM | 3084 | HZ2 | TRP | A | 218 | 28.322 | 15.706 | 31.700 | 1.00 | 0.00 | H |
| ATOM | 3085 | CH2 | TRP | A | 218 | 28.201 | 17.065 | 30.108 | 1.00 | 0.00 | C |
| ATOM | 3086 | HH2 | TRP | A | 218 | 27.935 | 17.851 | 30.666 | 1.00 | 0.00 | H |
| ATOM | 3087 | CZ3 | TRP | A | 218 | 28.342 | 17.205 | 28.715 | 1.00 | 0.00 | C |
| ATOM | 3088 | HZ3 | TRP | A | 218 | 28.178 | 18.093 | 28.286 | 1.00 | 0.00 | H |
| ATOM | 3089 | CE3 | TRP | A | 218 | 28.713 | 16.102 | 27.920 | 1.00 | 0.00 | C |
| ATOM | 3090 | HE3 | TRP | A | 218 | 28.794 | 16.215 | 26.930 | 1.00 | 0.00 | H |
| ATOM | 3091 | CD2 | TRP | A | 218 | 28.972 | 14.839 | 28.500 | 1.00 | 0.00 | C |
| ATOM | 3092 | C | TRP | A | 218 | 27.896 | 11.171 | 26.645 | 1.00 | 0.00 | C |
| ATOM | 3093 | O | TRP | A | 218 | 28.541 | 10.151 | 26.408 | 1.00 | 0.00 | O |
| ATOM | 3094 | N | GLU | A | 219 | 26.941 | 11.205 | 27.575 | 1.00 | 0.00 | N |
| ATOM | 3095 | H | GLU | A | 219 | 26.453 | 12.067 | 27.715 | 1.00 | 0.00 | H |
| ATOM | 3096 | CA | GLU | A | 219 | 26.562 | 10.060 | 28.404 | 1.00 | 0.00 | C |
| ATOM | 3097 | HA | GLU | A | 219 | 27.426 | 9.691 | 28.747 | 1.00 | 0.00 | H |
| ATOM | 3098 | CB | GLU | A | 219 | 25.644 | 10.528 | 29.537 | 1.00 | 0.00 | C |
| ATOM | 3099 | HB1 | GLU | A | 219 | 25.430 | 9.733 | 30.105 | 1.00 | 0.00 | H |
| ATOM | 3100 | HB2 | GLU | A | 219 | 24.804 | 10.879 | 29.124 | 1.00 | 0.00 | H |
| ATOM | 3101 | CG | GLU | A | 219 | 26.204 | 11.617 | 30.452 | 1.00 | 0.00 | C |
| ATOM | 3102 | HG1 | GLU | A | 219 | 25.446 | 12.005 | 30.976 | 1.00 | 0.00 | H |
| ATOM | 3103 | HG2 | GLU | A | 219 | 26.611 | 12.328 | 29.879 | 1.00 | 0.00 | H |
| ATOM | 3104 | CD | GLU | A | 219 | 27.263 | 11.145 | 31.437 | 1.00 | 0.00 | C |
| ATOM | 3105 | OE1 | GLU | A | 219 | 27.327 | 11.810 | 32.493 | 1.00 | 0.00 | O |
| ATOM | 3106 | OE2 | GLU | A | 219 | 28.041 | 10.208 | 31.155 | 1.00 | 0.00 | O |
| ATOM | 3107 | C | GLU | A | 219 | 25.813 | 8.998 | 27.589 | 1.00 | 0.00 | C |
| ATOM | 3108 | O | GLU | A | 219 | 25.122 | 9.317 | 26.612 | 1.00 | 0.00 | O |
| ATOM | 3109 | N | LYS | A | 220 | 25.946 | 7.737 | 27.932 | 1.00 | 0.00 | N |
| ATOM | 3110 | H | LYS | A | 220 | 26.641 | 7.476 | 28.602 | 1.00 | 0.00 | H |
| ATOM | 3111 | CA | LYS | A | 220 | 25.070 | 6.692 | 27.331 | 1.00 | 0.00 | C |
| ATOM | 3112 | HA | LYS | A | 220 | 25.219 | 6.729 | 26.343 | 1.00 | 0.00 | H |
| ATOM | 3113 | CB | LYS | A | 220 | 25.456 | 5.311 | 27.877 | 1.00 | 0.00 | C |
| ATOM | 3114 | HB1 | LYS | A | 220 | 24.735 | 4.659 | 27.641 | 1.00 | 0.00 | H |
| ATOM | 3115 | HB2 | LYS | A | 220 | 25.538 | 5.371 | 28.872 | 1.00 | 0.00 | H |
| ATOM | 3116 | CG | LYS | A | 220 | 26.796 | 4.814 | 27.290 | 1.00 | 0.00 | C |
| ATOM | 3117 | HG1 | LYS | A | 220 | 27.533 | 5.422 | 27.585 | 1.00 | 0.00 | H |
| ATOM | 3118 | HG2 | LYS | A | 220 | 26.742 | 4.816 | 26.291 | 1.00 | 0.00 | H |
| ATOM | 3119 | CD | LYS | A | 220 | 27.145 | 3.432 | 27.723 | 1.00 | 0.00 | C |
| ATOM | 3120 | HD1 | LYS | A | 220 | 26.418 | 2.804 | 27.446 | 1.00 | 0.00 | H |
| ATOM | 3121 | HD2 | LYS | A | 220 | 27.243 | 3.411 | 28.718 | 1.00 | 0.00 | H |
| ATOM | 3122 | CE | LYS | A | 220 | 28.466 | 3.053 | 27.047 | 1.00 | 0.00 | C |
| ATOM | 3123 | HE1 | LYS | A | 220 | 29.168 | 3.714 | 27.312 | 1.00 | 0.00 | H |
| ATOM | 3124 | HE2 | LYS | A | 220 | 28.342 | 3.078 | 26.055 | 1.00 | 0.00 | H |
| ATOM | 3125 | NZ | LYS | A | 220 | 29.024 | 1.769 | 27.330 | 1.00 | 0.00 | N |
| ATOM | 3126 | HZ1 | LYS | A | 220 | 29.879 | 1.658 | 26.824 | 1.00 | 0.00 | H |
| ATOM | 3127 | HZ2 | LYS | A | 220 | 29.205 | 1.693 | 28.311 | 1.00 | 0.00 | H |
| ATOM | 3128 | HZ3 | LYS | A | 220 | 28.379 | 1.057 | 27.053 | 1.00 | 0.00 | H |
| ATOM | 3129 | C | LYS | A | 220 | 23.597 | 6.976 | 27.587 | 1.00 | 0.00 | C |

| | | | | | | | | | | | |
|---|---|---|---|---|---|---|---|---|---|---|---|
| ATOM | 3130 | O | LYS | A | 220 | 22.820 | 6.630 | 26.667 | 1.00 | 0.00 | O |
| ATOM | 3131 | N | VAL | A | 221 | 23.269 | 7.555 | 28.750 | 1.00 | 0.00 | N |
| ATOM | 3132 | H | VAL | A | 221 | 23.985 | 7.748 | 29.421 | 1.00 | 0.00 | H |
| ATOM | 3133 | CA | VAL | A | 221 | 21.873 | 7.913 | 29.060 | 1.00 | 0.00 | C |
| ATOM | 3134 | HA | VAL | A | 221 | 21.419 | 7.836 | 28.172 | 1.00 | 0.00 | H |
| ATOM | 3135 | CB | VAL | A | 221 | 21.362 | 6.959 | 30.204 | 1.00 | 0.00 | C |
| ATOM | 3136 | HB | VAL | A | 221 | 20.428 | 7.276 | 30.369 | 1.00 | 0.00 | H |
| ATOM | 3137 | CG1 | VAL | A | 221 | 21.454 | 5.463 | 29.801 | 1.00 | 0.00 | C |
| ATOM | 3138 | HG11 | VAL | A | 221 | 21.121 | 4.894 | 30.553 | 1.00 | 0.00 | H |
| ATOM | 3139 | HG12 | VAL | A | 221 | 20.895 | 5.302 | 28.988 | 1.00 | 0.00 | H |
| ATOM | 3140 | HG13 | VAL | A | 221 | 22.406 | 5.229 | 29.602 | 1.00 | 0.00 | H |
| ATOM | 3141 | CG2 | VAL | A | 221 | 22.154 | 7.028 | 31.504 | 1.00 | 0.00 | C |
| ATOM | 3142 | HG21 | VAL | A | 221 | 21.760 | 6.393 | 32.168 | 1.00 | 0.00 | H |
| ATOM | 3143 | HG22 | VAL | A | 221 | 23.106 | 6.778 | 31.328 | 1.00 | 0.00 | H |
| ATOM | 3144 | HG23 | VAL | A | 221 | 22.115 | 7.958 | 31.869 | 1.00 | 0.00 | H |
| ATOM | 3145 | C | VAL | A | 221 | 21.634 | 9.334 | 29.488 | 1.00 | 0.00 | C |
| ATOM | 3146 | O | VAL | A | 221 | 22.639 | 9.986 | 29.911 | 1.00 | 0.00 | O |
| ATOM | 3147 | N | THR | A | 222 | 20.352 | 9.788 | 29.449 | 1.00 | 0.00 | N |
| ATOM | 3148 | H | THR | A | 222 | 19.632 | 9.166 | 29.141 | 1.00 | 0.00 | H |
| ATOM | 3149 | CA | THR | A | 222 | 19.973 | 11.191 | 29.851 | 1.00 | 0.00 | C |
| ATOM | 3150 | HA | THR | A | 222 | 20.533 | 11.816 | 29.307 | 1.00 | 0.00 | H |
| ATOM | 3151 | CB | THR | A | 222 | 18.446 | 11.406 | 29.566 | 1.00 | 0.00 | C |
| ATOM | 3152 | HB | THR | A | 222 | 18.231 | 12.317 | 29.918 | 1.00 | 0.00 | H |
| ATOM | 3153 | CG2 | THR | A | 222 | 18.032 | 11.269 | 28.119 | 1.00 | 0.00 | C |
| ATOM | 3154 | HG21 | THR | A | 222 | 17.047 | 11.422 | 28.038 | 1.00 | 0.00 | H |
| ATOM | 3155 | HG22 | THR | A | 222 | 18.521 | 11.944 | 27.567 | 1.00 | 0.00 | H |
| ATOM | 3156 | HG23 | THR | A | 222 | 18.254 | 10.350 | 27.794 | 1.00 | 0.00 | H |
| ATOM | 3157 | OG1 | THR | A | 222 | 17.629 | 10.505 | 30.244 | 1.00 | 0.00 | O |
| ATOM | 3158 | HG1 | THR | A | 222 | 16.672 | 10.692 | 30.023 | 1.00 | 0.00 | H |
| ATOM | 3159 | C | THR | A | 222 | 20.281 | 11.442 | 31.354 | 1.00 | 0.00 | C |
| ATOM | 3160 | O | THR | A | 222 | 19.933 | 10.649 | 32.230 | 1.00 | 0.00 | O |
| ATOM | 3161 | N | GLY | A | 223 | 20.872 | 12.601 | 31.674 | 1.00 | 0.00 | N |
| ATOM | 3162 | H | GLY | A | 223 | 21.178 | 13.193 | 30.928 | 1.00 | 0.00 | H |
| ATOM | 3163 | CA | GLY | A | 223 | 21.098 | 13.054 | 33.047 | 1.00 | 0.00 | C |
| ATOM | 3164 | HA1 | GLY | A | 223 | 21.963 | 12.665 | 33.364 | 1.00 | 0.00 | H |
| ATOM | 3165 | HA2 | GLY | A | 223 | 20.350 | 12.710 | 33.614 | 1.00 | 0.00 | H |
| ATOM | 3166 | C | GLY | A | 223 | 21.159 | 14.541 | 33.174 | 1.00 | 0.00 | C |
| ATOM | 3167 | O | GLY | A | 223 | 21.490 | 15.246 | 32.234 | 1.00 | 0.00 | O |
| ATOM | 3168 | N | HIS | A | 224 | 20.784 | 15.026 | 34.315 | 1.00 | 0.00 | N |
| ATOM | 3169 | H | HIS | A | 224 | 20.689 | 14.396 | 35.086 | 1.00 | 0.00 | H |
| ATOM | 3170 | CA | HIS | A | 224 | 20.490 | 16.446 | 34.544 | 1.00 | 0.00 | C |
| ATOM | 3171 | HA | HIS | A | 224 | 19.926 | 16.793 | 33.795 | 1.00 | 0.00 | H |
| ATOM | 3172 | CB | HIS | A | 224 | 19.701 | 16.579 | 35.840 | 1.00 | 0.00 | C |
| ATOM | 3173 | HB1 | HIS | A | 224 | 20.303 | 16.446 | 36.627 | 1.00 | 0.00 | H |
| ATOM | 3174 | HB2 | HIS | A | 224 | 18.963 | 15.904 | 35.867 | 1.00 | 0.00 | H |
| ATOM | 3175 | CG | HIS | A | 224 | 19.119 | 17.947 | 35.890 | 1.00 | 0.00 | C |
| ATOM | 3176 | ND1 | HIS | A | 224 | 19.748 | 19.072 | 36.434 | 1.00 | 0.00 | N |
| ATOM | 3177 | CE1 | HIS | A | 224 | 18.907 | 20.083 | 36.335 | 1.00 | 0.00 | C |
| ATOM | 3178 | HE1 | HIS | A | 224 | 19.091 | 21.018 | 36.638 | 1.00 | 0.00 | H |
| ATOM | 3179 | NE2 | HIS | A | 224 | 17.787 | 19.673 | 35.776 | 1.00 | 0.00 | N |
| ATOM | 3180 | HE2 | HIS | A | 224 | 16.984 | 20.241 | 35.594 | 1.00 | 0.00 | H |
| ATOM | 3181 | CD2 | HIS | A | 224 | 17.906 | 18.325 | 35.483 | 1.00 | 0.00 | C |
| ATOM | 3182 | HD2 | HIS | A | 224 | 17.215 | 17.746 | 35.050 | 1.00 | 0.00 | H |
| ATOM | 3183 | C | HIS | A | 224 | 21.806 | 17.228 | 34.568 | 1.00 | 0.00 | C |

| | | | | | | | | | | | |
|---|---|---|---|---|---|---|---|---|---|---|---|
| ATOM | 3184 | O | HIS | A | 224 | 22.836 | 16.749 | 35.046 | 1.00 | 0.00 | O |
| ATOM | 3185 | N | ASN | A | 225 | 21.721 | 18.520 | 34.111 | 1.00 | 0.00 | N |
| ATOM | 3186 | H | ASN | A | 225 | 20.841 | 18.843 | 33.764 | 1.00 | 0.00 | H |
| ATOM | 3187 | CA | ASN | A | 225 | 22.850 | 19.446 | 34.106 | 1.00 | 0.00 | C |
| ATOM | 3188 | HA | ASN | A | 225 | 23.528 | 19.053 | 33.485 | 1.00 | 0.00 | H |
| ATOM | 3189 | CB | ASN | A | 225 | 22.299 | 20.823 | 33.664 | 1.00 | 0.00 | C |
| ATOM | 3190 | HB1 | ASN | A | 225 | 21.682 | 21.162 | 34.374 | 1.00 | 0.00 | H |
| ATOM | 3191 | HB2 | ASN | A | 225 | 21.792 | 20.704 | 32.810 | 1.00 | 0.00 | H |
| ATOM | 3192 | CG | ASN | A | 225 | 23.368 | 21.883 | 33.429 | 1.00 | 0.00 | C |
| ATOM | 3193 | OD1 | ASN | A | 225 | 24.549 | 21.608 | 33.296 | 1.00 | 0.00 | O |
| ATOM | 3194 | ND2 | ASN | A | 225 | 22.970 | 23.113 | 33.205 | 1.00 | 0.00 | N |
| ATOM | 3195 | HD21 | ASN | A | 225 | 21.993 | 23.328 | 33.193 | 1.00 | 0.00 | H |
| ATOM | 3196 | HD22 | ASN | A | 225 | 23.643 | 23.835 | 33.047 | 1.00 | 0.00 | H |
| ATOM | 3197 | C | ASN | A | 225 | 23.425 | 19.635 | 35.556 | 1.00 | 0.00 | C |
| ATOM | 3198 | O | ASN | A | 225 | 24.616 | 19.887 | 35.726 | 1.00 | 0.00 | O |
| ATOM | 3199 | N | SER | A | 226 | 22.568 | 19.631 | 36.598 | 1.00 | 0.00 | N |
| ATOM | 3200 | H | SER | A | 226 | 21.722 | 19.113 | 36.474 | 1.00 | 0.00 | H |
| ATOM | 3201 | CA | SER | A | 226 | 22.738 | 20.304 | 37.899 | 1.00 | 0.00 | C |
| ATOM | 3202 | HA | SER | A | 226 | 23.645 | 20.037 | 38.224 | 1.00 | 0.00 | H |
| ATOM | 3203 | CB | SER | A | 226 | 22.698 | 21.837 | 37.665 | 1.00 | 0.00 | C |
| ATOM | 3204 | HB1 | SER | A | 226 | 21.764 | 22.156 | 37.501 | 1.00 | 0.00 | H |
| ATOM | 3205 | HB2 | SER | A | 226 | 23.281 | 22.103 | 36.897 | 1.00 | 0.00 | H |
| ATOM | 3206 | OG | SER | A | 226 | 23.203 | 22.461 | 38.879 | 1.00 | 0.00 | O |
| ATOM | 3207 | HG | SER | A | 226 | 23.195 | 23.455 | 38.771 | 1.00 | 0.00 | H |
| ATOM | 3208 | C | SER | A | 226 | 21.729 | 19.803 | 38.997 | 1.00 | 0.00 | C |
| ATOM | 3209 | O | SER | A | 226 | 21.076 | 20.609 | 39.689 | 1.00 | 0.00 | O |
| ATOM | 3210 | N | PRO | A | 227 | 21.585 | 18.453 | 39.228 | 1.00 | 0.00 | N |
| ATOM | 3211 | CD | PRO | A | 227 | 22.349 | 17.352 | 38.646 | 1.00 | 0.00 | C |
| ATOM | 3212 | HD1 | PRO | A | 227 | 23.323 | 17.553 | 38.749 | 1.00 | 0.00 | H |
| ATOM | 3213 | HD2 | PRO | A | 227 | 22.122 | 17.288 | 37.674 | 1.00 | 0.00 | H |
| ATOM | 3214 | CG | PRO | A | 227 | 21.975 | 16.043 | 39.387 | 1.00 | 0.00 | C |
| ATOM | 3215 | HG1 | PRO | A | 227 | 22.616 | 15.829 | 40.124 | 1.00 | 0.00 | H |
| ATOM | 3216 | HG2 | PRO | A | 227 | 21.909 | 15.262 | 38.766 | 1.00 | 0.00 | H |
| ATOM | 3217 | CB | PRO | A | 227 | 20.612 | 16.446 | 39.928 | 1.00 | 0.00 | C |
| ATOM | 3218 | HB1 | PRO | A | 227 | 20.402 | 15.924 | 40.755 | 1.00 | 0.00 | H |
| ATOM | 3219 | HB2 | PRO | A | 227 | 19.906 | 16.279 | 39.240 | 1.00 | 0.00 | H |
| ATOM | 3220 | CA | PRO | A | 227 | 20.648 | 17.902 | 40.257 | 1.00 | 0.00 | C |
| ATOM | 3221 | HA | PRO | A | 227 | 19.745 | 18.328 | 40.198 | 1.00 | 0.00 | H |
| ATOM | 3222 | C | PRO | A | 227 | 21.102 | 18.167 | 41.704 | 1.00 | 0.00 | C |
| ATOM | 3223 | O | PRO | A | 227 | 22.304 | 18.247 | 42.046 | 1.00 | 0.00 | O |
| ATOM | 3224 | N | LEU | A | 228 | 20.227 | 18.529 | 42.606 | 1.00 | 0.00 | N |
| ATOM | 3225 | H | LEU | A | 228 | 19.268 | 18.507 | 42.322 | 1.00 | 0.00 | H |
| ATOM | 3226 | CA | LEU | A | 228 | 20.487 | 18.974 | 44.017 | 1.00 | 0.00 | C |
| ATOM | 3227 | HA | LEU | A | 228 | 21.220 | 19.651 | 43.953 | 1.00 | 0.00 | H |
| ATOM | 3228 | CB | LEU | A | 228 | 19.304 | 19.749 | 44.640 | 1.00 | 0.00 | C |
| ATOM | 3229 | HB1 | LEU | A | 228 | 18.516 | 19.136 | 44.694 | 1.00 | 0.00 | H |
| ATOM | 3230 | HB2 | LEU | A | 228 | 19.084 | 20.524 | 44.047 | 1.00 | 0.00 | H |
| ATOM | 3231 | CG | LEU | A | 228 | 19.599 | 20.287 | 46.053 | 1.00 | 0.00 | C |
| ATOM | 3232 | HG | LEU | A | 228 | 19.725 | 19.460 | 46.601 | 1.00 | 0.00 | H |
| ATOM | 3233 | CD1 | LEU | A | 228 | 20.875 | 21.117 | 46.183 | 1.00 | 0.00 | C |
| ATOM | 3234 | HD11 | LEU | A | 228 | 20.981 | 21.420 | 47.130 | 1.00 | 0.00 | H |
| ATOM | 3235 | HD12 | LEU | A | 228 | 21.663 | 20.560 | 45.921 | 1.00 | 0.00 | H |
| ATOM | 3236 | HD13 | LEU | A | 228 | 20.817 | 21.914 | 45.582 | 1.00 | 0.00 | H |
| ATOM | 3237 | CD2 | LEU | A | 228 | 18.434 | 21.205 | 46.535 | 1.00 | 0.00 | C |

```
ATOM   3238 HD21 LEU A 228      18.637  21.547  47.452  1.00  0.00           H
ATOM   3239 HD22 LEU A 228      18.335  21.976  45.906  1.00  0.00           H
ATOM   3240 HD23 LEU A 228      17.583  20.680  46.556  1.00  0.00           H
ATOM   3241  C   LEU A 228      20.971  17.759  44.889  1.00  0.00           C
ATOM   3242  O   LEU A 228      21.797  17.945  45.782  1.00  0.00           O
ATOM   3243  N   TYR A 229      20.527  16.549  44.485  1.00  0.00           N
ATOM   3244  H   TYR A 229      20.014  16.506  43.628  1.00  0.00           H
ATOM   3245  CA  TYR A 229      20.743  15.285  45.216  1.00  0.00           C
ATOM   3246  HA  TYR A 229      21.594  15.398  45.730  1.00  0.00           H
ATOM   3247  CB  TYR A 229      19.630  15.069  46.236  1.00  0.00           C
ATOM   3248  HB1 TYR A 229      19.746  14.158  46.631  1.00  0.00           H
ATOM   3249  HB2 TYR A 229      18.756  15.113  45.752  1.00  0.00           H
ATOM   3250  CG  TYR A 229      19.538  16.021  47.375  1.00  0.00           C
ATOM   3251  CD1 TYR A 229      20.427  16.000  48.481  1.00  0.00           C
ATOM   3252  HD1 TYR A 229      21.222  15.393  48.471  1.00  0.00           H
ATOM   3253  CE1 TYR A 229      20.199  16.808  49.559  1.00  0.00           C
ATOM   3254  HE1 TYR A 229      20.844  16.814  50.323  1.00  0.00           H
ATOM   3255  CZ  TYR A 229      19.043  17.644  49.597  1.00  0.00           C
ATOM   3256  OH  TYR A 229      18.785  18.490  50.662  1.00  0.00           O
ATOM   3257  HH  TYR A 229      17.932  18.985  50.495  1.00  0.00           H
ATOM   3258  CE2 TYR A 229      18.232  17.772  48.451  1.00  0.00           C
ATOM   3259  HE2 TYR A 229      17.487  18.439  48.418  1.00  0.00           H
ATOM   3260  CD2 TYR A 229      18.501  16.939  47.363  1.00  0.00           C
ATOM   3261  HD2 TYR A 229      17.926  17.008  46.548  1.00  0.00           H
ATOM   3262  C   TYR A 229      20.865  14.077  44.198  1.00  0.00           C
ATOM   3263  O   TYR A 229      20.530  14.177  43.058  1.00  0.00           O
ATOM   3264  N   LYS A 230      21.235  12.939  44.653  1.00  0.00           N
ATOM   3265  H   LYS A 230      21.402  12.899  45.638  1.00  0.00           H
ATOM   3266  CA  LYS A 230      21.449  11.669  43.910  1.00  0.00           C
ATOM   3267  HA  LYS A 230      21.961  11.917  43.088  1.00  0.00           H
ATOM   3268  CB  LYS A 230      22.236  10.656  44.777  1.00  0.00           C
ATOM   3269  HB1 LYS A 230      23.036  11.129  45.146  1.00  0.00           H
ATOM   3270  HB2 LYS A 230      22.535   9.912  44.179  1.00  0.00           H
ATOM   3271  CG  LYS A 230      21.445  10.032  45.972  1.00  0.00           C
ATOM   3272  HG1 LYS A 230      20.778   9.366  45.636  1.00  0.00           H
ATOM   3273  HG2 LYS A 230      20.972  10.748  46.486  1.00  0.00           H
ATOM   3274  CD  LYS A 230      22.494   9.327  46.880  1.00  0.00           C
ATOM   3275  HD1 LYS A 230      23.123  10.000  47.268  1.00  0.00           H
ATOM   3276  HD2 LYS A 230      23.011   8.654  46.352  1.00  0.00           H
ATOM   3277  CE  LYS A 230      21.751   8.622  48.007  1.00  0.00           C
ATOM   3278  HE1 LYS A 230      21.004   8.089  47.610  1.00  0.00           H
ATOM   3279  HE2 LYS A 230      21.376   9.315  48.622  1.00  0.00           H
ATOM   3280  NZ  LYS A 230      22.576   7.693  48.829  1.00  0.00           N
ATOM   3281  HZ1 LYS A 230      22.006   7.280  49.539  1.00  0.00           H
ATOM   3282  HZ2 LYS A 230      22.952   6.975  48.244  1.00  0.00           H
ATOM   3283  HZ3 LYS A 230      23.324   8.201  49.256  1.00  0.00           H
ATOM   3284  C   LYS A 230      20.102  11.037  43.542  1.00  0.00           C
ATOM   3285  O   LYS A 230      19.149  11.183  44.273  1.00  0.00           O
ATOM   3286  N   ARG A 231      20.062  10.254  42.510  1.00  0.00           N
ATOM   3287  H   ARG A 231      20.875  10.242  41.928  1.00  0.00           H
ATOM   3288  CA  ARG A 231      18.899   9.341  42.084  1.00  0.00           C
ATOM   3289  HA  ARG A 231      18.060   9.781  42.404  1.00  0.00           H
ATOM   3290  CB  ARG A 231      18.888   9.208  40.557  1.00  0.00           C
ATOM   3291  HB1 ARG A 231      19.373   8.367  40.318  1.00  0.00           H
```

| | | | | | | | | | | |
|---|---|---|---|---|---|---|---|---|---|---|
| ATOM | 3292 | HB2 | ARG | A | 231 | 19.376 | 9.994 | 40.177 | 1.00 | 0.00 | H |
| ATOM | 3293 | CG | ARG | A | 231 | 17.460 | 9.156 | 39.931 | 1.00 | 0.00 | C |
| ATOM | 3294 | HG1 | ARG | A | 231 | 16.881 | 9.877 | 40.311 | 1.00 | 0.00 | H |
| ATOM | 3295 | HG2 | ARG | A | 231 | 17.034 | 8.266 | 40.094 | 1.00 | 0.00 | H |
| ATOM | 3296 | CD | ARG | A | 231 | 17.643 | 9.375 | 38.434 | 1.00 | 0.00 | C |
| ATOM | 3297 | HD1 | ARG | A | 231 | 18.158 | 10.216 | 38.270 | 1.00 | 0.00 | H |
| ATOM | 3298 | HD2 | ARG | A | 231 | 16.752 | 9.437 | 37.984 | 1.00 | 0.00 | H |
| ATOM | 3299 | NE | ARG | A | 231 | 18.337 | 8.365 | 37.774 | 1.00 | 0.00 | N |
| ATOM | 3300 | HE | ARG | A | 231 | 19.327 | 8.487 | 37.699 | 1.00 | 0.00 | H |
| ATOM | 3301 | CZ | ARG | A | 231 | 17.844 | 7.263 | 37.234 | 1.00 | 0.00 | C |
| ATOM | 3302 | NH1 | ARG | A | 231 | 16.611 | 7.013 | 37.263 | 1.00 | 0.00 | N |
| ATOM | 3303 | HH11 | ARG | A | 231 | 15.981 | 7.654 | 37.701 | 1.00 | 0.00 | H |
| ATOM | 3304 | HH12 | ARG | A | 231 | 16.261 | 6.173 | 36.848 | 1.00 | 0.00 | H |
| ATOM | 3305 | NH2 | ARG | A | 231 | 18.616 | 6.321 | 36.857 | 1.00 | 0.00 | N |
| ATOM | 3306 | HH21 | ARG | A | 231 | 19.606 | 6.411 | 36.968 | 1.00 | 0.00 | H |
| ATOM | 3307 | HH22 | ARG | A | 231 | 18.232 | 5.492 | 36.450 | 1.00 | 0.00 | H |
| ATOM | 3308 | C | ARG | A | 231 | 18.792 | 8.013 | 42.786 | 1.00 | 0.00 | C |
| ATOM | 3309 | O | ARG | A | 231 | 19.845 | 7.557 | 43.287 | 1.00 | 0.00 | O |
| ATOM | 3310 | N | GLN | A | 232 | 17.635 | 7.374 | 43.014 | 1.00 | 0.00 | N |
| ATOM | 3311 | H | GLN | A | 232 | 16.809 | 7.692 | 42.548 | 1.00 | 0.00 | H |
| ATOM | 3312 | CA | GLN | A | 232 | 17.536 | 6.215 | 43.930 | 1.00 | 0.00 | C |
| ATOM | 3313 | HA | GLN | A | 232 | 17.871 | 6.630 | 44.776 | 1.00 | 0.00 | H |
| ATOM | 3314 | CB | GLN | A | 232 | 16.072 | 5.784 | 44.077 | 1.00 | 0.00 | C |
| ATOM | 3315 | HB1 | GLN | A | 232 | 15.758 | 5.406 | 43.206 | 1.00 | 0.00 | H |
| ATOM | 3316 | HB2 | GLN | A | 232 | 15.521 | 6.585 | 44.310 | 1.00 | 0.00 | H |
| ATOM | 3317 | CG | GLN | A | 232 | 15.875 | 4.733 | 45.158 | 1.00 | 0.00 | C |
| ATOM | 3318 | HG1 | GLN | A | 232 | 14.948 | 4.831 | 45.521 | 1.00 | 0.00 | H |
| ATOM | 3319 | HG2 | GLN | A | 232 | 16.538 | 4.909 | 45.886 | 1.00 | 0.00 | H |
| ATOM | 3320 | CD | GLN | A | 232 | 16.068 | 3.277 | 44.627 | 1.00 | 0.00 | C |
| ATOM | 3321 | OE1 | GLN | A | 232 | 16.147 | 3.014 | 43.421 | 1.00 | 0.00 | O |
| ATOM | 3322 | NE2 | GLN | A | 232 | 15.930 | 2.303 | 45.538 | 1.00 | 0.00 | N |
| ATOM | 3323 | HE21 | GLN | A | 232 | 15.717 | 2.533 | 46.488 | 1.00 | 0.00 | H |
| ATOM | 3324 | HE22 | GLN | A | 232 | 16.040 | 1.346 | 45.268 | 1.00 | 0.00 | H |
| ATOM | 3325 | C | GLN | A | 232 | 18.499 | 4.989 | 43.684 | 1.00 | 0.00 | C |
| ATOM | 3326 | O | GLN | A | 232 | 18.763 | 4.248 | 44.600 | 1.00 | 0.00 | O |
| ATOM | 3327 | N | GLU | A | 233 | 18.885 | 4.756 | 42.448 | 1.00 | 0.00 | N |
| ATOM | 3328 | H | GLU | A | 233 | 18.479 | 5.315 | 41.725 | 1.00 | 0.00 | H |
| ATOM | 3329 | CA | GLU | A | 233 | 19.866 | 3.738 | 42.049 | 1.00 | 0.00 | C |
| ATOM | 3330 | HA | GLU | A | 233 | 19.764 | 2.934 | 42.634 | 1.00 | 0.00 | H |
| ATOM | 3331 | CB | GLU | A | 233 | 19.570 | 3.385 | 40.534 | 1.00 | 0.00 | C |
| ATOM | 3332 | HB1 | GLU | A | 233 | 18.583 | 3.437 | 40.382 | 1.00 | 0.00 | H |
| ATOM | 3333 | HB2 | GLU | A | 233 | 19.885 | 2.453 | 40.357 | 1.00 | 0.00 | H |
| ATOM | 3334 | CG | GLU | A | 233 | 20.251 | 4.312 | 39.497 | 1.00 | 0.00 | C |
| ATOM | 3335 | HG1 | GLU | A | 233 | 21.215 | 4.413 | 39.743 | 1.00 | 0.00 | H |
| ATOM | 3336 | HG2 | GLU | A | 233 | 19.804 | 5.206 | 39.525 | 1.00 | 0.00 | H |
| ATOM | 3337 | CD | GLU | A | 233 | 20.183 | 3.797 | 38.064 | 1.00 | 0.00 | C |
| ATOM | 3338 | OE1 | GLU | A | 233 | 21.056 | 4.039 | 37.226 | 1.00 | 0.00 | O |
| ATOM | 3339 | OE2 | GLU | A | 233 | 19.168 | 3.143 | 37.813 | 1.00 | 0.00 | O |
| ATOM | 3340 | C | GLU | A | 233 | 21.289 | 4.225 | 42.327 | 1.00 | 0.00 | C |
| ATOM | 3341 | O | GLU | A | 233 | 22.188 | 3.353 | 42.382 | 1.00 | 0.00 | O |
| ATOM | 3342 | N | GLU | A | 234 | 21.597 | 5.513 | 42.289 | 1.00 | 0.00 | N |
| ATOM | 3343 | H | GLU | A | 234 | 20.901 | 6.227 | 42.370 | 1.00 | 0.00 | H |
| ATOM | 3344 | CA | GLU | A | 234 | 23.076 | 5.842 | 42.115 | 1.00 | 0.00 | C |
| ATOM | 3345 | HA | GLU | A | 234 | 23.342 | 5.338 | 41.293 | 1.00 | 0.00 | H |

| | | | | | | | | | | | |
|---|---|---|---|---|---|---|---|---|---|---|---|
| ATOM | 3346 | CB | GLU | A | 234 | 23.233 | 7.385 | 41.966 | 1.00 | 0.00 | C |
| ATOM | 3347 | HB1 | GLU | A | 234 | 24.195 | 7.635 | 41.858 | 1.00 | 0.00 | H |
| ATOM | 3348 | HB2 | GLU | A | 234 | 22.852 | 7.859 | 42.760 | 1.00 | 0.00 | H |
| ATOM | 3349 | CG | GLU | A | 234 | 22.470 | 7.782 | 40.731 | 1.00 | 0.00 | C |
| ATOM | 3350 | HG1 | GLU | A | 234 | 21.509 | 7.551 | 40.881 | 1.00 | 0.00 | H |
| ATOM | 3351 | HG2 | GLU | A | 234 | 22.827 | 7.244 | 39.968 | 1.00 | 0.00 | H |
| ATOM | 3352 | CD | GLU | A | 234 | 22.594 | 9.308 | 40.396 | 1.00 | 0.00 | C |
| ATOM | 3353 | OE1 | GLU | A | 234 | 22.935 | 9.555 | 39.217 | 1.00 | 0.00 | O |
| ATOM | 3354 | OE2 | GLU | A | 234 | 22.604 | 10.241 | 41.226 | 1.00 | 0.00 | O |
| ATOM | 3355 | C | GLU | A | 234 | 24.007 | 5.370 | 43.300 | 1.00 | 0.00 | C |
| ATOM | 3356 | O | GLU | A | 234 | 23.625 | 5.681 | 44.456 | 1.00 | 0.00 | O |
| ATOM | 3357 | N | SER | A | 235 | 25.170 | 4.778 | 42.854 | 1.00 | 0.00 | N |
| ATOM | 3358 | H | SER | A | 235 | 25.377 | 4.831 | 41.877 | 1.00 | 0.00 | H |
| ATOM | 3359 | CA | SER | A | 235 | 26.135 | 4.067 | 43.730 | 1.00 | 0.00 | C |
| ATOM | 3360 | HA | SER | A | 235 | 26.062 | 4.422 | 44.662 | 1.00 | 0.00 | H |
| ATOM | 3361 | CB | SER | A | 235 | 25.750 | 2.553 | 43.778 | 1.00 | 0.00 | C |
| ATOM | 3362 | HB1 | SER | A | 235 | 26.071 | 2.071 | 42.963 | 1.00 | 0.00 | H |
| ATOM | 3363 | HB2 | SER | A | 235 | 24.761 | 2.433 | 43.869 | 1.00 | 0.00 | H |
| ATOM | 3364 | OG | SER | A | 235 | 26.314 | 1.895 | 44.856 | 1.00 | 0.00 | O |
| ATOM | 3365 | HG | SER | A | 235 | 26.038 | 0.934 | 44.841 | 1.00 | 0.00 | H |
| ATOM | 3366 | C | SER | A | 235 | 27.584 | 4.387 | 43.288 | 1.00 | 0.00 | C |
| ATOM | 3367 | O | SER | A | 235 | 27.839 | 4.808 | 42.166 | 1.00 | 0.00 | O |
| ATOM | 3368 | N | GLY | A | 236 | 28.575 | 4.146 | 44.220 | 1.00 | 0.00 | N |
| ATOM | 3369 | H | GLY | A | 236 | 28.318 | 3.718 | 45.086 | 1.00 | 0.00 | H |
| ATOM | 3370 | CA | GLY | A | 236 | 29.980 | 4.485 | 44.005 | 1.00 | 0.00 | C |
| ATOM | 3371 | HA1 | GLY | A | 236 | 30.365 | 3.793 | 43.395 | 1.00 | 0.00 | H |
| ATOM | 3372 | HA2 | GLY | A | 236 | 30.438 | 4.439 | 44.893 | 1.00 | 0.00 | H |
| ATOM | 3373 | C | GLY | A | 236 | 30.220 | 5.844 | 43.407 | 1.00 | 0.00 | C |
| ATOM | 3374 | O | GLY | A | 236 | 29.901 | 6.900 | 43.989 | 1.00 | 0.00 | O |
| ATOM | 3375 | N | ALA | A | 237 | 30.937 | 5.917 | 42.293 | 1.00 | 0.00 | N |
| ATOM | 3376 | H | ALA | A | 237 | 31.262 | 5.064 | 41.884 | 1.00 | 0.00 | H |
| ATOM | 3377 | CA | ALA | A | 237 | 31.279 | 7.174 | 41.630 | 1.00 | 0.00 | C |
| ATOM | 3378 | HA | ALA | A | 237 | 31.684 | 7.755 | 42.336 | 1.00 | 0.00 | H |
| ATOM | 3379 | CB | ALA | A | 237 | 32.288 | 6.875 | 40.515 | 1.00 | 0.00 | C |
| ATOM | 3380 | HB1 | ALA | A | 237 | 32.533 | 7.726 | 40.050 | 1.00 | 0.00 | H |
| ATOM | 3381 | HB2 | ALA | A | 237 | 33.109 | 6.463 | 40.910 | 1.00 | 0.00 | H |
| ATOM | 3382 | HB3 | ALA | A | 237 | 31.880 | 6.242 | 39.857 | 1.00 | 0.00 | H |
| ATOM | 3383 | C | ALA | A | 237 | 30.041 | 7.912 | 41.087 | 1.00 | 0.00 | C |
| ATOM | 3384 | O | ALA | A | 237 | 30.039 | 9.145 | 41.063 | 1.00 | 0.00 | O |
| ATOM | 3385 | N | ALA | A | 238 | 28.966 | 7.189 | 40.737 | 1.00 | 0.00 | N |
| ATOM | 3386 | H | ALA | A | 238 | 29.000 | 6.196 | 40.853 | 1.00 | 0.00 | H |
| ATOM | 3387 | CA | ALA | A | 238 | 27.747 | 7.786 | 40.194 | 1.00 | 0.00 | C |
| ATOM | 3388 | HA | ALA | A | 238 | 28.038 | 8.312 | 39.395 | 1.00 | 0.00 | H |
| ATOM | 3389 | CB | ALA | A | 238 | 26.769 | 6.685 | 39.761 | 1.00 | 0.00 | C |
| ATOM | 3390 | HB1 | ALA | A | 238 | 25.939 | 7.103 | 39.391 | 1.00 | 0.00 | H |
| ATOM | 3391 | HB2 | ALA | A | 238 | 27.197 | 6.116 | 39.059 | 1.00 | 0.00 | H |
| ATOM | 3392 | HB3 | ALA | A | 238 | 26.534 | 6.119 | 40.551 | 1.00 | 0.00 | H |
| ATOM | 3393 | C | ALA | A | 238 | 27.101 | 8.752 | 41.195 | 1.00 | 0.00 | C |
| ATOM | 3394 | O | ALA | A | 238 | 26.808 | 9.890 | 40.828 | 1.00 | 0.00 | O |
| ATOM | 3395 | N | ALA | A | 239 | 27.031 | 8.388 | 42.489 | 1.00 | 0.00 | N |
| ATOM | 3396 | H | ALA | A | 239 | 27.410 | 7.511 | 42.784 | 1.00 | 0.00 | H |
| ATOM | 3397 | CA | ALA | A | 239 | 26.400 | 9.278 | 43.464 | 1.00 | 0.00 | C |
| ATOM | 3398 | HA | ALA | A | 239 | 25.590 | 9.540 | 42.939 | 1.00 | 0.00 | H |
| ATOM | 3399 | CB | ALA | A | 239 | 26.003 | 8.579 | 44.790 | 1.00 | 0.00 | C |

| | | | | | | | | | | | |
|---|---|---|---|---|---|---|---|---|---|---|---|
| ATOM | 3400 | HB1 | ALA | A | 239 | 25.580 | 9.245 | 45.404 | 1.00 | 0.00 | H |
| ATOM | 3401 | HB2 | ALA | A | 239 | 25.353 | 7.843 | 44.598 | 1.00 | 0.00 | H |
| ATOM | 3402 | HB3 | ALA | A | 239 | 26.820 | 8.198 | 45.223 | 1.00 | 0.00 | H |
| ATOM | 3403 | C | ALA | A | 239 | 27.228 | 10.524 | 43.802 | 1.00 | 0.00 | C |
| ATOM | 3404 | O | ALA | A | 239 | 26.790 | 11.317 | 44.631 | 1.00 | 0.00 | O |
| ATOM | 3405 | N | SER | A | 240 | 28.418 | 10.733 | 43.220 | 1.00 | 0.00 | N |
| ATOM | 3406 | H | SER | A | 240 | 28.796 | 10.022 | 42.626 | 1.00 | 0.00 | H |
| ATOM | 3407 | CA | SER | A | 240 | 29.190 | 11.973 | 43.424 | 1.00 | 0.00 | C |
| ATOM | 3408 | HA | SER | A | 240 | 29.035 | 12.186 | 44.389 | 1.00 | 0.00 | H |
| ATOM | 3409 | CB | SER | A | 240 | 30.684 | 11.739 | 43.169 | 1.00 | 0.00 | C |
| ATOM | 3410 | HB1 | SER | A | 240 | 30.989 | 10.908 | 43.635 | 1.00 | 0.00 | H |
| ATOM | 3411 | HB2 | SER | A | 240 | 31.217 | 12.519 | 43.496 | 1.00 | 0.00 | H |
| ATOM | 3412 | OG | SER | A | 240 | 30.997 | 11.576 | 41.797 | 1.00 | 0.00 | O |
| ATOM | 3413 | HG | SER | A | 240 | 31.981 | 11.428 | 41.695 | 1.00 | 0.00 | H |
| ATOM | 3414 | C | SER | A | 240 | 28.675 | 13.163 | 42.598 | 1.00 | 0.00 | C |
| ATOM | 3415 | O | SER | A | 240 | 29.088 | 14.301 | 42.817 | 1.00 | 0.00 | O |
| ATOM | 3416 | N | LEU | A | 241 | 27.789 | 12.909 | 41.630 | 1.00 | 0.00 | N |
| ATOM | 3417 | H | LEU | A | 241 | 27.296 | 12.040 | 41.666 | 1.00 | 0.00 | H |
| ATOM | 3418 | CA | LEU | A | 241 | 27.491 | 13.817 | 40.520 | 1.00 | 0.00 | C |
| ATOM | 3419 | HA | LEU | A | 241 | 28.235 | 14.484 | 40.487 | 1.00 | 0.00 | H |
| ATOM | 3420 | CB | LEU | A | 241 | 27.461 | 12.995 | 39.226 | 1.00 | 0.00 | C |
| ATOM | 3421 | HB1 | LEU | A | 241 | 27.187 | 13.593 | 38.473 | 1.00 | 0.00 | H |
| ATOM | 3422 | HB2 | LEU | A | 241 | 26.790 | 12.261 | 39.329 | 1.00 | 0.00 | H |
| ATOM | 3423 | CG | LEU | A | 241 | 28.829 | 12.374 | 38.894 | 1.00 | 0.00 | C |
| ATOM | 3424 | HG | LEU | A | 241 | 29.249 | 12.010 | 39.726 | 1.00 | 0.00 | H |
| ATOM | 3425 | CD1 | LEU | A | 241 | 28.599 | 11.259 | 37.891 | 1.00 | 0.00 | C |
| ATOM | 3426 | HD11 | LEU | A | 241 | 29.474 | 10.836 | 37.656 | 1.00 | 0.00 | H |
| ATOM | 3427 | HD12 | LEU | A | 241 | 27.993 | 10.571 | 38.290 | 1.00 | 0.00 | H |
| ATOM | 3428 | HD13 | LEU | A | 241 | 28.178 | 11.635 | 37.065 | 1.00 | 0.00 | H |
| ATOM | 3429 | CD2 | LEU | A | 241 | 29.798 | 13.413 | 38.319 | 1.00 | 0.00 | C |
| ATOM | 3430 | HD21 | LEU | A | 241 | 30.674 | 12.975 | 38.116 | 1.00 | 0.00 | H |
| ATOM | 3431 | HD22 | LEU | A | 241 | 29.415 | 13.798 | 37.479 | 1.00 | 0.00 | H |
| ATOM | 3432 | HD23 | LEU | A | 241 | 29.938 | 14.144 | 38.987 | 1.00 | 0.00 | H |
| ATOM | 3433 | C | LEU | A | 241 | 26.226 | 14.653 | 40.748 | 1.00 | 0.00 | C |
| ATOM | 3434 | O | LEU | A | 241 | 25.292 | 14.642 | 39.945 | 1.00 | 0.00 | O |
| ATOM | 3435 | N | ASN | A | 242 | 26.179 | 15.330 | 41.893 | 1.00 | 0.00 | N |
| ATOM | 3436 | H | ASN | A | 242 | 26.958 | 15.251 | 42.515 | 1.00 | 0.00 | H |
| ATOM | 3437 | CA | ASN | A | 242 | 25.065 | 16.180 | 42.301 | 1.00 | 0.00 | C |
| ATOM | 3438 | HA | ASN | A | 242 | 24.754 | 16.663 | 41.482 | 1.00 | 0.00 | H |
| ATOM | 3439 | CB | ASN | A | 242 | 23.964 | 15.286 | 42.851 | 1.00 | 0.00 | C |
| ATOM | 3440 | HB1 | ASN | A | 242 | 23.742 | 14.607 | 42.151 | 1.00 | 0.00 | H |
| ATOM | 3441 | HB2 | ASN | A | 242 | 23.163 | 15.859 | 43.026 | 1.00 | 0.00 | H |
| ATOM | 3442 | CG | ASN | A | 242 | 24.292 | 14.540 | 44.134 | 1.00 | 0.00 | C |
| ATOM | 3443 | OD1 | ASN | A | 242 | 24.178 | 15.055 | 45.262 | 1.00 | 0.00 | O |
| ATOM | 3444 | ND2 | ASN | A | 242 | 24.672 | 13.310 | 43.973 | 1.00 | 0.00 | N |
| ATOM | 3445 | HD21 | ASN | A | 242 | 24.731 | 12.921 | 43.054 | 1.00 | 0.00 | H |
| ATOM | 3446 | HD22 | ASN | A | 242 | 24.906 | 12.751 | 44.768 | 1.00 | 0.00 | H |
| ATOM | 3447 | C | ASN | A | 242 | 25.492 | 17.241 | 43.317 | 1.00 | 0.00 | C |
| ATOM | 3448 | O | ASN | A | 242 | 26.561 | 17.101 | 43.916 | 1.00 | 0.00 | O |
| ATOM | 3449 | N | VAL | A | 243 | 24.764 | 18.361 | 43.394 | 1.00 | 0.00 | N |
| ATOM | 3450 | H | VAL | A | 243 | 23.890 | 18.416 | 42.912 | 1.00 | 0.00 | H |
| ATOM | 3451 | CA | VAL | A | 243 | 25.244 | 19.534 | 44.195 | 1.00 | 0.00 | C |
| ATOM | 3452 | HA | VAL | A | 243 | 26.083 | 19.859 | 43.759 | 1.00 | 0.00 | H |
| ATOM | 3453 | CB | VAL | A | 243 | 24.252 | 20.719 | 44.118 | 1.00 | 0.00 | C |

| | | | | | | | | | | | |
|---|---|---|---|---|---|---|---|---|---|---|---|
| ATOM | 3454 | HB | VAL | A | 243 | 23.407 | 20.226 | 44.328 | 1.00 | 0.00 | H |
| ATOM | 3455 | CG1 | VAL | A | 243 | 24.578 | 21.808 | 45.097 | 1.00 | 0.00 | C |
| ATOM | 3456 | HG11 | VAL | A | 243 | 23.911 | 22.547 | 45.008 | 1.00 | 0.00 | H |
| ATOM | 3457 | HG12 | VAL | A | 243 | 24.546 | 21.441 | 46.027 | 1.00 | 0.00 | H |
| ATOM | 3458 | HG13 | VAL | A | 243 | 25.494 | 22.162 | 44.909 | 1.00 | 0.00 | H |
| ATOM | 3459 | CG2 | VAL | A | 243 | 24.064 | 21.474 | 42.781 | 1.00 | 0.00 | C |
| ATOM | 3460 | HG21 | VAL | A | 243 | 23.393 | 22.206 | 42.903 | 1.00 | 0.00 | H |
| ATOM | 3461 | HG22 | VAL | A | 243 | 24.938 | 21.867 | 42.494 | 1.00 | 0.00 | H |
| ATOM | 3462 | HG23 | VAL | A | 243 | 23.740 | 20.837 | 42.081 | 1.00 | 0.00 | H |
| ATOM | 3463 | C | VAL | A | 243 | 25.555 | 19.073 | 45.676 | 1.00 | 0.00 | C |
| ATOM | 3464 | O | VAL | A | 243 | 26.592 | 19.396 | 46.232 | 1.00 | 0.00 | O |
| ATOM | 3465 | N | ASP | A | 244 | 24.614 | 18.315 | 46.312 | 1.00 | 0.00 | N |
| ATOM | 3466 | H | ASP | A | 244 | 23.756 | 18.112 | 45.840 | 1.00 | 0.00 | H |
| ATOM | 3467 | CA | ASP | A | 244 | 24.827 | 17.771 | 47.711 | 1.00 | 0.00 | C |
| ATOM | 3468 | HA | ASP | A | 244 | 24.776 | 18.608 | 48.256 | 1.00 | 0.00 | H |
| ATOM | 3469 | CB | ASP | A | 244 | 23.713 | 16.930 | 48.235 | 1.00 | 0.00 | C |
| ATOM | 3470 | HB1 | ASP | A | 244 | 23.667 | 16.112 | 47.662 | 1.00 | 0.00 | H |
| ATOM | 3471 | HB2 | ASP | A | 244 | 22.871 | 17.458 | 48.130 | 1.00 | 0.00 | H |
| ATOM | 3472 | CG | ASP | A | 244 | 23.838 | 16.493 | 49.705 | 1.00 | 0.00 | C |
| ATOM | 3473 | OD1 | ASP | A | 244 | 23.667 | 15.277 | 50.054 | 1.00 | 0.00 | O |
| ATOM | 3474 | OD2 | ASP | A | 244 | 24.226 | 17.338 | 50.545 | 1.00 | 0.00 | O |
| ATOM | 3475 | C | ASP | A | 244 | 26.215 | 17.060 | 47.929 | 1.00 | 0.00 | C |
| ATOM | 3476 | O | ASP | A | 244 | 26.978 | 17.461 | 48.799 | 1.00 | 0.00 | O |
| ATOM | 3477 | N | ALA | A | 245 | 26.446 | 16.124 | 47.071 | 1.00 | 0.00 | N |
| ATOM | 3478 | H | ALA | A | 245 | 25.736 | 15.954 | 46.388 | 1.00 | 0.00 | H |
| ATOM | 3479 | CA | ALA | A | 245 | 27.599 | 15.318 | 47.001 | 1.00 | 0.00 | C |
| ATOM | 3480 | HA | ALA | A | 245 | 27.547 | 14.937 | 47.924 | 1.00 | 0.00 | H |
| ATOM | 3481 | CB | ALA | A | 245 | 27.511 | 14.102 | 46.092 | 1.00 | 0.00 | C |
| ATOM | 3482 | HB1 | ALA | A | 245 | 28.378 | 13.604 | 46.117 | 1.00 | 0.00 | H |
| ATOM | 3483 | HB2 | ALA | A | 245 | 26.774 | 13.504 | 46.406 | 1.00 | 0.00 | H |
| ATOM | 3484 | HB3 | ALA | A | 245 | 27.324 | 14.399 | 45.156 | 1.00 | 0.00 | H |
| ATOM | 3485 | C | ALA | A | 245 | 28.906 | 16.211 | 46.671 | 1.00 | 0.00 | C |
| ATOM | 3486 | O | ALA | A | 245 | 29.930 | 15.935 | 47.259 | 1.00 | 0.00 | O |
| ATOM | 3487 | N | ALA | A | 246 | 28.844 | 17.169 | 45.777 | 1.00 | 0.00 | N |
| ATOM | 3488 | H | ALA | A | 246 | 27.982 | 17.282 | 45.283 | 1.00 | 0.00 | H |
| ATOM | 3489 | CA | ALA | A | 246 | 29.932 | 18.082 | 45.447 | 1.00 | 0.00 | C |
| ATOM | 3490 | HA | ALA | A | 246 | 30.730 | 17.496 | 45.307 | 1.00 | 0.00 | H |
| ATOM | 3491 | CB | ALA | A | 246 | 29.548 | 18.834 | 44.174 | 1.00 | 0.00 | C |
| ATOM | 3492 | HB1 | ALA | A | 246 | 30.281 | 19.468 | 43.927 | 1.00 | 0.00 | H |
| ATOM | 3493 | HB2 | ALA | A | 246 | 29.408 | 18.180 | 43.430 | 1.00 | 0.00 | H |
| ATOM | 3494 | HB3 | ALA | A | 246 | 28.703 | 19.346 | 44.331 | 1.00 | 0.00 | H |
| ATOM | 3495 | C | ALA | A | 246 | 30.296 | 19.056 | 46.585 | 1.00 | 0.00 | C |
| ATOM | 3496 | O | ALA | A | 246 | 31.483 | 19.221 | 46.794 | 1.00 | 0.00 | O |
| ATOM | 3497 | N | VAL | A | 247 | 29.311 | 19.626 | 47.239 | 1.00 | 0.00 | N |
| ATOM | 3498 | H | VAL | A | 247 | 28.397 | 19.492 | 46.855 | 1.00 | 0.00 | H |
| ATOM | 3499 | CA | VAL | A | 247 | 29.382 | 20.456 | 48.494 | 1.00 | 0.00 | C |
| ATOM | 3500 | HA | VAL | A | 247 | 29.989 | 21.246 | 48.412 | 1.00 | 0.00 | H |
| ATOM | 3501 | CB | VAL | A | 247 | 27.954 | 21.093 | 48.797 | 1.00 | 0.00 | C |
| ATOM | 3502 | HB | VAL | A | 247 | 27.235 | 20.400 | 48.839 | 1.00 | 0.00 | H |
| ATOM | 3503 | CG1 | VAL | A | 247 | 28.031 | 21.773 | 50.197 | 1.00 | 0.00 | C |
| ATOM | 3504 | HG11 | VAL | A | 247 | 27.146 | 22.184 | 50.417 | 1.00 | 0.00 | H |
| ATOM | 3505 | HG12 | VAL | A | 247 | 28.261 | 21.088 | 50.888 | 1.00 | 0.00 | H |
| ATOM | 3506 | HG13 | VAL | A | 247 | 28.735 | 22.483 | 50.184 | 1.00 | 0.00 | H |
| ATOM | 3507 | CG2 | VAL | A | 247 | 27.490 | 22.053 | 47.639 | 1.00 | 0.00 | C |

```
ATOM   3508 HG21 VAL A 247      26.593  22.434  47.861  1.00  0.00           H
ATOM   3509 HG22 VAL A 247      28.152  22.796  47.537  1.00  0.00           H
ATOM   3510 HG23 VAL A 247      27.431  21.539  46.783  1.00  0.00           H
ATOM   3511  C   VAL A 247      30.015  19.522  49.563  1.00  0.00           C
ATOM   3512  O   VAL A 247      31.023  19.860  50.171  1.00  0.00           O
ATOM   3513  N   GLN A 248      29.522  18.320  49.769  1.00  0.00           N
ATOM   3514  H   GLN A 248      28.723  18.043  49.235  1.00  0.00           H
ATOM   3515  CA  GLN A 248      30.104  17.304  50.790  1.00  0.00           C
ATOM   3516  HA  GLN A 248      30.015  17.627  51.732  1.00  0.00           H
ATOM   3517  CB  GLN A 248      29.384  15.962  50.590  1.00  0.00           C
ATOM   3518  HB1 GLN A 248      29.660  15.564  49.715  1.00  0.00           H
ATOM   3519  HB2 GLN A 248      28.395  16.112  50.592  1.00  0.00           H
ATOM   3520  CG  GLN A 248      29.780  15.008  51.757  1.00  0.00           C
ATOM   3521  HG1 GLN A 248      29.408  15.351  52.620  1.00  0.00           H
ATOM   3522  HG2 GLN A 248      30.776  14.945  51.824  1.00  0.00           H
ATOM   3523  CD  GLN A 248      29.159  13.598  51.408  1.00  0.00           C
ATOM   3524  OE1 GLN A 248      27.888  13.522  51.151  1.00  0.00           O
ATOM   3525  NE2 GLN A 248      29.859  12.576  51.373  1.00  0.00           N
ATOM   3526 HE21 GLN A 248      30.838  12.634  51.568  1.00  0.00           H
ATOM   3527 HE22 GLN A 248      29.445  11.694  51.150  1.00  0.00           H
ATOM   3528  C   GLN A 248      31.547  17.117  50.515  1.00  0.00           C
ATOM   3529  O   GLN A 248      32.381  17.084  51.398  1.00  0.00           O
ATOM   3530  N   GLN A 249      31.877  16.948  49.234  1.00  0.00           N
ATOM   3531  H   GLN A 249      31.154  17.001  48.545  1.00  0.00           H
ATOM   3532  CA  GLN A 249      33.283  16.682  48.776  1.00  0.00           C
ATOM   3533  HA  GLN A 249      33.620  15.909  49.314  1.00  0.00           H
ATOM   3534  CB  GLN A 249      33.312  16.341  47.259  1.00  0.00           C
ATOM   3535  HB1 GLN A 249      32.952  17.133  46.765  1.00  0.00           H
ATOM   3536  HB2 GLN A 249      32.710  15.556  47.115  1.00  0.00           H
ATOM   3537  CG  GLN A 249      34.676  15.991  46.626  1.00  0.00           C
ATOM   3538  HG1 GLN A 249      35.267  16.792  46.723  1.00  0.00           H
ATOM   3539  HG2 GLN A 249      34.519  15.808  45.655  1.00  0.00           H
ATOM   3540  CD  GLN A 249      35.434  14.779  47.212  1.00  0.00           C
ATOM   3541  OE1 GLN A 249      36.683  14.761  47.069  1.00  0.00           O
ATOM   3542  NE2 GLN A 249      34.907  13.962  47.951  1.00  0.00           N
ATOM   3543 HE21 GLN A 249      33.944  14.062  48.201  1.00  0.00           H
ATOM   3544 HE22 GLN A 249      35.439  13.193  48.306  1.00  0.00           H
ATOM   3545  C   GLN A 249      34.194  17.825  49.092  1.00  0.00           C
ATOM   3546  O   GLN A 249      35.263  17.709  49.660  1.00  0.00           O
ATOM   3547  N   TRP A 250      33.841  18.970  48.623  1.00  0.00           N
ATOM   3548  H   TRP A 250      32.993  19.016  48.095  1.00  0.00           H
ATOM   3549  CA  TRP A 250      34.620  20.251  48.816  1.00  0.00           C
ATOM   3550  HA  TRP A 250      35.554  19.929  48.660  1.00  0.00           H
ATOM   3551  CB  TRP A 250      34.220  21.379  47.785  1.00  0.00           C
ATOM   3552  HB1 TRP A 250      34.460  22.279  48.149  1.00  0.00           H
ATOM   3553  HB2 TRP A 250      33.238  21.345  47.600  1.00  0.00           H
ATOM   3554  CG  TRP A 250      34.898  21.250  46.494  1.00  0.00           C
ATOM   3555  CD1 TRP A 250      34.330  20.647  45.452  1.00  0.00           C
ATOM   3556  HD1 TRP A 250      33.456  20.162  45.476  1.00  0.00           H
ATOM   3557  NE1 TRP A 250      35.108  20.786  44.357  1.00  0.00           N
ATOM   3558  HE1 TRP A 250      34.832  20.510  43.436  1.00  0.00           H
ATOM   3559  CE2 TRP A 250      36.374  21.374  44.669  1.00  0.00           C
ATOM   3560  CZ2 TRP A 250      37.531  21.641  43.894  1.00  0.00           C
ATOM   3561  HZ2 TRP A 250      37.566  21.435  42.916  1.00  0.00           H
```

| | | | | | | | | | | | |
|---|---|---|---|---|---|---|---|---|---|---|---|
| ATOM | 3562 | CH2 | TRP | A | 250 | 38.603 | 22.198 | 44.554 | 1.00 | 0.00 | C |
| ATOM | 3563 | HH2 | TRP | A | 250 | 39.457 | 22.352 | 44.056 | 1.00 | 0.00 | H |
| ATOM | 3564 | CZ3 | TRP | A | 250 | 38.525 | 22.557 | 45.919 | 1.00 | 0.00 | C |
| ATOM | 3565 | HZ3 | TRP | A | 250 | 39.286 | 23.033 | 46.359 | 1.00 | 0.00 | H |
| ATOM | 3566 | CE3 | TRP | A | 250 | 37.355 | 22.236 | 46.661 | 1.00 | 0.00 | C |
| ATOM | 3567 | HE3 | TRP | A | 250 | 37.310 | 22.480 | 47.630 | 1.00 | 0.00 | H |
| ATOM | 3568 | CD2 | TRP | A | 250 | 36.280 | 21.596 | 46.076 | 1.00 | 0.00 | C |
| ATOM | 3569 | C | TRP | A | 250 | 34.672 | 20.793 | 50.228 | 1.00 | 0.00 | C |
| ATOM | 3570 | O | TRP | A | 250 | 35.590 | 21.456 | 50.664 | 1.00 | 0.00 | O |
| ATOM | 3571 | N | LEU | A | 251 | 33.772 | 20.390 | 51.092 | 1.00 | 0.00 | N |
| ATOM | 3572 | H | LEU | A | 251 | 32.988 | 19.868 | 50.756 | 1.00 | 0.00 | H |
| ATOM | 3573 | CA | LEU | A | 251 | 33.864 | 20.672 | 52.540 | 1.00 | 0.00 | C |
| ATOM | 3574 | HA | LEU | A | 251 | 34.460 | 21.471 | 52.626 | 1.00 | 0.00 | H |
| ATOM | 3575 | CB | LEU | A | 251 | 32.506 | 21.044 | 53.099 | 1.00 | 0.00 | C |
| ATOM | 3576 | HB1 | LEU | A | 251 | 32.631 | 21.121 | 54.088 | 1.00 | 0.00 | H |
| ATOM | 3577 | HB2 | LEU | A | 251 | 31.908 | 20.268 | 52.898 | 1.00 | 0.00 | H |
| ATOM | 3578 | CG | LEU | A | 251 | 31.770 | 22.336 | 52.621 | 1.00 | 0.00 | C |
| ATOM | 3579 | HG | LEU | A | 251 | 31.639 | 22.201 | 51.639 | 1.00 | 0.00 | H |
| ATOM | 3580 | CD1 | LEU | A | 251 | 30.484 | 22.530 | 53.440 | 1.00 | 0.00 | C |
| ATOM | 3581 | HD11 | LEU | A | 251 | 30.015 | 23.358 | 53.132 | 1.00 | 0.00 | H |
| ATOM | 3582 | HD12 | LEU | A | 251 | 29.885 | 21.740 | 53.312 | 1.00 | 0.00 | H |
| ATOM | 3583 | HD13 | LEU | A | 251 | 30.716 | 22.619 | 54.409 | 1.00 | 0.00 | H |
| ATOM | 3584 | CD2 | LEU | A | 251 | 32.511 | 23.631 | 52.755 | 1.00 | 0.00 | C |
| ATOM | 3585 | HD21 | LEU | A | 251 | 31.939 | 24.379 | 52.418 | 1.00 | 0.00 | H |
| ATOM | 3586 | HD22 | LEU | A | 251 | 32.734 | 23.789 | 53.717 | 1.00 | 0.00 | H |
| ATOM | 3587 | HD23 | LEU | A | 251 | 33.355 | 23.589 | 52.220 | 1.00 | 0.00 | H |
| ATOM | 3588 | C | LEU | A | 251 | 34.597 | 19.621 | 53.325 | 1.00 | 0.00 | C |
| ATOM | 3589 | O | LEU | A | 251 | 35.383 | 19.984 | 54.198 | 1.00 | 0.00 | O |
| ATOM | 3590 | N | GLN | A | 252 | 34.533 | 18.380 | 52.971 | 1.00 | 0.00 | N |
| ATOM | 3591 | H | GLN | A | 252 | 33.872 | 18.113 | 52.270 | 1.00 | 0.00 | H |
| ATOM | 3592 | CA | GLN | A | 252 | 35.370 | 17.377 | 53.543 | 1.00 | 0.00 | C |
| ATOM | 3593 | HA | GLN | A | 252 | 35.394 | 17.664 | 54.501 | 1.00 | 0.00 | H |
| ATOM | 3594 | CB | GLN | A | 252 | 34.730 | 15.960 | 53.375 | 1.00 | 0.00 | C |
| ATOM | 3595 | HB1 | GLN | A | 252 | 35.390 | 15.278 | 53.689 | 1.00 | 0.00 | H |
| ATOM | 3596 | HB2 | GLN | A | 252 | 34.537 | 15.815 | 52.404 | 1.00 | 0.00 | H |
| ATOM | 3597 | CG | GLN | A | 252 | 33.404 | 15.757 | 54.170 | 1.00 | 0.00 | C |
| ATOM | 3598 | HG1 | GLN | A | 252 | 32.933 | 14.954 | 53.805 | 1.00 | 0.00 | H |
| ATOM | 3599 | HG2 | GLN | A | 252 | 32.830 | 16.566 | 54.044 | 1.00 | 0.00 | H |
| ATOM | 3600 | CD | GLN | A | 252 | 33.648 | 15.545 | 55.704 | 1.00 | 0.00 | C |
| ATOM | 3601 | OE1 | GLN | A | 252 | 34.685 | 15.822 | 56.308 | 1.00 | 0.00 | O |
| ATOM | 3602 | NE2 | GLN | A | 252 | 32.657 | 15.080 | 56.423 | 1.00 | 0.00 | N |
| ATOM | 3603 | HE21 | GLN | A | 252 | 31.780 | 14.872 | 55.990 | 1.00 | 0.00 | H |
| ATOM | 3604 | HE22 | GLN | A | 252 | 32.778 | 14.933 | 57.405 | 1.00 | 0.00 | H |
| ATOM | 3605 | C | GLN | A | 252 | 36.816 | 17.420 | 53.047 | 1.00 | 0.00 | C |
| ATOM | 3606 | O | GLN | A | 252 | 37.747 | 16.921 | 53.734 | 1.00 | 0.00 | O |
| ATOM | 3607 | N | LYS | A | 253 | 37.069 | 18.124 | 51.970 | 1.00 | 0.00 | N |
| ATOM | 3608 | H | LYS | A | 253 | 36.297 | 18.441 | 51.420 | 1.00 | 0.00 | H |
| ATOM | 3609 | CA | LYS | A | 253 | 38.448 | 18.477 | 51.522 | 1.00 | 0.00 | C |
| ATOM | 3610 | HA | LYS | A | 253 | 39.067 | 17.757 | 51.835 | 1.00 | 0.00 | H |
| ATOM | 3611 | CB | LYS | A | 253 | 38.468 | 18.559 | 49.963 | 1.00 | 0.00 | C |
| ATOM | 3612 | HB1 | LYS | A | 253 | 37.908 | 19.338 | 49.679 | 1.00 | 0.00 | H |
| ATOM | 3613 | HB2 | LYS | A | 253 | 38.076 | 17.715 | 49.596 | 1.00 | 0.00 | H |
| ATOM | 3614 | CG | LYS | A | 253 | 39.891 | 18.737 | 49.365 | 1.00 | 0.00 | C |
| ATOM | 3615 | HG1 | LYS | A | 253 | 40.452 | 17.934 | 49.568 | 1.00 | 0.00 | H |

| | | | | | | | | | | | |
|---|---|---|---|---|---|---|---|---|---|---|---|
| ATOM | 3616 | HG2 | LYS | A | 253 | 40.325 | 19.553 | 49.748 | 1.00 | 0.00 | H |
| ATOM | 3617 | CD | LYS | A | 253 | 39.797 | 18.895 | 47.873 | 1.00 | 0.00 | C |
| ATOM | 3618 | HD1 | LYS | A | 253 | 39.237 | 19.697 | 47.668 | 1.00 | 0.00 | H |
| ATOM | 3619 | HD2 | LYS | A | 253 | 39.364 | 18.079 | 47.489 | 1.00 | 0.00 | H |
| ATOM | 3620 | CE | LYS | A | 253 | 41.142 | 19.072 | 47.235 | 1.00 | 0.00 | C |
| ATOM | 3621 | HE1 | LYS | A | 253 | 41.719 | 19.627 | 47.834 | 1.00 | 0.00 | H |
| ATOM | 3622 | HE2 | LYS | A | 253 | 41.030 | 19.537 | 46.357 | 1.00 | 0.00 | H |
| ATOM | 3623 | NZ | LYS | A | 253 | 41.838 | 17.828 | 46.986 | 1.00 | 0.00 | N |
| ATOM | 3624 | HZ1 | LYS | A | 253 | 42.723 | 18.020 | 46.562 | 1.00 | 0.00 | H |
| ATOM | 3625 | HZ2 | LYS | A | 253 | 41.981 | 17.347 | 47.851 | 1.00 | 0.00 | H |
| ATOM | 3626 | HZ3 | LYS | A | 253 | 41.291 | 17.257 | 46.373 | 1.00 | 0.00 | H |
| ATOM | 3627 | C | LYS | A | 253 | 38.899 | 19.804 | 52.174 | 1.00 | 0.00 | C |
| ATOM | 3628 | O | LYS | A | 253 | 40.042 | 20.254 | 52.060 | 1.00 | 0.00 | O |
| ATOM | 3629 | N | GLY | A | 254 | 37.949 | 20.432 | 52.936 | 1.00 | 0.00 | N |
| ATOM | 3630 | H | GLY | A | 254 | 36.981 | 20.232 | 52.786 | 1.00 | 0.00 | H |
| ATOM | 3631 | CA | GLY | A | 254 | 38.342 | 21.393 | 53.967 | 1.00 | 0.00 | C |
| ATOM | 3632 | HA1 | GLY | A | 254 | 39.314 | 21.240 | 54.144 | 1.00 | 0.00 | H |
| ATOM | 3633 | HA2 | GLY | A | 254 | 37.814 | 21.165 | 54.785 | 1.00 | 0.00 | H |
| ATOM | 3634 | C | GLY | A | 254 | 38.119 | 22.883 | 53.620 | 1.00 | 0.00 | C |
| ATOM | 3635 | O | GLY | A | 254 | 38.516 | 23.739 | 54.433 | 1.00 | 0.00 | O |
| ATOM | 3636 | N | THR | A | 255 | 37.479 | 23.168 | 52.483 | 1.00 | 0.00 | N |
| ATOM | 3637 | H | THR | A | 255 | 37.254 | 22.411 | 51.869 | 1.00 | 0.00 | H |
| ATOM | 3638 | CA | THR | A | 255 | 37.099 | 24.477 | 52.092 | 1.00 | 0.00 | C |
| ATOM | 3639 | HA | THR | A | 255 | 38.010 | 24.859 | 51.935 | 1.00 | 0.00 | H |
| ATOM | 3640 | CB | THR | A | 255 | 36.404 | 24.561 | 50.770 | 1.00 | 0.00 | C |
| ATOM | 3641 | HB | THR | A | 255 | 35.446 | 24.380 | 50.994 | 1.00 | 0.00 | H |
| ATOM | 3642 | CG2 | THR | A | 255 | 36.630 | 25.881 | 50.044 | 1.00 | 0.00 | C |
| ATOM | 3643 | HG21 | THR | A | 255 | 36.140 | 25.870 | 49.172 | 1.00 | 0.00 | H |
| ATOM | 3644 | HG22 | THR | A | 255 | 36.290 | 26.634 | 50.608 | 1.00 | 0.00 | H |
| ATOM | 3645 | HG23 | THR | A | 255 | 37.608 | 26.006 | 49.875 | 1.00 | 0.00 | H |
| ATOM | 3646 | OG1 | THR | A | 255 | 36.814 | 23.577 | 49.863 | 1.00 | 0.00 | O |
| ATOM | 3647 | HG1 | THR | A | 255 | 36.315 | 23.687 | 49.004 | 1.00 | 0.00 | H |
| ATOM | 3648 | C | THR | A | 255 | 36.298 | 25.234 | 53.216 | 1.00 | 0.00 | C |
| ATOM | 3649 | O | THR | A | 255 | 35.283 | 24.720 | 53.599 | 1.00 | 0.00 | O |
| ATOM | 3650 | N | PRO | A | 256 | 36.533 | 26.470 | 53.663 | 1.00 | 0.00 | N |
| ATOM | 3651 | CD | PRO | A | 256 | 37.827 | 27.237 | 53.544 | 1.00 | 0.00 | C |
| ATOM | 3652 | HD1 | PRO | A | 256 | 37.871 | 27.761 | 52.693 | 1.00 | 0.00 | H |
| ATOM | 3653 | HD2 | PRO | A | 256 | 38.622 | 26.634 | 53.606 | 1.00 | 0.00 | H |
| ATOM | 3654 | CG | PRO | A | 256 | 37.790 | 28.130 | 54.677 | 1.00 | 0.00 | C |
| ATOM | 3655 | HG1 | PRO | A | 256 | 38.423 | 28.891 | 54.539 | 1.00 | 0.00 | H |
| ATOM | 3656 | HG2 | PRO | A | 256 | 38.036 | 27.639 | 55.513 | 1.00 | 0.00 | H |
| ATOM | 3657 | CB | PRO | A | 256 | 36.364 | 28.595 | 54.707 | 1.00 | 0.00 | C |
| ATOM | 3658 | HB1 | PRO | A | 256 | 36.174 | 29.249 | 53.975 | 1.00 | 0.00 | H |
| ATOM | 3659 | HB2 | PRO | A | 256 | 36.128 | 29.004 | 55.588 | 1.00 | 0.00 | H |
| ATOM | 3660 | CA | PRO | A | 256 | 35.673 | 27.227 | 54.480 | 1.00 | 0.00 | C |
| ATOM | 3661 | HA | PRO | A | 256 | 35.560 | 26.622 | 55.268 | 1.00 | 0.00 | H |
| ATOM | 3662 | C | PRO | A | 256 | 34.302 | 27.578 | 53.818 | 1.00 | 0.00 | C |
| ATOM | 3663 | O | PRO | A | 256 | 34.187 | 27.916 | 52.620 | 1.00 | 0.00 | O |
| ATOM | 3664 | N | ALA | A | 257 | 33.274 | 27.456 | 54.600 | 1.00 | 0.00 | N |
| ATOM | 3665 | H | ALA | A | 257 | 33.369 | 27.345 | 55.589 | 1.00 | 0.00 | H |
| ATOM | 3666 | CA | ALA | A | 257 | 31.975 | 27.491 | 53.948 | 1.00 | 0.00 | C |
| ATOM | 3667 | HA | ALA | A | 257 | 32.042 | 26.904 | 53.142 | 1.00 | 0.00 | H |
| ATOM | 3668 | CB | ALA | A | 257 | 30.919 | 26.923 | 54.948 | 1.00 | 0.00 | C |
| ATOM | 3669 | HB1 | ALA | A | 257 | 30.015 | 26.937 | 54.521 | 1.00 | 0.00 | H |

```
ATOM   3670  HB2 ALA A 257      31.160  25.983  55.189  1.00  0.00           H
ATOM   3671  HB3 ALA A 257      30.906  27.486  55.775  1.00  0.00           H
ATOM   3672  C   ALA A 257      31.639  28.912  53.419  1.00  0.00           C
ATOM   3673  O   ALA A 257      31.127  29.053  52.294  1.00  0.00           O
ATOM   3674  N   SER A 258      32.106  29.891  54.208  1.00  0.00           N
ATOM   3675  H   SER A 258      32.579  29.615  55.045  1.00  0.00           H
ATOM   3676  CA  SER A 258      31.984  31.327  53.953  1.00  0.00           C
ATOM   3677  HA  SER A 258      31.011  31.414  53.739  1.00  0.00           H
ATOM   3678  CB  SER A 258      32.460  32.082  55.204  1.00  0.00           C
ATOM   3679  HB1 SER A 258      31.967  31.792  56.024  1.00  0.00           H
ATOM   3680  HB2 SER A 258      32.374  33.072  55.096  1.00  0.00           H
ATOM   3681  OG  SER A 258      33.812  31.777  55.377  1.00  0.00           O
ATOM   3682  HG  SER A 258      34.160  32.252  56.185  1.00  0.00           H
ATOM   3683  C   SER A 258      32.791  31.883  52.756  1.00  0.00           C
ATOM   3684  O   SER A 258      32.445  33.006  52.298  1.00  0.00           O
ATOM   3685  N   LYS A 259      33.675  31.067  52.210  1.00  0.00           N
ATOM   3686  H   LYS A 259      33.893  30.210  52.677  1.00  0.00           H
ATOM   3687  CA  LYS A 259      34.362  31.379  50.918  1.00  0.00           C
ATOM   3688  HA  LYS A 259      34.093  32.322  50.720  1.00  0.00           H
ATOM   3689  CB  LYS A 259      35.877  31.278  51.065  1.00  0.00           C
ATOM   3690  HB1 LYS A 259      36.290  31.141  50.165  1.00  0.00           H
ATOM   3691  HB2 LYS A 259      36.103  30.504  51.657  1.00  0.00           H
ATOM   3692  CG  LYS A 259      36.402  32.590  51.691  1.00  0.00           C
ATOM   3693  HG1 LYS A 259      36.013  32.685  52.607  1.00  0.00           H
ATOM   3694  HG2 LYS A 259      36.105  33.357  51.122  1.00  0.00           H
ATOM   3695  CD  LYS A 259      37.918  32.656  51.815  1.00  0.00           C
ATOM   3696  HD1 LYS A 259      38.293  32.368  50.934  1.00  0.00           H
ATOM   3697  HD2 LYS A 259      38.180  31.997  52.520  1.00  0.00           H
ATOM   3698  CE  LYS A 259      38.573  34.001  52.176  1.00  0.00           C
ATOM   3699  HE1 LYS A 259      38.245  34.274  53.080  1.00  0.00           H
ATOM   3700  HE2 LYS A 259      38.285  34.678  51.499  1.00  0.00           H
ATOM   3701  NZ  LYS A 259      40.085  34.025  52.221  1.00  0.00           N
ATOM   3702  HZ1 LYS A 259      40.397  34.944  52.464  1.00  0.00           H
ATOM   3703  HZ2 LYS A 259      40.409  33.371  52.905  1.00  0.00           H
ATOM   3704  HZ3 LYS A 259      40.449  33.776  51.324  1.00  0.00           H
ATOM   3705  C   LYS A 259      33.905  30.491  49.753  1.00  0.00           C
ATOM   3706  O   LYS A 259      34.193  30.896  48.616  1.00  0.00           O
ATOM   3707  N   LEU A 260      33.289  29.352  49.953  1.00  0.00           N
ATOM   3708  H   LEU A 260      33.166  29.044  50.896  1.00  0.00           H
ATOM   3709  CA  LEU A 260      32.761  28.489  48.862  1.00  0.00           C
ATOM   3710  HA  LEU A 260      33.365  28.490  48.065  1.00  0.00           H
ATOM   3711  CB  LEU A 260      32.611  27.022  49.479  1.00  0.00           C
ATOM   3712  HB1 LEU A 260      31.950  27.067  50.228  1.00  0.00           H
ATOM   3713  HB2 LEU A 260      33.502  26.744  49.838  1.00  0.00           H
ATOM   3714  CG  LEU A 260      32.117  25.903  48.440  1.00  0.00           C
ATOM   3715  HG  LEU A 260      31.342  26.243  47.907  1.00  0.00           H
ATOM   3716  CD1 LEU A 260      33.116  25.470  47.365  1.00  0.00           C
ATOM   3717 HD11 LEU A 260      32.697  24.774  46.782  1.00  0.00           H
ATOM   3718 HD12 LEU A 260      33.371  26.261  46.809  1.00  0.00           H
ATOM   3719 HD13 LEU A 260      33.932  25.092  47.802  1.00  0.00           H
ATOM   3720  CD2 LEU A 260      31.874  24.669  49.336  1.00  0.00           C
ATOM   3721 HD21 LEU A 260      31.558  23.905  48.773  1.00  0.00           H
ATOM   3722 HD22 LEU A 260      32.727  24.416  49.793  1.00  0.00           H
ATOM   3723 HD23 LEU A 260      31.180  24.887  50.022  1.00  0.00           H
```

```
ATOM   3724  C   LEU A 260      31.452  29.091  48.382  1.00  0.00           C
ATOM   3725  O   LEU A 260      30.547  29.491  49.112  1.00  0.00           O
ATOM   3726  N   ILE A 261      31.326  29.297  47.017  1.00  0.00           N
ATOM   3727  H   ILE A 261      32.109  28.996  46.473  1.00  0.00           H
ATOM   3728  CA  ILE A 261      30.194  29.904  46.220  1.00  0.00           C
ATOM   3729  HA  ILE A 261      29.416  30.211  46.768  1.00  0.00           H
ATOM   3730  CB  ILE A 261      30.782  31.148  45.502  1.00  0.00           C
ATOM   3731  HB  ILE A 261      31.397  30.737  44.829  1.00  0.00           H
ATOM   3732  CG2 ILE A 261      29.757  31.993  44.753  1.00  0.00           C
ATOM   3733 HG21 ILE A 261      30.216  32.770  44.322  1.00  0.00           H
ATOM   3734 HG22 ILE A 261      29.315  31.435  44.051  1.00  0.00           H
ATOM   3735 HG23 ILE A 261      29.068  32.328  45.395  1.00  0.00           H
ATOM   3736  CG1 ILE A 261      31.599  32.079  46.405  1.00  0.00           C
ATOM   3737 HG11 ILE A 261      32.061  32.743  45.816  1.00  0.00           H
ATOM   3738 HG12 ILE A 261      32.281  31.520  46.877  1.00  0.00           H
ATOM   3739  CD  ILE A 261      30.805  32.862  47.473  1.00  0.00           C
ATOM   3740  HD1 ILE A 261      31.432  33.434  48.002  1.00  0.00           H
ATOM   3741  HD2 ILE A 261      30.124  33.441  47.024  1.00  0.00           H
ATOM   3742  HD3 ILE A 261      30.345  32.218  48.085  1.00  0.00           H
ATOM   3743  C   ILE A 261      29.675  28.793  45.296  1.00  0.00           C
ATOM   3744  O   ILE A 261      30.463  28.042  44.701  1.00  0.00           O
ATOM   3745  N   LEU A 262      28.348  28.712  45.133  1.00  0.00           N
ATOM   3746  H   LEU A 262      27.784  29.417  45.563  1.00  0.00           H
ATOM   3747  CA  LEU A 262      27.664  27.665  44.370  1.00  0.00           C
ATOM   3748  HA  LEU A 262      28.354  26.967  44.179  1.00  0.00           H
ATOM   3749  CB  LEU A 262      26.515  27.114  45.286  1.00  0.00           C
ATOM   3750  HB1 LEU A 262      25.964  27.886  45.604  1.00  0.00           H
ATOM   3751  HB2 LEU A 262      26.928  26.653  46.071  1.00  0.00           H
ATOM   3752  CG  LEU A 262      25.559  26.128  44.655  1.00  0.00           C
ATOM   3753  HG  LEU A 262      25.297  26.571  43.797  1.00  0.00           H
ATOM   3754  CD1 LEU A 262      26.250  24.788  44.473  1.00  0.00           C
ATOM   3755 HD11 LEU A 262      25.614  24.139  44.055  1.00  0.00           H
ATOM   3756 HD12 LEU A 262      27.047  24.902  43.880  1.00  0.00           H
ATOM   3757 HD13 LEU A 262      26.544  24.441  45.364  1.00  0.00           H
ATOM   3758  CD2 LEU A 262      24.324  25.785  45.557  1.00  0.00           C
ATOM   3759 HD21 LEU A 262      23.735  25.131  45.082  1.00  0.00           H
ATOM   3760 HD22 LEU A 262      24.640  25.385  46.417  1.00  0.00           H
ATOM   3761 HD23 LEU A 262      23.808  26.620  45.748  1.00  0.00           H
ATOM   3762  C   LEU A 262      27.208  28.101  42.946  1.00  0.00           C
ATOM   3763  O   LEU A 262      26.269  28.874  42.746  1.00  0.00           O
ATOM   3764  N   GLY A 263      27.770  27.413  41.925  1.00  0.00           N
ATOM   3765  H   GLY A 263      28.471  26.732  42.136  1.00  0.00           H
ATOM   3766  CA  GLY A 263      27.393  27.624  40.510  1.00  0.00           C
ATOM   3767  HA1 GLY A 263      28.071  27.162  39.938  1.00  0.00           H
ATOM   3768  HA2 GLY A 263      27.417  28.607  40.327  1.00  0.00           H
ATOM   3769  C   GLY A 263      26.017  27.093  40.170  1.00  0.00           C
ATOM   3770  O   GLY A 263      25.732  25.911  40.146  1.00  0.00           O
ATOM   3771  N   MET A 264      25.135  28.071  39.988  1.00  0.00           N
ATOM   3772  H   MET A 264      25.493  28.989  40.158  1.00  0.00           H
ATOM   3773  CA  MET A 264      23.776  28.034  39.595  1.00  0.00           C
ATOM   3774  HA  MET A 264      23.454  27.113  39.813  1.00  0.00           H
ATOM   3775  CB  MET A 264      23.023  29.102  40.420  1.00  0.00           C
ATOM   3776  HB1 MET A 264      22.084  29.192  40.089  1.00  0.00           H
ATOM   3777  HB2 MET A 264      23.488  29.984  40.346  1.00  0.00           H
```

| | | | | | | | | | | | |
|---|---|---|---|---|---|---|---|---|---|---|---|
| ATOM | 3778 | CG | MET | A | 264 | 22.996 | 28.681 | 41.864 | 1.00 | 0.00 | C |
| ATOM | 3779 | HG1 | MET | A | 264 | 23.946 | 28.787 | 42.158 | 1.00 | 0.00 | H |
| ATOM | 3780 | HG2 | MET | A | 264 | 22.767 | 27.709 | 41.822 | 1.00 | 0.00 | H |
| ATOM | 3781 | SD | MET | A | 264 | 21.892 | 29.555 | 42.932 | 1.00 | 0.00 | S |
| ATOM | 3782 | CE | MET | A | 264 | 20.343 | 28.727 | 42.500 | 1.00 | 0.00 | C |
| ATOM | 3783 | HE1 | MET | A | 264 | 19.595 | 29.119 | 43.035 | 1.00 | 0.00 | H |
| ATOM | 3784 | HE2 | MET | A | 264 | 20.158 | 28.854 | 41.526 | 1.00 | 0.00 | H |
| ATOM | 3785 | HE3 | MET | A | 264 | 20.420 | 27.750 | 42.700 | 1.00 | 0.00 | H |
| ATOM | 3786 | C | MET | A | 264 | 23.647 | 28.296 | 38.053 | 1.00 | 0.00 | C |
| ATOM | 3787 | O | MET | A | 264 | 23.913 | 29.383 | 37.482 | 1.00 | 0.00 | O |
| ATOM | 3788 | N | PRO | A | 265 | 23.370 | 27.284 | 37.260 | 1.00 | 0.00 | N |
| ATOM | 3789 | CD | PRO | A | 265 | 23.286 | 25.993 | 37.716 | 1.00 | 0.00 | C |
| ATOM | 3790 | HD1 | PRO | A | 265 | 22.468 | 25.929 | 38.288 | 1.00 | 0.00 | H |
| ATOM | 3791 | HD2 | PRO | A | 265 | 24.097 | 25.818 | 38.274 | 1.00 | 0.00 | H |
| ATOM | 3792 | CG | PRO | A | 265 | 23.212 | 25.055 | 36.502 | 1.00 | 0.00 | C |
| ATOM | 3793 | HG1 | PRO | A | 265 | 22.321 | 24.603 | 36.460 | 1.00 | 0.00 | H |
| ATOM | 3794 | HG2 | PRO | A | 265 | 23.934 | 24.365 | 36.547 | 1.00 | 0.00 | H |
| ATOM | 3795 | CB | PRO | A | 265 | 23.411 | 25.960 | 35.283 | 1.00 | 0.00 | C |
| ATOM | 3796 | HB1 | PRO | A | 265 | 22.744 | 25.744 | 34.570 | 1.00 | 0.00 | H |
| ATOM | 3797 | HB2 | PRO | A | 265 | 24.336 | 25.860 | 34.916 | 1.00 | 0.00 | H |
| ATOM | 3798 | CA | PRO | A | 265 | 23.191 | 27.378 | 35.816 | 1.00 | 0.00 | C |
| ATOM | 3799 | HA | PRO | A | 265 | 23.867 | 28.014 | 35.444 | 1.00 | 0.00 | H |
| ATOM | 3800 | C | PRO | A | 265 | 21.824 | 27.980 | 35.437 | 1.00 | 0.00 | C |
| ATOM | 3801 | O | PRO | A | 265 | 20.788 | 27.497 | 35.831 | 1.00 | 0.00 | O |
| ATOM | 3802 | N | THR | A | 266 | 21.839 | 28.973 | 34.526 | 1.00 | 0.00 | N |
| ATOM | 3803 | H | THR | A | 266 | 22.724 | 29.375 | 34.292 | 1.00 | 0.00 | H |
| ATOM | 3804 | CA | THR | A | 266 | 20.633 | 29.515 | 33.843 | 1.00 | 0.00 | C |
| ATOM | 3805 | HA | THR | A | 266 | 19.840 | 29.421 | 34.445 | 1.00 | 0.00 | H |
| ATOM | 3806 | CB | THR | A | 266 | 20.742 | 31.006 | 33.596 | 1.00 | 0.00 | C |
| ATOM | 3807 | HB | THR | A | 266 | 20.051 | 31.304 | 32.938 | 1.00 | 0.00 | H |
| ATOM | 3808 | CG2 | THR | A | 266 | 20.585 | 31.876 | 34.836 | 1.00 | 0.00 | C |
| ATOM | 3809 | HG21 | THR | A | 266 | 20.671 | 32.839 | 34.581 | 1.00 | 0.00 | H |
| ATOM | 3810 | HG22 | THR | A | 266 | 19.685 | 31.718 | 35.242 | 1.00 | 0.00 | H |
| ATOM | 3811 | HG23 | THR | A | 266 | 21.296 | 31.643 | 35.499 | 1.00 | 0.00 | H |
| ATOM | 3812 | OG1 | THR | A | 266 | 21.970 | 31.213 | 33.025 | 1.00 | 0.00 | O |
| ATOM | 3813 | HG1 | THR | A | 266 | 22.092 | 32.188 | 32.841 | 1.00 | 0.00 | H |
| ATOM | 3814 | C | THR | A | 266 | 20.406 | 28.695 | 32.582 | 1.00 | 0.00 | C |
| ATOM | 3815 | O | THR | A | 266 | 19.855 | 29.172 | 31.555 | 1.00 | 0.00 | O |
| ATOM | 3816 | N | TYR | A | 267 | 20.799 | 27.415 | 32.556 | 1.00 | 0.00 | N |
| ATOM | 3817 | H | TYR | A | 267 | 21.203 | 26.996 | 33.369 | 1.00 | 0.00 | H |
| ATOM | 3818 | CA | TYR | A | 267 | 20.633 | 26.638 | 31.334 | 1.00 | 0.00 | C |
| ATOM | 3819 | HA | TYR | A | 267 | 19.780 | 27.025 | 30.983 | 1.00 | 0.00 | H |
| ATOM | 3820 | CB | TYR | A | 267 | 21.843 | 26.844 | 30.396 | 1.00 | 0.00 | C |
| ATOM | 3821 | HB1 | TYR | A | 267 | 21.956 | 27.825 | 30.241 | 1.00 | 0.00 | H |
| ATOM | 3822 | HB2 | TYR | A | 267 | 21.647 | 26.389 | 29.527 | 1.00 | 0.00 | H |
| ATOM | 3823 | CG | TYR | A | 267 | 23.171 | 26.299 | 30.918 | 1.00 | 0.00 | C |
| ATOM | 3824 | CD1 | TYR | A | 267 | 23.949 | 27.078 | 31.795 | 1.00 | 0.00 | C |
| ATOM | 3825 | HD1 | TYR | A | 267 | 23.621 | 27.977 | 32.084 | 1.00 | 0.00 | H |
| ATOM | 3826 | CE1 | TYR | A | 267 | 25.186 | 26.599 | 32.268 | 1.00 | 0.00 | C |
| ATOM | 3827 | HE1 | TYR | A | 267 | 25.722 | 27.156 | 32.902 | 1.00 | 0.00 | H |
| ATOM | 3828 | CZ | TYR | A | 267 | 25.677 | 25.343 | 31.858 | 1.00 | 0.00 | C |
| ATOM | 3829 | OH | TYR | A | 267 | 26.887 | 24.928 | 32.319 | 1.00 | 0.00 | O |
| ATOM | 3830 | HH | TYR | A | 267 | 27.096 | 24.025 | 31.942 | 1.00 | 0.00 | H |
| ATOM | 3831 | CE2 | TYR | A | 267 | 24.896 | 24.554 | 30.984 | 1.00 | 0.00 | C |

```
ATOM   3832  HE2 TYR A 267      25.226  23.656  30.694  1.00  0.00           H
ATOM   3833  CD2 TYR A 267      23.650  25.032  30.519  1.00  0.00           C
ATOM   3834  HD2 TYR A 267      23.102  24.468  29.901  1.00  0.00           H
ATOM   3835  C   TYR A 267      20.311  25.177  31.484  1.00  0.00           C
ATOM   3836  O   TYR A 267      20.465  24.589  32.532  1.00  0.00           O
ATOM   3837  N   GLY A 268      19.850  24.554  30.417  1.00  0.00           N
ATOM   3838  H   GLY A 268      19.509  25.092  29.646  1.00  0.00           H
ATOM   3839  CA  GLY A 268      19.825  23.081  30.334  1.00  0.00           C
ATOM   3840  HA1 GLY A 268      18.915  22.829  30.006  1.00  0.00           H
ATOM   3841  HA2 GLY A 268      19.949  22.741  31.266  1.00  0.00           H
ATOM   3842  C   GLY A 268      20.851  22.399  29.444  1.00  0.00           C
ATOM   3843  O   GLY A 268      21.382  23.043  28.544  1.00  0.00           O
ATOM   3844  N   ARG A 269      21.029  21.072  29.584  1.00  0.00           N
ATOM   3845  H   ARG A 269      20.621  20.613  30.373  1.00  0.00           H
ATOM   3846  CA  ARG A 269      21.799  20.272  28.622  1.00  0.00           C
ATOM   3847  HA  ARG A 269      22.066  20.888  27.880  1.00  0.00           H
ATOM   3848  CB  ARG A 269      23.070  19.768  29.307  1.00  0.00           C
ATOM   3849  HB1 ARG A 269      23.606  19.243  28.646  1.00  0.00           H
ATOM   3850  HB2 ARG A 269      22.812  19.177  30.072  1.00  0.00           H
ATOM   3851  CG  ARG A 269      23.923  20.940  29.836  1.00  0.00           C
ATOM   3852  HG1 ARG A 269      23.578  21.205  30.736  1.00  0.00           H
ATOM   3853  HG2 ARG A 269      23.834  21.710  29.205  1.00  0.00           H
ATOM   3854  CD  ARG A 269      25.408  20.604  29.973  1.00  0.00           C
ATOM   3855  HD1 ARG A 269      25.519  19.678  30.333  1.00  0.00           H
ATOM   3856  HD2 ARG A 269      25.852  21.256  30.588  1.00  0.00           H
ATOM   3857  NE  ARG A 269      26.064  20.678  28.653  1.00  0.00           N
ATOM   3858  HE  ARG A 269      25.632  20.164  27.912  1.00  0.00           H
ATOM   3859  CZ  ARG A 269      27.161  21.349  28.342  1.00  0.00           C
ATOM   3860  NH1 ARG A 269      27.637  21.283  27.134  1.00  0.00           N
ATOM   3861 HH11 ARG A 269      27.174  20.727  26.444  1.00  0.00           H
ATOM   3862 HH12 ARG A 269      28.466  21.789  26.895  1.00  0.00           H
ATOM   3863  NH2 ARG A 269      27.837  22.084  29.169  1.00  0.00           N
ATOM   3864 HH21 ARG A 269      27.537  22.173  30.119  1.00  0.00           H
ATOM   3865 HH22 ARG A 269      28.658  22.561  28.857  1.00  0.00           H
ATOM   3866  C   ARG A 269      20.955  19.185  27.928  1.00  0.00           C
ATOM   3867  O   ARG A 269      20.164  18.511  28.583  1.00  0.00           O
ATOM   3868  N   SER A 270      21.129  19.042  26.630  1.00  0.00           N
ATOM   3869  H   SER A 270      21.974  19.394  26.226  1.00  0.00           H
ATOM   3870  CA  SER A 270      20.116  18.370  25.706  1.00  0.00           C
ATOM   3871  HA  SER A 270      19.327  18.234  26.305  1.00  0.00           H
ATOM   3872  CB  SER A 270      19.783  19.273  24.411  1.00  0.00           C
ATOM   3873  HB1 SER A 270      19.649  20.234  24.654  1.00  0.00           H
ATOM   3874  HB2 SER A 270      18.977  18.937  23.923  1.00  0.00           H
ATOM   3875  OG  SER A 270      20.914  19.202  23.498  1.00  0.00           O
ATOM   3876  HG  SER A 270      20.724  19.757  22.688  1.00  0.00           H
ATOM   3877  C   SER A 270      20.535  16.984  25.243  1.00  0.00           C
ATOM   3878  O   SER A 270      21.721  16.619  25.192  1.00  0.00           O
ATOM   3879  N   PHE A 271      19.591  16.171  24.840  1.00  0.00           N
ATOM   3880  H   PHE A 271      18.651  16.494  24.952  1.00  0.00           H
ATOM   3881  CA  PHE A 271      19.757  14.814  24.231  1.00  0.00           C
ATOM   3882  HA  PHE A 271      20.641  14.938  23.779  1.00  0.00           H
ATOM   3883  CB  PHE A 271      19.739  13.636  25.277  1.00  0.00           C
ATOM   3884  HB1 PHE A 271      19.971  12.786  24.805  1.00  0.00           H
ATOM   3885  HB2 PHE A 271      18.818  13.563  25.661  1.00  0.00           H
```

| | | | | | | | | | | | |
|---|---|---|---|---|---|---|---|---|---|---|---|
| ATOM | 3886 | CG | PHE | A | 271 | 20.713 | 13.788 | 26.451 | 1.00 | 0.00 | C |
| ATOM | 3887 | CD1 | PHE | A | 271 | 20.299 | 14.580 | 27.480 | 1.00 | 0.00 | C |
| ATOM | 3888 | HD1 | PHE | A | 271 | 19.487 | 15.153 | 27.370 | 1.00 | 0.00 | H |
| ATOM | 3889 | CE1 | PHE | A | 271 | 21.010 | 14.598 | 28.704 | 1.00 | 0.00 | C |
| ATOM | 3890 | HE1 | PHE | A | 271 | 20.676 | 15.143 | 29.473 | 1.00 | 0.00 | H |
| ATOM | 3891 | CZ | PHE | A | 271 | 22.177 | 13.852 | 28.830 | 1.00 | 0.00 | C |
| ATOM | 3892 | HZ | PHE | A | 271 | 22.770 | 13.975 | 29.625 | 1.00 | 0.00 | H |
| ATOM | 3893 | CE2 | PHE | A | 271 | 22.517 | 12.920 | 27.826 | 1.00 | 0.00 | C |
| ATOM | 3894 | HE2 | PHE | A | 271 | 23.243 | 12.249 | 27.975 | 1.00 | 0.00 | H |
| ATOM | 3895 | CD2 | PHE | A | 271 | 21.827 | 12.957 | 26.641 | 1.00 | 0.00 | C |
| ATOM | 3896 | HD2 | PHE | A | 271 | 22.128 | 12.372 | 25.888 | 1.00 | 0.00 | H |
| ATOM | 3897 | C | PHE | A | 271 | 18.662 | 14.456 | 23.230 | 1.00 | 0.00 | C |
| ATOM | 3898 | O | PHE | A | 271 | 17.535 | 14.817 | 23.462 | 1.00 | 0.00 | O |
| ATOM | 3899 | N | THR | A | 272 | 18.943 | 13.720 | 22.197 | 1.00 | 0.00 | N |
| ATOM | 3900 | H | THR | A | 272 | 19.924 | 13.640 | 22.020 | 1.00 | 0.00 | H |
| ATOM | 3901 | CA | THR | A | 272 | 18.126 | 12.994 | 21.253 | 1.00 | 0.00 | C |
| ATOM | 3902 | HA | THR | A | 272 | 17.186 | 13.335 | 21.230 | 1.00 | 0.00 | H |
| ATOM | 3903 | CB | THR | A | 272 | 18.630 | 13.239 | 19.789 | 1.00 | 0.00 | C |
| ATOM | 3904 | HB | THR | A | 272 | 18.522 | 14.171 | 19.444 | 1.00 | 0.00 | H |
| ATOM | 3905 | CG2 | THR | A | 272 | 20.100 | 12.963 | 19.564 | 1.00 | 0.00 | C |
| ATOM | 3906 | HG21 | THR | A | 272 | 20.331 | 13.145 | 18.608 | 1.00 | 0.00 | H |
| ATOM | 3907 | HG22 | THR | A | 272 | 20.645 | 13.555 | 20.157 | 1.00 | 0.00 | H |
| ATOM | 3908 | HG23 | THR | A | 272 | 20.296 | 12.006 | 19.778 | 1.00 | 0.00 | H |
| ATOM | 3909 | OG1 | THR | A | 272 | 17.822 | 12.393 | 19.056 | 1.00 | 0.00 | O |
| ATOM | 3910 | HG1 | THR | A | 272 | 18.057 | 12.465 | 18.087 | 1.00 | 0.00 | H |
| ATOM | 3911 | C | THR | A | 272 | 17.966 | 11.602 | 21.802 | 1.00 | 0.00 | C |
| ATOM | 3912 | O | THR | A | 272 | 19.007 | 10.955 | 22.090 | 1.00 | 0.00 | O |
| ATOM | 3913 | N | LEU | A | 273 | 16.753 | 11.096 | 22.058 | 1.00 | 0.00 | N |
| ATOM | 3914 | H | LEU | A | 273 | 15.957 | 11.668 | 21.862 | 1.00 | 0.00 | H |
| ATOM | 3915 | CA | LEU | A | 273 | 16.518 | 9.765 | 22.604 | 1.00 | 0.00 | C |
| ATOM | 3916 | HA | LEU | A | 273 | 17.250 | 9.679 | 23.280 | 1.00 | 0.00 | H |
| ATOM | 3917 | CB | LEU | A | 273 | 15.032 | 9.768 | 23.152 | 1.00 | 0.00 | C |
| ATOM | 3918 | HB1 | LEU | A | 273 | 14.800 | 8.839 | 23.439 | 1.00 | 0.00 | H |
| ATOM | 3919 | HB2 | LEU | A | 273 | 14.423 | 10.050 | 22.411 | 1.00 | 0.00 | H |
| ATOM | 3920 | CG | LEU | A | 273 | 14.805 | 10.740 | 24.380 | 1.00 | 0.00 | C |
| ATOM | 3921 | HG | LEU | A | 273 | 15.245 | 11.625 | 24.229 | 1.00 | 0.00 | H |
| ATOM | 3922 | CD1 | LEU | A | 273 | 13.255 | 10.796 | 24.614 | 1.00 | 0.00 | C |
| ATOM | 3923 | HD11 | LEU | A | 273 | 13.056 | 11.401 | 25.385 | 1.00 | 0.00 | H |
| ATOM | 3924 | HD12 | LEU | A | 273 | 12.806 | 11.145 | 23.791 | 1.00 | 0.00 | H |
| ATOM | 3925 | HD13 | LEU | A | 273 | 12.915 | 9.877 | 24.815 | 1.00 | 0.00 | H |
| ATOM | 3926 | CD2 | LEU | A | 273 | 15.339 | 10.068 | 25.644 | 1.00 | 0.00 | C |
| ATOM | 3927 | HD21 | LEU | A | 273 | 15.201 | 10.674 | 26.427 | 1.00 | 0.00 | H |
| ATOM | 3928 | HD22 | LEU | A | 273 | 14.850 | 9.209 | 25.797 | 1.00 | 0.00 | H |
| ATOM | 3929 | HD23 | LEU | A | 273 | 16.315 | 9.881 | 25.535 | 1.00 | 0.00 | H |
| ATOM | 3930 | C | LEU | A | 273 | 16.622 | 8.566 | 21.543 | 1.00 | 0.00 | C |
| ATOM | 3931 | O | LEU | A | 273 | 16.101 | 8.623 | 20.372 | 1.00 | 0.00 | O |
| ATOM | 3932 | N | ALA | A | 274 | 17.246 | 7.459 | 21.975 | 1.00 | 0.00 | N |
| ATOM | 3933 | H | ALA | A | 274 | 17.677 | 7.475 | 22.877 | 1.00 | 0.00 | H |
| ATOM | 3934 | CA | ALA | A | 274 | 17.328 | 6.176 | 21.152 | 1.00 | 0.00 | C |
| ATOM | 3935 | HA | ALA | A | 274 | 17.930 | 6.324 | 20.367 | 1.00 | 0.00 | H |
| ATOM | 3936 | CB | ALA | A | 274 | 17.920 | 5.141 | 21.963 | 1.00 | 0.00 | C |
| ATOM | 3937 | HB1 | ALA | A | 274 | 17.979 | 4.295 | 21.433 | 1.00 | 0.00 | H |
| ATOM | 3938 | HB2 | ALA | A | 274 | 18.837 | 5.422 | 22.245 | 1.00 | 0.00 | H |
| ATOM | 3939 | HB3 | ALA | A | 274 | 17.355 | 4.988 | 22.773 | 1.00 | 0.00 | H |

| | | | | | | | | | | | |
|---|---|---|---|---|---|---|---|---|---|---|---|
| ATOM | 3940 | C | ALA | A | 274 | 15.911 | 5.705 | 20.700 | 1.00 | 0.00 | C |
| ATOM | 3941 | O | ALA | A | 274 | 15.745 | 5.167 | 19.651 | 1.00 | 0.00 | O |
| ATOM | 3942 | N | SER | A | 275 | 14.872 | 5.869 | 21.574 | 1.00 | 0.00 | N |
| ATOM | 3943 | H | SER | A | 275 | 15.105 | 5.970 | 22.541 | 1.00 | 0.00 | H |
| ATOM | 3944 | CA | SER | A | 275 | 13.473 | 5.910 | 21.231 | 1.00 | 0.00 | C |
| ATOM | 3945 | HA | SER | A | 275 | 13.389 | 6.217 | 20.283 | 1.00 | 0.00 | H |
| ATOM | 3946 | CB | SER | A | 275 | 12.885 | 4.509 | 21.313 | 1.00 | 0.00 | C |
| ATOM | 3947 | HB1 | SER | A | 275 | 13.063 | 4.092 | 22.204 | 1.00 | 0.00 | H |
| ATOM | 3948 | HB2 | SER | A | 275 | 13.247 | 3.920 | 20.590 | 1.00 | 0.00 | H |
| ATOM | 3949 | OG | SER | A | 275 | 11.489 | 4.613 | 21.143 | 1.00 | 0.00 | O |
| ATOM | 3950 | HG | SER | A | 275 | 11.078 | 3.703 | 21.193 | 1.00 | 0.00 | H |
| ATOM | 3951 | C | SER | A | 275 | 12.780 | 6.919 | 22.145 | 1.00 | 0.00 | C |
| ATOM | 3952 | O | SER | A | 275 | 13.157 | 6.993 | 23.313 | 1.00 | 0.00 | O |
| ATOM | 3953 | N | SER | A | 276 | 11.816 | 7.630 | 21.548 | 1.00 | 0.00 | N |
| ATOM | 3954 | H | SER | A | 276 | 11.678 | 7.456 | 20.573 | 1.00 | 0.00 | H |
| ATOM | 3955 | CA | SER | A | 276 | 10.937 | 8.637 | 22.159 | 1.00 | 0.00 | C |
| ATOM | 3956 | HA | SER | A | 276 | 11.475 | 9.215 | 22.772 | 1.00 | 0.00 | H |
| ATOM | 3957 | CB | SER | A | 276 | 10.321 | 9.524 | 21.018 | 1.00 | 0.00 | C |
| ATOM | 3958 | HB1 | SER | A | 276 | 9.626 | 10.153 | 21.367 | 1.00 | 0.00 | H |
| ATOM | 3959 | HB2 | SER | A | 276 | 9.930 | 8.969 | 20.284 | 1.00 | 0.00 | H |
| ATOM | 3960 | OG | SER | A | 276 | 11.329 | 10.300 | 20.453 | 1.00 | 0.00 | O |
| ATOM | 3961 | HG | SER | A | 276 | 10.944 | 10.868 | 19.725 | 1.00 | 0.00 | H |
| ATOM | 3962 | C | SER | A | 276 | 9.924 | 7.951 | 23.057 | 1.00 | 0.00 | C |
| ATOM | 3963 | O | SER | A | 276 | 9.180 | 8.562 | 23.805 | 1.00 | 0.00 | O |
| ATOM | 3964 | N | SER | A | 277 | 9.843 | 6.591 | 22.949 | 1.00 | 0.00 | N |
| ATOM | 3965 | H | SER | A | 277 | 10.381 | 6.122 | 22.249 | 1.00 | 0.00 | H |
| ATOM | 3966 | CA | SER | A | 277 | 8.945 | 5.770 | 23.878 | 1.00 | 0.00 | C |
| ATOM | 3967 | HA | SER | A | 277 | 8.015 | 6.117 | 23.752 | 1.00 | 0.00 | H |
| ATOM | 3968 | CB | SER | A | 277 | 9.186 | 4.297 | 23.491 | 1.00 | 0.00 | C |
| ATOM | 3969 | HB1 | SER | A | 277 | 8.885 | 4.103 | 22.557 | 1.00 | 0.00 | H |
| ATOM | 3970 | HB2 | SER | A | 277 | 8.736 | 3.667 | 24.124 | 1.00 | 0.00 | H |
| ATOM | 3971 | OG | SER | A | 277 | 10.573 | 4.081 | 23.563 | 1.00 | 0.00 | O |
| ATOM | 3972 | HG | SER | A | 277 | 10.776 | 3.133 | 23.318 | 1.00 | 0.00 | H |
| ATOM | 3973 | C | SER | A | 277 | 9.214 | 5.897 | 25.385 | 1.00 | 0.00 | C |
| ATOM | 3974 | O | SER | A | 277 | 8.375 | 5.476 | 26.199 | 1.00 | 0.00 | O |
| ATOM | 3975 | N | ASP | A | 278 | 10.354 | 6.395 | 25.859 | 1.00 | 0.00 | N |
| ATOM | 3976 | H | ASP | A | 278 | 11.006 | 6.732 | 25.180 | 1.00 | 0.00 | H |
| ATOM | 3977 | CA | ASP | A | 278 | 10.764 | 6.506 | 27.264 | 1.00 | 0.00 | C |
| ATOM | 3978 | HA | ASP | A | 278 | 9.937 | 6.582 | 27.820 | 1.00 | 0.00 | H |
| ATOM | 3979 | CB | ASP | A | 278 | 11.522 | 5.254 | 27.686 | 1.00 | 0.00 | C |
| ATOM | 3980 | HB1 | ASP | A | 278 | 12.420 | 5.231 | 27.246 | 1.00 | 0.00 | H |
| ATOM | 3981 | HB2 | ASP | A | 278 | 11.005 | 4.437 | 27.431 | 1.00 | 0.00 | H |
| ATOM | 3982 | CG | ASP | A | 278 | 11.688 | 5.308 | 29.190 | 1.00 | 0.00 | C |
| ATOM | 3983 | OD1 | ASP | A | 278 | 10.888 | 5.925 | 29.956 | 1.00 | 0.00 | O |
| ATOM | 3984 | OD2 | ASP | A | 278 | 12.501 | 4.546 | 29.710 | 1.00 | 0.00 | O |
| ATOM | 3985 | C | ASP | A | 278 | 11.503 | 7.769 | 27.516 | 1.00 | 0.00 | C |
| ATOM | 3986 | O | ASP | A | 278 | 12.707 | 7.941 | 27.289 | 1.00 | 0.00 | O |
| ATOM | 3987 | N | THR | A | 279 | 10.769 | 8.755 | 28.045 | 1.00 | 0.00 | N |
| ATOM | 3988 | H | THR | A | 279 | 9.835 | 8.525 | 28.317 | 1.00 | 0.00 | H |
| ATOM | 3989 | CA | THR | A | 279 | 11.202 | 10.148 | 28.266 | 1.00 | 0.00 | C |
| ATOM | 3990 | HA | THR | A | 279 | 11.966 | 10.282 | 27.634 | 1.00 | 0.00 | H |
| ATOM | 3991 | CB | THR | A | 279 | 10.142 | 11.204 | 27.899 | 1.00 | 0.00 | C |
| ATOM | 3992 | HB | THR | A | 279 | 10.499 | 12.104 | 28.151 | 1.00 | 0.00 | H |
| ATOM | 3993 | CG2 | THR | A | 279 | 9.712 | 11.107 | 26.444 | 1.00 | 0.00 | C |

```
ATOM   3994 HG21 THR A 279       9.026  11.809  26.252  1.00  0.00           H
ATOM   3995 HG22 THR A 279      10.506  11.246  25.852  1.00  0.00           H
ATOM   3996 HG23 THR A 279       9.321  10.203  26.271  1.00  0.00           H
ATOM   3997  OG1 THR A 279       8.990  11.040  28.618  1.00  0.00           O
ATOM   3998  HG1 THR A 279       8.329  11.742  28.353  1.00  0.00           H
ATOM   3999  C   THR A 279      11.672  10.305  29.700  1.00  0.00           C
ATOM   4000  O   THR A 279      11.984  11.424  30.038  1.00  0.00           O
ATOM   4001  N   ARG A 280      11.670   9.235  30.607  1.00  0.00           N
ATOM   4002  H   ARG A 280      11.302   8.352  30.316  1.00  0.00           H
ATOM   4003  CA  ARG A 280      12.199   9.378  31.986  1.00  0.00           C
ATOM   4004  HA  ARG A 280      11.622  10.097  32.374  1.00  0.00           H
ATOM   4005  CB  ARG A 280      11.999   8.090  32.799  1.00  0.00           C
ATOM   4006  HB1 ARG A 280      12.370   8.269  33.710  1.00  0.00           H
ATOM   4007  HB2 ARG A 280      12.545   7.383  32.350  1.00  0.00           H
ATOM   4008  CG  ARG A 280      10.531   7.533  32.973  1.00  0.00           C
ATOM   4009  HG1 ARG A 280      10.097   7.482  32.073  1.00  0.00           H
ATOM   4010  HG2 ARG A 280      10.016   8.161  33.556  1.00  0.00           H
ATOM   4011  CD  ARG A 280      10.515   6.094  33.631  1.00  0.00           C
ATOM   4012  HD1 ARG A 280       9.582   5.771  33.793  1.00  0.00           H
ATOM   4013  HD2 ARG A 280      11.028   6.074  34.489  1.00  0.00           H
ATOM   4014  NE  ARG A 280      11.161   5.197  32.679  1.00  0.00           N
ATOM   4015  HE  ARG A 280      11.387   5.577  31.782  1.00  0.00           H
ATOM   4016  CZ  ARG A 280      11.480   3.914  32.891  1.00  0.00           C
ATOM   4017  NH1 ARG A 280      11.960   3.287  31.933  1.00  0.00           N
ATOM   4018 HH11 ARG A 280      12.092   3.743  31.053  1.00  0.00           H
ATOM   4019 HH12 ARG A 280      12.214   2.326  32.041  1.00  0.00           H
ATOM   4020  NH2 ARG A 280      11.260   3.251  34.037  1.00  0.00           N
ATOM   4021 HH21 ARG A 280      10.829   3.717  34.809  1.00  0.00           H
ATOM   4022 HH22 ARG A 280      11.527   2.291  34.118  1.00  0.00           H
ATOM   4023  C   ARG A 280      13.711   9.834  32.026  1.00  0.00           C
ATOM   4024  O   ARG A 280      14.350   9.928  30.991  1.00  0.00           O
ATOM   4025  N   VAL A 281      14.287  10.223  33.162  1.00  0.00           N
ATOM   4026  H   VAL A 281      13.713  10.465  33.944  1.00  0.00           H
ATOM   4027  CA  VAL A 281      15.765  10.305  33.292  1.00  0.00           C
ATOM   4028  HA  VAL A 281      16.171  10.791  32.518  1.00  0.00           H
ATOM   4029  CB  VAL A 281      16.116  11.099  34.481  1.00  0.00           C
ATOM   4030  HB  VAL A 281      15.616  11.942  34.282  1.00  0.00           H
ATOM   4031  CG1 VAL A 281      15.644  10.586  35.847  1.00  0.00           C
ATOM   4032 HG11 VAL A 281      15.948  11.215  36.563  1.00  0.00           H
ATOM   4033 HG12 VAL A 281      14.646  10.530  35.855  1.00  0.00           H
ATOM   4034 HG13 VAL A 281      16.031   9.679  36.014  1.00  0.00           H
ATOM   4035  CG2 VAL A 281      17.614  11.268  34.573  1.00  0.00           C
ATOM   4036 HG21 VAL A 281      17.839  11.808  35.384  1.00  0.00           H
ATOM   4037 HG22 VAL A 281      18.047  10.369  34.640  1.00  0.00           H
ATOM   4038 HG23 VAL A 281      17.947  11.738  33.756  1.00  0.00           H
ATOM   4039  C   VAL A 281      16.327   8.917  33.310  1.00  0.00           C
ATOM   4040  O   VAL A 281      15.592   8.006  33.534  1.00  0.00           O
ATOM   4041  N   GLY A 282      17.671   8.762  32.918  1.00  0.00           N
ATOM   4042  H   GLY A 282      18.232   9.554  32.677  1.00  0.00           H
ATOM   4043  CA  GLY A 282      18.228   7.391  32.874  1.00  0.00           C
ATOM   4044  HA1 GLY A 282      17.909   6.922  33.698  1.00  0.00           H
ATOM   4045  HA2 GLY A 282      19.223   7.481  32.906  1.00  0.00           H
ATOM   4046  C   GLY A 282      17.825   6.558  31.617  1.00  0.00           C
ATOM   4047  O   GLY A 282      18.176   5.399  31.449  1.00  0.00           O
```

| | | | | | | | | | | | |
|---|---|---|---|---|---|---|---|---|---|---|---|
| ATOM | 4048 | N | ALA | A | 283 | 17.166 | 7.257 | 30.665 | 1.00 | 0.00 | N |
| ATOM | 4049 | H | ALA | A | 283 | 16.942 | 8.204 | 30.895 | 1.00 | 0.00 | H |
| ATOM | 4050 | CA | ALA | A | 283 | 16.733 | 6.797 | 29.322 | 1.00 | 0.00 | C |
| ATOM | 4051 | HA | ALA | A | 283 | 16.464 | 5.834 | 29.309 | 1.00 | 0.00 | H |
| ATOM | 4052 | CB | ALA | A | 283 | 15.417 | 7.513 | 28.959 | 1.00 | 0.00 | C |
| ATOM | 4053 | HB1 | ALA | A | 283 | 15.112 | 7.213 | 28.055 | 1.00 | 0.00 | H |
| ATOM | 4054 | HB2 | ALA | A | 283 | 14.718 | 7.289 | 29.638 | 1.00 | 0.00 | H |
| ATOM | 4055 | HB3 | ALA | A | 283 | 15.567 | 8.502 | 28.949 | 1.00 | 0.00 | H |
| ATOM | 4056 | C | ALA | A | 283 | 17.960 | 6.854 | 28.359 | 1.00 | 0.00 | C |
| ATOM | 4057 | O | ALA | A | 283 | 18.778 | 7.764 | 28.494 | 1.00 | 0.00 | O |
| ATOM | 4058 | N | PRO | A | 284 | 18.185 | 5.926 | 27.439 | 1.00 | 0.00 | N |
| ATOM | 4059 | CD | PRO | A | 284 | 17.129 | 4.831 | 27.261 | 1.00 | 0.00 | C |
| ATOM | 4060 | HD1 | PRO | A | 284 | 16.209 | 5.219 | 27.198 | 1.00 | 0.00 | H |
| ATOM | 4061 | HD2 | PRO | A | 284 | 17.161 | 4.169 | 28.010 | 1.00 | 0.00 | H |
| ATOM | 4062 | CG | PRO | A | 284 | 17.559 | 4.196 | 25.935 | 1.00 | 0.00 | C |
| ATOM | 4063 | HG1 | PRO | A | 284 | 17.097 | 4.631 | 25.163 | 1.00 | 0.00 | H |
| ATOM | 4064 | HG2 | PRO | A | 284 | 17.367 | 3.215 | 25.929 | 1.00 | 0.00 | H |
| ATOM | 4065 | CB | PRO | A | 284 | 19.029 | 4.428 | 25.846 | 1.00 | 0.00 | C |
| ATOM | 4066 | HB1 | PRO | A | 284 | 19.321 | 4.467 | 24.890 | 1.00 | 0.00 | H |
| ATOM | 4067 | HB2 | PRO | A | 284 | 19.528 | 3.701 | 26.317 | 1.00 | 0.00 | H |
| ATOM | 4068 | CA | PRO | A | 284 | 19.272 | 5.727 | 26.508 | 1.00 | 0.00 | C |
| ATOM | 4069 | HA | PRO | A | 284 | 20.140 | 5.729 | 27.005 | 1.00 | 0.00 | H |
| ATOM | 4070 | C | PRO | A | 284 | 19.203 | 6.823 | 25.438 | 1.00 | 0.00 | C |
| ATOM | 4071 | O | PRO | A | 284 | 18.155 | 7.083 | 24.803 | 1.00 | 0.00 | O |
| ATOM | 4072 | N | ALA | A | 285 | 20.361 | 7.438 | 25.130 | 1.00 | 0.00 | N |
| ATOM | 4073 | H | ALA | A | 285 | 21.203 | 7.113 | 25.561 | 1.00 | 0.00 | H |
| ATOM | 4074 | CA | ALA | A | 285 | 20.443 | 8.533 | 24.217 | 1.00 | 0.00 | C |
| ATOM | 4075 | HA | ALA | A | 285 | 19.639 | 8.614 | 23.628 | 1.00 | 0.00 | H |
| ATOM | 4076 | CB | ALA | A | 285 | 20.511 | 9.776 | 25.106 | 1.00 | 0.00 | C |
| ATOM | 4077 | HB1 | ALA | A | 285 | 20.571 | 10.593 | 24.533 | 1.00 | 0.00 | H |
| ATOM | 4078 | HB2 | ALA | A | 285 | 19.689 | 9.826 | 25.673 | 1.00 | 0.00 | H |
| ATOM | 4079 | HB3 | ALA | A | 285 | 21.318 | 9.722 | 25.694 | 1.00 | 0.00 | H |
| ATOM | 4080 | C | ALA | A | 285 | 21.652 | 8.265 | 23.290 | 1.00 | 0.00 | C |
| ATOM | 4081 | O | ALA | A | 285 | 22.500 | 7.399 | 23.471 | 1.00 | 0.00 | O |
| ATOM | 4082 | N | THR | A | 286 | 21.618 | 9.018 | 22.162 | 1.00 | 0.00 | N |
| ATOM | 4083 | H | THR | A | 286 | 20.872 | 9.684 | 22.133 | 1.00 | 0.00 | H |
| ATOM | 4084 | CA | THR | A | 286 | 22.497 | 9.027 | 20.958 | 1.00 | 0.00 | C |
| ATOM | 4085 | HA | THR | A | 286 | 23.261 | 8.472 | 21.287 | 1.00 | 0.00 | H |
| ATOM | 4086 | CB | THR | A | 286 | 21.874 | 8.338 | 19.752 | 1.00 | 0.00 | C |
| ATOM | 4087 | HB | THR | A | 286 | 22.381 | 8.594 | 18.929 | 1.00 | 0.00 | H |
| ATOM | 4088 | CG2 | THR | A | 286 | 21.811 | 6.800 | 20.035 | 1.00 | 0.00 | C |
| ATOM | 4089 | HG21 | THR | A | 286 | 21.403 | 6.333 | 19.250 | 1.00 | 0.00 | H |
| ATOM | 4090 | HG22 | THR | A | 286 | 22.736 | 6.451 | 20.187 | 1.00 | 0.00 | H |
| ATOM | 4091 | HG23 | THR | A | 286 | 21.254 | 6.633 | 20.848 | 1.00 | 0.00 | H |
| ATOM | 4092 | OG1 | THR | A | 286 | 20.581 | 8.817 | 19.454 | 1.00 | 0.00 | O |
| ATOM | 4093 | HG1 | THR | A | 286 | 20.221 | 8.332 | 18.657 | 1.00 | 0.00 | H |
| ATOM | 4094 | C | THR | A | 286 | 23.033 | 10.427 | 20.544 | 1.00 | 0.00 | C |
| ATOM | 4095 | O | THR | A | 286 | 22.873 | 10.909 | 19.384 | 1.00 | 0.00 | O |
| ATOM | 4096 | N | GLY | A | 287 | 23.586 | 11.167 | 21.475 | 1.00 | 0.00 | N |
| ATOM | 4097 | H | GLY | A | 287 | 23.703 | 10.761 | 22.381 | 1.00 | 0.00 | H |
| ATOM | 4098 | CA | GLY | A | 287 | 24.052 | 12.566 | 21.281 | 1.00 | 0.00 | C |
| ATOM | 4099 | HA1 | GLY | A | 287 | 24.346 | 12.675 | 20.331 | 1.00 | 0.00 | H |
| ATOM | 4100 | HA2 | GLY | A | 287 | 24.827 | 12.731 | 21.891 | 1.00 | 0.00 | H |
| ATOM | 4101 | C | GLY | A | 287 | 22.918 | 13.556 | 21.595 | 1.00 | 0.00 | C |

| | | | | | | | | | | | |
|---|---|---|---|---|---|---|---|---|---|---|---|
| ATOM | 4102 | O | GLY | A | 287 | 22.043 | 13.254 | 22.362 | 1.00 | 0.00 | O |
| ATOM | 4103 | N | SER | A | 288 | 23.025 | 14.768 | 21.005 | 1.00 | 0.00 | N |
| ATOM | 4104 | H | SER | A | 288 | 23.649 | 14.839 | 20.227 | 1.00 | 0.00 | H |
| ATOM | 4105 | CA | SER | A | 288 | 22.317 | 15.980 | 21.396 | 1.00 | 0.00 | C |
| ATOM | 4106 | HA | SER | A | 288 | 21.931 | 15.685 | 22.270 | 1.00 | 0.00 | H |
| ATOM | 4107 | CB | SER | A | 288 | 23.384 | 17.142 | 21.334 | 1.00 | 0.00 | C |
| ATOM | 4108 | HB1 | SER | A | 288 | 24.216 | 16.898 | 21.832 | 1.00 | 0.00 | H |
| ATOM | 4109 | HB2 | SER | A | 288 | 23.014 | 17.995 | 21.701 | 1.00 | 0.00 | H |
| ATOM | 4110 | OG | SER | A | 288 | 23.760 | 17.392 | 19.978 | 1.00 | 0.00 | O |
| ATOM | 4111 | HG | SER | A | 288 | 24.435 | 18.129 | 19.950 | 1.00 | 0.00 | H |
| ATOM | 4112 | C | SER | A | 288 | 21.075 | 16.437 | 20.643 | 1.00 | 0.00 | C |
| ATOM | 4113 | O | SER | A | 288 | 20.032 | 16.904 | 21.213 | 1.00 | 0.00 | O |
| ATOM | 4114 | N | GLY | A | 289 | 21.156 | 16.151 | 19.372 | 1.00 | 0.00 | N |
| ATOM | 4115 | H | GLY | A | 289 | 21.974 | 15.677 | 19.045 | 1.00 | 0.00 | H |
| ATOM | 4116 | CA | GLY | A | 289 | 20.067 | 16.502 | 18.354 | 1.00 | 0.00 | C |
| ATOM | 4117 | HA1 | GLY | A | 289 | 19.220 | 16.705 | 18.846 | 1.00 | 0.00 | H |
| ATOM | 4118 | HA2 | GLY | A | 289 | 19.923 | 15.716 | 17.753 | 1.00 | 0.00 | H |
| ATOM | 4119 | C | GLY | A | 289 | 20.410 | 17.696 | 17.493 | 1.00 | 0.00 | C |
| ATOM | 4120 | O | GLY | A | 289 | 21.606 | 18.056 | 17.368 | 1.00 | 0.00 | O |
| ATOM | 4121 | N | THR | A | 290 | 19.403 | 18.276 | 16.861 | 1.00 | 0.00 | N |
| ATOM | 4122 | H | THR | A | 290 | 18.550 | 17.758 | 16.799 | 1.00 | 0.00 | H |
| ATOM | 4123 | CA | THR | A | 290 | 19.417 | 19.608 | 16.241 | 1.00 | 0.00 | C |
| ATOM | 4124 | HA | THR | A | 290 | 20.189 | 19.549 | 15.608 | 1.00 | 0.00 | H |
| ATOM | 4125 | CB | THR | A | 290 | 18.129 | 19.958 | 15.448 | 1.00 | 0.00 | C |
| ATOM | 4126 | HB | THR | A | 290 | 17.346 | 19.802 | 16.050 | 1.00 | 0.00 | H |
| ATOM | 4127 | CG2 | THR | A | 290 | 18.054 | 21.363 | 14.845 | 1.00 | 0.00 | C |
| ATOM | 4128 | HG21 | THR | A | 290 | 17.186 | 21.472 | 14.361 | 1.00 | 0.00 | H |
| ATOM | 4129 | HG22 | THR | A | 290 | 18.116 | 22.043 | 15.576 | 1.00 | 0.00 | H |
| ATOM | 4130 | HG23 | THR | A | 290 | 18.811 | 21.492 | 14.204 | 1.00 | 0.00 | H |
| ATOM | 4131 | OG1 | THR | A | 290 | 18.180 | 19.154 | 14.287 | 1.00 | 0.00 | O |
| ATOM | 4132 | HG1 | THR | A | 290 | 17.372 | 19.330 | 13.724 | 1.00 | 0.00 | H |
| ATOM | 4133 | C | THR | A | 290 | 19.670 | 20.670 | 17.274 | 1.00 | 0.00 | C |
| ATOM | 4134 | O | THR | A | 290 | 18.944 | 20.743 | 18.221 | 1.00 | 0.00 | O |
| ATOM | 4135 | N | PRO | A | 291 | 20.750 | 21.477 | 17.226 | 1.00 | 0.00 | N |
| ATOM | 4136 | CD | PRO | A | 291 | 21.731 | 21.536 | 16.167 | 1.00 | 0.00 | C |
| ATOM | 4137 | HD1 | PRO | A | 291 | 21.271 | 21.823 | 15.327 | 1.00 | 0.00 | H |
| ATOM | 4138 | HD2 | PRO | A | 291 | 22.114 | 20.621 | 16.041 | 1.00 | 0.00 | H |
| ATOM | 4139 | CG | PRO | A | 291 | 22.825 | 22.514 | 16.521 | 1.00 | 0.00 | C |
| ATOM | 4140 | HG1 | PRO | A | 291 | 22.746 | 23.345 | 15.971 | 1.00 | 0.00 | H |
| ATOM | 4141 | HG2 | PRO | A | 291 | 23.725 | 22.103 | 16.377 | 1.00 | 0.00 | H |
| ATOM | 4142 | CB | PRO | A | 291 | 22.553 | 22.793 | 18.030 | 1.00 | 0.00 | C |
| ATOM | 4143 | HB1 | PRO | A | 291 | 22.738 | 23.752 | 18.244 | 1.00 | 0.00 | H |
| ATOM | 4144 | HB2 | PRO | A | 291 | 23.129 | 22.207 | 18.600 | 1.00 | 0.00 | H |
| ATOM | 4145 | CA | PRO | A | 291 | 21.045 | 22.466 | 18.267 | 1.00 | 0.00 | C |
| ATOM | 4146 | HA | PRO | A | 291 | 20.853 | 22.052 | 19.157 | 1.00 | 0.00 | H |
| ATOM | 4147 | C | PRO | A | 291 | 20.140 | 23.791 | 18.121 | 1.00 | 0.00 | C |
| ATOM | 4148 | O | PRO | A | 291 | 19.702 | 24.144 | 17.057 | 1.00 | 0.00 | O |
| ATOM | 4149 | N | GLY | A | 292 | 19.834 | 24.442 | 19.265 | 1.00 | 0.00 | N |
| ATOM | 4150 | H | GLY | A | 292 | 20.383 | 24.230 | 20.074 | 1.00 | 0.00 | H |
| ATOM | 4151 | CA | GLY | A | 292 | 18.782 | 25.416 | 19.417 | 1.00 | 0.00 | C |
| ATOM | 4152 | HA1 | GLY | A | 292 | 18.693 | 25.640 | 20.387 | 1.00 | 0.00 | H |
| ATOM | 4153 | HA2 | GLY | A | 292 | 17.927 | 25.016 | 19.087 | 1.00 | 0.00 | H |
| ATOM | 4154 | C | GLY | A | 292 | 19.193 | 26.663 | 18.576 | 1.00 | 0.00 | C |
| ATOM | 4155 | O | GLY | A | 292 | 20.379 | 27.101 | 18.676 | 1.00 | 0.00 | O |

| | | | | | | | | | | | |
|---|---|---|---|---|---|---|---|---|---|---|---|
| ATOM | 4156 | N | PRO | A | 293 | 18.312 | 27.256 | 17.782 | 1.00 | 0.00 | N |
| ATOM | 4157 | CD | PRO | A | 293 | 17.014 | 26.778 | 17.478 | 1.00 | 0.00 | C |
| ATOM | 4158 | HD1 | PRO | A | 293 | 16.620 | 26.374 | 18.304 | 1.00 | 0.00 | H |
| ATOM | 4159 | HD2 | PRO | A | 293 | 17.089 | 26.078 | 16.768 | 1.00 | 0.00 | H |
| ATOM | 4160 | CG | PRO | A | 293 | 16.207 | 28.002 | 16.996 | 1.00 | 0.00 | C |
| ATOM | 4161 | HG1 | PRO | A | 293 | 15.928 | 28.605 | 17.743 | 1.00 | 0.00 | H |
| ATOM | 4162 | HG2 | PRO | A | 293 | 15.406 | 27.752 | 16.453 | 1.00 | 0.00 | H |
| ATOM | 4163 | CB | PRO | A | 293 | 17.306 | 28.603 | 16.154 | 1.00 | 0.00 | C |
| ATOM | 4164 | HB1 | PRO | A | 293 | 17.130 | 29.571 | 15.977 | 1.00 | 0.00 | H |
| ATOM | 4165 | HB2 | PRO | A | 293 | 17.395 | 28.116 | 15.285 | 1.00 | 0.00 | H |
| ATOM | 4166 | CA | PRO | A | 293 | 18.542 | 28.447 | 16.956 | 1.00 | 0.00 | C |
| ATOM | 4167 | HA | PRO | A | 293 | 19.340 | 28.292 | 16.373 | 1.00 | 0.00 | H |
| ATOM | 4168 | C | PRO | A | 293 | 18.816 | 29.714 | 17.803 | 1.00 | 0.00 | C |
| ATOM | 4169 | O | PRO | A | 293 | 19.325 | 30.644 | 17.271 | 1.00 | 0.00 | O |
| ATOM | 4170 | N | PHE | A | 294 | 18.513 | 29.770 | 19.122 | 1.00 | 0.00 | N |
| ATOM | 4171 | H | PHE | A | 294 | 17.914 | 29.059 | 19.489 | 1.00 | 0.00 | H |
| ATOM | 4172 | CA | PHE | A | 294 | 19.005 | 30.811 | 20.066 | 1.00 | 0.00 | C |
| ATOM | 4173 | HA | PHE | A | 294 | 19.050 | 31.645 | 19.516 | 1.00 | 0.00 | H |
| ATOM | 4174 | CB | PHE | A | 294 | 18.089 | 31.115 | 21.279 | 1.00 | 0.00 | C |
| ATOM | 4175 | HB1 | PHE | A | 294 | 18.529 | 31.828 | 21.825 | 1.00 | 0.00 | H |
| ATOM | 4176 | HB2 | PHE | A | 294 | 18.012 | 30.279 | 21.822 | 1.00 | 0.00 | H |
| ATOM | 4177 | CG | PHE | A | 294 | 16.652 | 31.595 | 20.932 | 1.00 | 0.00 | C |
| ATOM | 4178 | CD1 | PHE | A | 294 | 16.301 | 32.108 | 19.647 | 1.00 | 0.00 | C |
| ATOM | 4179 | HD1 | PHE | A | 294 | 17.002 | 32.231 | 18.944 | 1.00 | 0.00 | H |
| ATOM | 4180 | CE1 | PHE | A | 294 | 14.988 | 32.433 | 19.383 | 1.00 | 0.00 | C |
| ATOM | 4181 | HE1 | PHE | A | 294 | 14.726 | 32.705 | 18.457 | 1.00 | 0.00 | H |
| ATOM | 4182 | CZ | PHE | A | 294 | 14.034 | 32.389 | 20.370 | 1.00 | 0.00 | C |
| ATOM | 4183 | HZ | PHE | A | 294 | 13.080 | 32.601 | 20.160 | 1.00 | 0.00 | H |
| ATOM | 4184 | CE2 | PHE | A | 294 | 14.404 | 32.051 | 21.646 | 1.00 | 0.00 | C |
| ATOM | 4185 | HE2 | PHE | A | 294 | 13.708 | 32.062 | 22.364 | 1.00 | 0.00 | H |
| ATOM | 4186 | CD2 | PHE | A | 294 | 15.694 | 31.696 | 21.983 | 1.00 | 0.00 | C |
| ATOM | 4187 | HD2 | PHE | A | 294 | 15.949 | 31.516 | 22.933 | 1.00 | 0.00 | H |
| ATOM | 4188 | C | PHE | A | 294 | 20.353 | 30.444 | 20.525 | 1.00 | 0.00 | C |
| ATOM | 4189 | O | PHE | A | 294 | 21.302 | 31.202 | 20.310 | 1.00 | 0.00 | O |
| ATOM | 4190 | N | THR | A | 295 | 20.441 | 29.313 | 21.208 | 1.00 | 0.00 | N |
| ATOM | 4191 | H | THR | A | 295 | 19.686 | 28.657 | 21.193 | 1.00 | 0.00 | H |
| ATOM | 4192 | CA | THR | A | 295 | 21.687 | 29.017 | 22.011 | 1.00 | 0.00 | C |
| ATOM | 4193 | HA | THR | A | 295 | 21.929 | 29.840 | 22.524 | 1.00 | 0.00 | H |
| ATOM | 4194 | CB | THR | A | 295 | 21.443 | 27.877 | 23.021 | 1.00 | 0.00 | C |
| ATOM | 4195 | HB | THR | A | 295 | 21.338 | 27.054 | 22.463 | 1.00 | 0.00 | H |
| ATOM | 4196 | CG2 | THR | A | 295 | 22.561 | 27.719 | 24.051 | 1.00 | 0.00 | C |
| ATOM | 4197 | HG21 | THR | A | 295 | 22.339 | 26.966 | 24.671 | 1.00 | 0.00 | H |
| ATOM | 4198 | HG22 | THR | A | 295 | 23.421 | 27.521 | 23.581 | 1.00 | 0.00 | H |
| ATOM | 4199 | HG23 | THR | A | 295 | 22.654 | 28.566 | 24.575 | 1.00 | 0.00 | H |
| ATOM | 4200 | OG1 | THR | A | 295 | 20.225 | 28.027 | 23.769 | 1.00 | 0.00 | O |
| ATOM | 4201 | HG1 | THR | A | 295 | 20.126 | 27.260 | 24.403 | 1.00 | 0.00 | H |
| ATOM | 4202 | C | THR | A | 295 | 22.850 | 28.750 | 21.111 | 1.00 | 0.00 | C |
| ATOM | 4203 | O | THR | A | 295 | 23.951 | 29.156 | 21.484 | 1.00 | 0.00 | O |
| ATOM | 4204 | N | LYS | A | 296 | 22.713 | 28.134 | 19.933 | 1.00 | 0.00 | N |
| ATOM | 4205 | H | LYS | A | 296 | 21.776 | 28.027 | 19.600 | 1.00 | 0.00 | H |
| ATOM | 4206 | CA | LYS | A | 296 | 23.780 | 27.589 | 19.059 | 1.00 | 0.00 | C |
| ATOM | 4207 | HA | LYS | A | 296 | 23.232 | 27.046 | 18.422 | 1.00 | 0.00 | H |
| ATOM | 4208 | CB | LYS | A | 296 | 24.538 | 28.726 | 18.338 | 1.00 | 0.00 | C |
| ATOM | 4209 | HB1 | LYS | A | 296 | 25.174 | 28.309 | 17.688 | 1.00 | 0.00 | H |

| | | | | | | | | | | |
|---|---|---|---|---|---|---|---|---|---|---|
| ATOM | 4210 | HB2 | LYS | A | 296 | 25.052 | 29.239 | 19.026 | 1.00 | 0.00 | H |
| ATOM | 4211 | CG | LYS | A | 296 | 23.656 | 29.726 | 17.563 | 1.00 | 0.00 | C |
| ATOM | 4212 | HG1 | LYS | A | 296 | 24.249 | 30.414 | 17.144 | 1.00 | 0.00 | H |
| ATOM | 4213 | HG2 | LYS | A | 296 | 23.036 | 30.170 | 18.210 | 1.00 | 0.00 | H |
| ATOM | 4214 | CD | LYS | A | 296 | 22.814 | 29.071 | 16.455 | 1.00 | 0.00 | C |
| ATOM | 4215 | HD1 | LYS | A | 296 | 22.272 | 28.331 | 16.853 | 1.00 | 0.00 | H |
| ATOM | 4216 | HD2 | LYS | A | 296 | 23.427 | 28.698 | 15.758 | 1.00 | 0.00 | H |
| ATOM | 4217 | CE | LYS | A | 296 | 21.866 | 30.078 | 15.793 | 1.00 | 0.00 | C |
| ATOM | 4218 | HE1 | LYS | A | 296 | 21.425 | 30.629 | 16.502 | 1.00 | 0.00 | H |
| ATOM | 4219 | HE2 | LYS | A | 296 | 21.169 | 29.582 | 15.275 | 1.00 | 0.00 | H |
| ATOM | 4220 | NZ | LYS | A | 296 | 22.543 | 31.007 | 14.858 | 1.00 | 0.00 | N |
| ATOM | 4221 | HZ1 | LYS | A | 296 | 21.870 | 31.633 | 14.464 | 1.00 | 0.00 | H |
| ATOM | 4222 | HZ2 | LYS | A | 296 | 23.236 | 31.531 | 15.353 | 1.00 | 0.00 | H |
| ATOM | 4223 | HZ3 | LYS | A | 296 | 22.980 | 30.484 | 14.126 | 1.00 | 0.00 | H |
| ATOM | 4224 | C | LYS | A | 296 | 24.757 | 26.569 | 19.683 | 1.00 | 0.00 | C |
| ATOM | 4225 | O | LYS | A | 296 | 25.173 | 25.662 | 18.968 | 1.00 | 0.00 | O |
| ATOM | 4226 | N | GLU | A | 297 | 25.081 | 26.647 | 20.976 | 1.00 | 0.00 | N |
| ATOM | 4227 | H | GLU | A | 297 | 24.665 | 27.376 | 21.519 | 1.00 | 0.00 | H |
| ATOM | 4228 | CA | GLU | A | 297 | 26.012 | 25.728 | 21.655 | 1.00 | 0.00 | C |
| ATOM | 4229 | HA | GLU | A | 297 | 26.841 | 25.780 | 21.099 | 1.00 | 0.00 | H |
| ATOM | 4230 | CB | GLU | A | 297 | 26.310 | 26.185 | 23.096 | 1.00 | 0.00 | C |
| ATOM | 4231 | HB1 | GLU | A | 297 | 27.089 | 25.652 | 23.426 | 1.00 | 0.00 | H |
| ATOM | 4232 | HB2 | GLU | A | 297 | 25.504 | 25.977 | 23.651 | 1.00 | 0.00 | H |
| ATOM | 4233 | CG | GLU | A | 297 | 26.645 | 27.667 | 23.315 | 1.00 | 0.00 | C |
| ATOM | 4234 | HG1 | GLU | A | 297 | 26.944 | 27.782 | 24.262 | 1.00 | 0.00 | H |
| ATOM | 4235 | HG2 | GLU | A | 297 | 25.813 | 28.200 | 23.161 | 1.00 | 0.00 | H |
| ATOM | 4236 | CD | GLU | A | 297 | 27.740 | 28.225 | 22.402 | 1.00 | 0.00 | C |
| ATOM | 4237 | OE1 | GLU | A | 297 | 27.622 | 29.433 | 22.076 | 1.00 | 0.00 | O |
| ATOM | 4238 | OE2 | GLU | A | 297 | 28.698 | 27.488 | 22.086 | 1.00 | 0.00 | O |
| ATOM | 4239 | C | GLU | A | 297 | 25.473 | 24.279 | 21.679 | 1.00 | 0.00 | C |
| ATOM | 4240 | O | GLU | A | 297 | 24.441 | 23.994 | 22.287 | 1.00 | 0.00 | O |
| ATOM | 4241 | N | GLY | A | 298 | 26.193 | 23.342 | 21.051 | 1.00 | 0.00 | N |
| ATOM | 4242 | H | GLY | A | 298 | 27.165 | 23.520 | 20.898 | 1.00 | 0.00 | H |
| ATOM | 4243 | CA | GLY | A | 298 | 25.638 | 22.064 | 20.572 | 1.00 | 0.00 | C |
| ATOM | 4244 | HA1 | GLY | A | 298 | 26.420 | 21.498 | 20.311 | 1.00 | 0.00 | H |
| ATOM | 4245 | HA2 | GLY | A | 298 | 25.096 | 22.279 | 19.759 | 1.00 | 0.00 | H |
| ATOM | 4246 | C | GLY | A | 298 | 24.758 | 21.261 | 21.546 | 1.00 | 0.00 | C |
| ATOM | 4247 | O | GLY | A | 298 | 23.623 | 20.930 | 21.179 | 1.00 | 0.00 | O |
| ATOM | 4248 | N | GLY | A | 299 | 25.273 | 20.979 | 22.748 | 1.00 | 0.00 | N |
| ATOM | 4249 | H | GLY | A | 299 | 26.159 | 21.384 | 22.975 | 1.00 | 0.00 | H |
| ATOM | 4250 | CA | GLY | A | 299 | 24.643 | 20.123 | 23.762 | 1.00 | 0.00 | C |
| ATOM | 4251 | HA1 | GLY | A | 299 | 25.362 | 19.544 | 24.146 | 1.00 | 0.00 | H |
| ATOM | 4252 | HA2 | GLY | A | 299 | 23.972 | 19.552 | 23.289 | 1.00 | 0.00 | H |
| ATOM | 4253 | C | GLY | A | 299 | 23.946 | 20.863 | 24.899 | 1.00 | 0.00 | C |
| ATOM | 4254 | O | GLY | A | 299 | 23.951 | 20.363 | 26.029 | 1.00 | 0.00 | O |
| ATOM | 4255 | N | MET | A | 300 | 23.455 | 22.083 | 24.664 | 1.00 | 0.00 | N |
| ATOM | 4256 | H | MET | A | 300 | 23.569 | 22.477 | 23.752 | 1.00 | 0.00 | H |
| ATOM | 4257 | CA | MET | A | 300 | 22.756 | 22.870 | 25.685 | 1.00 | 0.00 | C |
| ATOM | 4258 | HA | MET | A | 300 | 22.240 | 22.180 | 26.192 | 1.00 | 0.00 | H |
| ATOM | 4259 | CB | MET | A | 300 | 23.763 | 23.573 | 26.604 | 1.00 | 0.00 | C |
| ATOM | 4260 | HB1 | MET | A | 300 | 24.310 | 22.876 | 27.067 | 1.00 | 0.00 | H |
| ATOM | 4261 | HB2 | MET | A | 300 | 23.256 | 24.104 | 27.283 | 1.00 | 0.00 | H |
| ATOM | 4262 | CG | MET | A | 300 | 24.712 | 24.515 | 25.879 | 1.00 | 0.00 | C |
| ATOM | 4263 | HG1 | MET | A | 300 | 24.125 | 25.154 | 25.381 | 1.00 | 0.00 | H |

```
ATOM   4264  HG2 MET A 300      25.210  23.943  25.227  1.00  0.00           H
ATOM   4265  SD  MET A 300      25.808  25.348  27.044  1.00  0.00           S
ATOM   4266  CE  MET A 300      24.734  26.713  27.567  1.00  0.00           C
ATOM   4267  HE1 MET A 300      25.217  27.281  28.234  1.00  0.00           H
ATOM   4268  HE2 MET A 300      24.489  27.268  26.772  1.00  0.00           H
ATOM   4269  HE3 MET A 300      23.904  26.343  27.984  1.00  0.00           H
ATOM   4270  C   MET A 300      21.719  23.849  25.130  1.00  0.00           C
ATOM   4271  O   MET A 300      21.728  24.191  23.952  1.00  0.00           O
ATOM   4272  N   LEU A 301      20.781  24.248  25.994  1.00  0.00           N
ATOM   4273  H   LEU A 301      20.863  23.942  26.942  1.00  0.00           H
ATOM   4274  CA  LEU A 301      19.644  25.100  25.652  1.00  0.00           C
ATOM   4275  HA  LEU A 301      19.925  25.550  24.804  1.00  0.00           H
ATOM   4276  CB  LEU A 301      18.360  24.249  25.458  1.00  0.00           C
ATOM   4277  HB1 LEU A 301      17.583  24.864  25.324  1.00  0.00           H
ATOM   4278  HB2 LEU A 301      18.211  23.698  26.279  1.00  0.00           H
ATOM   4279  CG  LEU A 301      18.463  23.303  24.236  1.00  0.00           C
ATOM   4280  HG  LEU A 301      19.320  22.798  24.337  1.00  0.00           H
ATOM   4281  CD1 LEU A 301      17.241  22.422  24.165  1.00  0.00           C
ATOM   4282 HD11 LEU A 301      17.314  21.814  23.374  1.00  0.00           H
ATOM   4283 HD12 LEU A 301      17.175  21.876  25.000  1.00  0.00           H
ATOM   4284 HD13 LEU A 301      16.424  22.992  24.073  1.00  0.00           H
ATOM   4285  CD2 LEU A 301      18.406  24.052  22.922  1.00  0.00           C
ATOM   4286 HD21 LEU A 301      18.476  23.403  22.165  1.00  0.00           H
ATOM   4287 HD22 LEU A 301      17.539  24.547  22.858  1.00  0.00           H
ATOM   4288 HD23 LEU A 301      19.165  24.701  22.875  1.00  0.00           H
ATOM   4289  C   LEU A 301      19.348  26.095  26.839  1.00  0.00           C
ATOM   4290  O   LEU A 301      19.127  25.702  28.004  1.00  0.00           O
ATOM   4291  N   ALA A 302      19.410  27.387  26.501  1.00  0.00           N
ATOM   4292  H   ALA A 302      19.644  27.651  25.565  1.00  0.00           H
ATOM   4293  CA  ALA A 302      19.124  28.451  27.543  1.00  0.00           C
ATOM   4294  HA  ALA A 302      19.828  28.253  28.225  1.00  0.00           H
ATOM   4295  CB  ALA A 302      19.244  29.859  26.918  1.00  0.00           C
ATOM   4296  HB1 ALA A 302      19.053  30.550  27.615  1.00  0.00           H
ATOM   4297  HB2 ALA A 302      20.170  29.989  26.564  1.00  0.00           H
ATOM   4298  HB3 ALA A 302      18.586  29.949  26.170  1.00  0.00           H
ATOM   4299  C   ALA A 302      17.694  28.395  28.219  1.00  0.00           C
ATOM   4300  O   ALA A 302      16.756  27.870  27.602  1.00  0.00           O
ATOM   4301  N   TYR A 303      17.609  28.896  29.389  1.00  0.00           N
ATOM   4302  H   TYR A 303      18.456  29.114  29.875  1.00  0.00           H
ATOM   4303  CA  TYR A 303      16.338  29.168  30.056  1.00  0.00           C
ATOM   4304  HA  TYR A 303      16.007  28.233  30.184  1.00  0.00           H
ATOM   4305  CB  TYR A 303      16.529  29.741  31.445  1.00  0.00           C
ATOM   4306  HB1 TYR A 303      16.991  30.623  31.352  1.00  0.00           H
ATOM   4307  HB2 TYR A 303      17.113  29.112  31.959  1.00  0.00           H
ATOM   4308  CG  TYR A 303      15.221  29.951  32.231  1.00  0.00           C
ATOM   4309  CD1 TYR A 303      14.625  28.811  32.795  1.00  0.00           C
ATOM   4310  HD1 TYR A 303      15.094  27.935  32.684  1.00  0.00           H
ATOM   4311  CE1 TYR A 303      13.415  28.823  33.504  1.00  0.00           C
ATOM   4312  HE1 TYR A 303      12.981  27.987  33.841  1.00  0.00           H
ATOM   4313  CZ  TYR A 303      12.852  30.112  33.708  1.00  0.00           C
ATOM   4314  OH  TYR A 303      11.794  30.165  34.499  1.00  0.00           O
ATOM   4315  HH  TYR A 303      11.480  31.112  34.573  1.00  0.00           H
ATOM   4316  CE2 TYR A 303      13.505  31.291  33.311  1.00  0.00           C
ATOM   4317  HE2 TYR A 303      13.141  32.176  33.601  1.00  0.00           H
```

```
ATOM   4318  CD2 TYR A 303      14.652  31.247  32.520  1.00  0.00           C
ATOM   4319  HD2 TYR A 303      15.067  32.084  32.165  1.00  0.00           H
ATOM   4320  C   TYR A 303      15.243  29.902  29.192  1.00  0.00           C
ATOM   4321  O   TYR A 303      14.143  29.428  28.972  1.00  0.00           O
ATOM   4322  N   TYR A 304      15.643  30.987  28.517  1.00  0.00           N
ATOM   4323  H   TYR A 304      16.551  31.357  28.713  1.00  0.00           H
ATOM   4324  CA  TYR A 304      14.816  31.668  27.498  1.00  0.00           C
ATOM   4325  HA  TYR A 304      13.923  31.778  27.935  1.00  0.00           H
ATOM   4326  CB  TYR A 304      15.475  33.059  27.134  1.00  0.00           C
ATOM   4327  HB1 TYR A 304      15.915  33.394  27.967  1.00  0.00           H
ATOM   4328  HB2 TYR A 304      14.727  33.673  26.882  1.00  0.00           H
ATOM   4329  CG  TYR A 304      16.543  33.188  25.998  1.00  0.00           C
ATOM   4330  CD1 TYR A 304      16.166  33.511  24.662  1.00  0.00           C
ATOM   4331  HD1 TYR A 304      15.203  33.504  24.393  1.00  0.00           H
ATOM   4332  CE1 TYR A 304      17.210  33.856  23.681  1.00  0.00           C
ATOM   4333  HE1 TYR A 304      16.971  34.212  22.778  1.00  0.00           H
ATOM   4334  CZ  TYR A 304      18.550  33.669  24.054  1.00  0.00           C
ATOM   4335  OH  TYR A 304      19.650  33.933  23.319  1.00  0.00           O
ATOM   4336  HH  TYR A 304      20.467  33.699  23.846  1.00  0.00           H
ATOM   4337  CE2 TYR A 304      18.861  33.314  25.416  1.00  0.00           C
ATOM   4338  HE2 TYR A 304      19.816  33.272  25.710  1.00  0.00           H
ATOM   4339  CD2 TYR A 304      17.857  33.032  26.319  1.00  0.00           C
ATOM   4340  HD2 TYR A 304      18.100  32.705  27.232  1.00  0.00           H
ATOM   4341  C   TYR A 304      14.492  30.815  26.274  1.00  0.00           C
ATOM   4342  O   TYR A 304      13.447  30.974  25.647  1.00  0.00           O
ATOM   4343  N   GLU A 305      15.316  29.856  25.885  1.00  0.00           N
ATOM   4344  H   GLU A 305      16.172  29.745  26.389  1.00  0.00           H
ATOM   4345  CA  GLU A 305      15.054  28.925  24.741  1.00  0.00           C
ATOM   4346  HA  GLU A 305      14.618  29.502  24.051  1.00  0.00           H
ATOM   4347  CB  GLU A 305      16.376  28.325  24.205  1.00  0.00           C
ATOM   4348  HB1 GLU A 305      16.801  27.771  24.921  1.00  0.00           H
ATOM   4349  HB2 GLU A 305      16.995  29.067  23.946  1.00  0.00           H
ATOM   4350  CG  GLU A 305      16.108  27.439  22.973  1.00  0.00           C
ATOM   4351  HG1 GLU A 305      15.195  27.663  22.633  1.00  0.00           H
ATOM   4352  HG2 GLU A 305      16.127  26.486  23.276  1.00  0.00           H
ATOM   4353  CD  GLU A 305      17.129  27.604  21.795  1.00  0.00           C
ATOM   4354  OE1 GLU A 305      18.385  27.506  21.942  1.00  0.00           O
ATOM   4355  OE2 GLU A 305      16.698  27.815  20.647  1.00  0.00           O
ATOM   4356  C   GLU A 305      14.079  27.788  25.097  1.00  0.00           C
ATOM   4357  O   GLU A 305      13.235  27.537  24.193  1.00  0.00           O
ATOM   4358  N   VAL A 306      14.117  27.199  26.270  1.00  0.00           N
ATOM   4359  H   VAL A 306      14.849  27.432  26.911  1.00  0.00           H
ATOM   4360  CA  VAL A 306      13.066  26.163  26.679  1.00  0.00           C
ATOM   4361  HA  VAL A 306      12.902  25.570  25.891  1.00  0.00           H
ATOM   4362  CB  VAL A 306      13.551  25.238  27.803  1.00  0.00           C
ATOM   4363  HB  VAL A 306      12.783  24.656  28.069  1.00  0.00           H
ATOM   4364  CG1 VAL A 306      14.568  24.288  27.358  1.00  0.00           C
ATOM   4365 HG11 VAL A 306      14.846  23.713  28.127  1.00  0.00           H
ATOM   4366 HG12 VAL A 306      14.192  23.713  26.631  1.00  0.00           H
ATOM   4367 HG13 VAL A 306      15.362  24.788  27.011  1.00  0.00           H
ATOM   4368  CG2 VAL A 306      14.026  26.026  29.024  1.00  0.00           C
ATOM   4369 HG21 VAL A 306      14.333  25.390  29.732  1.00  0.00           H
ATOM   4370 HG22 VAL A 306      14.784  26.623  28.761  1.00  0.00           H
ATOM   4371 HG23 VAL A 306      13.272  26.580  29.377  1.00  0.00           H
```

| | | | | | | | | | | | |
|---|---|---|---|---|---|---|---|---|---|---|---|
| ATOM | 4372 | C | VAL | A | 306 | 11.738 | 26.803 | 26.962 | 1.00 | 0.00 | C |
| ATOM | 4373 | O | VAL | A | 306 | 10.669 | 26.282 | 26.784 | 1.00 | 0.00 | O |
| ATOM | 4374 | N | CYS | A | 307 | 11.809 | 28.047 | 27.449 | 1.00 | 0.00 | N |
| ATOM | 4375 | H | CYS | A | 307 | 12.710 | 28.411 | 27.685 | 1.00 | 0.00 | H |
| ATOM | 4376 | CA | CYS | A | 307 | 10.614 | 28.925 | 27.661 | 1.00 | 0.00 | C |
| ATOM | 4377 | HA | CYS | A | 307 | 10.018 | 28.483 | 28.331 | 1.00 | 0.00 | H |
| ATOM | 4378 | CB | CYS | A | 307 | 10.966 | 30.284 | 28.241 | 1.00 | 0.00 | C |
| ATOM | 4379 | HB1 | CYS | A | 307 | 10.165 | 30.829 | 27.991 | 1.00 | 0.00 | H |
| ATOM | 4380 | HB2 | CYS | A | 307 | 11.744 | 30.555 | 27.674 | 1.00 | 0.00 | H |
| ATOM | 4381 | SG | CYS | A | 307 | 11.352 | 30.429 | 30.037 | 1.00 | 0.00 | S |
| ATOM | 4382 | C | CYS | A | 307 | 9.909 | 29.068 | 26.347 | 1.00 | 0.00 | C |
| ATOM | 4383 | O | CYS | A | 307 | 8.733 | 28.807 | 26.347 | 1.00 | 0.00 | O |
| ATOM | 4384 | N | SER | A | 308 | 10.702 | 29.259 | 25.242 | 1.00 | 0.00 | N |
| ATOM | 4385 | H | SER | A | 308 | 11.696 | 29.255 | 25.353 | 1.00 | 0.00 | H |
| ATOM | 4386 | CA | SER | A | 308 | 10.132 | 29.471 | 23.889 | 1.00 | 0.00 | C |
| ATOM | 4387 | HA | SER | A | 308 | 9.307 | 30.029 | 23.981 | 1.00 | 0.00 | H |
| ATOM | 4388 | CB | SER | A | 308 | 11.263 | 30.147 | 23.067 | 1.00 | 0.00 | C |
| ATOM | 4389 | HB1 | SER | A | 308 | 11.003 | 30.262 | 22.108 | 1.00 | 0.00 | H |
| ATOM | 4390 | HB2 | SER | A | 308 | 12.117 | 29.630 | 23.121 | 1.00 | 0.00 | H |
| ATOM | 4391 | OG | SER | A | 308 | 11.532 | 31.406 | 23.557 | 1.00 | 0.00 | O |
| ATOM | 4392 | HG | SER | A | 308 | 12.261 | 31.825 | 23.016 | 1.00 | 0.00 | H |
| ATOM | 4393 | C | SER | A | 308 | 9.722 | 28.134 | 23.214 | 1.00 | 0.00 | C |
| ATOM | 4394 | O | SER | A | 308 | 9.015 | 28.205 | 22.181 | 1.00 | 0.00 | O |
| ATOM | 4395 | N | TRP | A | 309 | 9.915 | 26.938 | 23.831 | 1.00 | 0.00 | N |
| ATOM | 4396 | H | TRP | A | 309 | 10.480 | 26.961 | 24.655 | 1.00 | 0.00 | H |
| ATOM | 4397 | CA | TRP | A | 309 | 9.402 | 25.615 | 23.446 | 1.00 | 0.00 | C |
| ATOM | 4398 | HA | TRP | A | 309 | 9.444 | 25.597 | 22.447 | 1.00 | 0.00 | H |
| ATOM | 4399 | CB | TRP | A | 309 | 10.298 | 24.402 | 23.980 | 1.00 | 0.00 | C |
| ATOM | 4400 | HB1 | TRP | A | 309 | 9.786 | 23.549 | 23.875 | 1.00 | 0.00 | H |
| ATOM | 4401 | HB2 | TRP | A | 309 | 10.503 | 24.551 | 24.947 | 1.00 | 0.00 | H |
| ATOM | 4402 | CG | TRP | A | 309 | 11.599 | 24.118 | 23.375 | 1.00 | 0.00 | C |
| ATOM | 4403 | CD1 | TRP | A | 309 | 12.437 | 24.951 | 22.794 | 1.00 | 0.00 | C |
| ATOM | 4404 | HD1 | TRP | A | 309 | 12.312 | 25.938 | 22.696 | 1.00 | 0.00 | H |
| ATOM | 4405 | NE1 | TRP | A | 309 | 13.546 | 24.202 | 22.332 | 1.00 | 0.00 | N |
| ATOM | 4406 | HE1 | TRP | A | 309 | 14.350 | 24.564 | 21.861 | 1.00 | 0.00 | H |
| ATOM | 4407 | CE2 | TRP | A | 309 | 13.306 | 22.849 | 22.660 | 1.00 | 0.00 | C |
| ATOM | 4408 | CZ2 | TRP | A | 309 | 14.160 | 21.761 | 22.547 | 1.00 | 0.00 | C |
| ATOM | 4409 | HZ2 | TRP | A | 309 | 15.027 | 21.851 | 22.057 | 1.00 | 0.00 | H |
| ATOM | 4410 | CH2 | TRP | A | 309 | 13.797 | 20.519 | 23.131 | 1.00 | 0.00 | C |
| ATOM | 4411 | HH2 | TRP | A | 309 | 14.368 | 19.712 | 22.979 | 1.00 | 0.00 | H |
| ATOM | 4412 | CZ3 | TRP | A | 309 | 12.640 | 20.419 | 23.921 | 1.00 | 0.00 | C |
| ATOM | 4413 | HZ3 | TRP | A | 309 | 12.401 | 19.560 | 24.374 | 1.00 | 0.00 | H |
| ATOM | 4414 | CE3 | TRP | A | 309 | 11.828 | 21.556 | 24.058 | 1.00 | 0.00 | C |
| ATOM | 4415 | HE3 | TRP | A | 309 | 10.974 | 21.485 | 24.574 | 1.00 | 0.00 | H |
| ATOM | 4416 | CD2 | TRP | A | 309 | 12.192 | 22.819 | 23.476 | 1.00 | 0.00 | C |
| ATOM | 4417 | C | TRP | A | 309 | 7.931 | 25.531 | 24.000 | 1.00 | 0.00 | C |
| ATOM | 4418 | O | TRP | A | 309 | 7.678 | 24.927 | 25.048 | 1.00 | 0.00 | O |
| ATOM | 4419 | N | LYS | A | 310 | 7.030 | 26.275 | 23.363 | 1.00 | 0.00 | N |
| ATOM | 4420 | H | LYS | A | 310 | 7.343 | 26.771 | 22.553 | 1.00 | 0.00 | H |
| ATOM | 4421 | CA | LYS | A | 310 | 5.633 | 26.434 | 23.735 | 1.00 | 0.00 | C |
| ATOM | 4422 | HA | LYS | A | 310 | 5.650 | 26.697 | 24.700 | 1.00 | 0.00 | H |
| ATOM | 4423 | CB | LYS | A | 310 | 5.014 | 27.641 | 22.980 | 1.00 | 0.00 | C |
| ATOM | 4424 | HB1 | LYS | A | 310 | 4.924 | 27.420 | 22.009 | 1.00 | 0.00 | H |
| ATOM | 4425 | HB2 | LYS | A | 310 | 5.603 | 28.442 | 23.086 | 1.00 | 0.00 | H |

| | | | | | | | | | | | |
|---|---|---|---|---|---|---|---|---|---|---|---|
| ATOM | 4426 | CG | LYS | A | 310 | 3.623 | 27.950 | 23.564 | 1.00 | 0.00 | C |
| ATOM | 4427 | HG1 | LYS | A | 310 | 3.675 | 27.891 | 24.561 | 1.00 | 0.00 | H |
| ATOM | 4428 | HG2 | LYS | A | 310 | 2.975 | 27.269 | 23.223 | 1.00 | 0.00 | H |
| ATOM | 4429 | CD | LYS | A | 310 | 3.114 | 29.353 | 23.179 | 1.00 | 0.00 | C |
| ATOM | 4430 | HD1 | LYS | A | 310 | 2.243 | 29.479 | 23.653 | 1.00 | 0.00 | H |
| ATOM | 4431 | HD2 | LYS | A | 310 | 2.957 | 29.337 | 22.192 | 1.00 | 0.00 | H |
| ATOM | 4432 | CE | LYS | A | 310 | 4.023 | 30.605 | 23.501 | 1.00 | 0.00 | C |
| ATOM | 4433 | HE1 | LYS | A | 310 | 3.562 | 31.446 | 23.219 | 1.00 | 0.00 | H |
| ATOM | 4434 | HE2 | LYS | A | 310 | 4.897 | 30.528 | 23.022 | 1.00 | 0.00 | H |
| ATOM | 4435 | NZ | LYS | A | 310 | 4.344 | 30.770 | 24.905 | 1.00 | 0.00 | N |
| ATOM | 4436 | HZ1 | LYS | A | 310 | 4.919 | 31.580 | 25.023 | 1.00 | 0.00 | H |
| ATOM | 4437 | HZ2 | LYS | A | 310 | 3.500 | 30.881 | 25.429 | 1.00 | 0.00 | H |
| ATOM | 4438 | HZ3 | LYS | A | 310 | 4.835 | 29.963 | 25.232 | 1.00 | 0.00 | H |
| ATOM | 4439 | C | LYS | A | 310 | 4.835 | 25.074 | 23.717 | 1.00 | 0.00 | C |
| ATOM | 4440 | O | LYS | A | 310 | 5.097 | 24.238 | 22.890 | 1.00 | 0.00 | O |
| ATOM | 4441 | N | GLY | A | 311 | 3.854 | 24.867 | 24.684 | 1.00 | 0.00 | N |
| ATOM | 4442 | H | GLY | A | 311 | 3.618 | 25.610 | 25.311 | 1.00 | 0.00 | H |
| ATOM | 4443 | CA | GLY | A | 311 | 3.128 | 23.518 | 24.805 | 1.00 | 0.00 | C |
| ATOM | 4444 | HA1 | GLY | A | 311 | 2.715 | 23.314 | 23.917 | 1.00 | 0.00 | H |
| ATOM | 4445 | HA2 | GLY | A | 311 | 2.408 | 23.621 | 25.492 | 1.00 | 0.00 | H |
| ATOM | 4446 | C | GLY | A | 311 | 4.012 | 22.313 | 25.212 | 1.00 | 0.00 | C |
| ATOM | 4447 | O | GLY | A | 311 | 3.543 | 21.138 | 25.174 | 1.00 | 0.00 | O |
| ATOM | 4448 | N | ALA | A | 312 | 5.314 | 22.503 | 25.498 | 1.00 | 0.00 | N |
| ATOM | 4449 | H | ALA | A | 312 | 5.733 | 23.405 | 25.391 | 1.00 | 0.00 | H |
| ATOM | 4450 | CA | ALA | A | 312 | 6.076 | 21.403 | 25.952 | 1.00 | 0.00 | C |
| ATOM | 4451 | HA | ALA | A | 312 | 5.867 | 20.775 | 25.202 | 1.00 | 0.00 | H |
| ATOM | 4452 | CB | ALA | A | 312 | 7.514 | 21.765 | 25.949 | 1.00 | 0.00 | C |
| ATOM | 4453 | HB1 | ALA | A | 312 | 8.054 | 20.987 | 26.270 | 1.00 | 0.00 | H |
| ATOM | 4454 | HB2 | ALA | A | 312 | 7.795 | 22.005 | 25.020 | 1.00 | 0.00 | H |
| ATOM | 4455 | HB3 | ALA | A | 312 | 7.662 | 22.547 | 26.555 | 1.00 | 0.00 | H |
| ATOM | 4456 | C | ALA | A | 312 | 5.672 | 20.758 | 27.385 | 1.00 | 0.00 | C |
| ATOM | 4457 | O | ALA | A | 312 | 5.218 | 21.409 | 28.316 | 1.00 | 0.00 | O |
| ATOM | 4458 | N | THR | A | 313 | 5.930 | 19.450 | 27.481 | 1.00 | 0.00 | N |
| ATOM | 4459 | H | THR | A | 313 | 6.307 | 19.031 | 26.655 | 1.00 | 0.00 | H |
| ATOM | 4460 | CA | THR | A | 313 | 5.741 | 18.516 | 28.637 | 1.00 | 0.00 | C |
| ATOM | 4461 | HA | THR | A | 313 | 4.948 | 18.831 | 29.158 | 1.00 | 0.00 | H |
| ATOM | 4462 | CB | THR | A | 313 | 5.478 | 17.090 | 28.071 | 1.00 | 0.00 | C |
| ATOM | 4463 | HB | THR | A | 313 | 6.305 | 16.751 | 27.623 | 1.00 | 0.00 | H |
| ATOM | 4464 | CG2 | THR | A | 313 | 4.914 | 16.213 | 29.191 | 1.00 | 0.00 | C |
| ATOM | 4465 | HG21 | THR | A | 313 | 4.741 | 15.293 | 28.839 | 1.00 | 0.00 | H |
| ATOM | 4466 | HG22 | THR | A | 313 | 5.574 | 16.164 | 29.941 | 1.00 | 0.00 | H |
| ATOM | 4467 | HG23 | THR | A | 313 | 4.059 | 16.609 | 29.525 | 1.00 | 0.00 | H |
| ATOM | 4468 | OG1 | THR | A | 313 | 4.529 | 17.053 | 27.062 | 1.00 | 0.00 | O |
| ATOM | 4469 | HG1 | THR | A | 313 | 4.412 | 16.111 | 26.748 | 1.00 | 0.00 | H |
| ATOM | 4470 | C | THR | A | 313 | 6.883 | 18.584 | 29.658 | 1.00 | 0.00 | C |
| ATOM | 4471 | O | THR | A | 313 | 7.907 | 17.951 | 29.505 | 1.00 | 0.00 | O |
| ATOM | 4472 | N | LYS | A | 314 | 6.701 | 19.383 | 30.711 | 1.00 | 0.00 | N |
| ATOM | 4473 | H | LYS | A | 314 | 5.847 | 19.904 | 30.726 | 1.00 | 0.00 | H |
| ATOM | 4474 | CA | LYS | A | 314 | 7.612 | 19.591 | 31.860 | 1.00 | 0.00 | C |
| ATOM | 4475 | HA | LYS | A | 314 | 8.550 | 19.405 | 31.569 | 1.00 | 0.00 | H |
| ATOM | 4476 | CB | LYS | A | 314 | 7.520 | 21.053 | 32.255 | 1.00 | 0.00 | C |
| ATOM | 4477 | HB1 | LYS | A | 314 | 6.598 | 21.222 | 32.603 | 1.00 | 0.00 | H |
| ATOM | 4478 | HB2 | LYS | A | 314 | 7.673 | 21.605 | 31.435 | 1.00 | 0.00 | H |
| ATOM | 4479 | CG | LYS | A | 314 | 8.541 | 21.471 | 33.332 | 1.00 | 0.00 | C |

| | | | | | | | | | | | |
|---|---|---|---|---|---|---|---|---|---|---|---|
| ATOM | 4480 | HG1 | LYS | A | 314 | 9.468 | 21.392 | 32.965 | 1.00 | 0.00 | H |
| ATOM | 4481 | HG2 | LYS | A | 314 | 8.447 | 20.877 | 34.131 | 1.00 | 0.00 | H |
| ATOM | 4482 | CD | LYS | A | 314 | 8.256 | 22.944 | 33.732 | 1.00 | 0.00 | C |
| ATOM | 4483 | HD1 | LYS | A | 314 | 8.165 | 23.511 | 32.913 | 1.00 | 0.00 | H |
| ATOM | 4484 | HD2 | LYS | A | 314 | 9.000 | 23.296 | 34.300 | 1.00 | 0.00 | H |
| ATOM | 4485 | CE | LYS | A | 314 | 7.009 | 23.065 | 34.501 | 1.00 | 0.00 | C |
| ATOM | 4486 | HE1 | LYS | A | 314 | 7.143 | 22.597 | 35.374 | 1.00 | 0.00 | H |
| ATOM | 4487 | HE2 | LYS | A | 314 | 6.286 | 22.606 | 33.984 | 1.00 | 0.00 | H |
| ATOM | 4488 | NZ | LYS | A | 314 | 6.616 | 24.531 | 34.753 | 1.00 | 0.00 | N |
| ATOM | 4489 | HZ1 | LYS | A | 314 | 5.765 | 24.563 | 35.277 | 1.00 | 0.00 | H |
| ATOM | 4490 | HZ2 | LYS | A | 314 | 7.342 | 24.987 | 35.268 | 1.00 | 0.00 | H |
| ATOM | 4491 | HZ3 | LYS | A | 314 | 6.485 | 24.996 | 33.878 | 1.00 | 0.00 | H |
| ATOM | 4492 | C | LYS | A | 314 | 7.292 | 18.600 | 33.025 | 1.00 | 0.00 | C |
| ATOM | 4493 | O | LYS | A | 314 | 6.130 | 18.444 | 33.342 | 1.00 | 0.00 | O |
| ATOM | 4494 | N | GLN | A | 315 | 8.314 | 18.101 | 33.747 | 1.00 | 0.00 | N |
| ATOM | 4495 | H | GLN | A | 315 | 9.242 | 18.335 | 33.456 | 1.00 | 0.00 | H |
| ATOM | 4496 | CA | GLN | A | 315 | 8.173 | 17.254 | 34.910 | 1.00 | 0.00 | C |
| ATOM | 4497 | HA | GLN | A | 315 | 7.258 | 17.400 | 35.287 | 1.00 | 0.00 | H |
| ATOM | 4498 | CB | GLN | A | 315 | 8.199 | 15.782 | 34.602 | 1.00 | 0.00 | C |
| ATOM | 4499 | HB1 | GLN | A | 315 | 8.113 | 15.300 | 35.474 | 1.00 | 0.00 | H |
| ATOM | 4500 | HB2 | GLN | A | 315 | 9.092 | 15.584 | 34.198 | 1.00 | 0.00 | H |
| ATOM | 4501 | CG | GLN | A | 315 | 7.135 | 15.227 | 33.662 | 1.00 | 0.00 | C |
| ATOM | 4502 | HG1 | GLN | A | 315 | 7.478 | 14.366 | 33.285 | 1.00 | 0.00 | H |
| ATOM | 4503 | HG2 | GLN | A | 315 | 7.006 | 15.887 | 32.922 | 1.00 | 0.00 | H |
| ATOM | 4504 | CD | GLN | A | 315 | 5.777 | 14.948 | 34.259 | 1.00 | 0.00 | C |
| ATOM | 4505 | OE1 | GLN | A | 315 | 5.381 | 15.538 | 35.295 | 1.00 | 0.00 | O |
| ATOM | 4506 | NE2 | GLN | A | 315 | 4.967 | 14.073 | 33.647 | 1.00 | 0.00 | N |
| ATOM | 4507 | HE21 | GLN | A | 315 | 5.266 | 13.617 | 32.809 | 1.00 | 0.00 | H |
| ATOM | 4508 | HE22 | GLN | A | 315 | 4.064 | 13.876 | 34.029 | 1.00 | 0.00 | H |
| ATOM | 4509 | C | GLN | A | 315 | 9.272 | 17.677 | 35.859 | 1.00 | 0.00 | C |
| ATOM | 4510 | O | GLN | A | 315 | 10.258 | 18.262 | 35.492 | 1.00 | 0.00 | O |
| ATOM | 4511 | N | ARG | A | 316 | 9.156 | 17.301 | 37.216 | 1.00 | 0.00 | N |
| ATOM | 4512 | H | ARG | A | 316 | 8.240 | 17.225 | 37.611 | 1.00 | 0.00 | H |
| ATOM | 4513 | CA | ARG | A | 316 | 10.350 | 17.007 | 38.104 | 1.00 | 0.00 | C |
| ATOM | 4514 | HA | ARG | A | 316 | 10.994 | 17.744 | 37.897 | 1.00 | 0.00 | H |
| ATOM | 4515 | CB | ARG | A | 316 | 9.951 | 17.083 | 39.571 | 1.00 | 0.00 | C |
| ATOM | 4516 | HB1 | ARG | A | 316 | 10.379 | 16.312 | 40.044 | 1.00 | 0.00 | H |
| ATOM | 4517 | HB2 | ARG | A | 316 | 8.956 | 16.995 | 39.621 | 1.00 | 0.00 | H |
| ATOM | 4518 | CG | ARG | A | 316 | 10.382 | 18.416 | 40.273 | 1.00 | 0.00 | C |
| ATOM | 4519 | HG1 | ARG | A | 316 | 11.381 | 18.405 | 40.325 | 1.00 | 0.00 | H |
| ATOM | 4520 | HG2 | ARG | A | 316 | 10.000 | 18.393 | 41.197 | 1.00 | 0.00 | H |
| ATOM | 4521 | CD | ARG | A | 316 | 9.941 | 19.797 | 39.595 | 1.00 | 0.00 | C |
| ATOM | 4522 | HD1 | ARG | A | 316 | 8.969 | 19.952 | 39.774 | 1.00 | 0.00 | H |
| ATOM | 4523 | HD2 | ARG | A | 316 | 10.093 | 19.739 | 38.608 | 1.00 | 0.00 | H |
| ATOM | 4524 | NE | ARG | A | 316 | 10.664 | 21.040 | 40.065 | 1.00 | 0.00 | N |
| ATOM | 4525 | HE | ARG | A | 316 | 11.129 | 20.922 | 40.942 | 1.00 | 0.00 | H |
| ATOM | 4526 | CZ | ARG | A | 316 | 10.804 | 22.226 | 39.564 | 1.00 | 0.00 | C |
| ATOM | 4527 | NH1 | ARG | A | 316 | 10.181 | 22.572 | 38.558 | 1.00 | 0.00 | N |
| ATOM | 4528 | HH11 | ARG | A | 316 | 9.554 | 21.932 | 38.114 | 1.00 | 0.00 | H |
| ATOM | 4529 | HH12 | ARG | A | 316 | 10.303 | 23.492 | 38.186 | 1.00 | 0.00 | H |
| ATOM | 4530 | NH2 | ARG | A | 316 | 11.689 | 23.031 | 39.992 | 1.00 | 0.00 | N |
| ATOM | 4531 | HH21 | ARG | A | 316 | 12.300 | 22.754 | 40.734 | 1.00 | 0.00 | H |
| ATOM | 4532 | HH22 | ARG | A | 316 | 11.775 | 23.941 | 39.587 | 1.00 | 0.00 | H |
| ATOM | 4533 | C | ARG | A | 316 | 11.053 | 15.618 | 37.811 | 1.00 | 0.00 | C |

```
ATOM   4534  O   ARG A 316      10.404  14.637  37.409  1.00  0.00           O
ATOM   4535  N   ILE A 317      12.405  15.591  37.880  1.00  0.00           N
ATOM   4536  H   ILE A 317      12.963  16.346  37.536  1.00  0.00           H
ATOM   4537  CA  ILE A 317      12.992  14.436  38.465  1.00  0.00           C
ATOM   4538  HA  ILE A 317      12.505  13.675  38.037  1.00  0.00           H
ATOM   4539  CB  ILE A 317      14.441  14.270  38.122  1.00  0.00           C
ATOM   4540  HB  ILE A 317      14.885  15.042  38.577  1.00  0.00           H
ATOM   4541  CG2 ILE A 317      14.969  12.925  38.673  1.00  0.00           C
ATOM   4542 HG21 ILE A 317      15.936  12.825  38.438  1.00  0.00           H
ATOM   4543 HG22 ILE A 317      14.864  12.908  39.667  1.00  0.00           H
ATOM   4544 HG23 ILE A 317      14.449  12.172  38.270  1.00  0.00           H
ATOM   4545  CG1 ILE A 317      14.770  14.420  36.622  1.00  0.00           C
ATOM   4546 HG11 ILE A 317      14.250  15.191  36.253  1.00  0.00           H
ATOM   4547 HG12 ILE A 317      14.504  13.581  36.148  1.00  0.00           H
ATOM   4548  CD  ILE A 317      16.255  14.671  36.362  1.00  0.00           C
ATOM   4549  HD1 ILE A 317      16.411  14.760  35.378  1.00  0.00           H
ATOM   4550  HD2 ILE A 317      16.536  15.514  36.822  1.00  0.00           H
ATOM   4551  HD3 ILE A 317      16.790  13.904  36.716  1.00  0.00           H
ATOM   4552  C   ILE A 317      12.785  14.471  39.969  1.00  0.00           C
ATOM   4553  O   ILE A 317      13.316  15.410  40.634  1.00  0.00           O
ATOM   4554  N   GLN A 318      11.926  13.650  40.529  1.00  0.00           N
ATOM   4555  H   GLN A 318      11.479  12.967  39.951  1.00  0.00           H
ATOM   4556  CA  GLN A 318      11.578  13.681  41.991  1.00  0.00           C
ATOM   4557  HA  GLN A 318      11.289  14.626  42.142  1.00  0.00           H
ATOM   4558  CB  GLN A 318      10.393  12.689  42.284  1.00  0.00           C
ATOM   4559  HB1 GLN A 318      10.229  12.674  43.270  1.00  0.00           H
ATOM   4560  HB2 GLN A 318      10.661  11.776  41.977  1.00  0.00           H
ATOM   4561  CG  GLN A 318       9.048  13.120  41.535  1.00  0.00           C
ATOM   4562  HG1 GLN A 318       9.243  13.256  40.564  1.00  0.00           H
ATOM   4563  HG2 GLN A 318       8.712  13.975  41.931  1.00  0.00           H
ATOM   4564  CD  GLN A 318       7.906  12.118  41.618  1.00  0.00           C
ATOM   4565  OE1 GLN A 318       7.802  11.384  42.589  1.00  0.00           O
ATOM   4566  NE2 GLN A 318       6.995  12.123  40.679  1.00  0.00           N
ATOM   4567 HE21 GLN A 318       7.057  12.777  39.925  1.00  0.00           H
ATOM   4568 HE22 GLN A 318       6.236  11.473  40.715  1.00  0.00           H
ATOM   4569  C   GLN A 318      12.809  13.415  42.954  1.00  0.00           C
ATOM   4570  O   GLN A 318      12.956  14.195  43.847  1.00  0.00           O
ATOM   4571  N   ASP A 319      13.604  12.358  42.717  1.00  0.00           N
ATOM   4572  H   ASP A 319      13.464  11.807  41.894  1.00  0.00           H
ATOM   4573  CA  ASP A 319      14.709  11.993  43.683  1.00  0.00           C
ATOM   4574  HA  ASP A 319      14.252  11.939  44.571  1.00  0.00           H
ATOM   4575  CB  ASP A 319      15.356  10.648  43.183  1.00  0.00           C
ATOM   4576  HB1 ASP A 319      16.141  10.474  43.777  1.00  0.00           H
ATOM   4577  HB2 ASP A 319      15.675  10.817  42.251  1.00  0.00           H
ATOM   4578  CG  ASP A 319      14.628   9.309  43.104  1.00  0.00           C
ATOM   4579  OD1 ASP A 319      13.697   9.096  43.933  1.00  0.00           O
ATOM   4580  OD2 ASP A 319      14.993   8.465  42.237  1.00  0.00           O
ATOM   4581  C   ASP A 319      15.890  13.058  43.746  1.00  0.00           C
ATOM   4582  O   ASP A 319      16.748  13.011  44.649  1.00  0.00           O
ATOM   4583  N   GLN A 320      15.849  13.920  42.702  1.00  0.00           N
ATOM   4584  H   GLN A 320      15.073  13.892  42.072  1.00  0.00           H
ATOM   4585  CA  GLN A 320      16.926  14.897  42.479  1.00  0.00           C
ATOM   4586  HA  GLN A 320      17.535  14.798  43.266  1.00  0.00           H
ATOM   4587  CB  GLN A 320      17.609  14.505  41.167  1.00  0.00           C
```

| | | | | | | | | | | | |
|---|---|---|---|---|---|---|---|---|---|---|---|
| ATOM | 4588 | HB1 | GLN | A | 320 | 18.372 | 15.132 | 41.008 | 1.00 | 0.00 | H |
| ATOM | 4589 | HB2 | GLN | A | 320 | 16.944 | 14.595 | 40.426 | 1.00 | 0.00 | H |
| ATOM | 4590 | CG | GLN | A | 320 | 18.151 | 13.078 | 41.154 | 1.00 | 0.00 | C |
| ATOM | 4591 | HG1 | GLN | A | 320 | 17.401 | 12.437 | 40.992 | 1.00 | 0.00 | H |
| ATOM | 4592 | HG2 | GLN | A | 320 | 18.579 | 12.874 | 42.035 | 1.00 | 0.00 | H |
| ATOM | 4593 | CD | GLN | A | 320 | 19.176 | 12.919 | 40.063 | 1.00 | 0.00 | C |
| ATOM | 4594 | OE1 | GLN | A | 320 | 18.867 | 12.566 | 38.921 | 1.00 | 0.00 | O |
| ATOM | 4595 | NE2 | GLN | A | 320 | 20.457 | 13.136 | 40.344 | 1.00 | 0.00 | N |
| ATOM | 4596 | HE21 | GLN | A | 320 | 20.728 | 13.401 | 41.269 | 1.00 | 0.00 | H |
| ATOM | 4597 | HE22 | GLN | A | 320 | 21.150 | 13.034 | 39.630 | 1.00 | 0.00 | H |
| ATOM | 4598 | C | GLN | A | 320 | 16.523 | 16.396 | 42.539 | 1.00 | 0.00 | C |
| ATOM | 4599 | O | GLN | A | 320 | 17.417 | 17.233 | 42.453 | 1.00 | 0.00 | O |
| ATOM | 4600 | N | LYS | A | 321 | 15.233 | 16.615 | 42.720 | 1.00 | 0.00 | N |
| ATOM | 4601 | H | LYS | A | 321 | 14.672 | 15.789 | 42.665 | 1.00 | 0.00 | H |
| ATOM | 4602 | CA | LYS | A | 321 | 14.466 | 17.830 | 42.985 | 1.00 | 0.00 | C |
| ATOM | 4603 | HA | LYS | A | 321 | 13.517 | 17.515 | 42.966 | 1.00 | 0.00 | H |
| ATOM | 4604 | CB | LYS | A | 321 | 14.841 | 18.346 | 44.333 | 1.00 | 0.00 | C |
| ATOM | 4605 | HB1 | LYS | A | 321 | 14.414 | 19.241 | 44.458 | 1.00 | 0.00 | H |
| ATOM | 4606 | HB2 | LYS | A | 321 | 15.836 | 18.443 | 44.368 | 1.00 | 0.00 | H |
| ATOM | 4607 | CG | LYS | A | 321 | 14.380 | 17.399 | 45.445 | 1.00 | 0.00 | C |
| ATOM | 4608 | HG1 | LYS | A | 321 | 14.793 | 17.687 | 46.309 | 1.00 | 0.00 | H |
| ATOM | 4609 | HG2 | LYS | A | 321 | 14.679 | 16.470 | 45.226 | 1.00 | 0.00 | H |
| ATOM | 4610 | CD | LYS | A | 321 | 12.828 | 17.433 | 45.570 | 1.00 | 0.00 | C |
| ATOM | 4611 | HD1 | LYS | A | 321 | 12.407 | 17.085 | 44.732 | 1.00 | 0.00 | H |
| ATOM | 4612 | HD2 | LYS | A | 321 | 12.516 | 18.368 | 45.737 | 1.00 | 0.00 | H |
| ATOM | 4613 | CE | LYS | A | 321 | 12.418 | 16.552 | 46.740 | 1.00 | 0.00 | C |
| ATOM | 4614 | HE1 | LYS | A | 321 | 11.553 | 16.875 | 47.124 | 1.00 | 0.00 | H |
| ATOM | 4615 | HE2 | LYS | A | 321 | 13.125 | 16.573 | 47.447 | 1.00 | 0.00 | H |
| ATOM | 4616 | NZ | LYS | A | 321 | 12.242 | 15.161 | 46.295 | 1.00 | 0.00 | N |
| ATOM | 4617 | HZ1 | LYS | A | 321 | 11.973 | 14.594 | 47.073 | 1.00 | 0.00 | H |
| ATOM | 4618 | HZ2 | LYS | A | 321 | 11.531 | 15.123 | 45.593 | 1.00 | 0.00 | H |
| ATOM | 4619 | HZ3 | LYS | A | 321 | 13.103 | 14.821 | 45.916 | 1.00 | 0.00 | H |
| ATOM | 4620 | C | LYS | A | 321 | 14.585 | 18.947 | 41.927 | 1.00 | 0.00 | C |
| ATOM | 4621 | O | LYS | A | 321 | 14.646 | 20.145 | 42.239 | 1.00 | 0.00 | O |
| ATOM | 4622 | N | VAL | A | 322 | 14.724 | 18.559 | 40.619 | 1.00 | 0.00 | N |
| ATOM | 4623 | H | VAL | A | 322 | 14.627 | 17.589 | 40.394 | 1.00 | 0.00 | H |
| ATOM | 4624 | CA | VAL | A | 322 | 15.017 | 19.531 | 39.506 | 1.00 | 0.00 | C |
| ATOM | 4625 | HA | VAL | A | 322 | 14.733 | 20.406 | 39.898 | 1.00 | 0.00 | H |
| ATOM | 4626 | CB | VAL | A | 322 | 16.492 | 19.571 | 39.081 | 1.00 | 0.00 | C |
| ATOM | 4627 | HB | VAL | A | 322 | 16.558 | 19.985 | 38.173 | 1.00 | 0.00 | H |
| ATOM | 4628 | CG1 | VAL | A | 322 | 17.279 | 20.296 | 40.187 | 1.00 | 0.00 | C |
| ATOM | 4629 | HG11 | VAL | A | 322 | 18.246 | 20.337 | 39.936 | 1.00 | 0.00 | H |
| ATOM | 4630 | HG12 | VAL | A | 322 | 16.923 | 21.224 | 40.295 | 1.00 | 0.00 | H |
| ATOM | 4631 | HG13 | VAL | A | 322 | 17.179 | 19.798 | 41.048 | 1.00 | 0.00 | H |
| ATOM | 4632 | CG2 | VAL | A | 322 | 17.075 | 18.228 | 38.992 | 1.00 | 0.00 | C |
| ATOM | 4633 | HG21 | VAL | A | 322 | 18.033 | 18.298 | 38.713 | 1.00 | 0.00 | H |
| ATOM | 4634 | HG22 | VAL | A | 322 | 17.019 | 17.781 | 39.885 | 1.00 | 0.00 | H |
| ATOM | 4635 | HG23 | VAL | A | 322 | 16.570 | 17.690 | 38.317 | 1.00 | 0.00 | H |
| ATOM | 4636 | C | VAL | A | 322 | 14.272 | 19.119 | 38.215 | 1.00 | 0.00 | C |
| ATOM | 4637 | O | VAL | A | 322 | 13.937 | 17.923 | 38.124 | 1.00 | 0.00 | O |
| ATOM | 4638 | N | PRO | A | 323 | 13.922 | 20.095 | 37.336 | 1.00 | 0.00 | N |
| ATOM | 4639 | CD | PRO | A | 323 | 14.114 | 21.565 | 37.380 | 1.00 | 0.00 | C |
| ATOM | 4640 | HD1 | PRO | A | 323 | 15.090 | 21.779 | 37.337 | 1.00 | 0.00 | H |
| ATOM | 4641 | HD2 | PRO | A | 323 | 13.729 | 21.930 | 38.228 | 1.00 | 0.00 | H |

```
ATOM   4642  CG  PRO A 323      13.340  22.088  36.109  1.00  0.00           C
ATOM   4643  HG1 PRO A 323      13.954  22.201  35.328  1.00  0.00           H
ATOM   4644  HG2 PRO A 323      12.874  22.953  36.295  1.00  0.00           H
ATOM   4645  CB  PRO A 323      12.322  20.993  35.823  1.00  0.00           C
ATOM   4646  HB1 PRO A 323      12.078  20.983  34.853  1.00  0.00           H
ATOM   4647  HB2 PRO A 323      11.498  21.125  36.373  1.00  0.00           H
ATOM   4648  CA  PRO A 323      12.996  19.704  36.201  1.00  0.00           C
ATOM   4649  HA  PRO A 323      12.345  19.012  36.512  1.00  0.00           H
ATOM   4650  C   PRO A 323      13.712  19.053  34.982  1.00  0.00           C
ATOM   4651  O   PRO A 323      14.929  19.009  34.916  1.00  0.00           O
ATOM   4652  N   TYR A 324      12.871  18.596  34.040  1.00  0.00           N
ATOM   4653  H   TYR A 324      11.923  18.439  34.317  1.00  0.00           H
ATOM   4654  CA  TYR A 324      13.216  18.300  32.619  1.00  0.00           C
ATOM   4655  HA  TYR A 324      13.928  18.945  32.341  1.00  0.00           H
ATOM   4656  CB  TYR A 324      13.840  16.923  32.456  1.00  0.00           C
ATOM   4657  HB1 TYR A 324      14.546  16.798  33.153  1.00  0.00           H
ATOM   4658  HB2 TYR A 324      14.252  16.855  31.547  1.00  0.00           H
ATOM   4659  CG  TYR A 324      12.838  15.826  32.601  1.00  0.00           C
ATOM   4660  CD1 TYR A 324      12.424  15.521  33.890  1.00  0.00           C
ATOM   4661  HD1 TYR A 324      12.728  16.079  34.662  1.00  0.00           H
ATOM   4662  CE1 TYR A 324      11.566  14.411  34.104  1.00  0.00           C
ATOM   4663  HE1 TYR A 324      11.365  14.127  35.041  1.00  0.00           H
ATOM   4664  CZ  TYR A 324      10.998  13.708  33.050  1.00  0.00           C
ATOM   4665  OH  TYR A 324      10.071  12.680  33.251  1.00  0.00           O
ATOM   4666  HH  TYR A 324       9.789  12.308  32.367  1.00  0.00           H
ATOM   4667  CE2 TYR A 324      11.395  13.991  31.718  1.00  0.00           C
ATOM   4668  HE2 TYR A 324      11.018  13.494  30.936  1.00  0.00           H
ATOM   4669  CD2 TYR A 324      12.379  15.043  31.558  1.00  0.00           C
ATOM   4670  HD2 TYR A 324      12.746  15.212  30.643  1.00  0.00           H
ATOM   4671  C   TYR A 324      11.984  18.518  31.733  1.00  0.00           C
ATOM   4672  O   TYR A 324      10.804  18.536  32.163  1.00  0.00           O
ATOM   4673  N   ILE A 325      12.298  18.805  30.420  1.00  0.00           N
ATOM   4674  H   ILE A 325      13.268  18.860  30.183  1.00  0.00           H
ATOM   4675  CA  ILE A 325      11.327  19.037  29.339  1.00  0.00           C
ATOM   4676  HA  ILE A 325      10.415  18.744  29.626  1.00  0.00           H
ATOM   4677  CB  ILE A 325      11.283  20.623  29.061  1.00  0.00           C
ATOM   4678  HB  ILE A 325      12.200  20.882  29.364  1.00  0.00           H
ATOM   4679  CG2 ILE A 325      11.192  21.132  27.635  1.00  0.00           C
ATOM   4680 HG21 ILE A 325      11.175  22.132  27.638  1.00  0.00           H
ATOM   4681 HG22 ILE A 325      11.986  20.814  27.116  1.00  0.00           H
ATOM   4682 HG23 ILE A 325      10.356  20.785  27.210  1.00  0.00           H
ATOM   4683  CG1 ILE A 325      10.121  21.358  29.810  1.00  0.00           C
ATOM   4684 HG11 ILE A 325      10.079  20.997  30.742  1.00  0.00           H
ATOM   4685 HG12 ILE A 325       9.267  21.149  29.333  1.00  0.00           H
ATOM   4686  CD  ILE A 325      10.240  22.909  29.909  1.00  0.00           C
ATOM   4687  HD1 ILE A 325       9.451  23.273  30.403  1.00  0.00           H
ATOM   4688  HD2 ILE A 325      11.079  23.148  30.398  1.00  0.00           H
ATOM   4689  HD3 ILE A 325      10.267  23.301  28.989  1.00  0.00           H
ATOM   4690  C   ILE A 325      11.653  18.086  28.141  1.00  0.00           C
ATOM   4691  O   ILE A 325      12.811  17.584  28.111  1.00  0.00           O
ATOM   4692  N   PHE A 326      10.752  17.884  27.247  1.00  0.00           N
ATOM   4693  H   PHE A 326       9.859  18.314  27.377  1.00  0.00           H
ATOM   4694  CA  PHE A 326      10.953  17.055  26.038  1.00  0.00           C
ATOM   4695  HA  PHE A 326      11.897  17.197  25.740  1.00  0.00           H
```

| | | | | | | | | | | |
|---|---|---|---|---|---|---|---|---|---|---|
| ATOM | 4696 | CB | PHE | A | 326 | 10.671 | 15.521 | 26.446 | 1.00 | 0.00 | C |
| ATOM | 4697 | HB1 | PHE | A | 326 | 9.956 | 15.473 | 27.143 | 1.00 | 0.00 | H |
| ATOM | 4698 | HB2 | PHE | A | 326 | 11.506 | 15.095 | 26.795 | 1.00 | 0.00 | H |
| ATOM | 4699 | CG | PHE | A | 326 | 10.203 | 14.726 | 25.224 | 1.00 | 0.00 | C |
| ATOM | 4700 | CD1 | PHE | A | 326 | 8.878 | 14.382 | 24.931 | 1.00 | 0.00 | C |
| ATOM | 4701 | HD1 | PHE | A | 326 | 8.159 | 14.728 | 25.534 | 1.00 | 0.00 | H |
| ATOM | 4702 | CE1 | PHE | A | 326 | 8.489 | 13.569 | 23.826 | 1.00 | 0.00 | C |
| ATOM | 4703 | HE1 | PHE | A | 326 | 7.536 | 13.356 | 23.609 | 1.00 | 0.00 | H |
| ATOM | 4704 | CZ | PHE | A | 326 | 9.596 | 13.089 | 23.060 | 1.00 | 0.00 | C |
| ATOM | 4705 | HZ | PHE | A | 326 | 9.395 | 12.536 | 22.252 | 1.00 | 0.00 | H |
| ATOM | 4706 | CE2 | PHE | A | 326 | 10.935 | 13.335 | 23.356 | 1.00 | 0.00 | C |
| ATOM | 4707 | HE2 | PHE | A | 326 | 11.654 | 12.963 | 22.769 | 1.00 | 0.00 | H |
| ATOM | 4708 | CD2 | PHE | A | 326 | 11.276 | 14.111 | 24.486 | 1.00 | 0.00 | C |
| ATOM | 4709 | HD2 | PHE | A | 326 | 12.229 | 14.229 | 24.764 | 1.00 | 0.00 | H |
| ATOM | 4710 | C | PHE | A | 326 | 10.076 | 17.556 | 24.851 | 1.00 | 0.00 | C |
| ATOM | 4711 | O | PHE | A | 326 | 8.960 | 18.053 | 25.180 | 1.00 | 0.00 | O |
| ATOM | 4712 | N | ARG | A | 327 | 10.583 | 17.497 | 23.589 | 1.00 | 0.00 | N |
| ATOM | 4713 | H | ARG | A | 327 | 11.542 | 17.232 | 23.492 | 1.00 | 0.00 | H |
| ATOM | 4714 | CA | ARG | A | 327 | 9.822 | 17.796 | 22.332 | 1.00 | 0.00 | C |
| ATOM | 4715 | HA | ARG | A | 327 | 8.846 | 17.689 | 22.522 | 1.00 | 0.00 | H |
| ATOM | 4716 | CB | ARG | A | 327 | 10.207 | 19.250 | 21.946 | 1.00 | 0.00 | C |
| ATOM | 4717 | HB1 | ARG | A | 327 | 11.167 | 19.277 | 21.667 | 1.00 | 0.00 | H |
| ATOM | 4718 | HB2 | ARG | A | 327 | 10.072 | 19.850 | 22.734 | 1.00 | 0.00 | H |
| ATOM | 4719 | CG | ARG | A | 327 | 9.325 | 19.740 | 20.777 | 1.00 | 0.00 | C |
| ATOM | 4720 | HG1 | ARG | A | 327 | 8.391 | 19.404 | 20.898 | 1.00 | 0.00 | H |
| ATOM | 4721 | HG2 | ARG | A | 327 | 9.693 | 19.397 | 19.913 | 1.00 | 0.00 | H |
| ATOM | 4722 | CD | ARG | A | 327 | 9.328 | 21.290 | 20.770 | 1.00 | 0.00 | C |
| ATOM | 4723 | HD1 | ARG | A | 327 | 9.229 | 21.633 | 21.704 | 1.00 | 0.00 | H |
| ATOM | 4724 | HD2 | ARG | A | 327 | 8.573 | 21.628 | 20.209 | 1.00 | 0.00 | H |
| ATOM | 4725 | NE | ARG | A | 327 | 10.618 | 21.814 | 20.202 | 1.00 | 0.00 | N |
| ATOM | 4726 | HE | ARG | A | 327 | 11.305 | 21.162 | 19.880 | 1.00 | 0.00 | H |
| ATOM | 4727 | CZ | ARG | A | 327 | 10.874 | 23.111 | 20.114 | 1.00 | 0.00 | C |
| ATOM | 4728 | NH1 | ARG | A | 327 | 12.044 | 23.506 | 19.670 | 1.00 | 0.00 | N |
| ATOM | 4729 | HH11 | ARG | A | 327 | 12.731 | 22.831 | 19.402 | 1.00 | 0.00 | H |
| ATOM | 4730 | HH12 | ARG | A | 327 | 12.248 | 24.482 | 19.599 | 1.00 | 0.00 | H |
| ATOM | 4731 | NH2 | ARG | A | 327 | 10.035 | 24.051 | 20.432 | 1.00 | 0.00 | N |
| ATOM | 4732 | HH21 | ARG | A | 327 | 9.124 | 23.815 | 20.770 | 1.00 | 0.00 | H |
| ATOM | 4733 | HH22 | ARG | A | 327 | 10.300 | 25.011 | 20.339 | 1.00 | 0.00 | H |
| ATOM | 4734 | C | ARG | A | 327 | 10.178 | 16.805 | 21.169 | 1.00 | 0.00 | C |
| ATOM | 4735 | O | ARG | A | 327 | 11.154 | 16.985 | 20.451 | 1.00 | 0.00 | O |
| ATOM | 4736 | N | ASP | A | 328 | 9.315 | 15.816 | 20.978 | 1.00 | 0.00 | N |
| ATOM | 4737 | H | ASP | A | 328 | 8.587 | 15.738 | 21.659 | 1.00 | 0.00 | H |
| ATOM | 4738 | CA | ASP | A | 328 | 9.287 | 14.838 | 19.918 | 1.00 | 0.00 | C |
| ATOM | 4739 | HA | ASP | A | 328 | 8.523 | 14.273 | 20.229 | 1.00 | 0.00 | H |
| ATOM | 4740 | CB | ASP | A | 328 | 8.898 | 15.547 | 18.628 | 1.00 | 0.00 | C |
| ATOM | 4741 | HB1 | ASP | A | 328 | 9.598 | 16.246 | 18.482 | 1.00 | 0.00 | H |
| ATOM | 4742 | HB2 | ASP | A | 328 | 8.021 | 15.989 | 18.817 | 1.00 | 0.00 | H |
| ATOM | 4743 | CG | ASP | A | 328 | 8.732 | 14.817 | 17.284 | 1.00 | 0.00 | C |
| ATOM | 4744 | OD1 | ASP | A | 328 | 8.567 | 15.520 | 16.292 | 1.00 | 0.00 | O |
| ATOM | 4745 | OD2 | ASP | A | 328 | 8.661 | 13.581 | 17.137 | 1.00 | 0.00 | O |
| ATOM | 4746 | C | ASP | A | 328 | 10.449 | 13.743 | 19.822 | 1.00 | 0.00 | C |
| ATOM | 4747 | O | ASP | A | 328 | 10.150 | 12.547 | 19.843 | 1.00 | 0.00 | O |
| ATOM | 4748 | N | ASN | A | 329 | 11.695 | 14.227 | 19.959 | 1.00 | 0.00 | N |
| ATOM | 4749 | H | ASN | A | 329 | 11.828 | 15.216 | 19.890 | 1.00 | 0.00 | H |

| | | | | | | | | | | | |
|---|---|---|---|---|---|---|---|---|---|---|---|
| ATOM | 4750 | CA | ASN | A | 329 | 12.877 | 13.379 | 20.206 | 1.00 | 0.00 | C |
| ATOM | 4751 | HA | ASN | A | 329 | 12.563 | 12.578 | 20.715 | 1.00 | 0.00 | H |
| ATOM | 4752 | CB | ASN | A | 329 | 13.452 | 12.932 | 18.858 | 1.00 | 0.00 | C |
| ATOM | 4753 | HB1 | ASN | A | 329 | 12.834 | 12.254 | 18.460 | 1.00 | 0.00 | H |
| ATOM | 4754 | HB2 | ASN | A | 329 | 14.346 | 12.514 | 19.019 | 1.00 | 0.00 | H |
| ATOM | 4755 | CG | ASN | A | 329 | 13.638 | 14.030 | 17.845 | 1.00 | 0.00 | C |
| ATOM | 4756 | OD1 | ASN | A | 329 | 14.667 | 14.668 | 17.847 | 1.00 | 0.00 | O |
| ATOM | 4757 | ND2 | ASN | A | 329 | 12.793 | 14.371 | 16.909 | 1.00 | 0.00 | N |
| ATOM | 4758 | HD21 | ASN | A | 329 | 11.925 | 13.884 | 16.818 | 1.00 | 0.00 | H |
| ATOM | 4759 | HD22 | ASN | A | 329 | 13.015 | 15.119 | 16.284 | 1.00 | 0.00 | H |
| ATOM | 4760 | C | ASN | A | 329 | 13.886 | 14.002 | 21.150 | 1.00 | 0.00 | C |
| ATOM | 4761 | O | ASN | A | 329 | 14.673 | 13.258 | 21.692 | 1.00 | 0.00 | O |
| ATOM | 4762 | N | GLN | A | 330 | 13.811 | 15.313 | 21.378 | 1.00 | 0.00 | N |
| ATOM | 4763 | H | GLN | A | 330 | 13.046 | 15.842 | 21.010 | 1.00 | 0.00 | H |
| ATOM | 4764 | CA | GLN | A | 330 | 14.859 | 15.985 | 22.175 | 1.00 | 0.00 | C |
| ATOM | 4765 | HA | GLN | A | 330 | 15.661 | 15.389 | 22.219 | 1.00 | 0.00 | H |
| ATOM | 4766 | CB | GLN | A | 330 | 15.246 | 17.262 | 21.443 | 1.00 | 0.00 | C |
| ATOM | 4767 | HB1 | GLN | A | 330 | 14.635 | 17.996 | 21.739 | 1.00 | 0.00 | H |
| ATOM | 4768 | HB2 | GLN | A | 330 | 15.135 | 17.112 | 20.461 | 1.00 | 0.00 | H |
| ATOM | 4769 | CG | GLN | A | 330 | 16.681 | 17.691 | 21.708 | 1.00 | 0.00 | C |
| ATOM | 4770 | HG1 | GLN | A | 330 | 17.304 | 16.963 | 21.422 | 1.00 | 0.00 | H |
| ATOM | 4771 | HG2 | GLN | A | 330 | 16.800 | 17.868 | 22.685 | 1.00 | 0.00 | H |
| ATOM | 4772 | CD | GLN | A | 330 | 16.982 | 18.962 | 20.917 | 1.00 | 0.00 | C |
| ATOM | 4773 | OE1 | GLN | A | 330 | 16.133 | 19.772 | 20.685 | 1.00 | 0.00 | O |
| ATOM | 4774 | NE2 | GLN | A | 330 | 18.253 | 19.087 | 20.559 | 1.00 | 0.00 | N |
| ATOM | 4775 | HE21 | GLN | A | 330 | 18.915 | 18.381 | 20.811 | 1.00 | 0.00 | H |
| ATOM | 4776 | HE22 | GLN | A | 330 | 18.549 | 19.887 | 20.037 | 1.00 | 0.00 | H |
| ATOM | 4777 | C | GLN | A | 330 | 14.421 | 16.196 | 23.644 | 1.00 | 0.00 | C |
| ATOM | 4778 | O | GLN | A | 330 | 13.348 | 16.774 | 23.835 | 1.00 | 0.00 | O |
| ATOM | 4779 | N | TRP | A | 331 | 15.329 | 15.869 | 24.576 | 1.00 | 0.00 | N |
| ATOM | 4780 | H | TRP | A | 331 | 16.211 | 15.546 | 24.234 | 1.00 | 0.00 | H |
| ATOM | 4781 | CA | TRP | A | 331 | 15.191 | 15.925 | 25.997 | 1.00 | 0.00 | C |
| ATOM | 4782 | HA | TRP | A | 331 | 14.268 | 16.233 | 26.230 | 1.00 | 0.00 | H |
| ATOM | 4783 | CB | TRP | A | 331 | 15.611 | 14.453 | 26.463 | 1.00 | 0.00 | C |
| ATOM | 4784 | HB1 | TRP | A | 331 | 16.606 | 14.362 | 26.414 | 1.00 | 0.00 | H |
| ATOM | 4785 | HB2 | TRP | A | 331 | 15.184 | 13.780 | 25.859 | 1.00 | 0.00 | H |
| ATOM | 4786 | CG | TRP | A | 331 | 15.213 | 14.082 | 27.871 | 1.00 | 0.00 | C |
| ATOM | 4787 | CD1 | TRP | A | 331 | 14.154 | 13.309 | 28.109 | 1.00 | 0.00 | C |
| ATOM | 4788 | HD1 | TRP | A | 331 | 13.524 | 12.930 | 27.431 | 1.00 | 0.00 | H |
| ATOM | 4789 | NE1 | TRP | A | 331 | 14.098 | 13.134 | 29.481 | 1.00 | 0.00 | N |
| ATOM | 4790 | HE1 | TRP | A | 331 | 13.380 | 12.613 | 29.942 | 1.00 | 0.00 | H |
| ATOM | 4791 | CE2 | TRP | A | 331 | 15.122 | 13.745 | 30.138 | 1.00 | 0.00 | C |
| ATOM | 4792 | CZ2 | TRP | A | 331 | 15.445 | 13.766 | 31.478 | 1.00 | 0.00 | C |
| ATOM | 4793 | HZ2 | TRP | A | 331 | 14.871 | 13.273 | 32.131 | 1.00 | 0.00 | H |
| ATOM | 4794 | CH2 | TRP | A | 331 | 16.593 | 14.485 | 31.948 | 1.00 | 0.00 | C |
| ATOM | 4795 | HH2 | TRP | A | 331 | 16.808 | 14.558 | 32.922 | 1.00 | 0.00 | H |
| ATOM | 4796 | CZ3 | TRP | A | 331 | 17.394 | 15.078 | 30.969 | 1.00 | 0.00 | C |
| ATOM | 4797 | HZ3 | TRP | A | 331 | 18.238 | 15.530 | 31.258 | 1.00 | 0.00 | H |
| ATOM | 4798 | CE3 | TRP | A | 331 | 17.074 | 15.074 | 29.584 | 1.00 | 0.00 | C |
| ATOM | 4799 | HE3 | TRP | A | 331 | 17.654 | 15.551 | 28.924 | 1.00 | 0.00 | H |
| ATOM | 4800 | CD2 | TRP | A | 331 | 15.879 | 14.358 | 29.147 | 1.00 | 0.00 | C |
| ATOM | 4801 | C | TRP | A | 331 | 16.160 | 16.944 | 26.567 | 1.00 | 0.00 | C |
| ATOM | 4802 | O | TRP | A | 331 | 17.294 | 16.993 | 26.148 | 1.00 | 0.00 | O |
| ATOM | 4803 | N | VAL | A | 332 | 15.788 | 17.753 | 27.602 | 1.00 | 0.00 | N |

| | | | | | | | | | | | |
|---|---|---|---|---|---|---|---|---|---|---|---|
| ATOM | 4804 | H | VAL | A | 332 | 14.863 | 17.624 | 27.960 | 1.00 | 0.00 | H |
| ATOM | 4805 | CA | VAL | A | 332 | 16.586 | 18.771 | 28.232 | 1.00 | 0.00 | C |
| ATOM | 4806 | HA | VAL | A | 332 | 17.530 | 18.475 | 28.085 | 1.00 | 0.00 | H |
| ATOM | 4807 | CB | VAL | A | 332 | 16.443 | 20.138 | 27.575 | 1.00 | 0.00 | C |
| ATOM | 4808 | HB | VAL | A | 332 | 16.792 | 19.992 | 26.649 | 1.00 | 0.00 | H |
| ATOM | 4809 | CG1 | VAL | A | 332 | 14.953 | 20.652 | 27.449 | 1.00 | 0.00 | C |
| ATOM | 4810 | HG11 | VAL | A | 332 | 14.946 | 21.550 | 27.010 | 1.00 | 0.00 | H |
| ATOM | 4811 | HG12 | VAL | A | 332 | 14.424 | 20.007 | 26.897 | 1.00 | 0.00 | H |
| ATOM | 4812 | HG13 | VAL | A | 332 | 14.547 | 20.723 | 28.360 | 1.00 | 0.00 | H |
| ATOM | 4813 | CG2 | VAL | A | 332 | 17.319 | 21.188 | 28.295 | 1.00 | 0.00 | C |
| ATOM | 4814 | HG21 | VAL | A | 332 | 17.210 | 22.076 | 27.848 | 1.00 | 0.00 | H |
| ATOM | 4815 | HG22 | VAL | A | 332 | 17.036 | 21.259 | 29.251 | 1.00 | 0.00 | H |
| ATOM | 4816 | HG23 | VAL | A | 332 | 18.278 | 20.908 | 28.251 | 1.00 | 0.00 | H |
| ATOM | 4817 | C | VAL | A | 332 | 16.394 | 18.777 | 29.695 | 1.00 | 0.00 | C |
| ATOM | 4818 | O | VAL | A | 332 | 15.281 | 18.919 | 30.191 | 1.00 | 0.00 | O |
| ATOM | 4819 | N | GLY | A | 333 | 17.495 | 18.717 | 30.411 | 1.00 | 0.00 | N |
| ATOM | 4820 | H | GLY | A | 333 | 18.331 | 18.578 | 29.880 | 1.00 | 0.00 | H |
| ATOM | 4821 | CA | GLY | A | 333 | 17.701 | 18.820 | 31.865 | 1.00 | 0.00 | C |
| ATOM | 4822 | HA1 | GLY | A | 333 | 18.341 | 18.101 | 32.135 | 1.00 | 0.00 | H |
| ATOM | 4823 | HA2 | GLY | A | 333 | 16.819 | 18.666 | 32.311 | 1.00 | 0.00 | H |
| ATOM | 4824 | C | GLY | A | 333 | 18.255 | 20.181 | 32.287 | 1.00 | 0.00 | C |
| ATOM | 4825 | O | GLY | A | 333 | 19.448 | 20.432 | 32.151 | 1.00 | 0.00 | O |
| ATOM | 4826 | N | PHE | A | 334 | 17.395 | 21.014 | 32.919 | 1.00 | 0.00 | N |
| ATOM | 4827 | H | PHE | A | 334 | 16.555 | 20.614 | 33.285 | 1.00 | 0.00 | H |
| ATOM | 4828 | CA | PHE | A | 334 | 17.592 | 22.451 | 33.112 | 1.00 | 0.00 | C |
| ATOM | 4829 | HA | PHE | A | 334 | 18.582 | 22.573 | 33.040 | 1.00 | 0.00 | H |
| ATOM | 4830 | CB | PHE | A | 334 | 16.921 | 23.254 | 32.013 | 1.00 | 0.00 | C |
| ATOM | 4831 | HB1 | PHE | A | 334 | 17.116 | 22.772 | 31.159 | 1.00 | 0.00 | H |
| ATOM | 4832 | HB2 | PHE | A | 334 | 17.375 | 24.145 | 32.004 | 1.00 | 0.00 | H |
| ATOM | 4833 | CG | PHE | A | 334 | 15.348 | 23.490 | 32.112 | 1.00 | 0.00 | C |
| ATOM | 4834 | CD1 | PHE | A | 334 | 14.465 | 22.386 | 31.831 | 1.00 | 0.00 | C |
| ATOM | 4835 | HD1 | PHE | A | 334 | 14.845 | 21.562 | 31.412 | 1.00 | 0.00 | H |
| ATOM | 4836 | CE1 | PHE | A | 334 | 13.104 | 22.422 | 32.120 | 1.00 | 0.00 | C |
| ATOM | 4837 | HE1 | PHE | A | 334 | 12.501 | 21.638 | 31.975 | 1.00 | 0.00 | H |
| ATOM | 4838 | CZ | PHE | A | 334 | 12.639 | 23.663 | 32.638 | 1.00 | 0.00 | C |
| ATOM | 4839 | HZ | PHE | A | 334 | 11.676 | 23.704 | 32.906 | 1.00 | 0.00 | H |
| ATOM | 4840 | CE2 | PHE | A | 334 | 13.389 | 24.832 | 32.812 | 1.00 | 0.00 | C |
| ATOM | 4841 | HE2 | PHE | A | 334 | 12.984 | 25.687 | 33.137 | 1.00 | 0.00 | H |
| ATOM | 4842 | CD2 | PHE | A | 334 | 14.788 | 24.712 | 32.487 | 1.00 | 0.00 | C |
| ATOM | 4843 | HD2 | PHE | A | 334 | 15.371 | 25.523 | 32.531 | 1.00 | 0.00 | H |
| ATOM | 4844 | C | PHE | A | 334 | 17.234 | 22.934 | 34.493 | 1.00 | 0.00 | C |
| ATOM | 4845 | O | PHE | A | 334 | 16.470 | 22.277 | 35.183 | 1.00 | 0.00 | O |
| ATOM | 4846 | N | ASP | A | 335 | 17.653 | 24.127 | 34.938 | 1.00 | 0.00 | N |
| ATOM | 4847 | H | ASP | A | 335 | 18.362 | 24.603 | 34.417 | 1.00 | 0.00 | H |
| ATOM | 4848 | CA | ASP | A | 335 | 17.116 | 24.789 | 36.177 | 1.00 | 0.00 | C |
| ATOM | 4849 | HA | ASP | A | 335 | 16.662 | 24.043 | 36.664 | 1.00 | 0.00 | H |
| ATOM | 4850 | CB | ASP | A | 335 | 18.253 | 25.421 | 37.072 | 1.00 | 0.00 | C |
| ATOM | 4851 | HB1 | ASP | A | 335 | 17.858 | 26.087 | 37.705 | 1.00 | 0.00 | H |
| ATOM | 4852 | HB2 | ASP | A | 335 | 18.932 | 25.872 | 36.493 | 1.00 | 0.00 | H |
| ATOM | 4853 | CG | ASP | A | 335 | 18.964 | 24.377 | 37.887 | 1.00 | 0.00 | C |
| ATOM | 4854 | OD1 | ASP | A | 335 | 18.283 | 23.624 | 38.655 | 1.00 | 0.00 | O |
| ATOM | 4855 | OD2 | ASP | A | 335 | 20.224 | 24.373 | 37.987 | 1.00 | 0.00 | O |
| ATOM | 4856 | C | ASP | A | 335 | 16.108 | 25.913 | 35.807 | 1.00 | 0.00 | C |
| ATOM | 4857 | O | ASP | A | 335 | 16.182 | 26.605 | 34.796 | 1.00 | 0.00 | O |

| | | | | | | | | | | | |
|---|---|---|---|---|---|---|---|---|---|---|---|
| ATOM | 4858 | N | ASP | A | 336 | 15.100 | 25.966 | 36.722 | 1.00 | 0.00 | N |
| ATOM | 4859 | H | ASP | A | 336 | 15.130 | 25.309 | 37.475 | 1.00 | 0.00 | H |
| ATOM | 4860 | CA | ASP | A | 336 | 13.984 | 26.899 | 36.695 | 1.00 | 0.00 | C |
| ATOM | 4861 | HA | ASP | A | 336 | 14.239 | 27.476 | 35.919 | 1.00 | 0.00 | H |
| ATOM | 4862 | CB | ASP | A | 336 | 12.608 | 26.174 | 36.454 | 1.00 | 0.00 | C |
| ATOM | 4863 | HB1 | ASP | A | 336 | 12.715 | 25.557 | 35.675 | 1.00 | 0.00 | H |
| ATOM | 4864 | HB2 | ASP | A | 336 | 11.925 | 26.871 | 36.236 | 1.00 | 0.00 | H |
| ATOM | 4865 | CG | ASP | A | 336 | 12.030 | 25.339 | 37.595 | 1.00 | 0.00 | C |
| ATOM | 4866 | OD1 | ASP | A | 336 | 12.457 | 25.441 | 38.759 | 1.00 | 0.00 | O |
| ATOM | 4867 | OD2 | ASP | A | 336 | 10.895 | 24.878 | 37.360 | 1.00 | 0.00 | O |
| ATOM | 4868 | C | ASP | A | 336 | 13.882 | 27.737 | 38.013 | 1.00 | 0.00 | C |
| ATOM | 4869 | O | ASP | A | 336 | 14.693 | 27.596 | 38.923 | 1.00 | 0.00 | O |
| ATOM | 4870 | N | VAL | A | 337 | 12.850 | 28.566 | 38.069 | 1.00 | 0.00 | N |
| ATOM | 4871 | H | VAL | A | 337 | 12.187 | 28.544 | 37.321 | 1.00 | 0.00 | H |
| ATOM | 4872 | CA | VAL | A | 337 | 12.620 | 29.533 | 39.187 | 1.00 | 0.00 | C |
| ATOM | 4873 | HA | VAL | A | 337 | 13.509 | 29.989 | 39.159 | 1.00 | 0.00 | H |
| ATOM | 4874 | CB | VAL | A | 337 | 11.427 | 30.568 | 39.003 | 1.00 | 0.00 | C |
| ATOM | 4875 | HB | VAL | A | 337 | 11.291 | 31.196 | 39.769 | 1.00 | 0.00 | H |
| ATOM | 4876 | CG1 | VAL | A | 337 | 11.740 | 31.320 | 37.745 | 1.00 | 0.00 | C |
| ATOM | 4877 | HG11 | VAL | A | 337 | 11.021 | 31.993 | 37.571 | 1.00 | 0.00 | H |
| ATOM | 4878 | HG12 | VAL | A | 337 | 12.618 | 31.788 | 37.846 | 1.00 | 0.00 | H |
| ATOM | 4879 | HG13 | VAL | A | 337 | 11.790 | 30.680 | 36.978 | 1.00 | 0.00 | H |
| ATOM | 4880 | CG2 | VAL | A | 337 | 10.148 | 29.805 | 38.800 | 1.00 | 0.00 | C |
| ATOM | 4881 | HG21 | VAL | A | 337 | 9.391 | 30.448 | 38.683 | 1.00 | 0.00 | H |
| ATOM | 4882 | HG22 | VAL | A | 337 | 10.228 | 29.232 | 37.984 | 1.00 | 0.00 | H |
| ATOM | 4883 | HG23 | VAL | A | 337 | 9.974 | 29.227 | 39.597 | 1.00 | 0.00 | H |
| ATOM | 4884 | C | VAL | A | 337 | 12.456 | 28.887 | 40.525 | 1.00 | 0.00 | C |
| ATOM | 4885 | O | VAL | A | 337 | 12.997 | 29.446 | 41.430 | 1.00 | 0.00 | O |
| ATOM | 4886 | N | GLU | A | 338 | 11.673 | 27.780 | 40.647 | 1.00 | 0.00 | N |
| ATOM | 4887 | H | GLU | A | 338 | 11.164 | 27.487 | 39.837 | 1.00 | 0.00 | H |
| ATOM | 4888 | CA | GLU | A | 338 | 11.513 | 26.980 | 41.868 | 1.00 | 0.00 | C |
| ATOM | 4889 | HA | GLU | A | 338 | 11.205 | 27.677 | 42.515 | 1.00 | 0.00 | H |
| ATOM | 4890 | CB | GLU | A | 338 | 10.452 | 25.902 | 41.758 | 1.00 | 0.00 | C |
| ATOM | 4891 | HB1 | GLU | A | 338 | 10.140 | 25.723 | 42.691 | 1.00 | 0.00 | H |
| ATOM | 4892 | HB2 | GLU | A | 338 | 10.916 | 25.092 | 41.400 | 1.00 | 0.00 | H |
| ATOM | 4893 | CG | GLU | A | 338 | 9.193 | 26.113 | 40.899 | 1.00 | 0.00 | C |
| ATOM | 4894 | HG1 | GLU | A | 338 | 8.631 | 25.288 | 40.964 | 1.00 | 0.00 | H |
| ATOM | 4895 | HG2 | GLU | A | 338 | 9.483 | 26.246 | 39.951 | 1.00 | 0.00 | H |
| ATOM | 4896 | CD | GLU | A | 338 | 8.307 | 27.272 | 41.250 | 1.00 | 0.00 | C |
| ATOM | 4897 | OE1 | GLU | A | 338 | 7.371 | 27.489 | 40.433 | 1.00 | 0.00 | O |
| ATOM | 4898 | OE2 | GLU | A | 338 | 8.597 | 28.213 | 42.020 | 1.00 | 0.00 | O |
| ATOM | 4899 | C | GLU | A | 338 | 12.841 | 26.292 | 42.302 | 1.00 | 0.00 | C |
| ATOM | 4900 | O | GLU | A | 338 | 13.098 | 26.291 | 43.504 | 1.00 | 0.00 | O |
| ATOM | 4901 | N | SER | A | 339 | 13.622 | 25.775 | 41.344 | 1.00 | 0.00 | N |
| ATOM | 4902 | H | SER | A | 339 | 13.287 | 25.860 | 40.406 | 1.00 | 0.00 | H |
| ATOM | 4903 | CA | SER | A | 339 | 14.875 | 25.118 | 41.509 | 1.00 | 0.00 | C |
| ATOM | 4904 | HA | SER | A | 339 | 14.657 | 24.382 | 42.149 | 1.00 | 0.00 | H |
| ATOM | 4905 | CB | SER | A | 339 | 15.362 | 24.523 | 40.198 | 1.00 | 0.00 | C |
| ATOM | 4906 | HB1 | SER | A | 339 | 15.678 | 25.234 | 39.570 | 1.00 | 0.00 | H |
| ATOM | 4907 | HB2 | SER | A | 339 | 14.648 | 23.981 | 39.755 | 1.00 | 0.00 | H |
| ATOM | 4908 | OG | SER | A | 339 | 16.452 | 23.673 | 40.511 | 1.00 | 0.00 | O |
| ATOM | 4909 | HG | SER | A | 339 | 16.804 | 23.259 | 39.671 | 1.00 | 0.00 | H |
| ATOM | 4910 | C | SER | A | 339 | 15.943 | 26.076 | 42.098 | 1.00 | 0.00 | C |
| ATOM | 4911 | O | SER | A | 339 | 16.560 | 25.714 | 43.097 | 1.00 | 0.00 | O |

| | | | | | | | | | | | |
|---|---|---|---|---|---|---|---|---|---|---|---|
| ATOM | 4912 | N | PHE | A | 340 | 16.057 | 27.271 | 41.485 | 1.00 | 0.00 | N |
| ATOM | 4913 | H | PHE | A | 340 | 15.545 | 27.432 | 40.641 | 1.00 | 0.00 | H |
| ATOM | 4914 | CA | PHE | A | 340 | 16.917 | 28.357 | 42.021 | 1.00 | 0.00 | C |
| ATOM | 4915 | HA | PHE | A | 340 | 17.878 | 28.082 | 42.008 | 1.00 | 0.00 | H |
| ATOM | 4916 | CB | PHE | A | 340 | 16.665 | 29.628 | 41.083 | 1.00 | 0.00 | C |
| ATOM | 4917 | HB1 | PHE | A | 340 | 16.695 | 30.462 | 41.634 | 1.00 | 0.00 | H |
| ATOM | 4918 | HB2 | PHE | A | 340 | 15.769 | 29.549 | 40.646 | 1.00 | 0.00 | H |
| ATOM | 4919 | CG | PHE | A | 340 | 17.698 | 29.780 | 39.975 | 1.00 | 0.00 | C |
| ATOM | 4920 | CD1 | PHE | A | 340 | 18.026 | 28.728 | 39.092 | 1.00 | 0.00 | C |
| ATOM | 4921 | HD1 | PHE | A | 340 | 17.521 | 27.868 | 39.168 | 1.00 | 0.00 | H |
| ATOM | 4922 | CE1 | PHE | A | 340 | 19.021 | 28.831 | 38.119 | 1.00 | 0.00 | C |
| ATOM | 4923 | HE1 | PHE | A | 340 | 19.205 | 28.079 | 37.487 | 1.00 | 0.00 | H |
| ATOM | 4924 | CZ | PHE | A | 340 | 19.754 | 30.026 | 38.058 | 1.00 | 0.00 | C |
| ATOM | 4925 | HZ | PHE | A | 340 | 20.549 | 30.089 | 37.455 | 1.00 | 0.00 | H |
| ATOM | 4926 | CE2 | PHE | A | 340 | 19.382 | 31.133 | 38.835 | 1.00 | 0.00 | C |
| ATOM | 4927 | HE2 | PHE | A | 340 | 19.833 | 32.014 | 38.691 | 1.00 | 0.00 | H |
| ATOM | 4928 | CD2 | PHE | A | 340 | 18.370 | 31.013 | 39.825 | 1.00 | 0.00 | C |
| ATOM | 4929 | HD2 | PHE | A | 340 | 18.137 | 31.791 | 40.408 | 1.00 | 0.00 | H |
| ATOM | 4930 | C | PHE | A | 340 | 16.571 | 28.589 | 43.515 | 1.00 | 0.00 | C |
| ATOM | 4931 | O | PHE | A | 340 | 17.453 | 28.580 | 44.367 | 1.00 | 0.00 | O |
| ATOM | 4932 | N | LYS | A | 341 | 15.240 | 28.746 | 43.823 | 1.00 | 0.00 | N |
| ATOM | 4933 | H | LYS | A | 341 | 14.566 | 28.754 | 43.084 | 1.00 | 0.00 | H |
| ATOM | 4934 | CA | LYS | A | 341 | 14.782 | 28.903 | 45.214 | 1.00 | 0.00 | C |
| ATOM | 4935 | HA | LYS | A | 341 | 15.220 | 29.661 | 45.698 | 1.00 | 0.00 | H |
| ATOM | 4936 | CB | LYS | A | 341 | 13.237 | 29.236 | 45.091 | 1.00 | 0.00 | C |
| ATOM | 4937 | HB1 | LYS | A | 341 | 12.772 | 28.518 | 44.573 | 1.00 | 0.00 | H |
| ATOM | 4938 | HB2 | LYS | A | 341 | 13.112 | 30.114 | 44.629 | 1.00 | 0.00 | H |
| ATOM | 4939 | CG | LYS | A | 341 | 12.610 | 29.320 | 46.491 | 1.00 | 0.00 | C |
| ATOM | 4940 | HG1 | LYS | A | 341 | 13.271 | 29.717 | 47.128 | 1.00 | 0.00 | H |
| ATOM | 4941 | HG2 | LYS | A | 341 | 12.361 | 28.402 | 46.799 | 1.00 | 0.00 | H |
| ATOM | 4942 | CD | LYS | A | 341 | 11.377 | 30.171 | 46.499 | 1.00 | 0.00 | C |
| ATOM | 4943 | HD1 | LYS | A | 341 | 10.851 | 29.919 | 45.687 | 1.00 | 0.00 | H |
| ATOM | 4944 | HD2 | LYS | A | 341 | 11.684 | 31.119 | 46.416 | 1.00 | 0.00 | H |
| ATOM | 4945 | CE | LYS | A | 341 | 10.435 | 30.097 | 47.696 | 1.00 | 0.00 | C |
| ATOM | 4946 | HE1 | LYS | A | 341 | 10.011 | 29.192 | 47.728 | 1.00 | 0.00 | H |
| ATOM | 4947 | HE2 | LYS | A | 341 | 9.725 | 30.794 | 47.602 | 1.00 | 0.00 | H |
| ATOM | 4948 | NZ | LYS | A | 341 | 11.086 | 30.315 | 48.980 | 1.00 | 0.00 | N |
| ATOM | 4949 | HZ1 | LYS | A | 341 | 10.408 | 30.252 | 49.712 | 1.00 | 0.00 | H |
| ATOM | 4950 | HZ2 | LYS | A | 341 | 11.792 | 29.620 | 49.118 | 1.00 | 0.00 | H |
| ATOM | 4951 | HZ3 | LYS | A | 341 | 11.505 | 31.223 | 48.991 | 1.00 | 0.00 | H |
| ATOM | 4952 | C | LYS | A | 341 | 15.276 | 27.701 | 46.073 | 1.00 | 0.00 | C |
| ATOM | 4953 | O | LYS | A | 341 | 15.898 | 27.967 | 47.130 | 1.00 | 0.00 | O |
| ATOM | 4954 | N | THR | A | 342 | 14.992 | 26.496 | 45.609 | 1.00 | 0.00 | N |
| ATOM | 4955 | H | THR | A | 342 | 14.556 | 26.440 | 44.711 | 1.00 | 0.00 | H |
| ATOM | 4956 | CA | THR | A | 342 | 15.266 | 25.230 | 46.309 | 1.00 | 0.00 | C |
| ATOM | 4957 | HA | THR | A | 342 | 14.694 | 25.238 | 47.129 | 1.00 | 0.00 | H |
| ATOM | 4958 | CB | THR | A | 342 | 14.923 | 24.039 | 45.404 | 1.00 | 0.00 | C |
| ATOM | 4959 | HB | THR | A | 342 | 15.556 | 24.123 | 44.635 | 1.00 | 0.00 | H |
| ATOM | 4960 | CG2 | THR | A | 342 | 14.994 | 22.607 | 45.962 | 1.00 | 0.00 | C |
| ATOM | 4961 | HG21 | THR | A | 342 | 14.741 | 21.957 | 45.245 | 1.00 | 0.00 | H |
| ATOM | 4962 | HG22 | THR | A | 342 | 15.925 | 22.416 | 46.272 | 1.00 | 0.00 | H |
| ATOM | 4963 | HG23 | THR | A | 342 | 14.361 | 22.517 | 46.731 | 1.00 | 0.00 | H |
| ATOM | 4964 | OG1 | THR | A | 342 | 13.570 | 24.156 | 45.082 | 1.00 | 0.00 | O |
| ATOM | 4965 | HG1 | THR | A | 342 | 13.301 | 23.396 | 44.490 | 1.00 | 0.00 | H |

| | | | | | | | | | | | |
|---|---|---|---|---|---|---|---|---|---|---|---|
| ATOM | 4966 | C | THR | A | 342 | 16.747 | 25.143 | 46.674 | 1.00 | 0.00 | C |
| ATOM | 4967 | O | THR | A | 342 | 17.162 | 24.875 | 47.818 | 1.00 | 0.00 | O |
| ATOM | 4968 | N | LYS | A | 343 | 17.601 | 25.435 | 45.746 | 1.00 | 0.00 | N |
| ATOM | 4969 | H | LYS | A | 343 | 17.226 | 25.587 | 44.831 | 1.00 | 0.00 | H |
| ATOM | 4970 | CA | LYS | A | 343 | 19.085 | 25.571 | 45.885 | 1.00 | 0.00 | C |
| ATOM | 4971 | HA | LYS | A | 343 | 19.353 | 24.716 | 46.329 | 1.00 | 0.00 | H |
| ATOM | 4972 | CB | LYS | A | 343 | 19.700 | 25.809 | 44.466 | 1.00 | 0.00 | C |
| ATOM | 4973 | HB1 | LYS | A | 343 | 20.689 | 25.940 | 44.531 | 1.00 | 0.00 | H |
| ATOM | 4974 | HB2 | LYS | A | 343 | 19.285 | 26.610 | 44.035 | 1.00 | 0.00 | H |
| ATOM | 4975 | CG | LYS | A | 343 | 19.392 | 24.549 | 43.635 | 1.00 | 0.00 | C |
| ATOM | 4976 | HG1 | LYS | A | 343 | 18.399 | 24.439 | 43.594 | 1.00 | 0.00 | H |
| ATOM | 4977 | HG2 | LYS | A | 343 | 19.795 | 23.762 | 44.102 | 1.00 | 0.00 | H |
| ATOM | 4978 | CD | LYS | A | 343 | 19.933 | 24.580 | 42.191 | 1.00 | 0.00 | C |
| ATOM | 4979 | HD1 | LYS | A | 343 | 20.888 | 24.877 | 42.202 | 1.00 | 0.00 | H |
| ATOM | 4980 | HD2 | LYS | A | 343 | 19.391 | 25.222 | 41.649 | 1.00 | 0.00 | H |
| ATOM | 4981 | CE | LYS | A | 343 | 19.868 | 23.232 | 41.527 | 1.00 | 0.00 | C |
| ATOM | 4982 | HE1 | LYS | A | 343 | 18.934 | 23.031 | 41.233 | 1.00 | 0.00 | H |
| ATOM | 4983 | HE2 | LYS | A | 343 | 20.181 | 22.518 | 42.153 | 1.00 | 0.00 | H |
| ATOM | 4984 | NZ | LYS | A | 343 | 20.742 | 23.267 | 40.362 | 1.00 | 0.00 | N |
| ATOM | 4985 | HZ1 | LYS | A | 343 | 20.718 | 22.380 | 39.900 | 1.00 | 0.00 | H |
| ATOM | 4986 | HZ2 | LYS | A | 343 | 20.430 | 23.979 | 39.733 | 1.00 | 0.00 | H |
| ATOM | 4987 | HZ3 | LYS | A | 343 | 21.678 | 23.466 | 40.653 | 1.00 | 0.00 | H |
| ATOM | 4988 | C | LYS | A | 343 | 19.543 | 26.781 | 46.761 | 1.00 | 0.00 | C |
| ATOM | 4989 | O | LYS | A | 343 | 20.487 | 26.552 | 47.505 | 1.00 | 0.00 | O |
| ATOM | 4990 | N | VAL | A | 344 | 18.787 | 27.888 | 46.792 | 1.00 | 0.00 | N |
| ATOM | 4991 | H | VAL | A | 344 | 17.969 | 27.956 | 46.221 | 1.00 | 0.00 | H |
| ATOM | 4992 | CA | VAL | A | 344 | 19.177 | 29.022 | 47.688 | 1.00 | 0.00 | C |
| ATOM | 4993 | HA | VAL | A | 344 | 20.155 | 29.089 | 47.489 | 1.00 | 0.00 | H |
| ATOM | 4994 | CB | VAL | A | 344 | 18.676 | 30.467 | 47.386 | 1.00 | 0.00 | C |
| ATOM | 4995 | HB | VAL | A | 344 | 17.676 | 30.444 | 47.397 | 1.00 | 0.00 | H |
| ATOM | 4996 | CG1 | VAL | A | 344 | 19.093 | 31.503 | 48.388 | 1.00 | 0.00 | C |
| ATOM | 4997 | HG11 | VAL | A | 344 | 18.730 | 32.394 | 48.117 | 1.00 | 0.00 | H |
| ATOM | 4998 | HG12 | VAL | A | 344 | 18.736 | 31.258 | 49.289 | 1.00 | 0.00 | H |
| ATOM | 4999 | HG13 | VAL | A | 344 | 20.091 | 31.547 | 48.426 | 1.00 | 0.00 | H |
| ATOM | 5000 | CG2 | VAL | A | 344 | 19.250 | 30.911 | 46.065 | 1.00 | 0.00 | C |
| ATOM | 5001 | HG21 | VAL | A | 344 | 18.936 | 31.838 | 45.860 | 1.00 | 0.00 | H |
| ATOM | 5002 | HG22 | VAL | A | 344 | 20.249 | 30.901 | 46.115 | 1.00 | 0.00 | H |
| ATOM | 5003 | HG23 | VAL | A | 344 | 18.947 | 30.289 | 45.344 | 1.00 | 0.00 | H |
| ATOM | 5004 | C | VAL | A | 344 | 18.795 | 28.624 | 49.145 | 1.00 | 0.00 | C |
| ATOM | 5005 | O | VAL | A | 344 | 19.476 | 29.126 | 50.068 | 1.00 | 0.00 | O |
| ATOM | 5006 | N | SER | A | 345 | 17.734 | 27.892 | 49.317 | 1.00 | 0.00 | N |
| ATOM | 5007 | H | SER | A | 345 | 17.157 | 27.667 | 48.532 | 1.00 | 0.00 | H |
| ATOM | 5008 | CA | SER | A | 345 | 17.369 | 27.386 | 50.665 | 1.00 | 0.00 | C |
| ATOM | 5009 | HA | SER | A | 345 | 17.346 | 28.227 | 51.205 | 1.00 | 0.00 | H |
| ATOM | 5010 | CB | SER | A | 345 | 15.954 | 26.749 | 50.636 | 1.00 | 0.00 | C |
| ATOM | 5011 | HB1 | SER | A | 345 | 15.905 | 26.029 | 49.944 | 1.00 | 0.00 | H |
| ATOM | 5012 | HB2 | SER | A | 345 | 15.259 | 27.442 | 50.445 | 1.00 | 0.00 | H |
| ATOM | 5013 | OG | SER | A | 345 | 15.578 | 26.150 | 51.840 | 1.00 | 0.00 | O |
| ATOM | 5014 | HG | SER | A | 345 | 14.661 | 25.763 | 51.748 | 1.00 | 0.00 | H |
| ATOM | 5015 | C | SER | A | 345 | 18.450 | 26.362 | 51.300 | 1.00 | 0.00 | C |
| ATOM | 5016 | O | SER | A | 345 | 18.882 | 26.459 | 52.477 | 1.00 | 0.00 | O |
| ATOM | 5017 | N | TYR | A | 346 | 18.860 | 25.365 | 50.511 | 1.00 | 0.00 | N |
| ATOM | 5018 | H | TYR | A | 346 | 18.341 | 25.205 | 49.671 | 1.00 | 0.00 | H |
| ATOM | 5019 | CA | TYR | A | 346 | 20.031 | 24.464 | 50.774 | 1.00 | 0.00 | C |

| | | | | | | | | | | | |
|---|---|---|---|---|---|---|---|---|---|---|---|
| ATOM | 5020 | HA | TYR | A | 346 | 19.777 | 24.010 | 51.628 | 1.00 | 0.00 | H |
| ATOM | 5021 | CB | TYR | A | 346 | 20.087 | 23.367 | 49.718 | 1.00 | 0.00 | C |
| ATOM | 5022 | HB1 | TYR | A | 346 | 20.181 | 23.783 | 48.814 | 1.00 | 0.00 | H |
| ATOM | 5023 | HB2 | TYR | A | 346 | 19.243 | 22.831 | 49.752 | 1.00 | 0.00 | H |
| ATOM | 5024 | CG | TYR | A | 346 | 21.236 | 22.444 | 49.930 | 1.00 | 0.00 | C |
| ATOM | 5025 | CD1 | TYR | A | 346 | 21.088 | 21.237 | 50.619 | 1.00 | 0.00 | C |
| ATOM | 5026 | HD1 | TYR | A | 346 | 20.183 | 21.003 | 50.974 | 1.00 | 0.00 | H |
| ATOM | 5027 | CE1 | TYR | A | 346 | 22.125 | 20.349 | 50.837 | 1.00 | 0.00 | C |
| ATOM | 5028 | HE1 | TYR | A | 346 | 21.991 | 19.491 | 51.334 | 1.00 | 0.00 | H |
| ATOM | 5029 | CZ | TYR | A | 346 | 23.353 | 20.712 | 50.332 | 1.00 | 0.00 | C |
| ATOM | 5030 | OH | TYR | A | 346 | 24.403 | 19.946 | 50.455 | 1.00 | 0.00 | O |
| ATOM | 5031 | HH | TYR | A | 346 | 25.190 | 20.387 | 50.024 | 1.00 | 0.00 | H |
| ATOM | 5032 | CE2 | TYR | A | 346 | 23.529 | 21.861 | 49.568 | 1.00 | 0.00 | C |
| ATOM | 5033 | HE2 | TYR | A | 346 | 24.415 | 22.036 | 49.140 | 1.00 | 0.00 | H |
| ATOM | 5034 | CD2 | TYR | A | 346 | 22.465 | 22.793 | 49.386 | 1.00 | 0.00 | C |
| ATOM | 5035 | HD2 | TYR | A | 346 | 22.597 | 23.653 | 48.893 | 1.00 | 0.00 | H |
| ATOM | 5036 | C | TYR | A | 346 | 21.459 | 25.219 | 51.040 | 1.00 | 0.00 | C |
| ATOM | 5037 | O | TYR | A | 346 | 22.264 | 24.898 | 51.925 | 1.00 | 0.00 | O |
| ATOM | 5038 | N | LEU | A | 347 | 21.595 | 26.307 | 50.267 | 1.00 | 0.00 | N |
| ATOM | 5039 | H | LEU | A | 347 | 20.880 | 26.515 | 49.600 | 1.00 | 0.00 | H |
| ATOM | 5040 | CA | LEU | A | 347 | 22.735 | 27.191 | 50.361 | 1.00 | 0.00 | C |
| ATOM | 5041 | HA | LEU | A | 347 | 23.544 | 26.604 | 50.317 | 1.00 | 0.00 | H |
| ATOM | 5042 | CB | LEU | A | 347 | 22.719 | 28.166 | 49.125 | 1.00 | 0.00 | C |
| ATOM | 5043 | HB1 | LEU | A | 347 | 21.894 | 28.726 | 49.200 | 1.00 | 0.00 | H |
| ATOM | 5044 | HB2 | LEU | A | 347 | 22.664 | 27.600 | 48.302 | 1.00 | 0.00 | H |
| ATOM | 5045 | CG | LEU | A | 347 | 23.786 | 29.149 | 48.808 | 1.00 | 0.00 | C |
| ATOM | 5046 | HG | LEU | A | 347 | 24.613 | 28.605 | 48.950 | 1.00 | 0.00 | H |
| ATOM | 5047 | CD1 | LEU | A | 347 | 23.749 | 29.670 | 47.348 | 1.00 | 0.00 | C |
| ATOM | 5048 | HD11 | LEU | A | 347 | 24.492 | 30.324 | 47.209 | 1.00 | 0.00 | H |
| ATOM | 5049 | HD12 | LEU | A | 347 | 23.858 | 28.902 | 46.717 | 1.00 | 0.00 | H |
| ATOM | 5050 | HD13 | LEU | A | 347 | 22.873 | 30.120 | 47.177 | 1.00 | 0.00 | H |
| ATOM | 5051 | CD2 | LEU | A | 347 | 23.700 | 30.394 | 49.746 | 1.00 | 0.00 | C |
| ATOM | 5052 | HD21 | LEU | A | 347 | 24.428 | 31.039 | 49.512 | 1.00 | 0.00 | H |
| ATOM | 5053 | HD22 | LEU | A | 347 | 22.811 | 30.838 | 49.630 | 1.00 | 0.00 | H |
| ATOM | 5054 | HD23 | LEU | A | 347 | 23.806 | 30.103 | 50.697 | 1.00 | 0.00 | H |
| ATOM | 5055 | C | LEU | A | 347 | 22.881 | 27.833 | 51.746 | 1.00 | 0.00 | C |
| ATOM | 5056 | O | LEU | A | 347 | 23.995 | 27.916 | 52.226 | 1.00 | 0.00 | O |
| ATOM | 5057 | N | LYS | A | 348 | 21.742 | 28.344 | 52.221 | 1.00 | 0.00 | N |
| ATOM | 5058 | H | LYS | A | 348 | 20.924 | 28.311 | 51.646 | 1.00 | 0.00 | H |
| ATOM | 5059 | CA | LYS | A | 348 | 21.645 | 28.938 | 53.521 | 1.00 | 0.00 | C |
| ATOM | 5060 | HA | LYS | A | 348 | 22.470 | 29.491 | 53.636 | 1.00 | 0.00 | H |
| ATOM | 5061 | CB | LYS | A | 348 | 20.345 | 29.777 | 53.557 | 1.00 | 0.00 | C |
| ATOM | 5062 | HB1 | LYS | A | 348 | 20.171 | 29.995 | 54.517 | 1.00 | 0.00 | H |
| ATOM | 5063 | HB2 | LYS | A | 348 | 19.614 | 29.189 | 53.211 | 1.00 | 0.00 | H |
| ATOM | 5064 | CG | LYS | A | 348 | 20.294 | 31.126 | 52.746 | 1.00 | 0.00 | C |
| ATOM | 5065 | HG1 | LYS | A | 348 | 20.492 | 30.922 | 51.787 | 1.00 | 0.00 | H |
| ATOM | 5066 | HG2 | LYS | A | 348 | 20.998 | 31.736 | 53.110 | 1.00 | 0.00 | H |
| ATOM | 5067 | CD | LYS | A | 348 | 18.906 | 31.875 | 52.816 | 1.00 | 0.00 | C |
| ATOM | 5068 | HD1 | LYS | A | 348 | 18.990 | 32.775 | 52.389 | 1.00 | 0.00 | H |
| ATOM | 5069 | HD2 | LYS | A | 348 | 18.633 | 31.984 | 53.772 | 1.00 | 0.00 | H |
| ATOM | 5070 | CE | LYS | A | 348 | 17.796 | 31.140 | 52.109 | 1.00 | 0.00 | C |
| ATOM | 5071 | HE1 | LYS | A | 348 | 17.563 | 30.317 | 52.627 | 1.00 | 0.00 | H |
| ATOM | 5072 | HE2 | LYS | A | 348 | 18.102 | 30.882 | 51.193 | 1.00 | 0.00 | H |
| ATOM | 5073 | NZ | LYS | A | 348 | 16.576 | 32.019 | 51.997 | 1.00 | 0.00 | N |

| | | | | | | | | | | | |
|---|---|---|---|---|---|---|---|---|---|---|---|
| ATOM | 5074 | HZ1 | LYS | A | 348 | 15.848 | 31.520 | 51.527 | 1.00 | 0.00 | H |
| ATOM | 5075 | HZ2 | LYS | A | 348 | 16.266 | 32.277 | 52.912 | 1.00 | 0.00 | H |
| ATOM | 5076 | HZ3 | LYS | A | 348 | 16.805 | 32.843 | 51.478 | 1.00 | 0.00 | H |
| ATOM | 5077 | C | LYS | A | 348 | 21.702 | 27.925 | 54.625 | 1.00 | 0.00 | C |
| ATOM | 5078 | O | LYS | A | 348 | 22.351 | 28.153 | 55.650 | 1.00 | 0.00 | O |
| ATOM | 5079 | N | GLN | A | 349 | 21.084 | 26.699 | 54.389 | 1.00 | 0.00 | N |
| ATOM | 5080 | H | GLN | A | 349 | 20.524 | 26.604 | 53.566 | 1.00 | 0.00 | H |
| ATOM | 5081 | CA | GLN | A | 349 | 21.203 | 25.579 | 55.248 | 1.00 | 0.00 | C |
| ATOM | 5082 | HA | GLN | A | 349 | 20.812 | 26.011 | 56.061 | 1.00 | 0.00 | H |
| ATOM | 5083 | CB | GLN | A | 349 | 20.332 | 24.413 | 54.751 | 1.00 | 0.00 | C |
| ATOM | 5084 | HB1 | GLN | A | 349 | 20.673 | 24.122 | 53.857 | 1.00 | 0.00 | H |
| ATOM | 5085 | HB2 | GLN | A | 349 | 19.391 | 24.738 | 54.659 | 1.00 | 0.00 | H |
| ATOM | 5086 | CG | GLN | A | 349 | 20.358 | 23.197 | 55.728 | 1.00 | 0.00 | C |
| ATOM | 5087 | HG1 | GLN | A | 349 | 19.432 | 22.819 | 55.756 | 1.00 | 0.00 | H |
| ATOM | 5088 | HG2 | GLN | A | 349 | 20.602 | 23.548 | 56.632 | 1.00 | 0.00 | H |
| ATOM | 5089 | CD | GLN | A | 349 | 21.318 | 22.017 | 55.429 | 1.00 | 0.00 | C |
| ATOM | 5090 | OE1 | GLN | A | 349 | 21.161 | 21.333 | 54.400 | 1.00 | 0.00 | O |
| ATOM | 5091 | NE2 | GLN | A | 349 | 22.263 | 21.714 | 56.300 | 1.00 | 0.00 | N |
| ATOM | 5092 | HE21 | GLN | A | 349 | 22.354 | 22.244 | 57.143 | 1.00 | 0.00 | H |
| ATOM | 5093 | HE22 | GLN | A | 349 | 22.887 | 20.955 | 56.116 | 1.00 | 0.00 | H |
| ATOM | 5094 | C | GLN | A | 349 | 22.636 | 25.135 | 55.683 | 1.00 | 0.00 | C |
| ATOM | 5095 | O | GLN | A | 349 | 22.907 | 24.913 | 56.840 | 1.00 | 0.00 | O |
| ATOM | 5096 | N | LYS | A | 350 | 23.496 | 24.978 | 54.671 | 1.00 | 0.00 | N |
| ATOM | 5097 | H | LYS | A | 350 | 23.142 | 25.063 | 53.740 | 1.00 | 0.00 | H |
| ATOM | 5098 | CA | LYS | A | 350 | 24.939 | 24.687 | 54.839 | 1.00 | 0.00 | C |
| ATOM | 5099 | HA | LYS | A | 350 | 25.101 | 24.078 | 55.615 | 1.00 | 0.00 | H |
| ATOM | 5100 | CB | LYS | A | 350 | 25.330 | 24.153 | 53.452 | 1.00 | 0.00 | C |
| ATOM | 5101 | HB1 | LYS | A | 350 | 26.325 | 24.220 | 53.379 | 1.00 | 0.00 | H |
| ATOM | 5102 | HB2 | LYS | A | 350 | 24.905 | 24.752 | 52.773 | 1.00 | 0.00 | H |
| ATOM | 5103 | CG | LYS | A | 350 | 24.946 | 22.714 | 53.088 | 1.00 | 0.00 | C |
| ATOM | 5104 | HG1 | LYS | A | 350 | 25.288 | 22.541 | 52.164 | 1.00 | 0.00 | H |
| ATOM | 5105 | HG2 | LYS | A | 350 | 23.947 | 22.665 | 53.089 | 1.00 | 0.00 | H |
| ATOM | 5106 | CD | LYS | A | 350 | 25.506 | 21.593 | 54.050 | 1.00 | 0.00 | C |
| ATOM | 5107 | HD1 | LYS | A | 350 | 25.057 | 21.628 | 54.943 | 1.00 | 0.00 | H |
| ATOM | 5108 | HD2 | LYS | A | 350 | 26.495 | 21.684 | 54.170 | 1.00 | 0.00 | H |
| ATOM | 5109 | CE | LYS | A | 350 | 25.217 | 20.274 | 53.418 | 1.00 | 0.00 | C |
| ATOM | 5110 | HE1 | LYS | A | 350 | 25.805 | 20.167 | 52.616 | 1.00 | 0.00 | H |
| ATOM | 5111 | HE2 | LYS | A | 350 | 24.259 | 20.258 | 53.133 | 1.00 | 0.00 | H |
| ATOM | 5112 | NZ | LYS | A | 350 | 25.470 | 19.135 | 54.375 | 1.00 | 0.00 | N |
| ATOM | 5113 | HZ1 | LYS | A | 350 | 25.267 | 18.267 | 53.921 | 1.00 | 0.00 | H |
| ATOM | 5114 | HZ2 | LYS | A | 350 | 26.428 | 19.146 | 54.660 | 1.00 | 0.00 | H |
| ATOM | 5115 | HZ3 | LYS | A | 350 | 24.882 | 19.237 | 55.177 | 1.00 | 0.00 | H |
| ATOM | 5116 | C | LYS | A | 350 | 25.729 | 25.914 | 55.141 | 1.00 | 0.00 | C |
| ATOM | 5117 | O | LYS | A | 350 | 26.858 | 25.759 | 55.754 | 1.00 | 0.00 | O |
| ATOM | 5118 | N | GLY | A | 351 | 25.305 | 27.160 | 54.859 | 1.00 | 0.00 | N |
| ATOM | 5119 | H | GLY | A | 351 | 24.392 | 27.255 | 54.462 | 1.00 | 0.00 | H |
| ATOM | 5120 | CA | GLY | A | 351 | 26.067 | 28.385 | 55.084 | 1.00 | 0.00 | C |
| ATOM | 5121 | HA1 | GLY | A | 351 | 26.407 | 28.370 | 56.024 | 1.00 | 0.00 | H |
| ATOM | 5122 | HA2 | GLY | A | 351 | 25.448 | 29.161 | 54.966 | 1.00 | 0.00 | H |
| ATOM | 5123 | C | GLY | A | 351 | 27.233 | 28.505 | 54.117 | 1.00 | 0.00 | C |
| ATOM | 5124 | O | GLY | A | 351 | 28.372 | 28.797 | 54.594 | 1.00 | 0.00 | O |
| ATOM | 5125 | N | LEU | A | 352 | 27.029 | 28.149 | 52.843 | 1.00 | 0.00 | N |
| ATOM | 5126 | H | LEU | A | 352 | 26.232 | 27.575 | 52.656 | 1.00 | 0.00 | H |
| ATOM | 5127 | CA | LEU | A | 352 | 27.858 | 28.518 | 51.703 | 1.00 | 0.00 | C |

| | | | | | | | | | | | |
|---|---|---|---|---|---|---|---|---|---|---|---|
| ATOM | 5128 | HA | LEU | A | 352 | 28.766 | 28.141 | 51.884 | 1.00 | 0.00 | H |
| ATOM | 5129 | CB | LEU | A | 352 | 27.466 | 27.878 | 50.431 | 1.00 | 0.00 | C |
| ATOM | 5130 | HB1 | LEU | A | 352 | 28.257 | 27.959 | 49.825 | 1.00 | 0.00 | H |
| ATOM | 5131 | HB2 | LEU | A | 352 | 26.711 | 28.424 | 50.068 | 1.00 | 0.00 | H |
| ATOM | 5132 | CG | LEU | A | 352 | 27.024 | 26.375 | 50.493 | 1.00 | 0.00 | C |
| ATOM | 5133 | HG | LEU | A | 352 | 26.117 | 26.378 | 50.914 | 1.00 | 0.00 | H |
| ATOM | 5134 | CD1 | LEU | A | 352 | 26.919 | 25.688 | 49.136 | 1.00 | 0.00 | C |
| ATOM | 5135 | HD11 | LEU | A | 352 | 26.633 | 24.738 | 49.264 | 1.00 | 0.00 | H |
| ATOM | 5136 | HD12 | LEU | A | 352 | 26.245 | 26.165 | 48.572 | 1.00 | 0.00 | H |
| ATOM | 5137 | HD13 | LEU | A | 352 | 27.810 | 25.710 | 48.682 | 1.00 | 0.00 | H |
| ATOM | 5138 | CD2 | LEU | A | 352 | 28.061 | 25.555 | 51.275 | 1.00 | 0.00 | C |
| ATOM | 5139 | HD21 | LEU | A | 352 | 27.772 | 24.598 | 51.310 | 1.00 | 0.00 | H |
| ATOM | 5140 | HD22 | LEU | A | 352 | 28.949 | 25.619 | 50.819 | 1.00 | 0.00 | H |
| ATOM | 5141 | HD23 | LEU | A | 352 | 28.137 | 25.914 | 52.205 | 1.00 | 0.00 | H |
| ATOM | 5142 | C | LEU | A | 352 | 27.942 | 29.999 | 51.609 | 1.00 | 0.00 | C |
| ATOM | 5143 | O | LEU | A | 352 | 26.988 | 30.691 | 51.988 | 1.00 | 0.00 | O |
| ATOM | 5144 | N | GLY | A | 353 | 28.984 | 30.614 | 51.046 | 1.00 | 0.00 | N |
| ATOM | 5145 | H | GLY | A | 353 | 29.672 | 30.034 | 50.610 | 1.00 | 0.00 | H |
| ATOM | 5146 | CA | GLY | A | 353 | 29.213 | 32.069 | 51.008 | 1.00 | 0.00 | C |
| ATOM | 5147 | HA1 | GLY | A | 353 | 30.174 | 32.181 | 50.754 | 1.00 | 0.00 | H |
| ATOM | 5148 | HA2 | GLY | A | 353 | 29.078 | 32.385 | 51.947 | 1.00 | 0.00 | H |
| ATOM | 5149 | C | GLY | A | 353 | 28.392 | 32.979 | 50.099 | 1.00 | 0.00 | C |
| ATOM | 5150 | O | GLY | A | 353 | 28.566 | 34.173 | 50.045 | 1.00 | 0.00 | O |
| ATOM | 5151 | N | GLY | A | 354 | 27.456 | 32.389 | 49.315 | 1.00 | 0.00 | N |
| ATOM | 5152 | H | GLY | A | 354 | 27.350 | 31.405 | 49.458 | 1.00 | 0.00 | H |
| ATOM | 5153 | CA | GLY | A | 354 | 26.590 | 32.949 | 48.310 | 1.00 | 0.00 | C |
| ATOM | 5154 | HA1 | GLY | A | 354 | 26.889 | 33.887 | 48.133 | 1.00 | 0.00 | H |
| ATOM | 5155 | HA2 | GLY | A | 354 | 25.659 | 32.959 | 48.676 | 1.00 | 0.00 | H |
| ATOM | 5156 | C | GLY | A | 354 | 26.593 | 32.205 | 47.045 | 1.00 | 0.00 | C |
| ATOM | 5157 | O | GLY | A | 354 | 27.048 | 31.052 | 47.023 | 1.00 | 0.00 | O |
| ATOM | 5158 | N | ALA | A | 355 | 26.140 | 32.882 | 45.952 | 1.00 | 0.00 | N |
| ATOM | 5159 | H | ALA | A | 355 | 25.955 | 33.864 | 45.994 | 1.00 | 0.00 | H |
| ATOM | 5160 | CA | ALA | A | 355 | 25.934 | 32.129 | 44.719 | 1.00 | 0.00 | C |
| ATOM | 5161 | HA | ALA | A | 355 | 26.349 | 31.249 | 44.953 | 1.00 | 0.00 | H |
| ATOM | 5162 | CB | ALA | A | 355 | 24.372 | 31.974 | 44.387 | 1.00 | 0.00 | C |
| ATOM | 5163 | HB1 | ALA | A | 355 | 24.261 | 31.455 | 43.539 | 1.00 | 0.00 | H |
| ATOM | 5164 | HB2 | ALA | A | 355 | 23.919 | 31.490 | 45.135 | 1.00 | 0.00 | H |
| ATOM | 5165 | HB3 | ALA | A | 355 | 23.963 | 32.880 | 44.279 | 1.00 | 0.00 | H |
| ATOM | 5166 | C | ALA | A | 355 | 26.731 | 32.717 | 43.503 | 1.00 | 0.00 | C |
| ATOM | 5167 | O | ALA | A | 355 | 27.414 | 33.754 | 43.610 | 1.00 | 0.00 | O |
| ATOM | 5168 | N | MET | A | 356 | 26.671 | 31.919 | 42.399 | 1.00 | 0.00 | N |
| ATOM | 5169 | H | MET | A | 356 | 26.124 | 31.083 | 42.430 | 1.00 | 0.00 | H |
| ATOM | 5170 | CA | MET | A | 356 | 27.385 | 32.256 | 41.176 | 1.00 | 0.00 | C |
| ATOM | 5171 | HA | MET | A | 356 | 27.548 | 33.243 | 41.174 | 1.00 | 0.00 | H |
| ATOM | 5172 | CB | MET | A | 356 | 28.804 | 31.652 | 41.118 | 1.00 | 0.00 | C |
| ATOM | 5173 | HB1 | MET | A | 356 | 28.756 | 30.710 | 41.451 | 1.00 | 0.00 | H |
| ATOM | 5174 | HB2 | MET | A | 356 | 29.396 | 32.189 | 41.720 | 1.00 | 0.00 | H |
| ATOM | 5175 | CG | MET | A | 356 | 29.446 | 31.627 | 39.724 | 1.00 | 0.00 | C |
| ATOM | 5176 | HG1 | MET | A | 356 | 30.428 | 31.691 | 39.902 | 1.00 | 0.00 | H |
| ATOM | 5177 | HG2 | MET | A | 356 | 29.130 | 32.471 | 39.290 | 1.00 | 0.00 | H |
| ATOM | 5178 | SD | MET | A | 356 | 29.029 | 30.158 | 38.736 | 1.00 | 0.00 | S |
| ATOM | 5179 | CE | MET | A | 356 | 30.225 | 28.972 | 39.423 | 1.00 | 0.00 | C |
| ATOM | 5180 | HE1 | MET | A | 356 | 30.117 | 28.088 | 38.967 | 1.00 | 0.00 | H |
| ATOM | 5181 | HE2 | MET | A | 356 | 31.154 | 29.312 | 39.276 | 1.00 | 0.00 | H |

| | | | | | | | | | | |
|---|---|---|---|---|---|---|---|---|---|---|
| ATOM | 5182 | HE3 | MET | A | 356 | 30.061 | 28.864 | 40.404 | 1.00 | 0.00 | H |
| ATOM | 5183 | C | MET | A | 356 | 26.447 | 31.783 | 40.014 | 1.00 | 0.00 | C |
| ATOM | 5184 | O | MET | A | 356 | 25.819 | 30.713 | 40.218 | 1.00 | 0.00 | O |
| ATOM | 5185 | N | VAL | A | 357 | 26.443 | 32.508 | 38.855 | 1.00 | 0.00 | N |
| ATOM | 5186 | H | VAL | A | 357 | 27.040 | 33.309 | 38.812 | 1.00 | 0.00 | H |
| ATOM | 5187 | CA | VAL | A | 357 | 25.645 | 32.209 | 37.686 | 1.00 | 0.00 | C |
| ATOM | 5188 | HA | VAL | A | 357 | 25.305 | 31.315 | 37.979 | 1.00 | 0.00 | H |
| ATOM | 5189 | CB | VAL | A | 357 | 24.454 | 33.161 | 37.436 | 1.00 | 0.00 | C |
| ATOM | 5190 | HB | VAL | A | 357 | 24.812 | 34.083 | 37.288 | 1.00 | 0.00 | H |
| ATOM | 5191 | CG1 | VAL | A | 357 | 23.717 | 32.774 | 36.167 | 1.00 | 0.00 | C |
| ATOM | 5192 | HG11 | VAL | A | 357 | 22.951 | 33.400 | 36.022 | 1.00 | 0.00 | H |
| ATOM | 5193 | HG12 | VAL | A | 357 | 24.343 | 32.827 | 35.389 | 1.00 | 0.00 | H |
| ATOM | 5194 | HG13 | VAL | A | 357 | 23.372 | 31.840 | 36.254 | 1.00 | 0.00 | H |
| ATOM | 5195 | CG2 | VAL | A | 357 | 23.503 | 33.221 | 38.660 | 1.00 | 0.00 | C |
| ATOM | 5196 | HG21 | VAL | A | 357 | 22.746 | 33.844 | 38.464 | 1.00 | 0.00 | H |
| ATOM | 5197 | HG22 | VAL | A | 357 | 23.142 | 32.307 | 38.847 | 1.00 | 0.00 | H |
| ATOM | 5198 | HG23 | VAL | A | 357 | 24.007 | 33.550 | 39.458 | 1.00 | 0.00 | H |
| ATOM | 5199 | C | VAL | A | 357 | 26.463 | 32.074 | 36.401 | 1.00 | 0.00 | C |
| ATOM | 5200 | O | VAL | A | 357 | 27.048 | 33.071 | 35.966 | 1.00 | 0.00 | O |
| ATOM | 5201 | N | TRP | A | 358 | 26.332 | 30.926 | 35.734 | 1.00 | 0.00 | N |
| ATOM | 5202 | H | TRP | A | 358 | 26.003 | 30.127 | 36.237 | 1.00 | 0.00 | H |
| ATOM | 5203 | CA | TRP | A | 358 | 26.640 | 30.764 | 34.313 | 1.00 | 0.00 | C |
| ATOM | 5204 | HA | TRP | A | 358 | 27.123 | 31.587 | 34.014 | 1.00 | 0.00 | H |
| ATOM | 5205 | CB | TRP | A | 358 | 27.564 | 29.560 | 34.106 | 1.00 | 0.00 | C |
| ATOM | 5206 | HB1 | TRP | A | 358 | 27.038 | 28.718 | 34.229 | 1.00 | 0.00 | H |
| ATOM | 5207 | HB2 | TRP | A | 358 | 28.302 | 29.593 | 34.780 | 1.00 | 0.00 | H |
| ATOM | 5208 | CG | TRP | A | 358 | 28.180 | 29.528 | 32.744 | 1.00 | 0.00 | C |
| ATOM | 5209 | CD1 | TRP | A | 358 | 27.564 | 29.181 | 31.587 | 1.00 | 0.00 | C |
| ATOM | 5210 | HD1 | TRP | A | 358 | 26.626 | 28.843 | 31.510 | 1.00 | 0.00 | H |
| ATOM | 5211 | NE1 | TRP | A | 358 | 28.434 | 29.368 | 30.531 | 1.00 | 0.00 | N |
| ATOM | 5212 | HE1 | TRP | A | 358 | 28.208 | 29.177 | 29.576 | 1.00 | 0.00 | H |
| ATOM | 5213 | CE2 | TRP | A | 358 | 29.649 | 29.849 | 30.955 | 1.00 | 0.00 | C |
| ATOM | 5214 | CZ2 | TRP | A | 358 | 30.820 | 30.197 | 30.275 | 1.00 | 0.00 | C |
| ATOM | 5215 | HZ2 | TRP | A | 358 | 30.868 | 30.124 | 29.279 | 1.00 | 0.00 | H |
| ATOM | 5216 | CH2 | TRP | A | 358 | 31.927 | 30.650 | 31.008 | 1.00 | 0.00 | C |
| ATOM | 5217 | HH2 | TRP | A | 358 | 32.765 | 30.921 | 30.534 | 1.00 | 0.00 | H |
| ATOM | 5218 | CZ3 | TRP | A | 358 | 31.855 | 30.722 | 32.410 | 1.00 | 0.00 | C |
| ATOM | 5219 | HZ3 | TRP | A | 358 | 32.649 | 31.032 | 32.933 | 1.00 | 0.00 | H |
| ATOM | 5220 | CE3 | TRP | A | 358 | 30.673 | 30.361 | 33.082 | 1.00 | 0.00 | C |
| ATOM | 5221 | HE3 | TRP | A | 358 | 30.641 | 30.409 | 34.080 | 1.00 | 0.00 | H |
| ATOM | 5222 | CD2 | TRP | A | 358 | 29.532 | 29.932 | 32.371 | 1.00 | 0.00 | C |
| ATOM | 5223 | C | TRP | A | 358 | 25.321 | 30.608 | 33.535 | 1.00 | 0.00 | C |
| ATOM | 5224 | O | TRP | A | 358 | 24.612 | 29.630 | 33.745 | 1.00 | 0.00 | O |
| ATOM | 5225 | N | ALA | A | 359 | 24.884 | 31.544 | 32.693 | 1.00 | 0.00 | N |
| ATOM | 5226 | H | ALA | A | 359 | 24.031 | 31.325 | 32.219 | 1.00 | 0.00 | H |
| ATOM | 5227 | CA | ALA | A | 359 | 25.467 | 32.846 | 32.359 | 1.00 | 0.00 | C |
| ATOM | 5228 | HA | ALA | A | 359 | 26.013 | 33.119 | 33.151 | 1.00 | 0.00 | H |
| ATOM | 5229 | CB | ALA | A | 359 | 26.359 | 32.685 | 31.119 | 1.00 | 0.00 | C |
| ATOM | 5230 | HB1 | ALA | A | 359 | 26.763 | 33.569 | 30.882 | 1.00 | 0.00 | H |
| ATOM | 5231 | HB2 | ALA | A | 359 | 27.087 | 32.028 | 31.315 | 1.00 | 0.00 | H |
| ATOM | 5232 | HB3 | ALA | A | 359 | 25.808 | 32.354 | 30.353 | 1.00 | 0.00 | H |
| ATOM | 5233 | C | ALA | A | 359 | 24.380 | 33.920 | 32.137 | 1.00 | 0.00 | C |
| ATOM | 5234 | O | ALA | A | 359 | 23.193 | 33.627 | 31.989 | 1.00 | 0.00 | O |
| ATOM | 5235 | N | LEU | A | 360 | 24.810 | 35.179 | 32.071 | 1.00 | 0.00 | N |

| | | | | | | | | | | | |
|---|---|---|---|---|---|---|---|---|---|---|---|
| ATOM | 5236 | H | LEU | A | 360 | 25.797 | 35.329 | 32.121 | 1.00 | 0.00 | H |
| ATOM | 5237 | CA | LEU | A | 360 | 23.981 | 36.328 | 31.935 | 1.00 | 0.00 | C |
| ATOM | 5238 | HA | LEU | A | 360 | 23.355 | 36.121 | 32.687 | 1.00 | 0.00 | H |
| ATOM | 5239 | CB | LEU | A | 360 | 24.731 | 37.656 | 32.079 | 1.00 | 0.00 | C |
| ATOM | 5240 | HB1 | LEU | A | 360 | 24.095 | 38.405 | 31.894 | 1.00 | 0.00 | H |
| ATOM | 5241 | HB2 | LEU | A | 360 | 25.475 | 37.678 | 31.412 | 1.00 | 0.00 | H |
| ATOM | 5242 | CG | LEU | A | 360 | 25.319 | 37.833 | 33.494 | 1.00 | 0.00 | C |
| ATOM | 5243 | HG | LEU | A | 360 | 25.549 | 36.919 | 33.829 | 1.00 | 0.00 | H |
| ATOM | 5244 | CD1 | LEU | A | 360 | 26.623 | 38.547 | 33.565 | 1.00 | 0.00 | C |
| ATOM | 5245 | HD11 | LEU | A | 360 | 26.917 | 38.610 | 34.519 | 1.00 | 0.00 | H |
| ATOM | 5246 | HD12 | LEU | A | 360 | 27.308 | 38.044 | 33.037 | 1.00 | 0.00 | H |
| ATOM | 5247 | HD13 | LEU | A | 360 | 26.520 | 39.467 | 33.187 | 1.00 | 0.00 | H |
| ATOM | 5248 | CD2 | LEU | A | 360 | 24.290 | 38.542 | 34.382 | 1.00 | 0.00 | C |
| ATOM | 5249 | HD21 | LEU | A | 360 | 24.667 | 38.659 | 35.301 | 1.00 | 0.00 | H |
| ATOM | 5250 | HD22 | LEU | A | 360 | 24.075 | 39.438 | 33.993 | 1.00 | 0.00 | H |
| ATOM | 5251 | HD23 | LEU | A | 360 | 23.457 | 37.992 | 34.431 | 1.00 | 0.00 | H |
| ATOM | 5252 | C | LEU | A | 360 | 23.290 | 36.443 | 30.588 | 1.00 | 0.00 | C |
| ATOM | 5253 | O | LEU | A | 360 | 22.230 | 37.057 | 30.458 | 1.00 | 0.00 | O |
| ATOM | 5254 | N | ASP | A | 361 | 23.794 | 35.734 | 29.565 | 1.00 | 0.00 | N |
| ATOM | 5255 | H | ASP | A | 361 | 24.644 | 35.229 | 29.712 | 1.00 | 0.00 | H |
| ATOM | 5256 | CA | ASP | A | 361 | 23.167 | 35.662 | 28.252 | 1.00 | 0.00 | C |
| ATOM | 5257 | HA | ASP | A | 361 | 22.579 | 36.470 | 28.225 | 1.00 | 0.00 | H |
| ATOM | 5258 | CB | ASP | A | 361 | 24.346 | 35.608 | 27.232 | 1.00 | 0.00 | C |
| ATOM | 5259 | HB1 | ASP | A | 361 | 24.812 | 36.493 | 27.237 | 1.00 | 0.00 | H |
| ATOM | 5260 | HB2 | ASP | A | 361 | 23.971 | 35.433 | 26.321 | 1.00 | 0.00 | H |
| ATOM | 5261 | CG | ASP | A | 361 | 25.394 | 34.528 | 27.520 | 1.00 | 0.00 | C |
| ATOM | 5262 | OD1 | ASP | A | 361 | 25.314 | 33.438 | 26.915 | 1.00 | 0.00 | O |
| ATOM | 5263 | OD2 | ASP | A | 361 | 26.324 | 34.802 | 28.318 | 1.00 | 0.00 | O |
| ATOM | 5264 | C | ASP | A | 361 | 22.215 | 34.471 | 27.998 | 1.00 | 0.00 | C |
| ATOM | 5265 | O | ASP | A | 361 | 21.887 | 34.230 | 26.846 | 1.00 | 0.00 | O |
| ATOM | 5266 | N | LEU | A | 362 | 21.839 | 33.755 | 29.061 | 1.00 | 0.00 | N |
| ATOM | 5267 | H | LEU | A | 362 | 22.065 | 34.134 | 29.959 | 1.00 | 0.00 | H |
| ATOM | 5268 | CA | LEU | A | 362 | 21.123 | 32.460 | 29.049 | 1.00 | 0.00 | C |
| ATOM | 5269 | HA | LEU | A | 362 | 20.858 | 32.379 | 28.088 | 1.00 | 0.00 | H |
| ATOM | 5270 | CB | LEU | A | 362 | 22.069 | 31.310 | 29.452 | 1.00 | 0.00 | C |
| ATOM | 5271 | HB1 | LEU | A | 362 | 21.545 | 30.459 | 29.432 | 1.00 | 0.00 | H |
| ATOM | 5272 | HB2 | LEU | A | 362 | 22.386 | 31.485 | 30.384 | 1.00 | 0.00 | H |
| ATOM | 5273 | CG | LEU | A | 362 | 23.309 | 31.122 | 28.558 | 1.00 | 0.00 | C |
| ATOM | 5274 | HG | LEU | A | 362 | 23.780 | 32.004 | 28.532 | 1.00 | 0.00 | H |
| ATOM | 5275 | CD1 | LEU | A | 362 | 24.201 | 30.022 | 29.138 | 1.00 | 0.00 | C |
| ATOM | 5276 | HD11 | LEU | A | 362 | 25.005 | 29.903 | 28.555 | 1.00 | 0.00 | H |
| ATOM | 5277 | HD12 | LEU | A | 362 | 24.492 | 30.280 | 30.059 | 1.00 | 0.00 | H |
| ATOM | 5278 | HD13 | LEU | A | 362 | 23.689 | 29.164 | 29.179 | 1.00 | 0.00 | H |
| ATOM | 5279 | CD2 | LEU | A | 362 | 22.956 | 30.723 | 27.122 | 1.00 | 0.00 | C |
| ATOM | 5280 | HD21 | LEU | A | 362 | 23.796 | 30.615 | 26.590 | 1.00 | 0.00 | H |
| ATOM | 5281 | HD22 | LEU | A | 362 | 22.453 | 29.858 | 27.130 | 1.00 | 0.00 | H |
| ATOM | 5282 | HD23 | LEU | A | 362 | 22.388 | 31.435 | 26.708 | 1.00 | 0.00 | H |
| ATOM | 5283 | C | LEU | A | 362 | 19.802 | 32.419 | 29.870 | 1.00 | 0.00 | C |
| ATOM | 5284 | O | LEU | A | 362 | 18.984 | 31.486 | 29.764 | 1.00 | 0.00 | O |
| ATOM | 5285 | N | ASP | A | 363 | 19.552 | 33.468 | 30.664 | 1.00 | 0.00 | N |
| ATOM | 5286 | H | ASP | A | 363 | 20.301 | 34.109 | 30.831 | 1.00 | 0.00 | H |
| ATOM | 5287 | CA | ASP | A | 363 | 18.269 | 33.749 | 31.305 | 1.00 | 0.00 | C |
| ATOM | 5288 | HA | ASP | A | 363 | 17.809 | 32.883 | 31.500 | 1.00 | 0.00 | H |
| ATOM | 5289 | CB | ASP | A | 363 | 18.548 | 34.494 | 32.606 | 1.00 | 0.00 | C |

| | | | | | | | | | | | |
|---|---|---|---|---|---|---|---|---|---|---|---|
| ATOM | 5290 | HB1 | ASP | A | 363 | 18.804 | 35.435 | 32.386 | 1.00 | 0.00 | H |
| ATOM | 5291 | HB2 | ASP | A | 363 | 19.307 | 34.043 | 33.075 | 1.00 | 0.00 | H |
| ATOM | 5292 | CG | ASP | A | 363 | 17.377 | 34.539 | 33.552 | 1.00 | 0.00 | C |
| ATOM | 5293 | OD1 | ASP | A | 363 | 16.342 | 33.867 | 33.377 | 1.00 | 0.00 | O |
| ATOM | 5294 | OD2 | ASP | A | 363 | 17.437 | 35.331 | 34.514 | 1.00 | 0.00 | O |
| ATOM | 5295 | C | ASP | A | 363 | 17.389 | 34.570 | 30.356 | 1.00 | 0.00 | C |
| ATOM | 5296 | O | ASP | A | 363 | 17.921 | 34.878 | 29.298 | 1.00 | 0.00 | O |
| ATOM | 5297 | N | ASP | A | 364 | 16.160 | 34.860 | 30.738 | 1.00 | 0.00 | N |
| ATOM | 5298 | H | ASP | A | 364 | 15.808 | 34.438 | 31.573 | 1.00 | 0.00 | H |
| ATOM | 5299 | CA | ASP | A | 364 | 15.277 | 35.783 | 29.986 | 1.00 | 0.00 | C |
| ATOM | 5300 | HA | ASP | A | 364 | 15.421 | 35.633 | 29.008 | 1.00 | 0.00 | H |
| ATOM | 5301 | CB | ASP | A | 364 | 13.709 | 35.515 | 30.228 | 1.00 | 0.00 | C |
| ATOM | 5302 | HB1 | ASP | A | 364 | 13.437 | 35.816 | 31.142 | 1.00 | 0.00 | H |
| ATOM | 5303 | HB2 | ASP | A | 364 | 13.497 | 34.544 | 30.121 | 1.00 | 0.00 | H |
| ATOM | 5304 | CG | ASP | A | 364 | 12.853 | 36.233 | 29.283 | 1.00 | 0.00 | C |
| ATOM | 5305 | OD1 | ASP | A | 364 | 13.329 | 37.008 | 28.506 | 1.00 | 0.00 | O |
| ATOM | 5306 | OD2 | ASP | A | 364 | 11.643 | 36.035 | 29.263 | 1.00 | 0.00 | O |
| ATOM | 5307 | C | ASP | A | 364 | 15.759 | 37.185 | 30.371 | 1.00 | 0.00 | C |
| ATOM | 5308 | O | ASP | A | 364 | 15.134 | 37.973 | 31.106 | 1.00 | 0.00 | O |
| ATOM | 5309 | N | PHE | A | 365 | 16.699 | 37.711 | 29.621 | 1.00 | 0.00 | N |
| ATOM | 5310 | H | PHE | A | 365 | 17.007 | 37.191 | 28.824 | 1.00 | 0.00 | H |
| ATOM | 5311 | CA | PHE | A | 365 | 17.338 | 39.035 | 29.880 | 1.00 | 0.00 | C |
| ATOM | 5312 | HA | PHE | A | 365 | 17.361 | 39.149 | 30.873 | 1.00 | 0.00 | H |
| ATOM | 5313 | CB | PHE | A | 365 | 18.829 | 39.089 | 29.336 | 1.00 | 0.00 | C |
| ATOM | 5314 | HB1 | PHE | A | 365 | 19.420 | 38.472 | 29.856 | 1.00 | 0.00 | H |
| ATOM | 5315 | HB2 | PHE | A | 365 | 19.192 | 40.019 | 29.387 | 1.00 | 0.00 | H |
| ATOM | 5316 | CG | PHE | A | 365 | 18.872 | 38.668 | 27.926 | 1.00 | 0.00 | C |
| ATOM | 5317 | CD1 | PHE | A | 365 | 18.350 | 39.436 | 26.872 | 1.00 | 0.00 | C |
| ATOM | 5318 | HD1 | PHE | A | 365 | 17.907 | 40.308 | 27.082 | 1.00 | 0.00 | H |
| ATOM | 5319 | CE1 | PHE | A | 365 | 18.428 | 39.027 | 25.559 | 1.00 | 0.00 | C |
| ATOM | 5320 | HE1 | PHE | A | 365 | 17.999 | 39.566 | 24.835 | 1.00 | 0.00 | H |
| ATOM | 5321 | CZ | PHE | A | 365 | 19.119 | 37.842 | 25.250 | 1.00 | 0.00 | C |
| ATOM | 5322 | HZ | PHE | A | 365 | 19.247 | 37.566 | 24.297 | 1.00 | 0.00 | H |
| ATOM | 5323 | CE2 | PHE | A | 365 | 19.622 | 37.058 | 26.280 | 1.00 | 0.00 | C |
| ATOM | 5324 | HE2 | PHE | A | 365 | 20.018 | 36.164 | 26.072 | 1.00 | 0.00 | H |
| ATOM | 5325 | CD2 | PHE | A | 365 | 19.581 | 37.501 | 27.591 | 1.00 | 0.00 | C |
| ATOM | 5326 | HD2 | PHE | A | 365 | 20.059 | 36.988 | 28.304 | 1.00 | 0.00 | H |
| ATOM | 5327 | C | PHE | A | 365 | 16.453 | 40.140 | 29.250 | 1.00 | 0.00 | C |
| ATOM | 5328 | O | PHE | A | 365 | 16.572 | 41.336 | 29.511 | 1.00 | 0.00 | O |
| ATOM | 5329 | N | ALA | A | 366 | 15.487 | 39.754 | 28.372 | 1.00 | 0.00 | N |
| ATOM | 5330 | H | ALA | A | 366 | 15.403 | 38.767 | 28.232 | 1.00 | 0.00 | H |
| ATOM | 5331 | CA | ALA | A | 366 | 14.551 | 40.584 | 27.602 | 1.00 | 0.00 | C |
| ATOM | 5332 | HA | ALA | A | 366 | 14.930 | 41.509 | 27.560 | 1.00 | 0.00 | H |
| ATOM | 5333 | CB | ALA | A | 366 | 14.480 | 40.005 | 26.177 | 1.00 | 0.00 | C |
| ATOM | 5334 | HB1 | ALA | A | 366 | 13.848 | 40.551 | 25.627 | 1.00 | 0.00 | H |
| ATOM | 5335 | HB2 | ALA | A | 366 | 15.390 | 40.028 | 25.762 | 1.00 | 0.00 | H |
| ATOM | 5336 | HB3 | ALA | A | 366 | 14.156 | 39.060 | 26.217 | 1.00 | 0.00 | H |
| ATOM | 5337 | C | ALA | A | 366 | 13.150 | 40.684 | 28.316 | 1.00 | 0.00 | C |
| ATOM | 5338 | O | ALA | A | 366 | 12.392 | 41.661 | 28.007 | 1.00 | 0.00 | O |
| ATOM | 5339 | N | GLY | A | 367 | 12.787 | 39.671 | 29.115 | 1.00 | 0.00 | N |
| ATOM | 5340 | H | GLY | A | 367 | 13.480 | 38.965 | 29.262 | 1.00 | 0.00 | H |
| ATOM | 5341 | CA | GLY | A | 367 | 11.500 | 39.457 | 29.808 | 1.00 | 0.00 | C |
| ATOM | 5342 | HA1 | GLY | A | 367 | 11.271 | 40.307 | 30.283 | 1.00 | 0.00 | H |
| ATOM | 5343 | HA2 | GLY | A | 367 | 11.638 | 38.727 | 30.478 | 1.00 | 0.00 | H |

| | | | | | | | | | | | |
|---|---|---|---|---|---|---|---|---|---|---|---|
| ATOM | 5344 | C | GLY | A | 367 | 10.347 | 39.065 | 28.800 | 1.00 | 0.00 | C |
| ATOM | 5345 | O | GLY | A | 367 | 9.173 | 39.123 | 29.085 | 1.00 | 0.00 | O |
| ATOM | 5346 | N | PHE | A | 368 | 10.748 | 38.539 | 27.618 | 1.00 | 0.00 | N |
| ATOM | 5347 | H | PHE | A | 368 | 11.720 | 38.330 | 27.509 | 1.00 | 0.00 | H |
| ATOM | 5348 | CA | PHE | A | 368 | 9.868 | 38.257 | 26.497 | 1.00 | 0.00 | C |
| ATOM | 5349 | HA | PHE | A | 368 | 8.932 | 38.548 | 26.696 | 1.00 | 0.00 | H |
| ATOM | 5350 | CB | PHE | A | 368 | 10.702 | 38.994 | 25.374 | 1.00 | 0.00 | C |
| ATOM | 5351 | HB1 | PHE | A | 368 | 11.604 | 38.569 | 25.293 | 1.00 | 0.00 | H |
| ATOM | 5352 | HB2 | PHE | A | 368 | 10.809 | 39.959 | 25.615 | 1.00 | 0.00 | H |
| ATOM | 5353 | CG | PHE | A | 368 | 10.081 | 38.952 | 24.021 | 1.00 | 0.00 | C |
| ATOM | 5354 | CD1 | PHE | A | 368 | 10.646 | 38.060 | 23.073 | 1.00 | 0.00 | C |
| ATOM | 5355 | HD1 | PHE | A | 368 | 11.495 | 37.583 | 23.301 | 1.00 | 0.00 | H |
| ATOM | 5356 | CE1 | PHE | A | 368 | 10.046 | 37.828 | 21.821 | 1.00 | 0.00 | C |
| ATOM | 5357 | HE1 | PHE | A | 368 | 10.409 | 37.088 | 21.255 | 1.00 | 0.00 | H |
| ATOM | 5358 | CZ | PHE | A | 368 | 8.968 | 38.574 | 21.321 | 1.00 | 0.00 | C |
| ATOM | 5359 | HZ | PHE | A | 368 | 8.613 | 38.482 | 20.391 | 1.00 | 0.00 | H |
| ATOM | 5360 | CE2 | PHE | A | 368 | 8.443 | 39.463 | 22.270 | 1.00 | 0.00 | C |
| ATOM | 5361 | HE2 | PHE | A | 368 | 7.679 | 40.038 | 21.977 | 1.00 | 0.00 | H |
| ATOM | 5362 | CD2 | PHE | A | 368 | 8.906 | 39.629 | 23.644 | 1.00 | 0.00 | C |
| ATOM | 5363 | HD2 | PHE | A | 368 | 8.407 | 40.203 | 24.294 | 1.00 | 0.00 | H |
| ATOM | 5364 | C | PHE | A | 368 | 9.619 | 36.783 | 26.187 | 1.00 | 0.00 | C |
| ATOM | 5365 | O | PHE | A | 368 | 8.782 | 36.573 | 25.245 | 1.00 | 0.00 | O |
| ATOM | 5366 | N | SER | A | 369 | 10.407 | 35.872 | 26.781 | 1.00 | 0.00 | N |
| ATOM | 5367 | H | SER | A | 369 | 11.062 | 36.180 | 27.471 | 1.00 | 0.00 | H |
| ATOM | 5368 | CA | SER | A | 369 | 10.344 | 34.443 | 26.458 | 1.00 | 0.00 | C |
| ATOM | 5369 | HA | SER | A | 369 | 9.745 | 34.426 | 25.658 | 1.00 | 0.00 | H |
| ATOM | 5370 | CB | SER | A | 369 | 11.770 | 33.910 | 26.221 | 1.00 | 0.00 | C |
| ATOM | 5371 | HB1 | SER | A | 369 | 11.761 | 32.928 | 26.034 | 1.00 | 0.00 | H |
| ATOM | 5372 | HB2 | SER | A | 369 | 12.359 | 34.094 | 27.008 | 1.00 | 0.00 | H |
| ATOM | 5373 | OG | SER | A | 369 | 12.384 | 34.495 | 25.143 | 1.00 | 0.00 | O |
| ATOM | 5374 | HG | SER | A | 369 | 13.302 | 34.114 | 25.035 | 1.00 | 0.00 | H |
| ATOM | 5375 | C | SER | A | 369 | 9.677 | 33.570 | 27.470 | 1.00 | 0.00 | C |
| ATOM | 5376 | O | SER | A | 369 | 9.052 | 32.572 | 27.044 | 1.00 | 0.00 | O |
| ATOM | 5377 | N | CYS | A | 370 | 9.794 | 34.014 | 28.690 | 1.00 | 0.00 | N |
| ATOM | 5378 | H | CYS | A | 370 | 10.365 | 34.826 | 28.810 | 1.00 | 0.00 | H |
| ATOM | 5379 | CA | CYS | A | 370 | 9.173 | 33.451 | 29.938 | 1.00 | 0.00 | C |
| ATOM | 5380 | HA | CYS | A | 370 | 8.741 | 32.583 | 29.692 | 1.00 | 0.00 | H |
| ATOM | 5381 | CB | CYS | A | 370 | 10.304 | 33.211 | 30.940 | 1.00 | 0.00 | C |
| ATOM | 5382 | HB1 | CYS | A | 370 | 9.882 | 32.641 | 31.645 | 1.00 | 0.00 | H |
| ATOM | 5383 | HB2 | CYS | A | 370 | 10.490 | 34.120 | 31.313 | 1.00 | 0.00 | H |
| ATOM | 5384 | SG | CYS | A | 370 | 11.802 | 32.444 | 30.267 | 1.00 | 0.00 | S |
| ATOM | 5385 | C | CYS | A | 370 | 8.006 | 34.304 | 30.517 | 1.00 | 0.00 | C |
| ATOM | 5386 | O | CYS | A | 370 | 7.191 | 33.924 | 31.412 | 1.00 | 0.00 | O |
| ATOM | 5387 | N | ASN | A | 371 | 7.943 | 35.528 | 29.991 | 1.00 | 0.00 | N |
| ATOM | 5388 | H | ASN | A | 371 | 8.692 | 35.833 | 29.403 | 1.00 | 0.00 | H |
| ATOM | 5389 | CA | ASN | A | 371 | 6.828 | 36.446 | 30.235 | 1.00 | 0.00 | C |
| ATOM | 5390 | HA | ASN | A | 371 | 7.181 | 37.369 | 30.083 | 1.00 | 0.00 | H |
| ATOM | 5391 | CB | ASN | A | 371 | 5.715 | 36.069 | 29.209 | 1.00 | 0.00 | C |
| ATOM | 5392 | HB1 | ASN | A | 371 | 5.072 | 36.831 | 29.130 | 1.00 | 0.00 | H |
| ATOM | 5393 | HB2 | ASN | A | 371 | 5.231 | 35.257 | 29.536 | 1.00 | 0.00 | H |
| ATOM | 5394 | CG | ASN | A | 371 | 6.225 | 35.764 | 27.825 | 1.00 | 0.00 | C |
| ATOM | 5395 | OD1 | ASN | A | 371 | 6.339 | 34.642 | 27.336 | 1.00 | 0.00 | O |
| ATOM | 5396 | ND2 | ASN | A | 371 | 6.584 | 36.819 | 27.155 | 1.00 | 0.00 | N |
| ATOM | 5397 | HD21 | ASN | A | 371 | 6.507 | 37.725 | 27.570 | 1.00 | 0.00 | H |

```
ATOM   5398 HD22 ASN A 371       6.937  36.724  26.224  1.00  0.00           H
ATOM   5399  C   ASN A 371       6.372  36.554  31.700  1.00  0.00           C
ATOM   5400  O   ASN A 371       5.198  36.301  31.934  1.00  0.00           O
ATOM   5401  N   GLN A 372       7.392  36.779  32.615  1.00  0.00           N
ATOM   5402  H   GLN A 372       8.292  36.920  32.204  1.00  0.00           H
ATOM   5403  CA  GLN A 372       7.378  36.845  34.074  1.00  0.00           C
ATOM   5404  HA  GLN A 372       6.424  37.075  34.266  1.00  0.00           H
ATOM   5405  CB  GLN A 372       7.769  35.461  34.590  1.00  0.00           C
ATOM   5406  HB1 GLN A 372       7.158  34.807  34.144  1.00  0.00           H
ATOM   5407  HB2 GLN A 372       7.593  35.464  35.574  1.00  0.00           H
ATOM   5408  CG  GLN A 372       9.182  34.922  34.408  1.00  0.00           C
ATOM   5409  HG1 GLN A 372       9.769  35.343  35.100  1.00  0.00           H
ATOM   5410  HG2 GLN A 372       9.501  35.182  33.497  1.00  0.00           H
ATOM   5411  CD  GLN A 372       9.308  33.417  34.538  1.00  0.00           C
ATOM   5412  OE1 GLN A 372      10.327  32.873  34.125  1.00  0.00           O
ATOM   5413  NE2 GLN A 372       8.448  32.838  35.348  1.00  0.00           N
ATOM   5414 HE21 GLN A 372       7.767  33.389  35.831  1.00  0.00           H
ATOM   5415 HE22 GLN A 372       8.475  31.847  35.480  1.00  0.00           H
ATOM   5416  C   GLN A 372       8.227  37.992  34.764  1.00  0.00           C
ATOM   5417  O   GLN A 372       8.492  37.976  35.961  1.00  0.00           O
ATOM   5418  N   GLY A 373       8.641  38.941  33.970  1.00  0.00           N
ATOM   5419  H   GLY A 373       8.212  39.015  33.070  1.00  0.00           H
ATOM   5420  CA  GLY A 373       9.675  39.891  34.302  1.00  0.00           C
ATOM   5421  HA1 GLY A 373       9.746  39.901  35.299  1.00  0.00           H
ATOM   5422  HA2 GLY A 373       9.358  40.784  33.982  1.00  0.00           H
ATOM   5423  C   GLY A 373      11.061  39.548  33.664  1.00  0.00           C
ATOM   5424  O   GLY A 373      11.314  38.369  33.433  1.00  0.00           O
ATOM   5425  N   ARG A 374      11.904  40.456  33.289  1.00  0.00           N
ATOM   5426  H   ARG A 374      11.605  41.410  33.298  1.00  0.00           H
ATOM   5427  CA  ARG A 374      13.359  40.144  32.824  1.00  0.00           C
ATOM   5428  HA  ARG A 374      13.257  39.287  32.320  1.00  0.00           H
ATOM   5429  CB  ARG A 374      13.882  41.287  31.838  1.00  0.00           C
ATOM   5430  HB1 ARG A 374      13.221  41.439  31.103  1.00  0.00           H
ATOM   5431  HB2 ARG A 374      14.764  41.027  31.444  1.00  0.00           H
ATOM   5432  CG  ARG A 374      14.071  42.594  32.563  1.00  0.00           C
ATOM   5433  HG1 ARG A 374      14.604  42.436  33.395  1.00  0.00           H
ATOM   5434  HG2 ARG A 374      13.176  42.969  32.806  1.00  0.00           H
ATOM   5435  CD  ARG A 374      14.828  43.561  31.602  1.00  0.00           C
ATOM   5436  HD1 ARG A 374      15.594  43.072  31.185  1.00  0.00           H
ATOM   5437  HD2 ARG A 374      15.174  44.338  32.128  1.00  0.00           H
ATOM   5438  NE  ARG A 374      13.960  44.107  30.482  1.00  0.00           N
ATOM   5439  HE  ARG A 374      12.986  44.198  30.690  1.00  0.00           H
ATOM   5440  CZ  ARG A 374      14.333  44.482  29.251  1.00  0.00           C
ATOM   5441  NH1 ARG A 374      15.328  43.959  28.587  1.00  0.00           N
ATOM   5442 HH11 ARG A 374      15.871  43.225  28.995  1.00  0.00           H
ATOM   5443 HH12 ARG A 374      15.547  44.292  27.670  1.00  0.00           H
ATOM   5444  NH2 ARG A 374      13.697  45.465  28.645  1.00  0.00           N
ATOM   5445 HH21 ARG A 374      12.939  45.928  29.104  1.00  0.00           H
ATOM   5446 HH22 ARG A 374      13.973  45.748  27.726  1.00  0.00           H
ATOM   5447  C   ARG A 374      14.420  39.775  33.984  1.00  0.00           C
ATOM   5448  O   ARG A 374      14.233  40.135  35.152  1.00  0.00           O
ATOM   5449  N   TYR A 375      15.495  39.189  33.457  1.00  0.00           N
ATOM   5450  H   TYR A 375      15.741  39.352  32.501  1.00  0.00           H
ATOM   5451  CA  TYR A 375      16.317  38.300  34.294  1.00  0.00           C
```

| | | | | | | | | | | | |
|---|---|---|---|---|---|---|---|---|---|---|---|
| ATOM | 5452 | HA | TYR | A | 375 | 16.613 | 37.549 | 33.704 | 1.00 | 0.00 | H |
| ATOM | 5453 | CB | TYR | A | 375 | 17.442 | 39.211 | 34.809 | 1.00 | 0.00 | C |
| ATOM | 5454 | HB1 | TYR | A | 375 | 18.016 | 38.697 | 35.447 | 1.00 | 0.00 | H |
| ATOM | 5455 | HB2 | TYR | A | 375 | 17.040 | 39.995 | 35.281 | 1.00 | 0.00 | H |
| ATOM | 5456 | CG | TYR | A | 375 | 18.318 | 39.729 | 33.710 | 1.00 | 0.00 | C |
| ATOM | 5457 | CD1 | TYR | A | 375 | 19.234 | 38.943 | 32.963 | 1.00 | 0.00 | C |
| ATOM | 5458 | HD1 | TYR | A | 375 | 19.255 | 37.946 | 33.043 | 1.00 | 0.00 | H |
| ATOM | 5459 | CE1 | TYR | A | 375 | 20.073 | 39.600 | 32.142 | 1.00 | 0.00 | C |
| ATOM | 5460 | HE1 | TYR | A | 375 | 20.749 | 39.053 | 31.649 | 1.00 | 0.00 | H |
| ATOM | 5461 | CZ | TYR | A | 375 | 20.084 | 41.004 | 31.913 | 1.00 | 0.00 | C |
| ATOM | 5462 | OH | TYR | A | 375 | 21.074 | 41.540 | 31.220 | 1.00 | 0.00 | O |
| ATOM | 5463 | HH | TYR | A | 375 | 20.935 | 42.528 | 31.149 | 1.00 | 0.00 | H |
| ATOM | 5464 | CE2 | TYR | A | 375 | 19.190 | 41.755 | 32.601 | 1.00 | 0.00 | C |
| ATOM | 5465 | HE2 | TYR | A | 375 | 19.180 | 42.747 | 32.478 | 1.00 | 0.00 | H |
| ATOM | 5466 | CD2 | TYR | A | 375 | 18.276 | 41.155 | 33.487 | 1.00 | 0.00 | C |
| ATOM | 5467 | HD2 | TYR | A | 375 | 17.598 | 41.716 | 33.962 | 1.00 | 0.00 | H |
| ATOM | 5468 | C | TYR | A | 375 | 15.613 | 37.516 | 35.414 | 1.00 | 0.00 | C |
| ATOM | 5469 | O | TYR | A | 375 | 15.905 | 37.680 | 36.577 | 1.00 | 0.00 | O |
| ATOM | 5470 | N | PRO | A | 376 | 14.574 | 36.683 | 35.169 | 1.00 | 0.00 | N |
| ATOM | 5471 | CD | PRO | A | 376 | 14.110 | 36.258 | 33.822 | 1.00 | 0.00 | C |
| ATOM | 5472 | HD1 | PRO | A | 376 | 14.869 | 35.878 | 33.292 | 1.00 | 0.00 | H |
| ATOM | 5473 | HD2 | PRO | A | 376 | 13.712 | 37.031 | 33.327 | 1.00 | 0.00 | H |
| ATOM | 5474 | CG | PRO | A | 376 | 13.059 | 35.193 | 34.143 | 1.00 | 0.00 | C |
| ATOM | 5475 | HG1 | PRO | A | 376 | 13.475 | 34.288 | 34.235 | 1.00 | 0.00 | H |
| ATOM | 5476 | HG2 | PRO | A | 376 | 12.347 | 35.163 | 33.441 | 1.00 | 0.00 | H |
| ATOM | 5477 | CB | PRO | A | 376 | 12.496 | 35.671 | 35.470 | 1.00 | 0.00 | C |
| ATOM | 5478 | HB1 | PRO | A | 376 | 12.069 | 34.912 | 35.961 | 1.00 | 0.00 | H |
| ATOM | 5479 | HB2 | PRO | A | 376 | 11.821 | 36.393 | 35.322 | 1.00 | 0.00 | H |
| ATOM | 5480 | CA | PRO | A | 376 | 13.720 | 36.209 | 36.241 | 1.00 | 0.00 | C |
| ATOM | 5481 | HA | PRO | A | 376 | 13.585 | 36.987 | 36.854 | 1.00 | 0.00 | H |
| ATOM | 5482 | C | PRO | A | 376 | 14.294 | 35.097 | 37.157 | 1.00 | 0.00 | C |
| ATOM | 5483 | O | PRO | A | 376 | 14.057 | 35.084 | 38.370 | 1.00 | 0.00 | O |
| ATOM | 5484 | N | LEU | A | 377 | 15.168 | 34.248 | 36.625 | 1.00 | 0.00 | N |
| ATOM | 5485 | H | LEU | A | 377 | 15.302 | 34.286 | 35.635 | 1.00 | 0.00 | H |
| ATOM | 5486 | CA | LEU | A | 377 | 15.954 | 33.256 | 37.379 | 1.00 | 0.00 | C |
| ATOM | 5487 | HA | LEU | A | 377 | 15.294 | 32.655 | 37.831 | 1.00 | 0.00 | H |
| ATOM | 5488 | CB | LEU | A | 377 | 16.859 | 32.440 | 36.441 | 1.00 | 0.00 | C |
| ATOM | 5489 | HB1 | LEU | A | 377 | 17.596 | 32.051 | 36.994 | 1.00 | 0.00 | H |
| ATOM | 5490 | HB2 | LEU | A | 377 | 17.245 | 33.071 | 35.768 | 1.00 | 0.00 | H |
| ATOM | 5491 | CG | LEU | A | 377 | 16.177 | 31.282 | 35.672 | 1.00 | 0.00 | C |
| ATOM | 5492 | HG | LEU | A | 377 | 15.459 | 31.616 | 35.061 | 1.00 | 0.00 | H |
| ATOM | 5493 | CD1 | LEU | A | 377 | 17.211 | 30.566 | 34.841 | 1.00 | 0.00 | C |
| ATOM | 5494 | HD11 | LEU | A | 377 | 16.777 | 29.816 | 34.342 | 1.00 | 0.00 | H |
| ATOM | 5495 | HD12 | LEU | A | 377 | 17.619 | 31.207 | 34.191 | 1.00 | 0.00 | H |
| ATOM | 5496 | HD13 | LEU | A | 377 | 17.924 | 30.199 | 35.439 | 1.00 | 0.00 | H |
| ATOM | 5497 | CD2 | LEU | A | 377 | 15.552 | 30.332 | 36.662 | 1.00 | 0.00 | C |
| ATOM | 5498 | HD21 | LEU | A | 377 | 15.110 | 29.582 | 36.171 | 1.00 | 0.00 | H |
| ATOM | 5499 | HD22 | LEU | A | 377 | 16.261 | 29.964 | 37.264 | 1.00 | 0.00 | H |
| ATOM | 5500 | HD23 | LEU | A | 377 | 14.871 | 30.820 | 37.208 | 1.00 | 0.00 | H |
| ATOM | 5501 | C | LEU | A | 377 | 16.880 | 34.031 | 38.356 | 1.00 | 0.00 | C |
| ATOM | 5502 | O | LEU | A | 377 | 16.781 | 33.933 | 39.566 | 1.00 | 0.00 | O |
| ATOM | 5503 | N | ILE | A | 378 | 17.709 | 34.863 | 37.799 | 1.00 | 0.00 | N |
| ATOM | 5504 | H | ILE | A | 378 | 17.661 | 34.965 | 36.805 | 1.00 | 0.00 | H |
| ATOM | 5505 | CA | ILE | A | 378 | 18.707 | 35.662 | 38.510 | 1.00 | 0.00 | C |

| | | | | | | | | | | | |
|---|---|---|---|---|---|---|---|---|---|---|---|
| ATOM | 5506 | HA | ILE | A | 378 | 19.328 | 34.988 | 38.911 | 1.00 | 0.00 | H |
| ATOM | 5507 | CB | ILE | A | 378 | 19.605 | 36.519 | 37.593 | 1.00 | 0.00 | C |
| ATOM | 5508 | HB | ILE | A | 378 | 19.098 | 37.237 | 37.116 | 1.00 | 0.00 | H |
| ATOM | 5509 | CG2 | ILE | A | 378 | 20.567 | 37.399 | 38.292 | 1.00 | 0.00 | C |
| ATOM | 5510 | HG21 | ILE | A | 378 | 21.100 | 37.911 | 37.619 | 1.00 | 0.00 | H |
| ATOM | 5511 | HG22 | ILE | A | 378 | 20.069 | 38.038 | 38.878 | 1.00 | 0.00 | H |
| ATOM | 5512 | HG23 | ILE | A | 378 | 21.182 | 36.843 | 38.851 | 1.00 | 0.00 | H |
| ATOM | 5513 | CG1 | ILE | A | 378 | 20.181 | 35.589 | 36.579 | 1.00 | 0.00 | C |
| ATOM | 5514 | HG11 | ILE | A | 378 | 19.433 | 35.281 | 35.991 | 1.00 | 0.00 | H |
| ATOM | 5515 | HG12 | ILE | A | 378 | 20.563 | 34.806 | 37.069 | 1.00 | 0.00 | H |
| ATOM | 5516 | CD | ILE | A | 378 | 21.254 | 36.143 | 35.699 | 1.00 | 0.00 | C |
| ATOM | 5517 | HD1 | ILE | A | 378 | 21.562 | 35.435 | 35.063 | 1.00 | 0.00 | H |
| ATOM | 5518 | HD2 | ILE | A | 378 | 20.894 | 36.920 | 35.183 | 1.00 | 0.00 | H |
| ATOM | 5519 | HD3 | ILE | A | 378 | 22.024 | 36.444 | 36.262 | 1.00 | 0.00 | H |
| ATOM | 5520 | C | ILE | A | 378 | 18.031 | 36.447 | 39.637 | 1.00 | 0.00 | C |
| ATOM | 5521 | O | ILE | A | 378 | 18.374 | 36.506 | 40.761 | 1.00 | 0.00 | O |
| ATOM | 5522 | N | GLN | A | 379 | 16.969 | 37.133 | 39.288 | 1.00 | 0.00 | N |
| ATOM | 5523 | H | GLN | A | 379 | 16.693 | 37.127 | 38.327 | 1.00 | 0.00 | H |
| ATOM | 5524 | CA | GLN | A | 379 | 16.146 | 37.929 | 40.267 | 1.00 | 0.00 | C |
| ATOM | 5525 | HA | GLN | A | 379 | 16.765 | 38.562 | 40.731 | 1.00 | 0.00 | H |
| ATOM | 5526 | CB | GLN | A | 379 | 15.068 | 38.708 | 39.540 | 1.00 | 0.00 | C |
| ATOM | 5527 | HB1 | GLN | A | 379 | 14.337 | 38.882 | 40.200 | 1.00 | 0.00 | H |
| ATOM | 5528 | HB2 | GLN | A | 379 | 14.724 | 38.119 | 38.809 | 1.00 | 0.00 | H |
| ATOM | 5529 | CG | GLN | A | 379 | 15.392 | 40.029 | 38.906 | 1.00 | 0.00 | C |
| ATOM | 5530 | HG1 | GLN | A | 379 | 14.530 | 40.358 | 38.521 | 1.00 | 0.00 | H |
| ATOM | 5531 | HG2 | GLN | A | 379 | 16.030 | 39.819 | 38.165 | 1.00 | 0.00 | H |
| ATOM | 5532 | CD | GLN | A | 379 | 16.012 | 41.193 | 39.758 | 1.00 | 0.00 | C |
| ATOM | 5533 | OE1 | GLN | A | 379 | 16.341 | 40.937 | 40.914 | 1.00 | 0.00 | O |
| ATOM | 5534 | NE2 | GLN | A | 379 | 16.209 | 42.370 | 39.210 | 1.00 | 0.00 | N |
| ATOM | 5535 | HE21 | GLN | A | 379 | 15.964 | 42.523 | 38.253 | 1.00 | 0.00 | H |
| ATOM | 5536 | HE22 | GLN | A | 379 | 16.604 | 43.113 | 39.750 | 1.00 | 0.00 | H |
| ATOM | 5537 | C | GLN | A | 379 | 15.475 | 36.999 | 41.279 | 1.00 | 0.00 | C |
| ATOM | 5538 | O | GLN | A | 379 | 15.537 | 37.381 | 42.429 | 1.00 | 0.00 | O |
| ATOM | 5539 | N | THR | A | 380 | 14.916 | 35.824 | 40.928 | 1.00 | 0.00 | N |
| ATOM | 5540 | H | THR | A | 380 | 14.732 | 35.608 | 39.969 | 1.00 | 0.00 | H |
| ATOM | 5541 | CA | THR | A | 380 | 14.562 | 34.806 | 42.063 | 1.00 | 0.00 | C |
| ATOM | 5542 | HA | THR | A | 380 | 13.865 | 35.263 | 42.615 | 1.00 | 0.00 | H |
| ATOM | 5543 | CB | THR | A | 380 | 13.989 | 33.505 | 41.497 | 1.00 | 0.00 | C |
| ATOM | 5544 | HB | THR | A | 380 | 14.583 | 33.087 | 40.810 | 1.00 | 0.00 | H |
| ATOM | 5545 | CG2 | THR | A | 380 | 13.692 | 32.522 | 42.665 | 1.00 | 0.00 | C |
| ATOM | 5546 | HG21 | THR | A | 380 | 13.317 | 31.671 | 42.297 | 1.00 | 0.00 | H |
| ATOM | 5547 | HG22 | THR | A | 380 | 14.539 | 32.327 | 43.160 | 1.00 | 0.00 | H |
| ATOM | 5548 | HG23 | THR | A | 380 | 13.029 | 32.936 | 43.289 | 1.00 | 0.00 | H |
| ATOM | 5549 | OG1 | THR | A | 380 | 12.933 | 33.736 | 40.664 | 1.00 | 0.00 | O |
| ATOM | 5550 | HG1 | THR | A | 380 | 12.587 | 32.865 | 40.315 | 1.00 | 0.00 | H |
| ATOM | 5551 | C | THR | A | 380 | 15.724 | 34.471 | 43.017 | 1.00 | 0.00 | C |
| ATOM | 5552 | O | THR | A | 380 | 15.597 | 34.676 | 44.227 | 1.00 | 0.00 | O |
| ATOM | 5553 | N | LEU | A | 381 | 16.903 | 34.090 | 42.530 | 1.00 | 0.00 | N |
| ATOM | 5554 | H | LEU | A | 381 | 16.980 | 33.936 | 41.545 | 1.00 | 0.00 | H |
| ATOM | 5555 | CA | LEU | A | 381 | 18.091 | 33.884 | 43.355 | 1.00 | 0.00 | C |
| ATOM | 5556 | HA | LEU | A | 381 | 17.888 | 33.216 | 44.071 | 1.00 | 0.00 | H |
| ATOM | 5557 | CB | LEU | A | 381 | 19.149 | 33.355 | 42.340 | 1.00 | 0.00 | C |
| ATOM | 5558 | HB1 | LEU | A | 381 | 18.975 | 33.786 | 41.455 | 1.00 | 0.00 | H |
| ATOM | 5559 | HB2 | LEU | A | 381 | 19.035 | 32.365 | 42.257 | 1.00 | 0.00 | H |

| | | | | | | | | | | |
|---|---|---|---|---|---|---|---|---|---|---|
| ATOM | 5560 | CG | LEU | A | 381 | 20.619 | 33.626 | 42.711 | 1.00 | 0.00 | C |
| ATOM | 5561 | HG | LEU | A | 381 | 20.751 | 34.605 | 42.867 | 1.00 | 0.00 | H |
| ATOM | 5562 | CD1 | LEU | A | 381 | 21.100 | 33.063 | 44.004 | 1.00 | 0.00 | C |
| ATOM | 5563 | HD11 | LEU | A | 381 | 22.062 | 33.303 | 44.135 | 1.00 | 0.00 | H |
| ATOM | 5564 | HD12 | LEU | A | 381 | 20.557 | 33.441 | 44.754 | 1.00 | 0.00 | H |
| ATOM | 5565 | HD13 | LEU | A | 381 | 21.003 | 32.068 | 43.991 | 1.00 | 0.00 | H |
| ATOM | 5566 | CD2 | LEU | A | 381 | 21.412 | 33.002 | 41.557 | 1.00 | 0.00 | C |
| ATOM | 5567 | HD21 | LEU | A | 381 | 22.391 | 33.132 | 41.716 | 1.00 | 0.00 | H |
| ATOM | 5568 | HD22 | LEU | A | 381 | 21.210 | 32.024 | 41.504 | 1.00 | 0.00 | H |
| ATOM | 5569 | HD23 | LEU | A | 381 | 21.152 | 33.443 | 40.698 | 1.00 | 0.00 | H |
| ATOM | 5570 | C | LEU | A | 381 | 18.513 | 35.166 | 44.168 | 1.00 | 0.00 | C |
| ATOM | 5571 | O | LEU | A | 381 | 18.587 | 35.029 | 45.374 | 1.00 | 0.00 | O |
| ATOM | 5572 | N | ARG | A | 382 | 18.479 | 36.336 | 43.516 | 1.00 | 0.00 | N |
| ATOM | 5573 | H | ARG | A | 382 | 18.330 | 36.322 | 42.527 | 1.00 | 0.00 | H |
| ATOM | 5574 | CA | ARG | A | 382 | 18.648 | 37.630 | 44.169 | 1.00 | 0.00 | C |
| ATOM | 5575 | HA | ARG | A | 382 | 19.512 | 37.617 | 44.672 | 1.00 | 0.00 | H |
| ATOM | 5576 | CB | ARG | A | 382 | 18.685 | 38.667 | 43.073 | 1.00 | 0.00 | C |
| ATOM | 5577 | HB1 | ARG | A | 382 | 17.760 | 38.722 | 42.698 | 1.00 | 0.00 | H |
| ATOM | 5578 | HB2 | ARG | A | 382 | 19.309 | 38.322 | 42.372 | 1.00 | 0.00 | H |
| ATOM | 5579 | CG | ARG | A | 382 | 19.119 | 40.085 | 43.393 | 1.00 | 0.00 | C |
| ATOM | 5580 | HG1 | ARG | A | 382 | 19.222 | 40.570 | 42.524 | 1.00 | 0.00 | H |
| ATOM | 5581 | HG2 | ARG | A | 382 | 20.006 | 40.030 | 43.851 | 1.00 | 0.00 | H |
| ATOM | 5582 | CD | ARG | A | 382 | 18.169 | 40.902 | 44.287 | 1.00 | 0.00 | C |
| ATOM | 5583 | HD1 | ARG | A | 382 | 18.589 | 41.789 | 44.481 | 1.00 | 0.00 | H |
| ATOM | 5584 | HD2 | ARG | A | 382 | 18.024 | 40.407 | 45.144 | 1.00 | 0.00 | H |
| ATOM | 5585 | NE | ARG | A | 382 | 16.837 | 41.175 | 43.722 | 1.00 | 0.00 | N |
| ATOM | 5586 | HE | ARG | A | 382 | 16.723 | 40.883 | 42.773 | 1.00 | 0.00 | H |
| ATOM | 5587 | CZ | ARG | A | 382 | 15.764 | 41.741 | 44.254 | 1.00 | 0.00 | C |
| ATOM | 5588 | NH1 | ARG | A | 382 | 15.823 | 42.168 | 45.486 | 1.00 | 0.00 | N |
| ATOM | 5589 | HH11 | ARG | A | 382 | 16.668 | 42.065 | 46.011 | 1.00 | 0.00 | H |
| ATOM | 5590 | HH12 | ARG | A | 382 | 15.023 | 42.598 | 45.905 | 1.00 | 0.00 | H |
| ATOM | 5591 | NH2 | ARG | A | 382 | 14.630 | 41.883 | 43.678 | 1.00 | 0.00 | N |
| ATOM | 5592 | HH21 | ARG | A | 382 | 14.499 | 41.552 | 42.744 | 1.00 | 0.00 | H |
| ATOM | 5593 | HH22 | ARG | A | 382 | 13.875 | 42.325 | 44.162 | 1.00 | 0.00 | H |
| ATOM | 5594 | C | ARG | A | 382 | 17.677 | 37.903 | 45.276 | 1.00 | 0.00 | C |
| ATOM | 5595 | O | ARG | A | 382 | 18.033 | 38.277 | 46.412 | 1.00 | 0.00 | O |
| ATOM | 5596 | N | GLN | A | 383 | 16.359 | 37.742 | 45.099 | 1.00 | 0.00 | N |
| ATOM | 5597 | H | GLN | A | 383 | 16.095 | 37.501 | 44.165 | 1.00 | 0.00 | H |
| ATOM | 5598 | CA | GLN | A | 383 | 15.299 | 37.851 | 45.981 | 1.00 | 0.00 | C |
| ATOM | 5599 | HA | GLN | A | 383 | 15.438 | 38.749 | 46.398 | 1.00 | 0.00 | H |
| ATOM | 5600 | CB | GLN | A | 383 | 13.983 | 37.906 | 45.199 | 1.00 | 0.00 | C |
| ATOM | 5601 | HB1 | GLN | A | 383 | 13.902 | 37.083 | 44.636 | 1.00 | 0.00 | H |
| ATOM | 5602 | HB2 | GLN | A | 383 | 13.986 | 38.714 | 44.610 | 1.00 | 0.00 | H |
| ATOM | 5603 | CG | GLN | A | 383 | 12.784 | 37.983 | 46.127 | 1.00 | 0.00 | C |
| ATOM | 5604 | HG1 | GLN | A | 383 | 12.814 | 37.186 | 46.730 | 1.00 | 0.00 | H |
| ATOM | 5605 | HG2 | GLN | A | 383 | 11.959 | 37.944 | 45.563 | 1.00 | 0.00 | H |
| ATOM | 5606 | CD | GLN | A | 383 | 12.665 | 39.196 | 47.002 | 1.00 | 0.00 | C |
| ATOM | 5607 | OE1 | GLN | A | 383 | 13.069 | 40.320 | 46.701 | 1.00 | 0.00 | O |
| ATOM | 5608 | NE2 | GLN | A | 383 | 12.097 | 39.024 | 48.207 | 1.00 | 0.00 | N |
| ATOM | 5609 | HE21 | GLN | A | 383 | 11.776 | 38.118 | 48.484 | 1.00 | 0.00 | H |
| ATOM | 5610 | HE22 | GLN | A | 383 | 11.995 | 39.802 | 48.827 | 1.00 | 0.00 | H |
| ATOM | 5611 | C | GLN | A | 383 | 15.393 | 36.857 | 47.159 | 1.00 | 0.00 | C |
| ATOM | 5612 | O | GLN | A | 383 | 15.277 | 37.257 | 48.315 | 1.00 | 0.00 | O |
| ATOM | 5613 | N | GLU | A | 384 | 15.845 | 35.631 | 46.957 | 1.00 | 0.00 | N |

| | | | | | | | | | | | |
|---|---|---|---|---|---|---|---|---|---|---|---|
| ATOM | 5614 | H | GLU | A | 384 | 16.221 | 35.383 | 46.064 | 1.00 | 0.00 | H |
| ATOM | 5615 | CA | GLU | A | 384 | 15.801 | 34.632 | 48.028 | 1.00 | 0.00 | C |
| ATOM | 5616 | HA | GLU | A | 384 | 15.018 | 34.817 | 48.622 | 1.00 | 0.00 | H |
| ATOM | 5617 | CB | GLU | A | 384 | 15.576 | 33.306 | 47.399 | 1.00 | 0.00 | C |
| ATOM | 5618 | HB1 | GLU | A | 384 | 15.600 | 32.631 | 48.137 | 1.00 | 0.00 | H |
| ATOM | 5619 | HB2 | GLU | A | 384 | 16.344 | 33.149 | 46.779 | 1.00 | 0.00 | H |
| ATOM | 5620 | CG | GLU | A | 384 | 14.277 | 33.106 | 46.611 | 1.00 | 0.00 | C |
| ATOM | 5621 | HG1 | GLU | A | 384 | 14.339 | 32.267 | 46.070 | 1.00 | 0.00 | H |
| ATOM | 5622 | HG2 | GLU | A | 384 | 14.131 | 33.886 | 46.002 | 1.00 | 0.00 | H |
| ATOM | 5623 | CD | GLU | A | 384 | 13.112 | 33.000 | 47.651 | 1.00 | 0.00 | C |
| ATOM | 5624 | OE1 | GLU | A | 384 | 12.331 | 33.988 | 47.752 | 1.00 | 0.00 | O |
| ATOM | 5625 | OE2 | GLU | A | 384 | 13.038 | 32.146 | 48.543 | 1.00 | 0.00 | O |
| ATOM | 5626 | C | GLU | A | 384 | 17.149 | 34.683 | 48.843 | 1.00 | 0.00 | C |
| ATOM | 5627 | O | GLU | A | 384 | 17.339 | 33.959 | 49.786 | 1.00 | 0.00 | O |
| ATOM | 5628 | N | LEU | A | 385 | 18.032 | 35.574 | 48.427 | 1.00 | 0.00 | N |
| ATOM | 5629 | H | LEU | A | 385 | 17.926 | 35.934 | 47.500 | 1.00 | 0.00 | H |
| ATOM | 5630 | CA | LEU | A | 385 | 19.166 | 36.077 | 49.232 | 1.00 | 0.00 | C |
| ATOM | 5631 | HA | LEU | A | 385 | 19.288 | 35.435 | 49.989 | 1.00 | 0.00 | H |
| ATOM | 5632 | CB | LEU | A | 385 | 20.418 | 36.151 | 48.342 | 1.00 | 0.00 | C |
| ATOM | 5633 | HB1 | LEU | A | 385 | 21.125 | 36.654 | 48.839 | 1.00 | 0.00 | H |
| ATOM | 5634 | HB2 | LEU | A | 385 | 20.179 | 36.648 | 47.508 | 1.00 | 0.00 | H |
| ATOM | 5635 | CG | LEU | A | 385 | 21.015 | 34.826 | 47.924 | 1.00 | 0.00 | C |
| ATOM | 5636 | HG | LEU | A | 385 | 20.292 | 34.290 | 47.487 | 1.00 | 0.00 | H |
| ATOM | 5637 | CD1 | LEU | A | 385 | 22.144 | 35.068 | 46.958 | 1.00 | 0.00 | C |
| ATOM | 5638 | HD11 | LEU | A | 385 | 22.539 | 34.192 | 46.681 | 1.00 | 0.00 | H |
| ATOM | 5639 | HD12 | LEU | A | 385 | 21.796 | 35.546 | 46.152 | 1.00 | 0.00 | H |
| ATOM | 5640 | HD13 | LEU | A | 385 | 22.847 | 35.626 | 47.399 | 1.00 | 0.00 | H |
| ATOM | 5641 | CD2 | LEU | A | 385 | 21.558 | 34.056 | 49.122 | 1.00 | 0.00 | C |
| ATOM | 5642 | HD21 | LEU | A | 385 | 21.945 | 33.187 | 48.812 | 1.00 | 0.00 | H |
| ATOM | 5643 | HD22 | LEU | A | 385 | 22.270 | 34.596 | 49.571 | 1.00 | 0.00 | H |
| ATOM | 5644 | HD23 | LEU | A | 385 | 20.816 | 33.879 | 49.769 | 1.00 | 0.00 | H |
| ATOM | 5645 | C | LEU | A | 385 | 18.875 | 37.416 | 49.854 | 1.00 | 0.00 | C |
| ATOM | 5646 | O | LEU | A | 385 | 19.724 | 37.952 | 50.678 | 1.00 | 0.00 | O |
| ATOM | 5647 | N | SER | A | 386 | 17.690 | 37.936 | 49.634 | 1.00 | 0.00 | N |
| ATOM | 5648 | H | SER | A | 386 | 17.018 | 37.402 | 49.121 | 1.00 | 0.00 | H |
| ATOM | 5649 | CA | SER | A | 386 | 17.314 | 39.274 | 50.113 | 1.00 | 0.00 | C |
| ATOM | 5650 | HA | SER | A | 386 | 16.436 | 39.444 | 49.665 | 1.00 | 0.00 | H |
| ATOM | 5651 | CB | SER | A | 386 | 16.974 | 39.313 | 51.561 | 1.00 | 0.00 | C |
| ATOM | 5652 | HB1 | SER | A | 386 | 16.794 | 40.263 | 51.815 | 1.00 | 0.00 | H |
| ATOM | 5653 | HB2 | SER | A | 386 | 17.761 | 38.974 | 52.077 | 1.00 | 0.00 | H |
| ATOM | 5654 | OG | SER | A | 386 | 15.840 | 38.552 | 52.001 | 1.00 | 0.00 | O |
| ATOM | 5655 | HG | SER | A | 386 | 15.725 | 38.666 | 52.988 | 1.00 | 0.00 | H |
| ATOM | 5656 | C | SER | A | 386 | 18.359 | 40.364 | 49.695 | 1.00 | 0.00 | C |
| ATOM | 5657 | O | SER | A | 386 | 18.663 | 41.205 | 50.497 | 1.00 | 0.00 | O |
| ATOM | 5658 | N | LEU | A | 387 | 18.974 | 40.230 | 48.500 | 1.00 | 0.00 | N |
| ATOM | 5659 | H | LEU | A | 387 | 18.768 | 39.443 | 47.919 | 1.00 | 0.00 | H |
| ATOM | 5660 | CA | LEU | A | 387 | 19.926 | 41.217 | 48.060 | 1.00 | 0.00 | C |
| ATOM | 5661 | HA | LEU | A | 387 | 20.368 | 41.379 | 48.942 | 1.00 | 0.00 | H |
| ATOM | 5662 | CB | LEU | A | 387 | 20.922 | 40.688 | 46.927 | 1.00 | 0.00 | C |
| ATOM | 5663 | HB1 | LEU | A | 387 | 21.540 | 41.435 | 46.682 | 1.00 | 0.00 | H |
| ATOM | 5664 | HB2 | LEU | A | 387 | 20.377 | 40.431 | 46.129 | 1.00 | 0.00 | H |
| ATOM | 5665 | CG | LEU | A | 387 | 21.813 | 39.457 | 47.287 | 1.00 | 0.00 | C |
| ATOM | 5666 | HG | LEU | A | 387 | 21.194 | 38.735 | 47.594 | 1.00 | 0.00 | H |
| ATOM | 5667 | CD1 | LEU | A | 387 | 22.610 | 38.989 | 46.069 | 1.00 | 0.00 | C |

```
ATOM   5668 HD11 LEU A 387      23.173  38.202  46.321  1.00  0.00           H
ATOM   5669 HD12 LEU A 387      21.979  38.728  45.338  1.00  0.00           H
ATOM   5670 HD13 LEU A 387      23.199  39.732  45.751  1.00  0.00           H
ATOM   5671  CD2 LEU A 387      22.796  39.731  48.424  1.00  0.00           C
ATOM   5672 HD21 LEU A 387      23.334  38.908  48.605  1.00  0.00           H
ATOM   5673 HD22 LEU A 387      23.407  40.478  48.163  1.00  0.00           H
ATOM   5674 HD23 LEU A 387      22.290  39.986  49.248  1.00  0.00           H
ATOM   5675  C   LEU A 387      19.288  42.516  47.568  1.00  0.00           C
ATOM   5676  O   LEU A 387      18.312  42.438  46.859  1.00  0.00           O
ATOM   5677  N   PRO A 388      19.971  43.649  47.880  1.00  0.00           N
ATOM   5678  CD  PRO A 388      21.107  43.801  48.807  1.00  0.00           C
ATOM   5679  HD1 PRO A 388      21.974  43.716  48.315  1.00  0.00           H
ATOM   5680  HD2 PRO A 388      21.064  43.111  49.529  1.00  0.00           H
ATOM   5681  CG  PRO A 388      20.899  45.217  49.358  1.00  0.00           C
ATOM   5682  HG1 PRO A 388      21.782  45.679  49.437  1.00  0.00           H
ATOM   5683  HG2 PRO A 388      20.468  45.166  50.259  1.00  0.00           H
ATOM   5684  CB  PRO A 388      20.016  46.020  48.447  1.00  0.00           C
ATOM   5685  HB1 PRO A 388      20.572  46.650  47.905  1.00  0.00           H
ATOM   5686  HB2 PRO A 388      19.354  46.539  48.988  1.00  0.00           H
ATOM   5687  CA  PRO A 388      19.302  44.988  47.538  1.00  0.00           C
ATOM   5688  HA  PRO A 388      18.313  44.914  47.663  1.00  0.00           H
ATOM   5689  C   PRO A 388      19.433  45.408  46.095  1.00  0.00           C
ATOM   5690  O   PRO A 388      20.532  45.405  45.508  1.00  0.00           O
ATOM   5691  N   TYR A 389      18.231  45.566  45.448  1.00  0.00           N
ATOM   5692  H   TYR A 389      17.380  45.417  45.952  1.00  0.00           H
ATOM   5693  CA  TYR A 389      18.167  45.961  43.986  1.00  0.00           C
ATOM   5694  HA  TYR A 389      18.929  45.465  43.570  1.00  0.00           H
ATOM   5695  CB  TYR A 389      16.835  45.480  43.410  1.00  0.00           C
ATOM   5696  HB1 TYR A 389      16.092  45.994  43.839  1.00  0.00           H
ATOM   5697  HB2 TYR A 389      16.727  44.507  43.612  1.00  0.00           H
ATOM   5698  CG  TYR A 389      16.686  45.638  41.956  1.00  0.00           C
ATOM   5699  CD1 TYR A 389      15.389  45.954  41.360  1.00  0.00           C
ATOM   5700  HD1 TYR A 389      14.568  46.066  41.920  1.00  0.00           H
ATOM   5701  CE1 TYR A 389      15.351  46.096  39.937  1.00  0.00           C
ATOM   5702  HE1 TYR A 389      14.511  46.427  39.508  1.00  0.00           H
ATOM   5703  CZ  TYR A 389      16.428  45.794  39.139  1.00  0.00           C
ATOM   5704  OH  TYR A 389      16.312  45.911  37.767  1.00  0.00           O
ATOM   5705  HH  TYR A 389      17.178  45.656  37.336  1.00  0.00           H
ATOM   5706  CE2 TYR A 389      17.607  45.408  39.701  1.00  0.00           C
ATOM   5707  HE2 TYR A 389      18.389  45.208  39.111  1.00  0.00           H
ATOM   5708  CD2 TYR A 389      17.746  45.285  41.105  1.00  0.00           C
ATOM   5709  HD2 TYR A 389      18.603  44.945  41.492  1.00  0.00           H
ATOM   5710  C   TYR A 389      18.412  47.502  43.658  1.00  0.00           C
ATOM   5711  O   TYR A 389      18.679  47.826  42.484  1.00  0.00           O
ATOM   5712  N   LEU A 390      18.249  48.336  44.662  1.00  0.00           N
ATOM   5713  H   LEU A 390      17.820  47.952  45.480  1.00  0.00           H
ATOM   5714  CA  LEU A 390      18.608  49.734  44.740  1.00  0.00           C
ATOM   5715  HA  LEU A 390      18.257  50.238  43.951  1.00  0.00           H
ATOM   5716  CB  LEU A 390      17.896  50.328  45.982  1.00  0.00           C
ATOM   5717  HB1 LEU A 390      18.282  49.876  46.786  1.00  0.00           H
ATOM   5718  HB2 LEU A 390      16.926  50.097  45.903  1.00  0.00           H
ATOM   5719  CG  LEU A 390      17.992  51.866  46.220  1.00  0.00           C
ATOM   5720  HG  LEU A 390      17.452  52.038  47.044  1.00  0.00           H
ATOM   5721  CD1 LEU A 390      19.400  52.288  46.573  1.00  0.00           C
```

```
ATOM   5722 HD11 LEU A 390      19.425  53.277  46.718  1.00  0.00           H
ATOM   5723 HD12 LEU A 390      19.688  51.821  47.409  1.00  0.00           H
ATOM   5724 HD13 LEU A 390      20.018  52.045  45.825  1.00  0.00           H
ATOM   5725  CD2 LEU A 390      17.226  52.808  45.271  1.00  0.00           C
ATOM   5726 HD21 LEU A 390      17.382  53.757  45.546  1.00  0.00           H
ATOM   5727 HD22 LEU A 390      17.551  52.675  44.335  1.00  0.00           H
ATOM   5728 HD23 LEU A 390      16.248  52.604  45.317  1.00  0.00           H
ATOM   5729  C   LEU A 390      20.194  49.798  44.734  1.00  0.00           C
ATOM   5730  O   LEU A 390      20.807  49.192  45.628  1.00  0.00           O
ATOM   5731  N   PRO A 391      20.844  50.529  43.794  1.00  0.00           N
ATOM   5732  CD  PRO A 391      20.227  51.183  42.650  1.00  0.00           C
ATOM   5733  HD1 PRO A 391      19.480  51.759  42.981  1.00  0.00           H
ATOM   5734  HD2 PRO A 391      19.861  50.477  42.043  1.00  0.00           H
ATOM   5735  CG  PRO A 391      21.274  52.031  41.917  1.00  0.00           C
ATOM   5736  HG1 PRO A 391      21.260  52.985  42.218  1.00  0.00           H
ATOM   5737  HG2 PRO A 391      21.158  51.993  40.924  1.00  0.00           H
ATOM   5738  CB  PRO A 391      22.543  51.307  42.372  1.00  0.00           C
ATOM   5739  HB1 PRO A 391      23.307  51.952  42.375  1.00  0.00           H
ATOM   5740  HB2 PRO A 391      22.741  50.559  41.738  1.00  0.00           H
ATOM   5741  CA  PRO A 391      22.305  50.744  43.815  1.00  0.00           C
ATOM   5742  HA  PRO A 391      22.806  49.904  44.021  1.00  0.00           H
ATOM   5743  C   PRO A 391      22.776  51.657  44.970  1.00  0.00           C
ATOM   5744  O   PRO A 391      22.632  52.901  44.826  1.00  0.00           O
ATOM   5745  N   SER A 392      23.265  51.038  46.068  1.00  0.00           N
ATOM   5746  H   SER A 392      23.345  50.041  46.078  1.00  0.00           H
ATOM   5747  CA  SER A 392      23.681  51.811  47.247  1.00  0.00           C
ATOM   5748  HA  SER A 392      23.506  52.757  46.976  1.00  0.00           H
ATOM   5749  CB  SER A 392      22.841  51.348  48.400  1.00  0.00           C
ATOM   5750  HB1 SER A 392      22.982  50.371  48.563  1.00  0.00           H
ATOM   5751  HB2 SER A 392      21.873  51.518  48.215  1.00  0.00           H
ATOM   5752  OG  SER A 392      23.182  52.037  49.585  1.00  0.00           O
ATOM   5753  HG  SER A 392      22.609  51.711  50.337  1.00  0.00           H
ATOM   5754  C   SER A 392      25.171  51.778  47.637  1.00  0.00           C
ATOM   5755  O   SER A 392      25.763  50.701  47.692  1.00  0.00           O
ATOM   5756  N   GLY A 393      25.694  52.937  47.979  1.00  0.00           N
ATOM   5757  H   GLY A 393      25.171  53.770  47.800  1.00  0.00           H
ATOM   5758  CA  GLY A 393      27.018  53.058  48.614  1.00  0.00           C
ATOM   5759  HA1 GLY A 393      27.119  52.270  49.221  1.00  0.00           H
ATOM   5760  HA2 GLY A 393      27.001  53.897  49.157  1.00  0.00           H
ATOM   5761  C   GLY A 393      28.226  53.108  47.660  1.00  0.00           C
ATOM   5762  O   GLY A 393      28.186  52.781  46.447  1.00  0.00           O
ATOM   5763  N   THR A 394      29.397  53.463  48.206  1.00  0.00           N
ATOM   5764  H   THR A 394      29.397  53.854  49.127  1.00  0.00           H
ATOM   5765  CA  THR A 394      30.702  53.300  47.498  1.00  0.00           C
ATOM   5766  HA  THR A 394      30.546  52.747  46.680  1.00  0.00           H
ATOM   5767  CB  THR A 394      31.248  54.642  47.064  1.00  0.00           C
ATOM   5768  HB  THR A 394      31.421  55.137  47.916  1.00  0.00           H
ATOM   5769  CG2 THR A 394      32.467  54.511  46.126  1.00  0.00           C
ATOM   5770 HG21 THR A 394      32.789  55.422  45.869  1.00  0.00           H
ATOM   5771 HG22 THR A 394      33.199  54.019  46.598  1.00  0.00           H
ATOM   5772 HG23 THR A 394      32.202  54.007  45.304  1.00  0.00           H
ATOM   5773  OG1 THR A 394      30.330  55.413  46.360  1.00  0.00           O
ATOM   5774  HG1 THR A 394      30.752  56.283  46.104  1.00  0.00           H
ATOM   5775  C   THR A 394      31.681  52.576  48.380  1.00  0.00           C
```

| | | | | | | | | | | |
|---|---|---|---|---|---|---|---|---|---|---|
| ATOM | 5776 | O | THR | A | 394 | 31.037 | 50.972 | 47.639 | 1.00 | 0.00 | O |
| ATOM | 5777 | N | PRO | A | 395 | 32.711 | 52.257 | 48.296 | 1.00 | 0.00 | N |
| ATOM | 5778 | CD | PRO | A | 395 | 33.293 | 53.475 | 48.809 | 1.00 | 0.00 | C |
| ATOM | 5779 | HD1 | PRO | A | 395 | 32.651 | 53.904 | 49.445 | 1.00 | 0.00 | H |
| ATOM | 5780 | HD2 | PRO | A | 395 | 33.476 | 54.098 | 48.048 | 1.00 | 0.00 | H |
| ATOM | 5781 | CG | PRO | A | 395 | 34.595 | 53.044 | 49.518 | 1.00 | 0.00 | C |
| ATOM | 5782 | HG1 | PRO | A | 395 | 34.427 | 52.835 | 50.481 | 1.00 | 0.00 | H |
| ATOM | 5783 | HG2 | PRO | A | 395 | 35.294 | 53.756 | 49.448 | 1.00 | 0.00 | H |
| ATOM | 5784 | CB | PRO | A | 395 | 35.023 | 51.785 | 48.756 | 1.00 | 0.00 | C |
| ATOM | 5785 | HB1 | PRO | A | 395 | 35.605 | 51.195 | 49.316 | 1.00 | 0.00 | H |
| ATOM | 5786 | HB2 | PRO | A | 395 | 35.498 | 52.012 | 47.906 | 1.00 | 0.00 | H |
| ATOM | 5787 | CA | PRO | A | 395 | 33.627 | 51.154 | 48.496 | 1.00 | 0.00 | C |
| ATOM | 5788 | HA | PRO | A | 395 | 33.644 | 50.599 | 47.665 | 1.00 | 0.00 | H |
| ATOM | 5789 | C | PRO | A | 395 | 33.123 | 50.190 | 49.589 | 1.00 | 0.00 | C |
| ATOM | 5790 | O | PRO | A | 395 | 32.162 | 49.485 | 49.478 | 1.00 | 0.00 | O |
| ATOM | 5791 | N | GLU | A | 396 | 33.814 | 50.076 | 50.747 | 1.00 | 0.00 | N |
| ATOM | 5792 | H | GLU | A | 396 | 34.720 | 50.500 | 50.749 | 1.00 | 0.00 | H |
| ATOM | 5793 | CA | GLU | A | 396 | 33.431 | 49.432 | 51.975 | 1.00 | 0.00 | C |
| ATOM | 5794 | HA | GLU | A | 396 | 33.246 | 48.529 | 51.587 | 1.00 | 0.00 | H |
| ATOM | 5795 | CB | GLU | A | 396 | 34.565 | 49.416 | 53.044 | 1.00 | 0.00 | C |
| ATOM | 5796 | HB1 | GLU | A | 396 | 35.378 | 49.023 | 52.614 | 1.00 | 0.00 | H |
| ATOM | 5797 | HB2 | GLU | A | 396 | 34.264 | 48.825 | 53.792 | 1.00 | 0.00 | H |
| ATOM | 5798 | CG | GLU | A | 396 | 35.004 | 50.736 | 53.681 | 1.00 | 0.00 | C |
| ATOM | 5799 | HG1 | GLU | A | 396 | 35.284 | 50.542 | 54.621 | 1.00 | 0.00 | H |
| ATOM | 5800 | HG2 | GLU | A | 396 | 34.213 | 51.348 | 53.689 | 1.00 | 0.00 | H |
| ATOM | 5801 | CD | GLU | A | 396 | 36.139 | 51.487 | 53.026 | 1.00 | 0.00 | C |
| ATOM | 5802 | OE1 | GLU | A | 396 | 35.993 | 51.773 | 51.800 | 1.00 | 0.00 | O |
| ATOM | 5803 | OE2 | GLU | A | 396 | 37.160 | 51.831 | 53.688 | 1.00 | 0.00 | O |
| ATOM | 5804 | C | GLU | A | 396 | 32.147 | 49.943 | 52.623 | 1.00 | 0.00 | C |
| ATOM | 5805 | O | GLU | A | 396 | 31.603 | 49.151 | 53.386 | 1.00 | 0.00 | O |
| ATOM | 5806 | N | LEU | A | 397 | 31.500 | 51.126 | 52.247 | 1.00 | 0.00 | N |
| ATOM | 5807 | H | LEU | A | 397 | 31.871 | 51.605 | 51.452 | 1.00 | 0.00 | H |
| ATOM | 5808 | CA | LEU | A | 397 | 30.323 | 51.743 | 52.904 | 1.00 | 0.00 | C |
| ATOM | 5809 | HA | LEU | A | 397 | 30.222 | 51.175 | 53.721 | 1.00 | 0.00 | H |
| ATOM | 5810 | CB | LEU | A | 397 | 30.660 | 53.206 | 53.251 | 1.00 | 0.00 | C |
| ATOM | 5811 | HB1 | LEU | A | 397 | 29.863 | 53.570 | 53.733 | 1.00 | 0.00 | H |
| ATOM | 5812 | HB2 | LEU | A | 397 | 30.768 | 53.683 | 52.379 | 1.00 | 0.00 | H |
| ATOM | 5813 | CG | LEU | A | 397 | 31.924 | 53.538 | 54.123 | 1.00 | 0.00 | C |
| ATOM | 5814 | HG | LEU | A | 397 | 32.665 | 53.136 | 53.585 | 1.00 | 0.00 | H |
| ATOM | 5815 | CD1 | LEU | A | 397 | 32.044 | 55.074 | 54.343 | 1.00 | 0.00 | C |
| ATOM | 5816 | HD11 | LEU | A | 397 | 32.852 | 55.269 | 54.899 | 1.00 | 0.00 | H |
| ATOM | 5817 | HD12 | LEU | A | 397 | 32.130 | 55.532 | 53.458 | 1.00 | 0.00 | H |
| ATOM | 5818 | HD13 | LEU | A | 397 | 31.227 | 55.408 | 54.813 | 1.00 | 0.00 | H |
| ATOM | 5819 | CD2 | LEU | A | 397 | 31.941 | 52.946 | 55.593 | 1.00 | 0.00 | C |
| ATOM | 5820 | HD21 | LEU | A | 397 | 32.786 | 53.219 | 56.052 | 1.00 | 0.00 | H |
| ATOM | 5821 | HD22 | LEU | A | 397 | 31.156 | 53.298 | 56.103 | 1.00 | 0.00 | H |
| ATOM | 5822 | HD23 | LEU | A | 397 | 31.891 | 51.948 | 55.551 | 1.00 | 0.00 | H |
| ATOM | 5823 | C | LEU | A | 397 | 28.949 | 51.680 | 52.087 | 1.00 | 0.00 | C |
| ATOM | 5824 | O | LEU | A | 397 | 28.807 | 51.440 | 50.912 | 1.00 | 0.00 | O |
| ATOM | 5825 | N | GLU | A | 398 | 27.916 | 51.837 | 52.895 | 1.00 | 0.00 | N |
| ATOM | 5826 | H | GLU | A | 398 | 28.096 | 51.940 | 53.873 | 1.00 | 0.00 | H |
| ATOM | 5827 | CA | GLU | A | 398 | 26.480 | 51.870 | 52.429 | 1.00 | 0.00 | C |
| ATOM | 5828 | HA | GLU | A | 398 | 26.456 | 51.307 | 51.603 | 1.00 | 0.00 | H |
| ATOM | 5829 | CB | GLU | A | 398 | 25.567 | 51.272 | 53.505 | 1.00 | 0.00 | C |

| | | | | | | | | | | | |
|---|---|---|---|---|---|---|---|---|---|---|---|
| ATOM | 5830 | HB1 | GLU | A | 398 | 24.617 | 51.372 | 53.208 | 1.00 | 0.00 | H |
| ATOM | 5831 | HB2 | GLU | A | 398 | 25.705 | 51.778 | 54.356 | 1.00 | 0.00 | H |
| ATOM | 5832 | CG | GLU | A | 398 | 25.884 | 49.755 | 53.741 | 1.00 | 0.00 | C |
| ATOM | 5833 | HG1 | GLU | A | 398 | 26.828 | 49.669 | 54.059 | 1.00 | 0.00 | H |
| ATOM | 5834 | HG2 | GLU | A | 398 | 25.778 | 49.265 | 52.876 | 1.00 | 0.00 | H |
| ATOM | 5835 | CD | GLU | A | 398 | 24.931 | 49.112 | 54.806 | 1.00 | 0.00 | C |
| ATOM | 5836 | OE1 | GLU | A | 398 | 25.291 | 48.077 | 55.394 | 1.00 | 0.00 | O |
| ATOM | 5837 | OE2 | GLU | A | 398 | 23.756 | 49.576 | 55.000 | 1.00 | 0.00 | O |
| ATOM | 5838 | C | GLU | A | 398 | 25.990 | 53.286 | 52.038 | 1.00 | 0.00 | C |
| ATOM | 5839 | O | GLU | A | 398 | 24.791 | 53.523 | 51.694 | 1.00 | 0.00 | O |
| ATOM | 5840 | N | VAL | A | 399 | 26.909 | 54.275 | 52.074 | 1.00 | 0.00 | N |
| ATOM | 5841 | H | VAL | A | 399 | 27.825 | 54.007 | 52.373 | 1.00 | 0.00 | H |
| ATOM | 5842 | CA | VAL | A | 399 | 26.744 | 55.646 | 51.743 | 1.00 | 0.00 | C |
| ATOM | 5843 | HA | VAL | A | 399 | 25.927 | 55.798 | 51.187 | 1.00 | 0.00 | H |
| ATOM | 5844 | CB | VAL | A | 399 | 26.573 | 56.452 | 53.142 | 1.00 | 0.00 | C |
| ATOM | 5845 | HB | VAL | A | 399 | 26.377 | 57.416 | 52.961 | 1.00 | 0.00 | H |
| ATOM | 5846 | CG1 | VAL | A | 399 | 25.436 | 55.911 | 53.891 | 1.00 | 0.00 | C |
| ATOM | 5847 | HG11 | VAL | A | 399 | 25.335 | 56.411 | 54.751 | 1.00 | 0.00 | H |
| ATOM | 5848 | HG12 | VAL | A | 399 | 24.602 | 56.010 | 53.348 | 1.00 | 0.00 | H |
| ATOM | 5849 | HG13 | VAL | A | 399 | 25.595 | 54.943 | 54.085 | 1.00 | 0.00 | H |
| ATOM | 5850 | CG2 | VAL | A | 399 | 27.824 | 56.394 | 54.056 | 1.00 | 0.00 | C |
| ATOM | 5851 | HG21 | VAL | A | 399 | 27.648 | 56.909 | 54.895 | 1.00 | 0.00 | H |
| ATOM | 5852 | HG22 | VAL | A | 399 | 28.025 | 55.442 | 54.285 | 1.00 | 0.00 | H |
| ATOM | 5853 | HG23 | VAL | A | 399 | 28.605 | 56.794 | 53.577 | 1.00 | 0.00 | H |
| ATOM | 5854 | C | VAL | A | 399 | 27.923 | 55.992 | 50.812 | 1.00 | 0.00 | C |
| ATOM | 5855 | O | VAL | A | 399 | 28.834 | 55.169 | 50.619 | 1.00 | 0.00 | O |
| ATOM | 5856 | N | PRO | A | 400 | 28.024 | 57.251 | 50.315 | 1.00 | 0.00 | N |
| ATOM | 5857 | CD | PRO | A | 400 | 27.015 | 58.296 | 50.422 | 1.00 | 0.00 | C |
| ATOM | 5858 | HD1 | PRO | A | 400 | 27.237 | 58.900 | 51.187 | 1.00 | 0.00 | H |
| ATOM | 5859 | HD2 | PRO | A | 400 | 26.117 | 57.882 | 50.573 | 1.00 | 0.00 | H |
| ATOM | 5860 | CG | PRO | A | 400 | 27.063 | 59.043 | 49.096 | 1.00 | 0.00 | C |
| ATOM | 5861 | HG1 | PRO | A | 400 | 26.713 | 59.976 | 49.180 | 1.00 | 0.00 | H |
| ATOM | 5862 | HG2 | PRO | A | 400 | 26.560 | 58.562 | 48.378 | 1.00 | 0.00 | H |
| ATOM | 5863 | CB | PRO | A | 400 | 28.545 | 59.030 | 48.836 | 1.00 | 0.00 | C |
| ATOM | 5864 | HB1 | PRO | A | 400 | 28.971 | 59.820 | 49.276 | 1.00 | 0.00 | H |
| ATOM | 5865 | HB2 | PRO | A | 400 | 28.714 | 59.066 | 47.851 | 1.00 | 0.00 | H |
| ATOM | 5866 | CA | PRO | A | 400 | 29.075 | 57.693 | 49.448 | 1.00 | 0.00 | C |
| ATOM | 5867 | HA | PRO | A | 400 | 29.245 | 56.943 | 48.808 | 1.00 | 0.00 | H |
| ATOM | 5868 | C | PRO | A | 400 | 30.413 | 57.867 | 50.105 | 1.00 | 0.00 | C |
| ATOM | 5869 | O | PRO | A | 400 | 30.529 | 58.348 | 51.239 | 1.00 | 0.00 | O |
| ATOM | 5870 | N | LYS | A | 401 | 31.451 | 57.607 | 49.289 | 1.00 | 0.00 | N |
| ATOM | 5871 | H | LYS | A | 401 | 31.272 | 57.157 | 48.414 | 1.00 | 0.00 | H |
| ATOM | 5872 | CA | LYS | A | 401 | 32.798 | 57.947 | 49.620 | 1.00 | 0.00 | C |
| ATOM | 5873 | HA | LYS | A | 401 | 32.698 | 58.277 | 50.559 | 1.00 | 0.00 | H |
| ATOM | 5874 | CB | LYS | A | 401 | 33.770 | 56.784 | 49.691 | 1.00 | 0.00 | C |
| ATOM | 5875 | HB1 | LYS | A | 401 | 34.664 | 57.156 | 49.940 | 1.00 | 0.00 | H |
| ATOM | 5876 | HB2 | LYS | A | 401 | 33.820 | 56.380 | 48.778 | 1.00 | 0.00 | H |
| ATOM | 5877 | CG | LYS | A | 401 | 33.407 | 55.666 | 50.701 | 1.00 | 0.00 | C |
| ATOM | 5878 | HG1 | LYS | A | 401 | 32.511 | 55.282 | 50.479 | 1.00 | 0.00 | H |
| ATOM | 5879 | HG2 | LYS | A | 401 | 33.391 | 56.036 | 51.630 | 1.00 | 0.00 | H |
| ATOM | 5880 | CD | LYS | A | 401 | 34.455 | 54.569 | 50.618 | 1.00 | 0.00 | C |
| ATOM | 5881 | HD1 | LYS | A | 401 | 34.507 | 54.251 | 49.671 | 1.00 | 0.00 | H |
| ATOM | 5882 | HD2 | LYS | A | 401 | 34.170 | 53.814 | 51.208 | 1.00 | 0.00 | H |
| ATOM | 5883 | CE | LYS | A | 401 | 35.843 | 55.040 | 51.056 | 1.00 | 0.00 | C |

```
ATOM   5884  HE1 LYS A 401      35.835  55.240  52.036  1.00  0.00           H
ATOM   5885  HE2 LYS A 401      36.092  55.866  50.550  1.00  0.00           H
ATOM   5886  NZ  LYS A 401      36.875  53.997  50.795  1.00  0.00           N
ATOM   5887  HZ1 LYS A 401      37.770  54.332  51.091  1.00  0.00           H
ATOM   5888  HZ2 LYS A 401      36.646  53.167  51.303  1.00  0.00           H
ATOM   5889  HZ3 LYS A 401      36.902  53.793  49.817  1.00  0.00           H
ATOM   5890  C   LYS A 401      33.306  59.013  48.638  1.00  0.00           C
ATOM   5891  O   LYS A 401      33.368  58.734  47.446  1.00  0.00           O
ATOM   5892  N   PRO A 402      33.620  60.227  49.092  1.00  0.00           N
ATOM   5893  CD  PRO A 402      33.489  60.727  50.474  1.00  0.00           C
ATOM   5894  HD1 PRO A 402      34.133  60.231  51.056  1.00  0.00           H
ATOM   5895  HD2 PRO A 402      32.555  60.557  50.789  1.00  0.00           H
ATOM   5896  CG  PRO A 402      33.796  62.234  50.501  1.00  0.00           C
ATOM   5897  HG1 PRO A 402      34.741  62.399  50.782  1.00  0.00           H
ATOM   5898  HG2 PRO A 402      33.175  62.712  51.122  1.00  0.00           H
ATOM   5899  CB  PRO A 402      33.594  62.732  49.115  1.00  0.00           C
ATOM   5900  HB1 PRO A 402      34.245  63.464  48.917  1.00  0.00           H
ATOM   5901  HB2 PRO A 402      32.662  63.078  49.009  1.00  0.00           H
ATOM   5902  CA  PRO A 402      33.853  61.461  48.228  1.00  0.00           C
ATOM   5903  HA  PRO A 402      33.190  61.426  47.481  1.00  0.00           H
ATOM   5904  C   PRO A 402      35.323  61.475  47.579  1.00  0.00           C
ATOM   5905  O   PRO A 402      35.583  62.200  46.607  1.00  0.00           O
ATOM   5906  N   GLY A 403      36.176  60.586  47.956  1.00  0.00           N
ATOM   5907  H   GLY A 403      35.887  59.960  48.680  1.00  0.00           H
ATOM   5908  CA  GLY A 403      37.528  60.405  47.426  1.00  0.00           C
ATOM   5909  HA1 GLY A 403      38.040  59.835  48.068  1.00  0.00           H
ATOM   5910  HA2 GLY A 403      37.960  61.305  47.361  1.00  0.00           H
ATOM   5911  C   GLY A 403      37.500  59.742  46.052  1.00  0.00           C
ATOM   5912  O   GLY A 403      36.499  59.158  45.690  1.00  0.00           O
ATOM   5913  N   GLN A 404      38.566  59.968  45.275  1.00  0.00           N
ATOM   5914  H   GLN A 404      39.312  60.477  45.705  1.00  0.00           H
ATOM   5915  CA  GLN A 404      38.795  59.570  43.868  1.00  0.00           C
ATOM   5916  HA  GLN A 404      38.123  60.039  43.295  1.00  0.00           H
ATOM   5917  CB  GLN A 404      40.173  60.027  43.409  1.00  0.00           C
ATOM   5918  HB1 GLN A 404      40.852  59.580  43.991  1.00  0.00           H
ATOM   5919  HB2 GLN A 404      40.224  61.017  43.537  1.00  0.00           H
ATOM   5920  CG  GLN A 404      40.456  59.670  41.886  1.00  0.00           C
ATOM   5921  HG1 GLN A 404      39.835  60.208  41.316  1.00  0.00           H
ATOM   5922  HG2 GLN A 404      40.272  58.696  41.752  1.00  0.00           H
ATOM   5923  CD  GLN A 404      41.855  59.917  41.339  1.00  0.00           C
ATOM   5924  OE1 GLN A 404      42.636  60.632  41.992  1.00  0.00           O
ATOM   5925  NE2 GLN A 404      42.170  59.492  40.154  1.00  0.00           N
ATOM   5926 HE21 GLN A 404      41.494  59.003  39.602  1.00  0.00           H
ATOM   5927 HE22 GLN A 404      43.088  59.653  39.793  1.00  0.00           H
ATOM   5928  C   GLN A 404      38.588  58.058  43.722  1.00  0.00           C
ATOM   5929  O   GLN A 404      39.132  57.259  44.525  1.00  0.00           O
ATOM   5930  N   PRO A 405      37.843  57.598  42.684  1.00  0.00           N
ATOM   5931  CD  PRO A 405      37.100  58.432  41.689  1.00  0.00           C
ATOM   5932  HD1 PRO A 405      37.705  59.123  41.293  1.00  0.00           H
ATOM   5933  HD2 PRO A 405      36.320  58.881  42.125  1.00  0.00           H
ATOM   5934  CG  PRO A 405      36.627  57.506  40.632  1.00  0.00           C
ATOM   5935  HG1 PRO A 405      37.269  57.468  39.867  1.00  0.00           H
ATOM   5936  HG2 PRO A 405      35.723  57.768  40.296  1.00  0.00           H
ATOM   5937  CB  PRO A 405      36.585  56.211  41.387  1.00  0.00           C
```

| | | | | | | | | | | | |
|---|---|---|---|---|---|---|---|---|---|---|---|
| ATOM | 5938 | HB1 | PRO | A | 405 | 36.735 | 55.453 | 40.752 | 1.00 | 0.00 | H |
| ATOM | 5939 | HB2 | PRO | A | 405 | 35.690 | 56.110 | 41.822 | 1.00 | 0.00 | H |
| ATOM | 5940 | CA | PRO | A | 405 | 37.646 | 56.187 | 42.438 | 1.00 | 0.00 | C |
| ATOM | 5941 | HA | PRO | A | 405 | 37.385 | 55.757 | 43.302 | 1.00 | 0.00 | H |
| ATOM | 5942 | C | PRO | A | 405 | 38.905 | 55.403 | 41.935 | 1.00 | 0.00 | C |
| ATOM | 5943 | O | PRO | A | 405 | 39.865 | 56.012 | 41.443 | 1.00 | 0.00 | O |
| ATOM | 5944 | N | SER | A | 406 | 38.718 | 54.081 | 41.844 | 1.00 | 0.00 | N |
| ATOM | 5945 | H | SER | A | 406 | 37.894 | 53.714 | 42.276 | 1.00 | 0.00 | H |
| ATOM | 5946 | CA | SER | A | 406 | 39.589 | 53.123 | 41.181 | 1.00 | 0.00 | C |
| ATOM | 5947 | HA | SER | A | 406 | 40.502 | 53.408 | 41.472 | 1.00 | 0.00 | H |
| ATOM | 5948 | CB | SER | A | 406 | 39.172 | 51.722 | 41.607 | 1.00 | 0.00 | C |
| ATOM | 5949 | HB1 | SER | A | 406 | 39.724 | 51.025 | 41.149 | 1.00 | 0.00 | H |
| ATOM | 5950 | HB2 | SER | A | 406 | 38.204 | 51.559 | 41.416 | 1.00 | 0.00 | H |
| ATOM | 5951 | OG | SER | A | 406 | 39.355 | 51.560 | 42.969 | 1.00 | 0.00 | O |
| ATOM | 5952 | HG | SER | A | 406 | 39.078 | 50.637 | 43.235 | 1.00 | 0.00 | H |
| ATOM | 5953 | C | SER | A | 406 | 39.479 | 53.129 | 39.647 | 1.00 | 0.00 | C |
| ATOM | 5954 | O | SER | A | 406 | 40.194 | 52.431 | 38.916 | 1.00 | 0.00 | O |
| ATOM | 5955 | N | GLU | A | 407 | 38.620 | 53.950 | 39.016 | 1.00 | 0.00 | N |
| ATOM | 5956 | H | GLU | A | 407 | 37.989 | 54.476 | 39.587 | 1.00 | 0.00 | H |
| ATOM | 5957 | CA | GLU | A | 407 | 38.527 | 54.142 | 37.561 | 1.00 | 0.00 | C |
| ATOM | 5958 | HA | GLU | A | 407 | 39.463 | 54.218 | 37.217 | 1.00 | 0.00 | H |
| ATOM | 5959 | CB | GLU | A | 407 | 37.981 | 52.892 | 36.838 | 1.00 | 0.00 | C |
| ATOM | 5960 | HB1 | GLU | A | 407 | 38.536 | 52.107 | 37.113 | 1.00 | 0.00 | H |
| ATOM | 5961 | HB2 | GLU | A | 407 | 38.074 | 53.037 | 35.853 | 1.00 | 0.00 | H |
| ATOM | 5962 | CG | GLU | A | 407 | 36.517 | 52.549 | 37.121 | 1.00 | 0.00 | C |
| ATOM | 5963 | HG1 | GLU | A | 407 | 36.225 | 51.822 | 36.499 | 1.00 | 0.00 | H |
| ATOM | 5964 | HG2 | GLU | A | 407 | 35.955 | 53.362 | 36.969 | 1.00 | 0.00 | H |
| ATOM | 5965 | CD | GLU | A | 407 | 36.325 | 52.077 | 38.557 | 1.00 | 0.00 | C |
| ATOM | 5966 | OE1 | GLU | A | 407 | 35.849 | 52.915 | 39.358 | 1.00 | 0.00 | O |
| ATOM | 5967 | OE2 | GLU | A | 407 | 36.469 | 50.862 | 38.786 | 1.00 | 0.00 | O |
| ATOM | 5968 | C | GLU | A | 407 | 37.892 | 55.496 | 37.229 | 1.00 | 0.00 | C |
| ATOM | 5969 | O | GLU | A | 407 | 36.962 | 55.955 | 37.965 | 1.00 | 0.00 | O |
| ATOM | 5970 | N | PRO | A | 408 | 38.286 | 56.216 | 36.172 | 1.00 | 0.00 | N |
| ATOM | 5971 | CD | PRO | A | 408 | 39.245 | 55.797 | 35.097 | 1.00 | 0.00 | C |
| ATOM | 5972 | HD1 | PRO | A | 408 | 38.748 | 55.385 | 34.333 | 1.00 | 0.00 | H |
| ATOM | 5973 | HD2 | PRO | A | 408 | 39.903 | 55.137 | 35.460 | 1.00 | 0.00 | H |
| ATOM | 5974 | CG | PRO | A | 408 | 39.951 | 57.063 | 34.652 | 1.00 | 0.00 | C |
| ATOM | 5975 | HG1 | PRO | A | 408 | 40.332 | 56.977 | 33.731 | 1.00 | 0.00 | H |
| ATOM | 5976 | HG2 | PRO | A | 408 | 40.673 | 57.334 | 35.288 | 1.00 | 0.00 | H |
| ATOM | 5977 | CB | PRO | A | 408 | 38.752 | 58.009 | 34.698 | 1.00 | 0.00 | C |
| ATOM | 5978 | HB1 | PRO | A | 408 | 38.239 | 57.942 | 33.842 | 1.00 | 0.00 | H |
| ATOM | 5979 | HB2 | PRO | A | 408 | 39.071 | 58.949 | 34.823 | 1.00 | 0.00 | H |
| ATOM | 5980 | CA | PRO | A | 408 | 37.861 | 57.570 | 35.903 | 1.00 | 0.00 | C |
| ATOM | 5981 | HA | PRO | A | 408 | 37.949 | 58.150 | 36.713 | 1.00 | 0.00 | H |
| ATOM | 5982 | C | PRO | A | 408 | 36.423 | 57.677 | 35.712 | 1.00 | 0.00 | C |
| ATOM | 5983 | O | PRO | A | 408 | 35.807 | 56.894 | 34.964 | 1.00 | 0.00 | O |
| ATOM | 5984 | N | GLU | A | 409 | 35.823 | 58.644 | 36.451 | 1.00 | 0.00 | N |
| ATOM | 5985 | H | GLU | A | 409 | 36.412 | 59.266 | 36.967 | 1.00 | 0.00 | H |
| ATOM | 5986 | CA | GLU | A | 409 | 34.333 | 58.849 | 36.550 | 1.00 | 0.00 | C |
| ATOM | 5987 | HA | GLU | A | 409 | 34.243 | 59.457 | 37.339 | 1.00 | 0.00 | H |
| ATOM | 5988 | CB | GLU | A | 409 | 33.740 | 59.693 | 35.386 | 1.00 | 0.00 | C |
| ATOM | 5989 | HB1 | GLU | A | 409 | 32.824 | 59.995 | 35.649 | 1.00 | 0.00 | H |
| ATOM | 5990 | HB2 | GLU | A | 409 | 33.679 | 59.113 | 34.574 | 1.00 | 0.00 | H |
| ATOM | 5991 | CG | GLU | A | 409 | 34.556 | 60.938 | 35.013 | 1.00 | 0.00 | C |

| | | | | | | | | | | | |
|---|---|---|---|---|---|---|---|---|---|---|---|
| ATOM | 5992 | HG1 | GLU | A | 409 | 33.955 | 61.567 | 34.521 | 1.00 | 0.00 | H |
| ATOM | 5993 | HG2 | GLU | A | 409 | 35.302 | 60.652 | 34.412 | 1.00 | 0.00 | H |
| ATOM | 5994 | CD | GLU | A | 409 | 35.164 | 61.692 | 36.205 | 1.00 | 0.00 | C |
| ATOM | 5995 | OE1 | GLU | A | 409 | 34.465 | 61.995 | 37.206 | 1.00 | 0.00 | O |
| ATOM | 5996 | OE2 | GLU | A | 409 | 36.359 | 62.116 | 36.140 | 1.00 | 0.00 | O |
| ATOM | 5997 | C | GLU | A | 409 | 33.537 | 57.530 | 36.930 | 1.00 | 0.00 | C |
| ATOM | 5998 | O | GLU | A | 409 | 32.340 | 57.342 | 36.552 | 1.00 | 0.00 | O |
| ATOM | 5999 | N | HIS | A | 410 | 34.081 | 56.603 | 37.692 | 1.00 | 0.00 | N |
| ATOM | 6000 | H | HIS | A | 410 | 35.013 | 56.750 | 38.022 | 1.00 | 0.00 | H |
| ATOM | 6001 | CA | HIS | A | 410 | 33.392 | 55.363 | 38.084 | 1.00 | 0.00 | C |
| ATOM | 6002 | HA | HIS | A | 410 | 34.136 | 54.894 | 38.561 | 1.00 | 0.00 | H |
| ATOM | 6003 | CB | HIS | A | 410 | 32.194 | 55.666 | 38.994 | 1.00 | 0.00 | C |
| ATOM | 6004 | HB1 | HIS | A | 410 | 31.851 | 54.800 | 39.357 | 1.00 | 0.00 | H |
| ATOM | 6005 | HB2 | HIS | A | 410 | 31.483 | 56.100 | 38.441 | 1.00 | 0.00 | H |
| ATOM | 6006 | CG | HIS | A | 410 | 32.543 | 56.603 | 40.183 | 1.00 | 0.00 | C |
| ATOM | 6007 | ND1 | HIS | A | 410 | 32.402 | 57.985 | 40.294 | 1.00 | 0.00 | N |
| ATOM | 6008 | CE1 | HIS | A | 410 | 32.728 | 58.380 | 41.579 | 1.00 | 0.00 | C |
| ATOM | 6009 | HE1 | HIS | A | 410 | 32.710 | 59.313 | 41.938 | 1.00 | 0.00 | H |
| ATOM | 6010 | NE2 | HIS | A | 410 | 33.070 | 57.282 | 42.245 | 1.00 | 0.00 | N |
| ATOM | 6011 | HE2 | HIS | A | 410 | 33.392 | 57.266 | 43.192 | 1.00 | 0.00 | H |
| ATOM | 6012 | CD2 | HIS | A | 410 | 32.915 | 56.145 | 41.430 | 1.00 | 0.00 | C |
| ATOM | 6013 | HD2 | HIS | A | 410 | 33.048 | 55.192 | 41.702 | 1.00 | 0.00 | H |
| ATOM | 6014 | C | HIS | A | 410 | 33.006 | 54.384 | 36.992 | 1.00 | 0.00 | C |
| ATOM | 6015 | O | HIS | A | 410 | 32.466 | 53.359 | 37.325 | 1.00 | 0.00 | O |
| ATOM | 6016 | N | GLY | A | 411 | 33.261 | 54.577 | 35.742 | 1.00 | 0.00 | N |
| ATOM | 6017 | H | GLY | A | 411 | 33.617 | 55.477 | 35.488 | 1.00 | 0.00 | H |
| ATOM | 6018 | CA | GLY | A | 411 | 33.082 | 53.613 | 34.662 | 1.00 | 0.00 | C |
| ATOM | 6019 | HA1 | GLY | A | 411 | 32.103 | 53.439 | 34.554 | 1.00 | 0.00 | H |
| ATOM | 6020 | HA2 | GLY | A | 411 | 33.545 | 52.764 | 34.918 | 1.00 | 0.00 | H |
| ATOM | 6021 | C | GLY | A | 411 | 33.643 | 54.115 | 33.370 | 1.00 | 0.00 | C |
| ATOM | 6022 | O | GLY | A | 411 | 33.194 | 55.184 | 32.852 | 1.00 | 0.00 | O |
| ATOM | 6023 | N | PRO | A | 412 | 34.539 | 53.319 | 32.687 | 1.00 | 0.00 | N |
| ATOM | 6024 | CD | PRO | A | 412 | 35.446 | 52.256 | 33.149 | 1.00 | 0.00 | C |
| ATOM | 6025 | HD1 | PRO | A | 412 | 34.935 | 51.399 | 33.208 | 1.00 | 0.00 | H |
| ATOM | 6026 | HD2 | PRO | A | 412 | 35.794 | 52.500 | 34.054 | 1.00 | 0.00 | H |
| ATOM | 6027 | CG | PRO | A | 412 | 36.588 | 52.076 | 32.223 | 1.00 | 0.00 | C |
| ATOM | 6028 | HG1 | PRO | A | 412 | 36.844 | 51.111 | 32.168 | 1.00 | 0.00 | H |
| ATOM | 6029 | HG2 | PRO | A | 412 | 37.374 | 52.612 | 32.531 | 1.00 | 0.00 | H |
| ATOM | 6030 | CB | PRO | A | 412 | 35.992 | 52.612 | 30.875 | 1.00 | 0.00 | C |
| ATOM | 6031 | HB1 | PRO | A | 412 | 35.557 | 51.881 | 30.350 | 1.00 | 0.00 | H |
| ATOM | 6032 | HB2 | PRO | A | 412 | 36.693 | 53.053 | 30.314 | 1.00 | 0.00 | H |
| ATOM | 6033 | CA | PRO | A | 412 | 34.962 | 53.623 | 31.304 | 1.00 | 0.00 | C |
| ATOM | 6034 | HA | PRO | A | 412 | 35.326 | 54.546 | 31.426 | 1.00 | 0.00 | H |
| ATOM | 6035 | C | PRO | A | 412 | 33.873 | 53.717 | 30.188 | 1.00 | 0.00 | C |
| ATOM | 6036 | O | PRO | A | 412 | 32.717 | 53.473 | 30.412 | 1.00 | 0.00 | O |
| ATOM | 6037 | N | SER | A | 413 | 34.289 | 54.191 | 29.016 | 1.00 | 0.00 | N |
| ATOM | 6038 | H | SER | A | 413 | 35.218 | 54.560 | 29.020 | 1.00 | 0.00 | H |
| ATOM | 6039 | CA | SER | A | 413 | 33.590 | 54.262 | 27.688 | 1.00 | 0.00 | C |
| ATOM | 6040 | HA | SER | A | 413 | 33.086 | 53.421 | 27.491 | 1.00 | 0.00 | H |
| ATOM | 6041 | CB | SER | A | 413 | 32.624 | 55.515 | 27.784 | 1.00 | 0.00 | C |
| ATOM | 6042 | HB1 | SER | A | 413 | 33.095 | 56.330 | 28.120 | 1.00 | 0.00 | H |
| ATOM | 6043 | HB2 | SER | A | 413 | 31.824 | 55.332 | 28.356 | 1.00 | 0.00 | H |
| ATOM | 6044 | OG | SER | A | 413 | 32.163 | 55.795 | 26.483 | 1.00 | 0.00 | O |
| ATOM | 6045 | HG | SER | A | 413 | 31.549 | 56.584 | 26.510 | 1.00 | 0.00 | H |

| | | | | | | | | | | | |
|---|---|---|---|---|---|---|---|---|---|---|---|
| ATOM | 6046 | C | SER | A | 413 | 34.655 | 54.410 | 26.576 | 1.00 | 0.00 | C |
| ATOM | 6047 | O | SER | A | 413 | 35.646 | 55.177 | 26.708 | 1.00 | 0.00 | O |
| ATOM | 6048 | N | PRO | A | 414 | 34.534 | 53.599 | 25.489 | 1.00 | 0.00 | N |
| ATOM | 6049 | CD | PRO | A | 414 | 34.198 | 52.148 | 25.692 | 1.00 | 0.00 | C |
| ATOM | 6050 | HD1 | PRO | A | 414 | 33.207 | 52.017 | 25.709 | 1.00 | 0.00 | H |
| ATOM | 6051 | HD2 | PRO | A | 414 | 34.593 | 51.813 | 26.547 | 1.00 | 0.00 | H |
| ATOM | 6052 | CG | PRO | A | 414 | 34.775 | 51.411 | 24.559 | 1.00 | 0.00 | C |
| ATOM | 6053 | HG1 | PRO | A | 414 | 34.180 | 50.652 | 24.296 | 1.00 | 0.00 | H |
| ATOM | 6054 | HG2 | PRO | A | 414 | 35.681 | 51.059 | 24.795 | 1.00 | 0.00 | H |
| ATOM | 6055 | CB | PRO | A | 414 | 34.853 | 52.459 | 23.470 | 1.00 | 0.00 | C |
| ATOM | 6056 | HB1 | PRO | A | 414 | 33.968 | 52.574 | 23.018 | 1.00 | 0.00 | H |
| ATOM | 6057 | HB2 | PRO | A | 414 | 35.546 | 52.219 | 22.790 | 1.00 | 0.00 | H |
| ATOM | 6058 | CA | PRO | A | 414 | 35.256 | 53.719 | 24.300 | 1.00 | 0.00 | C |
| ATOM | 6059 | HA | PRO | A | 414 | 36.221 | 53.737 | 24.562 | 1.00 | 0.00 | H |
| ATOM | 6060 | C | PRO | A | 414 | 35.002 | 55.073 | 23.567 | 1.00 | 0.00 | C |
| ATOM | 6061 | O | PRO | A | 414 | 35.522 | 55.239 | 22.488 | 1.00 | 0.00 | O |
| ATOM | 6062 | N | GLY | A | 415 | 34.110 | 55.926 | 24.040 | 1.00 | 0.00 | N |
| ATOM | 6063 | H | GLY | A | 415 | 33.594 | 55.704 | 24.867 | 1.00 | 0.00 | H |
| ATOM | 6064 | CA | GLY | A | 415 | 33.878 | 57.205 | 23.345 | 1.00 | 0.00 | C |
| ATOM | 6065 | HA1 | GLY | A | 415 | 34.757 | 57.525 | 22.992 | 1.00 | 0.00 | H |
| ATOM | 6066 | HA2 | GLY | A | 415 | 33.531 | 57.859 | 24.017 | 1.00 | 0.00 | H |
| ATOM | 6067 | C | GLY | A | 415 | 32.884 | 57.089 | 22.193 | 1.00 | 0.00 | C |
| ATOM | 6068 | O | GLY | A | 415 | 32.751 | 57.988 | 21.374 | 1.00 | 0.00 | O |
| ATOM | 6069 | N | GLN | A | 416 | 32.203 | 55.924 | 22.169 | 1.00 | 0.00 | N |
| ATOM | 6070 | H | GLN | A | 416 | 32.320 | 55.268 | 22.914 | 1.00 | 0.00 | H |
| ATOM | 6071 | CA | GLN | A | 416 | 31.283 | 55.606 | 21.049 | 1.00 | 0.00 | C |
| ATOM | 6072 | HA | GLN | A | 416 | 31.507 | 56.254 | 20.321 | 1.00 | 0.00 | H |
| ATOM | 6073 | CB | GLN | A | 416 | 31.498 | 54.124 | 20.621 | 1.00 | 0.00 | C |
| ATOM | 6074 | HB1 | GLN | A | 416 | 30.777 | 53.876 | 19.974 | 1.00 | 0.00 | H |
| ATOM | 6075 | HB2 | GLN | A | 416 | 31.428 | 53.548 | 21.435 | 1.00 | 0.00 | H |
| ATOM | 6076 | CG | GLN | A | 416 | 32.862 | 53.839 | 19.950 | 1.00 | 0.00 | C |
| ATOM | 6077 | HG1 | GLN | A | 416 | 33.544 | 53.675 | 20.663 | 1.00 | 0.00 | H |
| ATOM | 6078 | HG2 | GLN | A | 416 | 33.129 | 54.637 | 19.410 | 1.00 | 0.00 | H |
| ATOM | 6079 | CD | GLN | A | 416 | 32.843 | 52.630 | 19.027 | 1.00 | 0.00 | C |
| ATOM | 6080 | OE1 | GLN | A | 416 | 33.030 | 52.708 | 17.819 | 1.00 | 0.00 | O |
| ATOM | 6081 | NE2 | GLN | A | 416 | 32.623 | 51.481 | 19.681 | 1.00 | 0.00 | N |
| ATOM | 6082 | HE21 | GLN | A | 416 | 32.493 | 51.487 | 20.672 | 1.00 | 0.00 | H |
| ATOM | 6083 | HE22 | GLN | A | 416 | 32.589 | 50.618 | 19.177 | 1.00 | 0.00 | H |
| ATOM | 6084 | C | GLN | A | 416 | 29.820 | 55.821 | 21.422 | 1.00 | 0.00 | C |
| ATOM | 6085 | O | GLN | A | 416 | 29.477 | 56.109 | 22.575 | 1.00 | 0.00 | O |
| ATOM | 6086 | N | ASP | A | 417 | 28.948 | 55.628 | 20.364 | 1.00 | 0.00 | N |
| ATOM | 6087 | H | ASP | A | 417 | 29.336 | 55.500 | 19.451 | 1.00 | 0.00 | H |
| ATOM | 6088 | CA | ASP | A | 417 | 27.524 | 55.601 | 20.502 | 1.00 | 0.00 | C |
| ATOM | 6089 | HA | ASP | A | 417 | 27.395 | 55.898 | 21.448 | 1.00 | 0.00 | H |
| ATOM | 6090 | CB | ASP | A | 417 | 26.885 | 56.571 | 19.480 | 1.00 | 0.00 | C |
| ATOM | 6091 | HB1 | ASP | A | 417 | 26.935 | 56.143 | 18.578 | 1.00 | 0.00 | H |
| ATOM | 6092 | HB2 | ASP | A | 417 | 27.423 | 57.414 | 19.479 | 1.00 | 0.00 | H |
| ATOM | 6093 | CG | ASP | A | 417 | 25.400 | 56.944 | 19.758 | 1.00 | 0.00 | C |
| ATOM | 6094 | OD1 | ASP | A | 417 | 24.836 | 56.607 | 20.828 | 1.00 | 0.00 | O |
| ATOM | 6095 | OD2 | ASP | A | 417 | 24.774 | 57.522 | 18.873 | 1.00 | 0.00 | O |
| ATOM | 6096 | C | ASP | A | 417 | 26.920 | 54.188 | 20.336 | 1.00 | 0.00 | C |
| ATOM | 6097 | O | ASP | A | 417 | 27.484 | 53.369 | 19.570 | 1.00 | 0.00 | O |
| ATOM | 6098 | N | THR | A | 418 | 25.819 | 53.957 | 20.989 | 1.00 | 0.00 | N |
| ATOM | 6099 | H | THR | A | 418 | 25.462 | 54.710 | 21.542 | 1.00 | 0.00 | H |

| | | | | | | | | | | | |
|---|---|---|---|---|---|---|---|---|---|---|---|
| ATOM | 6100 | CA | THR | A | 418 | 25.046 | 52.714 | 21.002 | 1.00 | 0.00 | C |
| ATOM | 6101 | HA | THR | A | 418 | 25.758 | 52.032 | 21.173 | 1.00 | 0.00 | H |
| ATOM | 6102 | CB | THR | A | 418 | 23.963 | 52.762 | 22.142 | 1.00 | 0.00 | C |
| ATOM | 6103 | HB | THR | A | 418 | 23.070 | 53.108 | 21.853 | 1.00 | 0.00 | H |
| ATOM | 6104 | CG2 | THR | A | 418 | 23.780 | 51.362 | 22.733 | 1.00 | 0.00 | C |
| ATOM | 6105 | HG21 | THR | A | 418 | 23.091 | 51.393 | 23.458 | 1.00 | 0.00 | H |
| ATOM | 6106 | HG22 | THR | A | 418 | 23.480 | 50.734 | 22.015 | 1.00 | 0.00 | H |
| ATOM | 6107 | HG23 | THR | A | 418 | 24.649 | 51.044 | 23.112 | 1.00 | 0.00 | H |
| ATOM | 6108 | OG1 | THR | A | 418 | 24.332 | 53.718 | 23.102 | 1.00 | 0.00 | O |
| ATOM | 6109 | HG1 | THR | A | 418 | 23.642 | 53.747 | 23.826 | 1.00 | 0.00 | H |
| ATOM | 6110 | C | THR | A | 418 | 24.361 | 52.376 | 19.636 | 1.00 | 0.00 | C |
| ATOM | 6111 | O | THR | A | 418 | 23.358 | 51.681 | 19.683 | 1.00 | 0.00 | O |
| ATOM | 6112 | N | PHE | A | 419 | 24.891 | 52.751 | 18.489 | 1.00 | 0.00 | N |
| ATOM | 6113 | H | PHE | A | 419 | 25.790 | 53.188 | 18.525 | 1.00 | 0.00 | H |
| ATOM | 6114 | CA | PHE | A | 419 | 24.269 | 52.582 | 17.136 | 1.00 | 0.00 | C |
| ATOM | 6115 | HA | PHE | A | 419 | 23.340 | 52.951 | 17.174 | 1.00 | 0.00 | H |
| ATOM | 6116 | CB | PHE | A | 419 | 25.154 | 53.391 | 16.099 | 1.00 | 0.00 | C |
| ATOM | 6117 | HB1 | PHE | A | 419 | 26.039 | 52.948 | 15.957 | 1.00 | 0.00 | H |
| ATOM | 6118 | HB2 | PHE | A | 419 | 25.298 | 54.332 | 16.405 | 1.00 | 0.00 | H |
| ATOM | 6119 | CG | PHE | A | 419 | 24.417 | 53.415 | 14.803 | 1.00 | 0.00 | C |
| ATOM | 6120 | CD1 | PHE | A | 419 | 24.737 | 52.527 | 13.794 | 1.00 | 0.00 | C |
| ATOM | 6121 | HD1 | PHE | A | 419 | 25.305 | 51.726 | 13.982 | 1.00 | 0.00 | H |
| ATOM | 6122 | CE1 | PHE | A | 419 | 24.259 | 52.765 | 12.524 | 1.00 | 0.00 | C |
| ATOM | 6123 | HE1 | PHE | A | 419 | 24.473 | 52.127 | 11.784 | 1.00 | 0.00 | H |
| ATOM | 6124 | CZ | PHE | A | 419 | 23.487 | 53.878 | 12.269 | 1.00 | 0.00 | C |
| ATOM | 6125 | HZ | PHE | A | 419 | 23.146 | 54.040 | 11.343 | 1.00 | 0.00 | H |
| ATOM | 6126 | CE2 | PHE | A | 419 | 23.179 | 54.771 | 13.273 | 1.00 | 0.00 | C |
| ATOM | 6127 | HE2 | PHE | A | 419 | 22.622 | 55.576 | 13.071 | 1.00 | 0.00 | H |
| ATOM | 6128 | CD2 | PHE | A | 419 | 23.637 | 54.556 | 14.552 | 1.00 | 0.00 | C |
| ATOM | 6129 | HD2 | PHE | A | 419 | 23.419 | 55.198 | 15.287 | 1.00 | 0.00 | H |
| ATOM | 6130 | C | PHE | A | 419 | 24.072 | 51.083 | 16.767 | 1.00 | 0.00 | C |
| ATOM | 6131 | O | PHE | A | 419 | 23.000 | 50.768 | 16.260 | 1.00 | 0.00 | O |
| ATOM | 6132 | N | CYS | A | 420 | 25.006 | 50.199 | 17.091 | 1.00 | 0.00 | N |
| ATOM | 6133 | H | CYS | A | 420 | 25.768 | 50.448 | 17.689 | 1.00 | 0.00 | H |
| ATOM | 6134 | CA | CYS | A | 420 | 24.885 | 48.852 | 16.547 | 1.00 | 0.00 | C |
| ATOM | 6135 | HA | CYS | A | 420 | 24.449 | 49.023 | 15.664 | 1.00 | 0.00 | H |
| ATOM | 6136 | CB | CYS | A | 420 | 26.261 | 48.236 | 16.327 | 1.00 | 0.00 | C |
| ATOM | 6137 | HB1 | CYS | A | 420 | 26.093 | 47.335 | 15.928 | 1.00 | 0.00 | H |
| ATOM | 6138 | HB2 | CYS | A | 420 | 26.658 | 48.128 | 17.239 | 1.00 | 0.00 | H |
| ATOM | 6139 | SG | CYS | A | 420 | 27.259 | 49.289 | 15.260 | 1.00 | 0.00 | S |
| ATOM | 6140 | C | CYS | A | 420 | 23.968 | 47.894 | 17.397 | 1.00 | 0.00 | C |
| ATOM | 6141 | O | CYS | A | 420 | 23.887 | 46.744 | 16.969 | 1.00 | 0.00 | O |
| ATOM | 6142 | N | GLN | A | 421 | 23.310 | 48.378 | 18.489 | 1.00 | 0.00 | N |
| ATOM | 6143 | H | GLN | A | 421 | 23.531 | 49.314 | 18.762 | 1.00 | 0.00 | H |
| ATOM | 6144 | CA | GLN | A | 421 | 22.345 | 47.700 | 19.282 | 1.00 | 0.00 | C |
| ATOM | 6145 | HA | GLN | A | 421 | 22.837 | 46.896 | 19.616 | 1.00 | 0.00 | H |
| ATOM | 6146 | CB | GLN | A | 421 | 21.892 | 48.628 | 20.426 | 1.00 | 0.00 | C |
| ATOM | 6147 | HB1 | GLN | A | 421 | 21.418 | 49.416 | 20.032 | 1.00 | 0.00 | H |
| ATOM | 6148 | HB2 | GLN | A | 421 | 22.700 | 48.940 | 20.925 | 1.00 | 0.00 | H |
| ATOM | 6149 | CG | GLN | A | 421 | 20.941 | 47.918 | 21.413 | 1.00 | 0.00 | C |
| ATOM | 6150 | HG1 | GLN | A | 421 | 20.336 | 47.306 | 20.904 | 1.00 | 0.00 | H |
| ATOM | 6151 | HG2 | GLN | A | 421 | 20.394 | 48.604 | 21.894 | 1.00 | 0.00 | H |
| ATOM | 6152 | CD | GLN | A | 421 | 21.678 | 47.100 | 22.442 | 1.00 | 0.00 | C |
| ATOM | 6153 | OE1 | GLN | A | 421 | 22.501 | 46.244 | 22.195 | 1.00 | 0.00 | O |

```
ATOM   6154  NE2 GLN A 421      21.188  47.131  23.666  1.00  0.00           N
ATOM   6155 HE21 GLN A 421      20.373  47.676  23.865  1.00  0.00           H
ATOM   6156 HE22 GLN A 421      21.631  46.610  24.395  1.00  0.00           H
ATOM   6157  C   GLN A 421      21.145  47.219  18.464  1.00  0.00           C
ATOM   6158  O   GLN A 421      20.403  48.061  17.932  1.00  0.00           O
ATOM   6159  N   GLY A 422      20.885  45.927  18.381  1.00  0.00           N
ATOM   6160  H   GLY A 422      21.488  45.292  18.864  1.00  0.00           H
ATOM   6161  CA  GLY A 422      19.772  45.377  17.628  1.00  0.00           C
ATOM   6162  HA1 GLY A 422      18.964  45.815  18.021  1.00  0.00           H
ATOM   6163  HA2 GLY A 422      19.770  44.404  17.860  1.00  0.00           H
ATOM   6164  C   GLY A 422      19.692  45.509  16.078  1.00  0.00           C
ATOM   6165  O   GLY A 422      18.721  44.968  15.478  1.00  0.00           O
ATOM   6166  N   LYS A 423      20.758  46.089  15.461  1.00  0.00           N
ATOM   6167  H   LYS A 423      21.381  46.623  16.033  1.00  0.00           H
ATOM   6168  CA  LYS A 423      21.075  45.999  14.021  1.00  0.00           C
ATOM   6169  HA  LYS A 423      20.197  45.999  13.542  1.00  0.00           H
ATOM   6170  CB  LYS A 423      21.844  47.210  13.548  1.00  0.00           C
ATOM   6171  HB1 LYS A 423      22.108  47.073  12.593  1.00  0.00           H
ATOM   6172  HB2 LYS A 423      22.665  47.310  14.109  1.00  0.00           H
ATOM   6173  CG  LYS A 423      20.972  48.477  13.670  1.00  0.00           C
ATOM   6174  HG1 LYS A 423      20.713  48.609  14.627  1.00  0.00           H
ATOM   6175  HG2 LYS A 423      20.148  48.366  13.114  1.00  0.00           H
ATOM   6176  CD  LYS A 423      21.761  49.713  13.177  1.00  0.00           C
ATOM   6177  HD1 LYS A 423      21.968  49.620  12.203  1.00  0.00           H
ATOM   6178  HD2 LYS A 423      22.614  49.802  13.692  1.00  0.00           H
ATOM   6179  CE  LYS A 423      20.860  50.955  13.416  1.00  0.00           C
ATOM   6180  HE1 LYS A 423      19.932  50.752  13.103  1.00  0.00           H
ATOM   6181  HE2 LYS A 423      21.225  51.729  12.898  1.00  0.00           H
ATOM   6182  NZ  LYS A 423      20.764  51.381  14.843  1.00  0.00           N
ATOM   6183  HZ1 LYS A 423      20.169  52.182  14.912  1.00  0.00           H
ATOM   6184  HZ2 LYS A 423      20.383  50.635  15.389  1.00  0.00           H
ATOM   6185  HZ3 LYS A 423      21.675  51.612  15.184  1.00  0.00           H
ATOM   6186  C   LYS A 423      21.771  44.698  13.680  1.00  0.00           C
ATOM   6187  O   LYS A 423      22.464  44.054  14.464  1.00  0.00           O
ATOM   6188  N   ALA A 424      21.525  44.338  12.389  1.00  0.00           N
ATOM   6189  H   ALA A 424      20.728  44.761  11.958  1.00  0.00           H
ATOM   6190  CA  ALA A 424      22.298  43.394  11.547  1.00  0.00           C
ATOM   6191  HA  ALA A 424      22.476  42.548  12.051  1.00  0.00           H
ATOM   6192  CB  ALA A 424      21.466  43.017  10.276  1.00  0.00           C
ATOM   6193  HB1 ALA A 424      21.991  42.380   9.711  1.00  0.00           H
ATOM   6194  HB2 ALA A 424      20.608  42.586  10.555  1.00  0.00           H
ATOM   6195  HB3 ALA A 424      21.268  43.844   9.750  1.00  0.00           H
ATOM   6196  C   ALA A 424      23.639  44.083  11.228  1.00  0.00           C
ATOM   6197  O   ALA A 424      23.742  45.274  11.051  1.00  0.00           O
ATOM   6198  N   ASP A 425      24.682  43.260  11.146  1.00  0.00           N
ATOM   6199  H   ASP A 425      24.531  42.286  11.312  1.00  0.00           H
ATOM   6200  CA  ASP A 425      26.035  43.706  10.826  1.00  0.00           C
ATOM   6201  HA  ASP A 425      26.232  44.457  11.456  1.00  0.00           H
ATOM   6202  CB  ASP A 425      27.031  42.561  11.056  1.00  0.00           C
ATOM   6203  HB1 ASP A 425      27.936  42.868  10.760  1.00  0.00           H
ATOM   6204  HB2 ASP A 425      26.746  41.781  10.500  1.00  0.00           H
ATOM   6205  CG  ASP A 425      27.135  42.098  12.513  1.00  0.00           C
ATOM   6206  OD1 ASP A 425      26.738  42.809  13.466  1.00  0.00           O
ATOM   6207  OD2 ASP A 425      27.645  40.980  12.738  1.00  0.00           O
```

| | | | | | | | | | | |
|---|---|---|---|---|---|---|---|---|---|---|
| ATOM | 6208 | C | ASP | A | 425 | 26.112 | 44.230 | 9.382 | 1.00 | 0.00 | C |
| ATOM | 6209 | O | ASP | A | 425 | 25.401 | 43.760 | 8.490 | 1.00 | 0.00 | O |
| ATOM | 6210 | N | GLY | A | 426 | 26.967 | 45.224 | 9.151 | 1.00 | 0.00 | N |
| ATOM | 6211 | H | GLY | A | 426 | 27.551 | 45.533 | 9.902 | 1.00 | 0.00 | H |
| ATOM | 6212 | CA | GLY | A | 426 | 27.094 | 45.884 | 7.858 | 1.00 | 0.00 | C |
| ATOM | 6213 | HA1 | GLY | A | 426 | 26.183 | 45.912 | 7.446 | 1.00 | 0.00 | H |
| ATOM | 6214 | HA2 | GLY | A | 426 | 27.699 | 45.322 | 7.294 | 1.00 | 0.00 | H |
| ATOM | 6215 | C | GLY | A | 426 | 27.654 | 47.300 | 7.947 | 1.00 | 0.00 | C |
| ATOM | 6216 | O | GLY | A | 426 | 27.890 | 47.847 | 9.027 | 1.00 | 0.00 | O |
| ATOM | 6217 | N | LEU | A | 427 | 27.844 | 47.906 | 6.775 | 1.00 | 0.00 | N |
| ATOM | 6218 | H | LEU | A | 427 | 27.765 | 47.358 | 5.942 | 1.00 | 0.00 | H |
| ATOM | 6219 | CA | LEU | A | 427 | 28.161 | 49.325 | 6.640 | 1.00 | 0.00 | C |
| ATOM | 6220 | HA | LEU | A | 427 | 28.834 | 49.526 | 7.352 | 1.00 | 0.00 | H |
| ATOM | 6221 | CB | LEU | A | 427 | 28.765 | 49.596 | 5.250 | 1.00 | 0.00 | C |
| ATOM | 6222 | HB1 | LEU | A | 427 | 28.899 | 50.583 | 5.159 | 1.00 | 0.00 | H |
| ATOM | 6223 | HB2 | LEU | A | 427 | 28.110 | 49.283 | 4.563 | 1.00 | 0.00 | H |
| ATOM | 6224 | CG | LEU | A | 427 | 30.113 | 48.906 | 4.970 | 1.00 | 0.00 | C |
| ATOM | 6225 | HG | LEU | A | 427 | 29.985 | 47.923 | 5.100 | 1.00 | 0.00 | H |
| ATOM | 6226 | CD1 | LEU | A | 427 | 30.560 | 49.208 | 3.541 | 1.00 | 0.00 | C |
| ATOM | 6227 | HD11 | LEU | A | 427 | 31.435 | 48.758 | 3.363 | 1.00 | 0.00 | H |
| ATOM | 6228 | HD12 | LEU | A | 427 | 29.874 | 48.868 | 2.898 | 1.00 | 0.00 | H |
| ATOM | 6229 | HD13 | LEU | A | 427 | 30.663 | 50.196 | 3.426 | 1.00 | 0.00 | H |
| ATOM | 6230 | CD2 | LEU | A | 427 | 31.205 | 49.385 | 5.925 | 1.00 | 0.00 | C |
| ATOM | 6231 | HD21 | LEU | A | 427 | 32.062 | 48.916 | 5.713 | 1.00 | 0.00 | H |
| ATOM | 6232 | HD22 | LEU | A | 427 | 31.330 | 50.372 | 5.821 | 1.00 | 0.00 | H |
| ATOM | 6233 | HD23 | LEU | A | 427 | 30.937 | 49.181 | 6.867 | 1.00 | 0.00 | H |
| ATOM | 6234 | C | LEU | A | 427 | 26.911 | 50.185 | 6.876 | 1.00 | 0.00 | C |
| ATOM | 6235 | O | LEU | A | 427 | 25.813 | 49.849 | 6.409 | 1.00 | 0.00 | O |
| ATOM | 6236 | N | TYR | A | 428 | 27.113 | 51.340 | 7.510 | 1.00 | 0.00 | N |
| ATOM | 6237 | H | TYR | A | 428 | 28.037 | 51.524 | 7.845 | 1.00 | 0.00 | H |
| ATOM | 6238 | CA | TYR | A | 428 | 26.094 | 52.361 | 7.757 | 1.00 | 0.00 | C |
| ATOM | 6239 | HA | TYR | A | 428 | 25.407 | 52.263 | 7.037 | 1.00 | 0.00 | H |
| ATOM | 6240 | CB | TYR | A | 428 | 25.456 | 52.121 | 9.137 | 1.00 | 0.00 | C |
| ATOM | 6241 | HB1 | TYR | A | 428 | 24.788 | 52.848 | 9.295 | 1.00 | 0.00 | H |
| ATOM | 6242 | HB2 | TYR | A | 428 | 26.182 | 52.180 | 9.823 | 1.00 | 0.00 | H |
| ATOM | 6243 | CG | TYR | A | 428 | 24.739 | 50.803 | 9.362 | 1.00 | 0.00 | C |
| ATOM | 6244 | CD1 | TYR | A | 428 | 25.279 | 49.795 | 10.183 | 1.00 | 0.00 | C |
| ATOM | 6245 | HD1 | TYR | A | 428 | 26.178 | 49.916 | 10.605 | 1.00 | 0.00 | H |
| ATOM | 6246 | CE1 | TYR | A | 428 | 24.541 | 48.613 | 10.405 | 1.00 | 0.00 | C |
| ATOM | 6247 | HE1 | TYR | A | 428 | 24.915 | 47.891 | 10.987 | 1.00 | 0.00 | H |
| ATOM | 6248 | CZ | TYR | A | 428 | 23.274 | 48.450 | 9.805 | 1.00 | 0.00 | C |
| ATOM | 6249 | OH | TYR | A | 428 | 22.628 | 47.202 | 9.854 | 1.00 | 0.00 | O |
| ATOM | 6250 | HH | TYR | A | 428 | 21.748 | 47.264 | 9.384 | 1.00 | 0.00 | H |
| ATOM | 6251 | CE2 | TYR | A | 428 | 22.778 | 49.440 | 8.980 | 1.00 | 0.00 | C |
| ATOM | 6252 | HE2 | TYR | A | 428 | 21.896 | 49.310 | 8.526 | 1.00 | 0.00 | H |
| ATOM | 6253 | CD2 | TYR | A | 428 | 23.484 | 50.603 | 8.776 | 1.00 | 0.00 | C |
| ATOM | 6254 | HD2 | TYR | A | 428 | 23.093 | 51.319 | 8.198 | 1.00 | 0.00 | H |
| ATOM | 6255 | C | TYR | A | 428 | 26.741 | 53.769 | 7.701 | 1.00 | 0.00 | C |
| ATOM | 6256 | O | TYR | A | 428 | 27.913 | 53.861 | 8.031 | 1.00 | 0.00 | O |
| ATOM | 6257 | N | PRO | A | 429 | 25.997 | 54.823 | 7.294 | 1.00 | 0.00 | N |
| ATOM | 6258 | CD | PRO | A | 429 | 24.669 | 54.725 | 6.663 | 1.00 | 0.00 | C |
| ATOM | 6259 | HD1 | PRO | A | 429 | 23.967 | 54.554 | 7.354 | 1.00 | 0.00 | H |
| ATOM | 6260 | HD2 | PRO | A | 429 | 24.658 | 53.988 | 5.987 | 1.00 | 0.00 | H |
| ATOM | 6261 | CG | PRO | A | 429 | 24.501 | 56.104 | 6.008 | 1.00 | 0.00 | C |

| | | | | | | | | | | | |
|---|---|---|---|---|---|---|---|---|---|---|---|
| ATOM | 6262 | HG1 | PRO | A | 429 | 23.549 | 56.404 | 6.072 | 1.00 | 0.00 | H |
| ATOM | 6263 | HG2 | PRO | A | 429 | 24.775 | 56.066 | 5.047 | 1.00 | 0.00 | H |
| ATOM | 6264 | CB | PRO | A | 429 | 25.413 | 57.066 | 6.782 | 1.00 | 0.00 | C |
| ATOM | 6265 | HB1 | PRO | A | 429 | 24.894 | 57.534 | 7.497 | 1.00 | 0.00 | H |
| ATOM | 6266 | HB2 | PRO | A | 429 | 25.809 | 57.742 | 6.161 | 1.00 | 0.00 | H |
| ATOM | 6267 | CA | PRO | A | 429 | 26.521 | 56.204 | 7.407 | 1.00 | 0.00 | C |
| ATOM | 6268 | HA | PRO | A | 429 | 27.418 | 56.255 | 6.968 | 1.00 | 0.00 | H |
| ATOM | 6269 | C | PRO | A | 429 | 26.721 | 56.642 | 8.845 | 1.00 | 0.00 | C |
| ATOM | 6270 | O | PRO | A | 429 | 26.105 | 56.104 | 9.756 | 1.00 | 0.00 | O |
| ATOM | 6271 | N | ASN | A | 430 | 27.605 | 57.648 | 9.007 | 1.00 | 0.00 | N |
| ATOM | 6272 | H | ASN | A | 430 | 28.060 | 58.059 | 8.217 | 1.00 | 0.00 | H |
| ATOM | 6273 | CA | ASN | A | 430 | 27.890 | 58.135 | 10.385 | 1.00 | 0.00 | C |
| ATOM | 6274 | HA | ASN | A | 430 | 27.739 | 57.386 | 11.030 | 1.00 | 0.00 | H |
| ATOM | 6275 | CB | ASN | A | 430 | 29.368 | 58.646 | 10.459 | 1.00 | 0.00 | C |
| ATOM | 6276 | HB1 | ASN | A | 430 | 29.517 | 59.289 | 9.708 | 1.00 | 0.00 | H |
| ATOM | 6277 | HB2 | ASN | A | 430 | 29.979 | 57.861 | 10.353 | 1.00 | 0.00 | H |
| ATOM | 6278 | CG | ASN | A | 430 | 29.824 | 59.348 | 11.686 | 1.00 | 0.00 | C |
| ATOM | 6279 | OD1 | ASN | A | 430 | 29.139 | 59.432 | 12.670 | 1.00 | 0.00 | O |
| ATOM | 6280 | ND2 | ASN | A | 430 | 31.023 | 59.887 | 11.759 | 1.00 | 0.00 | N |
| ATOM | 6281 | HD21 | ASN | A | 430 | 31.646 | 59.828 | 10.979 | 1.00 | 0.00 | H |
| ATOM | 6282 | HD22 | ASN | A | 430 | 31.311 | 60.355 | 12.594 | 1.00 | 0.00 | H |
| ATOM | 6283 | C | ASN | A | 430 | 26.936 | 59.293 | 10.670 | 1.00 | 0.00 | C |
| ATOM | 6284 | O | ASN | A | 430 | 26.987 | 60.271 | 9.961 | 1.00 | 0.00 | O |
| ATOM | 6285 | N | PRO | A | 431 | 26.138 | 59.363 | 11.726 | 1.00 | 0.00 | N |
| ATOM | 6286 | CD | PRO | A | 431 | 25.783 | 58.353 | 12.744 | 1.00 | 0.00 | C |
| ATOM | 6287 | HD1 | PRO | A | 431 | 26.256 | 58.534 | 13.606 | 1.00 | 0.00 | H |
| ATOM | 6288 | HD2 | PRO | A | 431 | 26.009 | 57.432 | 12.427 | 1.00 | 0.00 | H |
| ATOM | 6289 | CG | PRO | A | 431 | 24.266 | 58.573 | 12.856 | 1.00 | 0.00 | C |
| ATOM | 6290 | HG1 | PRO | A | 431 | 23.937 | 58.299 | 13.759 | 1.00 | 0.00 | H |
| ATOM | 6291 | HG2 | PRO | A | 431 | 23.782 | 58.052 | 12.153 | 1.00 | 0.00 | H |
| ATOM | 6292 | CB | PRO | A | 431 | 24.058 | 60.073 | 12.646 | 1.00 | 0.00 | C |
| ATOM | 6293 | HB1 | PRO | A | 431 | 24.093 | 60.555 | 13.522 | 1.00 | 0.00 | H |
| ATOM | 6294 | HB2 | PRO | A | 431 | 23.176 | 60.244 | 12.207 | 1.00 | 0.00 | H |
| ATOM | 6295 | CA | PRO | A | 431 | 25.219 | 60.518 | 11.731 | 1.00 | 0.00 | C |
| ATOM | 6296 | HA | PRO | A | 431 | 24.922 | 60.704 | 10.794 | 1.00 | 0.00 | H |
| ATOM | 6297 | C | PRO | A | 431 | 25.852 | 61.873 | 12.054 | 1.00 | 0.00 | C |
| ATOM | 6298 | O | PRO | A | 431 | 25.264 | 62.967 | 11.909 | 1.00 | 0.00 | O |
| ATOM | 6299 | N | ARG | A | 432 | 27.171 | 61.917 | 12.393 | 1.00 | 0.00 | N |
| ATOM | 6300 | H | ARG | A | 432 | 27.668 | 61.049 | 12.398 | 1.00 | 0.00 | H |
| ATOM | 6301 | CA | ARG | A | 432 | 27.944 | 63.145 | 12.759 | 1.00 | 0.00 | C |
| ATOM | 6302 | HA | ARG | A | 432 | 27.314 | 63.713 | 13.289 | 1.00 | 0.00 | H |
| ATOM | 6303 | CB | ARG | A | 432 | 29.170 | 62.640 | 13.488 | 1.00 | 0.00 | C |
| ATOM | 6304 | HB1 | ARG | A | 432 | 29.728 | 62.147 | 12.820 | 1.00 | 0.00 | H |
| ATOM | 6305 | HB2 | ARG | A | 432 | 28.853 | 62.004 | 14.192 | 1.00 | 0.00 | H |
| ATOM | 6306 | CG | ARG | A | 432 | 30.067 | 63.672 | 14.168 | 1.00 | 0.00 | C |
| ATOM | 6307 | HG1 | ARG | A | 432 | 30.436 | 64.256 | 13.445 | 1.00 | 0.00 | H |
| ATOM | 6308 | HG2 | ARG | A | 432 | 30.813 | 63.166 | 14.601 | 1.00 | 0.00 | H |
| ATOM | 6309 | CD | ARG | A | 432 | 29.380 | 64.593 | 15.260 | 1.00 | 0.00 | C |
| ATOM | 6310 | HD1 | ARG | A | 432 | 28.670 | 65.141 | 14.819 | 1.00 | 0.00 | H |
| ATOM | 6311 | HD2 | ARG | A | 432 | 30.072 | 65.198 | 15.653 | 1.00 | 0.00 | H |
| ATOM | 6312 | NE | ARG | A | 432 | 28.718 | 63.928 | 16.418 | 1.00 | 0.00 | N |
| ATOM | 6313 | HE | ARG | A | 432 | 27.812 | 63.534 | 16.265 | 1.00 | 0.00 | H |
| ATOM | 6314 | CZ | ARG | A | 432 | 29.223 | 63.819 | 17.608 | 1.00 | 0.00 | C |
| ATOM | 6315 | NH1 | ARG | A | 432 | 30.324 | 64.412 | 18.001 | 1.00 | 0.00 | N |

```
ATOM   6316 HH11 ARG A 432      30.832  64.994  17.366  1.00  0.00           H
ATOM   6317 HH12 ARG A 432      30.655  64.283  18.936  1.00  0.00           H
ATOM   6318  NH2 ARG A 432      28.538  63.330  18.625  1.00  0.00           N
ATOM   6319 HH21 ARG A 432      27.594  63.027  18.493  1.00  0.00           H
ATOM   6320 HH22 ARG A 432      28.963  63.262  19.528  1.00  0.00           H
ATOM   6321  C   ARG A 432      28.432  64.030  11.605  1.00  0.00           C
ATOM   6322  O   ARG A 432      28.497  65.244  11.735  1.00  0.00           O
ATOM   6323  N   GLU A 433      28.801  63.362  10.489  1.00  0.00           N
ATOM   6324  H   GLU A 433      28.455  62.437  10.333  1.00  0.00           H
ATOM   6325  CA  GLU A 433      29.698  63.969   9.506  1.00  0.00           C
ATOM   6326  HA  GLU A 433      29.465  64.929   9.348  1.00  0.00           H
ATOM   6327  CB  GLU A 433      31.141  63.868  10.038  1.00  0.00           C
ATOM   6328  HB1 GLU A 433      31.296  62.937  10.369  1.00  0.00           H
ATOM   6329  HB2 GLU A 433      31.250  64.513  10.794  1.00  0.00           H
ATOM   6330  CG  GLU A 433      32.193  64.190   8.960  1.00  0.00           C
ATOM   6331  HG1 GLU A 433      32.074  63.565   8.188  1.00  0.00           H
ATOM   6332  HG2 GLU A 433      33.107  64.071   9.348  1.00  0.00           H
ATOM   6333  CD  GLU A 433      32.079  65.604   8.445  1.00  0.00           C
ATOM   6334  OE1 GLU A 433      31.749  65.746   7.215  1.00  0.00           O
ATOM   6335  OE2 GLU A 433      32.333  66.546   9.275  1.00  0.00           O
ATOM   6336  C   GLU A 433      29.520  63.161   8.241  1.00  0.00           C
ATOM   6337  O   GLU A 433      29.563  61.936   8.236  1.00  0.00           O
ATOM   6338  N   ARG A 434      29.282  63.936   7.172  1.00  0.00           N
ATOM   6339  H   ARG A 434      29.386  64.928   7.238  1.00  0.00           H
ATOM   6340  CA  ARG A 434      28.868  63.321   5.903  1.00  0.00           C
ATOM   6341  HA  ARG A 434      28.160  62.676   6.191  1.00  0.00           H
ATOM   6342  CB  ARG A 434      28.288  64.359   4.921  1.00  0.00           C
ATOM   6343  HB1 ARG A 434      28.187  63.936   4.021  1.00  0.00           H
ATOM   6344  HB2 ARG A 434      28.917  65.134   4.856  1.00  0.00           H
ATOM   6345  CG  ARG A 434      26.913  64.867   5.394  1.00  0.00           C
ATOM   6346  HG1 ARG A 434      27.032  65.446   6.201  1.00  0.00           H
ATOM   6347  HG2 ARG A 434      26.332  64.087   5.626  1.00  0.00           H
ATOM   6348  CD  ARG A 434      26.246  65.683   4.275  1.00  0.00           C
ATOM   6349  HD1 ARG A 434      26.132  65.121   3.455  1.00  0.00           H
ATOM   6350  HD2 ARG A 434      26.798  66.487   4.054  1.00  0.00           H
ATOM   6351  NE  ARG A 434      24.929  66.140   4.698  1.00  0.00           N
ATOM   6352  HE  ARG A 434      24.638  65.965   5.639  1.00  0.00           H
ATOM   6353  CZ  ARG A 434      24.083  66.794   3.863  1.00  0.00           C
ATOM   6354  NH1 ARG A 434      24.417  67.099   2.586  1.00  0.00           N
ATOM   6355 HH11 ARG A 434      25.315  66.841   2.229  1.00  0.00           H
ATOM   6356 HH12 ARG A 434      23.766  67.582   2.001  1.00  0.00           H
ATOM   6357  NH2 ARG A 434      22.864  67.153   4.322  1.00  0.00           N
ATOM   6358 HH21 ARG A 434      22.600  66.936   5.262  1.00  0.00           H
ATOM   6359 HH22 ARG A 434      22.226  67.636   3.722  1.00  0.00           H
ATOM   6360  C   ARG A 434      29.960  62.511   5.239  1.00  0.00           C
ATOM   6361  O   ARG A 434      29.720  61.449   4.681  1.00  0.00           O
ATOM   6362  N   SER A 435      31.182  63.049   5.389  1.00  0.00           N
ATOM   6363  H   SER A 435      31.297  63.910   5.885  1.00  0.00           H
ATOM   6364  CA  SER A 435      32.348  62.359   4.817  1.00  0.00           C
ATOM   6365  HA  SER A 435      31.977  61.823   4.059  1.00  0.00           H
ATOM   6366  CB  SER A 435      33.361  63.422   4.361  1.00  0.00           C
ATOM   6367  HB1 SER A 435      32.896  64.170   3.887  1.00  0.00           H
ATOM   6368  HB2 SER A 435      34.046  63.018   3.755  1.00  0.00           H
ATOM   6369  OG  SER A 435      34.052  63.982   5.487  1.00  0.00           O
```

| | | | | | | | | | | | |
|---|---|---|---|---|---|---|---|---|---|---|---|
| ATOM | 6370 | HG | SER | A | 435 | 34.705 | 64.670 | 5.169 | 1.00 | 0.00 | H |
| ATOM | 6371 | C | SER | A | 435 | 33.001 | 61.328 | 5.721 | 1.00 | 0.00 | C |
| ATOM | 6372 | O | SER | A | 435 | 34.209 | 61.132 | 5.739 | 1.00 | 0.00 | O |
| ATOM | 6373 | N | SER | A | 436 | 32.116 | 60.684 | 6.496 | 1.00 | 0.00 | N |
| ATOM | 6374 | H | SER | A | 436 | 31.151 | 60.946 | 6.473 | 1.00 | 0.00 | H |
| ATOM | 6375 | CA | SER | A | 436 | 32.558 | 59.597 | 7.377 | 1.00 | 0.00 | C |
| ATOM | 6376 | HA | SER | A | 436 | 33.349 | 59.131 | 6.981 | 1.00 | 0.00 | H |
| ATOM | 6377 | CB | SER | A | 436 | 32.967 | 60.230 | 8.730 | 1.00 | 0.00 | C |
| ATOM | 6378 | HB1 | SER | A | 436 | 32.305 | 60.911 | 9.043 | 1.00 | 0.00 | H |
| ATOM | 6379 | HB2 | SER | A | 436 | 33.877 | 60.643 | 8.693 | 1.00 | 0.00 | H |
| ATOM | 6380 | OG | SER | A | 436 | 33.017 | 59.278 | 9.684 | 1.00 | 0.00 | O |
| ATOM | 6381 | HG | SER | A | 436 | 33.281 | 59.692 | 10.555 | 1.00 | 0.00 | H |
| ATOM | 6382 | C | SER | A | 436 | 31.471 | 58.536 | 7.466 | 1.00 | 0.00 | C |
| ATOM | 6383 | O | SER | A | 436 | 30.272 | 58.755 | 7.179 | 1.00 | 0.00 | O |
| ATOM | 6384 | N | PHE | A | 437 | 31.931 | 57.333 | 7.845 | 1.00 | 0.00 | N |
| ATOM | 6385 | H | PHE | A | 437 | 32.891 | 57.212 | 8.098 | 1.00 | 0.00 | H |
| ATOM | 6386 | CA | PHE | A | 437 | 30.999 | 56.197 | 7.882 | 1.00 | 0.00 | C |
| ATOM | 6387 | HA | PHE | A | 437 | 30.122 | 56.610 | 8.127 | 1.00 | 0.00 | H |
| ATOM | 6388 | CB | PHE | A | 437 | 30.903 | 55.532 | 6.502 | 1.00 | 0.00 | C |
| ATOM | 6389 | HB1 | PHE | A | 437 | 30.735 | 56.242 | 5.818 | 1.00 | 0.00 | H |
| ATOM | 6390 | HB2 | PHE | A | 437 | 30.136 | 54.890 | 6.511 | 1.00 | 0.00 | H |
| ATOM | 6391 | CG | PHE | A | 437 | 32.147 | 54.769 | 6.089 | 1.00 | 0.00 | C |
| ATOM | 6392 | CD1 | PHE | A | 437 | 33.266 | 55.452 | 5.574 | 1.00 | 0.00 | C |
| ATOM | 6393 | HD1 | PHE | A | 437 | 33.239 | 56.446 | 5.469 | 1.00 | 0.00 | H |
| ATOM | 6394 | CE1 | PHE | A | 437 | 34.421 | 54.739 | 5.208 | 1.00 | 0.00 | C |
| ATOM | 6395 | HE1 | PHE | A | 437 | 35.216 | 55.227 | 4.848 | 1.00 | 0.00 | H |
| ATOM | 6396 | CZ | PHE | A | 437 | 34.459 | 53.341 | 5.347 | 1.00 | 0.00 | C |
| ATOM | 6397 | HZ | PHE | A | 437 | 35.277 | 52.832 | 5.078 | 1.00 | 0.00 | H |
| ATOM | 6398 | CE2 | PHE | A | 437 | 33.343 | 52.660 | 5.865 | 1.00 | 0.00 | C |
| ATOM | 6399 | HE2 | PHE | A | 437 | 33.370 | 51.666 | 5.968 | 1.00 | 0.00 | H |
| ATOM | 6400 | CD2 | PHE | A | 437 | 32.191 | 53.372 | 6.238 | 1.00 | 0.00 | C |
| ATOM | 6401 | HD2 | PHE | A | 437 | 31.401 | 52.884 | 6.610 | 1.00 | 0.00 | H |
| ATOM | 6402 | C | PHE | A | 437 | 31.271 | 55.205 | 8.989 | 1.00 | 0.00 | C |
| ATOM | 6403 | O | PHE | A | 437 | 32.397 | 55.026 | 9.465 | 1.00 | 0.00 | O |
| ATOM | 6404 | N | TYR | A | 438 | 30.164 | 54.624 | 9.439 | 1.00 | 0.00 | N |
| ATOM | 6405 | H | TYR | A | 438 | 29.289 | 54.866 | 9.020 | 1.00 | 0.00 | H |
| ATOM | 6406 | CA | TYR | A | 438 | 30.179 | 53.637 | 10.532 | 1.00 | 0.00 | C |
| ATOM | 6407 | HA | TYR | A | 438 | 30.959 | 53.826 | 11.128 | 1.00 | 0.00 | H |
| ATOM | 6408 | CB | TYR | A | 438 | 28.869 | 53.756 | 11.318 | 1.00 | 0.00 | C |
| ATOM | 6409 | HB1 | TYR | A | 438 | 28.716 | 52.872 | 11.760 | 1.00 | 0.00 | H |
| ATOM | 6410 | HB2 | TYR | A | 438 | 28.145 | 53.924 | 10.649 | 1.00 | 0.00 | H |
| ATOM | 6411 | CG | TYR | A | 438 | 28.721 | 54.825 | 12.406 | 1.00 | 0.00 | C |
| ATOM | 6412 | CD1 | TYR | A | 438 | 27.451 | 55.012 | 12.940 | 1.00 | 0.00 | C |
| ATOM | 6413 | HD1 | TYR | A | 438 | 26.663 | 54.557 | 12.525 | 1.00 | 0.00 | H |
| ATOM | 6414 | CE1 | TYR | A | 438 | 27.268 | 55.861 | 14.091 | 1.00 | 0.00 | C |
| ATOM | 6415 | HE1 | TYR | A | 438 | 26.371 | 55.933 | 14.526 | 1.00 | 0.00 | H |
| ATOM | 6416 | CZ | TYR | A | 438 | 28.391 | 56.608 | 14.611 | 1.00 | 0.00 | C |
| ATOM | 6417 | OH | TYR | A | 438 | 28.301 | 57.404 | 15.713 | 1.00 | 0.00 | O |
| ATOM | 6418 | HH | TYR | A | 438 | 29.190 | 57.823 | 15.895 | 1.00 | 0.00 | H |
| ATOM | 6419 | CE2 | TYR | A | 438 | 29.584 | 56.437 | 13.944 | 1.00 | 0.00 | C |
| ATOM | 6420 | HE2 | TYR | A | 438 | 30.376 | 56.969 | 14.244 | 1.00 | 0.00 | H |
| ATOM | 6421 | CD2 | TYR | A | 438 | 29.745 | 55.556 | 12.866 | 1.00 | 0.00 | C |
| ATOM | 6422 | HD2 | TYR | A | 438 | 30.642 | 55.471 | 12.432 | 1.00 | 0.00 | H |
| ATOM | 6423 | C | TYR | A | 438 | 30.271 | 52.232 | 9.897 | 1.00 | 0.00 | C |

| | | | | | | | | | | | |
|---|---|---|---|---|---|---|---|---|---|---|---|
| ATOM | 6424 | O | TYR | A | 438 | 29.930 | 51.989 | 8.735 | 1.00 | 0.00 | O |
| ATOM | 6425 | N | SER | A | 439 | 30.654 | 51.255 | 10.700 | 1.00 | 0.00 | N |
| ATOM | 6426 | H | SER | A | 439 | 30.915 | 51.498 | 11.634 | 1.00 | 0.00 | H |
| ATOM | 6427 | CA | SER | A | 439 | 30.724 | 49.851 | 10.323 | 1.00 | 0.00 | C |
| ATOM | 6428 | HA | SER | A | 439 | 30.070 | 49.620 | 9.603 | 1.00 | 0.00 | H |
| ATOM | 6429 | CB | SER | A | 439 | 32.143 | 49.606 | 9.793 | 1.00 | 0.00 | C |
| ATOM | 6430 | HB1 | SER | A | 439 | 32.816 | 49.762 | 10.516 | 1.00 | 0.00 | H |
| ATOM | 6431 | HB2 | SER | A | 439 | 32.335 | 50.213 | 9.022 | 1.00 | 0.00 | H |
| ATOM | 6432 | OG | SER | A | 439 | 32.347 | 48.289 | 9.330 | 1.00 | 0.00 | O |
| ATOM | 6433 | HG | SER | A | 439 | 33.286 | 48.194 | 8.999 | 1.00 | 0.00 | H |
| ATOM | 6434 | C | SER | A | 439 | 30.366 | 49.042 | 11.567 | 1.00 | 0.00 | C |
| ATOM | 6435 | O | SER | A | 439 | 30.984 | 49.225 | 12.610 | 1.00 | 0.00 | O |
| ATOM | 6436 | N | CYS | A | 440 | 29.300 | 48.246 | 11.517 | 1.00 | 0.00 | N |
| ATOM | 6437 | H | CYS | A | 440 | 28.765 | 48.226 | 10.672 | 1.00 | 0.00 | H |
| ATOM | 6438 | CA | CYS | A | 440 | 28.866 | 47.397 | 12.630 | 1.00 | 0.00 | C |
| ATOM | 6439 | HA | CYS | A | 440 | 29.339 | 47.719 | 13.450 | 1.00 | 0.00 | H |
| ATOM | 6440 | CB | CYS | A | 440 | 27.348 | 47.519 | 12.833 | 1.00 | 0.00 | C |
| ATOM | 6441 | HB1 | CYS | A | 440 | 27.149 | 46.790 | 13.488 | 1.00 | 0.00 | H |
| ATOM | 6442 | HB2 | CYS | A | 440 | 26.980 | 47.271 | 11.937 | 1.00 | 0.00 | H |
| ATOM | 6443 | SG | CYS | A | 440 | 26.688 | 49.118 | 13.401 | 1.00 | 0.00 | S |
| ATOM | 6444 | C | CYS | A | 440 | 29.245 | 45.946 | 12.319 | 1.00 | 0.00 | C |
| ATOM | 6445 | O | CYS | A | 440 | 28.850 | 45.439 | 11.270 | 1.00 | 0.00 | O |
| ATOM | 6446 | N | ALA | A | 441 | 29.955 | 45.258 | 13.217 | 1.00 | 0.00 | N |
| ATOM | 6447 | H | ALA | A | 441 | 30.320 | 45.740 | 14.013 | 1.00 | 0.00 | H |
| ATOM | 6448 | CA | ALA | A | 441 | 30.219 | 43.822 | 13.079 | 1.00 | 0.00 | C |
| ATOM | 6449 | HA | ALA | A | 441 | 29.438 | 43.452 | 12.576 | 1.00 | 0.00 | H |
| ATOM | 6450 | CB | ALA | A | 441 | 31.491 | 43.592 | 12.261 | 1.00 | 0.00 | C |
| ATOM | 6451 | HB1 | ALA | A | 441 | 31.659 | 42.610 | 12.175 | 1.00 | 0.00 | H |
| ATOM | 6452 | HB2 | ALA | A | 441 | 31.379 | 43.992 | 11.351 | 1.00 | 0.00 | H |
| ATOM | 6453 | HB3 | ALA | A | 441 | 32.266 | 44.024 | 12.722 | 1.00 | 0.00 | H |
| ATOM | 6454 | C | ALA | A | 441 | 30.272 | 43.118 | 14.442 | 1.00 | 0.00 | C |
| ATOM | 6455 | O | ALA | A | 441 | 30.858 | 43.636 | 15.395 | 1.00 | 0.00 | O |
| ATOM | 6456 | N | ALA | A | 442 | 29.584 | 41.977 | 14.542 | 1.00 | 0.00 | N |
| ATOM | 6457 | H | ALA | A | 442 | 29.342 | 41.502 | 13.696 | 1.00 | 0.00 | H |
| ATOM | 6458 | CA | ALA | A | 442 | 29.162 | 41.377 | 15.803 | 1.00 | 0.00 | C |
| ATOM | 6459 | HA | ALA | A | 442 | 28.446 | 40.717 | 15.577 | 1.00 | 0.00 | H |
| ATOM | 6460 | CB | ALA | A | 442 | 30.324 | 40.586 | 16.421 | 1.00 | 0.00 | C |
| ATOM | 6461 | HB1 | ALA | A | 442 | 30.027 | 40.178 | 17.284 | 1.00 | 0.00 | H |
| ATOM | 6462 | HB2 | ALA | A | 442 | 30.608 | 39.863 | 15.791 | 1.00 | 0.00 | H |
| ATOM | 6463 | HB3 | ALA | A | 442 | 31.094 | 41.202 | 16.590 | 1.00 | 0.00 | H |
| ATOM | 6464 | C | ALA | A | 442 | 28.489 | 42.408 | 16.734 | 1.00 | 0.00 | C |
| ATOM | 6465 | O | ALA | A | 442 | 28.728 | 42.440 | 17.917 | 1.00 | 0.00 | O |
| ATOM | 6466 | N | GLY | A | 443 | 27.735 | 43.374 | 16.224 | 1.00 | 0.00 | N |
| ATOM | 6467 | H | GLY | A | 443 | 27.597 | 43.385 | 15.234 | 1.00 | 0.00 | H |
| ATOM | 6468 | CA | GLY | A | 443 | 27.089 | 44.428 | 17.018 | 1.00 | 0.00 | C |
| ATOM | 6469 | HA1 | GLY | A | 443 | 26.564 | 43.963 | 17.731 | 1.00 | 0.00 | H |
| ATOM | 6470 | HA2 | GLY | A | 443 | 26.463 | 44.904 | 16.400 | 1.00 | 0.00 | H |
| ATOM | 6471 | C | GLY | A | 443 | 28.038 | 45.467 | 17.680 | 1.00 | 0.00 | C |
| ATOM | 6472 | O | GLY | A | 443 | 27.574 | 46.364 | 18.377 | 1.00 | 0.00 | O |
| ATOM | 6473 | N | ARG | A | 444 | 29.365 | 45.405 | 17.466 | 1.00 | 0.00 | N |
| ATOM | 6474 | H | ARG | A | 444 | 29.740 | 44.583 | 17.037 | 1.00 | 0.00 | H |
| ATOM | 6475 | CA | ARG | A | 444 | 30.297 | 46.497 | 17.836 | 1.00 | 0.00 | C |
| ATOM | 6476 | HA | ARG | A | 444 | 29.952 | 47.028 | 18.610 | 1.00 | 0.00 | H |
| ATOM | 6477 | CB | ARG | A | 444 | 31.592 | 45.799 | 18.230 | 1.00 | 0.00 | C |

```
ATOM   6478  HB1 ARG A 444      31.852  45.238  17.444  1.00  0.00           H
ATOM   6479  HB2 ARG A 444      31.361  45.206  19.001  1.00  0.00           H
ATOM   6480  CG  ARG A 444      32.848  46.643  18.643  1.00  0.00           C
ATOM   6481  HG1 ARG A 444      33.349  46.185  19.377  1.00  0.00           H
ATOM   6482  HG2 ARG A 444      32.570  47.554  18.947  1.00  0.00           H
ATOM   6483  CD  ARG A 444      33.772  46.811  17.505  1.00  0.00           C
ATOM   6484  HD1 ARG A 444      33.265  47.227  16.750  1.00  0.00           H
ATOM   6485  HD2 ARG A 444      34.097  45.906  17.230  1.00  0.00           H
ATOM   6486  NE  ARG A 444      34.952  47.667  17.815  1.00  0.00           N
ATOM   6487  HE  ARG A 444      34.782  48.565  18.221  1.00  0.00           H
ATOM   6488  CZ  ARG A 444      36.180  47.329  17.596  1.00  0.00           C
ATOM   6489  NH1 ARG A 444      36.618  46.112  17.483  1.00  0.00           N
ATOM   6490 HH11 ARG A 444      35.984  45.343  17.566  1.00  0.00           H
ATOM   6491 HH12 ARG A 444      37.589  45.944  17.313  1.00  0.00           H
ATOM   6492  NH2 ARG A 444      37.012  48.295  17.438  1.00  0.00           N
ATOM   6493 HH21 ARG A 444      36.694  49.242  17.488  1.00  0.00           H
ATOM   6494 HH22 ARG A 444      37.977  48.100  17.265  1.00  0.00           H
ATOM   6495  C   ARG A 444      30.427  47.527  16.705  1.00  0.00           C
ATOM   6496  O   ARG A 444      30.662  47.191  15.542  1.00  0.00           O
ATOM   6497  N   LEU A 445      30.327  48.799  17.111  1.00  0.00           N
ATOM   6498  H   LEU A 445      30.095  48.935  18.074  1.00  0.00           H
ATOM   6499  CA  LEU A 445      30.514  50.005  16.325  1.00  0.00           C
ATOM   6500  HA  LEU A 445      30.068  49.789  15.457  1.00  0.00           H
ATOM   6501  CB  LEU A 445      29.773  51.201  16.938  1.00  0.00           C
ATOM   6502  HB1 LEU A 445      30.139  51.351  17.856  1.00  0.00           H
ATOM   6503  HB2 LEU A 445      28.804  50.963  17.000  1.00  0.00           H
ATOM   6504  CG  LEU A 445      29.907  52.524  16.129  1.00  0.00           C
ATOM   6505  HG  LEU A 445      30.857  52.778  15.946  1.00  0.00           H
ATOM   6506  CD1 LEU A 445      29.152  52.344  14.849  1.00  0.00           C
ATOM   6507 HD11 LEU A 445      29.218  53.179  14.303  1.00  0.00           H
ATOM   6508 HD12 LEU A 445      29.542  51.579  14.337  1.00  0.00           H
ATOM   6509 HD13 LEU A 445      28.192  52.154  15.052  1.00  0.00           H
ATOM   6510  CD2 LEU A 445      29.083  53.569  16.930  1.00  0.00           C
ATOM   6511 HD21 LEU A 445      29.124  54.453  16.465  1.00  0.00           H
ATOM   6512 HD22 LEU A 445      28.131  53.268  16.990  1.00  0.00           H
ATOM   6513 HD23 LEU A 445      29.463  53.659  17.850  1.00  0.00           H
ATOM   6514  C   LEU A 445      31.979  50.275  15.971  1.00  0.00           C
ATOM   6515  O   LEU A 445      32.860  50.091  16.784  1.00  0.00           O
ATOM   6516  N   PHE A 446      32.169  50.672  14.709  1.00  0.00           N
ATOM   6517  H   PHE A 446      31.412  50.658  14.056  1.00  0.00           H
ATOM   6518  CA  PHE A 446      33.487  51.124  14.296  1.00  0.00           C
ATOM   6519  HA  PHE A 446      33.865  51.496  15.143  1.00  0.00           H
ATOM   6520  CB  PHE A 446      34.540  50.044  13.990  1.00  0.00           C
ATOM   6521  HB1 PHE A 446      34.722  49.560  14.846  1.00  0.00           H
ATOM   6522  HB2 PHE A 446      35.369  50.518  13.693  1.00  0.00           H
ATOM   6523  CG  PHE A 446      34.279  48.978  12.958  1.00  0.00           C
ATOM   6524  CD1 PHE A 446      33.407  47.921  13.260  1.00  0.00           C
ATOM   6525  HD1 PHE A 446      32.894  47.928  14.118  1.00  0.00           H
ATOM   6526  CE1 PHE A 446      33.253  46.855  12.363  1.00  0.00           C
ATOM   6527  HE1 PHE A 446      32.582  46.138  12.552  1.00  0.00           H
ATOM   6528  CZ  PHE A 446      34.036  46.791  11.203  1.00  0.00           C
ATOM   6529  HZ  PHE A 446      33.964  46.006  10.588  1.00  0.00           H
ATOM   6530  CE2 PHE A 446      34.923  47.840  10.913  1.00  0.00           C
ATOM   6531  HE2 PHE A 446      35.475  47.805  10.080  1.00  0.00           H
```

| | | | | | | | | | | | |
|---|---|---|---|---|---|---|---|---|---|---|---|
| ATOM | 6532 | CD2 | PHE | A | 446 | 35.043 | 48.937 | 11.779 | 1.00 | 0.00 | C |
| ATOM | 6533 | HD2 | PHE | A | 446 | 35.668 | 49.686 | 11.559 | 1.00 | 0.00 | H |
| ATOM | 6534 | C | PHE | A | 446 | 33.357 | 52.267 | 13.304 | 1.00 | 0.00 | C |
| ATOM | 6535 | O | PHE | A | 446 | 32.365 | 52.348 | 12.581 | 1.00 | 0.00 | O |
| ATOM | 6536 | N | GLN | A | 447 | 34.274 | 53.230 | 13.368 | 1.00 | 0.00 | N |
| ATOM | 6537 | H | GLN | A | 447 | 35.091 | 53.085 | 13.926 | 1.00 | 0.00 | H |
| ATOM | 6538 | CA | GLN | A | 447 | 34.125 | 54.488 | 12.651 | 1.00 | 0.00 | C |
| ATOM | 6539 | HA | GLN | A | 447 | 33.434 | 54.288 | 11.956 | 1.00 | 0.00 | H |
| ATOM | 6540 | CB | GLN | A | 447 | 33.625 | 55.582 | 13.589 | 1.00 | 0.00 | C |
| ATOM | 6541 | HB1 | GLN | A | 447 | 34.276 | 55.648 | 14.345 | 1.00 | 0.00 | H |
| ATOM | 6542 | HB2 | GLN | A | 447 | 32.735 | 55.290 | 13.940 | 1.00 | 0.00 | H |
| ATOM | 6543 | CG | GLN | A | 447 | 33.463 | 56.996 | 12.956 | 1.00 | 0.00 | C |
| ATOM | 6544 | HG1 | GLN | A | 447 | 32.686 | 57.448 | 13.394 | 1.00 | 0.00 | H |
| ATOM | 6545 | HG2 | GLN | A | 447 | 33.276 | 56.883 | 11.980 | 1.00 | 0.00 | H |
| ATOM | 6546 | CD | GLN | A | 447 | 34.690 | 57.922 | 13.097 | 1.00 | 0.00 | C |
| ATOM | 6547 | OE1 | GLN | A | 447 | 35.358 | 58.193 | 12.108 | 1.00 | 0.00 | O |
| ATOM | 6548 | NE2 | GLN | A | 447 | 34.997 | 58.478 | 14.274 | 1.00 | 0.00 | N |
| ATOM | 6549 | HE21 | GLN | A | 447 | 34.429 | 58.296 | 15.076 | 1.00 | 0.00 | H |
| ATOM | 6550 | HE22 | GLN | A | 447 | 35.795 | 59.076 | 14.352 | 1.00 | 0.00 | H |
| ATOM | 6551 | C | GLN | A | 447 | 35.429 | 54.928 | 11.955 | 1.00 | 0.00 | C |
| ATOM | 6552 | O | GLN | A | 447 | 36.482 | 54.899 | 12.587 | 1.00 | 0.00 | O |
| ATOM | 6553 | N | GLN | A | 448 | 35.335 | 55.460 | 10.725 | 1.00 | 0.00 | N |
| ATOM | 6554 | H | GLN | A | 448 | 34.476 | 55.336 | 10.228 | 1.00 | 0.00 | H |
| ATOM | 6555 | CA | GLN | A | 448 | 36.413 | 56.215 | 10.057 | 1.00 | 0.00 | C |
| ATOM | 6556 | HA | GLN | A | 448 | 36.831 | 56.717 | 10.814 | 1.00 | 0.00 | H |
| ATOM | 6557 | CB | GLN | A | 448 | 37.456 | 55.275 | 9.416 | 1.00 | 0.00 | C |
| ATOM | 6558 | HB1 | GLN | A | 448 | 37.852 | 54.714 | 10.143 | 1.00 | 0.00 | H |
| ATOM | 6559 | HB2 | GLN | A | 448 | 38.170 | 55.843 | 9.007 | 1.00 | 0.00 | H |
| ATOM | 6560 | CG | GLN | A | 448 | 36.933 | 54.325 | 8.326 | 1.00 | 0.00 | C |
| ATOM | 6561 | HG1 | GLN | A | 448 | 37.710 | 53.955 | 7.816 | 1.00 | 0.00 | H |
| ATOM | 6562 | HG2 | GLN | A | 448 | 36.340 | 54.837 | 7.705 | 1.00 | 0.00 | H |
| ATOM | 6563 | CD | GLN | A | 448 | 36.138 | 53.164 | 8.910 | 1.00 | 0.00 | C |
| ATOM | 6564 | OE1 | GLN | A | 448 | 36.671 | 52.281 | 9.562 | 1.00 | 0.00 | O |
| ATOM | 6565 | NE2 | GLN | A | 448 | 34.832 | 53.140 | 8.757 | 1.00 | 0.00 | N |
| ATOM | 6566 | HE21 | GLN | A | 448 | 34.373 | 53.878 | 8.262 | 1.00 | 0.00 | H |
| ATOM | 6567 | HE22 | GLN | A | 448 | 34.298 | 52.384 | 9.135 | 1.00 | 0.00 | H |
| ATOM | 6568 | C | GLN | A | 448 | 35.895 | 57.245 | 9.028 | 1.00 | 0.00 | C |
| ATOM | 6569 | O | GLN | A | 448 | 34.819 | 57.082 | 8.448 | 1.00 | 0.00 | O |
| ATOM | 6570 | N | SER | A | 449 | 36.717 | 58.261 | 8.727 | 1.00 | 0.00 | N |
| ATOM | 6571 | H | SER | A | 449 | 37.466 | 58.457 | 9.360 | 1.00 | 0.00 | H |
| ATOM | 6572 | CA | SER | A | 449 | 36.594 | 59.112 | 7.524 | 1.00 | 0.00 | C |
| ATOM | 6573 | HA | SER | A | 449 | 35.688 | 59.532 | 7.577 | 1.00 | 0.00 | H |
| ATOM | 6574 | CB | SER | A | 449 | 37.700 | 60.182 | 7.511 | 1.00 | 0.00 | C |
| ATOM | 6575 | HB1 | SER | A | 449 | 38.598 | 59.746 | 7.458 | 1.00 | 0.00 | H |
| ATOM | 6576 | HB2 | SER | A | 449 | 37.648 | 60.739 | 8.340 | 1.00 | 0.00 | H |
| ATOM | 6577 | OG | SER | A | 449 | 37.635 | 61.085 | 6.425 | 1.00 | 0.00 | O |
| ATOM | 6578 | HG | SER | A | 449 | 38.385 | 61.743 | 6.493 | 1.00 | 0.00 | H |
| ATOM | 6579 | C | SER | A | 449 | 36.715 | 58.267 | 6.255 | 1.00 | 0.00 | C |
| ATOM | 6580 | O | SER | A | 449 | 37.508 | 57.326 | 6.211 | 1.00 | 0.00 | O |
| ATOM | 6581 | N | CYS | A | 450 | 36.029 | 58.664 | 5.182 | 1.00 | 0.00 | N |
| ATOM | 6582 | H | CYS | A | 450 | 35.317 | 59.359 | 5.282 | 1.00 | 0.00 | H |
| ATOM | 6583 | CA | CYS | A | 450 | 36.304 | 58.098 | 3.862 | 1.00 | 0.00 | C |
| ATOM | 6584 | HA | CYS | A | 450 | 36.539 | 57.136 | 4.002 | 1.00 | 0.00 | H |
| ATOM | 6585 | CB | CYS | A | 450 | 35.044 | 58.165 | 2.993 | 1.00 | 0.00 | C |

| | | | | | | | | | | | |
|---|---|---|---|---|---|---|---|---|---|---|---|
| ATOM | 6586 | HB1 | CYS | A | 450 | 34.997 | 59.114 | 2.680 | 1.00 | 0.00 | H |
| ATOM | 6587 | HB2 | CYS | A | 450 | 34.288 | 57.984 | 3.622 | 1.00 | 0.00 | H |
| ATOM | 6588 | SG | CYS | A | 450 | 35.081 | 56.992 | 1.614 | 1.00 | 0.00 | S |
| ATOM | 6589 | C | CYS | A | 450 | 37.511 | 58.808 | 3.216 | 1.00 | 0.00 | C |
| ATOM | 6590 | O | CYS | A | 450 | 37.594 | 60.031 | 3.317 | 1.00 | 0.00 | O |
| ATOM | 6591 | N | PRO | A | 451 | 38.425 | 58.104 | 2.518 | 1.00 | 0.00 | N |
| ATOM | 6592 | CD | PRO | A | 451 | 38.590 | 56.654 | 2.505 | 1.00 | 0.00 | C |
| ATOM | 6593 | HD1 | PRO | A | 451 | 37.732 | 56.220 | 2.231 | 1.00 | 0.00 | H |
| ATOM | 6594 | HD2 | PRO | A | 451 | 38.847 | 56.337 | 3.418 | 1.00 | 0.00 | H |
| ATOM | 6595 | CG | PRO | A | 451 | 39.698 | 56.381 | 1.491 | 1.00 | 0.00 | C |
| ATOM | 6596 | HG1 | PRO | A | 451 | 39.324 | 56.266 | 0.571 | 1.00 | 0.00 | H |
| ATOM | 6597 | HG2 | PRO | A | 451 | 40.223 | 55.567 | 1.741 | 1.00 | 0.00 | H |
| ATOM | 6598 | CB | PRO | A | 451 | 40.558 | 57.636 | 1.580 | 1.00 | 0.00 | C |
| ATOM | 6599 | HB1 | PRO | A | 451 | 41.098 | 57.765 | 0.749 | 1.00 | 0.00 | H |
| ATOM | 6600 | HB2 | PRO | A | 451 | 41.168 | 57.602 | 2.372 | 1.00 | 0.00 | H |
| ATOM | 6601 | CA | PRO | A | 451 | 39.500 | 58.732 | 1.735 | 1.00 | 0.00 | C |
| ATOM | 6602 | HA | PRO | A | 451 | 39.825 | 59.535 | 2.235 | 1.00 | 0.00 | H |
| ATOM | 6603 | C | PRO | A | 451 | 39.034 | 59.265 | 0.365 | 1.00 | 0.00 | C |
| ATOM | 6604 | O | PRO | A | 451 | 39.839 | 59.787 | -0.403 | 1.00 | 0.00 | O |
| ATOM | 6605 | N | THR | A | 452 | 37.750 | 59.106 | 0.030 | 1.00 | 0.00 | N |
| ATOM | 6606 | H | THR | A | 452 | 37.156 | 58.656 | 0.697 | 1.00 | 0.00 | H |
| ATOM | 6607 | CA | THR | A | 452 | 37.136 | 59.537 | -1.236 | 1.00 | 0.00 | C |
| ATOM | 6608 | HA | THR | A | 452 | 37.795 | 60.177 | -1.631 | 1.00 | 0.00 | H |
| ATOM | 6609 | CB | THR | A | 452 | 36.900 | 58.352 | -2.191 | 1.00 | 0.00 | C |
| ATOM | 6610 | HB | THR | A | 452 | 36.410 | 58.682 | -2.998 | 1.00 | 0.00 | H |
| ATOM | 6611 | CG2 | THR | A | 452 | 38.199 | 57.665 | -2.605 | 1.00 | 0.00 | C |
| ATOM | 6612 | HG21 | THR | A | 452 | 37.993 | 56.906 | -3.222 | 1.00 | 0.00 | H |
| ATOM | 6613 | HG22 | THR | A | 452 | 38.790 | 58.323 | -3.072 | 1.00 | 0.00 | H |
| ATOM | 6614 | HG23 | THR | A | 452 | 38.665 | 57.316 | -1.792 | 1.00 | 0.00 | H |
| ATOM | 6615 | OG1 | THR | A | 452 | 36.059 | 57.386 | -1.589 | 1.00 | 0.00 | O |
| ATOM | 6616 | HG1 | THR | A | 452 | 35.918 | 56.626 | -2.223 | 1.00 | 0.00 | H |
| ATOM | 6617 | C | THR | A | 452 | 35.798 | 60.220 | -0.971 | 1.00 | 0.00 | C |
| ATOM | 6618 | O | THR | A | 452 | 35.043 | 59.746 | -0.121 | 1.00 | 0.00 | O |
| ATOM | 6619 | N | GLY | A | 453 | 35.443 | 61.219 | -1.785 | 1.00 | 0.00 | N |
| ATOM | 6620 | H | GLY | A | 453 | 36.107 | 61.527 | -2.467 | 1.00 | 0.00 | H |
| ATOM | 6621 | CA | GLY | A | 453 | 34.139 | 61.893 | -1.741 | 1.00 | 0.00 | C |
| ATOM | 6622 | HA1 | GLY | A | 453 | 33.442 | 61.195 | -1.908 | 1.00 | 0.00 | H |
| ATOM | 6623 | HA2 | GLY | A | 453 | 34.128 | 62.560 | -2.486 | 1.00 | 0.00 | H |
| ATOM | 6624 | C | GLY | A | 453 | 33.826 | 62.610 | -0.420 | 1.00 | 0.00 | C |
| ATOM | 6625 | O | GLY | A | 453 | 34.649 | 62.642 | 0.493 | 1.00 | 0.00 | O |
| ATOM | 6626 | N | LEU | A | 454 | 32.627 | 63.202 | -0.323 | 1.00 | 0.00 | N |
| ATOM | 6627 | H | LEU | A | 454 | 32.003 | 63.145 | -1.102 | 1.00 | 0.00 | H |
| ATOM | 6628 | CA | LEU | A | 454 | 32.187 | 63.931 | 0.877 | 1.00 | 0.00 | C |
| ATOM | 6629 | HA | LEU | A | 454 | 32.974 | 63.765 | 1.471 | 1.00 | 0.00 | H |
| ATOM | 6630 | CB | LEU | A | 454 | 31.947 | 65.420 | 0.565 | 1.00 | 0.00 | C |
| ATOM | 6631 | HB1 | LEU | A | 454 | 31.310 | 65.781 | 1.246 | 1.00 | 0.00 | H |
| ATOM | 6632 | HB2 | LEU | A | 454 | 31.535 | 65.484 | -0.344 | 1.00 | 0.00 | H |
| ATOM | 6633 | CG | LEU | A | 454 | 33.193 | 66.308 | 0.576 | 1.00 | 0.00 | C |
| ATOM | 6634 | HG | LEU | A | 454 | 33.873 | 65.869 | -0.011 | 1.00 | 0.00 | H |
| ATOM | 6635 | CD1 | LEU | A | 454 | 32.756 | 67.700 | 0.090 | 1.00 | 0.00 | C |
| ATOM | 6636 | HD11 | LEU | A | 454 | 33.546 | 68.314 | 0.083 | 1.00 | 0.00 | H |
| ATOM | 6637 | HD12 | LEU | A | 454 | 32.381 | 67.628 | -0.834 | 1.00 | 0.00 | H |
| ATOM | 6638 | HD13 | LEU | A | 454 | 32.057 | 68.064 | 0.706 | 1.00 | 0.00 | H |
| ATOM | 6639 | CD2 | LEU | A | 454 | 33.846 | 66.470 | 1.963 | 1.00 | 0.00 | C |

| | | | | | | | | | | | |
|---|---|---|---|---|---|---|---|---|---|---|---|
| ATOM | 6640 | HD21 | LEU | A | 454 | 34.649 | 67.061 | 1.886 | 1.00 | 0.00 | H |
| ATOM | 6641 | HD22 | LEU | A | 454 | 33.189 | 66.881 | 2.595 | 1.00 | 0.00 | H |
| ATOM | 6642 | HD23 | LEU | A | 454 | 34.124 | 65.573 | 2.308 | 1.00 | 0.00 | H |
| ATOM | 6643 | C | LEU | A | 454 | 30.975 | 63.384 | 1.626 | 1.00 | 0.00 | C |
| ATOM | 6644 | O | LEU | A | 454 | 30.661 | 63.835 | 2.746 | 1.00 | 0.00 | O |
| ATOM | 6645 | N | VAL | A | 455 | 30.270 | 62.470 | 0.957 | 1.00 | 0.00 | N |
| ATOM | 6646 | H | VAL | A | 455 | 30.629 | 62.119 | 0.092 | 1.00 | 0.00 | H |
| ATOM | 6647 | CA | VAL | A | 455 | 29.000 | 61.969 | 1.444 | 1.00 | 0.00 | C |
| ATOM | 6648 | HA | VAL | A | 455 | 29.077 | 62.059 | 2.437 | 1.00 | 0.00 | H |
| ATOM | 6649 | CB | VAL | A | 455 | 27.835 | 62.851 | 0.948 | 1.00 | 0.00 | C |
| ATOM | 6650 | HB | VAL | A | 455 | 28.062 | 63.771 | 1.267 | 1.00 | 0.00 | H |
| ATOM | 6651 | CG1 | VAL | A | 455 | 27.699 | 62.920 | -0.573 | 1.00 | 0.00 | C |
| ATOM | 6652 | HG11 | VAL | A | 455 | 26.926 | 63.507 | -0.812 | 1.00 | 0.00 | H |
| ATOM | 6653 | HG12 | VAL | A | 455 | 28.538 | 63.297 | -0.966 | 1.00 | 0.00 | H |
| ATOM | 6654 | HG13 | VAL | A | 455 | 27.544 | 62.001 | -0.936 | 1.00 | 0.00 | H |
| ATOM | 6655 | CG2 | VAL | A | 455 | 26.499 | 62.420 | 1.542 | 1.00 | 0.00 | C |
| ATOM | 6656 | HG21 | VAL | A | 455 | 25.773 | 63.015 | 1.196 | 1.00 | 0.00 | H |
| ATOM | 6657 | HG22 | VAL | A | 455 | 26.308 | 61.474 | 1.280 | 1.00 | 0.00 | H |
| ATOM | 6658 | HG23 | VAL | A | 455 | 26.540 | 62.488 | 2.539 | 1.00 | 0.00 | H |
| ATOM | 6659 | C | VAL | A | 455 | 28.805 | 60.480 | 1.263 | 1.00 | 0.00 | C |
| ATOM | 6660 | O | VAL | A | 455 | 28.993 | 59.950 | 0.166 | 1.00 | 0.00 | O |
| ATOM | 6661 | N | PHE | A | 456 | 28.420 | 59.811 | 2.356 | 1.00 | 0.00 | N |
| ATOM | 6662 | H | PHE | A | 456 | 28.314 | 60.298 | 3.223 | 1.00 | 0.00 | H |
| ATOM | 6663 | CA | PHE | A | 456 | 28.152 | 58.370 | 2.294 | 1.00 | 0.00 | C |
| ATOM | 6664 | HA | PHE | A | 456 | 28.552 | 58.011 | 1.451 | 1.00 | 0.00 | H |
| ATOM | 6665 | CB | PHE | A | 456 | 28.775 | 57.709 | 3.543 | 1.00 | 0.00 | C |
| ATOM | 6666 | HB1 | PHE | A | 456 | 28.205 | 57.932 | 4.334 | 1.00 | 0.00 | H |
| ATOM | 6667 | HB2 | PHE | A | 456 | 29.691 | 58.088 | 3.675 | 1.00 | 0.00 | H |
| ATOM | 6668 | CG | PHE | A | 456 | 28.911 | 56.200 | 3.493 | 1.00 | 0.00 | C |
| ATOM | 6669 | CD1 | PHE | A | 456 | 30.075 | 55.607 | 2.980 | 1.00 | 0.00 | C |
| ATOM | 6670 | HD1 | PHE | A | 456 | 30.787 | 56.179 | 2.573 | 1.00 | 0.00 | H |
| ATOM | 6671 | CE1 | PHE | A | 456 | 30.245 | 54.213 | 3.036 | 1.00 | 0.00 | C |
| ATOM | 6672 | HE1 | PHE | A | 456 | 31.077 | 53.795 | 2.671 | 1.00 | 0.00 | H |
| ATOM | 6673 | CZ | PHE | A | 456 | 29.245 | 53.408 | 3.606 | 1.00 | 0.00 | C |
| ATOM | 6674 | HZ | PHE | A | 456 | 29.369 | 52.417 | 3.654 | 1.00 | 0.00 | H |
| ATOM | 6675 | CE2 | PHE | A | 456 | 28.071 | 53.994 | 4.109 | 1.00 | 0.00 | C |
| ATOM | 6676 | HE2 | PHE | A | 456 | 27.352 | 53.423 | 4.506 | 1.00 | 0.00 | H |
| ATOM | 6677 | CD2 | PHE | A | 456 | 27.915 | 55.390 | 4.050 | 1.00 | 0.00 | C |
| ATOM | 6678 | HD2 | PHE | A | 456 | 27.082 | 55.809 | 4.411 | 1.00 | 0.00 | H |
| ATOM | 6679 | C | PHE | A | 456 | 26.657 | 58.132 | 2.255 | 1.00 | 0.00 | C |
| ATOM | 6680 | O | PHE | A | 456 | 25.915 | 58.530 | 3.143 | 1.00 | 0.00 | O |
| ATOM | 6681 | N | SER | A | 457 | 26.263 | 57.437 | 1.174 | 1.00 | 0.00 | N |
| ATOM | 6682 | H | SER | A | 457 | 26.929 | 57.166 | 0.479 | 1.00 | 0.00 | H |
| ATOM | 6683 | CA | SER | A | 457 | 24.848 | 57.078 | 1.020 | 1.00 | 0.00 | C |
| ATOM | 6684 | HA | SER | A | 457 | 24.317 | 57.708 | 1.587 | 1.00 | 0.00 | H |
| ATOM | 6685 | CB | SER | A | 457 | 24.475 | 57.189 | -0.467 | 1.00 | 0.00 | C |
| ATOM | 6686 | HB1 | SER | A | 457 | 25.022 | 56.557 | -1.016 | 1.00 | 0.00 | H |
| ATOM | 6687 | HB2 | SER | A | 457 | 24.615 | 58.123 | -0.797 | 1.00 | 0.00 | H |
| ATOM | 6688 | OG | SER | A | 457 | 23.093 | 56.857 | -0.663 | 1.00 | 0.00 | O |
| ATOM | 6689 | HG | SER | A | 457 | 22.866 | 56.934 | -1.634 | 1.00 | 0.00 | H |
| ATOM | 6690 | C | SER | A | 457 | 24.601 | 55.666 | 1.505 | 1.00 | 0.00 | C |
| ATOM | 6691 | O | SER | A | 457 | 25.390 | 54.740 | 1.298 | 1.00 | 0.00 | O |
| ATOM | 6692 | N | ASN | A | 458 | 23.359 | 55.491 | 1.988 | 1.00 | 0.00 | N |
| ATOM | 6693 | H | ASN | A | 458 | 22.715 | 56.255 | 2.009 | 1.00 | 0.00 | H |

| | | | | | | | | | | | |
|---|---|---|---|---|---|---|---|---|---|---|---|
| ATOM | 6694 | CA | ASN | A | 458 | 22.954 | 54.167 | 2.486 | 1.00 | 0.00 | C |
| ATOM | 6695 | HA | ASN | A | 458 | 23.805 | 53.792 | 2.853 | 1.00 | 0.00 | H |
| ATOM | 6696 | CB | ASN | A | 458 | 21.852 | 54.321 | 3.548 | 1.00 | 0.00 | C |
| ATOM | 6697 | HB1 | ASN | A | 458 | 20.974 | 54.468 | 3.093 | 1.00 | 0.00 | H |
| ATOM | 6698 | HB2 | ASN | A | 458 | 22.062 | 55.108 | 4.129 | 1.00 | 0.00 | H |
| ATOM | 6699 | CG | ASN | A | 458 | 21.740 | 53.093 | 4.424 | 1.00 | 0.00 | C |
| ATOM | 6700 | OD1 | ASN | A | 458 | 22.492 | 52.130 | 4.354 | 1.00 | 0.00 | O |
| ATOM | 6701 | ND2 | ASN | A | 458 | 20.740 | 53.206 | 5.307 | 1.00 | 0.00 | N |
| ATOM | 6702 | HD21 | ASN | A | 458 | 20.179 | 54.034 | 5.324 | 1.00 | 0.00 | H |
| ATOM | 6703 | HD22 | ASN | A | 458 | 20.554 | 52.463 | 5.950 | 1.00 | 0.00 | H |
| ATOM | 6704 | C | ASN | A | 458 | 22.482 | 53.204 | 1.418 | 1.00 | 0.00 | C |
| ATOM | 6705 | O | ASN | A | 458 | 22.435 | 51.997 | 1.610 | 1.00 | 0.00 | O |
| ATOM | 6706 | N | SER | A | 459 | 22.170 | 53.808 | 0.259 | 1.00 | 0.00 | N |
| ATOM | 6707 | H | SER | A | 459 | 22.233 | 54.803 | 0.177 | 1.00 | 0.00 | H |
| ATOM | 6708 | CA | SER | A | 459 | 21.736 | 52.998 | -0.887 | 1.00 | 0.00 | C |
| ATOM | 6709 | HA | SER | A | 459 | 21.041 | 52.386 | -0.509 | 1.00 | 0.00 | H |
| ATOM | 6710 | CB | SER | A | 459 | 21.116 | 53.876 | -1.991 | 1.00 | 0.00 | C |
| ATOM | 6711 | HB1 | SER | A | 459 | 21.787 | 54.532 | -2.338 | 1.00 | 0.00 | H |
| ATOM | 6712 | HB2 | SER | A | 459 | 20.318 | 54.368 | -1.643 | 1.00 | 0.00 | H |
| ATOM | 6713 | OG | SER | A | 459 | 20.682 | 53.073 | -3.097 | 1.00 | 0.00 | O |
| ATOM | 6714 | HG | SER | A | 459 | 20.284 | 53.662 | -3.800 | 1.00 | 0.00 | H |
| ATOM | 6715 | C | SER | A | 459 | 22.849 | 52.126 | -1.433 | 1.00 | 0.00 | C |
| ATOM | 6716 | O | SER | A | 459 | 22.724 | 50.916 | -1.553 | 1.00 | 0.00 | O |
| ATOM | 6717 | N | CYS | A | 460 | 23.958 | 52.801 | -1.758 | 1.00 | 0.00 | N |
| ATOM | 6718 | H | CYS | A | 460 | 23.985 | 53.790 | -1.614 | 1.00 | 0.00 | H |
| ATOM | 6719 | CA | CYS | A | 460 | 25.130 | 52.132 | -2.317 | 1.00 | 0.00 | C |
| ATOM | 6720 | HA | CYS | A | 460 | 24.767 | 51.331 | -2.792 | 1.00 | 0.00 | H |
| ATOM | 6721 | CB | CYS | A | 460 | 25.826 | 53.085 | -3.289 | 1.00 | 0.00 | C |
| ATOM | 6722 | HB1 | CYS | A | 460 | 25.134 | 53.277 | -3.985 | 1.00 | 0.00 | H |
| ATOM | 6723 | HB2 | CYS | A | 460 | 26.559 | 52.536 | -3.691 | 1.00 | 0.00 | H |
| ATOM | 6724 | SG | CYS | A | 460 | 26.457 | 54.601 | -2.527 | 1.00 | 0.00 | S |
| ATOM | 6725 | C | CYS | A | 460 | 26.106 | 51.612 | -1.250 | 1.00 | 0.00 | C |
| ATOM | 6726 | O | CYS | A | 460 | 27.068 | 50.934 | -1.603 | 1.00 | 0.00 | O |
| ATOM | 6727 | N | LYS | A | 461 | 25.911 | 51.976 | 0.031 | 1.00 | 0.00 | N |
| ATOM | 6728 | H | LYS | A | 461 | 25.040 | 52.410 | 0.262 | 1.00 | 0.00 | H |
| ATOM | 6729 | CA | LYS | A | 461 | 26.889 | 51.782 | 1.122 | 1.00 | 0.00 | C |
| ATOM | 6730 | HA | LYS | A | 461 | 26.541 | 52.303 | 1.901 | 1.00 | 0.00 | H |
| ATOM | 6731 | CB | LYS | A | 461 | 26.966 | 50.296 | 1.522 | 1.00 | 0.00 | C |
| ATOM | 6732 | HB1 | LYS | A | 461 | 27.547 | 50.224 | 2.333 | 1.00 | 0.00 | H |
| ATOM | 6733 | HB2 | LYS | A | 461 | 27.385 | 49.793 | 0.766 | 1.00 | 0.00 | H |
| ATOM | 6734 | CG | LYS | A | 461 | 25.624 | 49.626 | 1.844 | 1.00 | 0.00 | C |
| ATOM | 6735 | HG1 | LYS | A | 461 | 25.764 | 48.636 | 1.867 | 1.00 | 0.00 | H |
| ATOM | 6736 | HG2 | LYS | A | 461 | 24.976 | 49.854 | 1.118 | 1.00 | 0.00 | H |
| ATOM | 6737 | CD | LYS | A | 461 | 25.019 | 50.060 | 3.185 | 1.00 | 0.00 | C |
| ATOM | 6738 | HD1 | LYS | A | 461 | 24.774 | 51.029 | 3.158 | 1.00 | 0.00 | H |
| ATOM | 6739 | HD2 | LYS | A | 461 | 25.670 | 49.901 | 3.928 | 1.00 | 0.00 | H |
| ATOM | 6740 | CE | LYS | A | 461 | 23.767 | 49.203 | 3.385 | 1.00 | 0.00 | C |
| ATOM | 6741 | HE1 | LYS | A | 461 | 23.961 | 48.270 | 3.081 | 1.00 | 0.00 | H |
| ATOM | 6742 | HE2 | LYS | A | 461 | 23.022 | 49.584 | 2.837 | 1.00 | 0.00 | H |
| ATOM | 6743 | NZ | LYS | A | 461 | 23.292 | 49.124 | 4.785 | 1.00 | 0.00 | N |
| ATOM | 6744 | HZ1 | LYS | A | 461 | 22.475 | 48.549 | 4.828 | 1.00 | 0.00 | H |
| ATOM | 6745 | HZ2 | LYS | A | 461 | 24.009 | 48.728 | 5.358 | 1.00 | 0.00 | H |
| ATOM | 6746 | HZ3 | LYS | A | 461 | 23.071 | 50.042 | 5.114 | 1.00 | 0.00 | H |
| ATOM | 6747 | C | LYS | A | 461 | 28.264 | 52.364 | 0.751 | 1.00 | 0.00 | C |

| | | | | | | | | | | | |
|---|---|---|---|---|---|---|---|---|---|---|---|
| ATOM | 6748 | O | LYS | A | 461 | 29.291 | 51.707 | 0.919 | 1.00 | 0.00 | O |
| ATOM | 6749 | N | CYS | A | 462 | 28.267 | 53.546 | 0.132 | 1.00 | 0.00 | N |
| ATOM | 6750 | H | CYS | A | 462 | 27.440 | 54.106 | 0.173 | 1.00 | 0.00 | H |
| ATOM | 6751 | CA | CYS | A | 462 | 29.419 | 54.062 | -0.602 | 1.00 | 0.00 | C |
| ATOM | 6752 | HA | CYS | A | 462 | 30.213 | 53.653 | -0.152 | 1.00 | 0.00 | H |
| ATOM | 6753 | CB | CYS | A | 462 | 29.312 | 53.617 | -2.070 | 1.00 | 0.00 | C |
| ATOM | 6754 | HB1 | CYS | A | 462 | 28.906 | 52.705 | -2.005 | 1.00 | 0.00 | H |
| ATOM | 6755 | HB2 | CYS | A | 462 | 30.267 | 53.537 | -2.356 | 1.00 | 0.00 | H |
| ATOM | 6756 | SG | CYS | A | 462 | 28.371 | 54.679 | -3.203 | 1.00 | 0.00 | S |
| ATOM | 6757 | C | CYS | A | 462 | 29.596 | 55.581 | -0.481 | 1.00 | 0.00 | C |
| ATOM | 6758 | O | CYS | A | 462 | 28.634 | 56.325 | -0.276 | 1.00 | 0.00 | O |
| ATOM | 6759 | N | CYS | A | 463 | 30.837 | 56.034 | -0.668 | 1.00 | 0.00 | N |
| ATOM | 6760 | H | CYS | A | 463 | 31.558 | 55.355 | -0.809 | 1.00 | 0.00 | H |
| ATOM | 6761 | CA | CYS | A | 463 | 31.228 | 57.442 | -0.683 | 1.00 | 0.00 | C |
| ATOM | 6762 | HA | CYS | A | 463 | 30.613 | 57.897 | -0.039 | 1.00 | 0.00 | H |
| ATOM | 6763 | CB | CYS | A | 463 | 32.683 | 57.591 | -0.220 | 1.00 | 0.00 | C |
| ATOM | 6764 | HB1 | CYS | A | 463 | 32.790 | 58.572 | -0.058 | 1.00 | 0.00 | H |
| ATOM | 6765 | HB2 | CYS | A | 463 | 33.223 | 57.324 | -1.018 | 1.00 | 0.00 | H |
| ATOM | 6766 | SG | CYS | A | 463 | 33.122 | 56.619 | 1.242 | 1.00 | 0.00 | S |
| ATOM | 6767 | C | CYS | A | 463 | 31.041 | 58.052 | -2.080 | 1.00 | 0.00 | C |
| ATOM | 6768 | O | CYS | A | 463 | 31.390 | 57.432 | -3.089 | 1.00 | 0.00 | O |
| ATOM | 6769 | N | THR | A | 464 | 30.524 | 59.280 | -2.114 | 1.00 | 0.00 | N |
| ATOM | 6770 | H | THR | A | 464 | 30.405 | 59.739 | -1.234 | 1.00 | 0.00 | H |
| ATOM | 6771 | CA | THR | A | 464 | 30.110 | 60.032 | -3.311 | 1.00 | 0.00 | C |
| ATOM | 6772 | HA | THR | A | 464 | 30.707 | 59.719 | -4.050 | 1.00 | 0.00 | H |
| ATOM | 6773 | CB | THR | A | 464 | 28.631 | 59.730 | -3.636 | 1.00 | 0.00 | C |
| ATOM | 6774 | HB | THR | A | 464 | 28.343 | 60.364 | -4.353 | 1.00 | 0.00 | H |
| ATOM | 6775 | CG2 | THR | A | 464 | 28.386 | 58.286 | -4.061 | 1.00 | 0.00 | C |
| ATOM | 6776 | HG21 | THR | A | 464 | 27.414 | 58.156 | -4.256 | 1.00 | 0.00 | H |
| ATOM | 6777 | HG22 | THR | A | 464 | 28.921 | 58.085 | -4.882 | 1.00 | 0.00 | H |
| ATOM | 6778 | HG23 | THR | A | 464 | 28.663 | 57.670 | -3.324 | 1.00 | 0.00 | H |
| ATOM | 6779 | OG1 | THR | A | 464 | 27.805 | 59.975 | -2.521 | 1.00 | 0.00 | O |
| ATOM | 6780 | HG1 | THR | A | 464 | 26.855 | 59.772 | -2.759 | 1.00 | 0.00 | H |
| ATOM | 6781 | C | THR | A | 464 | 30.301 | 61.545 | -3.101 | 1.00 | 0.00 | C |
| ATOM | 6782 | O | THR | A | 464 | 30.819 | 61.974 | -2.068 | 1.00 | 0.00 | O |
| ATOM | 6783 | N | TRP | A | 465 | 29.892 | 62.378 | -4.059 | 1.00 | 0.00 | N |
| ATOM | 6784 | H | TRP | A | 465 | 29.749 | 61.988 | -4.969 | 1.00 | 0.00 | H |
| ATOM | 6785 | CA | TRP | A | 465 | 29.633 | 63.814 | -3.901 | 1.00 | 0.00 | C |
| ATOM | 6786 | HA | TRP | A | 465 | 29.140 | 63.916 | -3.037 | 1.00 | 0.00 | H |
| ATOM | 6787 | CB | TRP | A | 465 | 30.941 | 64.622 | -3.763 | 1.00 | 0.00 | C |
| ATOM | 6788 | HB1 | TRP | A | 465 | 31.382 | 64.353 | -2.907 | 1.00 | 0.00 | H |
| ATOM | 6789 | HB2 | TRP | A | 465 | 30.702 | 65.592 | -3.724 | 1.00 | 0.00 | H |
| ATOM | 6790 | CG | TRP | A | 465 | 31.976 | 64.482 | -4.835 | 1.00 | 0.00 | C |
| ATOM | 6791 | CD1 | TRP | A | 465 | 32.872 | 63.476 | -4.941 | 1.00 | 0.00 | C |
| ATOM | 6792 | HD1 | TRP | A | 465 | 32.942 | 62.702 | -4.311 | 1.00 | 0.00 | H |
| ATOM | 6793 | NE1 | TRP | A | 465 | 33.677 | 63.673 | -6.047 | 1.00 | 0.00 | N |
| ATOM | 6794 | HE1 | TRP | A | 465 | 34.423 | 63.065 | -6.319 | 1.00 | 0.00 | H |
| ATOM | 6795 | CE2 | TRP | A | 465 | 33.318 | 64.814 | -6.725 | 1.00 | 0.00 | C |
| ATOM | 6796 | CZ2 | TRP | A | 465 | 33.774 | 65.395 | -7.916 | 1.00 | 0.00 | C |
| ATOM | 6797 | HZ2 | TRP | A | 465 | 34.566 | 65.017 | -8.395 | 1.00 | 0.00 | H |
| ATOM | 6798 | CH2 | TRP | A | 465 | 33.102 | 66.515 | -8.426 | 1.00 | 0.00 | C |
| ATOM | 6799 | HH2 | TRP | A | 465 | 33.386 | 66.915 | -9.298 | 1.00 | 0.00 | H |
| ATOM | 6800 | CZ3 | TRP | A | 465 | 32.028 | 67.076 | -7.710 | 1.00 | 0.00 | C |
| ATOM | 6801 | HZ3 | TRP | A | 465 | 31.556 | 67.880 | -8.071 | 1.00 | 0.00 | H |

```
ATOM   6802  CE3 TRP A 465      31.614  66.511  -6.488  1.00  0.00           C
ATOM   6803  HE3 TRP A 465      30.876  66.943  -5.970  1.00  0.00           H
ATOM   6804  CD2 TRP A 465      32.225  65.345  -5.984  1.00  0.00           C
ATOM   6805  C   TRP A 465      28.731  64.305  -5.047  1.00  0.00           C
ATOM   6806  O   TRP A 465      28.358  63.529  -5.928  1.00  0.00           O
ATOM   6807  N   ASN A 466      28.283  65.567  -4.890  1.00  0.00           N
ATOM   6808  H   ASN A 466      28.582  66.110  -4.105  1.00  0.00           H
ATOM   6809  CA  ASN A 466      27.352  66.147  -5.876  1.00  0.00           C
ATOM   6810  HA  ASN A 466      27.640  65.803  -6.770  1.00  0.00           H
ATOM   6811  CB  ASN A 466      25.901  65.700  -5.600  1.00  0.00           C
ATOM   6812  HB1 ASN A 466      25.580  66.144  -4.763  1.00  0.00           H
ATOM   6813  HB2 ASN A 466      25.887  64.708  -5.477  1.00  0.00           H
ATOM   6814  CG  ASN A 466      24.935  66.048  -6.716  1.00  0.00           C
ATOM   6815  OD1 ASN A 466      23.717  66.026  -6.607  1.00  0.00           O
ATOM   6816  ND2 ASN A 466      25.579  66.410  -7.835  1.00  0.00           N
ATOM   6817 HD21 ASN A 466      26.579  66.424  -7.854  1.00  0.00           H
ATOM   6818 HD22 ASN A 466      25.060  66.666  -8.650  1.00  0.00           H
ATOM   6819  C   ASN A 466      27.466  67.659  -5.877  1.00  0.00           C
ATOM   6820  OC1 ASN A 466      28.249  68.074  -6.341  1.00  0.00           O
ATOM   6821  OC2 ASN A 466      26.651  68.399  -5.340  1.00  0.00           O
TER
ENDMDL
```
